# Virally induced lipid droplets are a platform for innate immune signalling complexes

**DOI:** 10.1101/2023.04.20.537741

**Authors:** EA Monson, JL Laws, Z Telikani, AJ Milligan, AM Rozario, I Amarasinghe, ML Smith, V Tran, Q Dinh, N Williamson, A Mechler, C Johnson, MJ Hofer, S Nie, DR Whelan, KJ Helbig

## Abstract

Lipid droplets (LDs) are upregulated by host cells in the face of pathogen infection, however, the reason for this phenomenon remains largely unknown. Here, we demonstrate that virally induced LDs house a distinct and dynamic proteome containing key antiviral signalling pathway members, including the essential pattern recognition receptor; RIG-I, key adaptor proteins; STAT1 and STAT2 and prominent interferon inducible proteins; viperin and MX1. Changes in the LD proteome were underpinned by specific key changes in the lipidome of virally driven LDs, particularity in the phospholipid membrane. Following virus infection, key antiviral proteins formed complex protein-protein interactions on the LD surface, positioning this organelle as a key antiviral signalling platform for the first time. It is clear that dynamic regulation of both the proteome and the lipidome of LDs occurs rapidly following viral infection towards the initiation of a successful innate immune response.

## Introduction

Lipid droplets (LDs) were initially considered simply as a cellular energy source, but are now recognised as dynamic, cytoplasmic organelles that are critical in many signalling events (*1*– *3*). It has been well described that LDs are upregulated in the face of pathogen infection to multiple bacteria, parasites and more recently, viruses (reviewed in (*4*)). LD upregulation during virus infection is now known to be essential in supporting a successful innate immune response in host cells (*5, 6*). In particular, during zika virus (ZIKV) and herpesvirus (HSV-1) infection, LD upregulation has been linked to the production of type-I and III interferons (IFNs) (IFN-β and IFN-λ) (*5*), however, the mechanism underpinning this increase in IFN production still remains elusive, and could involve both changes in lipid and protein profiles. Successful antiviral responses rely on complex protein interactions that require platforms for their assembly (*7*–*9*). The LD is well described to house viperin, a key antiviral protein that also regulates multiple antiviral signalling cascades (*1, 2, 10*), however, the role of the LD in facilitating other protein complex formations remains unknown. The LD proteome is both diverse and dynamic in nature, changing constantly to cellular cues; with its make-up also being driven by changes in lipid species (*11*). Given the recent emergence of LDs as essential organelles in host driven antiviral immunity, it is possible that these organelles play undescribed roles as signalling platforms during host response to viral infection.

## Results

### Antiviral innate immune signalling proteins are recruited to LDs during virus infection

LDs are upregulated early following viral infection in mammalian cells and are essential for the production of a heightened antiviral IFN state (*5*); however, the mechanisms that underpin their ability to drive IFN production and limit early viral replication remain unknown. To investigate this further we selected a well-established neural model of lymphocytic choriomeningitis virus (LCMV) infection (*12*). Mice were infected intracranially with LCMV, and their brains harvested at 2- and 4-days post infection (dpi), modelling early-stage infection (Fig S1A-C), with virally infected mice displaying an enhanced level of LDs in their brains at both 2 and 4 dpi (Fig 1A, B). (Fig 1C, S1D). To characterise the enhanced antiviral capacity of virally driven LDs, we performed comparative mass spectrometry profiling of proteins differentially associated with LCMV-LDs or SMAM-LDs isolated from the brain tissues of mice. In the virally infected brains, 4324 proteins were identified on LDs (Fig 1D; Table S1; S1E), with only 30 of these found to be differentially upregulated, and 24 of these to be exclusively upregulated on LDs (0.5%; Fig 1E, S1E). Functional annotation enrichment analysis revealed that most of the upregulated LD proteins at 4 dpi belonged to ‘cellular defence response to viruses’ and ‘response to interferons’ (Fig 1F, G), the predominant mammalian antiviral cytokines. The brain contains a very heterogenous mix of cell types, and immunofluorescence analysis revealed that astrocytes were the largest contributor in the brain to the upregulated LD response to virus (Fig 1H, S2).

**Figure 1.**
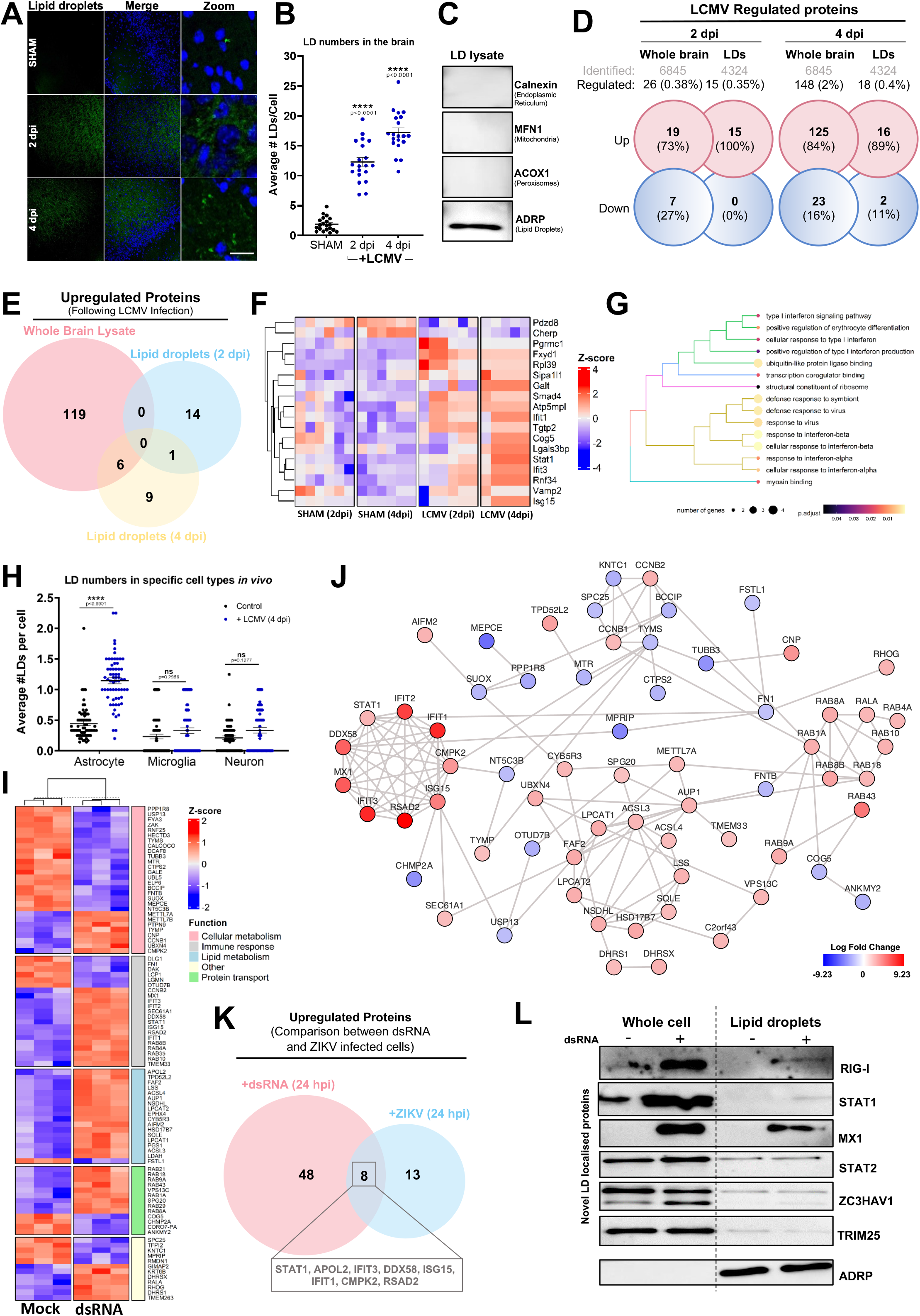
The proteome of lipid droplets changes significantly during virus infection *in vivo* and *in vitro*. **(A)** Mice were sham injected or intracranially infected with LCMV (500 PFU) for 2 or 4 days. Brains were harvested and sectioned for immunostaining. LDs were stained with Bodipy (493/503) (green) and nuclei with DAPI (blue). **(B)** LD numbers were analysed via ImageJ analysis software. Error bars represent ± SEM, n = 6 replicate mice per condition. **(C)** To confirm the purity of isolated LDs from mice brains western blot analysis of LD lysates was performed probing for organelle markers for the endoplasmic reticulum (Calnexin), mitochondria (MFN1), peroxisomes (ACOX1) and LDs (ADRP). Results indicate enrichment of LDs and no contamination of other cellular organelles. **(D)** Comparative analysis of protein ID and abundance were analysed by LC-MS/MS highlighting the number of proteins significantly regulated from both whole brain lysate and isolated LD lysate derived from mice brains following 4 dpi of LCMV. Red and blue tiles indicate up- and down regulated proteins, respectively **(E)** Significantly upregulated proteins following LCMV infection were compared with one another to highlight distinct proteomic profiles between LDs and whole brain lysates. Red circle reflects proteins exclusive to the whole brain lysate, blue circle reflects proteins exclusive to LDs 2dpi and the yellow circle reflects proteins exclusive to LDs 4dpi. **(F)** Hierarchical clustering of the 18 significantly regulated LD resident proteins 4 dpi of LCMV across replicates and their respective z-scores from the groups: control 2 dpi, control 4 dpi, LCMV 2dpi and LCMV 4dpi. Blue tiles refer to proteins with a z-score < 0 with red tiles referring to proteins with a z-score > 0. **(G)** Tree-plot displaying similarity clusters of the top 15 significantly enriched annotations from the gene ontology categories: Biological Process (BP), Molecular Function (MF) and Cellular Component (CC). Only annotations that surpassed an adjusted p-value of less than 0.05 were included. (**H**) Analysis of average number of LDs localised to each brain cell type with and without LCMV 4 dpi. Error bars represent values ± SEM. *P* values were determined by two-way ANOVA post-hoc pairwise comparisons with Bonferroni correction (n= 64 tissue sections over 6 mice). **(I)** Hierarchical clustering displaying the z-scores and gene ontology between replicates of the 92 significantly regulated astrocyte derived LD proteins following dsRNA stimulation at 24 hrs. Blue tiles refer to proteins with a z-score < 0 with red tiles referring to proteins with a z-score > 0. Gene ontology was performed using the Uniprot database within Perseus. Proteins were grouped into 5 main categories: cellular metabolism, immune response, lipid metabolism, protein transport and other. **(J)** Interaction diagram of significantly changed proteins following dsRNA viral mimic stimulation created using STRING confidence scores and interactions in cytoscape. Nodes are coloured in a continuous scale based off their log2 fold change, with red coloured nodes having the highest fold change and blue nodes having the greatest negative fold change. Proteins with no interactions with any other proteins were not represented within the network. All clusters are clustered using the ClusterONE plugin within cytoscape with a p < 0.05 significance threshold. **(K)** Similarities and differences between significantly enriched proteins from astrocyte derived LDs infected with either dsRNA or ZIKV at 24 hrs respectively. **(L)** Immunoblot analysis on whole cell lysate and isolated LD lysate from both mock and dsRNA stimulated astrocytes at 24 hrs confirmed the localisation of MX1, RIG-I, STAT1, STAT2, ZC3HAV1 and TRIM25 proteins on isolated LDs derived from cells that have been stimulated with dsRNA for 24 hrs. ADRP was used to confirm enrichment of purified LDs.

To gain a cell type specific LD proteomic profile response to viral infection in the absence of viral antagonism, we described the proteomic shift of LDs following viral mimic stimulation (dsRNA) in primary immortalised astrocyte cells. *In vitro* analysis of an immortalised primary astrocyte cell model following dsRNA stimulation at both 8 and 24 hrs revealed a significant upregulation of LDs at both time points, as we have seen previously (Fig S3A) (*5*). The proteomic profiles of isolated LDs revealed an upregulation of 10 and 56 proteins at 8 and 24 hrs respectively following stimulation of cells with dsRNA (Fig S3C and D). Functional categorisation of the entire LD proteome in comparison to the differentially regulated LD proteins following stimulation (24 hrs) revealed an increase in the distribution of proteins belonging to the ‘immune response’ and ‘lipid metabolism’ functional categories (Fig S3E, Fig 1I and Table 2); additionally, there were a significant number of enzymes present on the LD that facilitate protein and lipid post-translational modifications (Fig S3F). Further analysis revealed that protein members belonging to the ‘immune response’ cluster had a high level of predicted protein-protein associations with each other, and members of this cluster were also upregulated on the LD following an RNA viral infection (zika virus) at the same time point (24 hrs: Fig 1J, K, Table S2 and S3). Immunoblotting of LD fractions confirmed the presence of selected novel localised LD proteins, at steady state or upregulated following dsRNA stimulation (Fig 1L).

### The LD lipidome dynamically changes following activation of antiviral early innate signalling pathways

It is well established that viruses alter the cellular lipidome, often to enhance their own replication cycles (*13*). However, these changes are usually examined at later time points, excluding the possibility that early cellular lipid changes may occur to facilitate a pro-host response. Additionally, to our knowledge, the dynamics of lipids within the LD has never been examined following viral infection, therefore we applied a dual-omics approach to better understand the regulation of lipids and proteins on the LD during virus infection. Although there was limited changes to lipids in whole cell lysates following dsRNA stimulation of astrocyte cells, there was significant changes to the distribution of lipid categories of the LD lipidome as early as 8 hrs, with the greatest changes observed at 24 hrs, towards an increased abundance of glycerolipids, and a decrease in sterols (Fig 2A, S4A, B, Table S4). Further analysis of individual lipid classes revealed that following activation of antiviral pathways, LDs generally alter their lipidome to increase their distribution of long-chain polyunsaturated triacylglycerols, whilst decreasing their abundance of saturated cholesterol esters (Fig 2B, C, S4C-F). These lipid changes were underpinned by the LD proteome with the presence of important metabolic enzymes, in particular those responsible for the synthesis of long-chain polyunsaturated fatty acids (PUFAs), such as ACSL1, ACSL3 and ACSL4 (Fig 2D, E). A small but significant change was also observed in the structural lipids making up the LD phospholipid membrane, with an increase in PE (phosphatidylethanolamine) and PI (phosphatidylinositol) lipids (Fig 2F). Small changes in these membrane phospholipids are known to alter membrane curvature, stability and their ability to incorporate proteins (*14*–*18*); and additional analysis also revealed an increase of ether linkages in these structural lipids upregulated between 8 and 24 hrs (Fig 2G), known to support these functional changes, including support of cellular signalling at lipid membranes (*18*). Collectively, simultaneous proteomic and lipidomic analysis revealed that changes in the long and very long-chained PUFAs, as well as changes in the neutral lipids and membrane lipids were underpinned by dynamic proteomic changes to include important metabolic enzymes that drive these lipidomic alterations (Fig 2H).

**Figure 2:**
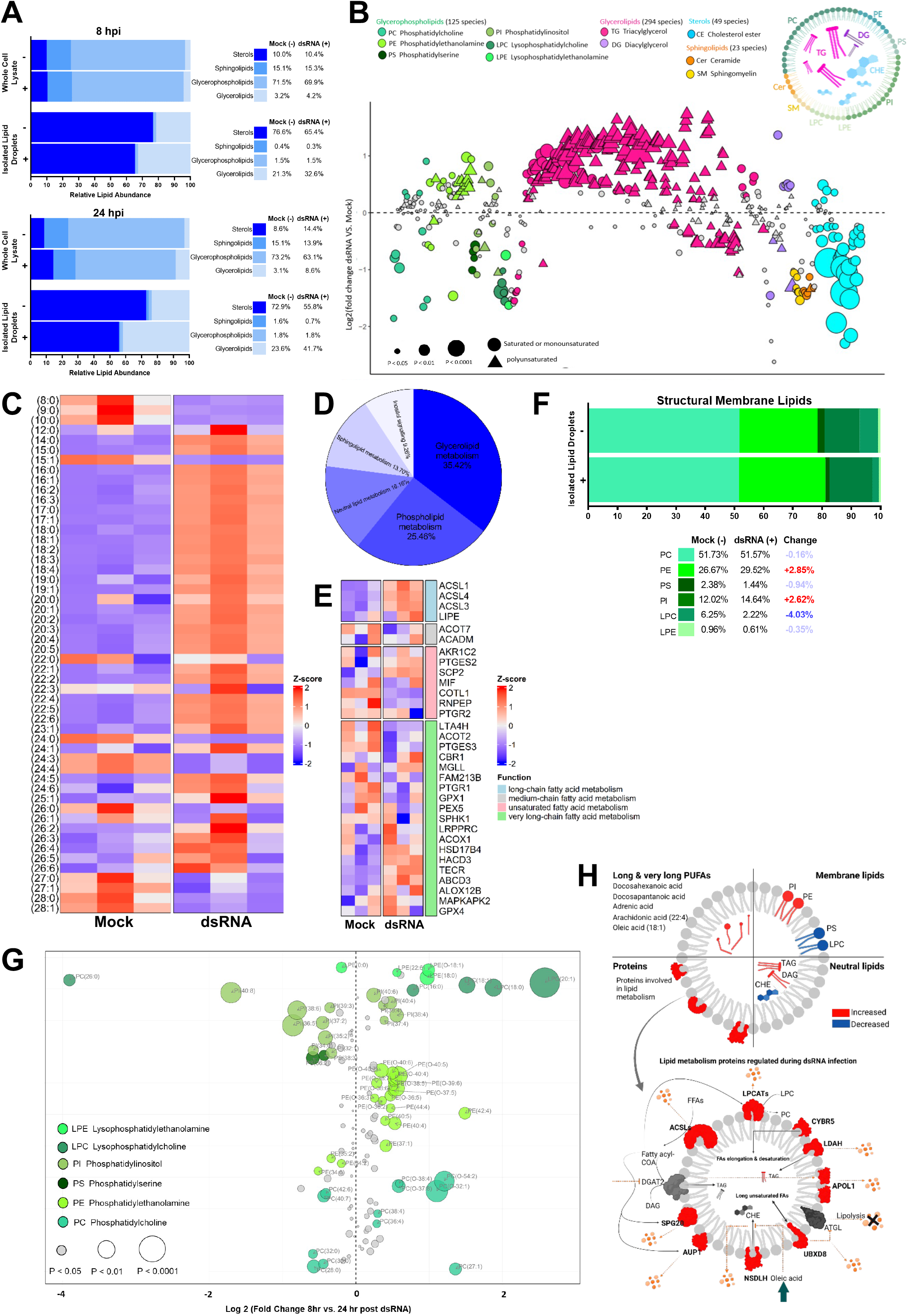
The lipidome of LDs changed significantly following dsRNA stimulation in primary immortalised astrocytes. **(A)** Relative abundance of major lipid categories identified in isolated whole cell lysates and LD fractions following dsRNA stimulation at 8 hrs and 24 hrs. LD fractions and whole cell lysates were analysed for changes in relative abundance of major lipid categories (Glycerolipids, Glycerophospholipids, Sphingolipids and Sterols) in following groups; Whole cell lysate post dsRNA stimulation at 8 and 24 hrs, LD fractions post dsRNA stimulation at 8 and 24 hrs. **(B)** Bubble plot of log2 fold changes in abundance of individual lipid species post dsRNA stimulation relative to mock at 24 hrs. Significance was determined by unpaired two-tailed students *t* test (n=3). Individual lipid species are coloured by the class of lipid that they belong to. PC phosphatidylcholine; PE phosphatidylethanolamine; PI phosphatidylinositol; PS phosphatidylserine; LPC lysophosphatidylcholine; LPE lysophosphatidylethanolamine; DAG diacylglycerol; TAG triacylglycerol; Cer ceramide; SM sphingomyelin; CE cholesterol ester. **(C)** Comparison of changes in relative abundance of fatty acids of all analysed lipid species in LD fraction isolated from dsRNA stimulated primary immortalized astrocytes; arranged from shortest to longest fatty acid chain lengths. **(D)** Distribution of identified LD resident proteins involved in lipid metabolism post dsRNA stimulation at 24 hrs based on the percentage abundance. Gene ontology was performed using the Uniprot database within Perseus. **(E)** Hierarchical clustering displaying the z-scores and gene ontology between replicates of the significantly regulated astrocyte derived LD proteins following dsRNA stimulation at 24 hrs. Blue tiles refer to proteins with a z-score < 0 with red tiles referring to proteins with a z-score > 0. Gene ontology was performed using the Uniprot database within Perseus. Proteins were grouped into 4 main categories (involved in regulating fatty acids metabolism). **(F)** Relative abundance of structural membrane phospholipids in LD fractions following dsRNA stimulation at 24 hrs. Changes in relative abundance of membrane phospholipids (PC phosphatidylcholine; PE phosphatidylethanolamine; PI phosphatidylinositol; PS phosphatidylserine; LPC lysophosphatidylcholine; LPE lysophosphatidylethanolamine) following dsRNA stimulation at 24 hrs in isolated LDs. **(G)** Bubble plot of log2 fold changes in relative abundance membrane phospholipids in isolated LDs following dsRNA stimulation from 8 hrs to 24 hrs. Individual phospholipid species (PC phosphatidylcholine; PE phosphatidylethanolamine; PI phosphatidylinositol; PS phosphatidylserine; LPC lysophosphatidylcholine; LPE lysophosphatidylethanolamine) characterized by abundance in LDs at 8 hrs relative to LDs at 24 hrs post dsRNA stimulation. Significance was determined by unpaired two-tailed students *t* test (n = 3). **(H)** Overview of changes in LD’s lipidome complemented by upregulation of LD‘s proteins involved in lipid metabolism. Proteins involved in lipid metabolism (long/very long/ poly unsaturated fatty acids (PUFAs) metabolism) were identified as significantly upregulated post dsRNA stimulation. These proteins could potentially play roles in LDs accumulation and protein recruitment to LDs (SPG20 & AUP1), fatty acids elongation and poly unsaturation (e.g., ACSLs, CYBR5 & UBXD8), increase in Triglycerides (e.g., APOL1 & LDAH), decrease in cholesterol esters (e.g., NSDHL) & also phospholipid synthesis (e.g., LPCAT), which were the main alterations observed in LDs lipidomic profile post dsRNA stimulation in primary immortalised astrocytes at 24 hrs.

### STAT proteins localise to LDs following virus infection

To date the only antiviral protein shown to localise to LDs in human cells is viperin (RSAD2), which is known to play roles in orchestrating a heightened antiviral environment (*2, 19, 20*). Our proteomic analysis of LDs in primary immortalised astrocyte cells revealed that multiple members of the early antiviral innate immune signalling pathways thought previously to be cytoplasmic, were present in the LD proteome, with expression of selected proteins being significantly upregulated following both pathway activation by dsRNA or ZIKV infection (Fig 3A, Table S2,3). Confocal imaging revealed the known LD localised protein, viperin (*21, 22*), to be highly abundant on the outside of LDs, with novel localised proteins; STAT1 and STAT2 being less abundant on LDs, and colocalising to only a subset of cytoplasmic LDs following activation of innate immune pathways by dsRNA (Fig 3B, S5). To better visualise and enumerate the populations of LDs these antiviral proteins are localised to, we isolated LDs from cells overexpressing proteins tagged with mCherry. Viperin was observed to localise to 54.8% of LDs on average, in comparison to STAT1 localising to 44.9% and STAT2 24.9%, which was also confirmed via immunoblotting (Fig 3C-E and S5A). Further confirmation of STAT1 localisation to the LD, and its phosphorylated forms (STAT1-Tyrosine701; ph-STAT1(T) and STAT1-Serine 727; ph-STAT1(S)) was performed using super resolution microscopy (single molecule localisation microscopy, SMLM), where we tracked STAT1 movement in the cell over a timeframe of 72 hpi (Fig 3F, G and S5B, C). There was a significant increase in colocalization events of LDs with STAT1 in all forms following stimulation (Fig 3F, G). However, when normalised to the increased cellular density of both LDs and STAT1 proteins, ph-STAT1 (T) demonstrated the most significantly enhanced interaction with the LD from as early as 8 hrs post activation of early innate antiviral signalling pathways (Fig 3G; S5B). This was further supported by a doubling of non-random co-localisation events compared to control conditions in which the degree of colocalization was equivalent to random levels of overlap (Fig S5B).

**Figure 3.**
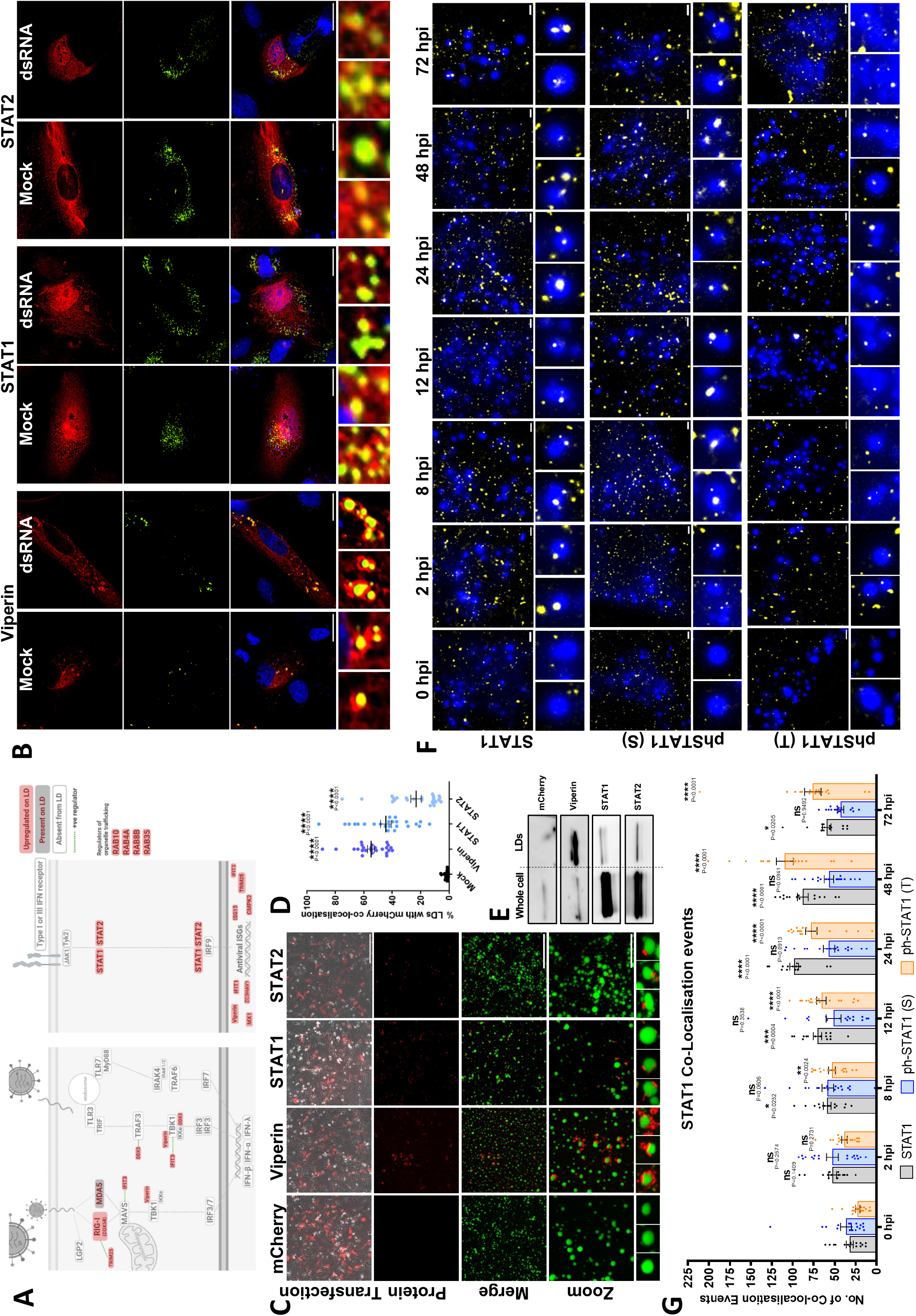
STAT proteins localise to LDs. **(A**) Schematic of the interferon signalling pathway following viral infection with proteins absent from the LD proteome in white, proteins present on the LD proteome in grey and proteins upregulated on the LD in red. **(B)** Astrocyte cells were transfected with a mCherry tagged viperin, STAT1 and STAT2 (red) and cells were stained with Bodipy (493/503) to visualise LDs (green) and DAPI to visualise cell nuclei (blue). The zoomed images indicate interaction between the respective mCherry tagged proteins and LDs. Scale bar, 50μm. **(C)** Confocal microscopy of astrocytes transfected with mCherry, viperin-mCherry, STAT1-mCherry and STAT2-mCherry respectively, with LDs then isolated and purified. Purified LDs were then imaged to determine the interaction between viperin, STAT1 and STAT2 with LDs. The zoom panel highlights the localisation of the aforementioned proteins with isolated and purified LDs. Scale bar, 400 μm for transfection panel, 50 μm for merge and 20 μm for zoom panels. **(D)** Isolated LDs were also imaged to allow the quantification of the percentage of LDs that have mCherry co-localisation with each data point representing the average percentage of co-localisation in a field of view of ∼500-3000 LDs (n=18-25) over 2 independent experiments. Error bars represent ± SEM. **(E)** To further validate the confocal microscopy of isolated LDs the purified LDs and the whole cell protein from the transfected astrocytes were immunoblotted for mCherry to show detection of mCherry tagged proteins **(F)** LD-STAT1 co-localisation events were visualised post dsRNA transfection via super resolution microscopy: Single-molecule localization microscopy (SMLM). The epifluorescence images of LDs were merged with the SMLM images of STAT1 (STAT1, phSTAT1 (S; serine)/ (T; tyrosine), and co-localisation events counted across a 72 hrs dsRNA stimulation time course. Cells were immunolabelled and imaged for LDs (Bodipy (493/503), epi, blue) and STAT1 (AlexaFluor647, SMLM, yellow). (n = 3 independent biological replicates with >150 cells imaged). Scale bars represent 1µm, with zoom inserts representing 1µm x 1µm in size. **(G)** LD-STAT1 co-localisation data was quantified as number of co-localisation events per cell. Captured images were merged and analysed using the ThunderSTORM plugin in ImageJ for co-localisation events. Error bars represent ± SEM, n= 6 cells over 3 independent assays.

### LDs form contacts with mitochondria to form signalosome complexes

LDs are dispersed throughout the cytoplasm of eukaryotic cells and move dynamically within cells to interact and communicate with other organelles, such as the endoplasmic reticulum, mitochondria, and peroxisomes, through the exchange of both lipids and proteins (*23*). There have been several hypotheses for why LDs may form contacts with other organelles, for example the delivery of LDs to lysosomes during autophagy to generate cholesterol (*24*), or the channelling of fatty acids liberated from lipolysis to sites of oxidation. Given the presence of both major viral RNA sensors; MDA5 at baseline, as well as upregulated RIG-I (DDX58) following both ZIKV infection and dsRNA stimulation we hypothesied that LDs may assist in signalosome formation to facilitate MAVS activation (Fig 3A, 1J, Supp Table 2 and 3). It is well established that the adaptor protein, MAVS localises to mitochondria (as well as to other organelles such as peroxisomes and the mitochondria-associated endoplasmic reticulum membrane (MAM) (*25*) and we wanted to examine its localisation with RIG-I and their potential to form a signalling complex at the LD. Confocal imaging analysis of astrocytes following activation of early innate signalling pathways with an RNA viral mimic demonstrated multiple instances of signalosome formation within individual cells, with RIG-I localised to LDs, forming complexes with MAVS localised to the mitochondria (Fig 4B, S6A). Live time imaging analysis of astrocytes also revealed that the frequency of LDs interacting with mitochondria increased significantly during both dsRNA and ZIKV infection, a phenomenon that did not occur when increasing LD numbers via oleic acid treatment of the cells (Fig 4C-G, S6B). This increase in interactions was driven by transient interactions of LD and mitochondria (Fig 4E, F), with some LDs interacting with mitochondria up to 15 times in a 10-minute timeframe, with an average of 6 interactions in virally infected cells compared to 1.8 in mock cells (Fig 4G).

**Figure 4:**
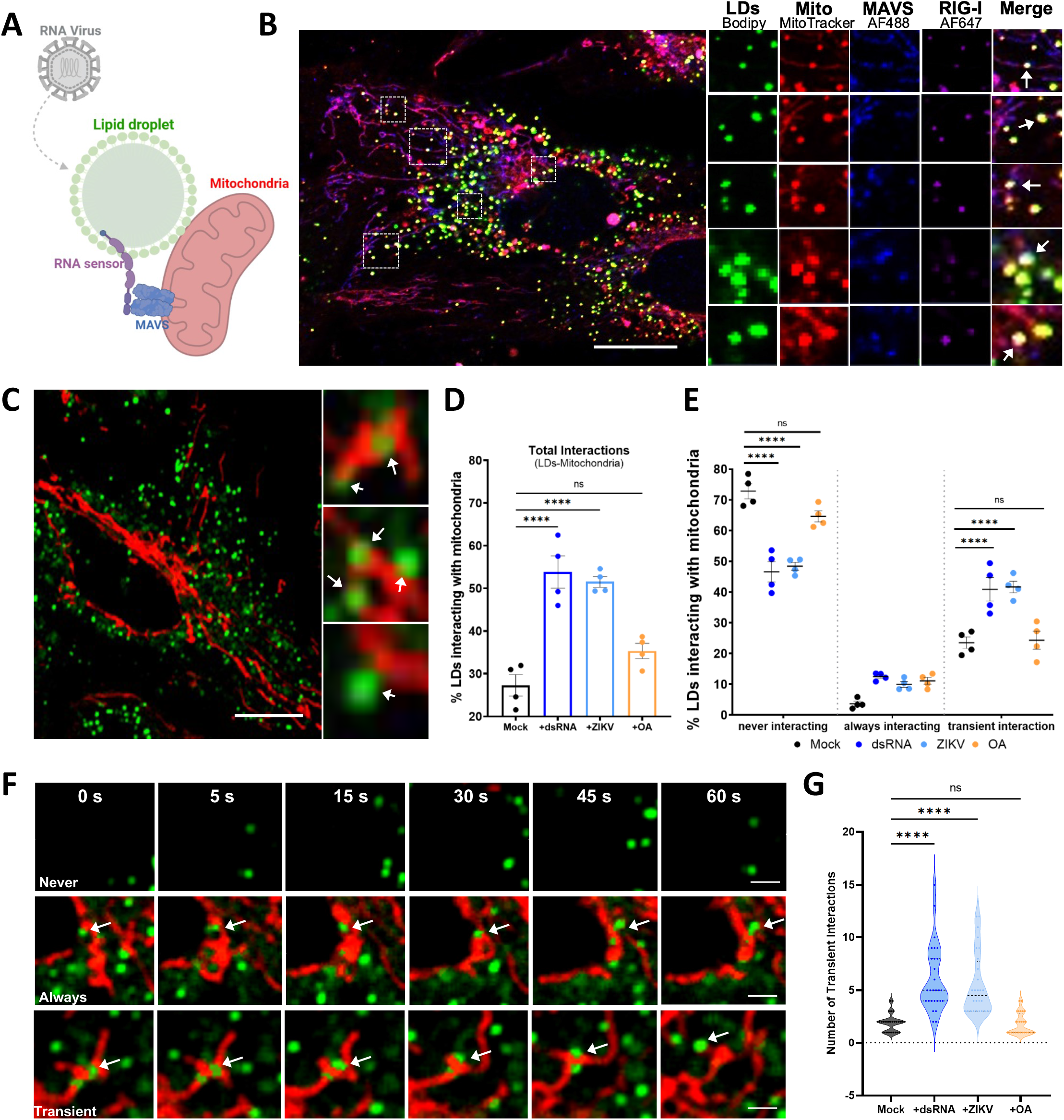
Lipid droplets form signalosomes with mitochondria. **(A)** Schematic depiction of the potential signalosome interaction between RIG-I localised to the LD surface and MAVS, localised to the mitochondria surface. **(B)** Primary immortalised astrocyte cells were transfected with a RIG-I over expression plasmid and stimulated with dsRNA for 24 hpi. Cells were live stained with MitoTracker Red prior to fixation and then all cells were stained with Autodot LD dye to visualise LDs (green), αRIG-I antibody (1:200) (purple) and αMAVS antibody (1:100) (blue). Scale bar, 50□μm. **(C)** Primary immortalised astrocyte cells were live stained with Bodipy (493/503) to visualise LDs (green) and MitoTracker Red to visualise mitochondria (red) and were live imaged on a Zeiss800 confocal over a 10 min timeframe. White arrows indicate interactions between the two organelles. Scale bar, 50□μm. **(D-G)** Cells were live stained, and infected with either ZIKV (MR766 strain) at an MOI1, stimulated with dsRNA viral mimic (Poly I:C) or treated with oleic acid (OA, 500 µM) for 8 hrs, and imaged for LD/mitochondria movement in live stimulated cells for 10 mins (120 frames). Images were analysed via Imaris image software for % interactions between LDs and mitochondria, and data was further analysed for numbers of transient interactions between the two organelles. n= 100 cells across 6 fields of view over 3 biological replicates, scale bar, 50 μm.

## Discussion

The role of LDs in viral infection has been dominantly studied from the point of view of viral pathogens usurping this organelle as a platform for assembly and manipulating its lipolysis and biogenesis to help enhance the viral life cycle (*4*). However, LD numbers are known to rapidly increase following viral infection, a cellular function that is required for optimal production of antiviral cytokines (*5*). Here we show that the LD proteome dynamically alters its makeup towards an enhanced profile of antiviral proteins and directly related antiviral signalling proteins, including the main RNA viral sensors, RIG-I and MDA5, as well as the main adaptor proteins in production of antiviral interferon stimulated genes, STAT1 and STAT2. Rapid proteomic changes to LDs following viral infection were underpinned by alterations in the lipidome of LDs towards a dominantly long-chain fatty acid profile in core lipids and small changes in the structural membrane lipids that likely facilitate a rapid proteome change (*26*). Organelle platforms are known to facilitate the formation of multiple signalosomes (*1, 10, 25*), and our analysis identified that at early time points following antiviral signalling pathway activation, LDs accumulated phosphorylated STATs, and interacted with mitochondria more frequently and for enhanced intervals, bringing together LD localised RIG-I and mitochondrial localised MAVS.

LD proteomes are understood to be dynamic in nature, but with the limited availability of LD proteome data, a better understanding of how these changes occurs is still to be established (*27, 28*). Very recently the LD proteome has been shown to be highly sensitive to bacterial LPS (*29*), however, the LPS changes driven in the LD proteome in this study have only a 17 protein overlap with those driven by viral RNA in our analysis, indicating that LD proteome changes are pathogen specific and likely to underpin specific functional capacity of the LD (Fig S7A,B). Overlapping proteome members were involved in organelle trafficking and lipid metabolism changes, indicating a commonality in the ability of the LD to traffic intracellularly and potentially mobilise altered lipid species as a core function of LDs in pathogen infection (Fig S7B). To our knowledge this is the first study to utilise a dual-omics approach to analyse the changing LD proteome simultaneously with its lipidomic changes in mammalian cells. This powerful tool was able to couple changes in metabolic enzymes in the LD proteome with changes seen in both the increases of lipids having long-chain fatty acyl chains, and the smaller alterations observed in the structural lipids of the LD membrane; it also highlighted the striking presence of multiple enzymes involved in post-translational modification of proteins, and potentially lipids within the LD proteome. Lipid alterations in virally driven LDs are likely to not only underpin the ability of the LD surface to alter its protein cargo but could also drive the production of bioactive lipid mediators such as the eicosanoid family, known to contribute to antipathogen immune defences (*4, 30*).

Signalosome formation is a requirement for efficient signalling intracellularly, to increase local concentrations of signalling components and promote weak interactions that may be required for enzyme activation. Signalosome formation often occurs at a cellular membrane platform such as the ER for STING, and mitochondria and peroxisome for MAVS (*25, 31, 32*); Alternatively, signalling molecules such as cGAS can induce self-organising centres to concentrate their reactions (*33*). Here we demonstrate that the LD can localise critical signalling proteins essential for the production of both interferon, the main antiviral cytokine, and interferon stimulated proteins, which underpin the successful antiviral cellular defences. Although some of these proteins remain constant at the LD surface (MDA5, STAT2), others accumulated transiently following viral infection (RIG-I, STAT1). Transient localisation of proteins to the LD surface is not well understood, however, post-translational modifications may underpin these events, and approximately 9% of our LD proteome following activation of early innate signalling pathways is composed of enzymes that facilitate these functions (as reviewed in (*34*)). Additionally, we were able to document small but significant changes to the external phospholipid membrane lipids towards a signature that can significantly enhance membrane curvature, protein binding capacity and cellular signalling at lipid membranes (*14*–*18*).

LDs are rapidly upregulated following viral infection and are critical in the early innate immune response facilitating a heightened antiviral environment (*5*). These studies highlight that viral driven LD upregulation coincides with dynamic changes to the LD lipidome and proteome to facilitate antiviral signalosome formation. Understanding the mechanism of LD facilitated antiviral defences may offer opportunities to tailor next generation antiviral therapeutics that may be more pan-antiviral in nature towards heightening the host cell antiviral environment.

## Methods

### Cells and culture conditions

Primary Immortalised Human Foetal Astrocytes were used throughout this study (referred to as primary immortalised astrocyte cells). These cells were maintained at 37□°C in a 5% CO_2_ air atmosphere in DMEM (Gibco, Cat; 12430054) containing 10% foetal bovine serum (FBS) (Gibco, Cat; 10099141), 100 units/mL penicillin and 100□μg/mL streptomycin (Sigma-Aldrich, cat; P0781).

### Lymphocytic Choriomeningitis Virus (LCMV) infection of mice

C57BL/6 mice were obtained from Australian BioResources (Moss Vale, NSW 2577) and housed under specific pathogen–free conditions in the animal facility at the University of Sydney, Sydney, Australia. Animal experiments were performed in accordance with the University of Sydney’s Animal Ethics Committee (1738/2020) and Institutional Biosafety Committee approval NLRD (22N004); mice were maintained under a 12 hr light/dark cycle at an ambient temperature of 20–23□°C and relative humidity of 40–60% and with ample food and water. All experiments were done in accordance with the Institutional Animal Care and Use Committee guidelines of the University of Sydney. All mice were aged between 8 and 16 weeks at the time of infection. Mice were anesthetized with 100 μg ketamine and 1 μg xylazine per gram bodyweight and intracranial infection was performed by injecting 500 PFU of LCMV (strain LCMV Armstrong 53b) diluted in 20 μL of phosphate-buffered saline (PBS) with 1% fetal bovine serum (FBS). Sham-infected mice were used as controls and received the same volume of PBS with 1% FBS but without virus. Mice were weighed at the times indicated below, and percent weight change was calculated. Mice were euthanized at 2- or 4-days post infection, and the brains were removed, flash frozen in optimum cutting temperature (OCT) medium.

### *In vitro* viral infection, viral mimics stimulation and plasmid transfection

Primary immortalised astrocyte cells were seeded at 5×10^6^ per T175cm^2^ flask plates prior to infection with Zika virus (MR766 strain) at an MOI of 1. Cells were washed once with PBS then infected with virus in serum free media for 4 hrs, followed by 20 hrs in DMEM supplemented with 10% foetal calf serum, at 37□°C containing 5% CO_2_. The viral mimic, poly I:C (dsRNA) (Invivogen) and all plasmid constructs used throughout the study were transfected into cells using PEI transfection reagent (Polyscience, Cat; 24765-1) at a concentration of 1□µg/ml. Viperin-mCherry and control mCherry plasmids were created as described previously (*35*). STAT1- and STAT2-mCherry tagged plasmids were kindly gifted to us from Associate Professor Greg Mosely (Monash University, Melbourne, VIC). RIG-I-wt was kindly gifted to us from Professor Stephen Polyak (University of Washington, Seattle, WA).

### Immunofluorescence microscopy

For cultured cells, briefly, cells were grown in 24-well plates on 12□mm glass coverslips coated with gelatine (0.2% [v/v]) were washed with PBS, fixed with 4% paraformaldehyde in PBS for 15□min at room temperature and permeabilised with 0.1% Triton X-100 in PBS for 10□min. Cells were blocked with 5% BSA for 1 hr, before antibody staining with αRIG-I (1:200; MA5-31715, Thermo Fisher Scientific), αMAVS (1:100; PA5-17256, Thermo Fisher Scientific). Cells were then incubated with Alexa Fluor 647 (1:200; A21236, Thermo Fisher) or Alexa Fluor 488 (1:200; A11008, Thermo Fisher Scientific) secondary antibody for 1□hr. Mitochondria were stained by incubating live cells with MitoTracker® Red (Thermo Fisher Scientific) at 100nM for 1 hr. LDs were stained by incubating cells with Bodipy (493/503) at 1□ng/mL for 1□hr and nuclei were stained with DAPI (Sigma-Aldrich, 1□µg/ml) for 5□min at room temperature. Samples were then washed with PBS and mounted with Vectashield Antifade Mounting Medium (Vector Laboratories). Preparation and staining of murine frozen brain sections were prepared following optimised protocols for tissue sections (*36*). Whole heads were sectioned sagittally. One section was snap-frozen and the other mounted immediately in OCT. Frozen sections were cut at 14□μM with a Leica CM 3050□S cryostat and mounted on microscope slides and stored at −80□°C. Sections were fixed with 4% paraformaldehyde in PBS for 15□min at room temperature. Sections were then washed with PBS, permeabilised with 0.1% Triton X-100 in PBS for 10□min, washed again and then blocked with 1% BSA for 30□mins. Sections were stained for specific cell types; astrocytes (αGFAP polyclonal antibody, 1:1000; PA3-16727, Invitrogen), neurons (αNeuN polyclonal antibody, 1:2500; PA5-78639, Invitrogen) and microglia (αTMEM119 monoclonal antibody, 1:10000; MA5-35043, Invitrogen). Sections were then washed and incubated with Alexa Fluor 555 secondary antibody at 1:200 for 1□hr. Bodipy (493/503) was used to stain for LDs at 1□ng/mL for 1□hr at room temperature, and nuclei were stained with DAPI for 5□min at room temperature. Images were then acquired using Zeiss 800 confocal microscope. Unless otherwise indicated images were processed using ImageJ analysis software.

### Lipid Droplet Isolation and Validation

#### Preparation of Lipid Droplet Fractions

Isolation of LD from cells and brain tissues was performed using a Lipid Droplet Isolation Kit (Cell Biolabs; Cat; MET-5011). For cells, 5x T-175cm^2^ flasks (5 × 10^6^ cells) of primary immortalised astrocyte cells were trypsinised, pelleted at 1000 g for 5 mins, washed 2 times with 1 x PBS. Mouse brain tissues were thawed on ice and 200mg of tissue surrounding the hippocampus was minced and put into sterile 1.5 mL microcentrifuge tube. Both cells and tissues were resuspended in 200 µl of reagent A (Cell Biolabs; Cat; MET-5011) and incubated on ice for 10 mins with occasional vortexing. 800 µl of 1 x reagent B (Cell Biolabs; Cat; MET-5011) was added to the cells/ tissues and further incubated on ice for 10 mins with occasional vortexing. Following incubation, cells/ tissues were carefully homogenised by being passed through a one inch 27-gauge needle attached to a 3 mL syringe five times. 600 µl of 1x reagent B was layered on top of the homogenates. Lysates were centrifuged for 3 hrs at 20,000 g at 4 °C. 100 µL of the top layer containing the floating LDs was taken per condition and stored at −80 °C for analysis of proteins and lipids.

#### Visualisation and analysis of LD-mCherry colocalization

Primary immortalised astrocytes were seeded at 5×10^6^ per T175cm^2^ flask prior to transfection with mCherry, viperin-mCherry, STAT1-mCherry and STAT2-mCherry. Cells were trypsinised and LDs were isolated. Following LD isolation, LDs were stained with Bodipy (493/503) at a concentration of 1□ng/mL. 10uls of purified LDs were spotted on a Nunc Lab-Tek II Chamber Slide System (Thermo Fisher Scientific) and were visualised via a Zeiss LSM 780 high-sensitivity laser scanning confocal microscope at 63x to determine both LDs stained with Bodipy (493/503) and mCherry expressing protein co-localisation. Image analysis was carried out using ImageJ, with LDs segmented using the Find Maxima function and a segmentation map was created. Segmentation maps were then used to separate interacting LDs and the Particle Analyser plugin was used to count LDs, create ROIs, and determine their sizes. To determine which LDs contained protein the LD area was isolated using an intensity threshold, and a binary image was created. The same method was used to find areas containing protein. The areas where both binaries overlapped was then determined via image calculator. This mask was then used in combination with the created ROIs to determine the presence or absence of protein in each LD.

#### Fluorescence and SMLM Image Acquisition and Processing

Switching buffer was applied to cells containing 80 µL 1M mercaptoethylamine (MEA), 20 µL 1 M potassium hydroxide (KOH) and 0.8 µL 1 mg/mL Bodipy (493/503) in PBS (pH 8.5) immediately prior to imaging. 8-well chamber slides were mounted on a custom SMLM setup based on (*37*). Briefly, the setup is built around an Olympus IX-83 inverted fluorescence microscope equipped with a 100X 1.49 oil immersion objective and a Photometrics Prime-95B sCMOS detector coupled to a pair of excitation lasers using appropriate dichroics and focal lenses (Semrock, Thorlabs). Diffraction-limited epifluorescence images of LDs were captured using 488 nm excitation at 8 mW (200 mW, Cobalt MLD), with 40 ms exposure. SMLM images were constructed by capturing 10,000 frames at 100 Hz with 200 mW 640 nm excitation (iBeam Smart, Toptica).

#### Lipid Droplet-STAT Protein Co-localisation Enumeration

For each time point, at least 6 fields of view were imaged at 100X magnification from varying locations across each well of the chamber slides. Images were imported to FIJI (ImageJ) and the LD channel converted to 16-bit before being scaled to 3000×3000, smoothed and binarized. The STAT channel was analysed using the ThunderSTORM plugin (*38*) for FIJI (*39*) to determine molecular coordinates from raw TIFF stacks and normalized gaussian renderings. These images were converted to 16-bit, smoothed and binarized before being overlayed with the LD channel to form the final merged LD-STAT images. To determine the number of colocalizations per cell, images were further analysed using the interaction factor analysis plugin (*40*) which is specifically designed to assess dense SMLM data and normalizes for coincidental co-localisation by generating Monte Carlo-based random renderings. This analysis was used to generate a ratio of real co-localisation to the number expected if only random interactions were present such that a ratio of 1 indicates entirely random overlap, where 2 indicates twice as many interactions as randomly modelled. These ratios are referred to throughout as ‘co-localisation factor’. Number of co-localisations for each image were determined using this object co-localisation analysis plugin of ImageJ.

### Western Blotting

Lysates were subjected to SDS-PAGE. The proteins were transferred to 0.2 µm nitrocellulose membranes (Bio-Strategy, Campbellfield, VIC, Australia) and probed with primary antibodies. The primary antibodies used were: mouse monoclonal αCalnexin (1:1000; sc-23954, Santa Cruz Biotechnology), mouse monoclonal αmfn1 (1:1000; sc-166644, Santa Cruz Biotechnology), mouse monoclonal αACOX1 (1:1000; sc-517306, Santa Cruz Biotechnology), rabbit monoclonal αPerilipin-2 (1:2000; ab108323, Abcam), mouse monoclonal αRIG-I (1:1000; sc-376845 Santa Cruz Biotechnology), rabbit monoclonal αMX1 (1:2000; ab207414, Abcam), rabbit polyclonal αSTAT1 (1:1000; 9172, Cell Signaling Technology), rabbit monoclonal αSTAT2 (1:1000; A3588, ABclonal), rabbit polyclonal αZC3HAV1 (1:5000; ab154680, Abcam), rabbit monoclonal αPhospho-STAT1 (Tyr701) (1:1000; #7649, Cell Signaling Technology), rabbit polyclonal αmCherry (1:1000; 5993, BioVision) and rabbit polyclonal αTRIM25 (1:1000; ab86365, Abcam). Following 3 × 5 min washes with TBS wash buffer, the membrane was incubated with HRP conjugated secondary antibodies (Goat αMouse IgG (H+L) Secondary Antibody, HRP, 31430, Thermo Fisher Scientific) and (Goat αRabbit IgG (H+L) Secondary Antibody, HRP, 31460, Thermo Fisher Scientific) for 1 hr diluted 1:10000. Following 5 × 10 min washes with TBS wash buffer, the membrane was incubated with GE (Amersham) or Femto (Thermo-scientific) Western Developer Reagent, dependent on the required sensitivity. The membranes were scanned using Amersham 600 chemiluminescence imager.

### Protein sample preparation for mass spectrometry

Proteins samples were precipitated from isolated LDs and whole cell via the S-trap micro protocol. Briefly 70 µl 2x lysis buffer (10% SDS in 50 mM TEAB) to 70 µl liquid sample [at a 1:1 ratio] so that final SDS is 5%. Samples were then sonicated for 5 mins to recover absorbed protein. Samples were then centrifuged for 8 mins at 13,000g. 500mM TCEP was added so that final concentration is 10 mM TCEP and incubate at 55 °C for 15 mins to reduce thiol groups. 500 mM IAA was added to reach final 50 mM IAA and incubate at RT for 30 mins to alkylate disulphides. 27.5% H_2_PO_4_ was then added so that the concentration is ∼2.5% phosphoric acid. Samples were vortexed and had pH checked to ensure acidity. 6X binding/wash buffer (100mM TEAB in 90% MeOH) was added to sample and mixed. Samples were centrifuged through the S-trap column at 4,000g for 30 secs to trap proteins. Protein was then cleaned by adding 150 µl of binding/washing buffer, and centrifuged 3 x at 4,000g for 30 secs discarding flowthrough. S-Trap column was then centrifuged at 4,000g for 1 min to fully remove binding/wash buffer. Protein was digested by adding 20 µl of digestion buffer (50mM TEAB + 1 µg Trypsin per 50 µg protein) and incubated at 37°C overnight. Proteins were eluted by adding 40 µl of 50 mM TEAB in water to the S-Trap and incubated for 30 mins. 40 µl of 0.2% formic acid in water was added to the S-Trap followed by centrifugation at 4,000g for 1 min. 40 µl of 50% acetonitrile (ACN) to the was added to the S-Trap and centrifuged at 4,000g for 1 min. Sample was placed in speedy vac to remove ACN and was followed by freeze drying the sample overnight.

### Lipid Extraction

Lipids were purified according to a modified protocol (*41*). Briefly, 10 µl of SPLASH Lipidomix (Avanti Polar Lipids) was spiked in each sample as internal standards. Lipids from the whole cell lysates and LD fractions were extracted by diluting lysates with methanol (with 0.01% BHT) so that the final concentration of the sample was 60% v/v MeOH containing 0.01% BHT. Lysates were further diluted with MeOH and ChCl_3_ so ratio of total H_2_O:CHCl_3_: MeOH was 0.74:1:2 and lysates were centrifuged at 14,000 x g for 15 mins to separate phases. Supernatants were collected and dried via speedvac centrifugation prior to analysis via LC-MS/MS.

### Quantitative proteomics and functional annotation analyses

Proteins were identified by mass spectrometry and relatively quantified by a liquid chromatography approach. Peptide samples were analysed by LC-MS/MS using an Ultimate 3000 UHPLC coupled to an Orbitrap Elite mass spectrometer (Thermo Fisher Scientific, San Jose, CA). Solvent A is 0.1% formic acid (FA) / 5% dimethyl sulfoxide (DMSO) in water and solvent B is 0.1% FA / 5% DMSO in acetonitrile (ACN). Each sample was injected onto a PepMap C18 trap column (75 μM X 2 cm, 3 μM, 100 Å, Thermo Fisher Scientific, San Jose, CA) at 5 μL/min for 6 min using 0.05% trifluoroacetic acid (TFA) / 3% ACN in water and then separated through a PepMap C18 analytical column (75 μM X 50 cm, 2 μM, 100 Å, Thermo Fisher Scientific, San Jose, CA) at a flow rate of 300 nL/min. The temperature of both columns was maintained at 50°C. During separation, the percentage of solvent B in mobile phase was increased from 3% to 23% in 89 min, from 23% to 40% in 10 min and from 40% to 80% in 5 min. Then the columns were cleaned at 80% solvent B for 5 min before decreasing the % B to 3% in 1 min and re-equilibrating for 8 min. The spray voltage, temperature of ion transfer tube and S-lens of the Orbitrap Elite mass spectrometer were set at 1.9 kV, 275 °C and 60% respectively. The full MS scans were acquired at m/z 300 – 1650, a resolving power of 120,000 at m/z 200, an auto gain control (AGC) target value of 1.0 × 10^6^ and a maximum injection time of 200 ms. The top 20 most abundant ions in the MS spectra were subjected to linear ion trap rapid collision induced dissociation (CID) at q value of 0.25, AGC target value of 5 × 10^3^, maximum injection time of 25 ms, isolation window of m/z 2 and NCE of 30%. Dynamic exclusion of 30 s was enabled. Data was searched by MaxQuant 1.4 against the UniProt homo sapien protein database. Trypsin was selected as enzyme. LFQ quantification and match between run were enabled and all other settings were default. The data sets (dsRNA, ZIKV, LCMV and respective controls) were imported into Perseus software (version 1.6.12.0). Label-free quantification (LFQ)/ or MS2 values were log2 transformed. Reverse database hits, potential contaminants, proteins only identified by site (a peptide carrying a modified residue), and proteins with 1 or less unique/razor peptides were removed from the matrices prior to *t* test statistical analysis. The permutation FDR corrected p-value less than 0.05 alongside a minimum of 3-5 measurements per group required for significance allowed identification of significantly upregulated or downregulated proteins following stimulation/infection of dsRNA, ZIKV or LCMV, and their respective controls. The transformed and filtered data was exported into the web-based software VolcanoseR, to visualise significantly changed proteins with a criterion of having a log^2^ fold change < -2 or > 2 and an FDR corrected P-value < 0.05. Hierarchical clustering was performed across all replicates and was visualised via the “ComplexHeatmap” package. Functional enrichment analysis was performed on significantly enriched proteins via the R package “Clusterprofiler” and visualised using “enrichplot” in the form of a tree-plot. The *t* test significant protein lists from each condition were visualised via their interactions using the STRING (Search Tool for the Retrieval of Interacting Genes/Proteins) database. The interaction data from STRING can be enhanced and adapted in cytoscape to produce enhanced network visualisation. Clustering of the network was performed using ClusterONE (Clustering with Overlapping Neighbourhood Expansion) with a p-value < 0.05 cut-off.

### Quantitative lipidomics

Samples were analysed by ultrahigh performance liquid chromatography (UHPLC) coupled to tandem mass spectrometry (MS/MS) employing a Vanquish UHPLC coupled to an Orbitrap Fusion Lumos mass spectrometer (Thermo Fisher Scientific, San Jose, CA, USA), with separate runs in positive and negative ion polarities. Solvent A was 6/4 (v/v) acetonitrile/water with 5 mM medronic acid and solvent B was 9/1 (v/v) isopropanol/acetonitrile. Both solvents A and B contained 10 mM ammonium acetate. 10 µl of each sample was injected into an Acquity UPLC HSS T3 C18 column (1 × 150 mm, 1.8 µm: Waters, Milford, MA, USA) at 50 °C at a flow rate of 60 μL/min for 3 min using 3% solvent B. During separation, the percentage of solvent B was increased from 3% to 70% in 5 min and from 70% to 99% in 16 min. Subsequently, the percentage of solvent B was maintained at 99% for 3 min. Finally, the percentage of solvent B was decreased to 3% in 0.1 min and maintained for 3.9 min. All MS experiments were performed using an electrospray ionization source. The spray voltages were 3.5 kV in positive ionisation-mode and 3.0 kV in negative ionisation-mode. In both polarities, the flow rates of sheath, auxiliary and sweep gases were 25 and 5 and 0 arbitrary unit(s), respectively. The ion transfer tube and vaporizer temperatures were maintained at 300 °C and 150 °C, respectively, and the ion funnel RF level was set at 50%. In the positive ionisation-mode from 3 to 24 min, top speed data-dependent scan with a cycle time of 1 s was used. Within each cycle, a full-scan MS-spectra were acquired firstly in the Orbitrap at a mass resolving power of 120,000 (at m/z 200) across an m/z range of 300–2000 using quadrupole isolation, an automatic gain control (AGC) target of 4e5 and a maximum injection time of 50 milliseconds, followed by higher-energy collisional dissociation (HCD)-MS/MS at a mass resolving power of 15,000 (at m/z 200), a normalised collision energy (NCE) of 27% at positive mode and 30% at negative mode, an m/z isolation window of 1, a maximum injection time of 35 milliseconds and an AGC target of 5e4. For the improved structural characterisation of glycerophosphocholine (PC) lipid cations, a data-dependent product ion (m/z 184.0733)-triggered collision-induced dissociation (CID)-MS/MS scan was performed in the cycle using a q-value of 0.25 and a NCE of 30%, with other settings being the same as that for HCD-MS/MS. For the improved structural characterisation of triacylglycerol (TG) lipid cations, the fatty acid + NH3 neutral loss product ions observed by HCD-MS/MS were used to trigger the acquisition of the top-3 data-dependent ion trap CID-MS3 scans in the cycle using a q-value of 0.25 and a NCE of 30%, with other settings being the same as that for HCD-MS/MS. Dynamic exclusion of 15 s was enabled and only ions with charge state of 1-3 were selected for fragmentation.

#### Lipid Identification and functional annotation analyses

LC-MS/MS data was searched through MS Dial 4.90. The mass accuracy settings are 0.005 Da and 0.025 Da for MS1 and MS2. The minimum peak height is 50000 and mass slice width is 0.05 Da. The identification score cut off is 80%. Post identification was done with a text file containing name and m/z of each standard in SPLASH® LIPIDOMIX® Mass Spec Standard (Cat. 330707, Avanti Polar Lipids, Birmingham, AL, USA). In positive mode, [M+H]+, [M+NH4]+ and [M+H-H2O]+ were selected as ion forms. In negative mode, [M-H]- and [M+CH3COO]-were selected as ion forms. All lipid classes available were selected for the search. PC, LPC, DG, TG, CE, SM were identified and quantified at positive mode while PE, LPE, PS, LPS, PG, LPG, PI, LPI, PA, LPA, Cer, CL were identified and quantified at negative mode. The retention time tolerance for alignment is 0.1 min. Lipids with maximum intensity less than 5-fold of average intensity in blank was removed. All other settings were default. All lipid LC-MS features were manually inspected and re-integrated when needed. These four types of lipids, 1) lipids with only sum composition except SM, 2) lipid identification due to peak tailing, 3) retention time outliner within each lipid class, 4) LPA and PA artifacts generated by in-source fragmentation of LPS and PS were also removed. The shorthand notation used for lipid classification and structural representation follows the nomenclature proposed previously (*42*). Quantification of lipid species in the unit of pmol from each sample was achieved by comparison of the LC peak areas of identified lipids against those of the corresponding internal lipid standards in the same lipid class and the quantity of each lipid standard at pmol. Since the lipid class of analyte and internal standard are identical but no co-ionization of analyte and IS are achieved, it’s categorized as level 3 quantification by Lipidomics Standards Initiative (lipidomicstandards.org/). Finally, the lipid species at the class, subclass or molecular species levels were normalized to either the total lipid concentration (i.e., mol% total lipid), or total lipid-class concentration (i.e., mol% total lipid class). For the lipid classes without correspondent stable isotope-labelled lipid standards, the LC peak areas of individual molecular species within these classes were normalised as follows: the MG species against the DG (18:1D7_15:0); the LPG against the PG (18:1D7_15:0), the LPA against the PA (18:1D7_15:0) and the LPS against the PS (18:1D7_15:0). All lipidomic data was imported to Excel for normalization and further processing. Significant differences between relative abundance of individual lipid species in two sample groups was acquired using unpaired two-tailed students *t* test (n= 3; biological replicates). Stack bar charts comparing relative abundance of major lipid categories and classes were made using Prism Version 8.4.3 software. Bubble plots of log2 fold changes in relative abundance of individual lipid species were made using R packages “ggplot2”. Relative quantification of fatty acids chain length among all 491 identified lipid species was analysed and plotted using R packages “Heatmap” and relative intensities were compared based on z-score. Principal component analysis (PCA) was performed using built in R packages (“stats”, “prcomp()”, “t”, “tibble”).

### Statistical Analysis and Reproducibility

Student’s *t* tests were used for statistical analysis between 2 groups, with experiments with 2 or more experimental groups statistically analysed using either an ordinary one, or two-way multiple comparison ANOVA or multiple *t* tests using the Holm-Sidak method for corrections for multiple comparisons with *P<* 0.05 considered to be significant. Omics data was analysed and plotted using R packages Version 4.3.0 as stated in relevant sections. All statistical analysis (unless otherwise indicated) was performed using Prism Version 8.4.3 software (GraphPad, La Jolla, United States). All experiments were performed in biological triplicate (unless otherwise stated), and technical duplicates were also performed for RT-PCRs. Error bars represent mean ± SEM, with a *P* value less than 0.05 considered to be significant. * *P*<0.05, ** *P*<0.01, *** *P*<0.001, **** *P*<0.0001.

## Funding

K.J.H and D.R.W are supported by a National Health and Medical Research Council (NHMRC) New Ideas Grant (APP1181434). E.A.M has funding from the CASS foundation and the Jack Brockhoff foundation. J.L.L currently holds an Australian Defence Scholarship.

## Acknowledgements

We would like to thank the La Trobe University Bioimaging Facility for their assistance with image acquisition, data collection and image analysis essential to this work. We also acknowledge the use of the Bio21 Mass Spectrometry and Proteomics Facility at the University of Melbourne and thank them for their continued support of our projects. We would like to thank Associate Professor Greg Mosely from Monash University (Melbourne, Australia) for gifting us the STAT1 and 2 mCherry constructs, Professor Stephen Polyak from the University of Washington (Seattle, WA) for gifting us the RIG-I-wt construct and Prof Paul Herzog from the Hudson Institute (Melbourne, Australia) for gifting us the MAVS antibody. All schematic diagrams throughout this manuscript were created using BioRender.

## Data availability

The data that support this study are available within the article and its Supplementary Information files are available from the authors upon request.

## Author Contributions

E.A.M. performed the majority of the experiments; with assistance from J.L.L, Z.T, M.L.S, A.J.M, I.A and A.R. Mice experiments were performed by M.H. Extraction of LDs from brain tissue was optimised by M.L.S with assistance from E.A.M, V.T and Q.D. E.A.M and S.N designed methods to extract lipids and proteins from LDs and optimised the preparation of samples. E.A.M and S.N assisted with proteomic and lipidomic analysis along with J.L.L, Z.T, K.H and A.M. Super resolution microscopy and analysis of STAT-LD co-localisations was performed by A.J.M with assistance from D.R.W and A.R. Image analysis of LD-mitochondria contacts was developed by C.J and E.A.M performed the analysis. E.A.M and K.J.H were responsible for the overall study design. E.A.M and K.J.H wrote the manuscript; all authors commented on the manuscript.

## Competing Interests

The authors declare no competing interests

## Supplementary figures

**Figure S1: The proteome of the brain changes following RNA virus infection**

(**A)** Study overview. Mice were sham injected or intracranially infected with LCMV (500 PFU) for 2 and 4 days. n= 6 replicates were taken from mice samples (SHAM 2dpi, LCMV 2dpi, SHAM 4dpi, LCMV 4dpi). (**B**) Monitoring changes in average body weight of mice models. Intracranially injected SHAM or LCMV mice were monitored for periods of 2- and 4-dpi and changes in body (%) were compared to day 0 (day of injection). (**C**) RT-qPCR analysis was used to quantify LCMV-NP, IFN-β and viperin in the brain tissues obtained post LCMV infection at 2 dpi and 4 dpi, data is expressed as the change in induction at 2 dpi relative to 4dpi. Error bars, mean values□±□SEM, P values determined by Student *t* test (n=□3 mice per group). **(D)** Representative images of LDs (Bodipy (493/503), green) following isolation from the brain. Images were taken on Nikon T*i*E fluorescence microscope; Scale bar represents 50 µm. **(E)** Heatmap displaying protein intensities of all identified proteins on LDs isolated from mice brain tissues from SHAM or LCMV infection at 2 dpi and 4 dpi. High abundance is shown in red, and low abundance is shown in blue.

**Figure S2: Lipid droplets are significantly upregulated in astrocyte cells in LCMV infected brains**.

Brain tissues were sectioned and stained for the major cell types; Astrocytes (αGFAP, 1:1000), Neurons (αNeuN, 1:2500) and Microglia (αTMEM119, 1:10000) all cell types are shown in red, LDs stained with Bodipy (493/503) (green) and DAPI to visualise the cell nuclei (blue). Tissues were imaged on a Zeiss 800 confocal microscope. Original magnification is 10X. Scale bar represents 300 µm, images representative of n=6.

**Figure S3: The LD proteome significantly changes in astrocyte cells following dsRNA stimulation**.

(**A)** Primary immortalised human astrocyte cells stimulated with dsRNA tagged with Rhodamine (red) for 8□and 24 hrs and stained with Bodipy (493/503) to visualise LDs (green) and DAPI to visualise the cell nuclei (blue). Cells were imaged on a a Zeiss 800 confocal microscope. Original magnification is 63X. Scale bar, 50□μm. (**B**) The purity of LDs isolated from primary immortalised astrocyte cells was assessed by immunoblot analysis. Equal amounts of protein from LD fractions and whole cell lysates were separated by SDS and blotted with indicated antibodies: perilipin 2 (ADRP) for LDs, MFN1 for mitochondria, Calnexin for ER and ACOX1 for peroxisome contaminants. (**C & D**) Volcano plot for differentially expressed genes when comparing dsRNA stimulated LDs for 8 hpi or 24 hpi with their respective controls. (n= 3 replicates per condition). **(E)** Hierarchical clustering of the 3870 identified LD proteins from astrocyte cells and their respective abundance in mock or dsRNA stimulation (24 hpi) conditions. Proteins are grouped according to gene ontology annotations (n= 3 replicates per condition). **(F)** Hierarchical clustering of the 325 identified Post-Translational Modification (PTM) regulatory enzymes in the LD proteome and their respective abundance in mock or dsRNA stimulation (24 hpi) conditions. Annotation reflects the PTM the protein regulates. (n= 3 replicates per condition).

**Figure S4: Changes in lipidome of LDs initiated early (8 hrs) following dsRNA stimulation in primary immortalised astrocytes**

(**A) &** (**B**) Principal component analysis (PCA) of lipid species identified in whole cell lysates and LD fractions isolated from primary immortalized astrocytes following dsRNA stimulation at 8 hrs and 24 hrs, respectively (n= 3, biological replicates). (**C**) Bubble plot of log2 fold changes in abundance of individual lipid species in dsRNA stimulated cells relative to mock at 8 hpi. Significance was determined by unpaired two-tailed students *t* test (n= 3). Individual lipid species are coloured by the class of lipid that they belong to. PC phosphatidylcholine; PE phosphatidylethanolamine; PI phosphatidylinositol; PS phosphatidylserine; LPC lysophosphatidylcholine; LPE lysophosphatidylethanolamine; DAG diacylglycerol; TAG triacylglycerol; Cer ceramide; SM sphingomyelin; CE cholesterol ester.

**(D)** Bubble plot of log2 fold changes in abundance of side chain fatty acids in isolated LDs following dsRNA stimulation 24 hpi. Fatty acid chain lengths were grouped based on carbon number of side chain fatty acids. **(E)** Comparison of relative intensity of fatty acids in major lipid classes identified in LD fraction isolated from dsRNA stimulated primary immortalized astrocytes; arranged from shortest to longest fatty acid chain lengths. **(F)** Distribution of side-chain fatty acids in isolated LDs from dsRNA stimulated primary immortalised astrocytes 24 hpi. Bar plot displaying distribution of fatty acids (number of carbons: number of double bonds, calculated based on presence on fatty acids) in LD fractions. Data is arranged from side chain fatty acids with highest to the lowest repeats.

**Figure S5: Lipid droplets house STAT proteins following innate immune activation of the cell by dsRNA**.

(**A)** Schematic diagram of experiments outlined in Fig 3C. Primary immortalised astrocyte cells were transfected with mCherry labelled proteins (viperin, STAT1 and STAT2) or mCherry control plasmid and their LDs were isolated. Isolated LD fractions were stained with Bodipy (493/503) to visualise isolated LD and were imaged on a Zeiss 780 confocal. (**B**) Quantification of the degree of randomness of STAT1 (STAT1, phSTAT1 (S; Serine 727, T; Tyrosine 701) co-localisation events per LD in primary immortalised astrocytes following dsRNA stimulation at 2, 8. 12. 24. 48 and 72 hpi. Images were analysed using the Object Co-localisation plugin in ImageJ. n= 18 cells over 3 biological replicates, error bars represent ± SEM. **(C)** Isolated LD fractions before and following dsRNA stimulation (24 hpi) were probed via immunoblot for the different activation states of STAT1 (STAT1 in all forms, ph-STAT1 (S; serine 727) and (T; tyrosine 701).

**Figure S6: Virally driven LDs form more contacts with mitochondria during infection**

(**A)** Primary immortalised astrocyte cells were transfected with a RIG-I over expression plasmid and stimulated with dsRNA for 24 hpi. Cells were live stained with MitoTracker Red prior to fixation and all cells were stained with Autodot LD dye to visualise LDs (green), αRIG-I antibody (1:200) (purple) and αMAVS antibody (1:100) (blue). Scale bar, 50□μm.

(**B)** Schematic of the analysis pipeline. Primary immortalised astrocytes were stained live with Bodipy (493/503) to visualise LDs, and Mito tracker Red to stain mitochondria prior to live imaging in cells. A total of 120 FoV images were captured at 500 ms exposure rates every 5 s (10 min in totally). Image stacks were aligned using linear stack alignment with SIFT plugin. Mitochondria channels were binarised in ImageJ, and the LD/Mito image stack was imported into Imaris image software. LDs were tracked using the “spots” function on Imaris and were coloured based on their intensity interactions with the mitochondria (red= interacting, purple= not interacting). Mean intensity (LDs to Mito), LD size and diameter were extracted and imported into Excel. Data was cleaned up by the exclusion of any data with large cell morphology shifts, >1 µm gaps/missing frames in LD paths and tracks which were < 10 frames. Interactions were analysed based on intensity (intensities over 100 were counted as interacting; with interactions lasting the duration of the movie counted as “always” interacting, intensities below 100 for the duration of the movie were counted as “never” interacting and those that had intensities over 100 at multiple stages of the movie were counted as “transient” interactions.

**Figure S7: Virally driven LDs house a specific proteome compared to LDs following bacterial infection**.

(**A)** LPS-induced LD proteome changes from (*29*) Vs dsRNA-induced LD proteome changes in this data set. (**B**) Of the 17 overlapped proteins, 6 were annotated to be involved in lipid metabolism (LDAH, ACSL4, LPCAT2, LPCAT1, FAF2, AUP1), 6 in organelle trafficking (RAB18, RAB21, RAB8A, RAB10, RAB1A, RAB35), 4 with diverse functions labelled as “other” (UBXN4, CYB5R3, RALA, HSD17B7) and only 1 was involved in immune signalling (RSAD2; viperin).

## Notes

### Competing Interest Statement

The authors have declared no competing interest.

