## Supplementary material for "Virally induced lipid droplets are a platform for innate immune signalling complexes": S Table 1

| Table S1: LD Proteins identified and their regulation in mice brains infected with LCMV | | | | |
|---|---|---|---|---|
| Protein Accessions | Gene Names | Protein Descriptions | LogFC (4dpi vs control) | LogFC (2dpi vs control) |
| **Proteins upregulated following LCMV Infection (2dpi)** | | | | |
| A0A0U1RNJ9 | Cd33 | Myeloid cell surface antigen CD33 (Fragment) | 0.629261748 | 4.419438998 |
| A0A0U1RPI1 | Fxyd1 | Phospholemman | 3.606335322 | 4.638598204 |
| A6H6A9 | Rabgap1l | Rab GTPase-activating protein 1-like | 0.586007643 | 2.076212247 |
| B1AWT7 | Ube2j1 | Ubiquitin-conjugating enzyme E2 J1 (Fragment) | -0.308402761 | 2.199696223 |
| B2RUR8 | Otud7b | OTU domain-containing protein 7B | 1.083388901 | 2.039957205 |
| D3YWQ0 | Dgki | Diacylglycerol kinase iota | 0.816776435 | 1.502887726 |
| D3Z3B8 | Dlg1 | Disks large homolog 1 | 0.309661341 | 2.218732198 |
| E9PZB2 | Rab11a | Ras-related protein Rab-11A (Fragment) | 1.135036453 | 2.578735828 |
| O35744 | Chil3 | Chitinase-like protein 3 | 0.391129589 | 2.373216867 |
| P05213 | Tuba1b | Tubulin alpha-1B chain | 0.229408169 | 2.876346906 |
| P81117 | Nucb2 | Nucleobindin-2 | 0.891626358 | 1.812079748 |
| Q3U9H3 | Bad | Bcl2-associated agonist of cell death | -0.805756156 | 1.476452669 |
| Q7TSG5 | Sh3d21 | SH3 domain-containing protein 21 | 4.953916995 | 7.14477396 |
| Q8K0C1 | Ipo13 | Importin-13 | -1.275793854 | 2.04897205 |
| Q99L04 | Dhrs1 | Dehydrogenase/reductase SDR family member 1 | -0.005300109 | 2.441308498 |
| **Proteins upregulated following LCMV Infection (4dpi)** | | | | |
| A0A087WSP5 | Stat1 | Signal transducer and activator of transcription | 1.962732442 | 0.577110767 |
| A0A0G2JGK8 | Rnf34 | E3 ubiquitin-protein ligase RNF34 (Fragment) | 3.154745436 | 2.038659414 |
| A0A0U1RPI1 | Fxyd1 | Phospholemman | 3.606335322 | 4.638598204 |
| A0A1Y7VLP0 | Atp5mpl | ATP synthase subunit ATP5MPL, mitochondrial (Fragment) | 3.186619457 | 1.789997021 |
| A2AMS3 | Galt | Galactose-1-phosphate uridylyltransferase | 4.634938876 | 1.785484314 |
| B0QZN5 | Vamp2 | Synaptobrevin-2 | 1.542623075 | -0.380085468 |
| O55022 | Pgrmc1 | Membrane-associated progesterone receptor component 1 | 0.98831803 | 1.608999093 |
| P62892 | Rpl39 | 60S ribosomal protein L39 | 2.189025847 | 4.81469202 |
| P97471 | Smad4 | Mothers against decapentaplegic homolog 4 | 1.489136728 | 1.324110587 |
| Q07797 | Lgals3bp | Galectin-3-binding protein | 2.887293911 | -1.094393412 |
| Q3T9E4 | Tgtp2 | T-cell-specific guanine nucleotide triphosphate-binding protein 2 | 2.512802982 | 1.308667501 |
| Q4VBF8 | Sipa1l1 | Signal-induced proliferation-associated 1-like protein 1 | 1.686963654 | 0.948061387 |
| Q64282 | Ifit1 | Interferon-induced protein with tetratricopeptide repeats 1 | 2.886704191 | 1.144900958 |
| Q64339 | Isg15 | Ubiquitin-like protein ISG15 | 1.496049245 | -0.536689281 |
| Q64345 | Ifit3 | Interferon-induced protein with tetratricopeptide repeats 3 | 3.204297447 | 1.341690779 |
| Q8C0L8 | Cog5 | Conserved oligomeric Golgi complex subunit 5 | 2.290706635 | 0.235586882 |
| **Proteins downregulated following LCMV Infection (4dpi)** | | | | |
| A0A1D5RL92 | Cherp | Calcium homeostasis endoplasmic reticulum protein | -4.082120482 | -0.757317543 |
| B9EJ80 | Pdzd8 | PDZ domain-containing protein 8 | -1.259453392 | -1.491836071 |
| **Proteins present/unchanged** | | | | |
| A0A023T778 | Magohb | Mago nashi protein | -0.399279467 | 0.697978338 |
| A0A067XG46 | Rpgr | X-linked retinitis pigmentosa GTPase regulator | 0.252903938 | 1.257616202 |
| A0A067XG53 | Cask | Peripheral plasma membrane protein CASK (Fragment) | 0.412899208 | 0.806010405 |
| A0A075B5K8 | Igkv1-99 | Immunoglobulin kappa variable 1-99 | -0.218547757 | -1.066729863 |
| A0A075B5P3 | Ighg2b | Immunoglobulin heavy constant gamma 2B (Fragment) | -0.24672273 | -0.115451813 |

| | | | | |
|---|---|---|---|---|
| A0A075B5P4 | Ighg1 | Ig gamma-1 chain C region secreted form (Fragment) | -0.374287637 | 0.562489986 |
| A0A075B5U4 | Ighv1-18 | Immunoglobulin heavy variable V1-18 | 0.201419846 | -1.127941291 |
| A0A075B6A3 | Igha | Immunoglobulin heavy constant alpha (Fragment) | -0.130847073 | -0.400971095 |
| A0A075B6D7 | Inpp5j | Inositol-polyphosphate 5-phosphatase | -0.982292334 | -0.326139609 |
| A0A076FR46 | Slc12a5 | KCC2a-S25 variant 1 | 0.57372036 | 0.773698807 |
| A0A087WNP6 | Cdv3 | Protein CDV3 | 0.225958729 | 0.48490874 |
| A0A087WNQ8 | Ogfrl1 | Opioid growth factor receptor-like protein 1 (Fragment) | -0.302777036 | 0.039024512 |
| A0A087WNT1 | Eloc | Elongin-C | -0.172437414 | -0.138698419 |
| A0A087WNU5 | Ank3 | Ankyrin-3 (Fragment) | 0.011158212 | 0.407452583 |
| A0A087WNU6 | Lrrfip1 | Leucine-rich repeat flightless-interacting protein 1 (Fragment) | -0.172695669 | 0.246498108 |
| A0A087WNV1 | Agfg1 | Arf-GAP domain and FG repeat-containing protein 1 | 0.321785323 | 0.398046652 |
| A0A087WNW3 | Ktn1 | Kinectin | -0.247770882 | -0.057782332 |
| A0A087WNW8 | Clasp1 | CLIP-associating protein 1 | -0.110112095 | -0.36299928 |
| A0A087WNZ9 | Rgs6 | Regulator of G-protein-signaling 6 | 0.036841202 | 0.183207512 |
| A0A087WP01 | Cdc34b | Cell division cycle 34B | 0.247512404 | -0.070758184 |
| A0A087WP80 | Lsamp | Limbic system-associated membrane protein | 0.330757046 | 0.739957015 |
| A0A087WPF0 | R3hdm1 | R3H domain-containing 1 (Fragment) | 0.265976016 | 0.492341359 |
| A0A087WPF8 | Abhd14b | Protein ABHD14B (Fragment) | 0.235820007 | 0.6337382 |
| A0A087WPH7 | Psmc3 | 26S proteasome regulatory subunit 6A (Fragment) | -0.254979547 | 1.426744302 |
| A0A087WPK3 | Lrrfip1 | Leucine-rich repeat flightless-interacting protein 1 | 0.685782941 | -0.275260448 |
| A0A087WPL5 | Dhx9 | DEAH box protein 9 | -0.062095674 | -0.187199593 |
| A0A087WPM2 | Ppfia4 | Protein tyrosine phosphatase, receptor type, f polypeptide (PTPRF),-interacting protein (liprin), alpha 4 | 1.333303432 | 2.124213219 |
| A0A087WPP8 | Abi2 | Abl interactor 2 | -0.133292135 | 0.344210148 |
| A0A087WQA0 | Bclaf1 | Bcl-2-associated transcription factor 1 | -0.375752703 | 0.915835539 |
| A0A087WQE8 | Kif1a | Kinesin-like protein KIF1A | -0.128772736 | -0.011798064 |
| A0A087WQS0 | Tns1 | Tensin 1 | 0.172861481 | 0.548649947 |
| A0A087WR20 | Ctsh | Pro-cathepsin H (Fragment) | -0.176285044 | 0.623088678 |
| A0A087WR50 | Fn1 | Fibronectin | -1.016707293 | 2.004517729 |
| A0A087WRE5 | Nudt16 | U8 snoRNA-decapping enzyme (Fragment) | -0.452610461 | 1.470948855 |
| A0A087WRE7 | Adhfe1 | Alcohol dehydrogenase iron-containing protein 1 (Fragment) | -0.198500093 | 1.23565038 |
| A0A087WRF2 | Agap1 | Arf-GAP with GTPase, ANK repeat and PH domain-containing protein 1 | -0.289013894 | 0.352169037 |
| A0A087WRG0 | Pfkfb2 | 6-phosphofructo-2-kinase | 0.364312522 | 0.346114 |
| A0A087WRH3 | Pcbp4 | Poly(rC)-binding protein 4 (Fragment) | 0.251422882 | 0.710804065 |
| A0A087WRY3 | Nucks1 | Nuclear ubiquitous casein and cyclin-dependent kinase substrate 1 | -0.489286709 | 0.172646205 |
| A0A087WS96 | Sh3bgrl2 | SH3 domain-binding glutamic acid-rich-like protein | -0.993918069 | -0.499370098 |
| A0A087WSM1 | Ica1l | Islet cell autoantigen 1-like protein | 1.014438725 | 0.879473209 |
| A0A087WSS1 | Relch | RAB11-binding protein RELCH | 0.766323853 | 0.300175985 |
| A0A087WST2 | Rbbp5 | Retinoblastoma-binding protein 5 | 0.069136429 | -0.296391169 |
| A0A0A0MQ68 | Gcdh | Glutaryl-CoA dehydrogenase, mitochondrial | -0.063674577 | -0.160012563 |
| A0A0A0MQ76 | Nop58 | Nucleolar protein 58 | -0.937366899 | -0.116714954 |
| A0A0A0MQ79 | Prrc2c | Protein PRRC2C | 0.226771927 | 0.336849213 |
| A0A0A0MQ90 | S100a13 | Protein S100-A13 | -1.513532607 | -0.224483967 |
| A0A0A0MQ97 | Nccrp1 | F-box only protein 50 | 0.306192112 | -1.41015927 |
| A0A0A0MQA3 | Serpina1a | Alpha-1-antitrypsin 1-1 | -0.001014932 | -0.204399427 |
| A0A0A0MQA5 | Tuba4a | Tubulin alpha chain (Fragment) | -0.236702919 | 0.422581355 |

| | | | | |
|---|---|---|---|---|
| A0A0A0MQC7 | Mapt | Microtubule-associated protein | -0.184980488 | 0.076634248 |
| A0A0A0MQE5 | Camsap1 | Calmodulin-regulated spectrin-associated protein 1 | 0.290532684 | 0.689745585 |
| A0A0A0MQE8 | Arhgap21 | Rho GTPase-activating protein 21 | 0.076136525 | 0.367480119 |
| A0A0A0MQF5 | Otud6b | Ubiquitinyl hydrolase 1 | -0.08080314 | -0.155087312 |
| A0A0A0MQG9 | Phyhd1 | Phytanoyl-CoA dioxygenase domain-containing protein 1 (Fragment) | -0.79817613 | -0.662985166 |
| A0A0A0MQM0 | Eif5a | Eukaryotic translation initiation factor 5A (Fragment) | 0.11637942 | 0.984079361 |
| A0A0A0MQN4 | Atg7 | Ubiquitin-like modifier-activating enzyme ATG7 | -0.26535759 | -0.151122729 |
| A0A0A6YVS6 | Syngap1 | Ras/Rap GTPase-activating protein SynGAP (Fragment) | -0.213917033 | 0.36410586 |
| A0A0A6YVU8 | Gm9774 | Predicted pseudogene 9774 | 0.07015775 | 0.162124793 |
| A0A0A6YVV4 | Map9 | Microtubule-associated protein 9 (Fragment) | 0.809472243 | 0.358170827 |
| A0A0A6YVV8 | Mbnl1 | Muscleblind-like protein 1 | -0.497520701 | 0.054853598 |
| A0A0A6YW06 | Enah | Protein enabled homolog | 0.059599272 | 0.696440061 |
| A0A0A6YW28 | Usp4 | Ubiquitin carboxyl-terminal hydrolase | -0.109519577 | 0.113963922 |
| A0A0A6YW36 | Rgs7 | Regulator of G-protein-signaling 7 | -0.025235335 | -0.800350825 |
| A0A0A6YW90 | Gria2 | Glutamate receptor | -0.087987995 | 0.399154186 |
| A0A0A6YWG7 | Rapgef2 | Cyclic nucleotide ras GEF | -0.050506147 | -0.080978711 |
| A0A0A6YWH2 | Aif1l | Allograft inflammatory factor 1-like | 0.491480446 | 2.116876205 |
| A0A0A6YWM5 | Rab3gap2 | Rab3 GTPase-activating protein non-catalytic subunit | 0.123027134 | 0.058285395 |
| A0A0A6YWM8 | Map4k4 | Mitogen-activated protein kinase kinase kinase kinase 4 | 0.02941672 | -0.039819876 |
| A0A0A6YWR6 | Zbtb18 | Zinc finger and BTB domain-containing protein 18 | -0.887672202 | 0.32284228 |
| A0A0A6YWX1 | Usp19 | Ubiquitin carboxyl-terminal hydrolase 19 | -0.215282122 | 0.538800716 |
| A0A0A6YWY8 | Pcdh9 | Protocadherin 9 | 0.133716265 | 0.300306797 |
| A0A0A6YX18 | Atp6v1h | V-type proton ATPase subunit H | -0.109573619 | 0.243670464 |
| A0A0A6YX26 | Rpl31 | 60S ribosomal protein L31 | 1.605349414 | 0.397088687 |
| A0A0A6YX56 | P4htm | Transmembrane prolyl 4-hydroxylase | 0.294295343 | 0.404798349 |
| A0A0A6YX71 | Dclk2 | Serine/threonine-protein kinase DCLK2 | -0.068907992 | 0.67270565 |
| A0A0A6YX73 | Prkar2a | cAMP-dependent protein kinase type II-alpha regulatory subunit | -0.020538902 | 0.483354886 |
| A0A0A6YXH3 | Osbpl9 | Oxysterol-binding protein | 0.234011555 | -0.139984449 |
| A0A0A6YXL6 | Lrba | Lipopolysaccharide-responsive and beige-like anchor protein | -0.074742699 | 0.001925786 |
| A0A0A6YXY1 | Ntmt1 | N-terminal Xaa-Pro-Lys N-methyltransferase 1 (Fragment) | 0.155838267 | -0.190240542 |
| A0A0A6YY35 | Rabgap1l | Rab GTPase-activating protein 1-like (Fragment) | -0.117428398 | 0.439267159 |
| A0A0A6YY67 | Camsap2 | Calmodulin-regulated spectrin-associated protein 2 | -0.218647957 | -0.06279103 |
| A0A0A6YY83 | Pcdh7 | Protocadherin 7 | -0.035046005 | 0.59113423 |
| A0A0A6YY91 | Ncam1 | Neural cell adhesion molecule 1 (Fragment) | 0.766114012 | 0.967232704 |
| A0A0B4J1E2 | Snw1 | SNW domain-containing protein 1 | 0.114885648 | -0.200311184 |
| A0A0D2X7Z2 | Eef1akmt1 | EEF1A lysine methyltransferase 1 | 0.054264641 | 0.328001499 |
| A0A0G2JDK0 | Mef2c | Myocyte-specific enhancer factor 2C (Fragment) | 0.743798892 | 1.515956879 |
| A0A0G2JDK2 | Gba | Glucosylceramidase | 1.312603951 | -0.1050512 |
| A0A0G2JDL2 | Celf3 | CUGBP Elav-like family member 3 | 1.342546558 | -1.046613852 |
| A0A0G2JDM6 | Ache | Carboxylic ester hydrolase | 0.532585716 | 0.00312829 |
| A0A0G2JDN0 | Ltf | Lactotransferrin | 1.041848787 | -1.441644986 |
| A0A0G2JDP0 | Hax1 | HCLS1-associated protein X-1 | 0.293709532 | 0.364200513 |
| A0A0G2JDV3 | Gbp6 | Guanylate-binding protein 6 | 0.934204865 | 2.020620863 |
| A0A0G2JDW2 | Myl3 | Myosin light chain 3 (Fragment) | 1.300574557 | 2.186175505 |
| A0A0G2JDW7 | Rps27 | 40S ribosomal protein S27 (Fragment) | 0.272797489 | 0.339296023 |

| | | | | |
|---|---|---|---|---|
| A0A0G2JDX4 | Tspan2 | Tetraspanin-2 (Fragment) | 0.623499902 | 1.344060421 |
| A0A0G2JDZ9 | Ankrd17 | Ankyrin repeat domain-containing protein 17 | 0.165129789 | 0.27520148 |
| A0A0G2JE00 | Magi2 | Membrane-associated guanylate kinase, WW and PDZ domain-containing protein 2 | 0.049046707 | 0.221478462 |
| A0A0G2JE35 | Arhgef9 | Collybistin | -0.462925561 | 1.901453654 |
| A0A0G2JE69 | Dph5 | Diphthine methyl ester synthase (Fragment) | 0.611192226 | 2.743178447 |
| A0A0G2JEC4 | Sh3glb1 | Endophilin-B1 | 0.078161462 | 0.395334244 |
| A0A0G2JED4 | Sh2b2 | SH2B adapter protein 2 | 1.538251432 | -0.404432535 |
| A0A0G2JEF3 | Selenof | Selenoprotein F (Fragment) | 0.513503234 | 0.47454373 |
| A0A0G2JEG8 | Amph | Amphiphysin | 0.140586281 | 0.064482053 |
| A0A0G2JEK2 | Crip1 | Cysteine-rich protein 1 | -0.188219515 | 0.559341431 |
| A0A0G2JEP0 | Fxr1 | Fragile X mental retardation syndrome-related protein 1 | -0.250522327 | 0.067033132 |
| A0A0G2JEP4 | Lrrfip2 | Leucine-rich repeat flightless-interacting protein 2 | 0.848807716 | 0.563265165 |
| A0A0G2JER9 | Dnajb6 | DnaJ homolog subfamily B member 6 | 1.279872576 | -0.055719376 |
| A0A0G2JES3 | Rpl9 | 60S ribosomal protein L9 (Fragment) | 0.706895828 | 3.585828145 |
| A0A0G2JEX1 | Nexn | Nexilin | -1.21823562 | 1.047833125 |
| A0A0G2JF67 | Pde5a | Phosphodiesterase | -0.664310741 | 1.914302031 |
| A0A0G2JFB0 | Rimbp2 | RIMS-binding protein 2 | 0.502890905 | 0.369078 |
| A0A0G2JFF1 | Ppp1cc | Serine/threonine-protein phosphatase (Fragment) | 0.115643915 | 0.187145551 |
| A0A0G2JFM9 | Fnbp1l | Formin-binding protein 1-like (Fragment) | 1.342770386 | 0.824930827 |
| A0A0G2JFT8 | Rufy3 | Protein RUFY3 | -0.203059546 | -0.368659973 |
| A0A0G2JFW6 | Rprd2 | Regulation of nuclear pre-mRNA domain-containing protein 2 (Fragment) | 1.07462031 | -0.536954244 |
| A0A0G2JFX0 | Dao | D-amino-acid oxidase (Fragment) | -0.700228691 | -1.083446662 |
| A0A0G2JFY5 | Fubp1 | Far upstream element-binding protein 1 | -0.064059798 | 0.389675935 |
| A0A0G2JFZ4 | Tsc22d2 | TSC22 domain family, member 2 | 0.004232979 | 0.12631019 |
| A0A0G2JG23 | Sez6l | Seizure 6-like protein | 0.073240471 | 0.426520824 |
| A0A0G2JGB7 | Lgi1 | Leucine-rich glioma-inactivated protein 1 | 0.825412846 | -0.060840448 |
| A0A0G2JGD2 | S100a4 | Protein S100-A4 (Fragment) | 0.786673864 | 1.584691525 |
| A0A0G2JGP4 | Nras | GTPase NRas (Fragment) | -0.206865819 | 0.665973663 |
| A0A0G2JGQ4 | Nub1 | NEDD8 ultimate buster 1 | 0.240894699 | 0.814849536 |
| A0A0G2JGX4 | Atp1a3 | Sodium/potassium-transporting ATPase subunit alpha | 0.388245138 | 0.590425173 |
| A0A0G2JGY9 | Plppr4 | 2-lysophosphatidate phosphatase PLPPR4 (Fragment) | 0.179473813 | 0.804774284 |
| A0A0H2UH22 | Cpeb2 | Cytoplasmic polyadenylation element-binding protein 2 | -0.2697251 | -0.339850744 |
| A0A0J9YTU3 | Specc1 | Cytospin-B | -0.945973778 | -0.075236162 |
| A0A0J9YTY0 | Septin11 | Septin | 0.249608167 | -0.077437719 |
| A0A0J9YU79 | Khk | Ketohexokinase | 0.127301216 | 0.029069821 |
| A0A0J9YUD5 | Nup205 | Nucleoporin 205 | 0.719761244 | 0.162629922 |
| A0A0J9YUN4 | Dnm1 | Dynamin GTPase | -0.173548063 | -0.043694814 |
| A0A0J9YUZ4 | Hmgb1 | High mobility group protein 1 (Fragment) | -0.304236221 | -0.046817621 |
| A0A0J9YVD2 | Ptprz1 | Receptor-type tyrosine-protein phosphatase zeta | 0.011312421 | 0.064927101 |
| A0A0M3HEP7 | Atp8a1 | Phospholipid-transporting ATPase | -0.525835387 | 0.12943697 |
| A0A0M3HEP9 | Txnrd2 | Thioredoxin reductase 2, mitochondrial | -0.148642063 | 0.302515666 |
| A0A0N4SUH8 | Nfu1 | NFU1 iron-sulfur cluster scaffold homolog, mitochondrial | -0.043440851 | 0.176780065 |
| A0A0N4SUV3 | Vamp1 | Vesicle-associated membrane protein 1 (Fragment) | 0.339401118 | 1.23592933 |
| A0A0N4SUZ0 | Magi1 | Membrane-associated guanylate kinase, WW and PDZ domain-containing protein 1 | 0.129950523 | 0.613678614 |
| A0A0N4SV10 | Etnk1 | Ethanolamine kinase 1 | -0.023089218 | 0.518803914 |

| | | | | |
|---|---|---|---|---|
| A0A0N4SV66 | H2aj | Histone H2A | -0.353565884 | 2.243923187 |
| A0A0N4SV80 | Zfp638 | Zinc finger protein 638 | 0.074125385 | 0.029159864 |
| A0A0N4SVA0 | Gm44596 | Predicted gene 44596 (Fragment) | -0.093698661 | 0.546304544 |
| A0A0N4SVB5 | Prrt3 | Proline-rich transmembrane protein 3 | 0.44083252 | 0.083357811 |
| A0A0N4SVC2 | Tra2a | Transformer-2 protein homolog alpha | 0.175877158 | 0.418307622 |
| A0A0N4SVL9 | Ppp6c | Serine/threonine-protein phosphatase | -0.031907082 | -0.369133631 |
| A0A0N4SVP8 | Eif4a3l2 | RNA helicase | 0.408362293 | 0.478581746 |
| A0A0N4SVQ1 | Ndufa4 | Cytochrome c oxidase subunit NDUFA4 | 0.482949257 | 1.198259513 |
| A0A0N4SVS6 | Cnbp | Cellular nucleic acid-binding protein | -0.375874011 | 0.553514322 |
| A0A0N4SVU2 | Hspa4l | Heat shock 70 kDa protein 4L (Fragment) | -0.366233238 | -0.511306683 |
| A0A0N4SW06 | Itfg2 | KICSTOR complex protein ITFG2 | 0.096791967 | 1.431134542 |
| A0A0N4SW73 | Rab11fip5 | Rab11 family-interacting protein 5 | 0.292791589 | 0.340116501 |
| A0A0N4SW93 | C2cd5 | C2 domain-containing protein 5 | -0.533672555 | -1.81529967 |
| A0A0N4SW94 | Myadm | Myeloid-associated differentiation marker (Fragment) | 0.572098764 | -0.186572393 |
| A0A0N4SW97 | Dcun1d2 | DCN1-like protein 2 | -0.139254061 | -0.270024459 |
| A0A0R3P9C8 | Ndufa9 | NADH dehydrogenase [ubiquinone] 1 alpha subcomplex subunit 9, mitochondrial | 0.724159336 | 0.268117269 |
| A0A0R3P9D0 | Plppr3 | Phospholipid phosphatase-related protein type 3 | 0.928214773 | 0.875529289 |
| A0A0R4IZW8 | Capns1 | Calcium-activated neutral proteinase small subunit | 0.072751522 | 0.658444246 |
| A0A0R4IZW9 | Sdf2 | Stromal cell-derived factor 2 | 0.070003064 | -1.384398778 |
| A0A0R4IZX5 | Ncan | Neurocan core protein | -0.107311376 | 0.162040393 |
| A0A0R4IZX8 | Pcp2 | Purkinje cell protein 2 | -1.501733144 | 1.020776272 |
| A0A0R4IZY0 | Thop1 | Thimet oligopeptidase | -0.159312884 | 0.119725068 |
| A0A0R4IZY9 | Dus3l | tRNA-dihydrouridine(47) synthase [NAD(P)(+)] | -0.558224456 | -2.259474595 |
| A0A0R4IZZ1 | Aplp1 | Amyloid-like protein 1 | 0.284129143 | 0.549317996 |
| A0A0R4J005 | Lims1 | LIM and senescent cell antigen-like-containing domain protein | -0.613995107 | 1.686850071 |
| A0A0R4J007 | Pald1 | Paladin | 0.223446608 | 0.530769507 |
| A0A0R4J008 | Hdac2 | Histone deacetylase 2 | -0.28524402 | -0.664449056 |
| A0A0R4J018 | Tpmt | Thiopurine S-methyltransferase | 0.125461801 | -0.39934508 |
| A0A0R4J023 | Auh | Methylglutaconyl-CoA hydratase, mitochondrial | 0.103395367 | 0.065388521 |
| A0A0R4J030 | Amer2 | APC membrane recruitment protein 2 | -0.04536012 | 0.28982989 |
| A0A0R4J034 | Pdxdc1 | Pyridoxal-dependent decarboxylase domain-containing protein 1 | -0.04427859 | 0.561196327 |
| A0A0R4J036 | Nefm | 160 kDa neurofilament protein | -0.121229617 | -0.182154338 |
| A0A0R4J038 | Kng1 | Bradykinin | -0.064362748 | 0.677614053 |
| A0A0R4J039 | Hrg | Histidine-rich glycoprotein | 0.223201338 | 1.683006128 |
| A0A0R4J041 | Mettl3 | N6-adenosine-methyltransferase subunit METTL3 | 0.177604771 | 2.192388058 |
| A0A0R4J047 | Luc7l | Putative RNA-binding protein Luc7-like 1 | -0.260381921 | 0.766726812 |
| A0A0R4J049 | Prmt5 | Protein arginine N-methyltransferase 5 | -0.045812798 | 1.202191671 |
| A0A0R4J050 | Acy1 | N-acyl-L-amino-acid amidohydrolase | -0.261273193 | 0.042262713 |
| A0A0R4J065 | Atg4b | Cysteine protease | -0.581203334 | -0.687096198 |
| A0A0R4J069 | Scly | Selenocysteine lyase | -0.194243336 | 0.355043729 |
| A0A0R4J078 | Ubxn4 | UBX domain-containing protein 4 | 0.177231153 | 1.07019043 |
| A0A0R4J079 | Acbd3 | Golgi resident protein GCP60 | -0.137116782 | 0.560741901 |
| A0A0R4J083 | Acadl | Long-chain specific acyl-CoA dehydrogenase, mitochondrial | -0.354424731 | 0.416700363 |
| A0A0R4J085 | Uap1 | UDP-N-acetylhexosamine pyrophosphorylase | -0.177417914 | 0.405960401 |
| A0A0R4J092 | Manba | Beta-mannosidase | -0.057615026 | 0.764948448 |

| | | | | |
|---|---|---|---|---|
| A0A0R4J094 | Fahd2a | Fumarylacetoacetate hydrolase domain-containing 2A | -0.173091348 | 0.199394703 |
| A0A0R4J099 | 2210016L21Rik | Protein CUSTOS | -0.187734032 | 0.61038359 |
| A0A0R4J0B1 | Mark2 | Non-specific serine/threonine protein kinase | 1.418452899 | 0.145557642 |
| A0A0R4J0B8 | Sh3gl3 | Endophilin-A3 | -0.008634885 | 0.549199422 |
| A0A0R4J0F6 | Gak | Cyclin-G-associated kinase | -0.068639215 | 0.513802528 |
| A0A0R4J0G0 | Pck2 | Phosphoenolpyruvate carboxykinase (GTP) | -0.141159503 | 0.808705648 |
| A0A0R4J0H0 | Eml4 | Echinoderm microtubule-associated protein-like 4 | 0.46833725 | -0.11864233 |
| A0A0R4J0I1 | Serpina3k | Serine protease inhibitor A3K | 0.179432487 | -0.039414406 |
| A0A0R4J0I9 | Lrp1 | Prolow-density lipoprotein receptor-related protein 1 | 0.080156612 | 0.510474205 |
| A0A0R4J0J0 | Palmd | Palmdelphin | 0.759060923 | 0.21517849 |
| A0A0R4J0J1 | Cpne9 | Copine-9 | 0.354483509 | 0.263121446 |
| A0A0R4J0K2 | Ckap5 | Cytoskeleton-associated protein 5 | -0.306914965 | -1.076149305 |
| A0A0R4J0L5 | Cog6 | Conserved oligomeric Golgi complex subunit 6 | -0.387890085 | -0.79448398 |
| A0A0R4J0M1 | Tbcc | Tubulin-folding cofactor C | -0.104402192 | 0.903127511 |
| A0A0R4J0M9 | Vcpip1 | Ubiquitinyl hydrolase 1 | 0.017945989 | 0.079065005 |
| A0A0R4J0N5 | Pdcd2 | Programmed cell death protein 2 | -0.207550748 | -0.338085254 |
| A0A0R4J0P1 | Acad8 | Isobutyryl-CoA dehydrogenase, mitochondrial | -0.29364783 | 0.402970791 |
| A0A0R4J0P6 | Parn | Poly(A)-specific ribonuclease PARN | -0.412233957 | 1.275817712 |
| A0A0R4J0Q2 | Inpp4a | Inositol polyphosphate-4-phosphatase type I A | -0.767276478 | 0.250760714 |
| A0A0R4J0Q5 | Lmnb2 | Lamin-B2 | -0.176859633 | 0.288709799 |
| A0A0R4J0Q9 | Cog7 | Component of oligomeric Golgi complex 7 | 0.196495183 | -0.591383298 |
| A0A0R4J0S1 | Cdc42ep1 | Cdc42 effector protein 1 | -0.525821527 | 0.12355725 |
| A0A0R4J0S3 | Rtn4ip1 | Reticulon-4-interacting protein 1, mitochondrial | -0.050922203 | 0.641299884 |
| A0A0R4J0S4 | Llgl1 | Lethal(2) giant larvae protein homolog 1 | -0.055091222 | 0.49071153 |
| A0A0R4J0T0 | Hscb | Iron-sulfur cluster co-chaperone protein HscB | -0.563947264 | 0.294226487 |
| A0A0R4J0T5 | Celf1 | CUGBP Elav-like family member 1 | 0.349815051 | -0.221694946 |
| A0A0R4J0T8 | Arfgap3 | ADP-ribosylation factor GTPase-activating protein 3 | -0.268349489 | -0.590050379 |
| A0A0R4J0T9 | Nyap2 | Neuronal tyrosine-phosphorylated phosphoinositide-3-kinase adapter 2 | -2.047251924 | 0.223974546 |
| A0A0R4J0U2 | Ppp4r2 | Serine/threonine-protein phosphatase 4 regulatory subunit 2 | -0.031244151 | 0.114225388 |
| A0A0R4J0U7 | Commd5 | COMM domain-containing protein 5 | -1.125710869 | -0.021818161 |
| A0A0R4J0V4 | Jmy | Junction-mediating and -regulatory protein | -0.098320198 | -1.6615809 |
| A0A0R4J0V5 | Polr2a | DNA-directed RNA polymerase subunit | -0.390635522 | 0.750858466 |
| A0A0R4J0W6 | Lrrc40 | Leucine-rich repeat-containing protein 40 | -0.147643407 | 0.130251249 |
| A0A0R4J0Z1 | Pdia4 | Protein disulfide-isomerase A4 | 0.181211503 | 0.377266884 |
| A0A0R4J0Z3 | Aqp4 | Aquaporin-4 | 1.346084785 | -0.077005068 |
| A0A0R4J107 | Apeh | Acyl-peptide hydrolase (Fragment) | -0.038908609 | 0.270156225 |
| A0A0R4J112 | Eif4g3 | Eukaryotic translation initiation factor 4 gamma 3 | -0.371094418 | 0.012763182 |
| A0A0R4J124 | Srpk2 | SRSF protein kinase 2 | 0.01460317 | -0.01541551 |
| A0A0R4J138 | Arsb | Arylsulfatase B | -0.011709181 | -0.195403099 |
| A0A0R4J140 | Cluh | Clustered mitochondria protein homolog | -0.279108493 | 0.056989034 |
| A0A0R4J194 | Cacnb1 | Calcium channel voltage-dependent subunit beta 1 | 2.227830219 | 0.141116619 |
| A0A0R4J195 | Rprd1b | Regulation of nuclear pre-mRNA domain-containing protein 1B | 0.894969336 | 1.605075479 |
| A0A0R4J1C5 | Atpaf2 | ATP synthase mitochondrial F1 complex assembly factor 2 | -0.019984738 | -0.851144314 |
| A0A0R4J1D0 | Cpne2 | Copine-2 | 0.723782444 | -0.530139844 |
| A0A0R4J1E3 | Dbn1 | Drebrin | -1.762441707 | -1.570336183 |

| | | | | |
|---|---|---|---|---|
| A0A0R4J1E8 | Tatdn1 | Putative deoxyribonuclease TATDN1 | 0.716219234 | 0.639386972 |
| A0A0R4J1F4 | Madd | MAP kinase-activating death domain protein | -0.253614362 | -0.076564153 |
| A0A0R4J1H6 | Golga3 | Golgin subfamily A member 3 | 0.261360391 | 0.166039944 |
| A0A0R4J1K1 | Cnot4 | CCR4-NOT transcription complex subunit 4 | 2.030350203 | 1.256304264 |
| A0A0R4J1N7 | Ank1 | Ankyrin-1 | -0.431394863 | 0.233255386 |
| A0A0R4J1N9 | Tfam | Transcription factor A, mitochondrial | 0.396853065 | 0.399344047 |
| A0A0R4J1P2 | Tpm3 | Tropomyosin alpha-3 chain | -1.212427298 | 0.526154836 |
| A0A0R4J1Q0 | Edc4 | Enhancer of mRNA-decapping protein 4 | 0.12969745 | 0.686668237 |
| A0A0R4J1R7 | Pcbd2 | 4a-hydroxytetrahydrobiopterin dehydratase | 2.281070805 | -0.24018844 |
| A0A0R4J1T1 | Sorbs2 | Sorbin and SH3 domain-containing protein 2 (Fragment) | -1.35740792 | -0.799355348 |
| A0A0R4J1T3 | Taok3 | Serine/threonine-protein kinase TAO3 (Fragment) | -0.215658156 | 0.671019713 |
| A0A0R4J1W7 | Cdc23 | Cell division cycle protein 23 homolog | 0.101385943 | 0.287578583 |
| A0A0R4J1Y3 | Osgep | Probable tRNA N6-adenosine threonylcarbamoyltransferase | -0.212768491 | 0.310908635 |
| A0A0R4J1Y6 | Sphkap | A-kinase anchor protein SPHKAP (Fragment) | -0.881016191 | 0.266247769 |
| A0A0R4J206 | Mocs2 | Molybdopterin synthase catalytic subunit | 1.3341657 | -1.065480868 |
| A0A0R4J215 | Gpatch11 | Coiled-coil domain-containing protein 75 | -1.30741415 | 0.445608139 |
| A0A0R4J254 | Xpo4 | Exportin-4 | -0.696536128 | 1.637423515 |
| A0A0R4J275 | Ndufa12 | NADH dehydrogenase [ubiquinone] 1 alpha subcomplex subunit 12 | 0.341732295 | 0.792796294 |
| A0A0R4J293 | Tgm1 | Protein-glutamine gamma-glutamyltransferase K | -0.077634462 | -0.452843984 |
| A0A0R4J2A6 | Anks1b | Ankyrin repeat and sterile alpha motif domain-containing protein 1B | 1.116694291 | -0.05683279 |
| A0A0R4J2B2 | Kctd12 | BTB/POZ domain-containing protein KCTD12 | -0.15115989 | -0.470854282 |
| A0A0R4J2C1 | Apbb1 | Amyloid-beta A4 precursor protein-binding family B member 1 | 0.196048737 | 0.260426839 |
| A0A0R4J2C2 | Syt2 | Synaptotagmin | -0.003095023 | 0.27061065 |
| A0A0R4J2D3 | Mia3 | Transport and Golgi organization protein 1 homolog (Fragment) | 0.823794429 | 1.196173032 |
| A0A0U1RNG8 | Kcmf1 | E3 ubiquitin-protein ligase KCMF1 | -0.289551703 | 0.77646176 |
| A0A0U1RPE4 | Gabra1 | Gamma-aminobutyric acid receptor subunit alpha-1 (Fragment) | -0.526673253 | 0.551427682 |
| A0A0U1RPL0 | Atxn2l | Ataxin-2-like protein | 0.474256166 | 0.159094493 |
| A0A0U1RPM0 | Apba2 | Amyloid-beta A4 precursor protein-binding family A member 2 | 0.014115683 | -0.069077492 |
| A0A0U1RQ20 | Pycard | Apoptosis-associated speck-like protein-containing a CARD (Fragment) | -0.068245252 | 0.118410269 |
| A0A0U1RQ27 | Mcee | Methylmalonyl-CoA epimerase, mitochondrial | -1.208110619 | -1.090526581 |
| A0A0U1RQ85 | Cbl | E3 ubiquitin-protein ligase CBL | 0.117162069 | 0.338274479 |
| A0A140LHA2 | Bub3 | Mitotic checkpoint protein BUB3 | 0.347136402 | 0.653261185 |
| A0A140LHB6 | Vstm2a | V-set and transmembrane domain-containing protein 2A (Fragment) | 0.170838261 | 0.79790465 |
| A0A140LHE2 | Smarca2 | Probable global transcription activator SNF2L2 (Fragment) | 0.68022836 | 0.307583809 |
| A0A140LIM2 | Zfand6 | AN1-type zinc finger protein 6 (Fragment) | 0.82376887 | -0.986974478 |
| A0A140LIW3 | Frmpd3 | FERM and PDZ domain-containing 3 | 0.241738002 | -0.658641497 |
| A0A140T8I9 | Pi4ka | Phosphatidylinositol 4-kinase alpha | 0.248304717 | 0.837256114 |
| A0A140T8J4 | Hebp1 | Heme-binding protein 1 | -0.084763718 | 0.006299814 |
| A0A140T8S0 | Megf10 | Multiple epidermal growth factor-like domains protein 10 (Fragment) | -0.930160427 | 0.740770976 |
| A0A171EBK8 | Rap1gap2 | Rap1 GTPase-activating protein 2 | -0.069770304 | 0.344141801 |
| A0A171KXD3 | Prmt1 | Protein arginine N-methyltransferase 1 | 0.120663293 | 1.204683304 |
| A0A1B0GQW6 | Rpl13a | 60S ribosomal protein L13a (Fragment) | -0.11808939 | 1.177076022 |
| A0A1B0GR11 | Taldo1 | Transaldolase | -0.202689234 | 0.291412989 |

| | | | | |
|---|---|---|---|---|
| A0A1B0GR85 | 2900026A02Rik | RIKEN cDNA 2900026A02 gene | 0.331648223 | -1.07004261 |
| A0A1B0GRJ9 | Pde2a | Phosphodiesterase | 0.029218165 | 0.160977681 |
| A0A1B0GRP7 | Plpbp | Pyridoxal phosphate homeostasis protein (Fragment) | -0.182043235 | 0.536360741 |
| A0A1B0GRR3 | Rps11 | 40S ribosomal protein S11 | 0.22641894 | -0.110356331 |
| A0A1B0GRU0 | Slc17a7 | Vesicular glutamate transporter 1 | 1.185874875 | -0.505761305 |
| A0A1B0GRU8 | Prpf40a | Pre-mRNA-processing factor 40 homolog A | 0.276918189 | 0.760723511 |
| A0A1B0GRV0 | Bpnt1 | 3'(2'),5'-bisphosphate nucleotidase 1 | -0.130119197 | 0.291034222 |
| A0A1B0GS08 | Stum | Protein stum homolog | 0.649059518 | 0.415476322 |
| A0A1B0GS13 | Bax | Apoptosis regulator BAX (Fragment) | 0.60745732 | 0.324103991 |
| A0A1B0GS41 | Gab1 | GRB2-associated-binding protein 1 | 0.123130735 | 0.736292044 |
| A0A1B0GS63 | Arfip2 | Arfaptin-2 | 0.116652711 | 0.431049824 |
| A0A1B0GSA5 | Ddhd2 | Phospholipase DDHD2 | 0.301286793 | 0.038252831 |
| A0A1B0GSR7 | Krt10 | Keratin, type I cytoskeletal 10 | 1.554887644 | 0.468542894 |
| A0A1B0GSX7 | Nup98 | Nuclear pore complex protein Nup96 | -0.636627229 | -0.032597542 |
| A0A1B0GT44 | Prmt9 | Protein arginine N-methyltransferase 9 | -1.101904249 | 1.005641699 |
| A0A1B0GT92 | Gys1 | Glycogen [starch] synthase | -0.186875693 | -0.085895856 |
| A0A1B0GX27 | Bcat2 | Branched-chain-amino-acid aminotransferase | -0.179178301 | -0.423910618 |
| A0A1C7CYU5 | Fam160a2 | FTS and Hook-interacting protein | 0.99852349 | 0.514133135 |
| A0A1C7CYV0 | Pls3 | Plastin-3 (Fragment) | 0.030390708 | 0.459962527 |
| A0A1C7ZMY3 | Shank2 | SH3 and multiple ankyrin repeat domains protein 2 | 0.145729129 | 0.236315568 |
| A0A1D5RL96 | Btbd8 | BTB/POZ domain-containing protein 8 | 0.407545058 | 0.571186701 |
| A0A1D5RLG0 | Hsdl1 | Inactive hydroxysteroid dehydrogenase-like protein 1 | -3.785180855 | -3.774385611 |
| A0A1D5RLG3 | Rab3gap1 | Rab3 GTPase-activating protein catalytic subunit | 0.0785278 | -0.043667952 |
| A0A1D5RLJ9 | Rtkn | Rhotekin (Fragment) | -0.303038947 | 0.421012243 |
| A0A1D5RLQ9 | Cdc42bpa | Non-specific serine/threonine protein kinase | 0.047018814 | 0.437726816 |
| A0A1D5RLV7 | Wdfy3 | WD repeat and FYVE domain-containing protein 3 | -0.429201285 | 0.054447492 |
| A0A1D5RLY2 | Vac14 | Protein VAC14 homolog | 0.162271945 | -0.365769386 |
| A0A1D5RM83 | Iqsec1 | IQ motif and SEC7 domain-containing protein 1 | 0.399982198 | -0.825315793 |
| A0A1D5RM85 | Rpl18a | 60S ribosomal protein L18a (Fragment) | -0.064865748 | -0.116894722 |
| A0A1D5RMG4 | Agap3 | Arf-GAP with GTPase, ANK repeat and PH domain-containing protein 3 | -0.137879976 | 1.162764549 |
| A0A1D5RMH8 | Ccdc158 | Coiled-coil domain-containing protein 158 (Fragment) | -0.738554605 | -0.866425355 |
| A0A1D5RMJ8 | Cnot1 | CCR4-NOT transcription complex subunit 1 | -0.000754897 | -0.126055876 |
| A0A1D5RMM8 | Pxn | Paxillin | 0.05368789 | 0.52906545 |
| A0A1L1SQ51 | Tln2 | Talin-2 | -0.085564359 | 0.12815237 |
| A0A1L1SQA8 | Rps25 | 40S ribosomal protein S25 | -0.372968324 | 1.344738801 |
| A0A1L1SQP9 | Tln2 | Talin-2 (Fragment) | -3.050969521 | 0.714813232 |
| A0A1L1SRH8 | Senp8 | Sentrin-specific protease 8 | 0.010280323 | 0.971808751 |
| A0A1L1SRJ9 | Commd4 | COMM domain-containing protein 4 (Fragment) | 0.866455396 | -0.503287156 |
| A0A1L1SRX2 | Ampd3 | AMP deaminase | 0.875169881 | 2.247891823 |
| A0A1L1SS23 | Rasgrf2 | Ras-specific guanine nucleotide-releasing factor 2 | 0.099864864 | -0.079574744 |
| A0A1L1SS44 | Aplp2 | Amyloid-like protein 2 (Fragment) | 0.08067948 | 0.567273458 |
| A0A1L1SSF2 | Adpgk | ADP-dependent glucokinase | -0.820430374 | -0.574351629 |
| A0A1L1SSH9 | Sparc | Osteonectin | -0.61137317 | 0.423311392 |
| A0A1L1SSS4 | Ncoa7 | Nuclear receptor coactivator 7 | -0.190537802 | -0.936212699 |
| A0A1L1SST5 | Spg21 | Maspardin | 0.445245139 | 0.980938752 |
| A0A1L1SSU2 | Dcun1d5 | DCN1-like protein (Fragment) | 0.569606177 | 0.631082455 |

| | | | | |
|---|---|---|---|---|
| A0A1L1ST53 | Tpd52l1 | Tumor protein D53 | 0.293279107 | -0.136026541 |
| A0A1L1STC5 | Gnb5 | Guanine nucleotide-binding protein subunit beta-5 | -0.62364219 | -0.987405618 |
| A0A1L1STE4 | Ilf3 | Interleukin enhancer-binding factor 3 | -0.193406264 | 0.098956108 |
| A0A1L1STF0 | Mindy2 | Ubiquitin carboxyl-terminal hydrolase | -0.007576656 | 0.68250529 |
| A0A1L1STF5 | Il18 | Interleukin-18 | -1.432932568 | 1.307332039 |
| A0A1L1SU77 | Siae | Sialate O-acetylesterase | 1.823915323 | 2.315786203 |
| A0A1L1SUM5 | Sergef | Secretion-regulating guanine nucleotide exchange factor | -0.543290742 | -0.809253375 |
| A0A1L1SUX8 | Thy1 | Thy-1 antigen (Fragment) | 0.669907538 | 1.101453781 |
| A0A1L1SV73 | Usp47 | Ubiquitin carboxyl-terminal hydrolase 47 | 0.040944672 | 0.068340619 |
| A0A1L1SVJ6 | Actn4 | Alpha-actinin-4 (Fragment) | 0.215311019 | 1.125598907 |
| A0A1L1SVK0 | Pafah1b2 | Platelet-activating factor acetylhydrolase IB subunit alpha2 (Fragment) | -0.189736748 | 0.362286091 |
| A0A1W2P6F6 | Myl6 | Myosin light polypeptide 6 | 0.098952103 | 0.65113306 |
| A0A1W2P6N7 | Smarcc2 | SWI/SNF complex subunit SMARCC2 | -1.593006738 | 0.253657579 |
| A0A1W2P6X3 | Fmnl1 | Formin-like protein 1 | -0.289961306 | -1.210434119 |
| A0A1W2P712 | Ralgapa1 | Ral GTPase-activating protein subunit alpha-1 | 0.474589507 | 2.095407009 |
| A0A1W2P768 | H3c14 | Histone H3.2 | -0.042046134 | 0.886113644 |
| A0A1W2P7A1 | Rps12 | 40S ribosomal protein S12 | -0.119946035 | 0.222383658 |
| A0A1W2P7A3 | Gm49918 | Predicted gene, 49918 | 1.642299589 | 0.83499829 |
| A0A1W2P7C1 | Ncoa1 | Nuclear receptor coactivator 1 (Fragment) | 1.064239025 | 1.395544608 |
| A0A1W2P7E1 | Asf1a | Histone chaperone ASF1A (Fragment) | -0.35369393 | -0.274246852 |
| A0A1W2P7H9 | Erh | Enhancer of rudimentary homolog | 0.909413147 | 1.257372856 |
| A0A1W2P7J9 | Gstt2 | Glutathione S-transferase theta-2 (Fragment) | -0.34668541 | 0.964372635 |
| A0A1W2P7K6 | Rab3ip | Rab-3A-interacting protein | -0.096019538 | 0.469173272 |
| A0A1W2P7Q6 | Xpot | Exportin-T | -0.206751188 | 0.575094541 |
| A0A1W2P7S5 | Trappc6b | Trafficking protein particle complex subunit 6B | -0.171194998 | -0.251948039 |
| A0A1W2P7V0 | Gopc | Golgi-associated PDZ and coiled-coil motif-containing protein (Fragment) | 0.072585201 | 0.53053538 |
| A0A1W2P7X0 | Abracl | ABRA C-terminal-like protein (Fragment) | -0.943293794 | -1.014265537 |
| A0A1W2P7Y9 | Nrcam | Neuronal cell adhesion molecule | -0.045634619 | 0.167431037 |
| A0A1W2P7Z1 | Pofut2 | Peptide-O-fucosyltransferase (Fragment) | 0.494699256 | 0.32902209 |
| A0A1W2P872 | Nova2 | NOVA alternative-splicing regulator 2 | -0.131707923 | 0.291708787 |
| A0A1Y7VIR0 | Ppp2r5c | Serine/threonine-protein phosphatase 2A 56 kDa regulatory subunit gamma isoform (Fragment) | 0.095122274 | 0.738939683 |
| A0A1Y7VJH3 | Ppp1r13b | Apoptosis-stimulating of p53 protein 1 (Fragment) | 0.499484762 | 0.901784738 |
| A0A1Y7VJX8 | Trim9 | E3 ubiquitin-protein ligase TRIM9 (Fragment) | 0.051665433 | -0.45371453 |
| A0A1Y7VJZ2 | Gstz1 | Maleylacetoacetate isomerase | -0.019395288 | 0.55119435 |
| A0A1Y7VK19 | Isca2 | Iron-sulfur cluster assembly 2 homolog, mitochondrial (Fragment) | -0.159851678 | 0.907809575 |
| A0A1Y7VK55 | Nfic | Nuclear factor 1 | 0.680540037 | -0.253773212 |
| A0A1Y7VK76 | Crppa | 2-C-methyl-D-erythritol 4-phosphate cytidylyltransferase-like protein | 0.384978008 | 0.761196931 |
| A0A1Y7VKT9 | Ubl5b | Ubiquitin-like protein 5 | 0.558692296 | 1.879656553 |
| A0A1Y7VL44 | Ttc7b | Tetratricopeptide repeat protein 7B | -0.563628642 | -0.426522573 |
| A0A1Y7VLY2 | Dock4 | Dedicator of cytokinesis protein 4 | -0.66896464 | -0.079534054 |
| A0A1Y7VM45 | Prpf39 | Pre-mRNA-processing factor 39 (Fragment) | -0.568178813 | 0.261556307 |
| A0A1Y7VMB1 | Trmt5 | tRNA (guanine(37)-N1)-methyltransferase (Fragment) | -0.043199857 | -0.609571616 |
| A0A1Y7VMC8 | Akr1c20 | Aldo-keto reductase family 1, member C20 (Fragment) | 0.584651089 | -0.746260325 |
| A0A1Y7VMN0 | Rock2 | Rho-associated protein kinase 2 (Fragment) | -0.110682551 | -0.08138148 |
| A0A217FL62 | Crhbp | Corticotropin-releasing factor-binding protein | -0.716483625 | 0.79258132 |

| | | | | |
|---|---|---|---|---|
| A0A217FL83 | Ccdc85c | Coiled-coil domain-containing protein 85C | -0.207225863 | -1.502905289 |
| A0A286YCG8 | Ssr1 | Signal sequence receptor subunit alpha | 0.868014654 | -0.825841586 |
| A0A286YCI0 | Ngly1 | N-glycanase 1 | -0.110232862 | -0.192371368 |
| A0A286YCJ4 | Spast | Spastin (Fragment) | -1.114478811 | -0.546057383 |
| A0A286YCL2 | Fcho2 | F-BAR domain only protein 2 (Fragment) | 0.731958707 | -1.124015331 |
| A0A286YCT6 | Epb41l3 | Band 4.1-like protein 3 (Fragment) | -0.885021464 | 0.510669549 |
| A0A286YCV9 | Kif13b | Kinesin family member 13B | 0.030071926 | 0.306076288 |
| A0A286YCW1 | Rad17 | Cell cycle checkpoint protein RAD17 | 1.291599512 | -2.742180347 |
| A0A286YCW8 | Camk2g | Calcium/calmodulin-dependent protein kinase type II subunit gamma (Fragment) | 0.996362686 | -1.142999967 |
| A0A286YDA2 | Nolc1 | Nucleolar and coiled-body phosphoprotein 1 | -0.397011948 | -0.806614081 |
| A0A286YDB3 | Tbc1d5 | TBC1 domain family member 5 | -0.123443826 | -0.447208087 |
| A0A286YDB6 | Synpr | Synaptoporin | -0.365396102 | -0.34311231 |
| A0A286YDB8 | Elp3 | Elongator complex protein 3 | 0.216473134 | -0.427563985 |
| A0A286YDC0 | Cdk7 | Cell division protein kinase 7 | -0.257435449 | 0.961126169 |
| A0A286YDH6 | Cadps | Calcium-dependent secretion activator 1 | -1.289963977 | -1.135382334 |
| A0A286YDI8 | Sec24c | Sec24-related gene family, member C (S. cerevisiae) | 0.040080134 | 0.340907097 |
| A0A286YDU2 | Atxn2 | Ataxin-2 (Fragment) | -0.129440022 | 0.915016651 |
| A0A286YDU3 | Ap3s1 | AP complex subunit sigma | 0.130272706 | -1.628833771 |
| A0A286YDY4 | Epb41l3 | Band 4.1-like protein 3 (Fragment) | 0.798857435 | -0.152874311 |
| A0A286YE31 | Ube2e1 | Ubiquitin-conjugating enzyme E2 E1 (Fragment) | -0.646554502 | 0.611675104 |
| A0A286YEB7 | Rps24 | 40S ribosomal protein S24 | 2.331934532 | 4.089190086 |
| A0A2C9F2A2 | Rin1 | Ras and Rab interactor 1 | 0.372831027 | -0.4094251 |
| A0A2C9F2D2 | Anxa7 | Annexin | -0.036933454 | 0.484616597 |
| A0A2I3BPC5 | Ppp3cc | Serine/threonine-protein phosphatase | -0.712299697 | 0.269411564 |
| A0A2I3BPG4 | Hlcs | Biotin--protein ligase | 0.003847535 | 0.141394456 |
| A0A2I3BPM7 | Samd4 | Sterile alpha motif domain-containing 4 | 1.363645363 | 2.600345214 |
| A0A2I3BPP1 | Tsc2 | Tuberin | 0.295001856 | 0.517891407 |
| A0A2I3BPT1 | App | Amyloid-beta A4 protein | 0.129074383 | 0.208628019 |
| A0A2I3BQ43 | Dgkh | Diacylglycerol kinase | 1.145620855 | -0.842889786 |
| A0A2I3BQN8 | Nudt18 | 8-oxo-dGDP phosphatase NUDT18 | 0.626186816 | 0.880333106 |
| A0A2I3BR11 | Bcl2l2 | Bcl-2-like protein 2 (Fragment) | -2.034122229 | -0.249231815 |
| A0A2I3BR29 | Fam107b | Protein FAM107B | 0.483436807 | 0.123046319 |
| A0A2I3BRD1 | Tbc1d23 | TBC1 domain family member 23 | -0.12751166 | -0.905998389 |
| A0A2I3BRL8 | Gm7324 | Predicted gene 7324 | 0.57862606 | 0.479046504 |
| A0A2I3BRT4 | Eppk1 | Epiplakin | 1.054972172 | -0.843844811 |
| A0A2I3BS14 | Cpq | Carboxypeptidase Q (Fragment) | 0.047802003 | 0.316666285 |
| A0A2K6EDK3 | Nudcd1 | NudC domain-containing protein 1 | 0.002977053 | 1.456998507 |
| A0A2R8VHH1 | Nup155 | Nuclear pore complex protein Nup155 | -2.347268232 | 2.70318831 |
| A0A2R8VHM0 | Kctd17 | Potassium channel tetramerisation domain-containing 17 (Fragment) | 0.410584831 | 1.179466963 |
| A0A2R8VHP3 | Gm5478 | Predicted pseudogene 5478 | -0.13753287 | 0.173230171 |
| A0A2R8VHV8 | Nell2 | Protein kinase C-binding protein NELL2 | 0.154283301 | -0.383341948 |
| A0A2R8VHX5 | Zfp385a | Zinc finger protein 385A | 0.769898357 | 1.238541782 |
| A0A2R8VI05 | Dnal4 | Dynein light chain | -0.092324384 | -0.645976067 |
| A0A2R8VI70 | Cyhr1 | Cysteine and histidine-rich protein 1 | 1.199558894 | -0.890597264 |
| A0A2R8VI79 | Rbfox1 | RNA binding protein fox-1 homolog 1 | 0.132932981 | -0.296902657 |

| | | | | |
|---|---|---|---|---|
| A0A2R8VJV3 | Copz1 | Coatomer subunit zeta | 0.20617129 | 0.583401203 |
| A0A2R8VJW0 | Aco2 | Aconitate hydratase, mitochondrial (Fragment) | -1.111684736 | -1.57807525 |
| A0A2R8VK70 | Cd47 | Integrin-associated protein | 0.923618126 | 0.98258241 |
| A0A2R8VKI7 | Tfcp2 | Alpha-globin transcription factor CP2 | 1.049488958 | -0.941081524 |
| A0A2R8W6E8 | Rcan2 | Calcipressin-2 (Fragment) | -0.59245259 | -0.792927742 |
| A0A2R8W6F8 | Mb | Myoglobin | -0.434094016 | 0.276383638 |
| A0A2R8W6V9 | Septin3 | Neuronal-specific septin-3 | 1.270085526 | 0.927614848 |
| A0A338P6E5 | Dlg4 | Disks large homolog 4 | 0.09831864 | 1.008413633 |
| A0A338P6F2 | Osbpl11 | Oxysterol-binding protein | 0.133486112 | 0.269319534 |
| A0A338P6F7 | Robo1 | Roundabout homolog 1 (Fragment) | -0.272422791 | -0.248588483 |
| A0A338P6I2 | Polr2h | DNA-directed RNA polymerases I, II, and III subunit RPABC3 | -0.597616927 | 0.075133642 |
| A0A338P6I7 | Ydjc | Carbohydrate deacetylase | -0.067116769 | 0.448776404 |
| A0A338P6J0 | Dlg2 | Disks large homolog 2 | 0.016360728 | 0.243879 |
| A0A338P6P6 | Acap2 | Arf-GAP with coiled-coil, ANK repeat and PH domain-containing protein 2 | 0.040865072 | 0.178428173 |
| A0A338P6P7 | Pacsin2 | Protein kinase C and casein kinase substrate in neurons protein 2 (Fragment) | 1.349741173 | 0.870787144 |
| A0A338P6R8 | Gm49601 | Septin | 0.44601059 | -0.926318169 |
| A0A338P6W3 | Ttc3 | RING-type E3 ubiquitin transferase | 0.973513254 | -0.215747197 |
| A0A338P7B7 | Dynlt1a | Dynein light chain Tctex-type 1A | -0.216559633 | 0.122500896 |
| A0A338P7E5 | Ube2l3 | Ubiquitin-conjugating enzyme E2 L3 | -0.05683438 | 0.530034065 |
| A0A384DV92 | Pde10a | Phosphodiesterase | 0.320027606 | -1.038953622 |
| A0A3B2W7H4 | Anks1 | Ankyrin repeat and SAM domain containing 1 | 0.649622409 | 0.085974852 |
| A0A3B2W7W2 | Ppard | Peroxisome proliferator-activated receptor delta | 1.256813304 | -1.036156178 |
| A0A3B2WB63 | Setd4 | SET domain-containing protein 4 (Fragment) | 0.939990489 | 0.251038869 |
| A0A3B2WBH9 | Tjp2 | Tight junction protein ZO-2 | -0.007216167 | 0.082759698 |
| A0A3B2WBL1 | Rpl10a | Ribosomal protein | 0.136619059 | 0.32434082 |
| A0A3B2WCD8 | Khsrp | Far upstream element-binding protein 2 | -0.02422212 | 0.25133276 |
| A0A3B2WCL5 | Ranbp3 | Ran-binding protein 3 | -0.083706983 | 0.440953573 |
| A0A3B2WCS4 | Ralbp1 | RalA-binding protein 1 | -0.513495286 | 0.347476323 |
| A0A3Q4EBU5 | Slc8a1 | Na(+)/Ca(2+)-exchange protein 1 | 0.468711662 | 0.51048549 |
| A0A3Q4EBV8 | Thoc1 | THO complex subunit 1 (Fragment) | -2.0024978 | -2.407884757 |
| A0A3Q4EG54 | Osbpl1a | Oxysterol-binding protein | -0.037331327 | 0.325626055 |
| A0A3Q4EGC9 | Ttc39c | Tetratricopeptide repeat protein 39C | -0.092677307 | 0.535298983 |
| A0A3Q4EH04 | Srsf7 | Serine/arginine-rich-splicing factor 7 | -0.116079998 | 0.09199063 |
| A0A3Q4EH84 | Cdo1 | Cysteine dioxygenase | 1.209431346 | 1.696821531 |
| A0A3Q4EHG2 | Commd10 | COMM domain-containing protein 10 | -0.555799421 | 1.529919227 |
| A0A3Q4EI12 | Rab18 | Ras-related protein Rab-18 | 0.410356267 | 0.645293077 |
| A0A3Q4L2S1 | Nubp2 | Cytosolic Fe-S cluster assembly factor NUBP2 | 0.754879125 | 1.255907694 |
| A0A494B8Y4 | Nrxn2 | Neurexin-2 | -0.545206006 | -0.057970206 |
| A0A494B923 | Papss2 | 3'-phosphoadenosine-5'-phosphosulfate synthase | 0.053338083 | 0.232125441 |
| A0A494B933 | Ppp1r14b | Protein phosphatase 1 regulatory subunit 14B (Fragment) | -2.505276632 | 1.049697558 |
| A0A494B953 | Nedd4l | HECT-type E3 ubiquitin transferase | -0.029699485 | -0.000802517 |
| A0A494B972 | Snx32 | Sorting nexin-32 | -0.62966706 | 0.474115213 |
| A0A494B9C3 | Vps13a | Vacuolar protein sorting-associated protein 13A (Fragment) | -0.027960873 | -0.458969911 |
| A0A494B9F4 | Ssh3 | Protein phosphatase Slingshot homolog 3 (Fragment) | 0.403980033 | 0.195830107 |
| A0A494B9F8 | Prelid3a | PRELI domain-containing protein 3A (Fragment) | 0.111482588 | 0.195164839 |

| | | | | |
|---|---|---|---|---|
| A0A494B9K4 | Add3 | Gamma-adducin | -0.951016553 | 5.455364863 |
| A0A494B9M7 | Trim36 | E3 ubiquitin-protein ligase Trim36 (Fragment) | -0.69433492 | 0.090321382 |
| A0A494B9Q5 | Cwf19l1 | CWF19-like protein 1 (Fragment) | -0.219980907 | 0.179566383 |
| A0A494B9R5 | Wdr4 | tRNA (guanine-N(7)-)-methyltransferase non-catalytic subunit WDR4 (Fragment) | -0.143873819 | 0.806221803 |
| A0A494B9V8 | Ptar1 | Protein prenyltransferase alpha subunit repeat-containing 1 | 0.671175162 | 1.185924768 |
| A0A494B9W4 | Rbm4b | RNA-binding protein 4B (Fragment) | -0.722940509 | 0.785491784 |
| A0A494B9X3 | Nt5c2 | Cytosolic purine 5'-nucleotidase | 0.008108584 | 0.373119831 |
| A0A494B9Z0 | Fau | 40S ribosomal protein S30 | 0.777359104 | 1.383694967 |
| A0A494BA07 | Pggt1b | Geranylgeranyl transferase type-1 subunit beta | 0.663158639 | 0.815979799 |
| A0A494BA44 | Rela | Transcription factor p65 | 0.104125849 | 0.50362285 |
| A0A494BA97 | Ostf1 | Osteoclast-stimulating factor 1 | 0.182400958 | 0.464647929 |
| A0A494BAB6 | Psmb8 | Proteasome subunit beta type-8 (Fragment) | -1.037332853 | 0.811374346 |
| A0A494BAI5 | Fech | Ferrochelatase | -0.236913331 | -0.225642999 |
| A0A494BAJ5 | Ccs | Superoxide dismutase [Cu-Zn] | 0.03541441 | 0.54240799 |
| A0A494BAJ6 | Arl3 | ADP-ribosylation factor-like protein 3 | -0.022437604 | 1.213310083 |
| A0A494BAT2 | Cuedc2 | CUE domain-containing protein 2 | 0.273274803 | 0.208295186 |
| A0A494BAW8 | Ndufs8 | NADH dehydrogenase [ubiquinone] iron-sulfur protein 8, mitochondrial (Fragment) | 2.562596989 | 1.015045166 |
| A0A494BB44 | Syt7 | Synaptotagmin-7 | 0.27067337 | -0.396242301 |
| A0A494BB75 | Wdr26 | WD repeat-containing protein 26 | 0.261314583 | -0.398409526 |
| A0A494BB95 | Eif1a | Eukaryotic translation initiation factor 4C | 0.287066714 | 0.361429532 |
| A0A494BBB0 | Capn1 | Calcium-activated neutral proteinase 1 (Fragment) | -0.105829398 | 0.506807804 |
| A0A494BBD3 | Scyl1 | N-terminal kinase-like protein (Fragment) | -0.756075446 | 0.188844681 |
| A0A498WFS2 | Ubxn1 | UBX domain-containing protein 1 | 0.012202835 | -0.084520499 |
| A0A498WGD8 | Txnl1 | Thioredoxin-like protein 1 | 0.014579964 | 0.234662374 |
| A0A498WGS3 | Mbp | Myelin basic protein | 0.126327896 | 0.296934764 |
| A0A4W9 | Negr1 | Neuronal growth regulator 1 | 0.397961203 | 0.677365144 |
| A0A571BD95 | Ube4b | Ubiquitin conjugation factor E4 B | -0.05138162 | -0.062662601 |
| A0A571BDG0 | Srcin1 | SRC kinase-signaling inhibitor 1 | 0.09483153 | 0.176485856 |
| A0A571BDP3 | Vcpkmt | Protein-lysine methyltransferase METTL21D | -0.846312364 | 0.142279148 |
| A0A571BDP7 | Rab3il1 | Guanine nucleotide exchange factor for Rab-3A | 0.805853907 | 0.497917811 |
| A0A571BE69 | Septin3 | Neuronal-specific septin-3 | 0.270205752 | 0.165736516 |
| A0A571BEC9 | Plin4 | Perilipin-4 | 1.468842951 | 0.259832859 |
| A0A571BEE9 | Iqcf3 | IQ motif-containing F2 | 0.204668427 | 0.480808894 |
| A0A571BEG7 | Patj | InaD-like protein | 0.003725402 | -0.396392186 |
| A0A571BEI3 | Gnas | Guanine nucleotide-binding protein G(s) subunit alpha isoforms short | 0.043936984 | -0.288488547 |
| A0A571BEL9 | Cct6a | T-complex protein 1 subunit zeta | 0.07538592 | -0.082473278 |
| A0A571BG24 | Limch1 | LIM and calponin homology domains-containing protein 1 | -0.012977537 | 0.718548139 |
| A0A571BGH0 | Kcnab2 | K(+) channel subunit beta-2 | 0.659386063 | -0.153963248 |
| A0A5F8MP75 | Mapk10 | Mitogen-activated protein kinase | -0.074258296 | 0.238583883 |
| A0A5F8MP96 | Septin4 | Septin-4 | -0.037040393 | 0.130995433 |
| A0A5F8MP98 | Tns3 | Tensin-3 | -0.020414702 | 0.453811646 |
| A0A5F8MPE1 | Epb41l3 | Band 4.1-like protein 3 | -0.300286802 | 0.563270569 |
| A0A5F8MPF0 | Kif1b | Kinesin-like protein KIF1B | -0.14514478 | -0.457024256 |
| A0A5F8MPI9 | Rapgef1 | Rap guanine nucleotide exchange factor (GEF) 1 | 0.335514832 | -0.097780387 |
| A0A5F8MPK9 | Sacm1l | Phosphatidylinositol-3-phosphatase SAC1 | -0.779443677 | -0.237669468 |

| | | | | |
|---|---|---|---|---|
| A0A5F8MPM1 | Shc3 | SHC-transforming protein 3 | -0.736398919 | -0.080246131 |
| A0A5F8MPN1 | Fry | Protein furry homolog | -0.055377563 | -0.041419824 |
| A0A5F8MPN4 | Mical3 | F-actin monooxygenase | 0.08186423 | 0.296858629 |
| A0A5F8MPP1 | Cracd | RIKEN cDNA C530008M17 gene | 0.716451899 | 0.274815877 |
| A0A5F8MPP4 | Usp25 | Ubiquitin carboxyl-terminal hydrolase 25 | -0.546410052 | 0.527373791 |
| A0A5F8MPR1 | Epb41l3 | Band 4.1-like protein 3 | -0.035259501 | 0.232803981 |
| A0A5F8MPS8 | Syt6 | Synaptotagmin-6 | 0.32832133 | -0.019660791 |
| A0A5F8MPT4 | Acap2 | Arf-GAP with coiled-coil, ANK repeat and PH domain-containing protein 2 (Fragment) | -0.50747606 | 0.157895247 |
| A0A5F8MPW1 | Agt | Angiotensin 1-10 | 0.034088612 | -0.095762491 |
| A0A5F8MPW9 | Peg3 | Paternally-expressed gene 3 protein | 0.036008898 | 0.535349528 |
| A0A5F8MPY3 | Ptk2 | Non-specific protein-tyrosine kinase | -0.093375556 | -0.311693033 |
| A0A5F8MPZ2 | Klc1 | Kinesin light chain | 0.090057087 | 0.333737532 |
| A0A5F8MQ05 | Bin2 | Bridging integrator 2 | 0.493156179 | 1.556426366 |
| A0A5F8MQ25 | Ppp6r2 | Serine/threonine-protein phosphatase 6 regulatory subunit 2 | 0.531122494 | 1.193391959 |
| A0A5H1ZRK8 | Igkc | Immunoglobulin kappa constant (Fragment) | -0.538733164 | -0.192242146 |
| A0A5H1ZRL3 | Nelfb | Negative elongation factor B | -0.171084531 | 0.708288829 |
| A0A5H1ZRL7 | Trp53bp1 | Transformation-related protein 53-binding protein 1 | -1.107916387 | 0.530875762 |
| A0A5H1ZRM8 | Scn2a | Sodium channel protein | -0.329204814 | 0.990187883 |
| A0A668KL36 | Ccdc50 | Coiled-coil domain-containing protein 50 | -0.078229109 | 1.534925302 |
| A0A668KL90 | Dgkz | Diacylglycerol kinase | 0.064947573 | -0.086197853 |
| A0A668KLA9 | Ppp2r2b | Serine/threonine-protein phosphatase 2A 55 kDa regulatory subunit B | 0.202634557 | -0.577982744 |
| A0A668KLC0 | Nectin1 | Nectin-1 | -0.155945746 | 2.286586006 |
| A0A668KLC6 | Map2 | Microtubule-associated protein | -0.105415789 | 0.166788419 |
| A0A668KLD3 | Akap12 | A-kinase anchor protein 12 | 0.005326176 | 0.236096382 |
| A0A668KLW3 | Csnk1e | Non-specific serine/threonine protein kinase (Fragment) | -0.936068344 | -0.817979455 |
| A0A668KM61 | Slc4a4 | Anion exchange protein | -0.259727637 | -0.105841001 |
| A0A6I8MWX8 | Dnajc7 | DnaJ homolog subfamily C member 7 | 0.023920918 | 0.427524567 |
| A0A6I8MWY5 | Golga2 | Golgin subfamily A member 2 | -0.017306487 | 0.364016215 |
| A0A6I8MWZ7 | Trim9 | E3 ubiquitin-protein ligase TRIM9 | 0.31881183 | -0.287693659 |
| A0A6I8MX08 | Ccdc88a | Girdin | -2.22622483 | -0.01368316 |
| A0A6I8MX12 | Rbms3 | RNA-binding motif, single-stranded-interacting protein 3 | 0.11835564 | 0.112122536 |
| A0A6I8MX15 | Tbc1d25 | TBC1 domain family member 25 | 1.277731244 | 0.634307384 |
| A0A6I8MX18 | Ago1 | Protein argonaute-1 | -0.10952479 | -0.90496095 |
| A0JNY3 | Gphn | Molybdopterin molybdenumtransferase | -0.016973146 | 0.020055612 |
| A0ZV96 | Oscp1 | Organic solute carrier protein 1 isoform | -0.293327936 | 0.110762596 |
| A1BN54 | Actn1 | Alpha actinin 1a | 0.077493604 | -0.214449247 |
| A1L3S7 | Gatad2b | Gatad2b protein | 0.624774933 | -1.894143422 |
| A1L3T7 | Ripor3 | RIPOR family member 3 | 0.551787821 | -0.270181338 |
| A2A432 | Cul4b | Cullin-4B | -0.13634723 | 0.180633227 |
| A2A4A6 | Plcg1 | 1-phosphatidylinositol 4,5-bisphosphate phosphodiesterase gamma | -0.004727077 | 0.261098226 |
| A2A4J8 | Vps25 | ESCRT-II complex subunit VPS25 | 0.479878553 | -0.000867526 |
| A2A513 | Krt10 | Keratin, type I cytoskeletal 10 | -0.071476237 | -0.457388878 |
| A2A547 | Rpl19 | Ribosomal protein L19 | 0.715569782 | 0.336678346 |
| A2A5R2 | Arfgef2 | Brefeldin A-inhibited guanine nucleotide-exchange protein 2 | -0.144917011 | -0.084076881 |
| A2A600 | Kpna2 | Importin subunit alpha-1 (Fragment) | -1.365704074 | 0.292519331 |

| | | | | |
|---|---|---|---|---|
| A2A690 | Tanc2 | Protein TANC2 | -0.484980869 | 1.43276612 |
| A2A699 | Fam171a2 | Protein FAM171A2 | 0.066480382 | 0.838665167 |
| A2A6H1 | Lasp1 | LIM and SH3 domain protein 1 (Fragment) | 0.925111898 | 1.302999814 |
| A2A6J4 | Lsp1 | Lymphocyte-specific protein 1 | 0.303824457 | 1.311939081 |
| A2A6M1 | Snf8 | Vacuolar-sorting protein SNF8 | -0.450152461 | -0.055325667 |
| A2A6Q8 | Myl4 | Myosin light chain 4 (Fragment) | -0.791927083 | 1.027144273 |
| A2A6T1 | Cdr2l | Cerebellar degeneration-related protein 2-like | 0.626521905 | 1.023847342 |
| A2A6U3 | Septin9 | Septin-9 | 0.46255792 | 0.805641651 |
| A2A7G9 | Fbxo6 | F-box only protein 6 (Fragment) | -0.788025602 | 0.489886284 |
| A2A7S7 | Yars | Tyrosine--tRNA ligase | 0.738619709 | 1.294579824 |
| A2A841 | Epb41 | Band 4.1 | -0.290997887 | 0.086646875 |
| A2A880 | Snx11 | Sorting nexin-11 (Fragment) | 0.408041573 | 0.233588696 |
| A2A8E2 | Czib | CXXC motif containing zinc binding protein | 0.259820239 | 0.72094361 |
| A2A8L5 | Ptprf | Receptor-type tyrosine-protein phosphatase F | -0.094760005 | 0.815007051 |
| A2A8R0 | Zfyve9 | Zinc finger FYVE domain-containing protein | -1.03018678 | 0.458866835 |
| A2A8V8 | Srrm1 | Serine/arginine repetitive matrix protein 1 | -0.854127884 | 0.097514868 |
| A2A9Q2 | Nrd1 | Nardilysin, N-arginine dibasic convertase, NRD convertase 1 | -0.002693717 | 0.28671217 |
| A2A9W7 | Gga3 | ADP-ribosylation factor-binding protein GGA3 | -0.044325701 | 0.548903147 |
| A2A9Z1 | Dmd | Dystrophin | -0.07986927 | 0.355388006 |
| A2AA71 | Sec24a | Protein transport protein Sec24A | -0.385884571 | 0.224940459 |
| A2AA85 | Szrd1 | SUZ domain-containing protein 1 (Fragment) | -0.439045016 | 0.466659387 |
| A2AAN0 | Exoc7 | Exocyst complex component 7 | -0.123917039 | 0.410077095 |
| A2ABY3 | Pcyt2 | Ethanolamine-phosphate cytidylyltransferase | -0.049990114 | 0.499038537 |
| A2ACG7 | Rpn2 | Dolichyl-diphosphooligosaccharide--protein glycosyltransferase subunit 2 | 0.998036194 | 1.254152457 |
| A2ACM0 | Rptor | Regulatory-associated protein of mTOR | -0.142672221 | -0.763456027 |
| A2AD03 | Chm | Rab proteins geranylgeranyltransferase component A | -0.153324095 | 0.389406522 |
| A2AD84 | Stk26 | Serine/threonine-protein kinase 26 | -0.831288179 | 1.868209839 |
| A2ADA6 | Acap3 | ArfGAP with coiled-coil, ankyrin repeat and PH domains 3 | -0.449972026 | 0.232860883 |
| A2ADE0 | Plekhm2 | Pleckstrin homology domain-containing family M member 2 | -0.831871001 | -0.685684363 |
| A2ADR8 | Ppp1r8 | Nuclear inhibitor of protein phosphatase 1 | -0.368952974 | 0.433221658 |
| A2ADY9 | Ddi2 | Protein DDI1 homolog 2 | 0.138183467 | 0.900878429 |
| A2AE27 | Ampd2 | AMP deaminase | -0.246444448 | -0.122730414 |
| A2AEC2 | Tceal3 | Transcription elongation factor A protein-like 3 (Fragment) | -0.34731528 | -0.247190952 |
| A2AEG6 | Gpm6b | Neuronal membrane glycoprotein M6-b | 0.563369497 | -0.25415198 |
| A2AEK1 | Cstf2 | AlphaCstF-64 variant 4 | 0.523144118 | 0.45196708 |
| A2AEK3 | Cstf2 | Cleavage stimulation factor subunit 2 (Fragment) | 0.093761826 | 0.367897987 |
| A2AEW8 | Gripap1 | GRIP1-associated protein 1 | 0.121966235 | 0.012678941 |
| A2AEX6 | Fhl1 | Four and a half LIM domains protein 1 | -0.223364131 | 0.502597809 |
| A2AF31 | Tmsb15b2 | Thymosin beta | -0.255992254 | 0.530865828 |
| A2AFG7 | L1cam | Neural cell adhesion molecule L1 | 1.153208001 | 0.421866417 |
| A2AFI8 | Reps2 | RalBP1-associated Eps domain-containing protein 2 | -0.300684452 | 0.613652229 |
| A2AFP5 | Rab9 | RAB9, member RAS oncogene family (Fragment) | 1.457502882 | 2.040010293 |
| A2AFQ0 | Huwe1 | HECT-type E3 ubiquitin transferase | -0.186397425 | -0.007352193 |
| A2AFQ2 | Hsd17b10 | 3-hydroxyacyl-CoA dehydrogenase type-2 | 0.053529898 | 0.653533618 |
| A2AG50 | Map7d2 | MAP7 domain-containing protein 2 | -0.12866691 | 0.534382979 |

| | | | | |
|---|---|---|---|---|
| A2AG83 | Psmd10 | 26S proteasome non-ATPase regulatory subunit 10 | -0.237086169 | 0.341403643 |
| A2AGI2 | Nlgn3 | Neuroligin-3 | 0.1405399 | 0.561839422 |
| A2AH25 | Arhgap1 | Rho GTPase-activating protein 1 | 0.238713805 | 0.380559127 |
| A2AH85 | Eftud2 | 116 kDa U5 small nuclear ribonucleoprotein component | -0.019237232 | 0.146274726 |
| A2AHM4 | Phactr3 | Phosphatase and actin regulator | 0.181861909 | -1.001020034 |
| A2AHW8 | Garnl3 | GTPase-activating Rap/Ran-GAP domain-like protein 3 | 0.049838607 | -0.413665215 |
| A2AHX9 | Bcl2l1 | Apoptosis regulator Bcl-X (Fragment) | 0.55358963 | 1.30852445 |
| A2AI52 | Usp8 | Ubiquitin carboxyl-terminal hydrolase 8 | -0.15963974 | 0.246562322 |
| A2AI87 | Phka1 | Phosphorylase b kinase regulatory subunit | 0.142987855 | -0.187144279 |
| A2AIG8 | Accs | 1-aminocyclopropane-1-carboxylate synthase-like protein 1 | 0.784345261 | 1.734897931 |
| A2AIH8 | Pir | Pirin (Fragment) | 0.509501457 | 0.230366707 |
| A2AIM4 | Tpm2 | Tropomyosin beta chain | -0.099159654 | 0.809217612 |
| A2AJ72 | Fubp3 | Far upstream element (FUSE)-binding protein 3 | -2.165618873 | -0.235084057 |
| A2AJ92 | Nsmf | NMDA receptor synaptonuclear-signaling and neuronal migration factor | 0.293296941 | 0.120418708 |
| A2AJH3 | Nmt2 | Glycylpeptide N-tetradecanoyltransferase | -0.422728093 | -1.500595411 |
| A2AJI0 | Map7d1 | MAP7 domain-containing protein 1 | 0.125728321 | 0.517615 |
| A2AJK8 | Ttc1 | Tetratricopeptide repeat protein 1 | 0.602186712 | 1.139582475 |
| A2AJP9 | Pdp1 | [Pyruvate dehydrogenase [acetyl-transferring]]-phosphatase 1, mitochondrial | -0.242440192 | 0.252227942 |
| A2AJW4 | Ppp1r3d | Protein phosphatase 1, regulatory subunit 3D | -0.170170116 | 0.939689954 |
| A2AKA9 | Brd3 | Bromodomain-containing protein 3 (Fragment) | -0.824901358 | -0.165122509 |
| A2AKD7 | Snta1 | Alpha-1-syntrophin | -0.195625814 | 0.425352414 |
| A2AKN8 | Mup8 | Major urinary protein 5 | 0.654021994 | 1.726814747 |
| A2AL12 | Hnrnpa3 | Heterogeneous nuclear ribonucleoprotein A3 | -0.014762338 | -0.384032408 |
| A2ALF0 | Dnajc8 | DnaJ homolog subfamily C member 8 | 0.01209027 | 0.766278108 |
| A2ALL9 | Atp2b3 | Calcium-transporting ATPase | 0.146243763 | 0.600821336 |
| A2ALS5 | Rap1gap | Rap1 GTPase-activating protein 1 | 0.025330575 | 0.52711757 |
| A2ALS7 | Rap1gap | Rap1 GTPase-activating protein 1 (Fragment) | 0.09320809 | 1.072034359 |
| A2ALU4 | Shroom2 | Protein Shroom2 | 0.095550696 | 0.320219835 |
| A2ALV3 | Sh3gl2 | Endophilin-A1 | 0.118831253 | 0.369029681 |
| A2ALV6 | Ecpas | Proteasome adapter and scaffold protein ECM29 | -0.126645184 | -0.307949861 |
| A2AMC3 | Pofut1 | GDP-fucose protein O-fucosyltransferase 1 | -0.454573965 | 0.623436451 |
| A2AMQ5 | Cds2 | Phosphatidate cytidylyltransferase | -0.306580098 | -0.425960223 |
| A2AMW0 | Capzb | F-actin-capping protein subunit beta | 0.470348962 | 0.339318117 |
| A2AMY5 | Ubap2 | Ubiquitin-associated protein 2 | 0.608851178 | -0.073849996 |
| A2AN08 | Ubr4 | E3 ubiquitin-protein ligase UBR4 | 0.085947259 | -0.09739399 |
| A2AN84 | Mpp1 | 55 kDa erythrocyte membrane protein | -0.339451567 | -0.358465195 |
| A2AP78 | Hmgb3 | High mobility group protein B3 (Fragment) | 1.307274723 | 2.643782298 |
| A2AP92 | Map3k7 | Mitogen-activated protein kinase kinase kinase 7 | 0.535824426 | -0.836916765 |
| A2APY7 | Ndufaf5 | Arginine-hydroxylase NDUFAF5, mitochondrial | -0.044134394 | -1.271446705 |
| A2AQ25 | Skt | Sickle tail protein | 0.328789361 | 0.042029222 |
| A2AQ45 | Fnbp1 | Formin-binding protein 1 | 0.084830729 | 0.427318573 |
| A2AQ87 | Shf | SH2 domain-containing adapter protein F (Fragment) | -0.371859392 | -0.207559745 |
| A2AQJ8 | Ganc | Neutral alpha-glucosidase C | 0.651177661 | 1.092983564 |
| A2AQN4 | Acss2 | Propionate--CoA ligase | -0.162496471 | 0.5337526 |
| A2AQU8 | Srxn1 | Sulfiredoxin | 1.172538408 | 0.62922891 |

| | | | | |
|---|---|---|---|---|
| A2ARP1 | Ppip5k1 | Inositol hexakisphosphate and diphosphoinositol-pentakisphosphate kinase 1 | -0.130147743 | 0.465597788 |
| A2ARP8 | Map1a | Microtubule-associated protein 1A | -0.044474792 | 0.176515261 |
| A2AS44 | Pkp4 | Plakophilin-4 (Fragment) | -0.04224577 | 0.868534247 |
| A2AS98 | Nckap1 | Nck-associated protein 1 | 0.087985865 | -0.179328124 |
| A2ASQ1 | Agrn | Agrin | -0.347237841 | 0.171952724 |
| A2ASW8 | Rapgef4 | Rap guanine nucleotide exchange factor 4 | -0.365773678 | -0.840049108 |
| A2AT91 | Plcb4 | Phosphoinositide phospholipase C (Fragment) | -0.257083289 | 0.202821573 |
| A2ATI9 | Gorasp2 | Golgi reassembly-stacking protein 2 | -0.011706034 | 0.41761144 |
| A2ATQ5 | Anapc1 | Anaphase-promoting complex subunit 1 | 0.081490962 | 0.705202659 |
| A2ATU9 | Hat1 | Histone acetyltransferase type B catalytic subunit | -0.348839283 | 0.740221262 |
| A2AU61 | Raly | RNA-binding protein Raly (Fragment) | 1.321760305 | -0.377600034 |
| A2AUE1 | Dnajc5 | DnaJ homolog subfamily C member 5 (Fragment) | -0.290173737 | 0.990932306 |
| A2AUK8 | Epb41l1 | Band 4.1-like protein 1 | 0.220271428 | -0.833435376 |
| A2AUR3 | Pter | Parathion hydrolase-related protein (Fragment) | -0.198889764 | 0.733472824 |
| A2AUR7 | Rsu1 | Ras suppressor protein 1 | 0.063486576 | 0.357803663 |
| A2AUX3 | Dab2ip | Disabled homolog 2-interacting protein | 0.133474 | 0.336397171 |
| A2AW05 | Ssrp1 | FACT complex subunit SSRP1 (Fragment) | 0.256374296 | 0.68660752 |
| A2AWA9 | Rabgap1 | Rab GTPase-activating protein 1 | -0.195187473 | 0.225073973 |
| A2AWI7 | Sh3glb2 | Endophilin-B2 | -0.179899025 | 0.650087674 |
| A2AWN8 | Ythdf1 | YTH domain-containing family protein 1 | -0.249823284 | -0.73157986 |
| A2BDX3 | Mocs3 | Adenylyltransferase and sulfurtransferase MOCS3 | -0.41638333 | 0.556907495 |
| A2BE93 | Set | Protein SET (Fragment) | 0.097362041 | 0.31124417 |
| A2BFF5 | Dync1i2 | Cytoplasmic dynein 1 intermediate chain 2 | 0.304466311 | 0.416698138 |
| A2BGI8 | Ppih | Peptidyl-prolyl cis-trans isomerase (Fragment) | 0.198440297 | 0.117498716 |
| A2BI12 | Psip1 | PC4 and SFRS1-interacting protein | -0.302034632 | 0.872873783 |
| A2RSH4 | Sv2c | Sv2c protein | -0.08223327 | 2.450417678 |
| A2RSJ4 | Uhrf1bp1l | UHRF1-binding protein 1-like | -1.813150863 | 0.550001303 |
| A2RSX7 | Tyw5 | tRNA wybutosine-synthesizing protein 5 | 0.023576832 | 0.632734934 |
| A2RSX9 | Arfip1 | ADP-ribosylation factor interacting protein 1 | 0.391689396 | 0.546777566 |
| A2RT62 | Fbxl16 | F-box/LRR-repeat protein 16 | -0.260214424 | 0.144642353 |
| A2RTH5 | Lcmt1 | Leucine carboxyl methyltransferase 1 | -0.003992685 | 0.649196307 |
| A2TJV2 | Palm3 | Paralemmin-3 | -0.506222534 | 0.892276446 |
| A3KFU5 | Pabpc4 | Polyadenylate-binding protein | 0.14306523 | 0.431160132 |
| A3KFX0 | Nt5c1a | Cytosolic 5'-nucleotidase 1A | -0.762782542 | 0.035535177 |
| A3KG44 | Ikbkg | NF-kappa-B essential modulator (Fragment) | -1.539408016 | -1.114723523 |
| A3KGB4 | Tbc1d8b | TBC1 domain family member 8B | 0.416160583 | -0.136718273 |
| A3KGE4 | Olfm1 | Noelin | 0.519220098 | 0.240429242 |
| A3KGL9 | Hmgn2 | Non-histone chromosomal protein HMG-17 | 0.81094106 | 0.614372889 |
| A3KGU7 | Sptan1 | Spectrin alpha chain, non-erythrocytic 1 | -0.001181412 | 0.02977403 |
| A3KMP2 | Ttc38 | Tetratricopeptide repeat protein 38 | -0.084749158 | 0.623865922 |
| A4GZ26 | Iqsec2 | ARF6 guanine nucleotide exchange factor IQArfGEF | -0.236826611 | 0.393807411 |
| A5A4Y9 | Ppp1r11 | E3 ubiquitin-protein ligase PPP1R11 | -0.398800882 | -0.056877136 |
| A6H5Y3 | Mtr | Methionine synthase | 0.153747781 | 0.241264502 |
| A6H611 | Mipep | Mitochondrial intermediate peptidase | 0.138467344 | -0.570545355 |
| A6H630 | Armt1 | Damage-control phosphatase ARMT1 | -0.124378204 | 0.484270255 |
| A6MDD2 | Ptprn | Protein tyrosine phosphatase receptor type N | 0.154910692 | 0.850373586 |

| | | | | |
|---|---|---|---|---|
| A6PWS5 | Gsn | Macrophage-capping protein (Fragment) | 0.237397957 | -0.089427948 |
| A6PWX9 | Uqcc1 | Ubiquinol-cytochrome-c reductase complex assembly factor 1 | 0.040972169 | -0.822698514 |
| A6X935 | Itih4 | Inter alpha-trypsin inhibitor, heavy chain 4 | 0.863133081 | 0.246792634 |
| A6X940 | Fermt2 | Fermitin family homolog 2 (Fragment) | -0.054863262 | 0.735371272 |
| A6ZI44 | Aldoa | Fructose-bisphosphate aldolase | -0.352186171 | -0.633533239 |
| A7M7S2 | Nprl3 | GATOR complex protein NPRL3 | -0.92432216 | 0.732102394 |
| A8DUK4 | Hbb-bs | Beta-globin | -0.076338387 | 0.373836517 |
| A8Y5H7 | Sec14l1 | SEC14-like protein 1 | -0.712550608 | 0.746400356 |
| A9DA50 | Lingo1 | Leucine-rich repeat and immunoglobulin-like domain-containing nogo receptor-interacting protein 1 | -0.598746808 | -0.902356942 |
| B0F2B4 | Nlgn4l | Neuroligin 4-like | 0.11968015 | 0.775277283 |
| B0QZN5 | Vamp2 | Synaptobrevin-2 | 1.542623075 | -0.380085468 |
| B0R091 | Chp1 | Calcineurin B homologous protein 1 | 0.175756677 | 1.00878183 |
| B0R0E3 | Scai | Protein SCAI (Fragment) | -0.078881931 | 0.705148379 |
| B0R0S4 | Rabepk | Rab9 effector protein with kelch motifs | 0.2903217 | -0.033342679 |
| B0V2N1 | Ptprs | Receptor-type tyrosine-protein phosphatase S | 0.200709693 | 0.910443465 |
| B1APT1 | Mpped2 | Metallophosphoesterase MPPED2 (Fragment) | -0.53937823 | 0.914458116 |
| B1APX2 | 5031439G07Rik | RIKEN cDNA 5031439G07 gene | 0.112972514 | 0.41741546 |
| B1AQ75 | Krt36 | Keratin, type I cuticular Ha6 | -0.754982313 | 0.720720609 |
| B1AQ77 | Krt15 | Keratin, type I cytoskeletal 15 | -1.961762651 | 2.28040425 |
| B1AQD4 | Rab34 | Ras-related protein Rab-34 (Fragment) | -2.089672629 | -0.180695613 |
| B1AQD9 | Unc119 | Protein unc-119 homolog A | 0.645230548 | 1.722054482 |
| B1AQF4 | Dusp3 | Dual specificity protein phosphatase | 0.075655619 | 1.076800505 |
| B1AQN2 | Ptprt | Protein-tyrosine-phosphatase | 0.316711044 | 1.135843595 |
| B1AQR8 | Lgals9 | Galectin | 1.368279807 | 0.572755496 |
| B1AQW2 | Mapt | Microtubule-associated protein | 0.16289622 | 0.300731341 |
| B1AQY2 | Arhgap23 | Rho GTPase-activating protein 23 | -0.129191176 | -0.36439085 |
| B1AQY9 | Septin8 | Septin | 0.016028372 | -0.160089811 |
| B1AQZ2 | Kif3a | Kinesin-like protein | 0.151517137 | -0.192092419 |
| B1AR13 | Cisd3 | CDGSH iron-sulfur domain-containing protein 3, mitochondrial | -0.433978399 | 0.226397355 |
| B1AR50 | Gabarap | Gamma-aminobutyric acid receptor-associated protein | -0.32078104 | 0.717419306 |
| B1AR74 | Bcas3 | Breast carcinoma-amplified sequence 3 homolog | 0.010046291 | 0.091691653 |
| B1ARU4 | Macf1 | Microtubule-actin cross-linking factor 1 | -0.20446434 | 0.254504522 |
| B1ARW4 | Ndufs5 | Complex I-15 kDa (Fragment) | 0.182364337 | 1.199585756 |
| B1AS06 | Dlgap3 | Disks large-associated protein 3 | 0.594862874 | -0.13965257 |
| B1ASE2 | Atp5h | ATP synthase subunit d, mitochondrial (Fragment) | 0.048984814 | 0.781044324 |
| B1ASZ3 | Gk | Glycerol kinase | -0.360974312 | -0.130535126 |
| B1AT82 | Prpsap1 | Phosphoribosyl pyrophosphate synthase-associated protein 1 | 0.051996326 | -0.073738575 |
| B1AT92 | Grb2 | Growth factor receptor-bound protein 2 | -0.021650823 | 0.393438021 |
| B1ATI9 | Gas7 | Growth arrest-specific protein 7 | -0.484098117 | -0.102282763 |
| B1ATR2 | Tex2 | Testis-expressed protein 2 | 0.695352046 | 1.857065042 |
| B1AU25 | Aifm1 | Apoptosis-inducing factor 1, mitochondrial | 0.072108046 | 1.588558674 |
| B1AU75 | Nasp | Nuclear autoantigenic sperm protein | -0.279902109 | 0.522009532 |
| B1AUN2 | Eif2b3 | Eukaryotic translation initiation factor 2B, subunit 3 | -0.445011075 | -0.47386233 |
| B1AUX2 | Hcfc1 | Host cell factor 1 | 0.035821025 | 0.952702363 |
| B1AUY7 | Naa10 | N-alpha-acetyltransferase 10 | -0.235548083 | 0.125985305 |

| | | | | |
|---|---|---|---|---|
| B1AVI2 | Stx17 | Syntaxin-17 (Fragment) | 0.302238878 | 0.303667704 |
| B1AVN9 | Phactr2 | Phosphatase and actin regulator | -0.32726593 | -0.173323631 |
| B1AVZ0 | Uprt | Uracil phosphoribosyltransferase homolog | 0.273714447 | 0.261995633 |
| B1AWC8 | Pde4b | Phosphodiesterase | 0.135869122 | 0.121807734 |
| B1AWD8 | Clta | Clathrin light chain | 0.06264712 | 0.247463067 |
| B1AWT2 | Rragd | Ras-related GTP-binding protein | 0.210507965 | 0.694772561 |
| B1AX58 | Pls3 | Plastin-3 | 0.306982994 | 0.660279274 |
| B1AX95 | Tprgl | Transformation-related protein 63-regulated-like | -0.718184662 | -1.441717505 |
| B1AXV0 | Frrs1l | DOMON domain-containing protein FRRS1L | 0.856144269 | 0.601203124 |
| B1AXZ4 | Elavl2 | ELAV-like protein | 0.507593918 | 0.307111263 |
| B1AY13 | Usp24 | Ubiquitin carboxyl-terminal hydrolase 24 | 0.046183904 | -0.570047855 |
| B1AZ46 | Baiap2 | Brain-specific angiogenesis inhibitor 1-associated protein 2 | 0.301694743 | 0.30962197 |
| B1AZF3 | Aatk | Serine/threonine-protein kinase LMTK1 | -0.958696524 | 0.867323716 |
| B1AZP2 | Dlgap4 | Disks large-associated protein 4 | 0.110092513 | 0.435723146 |
| B1B0B8 | Dnajc6 | Putative tyrosine-protein phosphatase auxilin (Fragment) | -1.653922304 | -0.90908138 |
| B1B0C7 | Hspg2 | Basement membrane-specific heparan sulfate proteoglycan core protein | 0.196591043 | 0.635542552 |
| B1B1A8 | Mylk | Myosin light chain kinase, smooth muscle | 0.684926764 | 0.943160852 |
| B1PSD9 | Pde4d | Phosphodiesterase | 0.086287626 | 0.918104649 |
| B2KF50 | Uhrf1bp1 | UHRF1 (ICBP90)-binding protein 1 | 0.236944008 | 0.122382164 |
| B2KGA7 | Zbtb8os | Protein archease | 0.099045054 | 0.163428148 |
| B2KGF0 | Rpe | Ribulose-phosphate 3-epimerase | -1.344395459 | 0.64585797 |
| B2M1R6 | Hnrnpk | Heterogeneous nuclear ribonucleoprotein K | -0.075163078 | 0.263679187 |
| B2M1R7 | Pcbp2 | Poly(rC)-binding protein 2 | 0.042164103 | 0.260468801 |
| B2RPU8 | Zbed5 | SCAN domain containing 3 | 0.504931037 | -0.652537028 |
| B2RQ57 | Abl2 | Tyrosine-protein kinase | 0.091466681 | 0.644965172 |
| B2RQ71 | Dip2c | Dip2c protein | 0.026718489 | 0.038311005 |
| B2RQ80 | Tnik | Tnik protein | 0.115777238 | 0.33471632 |
| B2RQC6 | Cad | CAD protein | 0.006538677 | -0.43772618 |
| B2RQK7 | Synpo2l | Synaptopodin 2-like protein | 0.198159695 | -0.030653159 |
| B2RQR5 | Sgsm1 | Small G protein signaling modulator 1 | -0.040308603 | 0.327858448 |
| B2RQS1 | Strn3 | Striatin-3 | -0.792660141 | -1.429387887 |
| B2RR82 | Itsn2 | Intersectin-2 | -1.665522846 | -0.2758406 |
| B2RRE2 | Myo18a | Myo18a protein | -0.194523716 | 0.264026801 |
| B2RRI2 | Sbno1 | Protein strawberry notch homolog 1 | -0.605196762 | 0.121755759 |
| B2RSH2 | Gnai1 | Guanine nucleotide-binding protein G(i) subunit alpha-1 | 0.86020902 | 0.483822187 |
| B2RSU7 | Gcc2 | GRIP and coiled-coil domain containing 2 | 0.269304784 | 0.683538437 |
| B2RT44 | Lonrf2 | LON peptidase N-terminal domain and ring finger 2 | -0.004216417 | 0.620596727 |
| B2RUG9 | Apc | Adenomatosis polyposis coli | 0.753913562 | 0.948782603 |
| B2RUJ5 | Apba1 | Amyloid-beta A4 precursor protein-binding family A member 1 | 0.106581624 | -0.175751845 |
| B2RUJ8 | Arhgap12 | Rho GTPase-activating protein 12 | 0.515750058 | -0.038178285 |
| B2RW11 | Ankrd34a | Ankyrin repeat domain 34A | 0.271974246 | -1.160861174 |
| B2RXC2 | Itpkb | Kinase | 0.351455561 | 0.515310605 |
| B2RXQ2 | Ppfia1 | Ppfia1 protein | -0.328396511 | -0.122710864 |
| B2RXT3 | Ogdhl | Ogdhl protein | 0.035902405 | -0.2457997 |
| B2RY56 | Rbm25 | RNA-binding protein 25 | -0.516565609 | 0.174284935 |

| | | | | |
|---|---|---|---|---|
| B7FAU5 | Emd | Emerin | -0.106598886 | 0.139223417 |
| B7FAU7 | Atp6ap1 | ATPase, H+ transporting, lysosomal accessory protein (Fragment) | 0.915168858 | 1.3399779 |
| B7FAU9 | Flna | Filamin, alpha | -0.012477938 | 0.274668535 |
| B7ZCA7 | Kazn | Kazrin (Fragment) | -0.226536433 | -0.305828075 |
| B7ZCU0 | Abi1 | Abl interactor 1 | 0.450049241 | -0.04956913 |
| B7ZNF6 | Ctnnd2 | Catenin delta-2 | 0.386522388 | 0.375603199 |
| B7ZNK6 | Aga | Aga protein | 0.609084956 | 0.760231972 |
| B7ZNL2 | Nap1l4 | Nap1l4 protein | 0.024847507 | 0.109532197 |
| B7ZNL3 | Tpm1 | Tpm1 protein | -0.389102491 | 0.67285792 |
| B7ZWG4 | Trim40 | E3 ubiquitin ligase TRIM40 | -0.082189576 | 0.718935966 |
| B8JJ84 | Stxbp6 | Syntaxin-binding protein 6 (Fragment) | 0.122496605 | -0.27086242 |
| B8QI34 | Ppfia2 | Liprin-alpha-2 | 0.124916999 | 0.121816794 |
| B9EHJ3 | Tjp1 | Tight junction protein ZO-1 | 0.293812243 | 0.434288502 |
| B9EIX2 | Cep170b | AW555464 protein | 0.114881929 | 0.217201551 |
| B9EJ80 | Pdzd8 | PDZ domain-containing protein 8 | -1.259453392 | -1.491836071 |
| B9EJA2 | Cttnbp2 | Cortactin-binding protein 2 | -0.117574056 | -0.660785993 |
| B9EJA3 | Ptprd | Protein-tyrosine-phosphatase | -0.051314608 | 0.001456261 |
| B9EKI3 | Tmf1 | TATA element modulatory factor | 0.441255697 | -1.349321405 |
| D0EM46 | Tesc | Calcineurin B homologous protein 3 | -0.811925538 | -0.316219648 |
| D3YTP3 | Mtx3 | Metaxin | -1.170320066 | -0.138771693 |
| D3YTP8 | Lsm4 | U6 snRNA-associated Sm-like protein LSm4 (Fragment) | -0.699580542 | -0.36960427 |
| D3YTQ3 | Hnrnpdl | Heterogeneous nuclear ribonucleoprotein D-like | 0.40541207 | -0.089191278 |
| D3YTQ9 | Rps15 | 40S ribosomal protein S15 | 0.854981486 | -0.61067009 |
| D3YTR4 | Mbd3 | Methyl-CpG-binding domain protein 3 | -0.723440552 | -0.596930424 |
| D3YTS1 | Sec11a | Signal peptidase complex catalytic subunit SEC11 | 0.378158172 | 0.965108077 |
| D3YTU2 | Ogfod1 | 2-oxoglutarate and iron-dependent oxygenase domain-containing protein 1 | -0.184596952 | -1.495804489 |
| D3YTU5 | Ankrd29 | Ankyrin repeat domain 29 | -0.25835336 | 0.315533161 |
| D3YTW7 | Adgrl3 | Adhesion G protein-coupled receptor L3 | -0.407607269 | 0.555202484 |
| D3YTZ7 | Josd2 | Ubiquitinyl hydrolase 1 | 0.733466498 | 0.951024532 |
| D3YU01 | Plekha1 | Pleckstrin homology domain-containing family A member 1 | 0.436187935 | 0.710143566 |
| D3YU12 | Nmral1 | NmrA-like family domain-containing protein 1 | 0.074700228 | 1.065309525 |
| D3YU63 | Enpp6 | Choline-specific glycerophosphodiester phosphodiesterase | 0.313357194 | 0.403234164 |
| D3YU89 | Cit | Citron Rho-interacting kinase | -0.054771932 | -2.6924541 |
| D3YUE4 | Fam151b | Family with sequence similarity 151, member B | -0.655801678 | 1.356457829 |
| D3YUJ3 | Ccnyl1 | Cyclin Y-like 1 | -1.888865439 | -0.755209287 |
| D3YUK4 | Ndufb10 | Complex I-PDSW (Fragment) | -0.542661889 | -0.198098818 |
| D3YUM1 | Ndufv1 | NADH dehydrogenase [ubiquinone] flavoprotein 1, mitochondrial | 0.078659852 | 0.597631454 |
| D3YUP1 | Carm1 | Coactivator-associated arginine methyltransferase 1 | 0.19920524 | -0.159452915 |
| D3YUP9 | Adam22 | Disintegrin and metalloproteinase domain-containing protein 22 | 0.863510831 | 0.719245911 |
| D3YUV9 | Eif4e2 | Eukaryotic translation initiation factor 4E type 2 | 0.481842041 | 0.794225852 |
| D3YUX2 | Mpdz | Multiple PDZ domain protein | -1.283325903 | -0.281440735 |
| D3YUY3 | Pip5k1a | Phosphatidylinositol 4-phosphate 5-kinase type-1 alpha | -1.548508199 | 1.62584122 |
| D3YUZ8 | Dlg2 | Disks large homolog 2 | -0.239100965 | 0.332990487 |
| D3YV13 | Dtx3 | E3 ubiquitin-protein ligase | -0.169813665 | 0.353267511 |

| | | | | |
|---|---|---|---|---|
| D3YVA2 | Ncald | Neurocalcin-delta (Fragment) | -0.49694465 | -0.941121419 |
| D3YVF0 | Akap5 | A-kinase anchor protein 5 | 0.101469008 | 0.186819553 |
| D3YVJ0 | Sugp2 | SURP and G-patch domain-containing protein 2 (Fragment) | -0.243119367 | -0.234513442 |
| D3YVQ6 | Rab17 | Ras-related protein Rab-17 (Fragment) | 0.258876038 | -0.213964462 |
| D3YVU0 | Usp46 | Ubiquitin carboxyl-terminal hydrolase 46 | 0.715009085 | -0.306484222 |
| D3YW09 | Sf3a2 | Splicing factor 3A subunit 2 | 0.466341591 | 1.629377921 |
| D3YW42 | Ccdc9 | Coiled-coil domain-containing protein 9 | 1.467312113 | 0.173315843 |
| D3YW97 | Ufm1 | Ubiquitin-fold modifier 1 | 0.35633386 | 0.788844268 |
| D3YWB9 | Cntnap4 | Contactin-associated protein-like 4 | 0.643416214 | 1.049937884 |
| D3YWF6 | Otub1 | Ubiquitinyl hydrolase 1 | -0.204339345 | 0.291104158 |
| D3YWN7 | Map7 | Ensconsin | 0.710756207 | 0.737743696 |
| D3YWT1 | Hnrnph3 | Heterogeneous nuclear ribonucleoprotein H3 | 0.084277344 | 0.614828904 |
| D3YWX2 | Ylpm1 | YLP motif-containing protein 1 | 0.229349232 | 0.549928188 |
| D3YWY4 | Grip1 | Glutamate receptor-interacting protein 1 | -0.223562972 | 0.815842867 |
| D3YX00 | Rgs8 | Regulator of G-protein signaling 8 (Fragment) | -0.887508074 | 0.865941366 |
| D3YX85 | Asap2 | Arf-GAP with SH3 domain, ANK repeat and PH domain-containing protein 2 | 0.042521413 | 0.098825137 |
| D3YXD1 | Ube2e2 | Ubiquitin-conjugating enzyme E2 E2 (Fragment) | -0.194056972 | 0.106277625 |
| D3YXH0 | Gm49948 | Predicted gene, 49948 | 0.151896604 | -0.146219095 |
| D3YXH5 | Sfxn5 | Sideroflexin-5 | -0.395698373 | 0.341136535 |
| D3YXK2 | Safb | Scaffold attachment factor B1 | 0.170631568 | 0.037380854 |
| D3YXP6 | Pmvk | Phosphomevalonate kinase | 0.011881733 | -0.233502865 |
| D3YXR5 | Pml | Protein PML | 0.582605298 | 0.593831062 |
| D3YXZ3 | Klc2 | Kinesin light chain | 0.078752168 | 0.14299202 |
| D3YY09 | Drap1 | Dr1-associated corepressor | -0.570265929 | 0.063704809 |
| D3YY41 | Pik3ca | Phosphatidylinositol 4,5-bisphosphate 3-kinase catalytic subunit alpha isoform | -0.587042046 | 1.292133808 |
| D3YY50 | Cutc | Copper homeostasis protein cutC homolog (Fragment) | 0.243699344 | 0.135244528 |
| D3YY73 | Sap18 | Histone deacetylase complex subunit SAP18 | 0.166985989 | -0.549477418 |
| D3YYA0 | Ash2l | Set1/Ash2 histone methyltransferase complex subunit ASH2 | -0.053486633 | 0.73479414 |
| D3YYC3 | Setdb1 | Histone-lysine N-methyltransferase SETDB1 | 0.312580999 | -1.897293886 |
| D3YYK8 | Mapre2 | Microtubule-associated protein RP/EB family member 2 (Fragment) | -0.07910916 | 0.166053613 |
| D3YYM6 | Rps5 | 40S ribosomal protein S5 (Fragment) | 0.160382048 | -0.415598234 |
| D3YYS6 | Mgll | Monoglyceride lipase | 0.097059218 | 0.095455011 |
| D3YYT0 | Cdh2 | Cadherin-2 | 0.060743237 | -0.606257121 |
| D3YYT1 | Glyr1 | Glyoxylate reductase 1 homolog | -0.775920184 | -0.309302966 |
| D3YYX8 | Fam241b | Protein FAM241B (Fragment) | 1.147998556 | 0.40593036 |
| D3YZ54 | Hacl1 | 2-hydroxyacyl-CoA lyase 1 | 1.958119615 | -3.022140265 |
| D3YZ62 | Myo5a | Unconventional myosin-Va | 0.175137774 | -0.708555857 |
| D3YZ98 | Pik3c3 | Phosphatidylinositol 3-kinase catalytic subunit type 3 | 0.021415488 | 0.340779622 |
| D3YZ99 | Ntan1 | Protein N-terminal asparagine amidohydrolase | -0.362843641 | -0.697272294 |
| D3YZA1 | Chtop | Chromatin target of PRMT1 protein (Fragment) | 0.228685602 | 0.836800098 |
| D3YZC9 | Sf1 | Splicing factor 1 | 0.211129379 | 0.013302962 |
| D3YZD8 | Aamdc | Mth938 domain-containing protein | 0.214942614 | 0.467641513 |
| D3YZI9 | Pgbd5 | PiggyBac transposable element-derived protein 5 | 0.297014078 | 0.284153779 |
| D3YZP3 | Lrrc4b | Leucine-rich repeat-containing protein 4B (Fragment) | -0.336007086 | 0.866320054 |
| D3YZP9 | Ccdc6 | Coiled-coil domain-containing protein 6 | -0.029740588 | 0.35587724 |

| | | | | |
|---|---|---|---|---|
| D3YZT7 | Plekhb1 | Pleckstrin homology domain-containing family B member 1 (Fragment) | 0.044522127 | 1.7083203 |
| D3YZU1 | Shank1 | SH3 and multiple ankyrin repeat domains protein 1 | -0.115484778 | -0.852548122 |
| D3YZW1 | Srgap1 | SLIT-ROBO Rho GTPase-activating protein 1 | 0.424391079 | 0.017962456 |
| D3YZZ4 | Tsc22d4 | TSC22 domain family protein 4 | -0.143705972 | 0.544979413 |
| D3YZZ5 | Tmed7 | Transmembrane p24-trafficking protein 7 | -0.295368719 | -0.789357682 |
| D3Z061 | Uba6 | E1 ubiquitin-activating enzyme | -0.020425638 | 0.427277247 |
| D3Z079 | Stxbp5 | Syntaxin-binding protein 5 | 0.261517207 | -0.434576035 |
| D3Z0D9 | Dnajc12 | DnaJ homolog subfamily C member 12 (Fragment) | -0.042338339 | 0.652920564 |
| D3Z0E6 | Bpnt1 | 3'(2'),5'-bisphosphate nucleotidase 1 | -0.066527907 | 0.054766973 |
| D3Z0F5 | Cops6 | COP9 signalosome complex subunit 6 | -0.060899448 | 0.273891131 |
| D3Z0K7 | Wipf2 | WAS/WASL-interacting protein family member 2 (Fragment) | 1.375595697 | -0.014568806 |
| D3Z0L4 | Chchd3 | MICOS complex subunit Mic19 (Fragment) | 0.639813995 | 1.534882863 |
| D3Z0M6 | Dync1i1 | Cytoplasmic dynein 1 intermediate chain 1 | 0.156994279 | -0.132007758 |
| D3Z0R1 | Tpmt | Thiopurine S-methyltransferase (Fragment) | 1.320555417 | 0.161495209 |
| D3Z0S7 | Stambpl1 | AMSH-like protease (Fragment) | 0.185044114 | 1.011996428 |
| D3Z0T5 | Cnpy4 | Protein canopy homolog 4 (Fragment) | 1.217753601 | 0.54657793 |
| D3Z0V7 | Tsc22d1 | TSC22 domain family protein 1 | -0.105657514 | -0.309145292 |
| D3Z118 | Ica1 | Islet cell autoantigen 1 | 0.089098136 | 0.415125052 |
| D3Z158 | Qars | Glutaminyl-tRNA synthetase | -0.40237023 | -0.502442519 |
| D3Z1E6 | Adam10 | Disintegrin and metalloproteinase domain-containing protein 10 (Fragment) | -0.972030354 | -2.019512335 |
| D3Z1T0 | Alkbh6 | Alpha-ketoglutarate-dependent dioxygenase alkB homolog 6 (Fragment) | 0.323524412 | 0.521934668 |
| D3Z1T8 | Ttc33 | Tetratricopeptide repeat protein 33 (Fragment) | 0.209919008 | 0.342451096 |
| D3Z220 | Ctsc | Dipeptidyl peptidase 1 (Fragment) | 0.625143305 | -0.036095142 |
| D3Z275 | Pcyox1 | Prenylcysteine oxidase (Fragment) | 1.173973052 | -1.90892005 |
| D3Z2E3 | Reps1 | RalBP1-associated Eps domain-containing protein 1 | 0.047595151 | 0.266216278 |
| D3Z2H2 | Ctnnd1 | Catenin delta-1 | 0.753322935 | -0.978681087 |
| D3Z2H9 | Tpm3-rs7 | Tropomyosin 3, related sequence 7 | -0.241419252 | 0.653862158 |
| D3Z2Z1 | Clip1 | CAP-Gly domain-containing linker protein 1 | 0.082895406 | 0.131074111 |
| D3Z325 | Tnfaip8 | Tumor necrosis factor alpha-induced protein 8 | -0.562356695 | 0.069946448 |
| D3Z396 | Ntm | Neurotrimin | 0.363442643 | 0.545339266 |
| D3Z3C1 | Faim | Fas apoptotic inhibitory molecule 1 | 0.3320913 | 0.044910272 |
| D3Z3J6 | Paip1 | Polyadenylate-binding protein-interacting protein 1 | 0.660652765 | 0.622205575 |
| D3Z3Y4 | Tial1 | Nucleolysin TIAR | 0.643277327 | 0.741466999 |
| D3Z3Z3 | Cacnb3 | Calcium channel voltage-dependent subunit beta 3 | 0.07956953 | 0.724007924 |
| D3Z479 | Snx15 | Sorting nexin-15 | -0.027142493 | 0.893409729 |
| D3Z497 | Gm7075 | Predicted gene 7075 | 0.359136264 | 0.389776707 |
| D3Z4C9 | Uqcc2 | Mitochondrial nucleoid factor 1 | -0.157006741 | -1.285612841 |
| D3Z4D2 | Rufy2 | RUN and FYVE domain-containing protein 2 | 0.172296047 | 1.48087724 |
| D3Z4H6 | Tia1 | Nucleolysin TIA-1 | 0.264732997 | 0.500288486 |
| D3Z4J9 | Eml1 | Echinoderm microtubule-associated protein-like 1 | 0.194668452 | 1.456417243 |
| D3Z4S6 | Tmem132a | Transmembrane protein 132A | -1.262767331 | -0.764804284 |
| D3Z4U0 | Zranb2 | Zinc finger Ran-binding domain-containing protein 2 | -0.034436925 | 0.415499369 |
| D3Z510 | Aldoa | Fructose-bisphosphate aldolase (Fragment) | -0.773086135 | -0.180924733 |
| D3Z553 | Churc1 | Protein Churchill | 0.101906459 | -0.212311904 |
| D3Z568 | Ndufb5 | Complex I-SGDH | -0.667021863 | -0.147011042 |

| | | | | |
|---|---|---|---|---|
| D3Z5N5 | Pikfyve | 1-phosphatidylinositol-3-phosphate 5-kinase | 0.282995542 | 0.29989481 |
| D3Z5R4 | Wipf3 | WAS/WASL-interacting protein family member 3 | 0.586577511 | 0.678836028 |
| D3Z5T2 | Armc10 | Armadillo repeat-containing protein 10 | 1.118143463 | -0.193224907 |
| D3Z600 | Pex5 | PTS1-BP | -0.693741004 | 1.601114591 |
| D3Z601 | Clstn3 | Calsyntenin-3 | -0.238055865 | -0.352195899 |
| D3Z632 | Aktip | AKT-interacting protein | 0.522279771 | -0.613769531 |
| D3Z645 | Vps29 | Vacuolar protein sorting-associated protein 29 | -0.035726198 | 0.422088464 |
| D3Z656 | Synj1 | Phosphoinositide 5-phosphatase | -0.103103383 | 0.201745033 |
| D3Z6B9 | Aldh1l2 | Formyltetrahydrofolate dehydrogenase | 0.042151642 | 1.211157163 |
| D3Z6G3 | Mapre3 | Microtubule-associated protein RP/EB family member 3 | -0.010488383 | 0.271989187 |
| D3Z6H3 | Dctn6 | Dynactin subunit 6 | 0.49593598 | 0.506857077 |
| D3Z6I4 | Cryzl1 | Quinone oxidoreductase-like protein 1 | 0.348788548 | 0.126627445 |
| D3Z6I8 | Tpm3 | Tropomyosin alpha-3 chain | -0.166926003 | 0.683095296 |
| D3Z6S5 | Plekhb1 | Pleckstrin homology domain-containing family B member 1 (Fragment) | 0.053974342 | 0.561212381 |
| D3Z723 | Nfyb | CAAT box DNA-binding protein subunit B (Fragment) | -0.84293588 | 0.020141125 |
| D3Z780 | Eif2b4 | Translation initiation factor eIF-2B subunit delta | -0.36188914 | 0.179672559 |
| D3Z794 | Sumo2 | Small ubiquitin-related modifier 2 | 0.08726174 | -0.151598612 |
| D3Z7E5 | Gsk3a | [Tau protein] kinase | 0.070436827 | 0.08066686 |
| D3Z7G4 | BC024978 | cDNA sequence BC024978 | -0.338547738 | -0.136951129 |
| D3Z7P3 | Gls | Glutaminase kidney isoform, mitochondrial | -0.114360809 | 0.330125809 |
| D3Z7T4 | Hdac10 | Polyamine deacetylase HDAC10 | -0.934302425 | 0.34957393 |
| D6RD11 | Ptprg | Receptor-type tyrosine-protein phosphatase gamma | 0.095443249 | -0.126246293 |
| D6RDE8 | Desi1 | Desumoylating isopeptidase 1 | -0.616169484 | 0.306684653 |
| D6REG4 | Grhpr | Glyoxylate reductase/hydroxypyruvate reductase | -2.247174899 | -1.551992178 |
| D6RET6 | Gpatch1 | G patch domain-containing protein 1 | 0.228976059 | 0.239309152 |
| D6REU0 | Mindy3 | Ubiquitin carboxyl-terminal hydrolase MINDY (Fragment) | -0.154637241 | 0.721156279 |
| D6RHA2 | Renbp | GlcNAc 2-epimerase | 0.529999606 | 1.134419441 |
| D6RIM8 | Ush1c | Harmonin | -1.038005606 | 0.042279243 |
| E0CX38 | Ttc39b | Tetratricopeptide repeat protein 39B (Fragment) | -2.429596509 | -0.159799735 |
| E0CX39 | Ggact | Gamma-glutamylaminecyclotransferase (Fragment) | -0.802078533 | 0.551413774 |
| E0CX41 | Lrrtm1 | Leucine-rich repeat transmembrane neuronal protein 1 (Fragment) | 0.207855352 | 0.727603833 |
| E0CX53 | Tbc1d4 | TBC1 domain family member 4 | -0.496336969 | 0.322293441 |
| E0CXA9 | Mob4 | MOB-like protein phocein | 0.301796627 | 0.597360611 |
| E0CXB9 | Ctnna2 | Alpha N-catenin | 0.074268945 | 0.063267072 |
| E0CXD4 | Pcdh1 | Protocadherin 1 | -0.048341592 | -0.723861376 |
| E0CXN5 | Gpd1 | Glycerol-3-phosphate dehydrogenase [NAD(+)] | -0.4514534 | -0.151088556 |
| E0CXS2 | Tle3 | Transducin-like enhancer protein 3 | 0.054561234 | 0.68358326 |
| E0CXZ1 | Fsd1l | FSD1-like protein (Fragment) | 0.335382303 | -0.209839503 |
| E0CY16 | Cadm1 | Cell adhesion molecule 1 | 0.875148614 | 1.146435897 |
| E0CY18 | Zfand2b | AN1-type zinc finger protein 2B (Fragment) | 0.025085926 | 0.563054085 |
| E0CY88 | Cyb5a | Cytochrome b5 | 0.015289275 | 0.381732464 |
| E0CY96 | Tsen15 | tRNA-splicing endonuclease subunit Sen15 | 0.234709279 | 2.9195895 |
| E0CYF5 | Maf1 | Repressor of RNA polymerase III transcription MAF1 | 0.590837129 | 0.31606706 |
| E0CYQ2 | Nudcd2 | NudC domain-containing protein 2 | -0.225459003 | 0.469850858 |
| E0CYT1 | Wnk2 | Non-specific serine/threonine protein kinase | -0.473080921 | 0.351848125 |

| | | | | |
|---|---|---|---|---|
| E0CYV0 | Pcmt1 | Protein-L-isoaspartate O-methyltransferase | -0.001788839 | 0.673175812 |
| E0CYV9 | | Uncharacterized protein C4orf54 homolog | -0.125936826 | 0.615150134 |
| E0CZ16 | Klhl3 | Kelch-like protein 3 | -0.677942403 | -1.466417472 |
| E0CZ50 | Ndrg4 | Protein NDRG4 (Fragment) | 0.577479712 | 0.965566953 |
| E0CZ72 | Kif2a | Kinesin-like protein | 0.001617908 | 0.421707312 |
| E0CZ78 | Ppp3cb | Serine/threonine-protein phosphatase | 0.054724058 | 0.319129944 |
| E0CZE0 | Nae1 | NEDD8-activating enzyme E1 regulatory subunit | -0.009311612 | 0.398003896 |
| E9PU87 | Sik3 | Non-specific serine/threonine protein kinase | -0.023568757 | 0.126956622 |
| E9PUA7 | Tpd52 | Tumor protein D52 | 0.152758376 | 0.181188424 |
| E9PUB7 | Msto1 | Protein misato homolog 1 | 0.242584769 | -0.467474937 |
| E9PUC5 | Psd3 | PH and SEC7 domain-containing protein 3 | 0.145323245 | -0.302665869 |
| E9PUD2 | Dnm1l | Dynamin-1-like protein | 0.17176431 | 0.225408236 |
| E9PUL5 | Prrt2 | Proline-rich transmembrane protein 2 | 1.306530412 | 1.08277909 |
| E9PUQ7 | Trmt2a | tRNA (uracil-5-)-methyltransferase homolog A | -0.242423503 | -0.591552575 |
| E9PUR6 | Nr3c1 | Glucocorticoid receptor | 1.021203804 | 1.287592729 |
| E9PUW7 | Xpo7 | Exportin-7 | 0.111370373 | 0.10513862 |
| E9PUZ5 | Pick1 | PRKCA-binding protein | -0.052961763 | 0.59633112 |
| E9PV14 | Epb41l1 | Band 4.1-like protein 1 | 0.00598952 | 0.058704535 |
| E9PV22 | Lrrc47 | Leucine-rich repeat-containing protein 47 | -0.171750641 | 0.476483186 |
| E9PV24 | Fga | Fibrinogen alpha chain | 0.305875524 | 1.235450427 |
| E9PV41 | Diaph1 | Protein diaphanous homolog 1 | -0.349197133 | 0.418114662 |
| E9PV44 | Atpif1 | ATP synthase F1 subunit epsilon | 0.394865735 | 0.096924305 |
| E9PV48 | Ifit3b | Interferon-induced protein with tetratricopeptide repeats 3B | 2.114112457 | 1.448904991 |
| E9PV69 | Cacul1 | CDK2-associated and cullin domain-containing protein 1 | -0.231595453 | 0.994760672 |
| E9PVA8 | Gcn1 | eIF-2-alpha kinase activator GCN1 | 0.184800593 | -0.590667884 |
| E9PVB6 | Limk1 | LIM domain kinase 1 | -0.429757849 | 1.409890811 |
| E9PVC6 | Eif4g1 | Eukaryotic translation initiation factor 4 gamma 1 | -0.129105568 | 0.168516795 |
| E9PVC7 | Cdk14 | Cyclin-dependent kinase 14 | 0.018623702 | -0.054185867 |
| E9PVM1 | Inpp4b | Phosphatidylinositol-3,4-bisphosphate 4-phosphatase | -0.562450473 | 0.522819281 |
| E9PVU0 | Myo6 | Unconventional myosin-6 | -0.325189527 | 0.65109849 |
| E9PW07 | Cbarp | Voltage-dependent calcium channel beta subunit-associated regulatory protein | 0.777912553 | -0.30905056 |
| E9PWB2 | Csnk1a1 | Casein kinase I isoform alpha | -0.399776967 | -0.165755113 |
| E9PWC8 | Papola | Poly(A) polymerase | -1.946269353 | -0.225525379 |
| E9PWV3 | Rps6ka1 | Ribosomal protein S6 kinase | -0.86680034 | 0.626224677 |
| E9PWY9 | Farsa | Phenylalanine--tRNA ligase | 0.05217514 | 0.229705334 |
| E9PX23 | Mta1 | Metastasis-associated protein MTA1 | -1.036461862 | 1.104308287 |
| E9PX28 | Pcdh10 | Protocadherin 10 | -0.437755299 | 0.441050529 |
| E9PX29 | Sptbn4 | Spectrin beta chain | 0.222992071 | -0.38105917 |
| E9PX33 | Rell2 | RELT-like protein 2 | 1.152536837 | 0.360189915 |
| E9PX53 | Ppp4r1 | Serine/threonine-protein phosphatase 4 regulatory subunit 1 | 0.371604395 | 0.386356036 |
| E9PX68 | Slc4a1ap | Solute carrier family 4 (anion exchanger), member 1, adaptor protein | 0.198630079 | 0.301463604 |
| E9PX89 | Bola3 | BolA-like protein 3 | -1.731350923 | -0.48628974 |
| E9PXF0 | Pcdh17 | Protocadherin 17 | -0.472223997 | 0.372342189 |
| E9PXP7 | Ston2 | Stonin-2 | -0.042972851 | 0.25112764 |
| E9PXX7 | Txndc5 | Thioredoxin domain-containing protein 5 | 0.134359264 | 0.033625444 |

| | | | | |
|---|---|---|---|---|
| E9PXY8 | Usp7 | Ubiquitin carboxyl-terminal hydrolase 7 | -0.153591887 | 0.213556449 |
| E9PY16 | Adap1 | ArfGAP with dual PH domains 1 | -0.200580883 | 0.307468096 |
| E9PYB0 | Ahnak2 | AHNAK nucleoprotein 2 (Fragment) | -0.197158813 | 0.342377345 |
| E9PYB1 | Tbc1d14 | TBC1 domain family member 14 | -0.672377745 | -0.286200205 |
| E9PYF4 | Lmo7 | LIM domain only 7 | 0.395397409 | 1.059808731 |
| E9PYG6 | Rasa1 | RAS p21 protein activator 1 | 0.159033553 | -0.302921613 |
| E9PYG9 | Samhd1 | Deoxynucleoside triphosphate triphosphohydrolase SAMHD1 | -1.284464582 | -0.690514882 |
| E9PYH0 | Vcan | Versican core protein | -0.269822852 | 0.241636753 |
| E9PYH2 | Acot7 | Cytosolic acyl coenzyme A thioester hydrolase | -0.2108874 | 0.368970235 |
| E9PYH3 | Etnppl | Ethanolamine-phosphate phospho-lyase | -0.167434057 | 0.142128944 |
| E9PYI8 | Usp14 | Ubiquitin carboxyl-terminal hydrolase | -0.090847365 | 0.229961395 |
| E9PYJ7 | Pitpnm2 | Membrane-associated phosphatidylinositol transfer protein 2 | 0.214567534 | -1.57217133 |
| E9PYJ9 | Ldb3 | LIM domain-binding protein 3 | -1.251803907 | 0.460076253 |
| E9PYN1 | Cadm1 | Cell adhesion molecule 1 | 1.26176432 | 1.263089816 |
| E9PYP8 | Crybb1 | Beta-B1 crystallin (Fragment) | 0.80580918 | 1.398866653 |
| E9PYT0 | Arhgap5 | Rho GTPase-activating protein 5 | -0.320492395 | -0.137629986 |
| E9PZ00 | Psap | Prosaposin | 0.035121981 | 1.003907998 |
| E9PZ43 | Map4 | Microtubule-associated protein | -0.17339201 | 0.376547337 |
| E9PZ91 | Fhit | Bis(5'-adenosyl)-triphosphatase | -0.036189715 | 0.102368196 |
| E9PZD8 | Cp | Ceruloplasmin | 0.375878874 | -0.212349693 |
| E9PZF5 | Anp32e | Acidic leucine-rich nuclear phosphoprotein 32 family member E (Fragment) | -1.666783778 | 0.973587195 |
| E9PZI9 | Cd200 | OX-2 membrane glycoprotein | 0.417914772 | 0.864418825 |
| E9PZM8 | Nptxr | Neuronal pentraxin receptor | -0.023175081 | 0.213883241 |
| E9Q027 | Cnot2 | CCR4-NOT transcription complex subunit 2 | -0.342257182 | -1.403880437 |
| E9Q0A7 | Oxr1 | Oxidation resistance protein 1 | -0.302724202 | -0.712670803 |
| E9Q0F0 | Krt78 | Keratin 78 | 1.869459089 | 1.310343107 |
| E9Q0H6 | Fabp7 | Fatty acid-binding protein, brain | -0.220606359 | 0.407995542 |
| E9Q0J5 | Kif21a | Kinesin-like protein KIF21A | -0.348422496 | -0.278953552 |
| E9Q0N0 | Itsn1 | Intersectin-1 | 0.023957475 | 0.345170657 |
| E9Q0W6 | Ablim2 | Actin-binding LIM protein 2 | -0.082216231 | -0.023026625 |
| E9Q0W8 | Snrpe | Small nuclear ribonucleoprotein E | -0.547350121 | 2.738630772 |
| E9Q150 | Dvl3 | Segment polarity protein dishevelled homolog DVL-3 | -0.146058432 | -0.144721349 |
| E9Q179 | Grsf1 | G-rich sequence factor 1 | -0.244702816 | 0.959696213 |
| E9Q1D5 | Tgfb1i1 | Transforming growth factor beta-1-induced transcript 1 protein | -0.6178243 | 0.247516155 |
| E9Q1G8 | Septin7 | Septin | 0.003487746 | 0.047307491 |
| E9Q1S3 | Sec23a | Protein transport protein SEC23 | -0.139892101 | -0.006847064 |
| E9Q1U6 | Ctif | CBP80/20-dependent translation initiation factor | 0.321402136 | 1.068336646 |
| E9Q1V2 | Cpne5 | Copine-5 (Fragment) | 0.798900286 | -0.58218956 |
| E9Q1W0 | Camk2d | Calcium/calmodulin-dependent protein kinase | 0.472340679 | -0.030430317 |
| E9Q1X8 | Cacna2d1 | Voltage-dependent calcium channel subunit alpha-2/delta-1 | 0.048839887 | 0.170226574 |
| E9Q1Y3 | Apob | Apolipoprotein B-100 (Fragment) | 1.522303168 | 1.22517697 |
| E9Q1Z0 | Krt90 | Keratin 90 | 0.04282608 | 1.304093361 |
| E9Q2U4 | Naa35 | N-alpha-acetyltransferase 35, NatC auxiliary subunit (Fragment) | 0.045090707 | -0.401596069 |
| E9Q2W9 | Actn4 | Alpha-actinin-4 (Fragment) | -0.416512299 | 2.595565955 |

| | | | | |
|---|---|---|---|---|
| E9Q2X2 | Nrxn3 | Neurexin-3-beta | 0.153958352 | 0.577393373 |
| E9Q350 | Baiap3 | BAI1-associated protein 3 | 0.407597542 | 0.540993373 |
| E9Q3E2 | Synpo | Synaptopodin | 0.492831548 | 1.093088627 |
| E9Q3G8 | Nup153 | Nucleoporin 153 | -0.238607407 | -0.081391335 |
| E9Q3M0 | Smad2 | Mothers against decapentaplegic homolog | 1.046684964 | -0.197927157 |
| E9Q3M3 | Dctn1 | Dynactin subunit 1 | -0.023321279 | -0.117148081 |
| E9Q3M9 | Cracdl | Capping protein-inhibiting regulator of actin-like | 0.231049252 | 0.274372101 |
| E9Q3Q6 | Alcam | Activated leukocyte cell adhesion molecule | 1.247978242 | 1.850096623 |
| E9Q3T8 | Trappc9 | Trafficking protein particle complex subunit 9 | 0.636457666 | -0.840210915 |
| E9Q3V0 | Slc6a9 | Sodium- and chloride-dependent glycine transporter 1 | -0.528608068 | -0.02865839 |
| E9Q3V9 | Adgrl1 | Adhesion G protein-coupled receptor L1 | -1.316715209 | -0.210947037 |
| E9Q446 | Rnf25 | E3 ubiquitin-protein ligase RNF25 | 0.032497152 | 0.54407994 |
| E9Q447 | Sptan1 | Spectrin alpha chain, non-erythrocytic 1 | 0.477566051 | -0.04047219 |
| E9Q452 | Tpm1 | Tropomyosin alpha-1 chain | -0.878902404 | 3.242145061 |
| E9Q456 | Tpm1 | Tropomyosin alpha-1 chain | 0.053320567 | 0.483448029 |
| E9Q481 | Ppp4r3a | Serine/threonine-protein phosphatase 4 regulatory subunit 3A | 0.163215097 | 1.564625104 |
| E9Q4M4 | Chchd6 | MICOS complex subunit Mic25 | -0.958640575 | 1.697242439 |
| E9Q4Z6 | Dzank1 | Double zinc ribbon and ankyrin repeat-containing protein 1 | 0.196011003 | 1.324254354 |
| E9Q512 | Trip11 | Thyroid hormone receptor interactor 11 | 0.12292881 | 1.123096625 |
| E9Q557 | Dsp | Desmoplakin | -0.067057578 | -0.425350666 |
| E9Q5B5 | Hk2 | Hexokinase | -0.650501092 | 1.034567197 |
| E9Q5D6 | Ranbp9 | Ran-binding protein 9 | 0.237727865 | 0.590034803 |
| E9Q616 | Ahnak | AHNAK nucleoprotein (desmoyokin) | 0.306936105 | 1.084345818 |
| E9Q6E0 | Mapk8ip3 | C-Jun-amino-terminal kinase-interacting protein 3 | 0.1254378 | 0.306477865 |
| E9Q6E5 | Srsf11 | Serine and arginine-rich-splicing factor 11 | -1.258412329 | 1.537530581 |
| E9Q6J5 | Bod1l | Biorientation of chromosomes in cell division protein 1-like 1 | 1.078193808 | 2.784983178 |
| E9Q6J8 | Dmwd | Dystrophia myotonica WD repeat-containing protein | 0.358809916 | -0.072815418 |
| E9Q6R7 | Utrn | Utrophin | -0.559914716 | -0.053329468 |
| E9Q6U4 | Pfdn4 | Prefoldin subunit 4 | -0.29070247 | 1.036845207 |
| E9Q7B0 | P4ha1 | Procollagen-proline 4-dioxygenase | 0.175246175 | 0.843600909 |
| E9Q7E9 | Pacs2 | Phosphofurin acidic cluster sorting protein 2 | -0.491463121 | 0.950230916 |
| E9Q7G0 | Numa1 | Nuclear mitotic apparatus protein 1 | -0.190406513 | -0.127514998 |
| E9Q7N5 | Gfm2 | Ribosome-releasing factor 2, mitochondrial | 0.138040876 | 1.19860061 |
| E9Q7U2 | Calcoco1 | Calcium-binding and coiled-coil domain-containing protein 1 | 0.319066525 | 0.326501369 |
| E9Q7V6 | Pde1c | Phosphodiesterase | -0.073755169 | -2.147297859 |
| E9Q827 | Arpp19 | cAMP-regulated phosphoprotein 19 | 0.046806717 | 0.139486949 |
| E9Q828 | Atp2b4 | Calcium-transporting ATPase | 0.680832227 | 0.136715094 |
| E9Q835 | Cadps2 | Calcium-dependent secretion activator 2 | -0.134222857 | -0.024109523 |
| E9Q855 | Scamp3 | Secretory carrier-associated membrane protein | -1.047333606 | 0.832546473 |
| E9Q8A3 | Pi4kb | Phosphatidylinositol 4-kinase beta | -0.602030691 | -0.531970978 |
| E9Q8N5 | Clasp2 | CLIP-associating protein 2 | -0.166131274 | 0.310947736 |
| E9Q8P8 | Fam131b | Protein FAM131B | 0.413003794 | -0.020643393 |
| E9Q912 | Rap1gds1 | Rap1 GTPase-GDP dissociation stimulator 1 | 0.022101657 | 0.577645938 |
| E9Q9B0 | Kif1c | Kinesin-like protein KIF1C | 1.680415535 | 0.337519328 |
| E9Q9C0 | Ablim1 | Actin-binding LIM protein 1 | 0.083023771 | -0.144944191 |

| | | | | |
|---|---|---|---|---|
| E9Q9C3 | Afdn | Afadin | 0.295952543 | 0.312191645 |
| E9Q9G6 | Kif1a | Kinesin-like protein KIF1A | 1.71158212 | -0.126292388 |
| E9Q9H0 | Dlg1 | Disks large homolog 1 | 0.621871297 | 0.67639327 |
| E9Q9J4 | Ppip5k2 | Inositol hexakisphosphate and diphosphoinositol-pentakisphosphate kinase | -1.218857034 | -0.101463477 |
| E9Q9T8 | Mybpc3 | Myosin-binding protein C, cardiac-type | 0.976547686 | 1.473906676 |
| E9QA16 | Cald1 | Caldesmon 1 | -0.29954319 | 0.518410365 |
| E9QAQ3 | Arhgap26 | Rho GTPase-activating protein 26 | 0.491612053 | 0.006011963 |
| E9QAQ5 | Gsk3b | Glycogen synthase kinase-3 beta | -0.186798414 | 0.14733394 |
| E9QAS4 | Chd4 | DNA helicase | 0.118448702 | 0.088713487 |
| E9QAT4 | Sec16a | Protein transport protein Sec16A | -0.129840247 | -1.107634703 |
| E9QB01 | Ncam1 | Neural cell adhesion molecule 1 | 0.059097735 | 1.654202143 |
| E9QB02 | Mars1 | Methionine--tRNA ligase, cytoplasmic | -0.113301373 | -0.369641145 |
| E9QK04 | Neo1 | Neogenin | -0.181356176 | 0.940560659 |
| E9QK34 | Nlgn1 | Neuroligin-1 | 0.295691236 | 0.843527953 |
| E9QK48 | Eml2 | Echinoderm microtubule-associated protein-like 2 | -0.208823395 | 0.351682822 |
| E9QK62 | Ngef | Ephexin-1 | 0.2051651 | 0.240073999 |
| E9QKB2 | Habp4 | Intracellular hyaluronan-binding protein 4 | 0.203250821 | 0.056102753 |
| E9QKI4 | Akt1s1 | Proline-rich AKT1 substrate 1 | 0.336010075 | -0.093487263 |
| E9QKZ2 | Ipo9 | Importin-9 | -0.175411828 | -0.023209254 |
| E9QL65 | Cog3 | Component of oligomeric Golgi complex 3 | 0.069864591 | -0.382162889 |
| E9QLA5 | Inf2 | Inverted formin-2 | -0.292257436 | 0.232996305 |
| E9QLB2 | Lyplal1 | Lysophospholipase-like protein 1 | 0.530910333 | 0.550733725 |
| E9QLK9 | Snap91 | Clathrin coat assembly protein AP180 | 0.184154288 | 0.636865139 |
| E9QM38 | Slc12a2 | Solute carrier family 12 member 2 | 0.045227432 | 0.428711573 |
| E9QM77 | Atxn2 | Ataxin-2 | -0.005675983 | -0.336526712 |
| E9QM90 | Relch | RAB11-binding protein RELCH | 0.04095637 | 0.079686642 |
| E9QMA2 | Fam172a | Cotranscriptional regulator FAM172A | 0.120346928 | -0.098663489 |
| E9QMB7 | Naa30 | N-alpha-acetyltransferase 30 | 0.614044412 | 1.683799744 |
| E9QMC2 | Grm5 | Metabotropic glutamate receptor 5 | -0.153860188 | 0.384651423 |
| E9QMI7 | Asap1 | Arf-GAP with SH3 domain, ANK repeat and PH domain-containing protein 1 | 0.048626932 | 0.249886672 |
| E9QMK9 | Dglucy | D-glutamate cyclase, mitochondrial | -0.02772665 | 0.493462086 |
| E9QN14 | Srgap3 | SLIT-ROBO Rho GTPase-activating protein 3 | -0.253054333 | -0.014811675 |
| E9QNA7 | Sorbs1 | Sorbin and SH3 domain-containing protein 1 | 0.03190705 | 0.354343891 |
| E9QND8 | Atl2 | Atlastin-2 | 1.643618711 | 0.718305111 |
| E9QNF7 | Cntnap2 | Contactin-associated protein-like 2 | 0.136537552 | 0.438218594 |
| E9QP54 | Ahi1 | Jouberin | 0.562206999 | 0.084366481 |
| E9QPD7 | Pcx | Pyruvate carboxylase | -0.265268008 | 0.077214877 |
| E9QPU1 | Vwf | von Willebrand factor | -0.092121442 | 1.007178624 |
| E9QPX3 | Ndufs4 | NADH dehydrogenase [ubiquinone] iron-sulfur protein 4, mitochondrial | 1.464600245 | 0.957255205 |
| E9QPZ3 | Flg2 | Filaggrin-2 | -0.170760791 | -1.984743675 |
| E9QQ10 | Akap9 | A-kinase anchor protein 9 | -0.391767279 | 0.514661789 |
| E9QQ38 | Ddhd1 | Phospholipase DDHD1 | 0.508356524 | 0.780264854 |
| F2Z3U3 | Raph1 | Ras association (RalGDS/AF-6) and pleckstrin homology domains 1 | 0.364110025 | 0.583548546 |
| F2Z3X3 | Cuedc1 | CUE domain-containing protein 1 | 0.322835191 | -0.314370314 |
| F2Z3X6 | Pdpk1 | 3-phosphoinositide-dependent protein kinase 1 | 0.188088894 | 2.09995079 |

| | | | | |
|---|---|---|---|---|
| F2Z455 | Fhl3 | Four and a half LIM domains protein 3 | 0.818180211 | 1.76018556 |
| F2Z456 | Cyb5r3 | NADH-cytochrome b5 reductase | -0.077125581 | -0.313477198 |
| F6QCI0 | Taf15 | TATA-box-binding protein-associated factor 15 (Fragment) | -0.208428987 | -0.225618919 |
| F6QHD1 | Mtdh | Protein LYRIC (Fragment) | -0.313699977 | 0.907543182 |
| F6R3V4 | Tcof1 | Treacle protein | -1.084212462 | 0.15182972 |
| F6RJ39 | Acin1 | Apoptotic chromatin condensation inducer in the nucleus (Fragment) | -0.387395032 | 0.546038469 |
| F6RPJ9 | Ide | Insulin-degrading enzyme (Fragment) | -0.016944472 | 0.217882156 |
| F6RXX6 | Wdr73 | WD repeat-containing protein 73 (Fragment) | 0.476255862 | 0.491328955 |
| F6S1R2 | Ahsa2 | Activator of 90 kDa heat shock protein ATPase homolog 2 (Fragment) | 0.107796987 | 0.208204587 |
| F6S2R9 | Dbr1 | Lariat debranching enzyme (Fragment) | -0.399103387 | -0.42171065 |
| F6SQH7 | Poldip2 | Polymerase delta-interacting protein 2 (Fragment) | 0.150190099 | -0.78406779 |
| F6T4L3 | Manf | Mesencephalic astrocyte-derived neurotrophic factor (Fragment) | 1.865675275 | 2.981311162 |
| F6TCF9 | Bag1 | BAG family molecular chaperone regulator 1 | 0.341965675 | 0.719354153 |
| F6TZK4 | Rims1 | Regulating synaptic membrane exocytosis protein 1 | 0.781731987 | 0.122082392 |
| F6UP77 | Amdhd2 | Amidohydrolase domain-containing protein 2 | -0.460055129 | -0.162506898 |
| F6V2U0 | Inpp4a | Phosphatidylinositol-3,4-bisphosphate 4-phosphatase | -0.103885619 | 0.049833457 |
| F6W8I0 | Yjefn3 | YjeF N-terminal domain-containing protein 3 | -0.285728137 | 0.334313234 |
| F6WNR1 | Plod1 | Procollagen-lysine,2-oxoglutarate 5-dioxygenase 1 (Fragment) | 0.374959135 | 1.059223493 |
| F6WR04 | Ctss | Cathepsin S | 0.596942298 | 0.714355946 |
| F6X4L9 | Coq5 | 2-methoxy-6-polyprenyl-1,4-benzoquinol methylase, mitochondrial (Fragment) | -1.559810607 | -0.093234062 |
| F6XC25 | Cc2d1b | Coiled-coil and C2 domain-containing protein 1B (Fragment) | 0.04311142 | 0.173382441 |
| F6XVP7 | Rnf31 | RBR-type E3 ubiquitin transferase (Fragment) | -0.673806318 | 1.816629062 |
| F6Y616 | Paip1 | Polyadenylate-binding protein-interacting protein 1 | 0.294208177 | -0.013043563 |
| F6YVP7 | Rps18-ps6 | 40S ribosomal protein S18 | -0.129029814 | -0.090261777 |
| F6Z1R4 | Cltc | Clathrin heavy chain 1 (Fragment) | 0.689259942 | 0.385492245 |
| F6Z9T5 | R3hdm2 | R3H domain-containing protein 2 (Fragment) | -0.256868966 | 0.780729055 |
| F6ZBL2 | Mib1 | RING-type E3 ubiquitin transferase (Fragment) | -0.28425436 | 0.985225677 |
| F6ZCF0 | Agtpbp1 | Cytosolic carboxypeptidase 1 (Fragment) | -0.747925536 | -0.703506947 |
| F6ZDS4 | Tpr | Nucleoprotein TPR | -0.068832652 | 0.186083953 |
| F6ZFU0 | Eef1d | Elongation factor 1-delta (Fragment) | 0.21911033 | -0.81987381 |
| F6ZHD8 | Gbe1 | 1,4-alpha-glucan branching enzyme | -0.479094982 | -0.688518524 |
| F7AA26 | Pakap | Paralemmin A kinase anchor protein (Fragment) | 0.117138354 | 0.62753137 |
| F7AI87 | Psmd4 | 26S proteasome non-ATPase regulatory subunit 4 (Fragment) | 0.108139769 | -0.31481266 |
| F7B1B6 | Coq8a | Atypical kinase COQ8A, mitochondrial (Fragment) | 0.69857502 | 1.755861024 |
| F7B227 | Ech1 | Delta(3,5)-Delta(2,4)-dienoyl-CoA isomerase, mitochondrial (Fragment) | 0.782033126 | 3.786350409 |
| F7BAB2 | Tmem132b | Transmembrane protein 132B | -0.041857942 | 1.267538548 |
| F7BGR7 | Gm21992 | Predicted gene 21992 | -0.754577192 | -0.327633699 |
| F7BJK1 | Pcdh1 | Protocadherin 1 (Fragment) | 0.012784513 | 0.487593492 |
| F7C3A0 | Plch2 | Phosphoinositide phospholipase C | -0.119537989 | -0.741344293 |
| F7CBP1 | Eif4g2 | Eukaryotic translation initiation factor 4 gamma 2 | -0.220004845 | 1.273269812 |
| F7CC56 | Rb1cc1 | RB1-inducible coiled-coil protein 1 (Fragment) | -0.094327068 | 1.850491891 |
| F7CHQ7 | Nop56 | Nucleolar protein 56 (Fragment) | -0.111877823 | 1.20169274 |
| F7CPX0 | Sgip1 | Endophilin-3-interacting protein (Fragment) | -1.387434419 | 0.184227626 |

| | | | | |
|---|---|---|---|---|
| F7CVQ1 | Ntng1 | Netrin-G1 | 1.09326992 | 2.508038362 |
| F7CZ64 | Cacng8 | Voltage-dependent calcium channel gamma-8 subunit | 0.593952847 | 1.288011154 |
| F7DBQ0 | Pdia6 | Protein disulfide-isomerase A6 | 0.350833448 | 0.952510834 |
| F7DE82 | Slirp | SRA stem-loop-interacting RNA-binding protein, mitochondrial (Fragment) | 0.658090528 | -0.157265345 |
| F8SLP9 | Pex5l | PEX5-related protein | -0.104985046 | 0.166307131 |
| F8VPK0 | Ttc37 | Tetratricopeptide repeat domain 37 | -0.174401887 | 0.307213147 |
| F8VPM7 | Erc1 | ELKS/Rab6-interacting/CAST family member 1 | 0.04486866 | 0.750863711 |
| F8VPN4 | Agl | 4-alpha-glucanotransferase | -0.406655153 | -0.28714323 |
| F8VPU2 | Farp1 | FERM, ARHGEF and pleckstrin domain-containing protein 1 | 0.101767858 | -0.00313139 |
| F8VPZ3 | Usp32 | Ubiquitinyl hydrolase 1 | 0.114364084 | 0.368168195 |
| F8VQ95 | Tacc1 | Transforming acidic coiled-coil-containing protein 1 | -0.021221129 | 0.436476549 |
| F8VQA4 | Pam | Peptidyl-glycine alpha-amidating monooxygenase | 0.208488846 | 0.496007284 |
| F8VQC1 | Srp72 | Signal recognition particle subunit SRP72 | -0.304424413 | 0.272103151 |
| F8VQE9 | Agap3 | Arf-GAP with GTPase, ANK repeat and PH domain-containing protein 3 | 0.044424597 | 0.053928057 |
| F8VQJ3 | Lamc1 | Laminin subunit gamma-1 | 0.007118766 | 1.431333303 |
| F8VQK3 | Gucy1a2 | Guanylate cyclase | -0.230126413 | 1.076542695 |
| F8VQK5 | Sash1 | SAM and SH3 domain-containing protein 1 | 0.02837162 | 0.252072175 |
| F8VQN6 | Arhgef12 | Rho guanine nucleotide exchange factor 12 | -1.191269048 | -1.144678593 |
| F8WGE7 | Txnl4a | Thioredoxin-like protein 4A (Fragment) | -1.41882995 | 0.194872379 |
| F8WGF2 | Nos1 | Constitutive NOS | -0.071640174 | 0.145501296 |
| F8WGT1 | Ahcyl2 | Putative adenosylhomocysteinase 3 | -0.332499981 | 0.256000996 |
| F8WGX0 | Fgd4 | FYVE, RhoGEF and PH domain-containing protein 4 (Fragment) | -0.07278525 | 0.841604153 |
| F8WGX5 | Dmxl1 | DmX-like protein 1 | 0.148953883 | -0.442022483 |
| F8WHB1 | Atp2b2 | Calcium-transporting ATPase | 0.328272629 | 0.403738976 |
| F8WHK6 | Phykpl | 5-phosphohydroxy-L-lysine phospho-lyase | -0.210749944 | 0.02428786 |
| F8WHL2 | Copa | Coatomer subunit alpha | -0.082553323 | 0.186122417 |
| F8WHM5 | Glg1 | Golgi apparatus protein 1 (Fragment) | -1.65264926 | 0.626550833 |
| F8WHQ1 | Tpd52 | Tumor protein D52 | 0.308770784 | 0.757246892 |
| F8WHR6 | Ddx46 | RNA helicase | 0.299473445 | 0.687530677 |
| F8WHU9 | Zpr1 | Zinc finger protein ZPR1 (Fragment) | 0.087820339 | 0.410243511 |
| F8WHW6 | Pip5k1c | Phosphatidylinositol 4-phosphate 5-kinase type-1 gamma | 0.017404175 | 0.066555659 |
| F8WI30 | Snx7 | Sorting nexin-7 | 0.642258835 | 0.032660325 |
| F8WIB1 | Arl1 | ADP-ribosylation factor-like protein 1 (Fragment) | 0.628427887 | 2.650480111 |
| F8WIE1 | Man2c1 | Alpha-mannosidase | -0.084967836 | 0.348267714 |
| F8WIE5 | Hectd1 | HECT-type E3 ubiquitin transferase | -0.138402685 | -1.609877507 |
| F8WIR1 | Ctsd | Cathepsin D | 0.223471324 | 0.85948197 |
| F8WIT2 | Anxa6 | Annexin | 0.21460104 | -0.61070776 |
| F8WIU1 | 1700037H04Rik | RIKEN cDNA 1700037H04 gene | -0.025988706 | -0.228703976 |
| F8WIV2 | Serpinb6a | Serine (or cysteine) peptidase inhibitor, clade B, member 6a | -0.05758311 | 0.38998874 |
| F8WIV5 | Dnm2 | Dynamin GTPase | -0.384056218 | -1.977491776 |
| F8WJ41 | Rps15a | 40S ribosomal protein S15a (Fragment) | 0.142113876 | 1.328332265 |
| F8WJA5 | Msi1 | RNA-binding protein Musashi homolog 1 (Fragment) | 0.861988576 | -0.978659153 |
| F8WJB9 | Evl | Ena/VASP-like protein | 0.098002561 | 0.859865824 |
| F8WJG3 | Tra2b | Transformer-2 protein homolog beta | -0.053116131 | 0.331488132 |
| F8WJI3 | Ccdc58 | Coiled-coil domain-containing protein 58 | -0.274327342 | 2.035378138 |

| | | | | |
|---|---|---|---|---|
| F8WJK8 | St13 | Hsc70-interacting protein | -0.025458272 | 0.489065806 |
| G3UVV4 | Hk1 | Hexokinase | 0.301854897 | -0.234386126 |
| G3UWD8 | Rchy1 | RING finger and CHY zinc finger domain-containing protein 1 | -0.226510398 | 1.152682145 |
| G3UWG2 | Tiam1 | T-lymphoma invasion and metastasis-inducing protein 1 | -0.370807552 | 0.642142137 |
| G3UWI9 | Sumo3 | Small ubiquitin-related modifier 3 | -0.097471428 | 0.471345266 |
| G3UWR2 | Gigyf2 | GRB10-interacting GYF protein 2 | -0.046199417 | -1.442749341 |
| G3UWV4 | Brsk2 | Serine/threonine-protein kinase BRSK2 | -0.17296985 | 0.121702353 |
| G3UX23 | Gne | Bifunctional UDP-N-acetylglucosamine 2-epimerase/N-acetylmannosamine kinase | -0.099616941 | 1.989917278 |
| G3UX48 | Prrc2a | Protein PRRC2A | -0.170355797 | 0.575384299 |
| G3UXB4 | Ctu2 | Cytoplasmic tRNA 2-thiolation protein 2 | 1.111281999 | -1.980234365 |
| G3UXL2 | Prps1l3 | Ribose-phosphate diphosphokinase | -0.053363005 | 0.122581959 |
| G3UXT7 | Fus | RNA-binding protein FUS (Fragment) | -0.020847925 | -0.481365522 |
| G3UXT9 | Usp13 | Ubiquitinyl hydrolase 1 | -0.195572344 | 0.736902555 |
| G3UXW9 | Gps1 | COP9 signalosome complex subunit 1 | 0.125577291 | 0.422270457 |
| G3UXZ5 | Psme1 | Proteasome activator complex subunit 1 (Fragment) | 0.228257116 | 1.41584142 |
| G3UY42 | Pabpn1 | Polyadenylate-binding protein 2 | -0.540039126 | 0.172237714 |
| G3UYD0 | Gtf2i | General transcription factor II-I | -0.066262404 | 0.780244986 |
| G3UYJ7 | Gm20441 | Predicted gene 20441 (Fragment) | -0.094546318 | -0.241056919 |
| G3UYP0 | Ube2i | SUMO-conjugating enzyme UBC9 (Fragment) | -0.184676234 | 0.585350672 |
| G3UYU4 | Flot1 | Flotillin | 0.683275127 | 0.996164163 |
| G3UYV7 | Rps28 | 40S ribosomal protein S28 (Fragment) | 1.499044577 | 0.819938342 |
| G3UZ48 | Syncrip | Heterogeneous nuclear ribonucleoprotein Q | -0.097039 | 0.863672098 |
| G3UZI7 | Arhgap6 | Rho GTPase-activating protein 6 | 0.184787718 | 1.000618696 |
| G3UZJ2 | Map2 | Microtubule-associated protein (Fragment) | -0.229207548 | -0.782203674 |
| G3UZM4 | Cadm2 | Cell adhesion molecule 2 (Fragment) | 0.41890955 | 0.764217059 |
| G3UZY2 | Txn2 | Thioredoxin, mitochondrial (Fragment) | 0.879795218 | 0.654774984 |
| G3X8R0 | Reep5 | Receptor expression-enhancing protein | 0.397062047 | 0.29651626 |
| G3X8R1 | Kptn | KICSTOR complex protein kaptin | 0.17391669 | -0.255693674 |
| G3X8R5 | Qrich1 | Glutamine-rich protein 1 | -0.112953854 | -0.457323869 |
| G3X8R8 | Mrtfb | Myocardin-related transcription factor B | 0.65614392 | -0.113791068 |
| G3X8T2 | Zc3h18 | Zinc finger CCCH domain-containing protein 18 | -1.204280376 | -0.460995038 |
| G3X8T3 | Ctsa | Carboxypeptidase | -0.272472254 | -0.01729552 |
| G3X8T9 | Serpina3n | Serine protease inhibitor A3N | 0.001933511 | 0.125876427 |
| G3X8U3 | 2210016F16Rik | Queuosine salvage protein | -0.096162192 | -0.358395735 |
| G3X8X7 | Vps16 | Vacuolar protein sorting-associated protein 16 homolog | 0.313540236 | 0.06535387 |
| G3X8Y3 | Naa15 | N-alpha-acetyltransferase 15, NatA auxiliary subunit | 0.041456795 | 0.386132081 |
| G3X915 | Lysmd2 | LysM and putative peptidoglycan-binding domain-containing protein 2 | 0.349496619 | 0.588909467 |
| G3X920 | Armc8 | Armadillo repeat-containing protein 8 | 0.740158081 | -0.759301345 |
| G3X928 | Sec23ip | SEC23-interacting protein | -0.208772437 | 0.013458252 |
| G3X956 | Supt16 | FACT complex subunit | 0.249666595 | -0.617623965 |
| G3X977 | Itih2 | Inter-alpha-trypsin inhibitor heavy chain H2 | -0.010946814 | 2.203129818 |
| G3X9A7 | Plcxd3 | PI-PLC X domain-containing protein 3 | -0.648420111 | 2.704713186 |
| G3X9G2 | Mink1 | Misshapen-like kinase 1 | 0.051409944 | -0.037977378 |
| G3X9H5 | Htt | Huntingtin | -0.042955971 | 0.020386219 |
| G3X9H7 | Mtss1 | Protein MTSS 1 | -0.49216795 | 0.68659989 |

| | | | | |
|---|---|---|---|---|
| G3X9K3 | Arfgef1 | Brefeldin A-inhibited guanine nucleotide-exchange protein 1 | -0.237276586 | -0.17705965 |
| G3X9N3 | Pnmal2 | PNMA-like 2 | -0.652589194 | 0.447915236 |
| G3X9U9 | Fis1 | Mitochondrial fission 1 protein | 0.346890163 | 0.793976784 |
| G3X9V8 | Serpinb3a | Serine (or cysteine) peptidase inhibitor, clade B (ovalbumin), member 3A | 0.414648501 | -0.171108087 |
| G3X9X1 | Kbtbd2 | Kelch repeat and BTB (POZ) domain-containing 2 | -0.265501308 | -0.091607889 |
| G3X9Y1 | Syt3 | Synaptotagmin-3 | -0.461193816 | 0.160360336 |
| G3X9Y5 | Ube4a | Ubiquitin conjugation factor E4 A | -0.106080627 | 1.150505702 |
| G3XA18 | Naaa | Ceramidase | 0.626933575 | 0.577642759 |
| G3XA53 | Omg | Oligodendrocyte-myelin glycoprotein | 0.419486523 | 0.39916261 |
| G5DDB7 | Plcd1 | Phosphoinositide phospholipase C | 0.044658788 | -0.090770562 |
| G5E814 | Ndufa11 | Complex I-B14.7 | -0.088218451 | -0.617021084 |
| G5E829 | Atp2b1 | Plasma membrane calcium-transporting ATPase 1 | 0.255764135 | 0.151251316 |
| G5E846 | Prph | Peripherin | 0.382384109 | 0.10259374 |
| G5E866 | Sf3b1 | Splicing factor 3B subunit 1 | 0.171454652 | -0.415837606 |
| G5E884 | Pak1 | Non-specific serine/threonine protein kinase | -0.015665627 | 0.493935585 |
| G5E895 | Akr1b10 | Aldo-keto reductase family 1, member B10 (aldose reductase) | -0.158604272 | 0.430887699 |
| G5E898 | Ppl | Periplakin | 1.50958306 | -0.474616607 |
| G5E8J3 | Wdr11 | WD repeat-containing protein 11 | 0.317912833 | 1.947225094 |
| G5E8J9 | Scyl2 | SCY1-like protein 2 | -0.283631357 | 1.104673982 |
| G5E8Q0 | Lyst | Lysosomal-trafficking regulator | -0.544764996 | -0.661077976 |
| G5E8Q4 | Cyth3 | Cytohesin-3 | 0.129953957 | 0.810625394 |
| G5E8R4 | Ppp6r3 | Serine/threonine-protein phosphatase 6 regulatory subunit 3 | -0.084940402 | 0.030444145 |
| G5E8R8 | Ubxn7 | UBX domain-containing protein 7 | 0.105128638 | 0.360200405 |
| G5E8T3 | Srp19 | Signal recognition particle 19 kDa protein | 0.297460874 | 1.009312948 |
| G5E8T6 | Magi3 | Membrane-associated guanylate kinase inverted 3 | 0.404620361 | 0.93717885 |
| G5E8T9 | Hagh | Hydroxyacylglutathione hydrolase, mitochondrial | -0.003566106 | -0.022252401 |
| G5E8Z3 | 2310050C09Rik | RIKEN cDNA 2310050C09 gene | 0.176428127 | 1.737647216 |
| G5E902 | Slc25a3 | Phosphate carrier protein, mitochondrial | 0.047842248 | 0.385863622 |
| G5E924 | Hnrnpl | Heterogeneous nuclear ribonucleoprotein L (Fragment) | 0.039087327 | -0.688585599 |
| H3BJ07 | Golph3l | Golgi phosphoprotein 3-like | -0.720842806 | 1.152881463 |
| H3BJ30 | Cpsf6 | Cleavage and polyadenylation specificity factor subunit 6 | -0.382250182 | 0.787430604 |
| H3BJB4 | Rnf141 | RING finger protein 141 | 0.172534053 | 0.295375188 |
| H3BJD6 | Ppp1r9a | Protein phosphatase 1, regulatory subunit 9A | 0.161471113 | -0.140070279 |
| H3BJH7 | Aftph | Aftiphilin | -0.691446749 | -0.466445764 |
| H3BJL6 | Esd | S-formylglutathione hydrolase | -0.237558492 | 0.428618431 |
| H3BJZ7 | Unc13a | Protein unc-13 homolog A | 0.138091214 | 0.234349092 |
| H3BK03 | Pon1 | Serum paraoxonase/arylesterase 1 (Fragment) | 0.169421546 | 0.171824932 |
| H3BKM2 | Zc3hc1 | Nuclear-interacting partner of ALK | -0.022427622 | 0.288799922 |
| H3BKQ3 | Evi5l | Ecotropic viral integration site 5-like | 0.05077076 | 0.22687006 |
| H3BL05 | Zfp428 | Zinc finger protein 428 (Fragment) | 1.022586759 | 2.200058142 |
| H3BLJ3 | Miga2 | Mitoguardin 2 (Fragment) | -1.375228691 | 0.320864201 |
| H3BLL2 | Atpaf1 | ATP synthase mitochondrial F1 complex assembly factor 1 | -0.080700175 | 0.579723199 |
| H7BWY4 | Dlg1 | Disks large homolog 1 | -0.022384548 | 0.23075374 |
| H7BWY8 | Tsr2 | Pre-rRNA-processing protein TSR2 homolog | 0.522621314 | 0.338652929 |
| H7BX26 | Cep170 | Centrosomal protein of 170 kDa | 0.154240767 | 0.188666821 |

| | | | | |
|---|---|---|---|---|
| H7BX88 | Crat | Carnitine O-acetyltransferase | 0.710173448 | -0.4589715 |
| H7BX95 | Srsf1 | Serine/arginine-rich splicing factor 1 | 0.024978447 | 0.067883492 |
| H9KUZ8 | Cep41 | Centrosomal protein of 41 kDa | -0.459155305 | 0.11476469 |
| I3ITR1 | AK157302 | cDNA sequence AK157302 | -0.051919079 | 0.531265418 |
| I7HJQ9 | Mtmr1 | Phosphatidylinositol-3,5-bisphosphate 3-phosphatase | 0.336324469 | -1.353881995 |
| I7HLV2 | Rpl10 | 60S ribosomal protein L10 (Fragment) | 0.915041637 | 1.44648091 |
| J3JS94 | Lage3 | EKC/KEOPS complex subunit Lage3 | -0.711678569 | 3.127202431 |
| J3KMQ6 | 5730455P16Rik | RIKEN cDNA 5730455P16 gene | -1.593260829 | 0.414651473 |
| J3QMG3 | Vdac3 | Voltage-dependent anion-selective channel protein 3 | 0.40981973 | 1.061455727 |
| J3QMM7 | Naxd | ATP-dependent (S)-NAD(P)H-hydrate dehydratase | -0.155425898 | 0.445957661 |
| J3QN31 | Adssl1 | Adenylosuccinate synthetase isozyme 1 | 0.154910215 | 0.255498091 |
| J3QN89 | Aamp | Angio-associated migratory protein | 0.253427696 | 0.510442734 |
| J3QNT7 | Epn2 | Epsin-2 | -0.089545441 | 0.302649816 |
| J3QNU6 | Arrb1 | Beta-arrestin-1 | -0.153336747 | 0.142257849 |
| J3QNW4 | Trappc13 | Trafficking protein particle complex subunit 13 | 0.082701937 | 0.197864532 |
| J3QP41 | Creg1 | Protein CREG1 | 0.327233251 | 0.162865321 |
| J3QP71 | Bsg | Basigin (Fragment) | 0.681484127 | 0.429511547 |
| J3QQ13 | Tnnt2 | Troponin T, cardiac muscle | 0.633075428 | 2.101530234 |
| K3W4R2 | Myh14 | Myosin-14 | -1.115376345 | -0.883843044 |
| K3W4T3 | Atp6v0a1 | V-type proton ATPase subunit a | 0.161269124 | 0.394172668 |
| K4DI58 | Cadm3 | Cell adhesion molecule 3 | 0.117865181 | 0.382606188 |
| M0QW59 | Ksr2 | Kinase suppressor of Ras 2 | 0.115145938 | 0.438570499 |
| M0QWN7 | Usp35 | Ubiquitin-specific peptidase 35 | 0.617516994 | 1.060511986 |
| O08529 | Capn2 | Calpain-2 catalytic subunit | -0.014859613 | 0.371307691 |
| O08539 | Bin1 | Myc box-dependent-interacting protein 1 | 0.076032575 | 0.230254491 |
| O08547 | Sec22b | Vesicle-trafficking protein SEC22b | 0.158684063 | 0.902941704 |
| O08553 | Dpysl2 | Dihydropyrimidinase-related protein 2 | -0.466673597 | 0.17599233 |
| O08576 | Rundc3a | RUN domain-containing protein 3A | -0.163488515 | -0.130143166 |
| O08582 | Gtpbp1 | GTP-binding protein 1 | 0.802128474 | 0.722802083 |
| O08583 | Alyref | THO complex subunit 4 | -1.330831814 | -0.533197085 |
| O08586 | Pten | Phosphatidylinositol 3,4,5-trisphosphate 3-phosphatase and dual-specificity protein phosphatase PTEN | 0.416439025 | 0.677483718 |
| O08599 | Stxbp1 | Syntaxin-binding protein 1 | 0.057635371 | 0.236503919 |
| O08648 | Map3k4 | Mitogen-activated protein kinase kinase kinase 4 | -0.264227613 | 0.752932151 |
| O08663 | Metap2 | Methionine aminopeptidase 2 | 0.332915433 | 2.185031732 |
| O08715 | Akap1 | A-kinase anchor protein 1, mitochondrial | -0.482383537 | 0.066386859 |
| O08749 | Dld | Dihydrolipoyl dehydrogenase, mitochondrial | -0.158743827 | 0.166609128 |
| O08759 | Ube3a | Ubiquitin-protein ligase E3A | 0.037498697 | 0.354602496 |
| O08795 | Prkcsh | Glucosidase 2 subunit beta | 0.029878871 | 0.211781661 |
| O08797 | Serpinb9 | SPI6 | 0.216161537 | -0.533789476 |
| O08800 | Serpinb8 | Serpin B8 | -0.98344895 | 0.146148046 |
| O08807 | Prdx4 | Peroxiredoxin-4 | 0.055832163 | 0.662142913 |
| O08842 | Gfra2 | GDNF family receptor alpha-2 | -0.161314138 | 0.465911071 |
| O08848 | RO60 | 60 kDa SS-A/Ro ribonucleoprotein | -0.095602481 | -0.122622967 |
| O08908 | Pik3r2 | Phosphatidylinositol 3-kinase regulatory subunit beta | -0.498326778 | 0.990093549 |
| O08915 | Aip | AH receptor-interacting protein | 0.050666841 | -0.115316868 |
| O08919 | Numbl | Numb-like protein | 0.103506925 | -0.106724103 |

| | | | | |
|---|---|---|---|---|
| O08992 | Sdcbp | Syntenin-1 | 0.027572441 | -0.115335941 |
| O08997 | Atox1 | Copper transport protein ATOX1 | -0.398499266 | 0.339844227 |
| O09061 | Psmb1 | Proteasome subunit beta type-1 | 0.330662251 | -0.142451127 |
| O09111 | Ndufb11 | NADH dehydrogenase [ubiquinone] 1 beta subcomplex subunit 11, mitochondrial | 0.945816898 | 1.528844039 |
| O09114 | Ptgds | Prostaglandin-H2 D-isomerase | -0.204550457 | 0.273510774 |
| O09126 | Sema4d | Semaphorin-4D | -0.430660343 | 0.70941178 |
| O09131 | Gsto1 | Glutathione S-transferase omega-1 | 0.092782593 | 0.382577896 |
| O09172 | Gclm | Glutamate--cysteine ligase regulatory subunit | -0.151321348 | 0.019010862 |
| O09174 | Amacr | Alpha-methylacyl-CoA racemase | -0.465266959 | -0.430849234 |
| O35075 | Vps26c | Vacuolar protein sorting-associated protein 26C | -0.473344962 | -0.06110096 |
| O35098 | Dpysl4 | Dihydropyrimidinase-related protein 4 | -0.247052892 | -0.037763278 |
| O35129 | Phb2 | Prohibitin-2 | 0.456246535 | 0.869860967 |
| O35136 | Ncam2 | Neural cell adhesion molecule 2 | -0.403987058 | 0.818100611 |
| O35215 | Ddt | D-dopachrome decarboxylase | -0.324197324 | 0.98827521 |
| O35226 | Psmd4 | 26S proteasome non-ATPase regulatory subunit 4 | -0.052910805 | 0.167188009 |
| O35286 | Dhx15 | Pre-mRNA-splicing factor ATP-dependent RNA helicase DHX15 | -0.3233833 | -0.023895899 |
| O35295 | Purb | Transcriptional activator protein Pur-beta | -0.009873549 | 0.048540751 |
| O35326 | Srsf5 | Serine/arginine-rich splicing factor 5 | -0.211158212 | -0.320109844 |
| O35343 | Kpna4 | Importin subunit alpha-3 | 0.086980279 | 0.710000674 |
| O35344 | Kpna3 | Importin subunit alpha-4 | -0.371232573 | -0.004330317 |
| O35345 | Kpna6 | Importin subunit alpha-7 | 0.011722279 | 0.34701093 |
| O35381 | Anp32a | Acidic leucine-rich nuclear phosphoprotein 32 family member A | -0.228367551 | 0.472676913 |
| O35393 | Efnb3 | Ephrin-B3 | -0.752737236 | 0.656757196 |
| O35405 | Pld3 | 5'-3' exonuclease PLD3 | 0.334710121 | 0.430230141 |
| O35465 | Fkbp8 | Peptidyl-prolyl cis-trans isomerase FKBP8 | -0.095804564 | 1.797583307 |
| O35526 | Stx1a | Syntaxin-1A | 0.087893105 | 0.172949155 |
| O35551 | Rabep1 | Rab GTPase-binding effector protein 1 | 0.22272288 | 0.433345159 |
| O35593 | Psmd14 | 26S proteasome non-ATPase regulatory subunit 14 | 0.249040286 | 0.654481411 |
| O35621 | Pmm1 | Phosphomannomutase 1 | -0.034969107 | 0.321457068 |
| O35633 | Slc32a1 | Vesicular inhibitory amino acid transporter | 0.03240935 | 0.598626932 |
| O35639 | Anxa3 | Annexin A3 | 0.302349822 | 0.431162198 |
| O35643 | Ap1b1 | AP-1 complex subunit beta-1 | -0.109374428 | 0.515314102 |
| O35658 | C1qbp | Complement component 1 Q subcomponent-binding protein, mitochondrial | -1.180834897 | 0.014660517 |
| O35668 | Hap1 | Huntingtin-associated protein 1 | 0.525188033 | -0.650424163 |
| O35684 | Serpini1 | Neuroserpin | -0.296295261 | -1.419798613 |
| O35685 | Nudc | Nuclear migration protein nudC | 0.018196805 | 0.387152672 |
| O35710 | Noct | Nocturnin | -2.087395206 | 0.544641177 |
| O35737 | Hnrnph1 | Heterogeneous nuclear ribonucleoprotein H | -0.256034692 | 0.491349856 |
| O35841 | Api5 | Apoptosis inhibitor 5 | -0.497258695 | -0.174554825 |
| O35857 | Timm44 | Mitochondrial import inner membrane translocase subunit TIM44 | -0.048966312 | 0.657142003 |
| O35864 | Cops5 | COP9 signalosome complex subunit 5 | 0.048963928 | 0.333380381 |
| O35887 | Calu | Calumenin | -0.194425678 | 0.170467695 |
| O35900 | Lsm2 | U6 snRNA-associated Sm-like protein LSm2 | 0.481997458 | 1.20734485 |
| O35943 | Fxn | Frataxin, mitochondrial | -0.950289027 | 0.994195779 |

| | | | | |
|---|---|---|---|---|
| O35954 | Pitpnm1 | Membrane-associated phosphatidylinositol transfer protein 1 | -0.091689301 | 0.338672002 |
| O35963 | Rab33b | Ras-related protein Rab-33B | -0.75110534 | 1.145147483 |
| O35969 | Gamt | Guanidinoacetate N-methyltransferase | 0.283497334 | 0.046528021 |
| O35988 | Sdc4 | Syndecan-4 | -0.419163545 | -0.185920715 |
| O54724 | Cavin1 | Caveolae-associated protein 1 | -0.089790471 | 0.963085334 |
| O54734 | Ddost | Dolichyl-diphosphooligosaccharide--protein glycosyltransferase 48 kDa subunit | 0.928531933 | 2.546474616 |
| O54774 | Ap3d1 | AP-3 complex subunit delta-1 | -0.544958941 | 0.505677223 |
| O54786 | Dffa | DNA fragmentation factor subunit alpha | -0.144357713 | 0.310469389 |
| O54833 | Csnk2a2 | Casein kinase II subunit alpha' | 0.058308601 | 1.489694754 |
| O54865 | Gucy1b1 | Guanylate cyclase soluble subunit beta-1 | 0.266300297 | 0.186065356 |
| O54950 | Prkag1 | 5'-AMP-activated protein kinase subunit gamma-1 | -0.379687023 | 1.404173056 |
| O54983 | Crym | Ketimine reductase mu-crystallin | 0.401785787 | 0.41679287 |
| O54984 | Get3 | ATPase GET3 | 0.058681138 | -0.001756191 |
| O54988 | Slk | STE20-like serine/threonine-protein kinase | 0.077967707 | 0.221606572 |
| O54991 | Cntnap1 | Contactin-associated protein 1 | 0.342967319 | 0.083848476 |
| O55013 | Trappc3 | Trafficking protein particle complex subunit 3 | 0.258305518 | -0.287698587 |
| O55029 | Copb2 | Coatomer subunit beta' | -0.118495909 | -0.01762565 |
| O55033 | Nck2 | Cytoplasmic protein NCK2 | 0.016269366 | 0.092126369 |
| O55042 | Snca | Alpha-synuclein | -0.390984408 | 0.443472862 |
| O55057 | Pde6d | Retinal rod rhodopsin-sensitive cGMP 3',5'-cyclic phosphodiesterase subunit delta | -0.910434469 | -0.90433979 |
| O55091 | Impact | Protein IMPACT | 0.300098991 | 0.51081419 |
| O55100 | Syngr1 | Synaptogyrin-1 | 0.796365229 | 0.564931075 |
| O55106 | Strn | Striatin | -0.000926018 | 0.138590177 |
| O55125 | Nipsnap1 | Protein NipSnap homolog 1 | -0.473043537 | 0.580135663 |
| O55126 | Nipsnap2 | Protein NipSnap homolog 2 | -0.176528454 | 0.118299166 |
| O55135 | Eif6 | Eukaryotic translation initiation factor 6 | 0.048408445 | 0.636095524 |
| O55137 | Acot1 | Acyl-coenzyme A thioesterase 1 | 0.018732897 | 0.456408978 |
| O55142 | Rpl35a | 60S ribosomal protein L35a | 0.7220987 | 0.8200442 |
| O55143 | Atp2a2 | Sarcoplasmic/endoplasmic reticulum calcium ATPase 2 | 0.11351099 | 0.375520706 |
| O55201 | Supt5h | Transcription elongation factor SPT5 | -0.175111326 | -0.162161509 |
| O55222 | Ilk | Integrin-linked protein kinase | 0.307758745 | 1.058634917 |
| O55229 | Chkb | Choline/ethanolamine kinase | -0.28044335 | 0.343409856 |
| O55234 | Psmb5 | Proteasome subunit beta type-5 | -0.014349874 | -0.35656627 |
| O70166 | Stmn3 | Stathmin-3 | 0.310775248 | 1.050722281 |
| O70172 | Pip4k2a | Phosphatidylinositol 5-phosphate 4-kinase type-2 alpha | -0.272453976 | 0.15151151 |
| O70194 | Eif3d | Eukaryotic translation initiation factor 3 subunit D | -0.219891389 | -0.265864372 |
| O70200 | Aif1 | Allograft inflammatory factor 1 | 0.217413934 | 1.262239774 |
| O70250 | Pgam2 | Phosphoglycerate mutase 2 | 0.483048662 | 0.462207953 |
| O70251 | Eef1b | Elongation factor 1-beta | 0.208835443 | 0.378860315 |
| O70252 | Hmox2 | Heme oxygenase 2 | 0.018737507 | -0.362518152 |
| O70310 | Nmt1 | Glycylpeptide N-tetradecanoyltransferase 1 | -0.022935168 | 0.315261841 |
| O70325 | Gpx4 | Phospholipid hydroperoxide glutathione peroxidase | -0.204111544 | 0.380202134 |
| O70362 | Gpld1 | Phosphatidylinositol-glycan-specific phospholipase D | 0.399417464 | -0.087665876 |
| O70378 | Emc8 | ER membrane protein complex subunit 8 | 0.431792037 | 1.16522185 |
| O70400 | Pdlim1 | PDZ and LIM domain protein 1 | -0.248276456 | 1.235264619 |

| | | | | |
|---|---|---|---|---|
| O70433 | Fhl2 | Four and a half LIM domains protein 2 | 0.017244371 | -0.156226476 |
| O70435 | Psma3 | Proteasome subunit alpha type-3 | 0.188617579 | -0.032406171 |
| O70439 | Stx7 | Syntaxin-7 | 0.673487377 | 0.887594382 |
| O70443 | Gnaz | Guanine nucleotide-binding protein G(z) subunit alpha | 0.058520031 | 2.045601686 |
| O70456 | Sfn | 14-3-3 protein sigma | 0.176348019 | -1.11411651 |
| O70481 | Ubr1 | E3 ubiquitin-protein ligase UBR1 | -0.118176238 | 0.255490303 |
| O70503 | Hsd17b12 | Very-long-chain 3-oxoacyl-CoA reductase | 0.238717779 | 0.923803171 |
| O70555 | Sprr2d | Small proline-rich protein 2D | -0.907555517 | 0.879453341 |
| O70589 | Cask | Peripheral plasma membrane protein CASK | -0.049368382 | 0.419278622 |
| O70591 | Pfdn2 | Prefoldin subunit 2 | -0.116118431 | 0.902745565 |
| O88271 | Cfdp1 | Craniofacial development protein 1 | -0.024094772 | -0.431301117 |
| O88342 | Wdr1 | WD repeat-containing protein 1 | -0.195766004 | 0.160710335 |
| O88398 | Avil | Advillin | 0.186067581 | -0.247975985 |
| O88441 | Mtx2 | Metaxin-2 | 1.415619532 | -0.120360533 |
| O88507 | Cntfr | Ciliary neurotrophic factor receptor subunit alpha | 0.30912501 | 0.592185974 |
| O88531 | Ppt1 | Palmitoyl-protein thioesterase 1 | -0.066914463 | 0.51098903 |
| O88532 | Zfr | Zinc finger RNA-binding protein | 0.101252588 | -0.004324118 |
| O88533 | Ddc | Aromatic-L-amino-acid decarboxylase | -0.037586657 | 0.387823741 |
| O88543 | Cops3 | COP9 signalosome complex subunit 3 | 0.09869868 | 0.292200724 |
| O88544 | Cops4 | COP9 signalosome complex subunit 4 | -0.123735396 | 0.279252211 |
| O88569 | Hnrnpa2b1 | Heterogeneous nuclear ribonucleoproteins A2/B1 | 0.012475109 | -0.173557917 |
| O88587 | Comt | Catechol O-methyltransferase | -0.000322342 | 0.478496393 |
| O88597 | Becn1 | Beclin-1 | -0.091322231 | 0.925404072 |
| O88685 | Psmc3 | 26S proteasome regulatory subunit 6A | 0.149187851 | 0.218552589 |
| O88696 | Clpp | ATP-dependent Clp protease proteolytic subunit, mitochondrial | -0.048175176 | 0.028381348 |
| O88712 | Ctbp1 | C-terminal-binding protein 1 | 0.034536139 | 0.388762633 |
| O88737 | Bsn | Protein bassoon | -0.05848182 | 0.026283582 |
| O88741 | Gdap1 | Ganglioside-induced differentiation-associated protein 1 | -0.192578284 | 0.334798654 |
| O88745 | Scrg1 | Scrapie-responsive protein 1 | -0.211347294 | 0.263912519 |
| O88746 | Tom1 | Target of Myb protein 1 | 0.108682982 | 0.109953562 |
| O88811 | Stam2 | Signal transducing adapter molecule 2 | 1.687392855 | 1.907828887 |
| O88834 | Shd | SH2 domain-containing adapter protein D | 0.454709085 | -0.815238953 |
| O88844 | Idh1 | Isocitrate dehydrogenase [NADP] cytoplasmic | -0.08847688 | 0.393741926 |
| O88845 | Akap10 | A-kinase anchor protein 10, mitochondrial | 0.813551617 | 0.584296385 |
| O88848 | Arl6 | ADP-ribosylation factor-like protein 6 | -0.422069867 | 0.197350661 |
| O88851 | Rbbp9 | Putative hydrolase RBBP9 | -0.176003933 | 0.645255407 |
| O88878 | Zfand5 | AN1-type zinc finger protein 5 | -0.525924269 | 0.925630967 |
| O88935 | Syn1 | Synapsin-1 | 0.023059336 | 0.042238235 |
| O88951 | Lin7b | Protein lin-7 homolog B | 0.000596809 | 0.185469945 |
| O88952 | Lin7c | Protein lin-7 homolog C | 0.058631961 | 0.491868655 |
| O88958 | Gnpda1 | Glucosamine-6-phosphate isomerase 1 | 0.295549806 | 0.56428194 |
| O88986 | Gcat | 2-amino-3-ketobutyrate coenzyme A ligase, mitochondrial | -0.441506608 | -0.805564721 |
| O89017 | Lgmn | Legumain | 0.385917473 | 0.311059157 |
| O89020 | Afm | Afamin | -0.123675028 | 1.388503551 |
| O89023 | Tpp1 | Tripeptidyl-peptidase 1 | -0.289880117 | 0.378053983 |

| | | | | |
|---|---|---|---|---|
| O89032 | Sh3pxd2a | SH3 and PX domain-containing protein 2A | 0.231120682 | 1.602313201 |
| O89053 | Coro1a | Coronin-1A | -0.139536222 | -0.059984525 |
| O89079 | Cope | Coatomer subunit epsilon | -0.460732937 | 0.288375696 |
| O89084 | Pde4a | cAMP-specific 3',5'-cyclic phosphodiesterase 4A | -0.288168271 | -0.192769845 |
| O89086 | Rbm3 | RNA-binding protein 3 | 0.495030212 | 0.998258114 |
| O89112 | Lancl1 | Glutathione S-transferase LANCL1 | 0.3843057 | -0.662811597 |
| P00158 | Mt-Cyb | Cytochrome b | -1.062253189 | -0.244059881 |
| P00375 | Dhfr | Dihydrofolate reductase | 0.164325015 | 0.823395093 |
| P00397 | Mtco1 | Cytochrome c oxidase subunit 1 | 1.65570488 | 2.339623928 |
| P00405 | Mtco2 | Cytochrome c oxidase subunit 2 | 0.918452199 | 1.238811334 |
| P00416 | mt-Co3 | Cytochrome c oxidase subunit 3 | 0.469437361 | 0.035234769 |
| P00493 | Hprt1 | Hypoxanthine-guanine phosphoribosyltransferase | -0.076236057 | 0.761370182 |
| P00687 | Amy1 | Alpha-amylase 1 | -0.063539871 | -0.950348854 |
| P00920 | Ca2 | Carbonic anhydrase 2 | -0.258575249 | -0.209691366 |
| P01027 | C3 | Complement C3 | -0.193345928 | -0.028371334 |
| P01029 | C4b | Complement C4-B | 0.36676089 | 1.458433549 |
| P01786 | | Ig heavy chain V region MOPC 47A | 1.060600662 | 1.174137115 |
| P01887 | B2m | Beta-2-microglobulin | 1.605828667 | 0.439097563 |
| P01898 | H2-Q10 | H-2 class I histocompatibility antigen, Q10 alpha chain | 1.041025702 | 0.312010765 |
| P02089 | Hbb-b2 | Hemoglobin subunit beta-2 | 1.48904988 | 3.187512398 |
| P02104 | Hbb-y | Hemoglobin subunit epsilon-Y2 | -0.528885365 | -0.225135962 |
| P02798 | Mt2 | Metallothionein-2 | 0.728082848 | 1.003993034 |
| P02802 | Mt1 | Metallothionein-1 | -0.499295203 | 0.917991797 |
| P03899 | Mtnd3 | NADH-ubiquinone oxidoreductase chain 3 | -1.09338665 | 0.62982591 |
| P03921 | Mtnd5 | NADH-ubiquinone oxidoreductase chain 5 | -0.042243544 | 0.546857834 |
| P03953 | Cfd | Complement factor D | 0.098210684 | -0.270299911 |
| P03958 | Ada | Adenosine deaminase | -0.148935604 | -0.364040693 |
| P03995 | Gfap | Glial fibrillary acidic protein | -0.05062542 | 0.069604556 |
| P04104 | Krt1 | Keratin, type II cytoskeletal 1 | 0.26933403 | -0.887114207 |
| P04444 | Hbb-bh1 | Hemoglobin subunit beta-H1 | 2.820903095 | -0.061558723 |
| P04627 | Araf | Serine/threonine-protein kinase A-Raf | 0.601532682 | -0.389026165 |
| P04925 | Prnp | Major prion protein | 0.279282792 | 1.433331649 |
| P05063 | Aldoc | Fructose-bisphosphate aldolase C | -0.25347875 | 0.151887576 |
| P05064 | Aldoa | Fructose-bisphosphate aldolase A | -0.171332932 | -0.13119634 |
| P05132 | Prkaca | cAMP-dependent protein kinase catalytic subunit alpha | 0.224732812 | -0.548138777 |
| P05201 | Got1 | Aspartate aminotransferase, cytoplasmic | -0.197146161 | 0.25625515 |
| P05202 | Got2 | Aspartate aminotransferase, mitochondrial | -0.09900767 | 0.183950742 |
| P05214 | Tuba3a | Tubulin alpha-3 chain | -0.28222688 | 1.218914032 |
| P05480 | Src | Neuronal proto-oncogene tyrosine-protein kinase Src | -0.049893475 | 0.246500015 |
| P05784 | Krt18 | Keratin, type I cytoskeletal 18 | 0.155745379 | 1.39228646 |
| P06151 | Ldha | L-lactate dehydrogenase A chain | -0.078173383 | 0.227469444 |
| P06728 | Apoa4 | Apolipoprotein A-IV | 0.589347967 | 0.182445983 |
| P06745 | Gpi | Glucose-6-phosphate isomerase | -0.146194712 | 0.251792272 |
| P06797 | Ctsl | Procathepsin L | -0.01719141 | 0.285837332 |
| P06801 | Me1 | NADP-dependent malic enzyme | -0.094173845 | -0.032996178 |
| P06837 | Gap43 | Neuromodulin | 0.226011626 | 0.629522006 |

| | | | | |
|---|---|---|---|---|
| P07309 | Ttr | Transthyretin | 0.085966492 | 0.397304058 |
| P07310 | Ckm | Creatine kinase M-type | -0.128106689 | 0.523180008 |
| P07356 | Anxa2 | Annexin A2 | 0.394739056 | 0.094103018 |
| P07724 | Alb | Albumin | -0.272167587 | 0.254073461 |
| P07742 | Rrm1 | Ribonucleoside-diphosphate reductase large subunit | 0.235034037 | 1.389417861 |
| P07744 | Krt4 | Keratin, type II cytoskeletal 4 | 2.035231113 | -1.550142288 |
| P07901 | Hsp90aa1 | Heat shock protein HSP 90-alpha | -0.021337255 | 0.410859744 |
| P07934 | Phkg1 | Phosphorylase b kinase gamma catalytic chain, skeletal muscle/heart isoform | -0.163131015 | -0.487791379 |
| P08030 | Aprt | Adenine phosphoribosyltransferase | 0.376563613 | 1.036896706 |
| P08113 | Hsp90b1 | Endoplasmin | 0.174792862 | 0.484453201 |
| P08122 | Col4a2 | Collagen alpha-2(IV) chain | 1.39534982 | 1.627841314 |
| P08226 | Apoe | Apolipoprotein E | 0.063515727 | 0.158571561 |
| P08228 | Sod1 | Superoxide dismutase [Cu-Zn] | -0.451693789 | 0.654221535 |
| P08249 | Mdh2 | Malate dehydrogenase, mitochondrial | -0.157092603 | 0.402814547 |
| P08414 | Camk4 | Calcium/calmodulin-dependent protein kinase type IV | -0.046240393 | -0.093536854 |
| P08551 | Nefl | Neurofilament light polypeptide | -0.130083402 | -0.21456782 |
| P08730 | Krt13 | Keratin, type I cytoskeletal 13 | -0.043874804 | -0.975160599 |
| P08752 | Gnai2 | Guanine nucleotide-binding protein G(i) subunit alpha-2 | 0.048349222 | 1.157387575 |
| P09103 | P4hb | Protein disulfide-isomerase | 0.122871526 | 0.390088399 |
| P09405 | Ncl | Nucleolin | -0.566867574 | 0.088680585 |
| P09411 | Pgk1 | Phosphoglycerate kinase 1 | -0.238711421 | 0.271891594 |
| P09470 | Ace | Angiotensin-converting enzyme | -0.479949411 | -0.068541686 |
| P09528 | Fth1 | Ferritin heavy chain | 0.010670058 | -0.912453175 |
| P09581 | Csf1r | Macrophage colony-stimulating factor 1 receptor | -0.548030059 | -0.245887518 |
| P09671 | Sod2 | Superoxide dismutase [Mn], mitochondrial | 0.050869942 | 0.588026365 |
| P0C027 | Nudt10 | Diphosphoinositol polyphosphate phosphohydrolase 3-alpha | 0.061667919 | 0.620011012 |
| P0C0S6 | H2az1 | Histone H2A.Z | -0.462993876 | 1.046596527 |
| P0C605 | Prkg1 | cGMP-dependent protein kinase 1 | -0.424259504 | 0.218149503 |
| P0DP26 | Calm1 | Calmodulin-1 | -0.216308784 | -0.254835447 |
| P10107 | Anxa1 | Annexin A1 | 0.776014487 | -0.700868289 |
| P10126 | Eef1a1 | Elongation factor 1-alpha 1 | 0.141205088 | 0.553596179 |
| P10493 | Nid1 | Nidogen-1 | 1.04450806 | 0.755716801 |
| P10518 | Alad | Delta-aminolevulinic acid dehydratase | -0.013751888 | 0.164219538 |
| P10605 | Ctsb | Cathepsin B | 0.117571068 | 0.279996077 |
| P10630 | Eif4a2 | Eukaryotic initiation factor 4A-II | -0.181097666 | 0.15656058 |
| P10639 | Txn | Thioredoxin | 0.007111422 | 0.544426282 |
| P10649 | Gstm1 | Glutathione S-transferase Mu 1 | -0.132546361 | 0.184826851 |
| P10711 | Tcea1 | Transcription elongation factor A protein 1 | -0.127805487 | 0.306819121 |
| P10833 | Rras | Ras-related protein R-Ras | -0.084652774 | 1.14826711 |
| P10852 | Slc3a2 | 4F2 cell-surface antigen heavy chain | 0.400394789 | 0.834116459 |
| P10853 | H2bc7 | Histone H2B type 1-F/J/L | 0.018522326 | 1.409386635 |
| P10922 | H1-0 | Histone H1.0 | 0.775108147 | 0.785033226 |
| P11031 | Sub1 | Activated RNA polymerase II transcriptional coactivator p15 | 0.306678518 | 1.051711559 |
| P11087 | Col1a1 | Collagen alpha-1(I) chain | -0.091184648 | 1.130838076 |
| P11352 | Gpx1 | Glutathione peroxidase 1 | -0.062326749 | 0.231468519 |

| | | | | |
|---|---|---|---|---|
| P11404 | Fabp3 | Fatty acid-binding protein, heart | 0.073978233 | 0.517574628 |
| P11438 | Lamp1 | Lysosome-associated membrane glycoprotein 1 | 0.757218965 | 0.430365086 |
| P11499 | Hsp90ab1 | Heat shock protein HSP 90-beta | 0.054619344 | 0.520529747 |
| P11679 | Krt8 | Keratin, type II cytoskeletal 8 | 0.607764117 | 2.140049775 |
| P11798 | Camk2a | Calcium/calmodulin-dependent protein kinase type II subunit alpha | 0.770886008 | -1.046689034 |
| P11881 | Itpr1 | Inositol 1,4,5-trisphosphate receptor type 1 | 0.354492283 | 0.384691397 |
| P11983 | Tcp1 | T-complex protein 1 subunit alpha | -0.035746511 | 0.211390018 |
| P12382 | Pfkl | ATP-dependent 6-phosphofructokinase, liver type | 0.020017783 | 0.49389569 |
| P12658 | Calb1 | Calbindin | -0.40813357 | 0.418479602 |
| P12787 | Cox5a | Cytochrome c oxidase subunit 5A, mitochondrial | 0.998534234 | 0.540482998 |
| P12815 | Pdcd6 | Programmed cell death protein 6 | -0.190123431 | 0.174988588 |
| P12849 | Prkar1b | cAMP-dependent protein kinase type I-beta regulatory subunit | -0.069717439 | -0.521846771 |
| P12960 | Cntn1 | Contactin-1 | 0.096670214 | 0.342846553 |
| P12970 | Rpl7a | 60S ribosomal protein L7a | 0.606055482 | 1.088693778 |
| P13020 | Gsn | Gelsolin | 0.055579758 | 0.31863149 |
| P13439 | Umps | Uridine 5'-monophosphate synthase | -0.144917234 | 0.022888025 |
| P13595 | Ncam1 | Neural cell adhesion molecule 1 | 0.092027632 | 0.683717728 |
| P13634 | Ca1 | Carbonic anhydrase 1 | 0.130770683 | 0.692938169 |
| P14069 | S100a6 | Protein S100-A6 | -0.570756912 | 0.21905454 |
| P14094 | Atp1b1 | Sodium/potassium-transporting ATPase subunit beta-1 | 0.222308286 | 0.566802025 |
| P14106 | C1qb | Complement C1q subcomponent subunit B | 1.482532533 | 0.199966749 |
| P14115 | Rpl27a | 60S ribosomal protein L27a | 1.02611119 | 1.240071456 |
| P14131 | Rps16 | 40S ribosomal protein S16 | 0.096933873 | 0.548692862 |
| P14148 | Rpl7 | 60S ribosomal protein L7 | 1.149133841 | 0.633025487 |
| P14152 | Mdh1 | Malate dehydrogenase, cytoplasmic | -0.225729561 | 0.188509941 |
| P14206 | Rpsa | 40S ribosomal protein SA | 0.273303541 | 0.711375237 |
| P14211 | Calr | Calreticulin | -0.002352142 | 0.302584966 |
| P14231 | Atp1b2 | Sodium/potassium-transporting ATPase subunit beta-2 | 0.651983452 | 0.711838881 |
| P14576 | Srp54 | Signal recognition particle 54 kDa protein | -0.219420306 | 0.266967614 |
| P14602 | Hspb1 | Heat shock protein beta-1 | 0.070707544 | 0.021940549 |
| P14685 | Psmd3 | 26S proteasome non-ATPase regulatory subunit 3 | -0.017399693 | -0.451497078 |
| P14733 | Lmnb1 | Lamin-B1 | -0.171036784 | 0.06773742 |
| P14824 | Anxa6 | Annexin A6 | -0.115636571 | 0.303462346 |
| P14869 | Rplp0 | 60S acidic ribosomal protein P0 | 0.524976985 | 0.278267543 |
| P14873 | Map1b | Microtubule-associated protein 1B | 0.085874112 | 0.234330495 |
| P15105 | Glul | Glutamine synthetase | -0.214500237 | 0.324184736 |
| P15209 | Ntrk2 | BDNF/NT-3 growth factors receptor | -0.144734033 | 0.389815013 |
| P15327 | Bpgm | Bisphosphoglycerate mutase | 0.006983884 | 0.64605395 |
| P15532 | Nme1 | Nucleoside diphosphate kinase A | -0.125820669 | 0.073252996 |
| P15626 | Gstm2 | Glutathione S-transferase Mu 2 | -0.263914712 | -0.737923145 |
| P15864 | H1-2 | Histone H1.2 | 0.612891261 | 2.696428458 |
| P16014 | Chgb | Secretogranin-1 | 0.181071027 | 0.692047119 |
| P16045 | Lgals1 | Galectin-1 | -0.741420555 | 0.948304017 |
| P16054 | Prkce | Protein kinase C epsilon type | -0.062180233 | 0.152661324 |
| P16110 | Lgals3 | Galectin-3 | -0.292216523 | -1.250799497 |

| | | | | |
|---|---|---|---|---|
| P16125 | Ldhb | L-lactate dehydrogenase B chain | -0.145785332 | 0.113844872 |
| P16254 | Srp14 | Signal recognition particle 14 kDa protein | 0.062364229 | 0.167673906 |
| P16330 | Cnp | 2',3'-cyclic-nucleotide 3'-phosphodiesterase | 0.102471161 | 0.183865547 |
| P16332 | Mmut | Methylmalonyl-CoA mutase, mitochondrial | -0.294233227 | -0.261628787 |
| P16460 | Ass1 | Argininosuccinate synthase | 0.246735764 | 0.681421439 |
| P16858 | Gapdh | Glyceraldehyde-3-phosphate dehydrogenase | -0.27050368 | 0.306292852 |
| P17047 | Lamp2 | Lysosome-associated membrane glycoprotein 2 | -0.68524828 | 0.295267582 |
| P17156 | Hspa2 | Heat shock-related 70 kDa protein 2 | -0.000612577 | -0.096178373 |
| P17182 | Eno1 | Alpha-enolase | -0.239548175 | 0.312334696 |
| P17183 | Eno2 | Gamma-enolase | -0.230463028 | 0.361897469 |
| P17426 | Ap2a1 | AP-2 complex subunit alpha-1 | -0.206492106 | 0.083750725 |
| P17427 | Ap2a2 | AP-2 complex subunit alpha-2 | -0.167939409 | -0.005316257 |
| P17563 | Selenbp1 | Methanethiol oxidase | -0.230865415 | 0.367822965 |
| P17742 | Ppia | Peptidyl-prolyl cis-trans isomerase A | -0.373936208 | 0.310642878 |
| P17751 | Tpi1 | Triosephosphate isomerase | -0.225227356 | 0.026906649 |
| P17809 | Slc2a1 | Solute carrier family 2, facilitated glucose transporter member 1 | 0.728599167 | 1.753584544 |
| P17879 | Hspa1b | Heat shock 70 kDa protein 1B | 0.195159976 | 0.50360775 |
| P17897 | Lyz1 | Lysozyme C-1 | 1.007395236 | -2.48305432 |
| P17918 | Pcna | Proliferating cell nuclear antigen | -0.190964222 | 1.115056356 |
| P18654 | Rps6ka3 | Ribosomal protein S6 kinase alpha-3 | 0.062345568 | 0.409915129 |
| P18760 | Cfl1 | Cofilin-1 | -0.217456881 | 0.301573753 |
| P18872 | Gnao1 | Guanine nucleotide-binding protein G(o) subunit alpha | 0.075161489 | 0.360910416 |
| P19001 | Krt19 | Keratin, type I cytoskeletal 19 | 0.091742229 | -1.263108412 |
| P19096 | Fasn | Fatty acid synthase | -0.114468002 | -0.026019891 |
| P19123 | Tnnc1 | Troponin C, slow skeletal and cardiac muscles | 0.859370645 | 1.271205743 |
| P19157 | Gstp1 | Glutathione S-transferase P 1 | -0.374886068 | 0.496670405 |
| P19221 | F2 | Prothrombin | 1.602256648 | 0.441500187 |
| P19246 | Nefh | Neurofilament heavy polypeptide | -0.155002753 | -0.039901574 |
| P19324 | Serpinh1 | Serpin H1 | -0.171093623 | -0.351467292 |
| P19536 | Cox5b | Cytochrome c oxidase subunit 5B, mitochondrial | 1.192107614 | 1.131280422 |
| P19783 | Cox4i1 | Cytochrome c oxidase subunit 4 isoform 1, mitochondrial | 0.159947999 | 0.789436181 |
| P20029 | Hspa5 | Endoplasmic reticulum chaperone BiP | 0.286622938 | 0.400379817 |
| P20060 | Hexb | Beta-hexosaminidase subunit beta | 0.72890679 | 0.147600333 |
| P20065 | Tmsb4x | Thymosin beta-4 | 1.084512075 | 1.322585066 |
| P20108 | Prdx3 | Thioredoxin-dependent peroxide reductase, mitochondrial | -0.036036682 | 0.44028314 |
| P20152 | Vim | Vimentin | -0.218101184 | 0.155210177 |
| P20917 | Mag | Myelin-associated glycoprotein | 0.079576937 | 0.418540637 |
| P21126 | Ubl4a | Ubiquitin-like protein 4A | 0.171206411 | 0.050150712 |
| P21278 | Gna11 | Guanine nucleotide-binding protein subunit alpha-11 | 0.012762674 | 1.080791791 |
| P21279 | Gnaq | Guanine nucleotide-binding protein G(q) subunit alpha | 0.292315038 | 0.474769115 |
| P21460 | Cst3 | Cystatin-C | -0.171705659 | -0.816563288 |
| P21550 | Eno3 | Beta-enolase | 0.125034205 | 0.743846575 |
| P21614 | Gc | Vitamin D-binding protein | -0.060178439 | -0.719511986 |
| P21661 | Pcsk2 | Neuroendocrine convertase 2 | -2.177170879 | -0.321675618 |
| P21981 | Tgm2 | Protein-glutamine gamma-glutamyltransferase 2 | 0.3300608 | 1.076089859 |

| | | | | |
|---|---|---|---|---|
| P22005 | Penk | Proenkephalin-A | 0.436130492 | -0.111650467 |
| P22599 | Serpina1b | Alpha-1-antitrypsin 1-2 | -0.033304532 | 0.746984164 |
| P22892 | Ap1g1 | AP-1 complex subunit gamma-1 | -0.129015255 | 0.001831373 |
| P22907 | Hmbs | Porphobilinogen deaminase | 0.695351919 | 0.736926874 |
| P23116 | Eif3a | Eukaryotic translation initiation factor 3 subunit A | -0.126282088 | 0.101517836 |
| P23198 | Cbx3 | Chromobox protein homolog 3 | -0.047237237 | 0.500951449 |
| P23242 | Gja1 | Gap junction alpha-1 protein | 0.105532551 | 0.55119888 |
| P23492 | Pnp | Purine nucleoside phosphorylase | 0.179631329 | 0.685622374 |
| P23591 | Gfus | GDP-L-fucose synthase | -0.092822425 | 0.499201139 |
| P23780 | Glb1 | Beta-galactosidase | -0.249188932 | -0.305577119 |
| P23927 | Cryab | Alpha-crystallin B chain | -0.143526077 | 0.32831049 |
| P23953 | Ces1c | Carboxylesterase 1C | 0.230072117 | 0.645494302 |
| P24270 | Cat | Catalase | -0.062924194 | 0.103896936 |
| P24369 | Ppib | Peptidyl-prolyl cis-trans isomerase B | 0.502679634 | 1.300244967 |
| P24472 | Gsta4 | Glutathione S-transferase A4 | -0.081086985 | 0.327266375 |
| P24527 | Lta4h | Leukotriene A-4 hydrolase | -0.260644086 | 0.172283808 |
| P24529 | Th | Tyrosine 3-monooxygenase | 0.12232612 | 0.412393411 |
| P24547 | Impdh2 | Inosine-5'-monophosphate dehydrogenase 2 | 0.046660614 | 0.409602483 |
| P24549 | Aldh1a1 | Retinal dehydrogenase 1 | -0.425841395 | -0.082693577 |
| P25444 | Rps2 | 40S ribosomal protein S2 | 0.32711064 | 0.420818965 |
| P25785 | Timp2 | Metalloproteinase inhibitor 2 | -0.051107725 | 0.022005717 |
| P26039 | Tln1 | Talin-1 | 0.063393529 | 0.489793142 |
| P26040 | Ezr | Ezrin | 1.294828701 | 0.561752001 |
| P26041 | Msn | Moesin | 0.13161087 | 0.341053486 |
| P26043 | Rdx | Radixin | 0.123127143 | 0.448288282 |
| P26231 | Ctnna1 | Catenin alpha-1 | -0.464597925 | -0.229278088 |
| P26339 | Chga | Chromogranin-A | 1.294433133 | 1.065138976 |
| P26350 | Ptma | Prothymosin alpha | 0.064125061 | 0.388354301 |
| P26369 | U2af2 | Splicing factor U2AF 65 kDa subunit | -0.043479983 | 1.050822258 |
| P26443 | Glud1 | Glutamate dehydrogenase 1, mitochondrial | 0.233645376 | 0.54453214 |
| P26450 | Pik3r1 | Phosphatidylinositol 3-kinase regulatory subunit alpha | -0.064439074 | 0.393959363 |
| P26516 | Psmd7 | 26S proteasome non-ATPase regulatory subunit 7 | 0.038908164 | 0.111666997 |
| P26638 | Sars1 | Serine--tRNA ligase, cytoplasmic | -0.13530105 | 0.499912103 |
| P26645 | Marcks | Myristoylated alanine-rich C-kinase substrate | 0.150670815 | 0.357582728 |
| P26883 | Fkbp1a | Peptidyl-prolyl cis-trans isomerase FKBP1A | -0.65018177 | -1.320852598 |
| P27048 | Snrpb | Small nuclear ribonucleoprotein-associated protein B | -0.584867859 | -0.46434164 |
| P27546 | Map4 | Microtubule-associated protein 4 | -0.078099378 | 0.151052634 |
| P27601 | Gna13 | Guanine nucleotide-binding protein subunit alpha-13 | 0.965583452 | 1.378543854 |
| P27612 | Plaa | Phospholipase A-2-activating protein | 0.01916399 | 0.403204123 |
| P27659 | Rpl3 | 60S ribosomal protein L3 | 0.778887526 | 1.119597594 |
| P27671 | Rasgrf1 | Ras-specific guanine nucleotide-releasing factor 1 | 0.011080138 | 0.019239744 |
| P27773 | Pdia3 | Protein disulfide-isomerase A3 | 0.143508403 | 0.223372777 |
| P28028 | Braf | Serine/threonine-protein kinase B-raf | -0.137211196 | -0.130599181 |
| P28184 | Mt3 | Metallothionein-3 | 0.96904583 | -1.950220724 |
| P28271 | Aco1 | Cytoplasmic aconitate hydratase | -0.34306008 | -0.267443021 |
| P28352 | Apex1 | DNA-(apurinic or apyrimidinic site) endonuclease | 0.158807564 | 0.844433149 |

| | | | | |
|---|---|---|---|---|
| P28474 | Adh5 | Alcohol dehydrogenase class-3 | -0.21217982 | 0.112415791 |
| P28651 | Ca8 | Carbonic anhydrase-related protein | -0.854382642 | 0.346086184 |
| P28652 | Camk2b | Calcium/calmodulin-dependent protein kinase type II subunit beta | 0.70198892 | -0.225674152 |
| P28656 | Nap1l1 | Nucleosome assembly protein 1-like 1 | 0.275594234 | 1.28499953 |
| P28658 | Atxn10 | Ataxin-10 | -0.006185627 | 1.067092896 |
| P28663 | Napb | Beta-soluble NSF attachment protein | 0.05025959 | 0.052288373 |
| P28665 | Mug1 | Murinoglobulin-1 | -0.330721315 | 0.239472707 |
| P28667 | Marcksl1 | MARCKS-related protein | 0.176609484 | 0.295095762 |
| P28738 | Kif5c | Kinesin heavy chain isoform 5C | -0.139442857 | 0.044466813 |
| P28798 | Grn | Progranulin | -1.092075125 | -0.273912589 |
| P28867 | Prkcd | Protein kinase C delta type | -0.009007422 | -0.299278736 |
| P29341 | Pabpc1 | Polyadenylate-binding protein 1 | -0.202553272 | 0.280166785 |
| P29351 | Ptpn6 | Tyrosine-protein phosphatase non-receptor type 6 | -0.129540952 | 0.646083832 |
| P29391 | Ftl1 | Ferritin light chain 1 | 0.046775309 | -0.435063998 |
| P29595 | Nedd8 | NEDD8 | 0.064294879 | 0.900621891 |
| P29699 | Ahsg | Alpha-2-HS-glycoprotein | 0.104222425 | 0.290623983 |
| P29758 | Oat | Ornithine aminotransferase, mitochondrial | -0.002786223 | 0.553077857 |
| P30275 | Ckmt1 | Creatine kinase U-type, mitochondrial | 0.366097959 | 0.17863973 |
| P30416 | Fkbp4 | Peptidyl-prolyl cis-trans isomerase FKBP4 | -0.190937964 | 0.278350512 |
| P31230 | Aimp1 | Aminoacyl tRNA synthase complex-interacting multifunctional protein 1 | -0.27844553 | 0.225539207 |
| P31324 | Prkar2b | cAMP-dependent protein kinase type II-beta regulatory subunit | 0.07519261 | 0.282753944 |
| P31648 | Slc6a1 | Sodium- and chloride-dependent GABA transporter 1 | -0.364514287 | 0.156420072 |
| P31650 | Slc6a11 | Sodium- and chloride-dependent GABA transporter 3 | 0.344874827 | 0.918220838 |
| P31750 | Akt1 | RAC-alpha serine/threonine-protein kinase | -0.051958593 | 1.312542915 |
| P31786 | Dbi | Acyl-CoA-binding protein | -0.01620849 | -1.529565096 |
| P31938 | Map2k1 | Dual specificity mitogen-activated protein kinase kinase 1 | 0.026335653 | 0.471402486 |
| P32020 | Scp2 | Sterol carrier protein 2 | 0.107705593 | 0.324877103 |
| P32067 | Ssb | Lupus La protein homolog | -0.710875893 | 0.431330045 |
| P32233 | Drg1 | Developmentally-regulated GTP-binding protein 1 | 0.881663895 | -0.578380366 |
| P32261 | Serpinc1 | Antithrombin-III | 0.181578922 | 0.423422496 |
| P32848 | Pvalb | Parvalbumin alpha | -0.339230728 | 0.008101781 |
| P32883 | Kras | GTPase KRas | -0.393134276 | 1.081197262 |
| P32921 | Wars1 | Tryptophan--tRNA ligase, cytoplasmic | -0.101203537 | 0.42104435 |
| P33146 | Cdh15 | Cadherin-15 | 0.078227647 | 0.353657881 |
| P33175 | Kif5a | Kinesin heavy chain isoform 5A | 0.054986509 | 0.282194773 |
| P34022 | Ranbp1 | Ran-specific GTPase-activating protein | -0.0728007 | 0.204844475 |
| P34057 | Rcvrn | Recoverin | 1.238245646 | 1.438652198 |
| P34884 | Mif | Macrophage migration inhibitory factor | 0.08811086 | 1.293747902 |
| P34914 | Ephx2 | Bifunctional epoxide hydrolase 2 | -0.297445838 | 0.309090455 |
| P35235 | Ptpn11 | Tyrosine-protein phosphatase non-receptor type 11 | -0.088921865 | 0.185941696 |
| P35278 | Rab5c | Ras-related protein Rab-5C | -0.156203016 | 1.204671701 |
| P35279 | Rab6a | Ras-related protein Rab-6A | -0.03951683 | 0.260578473 |
| P35282 | Rab21 | Ras-related protein Rab-21 | -0.186489773 | 0.228815238 |
| P35283 | Rab12 | Ras-related protein Rab-12 | -0.123810387 | -0.437419891 |
| P35290 | Rab24 | Ras-related protein Rab-24 | 1.283982213 | 0.732938925 |

| | | | | |
|---|---|---|---|---|
| P35486 | Pdha1 | Pyruvate dehydrogenase E1 component subunit alpha, somatic form, mitochondrial | 0.16618487 | 0.140349388 |
| P35492 | Hal | Histidine ammonia-lyase | 0.502372901 | -0.307895978 |
| P35505 | Fah | Fumarylacetoacetase | -0.21028649 | 0.586785158 |
| P35564 | Canx | Calnexin | 0.360431767 | 1.503422737 |
| P35585 | Ap1m1 | AP-1 complex subunit mu-1 | -0.112937609 | 0.251440048 |
| P35700 | Prdx1 | Peroxiredoxin-1 | 0.158147748 | 0.575284004 |
| P35762 | Cd81 | CD81 antigen | 1.092595736 | 0.847932339 |
| P35802 | Gpm6a | Neuronal membrane glycoprotein M6-a | 0.38434 | -0.511025429 |
| P35821 | Ptpn1 | Tyrosine-protein phosphatase non-receptor type 1 | -0.047255389 | 0.98409907 |
| P35979 | Rpl12 | 60S ribosomal protein L12 | 0.597351265 | 1.096646786 |
| P35980 | Rpl18 | 60S ribosomal protein L18 | 0.419060898 | 1.261842092 |
| P36552 | Cpox | Oxygen-dependent coproporphyrinogen-III oxidase, mitochondrial | 0.853947067 | -0.490911484 |
| P36916 | Gnl1 | Guanine nucleotide-binding protein-like 1 | -0.5187452 | 0.389568806 |
| P37040 | Por | NADPH--cytochrome P450 reductase | 0.575372219 | 0.879724026 |
| P37804 | Tagln | Transgelin | 0.38537426 | 0.905539513 |
| P38060 | Hmgcl | Hydroxymethylglutaryl-CoA lyase, mitochondrial | -0.076468468 | 0.301038583 |
| P38585 | Ttl | Tubulin--tyrosine ligase | 0.213660812 | 0.297920068 |
| P38647 | Hspa9 | Stress-70 protein, mitochondrial | 0.016838201 | 0.341707865 |
| P39053 | Dnm1 | Dynamin-1 | -0.232072608 | 0.303221703 |
| P39688 | Fyn | Tyrosine-protein kinase Fyn | 0.408495998 | 1.088960807 |
| P39749 | Fen1 | Flap endonuclease 1 | -0.001680628 | 1.108905792 |
| P40124 | Cap1 | Adenylyl cyclase-associated protein 1 | -0.148083814 | -0.293063482 |
| P40142 | Tkt | Transketolase | -0.141337522 | -0.006421725 |
| P40336 | Vps26a | Vacuolar protein sorting-associated protein 26A | 0.011759154 | 0.333348751 |
| P40936 | Inmt | Indolethylamine N-methyltransferase | -0.08762153 | 1.566632271 |
| P41105 | Rpl28 | 60S ribosomal protein L28 | 0.014866225 | 0.759185632 |
| P41241 | Csk | Tyrosine-protein kinase CSK | -0.190015856 | 0.525889079 |
| P42125 | Eci1 | Enoyl-CoA delta isomerase 1, mitochondrial | -0.216682466 | 0.007529259 |
| P42208 | Septin2 | Septin-2 | -0.027833017 | 1.070644379 |
| P42227 | Stat3 | Signal transducer and activator of transcription 3 | 0.269219176 | 1.317947149 |
| P42232 | Stat5b | Signal transducer and activator of transcription 5B | -0.342261251 | 0.297279199 |
| P42567 | Eps15 | Epidermal growth factor receptor substrate 15 | -0.105398877 | 0.635351817 |
| P42669 | Pura | Transcriptional activator protein Pur-alpha | -0.058438047 | 0.067533493 |
| P42932 | Cct8 | T-complex protein 1 subunit theta | 0.013902219 | -0.212830067 |
| P43006 | Slc1a2 | Excitatory amino acid transporter 2 | 0.368861643 | -0.127470652 |
| P43274 | H1-4 | Histone H1.4 | 0.898000908 | 1.576734702 |
| P43276 | H1-5 | Histone H1.5 | 0.96725839 | 1.554378033 |
| P43277 | H1-3 | Histone H1.3 | 0.392421754 | 0.895791531 |
| P45376 | Akr1b1 | Aldo-keto reductase family 1 member B1 | -0.131653404 | 0.743064562 |
| P45377 | Akr1b8 | Aldose reductase-related protein 2 | 0.669391727 | 1.716828346 |
| P45591 | Cfl2 | Cofilin-2 | -0.319935735 | 0.673777739 |
| P45878 | Fkbp2 | Peptidyl-prolyl cis-trans isomerase FKBP2 | -0.106051826 | -0.439376831 |
| P45952 | Acadm | Medium-chain specific acyl-CoA dehydrogenase, mitochondrial | -0.101488304 | 0.247536659 |
| P46061 | Rangap1 | Ran GTPase-activating protein 1 | 0.015689246 | 0.22008721 |
| P46096 | Syt1 | Synaptotagmin-1 | 0.107807954 | 0.239766121 |

| | | | | |
|---|---|---|---|---|
| P46412 | Gpx3 | Glutathione peroxidase 3 | -0.198948733 | 0.625515143 |
| P46414 | Cdkn1b | Cyclin-dependent kinase inhibitor 1B | -0.319066747 | 0.521396319 |
| P46460 | Nsf | Vesicle-fusing ATPase | 0.14968605 | 0.141664505 |
| P46467 | Vps4b | Vacuolar protein sorting-associated protein 4B | -0.009470463 | 0.462083499 |
| P46471 | Psmc2 | 26S proteasome regulatory subunit 7 | -0.007952181 | 0.341905594 |
| P46638 | Rab11b | Ras-related protein Rab-11B | 0.035586166 | 0.41226991 |
| P46656 | Fdx1 | Adrenodoxin, mitochondrial | 0.039172649 | 0.108689626 |
| P46660 | Ina | Alpha-internexin | -0.001653735 | -0.079626719 |
| P46664 | Adss2 | Adenylosuccinate synthetase isozyme 2 | -0.109684658 | 0.105612596 |
| P46737 | Brcc3 | Lys-63-specific deubiquitinase BRCC36 | 0.021538194 | -0.190776348 |
| P46935 | Nedd4 | E3 ubiquitin-protein ligase NEDD4 | -0.026198451 | -0.117178917 |
| P46938 | Yap1 | Transcriptional coactivator YAP1 | -0.122632281 | 2.566259861 |
| P47199 | Cryz | Quinone oxidoreductase | -0.028555234 | 0.317243258 |
| P47708 | Rph3a | Rabphilin-3A | -0.249715328 | 0.195940177 |
| P47738 | Aldh2 | Aldehyde dehydrogenase, mitochondrial | -0.173397827 | 0.314599037 |
| P47754 | Capza2 | F-actin-capping protein subunit alpha-2 | 0.107818 | 0.720427195 |
| P47757 | Capzb | F-actin-capping protein subunit beta | -0.031655947 | 0.145799796 |
| P47791 | Gsr | Glutathione reductase, mitochondrial | -0.360549132 | 0.215143522 |
| P47809 | Map2k4 | Dual specificity mitogen-activated protein kinase kinase 4 | -0.507258797 | 0.283551216 |
| P47811 | Mapk14 | Mitogen-activated protein kinase 14 | -0.587077395 | -0.096231778 |
| P47856 | Gfpt1 | Glutamine--fructose-6-phosphate aminotransferase [isomerizing] 1 | 0.201934083 | 0.638169289 |
| P47857 | Pfkm | ATP-dependent 6-phosphofructokinase, muscle type | -0.410300954 | -0.170937856 |
| P47867 | Scg3 | Secretogranin-3 | -0.14876976 | 0.684132894 |
| P47911 | Rpl6 | 60S ribosomal protein L6 | 0.483896446 | 0.337139924 |
| P47941 | Crkl | Crk-like protein | -0.259403674 | -0.836895466 |
| P47962 | Rpl5 | 60S ribosomal protein L5 | 0.380194505 | 0.517208576 |
| P47963 | Rpl13 | 60S ribosomal protein L13 | 1.499805101 | 1.986164888 |
| P48024 | Eif1 | Eukaryotic translation initiation factor 1 | -1.13132445 | -1.140945633 |
| P48036 | Anxa5 | Annexin A5 | -0.016639646 | 0.590493838 |
| P48318 | Gad1 | Glutamate decarboxylase 1 | 0.090751266 | 0.248206774 |
| P48320 | Gad2 | Glutamate decarboxylase 2 | -0.038915316 | 0.477234523 |
| P48428 | Tbca | Tubulin-specific chaperone A | -0.184338347 | -0.02326266 |
| P48678 | Lmna | Prelamin-A/C | 0.222703552 | 0.138002714 |
| P48722 | Hspa4l | Heat shock 70 kDa protein 4L | -0.198762194 | -0.011847178 |
| P48758 | Cbr1 | Carbonyl reductase [NADPH] 1 | -0.238350105 | 0.3165404 |
| P48774 | Gstm5 | Glutathione S-transferase Mu 5 | 0.032937813 | 0.469409943 |
| P48787 | Tnni3 | Troponin I, cardiac muscle | 1.811120987 | 2.037774245 |
| P48962 | Slc25a4 | ADP/ATP translocase 1 | 0.285762914 | 0.77706178 |
| P49312 | Hnrnpa1 | Heterogeneous nuclear ribonucleoprotein A1 | 0.252800973 | 0.082933267 |
| P49442 | Inpp1 | Inositol polyphosphate 1-phosphatase | -0.054961522 | 0.366717021 |
| P49443 | Ppm1a | Protein phosphatase 1A | -0.207273706 | 0.536889076 |
| P49586 | Pcyt1a | Choline-phosphate cytidylyltransferase A | -0.022052129 | -0.093705018 |
| P49615 | Cdk5 | Cyclin-dependent-like kinase 5 | -0.179774857 | 0.160309792 |
| P49710 | Hcls1 | Hematopoietic lineage cell-specific protein | 0.549690088 | 1.305760225 |
| P49722 | Psma2 | Proteasome subunit alpha type-2 | 0.201839574 | 0.040941238 |
| P49813 | Tmod1 | Tropomodulin-1 | -0.487683582 | -0.855977535 |

| | | | | |
|---|---|---|---|---|
| P49962 | Srp9 | Signal recognition particle 9 kDa protein | 0.044632085 | 0.503127575 |
| P50114 | S100b | Protein S100-B | -1.83589886 | 0.404090643 |
| P50171 | Hsd17b8 | Estradiol 17-beta-dehydrogenase 8 | 0.034206263 | 0.480832736 |
| P50247 | Ahcy | Adenosylhomocysteinase | -0.139949544 | 0.124201139 |
| P50396 | Gdi1 | Rab GDP dissociation inhibitor alpha | -0.045584615 | 0.159646352 |
| P50428 | Arsa | Arylsulfatase A | -0.415587362 | -1.177790165 |
| P50446 | Krt6a | Keratin, type II cytoskeletal 6A | -0.195473798 | -0.505029996 |
| P50516 | Atp6v1a | V-type proton ATPase catalytic subunit A | -0.036140569 | -0.137261391 |
| P50518 | Atp6v1e1 | V-type proton ATPase subunit E 1 | 0.141691494 | -0.090609868 |
| P50543 | S100a11 | Protein S100-A11 | 0.887811184 | -0.888086478 |
| P50544 | Acadvl | Very long-chain specific acyl-CoA dehydrogenase, mitochondrial | -0.318667126 | -0.731861115 |
| P50580 | Pa2g4 | Proliferation-associated protein 2G4 | 0.281743431 | 0.428482215 |
| P50608 | Fmod | Fibromodulin | -0.537857278 | -0.760465781 |
| P51125 | Cast | Calpastatin | -0.191491 | 0.548448881 |
| P51150 | Rab7a | Ras-related protein Rab-7a | 0.005400276 | 0.785197576 |
| P51163 | Uros | Uroporphyrinogen-III synthase | 0.113428593 | 0.309820334 |
| P51175 | Ppox | Protoporphyrinogen oxidase | -0.719789314 | 0.695518653 |
| P51569 | Gla | Alpha-galactosidase A | 1.087036896 | 2.138670921 |
| P51660 | Hsd17b4 | Peroxisomal multifunctional enzyme type 2 | 0.571626091 | 0.500676791 |
| P51855 | Gss | Glutathione synthetase | -0.003509521 | 0.369015853 |
| P51859 | Hdgf | Hepatoma-derived growth factor | -0.083716456 | 0.316403389 |
| P51863 | Atp6v0d1 | V-type proton ATPase subunit d 1 | 0.4264527 | 0.114328543 |
| P51881 | Slc25a5 | ADP/ATP translocase 2 | -0.011120415 | 0.506676515 |
| P51885 | Lum | Lumican | -0.112477303 | -1.709652265 |
| P51910 | Apod | Apolipoprotein D | 0.075411956 | 1.340729713 |
| P52196 | Tst | Thiosulfate sulfurtransferase | -0.201561578 | 0.066962083 |
| P52479 | Usp10 | Ubiquitin carboxyl-terminal hydrolase 10 | 0.000752481 | 1.377751827 |
| P52480 | Pkm | Pyruvate kinase PKM | -0.184886932 | 0.079807281 |
| P52483 | Ube2e3 | Ubiquitin-conjugating enzyme E2 E3 | 0.307995669 | 0.865104357 |
| P52503 | Ndufs6 | NADH dehydrogenase [ubiquinone] iron-sulfur protein 6, mitochondrial | 0.602750683 | 0.503163179 |
| P52623 | Uck1 | Uridine-cytidine kinase 1 | 0.276776123 | 0.133011341 |
| P52760 | Rida | 2-iminobutanoate/2-iminopropanoate deaminase | -0.185997836 | 0.243335088 |
| P53612 | Rabggtb | Geranylgeranyl transferase type-2 subunit beta | -0.01567742 | 1.420616468 |
| P53808 | Pctp | Phosphatidylcholine transfer protein | -0.21800464 | -0.82976071 |
| P53810 | Pitpna | Phosphatidylinositol transfer protein alpha isoform | -0.148088646 | 0.290834109 |
| P53986 | Slc16a1 | Monocarboxylate transporter 1 | -0.403999424 | 0.707763116 |
| P53994 | Rab2a | Ras-related protein Rab-2A | 0.027988148 | 0.263031165 |
| P53996 | Cnbp | Cellular nucleic acid-binding protein | 0.273150476 | -0.668231338 |
| P54071 | Idh2 | Isocitrate dehydrogenase [NADP], mitochondrial | 0.087255446 | -0.121822198 |
| P54116 | Stom | Stomatin | -0.459009806 | 0.013067404 |
| P54227 | Stmn1 | Stathmin | -0.008194224 | 0.953364054 |
| P54726 | Rad23a | UV excision repair protein RAD23 homolog A | -0.013531653 | 0.269801776 |
| P54728 | Rad23b | UV excision repair protein RAD23 homolog B | 0.171780427 | 0.621225675 |
| P54731 | Faf1 | FAS-associated factor 1 | 0.165338707 | 0.880370299 |
| P54775 | Psmc4 | 26S proteasome regulatory subunit 6B | -0.055318642 | 0.187267145 |

| | | | | |
|---|---|---|---|---|
| P54797 | Tango2 | Transport and Golgi organization 2 homolog | -0.074138228 | 0.59731245 |
| P54818 | Galc | Galactocerebrosidase | -0.550762971 | 1.245390574 |
| P54822 | Adsl | Adenylosuccinate lyase | -0.17229023 | 0.309924285 |
| P54823 | Ddx6 | Probable ATP-dependent RNA helicase DDX6 | 0.265205606 | 0.198432287 |
| P54830 | Ptpn5 | Tyrosine-protein phosphatase non-receptor type 5 | 0.461700249 | 0.423126539 |
| P54869 | Hmgcs2 | Hydroxymethylglutaryl-CoA synthase, mitochondrial | -0.240041892 | 0.93500185 |
| P54923 | Adprh | [Protein ADP-ribosylarginine] hydrolase | 0.058438714 | 0.469220956 |
| P55065 | Pltp | Phospholipid transfer protein | 0.142017873 | 0.275019646 |
| P55097 | Ctsk | Cathepsin K | 1.60566988 | 0.991831779 |
| P55258 | Rab8a | Ras-related protein Rab-8A | 0.030383333 | -0.228921254 |
| P55264 | Adk | Adenosine kinase | -0.55595789 | 0.472474575 |
| P55288 | Cdh11 | Cadherin-11 | -1.403388278 | 0.636556466 |
| P55302 | Lrpap1 | Alpha-2-macroglobulin receptor-associated protein | 0.218422635 | 0.478736401 |
| P55821 | Stmn2 | Stathmin-2 | -0.247087097 | 0.94499286 |
| P56135 | Atp5mf | ATP synthase subunit f, mitochondrial | 0.745576382 | 1.847927729 |
| P56183 | Rrp1 | Ribosomal RNA processing protein 1 homolog A | -0.176978318 | 1.008978446 |
| P56213 | Gfer | FAD-linked sulfhydryl oxidase ALR | 0.22721777 | 0.036838849 |
| P56371 | Rab4a | Ras-related protein Rab-4A | 0.280426598 | -0.178820292 |
| P56375 | Acyp2 | Acylphosphatase-2 | -0.408092054 | 0.455189705 |
| P56376 | Acyp1 | Acylphosphatase-1 | -0.20747935 | 0.598986308 |
| P56380 | Nudt2 | Bis(5'-nucleosyl)-tetraphosphatase [asymmetrical] | -0.148934778 | 0.942891439 |
| P56382 | Atp5f1e | ATP synthase subunit epsilon, mitochondrial | 0.46221091 | 0.365076383 |
| P56387 | Dynlt3 | Dynein light chain Tctex-type 3 | 0.272730414 | -0.542213758 |
| P56389 | Cda | Cytidine deaminase | -0.052257983 | 0.558028539 |
| P56391 | Cox6b1 | Cytochrome c oxidase subunit 6B1 | 0.473774529 | 0.809196154 |
| P56399 | Usp5 | Ubiquitin carboxyl-terminal hydrolase 5 | 0.015534337 | 0.23815759 |
| P56480 | Atp5f1b | ATP synthase subunit beta, mitochondrial | 0.426562564 | 1.226523399 |
| P56546 | Ctbp2 | C-terminal-binding protein 2 | 0.209954866 | -0.136478583 |
| P56564 | Slc1a3 | Excitatory amino acid transporter 1 | -0.126003361 | -0.810631275 |
| P56565 | S100a1 | Protein S100-A1 | -0.012217347 | -1.244222959 |
| P56812 | Pdcd5 | Programmed cell death protein 5 | 0.188050111 | 0.003202438 |
| P57722 | Pcbp3 | Poly(rC)-binding protein 3 | 0.666276582 | -0.802049001 |
| P57746 | Atp6v1d | V-type proton ATPase subunit D | -0.639799754 | 0.304841042 |
| P57759 | Erp29 | Endoplasmic reticulum resident protein 29 | 0.116979472 | 0.784715335 |
| P57776 | Eef1d | Elongation factor 1-delta | -0.060974375 | -0.096258322 |
| P57780 | Actn4 | Alpha-actinin-4 | 0.015510432 | 0.218326569 |
| P57784 | Snrpa1 | U2 small nuclear ribonucleoprotein A' | 0.26517334 | -1.008704503 |
| P58044 | Idi1 | Isopentenyl-diphosphate Delta-isomerase 1 | 0.217791176 | 1.799917301 |
| P58242 | Smpdl3b | Acid sphingomyelinase-like phosphodiesterase 3b | -0.165867138 | 1.723762671 |
| P58252 | Eef2 | Elongation factor 2 | -0.09394029 | 0.211345355 |
| P58281 | Opa1 | Dynamin-like 120 kDa protein, mitochondrial | 0.483344269 | 0.697668235 |
| P58389 | Ptpa | Serine/threonine-protein phosphatase 2A activator | 0.18290844 | 0.178475857 |
| P58404 | Strn4 | Striatin-4 | -0.050718435 | 0.202103774 |
| P58802 | Tbc1d10a | TBC1 domain family member 10A | -0.014623737 | 0.67024374 |
| P58871 | Tnks1bp1 | 182 kDa tankyrase-1-binding protein | -0.219860935 | 0.121256828 |
| P59016 | Vps33b | Vacuolar protein sorting-associated protein 33B | -0.481820901 | 0.074860573 |

| | | | | |
|---|---|---|---|---|
| P59017 | Bcl2l13 | Bcl-2-like protein 13 | -0.177801259 | 0.728975455 |
| P59114 | Pcif1 | mRNA (2'-O-methyladenosine-N(6)-)-methyltransferase | -0.567098904 | -0.782572826 |
| P59235 | Nup43 | Nucleoporin Nup43 | -0.075970459 | 0.919210593 |
| P59279 | Rab2b | Ras-related protein Rab-2B | -0.183953253 | 1.601442655 |
| P59325 | Eif5 | Eukaryotic translation initiation factor 5 | -0.185359828 | 0.426108519 |
| P59913 | Pcmtd1 | Protein-L-isoaspartate O-methyltransferase domain-containing protein 1 | 0.269958973 | 0.285673936 |
| P59999 | Arpc4 | Actin-related protein 2/3 complex subunit 4 | 0.274188487 | -0.72952954 |
| P60122 | Ruvbl1 | RuvB-like 1 | 0.418455474 | 0.734179656 |
| P60202 | Plp1 | Myelin proteolipid protein | 0.712860807 | 0.296552976 |
| P60229 | Eif3e | Eukaryotic translation initiation factor 3 subunit E | 0.02771794 | -0.025025209 |
| P60335 | Pcbp1 | Poly(rC)-binding protein 1 | -0.144084104 | 0.804029783 |
| P60469 | Ppfia3 | Liprin-alpha-3 | 0.090876452 | -0.000265757 |
| P60487 | Pdxp | Pyridoxal phosphate phosphatase | -0.122932879 | 0.256248792 |
| P60521 | Gabarapl2 | Gamma-aminobutyric acid receptor-associated protein-like 2 | -0.084665076 | 0.485667388 |
| P60670 | Nploc4 | Nuclear protein localization protein 4 homolog | 0.087031714 | -0.34415102 |
| P60710 | Actb | Actin, cytoplasmic 1 | -0.326164818 | 1.858858109 |
| P60761 | Nrgn | Neurogranin | 0.279084301 | 0.488138835 |
| P60764 | Rac3 | Ras-related C3 botulinum toxin substrate 3 | -0.604249732 | 0.056379795 |
| P60766 | Cdc42 | Cell division control protein 42 homolog | -0.39229517 | 0.320702712 |
| P60824 | Cirbp | Cold-inducible RNA-binding protein | -0.076797167 | 0.278722286 |
| P60840 | Ensa | Alpha-endosulfine | -0.047452641 | 0.171151638 |
| P60843 | Eif4a1 | Eukaryotic initiation factor 4A-I | -0.187513034 | 0.665005366 |
| P60867 | Rps20 | 40S ribosomal protein S20 | 0.297008387 | -0.071541627 |
| P60879 | Snap25 | Synaptosomal-associated protein 25 | 0.315681203 | 0.294629733 |
| P61021 | Rab5b | Ras-related protein Rab-5B | 0.181056182 | 0.849778016 |
| P61027 | Rab10 | Ras-related protein Rab-10 | -0.132317702 | -0.153843562 |
| P61028 | Rab8b | Ras-related protein Rab-8B | -0.28416516 | -0.252189159 |
| P61079 | Ube2d3 | Ubiquitin-conjugating enzyme E2 D3 | 0.334514014 | -1.420389255 |
| P61080 | Ube2d1 | Ubiquitin-conjugating enzyme E2 D1 | 0.400671069 | 0.284962813 |
| P61082 | Ube2m | NEDD8-conjugating enzyme Ubc12 | 0.108084551 | 0.309713999 |
| P61087 | Ube2k | Ubiquitin-conjugating enzyme E2 K | 0.051588821 | 0.664168358 |
| P61089 | Ube2n | Ubiquitin-conjugating enzyme E2 N | 0.085430845 | 1.690537135 |
| P61148 | Fgf1 | Fibroblast growth factor 1 | -0.270627244 | 0.639441649 |
| P61161 | Actr2 | Actin-related protein 2 | 0.215026855 | 0.595470111 |
| P61164 | Actr1a | Alpha-centractin | 0.202020772 | 0.131412665 |
| P61202 | Cops2 | COP9 signalosome complex subunit 2 | -0.122612635 | -0.711322625 |
| P61205 | Arf3 | ADP-ribosylation factor 3 | 0.272550011 | 0.843400955 |
| P61222 | Abce1 | ATP-binding cassette sub-family E member 1 | -0.315137831 | 0.627274195 |
| P61226 | Rap2b | Ras-related protein Rap-2b | 0.770266533 | 0.643039862 |
| P61255 | Rpl26 | 60S ribosomal protein L26 | 0.41777935 | 1.258166472 |
| P61264 | Stx1b | Syntaxin-1B | -0.059341685 | 0.099764188 |
| P61290 | Psme3 | Proteasome activator complex subunit 3 | -0.509000333 | 0.965364456 |
| P61294 | Rab6b | Ras-related protein Rab-6B | -0.271906153 | 0.359869003 |
| P61329 | Fgf12 | Fibroblast growth factor 12 | -0.007912699 | 0.252102375 |
| P61358 | Rpl27 | 60S ribosomal protein L27 | 0.117180061 | 1.339432557 |

| | | | | |
|---|---|---|---|---|
| P61458 | Pcbd1 | Pterin-4-alpha-carbinolamine dehydratase | 0.053279654 | 0.460821311 |
| P61460 | Depdc5 | GATOR complex protein DEPDC5 | 0.262430986 | -0.297043403 |
| P61750 | Arf4 | ADP-ribosylation factor 4 | 0.71553971 | 0.588727315 |
| P61759 | Vbp1 | Prefoldin subunit 3 | 0.001532904 | 0.540903886 |
| P61922 | Abat | 4-aminobutyrate aminotransferase, mitochondrial | -0.282381058 | 0.021940231 |
| P61963 | Dcaf7 | DDB1- and CUL4-associated factor 7 | 0.119618924 | 1.527744452 |
| P61965 | Wdr5 | WD repeat-containing protein 5 | 0.834467443 | 0.801035881 |
| P61967 | Ap1s1 | AP-1 complex subunit sigma-1A | -0.03799413 | 0.101875464 |
| P61971 | Nutf2 | Nuclear transport factor 2 | -0.00487903 | 0.727166653 |
| P61982 | Ywhag | 14-3-3 protein gamma | -0.114820608 | 0.051453908 |
| P62071 | Rras2 | Ras-related protein R-Ras2 | 0.322471015 | 1.037043254 |
| P62075 | Timm13 | Mitochondrial import inner membrane translocase subunit Tim13 | -0.127559185 | 0.076064746 |
| P62082 | Rps7 | 40S ribosomal protein S7 | 0.186075274 | -0.002177238 |
| P62137 | Ppp1ca | Serine/threonine-protein phosphatase PP1-alpha catalytic subunit | 0.350765483 | -0.233231703 |
| P62141 | Ppp1cb | Serine/threonine-protein phosphatase PP1-beta catalytic subunit | -0.086302249 | 0.545049032 |
| P62192 | Psmc1 | 26S proteasome regulatory subunit 4 | 0.076989237 | 0.543699265 |
| P62196 | Psmc5 | 26S proteasome regulatory subunit 8 | -0.054511229 | 0.152372042 |
| P62242 | Rps8 | 40S ribosomal protein S8 | 1.158489927 | 1.945862929 |
| P62254 | Ube2g1 | Ubiquitin-conjugating enzyme E2 G1 | 0.638184516 | 1.158349117 |
| P62257 | Ube2h | Ubiquitin-conjugating enzyme E2 H | 0.073253727 | 0.357262293 |
| P62259 | Ywhae | 14-3-3 protein epsilon | 0.067363294 | 0.170487086 |
| P62264 | Rps14 | 40S ribosomal protein S14 | 0.258406417 | 0.250741164 |
| P62267 | Rps23 | 40S ribosomal protein S23 | -0.087960275 | 1.035814285 |
| P62274 | Rps29 | 40S ribosomal protein S29 | 1.709483989 | 1.321295659 |
| P62301 | Rps13 | 40S ribosomal protein S13 | 0.090425014 | 0.723437786 |
| P62307 | Snrpf | Small nuclear ribonucleoprotein F | 0.030179087 | 0.895471732 |
| P62309 | Snrpg | Small nuclear ribonucleoprotein G | -0.043539333 | -0.203895887 |
| P62311 | Lsm3 | U6 snRNA-associated Sm-like protein LSm3 | 0.509805918 | 0.531298955 |
| P62313 | Lsm6 | U6 snRNA-associated Sm-like protein LSm6 | 0.04002622 | 0.197492282 |
| P62315 | Snrpd1 | Small nuclear ribonucleoprotein Sm D1 | -2.220475928 | 1.543688138 |
| P62317 | Snrpd2 | Small nuclear ribonucleoprotein Sm D2 | -0.200707404 | -0.199496587 |
| P62320 | Snrpd3 | Small nuclear ribonucleoprotein Sm D3 | 0.344630686 | 1.885327339 |
| P62331 | Arf6 | ADP-ribosylation factor 6 | 0.187789822 | -0.443870385 |
| P62334 | Psmc6 | 26S proteasome regulatory subunit 10B | -0.076846822 | -0.02811257 |
| P62340 | Tbpl1 | TATA box-binding protein-like 1 | -0.802224827 | 0.010446866 |
| P62488 | Polr2g | DNA-directed RNA polymerase II subunit RPB7 | 0.386271 | -0.430011908 |
| P62627 | Dynlrb1 | Dynein light chain roadblock-type 1 | 0.292647584 | 0.884298166 |
| P62631 | Eef1a2 | Elongation factor 1-alpha 2 | 0.105867418 | 0.054453691 |
| P62702 | Rps4x | 40S ribosomal protein S4, X isoform | -0.010751184 | 0.478504181 |
| P62715 | Ppp2cb | Serine/threonine-protein phosphatase 2A catalytic subunit beta isoform | 0.546670723 | 0.990924835 |
| P62737 | Acta2 | Actin, aortic smooth muscle | -0.083696938 | 0.944019794 |
| P62743 | Ap2s1 | AP-2 complex subunit sigma | -0.198406061 | 0.821135044 |
| P62746 | Rhob | Rho-related GTP-binding protein RhoB | 0.438656076 | 0.705902576 |
| P62748 | Hpcal1 | Hippocalcin-like protein 1 | -0.457098262 | -0.547112783 |
| P62751 | Rpl23a | 60S ribosomal protein L23a | 0.48746357 | 0.488154252 |

| | | | | |
|---|---|---|---|---|
| P62754 | Rps6 | 40S ribosomal protein S6 | 0.731599617 | 1.286006133 |
| P62761 | Vsnl1 | Visinin-like protein 1 | -0.04131705 | 0.158495267 |
| P62774 | Mtpn | Myotrophin | -0.383905125 | 2.540600777 |
| P62806 | H4c1 | Histone H4 | 0.251201757 | 0.947343508 |
| P62814 | Atp6v1b2 | V-type proton ATPase subunit B, brain isoform | -0.121786372 | 0.021104813 |
| P62821 | Rab1A | Ras-related protein Rab-1A | 0.090726153 | 0.508461793 |
| P62823 | Rab3c | Ras-related protein Rab-3C | 0.046688716 | 0.533686002 |
| P62827 | Ran | GTP-binding nuclear protein Ran | -0.074483871 | 0.491100629 |
| P62830 | Rpl23 | 60S ribosomal protein L23 | 0.408380667 | 1.317312241 |
| P62835 | Rap1a | Ras-related protein Rap-1A | -0.974037933 | -0.476837715 |
| P62838 | Ube2d2 | Ubiquitin-conjugating enzyme E2 D2 | -0.142076906 | 0.000166098 |
| P62855 | Rps26 | 40S ribosomal protein S26 | 1.528068415 | 0.396918456 |
| P62869 | Elob | Elongin-B | 0.078766155 | 0.630972068 |
| P62874 | Gnb1 | Guanine nucleotide-binding protein G(I)/G(S)/G(T) subunit beta-1 | 0.14257199 | 1.100424608 |
| P62878 | Rbx1 | E3 ubiquitin-protein ligase RBX1 | 0.282500394 | 1.042888323 |
| P62880 | Gnb2 | Guanine nucleotide-binding protein G(I)/G(S)/G(T) subunit beta-2 | 0.163182799 | 0.286009789 |
| P62881 | Gnb5 | Guanine nucleotide-binding protein subunit beta-5 | -0.365905698 | -0.044260025 |
| P62889 | Rpl30 | 60S ribosomal protein L30 | 0.608768082 | 0.256514867 |
| P62897 | Cycs | Cytochrome c, somatic | 0.322011121 | 1.214943886 |
| P62908 | Rps3 | 40S ribosomal protein S3 | 0.129188983 | 0.46692721 |
| P62915 | Gtf2b | Transcription initiation factor IIB | -0.21867555 | 1.421162764 |
| P62918 | Rpl8 | 60S ribosomal protein L8 | 0.706842391 | 0.330650012 |
| P62960 | Ybx1 | Y-box-binding protein 1 | 0.872094536 | 1.382329305 |
| P62962 | Pfn1 | Profilin-1 | -0.011416753 | 1.450697899 |
| P62965 | Crabp1 | Cellular retinoic acid-binding protein 1 | 0.121514924 | 1.032618205 |
| P62983 | Rps27a | Ubiquitin-40S ribosomal protein S27a | 0.521981748 | 0.933391253 |
| P63001 | Rac1 | Ras-related C3 botulinum toxin substrate 1 | -0.135193253 | 0.459005515 |
| P63005 | Pafah1b1 | Platelet-activating factor acetylhydrolase IB subunit beta | -0.140584373 | 0.04804039 |
| P63011 | Rab3a | Ras-related protein Rab-3A | -0.017836889 | 0.464076042 |
| P63017 | Hspa8 | Heat shock cognate 71 kDa protein | 0.012470754 | 0.288576762 |
| P63028 | Tpt1 | Translationally-controlled tumor protein | -0.051660887 | 0.284594218 |
| P63034 | Cyth2 | Cytohesin-2 | -0.160407098 | -0.68142271 |
| P63037 | Dnaja1 | DnaJ homolog subfamily A member 1 | -0.011120923 | 0.679514726 |
| P63038 | Hspd1 | 60 kDa heat shock protein, mitochondrial | -0.062397321 | 0.498402278 |
| P63040 | Cplx1 | Complexin-1 | 0.088062541 | 0.423443476 |
| P63046 | Sult4a1 | Sulfotransferase 4A1 | 0.048851776 | -0.439416726 |
| P63073 | Eif4e | Eukaryotic translation initiation factor 4E | 0.000813707 | 0.24024264 |
| P63085 | Mapk1 | Mitogen-activated protein kinase 1 | -0.107650503 | 0.395804723 |
| P63087 | Ppp1cc | Serine/threonine-protein phosphatase PP1-gamma catalytic subunit | 0.501543172 | 0.506887595 |
| P63089 | Ptn | Pleiotrophin | -0.246253045 | 0.204202493 |
| P63101 | Ywhaz | 14-3-3 protein zeta/delta | -0.066747856 | 0.274239222 |
| P63166 | Sumo1 | Small ubiquitin-related modifier 1 | -0.100358518 | 0.200290839 |
| P63168 | Dynll1 | Dynein light chain 1, cytoplasmic | -0.114503384 | -0.545798302 |
| P63248 | Pkia | cAMP-dependent protein kinase inhibitor alpha | -0.293442059 | 2.603204042 |
| P63260 | Actg1 | Actin, cytoplasmic 2 | -0.193407885 | 0.320606232 |

| | | | | |
|---|---|---|---|---|
| P63271 | Supt4h1a | Transcription elongation factor SPT4-A | -0.244142596 | 2.583221118 |
| P63276 | Rps17 | 40S ribosomal protein S17 | 0.626560656 | -0.288274924 |
| P63318 | Prkcg | Protein kinase C gamma type | -0.27020003 | -0.091176669 |
| P63321 | Rala | Ras-related protein Ral-A | -0.15093689 | 0.839944998 |
| P63325 | Rps10 | 40S ribosomal protein S10 | 0.148390102 | 0.238096078 |
| P63328 | Ppp3ca | Serine/threonine-protein phosphatase 2B catalytic subunit alpha isoform | 0.084114647 | 0.124356906 |
| P63330 | Ppp2ca | Serine/threonine-protein phosphatase 2A catalytic subunit alpha isoform | -0.107819303 | 0.383779526 |
| P67778 | Phb | Prohibitin | 0.73256712 | 1.110993862 |
| P67871 | Csnk2b | Casein kinase II subunit beta | -0.029492633 | 0.413680871 |
| P67984 | Rpl22 | 60S ribosomal protein L22 | 0.710328007 | 1.483055909 |
| P68040 | Rack1 | Receptor of activated protein C kinase 1 | 0.135279274 | 0.458945274 |
| P68181 | Prkacb | cAMP-dependent protein kinase catalytic subunit beta | 0.116043154 | -0.046500842 |
| P68254 | Ywhaq | 14-3-3 protein theta | -0.138318825 | 0.520077387 |
| P68372 | Tubb4b | Tubulin beta-4B chain | -0.243457476 | 0.670731227 |
| P68404 | Prkcb | Protein kinase C beta type | -0.166526286 | -0.137795925 |
| P68510 | Ywhah | 14-3-3 protein eta | 0.109756915 | 0.240396818 |
| P70122 | Sbds | Ribosome maturation protein SBDS | 0.346384303 | 0.529268265 |
| P70124 | Serpinb5 | Serpin B5 | 0.672485987 | -0.803449472 |
| P70158 | Smpdl3a | Acid sphingomyelinase-like phosphodiesterase 3a | -0.995065753 | 1.017706076 |
| P70168 | Kpnb1 | Importin subunit beta-1 | -0.050051435 | 0.337546984 |
| P70175 | Dlg3 | Disks large homolog 3 | -0.119460646 | 0.398260434 |
| P70188 | Kifap3 | Kinesin-associated protein 3 | -0.111930243 | -0.387784958 |
| P70195 | Psmb7 | Proteasome subunit beta type-7 | 0.245854505 | -0.298698107 |
| P70202 | Lxn | Latexin | -0.290959676 | 0.565421263 |
| P70206 | Plxna1 | Plexin-A1 | 0.440240447 | 0.237722874 |
| P70213 | Fv1 | Friend virus susceptibility protein 1 | -0.273944155 | -0.18729798 |
| P70232 | Chl1 | Neural cell adhesion molecule L1-like protein | -0.143312359 | 0.191780726 |
| P70236 | Map2k6 | Dual specificity mitogen-activated protein kinase kinase 6 | -0.048242442 | 0.126193841 |
| P70268 | Pkn1 | Serine/threonine-protein kinase N1 | 0.311504396 | 1.188934644 |
| P70271 | Pdlim4 | PDZ and LIM domain protein 4 | -0.144852638 | 0.63797156 |
| P70296 | Pebp1 | Phosphatidylethanolamine-binding protein 1 | -0.012945875 | 0.254225413 |
| P70297 | Stam | Signal transducing adapter molecule 1 | 0.009460608 | 0.36188364 |
| P70303 | Ctps2 | CTP synthase 2 | -0.945643997 | 1.448608716 |
| P70333 | Hnrnph2 | Heterogeneous nuclear ribonucleoprotein H2 | -0.316857433 | -0.148437182 |
| P70335 | Rock1 | Rho-associated protein kinase 1 | 0.194090207 | 0.038270791 |
| P70349 | Hint1 | Histidine triad nucleotide-binding protein 1 | -0.113317426 | 1.003761927 |
| P70353 | Nfyc | Nuclear transcription factor Y subunit gamma | 0.310064093 | 0.175771872 |
| P70362 | Ufd1 | Ubiquitin recognition factor in ER-associated degradation protein 1 | 0.064836756 | -0.420283635 |
| P70372 | Elavl1 | ELAV-like protein 1 | 0.355200227 | 0.840946992 |
| P70398 | Usp9x | Probable ubiquitin carboxyl-terminal hydrolase FAF-X | 0.014863141 | -0.05613025 |
| P70404 | Idh3g | Isocitrate dehydrogenase [NAD] subunit gamma 1, mitochondrial | -0.186968358 | 0.453625043 |
| P70425 | Rit2 | GTP-binding protein Rit2 | 0.223703798 | 0.126428445 |
| P70441 | Slc9a3r1 | Na(+)/H(+) exchange regulatory cofactor NHE-RF1 | -0.124572627 | 0.202847481 |
| P70444 | Bid | BH3-interacting domain death agonist | 0.178765583 | -0.15223829 |
| P70460 | Vasp | Vasodilator-stimulated phosphoprotein | -1.078110917 | -0.908931255 |

| | | | | |
|---|---|---|---|---|
| P70663 | Sparcl1 | SPARC-like protein 1 | -0.233650716 | 0.540614923 |
| P70670 | Naca | Nascent polypeptide-associated complex subunit alpha, muscle-specific form | -0.147342014 | 0.332995097 |
| P70671 | Irf3 | Interferon regulatory factor 3 | -0.012202628 | 0.284482797 |
| P70677 | Casp3 | Caspase-3 | 0.302594916 | 0.602585793 |
| P70697 | Urod | Uroporphyrinogen decarboxylase | 0.088706652 | 0.062906583 |
| P70698 | Ctps1 | CTP synthase 1 | 0.0407149 | 1.395764987 |
| P70699 | Gaa | Lysosomal alpha-glucosidase | 0.184969966 | 0.413565795 |
| P80313 | Cct7 | T-complex protein 1 subunit eta | -0.182961909 | -0.263136228 |
| P80314 | Cct2 | T-complex protein 1 subunit beta | 0.161492093 | 0.061627388 |
| P80315 | Cct4 | T-complex protein 1 subunit delta | -0.204196898 | -0.194692771 |
| P80316 | Cct5 | T-complex protein 1 subunit epsilon | -0.119109408 | -0.043001811 |
| P80318 | Cct3 | T-complex protein 1 subunit gamma | -0.194197973 | 0.019575914 |
| P80560 | Ptprn2 | Receptor-type tyrosine-protein phosphatase N2 | -0.190528234 | 0.059139411 |
| P81122 | Irs2 | Insulin receptor substrate 2 | -0.300784969 | 1.790836811 |
| P83741 | Wnk1 | Serine/threonine-protein kinase WNK1 | -0.162506485 | 0.201634089 |
| P83870 | Phf5a | PHD finger-like domain-containing protein 5A | 0.164826107 | -0.213778814 |
| P83887 | Tubg1 | Tubulin gamma-1 chain | -0.794791031 | -0.55627807 |
| P83917 | Cbx1 | Chromobox protein homolog 1 | 0.359005292 | -0.712172826 |
| P84075 | Hpca | Neuron-specific calcium-binding protein hippocalcin | -0.221969287 | 0.020348549 |
| P84084 | Arf5 | ADP-ribosylation factor 5 | -0.068305969 | 0.526763916 |
| P84086 | Cplx2 | Complexin-2 | 0.068852425 | -0.285877705 |
| P84091 | Ap2m1 | AP-2 complex subunit mu | -0.119952234 | 0.164007028 |
| P84096 | Rhog | Rho-related GTP-binding protein RhoG | 1.739169645 | 0.903624694 |
| P84104 | Srsf3 | Serine/arginine-rich splicing factor 3 | -0.1120327 | 0.22292916 |
| P85094 | Isoc2a | Isochorismatase domain-containing protein 2A | 0.309602102 | -0.053844293 |
| P97300 | Nptn | Neuroplastin | 1.355076981 | 0.079661051 |
| P97315 | Csrp1 | Cysteine and glycine-rich protein 1 | -0.268051593 | 0.327318827 |
| P97346 | Nxn | Nucleoredoxin | 0.287851683 | 0.81902345 |
| P97350 | Pkp1 | Plakophilin-1 | 0.277584521 | -1.053598722 |
| P97351 | Rps3a | 40S ribosomal protein S3a | -0.784561157 | -0.008680662 |
| P97355 | Sms | Spermine synthase | -0.288371245 | 0.225014846 |
| P97364 | Sephs2 | Selenide, water dikinase 2 | 0.02962052 | 0.607267539 |
| P97370 | Atp1b3 | Sodium/potassium-transporting ATPase subunit beta-3 | 0.358680614 | 0.630886396 |
| P97372 | Psme2 | Proteasome activator complex subunit 2 | 0.173154672 | -0.156421979 |
| P97379 | G3bp2 | Ras GTPase-activating protein-binding protein 2 | 0.095345211 | -0.236364047 |
| P97384 | Anxa11 | Annexin A11 | 0.239234734 | 0.42978255 |
| P97390 | Vps45 | Vacuolar protein sorting-associated protein 45 | 0.4498504 | 0.280612628 |
| P97427 | Crmp1 | Dihydropyrimidinase-related protein 1 | -0.055212657 | -0.026192029 |
| P97429 | Anxa4 | Annexin A4 | -0.094255606 | 0.36299324 |
| P97470 | Ppp4c | Serine/threonine-protein phosphatase 4 catalytic subunit | -0.096200975 | -0.244352818 |
| P97492 | Rgs14 | Regulator of G-protein signaling 14 | 0.664899317 | 0.36971728 |
| P97494 | Gclc | Glutamate--cysteine ligase catalytic subunit | -0.160208416 | 0.357885361 |
| P97765 | Wbp2 | WW domain-binding protein 2 | 0.265052001 | 0.477697055 |
| P97770 | Thumpd3 | THUMP domain-containing protein 3 | -0.348332659 | 0.758577824 |
| P97772 | Grm1 | Metabotropic glutamate receptor 1 | 0.34453338 | -1.110913436 |

| | | | | |
|---|---|---|---|---|
| P97797 | Sirpa | Tyrosine-protein phosphatase non-receptor type substrate 1 | 0.088338598 | 0.474825064 |
| P97807 | Fh | Fumarate hydratase, mitochondrial | -0.131367366 | 0.024573962 |
| P97820 | Map4k4 | Mitogen-activated protein kinase kinase kinase kinase 4 | 0.073131116 | 0.03234752 |
| P97823 | Lypla1 | Acyl-protein thioesterase 1 | 0.073585765 | 0.64590613 |
| P97825 | Jpt1 | Jupiter microtubule associated homolog 1 | 0.30329415 | 0.235000292 |
| P97855 | G3bp1 | Ras GTPase-activating protein-binding protein 1 | 0.118544515 | 0.635169665 |
| P97861 | Krt86 | Keratin, type II cuticular Hb6 | 0.244277604 | 0.222127597 |
| P97865 | Pex7 | Peroxisomal targeting signal 2 receptor | -1.40426 | 0.562073151 |
| P97930 | Dtymk | Thymidylate kinase | 0.092095311 | 0.615503152 |
| P97950 | Rab33a | Ras-related protein Rab-33A | 0.589786784 | 0.879936775 |
| P99024 | Tubb5 | Tubulin beta-5 chain | -0.120691299 | 1.310796738 |
| P99026 | Psmb4 | Proteasome subunit beta type-4 | 0.355834866 | 0.351157347 |
| P99027 | Rplp2 | 60S acidic ribosomal protein P2 | 0.420428181 | 2.888753017 |
| P99029 | Prdx5 | Peroxiredoxin-5, mitochondrial | -0.67273337 | 0.981354396 |
| Q00493 | Cpe | Carboxypeptidase E | 0.146933365 | 0.31060791 |
| Q00558 | F8a1 | 40-kDa huntingtin-associated protein | -0.354018847 | 1.140899181 |
| Q00560 | Il6st | Interleukin-6 receptor subunit beta | -0.887990538 | 0.515566667 |
| Q00612 | G6pdx | Glucose-6-phosphate 1-dehydrogenase X | -0.181514009 | 0.34051307 |
| Q00623 | Apoa1 | Apolipoprotein A-I | 0.440498638 | 0.298303922 |
| Q00897 | Serpina1d | Alpha-1-antitrypsin 1-4 | -0.574347973 | -0.38795344 |
| Q00898 | Serpina1e | Alpha-1-antitrypsin 1-5 | -0.721522268 | 1.348834833 |
| Q00915 | Rbp1 | Retinol-binding protein 1 | -0.722097778 | -0.236553669 |
| Q00PI9 | Hnrnpul2 | Heterogeneous nuclear ribonucleoprotein U-like protein 2 | 0.057000701 | -0.106324991 |
| Q01065 | Pde1b | Calcium/calmodulin-dependent 3',5'-cyclic nucleotide phosphodiesterase 1B | 0.083870951 | 0.158756097 |
| Q01149 | Col1a2 | Collagen alpha-2(I) chain | -0.001049169 | 1.134690046 |
| Q01768 | Nme2 | Nucleoside diphosphate kinase B | 0.038437875 | 0.196763992 |
| Q01853 | Vcp | Transitional endoplasmic reticulum ATPase | -0.028375689 | 0.023674329 |
| Q02053 | Uba1 | Ubiquitin-like modifier-activating enzyme 1 | -0.217234675 | 0.173918724 |
| Q02248 | Ctnnb1 | Catenin beta-1 | -0.117599932 | 0.303728739 |
| Q02257 | Jup | Junction plakoglobin | -0.053694884 | -0.070054531 |
| Q02566 | Myh6 | Myosin-6 | 0.655741374 | 1.853803635 |
| Q02614 | Sap30bp | SAP30-binding protein | -0.78824021 | -0.140131593 |
| Q02819 | Nucb1 | Nucleobindin-1 | -0.343046602 | 1.813287 |
| Q03137 | Epha4 | Ephrin type-A receptor 4 | 0.615823714 | -0.539937973 |
| Q03265 | Atp5f1a | ATP synthase subunit alpha, mitochondrial | 0.264060783 | 0.793463389 |
| Q03517 | Scg2 | Secretogranin-2 | 0.483873717 | 0.582992236 |
| Q04447 | Ckb | Creatine kinase B-type | -0.513795217 | 0.273057938 |
| Q04519 | Smpd1 | Sphingomyelin phosphodiesterase | 0.068945249 | -1.068977833 |
| Q04735 | Cdk16 | Cyclin-dependent kinase 16 | 0.592730077 | 0.530953089 |
| Q04736 | Yes1 | Tyrosine-protein kinase Yes | 0.059004593 | 0.411620299 |
| Q05186 | Rcn1 | Reticulocalbin-1 | 0.992077827 | 0.390999635 |
| Q05816 | Fabp5 | Fatty acid-binding protein 5 | -0.021812884 | 1.126167615 |
| Q05A62 | Dnal1 | Dynein light chain 1, axonemal | -0.114678129 | -0.005220731 |
| Q05BC3 | Eml1 | Echinoderm microtubule-associated protein-like 1 | 0.033310604 | 0.12057972 |
| Q05D44 | Eif5b | Eukaryotic translation initiation factor 5B | -0.879309877 | 1.092394193 |

| | | | | |
|---|---|---|---|---|
| Q06138 | Cab39 | Calcium-binding protein 39 | 0.115413888 | 0.484172821 |
| Q06185 | Atp5me | ATP synthase subunit e, mitochondrial | 1.488092613 | 1.350193818 |
| Q06335 | Aplp2 | Amyloid-like protein 2 | 0.056281726 | 0.551447392 |
| Q06890 | Clu | Clusterin | 0.576567078 | 0.558775425 |
| Q07235 | Serpine2 | Glia-derived nexin | -0.38305165 | -0.29273208 |
| Q07417 | Acads | Short-chain specific acyl-CoA dehydrogenase, mitochondrial | -0.564433193 | -0.133893172 |
| Q07456 | Ambp | Protein AMBP | -0.009042486 | 0.423632463 |
| Q08024 | Cbfb | Core-binding factor subunit beta | -0.060333411 | 0.510643482 |
| Q08189 | Tgm3 | Protein-glutamine gamma-glutamyltransferase E | 0.750759729 | -0.726226171 |
| Q08331 | Calb2 | Calretinin | -0.239700635 | 0.326965332 |
| Q08642 | Padi2 | Protein-arginine deiminase type-2 | -0.366161601 | -0.36520195 |
| Q08879 | Fbln1 | Fibulin-1 | -0.835966746 | 1.657803853 |
| Q08890 | Ids | Iduronate 2-sulfatase | 0.352259636 | 0.176979542 |
| Q0KK55 | Kndc1 | Kinase non-catalytic C-lobe domain-containing protein 1 | -0.491742865 | 0.107886473 |
| Q0KL01 | Ubxn2b | UBX domain-containing protein 2B | -0.143123341 | -0.620612621 |
| Q0KL02 | Trio | Triple functional domain protein | -0.120764033 | -0.274695714 |
| Q0QWG9 | Grid2ip | Delphilin | -0.844710477 | -0.425139268 |
| Q0V8T7 | Cntnap5c | Contactin-associated protein like 5-3 | -0.100957187 | 1.033930779 |
| Q0VAW6 | 8030462N17Rik | RIKEN cDNA 8030462N17 gene | -0.023114077 | 0.227754116 |
| Q0VBK2 | Krt80 | Keratin, type II cytoskeletal 80 | 0.938408883 | -0.553636551 |
| Q0VE82 | Cpne7 | Copine-7 | 0.244408703 | 0.360227585 |
| Q0VEJ0 | Cep76 | Centrosomal protein of 76 kDa | -1.043110212 | -0.474241098 |
| Q0VF59 | Dlgap2 | Disks large-associated protein 2 | 0.469916089 | 0.129291375 |
| Q0VGU4 | Vgf | Neurosecretory protein VGF | 1.04013923 | 0.789833864 |
| Q11011 | Npepps | Puromycin-sensitive aminopeptidase | -0.085638428 | 0.124757767 |
| Q11136 | Pepd | Xaa-Pro dipeptidase | 0.003742409 | 0.26828146 |
| Q149F3 | Gspt2 | Eukaryotic peptide chain release factor GTP-binding subunit ERF3B | 0.566288535 | 0.071033955 |
| Q14B01 | Rnf113a2 | Ring finger protein 113A2 | 0.121908633 | 0.416103363 |
| Q14B86 | Ltn1 | E3 ubiquitin-protein ligase listerin | -0.442992751 | -0.123310407 |
| Q14BB9 | Map6d1 | MAP6 domain-containing protein 1 | -0.307302666 | 0.110764186 |
| Q14BN8 | Rwdd2a | RWD domain-containing protein | 0.303249168 | 0.740673542 |
| Q14CH0 | Fam171b | Protein FAM171B | 0.4879378 | 0.323908329 |
| Q14CH7 | Aars2 | Alanine--tRNA ligase, mitochondrial | -0.544175784 | -0.307050387 |
| Q19LI2 | A1bg | Alpha-1B-glycoprotein | -0.703255526 | -0.428971052 |
| Q1HFZ0 | Nsun2 | RNA cytosine C(5)-methyltransferase NSUN2 | 0.339172586 | 1.224732876 |
| Q20BD0 | Hnrnpab | Heterogeneous nuclear ribonucleoprotein A/B | -0.091078091 | 1.342738469 |
| Q2M3X8 | Phactr1 | Phosphatase and actin regulator 1 | 0.123945204 | -0.406148434 |
| Q2TPA8 | Hsdl2 | Hydroxysteroid dehydrogenase-like protein 2 | -0.032976913 | 0.738257408 |
| Q32NY4 | Cnnm3 | Metal transporter CNNM3 | 0.533049615 | -1.179089546 |
| Q3B7Z2 | Osbp | Oxysterol-binding protein 1 | -0.101221434 | 0.27776893 |
| Q3SXD3 | Hddc2 | 5'-deoxynucleotidase HDDC2 | 1.394454193 | 0.581484715 |
| Q3TA40 | 6430548M08Rik | RIKEN cDNA 6430548M08 gene | -0.119063759 | 0.177993774 |
| Q3TB82 | Plekhf1 | Pleckstrin homology domain-containing family F member 1 | -2.035648982 | -1.42061011 |
| Q3TBL6 | Tnfaip8l3 | Tumor necrosis factor alpha-induced protein 8-like protein 3 | 0.826498063 | 1.93840154 |

| | | | | |
|---|---|---|---|---|
| Q3TBU7 | Agfg2 | Arf-GAP domain and FG repeat-containing protein 2 (Fragment) | -0.029671669 | -0.125158469 |
| Q3TC93 | Hs1bp3 | HCLS1-binding protein 3 | 0.013487943 | 0.438872814 |
| Q3TCD4 | Eci2 | Enoyl-CoA delta isomerase 2 | 0.125510057 | -0.319148223 |
| Q3TCH7 | Cul4a | Cullin-4A | -0.170789019 | -0.49673748 |
| Q3TCJ1 | Abraxas2 | BRISC complex subunit Abraxas 2 | -0.182358805 | 0.076465925 |
| Q3TCN2 | Plbd2 | Putative phospholipase B-like 2 | 0.161093203 | 1.244303226 |
| Q3TDD9 | Ppp1r21 | Protein phosphatase 1 regulatory subunit 21 | -0.329784584 | 0.102719307 |
| Q3TDK6 | Rogdi | Protein rogdi homolog | 0.251425457 | 0.036986351 |
| Q3TDN2 | Faf2 | FAS-associated factor 2 | -0.336998749 | 0.852300167 |
| Q3TDT0 | Trim3 | Tripartite motif-containing protein 3 | -0.397168891 | -0.729046981 |
| Q3TE40 | Rpa2 | Replication protein A 32 kDa subunit | 0.766550096 | 0.803604126 |
| Q3TES0 | Iqsec3 | IQ motif and SEC7 domain-containing protein 3 | -0.292388058 | -0.442463477 |
| Q3TFP0 | Srsf10 | Serine/arginine-rich-splicing factor 10 | -0.798009014 | -0.752335072 |
| Q3TFQ1 | Spryd7 | SPRY domain-containing protein 7 | 0.184689331 | 0.20237875 |
| Q3TGM7 | Hbs1l | HBS1-like protein | -0.243130557 | 0.69832627 |
| Q3TGS7 | Snx12 | Sorting nexin-12 | 0.069728088 | 0.827926636 |
| Q3TH34 | Dennd10 | DENN domain-containing protein 10 | 0.181554572 | 0.251578967 |
| Q3THE2 | Myl12b | Myosin regulatory light chain 12B | 0.361120796 | 0.620989323 |
| Q3THG9 | Aarsd1 | Alanyl-tRNA editing protein Aarsd1 | -0.099086062 | -0.490632216 |
| Q3THJ3 | Eif1ad | Probable RNA-binding protein EIF1AD | -0.140321509 | 1.244355043 |
| Q3THK3 | Gtf2f1 | General transcription factor IIF subunit 1 | -1.432381598 | 0.056001504 |
| Q3THK7 | Gmps | GMP synthase [glutamine-hydrolyzing] | -0.063239002 | 0.398876508 |
| Q3THS6 | Mat2a | S-adenosylmethionine synthase isoform type-2 | -0.266586622 | 0.636823813 |
| Q3TIR3 | Ric8a | Synembryn-A | 0.242648443 | 0.492080053 |
| Q3TIU4 | Pde12 | 2',5'-phosphodiesterase 12 | 0.143802993 | -0.274090131 |
| Q3TIV5 | Zc3h15 | Zinc finger CCCH domain-containing protein 15 | 0.283718904 | -0.463603973 |
| Q3TKY6 | Cwc27 | Spliceosome-associated protein CWC27 homolog | -1.255520169 | 0.512053887 |
| Q3TLK9 | Cnot7 | CCR4-NOT transcription complex subunit 7 | -0.456772105 | -1.29363958 |
| Q3TLS3 | Gdpgp1 | GDP-D-glucose phosphorylase 1 | 0.040045484 | 0.526658853 |
| Q3TLZ6 | Gatb | Glutamyl-tRNA(Gln) amidotransferase subunit B, mitochondrial | -0.058827591 | 0.501295249 |
| Q3TMH2 | Scrn3 | Secernin-3 | -0.196892166 | -0.651937326 |
| Q3TNA1 | Xylb | Xylulose kinase | 0.113940748 | 0.210469882 |
| Q3TPE9 | Ankmy2 | Ankyrin repeat and MYND domain-containing protein 2 | 0.083474731 | 1.023164272 |
| Q3TQI7 | | Telomere length and silencing protein 1 homolog | -0.437188021 | -1.049485425 |
| Q3TRJ4 | Krt26 | Keratin, type I cytoskeletal 26 | -0.752350044 | 0.848394076 |
| Q3TSA8 | Scamp1 | Secretory carrier-associated membrane protein | -0.746863461 | 0.290367762 |
| Q3TTL3 | Spata7 | SPATA7 isoform | 0.448578835 | 0.326272647 |
| Q3TTY5 | Krt2 | Keratin, type II cytoskeletal 2 epidermal | -0.123737017 | -0.441850344 |
| Q3TUQ7 | Prkaa1 | Acetyl-CoA carboxylase kinase | 0.043177128 | 0.236084779 |
| Q3TVK3 | Dnpep | Aspartyl aminopeptidase | -0.029962603 | -0.029198964 |
| Q3TW96 | Uap1l1 | UDP-N-acetylhexosamine pyrophosphorylase-like protein 1 | -0.129187902 | -0.437144121 |
| Q3TWW8 | Srsf6 | Serine/arginine-rich splicing factor 6 | -0.092206128 | 0.109816392 |
| Q3TXS7 | Psmd1 | 26S proteasome non-ATPase regulatory subunit 1 | 0.023600419 | 0.019213676 |
| Q3TXU5 | Dhps | Deoxyhypusine synthase | 0.029174868 | -0.969255924 |
| Q3TXV4 | Rab31 | Rab22B | 0.778900083 | 1.779582024 |

| | | | | |
|---|---|---|---|---|
| Q3TYD4 | Arsg | Arylsulfatase G | -0.588205338 | 0.28043143 |
| Q3TYD6 | Lmtk2 | Serine/threonine-protein kinase LMTK2 | -0.362819147 | 0.203213056 |
| Q3TYX3 | Smyd5 | SET and MYND domain-containing protein 5 | -0.022600238 | 0.797359625 |
| Q3TZ02 | Cyth1 | Cytohesin-1 | -0.206222979 | 0.01844883 |
| Q3TZT4 | Inpp5a | Inositol-polyphosphate 5-phosphatase | -0.485860093 | 0.120571136 |
| Q3U0B3 | Dhrs11 | Dehydrogenase/reductase SDR family member 11 | -0.236029847 | 0.048940818 |
| Q3U0D9 | Hace1 | E3 ubiquitin-protein ligase HACE1 | -0.306755892 | 0.237645308 |
| Q3U125 | Prxl2a | Peroxiredoxin-like 2 activated in M-CSF stimulated monocytes | -0.790282122 | 0.139913718 |
| Q3U186 | Rars2 | Probable arginine--tRNA ligase, mitochondrial | -0.767103354 | 0.366230647 |
| Q3U1F9 | Pag1 | Phosphoprotein associated with glycosphingolipid-enriched microdomains 1 | -0.561518574 | 0.543864091 |
| Q3U1J4 | Ddb1 | DNA damage-binding protein 1 | -0.098438295 | 0.362672011 |
| Q3U1V6 | Uevld | Ubiquitin-conjugating enzyme E2 variant 3 | 1.385970481 | 0.950797081 |
| Q3U276 | Sdhaf1 | Succinate dehydrogenase assembly factor 1, mitochondrial | 0.691940212 | 2.819277585 |
| Q3U2A8 | Vars2 | Valine--tRNA ligase, mitochondrial | -0.351973152 | -0.676755905 |
| Q3U2G2 | Hspa4 | Heat shock 70 kDa protein 4 | -0.064090157 | 0.109987259 |
| Q3U3J1 | Bckdha | 2-oxoisovalerate dehydrogenase subunit alpha | -0.084469986 | 0.357980569 |
| Q3U3Q1 | Ulk3 | Serine/threonine-protein kinase ULK3 | -0.164866225 | 0.843678951 |
| Q3U422 | Ndufv3 | Complex I-9kD | -1.081984552 | 0.168417295 |
| Q3U487 | Hectd3 | E3 ubiquitin-protein ligase HECTD3 | -0.513491408 | -0.210349083 |
| Q3U4F0 | Sfxn3 | Sideroflexin-3 | 1.123580424 | -0.365005175 |
| Q3U4S0 | Pank2 | Pantothenate kinase 2 | -2.402303823 | -0.151671251 |
| Q3U5F4 | Yrdc | YrdC domain-containing protein, mitochondrial | -0.190978273 | 0.556662877 |
| Q3U5Q7 | Cmpk2 | UMP-CMP kinase 2, mitochondrial | 0.318866444 | 1.005299568 |
| Q3U741 | Ddx17 | RNA helicase | 0.232795207 | 0.108191013 |
| Q3U7U3 | Fbxo7 | F-box only protein 7 | -1.238964335 | -0.162549814 |
| Q3UBG2 | Pid1 | PTB-containing, cubilin and LRP1-interacting protein | -0.391800594 | -1.785383066 |
| Q3UCV8 | Otulin | Ubiquitin thioesterase otulin | 0.157593314 | 0.80010732 |
| Q3UDD3 | Poldip3 | Polymerase delta-interacting protein 3 | -0.652666982 | 0.348596414 |
| Q3UDE2 | Ttll12 | Tubulin--tyrosine ligase-like protein 12 | -0.13829991 | 1.036218484 |
| Q3UDM0 | Mob1b | MOB kinase activator 1B | 0.005586815 | 0.028966586 |
| Q3UDP0 | Wdr41 | WD repeat-containing protein 41 | 0.100658258 | 0.36430486 |
| Q3UE37 | Ube2z | Ubiquitin-conjugating enzyme E2 Z | -0.084503841 | 0.623716036 |
| Q3UE92 | Xpnpep1 | Xaa-Pro aminopeptidase 1 | -0.040127913 | 0.344951789 |
| Q3UEB3 | Puf60 | Poly(U)-binding-splicing factor PUF60 | -0.111493619 | 0.250179768 |
| Q3UEB4 | Mvk | Mevalonate kinase | 0.171397209 | 0.099945704 |
| Q3UER8 | Fgg | Fibrinogen gamma chain | 0.211361504 | 0.966765563 |
| Q3UF75 | Parva | Alpha-parvin | -0.336162217 | -0.945709705 |
| Q3UF95 | Bag6 | BCL2-associated athanogene 6 | 0.113790385 | -0.019096057 |
| Q3UFK8 | Frmd8 | FERM domain-containing protein 8 | -0.122841263 | 0.119628112 |
| Q3UFS0 | Zyg11b | Protein zyg-11 homolog B | 0.266923491 | -0.400693099 |
| Q3UFY7 | Nt5c3b | 7-methylguanosine phosphate-specific 5'-nucleotidase | -0.21291647 | 0.423535506 |
| Q3UFY8 | Trmt10c | tRNA methyltransferase 10 homolog C | -0.933411598 | 1.081863085 |
| Q3UG98 | Nat9 | N-acetyltransferase 9 | 0.364887301 | 0.822521766 |
| Q3UGB5 | Dazap1 | DAZ-associated protein 1 | 0.039717166 | 0.213928064 |
| Q3UGC7 | Eif3j1 | Eukaryotic translation initiation factor 3 subunit J-A | 0.216396618 | 0.424779574 |

| | | | | |
|---|---|---|---|---|
| Q3UGR5 | Hdhd2 | Haloacid dehalogenase-like hydrolase domain-containing protein 2 | -0.408064842 | 0.058074156 |
| Q3UGS4 | Mcrip1 | Mapk-regulated corepressor-interacting protein 1 | 0.282719898 | -0.230889638 |
| Q3UGX2 | Sptb | Spectrin beta chain | -0.017538484 | 0.687710921 |
| Q3UGY8 | Arfgef3 | Brefeldin A-inhibited guanine nucleotide-exchange protein 3 | 0.072875595 | -0.5896945 |
| Q3UH45 | Tecpr2 | Tectonin beta-propeller repeat-containing 2 | -0.79183925 | -0.202919324 |
| Q3UH59 | Myh10 | Myosin-10 | -0.208946705 | -0.218184312 |
| Q3UH60 | Dip2b | Disco-interacting protein 2 homolog B | 0.440466913 | 0.240822474 |
| Q3UHB1 | Nt5dc3 | 5'-nucleotidase domain-containing protein 3 | -0.089696503 | -0.861685912 |
| Q3UHD1 | Adgrb1 | Adhesion G protein-coupled receptor B1 | -0.396431796 | -0.20866855 |
| Q3UHD2 | Gfod1 | Glucose-fructose oxidoreductase domain-containing protein 1 | 0.263493951 | -0.438294252 |
| Q3UHD6 | Snx27 | Sorting nexin-27 | 0.011089484 | 0.432454904 |
| Q3UHD9 | Agap2 | Arf-GAP with GTPase, ANK repeat and PH domain-containing protein 2 | -0.39040645 | -1.503403346 |
| Q3UHE1 | Pitpnm3 | Membrane-associated phosphatidylinositol transfer protein 3 | -1.608246414 | -0.381786346 |
| Q3UHG5 | Tspan7 | Tetraspanin | 0.521420193 | 0.582546711 |
| Q3UHG7 | Dennd11 | DENN domain-containing protein 11 | -0.175458368 | -0.052932739 |
| Q3UHI4 | Tmed8 | Protein TMED8 | 0.093061161 | 0.132503827 |
| Q3UHJ0 | Aak1 | AP2-associated protein kinase 1 | -0.150962162 | 0.13803641 |
| Q3UHL1 | Camkv | CaM kinase-like vesicle-associated protein | 0.258258979 | 0.51208957 |
| Q3UHX2 | Pdap1 | 28 kDa heat- and acid-stable phosphoprotein | 0.118971125 | 1.08447059 |
| Q3UI43 | Babam1 | BRISC and BRCA1-A complex member 1 | 0.183792877 | 0.146790981 |
| Q3UID4 | Prmt2 | Protein arginine N-methyltransferase 2 | 0.046450202 | 1.105227629 |
| Q3UJB0 | Sf3b2 | Splicing factor 3b, subunit 2 | 1.120556704 | 1.183805307 |
| Q3UJP5 | | Protein C8orf37 homolog | -0.057417234 | 0.202783267 |
| Q3UJU9 | Rmdn3 | Regulator of microtubule dynamics protein 3 | 0.032957586 | 0.645222505 |
| Q3UKC1 | Tax1bp1 | Tax1-binding protein 1 homolog | -0.046838474 | 0.37876606 |
| Q3UKJ7 | Smu1 | WD40 repeat-containing protein SMU1 | 0.185236963 | 0.341676474 |
| Q3ULD5 | Mccc2 | Methylcrotonoyl-CoA carboxylase beta chain, mitochondrial | 0.224406751 | 0.147016525 |
| Q3ULJ0 | Gpd1l | Glycerol-3-phosphate dehydrogenase 1-like protein | -0.188381131 | 0.07248052 |
| Q3UM45 | Ppp1r7 | Protein phosphatase 1 regulatory subunit 7 | 0.170038096 | 0.232277552 |
| Q3UMA3 | Hgs | Hepatocyte growth factor-regulated tyrosine kinase substrate | 0.038807265 | 0.319955031 |
| Q3UMB5 | Smcr8 | Guanine nucleotide exchange protein SMCR8 | 0.018030675 | 0.576981862 |
| Q3UMB9 | Washc4 | WASH complex subunit 4 | -0.027704398 | 0.20334816 |
| Q3UMG6 | Nme7 | Nucleoside diphosphate kinase 7 | 0.670742035 | -0.872821967 |
| Q3UMT1 | Ppp1r12c | Protein phosphatase 1 regulatory subunit 12C | 0.235103671 | 0.002599239 |
| Q3UMU9 | Hdgfl2 | Hepatoma-derived growth factor-related protein 2 | -0.646842289 | 0.920065403 |
| Q3UNA4 | Nxt2 | NTF2-related export protein 2 | 0.526837349 | 1.476325989 |
| Q3UNH4 | Gprin1 | G protein-regulated inducer of neurite outgrowth 1 | -0.012302907 | 0.365512371 |
| Q3UNZ8 | Cryzl2 | Quinone oxidoreductase-like protein 2 | -2.380975914 | 0.612192631 |
| Q3UPH1 | Prrc1 | Protein PRRC1 | 0.327046839 | 0.809178034 |
| Q3UPL0 | Sec31a | Protein transport protein Sec31A | -0.095709292 | -0.080723921 |
| Q3UQ44 | Iqgap2 | Ras GTPase-activating-like protein IQGAP2 | 0.735382366 | 0.16127642 |
| Q3UR70 | Tgfbrap1 | Transforming growth factor-beta receptor-associated protein 1 | -0.655873744 | -0.485376517 |
| Q3URE1 | Acsf3 | Malonate--CoA ligase ACSF3, mitochondrial | -0.165803083 | 0.062266032 |
| Q3USB7 | Plcl1 | Inactive phospholipase C-like protein 1 | -0.44700524 | 0.078433355 |

| | | | | |
|---|---|---|---|---|
| Q3USC7 | Gdap1l1 | Ganglioside-induced differentiation-associated protein 1-like 1 | -0.335959435 | 0.578817685 |
| Q3USJ8 | Fchsd2 | F-BAR and double SH3 domains protein 2 | -0.606815084 | 0.483625253 |
| Q3USR6 | Ngb | Neuroglobin | 0.651271089 | 0.590112527 |
| Q3UTZ4 | Tkfc | ATP-dependent dihydroxyacetone kinase | -0.237836361 | -0.63600413 |
| Q3UUF8 | Ankrd34b | Ankyrin repeat domain-containing protein 34B | 0.574482505 | 0.676150958 |
| Q3UUG6 | Tbc1d24 | TBC1 domain family member 24 | -0.268753719 | 0.047267278 |
| Q3UUI3 | Them4 | Acyl-coenzyme A thioesterase THEM4 | -0.119758161 | -0.691041946 |
| Q3UUJ4 | Strada | STE20-related kinase adapter protein alpha | -0.530984306 | -0.317846298 |
| Q3UV17 | Krt76 | Keratin, type II cytoskeletal 2 oral | 0.113939857 | -0.754339854 |
| Q3UVG3 | Fam91a1 | Protein FAM91A1 | -0.278489971 | 1.480139971 |
| Q3UVL4 | Vps51 | Vacuolar protein sorting-associated protein 51 homolog | 0.201398404 | 0.119614601 |
| Q3UX10 | Tubal3 | Tubulin alpha chain-like 3 | -0.892348512 | 0.107132037 |
| Q3UXU8 | Stn1 | CST complex subunit STN1 | 0.305704006 | 0.140078386 |
| Q3UY21 | Mog | Myelin-oligodendrocyte glycoprotein | 0.610088634 | 0.366547426 |
| Q3UYC0 | Ppm1h | Protein phosphatase 1H | 0.062272072 | 0.286158403 |
| Q3UYG8 | Macrod2 | ADP-ribose glycohydrolase MACROD2 | -0.047636255 | 0.645554066 |
| Q3UYV9 | Ncbp1 | Nuclear cap-binding protein subunit 1 | -0.119333776 | -0.139052232 |
| Q3V038 | Ttc9 | Tetratricopeptide repeat protein 9A | -0.134716702 | 0.754818439 |
| Q3V0K9 | Pls1 | Plastin-1 | 0.02492129 | 0.169998487 |
| Q3V117 | Acly | ATP-citrate synthase | -0.272734896 | 0.01715072 |
| Q3V1L6 | Mtmr11 | Myotubularin-related protein 11 | 1.080561002 | 2.335497061 |
| Q3V2R3 | Chn2 | Chimaerin | -0.122371356 | -0.283276876 |
| Q3V384 | Afg1l | AFG1-like ATPase | 1.271276824 | 1.568695068 |
| Q3V3R1 | Mthfd1l | Monofunctional C1-tetrahydrofolate synthase, mitochondrial | -0.148169263 | 0.073538939 |
| Q3V4D5 | Naa10 | N-alpha-acetyltransferase 10 | -0.514056524 | 1.47269249 |
| Q497I4 | Krt35 | Keratin, type I cuticular Ha5 | -0.524168841 | -0.120638688 |
| Q499X9 | Mars2 | Methionine--tRNA ligase, mitochondrial | -0.053892612 | -0.955404282 |
| Q4ACU6 | Shank3 | SH3 and multiple ankyrin repeat domains protein 3 | 0.101362769 | -0.589766502 |
| Q4FCQ7 | Ate1 | Arginyl-tRNA--protein transferase 1 | -0.116597939 | 0.718928178 |
| Q4KMM3 | Oxr1 | Oxidation resistance protein 1 | 0.000504684 | 0.083820025 |
| Q4LDD4 | Arap1 | Arf-GAP with Rho-GAP domain, ANK repeat and PH domain-containing protein 1 | -0.546057447 | -0.724205653 |
| Q4V9Z5 | Sez6l2 | Seizure 6-like protein 2 | -0.095600637 | -0.024755478 |
| Q4VA93 | Prkca | Protein kinase C | -0.220696354 | 0.14475139 |
| Q4VC33 | Maea | E3 ubiquitin-protein transferase MAEA | -0.408003712 | 0.54615434 |
| Q501J7 | Phactr4 | Phosphatase and actin regulator 4 | 0.104561869 | -0.466984342 |
| Q504M2 | Pdp2 | Pyruvate dehyrogenase phosphatase catalytic subunit 2 | -0.74644088 | 1.372515202 |
| Q50H33 | Kctd8 | BTB/POZ domain-containing protein KCTD8 | -0.031383991 | 1.063757539 |
| Q52KG3 | Magee2 | MAGE family member E2 | -1.155139256 | -0.024377346 |
| Q52KR3 | Prune2 | Protein prune homolog 2 | -0.059931405 | 0.145503203 |
| Q542V3 | Srsf4 | Serine/arginine-rich-splicing factor 4 | -0.317869631 | 0.054911772 |
| Q569Z6 | Thrap3 | Thyroid hormone receptor-associated protein 3 | -0.250601896 | 0.29513073 |
| Q571K4 | Tab3 | TGF-beta-activated kinase 1 and MAP3K7-binding protein 3 | -0.167784564 | 0.12465779 |
| Q58A65 | Spag9 | C-Jun-amino-terminal kinase-interacting protein 4 | 0.088077323 | 0.294014454 |
| Q5DQR4 | Stxbp5l | Syntaxin-binding protein 5-like | 0.657232745 | -0.00570337 |
| Q5DTY9 | Kctd16 | BTB/POZ domain-containing protein KCTD16 | -0.437437725 | 0.155576229 |

| | | | | |
|---|---|---|---|---|
| Q5DU31 | Ipcef1 | Interactor protein for cytohesin exchange factors 1 | -0.083106232 | -0.93986845 |
| Q5EBJ4 | Ermn | Ermin | 0.216660945 | 0.282595158 |
| Q5F258 | Git1 | ARF GTPase-activating protein GIT1 | 0.024382051 | 0.38715601 |
| Q5F2D9 | Arrb2 | Arrestin, beta 2 | 0.685405413 | -1.202902277 |
| Q5H8C4 | Vps13a | Vacuolar protein sorting-associated protein 13A | 0.055662791 | 0.160366217 |
| Q5ICG5 | Acsl6 | Long chain acyl-CoA synthetase 6 isoform 3 | -0.170104949 | -1.162170569 |
| Q5M8N0 | Cnrip1 | CB1 cannabinoid receptor-interacting protein 1 | 0.142385483 | 0.677652995 |
| Q5M8N4 | Sdr39u1 | Epimerase family protein SDR39U1 | 0.02258908 | -0.04493014 |
| Q5NCF2 | Trappc1 | Trafficking protein particle complex subunit 1 | -0.587413216 | -1.203792413 |
| Q5NCQ5 | Dph1 | 2-(3-amino-3-carboxypropyl)histidine synthase subunit 1 | -1.51109155 | -1.129100998 |
| Q5PR73 | Diras2 | GTP-binding protein Di-Ras2 | -0.024758148 | -0.129134178 |
| Q5QNQ6 | Osbp2 | Oxysterol-binding protein 2 | 0.362622865 | 0.419290384 |
| Q5RJI5 | Brsk1 | Serine/threonine-protein kinase BRSK1 | -0.792066987 | -0.636869113 |
| Q5RKN9 | Capza1 | F-actin-capping protein subunit alpha | -0.032212575 | 0.67148145 |
| Q5SQX6 | Cyfip2 | Cytoplasmic FMR1-interacting protein 2 | 0.155175432 | -0.263052781 |
| Q5SRF8 | Spred2 | Sprouty-related, EVH1 domain-containing protein 2 | -0.038145987 | -0.591141065 |
| Q5SRX1 | Tom1l2 | TOM1-like protein 2 | 0.095361519 | 0.91305383 |
| Q5SS83 | Flot2 | Flotillin | 0.138398711 | -0.109353781 |
| Q5SSL4 | Abr | Active breakpoint cluster region-related protein | 0.203100014 | 0.263734341 |
| Q5SSM3 | Arhgap44 | Rho GTPase-activating protein 44 | -0.023816331 | 0.392418861 |
| Q5SUF2 | Luc7l3 | Luc7-like protein 3 | -0.111021837 | 0.001686096 |
| Q5SUH6 | Clint1 | Clathrin interactor 1 | 0.010564518 | 0.604044755 |
| Q5SUR0 | Pfas | Phosphoribosylformylglycinamidine synthase | -0.402256838 | -0.005648931 |
| Q5SUS9 | Ewsr1 | RNA-binding protein EWS | 0.086464628 | 0.366342862 |
| Q5SV64 | Myh10 | Myosin-10 | -1.867384593 | -0.926577489 |
| Q5SV85 | Synrg | Synergin gamma | 0.259227053 | 0.089104811 |
| Q5SVR0 | Tbc1d9b | TBC1 domain family member 9B | 0.008559513 | 0.460865974 |
| Q5SWN2 | Rpa1 | Replication protein A subunit | 1.257920678 | 1.35778904 |
| Q5SWP3 | Nacad | NAC-alpha domain-containing protein 1 | -0.859398842 | 0.191498915 |
| Q5SWU9 | Acaca | Acetyl-CoA carboxylase 1 | 0.238879267 | 0.459612211 |
| Q5SWZ5 | Mprip | Myosin phosphatase Rho-interacting protein | 0.083282471 | -0.067721685 |
| Q5U3K5 | Rabl6 | Rab-like protein 6 | 0.108689149 | 0.218078613 |
| Q5U430 | Ubr3 | E3 ubiquitin-protein ligase UBR3 | -0.014216264 | -0.250422001 |
| Q5U4F6 | Dync2i2 | Cytoplasmic dynein 2 intermediate chain 2 | -0.122824796 | 0.203485012 |
| Q5U5V2 | Hykk | Hydroxylysine kinase | -0.764085706 | 0.392551263 |
| Q5XG69 | Fam169a | Soluble lamin-associated protein of 75 kDa | -0.074601046 | 0.521401087 |
| Q5XJY5 | Arcn1 | Coatomer subunit delta | -0.084339396 | -0.156887213 |
| Q5XPI3 | Rnf123 | E3 ubiquitin-protein ligase RNF123 | -0.638685894 | -0.180072625 |
| Q60590 | Orm1 | Alpha-1-acid glycoprotein 1 | 1.648274517 | 0.492646853 |
| Q60596 | Xrcc1 | DNA repair protein XRCC1 | 0.243697961 | 0.523093383 |
| Q60597 | Ogdh | 2-oxoglutarate dehydrogenase, mitochondrial | 0.034618537 | -0.124014537 |
| Q60598 | Cttn | Src substrate cortactin | 0.139608447 | 0.151805878 |
| Q60625 | Icam5 | Intercellular adhesion molecule 5 | 0.033598709 | -0.093420664 |
| Q60648 | Gm2a | Ganglioside GM2 activator | 0.432288074 | 0.334833145 |
| Q60668 | Hnrnpd | Heterogeneous nuclear ribonucleoprotein D0 | -0.111402798 | -0.0291152 |
| Q60676 | Ppp5c | Serine/threonine-protein phosphatase 5 | -0.04732008 | 0.148321311 |

| | | | | |
|---|---|---|---|---|
| Q60692 | Psmb6 | Proteasome subunit beta type-6 | 0.271641604 | 0.328903834 |
| Q60737 | Csnk2a1 | Casein kinase II subunit alpha | -0.228681723 | -0.003708363 |
| Q60749 | Khdrbs1 | KH domain-containing, RNA-binding, signal transduction-associated protein 1 | -0.34160312 | 0.946559111 |
| Q60771 | Cldn11 | Claudin-11 | 0.729746596 | -0.378482024 |
| Q60823 | Akt2 | RAC-beta serine/threonine-protein kinase | 0.283956146 | 0.21504577 |
| Q60829 | Ppp1r1b | Protein phosphatase 1 regulatory subunit 1B | 0.139095815 | 0.836201509 |
| Q60864 | Stip1 | Stress-induced-phosphoprotein 1 | 0.026280403 | 0.265788078 |
| Q60865 | Caprin1 | Caprin-1 | -0.099515915 | 0.192513625 |
| Q60900 | Elavl3 | ELAV-like protein 3 | 0.390656058 | -1.127483209 |
| Q60902 | Eps15l1 | Epidermal growth factor receptor substrate 15-like 1 | -0.016918437 | 0.524056753 |
| Q60930 | Vdac2 | Voltage-dependent anion-selective channel protein 2 | 0.306068198 | 1.038738251 |
| Q60932 | Vdac1 | Voltage-dependent anion-selective channel protein 1 | 0.459781933 | 0.440581163 |
| Q60960 | Kpna1 | Importin subunit alpha-5 | -0.037482103 | 0.091011365 |
| Q60963 | Pla2g7 | Platelet-activating factor acetylhydrolase | 0.158739535 | 0.37502416 |
| Q60967 | Papss1 | Bifunctional 3'-phosphoadenosine 5'-phosphosulfate synthase 1 | -0.195316251 | 0.182518164 |
| Q60972 | Rbbp4 | Histone-binding protein RBBP4 | -0.540466436 | -0.476973375 |
| Q60973 | Rbbp7 | Histone-binding protein RBBP7 | -0.040554206 | 0.857978821 |
| Q60994 | Adipoq | Adiponectin | 0.013436286 | 0.282592138 |
| Q60996 | Ppp2r5c | Serine/threonine-protein phosphatase 2A 56 kDa regulatory subunit gamma isoform | -0.299683348 | -0.352538904 |
| Q61024 | Asns | Asparagine synthetase [glutamine-hydrolyzing] | 0.018717511 | 0.357510885 |
| Q61029 | Tmpo | Lamina-associated polypeptide 2, isoforms beta/delta/epsilon/gamma | -0.011092758 | 0.394588153 |
| Q61035 | Hars1 | Histidine--tRNA ligase, cytoplasmic | 0.086113135 | 0.398314635 |
| Q61036 | Pak3 | Serine/threonine-protein kinase PAK 3 | -0.08743852 | 0.539464633 |
| Q61074 | Ppm1g | Protein phosphatase 1G | -0.13962698 | 0.242483775 |
| Q61081 | Cdc37 | Hsp90 co-chaperone Cdc37 | -0.051775646 | 0.317370097 |
| Q61140 | Bcar1 | Breast cancer anti-estrogen resistance protein 1 | -0.222053528 | 0.218450069 |
| Q61151 | Ppp2r5e | Serine/threonine-protein phosphatase 2A 56 kDa regulatory subunit epsilon isoform | -0.231884702 | 0.043034554 |
| Q61160 | Fadd | FAS-associated death domain protein | -0.070676676 | 0.829915524 |
| Q61161 | Map4k2 | Mitogen-activated protein kinase kinase kinase kinase 2 | 0.625900618 | -1.320657253 |
| Q61165 | Slc9a1 | Sodium/hydrogen exchanger 1 | -1.281450685 | 0.811481635 |
| Q61166 | Mapre1 | Microtubule-associated protein RP/EB family member 1 | -0.059779358 | 0.231291453 |
| Q61171 | Prdx2 | Peroxiredoxin-2 | 0.121822802 | 0.368530909 |
| Q61176 | Arg1 | Arginase-1 | 1.118613307 | -1.177727699 |
| Q61187 | Tsg101 | Tumor susceptibility gene 101 protein | 0.073305925 | 0.542191505 |
| Q61189 | Clns1a | Methylosome subunit pICln | 0.235052649 | 0.463134766 |
| Q61205 | Pafah1b3 | Platelet-activating factor acetylhydrolase IB subunit alpha1 | -0.138151614 | 0.862178802 |
| Q61233 | Lcp1 | Plastin-2 | 0.001509317 | 0.635955334 |
| Q61239 | Fnta | Protein farnesyltransferase/geranylgeranyltransferase type-1 subunit alpha | -0.052449099 | 0.357481798 |
| Q61247 | Serpinf2 | Alpha-2-antiplasmin | -0.001999505 | -0.223477046 |
| Q61249 | Igbp1 | Immunoglobulin-binding protein 1 | -0.060255559 | -2.032657186 |
| Q61282 | Acan | Aggrecan core protein | -0.465552457 | 0.307005405 |
| Q61330 | Cntn2 | Contactin-2 | -0.230323219 | 0.279037952 |
| Q61335 | Bcap31 | B-cell receptor-associated protein 31 | 1.550203037 | 0.992811203 |
| Q61361 | Bcan | Brevican core protein | -0.089367485 | 0.196333567 |

| | | | | |
|---|---|---|---|---|
| Q61411 | Hras | GTPase HRas | 0.534764067 | 1.127540032 |
| Q61425 | Hadh | Hydroxyacyl-coenzyme A dehydrogenase, mitochondrial | 0.02205286 | 0.222651958 |
| Q61481 | Pde1a | Calcium/calmodulin-dependent 3',5'-cyclic nucleotide phosphodiesterase 1A | -0.04878753 | 0.418049335 |
| Q61548 | Snap91 | Clathrin coat assembly protein AP180 | -0.138242213 | -0.020008087 |
| Q61553 | Fscn1 | Fascin | -0.051837413 | 0.064102809 |
| Q61578 | Fdxr | NADPH:adrenodoxin oxidoreductase, mitochondrial | 0.047588666 | 0.250081698 |
| Q61598 | Gdi2 | Rab GDP dissociation inhibitor beta | -0.109526253 | 0.388466835 |
| Q61599 | Arhgdib | Rho GDP-dissociation inhibitor 2 | 0.203770415 | 1.42996796 |
| Q61635 | Ifi47 | GTP-binding protein | 1.620029704 | 0.473732471 |
| Q61644 | Pacsin1 | Protein kinase C and casein kinase substrate in neurons protein 1 | 0.050508563 | 0.305254936 |
| Q61646 | Hp | Haptoglobin | 2.5277596 | 0.796781858 |
| Q61655 | Ddx19a | ATP-dependent RNA helicase DDX19A | 0.005546443 | 0.139835358 |
| Q61656 | Ddx5 | Probable ATP-dependent RNA helicase DDX5 | 0.089752706 | 0.510111809 |
| Q61686 | Cbx5 | Chromobox protein homolog 5 | -0.007805443 | 0.596599261 |
| Q61699 | Hsph1 | Heat shock protein 105 kDa | 0.011798096 | 0.172144254 |
| Q61701 | Elavl4 | ELAV-like protein 4 | -0.408457247 | -0.737043063 |
| Q61704 | Itih3 | Inter-alpha-trypsin inhibitor heavy chain H3 | -0.502815819 | -0.295855363 |
| Q61753 | Phgdh | D-3-phosphoglycerate dehydrogenase | -0.216708438 | 0.39527607 |
| Q61768 | Kif5b | Kinesin-1 heavy chain | -0.11201245 | -0.114533901 |
| Q61771 | Kif3b | Kinesin-like protein KIF3B | -0.290183735 | 0.374867439 |
| Q61781 | Krt14 | Keratin, type I cytoskeletal 14 | -0.253240649 | -0.513125102 |
| Q61792 | Lasp1 | LIM and SH3 domain protein 1 | 0.000950559 | 0.141186078 |
| Q61823 | Pdcd4 | Programmed cell death protein 4 | -0.127766037 | -0.913147807 |
| Q61838 | Pzp | Pregnancy zone protein | -0.078146394 | -0.033130805 |
| Q61937 | Npm1 | Nucleophosmin | -0.095135307 | 0.136285464 |
| Q62048 | Pea15 | Astrocytic phosphoprotein PEA-15 | 0.313465405 | 1.133684317 |
| Q62074 | Prkci | Protein kinase C iota type | -0.27377739 | 0.331498782 |
| Q62093 | Srsf2 | Serine/arginine-rich splicing factor 2 | -0.005049801 | 0.390544891 |
| Q62148 | Aldh1a2 | Retinal dehydrogenase 2 | -0.553503227 | 1.667802254 |
| Q62159 | Rhoc | Rho-related GTP-binding protein RhoC | -0.103485998 | 0.042755763 |
| Q62165 | Dag1 | Dystroglycan | 0.136855857 | 0.213456313 |
| Q62167 | Ddx3x | ATP-dependent RNA helicase DDX3X | -0.050329145 | -0.172820886 |
| Q62179 | Sema4b | Semaphorin-4B | 0.923847357 | -0.506009579 |
| Q62186 | Ssr4 | Translocon-associated protein subunit delta | 0.776605733 | -0.66972971 |
| Q62188 | Dpysl3 | Dihydropyrimidinase-related protein 3 | -0.037992732 | 0.057765643 |
| Q62189 | Snrpa | U1 small nuclear ribonucleoprotein A | -0.132491175 | -0.221188545 |
| Q62210 | Birc2 | Baculoviral IAP repeat-containing protein 2 | -0.421699651 | 0.163817167 |
| Q62241 | Snrpc | U1 small nuclear ribonucleoprotein C | -0.877845033 | 1.781199217 |
| Q62245 | Sos1 | Son of sevenless homolog 1 | -0.200782172 | 0.442842801 |
| Q62261 | Sptbn1 | Spectrin beta chain, non-erythrocytic 1 | -0.036161168 | -0.005887667 |
| Q62266 | Sprr1a | Cornifin-A | -0.658385372 | 0.805272261 |
| Q62267 | Sprr1b | Cornifin-B | -0.816629442 | 0.383344332 |
| Q62277 | Syp | Synaptophysin | 0.289754772 | 0.894567013 |
| Q62288 | Spock1 | Testican-1 | -0.068187364 | -0.041948318 |
| Q62318 | Trim28 | Transcription intermediary factor 1-beta | -0.176562246 | 0.396201293 |

| | | | | |
|---|---|---|---|---|
| Q62348 | Tsn | Translin | -0.120725282 | 0.290577253 |
| Q62376 | Snrnp70 | U1 small nuclear ribonucleoprotein 70 kDa | -0.344256306 | 0.05913798 |
| Q62383 | Supt6h | Transcription elongation factor SPT6 | -0.29882822 | -0.263657888 |
| Q62417 | Sorbs1 | Sorbin and SH3 domain-containing protein 1 | 0.91008056 | 2.582211574 |
| Q62418 | Dbnl | Drebrin-like protein | -0.020826403 | 0.074656487 |
| Q62419 | Sh3gl1 | Endophilin-A2 | 0.078762976 | 0.398543676 |
| Q62426 | Cstb | Cystatin-B | -0.030851905 | -0.230068048 |
| Q62433 | Ndrg1 | Protein NDRG1 | 0.160038948 | 0.986591657 |
| Q62443 | Nptx1 | Neuronal pentraxin-1 | -0.15396239 | 0.281872908 |
| Q62446 | Fkbp3 | Peptidyl-prolyl cis-trans isomerase FKBP3 | 0.259564304 | 0.697025458 |
| Q62465 | Vat1 | Synaptic vesicle membrane protein VAT-1 homolog | 0.230455875 | 1.149473667 |
| Q62523 | Zyx | Zyxin | -0.193188222 | 0.289279143 |
| Q63810 | Ppp3r1 | Calcineurin subunit B type 1 | -0.114520232 | -0.006973267 |
| Q63829 | Commd3 | COMM domain-containing protein 3 | -0.494344234 | 0.642578125 |
| Q63844 | Mapk3 | Mitogen-activated protein kinase 3 | -0.164977328 | 0.38954099 |
| Q63850 | Nup62 | Nuclear pore glycoprotein p62 | -1.16407512 | 0.430800756 |
| Q63918 | Cavin2 | Caveolae-associated protein 2 | -1.316295719 | 1.712164879 |
| Q63932 | Map2k2 | Dual specificity mitogen-activated protein kinase kinase 2 | -0.090090116 | 0.321085612 |
| Q64010 | Crk | Adapter molecule crk | 0.004981963 | 0.425193787 |
| Q640R3 | Hepacam | Hepatocyte cell adhesion molecule | 0.125786273 | 0.38853995 |
| Q64112 | Ifit2 | Interferon-induced protein with tetratricopeptide repeats 2 | 0.272515758 | 0.092274348 |
| Q64133 | Maoa | Amine oxidase [flavin-containing] A | -1.836486749 | 0.728792349 |
| Q64152 | Btf3 | Transcription factor BTF3 | 0.205029043 | 0.173571905 |
| Q641P0 | Actr3b | Actin-related protein 3B | 0.743331051 | 0.405194918 |
| Q64288 | Omp | Olfactory marker protein | -2.562327766 | 1.059277217 |
| Q64291 | Krt12 | Keratin, type I cytoskeletal 12 | 0.750220235 | -0.61457173 |
| Q64310 | Surf4 | Surfeit locus protein 4 | -1.127542337 | -0.402612845 |
| Q64332 | Syn2 | Synapsin-2 | 0.01811765 | 0.091341337 |
| Q64337 | Sqstm1 | Sequestosome-1 | -0.024139023 | 0.951058865 |
| Q64374 | Rgn | Regucalcin | 0.171337573 | 1.397078117 |
| Q64378 | Fkbp5 | Peptidyl-prolyl cis-trans isomerase FKBP5 | 0.142706553 | 0.458745956 |
| Q64433 | Hspe1 | 10 kDa heat shock protein, mitochondrial | 0.02600797 | 0.572164853 |
| Q64442 | Sord | Sorbitol dehydrogenase | -0.110304387 | 0.31583039 |
| Q64471 | Gstt1 | Glutathione S-transferase theta-1 | -0.296468163 | -0.033215364 |
| Q64514 | Tpp2 | Tripeptidyl-peptidase 2 | -0.116535568 | 0.028529008 |
| Q64520 | Guk1 | Guanylate kinase | 0.308597819 | 0.297250112 |
| Q64521 | Gpd2 | Glycerol-3-phosphate dehydrogenase, mitochondrial | 0.37159462 | 0.982705911 |
| Q64669 | Nqo1 | NAD(P)H dehydrogenase [quinone] 1 | -0.181941923 | 0.171369553 |
| Q64674 | Srm | Spermidine synthase | -0.005020364 | -0.445417086 |
| Q64727 | Vcl | Vinculin | 0.018231106 | 0.309179942 |
| Q64737 | Gart | Trifunctional purine biosynthetic protein adenosine-3 | -0.033677832 | 0.2045331 |
| Q66L47 | Gnal | Guanine nucleotide binding protein, alpha stimulating, olfactory type | 0.440175947 | 1.098476887 |
| Q68ED7 | Crtc1 | CREB-regulated transcription coactivator 1 | 0.620063241 | -0.016841571 |
| Q68FD5 | Cltc | Clathrin heavy chain 1 | -0.157138824 | 0.110179901 |
| Q68FE6 | Ripor1 | Rho family-interacting cell polarization regulator 1 | 0.981133207 | 0.948920886 |

| | | | | |
|---|---|---|---|---|
| Q68FG2 | Sptbn2 | Spectrin beta chain | -0.031339137 | -0.043606122 |
| Q68FH4 | Galk2 | N-acetylgalactosamine kinase | 0.03430287 | -0.097347101 |
| Q68FL1 | Adgrb3 | Adhesion G protein-coupled receptor B3 (Fragment) | -0.475616201 | -0.775553544 |
| Q68FM6 | Elfn2 | Protein phosphatase 1 regulatory subunit 29 | 0.586917639 | -1.657341838 |
| Q68FM7 | Arhgef11 | Rho guanine nucleotide exchange factor (GEF) 11 | -0.153026994 | -1.024278005 |
| Q69Z98 | Brsk2 | Serine/threonine-protein kinase BRSK2 | 0.323137506 | 0.996233702 |
| Q69ZC8 | Gpalpp1 | GPALPP motifs-containing protein 1 | 0.816065979 | 0.586922169 |
| Q69ZF3 | Gba2 | Non-lysosomal glucosylceramidase | 0.55989186 | 0.167634487 |
| Q69ZK0 | Prex1 | Phosphatidylinositol 3,4,5-trisphosphate-dependent Rac exchanger 1 protein | -0.272504934 | -0.002081076 |
| Q69ZW3 | Ehbp1 | EH domain-binding protein 1 | 0.23504413 | 1.232508024 |
| Q6A028 | Swap70 | Switch-associated protein 70 | -0.69634161 | 0.552115917 |
| Q6A037 | N4bp1 | NEDD4-binding protein 1 | -0.136841297 | -0.644175212 |
| Q6A068 | Cdc5l | Cell division cycle 5-like protein | 0.986919403 | -0.557964007 |
| Q6A0A2 | Larp4b | La-related protein 4B | -0.465100606 | 0.804011504 |
| Q6A0A9 | FAM120A | Constitutive coactivator of PPAR-gamma-like protein 1 | 1.055964979 | 0.516047955 |
| Q6DFW0 | | Guanine nucleotide exchange C9orf72 homolog | -0.184795825 | -0.380962054 |
| Q6DFY2 | Opcml | Opioid-binding protein/cell adhesion molecule-like | 0.700369644 | 0.527044455 |
| Q6DFZ1 | Gbf1 | Golgi-specific brefeldin A-resistance factor 1 | -0.509221331 | 0.375256538 |
| Q6DID3 | Scaf8 | SR-related and CTD-associated factor 8 | -0.798259226 | 0.038284938 |
| Q6GT24 | Prdx6 | Peroxiredoxin-6 | -0.011989021 | 0.487413724 |
| Q6I6G8 | Hecw2 | E3 ubiquitin-protein ligase HECW2 | -0.052433745 | -0.08161513 |
| Q6IFX2 | Krt42 | Keratin, type I cytoskeletal 42 | -0.221327082 | -1.003115336 |
| Q6IFZ6 | Krt77 | Keratin, type II cytoskeletal 1b | 0.063013522 | -0.177830378 |
| Q6IME9 | Krt72 | Keratin, type II cytoskeletal 72 | -0.070902952 | -0.629342556 |
| Q6IR34 | Gpsm1 | G-protein-signaling modulator 1 | 0.055440108 | 0.214421272 |
| Q6IRU2 | Tpm4 | Tropomyosin alpha-4 chain | -0.086931801 | 0.555776119 |
| Q6IRU5 | Cltb | Clathrin light chain B | 0.07327404 | 0.258076986 |
| Q6KAU4 | Mvb12b | Multivesicular body subunit 12B | -0.428875828 | 0.345953782 |
| Q6NSR8 | Npepl1 | Probable aminopeptidase NPEPL1 | -0.26645209 | 0.208477338 |
| Q6NT99 | Dusp23 | Dual specificity protein phosphatase 23 | -0.236228848 | 0.207702319 |
| Q6NVE8 | Wdr44 | WD repeat-containing protein 44 | 0.000288518 | 0.178058147 |
| Q6NVE9 | Pptc7 | Protein phosphatase PTC7 homolog | -0.223176702 | 0.429542859 |
| Q6NVF0 | Ocrl | Inositol polyphosphate 5-phosphatase OCRL | -0.096425311 | 0.007899761 |
| Q6NXH9 | Krt73 | Keratin, type II cytoskeletal 73 | -0.073678144 | -0.285926819 |
| Q6NZD2 | Snx1 | Sorting nexin-1 | -0.034701188 | 0.485878309 |
| Q6NZL0 | Soga3 | Protein SOGA3 | 0.197152615 | -0.097639243 |
| Q6NZM9 | Hdac4 | Histone deacetylase 4 | 0.352978452 | -0.193738461 |
| Q6NZR5 | Skiv2l | Superkiller viralicidic activity 2-like (S. cerevisiae) | 0.475995 | 1.933543205 |
| Q6P069 | Sri | Sorcin | -0.023439598 | 0.326384544 |
| Q6P1B9 | Bin1 | Bin1 protein | -0.040282027 | -0.524842898 |
| Q6P1E1 | Zmiz1 | Zinc finger MIZ domain-containing protein 1 | 0.738157972 | -0.169791381 |
| Q6P1F6 | Ppp2r2a | Serine/threonine-protein phosphatase 2A 55 kDa regulatory subunit B alpha isoform | 0.006680234 | 0.123216311 |
| Q6P1J1 | Crmp1 | Crmp1 protein | 0.085648473 | -1.066135724 |
| Q6P2B1 | Tnpo3 | Transportin-3 | -0.322579988 | 0.711047331 |
| Q6P3A8 | Bckdhb | 2-oxoisovalerate dehydrogenase subunit beta, mitochondrial | -0.699572277 | 0.411450704 |

| | | | | |
|---|---|---|---|---|
| Q6P4T2 | Snrnp200 | U5 small nuclear ribonucleoprotein 200 kDa helicase | -0.003151417 | -0.674342155 |
| Q6P542 | Abcf1 | ATP-binding cassette sub-family F member 1 | -0.395079358 | -0.04486831 |
| Q6P5B5 | Fxr2 | Fragile X mental retardation syndrome-related protein 2 | -0.420950794 | 0.505917708 |
| Q6P5E4 | Uggt1 | UDP-glucose:glycoprotein glucosyltransferase 1 | 0.263593006 | 0.545001189 |
| Q6P5E8 | Dgkq | Diacylglycerol kinase theta | 0.025664552 | -0.335867087 |
| Q6P5F9 | Xpo1 | Exportin-1 | -0.124911594 | 0.776514212 |
| Q6P5H2 | Nes | Nestin | 0.246646039 | -0.017240763 |
| Q6P5U7 | Nwd2 | NACHT and WD repeat domain-containing protein 2 | 0.229583327 | 0.424346447 |
| Q6P6I6 | Polr2m | DNA-directed RNA polymerase II subunit GRINL1A | 0.359180355 | 0.971937656 |
| Q6P8I4 | Pcnp | PEST proteolytic signal-containing nuclear protein | 0.235143185 | 0.775417964 |
| Q6P8J2 | Sat2 | Thialysine N-epsilon-acetyltransferase | -0.187627761 | 0.760414759 |
| Q6P8J7 | Ckmt2 | Creatine kinase S-type, mitochondrial | 1.635727024 | 1.244046688 |
| Q6P8K8 | Cpa4 | Carboxypeptidase A4 | 0.315567843 | -0.541555087 |
| Q6P8N8 | Clpx | ATP-dependent Clp protease ATP-binding subunit clpX-like, mitochondrial | 0.977558613 | 0.551648299 |
| Q6P8X1 | Snx6 | Sorting nexin-6 | -0.053203297 | 0.29050382 |
| Q6P9J5 | Kank4 | KN motif and ankyrin repeat domain-containing protein 4 | -0.329730765 | 1.043198586 |
| Q6P9K8 | Caskin1 | Caskin-1 | 0.059540685 | 0.369708856 |
| Q6P9Q6 | Fkbp15 | FK506-binding protein 15 | -0.710351912 | 0.23544693 |
| Q6P9R2 | Oxsr1 | Serine/threonine-protein kinase OSR1 | -0.050048478 | 0.322756608 |
| Q6P9S0 | Mtss2 | Protein MTSS 2 | 0.330000178 | 0.490110238 |
| Q6PAJ1 | Bcr | Breakpoint cluster region protein | -3.80E-05 | 0.1249005 |
| Q6PAK3 | Prmt8 | Protein arginine N-methyltransferase 8 | 0.10497853 | 0.062774658 |
| Q6PAM0 | Prkab2 | 5'-AMP-activated protein kinase subunit beta-2 | 0.543174712 | 1.779740969 |
| Q6PAR5 | Gapvd1 | GTPase-activating protein and VPS9 domain-containing protein 1 | -0.180804094 | -0.164512634 |
| Q6PAV2 | Herc4 | Probable E3 ubiquitin-protein ligase HERC4 | -0.365306918 | -0.488464673 |
| Q6PB44 | Ptpn23 | Tyrosine-protein phosphatase non-receptor type 23 | 0.01136783 | 0.303129514 |
| Q6PB66 | Lrpprc | Leucine-rich PPR motif-containing protein, mitochondrial | -0.044860999 | 0.265798251 |
| Q6PCN3 | Ttbk1 | Tau-tubulin kinase 1 | 0.14630305 | 0.586991469 |
| Q6PD03 | Ppp2r5a | Serine/threonine-protein phosphatase 2A 56 kDa regulatory subunit alpha isoform | 0.242453067 | 0.003877958 |
| Q6PD10 | Ip6k1 | Inositol hexakisphosphate kinase 1 | -0.344308503 | -0.056727091 |
| Q6PD19 | Armh3 | Armadillo-like helical domain-containing protein 3 | 0.119421577 | 0.835006078 |
| Q6PD24 | Ankrd13d | Ankyrin repeat domain-containing protein 13D | 0.488953749 | 2.220701377 |
| Q6PD28 | Ppp2r5b | Serine/threonine-protein phosphatase 2A 56 kDa regulatory subunit beta isoform | 0.218540064 | 0.503246625 |
| Q6PDJ6 | Fbxo42 | F-box only protein 42 | 0.424653657 | 0.74574693 |
| Q6PDL0 | Dync1li2 | Cytoplasmic dynein 1 light intermediate chain 2 | 0.320173518 | 0.2720294 |
| Q6PDS3 | Sarm1 | NAD(+) hydrolase SARM1 | -0.25917956 | -1.024649779 |
| Q6PDX6 | Rnf220 | E3 ubiquitin-protein ligase Rnf220 | -0.256131999 | 0.751439412 |
| Q6PDY2 | Ado | 2-aminoethanethiol dioxygenase | -0.152783171 | 0.483461062 |
| Q6PE15 | Abhd10 | Palmitoyl-protein thioesterase ABHD10, mitochondrial | 0.163672161 | 0.726527691 |
| Q6PEE3 | Rrm2b | Ribonucleoside-diphosphate reductase subunit M2 B | 0.181885624 | 0.296338399 |
| Q6PER3 | Mapre3 | Microtubule-associated protein RP/EB family member 3 | 0.106908353 | 0.342697461 |
| Q6PF96 | Etfdh | Electron transfer flavoprotein-ubiquinone oxidoreductase | 0.730936972 | 1.247580051 |
| Q6PGB6 | Naa50 | N-alpha-acetyltransferase 50 | 0.252284114 | -0.983263969 |
| Q6PGF2 | Morn4 | MORN repeat-containing protein 4 | 0.495602544 | 0.641591708 |
| Q6PGF7 | Exoc8 | Exocyst complex component 8 | 0.043000158 | -0.070786635 |

| | | | | |
|---|---|---|---|---|
| Q6PGH0 | Ubtd2 | Ubiquitin domain-containing protein 2 | 0.812969144 | 0.201040427 |
| Q6PGH2 | Jpt2 | Jupiter microtubule associated homolog 2 | 0.084830697 | -0.137568315 |
| Q6PGK3 | 2700097O09Rik | RIKEN cDNA 2700097O09 gene | -0.082906246 | -0.166299582 |
| Q6PGL7 | Washc2 | WASH complex subunit 2 | 0.069937611 | 0.293315411 |
| Q6PHN9 | Rab35 | Ras-related protein Rab-35 | 0.361570962 | 0.760713577 |
| Q6PHU5 | Sort1 | Sortilin | 0.567804845 | -0.300097307 |
| Q6PIE5 | Atp1a2 | Sodium/potassium-transporting ATPase subunit alpha-2 | 0.349588108 | 0.36712424 |
| Q6PIU9 | | Uncharacterized protein FLJ45252 homolog | -0.423905754 | 0.169822057 |
| Q6Q899 | Ddx58 | Antiviral innate immune response receptor RIG-I | -0.417561118 | 0.158212582 |
| Q6R891 | Ppp1r9b | Neurabin-2 | 0.15991478 | -0.277362982 |
| Q6SKR2 | N6amt1 | Methyltransferase N6AMT1 | -0.371687635 | -0.300957203 |
| Q6W8Q3 | Pcp4l1 | Purkinje cell protein 4-like protein 1 | 0.422820473 | 0.658443769 |
| Q6WKZ8 | Ubr2 | E3 ubiquitin-protein ligase UBR2 | 0.066646067 | 0.61778752 |
| Q6XE40 | Mpp3 | MAGUK p55 subfamily member 3 | -0.063879744 | 0.085825443 |
| Q6XLQ8 | Calu | Calumenin | -0.243348567 | 0.06573232 |
| Q6XPS7 | Tha1 | L-threonine aldolase | -0.10908521 | -0.762520154 |
| Q6ZPE2 | Sbf1 | Myotubularin-related protein 5 | 0.069088109 | -0.238424301 |
| Q6ZPJ3 | Ube2o | (E3-independent) E2 ubiquitin-conjugating enzyme UBE2O | -0.274082247 | 0.156109492 |
| Q6ZPU9 | Kifbp | KIF-binding protein | 0.018831285 | 0.063643138 |
| Q6ZQ29 | Taok2 | Serine/threonine-protein kinase TAO2 | 0.309956328 | 1.840564728 |
| Q6ZQ38 | Cand1 | Cullin-associated NEDD8-dissociated protein 1 | -0.032763036 | 0.215666453 |
| Q6ZQ58 | Larp1 | La-related protein 1 | 0.285857582 | -0.78552866 |
| Q6ZQI3 | Mlec | Malectin | -1.292472363 | 0.837581635 |
| Q6ZWM4 | Lsm8 | U6 snRNA-associated Sm-like protein LSm8 | -0.191112868 | -0.015534242 |
| Q6ZWN5 | Rps9 | 40S ribosomal protein S9 | 0.219314575 | 0.898462137 |
| Q6ZWR6 | Syne1 | Nesprin-1 | -0.009836102 | -0.248994509 |
| Q6ZWV7 | Rpl35 | 60S ribosomal protein L35 | 0.830810006 | 0.721066475 |
| Q6ZWX6 | Eif2s1 | Eukaryotic translation initiation factor 2 subunit 1 | 0.080535603 | 0.388485909 |
| Q6ZWY8 | Tmsb10 | Thymosin beta-10 | 0.166703574 | 1.116142273 |
| Q6ZWZ2 | Ube2r2 | Ubiquitin-conjugating enzyme E2 R2 | -0.131626002 | 0.77562205 |
| Q6ZWZ7 | Rpl17 | 60S ribosomal protein L17 | 0.78682181 | 1.577475548 |
| Q71M36 | Cspg5 | Chondroitin sulfate proteoglycan 5 | -0.563055325 | -0.645074526 |
| Q71RI9 | Kyat3 | Kynurenine--oxoglutarate transaminase 3 | -0.268520228 | -0.47812144 |
| Q76LS9 | Mindy1 | Ubiquitin carboxyl-terminal hydrolase MINDY-1 | -0.23294611 | 0.117382844 |
| Q76MZ3 | Ppp2r1a | Serine/threonine-protein phosphatase 2A 65 kDa regulatory subunit A alpha isoform | -0.072193209 | 0.243757884 |
| Q78IK2 | Atp5mk | ATP synthase membrane subunit K, mitochondrial | 1.066624928 | 1.114952087 |
| Q78J03 | Msrb2 | Methionine-R-sulfoxide reductase B2, mitochondrial | -0.040592353 | 0.260954698 |
| Q78JE5 | Fbxo22 | F-box only protein 22 | -0.13379453 | 0.32014211 |
| Q78JW9 | Ubfd1 | Ubiquitin domain-containing protein UBFD1 | 0.214680004 | 0.751449426 |
| Q78PG9 | Ccdc25 | Coiled-coil domain-containing protein 25 | 0.567833074 | 0.715143363 |
| Q78PY7 | Snd1 | Staphylococcal nuclease domain-containing protein 1 | -0.172128677 | 0.613730907 |
| Q78ZM0 | Snx3 | Sorting nexin-3 | 0.114730231 | 0.539102395 |
| Q791V5 | Mtch2 | Mitochondrial carrier homolog 2 | 0.231516234 | 1.392058531 |
| Q792Z1 | Try10 | Trypsin 10 | 0.06419398 | 0.162413597 |
| Q7M6W1 | Rtn1 | Reticulon | 0.268539174 | -0.244544824 |

| | | | | |
|---|---|---|---|---|
| Q7M6Y3 | Picalm | Phosphatidylinositol-binding clathrin assembly protein | -0.057958126 | 0.431426366 |
| Q7TMB8 | Cyfip1 | Cytoplasmic FMR1-interacting protein 1 | 0.059460068 | 0.115797361 |
| Q7TMC8 | Fcsk | L-fucose kinase | -0.910086346 | 0.698811213 |
| Q7TMM9 | Tubb2a | Tubulin beta-2A chain | 0.246436691 | 0.782925606 |
| Q7TMR0 | Prcp | Lysosomal Pro-X carboxypeptidase | -0.375557105 | -0.043436527 |
| Q7TMW6 | Ciao3 | Cytosolic iron-sulfur assembly component 3 | -1.593631474 | 0.257064899 |
| Q7TN29 | Smap2 | Stromal membrane-associated protein 2 | -0.074360625 | 0.533500512 |
| Q7TNC4 | Luc7l2 | Putative RNA-binding protein Luc7-like 2 | -0.948862012 | 1.355705579 |
| Q7TNE3 | Spag7 | Sperm-associated antigen 7 | 0.326091067 | 1.891360044 |
| Q7TNF0 | Doc2a | Double C2-like domain-containing protein alpha | -0.646186288 | -0.642726262 |
| Q7TNG8 | Ldhd | Probable D-lactate dehydrogenase, mitochondrial | -1.338798618 | 0.724160353 |
| Q7TNL5 | Ppp2r5d | Serine/threonine-protein phosphatase 2A 56 kDa regulatory subunit | -0.277805901 | 0.07485199 |
| Q7TNV0 | Dek | Protein DEK | -1.089197858 | 0.148350557 |
| Q7TPC1 | Cdsn | Corneodesmosin | 0.368588352 | -1.445136547 |
| Q7TPD2 | Fam185a | Protein FAM185A | -0.101557795 | 0.467683156 |
| Q7TPE5 | Slc7a6os | Probable RNA polymerase II nuclear localization protein SLC7A6OS | 0.508993149 | 0.708413601 |
| Q7TPM6 | Fsd1 | Fibronectin type III and SPRY domain-containing protein 1 | 0.010577901 | 0.392287413 |
| Q7TPR4 | Actn1 | Alpha-actinin-1 | 0.847716014 | 0.079220295 |
| Q7TQ95 | Lnpk | Endoplasmic reticulum junction formation protein lunapark | 0.555681292 | 0.946762562 |
| Q7TQD2 | Tppp | Tubulin polymerization-promoting protein | 0.013963445 | 0.291582743 |
| Q7TQG7 | Adam11 | Adam11 protein | -0.442648633 | 0.482263803 |
| Q7TQK5 | Ccdc93 | Coiled-coil domain-containing protein 93 | 0.319679737 | 0.355121295 |
| Q7TRZ9 | Olfr312 | Olfactory receptor | 0.949635792 | 0.483444532 |
| Q7TSE6 | Stk38l | Serine/threonine-protein kinase 38-like | -0.332013639 | 0.482535998 |
| Q7TSF1 | Dsg1b | Desmoglein-1-beta | 0.031404463 | -0.443448385 |
| Q7TSG2 | Ctdp1 | RNA polymerase II subunit A C-terminal domain phosphatase | 0.051988029 | -0.334081809 |
| Q7TSH2 | Phkb | Phosphorylase b kinase regulatory subunit beta | 0.388126914 | 0.403439522 |
| Q7TSI3 | Ppp6r1 | Serine/threonine-protein phosphatase 6 regulatory subunit 1 | 0.305022144 | 0.68235143 |
| Q7TSJ2 | Map6 | Microtubule-associated protein 6 | 0.273548571 | 0.485069275 |
| Q7TSJ7 | Mapk8 | Mitogen-activated protein kinase | 0.9000995 | -1.492467086 |
| Q7TSQ8 | Pdpr | Pyruvate dehydrogenase phosphatase regulatory subunit, mitochondrial | -0.29457531 | 0.421745459 |
| Q7TSS2 | Ube2q1 | Ubiquitin-conjugating enzyme E2 Q1 | -0.227155813 | 0.552832127 |
| Q7TSV4 | Pgm2 | Phosphoglucomutase-2 | -0.013216209 | 0.735293229 |
| Q7TT23 | | Uncharacterized protein C20orf194 homolog | -0.491174126 | 1.016781807 |
| Q7TT37 | Elp1 | Elongator complex protein 1 | -0.00745163 | -0.168318907 |
| Q7TT50 | Cdc42bpb | Serine/threonine-protein kinase MRCK beta | 0.00462335 | -0.27755324 |
| Q80SW1 | Ahcyl1 | S-adenosylhomocysteine hydrolase-like protein 1 | -0.022745768 | 0.410754045 |
| Q80SY6 | Adal | Adenosine deaminase-like protein | -0.953258006 | 1.063742797 |
| Q80T85 | Dcaf5 | DDB1- and CUL4-associated factor 5 | -0.6258358 | -0.077109973 |
| Q80TA6 | Mtmr12 | Myotubularin-related protein 12 | -0.241131496 | 1.931982597 |
| Q80TA9 | Epg5 | Ectopic P granules protein 5 homolog | 0.884714047 | 0.384875774 |
| Q80TB7 | Zswim6 | Zinc finger SWIM domain-containing protein 6 | 1.527328968 | -0.045001189 |
| Q80TB8 | Vat1l | Synaptic vesicle membrane protein VAT-1 homolog-like | 0.110993417 | 0.489087582 |
| Q80TJ1 | Cadps | Calcium-dependent secretion activator 1 | -0.083664322 | 0.12709713 |

| | | | | |
|---|---|---|---|---|
| Q80TL0 | Ppm1e | Protein phosphatase 1E | -0.270921485 | 0.221811295 |
| Q80TL4 | Phf24 | PHD finger protein 24 | -0.182445335 | 0.177793185 |
| Q80TL7 | Mon2 | Protein MON2 homolog | -0.064841715 | -0.945752621 |
| Q80TM9 | Nisch | Nischarin | -0.010954475 | 0.304620266 |
| Q80TQ2 | Cyld | Ubiquitin carboxyl-terminal hydrolase CYLD | 0.48758262 | -0.599018574 |
| Q80TZ3 | Dnajc6 | Putative tyrosine-protein phosphatase auxilin | -0.218188032 | -0.109421412 |
| Q80U04 | Pja2 | E3 ubiquitin-protein ligase Praja-2 | 0.210405223 | -1.239048958 |
| Q80U23 | Snph | Syntaphilin | -0.14553655 | 0.760373036 |
| Q80U35 | Arhgef17 | Rho guanine nucleotide exchange factor 17 | -1.231989638 | 0.624525944 |
| Q80U56 | Avl9 | Late secretory pathway protein AVL9 homolog | -0.137871424 | -0.124960105 |
| Q80U57 | Rims3 | Regulating synaptic membrane exocytosis protein 3 | -0.143714778 | 0.256457965 |
| Q80U72 | Scrib | Protein scribble homolog | 1.516682259 | 2.087606112 |
| Q80U78 | Pum1 | Pumilio homolog 1 | -0.660385227 | 1.180075089 |
| Q80U93 | Nup214 | Nuclear pore complex protein Nup214 | -1.082579613 | 0.479476452 |
| Q80U95 | Ube3c | Ubiquitin-protein ligase E3C | 0.95236365 | -0.196115335 |
| Q80UE5 | Epb41l2 | Band 4.1-like protein 2 | -0.093268744 | 0.179458777 |
| Q80UG2 | Plxna4 | Plexin-A4 | 0.020692539 | 0.103056749 |
| Q80UK0 | Sestd1 | SEC14 domain and spectrin repeat-containing protein 1 | 0.036922963 | -0.172175407 |
| Q80UP5 | Ankrd13a | Ankyrin repeat domain-containing protein 13A | 1.964610585 | -0.068935692 |
| Q80UU9 | Pgrmc2 | Membrane-associated progesterone receptor component 2 | 0.722681522 | 1.129594008 |
| Q80UW2 | Fbxo2 | F-box only protein 2 | -0.472193527 | -0.595115821 |
| Q80UW8 | Polr2e | DNA-directed RNA polymerases I, II, and III subunit RPABC1 | -0.432606379 | -0.265276273 |
| Q80VC9 | Camsap3 | Calmodulin-regulated spectrin-associated protein 3 | -0.000707372 | 0.187024752 |
| Q80VD1 | Fam98b | Protein FAM98B | 0.085677973 | 0.544098457 |
| Q80VE5 | Tbc1d22b | TBC1 domain family, member 22B | -0.652106317 | 0.132188479 |
| Q80VJ2 | Sra1 | Steroid receptor RNA activator 1 | 0.31187528 | 1.458879471 |
| Q80VJ3 | Dnph1 | 2'-deoxynucleoside 5'-phosphate N-hydrolase 1 | 0.237231541 | 1.101596514 |
| Q80VM5 | Dpp6 | Dipeptidyl aminopeptidase-like protein 6 | 0.436607456 | 0.177724202 |
| Q80VP0 | Tecpr1 | Tectonin beta-propeller repeat-containing protein 1 | 0.26524566 | 0.794640779 |
| Q80VP1 | Epn1 | Epsin-1 | -0.043510024 | 0.398200671 |
| Q80W21 | Gstm7 | Glutathione S-transferase Mu 7 | -0.212208303 | 0.708268801 |
| Q80W22 | Thnsl2 | Threonine synthase-like 2 | 0.023596287 | 0.59452343 |
| Q80W47 | Wipi2 | WD repeat domain phosphoinositide-interacting protein 2 | -0.079104265 | 0.587289492 |
| Q80WB5 | Ntaq1 | Protein N-terminal glutamine amidohydrolase | -0.172578144 | -0.400469621 |
| Q80WM4 | Hapln4 | Hyaluronan and proteoglycan link protein 4 | -0.149168142 | -0.26554362 |
| Q80X50 | Ubap2l | Ubiquitin-associated protein 2-like | 0.424148814 | 0.277990341 |
| Q80X68 | Csl | Citrate synthase | 0.362049421 | 1.775975704 |
| Q80X71 | Tmem106b | Transmembrane protein 106B | 0.46568807 | -0.06154712 |
| Q80X73 | Pelo | Protein pelota homolog | 0.048651759 | -0.011533101 |
| Q80X80 | C2cd2l | Phospholipid transfer protein C2CD2L | -0.30590814 | 0.779869397 |
| Q80X81 | Acat3 | Acetyl-Coenzyme A acetyltransferase 3 | 0.021242332 | -0.014291604 |
| Q80X90 | Flnb | Filamin-B | 0.336802769 | 0.455751896 |
| Q80X95 | Rraga | Ras-related GTP-binding protein A | -0.441330465 | 0.395596663 |
| Q80XE1 | Ric8b | Synembryn-B | 1.200025845 | 0.1464324 |
| Q80XI4 | Pip4k2b | Phosphatidylinositol 5-phosphate 4-kinase type-2 beta | 0.218952688 | 1.044852257 |

| | | | | |
|---|---|---|---|---|
| Q80XK6 | Atg2b | Autophagy-related protein 2 homolog B | 0.09910752 | -0.535398165 |
| Q80XN0 | Bdh1 | D-beta-hydroxybutyrate dehydrogenase, mitochondrial | 0.020101833 | 1.324875673 |
| Q80Y14 | Glrx5 | Glutaredoxin-related protein 5, mitochondrial | -0.839184348 | 0.923842112 |
| Q80Y55 | Bsdc1 | BSD domain-containing protein 1 | 0.122365316 | 1.150143981 |
| Q80Y81 | Elac2 | Zinc phosphodiesterase ELAC protein 2 | 0.005215804 | 0.907659849 |
| Q80YA7 | Dpp8 | Dipeptidyl peptidase 8 | 0.194452 | 0.002380848 |
| Q80YA9 | Cnksr2 | Connector enhancer of kinase suppressor of ras 2 | 0.473487155 | -0.216724555 |
| Q80YD1 | Supv3l1 | ATP-dependent RNA helicase SUPV3L1, mitochondrial | 0.048165735 | 1.815255006 |
| Q80YN3 | Bcas1 | Breast carcinoma-amplified sequence 1 homolog | 0.003467687 | 0.363075733 |
| Q80YV4 | Pank4 | 4'-phosphopantetheine phosphatase | 0.071705151 | 0.291451613 |
| Q80YX1 | Tnc | Tenascin | -0.417029572 | -0.181005319 |
| Q80ZI6 | Lrsam1 | E3 ubiquitin-protein ligase LRSAM1 | 0.103596433 | 0.478055 |
| Q80ZJ1 | Rap2a | Ras-related protein Rap-2a | 0.960349178 | 0.26412344 |
| Q80ZJ6 | Zer1 | Protein zer-1 homolog | -0.24441913 | -0.098625978 |
| Q80ZU0 | Arl5a | ADP-ribosylation factor-like protein 5A | 0.517166074 | 0.994525274 |
| Q80ZX0 | Sec24b | Sec24-related gene family, member B (S. cerevisiae) | 0.008097871 | 0.569176197 |
| Q80ZX8 | Spag1 | Sperm-associated antigen 1 | -0.26020778 | 0.5914011 |
| Q810A3 | Ttc9c | Tetratricopeptide repeat protein 9C | 0.149229749 | 0.261379083 |
| Q810A7 | Ddx42 | ATP-dependent RNA helicase DDX42 | -0.344580396 | 0.326231639 |
| Q810B6 | Ankfy1 | Rabankyrin-5 | -0.174981531 | 0.05206426 |
| Q810C1 | Slitrk1 | SLIT and NTRK-like protein 1 | 0.529592864 | -0.02374649 |
| Q810J8 | Zfyve1 | Zinc finger FYVE domain-containing protein 1 | -0.036912028 | 0.965298812 |
| Q810U3 | Nfasc | Neurofascin | 0.103088156 | 0.352467696 |
| Q811C2 | Atg4c | Cysteine protease ATG4C | -2.368456333 | -1.077321256 |
| Q811J3 | Ireb2 | Iron-responsive element-binding protein 2 | -0.415608597 | -0.20205005 |
| Q811P8 | Arhgap32 | Rho GTPase-activating protein 32 | -0.100567118 | -0.53407828 |
| Q8BFP9 | Pdk1 | [Pyruvate dehydrogenase (acetyl-transferring)] kinase isozyme 1, mitochondrial | 1.388220167 | -0.00076739 |
| Q8BFQ4 | Wdr82 | WD repeat-containing protein 82 | -0.071194299 | -0.19628032 |
| Q8BFQ8 | Gatd1 | Glutamine amidotransferase-like class 1 domain-containing protein 1 | 0.039971987 | 0.042501609 |
| Q8BFR4 | Gns | N-acetylglucosamine-6-sulfatase | -0.010961501 | -0.198413054 |
| Q8BFR5 | Tufm | Elongation factor Tu, mitochondrial | -0.112779299 | 0.324631055 |
| Q8BFS6 | Cpped1 | Serine/threonine-protein phosphatase CPPED1 | -0.375229073 | -0.034481366 |
| Q8BFU3 | Rnf214 | RING finger protein 214 | 0.109148916 | 0.477322896 |
| Q8BFV2 | Pcid2 | PCI domain-containing protein 2 | -0.005099869 | 0.352238496 |
| Q8BFW7 | Lpp | Lipoma-preferred partner homolog | -0.104405403 | 0.362423102 |
| Q8BFY6 | Pef1 | Peflin | 0.137917709 | 0.532099088 |
| Q8BFY9 | Tnpo1 | Transportin-1 | 0.241189671 | 0.79087925 |
| Q8BFZ3 | Actbl2 | Beta-actin-like protein 2 | -0.610813459 | 4.182333151 |
| Q8BFZ9 | Erlin2 | Erlin-2 | 1.182799021 | 0.837165674 |
| Q8BG02 | Ppp2r2c | Serine/threonine-protein phosphatase 2A 55 kDa regulatory subunit B gamma isoform | -0.871231778 | 1.686516672 |
| Q8BG18 | Necab1 | N-terminal EF-hand calcium-binding protein 1 | -0.004554272 | 0.504945119 |
| Q8BG32 | Psmd11 | 26S proteasome non-ATPase regulatory subunit 11 | 0.187581507 | -0.049268087 |
| Q8BG39 | Sv2b | Synaptic vesicle glycoprotein 2B | 0.82773908 | 0.004124959 |
| Q8BG40 | Katnb1 | Katanin p80 WD40 repeat-containing subunit B1 | -0.216057301 | 0.425825596 |
| Q8BG92 | Clvs2 | Clavesin-2 | 0.314795526 | 1.213814894 |

| | | | | |
|---|---|---|---|---|
| Q8BG93 | Nudt15 | Nucleotide triphosphate diphosphatase NUDT15 | -0.887290096 | 0.225235144 |
| Q8BG95 | Ppp1r12b | Protein phosphatase 1 regulatory subunit 12B | 0.161867491 | 1.026614507 |
| Q8BGB7 | Enoph1 | Enolase-phosphatase E1 | -0.081519159 | -0.900074323 |
| Q8BGC0 | Htatsf1 | HIV Tat-specific factor 1 homolog | 0.124332205 | 0.280557315 |
| Q8BGC4 | Zadh2 | Prostaglandin reductase-3 | -0.20078853 | 0.95217514 |
| Q8BGD8 | Coa6 | Cytochrome c oxidase assembly factor 6 homolog | 0.227926763 | 0.632167657 |
| Q8BGD9 | Eif4b | Eukaryotic translation initiation factor 4B | -0.098432255 | 0.087754091 |
| Q8BGF0 | AW551984 | Expressed sequence AW551984 | -0.716811975 | -0.447520574 |
| Q8BGG7 | Ubash3b | Ubiquitin-associated and SH3 domain-containing protein B | 0.251265717 | 1.562318484 |
| Q8BGH2 | Samm50 | Sorting and assembly machinery component 50 homolog | -0.027670701 | 1.820824941 |
| Q8BGJ5 | Ptbp1 | Polypyrimidine tract-binding protein 1 | 0.896607717 | 0.634516756 |
| Q8BGN2 | D3Ertd751e | UPF0462 protein C4orf33 homolog | -0.108764521 | 0.277912458 |
| Q8BGQ1 | Vipas39 | Spermatogenesis-defective protein 39 homolog | -1.161328411 | -0.086910407 |
| Q8BGQ7 | Aars1 | Alanine--tRNA ligase, cytoplasmic | -0.111465263 | 0.183178902 |
| Q8BGR6 | Arl15 | ADP-ribosylation factor-like protein 15 | 0.072501723 | 0.647528807 |
| Q8BGR9 | Ublcp1 | Ubiquitin-like domain-containing CTD phosphatase 1 | -0.304408805 | 0.689018567 |
| Q8BGS2 | Bola2 | BolA-like protein 2 | -0.837822628 | 0.225236098 |
| Q8BGT5 | Gpt2 | Alanine aminotransferase 2 | -0.13554519 | 0.192231496 |
| Q8BGT6 | Micall1 | MICAL-like protein 1 | -0.076082993 | 0.652941068 |
| Q8BGT8 | Phyhipl | Phytanoyl-CoA hydroxylase-interacting protein-like | -0.149734116 | 0.383816401 |
| Q8BGU5 | Ccny | Cyclin-Y | 0.204346307 | 2.183387756 |
| Q8BGW1 | Fto | Alpha-ketoglutarate-dependent dioxygenase FTO | -0.260881805 | 0.192595641 |
| Q8BGY2 | Eif5a2 | Eukaryotic translation initiation factor 5A-2 | -0.093474197 | 0.474980036 |
| Q8BGZ1 | Hpcal4 | Hippocalcin-like protein 4 | 0.002998606 | 0.56670475 |
| Q8BGZ7 | Krt75 | Keratin, type II cytoskeletal 75 | -0.054673386 | -0.150022507 |
| Q8BH44 | Coro2b | Coronin-2B | 0.057575448 | -0.804594358 |
| Q8BH55 | Thnsl1 | Threonine synthase-like 1 | 0.080729612 | 0.174511115 |
| Q8BH57 | Wdr48 | WD repeat-containing protein 48 | 0.05712293 | 0.123665333 |
| Q8BH58 | Tiprl | TIP41-like protein | 0.00228459 | 0.393550873 |
| Q8BH59 | Slc25a12 | Calcium-binding mitochondrial carrier protein Aralar1 | 0.289130179 | 1.49694411 |
| Q8BH61 | F13a1 | Coagulation factor XIII A chain | 0.619049358 | 0.242031892 |
| Q8BH66 | Atl1 | Atlastin-1 | 0.533389409 | 0.027565161 |
| Q8BH69 | Sephs1 | Selenide, water dikinase 1 | 0.210608832 | -0.148662408 |
| Q8BH70 | Fbxl4 | F-box/LRR-repeat protein 4 | -3.917089196 | -1.997200807 |
| Q8BH80 | Vapb | Vesicle-associated membrane protein, associated protein B and C | 0.00260458 | 0.796519121 |
| Q8BH95 | Echs1 | Enoyl-CoA hydratase, mitochondrial | 0.127539221 | 0.341389497 |
| Q8BHB9 | Clic6 | Chloride intracellular channel protein 6 | 0.121612263 | 0.259355704 |
| Q8BHE3 | Atcay | Caytaxin | -0.170726776 | 0.084390958 |
| Q8BHH2 | Rab9b | Ras-related protein Rab-9B | 1.070349439 | 2.964892069 |
| Q8BHI4 | Kbtbd3 | Kelch repeat and BTB domain-containing protein 3 | 0.160344156 | 0.119229635 |
| Q8BHJ5 | Tbl1xr1 | F-box-like/WD repeat-containing protein TBL1XR1 | 1.1738259 | 1.773448865 |
| Q8BHL3 | Tbc1d10b | TBC1 domain family member 10B | 0.569266129 | 0.514144739 |
| Q8BHL5 | Elmo2 | Engulfment and cell motility protein 2 | 0.082115618 | 0.51341629 |
| Q8BHL8 | Psmf1 | Proteasome inhibitor PI31 subunit | 0.327966118 | -0.781153679 |
| Q8BHN3 | Ganab | Neutral alpha-glucosidase AB | 0.133228588 | 0.437770526 |

| | | | | |
|---|---|---|---|---|
| Q8BHN7 | | Uncharacterized protein C12orf29 homolog | -1.140007019 | 0.203687509 |
| Q8BHS6 | Armcx3 | Armadillo repeat-containing X-linked protein 3 | 0.000464662 | 0.521597385 |
| Q8BHZ0 | Cyria | CYFIP-related Rac1 interactor A | 1.400999546 | 2.485837777 |
| Q8BI08 | Mal2 | Protein MAL2 | 0.044056543 | -2.496575435 |
| Q8BI72 | Cdkn2aip | CDKN2A-interacting protein | -0.328268115 | 0.399813652 |
| Q8BIF2 | Rbfox3 | RNA binding protein fox-1 homolog 3 | -0.47727677 | 0.842040539 |
| Q8BIJ6 | Iars2 | Isoleucine--tRNA ligase, mitochondrial | -0.329563109 | 0.204429309 |
| Q8BIJ7 | Rufy1 | RUN and FYVE domain-containing protein 1 | -0.488938999 | 0.026425838 |
| Q8BIP0 | Dars2 | Aspartate--tRNA ligase, mitochondrial | -0.13123134 | 1.167041063 |
| Q8BIQ6 | Zfp947 | Predicted gene, EG210853 | 0.466498788 | 0.204500993 |
| Q8BIQ9 | Prkag2 | 5'-AMP-activated protein kinase subunit gamma-2 | -0.773742739 | 0.886057695 |
| Q8BIV3 | Ranbp6 | Ran-binding protein 6 | -0.811003304 | 0.682578405 |
| Q8BIW1 | Prune1 | Exopolyphosphatase PRUNE1 | -0.066274484 | 0.216100534 |
| Q8BIZ1 | Anks1b | Ankyrin repeat and sterile alpha motif domain-containing protein 1B | 0.191554228 | -0.203473409 |
| Q8BJ71 | Nup93 | Nuclear pore complex protein Nup93 | 0.212963518 | -0.285391649 |
| Q8BJH1 | Zc2hc1a | Zinc finger C2HC domain-containing protein 1A | -0.228832912 | -0.055965583 |
| Q8BJU0 | Sgta | Small glutamine-rich tetratricopeptide repeat-containing protein alpha | -0.141247145 | 0.722181638 |
| Q8BJW6 | Eif2a | Eukaryotic translation initiation factor 2A | 1.087608624 | 0.716138124 |
| Q8BJY1 | Psmd5 | 26S proteasome non-ATPase regulatory subunit 5 | 0.0885547 | 1.239552339 |
| Q8BK64 | Ahsa1 | Activator of 90 kDa heat shock protein ATPase homolog 1 | -0.080615075 | 0.292329311 |
| Q8BK67 | Rcc2 | Protein RCC2 | 0.314540863 | 0.064171155 |
| Q8BKC5 | Ipo5 | Importin-5 | -0.06943922 | 0.359156291 |
| Q8BKG3 | Ptk7 | Inactive tyrosine-protein kinase 7 | 0.290457185 | -0.855557442 |
| Q8BKP1 | Tpd52l2 | Tumor protein D54 | -0.022621377 | -0.434201717 |
| Q8BKR5 | Ppp1r37 | Protein phosphatase 1 regulatory subunit 37 | -1.090636253 | -0.285182476 |
| Q8BKY8 | Mterf2 | Transcription termination factor 2, mitochondrial | 0.12603639 | 0.253593286 |
| Q8BKZ9 | Pdhx | Pyruvate dehydrogenase protein X component, mitochondrial | 0.291837025 | 0.173852126 |
| Q8BL65 | Ablim2 | Actin-binding LIM protein 2 | -0.184917132 | 0.349423885 |
| Q8BL66 | Eea1 | Early endosome antigen 1 | 0.015165933 | 0.30761528 |
| Q8BLF1 | Nceh1 | Neutral cholesterol ester hydrolase 1 | -1.710932954 | -0.812528928 |
| Q8BLR2 | Cpne4 | Copine-4 | -0.025202306 | 0.447949886 |
| Q8BLR9 | Hif1an | Hypoxia-inducible factor 1-alpha inhibitor | 0.054606088 | 0.467098236 |
| Q8BLY2 | Tars3 | Threonine--tRNA ligase 2, cytoplasmic | 0.056318537 | 1.272580783 |
| Q8BM72 | Hspa13 | Heat shock 70 kDa protein 13 | 0.514150842 | -0.234237989 |
| Q8BMA6 | Srp68 | Signal recognition particle subunit SRP68 | -0.00640281 | 0.476372878 |
| Q8BMF3 | Me3 | NADP-dependent malic enzyme, mitochondrial | -0.135707633 | -0.240033786 |
| Q8BMF4 | Dlat | Dihydrolipoyllysine-residue acetyltransferase component of pyruvate dehydrogenase complex, mitochondrial | 0.684811115 | 0.111936092 |
| Q8BMJ2 | Lars1 | Leucine--tRNA ligase, cytoplasmic | 0.370909437 | 0.777165731 |
| Q8BMJ3 | Eif1ax | Eukaryotic translation initiation factor 1A, X-chromosomal | 0.114420255 | 0.523802757 |
| Q8BMS1 | Hadha | Trifunctional enzyme subunit alpha, mitochondrial | -0.059902318 | 0.141976515 |
| Q8BMS9 | Rassf2 | Ras association domain-containing protein 2 | -0.587350623 | 0.457927704 |
| Q8BN51 | Tk2 | Thymidine kinase 2, mitochondrial | -0.02887907 | 0.447208087 |
| Q8BN59 | Larp6 | La-related protein 6 | 0.701721064 | 0.708542903 |
| Q8BNN1 | Spata2l | Spermatogenesis-associated protein 2-like protein | 0.694874287 | 0.25734059 |
| Q8BNU0 | Armc6 | Armadillo repeat-containing protein 6 | -0.004449272 | 0.088895162 |

| | | | | |
|---|---|---|---|---|
| Q8BNW9 | Kbtbd11 | Kelch repeat and BTB domain-containing protein 11 | 0.002860387 | 0.264926434 |
| Q8BNY6 | Ncs1 | Neuronal calcium sensor 1 | 0.090812747 | 0.475630124 |
| Q8BP00 | Iqcb1 | IQ calmodulin-binding motif-containing protein 1 | 0.095499579 | 0.04186511 |
| Q8BP27 | Sfr1 | Swi5-dependent recombination DNA repair protein 1 homolog | 0.215318998 | 0.788536231 |
| Q8BP40 | Acp6 | Lysophosphatidic acid phosphatase type 6 | 0.226416111 | 0.609262149 |
| Q8BP47 | NARS1 | Asparagine--tRNA ligase, cytoplasmic | 0.021622976 | 0.383698781 |
| Q8BP48 | Metap1 | Methionine aminopeptidase 1 | -0.538866297 | -0.172587077 |
| Q8BP56 | Pgghg | Protein-glucosylgalactosylhydroxylysine glucosidase | -0.330820417 | 0.260459105 |
| Q8BP67 | Rpl24 | 60S ribosomal protein L24 | 0.44317131 | 1.162758986 |
| Q8BP92 | Rcn2 | Reticulocalbin-2 | -0.429462592 | -0.485616843 |
| Q8BPA8 | Dpcd | Protein DPCD | -0.923490938 | -1.115410169 |
| Q8BPG6 | Sumf2 | Inactive C-alpha-formylglycine-generating enzyme 2 | -0.671592585 | -0.906045437 |
| Q8BPN8 | Dmxl2 | DmX-like protein 2 | 0.028433228 | 0.028101762 |
| Q8BPU7 | Elmo1 | Engulfment and cell motility protein 1 | 0.728471279 | 0.355749766 |
| Q8BQP9 | Rgs7bp | Regulator of G-protein signaling 7-binding protein | -1.045146306 | 1.088516633 |
| Q8BQV2 | Chat | Choline O-acetyltransferase | 0.738257535 | 0.962938468 |
| Q8BR63 | Fam177a1 | Protein FAM177A1 | 0.350548458 | 1.214540005 |
| Q8BR90 | | UPF0600 protein C5orf51 homolog | -0.733008226 | 1.01380221 |
| Q8BR92 | Palm2 | Paralemmin-2 | 0.080664698 | 0.444057782 |
| Q8BRF7 | Scfd1 | Sec1 family domain-containing protein 1 | 0.534486548 | 1.38579917 |
| Q8BRK8 | Prkaa2 | 5'-AMP-activated protein kinase catalytic subunit alpha-2 | 0.026793067 | -0.866528193 |
| Q8BS40 | Cptp | Ceramide-1-phosphate transfer protein | -0.506152185 | 0.636813482 |
| Q8BSK8 | Rps6kb1 | Ribosomal protein S6 kinase beta-1 | -0.230538591 | 2.057852268 |
| Q8BSL7 | Arf2 | ADP-ribosylation factor 2 | 0.248879147 | 0.205346743 |
| Q8BSZ2 | Ap3s2 | AP-3 complex subunit sigma-2 | 0.274848048 | 0.882814089 |
| Q8BT60 | Cpne3 | Copine-3 | 0.062125492 | 0.409797986 |
| Q8BTG3 | Tcp11l1 | T-complex protein 11-like protein 1 | 0.469493961 | 1.66875114 |
| Q8BTG7 | Ndrg4 | Protein NDRG4 | 0.087691752 | 0.616393248 |
| Q8BTI8 | Srrm2 | Serine/arginine repetitive matrix protein 2 | -0.397304789 | -0.112341722 |
| Q8BTI9 | Pik3cb | Phosphatidylinositol 4,5-bisphosphate 3-kinase catalytic subunit beta isoform | 0.247688675 | 0.644475301 |
| Q8BTR5 | Dusp28 | Dual specificity phosphatase 28 | 0.306079578 | -0.105065028 |
| Q8BTS4 | Nup54 | Nuclear pore complex protein Nup54 | -0.553299872 | -0.647776763 |
| Q8BTU1 | Cfap20 | Cilia- and flagella-associated protein 20 | -1.008282407 | -0.241309961 |
| Q8BTV2 | Cpsf7 | Cleavage and polyadenylation specificity factor subunit 7 | 0.495920022 | -0.491909663 |
| Q8BTW3 | Exosc6 | Exosome complex component MTR3 | 0.656770039 | 0.340975285 |
| Q8BTY1 | Kyat1 | Kynurenine--oxoglutarate transaminase 1 | -0.041560459 | 0.679425557 |
| Q8BTY8 | Scfd2 | Sec1 family domain-containing protein 2 | -0.109877268 | 0.788179954 |
| Q8BTZ7 | Gmppb | Mannose-1-phosphate guanyltransferase beta | 0.050008106 | 0.225201607 |
| Q8BU30 | Iars1 | Isoleucine--tRNA ligase, cytoplasmic | 0.342742856 | -2.303542147 |
| Q8BU31 | Rap2c | Ras-related protein Rap-2c | 0.51945672 | 0.301744143 |
| Q8BUK6 | Hook3 | Protein Hook homolog 3 | 0.284744008 | 0.448263963 |
| Q8BUY8 | Gprasp2 | G-protein coupled receptor-associated sorting protein 2 | 0.432216422 | 0.369824648 |
| Q8BV13 | Cops7b | COP9 signalosome complex subunit 7b | 0.620962207 | -0.281792641 |
| Q8BVG4 | Dpp9 | Dipeptidyl peptidase 9 | 0.330816619 | -0.28583622 |
| Q8BVI4 | Qdpr | Dihydropteridine reductase | -0.224942843 | 0.537818909 |

| | | | | |
|---|---|---|---|---|
| Q8BVL3 | Snx17 | Sorting nexin-17 | -0.298909632 | -0.3445762 |
| Q8BVQ5 | Ppme1 | Protein phosphatase methylesterase 1 | -0.105286312 | 0.218147119 |
| Q8BVU5 | Nudt9 | ADP-ribose pyrophosphatase, mitochondrial | -0.351451524 | -0.456127961 |
| Q8BW22 | Ss18l1 | Calcium-responsive transactivator | 0.303496726 | 0.887789965 |
| Q8BW86 | Arhgef33 | Rho guanine nucleotide exchange factor 33 | -1.105587228 | 0.273666064 |
| Q8BW96 | Camk1d | Calcium/calmodulin-dependent protein kinase type 1D | -0.372634029 | -0.114926497 |
| Q8BWF0 | Aldh5a1 | Succinate-semialdehyde dehydrogenase, mitochondrial | -0.16963431 | 0.051366488 |
| Q8BWP8 | B4gat1 | Beta-1,4-glucuronyltransferase 1 | -0.001312733 | 0.299232642 |
| Q8BWQ6 | Vps35l | VPS35 endosomal protein-sorting factor-like | -0.016478825 | -1.181271712 |
| Q8BWR2 | Pithd1 | PITH domain-containing protein 1 | -0.175303586 | 0.335299174 |
| Q8BWS5 | Gprin3 | G protein-regulated inducer of neurite outgrowth 3 | 0.318308417 | 0.396136443 |
| Q8BWT1 | Acaa2 | 3-ketoacyl-CoA thiolase, mitochondrial | -0.096791172 | 0.782644749 |
| Q8BWT5 | Dip2a | Disco-interacting protein 2 homolog A | -0.308169397 | -0.340144793 |
| Q8BWW3 | Pgm3 | Phosphoacetylglucosamine mutase | -0.134230487 | 0.31534481 |
| Q8BWY3 | Etf1 | Eukaryotic peptide chain release factor subunit 1 | 0.477100436 | 0.728235722 |
| Q8BWY4 | Mblac1 | Metallo-beta-lactamase domain-containing protein 1 | -0.131287225 | 0.625902494 |
| Q8BWZ3 | Naa25 | N-alpha-acetyltransferase 25, NatB auxiliary subunit | -0.117849223 | -0.296561718 |
| Q8BX02 | Kank2 | KN motif and ankyrin repeat domain-containing protein 2 | 0.363099257 | 0.312647661 |
| Q8BX70 | Vps13c | Vacuolar protein sorting-associated protein 13C | -0.022667726 | 0.899317582 |
| Q8BX94 | Osbpl2 | Oxysterol-binding protein-related protein 2 | -0.461566703 | 0.04561917 |
| Q8BXC6 | Commd2 | COMM domain-containing protein 2 | -0.127258428 | -0.103215694 |
| Q8BXN7 | Ppm1k | Protein phosphatase 1K, mitochondrial | -0.54669536 | 1.341994603 |
| Q8BXR9 | Osbpl6 | Oxysterol-binding protein-related protein 6 | 0.387300968 | 0.793394645 |
| Q8BYA0 | Tbcd | Tubulin-specific chaperone D | -0.149177933 | -0.169558843 |
| Q8BYH7 | Tbc1d17 | TBC1 domain family member 17 | -0.015610917 | 0.259803931 |
| Q8BYI9 | Tnr | Tenascin-R | -0.135923958 | 0.034891446 |
| Q8BYK6 | Ythdf3 | YTH domain-containing family protein 3 | 0.079560089 | 0.605550607 |
| Q8BYM8 | Cars2 | Probable cysteine--tRNA ligase, mitochondrial | -0.900481447 | 0.684776306 |
| Q8BYN3 | Itpk1 | Inositol-tetrakisphosphate 1-kinase | 0.194605128 | -0.473605951 |
| Q8BYW1 | Arhgap25 | Rho GTPase-activating protein 25 | 0.465131124 | -0.568192164 |
| Q8BYW9 | Eogt | EGF domain-specific O-linked N-acetylglucosamine transferase | -0.305868085 | -0.140700976 |
| Q8BYY4 | Ttc39b | Tetratricopeptide repeat protein 39B | 0.137214661 | -0.494461536 |
| Q8BZ98 | Dnm3 | Dynamin-3 | -0.495233472 | -0.289961338 |
| Q8BZA9 | Tigar | Fructose-2,6-bisphosphatase TIGAR | -0.814181964 | -0.432661057 |
| Q8BZB2 | Ppcdc | Phosphopantothenoylcysteine decarboxylase | -0.126968384 | 1.334772905 |
| Q8BZF8 | Pgm5 | Phosphoglucomutase-like protein 5 | 0.110592079 | 0.653280258 |
| Q8BZM1 | Glmn | Glomulin | 0.016647212 | 1.589565754 |
| Q8BZQ7 | Anapc2 | Anaphase-promoting complex subunit 2 | 3.799725056 | 1.666471799 |
| Q8BZR6 | Tom1l1 | TOM1-like protein 1 | -0.535867532 | 0.543732643 |
| Q8BZW8 | Nhlrc2 | NHL repeat-containing protein 2 | -0.025385157 | 0.471206506 |
| Q8BZZ3 | Wwp1 | NEDD4-like E3 ubiquitin-protein ligase WWP1 | 0.078899542 | 0.052646478 |
| Q8C033 | Arhgef10 | Rho guanine nucleotide exchange factor 10 | -0.466099644 | 1.083836953 |
| Q8C050 | Rps6ka5 | Ribosomal protein S6 kinase alpha-5 | -0.369280497 | 0.035712083 |
| Q8C052 | Map1s | Microtubule-associated protein 1S | 0.783590094 | -0.196927547 |
| Q8C078 | Camkk2 | Calcium/calmodulin-dependent protein kinase kinase 2 | -0.094486078 | 0.135866324 |
| Q8C079 | Strip1 | Striatin-interacting protein 1 | -0.456082821 | 0.28748099 |

| | | | | |
|---|---|---|---|---|
| Q8C080 | Snx16 | Sorting nexin-16 | 0.485235087 | 0.780821164 |
| Q8C0D5 | Efl1 | Elongation factor-like GTPase 1 | -0.21337862 | -0.37327973 |
| Q8C0E2 | Vps26b | Vacuolar protein sorting-associated protein 26B | -0.03842446 | -0.141214212 |
| Q8C0J2 | Atg16l1 | Autophagy-related protein 16-1 | 0.173893007 | 0.357227484 |
| Q8C0L6 | Paox | Peroxisomal N(1)-acetyl-spermine/spermidine oxidase | -0.072026189 | 0.20812877 |
| Q8C0L9 | Gpcpd1 | Glycerophosphocholine phosphodiesterase GPCPD1 | 0.069943619 | -0.147298336 |
| Q8C0M9 | Asrgl1 | Isoaspartyl peptidase/L-asparaginase | -0.318671354 | 0.244773547 |
| Q8C0P5 | Coro2a | Coronin-2A | 0.183656089 | -0.139715513 |
| Q8C0Y0 | Ppp4r4 | Serine/threonine-protein phosphatase 4 regulatory subunit 4 | -0.250545406 | -1.220035175 |
| Q8C166 | Cpne1 | Copine-1 | -0.072697862 | 0.032712142 |
| Q8C167 | Prepl | Prolyl endopeptidase-like | -0.114505895 | 0.349875609 |
| Q8C1Y8 | Ccz1 | Vacuolar fusion protein CCZ1 homolog | -0.56908954 | 0.542846243 |
| Q8C2E7 | Washc5 | WASH complex subunit 5 | -0.731758118 | -0.184702237 |
| Q8C2Q3 | Rbm14 | RNA-binding protein 14 | 0.078797563 | -0.005248388 |
| Q8C2Q8 | Atp5c1 | ATP synthase subunit gamma | 0.317393716 | 0.815160751 |
| Q8C3I8 | Hgh1 | Protein HGH1 homolog | 0.15349261 | 0.587465763 |
| Q8C3W1 | | Uncharacterized protein C1orf198 homolog | 0.436323166 | 0.687178453 |
| Q8C419 | Gpr158 | Probable G-protein coupled receptor 158 | 0.42585001 | 0.363578637 |
| Q8C460 | Eri3 | ERI1 exoribonuclease 3 | 0.232151572 | 1.042383353 |
| Q8C4B4 | Unc119b | Protein unc-119 homolog B | 0.547692839 | 1.860544523 |
| Q8C4Q6 | Aida | Axin interactor, dorsalization-associated protein | 0.517901198 | -0.315965494 |
| Q8C522 | Endod1 | Endonuclease domain-containing 1 protein | -0.203235086 | 0.610269388 |
| Q8C547 | Heatr5b | HEAT repeat-containing protein 5B | -0.054802163 | -0.202531179 |
| Q8C570 | Rae1 | mRNA export factor | 0.259538905 | -0.079579035 |
| Q8C5H8 | Nadk2 | NAD kinase 2, mitochondrial | -0.107414532 | 0.296300252 |
| Q8C5K5 | | Uncharacterized protein CXorf38 homolog | -1.560977586 | 0.141998927 |
| Q8C5P7 | Tdrp | Testis development-related protein | 0.782183997 | 1.05349652 |
| Q8C5W0 | Clmn | Calmin | -0.168809827 | 0.329178492 |
| Q8C5W3 | Tbcel | Tubulin-specific chaperone cofactor E-like protein | 0.006645393 | -0.384305318 |
| Q8C605 | Pfkp | ATP-dependent 6-phosphofructokinase | -0.178648822 | -0.002505302 |
| Q8C6B0 | Mettl7a1 | Methyltransferase-like 7A1 | 0.466946475 | -0.358151505 |
| Q8C6B2 | Rtkn | Rhotekin | 0.011434873 | 0.10260884 |
| Q8C6E0 | Cfap36 | Cilia- and flagella-associated protein 36 | -0.286559232 | 0.198441664 |
| Q8C729 | Fam126b | Protein FAM126B | 0.343119367 | 0.988874912 |
| Q8C754 | Vps52 | Vacuolar protein sorting-associated protein 52 homolog | -0.35836455 | -2.918374976 |
| Q8C788 | Snx18 | Sorting nexin | -0.103353691 | 1.95364809 |
| Q8C7D2 | Crbn | Protein cereblon | -0.236252117 | 0.056039174 |
| Q8C7E4 | Rnase4 | Ribonuclease 4 | -0.075614675 | 0.890564124 |
| Q8C845 | Efhd2 | EF-hand domain-containing protein D2 | -0.081234423 | 0.198036194 |
| Q8C854 | Myef2 | Myelin expression factor 2 | 0.478128242 | -0.238602797 |
| Q8C863 | Itch | E3 ubiquitin-protein ligase Itchy | -0.662436517 | -1.288058678 |
| Q8C878 | Uba3 | NEDD8-activating enzyme E1 catalytic subunit | -0.080610275 | 0.349834442 |
| Q8C8N2 | Scai | Protein SCAI | -0.105301189 | 0.414747715 |
| Q8C8T8 | Tsr2 | Pre-rRNA-processing protein TSR2 homolog | 0.364516449 | 0.628343423 |
| Q8CA72 | Gan | Gigaxonin | 0.203093942 | 0.373336951 |
| Q8CAA7 | Pgm2l1 | Glucose 1,6-bisphosphate synthase | -0.094875717 | 0.2136844 |

| | | | | |
|---|---|---|---|---|
| Q8CAB8 | Castor2 | Cytosolic arginine sensor for mTORC1 subunit 2 | -0.090663083 | 0.209300995 |
| Q8CAK1 | Iba57 | Putative transferase CAF17 homolog, mitochondrial | -0.015131219 | 0.631553332 |
| Q8CAK3 | Shfl | Shiftless antiviral inhibitor of ribosomal frameshifting protein homolog | 0.071394094 | 0.526209195 |
| Q8CAQ8 | Immt | MICOS complex subunit Mic60 | 0.3687452 | 0.692931811 |
| Q8CAY6 | Acat2 | Acetyl-CoA acetyltransferase, cytosolic | -0.145860736 | 0.122007688 |
| Q8CB27 | Yod1 | Ubiquitin thioesterase OTU1 | 0.94323225 | -0.357859453 |
| Q8CBC8 | Bcat1 | Branched-chain-amino-acid aminotransferase | -0.25806516 | 0.605383873 |
| Q8CBE3 | Wdr37 | WD repeat-containing protein 37 | 0.113282077 | 0.255950133 |
| Q8CBY8 | Dctn4 | Dynactin subunit 4 | 0.101091925 | 0.487111092 |
| Q8CC21 | Ttc19 | Tetratricopeptide repeat protein 19, mitochondrial | 0.560514005 | -0.325310548 |
| Q8CC86 | Naprt | Nicotinate phosphoribosyltransferase | -0.105428664 | 0.86686182 |
| Q8CC88 | Vwa8 | von Willebrand factor A domain-containing protein 8 | 0.126666164 | -0.926846186 |
| Q8CCB4 | Vps53 | Vacuolar protein sorting-associated protein 53 homolog | 0.035052872 | 0.079567432 |
| Q8CCF0 | Prpf31 | U4/U6 small nuclear ribonucleoprotein Prp31 | -1.05769186 | -0.173776309 |
| Q8CCP0 | Nemf | Nuclear export mediator factor Nemf | -1.042628765 | 0.816792965 |
| Q8CCT4 | Tceal5 | Transcription elongation factor A protein-like 5 | -0.294230747 | 0.219264984 |
| Q8CD19 | Lancl3 | LanC-like protein 3 | 0.953286076 | 0.415565968 |
| Q8CD92 | Ttc27 | Tetratricopeptide repeat protein 27 | 0.758433119 | 0.379905383 |
| Q8CDA1 | Inpp5f | Phosphatidylinositide phosphatase SAC2 | 0.063158258 | 0.4544185 |
| Q8CDM8 | Fam160b1 | Protein FAM160B1 | 0.268674469 | 0.028145631 |
| Q8CE50 | Snx30 | Sorting nexin-30 | -0.039411418 | 0.50533088 |
| Q8CE90 | Map2k7 | Dual specificity mitogen-activated protein kinase kinase 7 | -1.499874369 | 1.29652818 |
| Q8CF66 | Lamtor4 | Ragulator complex protein LAMTOR4 | 0.177798208 | 1.192450841 |
| Q8CF89 | Tab1 | TGF-beta-activated kinase 1 and MAP3K7-binding protein 1 | -0.124369685 | 0.435895602 |
| Q8CFR5 | Dtna | Dystrobrevin | -0.017016792 | 0.432409922 |
| Q8CFV9 | Rfk | Riboflavin kinase | -0.450981331 | -0.356180509 |
| Q8CG72 | Adprs | ADP-ribose glycohydrolase ARH3 | -0.414118512 | 1.12614584 |
| Q8CG76 | Akr7a2 | Aflatoxin B1 aldehyde reductase member 2 | -0.118587112 | 0.170492808 |
| Q8CGA0 | Ppm1f | Protein phosphatase 1F | -0.163475037 | 0.230916818 |
| Q8CGC7 | Eprs1 | Bifunctional glutamate/proline--tRNA ligase | -0.086045043 | -0.009633541 |
| Q8CGF6 | Wdr47 | WD repeat-containing protein 47 | 0.014553897 | 0.108405908 |
| Q8CGK3 | Lonp1 | Lon protease homolog, mitochondrial | -0.155716515 | 0.30804348 |
| Q8CGP0 | H2bu1 | Histone H2B type 3-B | 0.361374029 | 1.818208059 |
| Q8CGV2 | Tph2 | Tryptophan 5-hydroxylase 2 | -0.874079227 | 0.368160089 |
| Q8CGY8 | Ogt | UDP-N-acetylglucosamine--peptide N-acetylglucosaminyltransferase 110 kDa subunit | 0.120527617 | 0.374338786 |
| Q8CH02 | Sugp1 | SURP and G-patch domain-containing protein 1 | 0.998266474 | -0.214963913 |
| Q8CH18 | Ccar1 | Cell division cycle and apoptosis regulator protein 1 | -0.33401432 | -0.896574815 |
| Q8CH72 | Trim32 | E3 ubiquitin-protein ligase TRIM32 | -0.015533606 | 0.488903046 |
| Q8CHP5 | Pym1 | Partner of Y14 and mago | 0.386111387 | -0.785372893 |
| Q8CHP8 | Pgp | Glycerol-3-phosphate phosphatase | -0.080153529 | 0.545369784 |
| Q8CHQ0 | Fbxo4 | F-box only protein 4 | -0.164615059 | 0.110329946 |
| Q8CHS8 | Vps37a | Vacuolar protein sorting-associated protein 37A | 0.591400464 | -0.20203495 |
| Q8CHT0 | Aldh4a1 | Delta-1-pyrroline-5-carboxylate dehydrogenase, mitochondrial | -0.232975578 | -0.052076181 |
| Q8CHU3 | Epn2 | Epsin-2 | 0.153995482 | 0.550879161 |
| Q8CHW4 | Eif2b5 | Translation initiation factor eIF-2B subunit epsilon | 0.399991385 | 0.513252258 |

| | | | | |
|---|---|---|---|---|
| Q8CHX7 | Rftn2 | Raftlin-2 | -0.132441648 | 0.668132782 |
| Q8CHY3 | Dym | Dymeclin | -0.185787837 | 0.525414467 |
| Q8CI32 | Bag5 | BAG family molecular chaperone regulator 5 | 0.030359936 | 0.00157245 |
| Q8CI43 | Myl6b | Myosin light chain 6B | 0.650672754 | -1.584209919 |
| Q8CI51 | Pdlim5 | PDZ and LIM domain protein 5 | 0.205829334 | -0.198307673 |
| Q8CI61 | Bag4 | BAG family molecular chaperone regulator 4 | -0.154522419 | 0.268816789 |
| Q8CI70 | Lrrc20 | Leucine-rich repeat-containing protein 20 | -0.143248272 | -0.898879051 |
| Q8CI71 | Vps50 | Syndetin | 0.43571043 | 1.072530746 |
| Q8CI75 | Dis3l2 | DIS3-like exonuclease 2 | -0.090810776 | -0.324402968 |
| Q8CI94 | Pygb | Glycogen phosphorylase, brain form | -0.342697525 | 0.088403384 |
| Q8CIB5 | Fermt2 | Fermitin family homolog 2 | -0.306264559 | 0.23811388 |
| Q8CIH9 | Ppat | Amidophosphoribosyltransferase | -0.083696651 | -0.256107648 |
| Q8CII2 | Cdc123 | Cell division cycle protein 123 homolog | -0.151696841 | 0.499389489 |
| Q8CIN4 | Pak2 | Serine/threonine-protein kinase PAK 2 | 0.085457007 | 0.568425814 |
| Q8CIV8 | Tbce | Tubulin-specific chaperone E | -0.180818272 | -0.070213954 |
| Q8CJ40 | Crocc | Rootletin | 1.586733532 | 1.126062075 |
| Q8CJG0 | Ago2 | Protein argonaute-2 | 0.846697776 | 0.275514603 |
| Q8JZK9 | Hmgcs1 | Hydroxymethylglutaryl-CoA synthase, cytoplasmic | -0.128525925 | 0.747345765 |
| Q8JZL3 | Thtpa | Thiamine-triphosphatase | -0.215628783 | -0.123932362 |
| Q8JZM7 | Cdc73 | Parafibromin | 0.6394859 | -0.067053795 |
| Q8JZN5 | Acad9 | Complex I assembly factor ACAD9, mitochondrial | -0.170466455 | 0.302189986 |
| Q8JZP2 | Syn3 | Synapsin-3 | 0.209610844 | 0.197952429 |
| Q8JZQ2 | Afg3l2 | AFG3-like protein 2 | 0.608994675 | 1.011293093 |
| Q8JZQ9 | Eif3b | Eukaryotic translation initiation factor 3 subunit B | 0.020618661 | -0.030973434 |
| Q8JZR2 | Crk | Adapter molecule crk | -0.054295286 | 1.276694854 |
| Q8JZS0 | Lin7a | Protein lin-7 homolog A | 0.032500076 | 0.202963829 |
| Q8JZW4 | Cpne5 | Copine-5 | -0.225939655 | 0.689935843 |
| Q8JZW5 | Sh2d5 | SH2 domain-containing protein 5 | -0.253377628 | 1.004168193 |
| Q8JZX4 | Rbm17 | Splicing factor 45 | -1.199905841 | 0.311106841 |
| Q8JZX9 | Cdc42ep2 | Cdc42 effector protein 2 | 0.933548641 | 0.569840193 |
| Q8JZZ5 | Pitpnb | Phosphatidylinositol transfer protein beta isoform | -0.18499279 | 0.639811993 |
| Q8K004 | Spata2 | Spermatogenesis-associated protein 2 | 0.301771164 | -0.605805477 |
| Q8K010 | Oplah | 5-oxoprolinase | -0.148788452 | 0.143495242 |
| Q8K070 | Samd14 | Sterile alpha motif domain-containing protein 14 | -0.257280858 | 0.743020217 |
| Q8K0C9 | Gmds | GDP-mannose 4,6 dehydratase | -0.152428786 | 0.412470182 |
| Q8K0D5 | Gfm1 | Elongation factor G, mitochondrial | -0.07172486 | 0.626548449 |
| Q8K0E8 | Fgb | Fibrinogen beta chain | 0.306364409 | 1.24892664 |
| Q8K0G5 | Eipr1 | EARP and GARP complex-interacting protein 1 | -0.052587891 | 0.070620537 |
| Q8K0S0 | Phyhip | Phytanoyl-CoA hydroxylase-interacting protein | -0.176099714 | 0.050858816 |
| Q8K0T0 | Rtn1 | Reticulon-1 | 0.009141699 | 0.421490987 |
| Q8K0T4 | Katnal1 | Katanin p60 ATPase-containing subunit A-like 1 | 0.225892131 | 0.916118781 |
| Q8K0U4 | Hspa12a | Heat shock 70 kDa protein 12A | 0.005013021 | 0.120767593 |
| Q8K0V4 | Cnot3 | CCR4-NOT transcription complex subunit 3 | 0.356858571 | 0.00236702 |
| Q8K0X8 | Fez1 | Fasciculation and elongation protein zeta-1 | 0.779892763 | 0.785654863 |
| Q8K0Z7 | Taco1 | Translational activator of cytochrome c oxidase 1 | 0.21195701 | 0.430428505 |
| Q8K124 | Plekho2 | Pleckstrin homology domain-containing family O member 2 | 0.862951597 | 0.768147469 |

| | | | | |
|---|---|---|---|---|
| Q8K157 | Galm | Galactose mutarotase | -0.101082993 | 0.504035791 |
| Q8K183 | Pdxk | Pyridoxal kinase | -0.357738177 | -0.069083214 |
| Q8K1J6 | Trnt1 | CCA tRNA nucleotidyltransferase 1, mitochondrial | 0.06977307 | 0.223182519 |
| Q8K1M6 | Dnm1l | Dynamin-1-like protein | -0.104074923 | 0.146929423 |
| Q8K1R3 | Pnpt1 | Polyribonucleotide nucleotidyltransferase 1, mitochondrial | -0.696231842 | -0.304948966 |
| Q8K1R7 | Nek9 | Serine/threonine-protein kinase Nek9 | 0.012953377 | 0.266138077 |
| Q8K1Z0 | Coq9 | Ubiquinone biosynthesis protein COQ9, mitochondrial | -0.008801937 | 0.675560474 |
| Q8K212 | Pacs1 | Phosphofurin acidic cluster sorting protein 1 | 0.02344443 | 0.370788415 |
| Q8K215 | Lyrm4 | LYR motif-containing protein 4 | -0.083842119 | -0.278153578 |
| Q8K268 | Abcf3 | ATP-binding cassette sub-family F member 3 | 0.832872327 | -1.727993965 |
| Q8K274 | Fn3krp | Ketosamine-3-kinase | -0.001858075 | 0.22122256 |
| Q8K298 | Anln | Anillin | 0.148200353 | 1.345070521 |
| Q8K2B3 | Sdha | Succinate dehydrogenase [ubiquinone] flavoprotein subunit, mitochondrial | 0.076145299 | 0.288562775 |
| Q8K2C6 | Sirt5 | NAD-dependent protein deacylase sirtuin-5, mitochondrial | -0.542811807 | 0.502058824 |
| Q8K2C9 | Hacd3 | Very-long-chain (3R)-3-hydroxyacyl-CoA dehydratase 3 | 0.657298628 | 0.462436676 |
| Q8K2D3 | Edc3 | Enhancer of mRNA-decapping protein 3 | -0.266074371 | 0.851089001 |
| Q8K2D8 | Fibp | Acidic fibroblast growth factor intracellular-binding protein | -1.013722388 | 0.466014703 |
| Q8K2F8 | Lsm14a | Protein LSM14 homolog A | -0.629163965 | -0.093486627 |
| Q8K2I1 | Fntb | Protein farnesyltransferase subunit beta | 0.103578758 | 0.406298637 |
| Q8K2L8 | Trappc12 | Trafficking protein particle complex subunit 12 | -0.17356898 | 1.078652859 |
| Q8K2P6 | Rfesd | Rieske domain-containing protein | 0.283133666 | 0.290521781 |
| Q8K2Q7 | Brox | BRO1 domain-containing protein BROX | -0.042364375 | 0.675657113 |
| Q8K2Q9 | Shtn1 | Shootin-1 | 0.330892245 | 0.290789604 |
| Q8K2T8 | Paf1 | RNA polymerase II-associated factor 1 homolog | -1.156661844 | 0.643885692 |
| Q8K2V6 | Ipo11 | Importin-11 | 0.062808132 | -0.734014988 |
| Q8K310 | Matr3 | Matrin-3 | -0.012857087 | 0.189569155 |
| Q8K337 | Inpp5b | Type II inositol 1,4,5-trisphosphate 5-phosphatase | -0.457377497 | -0.015350342 |
| Q8K341 | Atat1 | Alpha-tubulin N-acetyltransferase 1 | -0.025021458 | 1.110980352 |
| Q8K354 | Cbr3 | Carbonyl reductase [NADPH] 3 | -0.086393197 | 0.497793198 |
| Q8K356 | Ly6h | Lymphocyte antigen 6 complex, locus H | 0.593301519 | 1.183066368 |
| Q8K382 | Dennd1a | DENN domain-containing protein 1A | -1.521039486 | -0.61286815 |
| Q8K394 | Plcl2 | Inactive phospholipase C-like protein 2 | -0.25971489 | 0.348484516 |
| Q8K3C3 | Lzic | Protein LZIC | 0.10966905 | -0.213016033 |
| Q8K3G9 | Appl2 | DCC-interacting protein 13-beta | -0.132514699 | 0.354853471 |
| Q8K3H0 | Appl1 | DCC-interacting protein 13-alpha | 0.059927622 | -0.010603269 |
| Q8K3K8 | Optn | Optineurin | -1.891958046 | 0.312669436 |
| Q8K3P0 | Sdr9c7 | Short-chain dehydrogenase/reductase family 9C member 7 | -0.770282968 | 1.445872227 |
| Q8K3W0 | Babam2 | BRISC and BRCA1-A complex member 2 | -0.236654727 | 0.970631917 |
| Q8K3X4 | Irf2bpl | Probable E3 ubiquitin-protein ligase IRF2BPL | 0.657615948 | 0.875715017 |
| Q8K406 | Lgi3 | Leucine-rich repeat LGI family member 3 | -0.902065404 | 0.649775187 |
| Q8K411 | Pitrm1 | Presequence protease, mitochondrial | -0.359428183 | 0.424261252 |
| Q8K4F5 | Abhd11 | Protein ABHD11 | -0.258556016 | 0.208482901 |
| Q8K4M5 | Commd1 | COMM domain-containing protein 1 | -0.932910283 | -0.06341966 |
| Q8K4R4 | Pitpnc1 | Cytoplasmic phosphatidylinositol transfer protein 1 | -0.013973999 | 0.518194675 |
| Q8K4Z3 | Naxe | NAD(P)H-hydrate epimerase | -0.223296293 | 0.389488856 |

| | | | | |
|---|---|---|---|---|
| Q8K4Z5 | Sf3a1 | Splicing factor 3A subunit 1 | -0.033933957 | -0.244813919 |
| Q8K596 | Slc8a2 | Sodium/calcium exchanger 2 | 0.771301301 | 1.341629028 |
| Q8N7N5 | Dcaf8 | DDB1- and CUL4-associated factor 8 | -0.411174806 | 0.234188239 |
| Q8QZR5 | Gpt | Alanine aminotransferase 1 | -0.06228714 | 0.063066483 |
| Q8QZS1 | Hibch | 3-hydroxyisobutyryl-CoA hydrolase, mitochondrial | -0.262197336 | 0.287148794 |
| Q8QZT1 | Acat1 | Acetyl-CoA acetyltransferase, mitochondrial | -0.003321012 | -0.092624029 |
| Q8QZT2 | Ccsap | Centriole, cilia and spindle-associated protein | 0.502982871 | -0.013242881 |
| Q8QZV4 | Stk32c | Serine/threonine-protein kinase 32C | 0.097459857 | 0.032892227 |
| Q8QZY1 | Eif3l | Eukaryotic translation initiation factor 3 subunit L | -0.044376882 | 0.177191257 |
| Q8QZY9 | Sf3b4 | Splicing factor 3B subunit 4 | 0.655044715 | 1.229158719 |
| Q8R010 | Aimp2 | Aminoacyl tRNA synthase complex-interacting multifunctional protein 2 | 1.686579593 | -0.051557779 |
| Q8R016 | Blmh | Bleomycin hydrolase | 0.006238015 | -0.130881945 |
| Q8R050 | Gspt1 | Eukaryotic peptide chain release factor GTP-binding subunit ERF3A | -0.065550709 | 0.161916256 |
| Q8R059 | Gale | UDP-glucose 4-epimerase | -0.013177077 | 0.137655735 |
| Q8R071 | Itpka | Inositol-trisphosphate 3-kinase A | -0.156593672 | -0.25872612 |
| Q8R086 | Suox | Sulfite oxidase, mitochondrial | 0.069276301 | 0.12142849 |
| Q8R0A0 | Gtf2f2 | General transcription factor IIF subunit 2 | -1.685119851 | -0.914644082 |
| Q8R0F6 | Ilkap | Integrin-linked kinase-associated serine/threonine phosphatase 2C | -0.085875448 | 0.13191398 |
| Q8R0F8 | Fahd1 | Acylpyruvase FAHD1, mitochondrial | -0.30597194 | -0.236639659 |
| Q8R0H9 | Gga1 | ADP-ribosylation factor-binding protein GGA1 | -0.184040546 | 0.427615643 |
| Q8R0Y6 | Aldh1l1 | Cytosolic 10-formyltetrahydrofolate dehydrogenase | -0.187041283 | -0.125972748 |
| Q8R104 | Sirt3 | NAD-dependent protein deacetylase sirtuin-3 | 0.434025987 | -0.452869892 |
| Q8R123 | Flad1 | FAD synthase | -0.131472588 | 0.148158391 |
| Q8R124 | Klhl36 | Kelch-like protein 36 | 0.310830243 | 0.911561966 |
| Q8R127 | Sccpdh | Saccharopine dehydrogenase-like oxidoreductase | 0.164060497 | 1.114615758 |
| Q8R151 | Znfx1 | NFX1-type zinc finger-containing protein 1 | -0.986621698 | 0.342249076 |
| Q8R164 | Bphl | Valacyclovir hydrolase | -0.174298477 | 0.510542552 |
| Q8R180 | Ero1a | ERO1-like protein alpha | 0.424495029 | -0.004249414 |
| Q8R191 | Syngr3 | Synaptogyrin-3 | 0.495824909 | 0.489670595 |
| Q8R1B0 | Stac2 | SH3 and cysteine-rich domain-containing protein 2 | -0.148241297 | 0.682179769 |
| Q8R1B4 | Eif3c | Eukaryotic translation initiation factor 3 subunit C | -0.170407645 | -0.090592066 |
| Q8R1F1 | Niban2 | Protein Niban 2 | 0.41926225 | -0.441527843 |
| Q8R1F6 | Hid1 | Protein HID1 | 0.012881438 | 0.739973863 |
| Q8R1G2 | Cmbl | Carboxymethylenebutenolidase homolog | -0.38439846 | 0.322692235 |
| Q8R1N4 | Nudcd3 | NudC domain-containing protein 3 | 0.124280357 | 0.348981857 |
| Q8R1Q8 | Dync1li1 | Cytoplasmic dynein 1 light intermediate chain 1 | 0.039755185 | 0.115817865 |
| Q8R1Q9 | Rbks | Ribokinase | -1.069792112 | 0.684732755 |
| Q8R1R3 | Stard7 | StAR-related lipid transfer protein 7, mitochondrial | -0.3581508 | 0.520936966 |
| Q8R1T1 | Chmp7 | Charged multivesicular body protein 7 | 0.002932326 | 0.734620094 |
| Q8R1X6 | Spart | Spartin | 0.058260155 | 1.057812691 |
| Q8R238 | Sdsl | Serine dehydratase-like | 0.080168152 | 2.272026539 |
| Q8R242 | Ctbs | Di-N-acetylchitobiase | 1.605978028 | 2.477790435 |
| Q8R2H9 | Phospho1 | Phosphoethanolamine/phosphocholine phosphatase | 0.007987404 | 0.348694801 |
| Q8R2K3 | Ssbp1 | Single-stranded DNA-binding protein, mitochondrial | 0.541190561 | 0.875566959 |
| Q8R2Q0 | Trim29 | Tripartite motif-containing protein 29 | -0.596175575 | -1.224279404 |

| | | | | |
|---|---|---|---|---|
| Q8R2R3 | Aagab | Alpha- and gamma-adaptin-binding protein p34 | -0.07053016 | 0.40758419 |
| Q8R2R9 | Ap3m2 | AP-3 complex subunit mu-2 | -0.105444908 | 0.050539017 |
| Q8R2U0 | Seh1l | Nucleoporin SEH1 | 0.024672381 | -0.893491586 |
| Q8R2U6 | Nudt4 | Diphosphoinositol polyphosphate phosphohydrolase 2 | -0.119299539 | 0.294368744 |
| Q8R2Y2 | Mcam | Cell surface glycoprotein MUC18 | -0.786964766 | 1.325664043 |
| Q8R307 | Vps18 | Vacuolar protein sorting-associated protein 18 homolog | 0.589009031 | 0.272706191 |
| Q8R317 | Ubqln1 | Ubiquilin-1 | -0.16323398 | 0.564133326 |
| Q8R326 | Pspc1 | Paraspeckle component 1 | 0.526135953 | 0.167268912 |
| Q8R332 | Nup58 | Nucleoporin p58/p45 | 1.945273336 | 3.000532468 |
| Q8R349 | Cdc16 | Cell division cycle protein 16 homolog | -0.597811476 | -0.101898829 |
| Q8R366 | Igsf8 | Immunoglobulin superfamily member 8 | 0.268263594 | 0.353216966 |
| Q8R3C0 | Mcmbp | Mini-chromosome maintenance complex-binding protein | -0.328615824 | 0.947628975 |
| Q8R3D1 | Tbc1d13 | TBC1 domain family member 13 | 0.0699687 | 0.066681385 |
| Q8R3E3 | Wipi1 | WD repeat domain phosphoinositide-interacting protein 1 | -0.016462421 | -0.406279405 |
| Q8R3F5 | Mcat | Malonyl-CoA-acyl carrier protein transacylase, mitochondrial | 0.762881978 | 0.32626311 |
| Q8R3H9 | Ttc4 | Tetratricopeptide repeat protein 4 | 0.20430212 | 0.138894081 |
| Q8R3P0 | Aspa | Aspartoacylase | 0.029443963 | 0.597601255 |
| Q8R3R8 | Gabarapl1 | Gamma-aminobutyric acid receptor-associated protein-like 1 | 0.239139811 | -0.162152926 |
| Q8R3V5 | Sh3glb2 | Endophilin-B2 | 1.615563679 | -0.437187513 |
| Q8R3W2 | 0610009B22Rik | RIKEN cDNA 0610009B22 gene | -0.41176974 | 0.682080746 |
| Q8R429 | Atp2a1 | Sarcoplasmic/endoplasmic reticulum calcium ATPase 1 | 0.64828585 | 2.47030735 |
| Q8R464 | Cadm4 | Cell adhesion molecule 4 | -0.388210042 | -0.1472435 |
| Q8R480 | Nup85 | Nuclear pore complex protein Nup85 | -0.121883202 | -0.428246816 |
| Q8R4N0 | Clybl | Citramalyl-CoA lyase, mitochondrial | -0.15930481 | 0.488135179 |
| Q8R4S0 | Ppp1r14c | Protein phosphatase 1 regulatory subunit 14C | 0.091178068 | -0.616526127 |
| Q8R4U7 | Luzp1 | Leucine zipper protein 1 | -0.645433521 | 0.448566119 |
| Q8R4X3 | Rbm12 | RNA-binding protein 12 | 0.199666087 | -0.1069959 |
| Q8R550 | Sh3kbp1 | SH3 domain-containing kinase-binding protein 1 | 0.035692533 | -0.192720095 |
| Q8R554 | Otud7a | OTU domain-containing protein 7A | 0.464116891 | 0.448504289 |
| Q8R570 | Snap47 | Synaptosomal-associated protein 47 | 0.149373086 | 0.183420022 |
| Q8R574 | Prpsap2 | Phosphoribosyl pyrophosphate synthase-associated protein 2 | 0.108938662 | -0.351229986 |
| Q8R5A3 | Apbb1ip | Amyloid beta A4 precursor protein-binding family B member 1-interacting protein | 0.824045086 | 0.77036794 |
| Q8R5A6 | Tbc1d22a | TBC1 domain family member 22A | -1.471911605 | 0.27564923 |
| Q8R5C5 | Actr1b | Beta-centractin | 0.637362989 | 0.12872982 |
| Q8R5H1 | Usp15 | Ubiquitin carboxyl-terminal hydrolase 15 | -0.074866962 | 0.297751427 |
| Q8R5H6 | Wasf1 | Wiskott-Aldrich syndrome protein family member 1 | 0.020515124 | -0.36484321 |
| Q8R5I6 | Gstm4 | Glutathione S-transferase Mu 4 | 0.398007902 | 0.680769285 |
| Q8R5J9 | Arl6ip5 | PRA1 family protein 3 | 0.09853878 | -0.156041463 |
| Q8VBT9 | Aspscr1 | Tether containing UBX domain for GLUT4 | -0.150788562 | 0.068645477 |
| Q8VBV7 | Cops8 | COP9 signalosome complex subunit 8 | -1.075989278 | -1.106131554 |
| Q8VBY2 | Camkk1 | Calcium/calmodulin-dependent protein kinase kinase 1 | -0.106699467 | 0.243233045 |
| Q8VC88 | Gca | Grancalcin | 0.075254122 | 0.410122077 |
| Q8VCA8 | Scrn2 | Secernin-2 | -0.663048426 | -0.456105391 |
| Q8VCC9 | Spon1 | Spondin-1 | -0.962943236 | -0.516453584 |
| Q8VCE6 | Nt5m | 5'(3')-deoxyribonucleotidase, mitochondrial | 0.08516407 | 0.380611579 |

| | | | | |
|---|---|---|---|---|
| Q8VCG1 | Dut | Deoxyuridine 5'-triphosphate nucleotidohydrolase | 0.211456712 | 0.43898042 |
| Q8VCI5 | Pex19 | Peroxisomal biogenesis factor 19 | 0.355650171 | -0.078274568 |
| Q8VCN5 | Cth | Cystathionine gamma-lyase | 0.3220143 | -0.098072052 |
| Q8VCR4 | Trmt112 | Multifunctional methyltransferase subunit TRM112-like protein | -0.32671601 | 0.646141688 |
| Q8VCT3 | Rnpep | Aminopeptidase B | -0.066480319 | 0.194350402 |
| Q8VCW8 | Acsf2 | Medium-chain acyl-CoA ligase ACSF2, mitochondrial | -0.255444717 | 0.273453712 |
| Q8VD33 | Sgtb | Small glutamine-rich tetratricopeptide repeat-containing protein beta | 0.022876485 | 0.155180454 |
| Q8VD37 | Sgip1 | SH3-containing GRB2-like protein 3-interacting protein 1 | -0.061391608 | 0.036458333 |
| Q8VD62 | Bles03 | UPF0696 protein C11orf68 homolog | 0.076718807 | 0.212815762 |
| Q8VD63 | Tspyl4 | Testis-specific Y-encoded-like protein 4 | -1.43747584 | 1.593574206 |
| Q8VD65 | Pik3r4 | Phosphoinositide 3-kinase regulatory subunit 4 | 0.497967307 | 0.132773161 |
| Q8VD75 | Hip1 | Huntingtin-interacting protein 1 | -0.248626073 | -0.867031654 |
| Q8VDC0 | Lars2 | Probable leucine--tRNA ligase, mitochondrial | 0.157423306 | 1.229387124 |
| Q8VDD5 | Myh9 | Myosin-9 | -0.134419982 | 0.251080354 |
| Q8VDD8 | Washc1 | WASH complex subunit 1 | -0.271800518 | 0.976522128 |
| Q8VDG5 | Ppcs | Phosphopantothenate--cysteine ligase | 0.093712552 | -0.228953362 |
| Q8VDH1 | Fbxo21 | F-box only protein 21 | 0.022930908 | 0.262555917 |
| Q8VDI7 | Ubac1 | Ubiquitin-associated domain-containing protein 1 | -0.387068876 | 0.500991503 |
| Q8VDJ3 | Hdlbp | Vigilin | -0.006914139 | 0.07251056 |
| Q8VDK1 | Nit1 | Deaminated glutathione amidase | -0.327186044 | 0.521464348 |
| Q8VDM4 | Psmd2 | 26S proteasome non-ATPase regulatory subunit 2 | 0.07387689 | 0.036984762 |
| Q8VDM6 | Hnrnpul1 | Heterogeneous nuclear ribonucleoprotein U-like protein 1 | -0.78108689 | -0.635503292 |
| Q8VDN2 | Atp1a1 | Sodium/potassium-transporting ATPase subunit alpha-1 | 0.193309625 | 0.376791636 |
| Q8VDP3 | Mical1 | [F-actin]-monooxygenase MICAL1 | 0.040965652 | 1.406604449 |
| Q8VDP4 | Ccar2 | Cell cycle and apoptosis regulator protein 2 | -0.155519231 | -0.067056815 |
| Q8VDQ1 | Ptgr2 | Prostaglandin reductase 2 | -0.096980063 | 0.380737305 |
| Q8VDQ8 | Sirt2 | NAD-dependent protein deacetylase sirtuin-2 | 0.125472705 | 0.659579277 |
| Q8VDS4 | Rprd1a | Regulation of nuclear pre-mRNA domain-containing protein 1A | -0.710939217 | 1.921153188 |
| Q8VDU5 | Snrk | SNF-related serine/threonine-protein kinase | -0.264336427 | 0.007811069 |
| Q8VDW0 | Ddx39a | ATP-dependent RNA helicase DDX39A | -0.981262461 | 0.627005974 |
| Q8VDZ4 | Zdhhc5 | Palmitoyltransferase ZDHHC5 | 0.378660901 | 0.222889741 |
| Q8VE11 | Mtmr6 | Myotubularin-related protein 6 | -0.300703589 | 0.761203289 |
| Q8VE38 | Oxnad1 | Oxidoreductase NAD-binding domain-containing protein 1 | 0.168594551 | 2.028571606 |
| Q8VE47 | Uba5 | Ubiquitin-like modifier-activating enzyme 5 | -0.189935207 | -1.359376431 |
| Q8VE62 | Paip1 | Polyadenylate-binding protein-interacting protein 1 | -0.022806327 | -0.210825761 |
| Q8VE70 | Pdcd10 | Programmed cell death protein 10 | 0.465170288 | 0.189881643 |
| Q8VE80 | Thoc3 | THO complex subunit 3 | -0.453183015 | 0.84502546 |
| Q8VE88 | Fam114a2 | Protein FAM114A2 | -0.259243647 | 0.490633806 |
| Q8VE95 | | UPF0598 protein C8orf82 homolog | 0.04192756 | 0.206084092 |
| Q8VE99 | Ccdc115 | Coiled-coil domain-containing protein 115 | 2.024008497 | 2.245189349 |
| Q8VEA4 | Chchd4 | Mitochondrial intermembrane space import and assembly protein 40 | 0.05266463 | 1.693985144 |
| Q8VEB4 | Pla2g15 | Phospholipase A2 group XV | -0.217740726 | -0.07675155 |
| Q8VEB6 | Elac1 | Zinc phosphodiesterase ELAC protein 1 | 0.01669302 | -0.072455565 |
| Q8VED2 | Bloc1s4 | Biogenesis of lysosome-related organelles complex 1 subunit 4 | -0.402323214 | -0.251909415 |

| | | | | |
|---|---|---|---|---|
| Q8VED5 | Krt79 | Keratin, type II cytoskeletal 79 | 0.263965797 | -0.456306458 |
| Q8VED9 | Lgalsl | Galectin-related protein | -0.160593383 | 0.59998401 |
| Q8VEE1 | Lmcd1 | LIM and cysteine-rich domains protein 1 | -0.007072894 | 0.335590045 |
| Q8VEH3 | Arl8a | ADP-ribosylation factor-like protein 8A | 0.298111216 | 0.91179959 |
| Q8VEH5 | Epm2aip1 | EPM2A-interacting protein 1 | -0.0451677 | 0.344143391 |
| Q8VEH6 | Cbwd1 | COBW domain-containing protein 1 | -1.694678656 | -1.414625486 |
| Q8VEJ9 | Vps4a | Vacuolar protein sorting-associated protein 4A | 0.050650247 | 0.241815249 |
| Q8VEK3 | Hnrnpu | Heterogeneous nuclear ribonucleoprotein U | -0.235235786 | 0.140583038 |
| Q8VH37 | Hdac8 | Histone deacetylase 8 | -0.074087461 | 0.387062709 |
| Q8VH51 | Rbm39 | RNA-binding protein 39 | -0.715822442 | 1.457015673 |
| Q8VHI6 | Wasf3 | Wiskott-Aldrich syndrome protein family member 3 | 0.2848423 | 0.746824582 |
| Q8VHJ5 | Mark1 | Serine/threonine-protein kinase MARK1 | 0.034186268 | 0.4150184 |
| Q8VHL1 | Setd7 | Histone-lysine N-methyltransferase SETD7 | -0.192851448 | 0.436105887 |
| Q8VHM5 | Hnrnpr | Heterogeneous nuclear ribonucleoprotein R | -0.446521568 | -0.439696153 |
| Q8VHP7 | Serpinb1b | Leukocyte elastase inhibitor B | -0.550955931 | 0.246768951 |
| Q8VHQ9 | Acot11 | Acyl-coenzyme A thioesterase 11 | -0.255144024 | 0.210988681 |
| Q8VHX6 | Flnc | Filamin-C | 0.053393777 | 0.917430083 |
| Q8VHY0 | Cspg4 | Chondroitin sulfate proteoglycan 4 | -0.567929618 | 0.870972157 |
| Q8VI63 | Mob2 | MOB kinase activator 2 | 0.390504328 | 1.311010838 |
| Q8VI75 | Ipo4 | Importin-4 | 0.202965069 | 0.676428318 |
| Q8VIJ6 | Sfpq | Splicing factor, proline- and glutamine-rich | 0.222809156 | -0.066669782 |
| Q8VIM9 | Irgq | Immunity-related GTPase family Q protein | -0.022054704 | 0.583837668 |
| Q8WTY4 | Ciapin1 | Anamorsin | -0.243173091 | 0.088416735 |
| Q8WUR0 | | Protein C19orf12 homolog | -0.005206935 | 1.120481014 |
| Q91UZ5 | Impa2 | Inositol monophosphatase 2 | 0.276477655 | -3.156291087 |
| Q91V09 | Wdr13 | WD repeat-containing protein 13 | -0.166521422 | 0.268857002 |
| Q91V35 | Ptpra | Receptor-type tyrosine-protein phosphatase alpha | -0.093621127 | -0.046324571 |
| Q91V36 | Nrbp2 | Nuclear receptor-binding protein 2 | 0.316383012 | 0.315980752 |
| Q91V41 | Rab14 | Ras-related protein Rab-14 | -0.033042717 | 1.182196935 |
| Q91V57 | Chn1 | N-chimaerin | 0.475673612 | 0.919750849 |
| Q91V64 | Isoc1 | Isochorismatase domain-containing protein 1 | 0.066522916 | 0.5937953 |
| Q91V76 | | Ester hydrolase C11orf54 homolog | -0.162936433 | 0.35240221 |
| Q91VA7 | Idh3b | Isocitrate dehydrogenase [NAD] subunit, mitochondrial | -0.076229858 | 0.272317886 |
| Q91VB8 | Hba-a1 | Alpha globin 1 | -0.260482661 | 0.467582703 |
| Q91VC7 | Ppp1r14a | Protein phosphatase 1 regulatory subunit 14A | 0.109999371 | 0.800837517 |
| Q91VD9 | Ndufs1 | NADH-ubiquinone oxidoreductase 75 kDa subunit, mitochondrial | 0.073567041 | 0.436040719 |
| Q91VF2 | Hnmt | Histamine N-methyltransferase | -0.094189771 | 0.329570134 |
| Q91VH2 | Snx9 | Sorting nexin-9 | -0.266657416 | 0.099229018 |
| Q91VH6 | Memo1 | Protein MEMO1 | 0.452783902 | 0.804620902 |
| Q91VI7 | Rnh1 | Ribonuclease inhibitor | -0.064398638 | 0.567121506 |
| Q91VJ4 | Stk38 | Serine/threonine-protein kinase 38 | -0.159309069 | -0.157721202 |
| Q91VJ5 | Pqbp1 | Polyglutamine-binding protein 1 | 0.040424093 | -0.167495728 |
| Q91VK1 | Bzw2 | Basic leucine zipper and W2 domain-containing protein 2 | 1.919921271 | 0.523825963 |
| Q91VM3 | Wdr45 | WD repeat domain phosphoinositide-interacting protein 4 | -0.180427742 | 0.519887447 |
| Q91VM9 | Ppa2 | Inorganic pyrophosphatase 2, mitochondrial | -0.10844628 | -0.724035422 |

| | | | | |
|---|---|---|---|---|
| Q91VR5 | Ddx1 | ATP-dependent RNA helicase DDX1 | 0.168309402 | 0.062530677 |
| Q91VR7 | Map1lc3a | Microtubule-associated proteins 1A/1B light chain 3A | -0.381587537 | 0.637643059 |
| Q91VT4 | Cbr4 | Carbonyl reductase family member 4 | -0.531456343 | 0.281889915 |
| Q91VU6 | Dcaf11 | DDB1- and CUL4-associated factor 11 | 0.910110346 | -0.215268532 |
| Q91VW3 | Sh3bgrl3 | SH3 domain-binding glutamic acid-rich-like protein 3 | -0.029626624 | 0.406696796 |
| Q91VZ6 | Smap1 | Stromal membrane-associated protein 1 | -0.467103163 | -0.517109712 |
| Q91W50 | Csde1 | Cold shock domain-containing protein E1 | -0.156543668 | -0.017767747 |
| Q91W61 | Fbxl15 | F-box/LRR-repeat protein 15 | -0.120945072 | 0.244757493 |
| Q91W67 | Ubl7 | Ubiquitin-like protein 7 | -1.075248369 | 0.873835882 |
| Q91W69 | Epn3 | Epsin-3 | 0.394874748 | -0.263289054 |
| Q91W86 | Vps11 | Vacuolar protein sorting-associated protein 11 homolog | 0.104053752 | -0.862790108 |
| Q91W96 | Anapc4 | Anaphase-promoting complex subunit 4 | -1.266385841 | -0.532330831 |
| Q91WC0 | Setd3 | Actin-histidine N-methyltransferase | -0.165757402 | 0.485581875 |
| Q91WD5 | Ndufs2 | NADH dehydrogenase [ubiquinone] iron-sulfur protein 2, mitochondrial | 0.545457649 | 2.030831973 |
| Q91WD9 | Scgn | Secretagogin | -0.697123051 | 0.725361824 |
| Q91WE2 | Psme3ip1 | PSME3-interacting protein | 0.258147653 | -0.734381199 |
| Q91WG2 | Rabep2 | Rab GTPase-binding effector protein 2 | -0.197931163 | 1.065500736 |
| Q91WG4 | Elp2 | Elongator complex protein 2 | 0.287255859 | 0.137126287 |
| Q91WG7 | Dgkg | Diacylglycerol kinase gamma | -0.355174383 | 1.388930837 |
| Q91WK1 | Spryd4 | SPRY domain-containing protein 4 | -0.068456936 | -0.383808931 |
| Q91WK2 | Eif3h | Eukaryotic translation initiation factor 3 subunit H | 0.455346743 | 0.376506805 |
| Q91WK5 | Gcsh | Glycine cleavage system H protein, mitochondrial | -0.584472656 | -0.893083731 |
| Q91WL8 | Wwox | WW domain-containing oxidoreductase | -1.1472085 | 0.004454613 |
| Q91WM1 | Strbp | Spermatid perinuclear RNA-binding protein | -0.812377135 | 2.152539372 |
| Q91WM2 | Hdhd5 | Haloacid dehalogenase-like hydrolase domain-containing 5 | -0.074610233 | -0.047791004 |
| Q91WQ3 | Yars1 | Tyrosine--tRNA ligase, cytoplasmic | -0.032011541 | 0.104057312 |
| Q91WS0 | Cisd1 | CDGSH iron-sulfur domain-containing protein 1 | 2.241358081 | 2.322579304 |
| Q91WT9 | Cbs | Cystathionine beta-synthase | -0.35200421 | 0.360882282 |
| Q91WU5 | As3mt | Arsenite methyltransferase | -0.26240956 | 0.005432447 |
| Q91WV0 | Dr1 | Protein Dr1 | -0.708668041 | 0.798562368 |
| Q91X51 | Gorasp1 | Golgi reassembly-stacking protein 1 | 0.342002773 | -0.541252772 |
| Q91X52 | Dcxr | L-xylulose reductase | -0.219271088 | -0.22631073 |
| Q91X72 | Hpx | Hemopexin | 0.920481269 | 0.460367521 |
| Q91X83 | Mat1a | S-adenosylmethionine synthase isoform type-1 | -0.554376825 | 0.701128483 |
| Q91X91 | Qprt | Nicotinate-nucleotide pyrophosphorylase [carboxylating] | -0.58975331 | -1.004924297 |
| Q91X96 | Rabif | Guanine nucleotide exchange factor MSS4 | 0.32444919 | 1.313590844 |
| Q91XD6 | Vps36 | Vacuolar protein-sorting-associated protein 36 | 0.141054726 | -0.022558212 |
| Q91XE4 | Acy3 | N-acyl-aromatic-L-amino acid amidohydrolase (carboxylate-forming) | -0.103434531 | 0.164122423 |
| Q91XF0 | Pnpo | Pyridoxine-5'-phosphate oxidase | 0.323683071 | 0.429743131 |
| Q91XH5 | Spr | Sepiapterin reductase | -0.271719869 | 0.457389037 |
| Q91XL1 | Lrg1 | Leucine-rich HEV glycoprotein | -0.578473345 | -0.223804633 |
| Q91XU0 | Wrnip1 | ATPase WRNIP1 | -0.195628802 | -0.030597051 |
| Q91XU3 | Pip4k2c | Phosphatidylinositol 5-phosphate 4-kinase type-2 gamma | 0.051455021 | -0.173671722 |
| Q91XV3 | Basp1 | Brain acid soluble protein 1 | 0.551816082 | 0.501047611 |
| Q91XW9 | Pcdhgc5 | Pcdhgc5 protein | -0.128622691 | 1.161222617 |

| | | | | |
|---|---|---|---|---|
| Q91YD9 | Wasl | Neural Wiskott-Aldrich syndrome protein | 0.482044983 | 0.22233359 |
| Q91YE3 | Egln1 | Egl nine homolog 1 | -0.063383547 | 0.40273269 |
| Q91YI0 | Asl | Argininosuccinate lyase | -0.053867277 | 0.08283631 |
| Q91YJ2 | Snx4 | Sorting nexin-4 | 0.046404012 | 0.337801139 |
| Q91YJ5 | Mtif2 | Translation initiation factor IF-2, mitochondrial | 0.370283794 | -0.042106311 |
| Q91YL2 | Rnf126 | E3 ubiquitin-protein ligase RNF126 | 0.105793603 | -0.295631886 |
| Q91YL3 | Uckl1 | Uridine-cytidine kinase-like 1 | -0.600328318 | -0.143395742 |
| Q91YM2 | Arhgap35 | Rho GTPase-activating protein 35 | 0.077625974 | 0.561376095 |
| Q91YM4 | Tbrg4 | FAST kinase domain-containing protein 4 | -0.148737812 | 2.606926123 |
| Q91YP0 | L2hgdh | L-2-hydroxyglutarate dehydrogenase, mitochondrial | 0.067795531 | 0.207067966 |
| Q91YP2 | Nln | Neurolysin, mitochondrial | -0.058258979 | -0.007057985 |
| Q91YQ3 | Csdc2 | Cold shock domain-containing protein C2 | 0.136333275 | -1.092834314 |
| Q91YQ5 | Rpn1 | Dolichyl-diphosphooligosaccharide--protein glycosyltransferase subunit 1 | 0.641856035 | 1.615575155 |
| Q91YR1 | Twf1 | Twinfilin-1 | 0.037419383 | 0.256459077 |
| Q91YS8 | Camk1 | Calcium/calmodulin-dependent protein kinase type 1 | -0.246380615 | -0.218213399 |
| Q91YT7 | Ythdf2 | YTH domain-containing family protein 2 | 1.767501211 | 0.534998735 |
| Q91YW3 | Dnajc3 | DnaJ homolog subfamily C member 3 | 0.290599759 | 0.588115692 |
| Q91Z25 | Arpc1b | Actin-related protein 2/3 complex subunit | 0.797653834 | 0.945922375 |
| Q91Z31 | Ptbp2 | Polypyrimidine tract-binding protein 2 | -0.238659922 | 0.106164138 |
| Q91Z40 | Gbp7 | Gbp6 protein | -1.358831914 | 0.559219758 |
| Q91Z53 | Grhpr | Glyoxylate reductase/hydroxypyruvate reductase | -0.279452705 | 0.250795046 |
| Q91Z61 | Diras1 | GTP-binding protein Di-Ras1 | -0.040148958 | -0.069457849 |
| Q91Z67 | Srgap2 | SLIT-ROBO Rho GTPase-activating protein 2 | -0.015944513 | -0.139618874 |
| Q91ZA3 | Pcca | Propionyl-CoA carboxylase alpha chain, mitochondrial | -0.048662313 | 0.344310602 |
| Q91ZJ5 | Ugp2 | UTP--glucose-1-phosphate uridylyltransferase | 0.028428396 | 0.129938126 |
| Q91ZM2 | Sh2b1 | SH2B adapter protein 1 | -0.090909608 | 0.477970441 |
| Q91ZP9 | Necab2 | N-terminal EF-hand calcium-binding protein 2 | 0.195916971 | 0.624421438 |
| Q91ZR1 | Rab4b | Ras-related protein Rab-4B | 0.273435084 | 0.812461217 |
| Q91ZZ3 | Sncb | Beta-synuclein | -0.202662977 | 0.835526784 |
| Q920A5 | Scpep1 | Retinoid-inducible serine carboxypeptidase | 0.169043668 | -0.31194973 |
| Q920E5 | Fdps | Farnesyl pyrophosphate synthase | -0.409920406 | 0.022694747 |
| Q920I9 | Wdr7 | WD repeat-containing protein 7 | 0.00076952 | -0.252443155 |
| Q920N7 | Syt12 | Synaptotagmin-12 | -0.189964994 | 0.270138741 |
| Q920P5 | Ak5 | Adenylate kinase isoenzyme 5 | 0.218412654 | -0.201496919 |
| Q920Q6 | Msi2 | RNA-binding protein Musashi homolog 2 | 0.308050744 | -0.280154546 |
| Q920Q8 | Ivns1abp | Influenza virus NS1A-binding protein homolog | -0.03919398 | 0.322771549 |
| Q920R0 | Als2 | Alsin | -0.570090485 | -1.025410493 |
| Q921C5 | Bicd2 | Protein bicaudal D homolog 2 | -0.375551383 | 0.915091912 |
| Q921F2 | Tardbp | TAR DNA-binding protein 43 | -0.044797166 | 0.091836452 |
| Q921F4 | Hnrnpll | Heterogeneous nuclear ribonucleoprotein L-like | -0.49257609 | 0.451447964 |
| Q921H8 | Acaa1a | 3-ketoacyl-CoA thiolase A, peroxisomal | 1.260103353 | -1.179642081 |
| Q921H9 | Coa7 | Cytochrome c oxidase assembly factor 7 | 0.460624282 | 1.674479961 |
| Q921I1 | Tf | Serotransferrin | -0.199655279 | 0.203074455 |
| Q921J2 | Rheb | GTP-binding protein Rheb | 0.003691864 | 0.339219093 |
| Q921L5 | Cog2 | Conserved oligomeric Golgi complex subunit 2 | -1.931968906 | -0.554466407 |
| Q921L6 | Cttn | Cttn protein | -0.202791977 | 0.392810742 |

| | | | | |
|---|---|---|---|---|
| Q921M3 | Sf3b3 | Splicing factor 3B subunit 3 | -0.088091246 | -0.237162272 |
| Q921M7 | Cyrib | CYFIP-related Rac1 interactor B | 0.293463898 | 0.703715324 |
| Q922B1 | Macrod1 | ADP-ribose glycohydrolase MACROD1 | -0.239308898 | 0.423594634 |
| Q922B2 | Dars1 | Aspartate--tRNA ligase, cytoplasmic | 0.134646257 | 0.203713576 |
| Q922D8 | Mthfd1 | C-1-tetrahydrofolate synthase, cytoplasmic | -0.180912145 | 0.05993557 |
| Q922F4 | Tubb6 | Tubulin beta-6 chain | -0.842893497 | 1.065086603 |
| Q922H1 | Prmt3 | Protein arginine N-methyltransferase 3 | 0.820294603 | -0.052745024 |
| Q922H2 | Pdk3 | [Pyruvate dehydrogenase (acetyl-transferring)] kinase isozyme 3, mitochondrial | -0.083305995 | 1.731898308 |
| Q922H4 | Gmppa | Mannose-1-phosphate guanyltransferase alpha | -0.012536971 | 0.124189218 |
| Q922H9 | Znf330 | Zinc finger protein 330 | -0.867245611 | 1.538286765 |
| Q922Q4 | Pycr2 | Pyrroline-5-carboxylate reductase 2 | -1.073475488 | 1.152445157 |
| Q922Q8 | Lrrc59 | Leucine-rich repeat-containing protein 59 | 2.806235298 | 0.81235981 |
| Q922R1 | | UPF0183 protein C16orf70 homolog | 0.774569003 | 0.458323161 |
| Q922U2 | Krt5 | Keratin, type II cytoskeletal 5 | 0.022529666 | -0.374526024 |
| Q922X9 | Prmt7 | Protein arginine N-methyltransferase 7 | -0.445237033 | 0.179262479 |
| Q923D2 | Blvrb | Flavin reductase (NADPH) | -0.349578857 | 0.354917685 |
| Q923M0 | Ppp1r16a | Protein phosphatase 1 regulatory subunit 16A | 0.002696451 | 0.062489669 |
| Q923T9 | Camk2g | Calcium/calmodulin-dependent protein kinase type II subunit gamma | 0.765346527 | -0.851905028 |
| Q924B0 | Impa1 | Inositol-1-monophosphatase | -0.160724322 | 0.642285665 |
| Q924C1 | Xpo5 | Exportin-5 | 0.083662287 | 0.055758317 |
| Q924M7 | Mpi | Mannose-6-phosphate isomerase | 0.156943576 | -0.129649639 |
| Q924S8 | Spred1 | Sprouty-related, EVH1 domain-containing protein 1 | 0.059069538 | 1.0322752 |
| Q924Y0 | Bbox1 | Gamma-butyrobetaine dioxygenase | 0.728749498 | 0.408624808 |
| Q924Z6 | Xpo6 | Exportin-6 | -0.295696735 | 0.618890127 |
| Q925E7 | Ppp2r2d | Serine/threonine-protein phosphatase 2A 55 kDa regulatory subunit B delta isoform | 0.243736013 | 0.31114165 |
| Q99J08 | Sec14l2 | SEC14-like protein 2 | -0.181751537 | 0.19464763 |
| Q99J09 | Wdr77 | Methylosome protein 50 | -0.145357482 | 1.022399267 |
| Q99J10 | Ctu1 | Cytoplasmic tRNA 2-thiolation protein 1 | -1.604828739 | -0.085523605 |
| Q99J36 | Thumpd1 | THUMP domain-containing protein 1 | 0.156942113 | 0.102045695 |
| Q99J39 | Mlycd | Malonyl-CoA decarboxylase, mitochondrial | -0.184640598 | -0.313372294 |
| Q99J45 | Nrbp1 | Nuclear receptor-binding protein | 0.248416456 | 0.722288291 |
| Q99J77 | Nans | Sialic acid synthase | -0.323771604 | -0.153578758 |
| Q99J83 | Atg5 | Autophagy protein 5 | -0.014353021 | 0.08017985 |
| Q99J95 | Cdk9 | Cyclin-dependent kinase 9 | -0.81996851 | 0.878895283 |
| Q99J99 | Mpst | 3-mercaptopyruvate sulfurtransferase | -0.078215408 | 0.504388014 |
| Q99JB2 | Stoml2 | Stomatin-like protein 2, mitochondrial | 1.358964221 | 2.467374007 |
| Q99JB8 | Pacsin3 | Protein kinase C and casein kinase II substrate protein 3 | 0.368613942 | -0.926425616 |
| Q99JF5 | Mvd | Diphosphomevalonate decarboxylase | -0.654095936 | 0.821608861 |
| Q99JI4 | Psmd6 | 26S proteasome non-ATPase regulatory subunit 6 | -0.08593003 | -0.49314642 |
| Q99JI6 | Rap1b | Ras-related protein Rap-1b | 0.33217624 | 0.72062397 |
| Q99JN2 | Klhl22 | Kelch-like protein 22 | -0.095862961 | -1.233324051 |
| Q99JP6 | Homer3 | Homer protein homolog 3 | -0.325329431 | -0.031007926 |
| Q99JR1 | Sfxn1 | Sideroflexin-1 | -0.652565765 | 0.661594947 |
| Q99JT9 | Adi1 | 1,2-dihydroxy-3-keto-5-methylthiopentene dioxygenase | -0.016499551 | -0.277483304 |
| Q99JX4 | Eif3m | Eukaryotic translation initiation factor 3 subunit M | 0.65942777 | 0.462137063 |

| | | | | |
|---|---|---|---|---|
| Q99JY0 | Hadhb | Trifunctional enzyme subunit beta, mitochondrial | 0.104572392 | 0.066486359 |
| Q99JY8 | Plpp3 | Phospholipid phosphatase 3 | 1.131997077 | -0.45391798 |
| Q99JY9 | Actr3 | Actin-related protein 3 | 0.18844951 | 0.323048433 |
| Q99JZ4 | Sar1a | GTP-binding protein SAR1a | 0.179868031 | 0.607690175 |
| Q99K28 | Arfgap2 | ADP-ribosylation factor GTPase-activating protein 2 | -0.182328701 | -0.379763285 |
| Q99K30 | Eps8l2 | Epidermal growth factor receptor kinase substrate 8-like protein 2 | -0.774282519 | -0.402053833 |
| Q99K46 | Usp11 | Ubiquitin carboxyl-terminal hydrolase 11 | -0.171924178 | 0.785933654 |
| Q99K48 | Nono | Non-POU domain-containing octamer-binding protein | -0.517571195 | -0.345868905 |
| Q99K67 | Aass | Alpha-aminoadipic semialdehyde synthase, mitochondrial | -1.710954905 | 0.020132383 |
| Q99K70 | Rragc | Ras-related GTP-binding protein C | -0.094811853 | 0.255571842 |
| Q99K85 | Psat1 | Phosphoserine aminotransferase | -0.229721705 | 0.240190506 |
| Q99KC8 | Vwa5a | von Willebrand factor A domain-containing protein 5A | -0.230655225 | 0.230234782 |
| Q99KE1 | Me2 | NAD-dependent malic enzyme, mitochondrial | -0.217239443 | -0.867346605 |
| Q99KG3 | Rbm10 | RNA-binding protein 10 | 0.330562814 | 0.095949173 |
| Q99KH8 | Stk24 | Serine/threonine-protein kinase 24 | -0.311646589 | 0.355312665 |
| Q99KI0 | Aco2 | Aconitate hydratase, mitochondrial | -0.304192861 | -0.004205704 |
| Q99KJ8 | Dctn2 | Dynactin subunit 2 | 0.154614576 | 0.172335625 |
| Q99KK7 | Dpp3 | Dipeptidyl peptidase 3 | -0.067798678 | 0.369371891 |
| Q99KK9 | Hars2 | Histidine--tRNA ligase, mitochondrial | 0.233411916 | 0.604642868 |
| Q99KL7 | Rab28 | Ras-related protein Rab-28 | 0.885478306 | 1.547576269 |
| Q99KN2 | Ciao1 | Probable cytosolic iron-sulfur protein assembly protein CIAO1 | 0.143888028 | -0.043831984 |
| Q99KP3 | Cryl1 | Lambda-crystallin homolog | -0.051560752 | 0.372198741 |
| Q99KP6 | Prpf19 | Pre-mRNA-processing factor 19 | 0.386783822 | 0.11731259 |
| Q99KQ4 | Nampt | Nicotinamide phosphoribosyltransferase | -0.060771306 | 0.337413629 |
| Q99KR3 | Lactb2 | Endoribonuclease LACTB2 | 0.18089625 | 0.722374916 |
| Q99KR7 | Ppif | Peptidyl-prolyl cis-trans isomerase F, mitochondrial | -0.26235555 | 0.087813536 |
| Q99KV1 | Dnajb11 | DnaJ homolog subfamily B member 11 | 0.746819305 | 0.676280975 |
| Q99KX1 | Mlf2 | Myeloid leukemia factor 2 | 0.273356501 | 0.327094873 |
| Q99L04 | Dhrs1 | Dehydrogenase/reductase SDR family member 1 | -0.005300109 | 2.441308498 |
| Q99L13 | Hibadh | 3-hydroxyisobutyrate dehydrogenase, mitochondrial | 0.023406982 | 0.38103199 |
| Q99L20 | Gstt3 | Glutathione S-transferase theta-3 | 0.296815427 | -0.00691843 |
| Q99L27 | Gmpr2 | GMP reductase 2 | -0.259720294 | 0.362232367 |
| Q99L45 | Eif2s2 | Eukaryotic translation initiation factor 2 subunit 2 | -0.026885414 | 0.038982709 |
| Q99LB2 | Dhrs4 | Dehydrogenase/reductase SDR family member 4 | 0.135066128 | 0.754166921 |
| Q99LB4 | Capg | Macrophage-capping protein | -0.024670283 | 0.19710207 |
| Q99LB6 | Mat2b | Methionine adenosyltransferase 2 subunit beta | -0.253015645 | 0.249679565 |
| Q99LB7 | Sardh | Sarcosine dehydrogenase, mitochondrial | -0.226671219 | 0.310807705 |
| Q99LC3 | Ndufa10 | NADH dehydrogenase [ubiquinone] 1 alpha subcomplex subunit 10, mitochondrial | 0.360196431 | 0.916084448 |
| Q99LC5 | Etfa | Electron transfer flavoprotein subunit alpha, mitochondrial | -0.177522977 | 0.364916801 |
| Q99LC8 | Eif2b1 | Translation initiation factor eIF-2B subunit alpha | 0.674225712 | 0.16754063 |
| Q99LD8 | Ddah2 | N(G),N(G)-dimethylarginine dimethylaminohydrolase 2 | -0.225774924 | 0.244609515 |
| Q99LD9 | Eif2b2 | Translation initiation factor eIF-2B subunit beta | -0.081607469 | -0.702469985 |
| Q99LE6 | Abcf2 | ATP-binding cassette sub-family F member 2 | -0.346790377 | 0.347804705 |
| Q99LF4 | Rtcb | RNA-splicing ligase RtcB homolog | -0.152346929 | -0.365087509 |
| Q99LG2 | Tnpo2 | Transportin-2 | 0.107543786 | -0.323392868 |

| | | | | |
|---|---|---|---|---|
| Q99LG4 | Ttc5 | Tetratricopeptide repeat protein 5 | 0.060321426 | -0.505596161 |
| Q99LP6 | Grpel1 | GrpE protein homolog 1, mitochondrial | -0.133936373 | 0.844027996 |
| Q99LS3 | Psph | Phosphoserine phosphatase | 0.115680854 | 0.454929034 |
| Q99LU0 | Chmp1b1 | Charged multivesicular body protein 1b-1 | -0.132370408 | -0.01162672 |
| Q99LX0 | Park7 | Parkinson disease protein 7 homolog | -0.138834572 | 0.405182203 |
| Q99M28 | Rnps1 | RNA-binding protein with serine-rich domain 1 | -2.10658172 | 0.929790656 |
| Q99M51 | Nck1 | Cytoplasmic protein NCK1 | -0.02155269 | 0.384947459 |
| Q99M71 | Epdr1 | Mammalian ependymin-related protein 1 | -0.055272357 | -0.00925223 |
| Q99M73 | Krt84 | Keratin, type II cuticular Hb4 | 0.274469662 | -0.131759644 |
| Q99M74 | Krt82 | Keratin, type II cuticular Hb2 | -0.296761831 | 0.027814309 |
| Q99M87 | Dnaja3 | DnaJ homolog subfamily A member 3, mitochondrial | 0.725026004 | 1.263864835 |
| Q99ME2 | Wdr6 | WD repeat-containing protein 6 | -0.044686413 | -0.160844803 |
| Q99MK8 | Grk2 | Beta-adrenergic receptor kinase 1 | 0.206350676 | 0.220325947 |
| Q99MN1 | Kars1 | Lysine--tRNA ligase | 0.053460058 | -0.480575403 |
| Q99MN9 | Pccb | Propionyl-CoA carboxylase beta chain, mitochondrial | -0.435840257 | -0.210323334 |
| Q99MP8 | Brap | BRCA1-associated protein | -0.050488885 | -0.117477973 |
| Q99MR0 | Actl6b | Actin-like protein 6B | 0.008177598 | 0.583906015 |
| Q99MR1 | Gigyf1 | GRB10-interacting GYF protein 1 | 0.420804469 | -0.045562744 |
| Q99MR6 | Srrt | Serrate RNA effector molecule homolog | -0.315454992 | 0.12404569 |
| Q99MR8 | Mccc1 | Methylcrotonoyl-CoA carboxylase subunit alpha, mitochondrial | 0.261317507 | -0.849467278 |
| Q99N12 | Dusp19 | Dual specificity phosphatase 19 | 0.019124158 | 1.026458104 |
| Q99NB1 | Acss1 | Acetyl-coenzyme A synthetase 2-like, mitochondrial | -0.069573784 | 0.247955481 |
| Q99NB8 | Ubqln4 | Ubiquilin-4 | -0.486171436 | 1.158451557 |
| Q99NF7 | Ppm1b | Protein-serine/threonine phosphatase | -0.168075943 | 0.206560453 |
| Q99P31 | Hspbp1 | Hsp70-binding protein 1 | -0.540505886 | 0.132482529 |
| Q99P58 | Rab27b | Ras-related protein Rab-27B | 0.428013452 | 1.475896994 |
| Q99P72 | Rtn4 | Reticulon-4 | 0.054315631 | 0.413041751 |
| Q99PG2 | Ogfr | Opioid growth factor receptor | 0.554747295 | -0.522898833 |
| Q99PL6 | Ubxn6 | UBX domain-containing protein 6 | 0.248359426 | -0.010479609 |
| Q99PS0 | Krt23 | Keratin, type I cytoskeletal 23 | 0.355544567 | -1.438218911 |
| Q99PT1 | Arhgdia | Rho GDP-dissociation inhibitor 1 | 0.055879656 | 0.25243632 |
| Q99PU5 | Acsbg1 | Long-chain-fatty-acid--CoA ligase ACSBG1 | 0.135233625 | 0.08968242 |
| Q99PV0 | Prpf8 | Pre-mRNA-processing-splicing factor 8 | 0.494051647 | 0.208118916 |
| Q9CPN9 | 2210010C04Rik | RIKEN cDNA 2210010C04 gene | -0.25067145 | 2.414047241 |
| Q9CPP6 | Ndufa5 | NADH dehydrogenase [ubiquinone] 1 alpha subcomplex subunit 5 | 0.361294429 | 1.64205424 |
| Q9CPQ1 | Cox6c | Cytochrome c oxidase subunit 6C | 0.480538082 | 1.054711342 |
| Q9CPQ8 | Atp5mg | ATP synthase subunit g, mitochondrial | 0.84404761 | 2.344562372 |
| Q9CPR1 | Rwdd4 | RWD domain-containing protein 4 | -0.081606611 | -0.197167397 |
| Q9CPS5 | Psmd8 | 26S proteasome non-ATPase regulatory subunit 8 | 0.240795708 | 0.246686459 |
| Q9CPS6 | Hint3 | Histidine triad nucleotide-binding protein 3 | 0.202594058 | -0.637357712 |
| Q9CPT3 | Nanp | N-acylneuraminate-9-phosphatase | 0.034366481 | 0.856114546 |
| Q9CPT4 | Mydgf | Myeloid-derived growth factor | -0.037504292 | -0.361679872 |
| Q9CPU0 | Glo1 | Lactoylglutathione lyase | 0.032203865 | 0.399954478 |
| Q9CPU4 | Mgst3 | Microsomal glutathione S-transferase 3 | 0.766653824 | 0.017497222 |
| Q9CPV4 | Glod4 | Glyoxalase domain-containing protein 4 | -0.180279223 | 0.612804731 |

| | | | | |
|---|---|---|---|---|
| Q9CPW2 | Fdx2 | Ferredoxin-2, mitochondrial | -0.293359534 | 0.012481689 |
| Q9CPW4 | Arpc5 | Actin-related protein 2/3 complex subunit 5 | 0.289819082 | 1.070185343 |
| Q9CPW7 | Zmat2 | Zinc finger matrin-type protein 2 | -0.937470055 | -0.272417863 |
| Q9CPX6 | Atg3 | Ubiquitin-like-conjugating enzyme ATG3 | 0.134826183 | 0.417051792 |
| Q9CPY7 | Lap3 | Cytosol aminopeptidase | -0.110540136 | 0.22821792 |
| Q9CPZ8 | Cmc1 | COX assembly mitochondrial protein homolog | 1.156425937 | 0.030606747 |
| Q9CQ10 | Chmp3 | Charged multivesicular body protein 3 | 0.200082652 | -0.043589433 |
| Q9CQ20 | Mid1ip1 | Mid1-interacting protein 1 | -0.796034781 | -0.538640658 |
| Q9CQ26 | Stambp | STAM-binding protein | 0.08920873 | 0.409808795 |
| Q9CQ45 | Nenf | Neudesin | -1.866147423 | -1.395198504 |
| Q9CQ49 | Ncbp2 | Nuclear cap-binding protein subunit 2 | -0.314728292 | -0.441348871 |
| Q9CQ60 | Pgls | 6-phosphogluconolactonase | -0.140453688 | 0.997837067 |
| Q9CQ62 | Decr1 | 2,4-dienoyl-CoA reductase [(3E)-enoyl-CoA-producing], mitochondrial | 0.29505415 | 1.783194701 |
| Q9CQ65 | Mtap | S-methyl-5'-thioadenosine phosphorylase | -0.188666725 | 0.608890216 |
| Q9CQ69 | Uqcrq | Cytochrome b-c1 complex subunit 8 | 0.347401206 | 1.497856458 |
| Q9CQ75 | Ndufa2 | NADH dehydrogenase [ubiquinone] 1 alpha subcomplex subunit 2 | 0.152480952 | 0.814250787 |
| Q9CQ86 | Mien1 | Migration and invasion enhancer 1 | -0.256930415 | -0.143651644 |
| Q9CQA1 | Trappc5 | Trafficking protein particle complex subunit 5 | 1.076625156 | -0.997790972 |
| Q9CQA3 | Sdhb | Succinate dehydrogenase [ubiquinone] iron-sulfur subunit, mitochondrial | 0.00615139 | 0.24949042 |
| Q9CQB4 | Uqcrb | Cytochrome b-c1 complex subunit 7 | 0.189001211 | 0.778785706 |
| Q9CQB7 | Lyrm1 | LYR motif-containing protein 1 | 0.709292126 | -0.328081926 |
| Q9CQC6 | Bzw1 | Basic leucine zipper and W2 domain-containing protein 1 | 0.076113002 | 0.431104819 |
| Q9CQC7 | Ndufb4 | NADH dehydrogenase [ubiquinone] 1 beta subcomplex subunit 4 | -0.813363171 | 1.473211567 |
| Q9CQC9 | Sar1b | GTP-binding protein SAR1b | 0.346105703 | 1.459559282 |
| Q9CQD1 | Rab5a | Ras-related protein Rab-5A | 0.136680603 | 0.657502651 |
| Q9CQD4 | Chmp1b2 | Charged multivesicular body protein 1b-2 | 0.984914017 | -0.242534161 |
| Q9CQE1 | Nipsnap3b | Protein NipSnap homolog 3B | 0.428200277 | -0.193992615 |
| Q9CQE5 | Rgs10 | Regulator of G-protein signaling 10 | 0.013813972 | 0.756342093 |
| Q9CQE8 | RTRAF | RNA transcription, translation and transport factor protein | 0.45432663 | 0.634683132 |
| Q9CQF3 | Nudt21 | Cleavage and polyadenylation specificity factor subunit 5 | 0.219377454 | 0.31084919 |
| Q9CQF4 | Mtres1 | Mitochondrial transcription rescue factor 1 | 0.582178911 | -0.041949749 |
| Q9CQG1 | Chac2 | Putative glutathione-specific gamma-glutamylcyclotransferase 2 | 0.049762821 | -1.311399122 |
| Q9CQH7 | Btf3l4 | Transcription factor BTF3 homolog 4 | 0.103276475 | 0.208954016 |
| Q9CQI3 | Gmfb | Glia maturation factor beta | 0.013359006 | 0.298250039 |
| Q9CQI6 | Cotl1 | Coactosin-like protein | 0.02097791 | 0.469311078 |
| Q9CQI7 | Snrpb2 | U2 small nuclear ribonucleoprotein B'' | 0.385619736 | 1.430719614 |
| Q9CQJ6 | Denr | Density-regulated protein | 0.419896412 | 0.350137075 |
| Q9CQJ8 | Ndufb9 | NADH dehydrogenase [ubiquinone] 1 beta subcomplex subunit 9 | 1.110575453 | 1.925425847 |
| Q9CQK7 | Rwdd1 | RWD domain-containing protein 1 | 0.190943877 | -0.160598596 |
| Q9CQM5 | Txndc17 | Thioredoxin domain-containing protein 17 | -0.192596944 | -0.218744119 |
| Q9CQM9 | Glrx3 | Glutaredoxin-3 | -0.050147788 | 0.699048837 |
| Q9CQN1 | Trap1 | Heat shock protein 75 kDa, mitochondrial | -0.040216064 | 0.70780309 |
| Q9CQN6 | Tmem14c | Transmembrane protein 14C | 1.748123503 | 0.541096369 |
| Q9CQQ7 | Atp5pb | ATP synthase F(0) complex subunit B1, mitochondrial | 0.465152359 | 0.868288676 |

| | | | | |
|---|---|---|---|---|
| Q9CQQ8 | Lsm7 | U6 snRNA-associated Sm-like protein LSm7 | 0.015574455 | 0.218568643 |
| Q9CQR2 | Rps21 | 40S ribosomal protein S21 | 0.248255952 | 0.735985279 |
| Q9CQR4 | Acot13 | Acyl-coenzyme A thioesterase 13 | 0.13105243 | 0.732195695 |
| Q9CQS8 | Sec61b | Protein transport protein Sec61 subunit beta | 1.065995185 | 1.406854312 |
| Q9CQT0 | Thg1l | tRNA(His) guanylyltransferase | -0.013272031 | 0.095161756 |
| Q9CQT1 | Mri1 | Methylthioribose-1-phosphate isomerase | -0.048339812 | 0.114230951 |
| Q9CQU0 | Txndc12 | Thioredoxin domain-containing protein 12 | 0.309875202 | -0.175206184 |
| Q9CQV6 | Map1lc3b | Microtubule-associated proteins 1A/1B light chain 3B | 0.197794278 | 0.563353697 |
| Q9CQV8 | Ywhab | 14-3-3 protein beta/alpha | -0.020331891 | 0.281334241 |
| Q9CQW1 | Ykt6 | Synaptobrevin homolog YKT6 | 0.030015691 | 0.716917515 |
| Q9CQW2 | Arl8b | ADP-ribosylation factor-like protein 8B | 0.904728572 | 1.021373749 |
| Q9CQX6 | Gm16286 | Predicted gene 16286 | -0.024098365 | 0.112018426 |
| Q9CQX8 | Mrps36 | 28S ribosomal protein S36, mitochondrial | 0.568525187 | -1.347451687 |
| Q9CQY1 | Atg12 | Ubiquitin-like protein ATG12 | 2.461484957 | -0.895286719 |
| Q9CQY2 | Ramac | RNA guanine-N7 methyltransferase activating subunit | 0.34870952 | 0.490951379 |
| Q9CQZ5 | Ndufa6 | NADH dehydrogenase [ubiquinone] 1 alpha subcomplex subunit 6 | -0.78185962 | -0.439368725 |
| Q9CQZ6 | Ndufb3 | NADH dehydrogenase [ubiquinone] 1 beta subcomplex subunit 3 | -0.103890888 | 0.555858135 |
| Q9CR00 | Psmd9 | 26S proteasome non-ATPase regulatory subunit 9 | 0.183867296 | 0.33244578 |
| Q9CR09 | Ufc1 | Ubiquitin-fold modifier-conjugating enzyme 1 | 0.03954579 | 0.255721887 |
| Q9CR16 | Ppid | Peptidyl-prolyl cis-trans isomerase D | 0.047241402 | 0.076734225 |
| Q9CR25 | Dph2 | 2-(3-amino-3-carboxypropyl)histidine synthase subunit 2 | -0.436863041 | 0.58396101 |
| Q9CR26 | Vta1 | Vacuolar protein sorting-associated protein VTA1 homolog | -0.080560621 | 0.425917943 |
| Q9CR27 | Washc3 | WASH complex subunit 3 | -0.308755302 | 1.028100808 |
| Q9CR29 | Ccdc43 | Coiled-coil domain-containing protein 43 | -0.042462794 | -2.774187041 |
| Q9CR51 | Atp6v1g1 | V-type proton ATPase subunit G 1 | -0.176804352 | -1.327363133 |
| Q9CR57 | Rpl14 | 60S ribosomal protein L14 | -0.01302592 | -1.681727807 |
| Q9CR61 | Ndufb7 | NADH dehydrogenase [ubiquinone] 1 beta subcomplex subunit 7 | -0.493537585 | 0.770555973 |
| Q9CR62 | Slc25a11 | Mitochondrial 2-oxoglutarate/malate carrier protein | -0.186312103 | 0.238960107 |
| Q9CR68 | Uqcrfs1 | Cytochrome b-c1 complex subunit Rieske, mitochondrial | 0.261266295 | -0.027386824 |
| Q9CR86 | Carhsp1 | Calcium-regulated heat stable protein 1 | 0.24569149 | 0.56755654 |
| Q9CR95 | Necap1 | Adaptin ear-binding coat-associated protein 1 | 0.134851837 | 0.556170781 |
| Q9CR98 | Fam136a | Protein FAM136A | 0.170453517 | 0.339132309 |
| Q9CRA5 | Golph3 | Golgi phosphoprotein 3 | -0.518810145 | 0.628672918 |
| Q9CRA7 | Dmac2l | ATP synthase subunit s, mitochondrial | -0.013685767 | 1.601581573 |
| Q9CRB2 | Nhp2 | H/ACA ribonucleoprotein complex subunit 2 | -0.80028197 | 0.326580048 |
| Q9CRB4 | 1700029F12Rik | RIKEN cDNA 1700029F12 gene | -1.373569775 | 3.493388335 |
| Q9CRB6 | Tppp3 | Tubulin polymerization-promoting protein family member 3 | 0.351796532 | 0.387441794 |
| Q9CRC9 | Gnpda2 | Glucosamine-6-phosphate isomerase 2 | -0.008902232 | -0.011956692 |
| Q9CRD4 | Dbndd2 | Dysbindin domain-containing protein 2 | 0.489551004 | 0.650629997 |
| Q9CS42 | Prps2 | Ribose-phosphate pyrophosphokinase 2 | -0.030240409 | -0.260152181 |
| Q9CS84 | Nrxn1 | Neurexin-1 | -0.459629409 | 0.758566221 |
| Q9CVB6 | Arpc2 | Actin-related protein 2/3 complex subunit 2 | 0.235042826 | 0.809438388 |
| Q9CVD2 | Atxn3 | Ataxin-3 | 0.655301762 | 1.168317 |
| Q9CW07 | Ppp1r3g | Protein phosphatase 1 regulatory subunit 3G | 0.260245164 | 0.399226348 |
| Q9CW46 | Raver1 | Ribonucleoprotein PTB-binding 1 | 0.223395697 | 0.333305518 |

| | | | | |
|---|---|---|---|---|
| Q9CW79 | Golga1 | Golgin subfamily A member 1 | 2.595744451 | -1.648503304 |
| Q9CWD3 | Nudt17 | Nucleoside diphosphate-linked moiety X motif 17 | -0.354892 | 0.079901218 |
| Q9CWF2 | Tubb2b | Tubulin beta-2B chain | 0.08187453 | -0.520106634 |
| Q9CWG8 | Ndufaf7 | Protein arginine methyltransferase NDUFAF7, mitochondrial | -0.155256526 | 0.030196985 |
| Q9CWI3 | Bccip | BRCA2 and CDKN1A-interacting protein | -0.957467111 | -0.077167352 |
| Q9CWJ9 | Atic | Bifunctional purine biosynthesis protein ATIC | 0.001161512 | 0.605788231 |
| Q9CWK3 | Cd2bp2 | CD2 antigen cytoplasmic tail-binding protein 2 | 0.332051309 | -0.382962227 |
| Q9CWK8 | Snx2 | Sorting nexin-2 | -0.043019263 | 0.0327185 |
| Q9CWL8 | Ctnnbl1 | Beta-catenin-like protein 1 | -0.909023857 | 0.254550139 |
| Q9CWN7 | Cnot11 | CCR4-NOT transcription complex subunit 11 | 0.453932667 | 0.562881708 |
| Q9CWS0 | Ddah1 | N(G),N(G)-dimethylarginine dimethylaminohydrolase 1 | -0.162765185 | 0.258476893 |
| Q9CWZ3 | Rbm8a | RNA-binding protein 8A | 0.211080265 | -0.439043522 |
| Q9CWZ7 | Napg | Gamma-soluble NSF attachment protein | 0.005681419 | 0.361488342 |
| Q9CX00 | Ist1 | IST1 homolog | 0.516192627 | 0.870552699 |
| Q9CX34 | Sugt1 | Protein SGT1 homolog | -0.067481931 | 0.40738074 |
| Q9CX80 | Cygb | Cytoglobin | 0.081089783 | 0.344236692 |
| Q9CX86 | Hnrnpa0 | Heterogeneous nuclear ribonucleoprotein A0 | 0.232659054 | -0.565771262 |
| Q9CXF4 | Tbc1d15 | TBC1 domain family member 15 | -0.310621548 | -0.552865505 |
| Q9CXG3 | Ppil4 | Peptidyl-prolyl cis-trans isomerase-like 4 | -0.314405314 | -0.07925574 |
| Q9CXJ1 | Ears2 | Probable glutamate--tRNA ligase, mitochondrial | -0.755601756 | 0.799687227 |
| Q9CXT8 | Pmpcb | Mitochondrial-processing peptidase subunit beta | 0.1413421 | 0.341469765 |
| Q9CXW3 | Cacybp | Calcyclin-binding protein | 0.12934138 | 0.187545458 |
| Q9CXW4 | Rpl11 | 60S ribosomal protein L11 | 0.489271196 | 0.630515893 |
| Q9CXY6 | Ilf2 | Interleukin enhancer-binding factor 2 | -0.405085627 | 2.141797225 |
| Q9CY27 | Tecr | Very-long-chain enoyl-CoA reductase | 0.376054223 | 0.551691373 |
| Q9CY34 | Ube2f | NEDD8-conjugating enzyme UBE2F | 0.293479093 | 1.438507557 |
| Q9CY58 | Serbp1 | Plasminogen activator inhibitor 1 RNA-binding protein | 0.343562794 | 0.610083262 |
| Q9CY62 | Rnf181 | E3 ubiquitin-protein ligase RNF181 | -0.511585522 | 1.289090157 |
| Q9CY64 | Blvra | Biliverdin reductase A | -0.10236702 | 0.428557237 |
| Q9CY97 | Ssu72 | RNA polymerase II subunit A C-terminal domain phosphatase SSU72 | -0.189237563 | 0.323459307 |
| Q9CYA0 | Creld2 | Protein disulfide isomerase Creld2 | 0.572075017 | 0.57668829 |
| Q9CYA6 | Zcchc8 | Zinc finger CCHC domain-containing protein 8 | -1.034908803 | -0.055484533 |
| Q9CYG7 | Tomm34 | Mitochondrial import receptor subunit TOM34 | 0.05003589 | 0.158270518 |
| Q9CYT6 | Cap2 | Adenylyl cyclase-associated protein 2 | -0.026615842 | -0.339025497 |
| Q9CYU6 | Dph7 | Diphthine methyltransferase | -0.138802115 | -0.04280297 |
| Q9CYW4 | Hdhd3 | Haloacid dehalogenase-like hydrolase domain-containing protein 3 | 0.063792578 | 0.071290175 |
| Q9CZ04 | Cops7a | COP9 signalosome complex subunit 7a | -0.255926228 | 0.406137943 |
| Q9CZ13 | Uqcrc1 | Cytochrome b-c1 complex subunit 1, mitochondrial | 0.111979644 | 0.594029586 |
| Q9CZ30 | Ola1 | Obg-like ATPase 1 | -0.020484289 | 0.135638873 |
| Q9CZ44 | Nsfl1c | NSFL1 cofactor p47 | -0.096943728 | 0.028851032 |
| Q9CZ49 | Klhl35 | Kelch-like protein 35 | -2.228120359 | 0.957502842 |
| Q9CZC8 | Scrn1 | Secernin-1 | -0.130798213 | 0.201165199 |
| Q9CZD3 | Gars1 | Glycine--tRNA ligase | 0.026540947 | 0.348540147 |
| Q9CZH3 | Psmg3 | Proteasome assembly chaperone 3 | -6.403993758 | -5.285201232 |
| Q9CZM2 | Rpl15 | 60S ribosomal protein L15 | 1.08100996 | 0.401910146 |

| | | | | |
|---|---|---|---|---|
| Q9CZN7 | Shmt2 | Serine hydroxymethyltransferase, mitochondrial | 0.246083132 | 1.246240775 |
| Q9CZN8 | Qrsl1 | Glutamyl-tRNA(Gln) amidotransferase subunit A, mitochondrial | -0.240541903 | -0.122689088 |
| Q9CZP7 | Cdc37l1 | Hsp90 co-chaperone Cdc37-like 1 | 0.280356057 | 0.55075264 |
| Q9CZR8 | Tsfm | Elongation factor Ts, mitochondrial | 0.151749198 | 1.755477111 |
| Q9CZS1 | Aldh1b1 | Aldehyde dehydrogenase X, mitochondrial | 0.393769519 | 0.283890565 |
| Q9CZT8 | Rab3b | Ras-related protein Rab-3B | 0.236980597 | 1.079191844 |
| Q9CZU3 | Mtrex | Exosome RNA helicase MTR4 | 0.21383791 | 0.383194606 |
| Q9CZU6 | Cs | Citrate synthase, mitochondrial | -0.140885862 | 0.325666746 |
| Q9CZW4 | Acsl3 | Long-chain-fatty-acid--CoA ligase 3 | -0.32626044 | 1.463312149 |
| Q9CZW5 | Tomm70 | Mitochondrial import receptor subunit TOM70 | 0.327673054 | 0.716770808 |
| Q9CZX8 | Rps19 | 40S ribosomal protein S19 | 0.387564405 | 0.096848965 |
| Q9CZY3 | Ube2v1 | Ubiquitin-conjugating enzyme E2 variant 1 | 0.261027908 | 0.378947576 |
| Q9D020 | Nt5c3a | Cytosolic 5'-nucleotidase 3A | 0.072579543 | 1.000203609 |
| Q9D051 | Pdhb | Pyruvate dehydrogenase E1 component subunit beta, mitochondrial | 0.431170813 | 0.70884339 |
| Q9D061 | Acbd6 | Acyl-CoA-binding domain-containing protein 6 | 0.108611425 | 0.475955486 |
| Q9D071 | Mms19 | MMS19 nucleotide excision repair protein homolog | 0.155616665 | 1.628985564 |
| Q9D0A3 | Arpin | Arpin | 0.019963042 | 0.295549711 |
| Q9D0B5 | Tstd3 | Thiosulfate sulfurtransferase/rhodanese-like domain-containing protein 3 | -0.89857591 | 0.167888165 |
| Q9D0B6 | Pbdc1 | Protein PBDC1 | 0.647399426 | 0.7018013 |
| Q9D0E1 | Hnrnpm | Heterogeneous nuclear ribonucleoprotein M | 0.118320243 | -0.681218147 |
| Q9D0E3 | Lysmd1 | LysM and putative peptidoglycan-binding domain-containing protein 1 | -0.208758068 | -0.2833999 |
| Q9D0F9 | Pgm1 | Phosphoglucomutase-1 | -0.219494756 | 0.012053172 |
| Q9D0I6 | Wdsub1 | WD repeat, SAM and U-box domain-containing protein 1 | -0.526336416 | 1.187075456 |
| Q9D0I9 | Rars1 | Arginine--tRNA ligase, cytoplasmic | -0.023154608 | 0.097572327 |
| Q9D0J4 | Arl2 | ADP-ribosylation factor-like protein 2 | -0.221982797 | 0.877692858 |
| Q9D0J8 | Ptms | Parathymosin | -0.131187216 | -0.691634337 |
| Q9D0K2 | Oxct1 | Succinyl-CoA:3-ketoacid coenzyme A transferase 1, mitochondrial | 0.005765851 | 0.189682007 |
| Q9D0L8 | Rnmt | mRNA cap guanine-N7 methyltransferase | -0.188956865 | 0.064000607 |
| Q9D0M3 | Cyc1 | Cytochrome c1, heme protein, mitochondrial | 0.138555654 | 0.328148047 |
| Q9D0M5 | Dynll2 | Dynein light chain 2, cytoplasmic | 0.562149111 | 1.21783034 |
| Q9D0R2 | Tars1 | Threonine--tRNA ligase 1, cytoplasmic | 0.012758986 | 1.36641264 |
| Q9D0R8 | Lsm12 | Protein LSM12 homolog | 0.014750195 | 0.05562067 |
| Q9D0S9 | Hint2 | Histidine triad nucleotide-binding protein 2, mitochondrial | 0.036141205 | 0.777694861 |
| Q9D0T1 | Snu13 | NHP2-like protein 1 | 0.085502275 | -0.552443504 |
| Q9D0W5 | Ppil1 | Peptidyl-prolyl cis-trans isomerase-like 1 | -0.287996515 | -0.414646467 |
| Q9D142 | Nudt14 | Uridine diphosphate glucose pyrophosphatase NUDT14 | 0.29092439 | -0.316794237 |
| Q9D154 | Serpinb1a | Leukocyte elastase inhibitor A | -0.069240443 | 0.345786413 |
| Q9D172 | Gatd3a | Glutamine amidotransferase-like class 1 domain-containing protein 3A, mitochondrial | -0.097226143 | 0.466411273 |
| Q9D1A2 | Cndp2 | Cytosolic non-specific dipeptidase | -0.007393837 | 0.188247681 |
| Q9D1C8 | Vps28 | Vacuolar protein sorting-associated protein 28 homolog | -0.273992443 | 0.796913306 |
| Q9D1E6 | Tbcb | Tubulin-folding cofactor B | -0.172103691 | 0.379100482 |
| Q9D1G1 | Rab1b | Ras-related protein Rab-1B | -0.044823106 | -0.175121625 |
| Q9D1H7 | Get4 | Golgi to ER traffic protein 4 homolog | 0.29963239 | 0.073179086 |
| Q9D1J1 | Necap2 | Adaptin ear-binding coat-associated protein 2 | -0.395791086 | -1.324465195 |

| | | | | |
|---|---|---|---|---|
| Q9D1J3 | Sarnp | SAP domain-containing ribonucleoprotein | 0.269025485 | 0.483207226 |
| Q9D1K2 | Atp6v1f | V-type proton ATPase subunit F | 0.001714547 | -1.262481372 |
| Q9D1M0 | Sec13 | Protein SEC13 homolog | 0.079809984 | 0.469106038 |
| Q9D1M4 | Eef1e1 | Eukaryotic translation elongation factor 1 epsilon-1 | 1.1593853 | -1.218819141 |
| Q9D1P4 | Chordc1 | Cysteine and histidine-rich domain-containing protein 1 | -0.237991778 | -0.060557365 |
| Q9D1Q6 | Erp44 | Endoplasmic reticulum resident protein 44 | -0.17235829 | 0.319630305 |
| Q9D1R9 | Rpl34 | 60S ribosomal protein L34 | 0.846097755 | 0.836380323 |
| Q9D273 | Mmab | Corrinoid adenosyltransferase | 0.267037169 | 0.214512189 |
| Q9D2C2 | Saal1 | Protein SAAL1 | -0.344705264 | -0.209294637 |
| Q9D2G2 | Dlst | Dihydrolipoyllysine-residue succinyltransferase component of 2-oxoglutarate dehydrogenase complex, mitochondrial | 0.641023318 | 0.310432911 |
| Q9D2M8 | Ube2v2 | Ubiquitin-conjugating enzyme E2 variant 2 | 0.13579642 | 0.544256687 |
| Q9D2N9 | Vps33a | Vacuolar protein sorting-associated protein 33A | -0.616489792 | 0.27063338 |
| Q9D2Q8 | S100a14 | Protein S100-A14 | 0.45364062 | -0.846760114 |
| Q9D2R0 | Aacs | Acetoacetyl-CoA synthetase | -0.150839806 | 0.427500884 |
| Q9D2V7 | Coro7 | Coronin-7 | 0.074939982 | 0.257266998 |
| Q9D358 | Acp1 | Low molecular weight phosphotyrosine protein phosphatase | 0.173071734 | -0.569061597 |
| Q9D385 | Arl2bp | ADP-ribosylation factor-like protein 2-binding protein | -1.18689607 | 0.116432826 |
| Q9D394 | Rufy3 | Protein RUFY3 | -0.145520083 | 0.262126446 |
| Q9D3D0 | Ttpal | Alpha-tocopherol transfer protein-like | -0.045560551 | 1.252351681 |
| Q9D3D9 | Atp5f1d | ATP synthase subunit delta, mitochondrial | 0.478252093 | 0.575553576 |
| Q9D404 | Oxsm | 3-oxoacyl-[acyl-carrier-protein] synthase, mitochondrial | 0.209889062 | 1.942657113 |
| Q9D4C9 | Clvs1 | Clavesin-1 | 0.181151772 | 1.328204632 |
| Q9D4H2 | Gcc1 | GRIP and coiled-coil domain-containing protein 1 | 0.054934947 | -0.324516455 |
| Q9D4H8 | Cul2 | Cullin-2 | -0.012224611 | -0.144267718 |
| Q9D4I9 | Rab23 | Ras-related protein Rab-23 | 0.052057997 | 0.387166023 |
| Q9D4J1 | Efhd1 | EF-hand domain-containing protein D1 | -0.240897465 | 1.245442788 |
| Q9D554 | Sf3a3 | Splicing factor 3A subunit 3 | 0.470491123 | -0.737774849 |
| Q9D5J6 | Shpk | Sedoheptulokinase | 0.036063894 | 1.455506961 |
| Q9D5V5 | Cul5 | Cullin-5 | -0.045363712 | 0.130580584 |
| Q9D5V6 | Syap1 | Synapse-associated protein 1 | 0.345703824 | -0.368486722 |
| Q9D646 | Krt34 | Keratin, type I cuticular Ha4 | 1.106880093 | -0.797065894 |
| Q9D662 | Sec23b | Protein transport protein Sec23B | -0.04701821 | -0.086735725 |
| Q9D6E4 | Ttc9b | Tetratricopeptide repeat protein 9B | 0.941756916 | 0.738278071 |
| Q9D6F9 | Tubb4a | Tubulin beta-4A chain | -0.421510887 | 0.150794983 |
| Q9D6H2 | Hspb11 | Intraflagellar transport protein 25 homolog | 0.508115896 | 0.073940118 |
| Q9D6J6 | Ndufv2 | NADH dehydrogenase [ubiquinone] flavoprotein 2, mitochondrial | 0.593388398 | 1.171664238 |
| Q9D6L8 | Ppil3 | Peptidyl-prolyl cis-trans isomerase-like 3 | 0.385405636 | -0.014722029 |
| Q9D6M3 | Slc25a22 | Mitochondrial glutamate carrier 1 | 0.213526789 | 0.571264267 |
| Q9D6P8 | Calml3 | Calmodulin-like protein 3 | 0.113606866 | -1.796571732 |
| Q9D6R2 | Idh3a | Isocitrate dehydrogenase [NAD] subunit alpha, mitochondrial | -0.095474625 | 0.54514726 |
| Q9D6S7 | Mrrf | Ribosome-recycling factor, mitochondrial | -0.21058197 | 0.989789883 |
| Q9D6W8 | Borcs6 | BLOC-1-related complex subunit 6 | 0.457356008 | 0.105734984 |
| Q9D6Y7 | Msra | Mitochondrial peptide methionine sulfoxide reductase | -0.229800701 | 0.4281497 |
| Q9D706 | Rpap3 | RNA polymerase II-associated protein 3 | -0.201555665 | -0.537773291 |

| | | | | |
|---|---|---|---|---|
| Q9D708 | S100a16 | Protein S100-A16 | 0.090470664 | -0.390022278 |
| Q9D7A8 | Armc1 | Armadillo repeat-containing protein 1 | 0.28950386 | 0.214905262 |
| Q9D7H3 | RtcA | RNA 3'-terminal phosphate cyclase | 0.320702998 | -0.627337933 |
| Q9D7I5 | Lhpp | Phospholysine phosphohistidine inorganic pyrophosphate phosphatase | -0.270536677 | 1.715363503 |
| Q9D7N9 | Apmap | Adipocyte plasma membrane-associated protein | -0.40054547 | 1.193078597 |
| Q9D7P6 | Iscu | Iron-sulfur cluster assembly enzyme ISCU, mitochondrial | -0.021397177 | 0.351029078 |
| Q9D7S7 | Rpl22l1 | 60S ribosomal protein L22-like 1 | -0.081073475 | -0.81424586 |
| Q9D7S9 | Chmp5 | Charged multivesicular body protein 5 | 0.22115469 | 0.54470555 |
| Q9D7X8 | Ggct | Gamma-glutamylcyclotransferase | 0.068821907 | 0.463652929 |
| Q9D819 | Ppa1 | Inorganic pyrophosphatase | -0.005095673 | 0.71156311 |
| Q9D820 | Prorsd1 | Prolyl-tRNA synthetase associated domain-containing protein 1 | 0.064640919 | 0.699301163 |
| Q9D832 | Dnajb4 | DnaJ homolog subfamily B member 4 | 0.769522953 | 0.659254233 |
| Q9D853 | Eef1akmt2 | EEF1A lysine methyltransferase 2 | 0.279551411 | 0.225643794 |
| Q9D883 | U2af1 | Splicing factor U2AF 35 kDa subunit | 0.110047817 | 0.676963647 |
| Q9D892 | Itpa | Inosine triphosphate pyrophosphatase | 0.282217725 | 0.7628088 |
| Q9D898 | Arpc5l | Actin-related protein 2/3 complex subunit 5-like protein | 0.225609239 | -0.394990762 |
| Q9D8B3 | Chmp4b | Charged multivesicular body protein 4b | 2.98E-01 | 0.357741356 |
| Q9D8E6 | Rpl4 | 60S ribosomal protein L4 | 0.438063272 | 1.28915596 |
| Q9D8L5 | Ccdc91 | Coiled-coil domain-containing protein 91 | -0.699565379 | 0.888944467 |
| Q9D8N0 | Eef1g | Elongation factor 1-gamma | -0.037552357 | 0.115379333 |
| Q9D8S4 | Rexo2 | Oligoribonuclease, mitochondrial | 0.133328978 | 0.776655992 |
| Q9D8S9 | Bola1 | BolA-like protein 1 | -0.068758424 | 0.205066363 |
| Q9D8U8 | Snx5 | Sorting nexin-5 | 0.122349866 | -0.166023254 |
| Q9D8W5 | Psmd12 | 26S proteasome non-ATPase regulatory subunit 12 | -0.126191616 | -0.002003034 |
| Q9D964 | Gatm | Glycine amidinotransferase, mitochondrial | -0.088460096 | 0.231538773 |
| Q9D967 | Mdp1 | Magnesium-dependent phosphatase 1 | 0.550641696 | 0.286132813 |
| Q9D9H8 | | UPF0565 protein C2orf69 homolog | -0.269767348 | -0.200796604 |
| Q9D9M5 | Phospho2 | Pyridoxal phosphate phosphatase PHOSPHO2 | -0.450953929 | 0.246092161 |
| Q9D9V3 | Echdc1 | Ethylmalonyl-CoA decarboxylase | 0.035974948 | 0.358151277 |
| Q9DA03 | Lyrm7 | Complex III assembly factor LYRM7 | -0.420396105 | -0.34763368 |
| Q9DAI2 | Ift22 | Intraflagellar transport protein 22 homolog | 0.349792067 | 1.394398212 |
| Q9DAK9 | Phpt1 | 14 kDa phosphohistidine phosphatase | -0.104124896 | 0.527506034 |
| Q9DAM7 | Tmem263 | Transmembrane protein 263 | 0.4941974 | 1.68179671 |
| Q9DAR7 | Dcps | m7GpppX diphosphatase | -0.072466087 | 0.499771118 |
| Q9DAT5 | Trmu | Mitochondrial tRNA-specific 2-thiouridylase 1 | -0.10179491 | 0.478078445 |
| Q9DAU1 | Cnpy3 | Protein canopy homolog 3 | 0.614664586 | 0.501169999 |
| Q9DAW9 | Cnn3 | Calponin-3 | -0.093041007 | 0.25798734 |
| Q9DAZ9 | Zfyve19 | Abscission/NoCut checkpoint regulator | 0.580169042 | 0.528602282 |
| Q9DB05 | Napa | Alpha-soluble NSF attachment protein | -0.148433558 | -0.284953117 |
| Q9DB15 | Mrpl12 | 39S ribosomal protein L12, mitochondrial | 0.69306167 | 2.44163688 |
| Q9DB16 | Cab39l | Calcium-binding protein 39-like | 0.230880578 | 0.854097684 |
| Q9DB20 | Atp5po | ATP synthase subunit O, mitochondrial | 0.168240611 | 1.7723972 |
| Q9DB27 | Mcts1 | Malignant T-cell-amplified sequence 1 | 0.291719913 | 0.000289281 |
| Q9DB29 | Iah1 | Isoamyl acetate-hydrolyzing esterase 1 homolog | 0.201455911 | 0.563068708 |
| Q9DB32 | Haghl | Hydroxyacylglutathione hydrolase-like protein | -0.205292956 | 0.322115739 |

| | | | | |
|---|---|---|---|---|
| Q9DB34 | Chmp2a | Charged multivesicular body protein 2a | 0.014757665 | -0.114843686 |
| Q9DB50 | Ap1s2 | AP-1 complex subunit sigma-2 | 0.118074767 | 0.044264476 |
| Q9DB60 | Prxl2b | Prostamide/prostaglandin F synthase | -0.091587766 | 0.942902406 |
| Q9DB77 | Uqcrc2 | Cytochrome b-c1 complex subunit 2, mitochondrial | 0.148736095 | 0.4004275 |
| Q9DBB8 | Dhdh | Trans-1,2-dihydrobenzene-1,2-diol dehydrogenase | -0.22556111 | 0.437476953 |
| Q9DBC0 | Selenoo | Protein adenylyltransferase SelO, mitochondrial | -0.149439367 | 0.088007609 |
| Q9DBC3 | Cmtr1 | Cap-specific mRNA (nucleoside-2'-O-)-methyltransferase 1 | -0.659578164 | 0.60668246 |
| Q9DBC7 | Prkar1a | cAMP-dependent protein kinase type I-alpha regulatory subunit | -0.203060913 | 0.085992813 |
| Q9DBE0 | Csad | Cysteine sulfinic acid decarboxylase | -0.216751226 | 0.630828698 |
| Q9DBF1 | Aldh7a1 | Alpha-aminoadipic semialdehyde dehydrogenase | -0.188617039 | 0.269646486 |
| Q9DBG3 | Ap2b1 | AP-2 complex subunit beta | -0.047379112 | 0.334677696 |
| Q9DBG5 | Plin3 | Perilipin-3 | -0.191439374 | 0.417452494 |
| Q9DBG9 | Tax1bp3 | Tax1-binding protein 3 | -0.400149314 | -0.113520304 |
| Q9DBJ1 | Pgam1 | Phosphoglycerate mutase 1 | -0.392658806 | 0.319819768 |
| Q9DBL1 | Acadsb | Short/branched chain specific acyl-CoA dehydrogenase, mitochondrial | -0.192298253 | 0.506065687 |
| Q9DBL7 | Coasy | Bifunctional coenzyme A synthase | -0.081973044 | 0.391194185 |
| Q9DBP5 | Cmpk1 | UMP-CMP kinase | -0.165692584 | 0.26763312 |
| Q9DBR0 | Akap8 | A-kinase anchor protein 8 | -0.415796026 | 0.257771651 |
| Q9DBR1 | Xrn2 | 5'-3' exoribonuclease 2 | -1.353430907 | -0.236963908 |
| Q9DBR7 | Ppp1r12a | Protein phosphatase 1 regulatory subunit 12A | -0.039984989 | -0.010243893 |
| Q9DBS5 | Klc4 | Kinesin light chain 4 | -0.207887236 | 0.333625793 |
| Q9DBX2 | Pdcl | Phosducin-like protein | -0.507782078 | 0.935072581 |
| Q9DBZ5 | Eif3k | Eukaryotic translation initiation factor 3 subunit K | 0.186619504 | -0.749997934 |
| Q9DC07 | Nebl | LIM zinc-binding domain-containing Nebulette | 0.041829491 | 0.401135445 |
| Q9DC22 | Dcaf6 | DDB1- and CUL4-associated factor 6 | -1.140384706 | -0.146187147 |
| Q9DC50 | Crot | Peroxisomal carnitine O-octanoyltransferase | 0.308085155 | 0.243565083 |
| Q9DC51 | Gnai3 | Guanine nucleotide-binding protein G(i) subunit alpha-3 | -0.80110124 | -0.568980853 |
| Q9DC61 | Pmpca | Mitochondrial-processing peptidase subunit alpha | 0.222509766 | 0.516581535 |
| Q9DC63 | Fbxo3 | F-box only protein 3 | -0.33538847 | -1.181020578 |
| Q9DC70 | Ndufs7 | NADH dehydrogenase [ubiquinone] iron-sulfur protein 7, mitochondrial | -1.050713476 | -0.581745942 |
| Q9DCB4 | Arpp21 | cAMP-regulated phosphoprotein 21 | 0.443482176 | 0.408137639 |
| Q9DCC4 | Pycr3 | Pyrroline-5-carboxylate reductase 3 | -0.189449946 | -0.047275225 |
| Q9DCD0 | Pgd | 6-phosphogluconate dehydrogenase, decarboxylating | 0.06582648 | 0.48983558 |
| Q9DCH4 | Eif3f | Eukaryotic translation initiation factor 3 subunit F | 0.060839144 | -0.238871892 |
| Q9DCJ1 | Mlst8 | Target of rapamycin complex subunit LST8 | 1.424321306 | 1.705774943 |
| Q9DCJ5 | Ndufa8 | NADH dehydrogenase [ubiquinone] 1 alpha subcomplex subunit 8 | -0.316278712 | 0.158759594 |
| Q9DCJ9 | Npl | N-acetylneuraminate lyase | -0.131410599 | -0.803738356 |
| Q9DCL8 | Ppp1r2 | Protein phosphatase inhibitor 2 | -0.174108028 | 0.110277494 |
| Q9DCL9 | Paics | Multifunctional protein ADE2 | 0.111182658 | 0.640442053 |
| Q9DCM0 | Ethe1 | Persulfide dioxygenase ETHE1, mitochondrial | -0.074747817 | -0.126637618 |
| Q9DCM2 | Gstk1 | Glutathione S-transferase kappa 1 | 0.348800087 | 2.00746568 |
| Q9DCN1 | Nudt12 | NAD-capped RNA hydrolase NUDT12 | 1.250739352 | -1.69688344 |
| Q9DCR2 | Ap3s1 | AP-3 complex subunit sigma-1 | 0.203468609 | 0.545028369 |
| Q9DCS2 | Mettl26 | Methyltransferase-like 26 | -0.160371017 | 0.345266183 |
| Q9DCS3 | Mecr | Enoyl-[acyl-carrier-protein] reductase, mitochondrial | -0.157481956 | 0.83151118 |

| | | | | |
|---|---|---|---|---|
| Q9DCT1 | Akr1e2 | 1,5-anhydro-D-fructose reductase | 0.206502183 | 0.478045146 |
| Q9DCT2 | Ndufs3 | NADH dehydrogenase [ubiquinone] iron-sulfur protein 3, mitochondrial | -2.032703447 | -0.096115748 |
| Q9DCT8 | Crip2 | Cysteine-rich protein 2 | -0.177671719 | 0.479729176 |
| Q9DCV4 | Rmdn1 | Regulator of microtubule dynamics protein 1 | 0.499997902 | 0.616429806 |
| Q9DCW4 | Etfb | Electron transfer flavoprotein subunit beta | -0.057212766 | 0.444683711 |
| Q9DCZ1 | Gmpr | GMP reductase 1 | -0.026262919 | 0.459355354 |
| Q9DCZ4 | Apoo | MICOS complex subunit Mic26 | 0.387436072 | 0.024131298 |
| Q9DD02 | Hikeshi | Protein Hikeshi | -0.948594602 | 0.078399499 |
| Q9DD18 | Dtd1 | D-aminoacyl-tRNA deacylase 1 | -0.308723132 | 0.5612216 |
| Q9EP53 | Tsc1 | Hamartin | 0.340252145 | 0.792668978 |
| Q9EPL2 | Clstn1 | Calsyntenin-1 | 0.006975269 | 0.283500036 |
| Q9EPL8 | Ipo7 | Importin-7 | -0.144386959 | 0.40314436 |
| Q9EPN1 | Nbea | Neurobeachin | -0.026334635 | -0.24891154 |
| Q9EPQ7 | Stard5 | StAR-related lipid transfer protein 5 | -0.32086366 | -0.264168898 |
| Q9EPU0 | Upf1 | Regulator of nonsense transcripts 1 | 0.404827309 | 1.23537159 |
| Q9EQ06 | Hsd17b11 | Estradiol 17-beta-dehydrogenase 11 | -0.130569935 | 1.354497433 |
| Q9EQ08 | Sgsh | Heparan N-sulfatase | -0.25626564 | -0.18197902 |
| Q9EQ20 | Aldh6a1 | Methylmalonate-semialdehyde dehydrogenase [acylating], mitochondrial | -0.037593969 | 0.348591169 |
| Q9EQ80 | Nif3l1 | NIF3-like protein 1 | -0.281942972 | 0.245337327 |
| Q9EQC8 | Prcc | Papillary Renal Cell carcinoma (Translocation-associated) | -1.636943404 | -0.179983934 |
| Q9EQF6 | Dpysl5 | Dihydropyrimidinase-related protein 5 | 0.090829849 | 0.210102717 |
| Q9EQG7 | Enpp5 | Ectonucleotide pyrophosphatase/phosphodiesterase family member 5 | -0.309972223 | -0.760838032 |
| Q9EQG9 | Cert1 | Ceramide transfer protein | 0.526047198 | 1.425781647 |
| Q9EQH2 | Erap1 | Endoplasmic reticulum aminopeptidase 1 | 0.822447968 | 0.053230286 |
| Q9EQH3 | Vps35 | Vacuolar protein sorting-associated protein 35 | -0.048638153 | 0.344895681 |
| Q9EQP2 | Ehd4 | EH domain-containing protein 4 | 1.34933246 | 2.687906027 |
| Q9EQQ9 | Oga | Protein O-GlcNAcase | -0.097235902 | 0.411819458 |
| Q9ER00 | Stx12 | Syntaxin-12 | 0.177465789 | 1.029426257 |
| Q9ER05 | Ctrl | Chymopasin | 0.215825049 | 0.240746021 |
| Q9ER35 | Fn3k | Fructosamine-3-kinase | 0.088849449 | 0.137296995 |
| Q9ER58 | Spock2 | Testican-2 | 0.245934391 | 0.693342527 |
| Q9ER72 | Cars1 | Cysteine--tRNA ligase, cytoplasmic | 0.059147135 | 0.31087478 |
| Q9ER73 | Elp4 | Elongator complex protein 4 | -0.030184778 | 0.901927551 |
| Q9ERD7 | Tubb3 | Tubulin beta-3 chain | -0.235023371 | 0.653243383 |
| Q9ERE7 | Mesd | LRP chaperone MESD | 0.047599665 | 0.485406081 |
| Q9ERF3 | Wdr61 | WD repeat-containing protein 61 | -0.094087442 | 0.046302954 |
| Q9ERG2 | Strn3 | Striatin-3 | 0.048267269 | 0.150609652 |
| Q9ERI6 | Rdh14 | Retinol dehydrogenase 14 | 0.350264041 | 2.22238938 |
| Q9ERK4 | Cse1l | Exportin-2 | -0.012914054 | 0.195955276 |
| Q9ERL9 | Gucy1a1 | Guanylate cyclase soluble subunit alpha-1 | -0.015156778 | 0.537081401 |
| Q9ERQ8 | Ca7 | Carbonic anhydrase 7 | -0.126340675 | 0.377914111 |
| Q9ERR1 | Ndel1 | Nuclear distribution protein nudE-like 1 | -0.089865812 | -0.022618294 |
| Q9ERS2 | Ndufa13 | NADH dehydrogenase [ubiquinone] 1 alpha subcomplex subunit 13 | -0.44875164 | 1.048749288 |
| Q9ERU9 | Ranbp2 | E3 SUMO-protein ligase RanBP2 | -0.175432173 | 1.097095788 |
| Q9ES28 | Arhgef7 | Rho guanine nucleotide exchange factor 7 | -0.103737259 | -0.195126216 |

| | | | | |
|---|---|---|---|---|
| Q9ES56 | Trappc4 | Trafficking protein particle complex subunit 4 | 0.43907938 | 1.262586912 |
| Q9ES74 | Nek7 | Serine/threonine-protein kinase Nek7 | -0.887618351 | 0.985719522 |
| Q9ES97 | Rtn3 | Reticulon-3 | 0.102406534 | 0.338472366 |
| Q9ESJ4 | Nckipsd | NCK-interacting protein with SH3 domain | -0.086725521 | 0.111334801 |
| Q9ESM3 | Hapln2 | Hyaluronan and proteoglycan link protein 2 | 0.019044272 | 1.064477523 |
| Q9ESN6 | Trim2 | Tripartite motif-containing protein 2 | 0.18714625 | 0.391760031 |
| Q9ESP1 | Sdf2l1 | Stromal cell-derived factor 2-like protein 1 | -1.404926586 | 0.814697107 |
| Q9EST1 | Gsdma | Gasdermin-A | 0.241076724 | -0.05045716 |
| Q9EST4 | Psmg2 | Proteasome assembly chaperone 2 | 0.158635871 | 0.847113848 |
| Q9EST5 | Anp32b | Acidic leucine-rich nuclear phosphoprotein 32 family member B | -0.086920706 | -0.2542003 |
| Q9ESW8 | Pgpep1 | Pyroglutamyl-peptidase 1 | -0.272013124 | 2.4098924 |
| Q9ET01 | Pygl | Glycogen phosphorylase, liver form | -0.486029243 | -0.21677955 |
| Q9ET22 | Dpp7 | Dipeptidyl peptidase 2 | -0.034938367 | 0.519420783 |
| Q9ET26 | Rnf114 | E3 ubiquitin-protein ligase RNF114 | -0.171035099 | 0.740194003 |
| Q9JHG0 | Cbln3 | Cerebellin-3 | -0.582908376 | -0.858700434 |
| Q9JHG6 | Rcan1 | Calcipressin-1 | -0.23984979 | -0.425274531 |
| Q9JHI5 | Ivd | Isovaleryl-CoA dehydrogenase, mitochondrial | -0.254151217 | -0.075072765 |
| Q9JHK4 | Rabggta | Geranylgeranyl transferase type-2 subunit alpha | -0.034773477 | 0.486203194 |
| Q9JHQ5 | Lztfl1 | Leucine zipper transcription factor-like protein 1 | 0.123950068 | 0.505829016 |
| Q9JHS3 | Lamtor2 | Ragulator complex protein LAMTOR2 | -0.122008546 | -0.153224309 |
| Q9JHS9 | Cwc15 | Spliceosome-associated protein CWC15 homolog | 0.399394258 | 0.333708445 |
| Q9JHU4 | Dync1h1 | Cytoplasmic dynein 1 heavy chain 1 | -0.128191439 | -0.362201055 |
| Q9JHU9 | Isyna1 | Inositol-3-phosphate synthase 1 | -0.004401875 | 0.370164553 |
| Q9JHW2 | Nit2 | Omega-amidase NIT2 | 0.112141577 | 0.425374667 |
| Q9JHX6 | Allc | Probable allantoicase | 2.027825546 | -0.242522796 |
| Q9JI11 | Stk4 | Serine/threonine-protein kinase 4 | -0.057840951 | 0.333349546 |
| Q9JI46 | Nudt3 | Diphosphoinositol polyphosphate phosphohydrolase 1 | -0.174658585 | 0.526937167 |
| Q9JI75 | Nqo2 | Ribosyldihydronicotinamide dehydrogenase [quinone] | -0.28498284 | 0.240621726 |
| Q9JI90 | Rnf14 | E3 ubiquitin-protein ligase RNF14 | -0.037613869 | 0.771457036 |
| Q9JI91 | Actn2 | Alpha-actinin-2 | 0.491034381 | 0.261003494 |
| Q9JIA7 | Sphk2 | Sphingosine kinase 2 | -0.68496453 | 0.464122534 |
| Q9JIF7 | Copb1 | Coatomer subunit beta | 0.020428944 | 0.065823714 |
| Q9JIG4 | Ppp1r3f | Protein phosphatase 1 regulatory subunit 3F | 0.664507993 | -0.176555951 |
| Q9JIG7 | Ccdc22 | Coiled-coil domain-containing protein 22 | -0.125241788 | -0.714517752 |
| Q9JIG8 | Praf2 | PRA1 family protein 2 | 0.742333825 | -1.3349576 |
| Q9JIH2 | Nup50 | Nuclear pore complex protein Nup50 | 0.415918636 | -0.650874138 |
| Q9JII6 | Akr1a1 | Aldo-keto reductase family 1 member A1 | -0.108078448 | 0.221992175 |
| Q9JIS5 | Sv2a | Synaptic vesicle glycoprotein 2A | 0.508957481 | 0.356059869 |
| Q9JIY5 | Htra2 | Serine protease HTRA2, mitochondrial | 0.282515748 | 0.199985027 |
| Q9JJ28 | Flii | Protein flightless-1 homolog | -0.219226011 | -0.03911225 |
| Q9JJA4 | Wdr12 | Ribosome biogenesis protein WDR12 | -1.53826437 | -0.079484105 |
| Q9JJC6 | Rilpl1 | RILP-like protein 1 | 0.002129873 | 0.427700837 |
| Q9JJF0 | Nap1l5 | Nucleosome assembly protein 1-like 5 | 1.646271833 | 1.48287948 |
| Q9JJG0 | Tacc2 | Transforming acidic coiled-coil-containing protein 2 | 0.07008969 | 0.027644475 |
| Q9JJI8 | Rpl38 | 60S ribosomal protein L38 | 1.02360398 | 1.3812747 |
| Q9JJK2 | Lancl2 | LanC-like protein 2 | -0.086822478 | 0.746965885 |

| | | | | |
|---|---|---|---|---|
| Q9JJL8 | Sars2 | Serine--tRNA ligase, mitochondrial | -0.193818442 | -0.396547318 |
| Q9JJU8 | Sh3bgrl | SH3 domain-binding glutamic acid-rich-like protein | -0.247832648 | 0.172148863 |
| Q9JJV2 | Pfn2 | Profilin-2 | -0.004963621 | 0.734194438 |
| Q9JJX7 | Tdp2 | Tyrosyl-DNA phosphodiesterase 2 | 1.665159289 | -0.094973564 |
| Q9JJZ2 | Tuba8 | Tubulin alpha-8 chain | -0.061418883 | 1.11220932 |
| Q9JK23 | Psmg1 | Proteasome assembly chaperone 1 | -0.178116163 | 2.336922169 |
| Q9JK38 | Gnpnat1 | Glucosamine 6-phosphate N-acetyltransferase | -0.673195918 | 1.324674765 |
| Q9JK42 | Pdk2 | [Pyruvate dehydrogenase (acetyl-transferring)] kinase isozyme 2, mitochondrial | -0.068828487 | 0.433629036 |
| Q9JK81 | Myg1 | MYG1 exonuclease | -0.278246403 | 0.137531122 |
| Q9JK92 | Hspb8 | Heat shock protein beta-8 | 0.202277247 | -2.09048605 |
| Q9JKB1 | Uchl3 | Ubiquitin carboxyl-terminal hydrolase isozyme L3 | -0.306162262 | 0.850188573 |
| Q9JKB3 | Ybx3 | Y-box-binding protein 3 | 0.826678944 | -0.543345928 |
| Q9JKC6 | Cend1 | Cell cycle exit and neuronal differentiation protein 1 | 0.869669151 | 0.474733988 |
| Q9JKC8 | Ap3m1 | AP-3 complex subunit mu-1 | 0.761103185 | 0.877383709 |
| Q9JKD3 | Scamp5 | Secretory carrier-associated membrane protein 5 | -0.016260433 | -0.636369864 |
| Q9JKF1 | Iqgap1 | Ras GTPase-activating-like protein IQGAP1 | -0.759422874 | 0.036413352 |
| Q9JKK0 | Rcan3 | Calcipressin-3 | 0.249962521 | -0.44731768 |
| Q9JKK7 | Tmod2 | Tropomodulin-2 | 0.323860677 | -0.009931405 |
| Q9JKL4 | Ndufaf3 | NADH dehydrogenase [ubiquinone] 1 alpha subcomplex assembly factor 3 | 0.325108083 | 0.488463402 |
| Q9JKN6 | Nova1 | RNA-binding protein Nova-1 | 0.084812546 | 0.545608838 |
| Q9JKR6 | Hyou1 | Hypoxia up-regulated protein 1 | 0.207761002 | 0.214867751 |
| Q9JKX6 | Nudt5 | ADP-sugar pyrophosphatase | 0.066856543 | 0.232856115 |
| Q9JKY0 | Cnot9 | CCR4-NOT transcription complex subunit 9 | 0.820779737 | 3.573826075 |
| Q9JKY5 | Hip1r | Huntingtin-interacting protein 1-related protein | -0.266636435 | 0.292839686 |
| Q9JL04 | Fmn2 | Formin-2 | -0.160192617 | -0.488301913 |
| Q9JL35 | Hmgn5 | High mobility group nucleosome-binding domain-containing protein 5 | -1.529198027 | 0.695908388 |
| Q9JL62 | Gltp | Glycolipid transfer protein | 0.275670465 | 0.695176601 |
| Q9JLB0 | Mpp6 | MAGUK p55 subfamily member 6 | -0.131676865 | 0.246022542 |
| Q9JLB2 | Mpp5 | MAGUK p55 subfamily member 5 | -0.783116945 | 0.263886929 |
| Q9JLI8 | Sart3 | Squamous cell carcinoma antigen recognized by T-cells 3 | -0.416350651 | -0.178068002 |
| Q9JLJ2 | Aldh9a1 | 4-trimethylaminobutyraldehyde dehydrogenase | -0.279429054 | 0.36068964 |
| Q9JLM8 | Dclk1 | Serine/threonine-protein kinase DCLK1 | 0.201310031 | -0.102779071 |
| Q9JLN9 | Mtor | Serine/threonine-protein kinase mTOR | 0.252703412 | -1.310521762 |
| Q9JLQ0 | Cd2ap | CD2-associated protein | -1.148249563 | 0.75780344 |
| Q9JLV1 | Bag3 | BAG family molecular chaperone regulator 3 | 0.302851137 | 0.46707805 |
| Q9JLV5 | Cul3 | Cullin-3 | -0.12078495 | 0.158657392 |
| Q9JM13 | Rabgef1 | Rab5 GDP/GTP exchange factor | 0.035465018 | 0.3195858 |
| Q9JM14 | Nt5c | 5'(3')-deoxyribonucleotidase, cytosolic type | -1.926533381 | -3.346525788 |
| Q9JM63 | Kcnj10 | ATP-sensitive inward rectifier potassium channel 10 | 0.132454999 | 0.423060338 |
| Q9JM76 | Arpc3 | Actin-related protein 2/3 complex subunit 3 | 0.220098432 | -0.136130015 |
| Q9JM96 | Cdc42ep4 | Cdc42 effector protein 4 | 0.204014015 | 0.628968716 |
| Q9JMA2 | Qtrt1 | Queuine tRNA-ribosyltransferase catalytic subunit 1 | -0.543324089 | 0.55806764 |
| Q9JMC3 | Dnaja4 | DnaJ homolog subfamily A member 4 | 0.324016794 | 0.207468351 |
| Q9JMD0 | Znf207 | BUB3-interacting and GLEBS motif-containing protein ZNF207 | 0.298949019 | 0.068638007 |
| Q9JMD3 | Stard10 | START domain-containing protein 10 | 0.382273801 | 0.687183062 |

| | | | | |
|---|---|---|---|---|
| Q9JME5 | Ap3b2 | AP-3 complex subunit beta-2 | -0.024757067 | 0.013994376 |
| Q9JME7 | Trappc2l | Trafficking protein particle complex subunit 2-like protein | 0.332265186 | 0.966573715 |
| Q9JMG1 | Edf1 | Endothelial differentiation-related factor 1 | -0.777645048 | 0.858087619 |
| Q9JMG7 | Hdgfl3 | Hepatoma-derived growth factor-related protein 3 | 0.25568711 | 0.554837068 |
| Q9JMH6 | Txnrd1 | Thioredoxin reductase 1, cytoplasmic | -0.107462025 | 0.197085539 |
| Q9QUH0 | Glrx | Glutaredoxin-1 | -0.179892445 | 1.844085693 |
| Q9QUI0 | Rhoa | Transforming protein RhoA | 0.19581941 | 0.481010119 |
| Q9QUK6 | Tlr4 | Toll-like receptor 4 | -0.179458046 | -0.710991859 |
| Q9QUM9 | Psma6 | Proteasome subunit alpha type-6 | 0.010075633 | 0.038008849 |
| Q9QUN9 | Dkk3 | Dickkopf-related protein 3 | 0.219362895 | 1.535451392 |
| Q9QUP5 | Hapln1 | Hyaluronan and proteoglycan link protein 1 | -0.436365827 | 0.170202573 |
| Q9QUR6 | Prep | Prolyl endopeptidase | -0.077428595 | 0.631149292 |
| Q9QUR7 | Pin1 | Peptidyl-prolyl cis-trans isomerase NIMA-interacting 1 | -0.275087674 | 0.544430097 |
| Q9QUR8 | Sema7a | Semaphorin-7A | 0.042053477 | 0.733533859 |
| Q9QVP9 | Ptk2b | Protein-tyrosine kinase 2-beta | 0.130511824 | 0.316868464 |
| Q9QWL7 | Krt17 | Keratin, type I cytoskeletal 17 | -0.501928329 | -0.753131549 |
| Q9QWR8 | Naga | Alpha-N-acetylgalactosaminidase | -0.023212465 | -0.052299658 |
| Q9QX60 | Dguok | Deoxyguanosine kinase, mitochondrial | 0.418687153 | 1.00664409 |
| Q9QXB9 | Drg2 | Developmentally-regulated GTP-binding protein 2 | -0.191457431 | 0.104245822 |
| Q9QXC1 | Fetub | Fetuin-B | -0.080801964 | -0.516563972 |
| Q9QXD8 | Limd1 | LIM domain-containing protein 1 | -1.030058479 | -0.258509795 |
| Q9QXE7 | Tbl1x | F-box-like/WD repeat-containing protein TBL1X | -0.355267588 | -0.323601087 |
| Q9QXK3 | Copg2 | Coatomer subunit gamma-2 | 0.105205345 | 0.370618025 |
| Q9QXS1 | Plec | Plectin | -0.043397427 | 0.000507196 |
| Q9QXS6 | Dbn1 | Drebrin | 0.11012586 | 0.142439206 |
| Q9QXT0 | Cnpy2 | Protein canopy homolog 2 | 0.031528727 | 0.25289615 |
| Q9QXV0 | Pcsk1n | ProSAAS | 0.821096961 | 1.313718001 |
| Q9QXY6 | Ehd3 | EH domain-containing protein 3 | 0.174005795 | 0.404179732 |
| Q9QY23 | Pkp3 | Plakophilin-3 | -0.587987455 | -0.608235359 |
| Q9QYB1 | Clic4 | Chloride intracellular channel protein 4 | 0.00984513 | 0.140503407 |
| Q9QYB5 | Add3 | Gamma-adducin | 0.028942521 | 0.0491786 |
| Q9QYB8 | Add2 | Beta-adducin | 0.087429301 | 0.213655949 |
| Q9QYC0 | Add1 | Alpha-adducin | -0.013973236 | -0.054872831 |
| Q9QYF9 | Ndrg3 | Protein NDRG3 | -0.272017097 | -0.472156207 |
| Q9QYG0 | Ndrg2 | Protein NDRG2 | -0.057124647 | 0.398996989 |
| Q9QYH6 | Maged1 | Melanoma-associated antigen D1 | -0.082194646 | 0.582821369 |
| Q9QYI5 | Dnajb2 | DnaJ homolog subfamily B member 2 | 0.17914896 | 0.391619205 |
| Q9QYJ0 | Dnaja2 | DnaJ homolog subfamily A member 2 | 0.207793935 | 0.386370341 |
| Q9QYJ3 | Dnajb1 | DnaJ homolog subfamily B member 1 | 0.195202128 | 0.371157964 |
| Q9QYK9 | Pnck | Calcium/calmodulin-dependent protein kinase type 1B | -0.224699084 | 2.142407209 |
| Q9QYP6 | Azi2 | 5-azacytidine-induced protein 2 | -1.363344701 | -0.702035904 |
| Q9QYR9 | Acot2 | Acyl-coenzyme A thioesterase 2, mitochondrial | -0.253022321 | -0.469224771 |
| Q9QYS9 | Qki | Protein quaking | 0.120967865 | -0.548602422 |
| Q9QYX7 | Pclo | Protein piccolo | 0.066287772 | 0.140352726 |
| Q9QZ06 | Tollip | Toll-interacting protein | 0.11044213 | 0.66782093 |
| Q9QZ08 | Nagk | N-acetyl-D-glucosamine kinase | -0.281176408 | 0.632123947 |

| | | | | |
|---|---|---|---|---|
| Q9QZ73 | Dcun1d1 | DCN1-like protein 1 | -0.136038494 | 0.915631612 |
| Q9QZA0 | Ca5b | Carbonic anhydrase 5B, mitochondrial | -0.347280248 | 0.135289749 |
| Q9QZB7 | Actr10 | Actin-related protein 10 | -0.666127364 | -0.966959318 |
| Q9QZB9 | Dctn5 | Dynactin subunit 5 | 0.658474795 | -0.009749095 |
| Q9QZD9 | Eif3i | Eukaryotic translation initiation factor 3 subunit I | 0.06793677 | 0.187952677 |
| Q9QZE5 | Copg1 | Coatomer subunit gamma-1 | -0.203674571 | 0.289383888 |
| Q9QZE7 | Tsnax | Translin-associated protein X | -0.202720515 | 0.872801304 |
| Q9QZH3 | Ppie | Peptidyl-prolyl cis-trans isomerase E | -0.33622268 | -0.465042909 |
| Q9QZM0 | Ubqln2 | Ubiquilin-2 | 0.330653286 | 0.748523394 |
| Q9QZQ8 | Macroh2a1 | Core histone macro-H2A.1 | 0.702847481 | 0.228655656 |
| Q9QZX7 | Srr | Serine racemase | 0.152975178 | 0.435303688 |
| Q9R013 | Ctsf | Cathepsin F | 0.235879993 | 0.015516917 |
| Q9R060 | Nubp1 | Cytosolic Fe-S cluster assembly factor NUBP1 | -0.157013385 | 0.191035112 |
| Q9R062 | Gyg1 | Glycogenin-1 | -0.322547181 | 0.874169827 |
| Q9R069 | Bcam | Basal cell adhesion molecule | -0.093005657 | 0.451111158 |
| Q9R0D8 | Wdr54 | WD repeat-containing protein 54 | 0.000596174 | -0.463736216 |
| Q9R0H5 | Krt71 | Keratin, type II cytoskeletal 71 | 0.264483388 | 0.378551801 |
| Q9R0N0 | Galk1 | Galactokinase | 0.081622632 | 0.792551041 |
| Q9R0P4 | Smap | Small acidic protein | 0.265786457 | -0.313777288 |
| Q9R0P5 | Dstn | Destrin | -0.131034851 | 0.65547657 |
| Q9R0P9 | Uchl1 | Ubiquitin carboxyl-terminal hydrolase isozyme L1 | -0.383563105 | -0.007361412 |
| Q9R0Q6 | Arpc1a | Actin-related protein 2/3 complex subunit 1A | 0.198253568 | 0.155891418 |
| Q9R0Q7 | Ptges3 | Prostaglandin E synthase 3 | -0.0719649 | 0.431777318 |
| Q9R0X4 | Acot9 | Acyl-coenzyme A thioesterase 9, mitochondrial | -0.326701196 | 0.024362564 |
| Q9R0Y5 | Ak1 | Adenylate kinase isoenzyme 1 | -0.090215556 | 0.64356041 |
| Q9R111 | Gda | Guanine deaminase | 0.125234222 | 0.289913813 |
| Q9R171 | Cbln1 | Cerebellin-1 | -0.499575806 | 0.222829024 |
| Q9R190 | Mta2 | Metastasis-associated protein MTA2 | -0.556608327 | -1.144588629 |
| Q9R1E6 | Enpp2 | Ectonucleotide pyrophosphatase/phosphodiesterase family member 2 | -0.176805814 | -0.43230025 |
| Q9R1K9 | Cetn2 | Centrin-2 | 0.432979266 | 0.581146558 |
| Q9R1L5 | Mast1 | Microtubule-associated serine/threonine-protein kinase 1 | -0.117659569 | 0.360205491 |
| Q9R1P0 | Psma4 | Proteasome subunit alpha type-4 | 0.166354275 | 0.275808493 |
| Q9R1P1 | Psmb3 | Proteasome subunit beta type-3 | 0.102499262 | 0.540176074 |
| Q9R1P3 | Psmb2 | Proteasome subunit beta type-2 | 0.275844828 | -0.327233791 |
| Q9R1P4 | Psma1 | Proteasome subunit alpha type-1 | -0.109064356 | -0.005343437 |
| Q9R1Q8 | Tagln3 | Transgelin-3 | 0.116392072 | 0.323883692 |
| Q9R1T2 | Sae1 | SUMO-activating enzyme subunit 1 | -0.045749124 | 0.567392349 |
| Q9R1T4 | Septin6 | Septin-6 | 0.033084774 | 0.195815245 |
| Q9R1Z7 | Pts | 6-pyruvoyl tetrahydrobiopterin synthase | -0.058268388 | -0.178336302 |
| Q9WTK7 | Stk11 | Serine/threonine-protein kinase STK11 | -0.003016249 | 0.295464357 |
| Q9WTL4 | Insrr | Insulin receptor-related protein | 0.033352343 | -1.203959147 |
| Q9WTL7 | Lypla2 | Acyl-protein thioesterase 2 | -0.212176005 | 0.4801507 |
| Q9WTM5 | Ruvbl2 | RuvB-like 2 | 0.091560809 | 0.49754556 |
| Q9WTN0 | Ggps1 | Geranylgeranyl pyrophosphate synthase | 0.157161681 | -0.00158151 |
| Q9WTP6 | Ak2 | Adenylate kinase 2, mitochondrial | -0.002410825 | 0.850938002 |

| | | | | |
|---|---|---|---|---|
| Q9WTP7 | Ak3 | GTP:AMP phosphotransferase AK3, mitochondrial | -0.021206188 | 0.309358279 |
| Q9WTR5 | Cdh13 | Cadherin-13 | 0.545769882 | 0.499493281 |
| Q9WTT4 | Atp6v1g2 | V-type proton ATPase subunit G 2 | -0.646126842 | 0.450149695 |
| Q9WTU6 | Mapk9 | Mitogen-activated protein kinase 9 | 0.204345036 | -0.558873812 |
| Q9WTX5 | Skp1 | S-phase kinase-associated protein 1 | -0.210536003 | -0.205556393 |
| Q9WTX6 | Cul1 | Cullin-1 | -0.206349754 | 0.063896656 |
| Q9WTX8 | Mad1l1 | Mitotic spindle assembly checkpoint protein MAD1 | -0.239301936 | -0.422130664 |
| Q9WU22 | Ptpn4 | Tyrosine-protein phosphatase non-receptor type 4 | -0.585439173 | -0.02038463 |
| Q9WU63 | Hebp2 | Heme-binding protein 2 | 0.05216287 | 0.388056755 |
| Q9WU78 | Pdcd6ip | Programmed cell death 6-interacting protein | 0.056596025 | 0.292442163 |
| Q9WUA2 | Farsb | Phenylalanine--tRNA ligase beta subunit | 0.362950198 | 0.189582666 |
| Q9WUA6 | Akt3 | RAC-gamma serine/threonine-protein kinase | -0.051398373 | 0.519808769 |
| Q9WUB0 | Rbck1 | RanBP-type and C3HC4-type zinc finger-containing protein 1 | 0.329178015 | 1.16882213 |
| Q9WUB3 | Pygm | Glycogen phosphorylase, muscle form | -0.390962728 | -0.007173538 |
| Q9WUD1 | Stub1 | E3 ubiquitin-protein ligase CHIP | -0.397202079 | 0.210207462 |
| Q9WUK2 | Eif4h | Eukaryotic translation initiation factor 4H | -0.041253344 | 0.289371014 |
| Q9WUM3 | Coro1b | Coronin-1B | -0.077812862 | 0.048947175 |
| Q9WUM4 | Coro1c | Coronin-1C | -0.202528572 | -0.206147671 |
| Q9WUM5 | Suclg1 | Succinate--CoA ligase [ADP/GDP-forming] subunit alpha, mitochondrial | -0.372786649 | 0.267484347 |
| Q9WUN2 | Tbk1 | Serine/threonine-protein kinase TBK1 | 0.387587674 | -0.467565219 |
| Q9WUP7 | Uchl5 | Ubiquitin carboxyl-terminal hydrolase isozyme L5 | 0.139389515 | 0.218953133 |
| Q9WUR9 | Ak4 | Adenylate kinase 4, mitochondrial | -0.221575292 | 0.358460585 |
| Q9WUT3 | Rps6ka2 | Ribosomal protein S6 kinase alpha-2 | 0.039433225 | 0.324471951 |
| Q9WUU7 | Ctsz | Cathepsin Z | -0.129587905 | 0.469438871 |
| Q9WV03 | Fam50a | Protein FAM50A | -1.779832681 | 0.531392415 |
| Q9WV34 | Mpp2 | MAGUK p55 subfamily member 2 | -0.129451529 | 0.300715923 |
| Q9WV54 | Asah1 | Acid ceramidase | 0.107243474 | 0.437470436 |
| Q9WV55 | Vapa | Vesicle-associated membrane protein-associated protein A | 0.006238111 | 0.300566832 |
| Q9WV69 | Dmtn | Dematin | 0.115431023 | 0.182893912 |
| Q9WV85 | Nme3 | Nucleoside diphosphate kinase 3 | -0.247980372 | -0.660310268 |
| Q9WVA4 | Tagln2 | Transgelin-2 | 0.28787082 | 0.684575558 |
| Q9WVE8 | Pacsin2 | Protein kinase C and casein kinase substrate in neurons protein 2 | 0.11091493 | 0.54792436 |
| Q9WVF8 | Tusc2 | Tumor suppressor candidate 2 | 0.794257323 | 0.213198821 |
| Q9WVJ2 | Psmd13 | 26S proteasome non-ATPase regulatory subunit 13 | 0.03730011 | 0.044627666 |
| Q9WVK4 | Ehd1 | EH domain-containing protein 1 | -0.060761229 | 0.280759494 |
| Q9WVQ5 | Apip | Methylthioribulose-1-phosphate dehydratase | -0.637831624 | -0.352098147 |
| Q9WVS7 | Map2k5 | Dual specificity mitogen-activated protein kinase kinase 5 | 1.217860349 | -0.660912832 |
| Q9Z0E0 | Ncdn | Neurochondrin | -0.056465848 | 0.237573624 |
| Q9Z0E6 | Gbp2 | Guanylate-binding protein 2 | 2.101806935 | 1.358904998 |
| Q9Z0F7 | Sncg | Gamma-synuclein | -0.280657641 | 0.932051341 |
| Q9Z0G0 | Gipc1 | PDZ domain-containing protein GIPC1 | 0.030072212 | 0.366030216 |
| Q9Z0H8 | Clip2 | CAP-Gly domain-containing linker protein 2 | 0.483325259 | 0.240335782 |
| Q9Z0J0 | Npc2 | NPC intracellular cholesterol transporter 2 | -0.035866229 | 0.442562262 |
| Q9Z0M5 | Lipa | Lysosomal acid lipase/cholesteryl ester hydrolase | 0.007903035 | -0.032187144 |
| Q9Z0N1 | Eif2s3x | Eukaryotic translation initiation factor 2 subunit 3, X-linked | -0.188408184 | 0.226551215 |

| | | | | |
|---|---|---|---|---|
| Q9Z0N2 | Eif2s3y | Eukaryotic translation initiation factor 2 subunit 3, Y-linked | -0.339260165 | 0.461838563 |
| Q9Z0P4 | Palm | Paralemmin-1 | 0.172211329 | 0.210491339 |
| Q9Z0P5 | Twf2 | Twinfilin-2 | 0.013264592 | 0.1872859 |
| Q9Z0Y1 | Dctn3 | Dynactin subunit 3 | 0.340049394 | 0.662602901 |
| Q9Z110 | Aldh18a1 | Delta-1-pyrroline-5-carboxylate synthase | -0.340940158 | 0.195933342 |
| Q9Z140 | Cpne6 | Copine-6 | 0.033953921 | 0.089443684 |
| Q9Z1A1 | Tfg | TFG protein | -0.784008185 | 0.944347541 |
| Q9Z1B3 | Plcb1 | 1-phosphatidylinositol 4,5-bisphosphate phosphodiesterase beta-1 | -0.028358523 | -0.020558675 |
| Q9Z1D1 | Eif3g | Eukaryotic translation initiation factor 3 subunit G | 0.02277441 | -0.624872526 |
| Q9Z1F9 | Uba2 | SUMO-activating enzyme subunit 2 | -0.062684282 | 0.130392551 |
| Q9Z1G3 | Atp6v1c1 | V-type proton ATPase subunit C 1 | 0.027619712 | -0.079423428 |
| Q9Z1J3 | Nfs1 | Cysteine desulfurase, mitochondrial | -0.061322816 | 0.287339052 |
| Q9Z1K5 | Arih1 | E3 ubiquitin-protein ligase ARIH1 | -0.061306381 | -0.110074997 |
| Q9Z1K6 | Arih2 | E3 ubiquitin-protein ligase ARIH2 | 0.225560443 | 0.83944122 |
| Q9Z1N5 | Ddx39b | Spliceosome RNA helicase Ddx39b | -0.193946457 | 0.451323032 |
| Q9Z1P6 | Ndufa7 | NADH dehydrogenase [ubiquinone] 1 alpha subcomplex subunit 7 | 0.414545504 | 0.941530704 |
| Q9Z1Q5 | Clic1 | Chloride intracellular channel protein 1 | -0.025916513 | 0.532421112 |
| Q9Z1Q9 | Vars1 | Valine--tRNA ligase | -0.301335462 | -0.294450601 |
| Q9Z1T1 | Ap3b1 | AP-3 complex subunit beta-1 | -0.743868192 | 0.833872954 |
| Q9Z1W9 | Stk39 | STE20/SPS1-related proline-alanine-rich protein kinase | 0.023984845 | -0.188644568 |
| Q9Z1Z0 | Uso1 | General vesicular transport factor p115 | 0.005657291 | 0.286012173 |
| Q9Z1Z2 | Strap | Serine-threonine kinase receptor-associated protein | -0.164052836 | -0.060979525 |
| Q9Z204 | Hnrnpc | Heterogeneous nuclear ribonucleoproteins C1/C2 | 0.315890153 | 0.139149666 |
| Q9Z247 | Fkbp9 | Peptidyl-prolyl cis-trans isomerase FKBP9 | 1.32240928 | 0.173908234 |
| Q9Z268 | Rasal1 | RasGAP-activating-like protein 1 | 0.126753139 | 0.392324448 |
| Q9Z275 | Rlbp1 | Retinaldehyde-binding protein 1 | -0.11244688 | 0.579019547 |
| Q9Z2C9 | Mtmr7 | Myotubularin-related protein 7 | -0.217960008 | 0.065370242 |
| Q9Z2D0 | Mtmr9 | Myotubularin-related protein 9 | -0.012821293 | 0.246071339 |
| Q9Z2D1 | Mtmr2 | Myotubularin-related protein 2 | -0.158534336 | -0.43541654 |
| Q9Z2D3 | Gsdme | Gasdermin-E | -0.695371914 | 0.88467137 |
| Q9Z2D6 | Mecp2 | Methyl-CpG-binding protein 2 | -0.330589612 | 0.762502432 |
| Q9Z2E4 | Ppp1r17 | Protein phosphatase 1 regulatory subunit 17 | -0.678512351 | -1.181954543 |
| Q9Z2I2 | Fkbp1b | Peptidyl-prolyl cis-trans isomerase FKBP1B | 0.566219298 | 0.532694658 |
| Q9Z2I8 | Suclg2 | Succinate--CoA ligase [GDP-forming] subunit beta, mitochondrial | -0.292761485 | 0.260965665 |
| Q9Z2I9 | Sucla2 | Succinate--CoA ligase [ADP-forming] subunit beta, mitochondrial | -0.101134173 | 0.410435994 |
| Q9Z2K1 | Krt16 | Keratin, type I cytoskeletal 16 | -0.357904371 | -0.78757604 |
| Q9Z2L6 | Minpp1 | Multiple inositol polyphosphate phosphatase 1 | -1.237956429 | 5.216188272 |
| Q9Z2L7 | Crlf3 | Cytokine receptor-like factor 3 | 0.206773313 | 0.837538401 |
| Q9Z2M7 | Pmm2 | Phosphomannomutase 2 | -0.01315012 | 0.429888884 |
| Q9Z2N8 | Actl6a | Actin-like protein 6A | -0.469277318 | 0.537790934 |
| Q9Z2Q6 | Septin5 | Septin-5 | 0.142554601 | 0.04691871 |
| Q9Z2T6 | Krt85 | Keratin, type II cuticular Hb5 | 0.769386832 | 0.86545976 |
| Q9Z2U0 | Psma7 | Proteasome subunit alpha type-7 | -0.025514221 | 0.602883021 |
| Q9Z2U1 | Psma5 | Proteasome subunit alpha type-5 | -0.049609598 | 0.440932433 |
| Q9Z2V5 | Hdac6 | Histone deacetylase 6 | 0.015191809 | 0.200625102 |

| | | | | |
|---|---|---|---|---|
| Q9Z2W1 | Stk25 | Serine/threonine-protein kinase 25 | -0.691721789 | -1.154807647 |
| Q9Z2X1 | Hnrnpf | Heterogeneous nuclear ribonucleoprotein F | -0.00416975 | 0.234418551 |
| Q9Z2Y3 | Homer1 | Homer protein homolog 1 | 0.375238736 | 0.257457733 |
| Q9Z315 | Sart1 | U4/U6.U5 tri-snRNP-associated protein 1 | 0.137492593 | 0.962741693 |
| Q9Z320 | Krt27 | Keratin, type I cytoskeletal 27 | -0.243314457 | 0.329675674 |
| S0DHL8 | Carmil2 | Capping protein, Arp2/3 and myosin-I linker protein 2 | 0.026145236 | -0.097528617 |
| S4R1M4 | Ank2 | Ankyrin-2 (Fragment) | -0.604065895 | 0.822883129 |
| S4R1M9 | Osbpl10 | Oxysterol-binding protein-related protein 10 | -0.1689785 | 0.271247864 |
| S4R1P5 | Dst | Dystonin | -0.315546258 | 0.222805341 |
| S4R1W8 | Gapdh | Glyceraldehyde-3-phosphate dehydrogenase (Fragment) | -1.114828555 | -0.398109277 |
| S4R2F3 | Ank2 | Ankyrin-2 | -0.189231555 | 0.018699646 |
| S4R2K9 | Ank3 | Ankyrin-3 (Fragment) | 0.14554828 | 0.046623389 |
| S4R2R5 | Ank2 | Ankyrin-2 | -0.72490619 | -1.376724084 |
| S4R2T7 | Ank2 | Ankyrin-2 (Fragment) | -0.506365617 | -0.577418804 |
| V9GX43 | Celf2 | CUGBP Elav-like family member 2 (Fragment) | 0.068168036 | -0.350434462 |
| V9GXM1 | Arfgap1 | ADP-ribosylation factor GTPase-activating protein 1 | 0.210029666 | 0.347786585 |
