## Supplementary material for "Virally induced lipid droplets are a platform for innate immune signalling complexes": S Table 2

| Table S2: LD Proteins identified and their regulation in Astrocytes stimulated with dsRNA | | | | | |
|---|---|---|---|---|---|
| Protein Accessions | Gene Names | Protein Descriptions | LogFC (8hr vs control) | LogFC (24hr vs control) | Gene Ontology |
| Proteins upregulated following dsRNA stimulation (8hrs) | | | | | |
| O14879 | IFIT3 | Interferon-induced protein with tetratricopeptide repeats 3 | 5.013711929 | 7.726207097 | Immune response |
| P09913 | IFIT2 | Interferon-induced protein with tetratricopeptide repeats 2 | 3.993145625 | 6.81721433 | Immune response |
| P09914 | IFIT1 | Interferon-induced protein with tetratricopeptide repeats 1 | 5.066923777 | 7.688687007 | Immune response |
| P11388 | TOP2A | DNA topoisomerase 2-alpha | 2.312225978 | 0.237378438 | Cellular metabolism |
| P20591 | MX1 | Interferon-induced GTP-binding protein Mx1 | 6.233834585 | 5.814037959 | Immune response |
| Q5TEJ8 | THEMIS2 | Protein THEMIS2 | 2.014291763 | 2.219636281 | Other |
| Q8WXG1 | RSAD2 | Radical S-adenosyl methionine domain-containing protein 2 | 2.606515249 | 9.234076182 | Immune response |
| Q96RL7-4 | VPS13A | Vacuolar protein sorting-associated protein 13A | 2.331557592 | 4.188173294 | Transport |
| Q9BYX4 | IFIH1 | Interferon-induced helicase C domain-containing protein 1 | 4.037366231 | 3.102505366 | Immune response |
| Q9NZ01 | TECR | Very-long-chain enoyl-CoA reductase | 2.206687291 | 1.558286031 | Lipid metabolism |
| Proteins upregulated following dsRNA stimulation (24hrs) | | | | | |
| A0A087WZT2 | METTL7B | Methyltransferase-like protein 7B | 0.286787033 | 2.807020187 | Cellular metabolism |
| A0A0A0MR09 | PTPN9 | Protein-tyrosine-phosphatase | 0 | 4.445039113 | Cellular metabolism |
| P42224 | STAT1 | Signal transducer and activator of transcription 1-alpha/beta | 0.283876419 | 2.480937958 | Immune response |
| A0A7P0TA76 | ACSL3 | Long-chain-fatty-acid--CoA ligase 3 | 0.727120082 | 2.419090907 | Lipid metabolism |
| B4DR61 | SEC61A1 | Protein transport protein Sec61 subunit alpha isoform 1 | 0.3150177 | 2.195833206 | Transport |
| B5MDU6 | LDAH | UPF0554 protein C2orf43 | 0.702131907 | 2.41792806 | Other |
| P57088 | TMEM33 | Transmembrane protein 33 | 0.889170965 | 2.028036118 | Other |
| Q9H8H3 | METTL7A | Methyltransferase-like protein 7A | 1.52543513 | 2.3282547 | Cellular metabolism |
| H0YNE9 | RAB8B | Ras-related protein Rab-8B | 1.66991806 | 3.09115092 | Transport |
| J3KQL8 | APOL2 | Apolipoprotein L2 | -0.130949656 | 4.084885279 | Lipid metabolism |
| O14879 | IFIT3 | Interferon-induced protein with tetratricopeptide repeats 3 | 5.013711929 | 7.726207097 | Immune response |
| O14966 | RAB29 | Ras-related protein Rab-7L1 | 0.494392395 | 2.78060023 | Transport |
| O43399 | TPD52L2 | Tumor protein D54 | 1.119777679 | 3.217659632 | Other |
| O60488-2 | ACSL4 | Long-chain-fatty-acid--CoA ligase 4 | 0.586028417 | 2.232774099 | Lipid metabolism |
| O95067 | CCNB2 | G2/mitotic-specific cyclin-B2 | -1.155277888 | 2.62821579 | Other |
| O95786 | DDX58 | Probable ATP-dependent RNA helicase DDX58 | 0.822710037 | 5.621814092 | Immune response |
| P00387-2 | CYB5R3 | NADH-cytochrome b5 reductase 3 | 0.636646271 | 2.442036947 | Lipid metabolism |
| P04259 | KRT6B | Keratin, type II cytoskeletal 6B | 0 | 2.495785395 | Other |
| P05161 | ISG15 | Ubiquitin-like protein ISG15 | 1.584328334 | 4.487184525 | Immune response |
| P09543-2 | CNP | 2,3-cyclic-nucleotide 3-phosphodiesterase | 0.184504827 | 3.659049988 | Cellular metabolism |
| P09913 | IFIT2 | Interferon-induced protein with tetratricopeptide repeats 2 | 3.993145625 | 6.81721433 | Immune response |
| P09914 | IFIT1 | Interferon-induced protein with tetratricopeptide repeats 1 | 5.066923777 | 7.688687007 | Immune response |
| P11233 | RALA | Ras-related protein Ral-A | 0.525677999 | 2.387378693 | Transport |
| P14635 | CCNB1 | G2/mitotic-specific cyclin-B1 | 0.258325577 | 2.315791448 | Cellular metabolism |
| P19971 | TYMP | Thymidine phosphorylase | -0.638599396 | 2.089799881 | Cellular metabolism |
| P20338 | RAB4A | Ras-related protein Rab-4A | 0.757256826 | 2.411489487 | Transport |
| P20591 | MX1 | Interferon-induced GTP-binding protein Mx1 | 6.233834585 | 5.814037959 | Immune response |
| P48449 | LSS | Lanosterol synthase | 0.761606852 | 2.56626002 | Lipid metabolism |
| P51151 | RAB9A | Ras-related protein Rab-9A | 0.3901577 | 2.842123667 | Transport |
| P56937-3 | HSD17B7 | 3-keto-steroid reductase | 0.673898061 | 2.960888545 | Lipid metabolism |

| | | | | | |
|---|---|---|---|---|---|
| P61006 | RAB8A | Ras-related protein Rab-8A | 1.54510498 | 2.851423264 | Transport |
| P61026 | RAB10 | Ras-related protein Rab-10 | 0.375212987 | 2.573587418 | Transport |
| P62820 | RAB1A | Ras-related protein Rab-1A | -0.106302897 | 2.697032928 | Transport |
| P84095 | RHOG | Rho-related GTP-binding protein RhoG | 0.730822245 | 2.490133286 | Other |
| Q14534 | SQLE | Squalene monooxygenase | -0.059093475 | 2.273886363 | Lipid metabolism |
| Q15286 | RAB35 | Ras-related protein Rab-35 | -0.402750651 | 2.367927551 | Transport |
| Q15738 | NSDHL | Sterol-4-alpha-carboxylate 3-dehydrogenase, decarboxylating | 0.742466609 | 2.76497523 | Lipid metabolism |
| Q32NB8 | PGS1 | CDP-diacylglycerol--glycerol-3-phosphate 3-phosphatidyltransferase, mitochondrial | 0.582439423 | 2.526611964 | Lipid metabolism |
| Q5EBM0-2 | CMPK2 | UMP-CMP kinase 2, mitochondrial | 0 | 3.603342692 | Cellular metabolism |
| Q709C8-3 | VPS13C | Vacuolar protein sorting-associated protein 13C | 0.90010198 | 2.431894302 | Transport |
| Q7L5N7 | LPCAT2 | Lysophosphatidylcholine acyltransferase 2 | 0.432391485 | 2.743785222 | Lipid metabolism |
| Q86YS6 | RAB43 | Ras-related protein Rab-43 | -0.954863866 | 4.741502126 | Transport |
| Q8IUS5 | EPHX4 | Epoxide hydrolase 4 | 0.459910711 | 2.999158223 | Other |
| Q8N0X7 | SPG20 | Spartin | 0.08765475 | 2.009578705 | Other |
| Q8N5I4 | DHRSX | Dehydrogenase/reductase SDR family member on chromosome X | 0.88176473 | 2.563165029 | Other |
| Q8NF37 | LPCAT1 | Lysophosphatidylcholine acyltransferase 1 | -0.035873413 | 2.822474798 | Lipid metabolism |
| Q8WUH6 | TMEM263 | Transmembrane protein 263 | 3.245290756 | 4.188417435 | Other |
| Q8WXG1 | RSAD2 | Radical S-adenosyl methionine domain-containing protein 2 | 2.606515249 | 9.234076182 | Immune response |
| Q92575 | UBXN4 | UBX domain-containing protein 4 | 1.123182933 | 2.762897491 | Cellular metabolism |
| Q96CS3 | FAF2 | FAS-associated factor 2 | 0.444726944 | 2.882000605 | Other |
| Q96LJ7 | DHRS1 | Dehydrogenase/reductase SDR family member 1 | 0.795609792 | 2.224393845 | Other |
| Q9BRQ8 | AIFM2 | Apoptosis-inducing factor 2 | 0.505276998 | 2.524138133 | Other |
| Q9NP72 | RAB18 | Ras-related protein Rab-18 | 0.409819921 | 2.415627797 | Transport |
| Q9UG22 | GIMAP2 | GTPase IMAP family member 2 | 0 | 5.225255966 | Other |
| Q9UL25 | RAB21 | Ras-related protein Rab-21 | -0.173776627 | 3.232548396 | Transport |
| Q9Y679 | AUP1 | Ancient ubiquitous protein 1 | 0.812328974 | 2.536839803 | Other |
| **Proteins downregulated following dsRNA stimulation (8hrs)** | | | | | |
| C9JIZ6 | PSAP | Prosaposin | -4.592389425 | -1.114802043 | Lipid metabolism |
| F8WEC0 | MMADHC | Methylmalonic aciduria and homocystinuria type D protein, mitochondrial | -3.642526627 | -1.106088003 | Cellular metabolism |
| O43164-2 | PJA2 | E3 ubiquitin-protein ligase Praja-2 | -2.62583669 | 0 | Cellular metabolism |
| Q9P2I0 | CPSF2 | Cleavage and polyadenylation specificity factor subunit 2 | -2.977603277 | -2.727367401 | Transport |
| **Proteins downregulated following dsRNA stimulation (24hrs)** | | | | | |
| A0A0A6YYL4 | CORO7-PAM16 | Coronin | -0.315741221 | -2.159619013 | Transport |
| Q12959-2 | DLG1 | Disks large homolog 1 | 0.79543368 | -2.682020187 | Immune response |
| A0A7P0TAV2 | MTR | Methionine synthase | -0.538960139 | -2.494082133 | Cellular metabolism |
| Q92995 | USP13 | Ubiquitin carboxyl-terminal hydrolase 13 | -0.179007848 | -2.155618032 | Cellular metabolism |
| C9IYN7 | ELP6 | Elongator complex protein 6 | -0.114273071 | -2.086010615 | Cellular metabolism |
| H0YC67 | RMDN1 | Regulator of microtubule dynamics protein 1 | -0.186487834 | -2.665707906 | Other |

| | | | | | |
|---|---|---|---|---|---|
| O43633 | CHMP2A | Charged multivesicular body protein 2a | -0.059028625 | -3.808405558 | Transport |
| P02751-17 | FN1 | Fibronectin | -0.797923406 | -2.031629562 | Transport |
| P04818 | TYMS | Thymidylate synthase | -0.802684148 | -2.216362 | Cellular metabolism |
| P13796 | LCP1 | Plastin-2 | -0.177000682 | -2.621720632 | Other |
| P48307-2 | TFPI2 | Tissue factor pathway inhibitor 2 | -0.222955068 | -3.417169571 | Other |
| P49356 | FNTB | Protein farnesyltransferase subunit beta | -0.164211273 | -2.410591125 | Cellular metabolism |
| P50748 | KNTC1 | Kinetochore-associated protein 1 | -0.040159225 | -2.652329763 | Other |
| P51687 | SUOX | Sulfite oxidase, mitochondrial | -1.483929952 | -2.393257141 | Cellular metabolism |
| Q12841 | FSTL1 | Follistatin-related protein 1 | -1.952020009 | -2.032182058 | Other |
| Q12972 | PPP1R8 | Nuclear inhibitor of protein phosphatase 1 | -0.114404678 | -2.955382029 | Cellular metabolism |
| Q13509 | TUBB3 | Tubulin beta-3 chain | -0.180454254 | -3.764647166 | Cellular metabolism |
| Q14376 | GALE | UDP-glucose 4-epimerase | -0.172948837 | -4.287476858 | Cellular metabolism |
| Q3LXA3 | DAK | Bifunctional ATP-dependent dihydroxyacetone kinase/FAD-AMP lyase (cyclizing) | -0.039713542 | -2.172501882 | Immune response |
| Q5T447 | HECTD3 | E3 ubiquitin-protein ligase HECTD3 | -0.072162628 | -2.248969396 | Cellular metabolism |
| Q5TAQ9 | DCAF8 | DDB1- and CUL4-associated factor 8 | -0.95735995 | -2.179983775 | Cellular metabolism |
| Q6GQQ9-2 | OTUD7B | OTU domain-containing protein 7B | 0.044070562 | -2.68447876 | Immune response |
| Q6WCQ1 | MPRIP | Myosin phosphatase Rho-interacting protein | 0.547382355 | -4.214472453 | Other |
| Q7L2J0 | MEPCE | 7SK snRNA methylphosphate capping enzyme | -0.409100215 | -5.175966263 | Cellular metabolism |
| Q8IV38 | ANKMY2 | Ankyrin repeat and MYND domain-containing protein 2 | -0.069720586 | -2.612548828 | Other |
| Q969T7-2 | NT5C3B | 7-methylguanosine phosphate-specific 5-nucleotidase | -0.059551875 | -2.198008219 | Cellular metabolism |
| Q96BH1 | RNF25 | E3 ubiquitin-protein ligase RNF25 | -0.05690066 | -2.953262329 | Cellular metabolism |
| Q99504-5 | EYA3 | Eyes absent homolog 3 | -0.143803279 | -2.780726751 | Cellular metabolism |
| Q99538-3 | LGMN | Legumain | -0.624713898 | -3.459128698 | Immune response |
| Q9BZL1 | UBL5 | Ubiquitin-like protein 5 | -0.665559133 | -4.066532771 | Cellular metabolism |
| Q9HBM1 | SPC25 | Kinetochore protein Spc25 | 0.146274567 | -2.183904648 | Other |
| Q9NRF8 | CTPS2 | CTP synthase 2 | 0.081635157 | -2.513303121 | Cellular metabolism |
| Q9NYL2-2 | ZAK | Mitogen-activated protein kinase kinase kinase MLT | -0.340040207 | -3.511322657 | Cellular metabolism |
| Q9P1Z2-2 | CALCOCO1 | Calcium-binding and coiled-coil domain-containing protein 1 | -0.535254796 | -4.825668971 | Cellular metabolism |
| Q9P287 | BCCIP | BRCA2 and CDKN1A-interacting protein | -0.273838043 | -2.517972946 | Cellular metabolism |
| Q9UP83-3 | COG5 | Conserved oligomeric Golgi complex subunit 5 | -0.465450287 | -2.356347402 | Transport |
| **Proteins present/unchanged** | | | | | |
| A0A024R4E5 | HDLBP | Vigilin | 0.17566363 | -0.257462819 | Lipid metabolism |

| | | | | | |
|---|---|---|---|---|---|
| A0A024RA52 | PSMA2 | Proteasome subunit alpha type | 0.140261332 | 0.464188894 | Immune response |
| A0A024RBG1 | NUDT4 | Diphosphoinositol polyphosphate phosphohydrolase 2 | -0.35189565 | -0.606040955 | Transport |
| A0A024RBT2 | DIABLO | Diablo homolog, mitochondrial | 0.326915105 | -2.361492157 | Other |
| G8JLL2 | PPT2 | Lysosomal thioesterase PPT2 | -0.260935465 | 0 | Other |
| P46379-3 | BAG6 | Large proline-rich protein BAG6 | -0.268374125 | -1.860606511 | Transport |
| A0A075B730 | EPPK1 | Epiplakin | 0.289014816 | 0.583296458 | Other |
| A0A075B749 | PDE4DIP | Myomegalin | -0.783456167 | -0.413244883 | Other |
| H0YL34 | SYNM | Synemin | 0.926755269 | 0 | Other |
| A0A075B7B5 | SRGAP2 | SLIT-ROBO Rho GTPase-activating protein 2 | -0.02919515 | 0 | Other |
| Q9NW36 | HELLS | Lymphoid-specific helicase | -0.424547831 | 0 | Cellular metabolism |
| Q16881 | TXNRD1 | Thioredoxin reductase 1, cytoplasmic | -0.044615428 | -0.140113195 | Lipid metabolism |
| K7EKX7 | GPX4 | Glutathione peroxidase | -0.150527318 | 0.188684464 | Lipid metabolism |
| A0A087WVR3 | UHRF1 | E3 ubiquitin-protein ligase UHRF1 | 0.834142049 | -0.320851008 | Cellular metabolism |
| A0A087WU53 | MAGT1 | Magnesium transporter protein 1 | 0.776774089 | 0 | Transport |
| A0A087WUB9 | CTNNBL1 | Beta-catenin-like protein 1 | 0.01120313 | -0.717511495 | Cellular metabolism |
| A0A087WUC6 | SPCS2 | Signal peptidase complex subunit 2 | 1.223227819 | -0.255293528 | Transport |
| M0R3C7 | SYMPK | Symplekin | -0.031393687 | 0 | Cellular metabolism |
| A0A2P0CT92 | CNTNAP3 | Contactin-associated protein-like 3B | -0.522785823 | -1.587509155 | Other |
| A0A087WUT6 | EIF5B | Eukaryotic translation initiation factor 5B | 0.031629562 | -0.598887126 | Cellular metabolism |
| A0A087WV25 | FGFR1OP | FGFR1 oncogene partner | 0.267000198 | -1.723405838 | Cellular metabolism |
| A0A0A0MT31 | PRH1 | Salivary acidic proline-rich phosphoprotein 1/2 | 0 | 0.819546382 | Other |
| A0A087WV97 | SMAP2 | Stromal membrane-associated protein 2 | -0.394804637 | 0 | Other |
| A0A087WVM4 | MTHFD1L | Monofunctional C1-tetrahydrofolate synthase, mitochondrial | 0 | 0.65195783 | Cellular metabolism |
| A0A087WVM7 | EMG1 | Ribosomal RNA small subunit methyltransferase NEP1 | 0.135724386 | 0 | Cellular metabolism |
| A0A087WVQ6 | CLTC | Clathrin heavy chain | -0.093022029 | -0.554243724 | Transport |
| A0A087WVS8 | MOB2 | MOB kinase activator 2 | -0.24114418 | -0.789047877 | Other |
| A0A087WW06 | TTC28 | Tetratricopeptide repeat protein 28 | 0.107726415 | 0 | Other |
| A0A087WW40 | SH3GLB1 | Endophilin-B1 | -0.287664413 | 1.157193502 | Lipid metabolism |
| A0A087WW88 | CHMP2B | Charged multivesicular body protein 2b | -0.371014913 | 1.09527715 | Transport |
| O60333-4 | KIF1B | Kinesin-like protein KIF1B | -0.160410563 | 0 | Transport |
| A0A087WWF6 | POLD2 | DNA polymerase delta subunit 2 | -0.255033493 | 0.052497864 | Cellular metabolism |
| Q9P2T1-2 | GMPR2 | GMP reductase 2 | -0.080628713 | 0.300285339 | Cellular metabolism |
| A0A087WXM6 | RPL17 | 60S ribosomal protein L17 | 0.265585581 | -0.069429398 | Transport |
| A0A499FIX8 | ESYT2 | Extended synaptotagmin-2 | 0.832958857 | 1.470630646 | Transport |

| | | | | | |
|---|---|---|---|---|---|
| A0A087WY55 | VTA1 | Vacuolar protein sorting-associated protein VTA1 homolog | 0.091230392 | -1.921020508 | Transport |
| A0A087WY71 | AP2M1 | AP-2 complex subunit mu | -0.230031331 | -1.447505951 | Transport |
| P61077-3 | UBE2D3 | Ubiquitin-conjugating enzyme E2 D3 | 0.016670227 | 0.307216644 | Immune response |
| A0A087X0V7 | APBB1 | Amyloid beta A4 precursor protein-binding family B member 1 | -1.210888545 | 0 | Cellular metabolism |
| A0A087WYN9 | DHX29 | ATP-dependent RNA helicase DHX29 | 0.901756287 | 0 | Cellular metabolism |
| A0A087WYU2 | C16orf13 | UPF0585 protein C16orf13 | -0.076285044 | -0.223099391 | Other |
| A0A087WYW3 | NKTR | NK-tumor recognition protein | 0.567507426 | -1.246109009 | Cellular metabolism |
| A0A087WZ13 | RAVER1 | Ribonucleoprotein PTB-binding 1 | 1.08462588 | 0.939468384 | Other |
| A0A087WZK9 | EIF3H | Eukaryotic translation initiation factor 3 subunit H | 0.19237264 | -0.046331406 | Cellular metabolism |
| A0A087WZL0 | PHLDB1 | Pleckstrin homology-like domain family B member 1 | -1.242982864 | 0 | Other |
| O43657 | TSPAN6 | Tetraspanin-6 | 0 | -0.417665482 | Other |
| A0A087WZV0 | PPIP5K2 | Inositol hexakisphosphate and diphosphoinositol-pentakisphosphate kinase 2 | -0.315049489 | 0 | Cellular metabolism |
| J3KPF9 | KIF3A | Kinesin-like protein | 0.185307821 | -0.522301992 | Transport |
| Q9Y3A5 | SBDS | Ribosome maturation protein SBDS | -0.020877838 | -1.361593246 | Cellular metabolism |
| A0A087X038 | PLEKHA2 | Pleckstrin homology domain-containing family A member 2 | 0.239736557 | 0 | Other |
| A0A087X054 | HYOU1 | Hypoxia up-regulated protein 1 | 0.355869929 | -0.714649836 | Transport |
| A0A7I2V673 | CTSH | Pro-cathepsin H | -0.282212575 | 0.235755285 | Immune response |
| A0A7I2V5U5 | TJP1 | Tight junction protein ZO-1 | 0.137189865 | -0.605033239 | Other |
| A0A087X0M4 | SLC4A1AP | Kanadaptin | -0.273056666 | -1.919502894 | Other |
| A0A087X0Q1 | YTHDF3 | YTH domain-containing family protein 3 | 0.039080302 | 0.580335617 | Other |
| A0A087X0Q9 | SELO | Selenoprotein O | -0.193011602 | 0 | Other |
| A0A087X0R6 | SNX12 | Sorting nexin-12 | -0.087827682 | -0.90587616 | Transport |
| A0A087X0S5 | COL6A1 | Collagen alpha-1(VI) chain | -0.57183075 | -1.276809692 | Other |
| A0A087X0W7 | ACOT2 | Acyl-coenzyme A thioesterase 1 | 0.730566025 | -0.624527613 | Lipid metabolism |
| A0A087X142 | 39692 | Septin-8 | -0.088423411 | -1.211801529 | Other |
| F5H211 | ATXN3 | Ataxin-3 | -0.289236705 | -1.356632868 | Cellular metabolism |
| A0A087X1B2 | USP39 | U4/U6.U5 tri-snRNP-associated protein 2 | 0.388008753 | 0.156697591 | Cellular metabolism |
| A0A087X1B7 | CHTOP | Chromatin target of PRMT1 protein | 0 | -0.712792079 | Transport |
| A0A087X1E4 | ARFIP2 | Arfaptin-2 | 0.243080139 | 0.35362498 | Other |
| A0A7I2V2W4 | NDUFA5 | NADH dehydrogenase [ubiquinone] 1 alpha subcomplex subunit 5 | 0 | -0.337657928 | Cellular metabolism |
| O60613 | 42248 | 15 kDa selenoprotein | 0.713781993 | -2.336734136 | Cellular metabolism |
| A0A087X1Z3 | PSME2 | Proteasome activator complex subunit 2 | 0.116381327 | 0.599926631 | Immune response |
| A0A087X208 | AGRN | Agrin | -0.74582545 | -0.837837855 | Lipid metabolism |
| A0A087X211 | KIAA1524 | Protein CIP2A | -0.283532461 | -1.724119822 | Other |

| | | | | | |
|---|---|---|---|---|---|
| A0A087X256 | KIAA1033 | WASH complex subunit 7 | -0.247193019 | -1.267491023 | Transport |
| A0A087X2I1 | PSMC6 | 26S protease regulatory subunit 10B | 0.270914078 | -0.283428192 | Immune response |
| A0A087X2I5 | LSM7 | U6 snRNA-associated Sm-like protein LSm7 | 0.184853236 | -0.61536471 | Cellular metabolism |
| A0A096LNH5 | C21orf33 | | -0.095877965 | -0.585121791 | Other |
| P30566 | ADSL | Adenylosuccinate lyase | 0.004747391 | -0.086779912 | Cellular metabolism |
| A0A096LP25 | AAK1 | AP2-associated protein kinase 1 | 0.273111979 | -1.191093445 | Transport |
| A0A0A0MR88 | FAM21C | WASH complex subunit FAM21C | -0.622633616 | 0 | Transport |
| A0A0A0MQS1 | PYCRL | Pyrroline-5-carboxylate reductase | -0.312360764 | -1.702226003 | Cellular metabolism |
| F5H0Q6 | CLASRP | CLK4-associating serine/arginine rich protein | 0.436852773 | -0.71182696 | Cellular metabolism |
| U3KQC1 | WDR18 | WD repeat-containing protein 18 | 0.083363851 | 0 | Other |
| A0A0A0MQU4 | SCLY | Selenocysteine lyase | 0.279101054 | -0.686572393 | Lipid metabolism |
| A0A0A0MQV6 | FGF2 | Fibroblast growth factor 2 | -0.087164561 | -0.440116247 | Immune response |
| Q9NR56-6 | MBNL1 | Muscleblind-like protein 1 | -0.339380264 | -0.853663762 | Cellular metabolism |
| A0A0A0MR02 | VDAC2 | Voltage-dependent anion-selective channel protein 2 | 0.455715815 | 0.906815211 | Transport |
| Q5T092 | RER1 | Protein RER1 | 0.457357407 | 0 | Transport |
| A0A0A0MR85 | GSTM4 | Glutathione S-transferase Mu 4 | -0.031924566 | 0 | Cellular metabolism |
| A0A0A0MRA5 | HNRNPUL1 | Heterogeneous nuclear ribonucleoprotein U-like protein 1 | -0.191356659 | -1.000818888 | Cellular metabolism |
| A0A0A0MRH0 | NLE1 | Notchless protein homolog 1 | 0.282892863 | -1.775761286 | Other |
| A0A0A0MRH6 | MIA3 | Melanoma inhibitory activity protein 3 | 0 | 1.0064195 | Transport |
| A0A0A0MRI1 | GPALPP1 | GPALPP motifs-containing protein 1 | 0.039002736 | -1.307305654 | Other |
| A0A0A0MRJ6 | PCMT1 | Protein-L-isoaspartate O-methyltransferase | -0.024304708 | -0.62987709 | Cellular metabolism |
| A0A0A0MRM8 | MYO6 | Unconventional myosin-VI | -0.236379623 | -1.614217122 | Transport |
| A0A0A0MRP6 | SMARCA1 | Probable global transcription activator SNF2L1 | -0.172509511 | -0.799551646 | Cellular metabolism |
| P09234 | SNRPC | U1 small nuclear ribonucleoprotein C | -1.468379974 | -1.800099691 | Cellular metabolism |
| Q8IZP0-10 | ABI1 | Abl interactor 1 | -0.783494314 | -2.038239797 | Immune response |
| J3QKK8 | FAM213B | Prostamide/prostaglandin F synthase | -0.075608571 | 0 | Lipid metabolism |
| A0A0A0MRX2 | DDRGK1 | DDRGK domain-containing protein 1 | 0 | 1.340094884 | Other |
| A0A0A0MRY0 | TBC1D22A | TBC1 domain family member 22A | -0.175102234 | 0 | Other |
| E7EMV7 | TNIP1 | TNFAIP3-interacting protein 1 | -0.162210464 | -0.804243088 | Cellular metabolism |
| A0A0A0MRZ8 | IGKV3D-11 | Ig kappa chain V-III region VG | 0.331816355 | 0 | Immune response |
| P01857 | IGHG1 | Ig gamma-1 chain C region | 0.048651377 | 0.768467585 | Immune response |
| A0A6I8PIQ6 | COG4 | Conserved oligomeric Golgi complex subunit 4 | -0.791505178 | -0.772883733 | Transport |
| A0A0A0MSA7 | EIF4G3 | Eukaryotic translation initiation factor 4 gamma 3 | -0.584050496 | 0.375454585 | Other |
| A0A0A0MSB8 | EXOC7 | Exocyst complex component 7 | -0.100008011 | 0.181709925 | Transport |

| | | | | | |
|---|---|---|---|---|---|
| A0A0A0MSE2 | HADH | Hydroxyacyl-coenzyme A dehydrogenase, mitochondrial | -1.197013219 | 1.776822408 | Lipid metabolism |
| A0A0A0MSI6 | FBXO44 | F-box only protein 44 | 0.135153453 | 0 | Other |
| F8W9B8 | EXOC5 | Exocyst complex component 5 | 0 | -1.427240372 | Transport |
| A0A0A0MSJ0 | DDX42 | ATP-dependent RNA helicase DDX42 | 0.389195124 | -0.796592712 | Cellular metabolism |
| A0A0A0MSJ9 | RASA2 | Ras GTPase-activating protein 2 | 0.305639903 | 0 | Other |
| A0A0A0MSK4 | GPSM1 | G-protein-signaling modulator 1 | -0.105750402 | 0 | Other |
| A0A0A0MSK5 | TOR1AIP1 | Torsin-1A-interacting protein 1 | 1.096115112 | 0.155959447 | Other |
| A0A0A0MSW4 | PITPNB | Phosphatidylinositol transfer protein beta isoform | -0.024097443 | -0.119901657 | Lipid metabolism |
| A0A0A0MT32 | LIPA | Lysosomal acid lipase/cholesteryl ester hydrolase | 0.421398163 | -0.649365107 | Lipid metabolism |
| H0YLC5 | POU2F3 | POU domain protein | -0.122962952 | 0 | Immune response |
| P51532-5 | SMARCA4 | Transcription activator BRG1 | 0.659378688 | 0 | Cellular metabolism |
| A0A0A0MTB8 | WDR36 | WD repeat-containing protein 36 | 0 | 1.284959157 | Cellular metabolism |
| A0A0A0MTB9 | RBP1 | Retinol-binding protein 1 | -0.230829875 | 0 | Lipid metabolism |
| A0A0A0MTH3 | ILK | Integrin-linked protein kinase | -0.338259379 | -0.737011592 | Cellular metabolism |
| A0A0A0MTH9 | BTAF1 | TATA-binding protein-associated factor 172 | -0.737513224 | 0 | Other |
| A0A0A0MTI1 | DHCR24 | Delta(24)-sterol reductase | 0 | 3.013046265 | Lipid metabolism |
| A0A0A0MTJ9 | NCEH1 | Neutral cholesterol ester hydrolase 1 | 0.933279673 | 1.122283936 | Lipid metabolism |
| A0A0A0MTL5 | SKP2 | S-phase kinase-associated protein 2 | -0.515377045 | 0 | Cellular metabolism |
| A0A0A0MTN0 | CUL2 | Cullin-2 | 0.065832138 | -0.215523402 | Cellular metabolism |
| A0A0A0MTS7 | TTN | Titin | -0.66456604 | -0.77299118 | Transport |
| A0A0A6YYA0 | TMED7-TICAM2 | Transmembrane emp24 domain-containing protein 7 | 1.320725123 | 0.662830989 | Immune response |
| A0A0A6YYH1 | C15orf38-AP3S2 | Arpin | -0.357488632 | -0.747125626 | Transport |
| A0A0B4J1S8 | PI4KB | Phosphatidylinositol 4-kinase beta | -0.575393041 | -0.605648041 | Lipid metabolism |
| F8VV32 | LYZ | Lysozyme | 1.212082545 | 1.118715922 | Other |
| A0A0B4J294 | TRAPPC2L | Trafficking protein particle complex subunit 2-like protein | -0.181483587 | 0 | Transport |
| A0A0B4J2A0 | TRPV1 | Sedoheptulokinase | 0.123121262 | 0.18552653 | Cellular metabolism |
| A0A0B4J2A4 | ACAA2 | 3-ketoacyl-CoA thiolase, mitochondrial | -0.194645564 | -0.188964844 | Lipid metabolism |
| G3V5T0 | GSTZ1 | Maleylacetoacetate isomerase | -0.269969304 | -1.336175283 | Cellular metabolism |
| A0A0C4DFN3 | MGLL | Monoglyceride lipase | 0.617515564 | 0.373486201 | Lipid metabolism |
| A0A0C4DFX9 | NELFA | Negative elongation factor A | -0.192709605 | -2.867907842 | Cellular metabolism |
| A0A0C4DFZ2 | ARSA | Arylsulfatase A | -2.605876287 | 0.502892812 | Lipid metabolism |
| C9J9K3 | RPSA | 40S ribosomal protein SA | 0.097067515 | -0.053600947 | Transport |
| F5GYV5 | ARL6IP4 | ADP-ribosylation factor-like protein 6-interacting protein 4 | -0.286317825 | -1.416952133 | Cellular metabolism |
| F1LLU7 | ECI2 | Enoyl-CoA delta isomerase 2, mitochondrial | -0.252585093 | 0 | Lipid metabolism |
| A0A0C4DGG1 | PACSIN3 | Protein kinase C and casein kinase substrate in neurons protein 3 | -0.116043091 | -1.763308207 | Transport |

| | | | | | |
|---|---|---|---|---|---|
| A0A2R8YFK9 | CHD4 | Chromodomain-helicase-DNA-binding protein 4 | 0.353003184 | 0.689417521 | Cellular metabolism |
| I3L1H9 | ZG16B | Zymogen granule protein 16 homolog B | 0 | 0.623362859 | Other |
| J3QRU8 | GIT1 | ARF GTPase-activating protein GIT1 | -0.448109945 | 0 | Other |
| A0A0C4DGQ5 | CAPNS1 | Calpain small subunit 1 | -0.058828354 | -0.585320791 | Other |
| Q6IAN0 | DHRS7B | Dehydrogenase/reductase SDR family member 7B | -0.264373144 | 1.048919042 | Other |
| A0A0C4DGS1 | DDOST | Dolichyl-diphosphooligosaccharide--protein glycosyltransferase 48 kDa subunit | 1.214260101 | 0.953163147 | Immune response |
| A0A0C4DGX8 | ATP6AP1 | V-type proton ATPase subunit S1 | 0 | 1.260363897 | Transport |
| A0A0C4DH22 | EPB41L1 | Band 4.1-like protein 1 | -0.569088618 | 0 | Other |
| P21127 | CDK11B | Cyclin-dependent kinase 11B | -0.009633382 | -0.359846751 | Immune response |
| B4DLR2 | FAP | Prolyl endopeptidase FAP | 0.572282791 | -1.166087468 | Other |
| A0A0D9SEU5 | RANBP10 | Ran-binding protein 10 | 0.280597687 | -0.873846054 | Other |
| O95819-6 | MAP4K4 | Mitogen-activated protein kinase kinase kinase kinase 4 | 0.4739151 | -1.408739726 | Cellular metabolism |
| A0A0D9SF44 | R3HCC1 | R3H and coiled-coil domain-containing protein 1 | -0.223321915 | 0 | Other |
| A0A2R8YFS5 | DDX3X | ATP-dependent RNA helicase DDX3X | 0.03198115 | 0.518883387 | Immune response |
| Q8N6H7 | ARFGAP2 | ADP-ribosylation factor GTPase-activating protein 2 | -0.044852575 | -0.279994965 | Transport |
| A0A0D9SFB1 | DNM1 | Dynamin-1 | 0.072064082 | 0 | Transport |
| A0A1B0GWF2 | STXBP1 | Syntaxin-binding protein 1 | 0.083897909 | -0.127459844 | Transport |
| A0A0D9SGE8 | PHF6 | PHD finger protein 6 | 0.406642278 | -0.009707769 | Cellular metabolism |
| A0A0G2JH68 | DIAPH1 | Protein diaphanous homolog 1 | -0.059940974 | -0.880095164 | Other |
| A0A0G2JHC2 | PPP1R18 | Phostensin | 0.022833506 | -0.047648748 | Other |
| A0A0G2JIW1 | HSPA1B | Heat shock 70 kDa protein 1B | -0.196090062 | -0.143778483 | Other |
| A2ACR1 | PSMB9 | Proteasome subunit beta type | 0.327791214 | 1.283911387 | Immune response |
| P25440-2 | BRD2 | Bromodomain-containing protein 2 | -0.808958054 | -0.801546733 | Cellular metabolism |
| A0A0G2JNZ5 | GBA | Glucosylceramidase | -0.960518519 | 0.033834457 | Lipid metabolism |
| A0A0G2JMS7 | SCRIB | Protein scribble homolog | -0.543348948 | 1.631413778 | Transport |
| Q12979 | ABR | Active breakpoint cluster region-related protein | 0.062973658 | -0.779768626 | Other |
| A0A0U1RQQ2 | SMARCB1 | SWI/SNF-related matrix-associated actin-dependent regulator of chromatin subfamily B member 1 | 0 | 0.158809662 | Cellular metabolism |
| A0A0J9YYL3 | PUF60 | Poly(U)-binding-splicing factor PUF60 | 0.146983465 | -0.344153086 | Cellular metabolism |
| A0A0J9YWB6 | CTDP1 | RNA polymerase II subunit A C-terminal domain phosphatase | 0.392026901 | 0.014222463 | Cellular metabolism |
| C9J2C4 | DNAJB6 | DnaJ homolog subfamily B member 6 | -0.394346237 | -0.23722585 | Cellular metabolism |
| P43243 | MATR3 | Matrin-3 | 0.457734426 | 0.259716034 | Other |
| A0A0R4J2F3 | | | 0.184206009 | 0 | #N/A |
| A0A0R4J2F6 | IFRD2 | Interferon-related developmental regulator 2 | 0.645559947 | 0 | Other |
| H0YKX3 | KIAA0101 | PCNA-associated factor | -1.322102229 | -2.076145172 | Cellular metabolism |

| | | | | | |
|---|---|---|---|---|---|
| Q9UNN8 | PROCR | Endothelial protein C receptor | -0.850609461 | -0.15460968 | Immune response |
| Q9Y484-2 | WDR45 | WD repeat domain phosphoinositide-interacting protein 4 | 0.226132075 | 1.071107229 | Cellular metabolism |
| A0A0U1RRM4 | PTBP1 | Polypyrimidine tract-binding protein 1 | 0.648769379 | 0.861574809 | Cellular metabolism |
| Q53Z42 | HLA-A | | 0.358809789 | 0.731901169 | Immune response |
| A0A140T9U8 | HLA-C | HLA class I histocompatibility antigen, Cw-7 alpha chain | -0.135563533 | 0.951289495 | Immune response |
| A0A140T9R1 | FLOT1 | Flotillin-1 | -0.40416654 | -0.114418666 | Transport |
| Q03518 | TAP1 | Antigen peptide transporter 1 | 0 | 2.522672653 | Transport |
| A0A1B0GTC3 | FTO | Alpha-ketoglutarate-dependent dioxygenase FTO | 0.009601593 | -0.570575078 | Cellular metabolism |
| A0A6I8PIT5 | HSPA12A | Heat shock 70 kDa protein 12A | -0.412260691 | -0.494978587 | Other |
| A0A1B0GV06 | ASAH1 | Acid ceramidase | -1.07905515 | 0.584311167 | Lipid metabolism |
| A0A1B0GTU4 | PXN | Paxillin | -0.154010773 | -1.377494176 | Cellular metabolism |
| A0A1B0GU86 | ACY1 | Aminoacylase-1 | -0.180720011 | -2.045731227 | Cellular metabolism |
| A0A1B0GUA3 | KIAA1279 | KIF1-binding protein | -0.310083389 | -0.803506215 | Transport |
| A0A1B0GUA8 | KIF13A | Kinesin-like protein KIF13A | 0.05067126 | -1.726423264 | Transport |
| A0A1B0GV47 | KIF21A | Kinesin-like protein KIF21A | 0.055004756 | 1.415619532 | Other |
| F8W787 | CTSD | Cathepsin D | -0.280762355 | -0.16525205 | Cellular metabolism |
| A0A1B0GVS8 | | | 0.351427714 | 2.052069346 | #N/A |
| A0A1B0GWI2 | CLN5 | Ceroid-lipofuscinosis neuronal protein 5 | -0.295988083 | -0.70259285 | Cellular metabolism |
| R4GND1 | UBE2E3 | Ubiquitin-conjugating enzyme E2 E2 | -0.568222046 | -0.836111069 | Cellular metabolism |
| A0A1C7CYX9 | DPYSL2 | Dihydropyrimidinase-related protein 2 | 0.062168121 | -0.506104787 | Transport |
| Q9BWM7 | SFXN3 | Sideroflexin-3 | 0 | -0.059725444 | Other |
| A0A1W2PPH1 | ME2 | NAD-dependent malic enzyme, mitochondrial | 0.385229747 | 0.099676132 | Cellular metabolism |
| A0A1W2PPZ5 | TCEA1 | Transcription elongation factor A protein 1 | -0.01063029 | -1.006734848 | Cellular metabolism |
| E9PNM1 | FDFT1 | Squalene synthase | 0.363299052 | 0 | Lipid metabolism |
| A0A1W2PQV2 | GCSH | Glycine cleavage system H protein, mitochondrial | -0.203573227 | -0.411794662 | Cellular metabolism |
| F2Z2I2 | PFKFB3 | 6-phosphofructo-2-kinase/fructose-2,6-bisphosphatase 3 | 1.589506149 | 0 | Cellular metabolism |
| A0A1W2PS43 | SCARB2 | Lysosome membrane protein 2 | -0.322051366 | 0.804573695 | Transport |
| B4DP61 | SMN2 | Survival motor neuron protein | -0.013855616 | -2.711058299 | Cellular metabolism |
| A0A1W2PS05 | PIK3C3 | Phosphatidylinositol 3-kinase catalytic subunit type 3 | -0.659950892 | -0.365280151 | Immune response |
| A0A494C0S2 | BIVM-ERCC5 | DNA repair protein complementing XP-G cells | 0.790757497 | 0 | Cellular metabolism |
| A0A1X7SBZ2 | DDX17 | Probable ATP-dependent RNA helicase DDX17 | 0.092955907 | 0.404556274 | Cellular metabolism |
| A0A286YES1 | IGHG3 | Ig gamma-3 chain C region | 0 | -2.540104548 | Immune response |
| A0A286YEY1 | IGHA1 | Ig alpha-1 chain C region | 1.105089188 | 1.107093811 | Immune response |
| A0A286YEY4 | IGHG2 | Ig gamma-2 chain C region | 0 | 0.32408905 | Immune response |
| A0A286YF22 | PHGDH | D-3-phosphoglycerate dehydrogenase | -0.032735189 | -0.54614385 | Cellular metabolism |
| A0A2C9F2P4 | PPT1 | Palmitoyl-protein thioesterase 1 | -0.124781926 | -0.537184397 | Lipid metabolism |

| | | | | | |
|---|---|---|---|---|---|
| A0A286YFF8 | MON2 | Protein MON2 homolog | -0.009711583 | -1.304138819 | Transport |
| A0A2Q3DP09 | ARMC9 | LisH domain-containing protein ARMC9 | -0.154440562 | -0.181089401 | Other |
| A0A2R8Y4L3 | CCDC82 | Coiled-coil domain-containing protein 82 | -0.930724462 | 0 | Other |
| Q969G3 | SMARCE1 | SWI/SNF-related matrix-associated actin-dependent regulator of chromatin subfamily E member 1 | -0.233814875 | 0 | Other |
| A0A2R8Y4W2 | EPS8 | Epidermal growth factor receptor kinase substrate 8 | -0.243492126 | 0 | Other |
| H3BV07 | TBC1D24 | TBC1 domain family member 24 | 0.02197965 | 0.205168406 | Other |
| A0A2R8Y804 | CTNNB1 | Catenin beta-1 | 0.738552094 | -0.369499842 | Immune response |
| A0A2R8Y566 | KIAA1468 | LisH domain and HEAT repeat-containing protein KIAA1468 | -0.226250331 | 0.541914622 | Other |
| A0A2R8Y575 | PLEKHM2 | Pleckstrin homology domain-containing family M member 2 | -0.123243332 | 0 | Other |
| A0A2R8Y595 | CTCF | Transcriptional repressor CTCF | 0.352692286 | -0.674558004 | Cellular metabolism |
| A0A2R8YEZ4 | POLR1C | DNA-directed RNA polymerases I and III subunit RPAC1 | 0.059112549 | -1.104357402 | Immune response |
| A0A2R8YDQ9 | SUCLA2 | Succinyl-CoA ligase [ADP-forming] subunit beta, mitochondrial | -0.152757009 | 0.387224833 | Cellular metabolism |
| Q3V6T2 | CCDC88A | Girdin | 1.148773829 | 0 | Cellular metabolism |
| Q9Y5P4-3 | COL4A3BP | Collagen type IV alpha-3-binding protein | -0.275346756 | -1.164999644 | Immune response |
| A0A2R8Y7R9 | PPFIA1 | Liprin-alpha-1 | -0.177703857 | 0 | Transport |
| P62263 | RPS14 | 40S ribosomal protein S14 | 0.334749858 | 0.208106359 | Transport |
| E7ETK0 | RPS24 | 40S ribosomal protein S24 | 0.076835632 | 1.893151601 | Transport |
| H3BSK8 | RBBP6 | E3 ubiquitin-protein ligase RBBP6 | 0.693210602 | 0.73281161 | Cellular metabolism |
| A0A2R8Y891 | PFKM | ATP-dependent 6-phosphofructokinase, muscle type | 0.021125793 | -0.602055868 | Cellular metabolism |
| Q5TBB1 | RNASEH2B | Ribonuclease H2 subunit B | -0.050175985 | 0.038394928 | Cellular metabolism |
| A0A2R8YGD1 | TPP1 | Tripeptidyl-peptidase 1 | -0.397081375 | -0.03184255 | Lipid metabolism |
| P68400 | CSNK2A1 | Casein kinase II subunit alpha | -0.022383372 | -0.749425252 | Cellular metabolism |
| A0A2R8YDE6 | HDAC6 | Histone deacetylase 6 | -0.468203227 | -2.218348821 | Transport |
| A0A2R8YE10 | VPS45 | Vacuolar protein sorting-associated protein 45 | -0.063439687 | 0 | Transport |
| A0A2R8YEQ9 | KCNQ1 | Potassium voltage-gated channel subfamily KQT member 1 | 0.264863968 | -1.221008937 | Transport |
| A0A2R8YFH5 | SEC23B | Protein transport protein Sec23B | -0.2115345 | 0.689001083 | Transport |
| Q14155-6 | ARHGEF7 | Rho guanine nucleotide exchange factor 7 | -1.152479808 | 0 | Other |
| A0A2R8YGH5 | AP1S1 | AP-1 complex subunit sigma-1A | 0.132591883 | -1.835312525 | Transport |
| A0A2U3TZH3 | EEF1A2 | Elongation factor 1-alpha 2 | -0.270495733 | 0.65877533 | Other |
| E9PNW4 | CD59 | CD59 glycoprotein | 1.018879573 | 0.394200643 | Immune response |
| A0A2U3TZL8 | KIF23 | Kinesin-like protein | -0.058497111 | 0.905101776 | Other |
| A0A3B3IRV1 | HDAC8 | Histone deacetylase 8 | -0.090096156 | 0 | Cellular metabolism |
| A0A3B3IRK6 | MOGS | Mannosyl-oligosaccharide glucosidase | 0 | 1.839258194 | Cellular metabolism |
| A0A3B3IRL5 | IMPDH1 | Inosine-5-monophosphate dehydrogenase | -0.762870153 | -0.740343094 | Cellular metabolism |

| | | | | | |
|---|---|---|---|---|---|
| A0A7P0Z4F9 | ADAR | Double-stranded RNA-specific adenosine deaminase | 0 | 1.827884038 | Immune response |
| A0A3B3IRR6 | C8orf33 | UPF0488 protein C8orf33 | 0 | 0.004460653 | Other |
| C9J3L8 | SSR1 | Translocon-associated protein subunit alpha | 0.526847839 | 1.453886668 | Transport |
| A0A7I2V2P6 | IDE | Insulin-degrading enzyme | -0.170553843 | -0.642885208 | Cellular metabolism |
| A0A3B3ITJ2 | UROS | Uroporphyrinogen-III synthase | -0.398342133 | 0 | Cellular metabolism |
| A0A3B3ISS6 | GPNMB | Transmembrane glycoprotein NMB | 0.095172246 | 0.366881053 | Other |
| F6QUN3 | ERI3 | ERI1 exoribonuclease 3 | 0.506474177 | 0 | Cellular metabolism |
| E9PG39 | AGK | Acylglycerol kinase, mitochondrial | 0 | 2.413574219 | Lipid metabolism |
| A0A3B3ITH3 | IMPACT | Protein IMPACT | 0.113604863 | 0 | Other |
| M0QXS5 | HNRNPL | Heterogeneous nuclear ribonucleoprotein L | 0.471209208 | 0.209111532 | Cellular metabolism |
| A0A3B3ITT5 | RPL29 | 60S ribosomal protein L29 | 0.359044393 | 0.146323522 | Transport |
| A0A3B3ITW3 | ABCD3 | ATP-binding cassette sub-family D member 3 | 1.200129827 | 1.132050196 | Lipid metabolism |
| E9PN17 | ATP5L | ATP synthase subunit g, mitochondrial | 0 | 1.278592428 | Transport |
| A0A3B3ITZ9 | THRAP3 | Thyroid hormone receptor-associated protein 3 | 0.150319417 | -0.003189087 | Cellular metabolism |
| H0YDP6 | GEMIN2 | Gem-associated protein 2 | 0.421544393 | 0 | Cellular metabolism |
| A0A3B3IU45 | MCM4 | DNA replication licensing factor MCM4 | 0.082855225 | -0.550136566 | Cellular metabolism |
| F8W717 | EML1 | Echinoderm microtubule-associated protein-like 1 | 0.43432045 | 0 | Other |
| Q13144 | EIF2B5 | Translation initiation factor eIF-2B subunit epsilon | -0.095921199 | -1.477411906 | Cellular metabolism |
| A0A3B3IUB5 | HM13 | Minor histocompatibility antigen H13 | 0.977595647 | 1.455868403 | Other |
| A0A3B3IUD7 | PDCD11 | Protein RRP5 homolog | 0.10310936 | 0.669846853 | Cellular metabolism |
| D6RH03 | IL6ST | Interleukin-6 receptor subunit beta | -1.879599889 | -2.0111866 | Cellular metabolism |
| A0A494BZY7 | METTL10 | Protein-lysine N-methyltransferase METTL10 | -0.74082311 | 0 | Other |
| Q9UHD2 | TBK1 | Serine/threonine-protein kinase TBK1 | -0.062383016 | -1.983520508 | Immune response |
| A0A494C066 | CCBL1 | Kynurenine--oxoglutarate transaminase 1 | -0.186037064 | -1.043221156 | Cellular metabolism |
| A0A494C068 | PRKAG2 | 5-AMP-activated protein kinase subunit gamma-2 | 0.164218267 | -0.557901382 | Lipid metabolism |
| O95394 | PGM3 | Phosphoacetylglucosamine mutase | -0.008544286 | -0.725150426 | Cellular metabolism |
| A0A7P0T9J1 | DIMT1 | Probable dimethyladenosine transferase | 0.810842514 | 0 | Other |
| A0A494C0J7 | TGM3 | Protein-glutamine gamma-glutamyltransferase E | -0.484154383 | 0.943608602 | Cellular metabolism |
| A0A494C0R8 | CLUH | | -0.026243846 | -0.917714437 | Other |
| A0A494C0W0 | PARN | Poly(A)-specific ribonuclease PARN | 0.822319667 | 0.099173864 | Cellular metabolism |
| A0A494C108 | KRI1 | Protein KRI1 homolog | 0.624575933 | -2.815901438 | Other |
| A0A494C164 | STAT2 | Signal transducer and activator of transcription 2 | -0.058533986 | 0.075330734 | Cellular metabolism |
| A0A494C1F2 | NUP214 | Nuclear pore complex protein Nup214 | -0.087000529 | 0.062707265 | Transport |
| Q14739 | LBR | Lamin-B receptor | 0 | 0.82483228 | Lipid metabolism |
| A0A494C1L5 | LRBA | Lipopolysaccharide-responsive and beige-like anchor protein | -0.010899226 | -1.305740992 | Transport |
| P26885 | FKBP2 | Peptidyl-prolyl cis-trans isomerase FKBP2 | 0.114811579 | 0.043209712 | Cellular metabolism |
| A0A499FI31 | SART3 | Squamous cell carcinoma antigen recognized by T-cells 3 | 0.083134969 | -0.649876912 | Cellular metabolism |
| A0A499FIZ0 | WDR26 | WD repeat-containing protein 26 | 0.261281967 | -0.505353292 | Other |

| | | | | | |
|---|---|---|---|---|---|
| A0A499FJS7 | IKBKB | Inhibitor of nuclear factor kappa-B kinase subunit beta | 0.205251058 | -0.83685112 | Immune response |
| U3KPX3 | CSNK1A1 | Casein kinase I isoform alpha | -0.372851054 | -0.348114649 | Transport |
| A0A590UJ73 | TSEN34 | tRNA-splicing endonuclease subunit Sen34 | -1.225775401 | 0.126622518 | Cellular metabolism |
| A0A590UJW8 | ITCH | E3 ubiquitin-protein ligase Itchy homolog | -0.585110982 | 0 | Immune response |
| A0A590UK80 | SRSF6 | Serine/arginine-rich splicing factor 6 | 0.264258067 | -0.286764781 | Transport |
| A0A5F9UK94 | AMPD2 | AMP deaminase 2 | -0.096572876 | -1.250473658 | Cellular metabolism |
| A0A5F9UP49 | SDF4 | 45 kDa calcium-binding protein | -0.205361684 | -0.700897217 | Transport |
| A0A5F9ZH29 | CDC45 | Cell division control protein 45 homolog | -1.042687734 | 0 | Cellular metabolism |
| A0A5F9ZH43 | XIRP2 | Xin actin-binding repeat-containing protein 2 | -0.224349976 | 0 | Other |
| A0A5F9ZHB6 | PRIM1 | DNA primase small subunit | 0.346338272 | -0.489421209 | Cellular metabolism |
| A0A5F9ZHG7 | RAPGEF6 | Rap guanine nucleotide exchange factor 6 | -0.079029719 | 0 | Other |
| P51648-2 | ALDH3A2 | Fatty aldehyde dehydrogenase | 0 | 1.776732763 | Lipid metabolism |
| A0A5F9ZHJ7 | ACAT1 | Acetyl-CoA acetyltransferase, mitochondrial | -0.018796921 | 0.161838531 | Lipid metabolism |
| E9PGZ4 | SACM1L | Phosphatidylinositide phosphatase SAC1 | 0.98314031 | 1.871628443 | Lipid metabolism |
| E9PQY3 | ACP2 | Lysosomal acid phosphatase | -0.404671351 | 0.459599177 | Cellular metabolism |
| A0A5F9ZHU7 | PRIM2 | DNA primase large subunit | -0.272445043 | 0 | Cellular metabolism |
| C9JX57 | C21orf59 | UPF0769 protein C21orf59 | -0.31783994 | 0 | Other |
| A0A5F9ZI21 | ARHGEF2 | Rho guanine nucleotide exchange factor 2 | 0.839894613 | 1.071320852 | Immune response |
| A0A5F9ZI63 | EED | Polycomb protein EED | 0.453339259 | 0.894913991 | Cellular metabolism |
| A0A5H1ZRP4 | NELFB | Negative elongation factor B | -0.286010742 | -0.394380569 | Cellular metabolism |
| A0A5H1ZRS1 | PCM1 | Pericentriolar material 1 protein | 0.143245697 | 0.274247487 | Transport |
| H0YH69 | ETNK1 | Ethanolamine kinase 1 | -1.999165853 | 0 | Lipid metabolism |
| A0A5K1VW74 | AKT3 | RAC-gamma serine/threonine-protein kinase | -0.283260345 | 0 | Cellular metabolism |
| Q13620 | CUL4B | Cullin-4B | -0.154659271 | -0.369928996 | Cellular metabolism |
| O60716 | CTNND1 | Catenin delta-1 | 1.02286466 | -0.227543513 | Cellular metabolism |
| Q92538 | GBF1 | Golgi-specific brefeldin A-resistance guanine nucleotide exchange factor 1 | -0.154249827 | -1.265028 | Transport |
| E7EPV7 | SNCA | Alpha-synuclein | -0.648870468 | -1.196715673 | Lipid metabolism |
| A0A669KBL1 | USP7 | Ubiquitin carboxyl-terminal hydrolase 7 | 0.186040878 | -0.363162994 | Cellular metabolism |
| A0A6I8PIV1 | LARS | Leucine--tRNA ligase, cytoplasmic | 0.063318253 | -0.818532308 | Cellular metabolism |
| A0A6I8PRS5 | GPATCH11 | G patch domain-containing protein 11 | -0.228136063 | -0.494425456 | Other |
| Q99615 | DNAJC7 | DnaJ homolog subfamily C member 7 | -0.254244486 | -0.21319898 | Cellular metabolism |
| I3L1H5 | DPH1 | Diphthamide biosynthesis protein 1 | -0.381605784 | 0.247406642 | Cellular metabolism |
| A0A6Q8KRG2 | GOLGA2 | Golgin subfamily A member 2 | 0.042417526 | -0.97686259 | Other |
| O95163 | IKBKAP | Elongator complex protein 1 | 0.115336736 | -0.69601059 | Immune response |

| | | | | | |
|---|---|---|---|---|---|
| A0A6Q8PGH7 | KIF2A | Kinesin-like protein KIF2A | 0.314211528 | -0.259695689 | Other |
| O95831 | AIFM1 | Apoptosis-inducing factor 1, mitochondrial | 0 | 0.547320048 | Cellular metabolism |
| A0A6Q8PHJ5 | NEDD4L | E3 ubiquitin-protein ligase NEDD4-like | -0.345345815 | -1.232447942 | Transport |
| A0A6Q8PFJ0 | LMNA | Prelamin-A/C | -0.267338435 | 0.00078392 | Transport |
| A0A6Q8PFK4 | HLCS | Biotin--protein ligase | -0.229729335 | 0 | Cellular metabolism |
| Q6UWE0 | LRSAM1 | E3 ubiquitin-protein ligase LRSAM1 | 0.067885717 | -1.481121699 | Transport |
| A0A6Q8PFU3 | PAFAH1B1 | Platelet-activating factor acetylhydrolase IB subunit alpha | 0.262657166 | -0.573818207 | Lipid metabolism |
| A0A6Q8PFY7 | TFG | Protein TFG | -0.158711116 | -0.210729599 | Other |
| A0A6Q8PGB0 | POLR2A | DNA-directed RNA polymerase II subunit RPB1 | -0.74296697 | -1.128729502 | Cellular metabolism |
| Q27J81 | INF2 | Inverted formin-2 | -0.239992142 | -0.588486989 | Other |
| P49588 | AARS | Alanine--tRNA ligase, cytoplasmic | -0.275938034 | -0.808434168 | Cellular metabolism |
| A0A6Q8PHS7 | MTMR2 | Myotubularin-related protein 2 | -0.441029231 | -0.470790863 | Lipid metabolism |
| A0A6Q8PGU6 | SPATA5 | Spermatogenesis-associated protein 5 | 0.43004926 | 0.384862264 | Other |
| A0A6Q8PH09 | FANCI | Fanconi anemia group I protein | -0.747762044 | 0.277667999 | Cellular metabolism |
| A0A6Q8PHK0 | TRIP12 | E3 ubiquitin-protein ligase TRIP12 | -0.294221242 | -1.456385295 | Cellular metabolism |
| P49902 | NT5C2 | Cytosolic purine 5-nucleotidase | -0.030243556 | -1.421209971 | Cellular metabolism |
| A0A7I2U302 | MME | Neprilysin | -0.436340968 | 1.339599609 | Cellular metabolism |
| O14964 | HGS | Hepatocyte growth factor-regulated tyrosine kinase substrate | -0.138825099 | -0.474134445 | Transport |
| H7C4K3 | ITGB1 | Integrin beta-1 | 0.532161077 | -0.053609212 | Transport |
| F8VZJ2 | NACA | Nascent polypeptide-associated complex subunit alpha | -0.190146128 | -0.508465449 | Transport |
| A0A7I2V466 | CTSC | Dipeptidyl peptidase 1 | -0.073044459 | -0.722500483 | Immune response |
| A0A7I2V2R3 | HNRNPA3 | Heterogeneous nuclear ribonucleoprotein A3 | 0.019584656 | -0.030946096 | Cellular metabolism |
| A0A7I2V2T3 | IMPDH2 | Inosine-5-monophosphate dehydrogenase 2 | 0.034395854 | -0.756538391 | Cellular metabolism |
| A0A7I2V4T9 | SGTA | Small glutamine-rich tetratricopeptide repeat-containing protein alpha | -0.007870992 | -0.675029119 | Other |
| A0A7I2V2U6 | MTHFD2 | Bifunctional methylenetetrahydrofolate dehydrogenase/cyclohydrolase, mitochondrial | -0.663002014 | 0.650901159 | Cellular metabolism |
| Q5JPE7 | NOMO2 | Nodal modulator 2 | -0.000699361 | 1.213239034 | Other |
| A0A7I2V351 | KPNA2 | Importin subunit alpha-1 | -0.295717239 | -0.307813644 | Transport |
| Q8WVM8 | SCFD1 | Sec1 family domain-containing protein 1 | -0.130236308 | -0.12399292 | Transport |
| P60228 | EIF3E | Eukaryotic translation initiation factor 3 subunit E | 0.133949916 | -0.378740946 | Cellular metabolism |
| P09651-2 | HNRNPA1 | Heterogeneous nuclear ribonucleoprotein A1 | 0.21950086 | 0.698850632 | Transport |
| P40763 | STAT3 | Signal transducer and activator of transcription 3 | -1.007226944 | -1.921508153 | Transport |
| A0A7I2V423 | H2AFY2 | Core histone macro-H2A.2 | 0.737838109 | -0.318106333 | Other |

| | | | | | |
|---|---|---|---|---|---|
| A0A7I2V428 | NCL | Nucleolin | 0.184666316 | -0.073894501 | Other |
| A0A7I2V430 | DDX1 | ATP-dependent RNA helicase DDX1 | 0.21609052 | -0.057576497 | Cellular metabolism |
| P26358-2 | DNMT1 | DNA (cytosine-5)-methyltransferase 1 | -1.094662348 | 0.302611669 | Cellular metabolism |
| A0A7I2V4J2 | NONO | Non-POU domain-containing octamer-binding protein | 0.81563441 | 0.586298625 | Cellular metabolism |
| P10619 | CTSA | Lysosomal protective protein | 0.058596929 | -0.249121348 | Lipid metabolism |
| A0A7I2V4L6 | YKT6 | Synaptobrevin homolog YKT6 | -0.289670308 | -0.069154104 | Transport |
| A0A7I2V4L7 | PABPC4 | Polyadenylate-binding protein | 0.003195445 | 0.18541654 | Cellular metabolism |
| A0A7I2V4X8 | NDUFS2 | NADH dehydrogenase [ubiquinone] iron-sulfur protein 2, mitochondrial | 0 | 1.336042404 | Cellular metabolism |
| A0A7I2V539 | NMT1 | Glycylpeptide N-tetradecanoyltransferase 1 | -0.081180573 | -0.099532445 | Cellular metabolism |
| O95299 | NDUFA10 | NADH dehydrogenase [ubiquinone] 1 alpha subcomplex subunit 10, mitochondrial | 0.246529261 | 1.738090515 | Cellular metabolism |
| Q5QP56 | BCL2L1 | Bcl-2-like protein 1 | -0.200670878 | 0.601769129 | Immune response |
| Q5JRX3-2 | PITRM1 | Presequence protease, mitochondrial | -0.490735372 | -0.888432185 | Other |
| C9JN15 | PPIG | Peptidyl-prolyl cis-trans isomerase | -0.734308243 | 0 | Cellular metabolism |
| A0A7I2V5R1 | NEK9 | Serine/threonine-protein kinase Nek9 | -0.044353485 | -0.590130488 | Cellular metabolism |
| A0A7I2V5T4 | ACO2 | Aconitate hydratase, mitochondrial | 0.025248845 | -0.124141057 | Cellular metabolism |
| A0A7I2V5X6 | PTGES2 | Prostaglandin E synthase 2 | 0.040763219 | 1.54240799 | Lipid metabolism |
| Q13347 | EIF3I | Eukaryotic translation initiation factor 3 subunit I | 0.176809311 | -0.392331441 | Cellular metabolism |
| J3QR09 | RPL19 | Ribosomal protein L19 | -0.199419022 | 2.493065516 | Transport |
| A0A7N4I394 | PRPF40A | Pre-mRNA-processing factor 40 homolog A | 0.617916743 | 0.108109792 | Cellular metabolism |
| G5E9W7 | MRPS22 | 28S ribosomal protein S22, mitochondrial | -1.599499385 | 1.580657959 | Cellular metabolism |
| O96017-9 | CHEK2 | Serine/threonine-protein kinase Chk2 | 0.318350474 | 1.49870491 | Cellular metabolism |
| A0A7P0T9L9 | ARFGEF2 | Brefeldin A-inhibited guanine nucleotide-exchange protein 2 | -0.033845901 | 0 | Transport |
| A0A7P0T8A1 | CTSS | Cathepsin S | 0 | -1.678487778 | Immune response |
| Q99986 | VRK1 | Serine/threonine-protein kinase VRK1 | -0.248566945 | 0.210333506 | Transport |
| A0A7P0T8U3 | ACAD9 | Acyl-CoA dehydrogenase family member 9, mitochondrial | 0 | -0.058232625 | Other |
| E9PHM6 | DST | Dystonin | 0.393324534 | 0 | Other |
| A0A7P0T893 | AKAP8 | A-kinase anchor protein 8 | 0.24621582 | -0.978569031 | Cellular metabolism |
| A0A7P0T8A6 | MICALL1 | MICAL-like protein 1 | -0.219196955 | 0 | Transport |
| A0A7P0Z4A2 | SNX9 | Sorting nexin-9 | 0.091181437 | -1.019219716 | Transport |
| Q96Q42 | ALS2 | Alsin | 0.084982554 | 0 | Transport |
| A0A7P0T8J7 | AAR2 | Protein AAR2 homolog | 0.849527995 | 0.304316203 | Other |
| G3V220 | TCERG1 | Transcription elongation regulator 1 | 0.695582708 | 0.297662099 | Cellular metabolism |
| A0A7P0T9X2 | ALG5 | Dolichyl-phosphate beta-glucosyltransferase | 0.668657939 | 3.211461385 | Cellular metabolism |
| A0A7P0T9D8 | CARKD | ATP-dependent (S)-NAD(P)H-hydrate dehydratase | 0.140476863 | -0.179199855 | Cellular metabolism |

| | | | | | |
|---|---|---|---|---|---|
| A0A7P0T915 | SIRT5 | NAD-dependent protein deacylase sirtuin-5, mitochondrial | -0.025677363 | 0 | Cellular metabolism |
| P27824-2 | CANX | Calnexin | 0.43380928 | 0.96541214 | Transport |
| Q9UBR2 | CTSZ | Cathepsin Z | -0.145433426 | -0.729701996 | Cellular metabolism |
| O43379-4 | WDR62 | WD repeat-containing protein 62 | -0.204122543 | 0 | Other |
| A0A7P0T9G6 | TUBG1 | Tubulin gamma-1 chain | -0.581842422 | 1.871583939 | Other |
| P43490 | NAMPT | Nicotinamide phosphoribosyltransferase | -0.022680918 | -0.363824844 | Cellular metabolism |
| P07237 | P4HB | Protein disulfide-isomerase | 0.236988703 | -0.326690674 | Cellular metabolism |
| A0A7P0TA78 | HTT | Huntingtin | 0.141130447 | 0 | Transport |
| A0A7P0TA85 | BSG | Basigin | 1.168517431 | 0.907388051 | Transport |
| A0A7P0TAE3 | AKAP8L | A-kinase anchor protein 8-like | -0.263040543 | -0.438563665 | Other |
| A0A7P0TAG0 | MMP14 | Matrix metalloproteinase-14 | -1.083159129 | -0.856712977 | Other |
| H0YDD4 | DLAT | Acetyltransferase component of pyruvate dehydrogenase complex | 0.630049388 | 0 | Cellular metabolism |
| P08195-4 | SLC3A2 | 4F2 cell-surface antigen heavy chain | 0.106487274 | 0.840919495 | Transport |
| P55072 | VCP | Transitional endoplasmic reticulum ATPase | 0.047398885 | 0.044665019 | Transport |
| A0A7P0TBA9 | EIF2AK2 | Interferon-induced, double-stranded RNA-activated protein kinase | 0.711507161 | 0.772591273 | Immune response |
| A0A7P0Z415 | CTSV | Cathepsin L2 | 0 | -1.136917114 | Immune response |
| A0A7P0Z497 | PPIB | Peptidyl-prolyl cis-trans isomerase B | 0.293884913 | 0.413626989 | Cellular metabolism |
| E7EX90 | DCTN1 | Dynactin subunit 1 | 0.374755859 | -0.790117264 | Transport |
| A0AVT1 | UBA6 | Ubiquitin-like modifier-activating enzyme 6 | -0.091346741 | -0.401789347 | Cellular metabolism |
| A0MZ66 | KIAA1598 | Shootin-1 | -0.481425603 | 1.357098262 | Other |
| M0R026 | ILVBL | Acetolactate synthase-like protein | 0 | 0.101247152 | Other |
| A1X283 | SH3PXD2B | SH3 and PX domain-containing protein 2B | 0.562388102 | 0.368687948 | Cellular metabolism |
| A2AB05 | MDC1 | Mediator of DNA damage checkpoint protein 1 | 0.029865901 | -2.727043152 | Cellular metabolism |
| Q5QPD2 | AURKA | Aurora kinase A | -1.567861557 | 0 | Cellular metabolism |
| A3KN83-3 | SBNO1 | Protein strawberry notch homolog 1 | -0.273206075 | 0 | Cellular metabolism |
| A5YKK6 | CNOT1 | CCR4-NOT transcription complex subunit 1 | 0.266373952 | 0.496676127 | Cellular metabolism |
| H3BTL1 | MAP1LC3B | Microtubule-associated proteins 1A/1B light chain 3B | -0.42312177 | -0.196860631 | Cellular metabolism |
| A6ND36 | FAM83G | Protein FAM83G | 0.080256144 | 0 | Other |
| A6NDG6 | PGP | Phosphoglycolate phosphatase | -0.112073898 | -0.219409307 | Cellular metabolism |
| A6NDU8 | C5orf51 | UPF0600 protein C5orf51 | 0.117363612 | -1.121470133 | Other |
| G3V2I3 | RCCD1 | RCC1 domain-containing protein 1 | -0.96186765 | 0 | Other |
| A6NEM2 | HCFC1 | Host cell factor 1 | 0.202232361 | 0.099722544 | Cellular metabolism |
| A6NFX8 | NUDT5 | ADP-sugar pyrophosphatase | -0.071252823 | -0.398993174 | Cellular metabolism |
| A6NH27 | ZWINT | ZW10 interactor | -1.311109543 | 0 | Other |
| J3KTL8 | SMCHD1 | Structural maintenance of chromosomes flexible hinge domain-containing protein 1 | 0 | 0.273351669 | Other |

| | | | | | |
|---|---|---|---|---|---|
| A6NHX0 | GATSL2 | GATS-like protein 2 | -0.172248205 | 0 | Other |
| A6NIE6 | RRN3P2 | Putative RRN3-like protein RRN3P2 | 1.226430257 | 1.161841075 | Cellular metabolism |
| A6NIH7 | UNC119B | Protein unc-119 homolog B | -0.108489354 | -1.022572835 | Transport |
| A6NMA8 | CRYZL1 | Quinone oxidoreductase-like protein 1 | 0.068950017 | -1.011322657 | Other |
| P09884 | POLA1 | DNA polymerase alpha catalytic subunit | -0.276435216 | 0 | Cellular metabolism |
| A6NMQ3 | ENSA | Alpha-endosulfine | -1.005482992 | -1.114678701 | Transport |
| A6NP16 | AMOT | Angiomotin | -1.69352595 | 1.027143478 | Other |
| A6ZKI3 | FAM127A | Protein FAM127A | -1.119913101 | 0 | Other |
| A8K2U0 | A2ML1 | Alpha-2-macroglobulin-like protein 1 | -0.069670995 | 1.05130132 | Other |
| A8MTY9 | DSCR3 | Down syndrome critical region protein 3 | -0.532419205 | 0 | Transport |
| A8MUA9 | SUMO3 | Small ubiquitin-related modifier 3 | 0.032164892 | 0.201888402 | Cellular metabolism |
| A8MUM1 | TSSC1 | Protein TSSC1 | -0.048299154 | -1.129736582 | Other |
| P62308 | SNRPG | Small nuclear ribonucleoprotein G | 0.244082769 | -0.058710734 | Cellular metabolism |
| A8MX75 | ERCC2 | TFIIH basal transcription factor complex helicase XPD subunit | -0.696642558 | 0 | Cellular metabolism |
| A8MXB7 | SNX24 | Sorting nexin-24 | -0.321853002 | 0 | Transport |
| C9JMC5 | ALDH3A1 | Aldehyde dehydrogenase, dimeric NADP-preferring | 0 | 2.199584961 | Cellular metabolism |
| A9Z1X7 | SRRM1 | Serine/arginine repetitive matrix protein 1 | 0.337441762 | -0.562327067 | Transport |
| Q9NWS6 | FAM118A | Protein FAM118A | 0.325138092 | 0 | Other |
| B0QYK0 | EWSR1 | RNA-binding protein EWS | 0.071611404 | 0.288955053 | Cellular metabolism |
| H3BQQ9 | UBE2I | SUMO-conjugating enzyme UBC9 | 0.215764999 | 0.161534627 | Cellular metabolism |
| B1AH87 | TSPO | Translocator protein | 0 | 0.73560969 | Lipid metabolism |
| B1AJQ6 | STX12 | Syntaxin-12 | 0 | 0.236415227 | Transport |
| P29323-2 | EPHB2 | Ephrin type-B receptor 2 | 0 | -0.882396062 | Cellular metabolism |
| B1AL69 | CDC37L1 | Hsp90 co-chaperone Cdc37-like 1 | -0.296890895 | 0 | Cellular metabolism |
| P60891 | PRPS1 | Ribose-phosphate pyrophosphokinase 1 | 0.106444677 | 0.088176092 | Cellular metabolism |
| B1AMU7 | EXOSC1 | Exosome complex component CSL4 | 1.487824122 | 0 | Cellular metabolism |
| B1AN48 | SPRR3 | Small proline-rich protein 3 | 0 | -0.262137731 | Cellular metabolism |
| B1APM4 | SOAT1 | Sterol O-acyltransferase 1 | 2.004689535 | 2.610294978 | Lipid metabolism |
| B2R5W2 | HNRNPC | Heterogeneous nuclear ribonucleoproteins C1/C2 | 0.539871852 | 0.094637553 | Cellular metabolism |
| B2WTI3 | JMJD6 | Bifunctional arginine demethylase and lysyl-hydroxylase JMJD6 | -0.245400747 | 0.27526029 | Cellular metabolism |
| C9J5N1 | PTGES3L-AARSD1 | Alanyl-tRNA editing protein Aarsd1 | -0.320279439 | -0.796341578 | Cellular metabolism |
| B3KTM8 | MORF4L1 | Mortality factor 4-like protein 1 | -1.213912328 | -0.459065119 | Cellular metabolism |
| B3KVZ3 | CENPH | Centromere protein H | 0.218269348 | -2.103666306 | Other |
| B4DDF4 | CNN2 | Calponin | 0.376387914 | -0.153883616 | Other |
| U3KQP3 | NDUFA13 | NADH dehydrogenase [ubiquinone] 1 alpha subcomplex subunit 13 | 0 | 0.829809189 | Transport |

| | | | | | |
|---|---|---|---|---|---|
| B4DHR0 | RABEP2 | Rab GTPase-binding effector protein 2 | 0.59223938 | -0.450344086 | Transport |
| B4DJ81 | NDUFS1 | NADH-ubiquinone oxidoreductase 75 kDa subunit, mitochondrial | 2.178972244 | 0.695549011 | Cellular metabolism |
| O75390 | CS | Citrate synthase, mitochondrial | -0.030169169 | 0.227198919 | Cellular metabolism |
| B4DK69 | AKR1C2 | Aldo-keto reductase family 1 member C2 | -0.646179199 | 0.069276174 | Lipid metabolism |
| B4DLN1 | MRPL12 | 39S ribosomal protein L12, mitochondrial | 2.310069402 | 1.349509557 | Transport |
| B4DLN5 | SH2B1 | SH2B adapter protein 1 | 0.355909348 | 0 | Other |
| H0Y4S7 | RBCK1 | RanBP-type and C3HC4-type zinc finger-containing protein 1 | -0.074899673 | 0 | Cellular metabolism |
| B4DR80 | STK24 | Serine/threonine-protein kinase 24 | 0.359064102 | -1.164168676 | Cellular metabolism |
| B4DTF2 | ANXA8L1 | Annexin | 0 | -1.107526143 | Transport |
| B4DUC8 | MTAP | S-methyl-5-thioadenosine phosphorylase | -0.233394623 | -0.905045827 | Cellular metabolism |
| B4DXZ6 | FXR1 | Fragile X mental retardation syndrome-related protein 1 | 0.301434835 | -0.366544724 | Other |
| B4E0T2 | PEX5 | Peroxisomal targeting signal 1 receptor | -0.808734894 | 0 | Lipid metabolism |
| Q8NBY1 | STK26 | Serine/threonine-protein kinase 26 | -0.063237508 | -0.267981847 | Cellular metabolism |
| B5MBZ0 | EML4 | Echinoderm microtubule-associated protein-like 4 | 0.092241923 | -0.823989232 | Other |
| B5MCF9 | PES1 | Pescadillo homolog | 0.19017601 | 0.471429825 | Cellular metabolism |
| B5MDQ0 | ERCC6L | DNA excision repair protein ERCC-6-like | -0.305809021 | 0.297012329 | Other |
| E7EW69 | 40422 | Septin-10 | 0.132646561 | -0.133771261 | Other |
| B7WP74 | CWC22 | Pre-mRNA-splicing factor CWC22 homolog | -0.252203623 | 0.914747874 | Cellular metabolism |
| B7WPG3 | HNRNPLL | Heterogeneous nuclear ribonucleoprotein L-like | 1.422981898 | 0 | Cellular metabolism |
| B7Z2Y2 | COG2 | Conserved oligomeric Golgi complex subunit 2 | -0.592089971 | 0 | Transport |
| B7Z3Q4 | AGPS | Alkyldihydroxyacetonephosphate synthase, peroxisomal | 0 | 3.016913096 | Lipid metabolism |
| H7C2W9 | RPL31 | 60S ribosomal protein L31 | 0.143193563 | 0.357250849 | Transport |
| B7ZAX5 | GALK2 | N-acetylgalactosamine kinase | -0.409842173 | 0 | Cellular metabolism |
| B7ZBQ3 | MED20 | Mediator of RNA polymerase II transcription subunit 20 | 0.092173258 | 0 | Cellular metabolism |
| B7ZC38 | SH3GLB2 | Endophilin-B2 | -0.06351916 | -0.090194702 | Other |
| B7ZL14 | FNBP1 | Formin-binding protein 1 | -0.459712982 | 0.622680028 | Transport |
| C9JEL3 | EIF4E2 | Eukaryotic translation initiation factor 4E type 2 | 0.681142807 | 0 | Other |
| F5H4E8 | METTL5 | Methyltransferase-like protein 5 | -0.073367437 | 0.181007385 | Other |
| B8ZZN6 | SUMO1 | Small ubiquitin-related modifier 1 | -0.193759918 | 0.397562027 | Cellular metabolism |
| B8ZZQ6 | PTMA | Prothymosin alpha | 0.065076828 | 0.366464615 | Cellular metabolism |
| B8ZZU8 | TCEB2 | Transcription elongation factor B polypeptide 2 | -0.048154831 | -0.43155098 | Cellular metabolism |
| H7C0I1 | AVL9 | Late secretory pathway protein AVL9 homolog | 0.042873383 | 0.29498291 | Other |
| C4AMC7 | WASH3P | Putative WAS protein family homolog 3 | 0.295293172 | -0.609100342 | Transport |
| C9IZD3 | ARHGAP17 | Rho GTPase-activating protein 17 | -0.106663386 | 0 | Other |

| | | | | | |
|---|---|---|---|---|---|
| C9J0I9 | ZC3HC1 | Nuclear-interacting partner of ALK | 0.042568843 | -0.847096125 | Cellular metabolism |
| C9J0K6 | SRI | Sorcin | -0.208386739 | -0.396268209 | Transport |
| C9J126 | CDH2 | Cadherin-2 | 0 | 1.148264567 | Other |
| C9J2Y9 | POLR2B | DNA-directed RNA polymerase | -0.104122162 | -0.619850159 | Cellular metabolism |
| Q8IUF1 | CBWD2 | COBW domain-containing protein 2 | -0.694098155 | 0 | Other |
| C9J4Z3 | RPL37A | 60S ribosomal protein L37a | 0.296823502 | 0.017359416 | Transport |
| C9J715 | NISCH | Nischarin | -0.086259842 | 0 | Transport |
| F8WDQ9 | UBE2F | NEDD8-conjugating enzyme UBE2F | -0.55604744 | 0 | Cellular metabolism |
| C9JA08 | NMD3 | 60S ribosomal export protein NMD3 | -0.057886759 | 0.172510783 | Transport |
| C9JNK6 | MTX2 | Metaxin-2 | 0 | 1.562407176 | Transport |
| C9JB30 | MAPRE3 | Microtubule-associated protein RP/EB family member 3 | 0.389673233 | 0 | Other |
| C9JBI3 | PSPH | Phosphoserine phosphatase | -0.273591359 | -0.542027791 | Cellular metabolism |
| C9JCC6 | DRAP1 | Dr1-associated corepressor | -0.0779953 | 0.126752218 | Cellular metabolism |
| C9JG97 | AAMP | Angio-associated migratory cell protein | -0.207778931 | -1.112873713 | Other |
| C9JEJ2 | PCYT1A | Choline-phosphate cytidylyltransferase A | -0.245351156 | -0.301024755 | Lipid metabolism |
| C9JEZ4 | CDC42EP3 | Cdc42 effector protein 3 | -0.221079508 | 0.651631037 | Other |
| P05090 | APOD | Apolipoprotein D | 0 | 1.194588979 | Lipid metabolism |
| C9JF58 | OSCP1 | Protein OSCP1 | -0.05610466 | -0.701494217 | Transport |
| C9JFE4 | GPS1 | COP9 signalosome complex subunit 1 | 0.043797175 | -0.382298787 | Cellular metabolism |
| C9JFR7 | CYCS | Cytochrome c | -0.366037369 | 0.52411588 | Cellular metabolism |
| C9JFV4 | PELP1 | Proline-, glutamic acid- and leucine-rich protein 1 | 0.060642242 | 1.098126729 | Cellular metabolism |
| C9JIF9 | APEH | Acylamino-acid-releasing enzyme | 0.094063441 | -0.087481817 | Cellular metabolism |
| C9JJ19 | MRPS34 | 28S ribosomal protein S34, mitochondrial | -0.312912623 | 0.352067947 | Cellular metabolism |
| C9JJT5 | ATP5J2-PTCD1 | ATP synthase subunit f, mitochondrial | -0.263320287 | 0.770674388 | Transport |
| C9JLU1 | POLR2H | DNA-directed RNA polymerases I, II, and III subunit RPABC3 | 0.346542994 | -0.34416453 | Immune response |
| C9JLV4 | APAF1 | Apoptotic protease-activating factor 1 | -0.237571716 | -1.325679779 | Other |
| I3L2L5 | FAM195B | Protein FAM195B | -0.241639455 | 0 | Other |
| C9JXB8 | RPL24 | 60S ribosomal protein L24 | 0.217758814 | -0.11198616 | Transport |
| C9JP16 | CRTAP | Cartilage-associated protein | -0.158498764 | -0.420136134 | Cellular metabolism |
| G5E9W8 | GYG1 | Glycogenin-1 | -0.135724386 | 0.452545166 | Cellular metabolism |
| C9JQV0 | C7orf50 | Uncharacterized protein C7orf50 | 0.700800578 | 0.710149765 | Other |
| C9JRD2 | DNAJB2 | DnaJ homolog subfamily B member 2 | 0.197066625 | 0 | Cellular metabolism |
| C9JRZ6 | CHCHD3 | MICOS complex subunit MIC19 | 0 | -0.294780095 | Other |
| C9JTD3 | STARD7 | StAR-related lipid transfer protein 7, mitochondrial | 0.178012212 | -1.467383703 | Other |
| C9JUM8 | IFI44 | Interferon-induced protein 44 | 0.66476059 | 0 | Other |

| | | | | | |
|---|---|---|---|---|---|
| C9JX88 | PSMC2 | 26S protease regulatory subunit 7 | 0.072968801 | -0.429579417 | Immune response |
| C9JYM0 | POP7 | Ribonuclease P protein subunit p20 | 0.921763738 | 0 | Cellular metabolism |
| C9JZY6 | UBE2H | Ubiquitin-conjugating enzyme E2 H | -0.091726939 | -0.59962972 | Cellular metabolism |
| C9K0J5 | RAPH1 | Ras-associated and pleckstrin homology domains-containing protein 1 | 0.017795563 | 0 | Other |
| D3DQV9 | EIF4G2 | Eukaryotic translation initiation factor 4 gamma 2 | 0.191571554 | -0.168534597 | Other |
| F6U1T9 | PPP3R1 | Calcineurin subunit B type 1 | 0.07650884 | 0.201840719 | Immune response |
| D3YTB1 | RPL32 | 60S ribosomal protein L32 | 0.078227997 | 0.032004038 | Transport |
| Q2L6G2 | HLA-B | | 0 | 2.560462952 | Immune response |
| Q13557-12 | CAMK2D | Calcium/calmodulin-dependent protein kinase type II subunit delta | -0.315017064 | 0.833620071 | Transport |
| D6R9P3 | HNRNPAB | Heterogeneous nuclear ribonucleoprotein A/B | 0.230122248 | 0.73990949 | Other |
| D6R9U7 | POLR3G | DNA-directed RNA polymerase III subunit RPC7 | -0.934155146 | -0.907904307 | Immune response |
| D6RIZ4 | MFSD10 | Major facilitator superfamily domain-containing protein 10 | 1.618548075 | -0.334842682 | Transport |
| D6RB01 | FAM169A | Soluble lamin-associated protein of 75 kDa | -0.326782862 | 0 | Other |
| D6RB92 | PSMG4 | Proteasome assembly chaperone 4 | 0.127063115 | 0 | Other |
| D6RCD7 | HSPA9 | Stress-70 protein, mitochondrial | -0.069290161 | 0.013337453 | Transport |
| D6RCM8 | TNFAIP8 | Tumor necrosis factor alpha-induced protein 8 | 0.036482493 | -1.001302083 | Other |
| Q9H173 | SIL1 | Nucleotide exchange factor SIL1 | 0.047578176 | -0.998351415 | Transport |
| D6RGI3 | 40787 | Septin-11 | 0.120093028 | -0.291410446 | Other |
| D6REX3 | SEC31A | Protein transport protein Sec31A | 0.016846975 | -0.299044291 | Transport |
| D6RFG8 | DCK | Deoxycytidine kinase | -0.279084524 | -0.475021998 | Cellular metabolism |
| D6RFH4 | CYB5B | Cytochrome b5 type B | 0 | -0.776444117 | Other |
| D6RIY6 | EXOSC9 | Exosome complex component RRP45 | 0.383716583 | 0 | Immune response |
| Q9Y6G5 | COMMD10 | COMM domain-containing protein 10 | 1.227436701 | 0 | Cellular metabolism |
| D6W601 | ADAT3 | Probable inactive tRNA-specific adenosine deaminase-like protein 3 | -0.146666845 | 0 | Cellular metabolism |
| E2QRF9 | GMNN | Geminin | 0.097634633 | 0.20065244 | Other |
| E5RFF9 | GINS4 | DNA replication complex GINS protein SLD5 | -0.067860285 | 1.044146856 | Cellular metabolism |
| E5RFZ8 | SLC39A14 | Zinc transporter ZIP14 | 0 | 1.103667577 | Transport |
| E5RG17 | TATDN1 | Putative deoxyribonuclease TATDN1 | -0.075546265 | -0.52004687 | Cellular metabolism |
| E5RGC1 | FBXW11 | F-box/WD repeat-containing protein 11 | -0.069386164 | -1.120105108 | Immune response |
| E5RGQ3 | RWDD1 | RWD domain-containing protein 1 | 0.340113322 | -0.183480581 | Other |
| E5RHF4 | MRPL15 | 39S ribosomal protein L15, mitochondrial | 0 | 0.77916654 | Cellular metabolism |
| E5RHG8 | TCEB1 | Transcription elongation factor B polypeptide 1 | 0.137819926 | -0.589114507 | Cellular metabolism |
| E5RHW4 | ERLIN2 | Erlin-2 | 0.929735184 | 0.803583145 | Cellular metabolism |
| E5RI99 | RPL30 | 60S ribosomal protein L30 | 0.607964834 | 1.170572917 | Transport |
| E5RIA4 | DNPEP | Aspartyl aminopeptidase | 0.115448634 | 0.372039159 | Cellular metabolism |
| O00244 | ATOX1 | Copper transport protein ATOX1 | -0.294834773 | -0.190463384 | Transport |

| | | | | | |
|---|---|---|---|---|---|
| E5RIW3 | TBCA | Tubulin-specific chaperone A | -0.538203557 | -0.008546193 | Cellular metabolism |
| E5RK69 | ANXA6 | Annexin | -0.160760244 | -0.446208954 | Other |
| Q7L5N1 | COPS6 | COP9 signalosome complex subunit 6 | 0.109472275 | -0.582335154 | Cellular metabolism |
| E7EMD6 | AKAP10 | A-kinase anchor protein 10, mitochondrial | 0.384503047 | 0 | Other |
| E7EMP6 | HMCES | Embryonic stem cell-specific 5-hydroxymethylcytosine-binding protein | -0.347562154 | 0 | Other |
| J3KMY5 | NPC2 | Epididymal secretory protein E1 | 0.003708522 | -0.366108576 | Lipid metabolism |
| E7EV99 | ADD1 | Alpha-adducin | -0.272119522 | -1.37876447 | Transport |
| E7EPB3 | RPL14 | 60S ribosomal protein L14 | 0.526937485 | 0.857206345 | Transport |
| G3V2F4 | TDP1 | Tyrosyl-DNA phosphodiesterase 1 | -0.069379171 | -1.53930982 | Cellular metabolism |
| G3V1Q4 | 39326 | Septin-7 | 0.045162201 | -0.096572876 | Other |
| E7EPM6 | ACSL1 | Long-chain-fatty-acid--CoA ligase 1 | 0.771936417 | 4.686363856 | Lipid metabolism |
| E7EPN9 | PRRC2C | Protein PRRC2C | 0.066116333 | -0.400510788 | Other |
| E7EPT4 | NDUFV2 | NADH dehydrogenase [ubiquinone] flavoprotein 2, mitochondrial | 0.669488271 | 1.19145902 | Cellular metabolism |
| E7EQ69 | NAA50 | N-alpha-acetyltransferase 50 | -0.203653336 | -1.250801086 | Cellular metabolism |
| E7EQB2 | LTF | Lactotransferrin | 0 | 0.479963303 | Immune response |
| H7C1W2 | IDH3G | Isocitrate dehydrogenase [NAD] subunit, mitochondrial | 0.449740728 | 0 | Cellular metabolism |
| E7EQR4 | EZR | Ezrin | -0.13133812 | -0.097119013 | Transport |
| E9PES6 | HMGB3 | High mobility group protein B3 | -0.244885763 | -1.719377518 | Cellular metabolism |
| E9PCS8 | MTA3 | Metastasis-associated protein MTA3 | 0 | 0.955667496 | Other |
| E7ERI8 | CLASP2 | CLIP-associating protein 2 | 0.401447296 | 0.796643575 | Other |
| E7ERK9 | EIF2B4 | Translation initiation factor eIF-2B subunit delta | 0.13114357 | -0.532245 | Cellular metabolism |
| E7ERS3 | ZC3H18 | Zinc finger CCCH domain-containing protein 18 | 0.20772171 | -0.263881048 | Other |
| E7ESC7 | MAEA | Macrophage erythroblast attacher | 0.172765732 | -0.780553182 | Other |
| E7ESL0 | MRPL22 | 39S ribosomal protein L22, mitochondrial | 0 | 0.193071365 | Cellular metabolism |
| E7ESY4 | MTA1 | Metastasis-associated protein MTA1 | 0.382158279 | 0.848740896 | Cellular metabolism |
| E7ETU5 | RBMS1 | RNA-binding motif, single-stranded-interacting protein 1 | -0.206154505 | 0 | Cellular metabolism |
| E7ETZ4 | BZW2 | Basic leucine zipper and W2 domain-containing protein 2 | -0.083056768 | -0.19043986 | Other |
| Q9NX74 | DUS2 | tRNA-dihydrouridine(20) synthase [NAD(P)+]-like | 0.026274999 | 0 | Other |
| E7EVA0 | MAP4 | Microtubule-associated protein | 0.202131907 | -0.059277852 | Other |
| E7EVC7 | ATG16L1 | Autophagy-related protein 16-1 | 0.322767258 | 0.379067739 | Transport |
| E7EW05 | SDAD1 | Protein SDA1 homolog | -0.491674423 | -4.975530624 | Transport |
| E9PID8 | CSTF2 | Cleavage stimulation factor subunit 2 | 0.258273443 | 0.461452484 | Cellular metabolism |
| E7EWW0 | C16orf62 | UPF0505 protein C16orf62 | -0.269095103 | 0 | Transport |
| E7EX17 | EIF4B | Eukaryotic translation initiation factor 4B | 0.126501083 | -0.336154938 | Cellular metabolism |
| E9PDF2 | OGDH | 2-oxoglutarate dehydrogenase, mitochondrial | -0.207885742 | 0.921470006 | Cellular metabolism |

| | | | | | |
|---|---|---|---|---|---|
| E9PCY5 | TOP2B | DNA topoisomerase 2 | 0 | -0.182435989 | Cellular metabolism |
| E9PDF6 | MYO1B | Unconventional myosin-Ib | -0.321621577 | -0.2107385 | Transport |
| E9PDJ9 | LIMCH1 | LIM and calponin homology domains-containing protein 1 | -0.322710037 | 1.652984619 | Other |
| E9PDP5 | ANKHD1 | Ankyrin repeat and KH domain-containing protein 1 | -0.702309926 | 0.500574748 | Other |
| E9PEB5 | FUBP1 | Far upstream element-binding protein 1 | 0.513690948 | 0.062671026 | Cellular metabolism |
| E9PER6 | PDPK1 | 3-phosphoinositide-dependent protein kinase 1 | -0.059879939 | -0.990136464 | Immune response |
| E9PF10 | NUP155 | Nuclear pore complex protein Nup155 | -0.33911705 | -0.021050771 | Transport |
| E9PFK9 | RABGEF1 | Rab5 GDP/GTP exchange factor | -0.145483653 | -0.981575648 | Transport |
| E9PFN5 | GSTK1 | Glutathione S-transferase kappa 1 | 0 | 0.554033915 | Cellular metabolism |
| P05067 | APP | Amyloid beta A4 protein | 0.062919617 | -2.410638809 | Immune response |
| E9PGC0 | RASA1 | Ras GTPase-activating protein 1 | -0.180796305 | 0.50568517 | Other |
| E9PGC8 | MAP1A | Microtubule-associated protein 1A | 0.185528437 | -1.663105011 | Other |
| Q04637-4 | EIF4G1 | Eukaryotic translation initiation factor 4 gamma 1 | 0.358539581 | -0.202217738 | Cellular metabolism |
| E9PGT3 | RPS6KA1 | Ribosomal protein S6 kinase | -0.506916046 | -1.914598465 | Immune response |
| E9PGT6 | COPS8 | COP9 signalosome complex subunit 8 | -0.114569346 | -0.840199788 | Cellular metabolism |
| E9PHA2 | NCAPH | Condensin complex subunit 2 | 0.125619888 | 0.103556315 | Cellular metabolism |
| P07858 | CTSB | Cathepsin B | -0.080212275 | -0.479059219 | Immune response |
| Q5T6K7 | NFYC | Nuclear transcription factor Y subunit gamma | 0.086126328 | -0.407660166 | Cellular metabolism |
| E9PIE4 | MTCH2 | Mitochondrial carrier homolog 2 | -0.019271215 | 1.075328191 | Transport |
| E9PIM6 | THY1 | Thy-1 membrane glycoprotein | 0 | 1.447271983 | Other |
| E9PJD7 | CYHR1 | Cysteine and histidine-rich protein 1 | -0.467774709 | 0 | Other |
| E9PJM3 | FBXO3 | F-box only protein 3 | -0.779186885 | 1.025935491 | Cellular metabolism |
| Q5H9R7-5 | PPP6R3 | Serine/threonine-protein phosphatase 6 regulatory subunit 3 | -0.203543981 | -1.017264048 | Other |
| E9PKP7 | UBTF | Nucleolar transcription factor 1 | -0.523223241 | 0.421234767 | Cellular metabolism |
| E9PKV2 | MRPL17 | 39S ribosomal protein L17, mitochondrial | -0.121855418 | 1.898513158 | Cellular metabolism |
| E9PL57 | NEDD8-MDP1 | | 0.212575912 | 1.403032303 | Other |
| E9PLI6 | EIF1AD | Probable RNA-binding protein EIF1AD | -0.03144455 | -0.556114833 | Other |
| P55786 | NPEPPS | Puromycin-sensitive aminopeptidase | -0.050294876 | -0.549858093 | Cellular metabolism |
| E9PLP8 | CSTF3 | Cleavage stimulation factor subunit 3 | 0 | 0.591427485 | Cellular metabolism |
| E9PM35 | ARRB1 | Beta-arrestin-1 | -0.195360819 | 0 | Transport |
| E9PM92 | C11orf58 | Small acidic protein | -0.195167542 | -0.554475149 | Other |
| E9PMH5 | BIRC2 | Baculoviral IAP repeat-containing protein 2 | -1.183395386 | 0 | Immune response |
| E9PMJ2 | FAM118B | Protein FAM118B | 0.323905309 | -0.718993505 | Other |

| | | | | | |
|---|---|---|---|---|---|
| E9PMQ6 | HSF1 | Heat shock factor protein 1 | 0.072470347 | 0.174412409 | Cellular metabolism |
| K4DIA7 | CD151 | Tetraspanin | 0 | 2.292458852 | Other |
| E9PMR6 | ARHGEF12 | Rho guanine nucleotide exchange factor 12 | -0.509786606 | 0 | Other |
| E9PN81 | RNASEH2C | Ribonuclease H2 subunit C | -0.515122096 | 0 | Cellular metabolism |
| E9PNF7 | PRCP | Lysosomal Pro-X carboxypeptidase | -0.268411001 | -0.26439476 | Other |
| E9PQ73 | FAT3 | Protocadherin Fat 3 | -0.684537252 | 0 | Other |
| E9PQP1 | NDUFV1 | NADH dehydrogenase [ubiquinone] flavoprotein 1, mitochondrial | 0 | -0.248763402 | Cellular metabolism |
| G3V1E1 | SMPD1 | Sphingomyelin phosphodiesterase | -1.358001073 | -0.589061101 | Lipid metabolism |
| E9PQY2 | PFDN4 | Prefoldin subunit 4 | -0.007132848 | 0 | Cellular metabolism |
| E9PR30 | FAU | 40S ribosomal protein S30 | 0.323453903 | 0.02343305 | Immune response |
| E9PS17 | SCYL1 | N-terminal kinase-like protein | 0.309815725 | 0.114111582 | Transport |
| F2Z2E2 | IQGAP3 | Ras GTPase-activating-like protein IQGAP3 | 0.052070618 | 0.278879801 | Cellular metabolism |
| F2Z2W7 | TRMT2A | tRNA (uracil-5-)-methyltransferase homolog A | 0.114692688 | 0 | Cellular metabolism |
| F5GWI4 | ADA | Adenosine deaminase | 0.031345367 | -1.886721929 | Transport |
| F5GWT4 | WNK1 | Serine/threonine-protein kinase WNK1 | 0.095685959 | -0.694043477 | Transport |
| F5H1S8 | MLEC | Malectin | 0 | -0.266150157 | Cellular metabolism |
| H0YFI1 | LAMTOR1 | Ragulator complex protein LAMTOR1 | 0 | 0.774424235 | Other |
| F5GX77 | TRMT112 | Multifunctional methyltransferase subunit TRM112-like protein | -0.040379842 | 0 | Cellular metabolism |
| F5GXX5 | DAD1 | Dolichyl-diphosphooligosaccharide--protein glycosyltransferase subunit DAD1 | 0 | 1.682388941 | Cellular metabolism |
| P09486 | SPARC | SPARC | 0.272542953 | -1.736279806 | Transport |
| J3QL14 | ATP6V0D1 | V-type proton ATPase subunit d 1 | -1.484182358 | 0 | Transport |
| H0YGN0 | CCDC53 | WASH complex subunit CCDC53 | -0.762893677 | 0 | Transport |
| F5H039 | GPHN | Gephyrin | 0.260225932 | -0.628402074 | Cellular metabolism |
| F5H0F9 | ANAPC5 | Anaphase-promoting complex subunit 5 | -0.56425031 | 0 | Cellular metabolism |
| F5H0R1 | TUT1 | Speckle targeted PIP5K1A-regulated poly(A) polymerase | -1.049064 | 0.307697932 | Cellular metabolism |
| F5H1S9 | PUS1 | tRNA pseudouridine synthase | 0.161050797 | -0.228065491 | Cellular metabolism |
| F5H8G1 | STARD10 | PCTP-like protein | -0.061276754 | -0.453076045 | Transport |
| F5H2U8 | HMGA2 | High mobility group protein HMGI-C | 0 | -0.567658106 | Cellular metabolism |
| V9GYY3 | MTHFD1 | C-1-tetrahydrofolate synthase, cytoplasmic | 0.22515742 | 0.000459671 | Cellular metabolism |
| F5H365 | SEC23A | Protein transport protein Sec23A | -0.040250142 | 0.382073085 | Transport |
| F5H3P3 | ARHGDIB | Rho GDP-dissociation inhibitor 2 | -0.540613174 | -1.740632375 | Immune response |
| F5H437 | CCND1 | G1/S-specific cyclin-D1 | -1.30790329 | -1.337553024 | Transport |
| F5H442 | TSG101 | Tumor susceptibility gene 101 protein | -0.14525795 | 0.595556895 | Transport |
| F8VY35 | NAP1L1 | Nucleosome assembly protein 1-like 1 | -0.062092463 | -0.823890686 | Cellular metabolism |
| F5H5I6 | GRSF1 | G-rich sequence factor 1 | -0.292636871 | 0.275438944 | Cellular metabolism |

| | | | | | |
|---|---|---|---|---|---|
| F5H5R8 | NAT1 | Arylamine N-acetyltransferase 1 | -1.228822072 | 0 | Cellular metabolism |
| F5H669 | CPSF7 | Cleavage and polyadenylation specificity factor subunit 7 | -0.277988434 | 0.03902626 | Cellular metabolism |
| F5H6E2 | MYO1C | Unconventional myosin-Ic | -0.595340729 | 0.830842336 | Immune response |
| F5H7P7 | PPWD1 | Peptidylprolyl isomerase domain and WD repeat-containing protein 1 | 0.503236771 | 0 | Cellular metabolism |
| F5H7W8 | C12orf43 | Uncharacterized protein C12orf43 | 0.117667516 | -0.20453008 | Other |
| F5H875 | CHMP1A | Charged multivesicular body protein 1a | 0.139549255 | 0 | Transport |
| F5H8D7 | XRCC1 | | -0.026147842 | -0.618082682 | Cellular metabolism |
| H0YBF6 | ASH2L | Set1/Ash2 histone methyltransferase complex subunit ASH2 | -1.020118078 | 0 | Cellular metabolism |
| F5H8H2 | MVK | Mevalonate kinase | -0.292919159 | -0.13000234 | Lipid metabolism |
| Q96FS4 | SIPA1 | Signal-induced proliferation-associated protein 1 | -0.327692668 | -1.857043584 | Other |
| F6TX30 | ASB6 | Ankyrin repeat and SOCS box protein 6 | -0.196287791 | 0 | Cellular metabolism |
| H0YB70 | HYI | Putative hydroxypyruvate isomerase | -0.195052465 | 0 | Other |
| F6VZ39 | RBM38 | RNA-binding protein 38 | 0.759234111 | 0 | Cellular metabolism |
| F8VNT9 | CD63 | Tetraspanin | 0.059060415 | 0.862645467 | Transport |
| F8VPD4 | CAD | CAD protein | 0.155561447 | -0.575588862 | Transport |
| F8VQE1 | LIMA1 | LIM domain and actin-binding protein 1 | 0.206234614 | 0.026602427 | Other |
| F8VRD2 | RACGAP1 | Rac GTPase-activating protein 1 | 0.874434153 | 0 | Transport |
| F8VVA7 | COPZ1 | Coatomer subunit zeta-1 | 0.183680216 | -0.18458875 | Transport |
| Q16527 | CSRP2 | Cysteine and glycine-rich protein 2 | 0.160737991 | -0.442596436 | Other |
| F8VXC8 | SMARCC2 | SWI/SNF complex subunit SMARCC2 | -0.585892359 | 0.296504974 | Cellular metabolism |
| F8WAK2 | GIT2 | ARF GTPase-activating protein GIT2 | 0.013859431 | -0.976620992 | Other |
| F8VYE8 | PPP1CC | Serine/threonine-protein phosphatase | -0.343509038 | 0.27565066 | Lipid metabolism |
| F8VZ44 | AAAS | Aladin | -0.661142349 | 4.173603694 | Transport |
| F8W1A4 | AK2 | Adenylate kinase 2, mitochondrial | -0.067688624 | -0.007858276 | Cellular metabolism |
| F8W6G1 | NRBP1 | | -0.444290797 | -1.754671097 | Other |
| F8W726 | UBAP2L | Ubiquitin-associated protein 2-like | -0.035100937 | -0.283596675 | Other |
| F8W785 | GOLIM4 | Golgi integral membrane protein 4 | -0.249306997 | -0.416522344 | Transport |
| Q15811 | ITSN1 | Intersectin-1 | -0.217483521 | -1.186197917 | Transport |
| F8W8D3 | SLBP | Histone RNA hairpin-binding protein | 0.017738342 | -1.243503571 | Transport |
| F8W8H5 | RAB24 | Ras-related protein Rab-24 | 0.972761154 | 0.529258728 | Transport |
| F8W930 | IGF2BP2 | Insulin-like growth factor 2 mRNA-binding protein 2 | 0.571752548 | -0.202750524 | Transport |
| F8W9F9 | WNK2 | Serine/threonine-protein kinase WNK2 | 0.837078094 | 1.517860413 | Cellular metabolism |
| F8W9X7 | CCDC93 | Coiled-coil domain-containing protein 93 | 0.367193222 | 0 | Transport |
| F8WA11 | CLASP1 | CLIP-associating protein 1 | 0.8040905 | -0.619174957 | Other |
| F8WA39 | MTMR1 | Myotubularin-related protein 1 | 1.863121668 | 2.822002411 | Lipid metabolism |

| | | | | | |
|---|---|---|---|---|---|
| F8WB06 | ATXN2 | Ataxin-2 | 0.085873286 | -0.267052968 | Transport |
| F8WBA3 | PRKD1 | Serine/threonine-protein kinase | 0.283605576 | 0 | Immune response |
| F8WCF6 | ARPC4-TTLL3 | Actin-related protein 2/3 complex subunit 4 | 0.216328303 | 0.218545914 | Immune response |
| F8WJN3 | CPSF6 | Cleavage and polyadenylation specificity factor subunit 6 | 0.961340586 | -0.547817866 | Cellular metabolism |
| G3V198 | NUP160 | Nuclear pore complex protein Nup160 | -0.517825445 | 0.083190282 | Transport |
| G3V1A6 | GSDMD | Gasdermin-D | -0.696429571 | -1.066513062 | Other |
| G3V256 | ZC3H14 | Zinc finger CCCH domain-containing protein 14 | 0.885084152 | 0.030274709 | Cellular metabolism |
| G3V424 | DNAL1 | Dynein light chain 1, axonemal | -0.725151062 | 0 | Other |
| G3V5N8 | ZFYVE1 | Zinc finger FYVE domain-containing protein 1 | -0.307631811 | 0 | Other |
| G3V5T9 | CDK2 | Cyclin-dependent kinase 2 | -0.332688649 | -0.907816569 | Transport |
| G3XAN4 | TRAM1 | Translocating chain-associated membrane protein 1 | 0 | 0.111689885 | Transport |
| G5E9A6 | USP11 | Ubiquitin carboxyl-terminal hydrolase | -1.239374797 | -1.546573003 | Cellular metabolism |
| G5E9C8 | SOS1 | Son of sevenless homolog 1 | -0.471532186 | 0 | Immune response |
| P35080-2 | PFN2 | Profilin-2 | -0.111935298 | -0.56820488 | Other |
| G5E9Z2 | CLPTM1L | Cleft lip and palate transmembrane protein 1-like protein | 0 | 1.180238088 | Other |
| G5EA06 | MRPS27 | 28S ribosomal protein S27, mitochondrial | -1.117667516 | 1.198927561 | Cellular metabolism |
| G5EA30 | CELF1 | CUGBP Elav-like family member 1 | 0.262289683 | 0 | Cellular metabolism |
| G5EA36 | CDC27 | Cell division cycle protein 27 homolog | -0.649814606 | 0 | Cellular metabolism |
| G8JL96 | PTPRS | Protein-tyrosine-phosphatase | -0.146799723 | -1.824396133 | Cellular metabolism |
| P31943 | HNRNPH1 | Heterogeneous nuclear ribonucleoprotein H | 0.460713069 | 0.16832606 | Cellular metabolism |
| G8JLD3 | ERC1 | ELKS/Rab6-interacting/CAST family member 1 | 0.63714091 | -0.910572052 | Transport |
| Q2L6G8 | CDSN | Corneodesmosin | 1.273302078 | 1.109039307 | Other |
| X6RLT1 | NELFCD | Negative elongation factor C/D | 0.00096639 | -1.270032247 | Cellular metabolism |
| H0Y398 | ZFAND3 | AN1-type zinc finger protein 3 | -0.583550771 | 0 | Other |
| H7C3X8 | SIK3 | Serine/threonine-protein kinase SIK3 | 0.454416911 | 0 | Cellular metabolism |
| H0Y512 | APMAP | Adipocyte plasma membrane-associated protein | 0.04069074 | 1.207760493 | Other |
| H0Y517 | CC2D1B | Coiled-coil and C2 domain-containing protein 1B | 0.304332733 | 0 | Cellular metabolism |
| H0Y5B4 | RPL36A | 60S ribosomal protein L36a | 0.136358897 | -0.043263753 | Transport |
| H0Y6D7 | FAM46A | Protein FAM46A | 0.413361867 | 0 | Other |
| H0Y8X4 | DNPH1 | 2-deoxynucleoside 5-phosphate N-hydrolase 1 | -0.040866216 | -0.006819407 | Cellular metabolism |
| H0Y938 | COPB2 | Coatomer subunit beta | 0.050832748 | -0.719561259 | Transport |
| H0Y993 | DEK | Protein DEK | -0.028570175 | -0.281211217 | Cellular metabolism |
| H0Y9Y3 | SYNPO2 | Synaptopodin-2 | 0 | 2.630775452 | Other |
| H0YAA3 | WDR41 | WD repeat-containing protein 41 | -0.280691783 | -1.084070206 | Other |
| H0YAN8 | ARHGEF10 | Rho guanine nucleotide exchange factor 10 | -0.209988276 | -1.950946172 | Other |

| | | | | | |
|---|---|---|---|---|---|
| P11940 | PABPC1 | Polyadenylate-binding protein 1 | -0.055672963 | 0.223363241 | Cellular metabolism |
| H0YAZ3 | SORBS3 | Vinexin | 0.166339874 | 0 | Other |
| H0YBV6 | MAK16 | Protein MAK16 homolog | 1.610655467 | 2.18540446 | Other |
| P16070 | CD44 | CD44 antigen | -0.879940669 | 0.192935308 | Other |
| H0YD93 | RUFY2 | RUN and FYVE domain-containing protein 2 | -0.315828959 | 0 | Other |
| H0YDJ3 | PRPF4B | Serine/threonine-protein kinase PRP4 homolog | 0.714923223 | 0.153413773 | Cellular metabolism |
| H0YDT6 | EIF3F | Eukaryotic translation initiation factor 3 subunit F | 0.056728363 | -0.010193507 | Cellular metabolism |
| P53041 | PPP5C | Serine/threonine-protein phosphatase 5 | 0.178417842 | -0.178492228 | Cellular metabolism |
| H0YEB6 | SSSCA1 | Sjoegren syndrome/scleroderma autoantigen 1 | -0.414715449 | 0 | Other |
| H0YFD6 | HADHA | Trifunctional enzyme subunit alpha, mitochondrial | 0.950234095 | 0.879761378 | Lipid metabolism |
| H0YG72 | ITFG2 | Integrin-alpha FG-GAP repeat-containing protein 2 | -0.448154449 | 0 | Other |
| H0YGR4 | REXO2 | Oligoribonuclease, mitochondrial | 0.069273631 | -0.530838013 | Cellular metabolism |
| H0YGS7 | TRIM65 | Tripartite motif-containing protein 65 | -0.0928243 | 0 | Other |
| H0YHA7 | RPL18 | 60S ribosomal protein L18 | 0.343808492 | 0.538896561 | Transport |
| H0YHS6 | YARS2 | Tyrosine--tRNA ligase | 0 | 0.566095352 | Cellular metabolism |
| H0YIE9 | FBXO21 | F-box only protein 21 | -0.569018682 | 0 | Cellular metabolism |
| H0YJE4 | DHRS7 | Dehydrogenase/reductase SDR family member 7 | 0.073485057 | 0.074925741 | Other |
| H0YJ75 | PPP2R5C | Serine/threonine-protein phosphatase 2A 56 kDa regulatory subunit gamma isoform | -0.347348531 | -1.008394241 | Cellular metabolism |
| H0YJ92 | MNAT1 | CDK-activating kinase assembly factor MAT1 | -0.589351018 | -1.476407369 | Cellular metabolism |
| H0YLP6 | RPL28 | 60S ribosomal protein L28 | 0.212053935 | 0.333504359 | Transport |
| H0YKX0 | RNF31 | E3 ubiquitin-protein ligase RNF31 | -0.128602346 | 1.036117554 | Cellular metabolism |
| H0YL72 | IDH3A | Isocitrate dehydrogenase [NAD] subunit alpha, mitochondrial | -0.454575857 | -0.003870646 | Cellular metabolism |
| H0YM23 | ANKRD17 | Ankyrin repeat domain-containing protein 17 | 0.15988032 | -0.469521205 | Immune response |
| H0YME5 | EIF2AK4 | Eukaryotic translation initiation factor 2-alpha kinase 4 | 0.08906428 | -0.953293482 | Cellular metabolism |
| H0YMV8 | RPS27L | 40S ribosomal protein S27 | 0.221861521 | -0.256205877 | Cellular metabolism |
| H0YMW2 | AKAP13 | A-kinase anchor protein 13 | 1.109835307 | 0.095322291 | Transport |
| H0YN26 | ANP32A | Acidic leucine-rich nuclear phosphoprotein 32 family member A | 0.253571828 | -0.472867966 | Transport |
| H0YN81 | WDR61 | WD repeat-containing protein 61 | 0.890476863 | 1.111715317 | Cellular metabolism |
| H0YNG3 | SEC11A | Signal peptidase complex catalytic subunit SEC11 | 1.233596166 | 1.117139181 | Transport |
| Q5SZE1 | CERS2 | Ceramide synthase 2 | 0.701503754 | -0.00200208 | Lipid metabolism |
| H3BLV9 | SRPK1 | SRSF protein kinase 1 | 0.492791494 | 0.858887355 | Immune response |
| H3BM14 | NUB1 | NEDD8 ultimate buster 1 | -0.045435588 | 0 | Cellular metabolism |
| H3BM38 | SNAP23 | Synaptosomal-associated protein | -2.152082443 | 0.131717682 | Transport |
| H3BV80 | RNPS1 | RNA-binding protein with serine-rich domain 1 | 0.005140305 | -0.334924062 | Transport |
| H3BN98 | RPS15A | | 0 | 0.030162175 | Transport |
| H3BNC9 | RPS17 | 40S ribosomal protein S17 | 0.100276311 | -0.019927979 | Transport |

| | | | | | |
|---|---|---|---|---|---|
| H3BNX8 | COX5A | Cytochrome c oxidase subunit 5A, mitochondrial | -0.17642657 | -0.492479324 | Cellular metabolism |
| H3BP20 | HEXA | Beta-hexosaminidase | -0.3211085 | -0.374383291 | Lipid metabolism |
| H3BP71 | RNF40 | E3 ubiquitin-protein ligase BRE1B | 0.459646861 | -0.180799484 | Cellular metabolism |
| H3BQK9 | MACF1 | Microtubule-actin cross-linking factor 1, isoforms 1/2/3/5 | 0.56034406 | -1.541978836 | Transport |
| H3BPK3 | HAGH | Hydroxyacylglutathione hydrolase, mitochondrial | -0.118589401 | -1.447572708 | Cellular metabolism |
| H3BUD2 | DCTN5 | Dynactin subunit 5 | -0.069730759 | 0 | Other |
| H3BRD2 | GFER | Sulfhydryl oxidase | 0.0151577 | -1.385700862 | Transport |
| H3BRF9 | ZFYVE19 | Abscission/NoCut checkpoint regulator | -0.123013179 | -1.243423462 | Other |
| O14562 | UBFD1 | Ubiquitin domain-containing protein UBFD1 | -0.040273666 | -0.240823746 | Other |
| H3BRQ0 | PPCDC | Phosphopantothenoylcysteine decarboxylase | 0.281194687 | 0 | Cellular metabolism |
| H3BS72 | HACD3 | Very-long-chain (3R)-3-hydroxyacyl-CoA dehydratase 3 | 0.780527751 | 1.75652504 | Lipid metabolism |
| H3BSW6 | CTU2 | Cytoplasmic tRNA 2-thiolation protein 2 | 0.071879705 | -0.430453618 | Cellular metabolism |
| H3BUB4 | SNUPN | Snurportin-1 | 0.151344935 | -1.387286504 | Transport |
| H3BUU8 | VAC14 | | -0.470863342 | 2.729377111 | Other |
| H6UMI1 | GABARAP | Gamma-aminobutyric acid receptor-associated protein | -1.475170135 | 0 | Transport |
| H7C2Z6 | GCA | Grancalcin | 0.014293035 | 0.067841848 | Other |
| H7BXY3 | DHX30 | Putative ATP-dependent RNA helicase DHX30 | -0.140371323 | 0.807036082 | Cellular metabolism |
| Q5JRG1 | NUPL1 | Nucleoporin p58/p45 | 0.130659739 | 0 | Transport |
| H7BYT1 | CSNK1D | Casein kinase I isoform delta | -0.069298426 | -0.582867304 | Transport |
| H7BZW6 | SAP18 | Histone deacetylase complex subunit SAP18 | 0.346354167 | 0.430176417 | Cellular metabolism |
| H7C096 | BNIP2 | BCL2/adenovirus E1B 19 kDa protein-interacting protein 2 | -0.396219889 | 0 | Transport |
| H7C0H8 | TMEM214 | Transmembrane protein 214 | 0 | 0.123342514 | Other |
| H7C144 | ACTN4 | | -0.299730937 | 0 | Other |
| H7C170 | WBSCR22 | Probable 18S rRNA (guanine-N(7))-methyltransferase | -0.037761052 | 0 | Cellular metabolism |
| H7C2Q8 | EBNA1BP2 | Probable rRNA-processing protein EBP2 | 1.005259196 | 0.218102137 | Other |
| H7C3G1 | ECT2 | Protein ECT2 | -0.663885752 | 0.232320786 | Transport |
| H7C3X5 | STX5 | Syntaxin-5 | -0.165136337 | 0 | Transport |
| H7C4F6 | KPNA4 | Importin subunit alpha-3 | -0.054777781 | -0.016383489 | Transport |
| H7C5E4 | XRN1 | 5-3 exoribonuclease 1 | 2.958938599 | 0 | Cellular metabolism |
| I3L0H8 | DDX19A | ATP-dependent RNA helicase DDX19A | 0.154669444 | -0.276203791 | Transport |
| I3L0N3 | NSF | Vesicle-fusing ATPase | -0.215316137 | 1.471110026 | Transport |
| J3KSI4 | MPDU1 | Mannose-P-dolichol utilization defect 1 protein | 0.961945216 | 2.078178406 | Transport |
| I3L1Y9 | FLYWCH2 | FLYWCH family member 2 | 0.075706482 | 0.589768728 | Other |
| I3L2B2 | EPN2 | Epsin-2 | 0.084486008 | 0.159461975 | Transport |
| I3L2C7 | GEMIN4 | Gem-associated protein 4 | 1.097578684 | 0 | Cellular metabolism |

| | | | | | |
|---|---|---|---|---|---|
| I3L2Z9 | NEURL4 | Neuralized-like protein 4 | -0.614729563 | 0 | Other |
| I3L4G0 | TOM1L1 | TOM1-like protein 1 | 0.214765549 | 0.280390422 | Transport |
| I3L4A1 | CHMP6 | Charged multivesicular body protein 6 | 0.927029292 | 1.415712992 | Transport |
| Q9UQB8 | BAIAP2 | Brain-specific angiogenesis inhibitor 1-associated protein 2 | 0.355950038 | 0 | Immune response |
| I3L4V0 | NAA38 | N-alpha-acetyltransferase 38, NatC auxiliary subunit | 0.292315165 | 0 | Other |
| J3KN16 | KIAA0368 | Proteasome-associated protein ECM29 homolog | 0.067479451 | 0.44966507 | Cellular metabolism |
| J3KNF4 | CCS | Copper chaperone for superoxide dismutase | -0.084229787 | -1.016387304 | Transport |
| J3KNN3 | PHKG2 | Phosphorylase b kinase gamma catalytic chain, liver/testis isoform | 0.391787847 | 0 | Cellular metabolism |
| J3QL05 | SRSF2 | Serine/arginine-rich splicing factor 2 | 0.304337819 | -0.709125519 | Transport |
| Q5QJE6 | DNTTIP2 | Deoxynucleotidyltransferase terminal-interacting protein 2 | 0 | -0.792191188 | Cellular metabolism |
| J3KPD3 | RBM7 | RNA-binding protein 7 | 0.577687581 | -0.52150472 | Other |
| J3KPJ3 | CAMKK1 | Calcium/calmodulin-dependent protein kinase kinase 1 | -0.384607951 | 0 | Other |
| J3KPP4 | LUC7L3 | Luc7-like protein 3 | 0.802335103 | -0.049489975 | Cellular metabolism |
| V9GYN7 | DNAJB12 | DnaJ homolog subfamily B member 12 | 0 | 1.92663002 | Other |
| J3KQ32 | OLA1 | Obg-like ATPase 1 | -0.061770757 | -0.718179067 | Other |
| J3KQ48 | PTRH2 | Peptidyl-tRNA hydrolase 2, mitochondrial | 1.161611557 | 2.487872442 | Other |
| J3KQJ1 | SUMF2 | Sulfatase-modifying factor 2 | 0.214913686 | -0.534633636 | Cellular metabolism |
| J3KR35 | CCDC12 | Coiled-coil domain-containing protein 12 | 0.2057031 | 0 | Other |
| J3KR44 | OTUB1 | Ubiquitin thioesterase OTUB1 | -0.145142237 | -0.350699743 | Cellular metabolism |
| Q9BTW9 | TBCD | Tubulin-specific chaperone D | -0.085797628 | -0.558394114 | Cellular metabolism |
| J3KS22 | DCXR | L-xylulose reductase | -0.149729411 | 0.002794902 | Cellular metabolism |
| J3KTA1 | FBXL20 | F-box/LRR-repeat protein 20 | 1.024988174 | 1.296203613 | Other |
| J3KTA4 | DDX5 | Probable ATP-dependent RNA helicase DDX5 | 0.341914495 | 0.473829269 | Cellular metabolism |
| J3KTL2 | SRSF1 | Serine/arginine-rich splicing factor 1 | 0.187283834 | -0.473211924 | Transport |
| J3QQV2 | SLC16A3 | Monocarboxylate transporter 4 | 0 | 3.589952469 | Transport |
| J3QK89 | CHERP | Calcium homeostasis endoplasmic reticulum protein | 0.0451533 | 0.047112147 | Transport |
| J3QKW2 | MRPS7 | 28S ribosomal protein S7, mitochondrial | 0 | -2.213933309 | Cellular metabolism |
| J3QL71 | SCRN2 | Secernin-2 | -0.108778636 | -1.75996081 | Transport |
| J3QLE5 | SNRPN | Small nuclear ribonucleoprotein-associated proteins B and B | 0.225699743 | -0.338668187 | Cellular metabolism |
| J3QLW7 | NIP7 | 60S ribosome subunit biogenesis protein NIP7 homolog | 0.359897614 | 0.931330999 | Other |
| J3QQJ0 | SAP30BP | SAP30-binding protein | -0.232484818 | -0.593030294 | Cellular metabolism |
| J3QQJ5 | TRAPPC8 | Trafficking protein particle complex subunit 8 | 0.914422989 | 0 | Transport |
| J3QQY2 | TMCO1 | Transmembrane and coiled-coil domain-containing protein 1 | 0 | 1.011351903 | Other |
| J3QR07 | YTHDC1 | YTH domain-containing protein 1 | 0.736567179 | 2.141426722 | Other |
| J3QRG6 | CDKN2A | Cyclin-dependent kinase inhibitor 2A | -0.193531036 | -0.640845617 | Cellular metabolism |
| J3QRS3 | MYL12A | Myosin regulatory light chain 12A | -0.131540934 | -0.310017904 | Transport |

| | | | | | |
|---|---|---|---|---|---|
| J3QRS9 | ZNF207 | BUB3-interacting and GLEBS motif-containing protein ZNF207 | 0.094821294 | 1.834182103 | Other |
| J3QRU1 | YES1 | Non-specific protein-tyrosine kinase | 0 | 1.444013596 | Immune response |
| J3QSV6 | RSL1D1 | Ribosomal L1 domain-containing protein 1 | 0.096871694 | 0.703215281 | Other |
| J3QT28 | BUB3 | Mitotic checkpoint protein BUB3 | 0.084826787 | -0.031482061 | Cellular metabolism |
| J3QTA2 | BAG1 | BAG family molecular chaperone regulator 1 | 0.120527903 | 0 | Cellular metabolism |
| J9JIC5 | C17orf75 | Protein Njmu-R1 | 0.104441961 | 0 | Other |
| K4DI92 | RWDD4 | RWD domain-containing protein 4 | -0.167105357 | -0.121344248 | Other |
| K7EIE8 | MBD3 | Methyl-CpG-binding domain protein 3 | -0.136504491 | 0 | Cellular metabolism |
| Q13485 | SMAD4 | Mothers against decapentaplegic homolog 4 | -0.166933695 | 0 | Cellular metabolism |
| Q9BV68 | RNF126 | E3 ubiquitin-protein ligase RNF126 | -0.100703557 | -2.01084582 | Transport |
| K7EJE8 | LONP1 | Lon protease homolog, mitochondrial | 0.156924566 | 0.102307002 | Cellular metabolism |
| K7EQ55 | DAZAP1 | DAZ-associated protein 1 | 0.387324015 | 0.030834198 | Other |
| O60437 | PPL | Periplakin | 0 | 0.754002889 | Other |
| P36954 | POLR2I | DNA-directed RNA polymerase II subunit RPB9 | 0.145799637 | 0 | Cellular metabolism |
| Q96S99 | PLEKHF1 | Pleckstrin homology domain-containing family F member 1 | -0.016703288 | 0 | Other |
| K7ELC2 | RPS15 | 40S ribosomal protein S15 | 0.135332743 | 1.109437307 | Transport |
| K7ELG9 | LSM12 | Protein LSM12 homolog | 0.342496236 | -0.668837865 | Other |
| K7ELL7 | PRKCSH | Glucosidase 2 subunit beta | 0.251246134 | -0.249795278 | Immune response |
| K7ELQ9 | TMEM205 | Transmembrane protein 205 | 0 | 4.123977025 | Other |
| P41567 | EIF1 | Eukaryotic translation initiation factor 1 | 0.032076518 | -0.070870717 | Other |
| K7EMC9 | WBP2 | WW domain-binding protein 2 | -0.406681061 | -1.899916967 | Other |
| K7ENF0 | NARS | | 0.155558904 | 0 | Other |
| K7ENR6 | PSMG2 | Proteasome assembly chaperone 2 | 0.010595322 | -0.160931905 | Other |
| K7EQ03 | RBM42 | RNA-binding protein 42 | -0.083873749 | 0.013059616 | Other |
| K7EQL6 | USP32 | Ubiquitin carboxyl-terminal hydrolase | -0.295458476 | -2.553045909 | Cellular metabolism |
| K7ER15 | HDHD2 | Haloacid dehalogenase-like hydrolase domain-containing protein 2 | 0.133158366 | -1.242331823 | Other |
| K7ERD7 | POP4 | Ribonuclease P protein subunit p29 | -0.833054225 | -0.081997554 | Cellular metabolism |
| K7ES61 | MRPL4 | 39S ribosomal protein L4, mitochondrial | -0.030143102 | 1.219299316 | Cellular metabolism |
| K7ES84 | EEF2KMT | Protein-lysine N-methyltransferase EEF2KMT | -1.334908168 | 0 | Cellular metabolism |
| O95571 | ETHE1 | Persulfide dioxygenase ETHE1, mitochondrial | -0.00852712 | -0.172110875 | Cellular metabolism |
| M0QXF9 | BCAT2 | Branched-chain-amino-acid aminotransferase | 0.496496836 | -1.72650528 | Cellular metabolism |
| M0QXZ5 | ZNF428 | Zinc finger protein 428 | -0.176615397 | -0.061913172 | Other |
| M0QY97 | ZC3H4 | Zinc finger CCCH domain-containing protein 4 | -0.030168533 | 0 | Other |
| M0QYU8 | KLK7 | Kallikrein-7 | -1.242010752 | -0.713452657 | Other |
| M0R0N4 | AP2S1 | AP-2 complex subunit sigma | -0.582042058 | -0.159301122 | Transport |

| | | | | | |
|---|---|---|---|---|---|
| M0QZD8 | | Putative uncharacterized protein LOC400499 | -0.187356313 | -0.02736028 | #N/A |
| M0QZR4 | ARHGEF1 | Rho guanine nucleotide exchange factor 1 | 0.185569763 | -0.654190699 | Other |
| M0R0C6 | KHSRP | Far upstream element-binding protein 2 | 0.612661362 | 0.134335836 | Transport |
| M0R0F0 | RPS5 | 40S ribosomal protein S5 | 0.079415639 | 0.174512227 | Transport |
| M0R300 | MYO9B | Unconventional myosin-IXb | -0.558462779 | 0 | Other |
| M0R117 | RPL18A | 60S ribosomal protein L18a | 0.396748861 | 0.408236186 | Transport |
| M0R208 | CLPP | ATP-dependent Clp protease proteolytic subunit | 0.232316971 | 0 | Other |
| M0R2B7 | POLD1 | DNA polymerase | 0.17379125 | -0.477490743 | Cellular metabolism |
| O00115 | DNASE2 | Deoxyribonuclease-2-alpha | -0.022929509 | -1.861526489 | Cellular metabolism |
| O00148 | DDX39A | ATP-dependent RNA helicase DDX39A | 0.114692688 | -0.355599721 | Transport |
| O00151 | PDLIM1 | PDZ and LIM domain protein 1 | 0.015736898 | -0.478305181 | Other |
| O00154-4 | ACOT7 | Cytosolic acyl coenzyme A thioester hydrolase | -0.082042058 | -1.039178848 | Lipid metabolism |
| O00170 | AIP | AH receptor-interacting protein | -0.106085459 | -0.650116603 | Transport |
| O00178 | GTPBP1 | GTP-binding protein 1 | 0.105190277 | 0.399935404 | Immune response |
| O00203 | AP3B1 | AP-3 complex subunit beta-1 | 0.031003952 | -0.350311915 | Transport |
| O00204-2 | SULT2B1 | Sulfotransferase family cytosolic 2B member 1 | 0 | 0.203802109 | Lipid metabolism |
| O00231 | PSMD11 | 26S proteasome non-ATPase regulatory subunit 11 | -0.121909459 | -0.658651352 | Immune response |
| O00232 | PSMD12 | 26S proteasome non-ATPase regulatory subunit 12 | -0.10283788 | 0.332152685 | Immune response |
| O00233 | PSMD9 | 26S proteasome non-ATPase regulatory subunit 9 | -0.138083776 | -0.892834345 | Immune response |
| O00264 | PGRMC1 | Membrane-associated progesterone receptor component 1 | 0.780009588 | 1.213599523 | Other |
| O00267-2 | SUPT5H | Transcription elongation factor SPT5 | -0.040339788 | -1.757763545 | Cellular metabolism |
| O00273 | DFFA | DNA fragmentation factor subunit alpha | -0.348006566 | -1.345989227 | Cellular metabolism |
| O00273-2 | DFFA | DNA fragmentation factor subunit alpha | -0.286417007 | -0.288469315 | Cellular metabolism |
| O00291-3 | HIP1 | Huntingtin-interacting protein 1 | 0 | -1.640300751 | Transport |
| O00299 | CLIC1 | Chloride intracellular channel protein 1 | -0.191020966 | -1.035905838 | Transport |
| O00391 | QSOX1 | Sulfhydryl oxidase 1 | 0 | 0.547871272 | Cellular metabolism |
| O00399 | DCTN6 | Dynactin subunit 6 | -0.079734166 | 0 | Other |
| O00410 | IPO5 | Importin-5 | -0.154778163 | -0.366820017 | Transport |
| O00418 | EEF2K | Eukaryotic elongation factor 2 kinase | -0.600905101 | 0 | Cellular metabolism |
| O00425 | IGF2BP3 | Insulin-like growth factor 2 mRNA-binding protein 3 | 0.88417689 | 1.141710281 | Transport |
| O00429 | DNM1L | Dynamin-1-like protein | 0.098133087 | -1.732849121 | Transport |
| O00442 | RTCA | RNA 3-terminal phosphate cyclase | -0.320854823 | -0.390528997 | Cellular metabolism |
| O00462 | MANBA | Beta-mannosidase | -0.966130575 | -1.861592611 | Cellular metabolism |
| O00469-2 | PLOD2 | Procollagen-lysine,2-oxoglutarate 5-dioxygenase 2 | 0.093161901 | -0.196553548 | Cellular metabolism |
| O00487 | PSMD14 | 26S proteasome non-ATPase regulatory subunit 14 | 0.125970205 | 1.087580363 | Immune response |

| | | | | | |
|---|---|---|---|---|---|
| O00499-9 | BIN1 | Myc box-dependent-interacting protein 1 | -0.01485316 | -1.779427846 | Transport |
| O00505 | KPNA3 | Importin subunit alpha-4 | -0.005083084 | 0.07190005 | Transport |
| O00560 | SDCBP | Syntenin-1 | -0.410405477 | 0.528262456 | Transport |
| O00567 | NOP56 | Nucleolar protein 56 | 0.181198756 | 0.755090078 | Cellular metabolism |
| O00592-2 | PODXL | Podocalyxin | -1.129264832 | 0.826770147 | Other |
| O00622 | CYR61 | Protein CYR61 | -1.764709473 | 0 | Cellular metabolism |
| O00625 | PIR | Pirin | -0.15232722 | -1.552216848 | Cellular metabolism |
| O00743 | PPP6C | Serine/threonine-protein phosphatase 6 catalytic subunit | -0.287308375 | -1.596633275 | Cellular metabolism |
| O00754-2 | MAN2B1 | Lysosomal alpha-mannosidase | 0.249106725 | -0.334619522 | Cellular metabolism |
| O00762 | UBE2C | Ubiquitin-conjugating enzyme E2 C | -0.845656713 | 0.173362096 | Cellular metabolism |
| O00764 | PDXK | Pyridoxal kinase | -0.198558807 | -1.223503749 | Cellular metabolism |
| O14530 | TXNDC9 | Thioredoxin domain-containing protein 9 | -0.595668157 | -0.804528554 | Other |
| Q5T0Q6 | DPYSL4 | Dihydropyrimidinase-related protein 4 | -1.259992599 | -2.689327876 | Other |
| O14545 | TRAFD1 | TRAF-type zinc finger domain-containing protein 1 | -0.056915283 | 0 | Other |
| O14579 | COPE | Coatomer subunit epsilon | -0.25188446 | 0.670739492 | Transport |
| O14617 | AP3D1 | AP-3 complex subunit delta-1 | -0.029712677 | -1.007827123 | Transport |
| O14641 | DVL2 | Segment polarity protein dishevelled homolog DVL-2 | 0.018863678 | -1.469900131 | Transport |
| O14646-2 | CHD1 | Chromodomain-helicase-DNA-binding protein 1 | -0.51155599 | -0.325709025 | Lipid metabolism |
| O14656 | TOR1A | Torsin-1A | 0 | 0.616768519 | Transport |
| O14657 | TOR1B | Torsin-1B | 0.1641922 | 0 | Cellular metabolism |
| O14732-2 | IMPA2 | Inositol monophosphatase 2 | 0 | 1.543703079 | Lipid metabolism |
| O14737 | PDCD5 | Programmed cell death protein 5 | -0.733186086 | 0.021155039 | Other |
| O14744 | PRMT5 | Protein arginine N-methyltransferase 5 | 0.21248436 | -1.549578349 | Cellular metabolism |
| O14745 | SLC9A3R1 | Na(+)/H(+) exchange regulatory cofactor NHE-RF1 | -0.235749563 | -0.056154251 | Transport |
| O14772 | FPGT | Fucose-1-phosphate guanylyltransferase | -0.59192721 | -2.173070908 | Cellular metabolism |
| O14777 | NDC80 | Kinetochore protein NDC80 homolog | -0.53437678 | -0.927131017 | Other |
| O14787-2 | TNPO2 | Transportin-2 | -0.209652583 | 0.066230138 | Transport |
| O14818 | PSMA7 | Proteasome subunit alpha type-7 | 0.183973312 | 0.094950994 | Immune response |
| O14828 | SCAMP3 | Secretory carrier-associated membrane protein 3 | 0 | 1.637799581 | Transport |
| O14907 | TAX1BP3 | Tax1-binding protein 3 | -0.038764318 | -1.027009328 | Other |
| O14908 | GIPC1 | PDZ domain-containing protein GIPC1 | -0.286314011 | -1.012091319 | Transport |
| O14929 | HAT1 | Histone acetyltransferase type B catalytic subunit | -0.221114477 | -1.264432907 | Cellular metabolism |
| O14933 | UBE2L6 | Ubiquitin/ISG15-conjugating enzyme E2 L6 | 0.106178284 | 0 | Immune response |
| O14949 | UQCRQ | Cytochrome b-c1 complex subunit 8 | 0 | 4.715264638 | Cellular metabolism |

| | | | | | |
|---|---|---|---|---|---|
| O14967 | CLGN | Calmegin | 0 | 0.532146454 | Cellular metabolism |
| O14974-3 | PPP1R12A | Protein phosphatase 1 regulatory subunit 12A | 0.224126816 | -0.527450562 | Cellular metabolism |
| O14976-2 | GAK | Cyclin-G-associated kinase | 0.339806239 | 0.351661046 | Transport |
| O14979-3 | HNRNPDL | Heterogeneous nuclear ribonucleoprotein D-like | 0.544867833 | 0.200425466 | Cellular metabolism |
| O14980 | XPO1 | Exportin-1 | 0.017319997 | 0.090177536 | Transport |
| O15018-2 | PDZD2 | PDZ domain-containing protein 2 | -0.857379913 | -1.825726191 | Other |
| O15027-2 | SEC16A | Protein transport protein Sec16A | 0.171262105 | -1.740266164 | Transport |
| O15042-2 | U2SURP | U2 snRNP-associated SURP motif-containing protein | 1.483695984 | 0.118798574 | Cellular metabolism |
| O15066 | KIF3B | Kinesin-like protein KIF3B | 0.044070562 | -2.27197965 | Transport |
| O15067 | PFAS | Phosphoribosylformylglycinamidine synthase | 0.061170578 | -0.3018411 | Cellular metabolism |
| O15084 | ANKRD28 | Serine/threonine-protein phosphatase 6 regulatory ankyrin repeat subunit A | -0.304137548 | -0.239106496 | Other |
| O15111 | CHUK | Inhibitor of nuclear factor kappa-B kinase subunit alpha | -0.032936732 | 0 | Immune response |
| O15116 | LSM1 | U6 snRNA-associated Sm-like protein LSm1 | 0.115582784 | 0 | Cellular metabolism |
| O15118 | NPC1 | Niemann-Pick C1 protein | 0 | -0.53136762 | Lipid metabolism |
| O15143 | ARPC1B | Actin-related protein 2/3 complex subunit 1B | 0.174073537 | -0.377305984 | Other |
| O15144 | ARPC2 | Actin-related protein 2/3 complex subunit 2 | 0.12359492 | -0.018963496 | Immune response |
| O15145 | ARPC3 | Actin-related protein 2/3 complex subunit 3 | -0.327071508 | 0.238136292 | Immune response |
| O15173-2 | PGRMC2 | Membrane-associated progesterone receptor component 2 | 0.524036407 | 1.98764801 | Other |
| O15212 | PFDN6 | Prefoldin subunit 6 | -0.715072632 | -0.628551483 | Cellular metabolism |
| O15231-3 | ZNF185 | | -0.031129201 | -1.754585902 | Other |
| O15231-6 | ZNF185 | | 0.41059049 | -0.604283651 | Other |
| Q5T8U5 | SURF4 | Surfeit locus protein 4 | 0.046288808 | 0.65034612 | Other |
| O15269 | SPTLC1 | Serine palmitoyltransferase 1 | 0 | 1.066480001 | Lipid metabolism |
| O15294 | OGT | UDP-N-acetylglucosamine--peptide N-acetylglucosaminyltransferase 110 kDa subunit | 0.012865067 | -1.100570043 | Cellular metabolism |
| O15305 | PMM2 | Phosphomannomutase 2 | -0.013992945 | -0.266726176 | Cellular metabolism |
| O15355 | PPM1G | Protein phosphatase 1G | -0.046892802 | -0.44858106 | Cellular metabolism |
| O15357 | INPPL1 | Phosphatidylinositol 3,4,5-trisphosphate 5-phosphatase 2 | -0.06310463 | -0.430150986 | Lipid metabolism |
| O15371-3 | EIF3D | Eukaryotic translation initiation factor 3 subunit D | 0.229426066 | -0.838704427 | Cellular metabolism |
| O15397 | IPO8 | Importin-8 | -0.384506861 | -1.785415649 | Transport |
| O15400-2 | STX7 | Syntaxin-7 | 0.228542964 | 0.472321192 | Transport |
| O15405-2 | TOX3 | TOX high mobility group box family member 3 | -0.341945012 | 0 | Cellular metabolism |
| O15446 | CD3EAP | DNA-directed RNA polymerase I subunit RPA34 | 1.741779327 | -0.726731618 | Cellular metabolism |
| O15460-2 | P4HA2 | Prolyl 4-hydroxylase subunit alpha-2 | 0.193617503 | -0.895467122 | Cellular metabolism |
| O15511 | ARPC5 | Actin-related protein 2/3 complex subunit 5 | 0.284427007 | -0.064212799 | Immune response |
| O15541 | RNF113A | RING finger protein 113A | 0.039637884 | -1.229276657 | Other |

| | | | | | |
|---|---|---|---|---|---|
| O43143 | DHX15 | Pre-mRNA-splicing factor ATP-dependent RNA helicase DHX15 | 0.133720398 | -0.233343124 | Cellular metabolism |
| O43148 | RNMT | mRNA cap guanine-N7 methyltransferase | -0.058844884 | -0.358231227 | Cellular metabolism |
| O43149 | ZZEF1 | Zinc finger ZZ-type and EF-hand domain-containing protein 1 | 0.528746923 | 0 | Other |
| O43150 | ASAP2 | Arf-GAP with SH3 domain, ANK repeat and PH domain-containing protein 2 | -0.011423747 | 0 | Other |
| O43156 | TTI1 | TELO2-interacting protein 1 homolog | 0.202487946 | -0.590701421 | Other |
| O43172-2 | PRPF4 | U4/U6 small nuclear ribonucleoprotein Prp4 | 0.325491587 | -0.221502304 | Cellular metabolism |
| O43237 | DYNC1LI2 | Cytoplasmic dynein 1 light intermediate chain 2 | -0.053072611 | -0.499539693 | Transport |
| O43242 | PSMD3 | 26S proteasome non-ATPase regulatory subunit 3 | 0.040788651 | -0.234022141 | Immune response |
| O43252 | PAPSS1 | Bifunctional 3-phosphoadenosine 5-phosphosulfate synthase 1 | -0.119689306 | -1.23505338 | Cellular metabolism |
| O43264 | ZW10 | Centromere/kinetochore protein zw10 homolog | -0.306575775 | 0.879217784 | Transport |
| O43290 | SART1 | U4/U6.U5 tri-snRNP-associated protein 1 | -0.174591064 | -0.098258336 | Cellular metabolism |
| O43294 | TGFB1I1 | Transforming growth factor beta-1-induced transcript 1 protein | -0.479557037 | -1.330478032 | Cellular metabolism |
| O43318-2 | MAP3K7 | Mitogen-activated protein kinase kinase kinase 7 | -0.095856984 | -0.903472265 | Immune response |
| O43324 | EEF1E1 | Eukaryotic translation elongation factor 1 epsilon-1 | 0.194004695 | 0.703399658 | Cellular metabolism |
| O43353 | RIPK2 | Receptor-interacting serine/threonine-protein kinase 2 | 0.005817413 | 0.628248851 | Immune response |
| O43390 | HNRNPR | Heterogeneous nuclear ribonucleoprotein R | 0.319396337 | 0.884700139 | Cellular metabolism |
| O43395 | PRPF3 | U4/U6 small nuclear ribonucleoprotein Prp3 | 0.725557963 | 0.206199646 | Cellular metabolism |
| O43396 | TXNL1 | Thioredoxin-like protein 1 | -0.155327479 | -0.261315028 | Cellular metabolism |
| O43427-2 | FIBP | Acidic fibroblast growth factor intracellular-binding protein | -0.212836583 | 0 | Other |
| O43447 | PPIH | Peptidyl-prolyl cis-trans isomerase H | 0.016305288 | -1.142792384 | Cellular metabolism |
| O43482 | OIP5 | Protein Mis18-beta | -0.412711461 | 0 | Other |
| O43488 | AKR7A2 | Aflatoxin B1 aldehyde reductase member 2 | -0.179334641 | -0.895278931 | Cellular metabolism |
| O43491 | EPB41L2 | Band 4.1-like protein 2 | -0.069074631 | -0.752919515 | Other |
| O43583 | DENR | Density-regulated protein | 0.007562637 | -0.206258774 | Other |
| O43592 | XPOT | Exportin-T | -0.047531128 | -0.747564316 | Transport |
| O43615 | TIMM44 | Mitochondrial import inner membrane translocase subunit TIM44 | -0.704795202 | 1.424070994 | Transport |
| O43617 | TRAPPC3 | Trafficking protein particle complex subunit 3 | -0.041987737 | -0.048995972 | Transport |
| O43639 | NCK2 | Cytoplasmic protein NCK2 | 0.408953985 | 0 | Other |
| O43660 | PLRG1 | Pleiotropic regulator 1 | 0.048187892 | -0.317363103 | Cellular metabolism |
| O43663-3 | PRC1 | Protein regulator of cytokinesis 1 | -1.027994156 | 0 | Other |
| O43665 | RGS10 | Regulator of G-protein signaling 10 | -0.040219625 | -0.389990489 | Other |
| O43681 | ASNA1 | ATPase ASNA1 | -0.062040965 | -0.350595474 | Transport |
| O43707 | ACTN4 | Alpha-actinin-4 | -0.093605677 | -0.402488708 | Other |

| | | | | | |
|---|---|---|---|---|---|
| O43719 | HTATSF1 | HIV Tat-specific factor 1 | -0.056021372 | -1.459405263 | Cellular metabolism |
| O43747 | AP1G1 | AP-1 complex subunit gamma-1 | 0.074196498 | 0.597316742 | Transport |
| O43776 | NARS | Asparagine--tRNA ligase, cytoplasmic | -0.008787155 | -0.497225444 | Other |
| O43809 | NUDT21 | Cleavage and polyadenylation specificity factor subunit 5 | 0.787130992 | 0.931540807 | Cellular metabolism |
| O43813 | LANCL1 | LanC-like protein 1 | -0.000214895 | -0.009635925 | Other |
| O43815 | STRN | Striatin | 0.133839925 | -0.125817617 | Other |
| O43818 | RRP9 | U3 small nucleolar RNA-interacting protein 2 | 0.466264089 | -0.033964793 | Cellular metabolism |
| O43847 | NRD1 | Nardilysin | 0.068024317 | -0.809659322 | Other |
| O43852 | CALU | Calumenin | 0.106737137 | -0.390624364 | Transport |
| O43852-2 | CALU | Calumenin | 0.124839147 | -0.138050715 | Transport |
| O43865 | AHCYL1 | Putative adenosylhomocysteinase 2 | -0.216343562 | -2.191794713 | Transport |
| O43924 | PDE6D | Retinal rod rhodopsin-sensitive cGMP 3,5-cyclic phosphodiesterase subunit delta | -0.004262288 | -1.679004669 | Other |
| O60231 | DHX16 | Putative pre-mRNA-splicing factor ATP-dependent RNA helicase DHX16 | 0.221618017 | -0.460314433 | Cellular metabolism |
| O60256 | PRPSAP2 | Phosphoribosyl pyrophosphate synthase-associated protein 2 | 0.284504573 | -1.260564168 | Cellular metabolism |
| O60264 | SMARCA5 | SWI/SNF-related matrix-associated actin-dependent regulator of chromatin subfamily A member 5 | -0.022901535 | 0.008546829 | Cellular metabolism |
| O60271-4 | SPAG9 | C-Jun-amino-terminal kinase-interacting protein 4 | -0.181886673 | -0.652170817 | Transport |
| O60294 | LCMT2 | tRNA wybutosine-synthesizing protein 4 | 0.340831121 | -0.18633016 | Cellular metabolism |
| O60306 | AQR | Intron-binding protein aquarius | 0.209519068 | 0 | Cellular metabolism |
| O60313 | OPA1 | Dynamin-like 120 kDa protein, mitochondrial | 0 | 1.038837433 | Transport |
| O60341 | KDM1A | Lysine-specific histone demethylase 1A | -0.779704412 | 0.457272847 | Cellular metabolism |
| O60343 | TBC1D4 | TBC1 domain family member 4 | 0.208098094 | 0 | Transport |
| O60443 | DFNA5 | Non-syndromic hearing impairment protein 5 | -0.335477193 | -0.827823003 | Other |
| O60447-2 | EVI5 | Ecotropic viral integration site 5 protein homolog | -0.449853261 | 0 | Transport |
| O60493 | SNX3 | Sorting nexin-3 | -0.245636622 | -0.239042282 | Transport |
| O60502 | MGEA5 | Protein O-GlcNAcase | -0.419136683 | -0.863681157 | Transport |
| O60506 | SYNCRIP | Heterogeneous nuclear ribonucleoprotein Q | 0.524442037 | 0.244323095 | Cellular metabolism |
| O60518 | RANBP6 | Ran-binding protein 6 | 0.417137782 | 0 | Transport |
| O60524-4 | NEMF | Nuclear export mediator factor NEMF | -0.350377401 | -0.777959188 | Transport |
| O60547-2 | GMDS | GDP-mannose 4,6 dehydratase | -0.185206095 | 0.413458506 | Cellular metabolism |
| Q5VUC6 | NMT2 | Glycylpeptide N-tetradecanoyltransferase | 0.223124822 | -2.162625631 | Cellular metabolism |
| O60566 | BUB1B | Mitotic checkpoint serine/threonine-protein kinase BUB1 beta | -0.338469823 | -1.402954102 | Cellular metabolism |
| O60568 | PLOD3 | Procollagen-lysine,2-oxoglutarate 5-dioxygenase 3 | -0.379751205 | -0.025225957 | Cellular metabolism |
| O60664 | PLIN3 | Perilipin-3 | 0.652338028 | 2.780045191 | Transport |
| O60671 | RAD1 | Cell cycle checkpoint protein RAD1 | -0.683046341 | -0.613131205 | Cellular metabolism |

| | | | | | |
|---|---|---|---|---|---|
| O60678 | PRMT3 | Protein arginine N-methyltransferase 3 | -0.237574895 | -1.954165141 | Cellular metabolism |
| O60684 | KPNA6 | Importin subunit alpha-7 | -0.134134928 | 0.043239594 | Transport |
| O60701 | UGDH | UDP-glucose 6-dehydrogenase | 0.01226298 | -0.40769132 | Cellular metabolism |
| O60711 | LPXN | Leupaxin | -0.313621521 | 0 | Cellular metabolism |
| O60749 | SNX2 | Sorting nexin-2 | 0.053483327 | -0.274398168 | Transport |
| O60763 | USO1 | General vesicular transport factor p115 | -0.055039088 | -0.421073914 | Transport |
| O60784 | TOM1 | Target of Myb protein 1 | 0.023359934 | -1.244852066 | Transport |
| U3KQK0 | HIST1H2BN | Histone H2B | 0.266064326 | 0.816699982 | Immune response |
| O60826 | CCDC22 | Coiled-coil domain-containing protein 22 | -0.448254267 | 1.712069194 | Transport |
| O60828-2 | PQBP1 | Polyglutamine-binding protein 1 | 0.046258926 | 0.319335938 | Cellular metabolism |
| O60832 | DKC1 | H/ACA ribonucleoprotein complex subunit 4 | -0.04736646 | -0.324539185 | Cellular metabolism |
| O60869-2 | EDF1 | Endothelial differentiation-related factor 1 | -0.201229095 | -0.774183273 | Cellular metabolism |
| O60884 | DNAJA2 | DnaJ homolog subfamily A member 2 | -0.107185364 | 0.005594254 | Cellular metabolism |
| O60885 | BRD4 | Bromodomain-containing protein 4 | -1.131905238 | -1.737183889 | Cellular metabolism |
| O60888-3 | CUTA | Protein CutA | -0.186947505 | 1.887091319 | Other |
| O60921 | HUS1 | Checkpoint protein HUS1 | -0.034852982 | 0 | Cellular metabolism |
| O60927 | PPP1R11 | Protein phosphatase 1 regulatory subunit 11 | 0.048829397 | -2.338464101 | Other |
| O60942-2 | RNGTT | mRNA-capping enzyme | 0.093479792 | -0.988295237 | Cellular metabolism |
| O75083 | WDR1 | WD repeat-containing protein 1 | -0.044760386 | -0.930852254 | Transport |
| O75113 | N4BP1 | NEDD4-binding protein 1 | 0.922810872 | -0.013288498 | Other |
| O75116 | ROCK2 | Rho-associated protein kinase 2 | 0.306829453 | -1.498809179 | Cellular metabolism |
| O75131 | CPNE3 | Copine-3 | -0.004956563 | -0.504454295 | Lipid metabolism |
| O75143-3 | ATG13 | Autophagy-related protein 13 | -0.387137095 | -1.552349091 | Cellular metabolism |
| O75155-2 | CAND2 | Cullin-associated NEDD8-dissociated protein 2 | -0.102400462 | 0 | Cellular metabolism |
| O75157-2 | TSC22D2 | TSC22 domain family protein 2 | -0.238444646 | -0.112480799 | Transport |
| O75170-2 | PPP6R2 | Serine/threonine-protein phosphatase 6 regulatory subunit 2 | 0.089438121 | -0.727052689 | Other |
| O75175 | CNOT3 | CCR4-NOT transcription complex subunit 3 | 0.49242719 | 0 | Cellular metabolism |
| O75191 | XYLB | Xylulose kinase | -0.336926142 | 0 | Cellular metabolism |
| O75223 | GGCT | Gamma-glutamylcyclotransferase | -0.091923396 | 0.140513102 | Cellular metabolism |
| O75312 | ZPR1 | Zinc finger protein ZPR1 | -0.042002996 | -1.357156118 | Cellular metabolism |
| O75330-4 | HMMR | Hyaluronan mediated motility receptor | -0.067893346 | 0 | Other |
| O75340 | PDCD6 | Programmed cell death protein 6 | 0.101165136 | -0.918542226 | Transport |
| O75342 | ALOX12B | Arachidonate 12-lipoxygenase, 12R-type | 0 | 0.499202092 | Lipid metabolism |

| | | | | | |
|---|---|---|---|---|---|
| O75348 | ATP6V1G1 | V-type proton ATPase subunit G 1 | -0.327708562 | 0.195869446 | Transport |
| O75351 | VPS4B | Vacuolar protein sorting-associated protein 4B | -0.301087697 | -0.453121821 | Transport |
| O75367-2 | H2AFY | Core histone macro-H2A.1 | -0.017825445 | 0.969579697 | Other |
| O75368 | SH3BGRL | SH3 domain-binding glutamic acid-rich-like protein | -0.072588603 | -0.252780279 | Other |
| O75369-8 | FLNB | Filamin-B | 0.051005046 | -0.47841835 | Other |
| O75376-2 | NCOR1 | Nuclear receptor corepressor 1 | 0.194969177 | 0.746262868 | Lipid metabolism |
| O75391 | SPAG7 | Sperm-associated antigen 7 | -0.016560872 | -0.455797195 | Other |
| O75396 | SEC22B | Vesicle-trafficking protein SEC22b | 0.69397227 | 1.637915293 | Transport |
| O75410-2 | TACC1 | Transforming acidic coiled-coil-containing protein 1 | -0.253490448 | -0.3979098 | Other |
| O75436 | VPS26A | Vacuolar protein sorting-associated protein 26A | -0.00863711 | -1.107159932 | Transport |
| O75439 | PMPCB | Mitochondrial-processing peptidase subunit beta | -0.341923396 | -0.428141912 | Transport |
| O75449 | KATNA1 | Katanin p60 ATPase-containing subunit A1 | 0.230661392 | -0.667844772 | Other |
| O75475 | PSIP1 | PC4 and SFRS1-interacting protein | -0.118151983 | -0.583971659 | Cellular metabolism |
| O75477 | ERLIN1 | Erlin-1 | 0 | 1.057285945 | Cellular metabolism |
| O75489 | NDUFS3 | NADH dehydrogenase [ubiquinone] iron-sulfur protein 3, mitochondrial | 0 | -0.06443278 | Cellular metabolism |
| Q5JRI1 | SRSF10 | Serine/arginine-rich splicing factor 10 | 0.343186696 | 1.509784698 | Transport |
| O75531 | BANF1 | Barrier-to-autointegration factor | 0 | 0.672843297 | Cellular metabolism |
| O75533 | SF3B1 | Splicing factor 3B subunit 1 | 0.180390676 | -0.452936172 | Cellular metabolism |
| O75534 | CSDE1 | Cold shock domain-containing protein E1 | 0.298482895 | -0.465905507 | Cellular metabolism |
| O75554 | WBP4 | WW domain-binding protein 4 | -0.023904165 | 0.203196208 | Cellular metabolism |
| O75569-3 | PRKRA | Interferon-inducible double-stranded RNA-dependent protein kinase activator A | 1.1980292 | 0.827960968 | Immune response |
| O75607 | NPM3 | Nucleoplasmin-3 | -0.266076406 | -0.497505188 | Cellular metabolism |
| O75608-2 | LYPLA1 | Acyl-protein thioesterase 1 | -0.265340805 | -0.366158803 | Lipid metabolism |
| O75635-2 | SERPINB7 | Serpin B7 | 0 | 0.144760132 | Other |
| O75643 | SNRNP200 | U5 small nuclear ribonucleoprotein 200 kDa helicase | 0.02624766 | -0.220153809 | Cellular metabolism |
| O75663 | TIPRL | TIP41-like protein | -0.16033109 | -0.41832606 | Other |
| O75688 | PPM1B | Protein phosphatase 1B | -0.141962051 | -0.673565547 | Cellular metabolism |
| O75691 | UTP20 | Small subunit processome component 20 homolog | -0.220610301 | 0 | Cellular metabolism |
| O75717 | WDHD1 | WD repeat and HMG-box DNA-binding protein 1 | -0.132441839 | -1.904214859 | Cellular metabolism |
| O75746 | SLC25A12 | Calcium-binding mitochondrial carrier protein Aralar1 | 0 | 0.059707006 | Transport |
| O75792 | RNASEH2A | Ribonuclease H2 subunit A | -0.616146088 | 0.278885523 | Cellular metabolism |
| O75794 | CDC123 | Cell division cycle protein 123 homolog | -1.016450882 | -1.529069901 | Other |
| O75815-2 | BCAR3 | Breast cancer anti-estrogen resistance protein 3 | -0.50226593 | -0.604107539 | Other |
| X6RLL4 | RPP40 | Ribonuclease P protein subunit p40 | 0.531824748 | 0.278687795 | Cellular metabolism |

| | | | | | |
|---|---|---|---|---|---|
| O75821 | EIF3G | Eukaryotic translation initiation factor 3 subunit G | 0.278129578 | -0.039639791 | Cellular metabolism |
| O75822 | EIF3J | Eukaryotic translation initiation factor 3 subunit J | 0.016340256 | -0.1430734 | Cellular metabolism |
| O75828 | CBR3 | Carbonyl reductase [NADPH] 3 | 0.103423436 | 0.116315206 | Cellular metabolism |
| O75832 | PSMD10 | 26S proteasome non-ATPase regulatory subunit 10 | 0.159424464 | -0.233025233 | Immune response |
| O75843 | AP1G2 | AP-1 complex subunit gamma-like 2 | 0.257562002 | 0 | Transport |
| O75844 | ZMPSTE24 | CAAX prenyl protease 1 homolog | 1.33527565 | 1.269480387 | Cellular metabolism |
| O75874 | IDH1 | Isocitrate dehydrogenase [NADP] cytoplasmic | -0.204607646 | -0.585714976 | Lipid metabolism |
| O75882-3 | ATRN | Attractin | 0.83512942 | -2.120599747 | Other |
| O75884 | RBBP9 | Putative hydrolase RBBP9 | -0.774363836 | 0.12316068 | Other |
| O75886 | STAM2 | Signal transducing adapter molecule 2 | -0.030813217 | -0.193305969 | Transport |
| O75896 | TUSC2 | Tumor suppressor candidate 2 | -0.192565918 | 0 | Transport |
| O75911 | DHRS3 | Short-chain dehydrogenase/reductase 3 | -0.460678736 | 2.695272446 | Lipid metabolism |
| O75925 | PIAS1 | E3 SUMO-protein ligase PIAS1 | -0.454547882 | 0 | Cellular metabolism |
| X6RCK5 | DCTN3 | Dynactin subunit 3 | 0.234146754 | -1.238984426 | Other |
| O75937 | DNAJC8 | DnaJ homolog subfamily C member 8 | 0.037539164 | -0.298442841 | Cellular metabolism |
| O75940 | SMNDC1 | Survival of motor neuron-related-splicing factor 30 | -0.114423752 | -1.304688136 | Cellular metabolism |
| O75947 | ATP5H | ATP synthase subunit d, mitochondrial | 0 | -0.192920685 | Transport |
| O75962-5 | TRIO | Triple functional domain protein | 0.436166128 | 0.457776388 | Other |
| O76003 | GLRX3 | Glutaredoxin-3 | -0.146505992 | -0.792013804 | Other |
| O76031 | CLPX | ATP-dependent Clp protease ATP-binding subunit clpX-like, mitochondrial | 0 | 0.421988805 | Cellular metabolism |
| O76054 | SEC14L2 | SEC14-like protein 2 | -0.338859558 | -0.787960052 | Transport |
| O76071 | CIAO1 | Probable cytosolic iron-sulfur protein assembly protein CIAO1 | 0.27964592 | -0.865139643 | Cellular metabolism |
| O76094 | SRP72 | Signal recognition particle subunit SRP72 | 0.064877828 | 0.1190389 | Transport |
| O94760 | DDAH1 | N(G),N(G)-dimethylarginine dimethylaminohydrolase 1 | 0.024463018 | -0.641785304 | Cellular metabolism |
| O94763-2 | URI1 | Unconventional prefoldin RPB5 interactor 1 | 0.332998912 | 0 | Other |
| O94776 | MTA2 | Metastasis-associated protein MTA2 | 0.238119125 | -0.318340937 | Cellular metabolism |
| O94804 | STK10 | Serine/threonine-protein kinase 10 | 0.397652308 | 0.283001582 | Cellular metabolism |
| O94808 | GFPT2 | Glutamine--fructose-6-phosphate aminotransferase [isomerizing] 2 | 0.100411097 | -1.485110601 | Cellular metabolism |
| O94817 | ATG12 | Ubiquitin-like protein ATG12 | -0.355450948 | 0 | Immune response |
| O94822 | LTN1 | E3 ubiquitin-protein ligase listerin | -0.373881658 | -1.143815994 | Cellular metabolism |
| O94826 | TOMM70A | Mitochondrial import receptor subunit TOM70 | 0 | 0.686726888 | Transport |
| O94829 | IPO13 | Importin-13 | -0.140071233 | -0.520153681 | Transport |
| O94830 | DDHD2 | Phospholipase DDHD2 | -0.244433085 | 0 | Lipid metabolism |
| O94855 | SEC24D | Protein transport protein Sec24D | -0.323720296 | 0 | Transport |

| | | | | | |
|---|---|---|---|---|---|
| O94874-3 | UFL1 | E3 UFM1-protein ligase 1 | 0 | 1.474365234 | Cellular metabolism |
| O94888 | UBXN7 | UBX domain-containing protein 7 | -0.17768987 | -0.898745219 | Other |
| O94903 | PROSC | Proline synthase co-transcribed bacterial homolog protein | 0.049304326 | -0.647267024 | Other |
| O94906 | PRPF6 | Pre-mRNA-processing factor 6 | 0.7575442 | 0.159683863 | Cellular metabolism |
| O94915 | FRYL | Protein furry homolog-like | -0.217856089 | 0 | Cellular metabolism |
| O94925-3 | GLS | Glutaminase kidney isoform, mitochondrial | -0.305697759 | -0.556901296 | Transport |
| O94966-4 | USP19 | Ubiquitin carboxyl-terminal hydrolase 19 | -0.81235822 | -1.213471731 | Cellular metabolism |
| O94967-2 | WDR47 | WD repeat-containing protein 47 | 0.12983195 | -1.040359497 | Other |
| O94973 | AP2A2 | AP-2 complex subunit alpha-2 | -0.11788559 | -0.005373001 | Transport |
| O94992 | HEXIM1 | Protein HEXIM1 | 0.077563604 | 0.481608073 | Cellular metabolism |
| O95059 | RPP14 | Ribonuclease P protein subunit p14 | 0.367673238 | 0 | Cellular metabolism |
| O95071-2 | UBR5 | E3 ubiquitin-protein ligase UBR5 | -0.279268901 | -0.720862071 | Cellular metabolism |
| O95104-2 | SCAF4 | Splicing factor, arginine/serine-rich 15 | 0.435883204 | 0.76954778 | Other |
| O95155-2 | UBE4B | Ubiquitin conjugation factor E4 B | -0.12853686 | -0.802103043 | Cellular metabolism |
| O95197-3 | RTN3 | Reticulon-3 | 0.275059382 | -0.284280777 | Transport |
| O95202 | LETM1 | LETM1 and EF-hand domain-containing protein 1, mitochondrial | 0 | 0.573896408 | Other |
| O95210 | STBD1 | Starch-binding domain-containing protein 1 | 0 | 3.034735998 | Other |
| O95218-2 | ZRANB2 | Zinc finger Ran-binding domain-containing protein 2 | -0.151433945 | -0.335983912 | Cellular metabolism |
| O95219 | SNX4 | Sorting nexin-4 | -1.013732274 | -0.29815038 | Transport |
| O95260-2 | ATE1 | Arginyl-tRNA--protein transferase 1 | -0.480738958 | -1.228729248 | Cellular metabolism |
| O95273-2 | CCNDBP1 | Cyclin-D1-binding protein 1 | 0.08415095 | -1.915564219 | Other |
| O95292 | VAPB | Vesicle-associated membrane protein-associated protein B/C | -0.359160741 | 1.153568904 | Lipid metabolism |
| O95302 | FKBP9 | Peptidyl-prolyl cis-trans isomerase FKBP9 | 0.212966919 | -0.551226934 | Cellular metabolism |
| O95336 | PGLS | 6-phosphogluconolactonase | -0.034200033 | 0.185031891 | Cellular metabolism |
| O95340 | PAPSS2 | Bifunctional 3-phosphoadenosine 5-phosphosulfate synthase 2 | 0.017835617 | -1.059670766 | Cellular metabolism |
| O95347 | SMC2 | Structural maintenance of chromosomes protein 2 | 0.109256744 | 0.968269348 | Cellular metabolism |
| O95352 | ATG7 | Ubiquitin-like modifier-activating enzyme ATG7 | 0.005308151 | -1.376045863 | Transport |
| O95361 | TRIM16 | Tripartite motif-containing protein 16 | -0.368104299 | -0.991213481 | Cellular metabolism |
| O95372 | LYPLA2 | Acyl-protein thioesterase 2 | -0.009940465 | -1.380961736 | Lipid metabolism |
| O95373 | IPO7 | Importin-7 | -0.170951207 | -0.530061722 | Immune response |
| O95376 | ARIH2 | E3 ubiquitin-protein ligase ARIH2 | -0.090262731 | -2.514399211 | Cellular metabolism |
| O95391 | SLU7 | Pre-mRNA-splicing factor SLU7 | 0.356417974 | 0.810157776 | Transport |
| O95396 | MOCS3 | Adenylyltransferase and sulfurtransferase MOCS3 | -0.176790873 | -0.644244512 | Cellular metabolism |
| O95400 | CD2BP2 | CD2 antigen cytoplasmic tail-binding protein 2 | 0.116378148 | -0.030275981 | Cellular metabolism |

| | | | | | |
|---|---|---|---|---|---|
| O95429 | BAG4 | BAG family molecular chaperone regulator 4 | -0.78587532 | -1.174985886 | Cellular metabolism |
| O95433 | AHSA1 | Activator of 90 kDa heat shock protein ATPase homolog 1 | -0.028800964 | -0.496721268 | Other |
| O95456 | PSMG1 | Proteasome assembly chaperone 1 | -0.13616689 | 1.065162659 | Other |
| O95486 | SEC24A | Protein transport protein Sec24A | -0.070705414 | -0.1774381 | Transport |
| O95487-2 | SEC24B | Protein transport protein Sec24B | -0.105558395 | 0.194173177 | Transport |
| O95551 | TDP2 | Tyrosyl-DNA phosphodiesterase 2 | -0.465132395 | -1.568532944 | Cellular metabolism |
| O95628-5 | CNOT4 | CCR4-NOT transcription complex subunit 4 | 0.429213842 | -0.22002538 | Cellular metabolism |
| O95630 | STAMBP | STAM-binding protein | -0.188281377 | -0.208503723 | Cellular metabolism |
| O95671 | ASMTL | N-acetylserotonin O-methyltransferase-like protein | -0.082956314 | -1.211419423 | Other |
| O95716 | RAB3D | Ras-related protein Rab-3D | -0.053949356 | 1.468544642 | Transport |
| O95721 | SNAP29 | Synaptosomal-associated protein 29 | -0.468099594 | 0.711271286 | Transport |
| O95747 | OXSR1 | Serine/threonine-protein kinase OSR1 | -0.009970347 | -0.085203171 | Cellular metabolism |
| O95749 | GGPS1 | Geranylgeranyl pyrophosphate synthase | -0.04903539 | -0.928105036 | Lipid metabolism |
| O95757 | HSPA4L | Heat shock 70 kDa protein 4L | 0.022628148 | -1.619396845 | Cellular metabolism |
| O95758-1 | PTBP3 | Polypyrimidine tract-binding protein 3 | -1.004773458 | 0 | Cellular metabolism |
| O95777 | LSM8 | U6 snRNA-associated Sm-like protein LSm8 | 0.177080154 | 0.261500041 | Cellular metabolism |
| O95782-2 | AP2A1 | AP-2 complex subunit alpha-1 | -0.109998703 | 0.36126709 | Transport |
| O95793 | STAU1 | Double-stranded RNA-binding protein Staufen homolog 1 | 0.133954366 | -0.195486069 | Transport |
| O95801 | TTC4 | Tetratricopeptide repeat protein 4 | -0.58171018 | -0.684185664 | Other |
| O95816 | BAG2 | BAG family molecular chaperone regulator 2 | -0.213739395 | 0.878614426 | Cellular metabolism |
| O95817 | BAG3 | BAG family molecular chaperone regulator 3 | -0.004018784 | -0.475699743 | Cellular metabolism |
| O95834 | EML2 | Echinoderm microtubule-associated protein-like 2 | -0.298032761 | -0.191582998 | Other |
| O95861 | BPNT1 | 3(2),5-bisphosphate nucleotidase 1 | -0.217350642 | -1.466058731 | Lipid metabolism |
| O95865 | DDAH2 | N(G),N(G)-dimethylarginine dimethylaminohydrolase 2 | -0.326234182 | -0.488739014 | Cellular metabolism |
| O95881 | TXNDC12 | Thioredoxin domain-containing protein 12 | 0.355174383 | -0.360455195 | Other |
| O95905 | ECD | Protein SGT1 | -0.284210841 | -0.485626856 | Cellular metabolism |
| O95999 | BCL10 | B-cell lymphoma/leukemia 10 | 0.038661321 | -1.42636172 | Immune response |
| O96000 | NDUFB10 | NADH dehydrogenase [ubiquinone] 1 beta subcomplex subunit 10 | 0 | 0.555282593 | Cellular metabolism |
| O96007 | MOCS2 | Molybdopterin synthase catalytic subunit | -0.43138059 | -1.815998077 | Cellular metabolism |
| O96008 | TOMM40 | Mitochondrial import receptor subunit TOM40 homolog | 0.279638926 | 1.10353597 | Transport |
| O96019 | ACTL6A | Actin-like protein 6A | -0.112549464 | -0.515977224 | Cellular metabolism |
| P00338 | LDHA | L-lactate dehydrogenase A chain | -0.209314982 | -0.424856186 | Cellular metabolism |
| P00352 | ALDH1A1 | Retinal dehydrogenase 1 | 0.106103897 | 0.524581909 | Lipid metabolism |

| | | | | | |
|---|---|---|---|---|---|
| P00367 | GLUD1 | Glutamate dehydrogenase 1, mitochondrial | 1.493338903 | -0.793364843 | Cellular metabolism |
| P00374 | DHFR | Dihydrofolate reductase | -0.311314901 | -2.393125534 | Cellular metabolism |
| P00390-2 | GSR | Glutathione reductase, mitochondrial | 0.054885228 | -0.599942525 | Cellular metabolism |
| P00403 | MT-CO2 | Cytochrome c oxidase subunit 2 | -0.108661016 | 1.623432159 | Cellular metabolism |
| P00441 | SOD1 | Superoxide dismutase [Cu-Zn] | 0.259873708 | -0.634426753 | Transport |
| P00491 | PNP | Purine nucleoside phosphorylase | -0.010892868 | -0.249302546 | Immune response |
| P00492 | HPRT1 | Hypoxanthine-guanine phosphoribosyltransferase | -0.108491898 | -0.572539647 | Cellular metabolism |
| P00505 | GOT2 | Aspartate aminotransferase, mitochondrial | -0.080502192 | 0.335023244 | Transport |
| P00533 | EGFR | Epidermal growth factor receptor | -0.683371226 | 0.362639109 | Immune response |
| P00558 | PGK1 | Phosphoglycerate kinase 1 | -0.054613113 | -0.195662816 | Cellular metabolism |
| Q5T9B7 | AK1 | Adenylate kinase isoenzyme 1 | -0.07088534 | -0.561625163 | Cellular metabolism |
| P00918 | CA2 | Carbonic anhydrase 2 | 0.139822642 | 0.950192769 | Transport |
| P00966 | ASS1 | Argininosuccinate synthase | -0.273639679 | 0.003278732 | Cellular metabolism |
| P01011 | SERPINA3 | Alpha-1-antichymotrypsin | 0 | -0.468813578 | Other |
| P01036 | CST4 | Cystatin-S | 0 | 1.579545339 | Other |
| P01040 | CSTA | Cystatin-A | -0.92554601 | 1.57597669 | Cellular metabolism |
| P01111 | NRAS | GTPase NRas | -0.647357305 | 0.021850586 | Immune response |
| P01584 | IL1B | Interleukin-1 beta | -0.453588486 | 0.35077095 | Immune response |
| P01833 | PIGR | Polymeric immunoglobulin receptor | 0 | 1.368231455 | Transport |
| P01834 | IGKC | Ig kappa chain C region | -0.119368871 | 0.956465403 | Immune response |
| P02452 | COL1A1 | Collagen alpha-1(I) chain | -1.938363393 | -2.588878632 | Transport |
| P02786 | TFRC | Transferrin receptor protein 1 | 0.162296295 | 0.759485881 | Transport |
| P02787 | TF | Serotransferrin | 0.932098389 | 0.041438421 | Transport |
| P02792 | FTL | Ferritin light chain | -0.268616994 | 0.732837677 | Transport |
| P02794 | FTH1 | Ferritin heavy chain | 0.363049189 | 0.283465068 | Immune response |
| P02795 | MT2A | Metallothionein-2 | -0.826093674 | -0.640310923 | Other |
| P04040 | CAT | Catalase | 0.030464808 | 0.604321798 | Lipid metabolism |
| P04049 | RAF1 | RAF proto-oncogene serine/threonine-protein kinase | -0.141338984 | 1.035900116 | Immune response |
| P04066 | FUCA1 | Tissue alpha-L-fucosidase | 0.196339289 | 0.526058197 | Cellular metabolism |
| P04075 | ALDOA | Fructose-bisphosphate aldolase A | -0.027881622 | -0.222904841 | Transport |
| P04080 | CSTB | Cystatin-B | -0.054083506 | -0.379337311 | Other |
| P04083 | ANXA1 | Annexin A1 | -0.138599396 | -0.448575338 | Transport |
| P04114 | APOB | Apolipoprotein B-100 | -0.90500323 | -0.848734538 | Lipid metabolism |
| P04150-3 | NR3C1 | Glucocorticoid receptor | -0.05109024 | 0 | Lipid metabolism |

| | | | | | |
|---|---|---|---|---|---|
| P04179 | SOD2 | Superoxide dismutase [Mn], mitochondrial | 0.048652649 | -0.307378133 | Cellular metabolism |
| P04181 | OAT | Ornithine aminotransferase, mitochondrial | -0.417289734 | -0.464261373 | Cellular metabolism |
| P04183 | TK1 | Thymidine kinase, cytosolic | -0.199517568 | -1.2462279 | Cellular metabolism |
| P04350 | TUBB4A | Tubulin beta-4A chain | 0.081490835 | -1.988457362 | Cellular metabolism |
| P04406 | GAPDH | Glyceraldehyde-3-phosphate dehydrogenase | 0.022356669 | 0.188273112 | Cellular metabolism |
| P04424 | ASL | Argininosuccinate lyase | -0.077171961 | -1.038647334 | Cellular metabolism |
| Q5SRN5 | HLA-A | HLA class I histocompatibility antigen, A-3 alpha chain | -0.323279063 | 3.662110647 | Immune response |
| P04637 | TP53 | Cellular tumor antigen p53 | 0.153495153 | -0.057068507 | Transport |
| P04733 | MT1F | Metallothionein-1F | -2.329435984 | 0.179908752 | Other |
| P04792 | HSPB1 | Heat shock protein beta-1 | -0.143381119 | -0.582456589 | Other |
| P04843 | RPN1 | Dolichyl-diphosphooligosaccharide--protein glycosyltransferase subunit 1 | 1.423274994 | 1.457979838 | Transport |
| P04844-2 | RPN2 | Dolichyl-diphosphooligosaccharide--protein glycosyltransferase subunit 2 | 1.253374736 | 1.264943441 | Transport |
| P04899 | GNAI2 | Guanine nucleotide-binding protein G(i) subunit alpha-2 | -0.287179311 | 0.510261536 | Cellular metabolism |
| P05023-4 | ATP1A1 | Sodium/potassium-transporting ATPase subunit alpha-1 | 0.735779444 | 0.749560674 | Transport |
| P05026 | ATP1B1 | Sodium/potassium-transporting ATPase subunit beta-1 | 0 | 0.217157364 | Transport |
| P05089 | ARG1 | Arginase-1 | -0.522538503 | 1.09631602 | Cellular metabolism |
| P05091 | ALDH2 | Aldehyde dehydrogenase, mitochondrial | -0.445383072 | -1.073775609 | Cellular metabolism |
| P05109 | S100A8 | Protein S100-A8 | -0.961432139 | 1.497512817 | Immune response |
| P05114 | HMGN1 | Non-histone chromosomal protein HMG-14 | 0.31080691 | -2.252096812 | Cellular metabolism |
| P05120 | SERPINB2 | Plasminogen activator inhibitor 2 | -0.199047089 | 0.107736588 | Other |
| P05121 | SERPINE1 | Plasminogen activator inhibitor 1 | -0.578121185 | -0.709524155 | Transport |
| P05141 | SLC25A5 | ADP/ATP translocase 2 | 0.161338806 | 0.982447306 | Transport |
| P05198 | EIF2S1 | Eukaryotic translation initiation factor 2 subunit 1 | 0.002763748 | -0.267407099 | Transport |
| P05204 | HMGN2 | Non-histone chromosomal protein HMG-17 | 0.020641963 | -3.566450119 | Other |
| P05362 | ICAM1 | Intercellular adhesion molecule 1 | -0.373526255 | 0.219962438 | Immune response |
| P05386 | RPLP1 | 60S acidic ribosomal protein P1 | 0.432195028 | 0.629042943 | Transport |
| P05387 | RPLP2 | 60S acidic ribosomal protein P2 | 0.090453466 | 0.770027161 | Transport |
| P05388 | RPLP0 | 60S acidic ribosomal protein P0 | 0.489421844 | 0.331540426 | Transport |
| P05455 | SSB | Lupus La protein | -0.000770569 | -0.298495611 | Cellular metabolism |
| P05783 | KRT18 | Keratin, type I cytoskeletal 18 | 0.183185577 | 0.30567805 | Transport |
| P05976-2 | MYL1 | Myosin light chain 1/3, skeletal muscle isoform | 0.033425649 | 0 | Other |
| P06132 | UROD | Uroporphyrinogen decarboxylase | -0.078922272 | -0.954280853 | Cellular metabolism |
| P06280 | GLA | Alpha-galactosidase A | -0.036827723 | -0.71587944 | Lipid metabolism |
| P06493 | CDK1 | Cyclin-dependent kinase 1 | 0.051987966 | 1.640359879 | Immune response |

| | | | | | |
|---|---|---|---|---|---|
| P06576 | ATP5B | ATP synthase subunit beta, mitochondrial | -0.225020091 | 0.878505707 | Lipid metabolism |
| P06702 | S100A9 | Protein S100-A9 | -1.285931269 | 0.984303157 | Immune response |
| R4GN98 | S100A6 | Protein S100 | -0.965457916 | 0.446166356 | Other |
| P06730 | EIF4E | Eukaryotic translation initiation factor 4E | 0.371519089 | -0.393060684 | Transport |
| P06733 | ENO1 | Alpha-enolase | -0.033340454 | -0.708286285 | Cellular metabolism |
| P06737 | PYGL | Glycogen phosphorylase, liver form | -0.036375046 | -0.16465505 | Cellular metabolism |
| P06744 | GPI | Glucose-6-phosphate isomerase | -0.117969513 | -0.201751709 | Immune response |
| P06756-3 | ITGAV | Integrin alpha-V | 0 | -0.521027883 | Transport |
| Q16778 | HIST2H2BE | Histone H2B type 2-E | 0.354780833 | 2.495778402 | Immune response |
| P07093-2 | SERPINE2 | Glia-derived nexin | 0.151025772 | 0 | Transport |
| P07099 | EPHX1 | Epoxide hydrolase 1 | -0.387444814 | 1.048540115 | Cellular metabolism |
| P07108 | DBI | Acyl-CoA-binding protein | -0.795084 | 0.298356374 | Lipid metabolism |
| P07195 | LDHB | L-lactate dehydrogenase B chain | -0.006692251 | -0.265645345 | Cellular metabolism |
| P07196 | NEFL | Neurofilament light polypeptide | -1.215734482 | 0 | Transport |
| P07203 | GPX1 | Glutathione peroxidase 1 | 0.167180379 | -0.877695084 | Lipid metabolism |
| P07305 | H1F0 | Histone H1.0 | 0 | 0.167938232 | Cellular metabolism |
| P07311 | ACYP1 | Acylphosphatase-1 | -0.047229131 | -0.229640961 | Cellular metabolism |
| P07355 | ANXA2 | Annexin A2 | -0.105019887 | -0.592045466 | Transport |
| P07384 | CAPN1 | Calpain-1 catalytic subunit | -0.137349447 | -0.655290604 | Cellular metabolism |
| Q5JP53 | TUBB | Tubulin beta chain | -0.414346695 | 0.213422139 | Immune response |
| P07476 | IVL | Involucrin | 0 | 0.068815867 | Cellular metabolism |
| P07686 | HEXB | Beta-hexosaminidase subunit beta | 0.026173274 | -0.557354609 | Lipid metabolism |
| P07711 | CTSL | Cathepsin L1 | -0.067622503 | -1.201862971 | Immune response |
| P07737 | PFN1 | Profilin-1 | 0.088321686 | -0.566266378 | Transport |
| P07738 | BPGM | Bisphosphoglycerate mutase | -0.347085317 | 0 | Cellular metabolism |
| P07741 | APRT | Adenine phosphoribosyltransferase | -0.279764811 | -0.418685277 | Transport |
| P07814 | EPRS | Bifunctional glutamate/proline--tRNA ligase | -0.002553304 | -0.314822515 | Cellular metabolism |
| P07900 | HSP90AA1 | Heat shock protein HSP 90-alpha | -0.124158223 | -0.476772308 | Immune response |
| P07902 | GALT | Galactose-1-phosphate uridylyltransferase | -0.386955261 | -0.204297384 | Cellular metabolism |
| P07942 | LAMB1 | Laminin subunit beta-1 | 0.313392639 | -0.824570338 | Other |
| P07954-2 | FH | Fumarate hydratase, mitochondrial | -0.062041601 | -0.089553197 | Cellular metabolism |
| P08133 | ANXA6 | Annexin A6 | -0.107896169 | -0.648983637 | Other |
| Q5JR08 | RHOC | Rho-related GTP-binding protein RhoC | 0.058782578 | -0.022163391 | Transport |

| | | | | | |
|---|---|---|---|---|---|
| P08238 | HSP90AB1 | Heat shock protein HSP 90-beta | 0.008416494 | -0.375261307 | Immune response |
| P08243-2 | ASNS | Asparagine synthetase [glutamine-hydrolyzing] | -0.250443776 | -1.28201739 | Cellular metabolism |
| P08397-2 | HMBS | Porphobilinogen deaminase | -0.236277898 | -0.60360527 | Cellular metabolism |
| P08559-3 | PDHA1 | Pyruvate dehydrogenase E1 component subunit alpha, somatic form, mitochondrial | 0.168670019 | -0.622702916 | Cellular metabolism |
| P08574 | CYC1 | Cytochrome c1, heme protein, mitochondrial | 0 | 1.017564138 | Cellular metabolism |
| P08579 | SNRPB2 | U2 small nuclear ribonucleoprotein B | 0.6143411 | -1.611596425 | Cellular metabolism |
| P08621 | SNRNP70 | U1 small nuclear ribonucleoprotein 70 kDa | 0.372570674 | 0.207768122 | Cellular metabolism |
| P08648 | ITGA5 | Integrin alpha-5 | 0 | 0.676094055 | Other |
| P08670 | VIM | Vimentin | 0.249267578 | 0.017710368 | Transport |
| P08754 | GNAI3 | Guanine nucleotide-binding protein G(k) subunit alpha | -0.294380824 | 0.501722336 | Transport |
| P08758 | ANXA5 | Annexin A5 | -0.084542592 | -0.323243459 | Other |
| P09012 | SNRPA | U1 small nuclear ribonucleoprotein A | 0.403141022 | -0.052160263 | Cellular metabolism |
| P09104 | ENO2 | Gamma-enolase | 0.032589595 | -0.732609431 | Cellular metabolism |
| P09132 | SRP19 | Signal recognition particle 19 kDa protein | -0.549183528 | -0.310351054 | Transport |
| P09211 | GSTP1 | Glutathione S-transferase P | -0.201314926 | -0.198829015 | Cellular metabolism |
| P09382 | LGALS1 | Galectin-1 | -0.046595256 | -0.057017644 | Other |
| P09417 | QDPR | Dihydropteridine reductase | -0.03518041 | -0.870061239 | Cellular metabolism |
| P09429 | HMGB1 | High mobility group protein B1 | -0.65501976 | 0.043271383 | Immune response |
| P09488-2 | GSTM1 | Glutathione S-transferase Mu 1 | -0.291615168 | -2.063316981 | Cellular metabolism |
| P09496-2 | CLTA | Clathrin light chain A | -0.027198156 | 0.368380864 | Transport |
| P09497-2 | CLTB | Clathrin light chain B | -0.626427333 | 0.503401438 | Transport |
| P09525 | ANXA4 | Annexin A4 | 0.040805181 | -0.30177625 | Other |
| P09601 | HMOX1 | Heme oxygenase 1 | 0.626667658 | 2.447461446 | Transport |
| P09622 | DLD | Dihydrolipoyl dehydrogenase, mitochondrial | 0.08063825 | 0.439793905 | Cellular metabolism |
| P09661 | SNRPA1 | U2 small nuclear ribonucleoprotein A | 0.431536992 | 0.015712102 | Cellular metabolism |
| P09669 | COX6C | Cytochrome c oxidase subunit 6C | 0 | 0.466625849 | Cellular metabolism |
| P09874 | PARP1 | Poly [ADP-ribose] polymerase 1 | 0.513897578 | -0.214399338 | Cellular metabolism |
| P09936 | UCHL1 | Ubiquitin carboxyl-terminal hydrolase isozyme L1 | -0.116275152 | -0.476212184 | Transport |
| P09960 | LTA4H | Leukotriene A-4 hydrolase | -0.038598378 | -0.524782817 | Lipid metabolism |
| P09972 | ALDOC | Fructose-bisphosphate aldolase C | -0.110172908 | -3.395149867 | Cellular metabolism |
| Q71UI9 | H2AFV | Histone H2A.V | 0.341871897 | 0.511914571 | Other |
| Q6ICJ4 | Em:AP000351.3 | Glutathione S-transferase theta-2B | -0.109572093 | 0.114201228 | Cellular metabolism |
| P0CG39 | POTEJ | POTE ankyrin domain family member J | -0.075463613 | -0.01751709 | Other |
| P0DN76 | U2AF1 | Splicing factor U2AF 35 kDa subunit | 0.706512451 | 0.228193919 | Transport |
| P0DN79 | CBS | Cystathionine beta-synthase | -0.09524854 | -0.614512126 | Cellular metabolism |

| | | | | | |
|---|---|---|---|---|---|
| P0DP25 | CALM2 | | 0.066261292 | -0.627662659 | Other |
| P0DPA2 | | | 0 | 0.616422653 | #N/A |
| P0DUB6 | AMY2B | Alpha-amylase 2B | 0.24446551 | 0.304658254 | Other |
| P10155 | TROVE2 | 60 kDa SS-A/Ro ribonucleoprotein | -0.809924444 | -1.132392883 | Cellular metabolism |
| P10253 | GAA | Lysosomal alpha-glucosidase | -0.040206909 | -2.218690236 | Cellular metabolism |
| P10301 | RRAS | Ras-related protein R-Ras | 0.049971898 | 0.688179652 | Other |
| P10398 | ARAF | Serine/threonine-protein kinase A-Raf | 0.510779699 | 0 | Cellular metabolism |
| P10599 | TXN | Thioredoxin | -0.247162501 | -0.39629364 | Immune response |
| P10606 | COX5B | Cytochrome c oxidase subunit 5B, mitochondrial | 0 | 0.827841441 | Cellular metabolism |
| P10644 | PRKAR1A | cAMP-dependent protein kinase type I-alpha regulatory subunit | -0.156618118 | -1.136458715 | Immune response |
| P10768 | ESD | S-formylglutathione hydrolase | -0.030223211 | -0.72056516 | Cellular metabolism |
| P10809 | HSPD1 | 60 kDa heat shock protein, mitochondrial | -0.083035151 | 0.346396764 | Immune response |
| P11021 | HSPA5 | 78 kDa glucose-regulated protein | 0.261168798 | 0.111619949 | Transport |
| P11047 | LAMC1 | Laminin subunit gamma-1 | 0.050195058 | -1.045113246 | Other |
| P11166 | SLC2A1 | Solute carrier family 2, facilitated glucose transporter member 1 | 0.271069209 | -0.04537646 | Transport |
| P11172 | UMPS | Uridine 5-monophosphate synthase | -0.213862737 | -0.507236481 | Transport |
| P11177-2 | PDHB | Pyruvate dehydrogenase E1 component subunit beta, mitochondrial | -0.057417552 | 0.185504913 | Cellular metabolism |
| P11216 | PYGB | Glycogen phosphorylase, brain form | -0.398328781 | -1.073907852 | Cellular metabolism |
| P11217 | PYGM | Glycogen phosphorylase, muscle form | -0.032619476 | 0.789653142 | Cellular metabolism |
| P11274-2 | BCR | Breakpoint cluster region protein | -1.001975377 | 0 | Cellular metabolism |
| P11279 | LAMP1 | Lysosome-associated membrane glycoprotein 1 | 0.059996923 | 0.363414764 | Transport |
| P11310-2 | ACADM | Medium-chain specific acyl-CoA dehydrogenase, mitochondrial | -0.246052424 | 0.241948446 | Lipid metabolism |
| P11387 | TOP1 | DNA topoisomerase 1 | -2.591597875 | 0.276189168 | Cellular metabolism |
| P11413 | G6PD | Glucose-6-phosphate 1-dehydrogenase | -0.073794047 | -1.095627467 | Lipid metabolism |
| P11441 | UBL4A | Ubiquitin-like protein 4A | 0.546513875 | 0 | Transport |
| P11717 | IGF2R | Cation-independent mannose-6-phosphate receptor | -0.042462031 | 0.229347865 | Transport |
| P11766 | ADH5 | Alcohol dehydrogenase class-3 | -0.011044184 | -0.772339503 | Lipid metabolism |
| P11802 | CDK4 | Cyclin-dependent kinase 4 | -0.341659546 | -0.940059662 | Cellular metabolism |
| P11908 | PRPS2 | Ribose-phosphate pyrophosphokinase 2 | 0.218701681 | 0.026417414 | Cellular metabolism |
| P12004 | PCNA | Proliferating cell nuclear antigen | 0.170381546 | -0.136290232 | Cellular metabolism |
| P12081 | HARS | Histidine--tRNA ligase, cytoplasmic | 0.018239339 | -0.340606054 | Cellular metabolism |
| P12110 | COL6A2 | Collagen alpha-2(VI) chain | 0 | -1.215363185 | Other |
| P12236 | SLC25A6 | ADP/ATP translocase 3 | 3.232500076 | 1.709298452 | Transport |
| P12270 | TPR | Nucleoprotein TPR | 0.322064718 | 0.031178157 | Transport |
| P12273 | PIP | Prolactin-inducible protein | 1.549468358 | 0.193780263 | Transport |

| | | | | | |
|---|---|---|---|---|---|
| P12277 | CKB | Creatine kinase B-type | 0.021556218 | 0.021570841 | Cellular metabolism |
| P12429 | ANXA3 | Annexin A3 | -0.130999883 | -0.853188197 | Transport |
| P12532 | CKMT1A | Creatine kinase U-type, mitochondrial | 0 | -0.084581375 | Cellular metabolism |
| P12955 | PEPD | Xaa-Pro dipeptidase | 0.001195908 | -0.603955587 | Cellular metabolism |
| P12956 | XRCC6 | X-ray repair cross-complementing protein 6 | 0.19991684 | -0.216290156 | Immune response |
| P13010 | XRCC5 | X-ray repair cross-complementing protein 5 | 0.095312119 | 0.145664215 | Immune response |
| P13073 | COX4I1 | Cytochrome c oxidase subunit 4 isoform 1, mitochondrial | 0 | 0.025010427 | Cellular metabolism |
| P13473-2 | LAMP2 | Lysosome-associated membrane glycoprotein 2 | -0.128465652 | 0.160938899 | Transport |
| P13489 | RNH1 | Ribonuclease inhibitor | 0.00212733 | -0.565306346 | Cellular metabolism |
| P13639 | EEF2 | Elongation factor 2 | 0.034418742 | -0.027615229 | Cellular metabolism |
| P13667 | PDIA4 | Protein disulfide-isomerase A4 | 0.088148753 | -0.08177503 | Transport |
| P13674 | P4HA1 | Prolyl 4-hydroxylase subunit alpha-1 | 0.552820841 | -0.016347885 | Cellular metabolism |
| P13693 | TPT1 | Translationally-controlled tumor protein | -0.079390844 | 0.232350667 | Transport |
| P13716 | ALAD | Delta-aminolevulinic acid dehydratase | 0.060497284 | 0.36051623 | Cellular metabolism |
| P13797 | PLS3 | Plastin-3 | -0.1065286 | -0.244897842 | Other |
| P13804 | ETFA | Electron transfer flavoprotein subunit alpha, mitochondrial | -0.175295512 | -0.262607574 | Cellular metabolism |
| P13861-2 | PRKAR2A | cAMP-dependent protein kinase type II-alpha regulatory subunit | 0.128707886 | -0.398932775 | Immune response |
| P13984 | GTF2F2 | General transcription factor IIF subunit 2 | -0.311520894 | -0.624133428 | Cellular metabolism |
| P14174 | MIF | Macrophage migration inhibitory factor | 0.019388199 | -0.143715541 | Immune response |
| P14324-2 | FDPS | Farnesyl pyrophosphate synthase | -0.067875544 | -0.932216644 | Lipid metabolism |
| P14550 | AKR1A1 | Alcohol dehydrogenase [NADP(+)] | -0.21081988 | -0.597026825 | Cellular metabolism |
| P14618 | PKM | Pyruvate kinase PKM | -0.120435715 | -0.51047643 | Cellular metabolism |
| P14618-2 | PKM | Pyruvate kinase PKM | -0.012083054 | -0.626608531 | Cellular metabolism |
| P14625 | HSP90B1 | Endoplasmin | 0.219246546 | 0.001012802 | Immune response |
| P14868 | DARS | Aspartate--tRNA ligase, cytoplasmic | 0.129805883 | -0.360846202 | Cellular metabolism |
| P14921-4 | ETS1 | Protein C-ets-1 | -2.15549914 | 0 | Immune response |
| P14923 | JUP | Junction plakoglobin | -0.094846725 | 1.172299067 | Other |
| P15104 | GLUL | Glutamine synthetase | 0.046630224 | 0.199112574 | Transport |
| P15121 | AKR1B1 | Aldose reductase | -0.145787557 | -0.802213669 | Lipid metabolism |
| P15144 | ANPEP | Aminopeptidase N | 0.050761541 | 0.384933472 | Cellular metabolism |
| P15153 | RAC2 | Ras-related C3 botulinum toxin substrate 2 | 0.038142522 | -0.00840505 | Other |
| P15170-2 | GSPT1 | Eukaryotic peptide chain release factor GTP-binding subunit ERF3A | 0.003619512 | -0.576599121 | Cellular metabolism |
| P15374 | UCHL3 | Ubiquitin carboxyl-terminal hydrolase isozyme L3 | -0.135307948 | -0.099875132 | Cellular metabolism |
| P15407 | FOSL1 | Fos-related antigen 1 | -1.610742569 | 0 | Cellular metabolism |

| | | | | | |
|---|---|---|---|---|---|
| P15559-2 | NQO1 | NAD(P)H dehydrogenase [quinone] 1 | -0.216554006 | -1.694863002 | Cellular metabolism |
| P15586 | GNS | N-acetylglucosamine-6-sulfatase | 0.050044378 | -0.59294637 | Cellular metabolism |
| P15880 | RPS2 | 40S ribosomal protein S2 | 0.073746999 | 0.464727402 | Transport |
| P15924 | DSP | Desmoplakin | -0.591159821 | 1.317842484 | Cellular metabolism |
| P15927 | RPA2 | Replication protein A 32 kDa subunit | 0.036170959 | 0.018458049 | Cellular metabolism |
| Q5TD07 | NQO2 | Ribosyldihydronicotinamide dehydrogenase [quinone] | -0.237711589 | -0.915480932 | Other |
| P16152 | CBR1 | Carbonyl reductase [NADPH] 1 | -0.192780177 | 0.061517715 | Lipid metabolism |
| P16278-3 | GLB1 | Beta-galactosidase | 0.016907374 | -1.440045675 | Lipid metabolism |
| P16298 | PPP3CB | Serine/threonine-protein phosphatase 2B catalytic subunit beta isoform | -0.446320216 | -1.612169902 | Immune response |
| P16333 | NCK1 | Cytoplasmic protein NCK1 | -0.00467364 | -1.581908544 | Immune response |
| P16383 | GCFC2 | GC-rich sequence DNA-binding factor 2 | 0.055805206 | 1.012564977 | Other |
| P16401 | HIST1H1B | Histone H1.5 | 0.820387522 | 0.531606674 | Other |
| P16402 | HIST1H1D | Histone H1.3 | 0.112099965 | 0.119023005 | Other |
| P16435 | POR | NADPH--cytochrome P450 reductase | 1.58784612 | 1.252599716 | Lipid metabolism |
| P16615-5 | ATP2A2 | Sarcoplasmic/endoplasmic reticulum calcium ATPase 2 | 1.630601883 | 0.827295303 | Transport |
| P16930 | FAH | Fumarylacetoacetase | 0.143007278 | -1.013923009 | Cellular metabolism |
| P16949 | STMN1 | Stathmin | -0.24116389 | -0.538426081 | Other |
| P16989 | YBX3 | Y-box-binding protein 3 | 0.164651235 | -0.21172142 | Cellular metabolism |
| P17050 | NAGA | Alpha-N-acetylgalactosaminidase | 0.157225927 | -0.359834671 | Lipid metabolism |
| P17096 | HMGA1 | High mobility group protein HMG-I/HMG-Y | 0.23195076 | 0.102362951 | Cellular metabolism |
| P17174 | GOT1 | Aspartate aminotransferase, cytoplasmic | -0.060873667 | -0.566967646 | Cellular metabolism |
| P17252 | PRKCA | Protein kinase C alpha type | -0.104523977 | -0.441347122 | Immune response |
| P17612-2 | PRKACA | cAMP-dependent protein kinase catalytic subunit alpha | 0.008680979 | 0.31080691 | Immune response |
| P17655 | CAPN2 | Calpain-2 catalytic subunit | -0.09184583 | -0.592672348 | Other |
| P17706-2 | PTPN2 | Tyrosine-protein phosphatase non-receptor type 2 | 0.150719961 | -0.148735682 | Cellular metabolism |
| P17812 | CTPS1 | CTP synthase 1 | 0.075503667 | -0.411750158 | Cellular metabolism |
| P17858 | PFKL | ATP-dependent 6-phosphofructokinase, liver type | 0.100397746 | 0.670846939 | Cellular metabolism |
| P17900 | GM2A | Ganglioside GM2 activator | -0.643929799 | 0.635677338 | Lipid metabolism |
| P17931 | LGALS3 | Galectin-3 | -0.124710719 | 0.407084147 | Immune response |
| P17987 | TCP1 | T-complex protein 1 subunit alpha | 0.041425705 | -0.035321554 | Transport |
| P18031 | PTPN1 | Tyrosine-protein phosphatase non-receptor type 1 | -0.453725179 | 2.08511289 | Cellular metabolism |
| P18077 | RPL35A | 60S ribosomal protein L35a | 0.473908106 | 0.662161509 | Transport |
| P18085 | ARF4 | ADP-ribosylation factor 4 | -0.202580134 | 0.102741877 | Transport |
| P18124 | RPL7 | 60S ribosomal protein L7 | 0.451548894 | 0.603342056 | Transport |
| P18206-2 | VCL | Vinculin | -0.025753657 | -0.511364619 | Transport |
| P18583-6 | SON | Protein SON | 0.23802503 | 0 | Cellular metabolism |

| | | | | | |
|---|---|---|---|---|---|
| P18615 | NELFE | Negative elongation factor E | 0.166468938 | -0.787466685 | Cellular metabolism |
| P18669 | PGAM1 | Phosphoglycerate mutase 1 | -0.042173386 | -0.396129608 | Cellular metabolism |
| P18754 | RCC1 | Regulator of chromosome condensation | 0.562104543 | -0.898086548 | Other |
| P18858 | LIG1 | DNA ligase 1 | -0.429480235 | -1.905207316 | Cellular metabolism |
| P19174-2 | PLCG1 | 1-phosphatidylinositol 4,5-bisphosphate phosphodiesterase gamma-1 | 0.055969874 | -2.203561783 | Immune response |
| P19367 | HK1 | Hexokinase-1 | 0.138255437 | 0.203066508 | Transport |
| P19387 | POLR2C | DNA-directed RNA polymerase II subunit RPB3 | -0.081238429 | 0.002399445 | Cellular metabolism |
| P19388 | POLR2E | DNA-directed RNA polymerases I, II, and III subunit RPABC1 | -0.15513738 | -0.345397313 | Immune response |
| P19474 | TRIM21 | E3 ubiquitin-protein ligase TRIM21 | 1.187034607 | 2.555693309 | Immune response |
| P19623 | SRM | Spermidine synthase | -0.042585373 | -0.55346489 | Cellular metabolism |
| P19784 | CSNK2A2 | Casein kinase II subunit alpha | 0.159856796 | -0.752955755 | Cellular metabolism |
| P19838 | NFKB1 | Nuclear factor NF-kappa-B p105 subunit | -0.677125931 | -0.679631551 | Immune response |
| P20042 | EIF2S2 | Eukaryotic translation initiation factor 2 subunit 2 | 0.111509959 | -0.31037267 | Transport |
| P20073-2 | ANXA7 | Annexin A7 | -0.11060524 | -0.381953557 | Cellular metabolism |
| P20248 | CCNA2 | Cyclin-A2 | 0.034170787 | 0 | Other |
| P20290-2 | BTF3 | Transcription factor BTF3 | -0.510485331 | -0.688725154 | Transport |
| P20337 | RAB3B | Ras-related protein Rab-3B | 0.140084585 | 1.71439298 | Transport |
| P20339-2 | RAB5A | Ras-related protein Rab-5A | -0.057856878 | 2.327130636 | Transport |
| P20340-2 | RAB6A | Ras-related protein Rab-6A | -0.076402028 | 1.049633662 | Transport |
| P20618 | PSMB1 | Proteasome subunit beta type-1 | 0.299163818 | 0.207555771 | Immune response |
| P20645 | M6PR | Cation-dependent mannose-6-phosphate receptor | 0 | 1.1272494 | Transport |
| P20700 | LMNB1 | Lamin-B1 | -0.063963572 | -0.030352275 | Other |
| P20962 | PTMS | Parathymosin | 1.583220164 | 0.118379593 | Cellular metabolism |
| P21266 | GSTM3 | Glutathione S-transferase Mu 3 | -0.062897364 | -0.497040431 | Cellular metabolism |
| P21281 | ATP6V1B2 | V-type proton ATPase subunit B, brain isoform | 0.018351873 | 0.443019231 | Transport |
| P21283 | ATP6V1C1 | V-type proton ATPase subunit C 1 | 0.03427887 | 0.268371582 | Transport |
| P21291 | CSRP1 | Cysteine and glycine-rich protein 1 | 0.452107747 | -1.453976949 | Other |
| P21333-2 | FLNA | Filamin-A | 0.191155752 | -0.447783788 | Transport |
| P21399 | ACO1 | Cytoplasmic aconitate hydratase | 0.238272985 | -1.426042557 | Transport |
| P21589-2 | NT5E | 5-nucleotidase | 0.324016571 | 0 | Cellular metabolism |
| P21796 | VDAC1 | Voltage-dependent anion-selective channel protein 1 | 0.564287186 | 1.152840296 | Transport |
| P21912 | SDHB | Succinate dehydrogenase [ubiquinone] iron-sulfur subunit, mitochondrial | 0 | 2.380338669 | Cellular metabolism |
| P21964-2 | COMT | Catechol O-methyltransferase | -0.150498708 | 0.742776871 | Lipid metabolism |
| P21980 | TGM2 | Protein-glutamine gamma-glutamyltransferase 2 | -0.010754903 | -0.3424263 | Transport |
| P22059 | OSBP | Oxysterol-binding protein 1 | 0.004423777 | -1.08635203 | Transport |
| P22087 | FBL | rRNA 2-O-methyltransferase fibrillarin | 0.201557795 | 0.847094218 | Cellular metabolism |

| | | | | | |
|---|---|---|---|---|---|
| P22102 | GART | Trifunctional purine biosynthetic protein adenosine-3 | -0.024413427 | -0.480765661 | Cellular metabolism |
| P22234 | PAICS | Multifunctional protein ADE2 | 0.016074498 | -0.251615524 | Cellular metabolism |
| P22307-6 | SCP2 | Non-specific lipid-transfer protein | 0.150278727 | 0.985346476 | Lipid metabolism |
| P22314-2 | UBA1 | Ubiquitin-like modifier-activating enzyme 1 | -0.038412094 | -0.635459264 | Cellular metabolism |
| Q32Q12 | NME1-NME2 | Nucleoside diphosphate kinase | 0.025309881 | -0.353295644 | Cellular metabolism |
| P22528 | SPRR1B | Cornifin-B | 0 | -0.716740926 | Cellular metabolism |
| P22531 | SPRR2E | Small proline-rich protein 2E | 0 | 0.123877207 | Other |
| P22626 | HNRNPA2B1 | Heterogeneous nuclear ribonucleoproteins A2/B1 | 0.450318654 | 0.373890559 | Transport |
| P22681 | CBL | E3 ubiquitin-protein ligase CBL | -0.530852 | 0.967058818 | Immune response |
| P22695 | UQCRC2 | Cytochrome b-c1 complex subunit 2, mitochondrial | 0.220125198 | -0.622606277 | Cellular metabolism |
| P22735 | TGM1 | Protein-glutamine gamma-glutamyltransferase K | -0.13787206 | 0.72286733 | Cellular metabolism |
| P23246 | SFPQ | Splicing factor, proline- and glutamine-rich | 0.448013306 | -0.060365041 | Cellular metabolism |
| P23381 | WARS | Tryptophan--tRNA ligase, cytoplasmic | 0.046247482 | 0.444335938 | Cellular metabolism |
| P23396 | RPS3 | 40S ribosomal protein S3 | 0.179430008 | 0.133525848 | Transport |
| P23443-3 | RPS6KB1 | Ribosomal protein S6 kinase beta-1 | -0.454977671 | -1.700315475 | Immune response |
| P23490 | LOR | Loricrin | 0 | -0.074623108 | Cellular metabolism |
| P23497 | SP100 | Nuclear autoantigen Sp-100 | 0.039281209 | 1.396579742 | Cellular metabolism |
| P23526 | AHCY | Adenosylhomocysteinase | -0.001070658 | -0.242420832 | Cellular metabolism |
| P23528 | CFL1 | Cofilin-1 | -0.101156871 | -0.461595535 | Immune response |
| P23743 | DGKA | Diacylglycerol kinase alpha | 0.045127233 | -0.248427073 | Other |
| P23919 | DTYMK | Thymidylate kinase | -0.13315328 | -1.538771947 | Cellular metabolism |
| P23921 | RRM1 | Ribonucleoside-diphosphate reductase large subunit | -0.275513967 | -1.34301885 | Cellular metabolism |
| P24386 | CHM | Rab proteins geranylgeranyltransferase component A 1 | -0.415451686 | -1.389235814 | Transport |
| P24534 | EEF1B2 | Elongation factor 1-beta | -0.057370504 | -0.124491374 | Cellular metabolism |
| P24539 | ATP5F1 | ATP synthase F(0) complex subunit B1, mitochondrial | 0.964546204 | -0.81941096 | Transport |
| P24666 | ACP1 | Low molecular weight phosphotyrosine protein phosphatase | -0.060282389 | -0.516916275 | Other |
| P24844 | MYL9 | Myosin regulatory light polypeptide 9 | 0.013907115 | -1.130784988 | Other |
| P25098 | ADRBK1 | Beta-adrenergic receptor kinase 1 | -0.029803594 | 0 | Immune response |
| P25205 | MCM3 | DNA replication licensing factor MCM3 | -0.098178864 | -0.168587367 | Cellular metabolism |
| P25311 | AZGP1 | Zinc-alpha-2-glycoprotein | -1.984102249 | 0.666834513 | Immune response |
| P25325-2 | MPST | 3-mercaptopyruvate sulfurtransferase | -0.192269007 | -1.173566818 | Cellular metabolism |
| P25398 | RPS12 | 40S ribosomal protein S12 | 0.217873255 | 0.095535278 | Transport |
| P25685 | DNAJB1 | DnaJ homolog subfamily B member 1 | 0.140844981 | -0.467334747 | Cellular metabolism |
| P25705 | ATP5A1 | ATP synthase subunit alpha, mitochondrial | 0.068996429 | 0.790094376 | Lipid metabolism |
| P25786 | PSMA1 | Proteasome subunit alpha type-1 | 0.139604568 | 0.088223775 | Immune response |

| | | | | | |
|---|---|---|---|---|---|
| P25788-2 | PSMA3 | Proteasome subunit alpha type-3 | 0.139303843 | 0.267265956 | Immune response |
| P25789 | PSMA4 | Proteasome subunit alpha type-4 | 0.205165863 | 0.115524292 | Immune response |
| P26006 | ITGA3 | Integrin alpha-3 | 0 | 0.037610372 | Other |
| P26038 | MSN | Moesin | -0.21155866 | -0.395820618 | Other |
| P26196 | DDX6 | Probable ATP-dependent RNA helicase DDX6 | 0.203252792 | 0.034926732 | Cellular metabolism |
| P26368-2 | U2AF2 | Splicing factor U2AF 65 kDa subunit | 0.593551636 | 0.492975871 | Transport |
| P26373 | RPL13 | 60S ribosomal protein L13 | 0.328821818 | 0.817145665 | Transport |
| P26374 | CHML | Rab proteins geranylgeranyltransferase component A 2 | -0.247549693 | -0.068639755 | Transport |
| P26583 | HMGB2 | High mobility group protein B2 | -0.244260152 | -0.124202092 | Cellular metabolism |
| P26639 | TARS | Threonine--tRNA ligase, cytoplasmic | -0.136318843 | -0.247938792 | Cellular metabolism |
| P26640 | VARS | Valine--tRNA ligase | -0.0540212 | -0.792149862 | Cellular metabolism |
| P26641 | EEF1G | Elongation factor 1-gamma | -0.283793132 | -0.184619904 | Cellular metabolism |
| P27144 | AK4 | Adenylate kinase 4, mitochondrial | -0.338294347 | 0 | Cellular metabolism |
| P27348 | YWHAQ | 14-3-3 protein theta | -0.028638204 | -0.355945587 | Transport |
| P27361 | MAPK3 | Mitogen-activated protein kinase 3 | -0.098103841 | 0.637325923 | Immune response |
| P27482 | CALML3 | Calmodulin-like protein 3 | 0 | 1.029067993 | Other |
| X1WI28 | RPL10 | 60S ribosomal protein L10 | 0.296834946 | 0.843434016 | Transport |
| P27694 | RPA1 | Replication protein A 70 kDa DNA-binding subunit | -0.15986824 | -0.331045787 | Cellular metabolism |
| P27695 | APEX1 | DNA-(apurinic or apyrimidinic site) lyase | -0.005965551 | -0.59308815 | Cellular metabolism |
| P27797 | CALR | Calreticulin | 0.038693746 | -0.140827815 | Transport |
| P27986 | PIK3R1 | Phosphatidylinositol 3-kinase regulatory subunit alpha | -0.319430033 | -1.74129804 | Immune response |
| X5D2R7 | PSM8 | Proteasome subunit beta type | 0.292184194 | 1.648817698 | Immune response |
| P28066 | PSMA5 | Proteasome subunit alpha type-5 | 0.366105398 | 0.767997106 | Immune response |
| P28070 | PSMB4 | Proteasome subunit beta type-4 | 0.152771632 | 0.236879349 | Immune response |
| P28072 | PSMB6 | Proteasome subunit beta type-6 | 0.060200373 | 0.141403834 | Immune response |
| P28074 | PSMB5 | Proteasome subunit beta type-5 | 0.130357107 | 0.094745636 | Immune response |
| P28161 | GSTM2 | Glutathione S-transferase Mu 2 | -0.150143305 | -0.477537791 | Cellular metabolism |
| P28482 | MAPK1 | Mitogen-activated protein kinase 1 | -0.132440567 | -0.259683609 | Immune response |
| P28799-3 | GRN | Granulins | -0.327643712 | -0.138865789 | Other |
| P28838-2 | LAP3 | Cytosol aminopeptidase | 0.063381831 | 0.372694016 | Other |
| P29083 | GTF2E1 | General transcription factor IIE subunit 1 | 0.281960169 | 0.352364222 | Cellular metabolism |
| P29084 | GTF2E2 | Transcription initiation factor IIE subunit beta | 0.038392385 | 0 | Cellular metabolism |
| P29144 | TPP2 | Tripeptidyl-peptidase 2 | -0.277322133 | -0.214558919 | Cellular metabolism |
| P29218 | IMPA1 | Inositol monophosphatase 1 | -0.034490585 | -0.225362142 | Lipid metabolism |
| P29279-2 | CTGF | Connective tissue growth factor | 0 | 0.256567637 | Lipid metabolism |
| P29317 | EPHA2 | Ephrin type-A receptor 2 | 0 | 1.1279761 | Cellular metabolism |
| P29353-2 | SHC1 | SHC-transforming protein 1 | -0.126218796 | 0.058572133 | Immune response |

| | | | | | |
|---|---|---|---|---|---|
| P29373 | CRABP2 | Cellular retinoic acid-binding protein 2 | -0.422434489 | -1.031813939 | Lipid metabolism |
| P29401 | TKT | Transketolase | -0.072773616 | 0.035042445 | Cellular metabolism |
| P29466-2 | CASP1 | Caspase-1 | 0.015307109 | 0.864625931 | Immune response |
| P29508 | SERPINB3 | Serpin B3 | -0.846551895 | 0.760684967 | Other |
| P29590 | PML | Protein PML | 0.305789312 | 1.129463196 | Immune response |
| P29692 | EEF1D | Elongation factor 1-delta | -0.22470665 | -0.106945674 | Cellular metabolism |
| P29966 | MARCKS | Myristoylated alanine-rich C-kinase substrate | -0.286113739 | -0.082765579 | Cellular metabolism |
| P29992 | GNA11 | Guanine nucleotide-binding protein subunit alpha-11 | 0 | 1.113634109 | Other |
| P30040 | ERP29 | Endoplasmic reticulum resident protein 29 | 0.539865494 | 0.075120926 | Transport |
| P30041 | PRDX6 | Peroxiredoxin-6 | -0.160547892 | -0.262016932 | Lipid metabolism |
| P30043 | BLVRB | Flavin reductase (NADPH) | -0.022685369 | -0.356672923 | Cellular metabolism |
| P30044-2 | PRDX5 | Peroxiredoxin-5, mitochondrial | -0.110505422 | -0.012983958 | Cellular metabolism |
| P30046 | DDT | D-dopachrome decarboxylase | 0.006916682 | -0.327900569 | Cellular metabolism |
| P30048-2 | PRDX3 | Thioredoxin-dependent peroxide reductase, mitochondrial | -0.060572306 | 0.286719004 | Cellular metabolism |
| P30049 | ATP5D | ATP synthase subunit delta, mitochondrial | -0.610108693 | 1.030820211 | Transport |
| P30050 | RPL12 | 60S ribosomal protein L12 | 0.564798991 | 0.491746267 | Transport |
| P30084 | ECHS1 | Enoyl-CoA hydratase, mitochondrial | 0.348912557 | -0.444447835 | Lipid metabolism |
| P30085 | CMPK1 | UMP-CMP kinase | -0.15680631 | -0.348653158 | Cellular metabolism |
| P30086 | PEBP1 | Phosphatidylethanolamine-binding protein 1 | -0.105695089 | -0.948845545 | Other |
| P30101 | PDIA3 | Protein disulfide-isomerase A3 | 0.281826019 | -0.070656459 | Transport |
| P30153 | PPP2R1A | Serine/threonine-protein phosphatase 2A 65 kDa regulatory subunit A alpha isoform | -0.11270078 | -0.347429276 | Lipid metabolism |
| P30154 | PPP2R1B | Serine/threonine-protein phosphatase 2A 65 kDa regulatory subunit A beta isoform | -0.093546549 | -0.552297592 | Other |
| P30307-2 | CDC25C | M-phase inducer phosphatase 3 | -0.200774511 | 0 | Cellular metabolism |
| P30405 | PPIF | Peptidyl-prolyl cis-trans isomerase F, mitochondrial | -0.328414917 | 1.184041341 | Cellular metabolism |
| P30519 | HMOX2 | Heme oxygenase 2 | 1.326580683 | 1.392173767 | Transport |
| P30520 | ADSS | Adenylosuccinate synthetase isozyme 2 | -0.20001475 | -1.195648193 | Cellular metabolism |
| P30533 | LRPAP1 | Alpha-2-macroglobulin receptor-associated protein | 0.128047307 | -0.406286875 | Transport |
| P30622-2 | CLIP1 | CAP-Gly domain-containing linker protein 1 | -0.254192988 | -1.939609528 | Other |
| P30711 | GSTT1 | Glutathione S-transferase theta-1 | -0.273230871 | 0 | Cellular metabolism |
| P30740 | SERPINB1 | Leukocyte elastase inhibitor | -0.716113408 | 0.352060954 | Other |
| P30837 | ALDH1B1 | Aldehyde dehydrogenase X, mitochondrial | 0.084681829 | 0.844403585 | Other |
| P31025 | LCN1 | Lipocalin-1 | -0.843421936 | 1.966210047 | Transport |
| P31040-2 | SDHA | Succinate dehydrogenase [ubiquinone] flavoprotein subunit, mitochondrial | 0.077325185 | 0.297073364 | Cellular metabolism |

| | | | | | |
|---|---|---|---|---|---|
| P31146 | CORO1A | Coronin-1A | 0.089009603 | -0.489330292 | Immune response |
| P31150 | GDI1 | Rab GDP dissociation inhibitor alpha | -0.120633443 | -0.703759511 | Transport |
| P31151 | S100A7 | Protein S100-A7 | -1.732427597 | 0.24177742 | Immune response |
| P31153 | MAT2A | S-adenosylmethionine synthase isoform type-2 | 0.092721303 | -0.485642115 | Cellular metabolism |
| P31323 | PRKAR2B | cAMP-dependent protein kinase type II-beta regulatory subunit | -0.001293182 | 0 | Immune response |
| P31350 | RRM2 | Ribonucleoside-diphosphate reductase subunit M2 | -0.393262227 | -1.250836054 | Cellular metabolism |
| P31689 | DNAJA1 | DnaJ homolog subfamily A member 1 | -0.077346802 | 0.185448329 | Cellular metabolism |
| P31749 | AKT1 | RAC-alpha serine/threonine-protein kinase | -0.117368698 | -0.193524679 | Immune response |
| P31751 | AKT2 | RAC-beta serine/threonine-protein kinase | -0.127957662 | -1.736606598 | Transport |
| P31930 | UQCRC1 | Cytochrome b-c1 complex subunit 1, mitochondrial | -0.871864319 | 0.802642186 | Cellular metabolism |
| P31937 | HIBADH | 3-hydroxyisobutyrate dehydrogenase, mitochondrial | -0.099004745 | -0.038602193 | Cellular metabolism |
| P31939 | ATIC | Bifunctional purine biosynthesis protein PURH | -0.075317383 | -0.586538951 | Cellular metabolism |
| P31942-2 | HNRNPH3 | Heterogeneous nuclear ribonucleoprotein H3 | 0.291374207 | 0.192267736 | Cellular metabolism |
| P31944 | CASP14 | Caspase-14 | -0.843667348 | 1.317843119 | Other |
| P31946 | YWHAB | 14-3-3 protein beta/alpha | 0.119844437 | -0.180979411 | Immune response |
| P31947-2 | SFN | 14-3-3 protein sigma | 0.036218007 | -0.119497935 | Cellular metabolism |
| P31948 | STIP1 | Stress-induced-phosphoprotein 1 | -0.003538767 | -0.594544729 | Other |
| P31949 | S100A11 | Protein S100-A11 | -0.026878357 | 0.604734421 | Other |
| P32019-3 | INPP5B | Type II inositol 1,4,5-trisphosphate 5-phosphatase | 0 | -0.35496966 | Lipid metabolism |
| P32119 | PRDX2 | Peroxiredoxin-2 | -0.139701207 | 0.371656418 | Cellular metabolism |
| P32121-5 | ARRB2 | Beta-arrestin-2 | 0.224770864 | -1.167118708 | Transport |
| P32321 | DCTD | Deoxycytidylate deaminase | -0.174726486 | -0.493332545 | Cellular metabolism |
| P32455 | GBP1 | Interferon-induced guanylate-binding protein 1 | 0.124472936 | 1.356300354 | Other |
| P32456 | GBP2 | Interferon-induced guanylate-binding protein 2 | -0.104362488 | -1.188854853 | Immune response |
| P32929 | CTH | Cystathionine gamma-lyase | -0.309345881 | 0 | Cellular metabolism |
| P32969 | RPL9 | 60S ribosomal protein L9 | 0.562023163 | 1.06666247 | Transport |
| P33176 | KIF5B | Kinesin-1 heavy chain | 0.246110916 | -0.570924759 | Transport |
| P33316-2 | DUT | Deoxyuridine 5-triphosphate nucleotidohydrolase, mitochondrial | -0.265402476 | -0.848156611 | Cellular metabolism |
| P33992 | MCM5 | DNA replication licensing factor MCM5 | 0.021291097 | -1.142563502 | Cellular metabolism |
| P33993 | MCM7 | DNA replication licensing factor MCM7 | 0.046983719 | -0.322235107 | Cellular metabolism |
| P34059 | GALNS | N-acetylgalactosamine-6-sulfatase | -0.194882711 | 0 | Cellular metabolism |
| P34896-2 | SHMT1 | Serine hydroxymethyltransferase, cytosolic | 0.10516866 | -0.746636709 | Cellular metabolism |
| P34897-3 | SHMT2 | Serine hydroxymethyltransferase, mitochondrial | 0.000429153 | -0.177155813 | Cellular metabolism |

| | | | | | |
|---|---|---|---|---|---|
| Q53FA3 | HSPA1L | Heat shock 70 kDa protein 1-like | -0.24282074 | -1.472186406 | Cellular metabolism |
| P34932 | HSPA4 | Heat shock 70 kDa protein 4 | -0.041660309 | -0.307739894 | Transport |
| P34949 | MPI | Mannose-6-phosphate isomerase | 0.001785278 | -0.649501165 | Cellular metabolism |
| P35080 | PFN2 | Profilin-2 | -0.256400426 | -0.00741895 | Other |
| P35221-2 | CTNNA1 | Catenin alpha-1 | -0.205228806 | -0.683788935 | Other |
| P35232 | PHB | Prohibitin | 0.149306615 | 1.05957222 | Cellular metabolism |
| P35237 | SERPINB6 | Serpin B6 | -0.157507579 | -0.491473516 | Other |
| P35241 | RDX | Radixin | -0.163681666 | -0.241480509 | Other |
| P35244 | RPA3 | Replication protein A 14 kDa subunit | 0.057200114 | -0.25413386 | Cellular metabolism |
| P35249 | RFC4 | Replication factor C subunit 4 | 0.071510951 | -0.367216746 | Cellular metabolism |
| P35250 | RFC2 | Replication factor C subunit 2 | -0.148100535 | 0.129367193 | Cellular metabolism |
| P35268 | RPL22 | 60S ribosomal protein L22 | 0.103647232 | 0.18016688 | Transport |
| P35269 | GTF2F1 | General transcription factor IIF subunit 1 | -0.136781057 | -0.463064194 | Cellular metabolism |
| P35270 | SPR | Sepiapterin reductase | -0.568490346 | -0.864788055 | Cellular metabolism |
| P35573 | AGL | Glycogen debranching enzyme | 0 | 0.219695409 | Cellular metabolism |
| P35579 | MYH9 | Myosin-9 | 0.010831197 | -0.435604095 | Transport |
| P35580 | MYH10 | Myosin-10 | 0.057049433 | -0.3287913 | Transport |
| P35612-4 | ADD2 | Beta-adducin | -0.133179983 | 0 | Transport |
| P35637 | FUS | RNA-binding protein FUS | 0.135011037 | 0.277960459 | Cellular metabolism |
| P35754 | GLRX | Glutaredoxin-1 | -0.2751592 | 0.497158686 | Cellular metabolism |
| P35813 | PPM1A | Protein phosphatase 1A | -0.252134323 | -2.380491892 | Cellular metabolism |
| P35914 | HMGCL | Hydroxymethylglutaryl-CoA lyase, mitochondrial | 0.178194046 | 0 | Lipid metabolism |
| P36404 | ARL2 | ADP-ribosylation factor-like protein 2 | -0.310690562 | -0.946041107 | Cellular metabolism |
| P36405 | ARL3 | ADP-ribosylation factor-like protein 3 | -0.167046229 | -1.247465769 | Transport |
| P36507 | MAP2K2 | Dual specificity mitogen-activated protein kinase kinase 2 | 0.040887833 | -1.250186284 | Immune response |
| P36542-2 | ATP5C1 | ATP synthase subunit gamma, mitochondrial | -0.305217743 | 1.236337662 | Transport |
| P36543 | ATP6V1E1 | V-type proton ATPase subunit E 1 | 0.797966003 | 0.247479121 | Transport |
| P36551 | CPOX | Oxygen-dependent coproporphyrinogen-III oxidase, mitochondrial | -0.187980652 | -1.332300822 | Cellular metabolism |
| P36578 | RPL4 | 60S ribosomal protein L4 | 0.256837845 | 0.485432943 | Transport |
| P36639-4 | NUDT1 | 7,8-dihydro-8-oxoguanine triphosphatase | -0.377445221 | -0.534501394 | Cellular metabolism |
| P36871 | PGM1 | Phosphoglucomutase-1 | -0.184770584 | -0.551664988 | Cellular metabolism |
| P36915 | GNL1 | Guanine nucleotide-binding protein-like 1 | -0.036967595 | -0.775157293 | Immune response |
| P36952 | SERPINB5 | Serpin B5 | 0 | 0.301763535 | Other |
| P36957 | DLST | Dihydrolipoyllysine-residue succinyltransferase component of 2-oxoglutarate dehydrogenase complex, mitochondrial | 0.040281296 | -0.330105464 | Cellular metabolism |
| P37108 | SRP14 | Signal recognition particle 14 kDa protein | -0.005352656 | -0.207539876 | Transport |

| | | | | | |
|---|---|---|---|---|---|
| P37198 | NUP62 | Nuclear pore glycoprotein p62 | -0.290235519 | 0.650306702 | Transport |
| P37235 | HPCAL1 | Hippocalcin-like protein 1 | -0.299690247 | -0.364754995 | Transport |
| P37802 | TAGLN2 | Transgelin-2 | -0.028550466 | -0.282978694 | Other |
| P37837 | TALDO1 | Transaldolase | -0.06737264 | -0.392591476 | Cellular metabolism |
| P38117-2 | ETFB | Electron transfer flavoprotein subunit beta | -0.397374471 | -0.338785807 | Cellular metabolism |
| P38159 | RBMX | RNA-binding motif protein, X chromosome | 0.350001653 | 0.29446729 | Cellular metabolism |
| P38432 | COIL | Coilin | -0.417013804 | -0.105602264 | Other |
| P38435 | GGCX | Vitamin K-dependent gamma-carboxylase | -0.105794271 | 0 | Cellular metabolism |
| P38606 | ATP6V1A | V-type proton ATPase catalytic subunit A | 0.05385526 | -0.298770905 | Transport |
| P38919 | EIF4A3 | Eukaryotic initiation factor 4A-III | 0.005283991 | 0.662281672 | Transport |
| P39019 | RPS19 | 40S ribosomal protein S19 | -0.089427948 | -0.08937645 | Transport |
| P39023 | RPL3 | 60S ribosomal protein L3 | 0.319084803 | 0.745928446 | Transport |
| P39748 | FEN1 | Flap endonuclease 1 | 0.085911433 | -0.1982282 | Cellular metabolism |
| P40121 | CAPG | Macrophage-capping protein | -0.253576914 | -0.449881872 | Other |
| P40123 | CAP2 | Adenylyl cyclase-associated protein 2 | 0.066307704 | -0.55598259 | Other |
| P40222 | TXLNA | Alpha-taxilin | 0.384724935 | -1.261660894 | Transport |
| P40227 | CCT6A | T-complex protein 1 subunit zeta | 0.096096675 | 0.233898163 | Cellular metabolism |
| P40261 | NNMT | Nicotinamide N-methyltransferase | -0.295256933 | -1.282538732 | Cellular metabolism |
| P40337-3 | VHL | Von Hippel-Lindau disease tumor suppressor | -0.419044495 | -2.108106613 | Cellular metabolism |
| P40429 | RPL13A | 60S ribosomal protein L13a | 0.618082682 | 0.218980789 | Transport |
| P40616-2 | ARL1 | ADP-ribosylation factor-like protein 1 | -0.433324178 | -0.370016098 | Transport |
| P40925 | MDH1 | Malate dehydrogenase, cytoplasmic | -0.086137136 | -0.484018326 | Cellular metabolism |
| P40926 | MDH2 | Malate dehydrogenase, mitochondrial | -0.001423518 | 0.318735758 | Cellular metabolism |
| P40937 | RFC5 | Replication factor C subunit 5 | 0.103432337 | 0.484595617 | Cellular metabolism |
| P40938 | RFC3 | Replication factor C subunit 3 | 0.69745636 | 0 | Cellular metabolism |
| P41091 | EIF2S3 | Eukaryotic translation initiation factor 2 subunit 3 | 0.133483251 | -0.199333827 | Transport |
| P41208 | CETN2 | Centrin-2 | -0.480319977 | -3.748852412 | Cellular metabolism |
| P41223 | BUD31 | Protein BUD31 homolog | 0.05569013 | -1.146572749 | Cellular metabolism |
| P41227 | NAA10 | N-alpha-acetyltransferase 10 | -0.083300908 | -0.433504105 | Cellular metabolism |
| P41229 | KDM5C | Lysine-specific demethylase 5C | -1.42883873 | -1.511946996 | Cellular metabolism |
| P41236 | PPP1R2 | Protein phosphatase inhibitor 2 | 0.019215902 | -0.45764478 | Cellular metabolism |
| P41240 | CSK | Tyrosine-protein kinase CSK | -0.073898951 | -0.565446218 | Immune response |
| P41250-2 | GARS | Glycine--tRNA ligase | 0.033043543 | -0.650126139 | Cellular metabolism |
| P41252 | IARS | Isoleucine--tRNA ligase, cytoplasmic | 0.011583328 | -0.350962957 | Cellular metabolism |

| | | | | | |
|---|---|---|---|---|---|
| P41743 | PRKCI | Protein kinase C iota type | -0.323188782 | -1.470340729 | Transport |
| P42025 | ACTR1B | Beta-centractin | 0.495030721 | 0 | Other |
| P42126-2 | ECI1 | Enoyl-CoA delta isomerase 1, mitochondrial | -0.207078298 | -0.268451055 | Lipid metabolism |
| P42166 | TMPO | Lamina-associated polypeptide 2, isoform alpha | 0.171389898 | 0.164284388 | Other |
| P42167 | TMPO | Lamina-associated polypeptide 2, isoforms beta/gamma | 0.225232442 | 0.306526184 | Other |
| P42285 | SKIV2L2 | Superkiller viralicidic activity 2-like 2 | 0.20530955 | -0.002800624 | Cellular metabolism |
| P42357 | HAL | Histidine ammonia-lyase | -0.312458038 | 0.51300176 | Cellular metabolism |
| P42566 | EPS15 | Epidermal growth factor receptor substrate 15 | -0.066340764 | -0.802587509 | Transport |
| P42574 | CASP3 | Caspase-3 | -0.060388565 | -0.288795471 | Cellular metabolism |
| P42575 | CASP2 | Caspase-2 | -0.870607376 | -1.491826375 | Other |
| P42677 | RPS27 | 40S ribosomal protein S27 | 0.181711197 | 0.07084465 | Transport |
| P42704 | LRPPRC | Leucine-rich PPR motif-containing protein, mitochondrial | 9.53674E-06 | 0.1861763 | Lipid metabolism |
| P42766 | RPL35 | 60S ribosomal protein L35 | 0.483846029 | 0.504966736 | Transport |
| P43121 | MCAM | Cell surface glycoprotein MUC18 | 0 | -0.315017064 | Other |
| P43246 | MSH2 | DNA mismatch repair protein Msh2 | -0.177089691 | -0.357175191 | Immune response |
| P43304 | GPD2 | Glycerol-3-phosphate dehydrogenase, mitochondrial | 0 | 1.444108963 | Lipid metabolism |
| P43487-2 | RANBP1 | Ran-specific GTPase-activating protein | -0.033123652 | -0.533268611 | Transport |
| P43686 | PSMC4 | 26S protease regulatory subunit 6B | 0.050455093 | -0.123076757 | Immune response |
| P43897 | TSFM | Elongation factor Ts, mitochondrial | -0.023817062 | 0.382476171 | Cellular metabolism |
| P45877 | PPIC | Peptidyl-prolyl cis-trans isomerase C | -0.010035197 | 0.804335912 | Cellular metabolism |
| P45973 | CBX5 | Chromobox protein homolog 5 | -0.530015945 | -0.974937439 | Other |
| P45974-2 | USP5 | Ubiquitin carboxyl-terminal hydrolase 5 | -0.077328364 | -0.47539711 | Cellular metabolism |
| P45983-3 | MAPK8 | Mitogen-activated protein kinase 8 | -0.504684448 | 1.214506785 | Immune response |
| P45984 | MAPK9 | Mitogen-activated protein kinase 9 | 0.045070648 | 0 | Immune response |
| P45985-2 | MAP2K4 | Dual specificity mitogen-activated protein kinase kinase 4 | -0.104656855 | 0.608720144 | Immune response |
| P46013-2 | MKI67 | Antigen KI-67 | 0.120980581 | -1.870936076 | Other |
| P46060 | RANGAP1 | Ran GTPase-activating protein 1 | -0.115351359 | -0.611209869 | Other |
| P46063 | RECQL | ATP-dependent DNA helicase Q1 | 0.169496536 | 0.188806534 | Cellular metabolism |
| P46087-2 | NOP2 | Probable 28S rRNA (cytosine(4447)-C(5))-methyltransferase | 1.151378632 | 0.455928167 | Cellular metabolism |
| P46108 | CRK | Adapter molecule crk | -0.069401423 | -0.667792638 | Immune response |
| P46109 | CRKL | Crk-like protein | -0.129769643 | -1.961193085 | Cellular metabolism |
| P46734 | MAP2K3 | Dual specificity mitogen-activated protein kinase kinase 3 | -0.222306569 | -0.5225811 | Immune response |
| P46736-2 | BRCC3 | Lys-63-specific deubiquitinase BRCC36 | -0.284926732 | -0.27977562 | Cellular metabolism |
| P46776 | RPL27A | 60S ribosomal protein L27a | 0.319046656 | 0.141231537 | Transport |
| P46777 | RPL5 | 60S ribosomal protein L5 | -0.379762014 | 0.097206116 | Transport |

| | | | | | |
|---|---|---|---|---|---|
| P46778 | RPL21 | 60S ribosomal protein L21 | 0.337846756 | 0.385783513 | Transport |
| P46781 | RPS9 | 40S ribosomal protein S9 | 0.245849609 | 0.199239731 | Transport |
| P46783 | RPS10 | 40S ribosomal protein S10 | 0.102689107 | -3.850254059 | Transport |
| P46821 | MAP1B | Microtubule-associated protein 1B | 0.226787567 | -0.602946599 | Transport |
| P46926 | GNPDA1 | Glucosamine-6-phosphate isomerase 1 | -0.116183599 | -0.312036514 | Cellular metabolism |
| P46934-4 | NEDD4 | E3 ubiquitin-protein ligase NEDD4 | -0.232285182 | -0.920423508 | Immune response |
| P46937-9 | YAP1 | Transcriptional coactivator YAP1 | -0.461809158 | -0.767725627 | Transport |
| P46940 | IQGAP1 | Ras GTPase-activating-like protein IQGAP1 | -0.049964905 | -0.488395055 | Cellular metabolism |
| P46977 | STT3A | Dolichyl-diphosphooligosaccharide--protein glycosyltransferase subunit STT3A | 0.402932485 | 0.848255157 | Cellular metabolism |
| P47224 | RABIF | Guanine nucleotide exchange factor MSS4 | 0.021909078 | 0.59409523 | Transport |
| P47755 | CAPZA2 | F-actin-capping protein subunit alpha-2 | -0.191932678 | -0.299397151 | Immune response |
| P47756-2 | CAPZB | F-actin-capping protein subunit beta | -0.09353447 | -0.624158859 | Other |
| P47813 | EIF1AX | Eukaryotic translation initiation factor 1A, X-chromosomal | 0.157335917 | -0.18724823 | Cellular metabolism |
| P47895 | ALDH1A3 | Aldehyde dehydrogenase family 1 member A3 | 0.261078517 | -1.233045578 | Lipid metabolism |
| P47897 | QARS | Glutamine--tRNA ligase | -0.009919484 | -0.332560221 | Cellular metabolism |
| P47929 | LGALS7 | Galectin-7 | -1.176610947 | 1.942367554 | Other |
| P47985 | UQCRFS1 | Cytochrome b-c1 complex subunit Rieske, mitochondrial | 0 | 0.686604818 | Cellular metabolism |
| P48047 | ATP5O | ATP synthase subunit O, mitochondrial | 0 | 0.977700551 | Transport |
| P48059 | LIMS1 | LIM and senescent cell antigen-like-containing domain protein 1 | -0.093992233 | -0.064158122 | Other |
| P48147 | PREP | Prolyl endopeptidase | 0.022104899 | -0.538152059 | Other |
| P48163 | ME1 | NADP-dependent malic enzyme | -0.125019709 | -0.247425079 | Lipid metabolism |
| P48200 | IREB2 | Iron-responsive element-binding protein 2 | 0.02863884 | 0 | Transport |
| P48382-2 | RFX5 | DNA-binding protein RFX5 | -0.025135676 | 0 | Cellular metabolism |
| P48444 | ARCN1 | Coatomer subunit delta | 0.188404083 | -0.517634074 | Transport |
| P48454-2 | PPP3CC | Serine/threonine-protein phosphatase 2B catalytic subunit gamma isoform | 0.584439596 | 0 | Other |
| P48506 | GCLC | Glutamate--cysteine ligase catalytic subunit | -0.206766129 | -1.143103282 | Cellular metabolism |
| P48507 | GCLM | Glutamate--cysteine ligase regulatory subunit | 0.123699824 | -0.610862732 | Cellular metabolism |
| R4GMR5 | PSMD8 | 26S proteasome non-ATPase regulatory subunit 8 | 0.102111181 | 0.219328562 | Immune response |
| P48634 | PRRC2A | Protein PRRC2A | 0.991449992 | -1.553694407 | Other |
| P48637 | GSS | Glutathione synthetase | -0.083167394 | -0.51797994 | Cellular metabolism |
| P48643 | CCT5 | T-complex protein 1 subunit epsilon | -0.001829147 | 0.040838242 | Cellular metabolism |
| P48681 | NES | Nestin | 0.057402929 | -1.586169561 | Other |
| P48723 | HSPA13 | Heat shock 70 kDa protein 13 | -0.696592331 | 0.281387965 | Other |
| P48735 | IDH2 | Isocitrate dehydrogenase [NADP], mitochondrial | -0.239156723 | 0.41725413 | Cellular metabolism |

| | | | | | |
|---|---|---|---|---|---|
| P49006 | MARCKSL1 | MARCKS-related protein | -1.646941503 | -0.0122509 | Other |
| P49137 | MAPKAPK2 | MAP kinase-activated protein kinase 2 | -0.821804047 | 0.856023153 | Immune response |
| P49189 | ALDH9A1 | 4-trimethylaminobutyraldehyde dehydrogenase | -0.103851318 | -0.404752096 | Cellular metabolism |
| P49207 | RPL34 | 60S ribosomal protein L34 | 0.416467031 | 0.629387538 | Transport |
| P49247 | RPIA | Ribose-5-phosphate isomerase | 0.128143946 | 0 | Cellular metabolism |
| P49257 | LMAN1 | Protein ERGIC-53 | 0.65900294 | 1.445659002 | Transport |
| P49321 | NASP | Nuclear autoantigenic sperm protein | -0.111244837 | -0.252456665 | Transport |
| P49327 | FASN | Fatty acid synthase | -0.052357992 | -0.417507807 | Lipid metabolism |
| P49354 | FNTA | Protein farnesyltransferase/geranylgeranyltransferase type-1 subunit alpha | -0.108118693 | 0.050022761 | Cellular metabolism |
| P49366 | DHPS | Deoxyhypusine synthase | -0.529558818 | -0.966677348 | Cellular metabolism |
| P49368 | CCT3 | T-complex protein 1 subunit gamma | 0.125766118 | -0.029007594 | Cellular metabolism |
| P49411 | TUFM | Elongation factor Tu, mitochondrial | -0.141324361 | 0.49181811 | Cellular metabolism |
| P49418-2 | AMPH | Amphiphysin | -0.554852168 | 0 | Transport |
| P49419-2 | ALDH7A1 | Alpha-aminoadipic semialdehyde dehydrogenase | -0.07796224 | -0.626056035 | Cellular metabolism |
| P49427 | CDC34 | Ubiquitin-conjugating enzyme E2 R1 | -0.381438573 | -0.772567113 | Cellular metabolism |
| P49441 | INPP1 | Inositol polyphosphate 1-phosphatase | -0.156088511 | -1.314869563 | Lipid metabolism |
| P49458 | SRP9 | Signal recognition particle 9 kDa protein | 0.304590861 | 0.05791537 | Transport |
| P49459 | UBE2A | Ubiquitin-conjugating enzyme E2 A | -0.516333898 | 0.5877889 | Cellular metabolism |
| P49589-3 | CARS | Cysteine--tRNA ligase, cytoplasmic | -0.089033127 | -0.367202759 | Cellular metabolism |
| Q5T5C7 | SARS | Serine--tRNA ligase, cytoplasmic | 0.03578186 | -1.517351786 | Cellular metabolism |
| P49593 | PPM1F | Protein phosphatase 1F | -0.118805567 | 0.030273438 | Cellular metabolism |
| P49662-2 | CASP4 | Caspase-4 | -0.19052124 | 0.155250549 | Other |
| P49720 | PSMB3 | Proteasome subunit beta type-3 | 0.198663076 | 0.350115458 | Immune response |
| P49721 | PSMB2 | Proteasome subunit beta type-2 | 0.217485428 | 0.011458715 | Immune response |
| P49736 | MCM2 | DNA replication licensing factor MCM2 | -0.022219976 | -0.633330027 | Cellular metabolism |
| P49748-2 | ACADVL | Very long-chain specific acyl-CoA dehydrogenase, mitochondrial | 0.808929443 | 1.0840861 | Lipid metabolism |
| P49750-1 | YLPM1 | YLP motif-containing protein 1 | 0.351010005 | 1.052110672 | Other |
| P49755 | TMED10 | Transmembrane emp24 domain-containing protein 10 | -1.269621531 | -0.914955775 | Transport |
| P49756 | RBM25 | RNA-binding protein 25 | 0.562543233 | 0.314186096 | Cellular metabolism |
| P49770 | EIF2B2 | Translation initiation factor eIF-2B subunit beta | -0.186174393 | -1.304087321 | Cellular metabolism |
| P49773 | HINT1 | Histidine triad nucleotide-binding protein 1 | -0.081041972 | -0.450025558 | Cellular metabolism |
| P49790 | NUP153 | Nuclear pore complex protein Nup153 | 0.127330144 | 0.213202794 | Transport |
| P49792 | RANBP2 | E3 SUMO-protein ligase RanBP2 | -0.057754517 | -0.748320897 | Transport |
| P49840 | GSK3A | Glycogen synthase kinase-3 alpha | 0.100153605 | -0.258556366 | Immune response |
| P49841-2 | GSK3B | Glycogen synthase kinase-3 beta | 0.356571198 | -1.85532252 | Immune response |

| | | | | | |
|---|---|---|---|---|---|
| P49903 | SEPHS1 | Selenide, water dikinase 1 | 0.225136439 | 0.033281326 | Cellular metabolism |
| P49914 | MTHFS | 5-formyltetrahydrofolate cyclo-ligase | 0.057523092 | -1.155046463 | Cellular metabolism |
| P49915 | GMPS | GMP synthase [glutamine-hydrolyzing] | -0.109359741 | -0.318659465 | Cellular metabolism |
| P49959 | MRE11A | Double-strand break repair protein MRE11A | 0.421302795 | 1.587370555 | Immune response |
| P50395 | GDI2 | Rab GDP dissociation inhibitor beta | -0.265125275 | -0.391108831 | Transport |
| P50402 | EMD | Emerin | 0.83398056 | 0.484688441 | Other |
| P50452 | SERPINB8 | Serpin B8 | -0.254013062 | -0.46387736 | Other |
| P50453 | SERPINB9 | Serpin B9 | -0.313109716 | -0.745102564 | Immune response |
| P50454 | SERPINH1 | Serpin H1 | 0.488565445 | 0.963797251 | Other |
| P50479 | PDLIM4 | PDZ and LIM domain protein 4 | 0.068150838 | -1.280167262 | Other |
| P50502 | ST13 | Hsc70-interacting protein | -0.03933843 | -0.676432927 | Cellular metabolism |
| P50548 | ERF | ETS domain-containing transcription factor ERF | 0.25812912 | 0 | Cellular metabolism |
| P50552 | VASP | Vasodilator-stimulated phosphoprotein | 0.11127917 | -3.468651454 | Other |
| P50570-2 | DNM2 | Dynamin-2 | 0.235432307 | -0.193303426 | Transport |
| P50579 | METAP2 | Methionine aminopeptidase 2 | 0.549671173 | -0.180388133 | Cellular metabolism |
| P50583 | NUDT2 | Bis(5-nucleosyl)-tetraphosphatase [asymmetrical] | -0.164812088 | 0.255407333 | Cellular metabolism |
| P50613 | CDK7 | Cyclin-dependent kinase 7 | 0.002613068 | -0.629605611 | Cellular metabolism |
| P50990 | CCT8 | T-complex protein 1 subunit theta | 0.031175613 | -0.016838074 | Cellular metabolism |
| P50991 | CCT4 | T-complex protein 1 subunit delta | 0.072601318 | 0.121546427 | Cellular metabolism |
| P50995-2 | ANXA11 | Annexin A11 | -0.143977483 | -0.380537033 | Transport |
| P51003 | PAPOLA | Poly(A) polymerase alpha | -0.078196843 | -1.583894094 | Cellular metabolism |
| P51116 | FXR2 | Fragile X mental retardation syndrome-related protein 2 | 0.46540451 | 1.12009112 | Other |
| P51148 | RAB5C | Ras-related protein Rab-5C | 0.260230382 | 2.468084335 | Transport |
| P51149 | RAB7A | Ras-related protein Rab-7a | -0.07423083 | 1.917981466 | Transport |
| P51153 | RAB13 | Ras-related protein Rab-13 | -1.174352646 | 2.69669342 | Transport |
| P51157 | RAB28 | Ras-related protein Rab-28 | -0.081804911 | 0 | Other |
| P51159 | RAB27A | Ras-related protein Rab-27A | -0.241861343 | 3.703074773 | Transport |
| P51178 | PLCD1 | 1-phosphatidylinositol 4,5-bisphosphate phosphodiesterase delta-1 | -0.19216156 | 1.755260468 | Lipid metabolism |
| P51398-2 | DAP3 | 28S ribosomal protein S29, mitochondrial | -0.861623128 | -0.160399755 | Cellular metabolism |
| P51452 | DUSP3 | Dual specificity protein phosphatase 3 | -0.300321579 | -0.875617981 | Immune response |
| P51570 | GALK1 | Galactokinase | -0.231840769 | -1.480703354 | Cellular metabolism |
| P51571 | SSR4 | Translocon-associated protein subunit delta | 0.897940954 | 0.792267481 | Transport |
| P51572 | BCAP31 | B-cell receptor-associated protein 31 | 1.298712413 | 0.693843206 | Transport |
| P51580 | TPMT | Thiopurine S-methyltransferase | -0.283574422 | -1.125717163 | Cellular metabolism |
| P51659 | HSD17B4 | Peroxisomal multifunctional enzyme type 2 | 1.466173808 | 0.953257879 | Lipid metabolism |

| | | | | | |
|---|---|---|---|---|---|
| P51665 | PSMD7 | 26S proteasome non-ATPase regulatory subunit 7 | 0.251845678 | 0.044005712 | Immune response |
| P51688 | SGSH | N-sulphoglucosamine sulphohydrolase | 0.371000926 | 0.274330139 | Cellular metabolism |
| P51692 | STAT5B | Signal transducer and activator of transcription 5B | -0.29138883 | -1.334819794 | Lipid metabolism |
| P51812 | RPS6KA3 | Ribosomal protein S6 kinase alpha-3 | -0.429146449 | -2.055088043 | Immune response |
| P51858 | HDGF | Hepatoma-derived growth factor | 0.052095413 | -0.756434759 | Cellular metabolism |
| P51911 | CNN1 | Calponin-1 | 0.292701085 | 0.698159536 | Other |
| P51970 | NDUFA8 | NADH dehydrogenase [ubiquinone] 1 alpha subcomplex subunit 8 | -0.293223063 | -0.132738113 | Cellular metabolism |
| P52209-2 | PGD | 6-phosphogluconate dehydrogenase, decarboxylating | -0.08993721 | -0.29842186 | Cellular metabolism |
| P52272-2 | HNRNPM | Heterogeneous nuclear ribonucleoprotein M | 0.648593267 | 0.353360494 | Cellular metabolism |
| P52294 | KPNA1 | Importin subunit alpha-5 | 0.018112183 | -0.2961483 | Transport |
| P52298 | NCBP2 | Nuclear cap-binding protein subunit 2 | 0.180060069 | -0.505530039 | Transport |
| P52306 | RAP1GDS1 | Rap1 GTPase-GDP dissociation stimulator 1 | -0.013654073 | -0.644838969 | Other |
| P52565 | ARHGDIA | Rho GDP-dissociation inhibitor 1 | -0.195573171 | -0.044826508 | Other |
| P52594-2 | AGFG1 | Arf-GAP domain and FG repeat-containing protein 1 | -0.300198873 | -1.201911926 | Transport |
| P52597 | HNRNPF | Heterogeneous nuclear ribonucleoprotein F | 0.113968531 | 0.074901581 | Cellular metabolism |
| P52657 | GTF2A2 | Transcription initiation factor IIA subunit 2 | 0.067929586 | 1.147472382 | Cellular metabolism |
| P52701 | MSH6 | DNA mismatch repair protein Msh6 | -0.186236699 | -0.689287186 | Cellular metabolism |
| P52732 | KIF11 | Kinesin-like protein KIF11 | -0.081045787 | -0.516263962 | Other |
| P52735-3 | VAV2 | Guanine nucleotide exchange factor VAV2 | 0.238729477 | -0.490029653 | Immune response |
| P52758 | HRSP12 | Ribonuclease UK114 | -0.234194438 | -0.021452586 | Other |
| P52788 | SMS | Spermine synthase | -0.279531479 | -0.965247472 | Cellular metabolism |
| P52789 | HK2 | Hexokinase-2 | 0.103822072 | -0.279482524 | Transport |
| P52888 | THOP1 | Thimet oligopeptidase | -0.000204086 | -0.660931905 | Cellular metabolism |
| P52907 | CAPZA1 | F-actin-capping protein subunit alpha-1 | 0.010161082 | -0.672048569 | Immune response |
| P52943 | CRIP2 | Cysteine-rich protein 2 | 0.456436793 | 0.016186396 | Other |
| P52948-6 | NUP98 | Nuclear pore complex protein Nup98-Nup96 | -0.051602681 | 0.312240601 | Transport |
| P53004 | BLVRA | Biliverdin reductase A | -0.08940951 | -0.189296722 | Cellular metabolism |
| P53007 | SLC25A1 | Tricarboxylate transport protein, mitochondrial | 0 | 0.130597432 | Lipid metabolism |
| P53350 | PLK1 | Serine/threonine-protein kinase PLK1 | -0.351282756 | 0 | Cellular metabolism |
| P53367 | ARFIP1 | Arfaptin-1 | 0.051079432 | 0.966295242 | Transport |
| P53384-2 | NUBP1 | Cytosolic Fe-S cluster assembly factor NUBP1 | 0.018050512 | -0.539017995 | Cellular metabolism |
| P53396 | ACLY | ATP-citrate synthase | 0.060068766 | -1.025344213 | Lipid metabolism |
| P53582 | METAP1 | Methionine aminopeptidase 1 | -0.087799708 | -1.016270955 | Cellular metabolism |
| P53597 | SUCLG1 | Succinyl-CoA ligase [ADP/GDP-forming] subunit alpha, mitochondrial | -0.111499786 | -1.405073802 | Cellular metabolism |
| P53602 | MVD | Diphosphomevalonate decarboxylase | -0.446663539 | -3.4161129 | Lipid metabolism |

| | | | | | |
|---|---|---|---|---|---|
| P53609 | PGGT1B | Geranylgeranyl transferase type-1 subunit beta | -0.212621689 | -0.127681096 | Cellular metabolism |
| P53611 | RABGGTB | Geranylgeranyl transferase type-2 subunit beta | -0.196987152 | -0.678771337 | Cellular metabolism |
| P53618 | COPB1 | Coatomer subunit beta | 0.090921402 | -0.020524979 | Transport |
| P53621 | COPA | Coatomer subunit alpha | 0.01802063 | -0.550120672 | Transport |
| P53677 | AP3M2 | AP-3 complex subunit mu-2 | 0.003595352 | 0 | Transport |
| P53701 | HCCS | Cytochrome c-type heme lyase | 0 | 0.918928782 | Other |
| P53778-2 | MAPK12 | Mitogen-activated protein kinase 12 | 0.124537786 | -1.446791967 | Cellular metabolism |
| P53803 | POLR2K | DNA-directed RNA polymerases I, II, and III subunit RPABC4 | -1.245697021 | 0 | Immune response |
| P53814-5 | SMTN | Smoothelin | -0.031284332 | -0.113150915 | Other |
| P53985 | SLC16A1 | Monocarboxylate transporter 1 | 0 | 0.841920853 | Lipid metabolism |
| P53990-2 | IST1 | IST1 homolog | -0.310974757 | 0.348866145 | Other |
| P53992 | SEC24C | Protein transport protein Sec24C | -0.361932755 | -2.142880758 | Transport |
| P53999 | SUB1 | Activated RNA polymerase II transcriptional coactivator p15 | -1.387250264 | -0.283248266 | Other |
| P54105 | CLNS1A | Methylosome subunit pICln | -0.247790655 | -2.382225037 | Transport |
| P54136 | RARS | Arginine--tRNA ligase, cytoplasmic | 0.02031072 | -0.366224925 | Cellular metabolism |
| P54577 | YARS | Tyrosine--tRNA ligase, cytoplasmic | -0.041413625 | -0.502999624 | Cellular metabolism |
| P54578 | USP14 | Ubiquitin carboxyl-terminal hydrolase 14 | -0.163301468 | -0.568197886 | Cellular metabolism |
| P54619-2 | PRKAG1 | 5-AMP-activated protein kinase subunit gamma-1 | -0.227518717 | 0.039563497 | Lipid metabolism |
| P54652 | HSPA2 | Heat shock-related 70 kDa protein 2 | 0.003715515 | -0.083960851 | Cellular metabolism |
| P54687 | BCAT1 | Branched-chain-amino-acid aminotransferase, cytosolic | 0.012339274 | -0.996311188 | Cellular metabolism |
| P54709 | ATP1B3 | Sodium/potassium-transporting ATPase subunit beta-3 | 0.468789419 | -0.752600352 | Transport |
| P54725-3 | RAD23A | UV excision repair protein RAD23 homolog A | -0.358390172 | -0.632567724 | Cellular metabolism |
| P54727 | RAD23B | UV excision repair protein RAD23 homolog B | -0.074372609 | -0.289515177 | Cellular metabolism |
| P54802 | NAGLU | Alpha-N-acetylglucosaminidase | 0.596621195 | 0 | Other |
| P54886-2 | ALDH18A1 | Delta-1-pyrroline-5-carboxylate synthase | -0.032399495 | 0.999711355 | Cellular metabolism |
| P54920 | NAPA | Alpha-soluble NSF attachment protein | -0.098900477 | 0.815437953 | Transport |
| P55010 | EIF5 | Eukaryotic translation initiation factor 5 | 0.081667582 | -0.326149623 | Cellular metabolism |
| P55036 | PSMD4 | 26S proteasome non-ATPase regulatory subunit 4 | 0.028399785 | -0.22996521 | Immune response |
| P55039 | DRG2 | Developmentally-regulated GTP-binding protein 2 | -0.324150721 | -1.097040812 | Other |
| P55042 | RRAD | GTP-binding protein RAD | 0.174945831 | 0.703726451 | Immune response |
| P55060-3 | CSE1L | Exportin-2 | -0.051608404 | -0.264287313 | Transport |
| P55081 | MFAP1 | Microfibrillar-associated protein 1 | -0.132802327 | 0.113710403 | Other |
| P55084-2 | HADHB | Trifunctional enzyme subunit beta, mitochondrial | 0.687308629 | 0.393512726 | Lipid metabolism |
| P55145 | MANF | Mesencephalic astrocyte-derived neurotrophic factor | 0.32114474 | 0.064634323 | Other |

| | | | | | |
|---|---|---|---|---|---|
| P55210 | CASP7 | Caspase-7 | -0.098510742 | 0.166542689 | Other |
| P55212 | CASP6 | Caspase-6 | -0.397376378 | 0.308556239 | Other |
| P55263 | ADK | Adenosine kinase | -0.044054667 | -0.718346278 | Cellular metabolism |
| P55327-2 | TPD52 | Tumor protein D52 | 0.146198908 | 1.685042063 | Transport |
| P55735 | SEC13 | Protein SEC13 homolog | -0.007535299 | -0.320501963 | Transport |
| P55769 | NHP2L1 | NHP2-like protein 1 | 0.152073542 | 0.010445913 | Cellular metabolism |
| P55795 | HNRNPH2 | Heterogeneous nuclear ribonucleoprotein H2 | 0.523384094 | 1.073582967 | Cellular metabolism |
| P55809 | OXCT1 | Succinyl-CoA:3-ketoacid coenzyme A transferase 1, mitochondrial | -0.110680898 | -0.378742218 | Lipid metabolism |
| P55884 | EIF3B | Eukaryotic translation initiation factor 3 subunit B | 0.136334737 | -0.380045573 | Cellular metabolism |
| P55957 | BID | BH3-interacting domain death agonist | 0.111324946 | 0.168651581 | Transport |
| P56182 | RRP1 | Ribosomal RNA processing protein 1 homolog A | 1.066820145 | 0.461503347 | Cellular metabolism |
| P56192 | MARS | Methionine--tRNA ligase, cytoplasmic | 0.295166651 | -0.290116628 | Cellular metabolism |
| P56211 | ARPP19 | cAMP-regulated phosphoprotein 19 | 0 | 0.094863256 | Other |
| P56377 | AP1S2 | AP-1 complex subunit sigma-2 | -0.397951762 | 0 | Transport |
| P56537 | EIF6 | Eukaryotic translation initiation factor 6 | -0.153530121 | -0.078002294 | Transport |
| P56545 | CTBP2 | C-terminal-binding protein 2 | -0.170007706 | -1.000912984 | Cellular metabolism |
| P56945-6 | BCAR1 | Breast cancer anti-estrogen resistance protein 1 | -0.286079407 | 1.204711278 | Other |
| P57081 | WDR4 | tRNA (guanine-N(7)-)-methyltransferase non-catalytic subunit WDR4 | 0.096024195 | -1.28768158 | Cellular metabolism |
| P57740 | NUP107 | Nuclear pore complex protein Nup107 | -0.596717199 | 0.014822006 | Transport |
| P57768-2 | SNX16 | Sorting nexin-16 | -0.525444667 | 0 | Transport |
| P58004 | SESN2 | Sestrin-2 | -1.138678233 | -1.785134633 | Lipid metabolism |
| P58546 | MTPN | Myotrophin | -0.041189194 | 0.05921491 | Cellular metabolism |
| P60059 | SEC61G | Protein transport protein Sec61 subunit gamma | 0 | 0.40801239 | Transport |
| P60174 | TPI1 | Triosephosphate isomerase | -0.035827637 | -0.234560649 | Cellular metabolism |
| P60370 | KRTAP10-5 | Keratin-associated protein 10-5 | 0 | -1.253166835 | Other |
| P60468 | SEC61B | Protein transport protein Sec61 subunit beta | 0.72631518 | 1.259853999 | Transport |
| P60484 | PTEN | Phosphatidylinositol 3,4,5-trisphosphate 3-phosphatase and dual-specificity protein phosphatase PTEN | 0.470080058 | -1.627058665 | Immune response |
| P60510 | PPP4C | Serine/threonine-protein phosphatase 4 catalytic subunit | -0.217116674 | -0.752965927 | Cellular metabolism |
| P60520 | GABARAPL2 | Gamma-aminobutyric acid receptor-associated protein-like 2 | -0.225582123 | -0.647469203 | Transport |
| P60604-2 | UBE2G2 | Ubiquitin-conjugating enzyme E2 G2 | 0.960804621 | 3.740049362 | Cellular metabolism |
| P60660-2 | MYL6 | Myosin light polypeptide 6 | -0.053368886 | -0.444484711 | Other |
| P60709 | ACTB | Actin, cytoplasmic 1 | -0.078830719 | -0.774394989 | Immune response |
| P60842 | EIF4A1 | Eukaryotic initiation factor 4A-I | -0.029436747 | -0.38846461 | Cellular metabolism |
| P60866 | RPS20 | 40S ribosomal protein S20 | 0.253991445 | -0.410601298 | Transport |

| | | | | | |
|---|---|---|---|---|---|
| P60900 | PSMA6 | Proteasome subunit alpha type-6 | 0.209046046 | 0.404397329 | Immune response |
| P60903 | S100A10 | Protein S100-A10 | -1.108111699 | 0 | Other |
| P60953 | CDC42 | Cell division control protein 42 homolog | -0.629343033 | -0.516330083 | Immune response |
| P60981 | DSTN | Destrin | -0.164293289 | -0.929128011 | Other |
| P60983 | GMFB | Glia maturation factor beta | -0.090712865 | -0.925833384 | Cellular metabolism |
| P61009 | SPCS3 | Signal peptidase complex subunit 3 | -0.513526917 | 1.5669384 | Transport |
| P61011-2 | SRP54 | Signal recognition particle 54 kDa protein | 0.493019104 | -0.524920781 | Transport |
| P61019 | RAB2A | Ras-related protein Rab-2A | 0.261918386 | 2.038485845 | Transport |
| P61020 | RAB5B | Ras-related protein Rab-5B | 0.076665878 | 3.786727269 | Transport |
| P61024 | CKS1B | Cyclin-dependent kinases regulatory subunit 1 | -0.242295583 | 0 | Other |
| P61081 | UBE2M | NEDD8-conjugating enzyme Ubc12 | 0.002490997 | -0.468357722 | Immune response |
| P61086 | UBE2K | Ubiquitin-conjugating enzyme E2 K | -0.171789805 | -0.254977544 | Cellular metabolism |
| P61088 | UBE2N | Ubiquitin-conjugating enzyme E2 N | 0.072535197 | 0.665193558 | Immune response |
| P61106 | RAB14 | Ras-related protein Rab-14 | -0.110120138 | 1.157325109 | Transport |
| P61158 | ACTR3 | Actin-related protein 3 | 0.269444784 | 0.065649668 | Immune response |
| P61160 | ACTR2 | Actin-related protein 2 | 0.075751623 | 0.04767863 | Immune response |
| P61163 | ACTR1A | Alpha-centractin | 0.21771876 | -1.034360886 | Transport |
| P61201 | COPS2 | COP9 signalosome complex subunit 2 | 0.172562917 | -0.096768061 | Cellular metabolism |
| P61221 | ABCE1 | ATP-binding cassette sub-family E member 1 | 0.130294164 | 0.441013336 | Transport |
| P61224 | RAP1B | Ras-related protein Rap-1b | 0.340777715 | 0.99352328 | Transport |
| P61247 | RPS3A | 40S ribosomal protein S3a | 0.193335851 | 0.04509608 | Transport |
| P61254 | RPL26 | 60S ribosomal protein L26 | 0.297279358 | 0.099822362 | Transport |
| P61289 | PSME3 | Proteasome activator complex subunit 3 | 0.076592127 | -0.676917394 | Immune response |
| P61313 | RPL15 | 60S ribosomal protein L15 | 0.246927261 | 1.100676219 | Transport |
| P61326 | MAGOH | Protein mago nashi homolog | -0.279778163 | -0.392796199 | Transport |
| P61353 | RPL27 | 60S ribosomal protein L27 | 0.469395955 | 0.896970113 | Transport |
| P61457 | PCBD1 | Pterin-4-alpha-carbinolamine dehydratase | 0.114142736 | -1.290988286 | Cellular metabolism |
| P61586 | RHOA | Transforming protein RhoA | -0.051749547 | -0.19397672 | Other |
| P61587 | RND3 | Rho-related GTP-binding protein RhoE | 0.899080912 | 1.531133652 | Other |
| P61604 | HSPE1 | 10 kDa heat shock protein, mitochondrial | -0.118525823 | -0.059837977 | Cellular metabolism |
| P61758 | VBP1 | Prefoldin subunit 3 | -0.189515432 | -0.535116831 | Cellular metabolism |
| P61769 | B2M | Beta-2-microglobulin | 0.241371791 | 1.338034948 | Immune response |
| P61956-2 | SUMO2 | Small ubiquitin-related modifier 2 | 0.024926503 | 0.007633209 | Cellular metabolism |
| P61960 | UFM1 | Ubiquitin-fold modifier 1 | -0.318058014 | -0.302270253 | Cellular metabolism |
| P61962 | DCAF7 | DDB1- and CUL4-associated factor 7 | 0.044471105 | -0.687995275 | Cellular metabolism |
| P61964 | WDR5 | WD repeat-containing protein 5 | 0.704659144 | -0.277594248 | Cellular metabolism |

| | | | | | |
|---|---|---|---|---|---|
| P61970 | NUTF2 | Nuclear transport factor 2 | -0.061524073 | 0.216536204 | Transport |
| P61978-2 | HNRNPK | Heterogeneous nuclear ribonucleoprotein K | 0.392664591 | -0.053371429 | Cellular metabolism |
| P61981 | YWHAG | 14-3-3 protein gamma | 0.161820094 | -0.351184209 | Transport |
| P62070 | RRAS2 | Ras-related protein R-Ras2 | -0.156690598 | 0.268006007 | Other |
| P62081 | RPS7 | 40S ribosomal protein S7 | 0.211263657 | 0.405677795 | Transport |
| P62136 | PPP1CA | Serine/threonine-protein phosphatase PP1-alpha catalytic subunit | 0.158128103 | -0.288110733 | Lipid metabolism |
| P62140 | PPP1CB | Serine/threonine-protein phosphatase PP1-beta catalytic subunit | 0.096178055 | 0.190338771 | Lipid metabolism |
| P62166 | NCS1 | Neuronal calcium sensor 1 | 0.50839742 | 0.784792582 | Other |
| P62191 | PSMC1 | 26S protease regulatory subunit 4 | -0.045360565 | -0.484434764 | Immune response |
| P62195 | PSMC5 | 26S protease regulatory subunit 8 | 0.11488533 | 0.034645716 | Immune response |
| P62241 | RPS8 | 40S ribosomal protein S8 | 0.200394313 | 0.393892288 | Transport |
| P62244 | RPS15A | 40S ribosomal protein S15a | 0.401095708 | -0.001205444 | Transport |
| P62249 | RPS16 | 40S ribosomal protein S16 | 0.068988164 | 0.478527069 | Transport |
| P62253 | UBE2G1 | Ubiquitin-conjugating enzyme E2 G1 | -0.359924952 | -0.680713018 | Cellular metabolism |
| P62258 | YWHAE | 14-3-3 protein epsilon | 0.131151199 | -0.33823204 | Transport |
| P62266 | RPS23 | 40S ribosomal protein S23 | 0.212802887 | 0.222012202 | Transport |
| P62269 | RPS18 | 40S ribosomal protein S18 | 0.126019796 | 0.274247487 | Transport |
| P62273 | RPS29 | 40S ribosomal protein S29 | 0.238312403 | -0.15567716 | Transport |
| P62277 | RPS13 | 40S ribosomal protein S13 | 0.190699895 | 0.325305303 | Transport |
| P62280 | RPS11 | 40S ribosomal protein S11 | 0.200508753 | 0.053845723 | Transport |
| P62304 | SNRPE | Small nuclear ribonucleoprotein E | -0.006442388 | 0.396392822 | Cellular metabolism |
| P62306 | SNRPF | Small nuclear ribonucleoprotein F | 0.415546417 | -0.348781586 | Cellular metabolism |
| P62312 | LSM6 | U6 snRNA-associated Sm-like protein LSm6 | 0.510872523 | 0.302858988 | Cellular metabolism |
| P62314 | SNRPD1 | Small nuclear ribonucleoprotein Sm D1 | 0.18576177 | -0.125419617 | Cellular metabolism |
| P62316 | SNRPD2 | Small nuclear ribonucleoprotein Sm D2 | 0.067405065 | -0.173531214 | Cellular metabolism |
| P62318 | SNRPD3 | Small nuclear ribonucleoprotein Sm D3 | 0.415718079 | -0.427706401 | Cellular metabolism |
| P62328 | TMSB4X | Thymosin beta-4 | -2.660083135 | 0.709637324 | Transport |
| P62330 | ARF6 | ADP-ribosylation factor 6 | 0 | 0.818874359 | Transport |
| P62380 | TBPL1 | TATA box-binding protein-like protein 1 | -0.528306961 | -0.787027995 | Cellular metabolism |
| P62424 | RPL7A | 60S ribosomal protein L7a | 0.286034266 | 0.25421079 | Transport |
| P62487 | POLR2G | DNA-directed RNA polymerase II subunit RPB7 | -0.42241923 | 0.050946554 | Cellular metabolism |
| P62495 | ETF1 | Eukaryotic peptide chain release factor subunit 1 | 0.163883209 | -0.154034932 | Cellular metabolism |
| P62633-2 | CNBP | Cellular nucleic acid-binding protein | 0.251593272 | -0.765068054 | Lipid metabolism |
| P62699 | YPEL5 | Protein yippee-like 5 | 0.318794886 | 0 | Other |
| P62701 | RPS4X | 40S ribosomal protein S4, X isoform | 0.239611944 | 0.31737709 | Transport |
| P62714 | PPP2CB | Serine/threonine-protein phosphatase 2A catalytic subunit beta isoform | 0.112260818 | 0.636069616 | Cellular metabolism |
| P62750 | RPL23A | 60S ribosomal protein L23a | 0.203838348 | 0.323758443 | Transport |

| | | | | | |
|---|---|---|---|---|---|
| P62753 | RPS6 | 40S ribosomal protein S6 | 0.255137761 | 0.291369756 | Transport |
| P62805 | HIST1H4A | Histone H4 | 0.169919968 | 0.554894765 | Cellular metabolism |
| P62826 | RAN | GTP-binding nuclear protein Ran | 0.080120087 | -0.499221802 | Transport |
| P62829 | RPL23 | 60S ribosomal protein L23 | 0.088199615 | -0.172281265 | Transport |
| P62851 | RPS25 | 40S ribosomal protein S25 | 0.34081459 | 0.177009583 | Transport |
| P62854 | RPS26 | 40S ribosomal protein S26 | 0.102324804 | -0.663365046 | Transport |
| P62857 | RPS28 | 40S ribosomal protein S28 | 0.108309428 | -0.318453471 | Transport |
| P62873-2 | GNB1 | Guanine nucleotide-binding protein G(I)/G(S)/G(T) subunit beta-1 | -0.081814448 | 1.061115901 | Cellular metabolism |
| P62877 | RBX1 | E3 ubiquitin-protein ligase RBX1 | -0.011436462 | -0.18807284 | Cellular metabolism |
| P62879 | GNB2 | Guanine nucleotide-binding protein G(I)/G(S)/G(T) subunit beta-2 | 0 | 1.53940328 | Cellular metabolism |
| Q59GN2 | RPL39P5 | Putative 60S ribosomal protein L39-like 5 | 0.262132009 | 1.71933492 | Immune response |
| P62906 | RPL10A | 60S ribosomal protein L10a | 0.282843908 | 0.333246231 | Transport |
| P62913 | RPL11 | 60S ribosomal protein L11 | 0.165550868 | 0.594284693 | Transport |
| P62917 | RPL8 | 60S ribosomal protein L8 | 0.456018448 | 0.345273336 | Transport |
| P62937 | PPIA | Peptidyl-prolyl cis-trans isomerase A | -0.086567561 | -0.087300618 | Transport |
| P62942 | FKBP1A | Peptidyl-prolyl cis-trans isomerase FKBP1A | -0.164825439 | -0.384717941 | Transport |
| P62979 | RPS27A | Ubiquitin-40S ribosomal protein S27a | -0.049224854 | 0.475342433 | Immune response |
| P62987 | UBA52 | Ubiquitin-60S ribosomal protein L40 | 0.625019073 | -0.302748998 | Immune response |
| P62993 | GRB2 | Growth factor receptor-bound protein 2 | -0.335350037 | -0.816229502 | Immune response |
| P62995-3 | TRA2B | Transformer-2 protein homolog beta | 0.509096146 | 0.137639364 | Cellular metabolism |
| P63000-2 | RAC1 | Ras-related C3 botulinum toxin substrate 1 | 0.00336202 | 0.538357417 | Immune response |
| P63010 | AP2B1 | AP-2 complex subunit beta | 0.110117594 | -0.20729955 | Transport |
| Q5JWF2 | GNAS | Guanine nucleotide-binding protein G(s) subunit alpha isoforms XLas | -0.371284485 | 1.138346354 | Transport |
| P63104 | YWHAZ | 14-3-3 protein zeta/delta | 0.014691671 | -0.160452525 | Transport |
| P63151 | PPP2R2A | Serine/threonine-protein phosphatase 2A 55 kDa regulatory subunit B alpha isoform | -0.192553202 | -0.181037267 | Cellular metabolism |
| P63167 | DYNLL1 | Dynein light chain 1, cytoplasmic | -0.033104579 | -1.071966807 | Transport |
| P63172 | DYNLT1 | Dynein light chain Tctex-type 1 | -0.15465037 | 0.221525828 | Transport |
| P63173 | RPL38 | 60S ribosomal protein L38 | 0.352097829 | -0.285473506 | Transport |
| P63208 | SKP1 | S-phase kinase-associated protein 1 | -0.727181117 | -0.162581126 | Immune response |
| Q8WVC2 | RPS21 | 40S ribosomal protein S21 | 0.064332962 | -0.343027115 | Transport |
| P63241 | EIF5A | Eukaryotic translation initiation factor 5A-1 | 0.03034846 | 0.593557358 | Transport |
| P63244 | GNB2L1 | Guanine nucleotide-binding protein subunit beta-2-like 1 | 0.106983185 | 0.072615306 | Other |
| P63261 | ACTG1 | Actin, cytoplasmic 2 | -0.047902425 | -0.102982839 | Immune response |
| P63272 | SUPT4H1 | Transcription elongation factor SPT4 | -0.452236176 | 0 | Cellular metabolism |
| P63313 | TMSB10 | Thymosin beta-10 | -0.385031382 | 0.314158122 | Other |

| | | | | | |
|---|---|---|---|---|---|
| P67775 | PPP2CA | Serine/threonine-protein phosphatase 2A catalytic subunit alpha isoform | -0.213546753 | -0.184346517 | Lipid metabolism |
| P67809 | YBX1 | Nuclease-sensitive element-binding protein 1 | 0.119146347 | -0.229138692 | Cellular metabolism |
| Q5SRQ6 | CSNK2B | Casein kinase II subunit beta | 0.001675288 | -0.224343618 | Cellular metabolism |
| P68032 | ACTC1 | Actin, alpha cardiac muscle 1 | 0.023840586 | -0.860971451 | Other |
| P68036 | UBE2L3 | Ubiquitin-conjugating enzyme E2 L3 | 0.196309408 | -0.575728734 | Cellular metabolism |
| P68104 | EEF1A1 | Elongation factor 1-alpha 1 | -0.021853129 | -0.120677948 | Cellular metabolism |
| P68366-2 | TUBA4A | Tubulin alpha-4A chain | -0.115557988 | -0.096572876 | Transport |
| P68371 | TUBB4B | Tubulin beta-4B chain | -0.083618164 | 0.261376063 | Immune response |
| P68402 | PAFAH1B2 | Platelet-activating factor acetylhydrolase IB subunit beta | -0.123647054 | -0.318929036 | Lipid metabolism |
| P68431 | HIST1H3A | Histone H3.1 | 0.142871857 | 0.486032486 | Cellular metabolism |
| P68871 | HBB | Hemoglobin subunit beta | 1.821652095 | 0.372554143 | Transport |
| P69905 | HBA1 | Hemoglobin subunit alpha | 2.288905462 | 0.800826391 | Transport |
| P78318 | IGBP1 | Immunoglobulin-binding protein 1 | -0.017288844 | -0.49372228 | Other |
| P78345 | RPP38 | Ribonuclease P protein subunit p38 | 2.302272797 | -0.174254735 | Cellular metabolism |
| P78346 | RPP30 | Ribonuclease P protein subunit p30 | 0.427818934 | 0.677570979 | Cellular metabolism |
| P78347-2 | GTF2I | General transcription factor II-I | 0.002833049 | -1.043950399 | Cellular metabolism |
| P78356 | PIP4K2B | Phosphatidylinositol 5-phosphate 4-kinase type-2 beta | 0.376473109 | -0.247828166 | Lipid metabolism |
| P78371 | CCT2 | T-complex protein 1 subunit beta | -0.022883097 | -0.051395416 | Cellular metabolism |
| P78406 | RAE1 | mRNA export factor | 0.080733617 | 0.518770854 | Transport |
| P78417 | GSTO1 | Glutathione S-transferase omega-1 | -0.029993693 | -0.527084351 | Cellular metabolism |
| P78527 | PRKDC | DNA-dependent protein kinase catalytic subunit | 0.368745804 | 0.301460902 | Immune response |
| P78560 | CRADD | Death domain-containing protein CRADD | -0.075386683 | 0 | Other |
| X6R8F3 | LCN2 | Neutrophil gelatinase-associated lipocalin | 0 | 1.134410222 | Immune response |
| P80217 | IFI35 | Interferon-induced 35 kDa protein | 0.817179998 | -1.449041367 | Other |
| P80303-2 | NUCB2 | Nucleobindin-2 | 0.346299489 | 0.070584615 | Other |
| P80723 | BASP1 | Brain acid soluble protein 1 | -1.934964498 | 1.237873713 | Other |
| P81605 | DCD | Dermcidin | 1.367125829 | 1.048909505 | Other |
| P82094 | TMF1 | TATA element modulatory factor | -1.201557159 | -1.335598628 | Transport |
| P82909 | MRPS36 | 28S ribosomal protein S36, mitochondrial | 0 | -1.204619726 | Cellular metabolism |
| P82933 | MRPS9 | 28S ribosomal protein S9, mitochondrial | 0.066490173 | 1.964431127 | Cellular metabolism |
| P82979 | SARNP | SAP domain-containing ribonucleoprotein | -0.27040418 | -0.126417796 | Transport |
| P83436 | COG7 | Conserved oligomeric Golgi complex subunit 7 | 0.921034495 | -1.679892222 | Transport |
| P83876 | TXNL4A | Thioredoxin-like protein 4A | -0.3348821 | -0.10193634 | Cellular metabolism |
| P83916 | CBX1 | Chromobox protein homolog 1 | 0.042248408 | -0.686384837 | Other |
| P84022 | SMAD3 | Mothers against decapentaplegic homolog 3 | -0.051270803 | -0.012479146 | Immune response |

| | | | | | |
|---|---|---|---|---|---|
| P84077 | ARF1 | ADP-ribosylation factor 1 | -0.16044871 | 0.024927139 | Lipid metabolism |
| P84085 | ARF5 | ADP-ribosylation factor 5 | -0.231899897 | -1.67280515 | Transport |
| P84090 | ERH | Enhancer of rudimentary homolog | 0.069006602 | 0.744393031 | Cellular metabolism |
| P84103 | SRSF3 | Serine/arginine-rich splicing factor 3 | 0.476453145 | 0.060333252 | Transport |
| P85037 | FOXK1 | Forkhead box protein K1 | 0.293401082 | -0.152481079 | Other |
| P86790 | CCZ1B | Vacuolar fusion protein CCZ1 homolog B | 0.237319311 | 0 | Transport |
| P98082-3 | DAB2 | Disabled homolog 2 | -0.095343272 | -0.719456355 | Transport |
| P98170 | XIAP | E3 ubiquitin-protein ligase XIAP | -0.657648087 | 0.414429983 | Cellular metabolism |
| P98175-2 | RBM10 | RNA-binding protein 10 | 0.269833247 | -0.159775416 | Cellular metabolism |
| P98179 | RBM3 | RNA-binding protein 3 | 0.392472585 | -0.200892766 | Cellular metabolism |
| Q00059 | TFAM | Transcription factor A, mitochondrial | 0 | 1.888806025 | Cellular metabolism |
| Q00169 | PITPNA | Phosphatidylinositol transfer protein alpha isoform | -0.125293732 | -0.330718994 | Lipid metabolism |
| Q00325-2 | SLC25A3 | Phosphate carrier protein, mitochondrial | -0.016527176 | 0.526585261 | Transport |
| Q00403 | GTF2B | Transcription initiation factor IIB | -0.846037547 | 0.387726466 | Cellular metabolism |
| Q00534 | CDK6 | Cyclin-dependent kinase 6 | -0.272399267 | -1.144489924 | Cellular metabolism |
| Q00535 | CDK5 | Cyclin-dependent-like kinase 5 | -0.236036301 | -1.126943588 | Transport |
| Q00577 | PURA | Transcriptional activator protein Pur-alpha | 1.055891037 | 0.175022761 | Cellular metabolism |
| Q00653-4 | NFKB2 | Nuclear factor NF-kappa-B p100 subunit | -0.244158427 | -1.69841512 | Immune response |
| Q00688 | FKBP3 | Peptidyl-prolyl cis-trans isomerase FKBP3 | 0.097622554 | 0.538885117 | Cellular metabolism |
| Q00765 | REEP5 | Receptor expression-enhancing protein 5 | -0.654059092 | 1.534184138 | Other |
| Q00796 | SORD | Sorbitol dehydrogenase | 0.060087204 | 0.892554601 | Cellular metabolism |
| Q00839 | HNRNPU | Heterogeneous nuclear ribonucleoprotein U | 0.312826792 | 0.226199468 | Cellular metabolism |
| Q00978 | IRF9 | Interferon regulatory factor 9 | 1.319004695 | 3.24610583 | Cellular metabolism |
| Q01082 | SPTBN1 | Spectrin beta chain, non-erythrocytic 1 | -0.249005636 | -0.670434316 | Transport |
| Q01085 | TIAL1 | Nucleolysin TIAR | 0.094301224 | 1.373653412 | Other |
| Q01105-2 | SET | Protein SET | 0.049910227 | -0.628487269 | Transport |
| Q01432-6 | AMPD3 | AMP deaminase 3 | -0.294062297 | 0 | Cellular metabolism |
| Q01459 | CTBS | Di-N-acetylchitobiase | 0.098412832 | 0 | Other |
| Q01469 | FABP5 | Fatty acid-binding protein, epidermal | -0.26862208 | -0.214724223 | Lipid metabolism |
| Q01518 | CAP1 | Adenylyl cyclase-associated protein 1 | 0.06174914 | -0.425902049 | Transport |
| Q01581 | HMGCS1 | Hydroxymethylglutaryl-CoA synthase, cytoplasmic | -0.303467433 | -1.530420939 | Lipid metabolism |
| Q01650 | SLC7A5 | Large neutral amino acids transporter small subunit 1 | 0 | 1.552445094 | Transport |
| Q01658 | DR1 | Protein Dr1 | 0.098635356 | -1.122241338 | Cellular metabolism |
| Q01780 | EXOSC10 | Exosome component 10 | 0.280066172 | 0.922104518 | Cellular metabolism |
| Q01804 | OTUD4 | OTU domain-containing protein 4 | 1.380639394 | 0.027839661 | Cellular metabolism |
| Q01813 | PFKP | ATP-dependent 6-phosphofructokinase, platelet type | -0.028370539 | -0.307407379 | Cellular metabolism |

| | | | | | |
|---|---|---|---|---|---|
| Q01968-2 | OCRL | Inositol polyphosphate 5-phosphatase OCRL-1 | -0.453481674 | -0.656843185 | Lipid metabolism |
| Q01970-2 | PLCB3 | 1-phosphatidylinositol 4,5-bisphosphate phosphodiesterase beta-3 | -0.218640645 | 0.02617836 | Lipid metabolism |
| Q01995 | TAGLN | Transgelin | -0.247833888 | -0.603522619 | Other |
| Q02413 | DSG1 | Desmoglein-1 | 0.327471415 | 1.109514236 | Other |
| Q02539 | HIST1H1A | Histone H1.1 | 0.375363668 | -1.459962209 | Other |
| Q02750 | MAP2K1 | Dual specificity mitogen-activated protein kinase kinase 1 | 0.133856455 | -0.162570953 | Immune response |
| Q02790 | FKBP4 | Peptidyl-prolyl cis-trans isomerase FKBP4 | -0.071844737 | -0.620042165 | Transport |
| Q02809 | PLOD1 | Procollagen-lysine,2-oxoglutarate 5-dioxygenase 1 | 0.814932505 | 0.079241435 | Cellular metabolism |
| Q02818 | NUCB1 | Nucleobindin-1 | 1.084236145 | -0.839602788 | Other |
| Q02878 | RPL6 | 60S ribosomal protein L6 | 0.359629313 | 0.401110967 | Transport |
| Q02952 | AKAP12 | A-kinase anchor protein 12 | -0.335845947 | -0.614290237 | Transport |
| Q02978 | SLC25A11 | Mitochondrial 2-oxoglutarate/malate carrier protein | 1.188026428 | 0.710408529 | Transport |
| Q03111 | MLLT1 | Protein ENL | 0.163533529 | -0.716332753 | Cellular metabolism |
| Q03135 | CAV1 | Caveolin-1 | 0.264848073 | 0.829750061 | Lipid metabolism |
| Q03252 | LMNB2 | Lamin-B2 | 0.287897746 | -0.160157522 | Other |
| Q03405-2 | PLAUR | Urokinase plasminogen activator surface receptor | 0.071036021 | -0.554536819 | Cellular metabolism |
| Q04323 | UBXN1 | UBX domain-containing protein 1 | 0.006411235 | -1.132632573 | Cellular metabolism |
| Q04446 | GBE1 | 1,4-alpha-glucan-branching enzyme | -0.047196706 | -0.843273163 | Cellular metabolism |
| Q04721 | NOTCH2 | Neurogenic locus notch homolog protein 2 | -0.66522789 | -2.188123067 | Immune response |
| Q04760 | GLO1 | Lactoylglutathione lyase | -0.00170962 | -0.339946111 | Cellular metabolism |
| Q04837 | SSBP1 | Single-stranded DNA-binding protein, mitochondrial | 0.132186254 | 0.540522257 | Cellular metabolism |
| Q04864 | REL | Proto-oncogene c-Rel | -0.808862686 | -1.103693008 | Immune response |
| Q04917 | YWHAH | 14-3-3 protein eta | 0.05789566 | -0.294792811 | Lipid metabolism |
| Q05048 | CSTF1 | Cleavage stimulation factor subunit 1 | 0.676287969 | 0 | Cellular metabolism |
| Q05086-2 | UBE3A | Ubiquitin-protein ligase E3A | -0.235253652 | -1.899733226 | Transport |
| Q05209 | PTPN12 | Tyrosine-protein phosphatase non-receptor type 12 | 0.058278402 | -0.335621516 | Cellular metabolism |
| Q05397-5 | PTK2 | Focal adhesion kinase 1 | 0.24641037 | -0.04053243 | Immune response |
| Q05469-2 | LIPE | Hormone-sensitive lipase | 0 | 1.437105179 | Lipid metabolism |
| Q5T760 | SRSF11 | Serine/arginine-rich splicing factor 11 | 0.447041829 | -0.631144206 | Transport |
| Q05682-5 | CALD1 | Caldesmon | 0.009366353 | -0.466158549 | Other |
| Q06124-1 | PTPN11 | Tyrosine-protein phosphatase non-receptor type 11 | -0.034151077 | -1.18530337 | Immune response |
| Q06190 | PPP2R3A | Serine/threonine-protein phosphatase 2A regulatory subunit B subunit alpha | -0.715240479 | -0.886964798 | Cellular metabolism |
| Q06203 | PPAT | Amidophosphoribosyltransferase | -0.052251816 | -0.062929789 | Transport |
| Q06210-2 | GFPT1 | Glutamine--fructose-6-phosphate aminotransferase [isomerizing] 1 | -0.037446976 | -0.207518895 | Cellular metabolism |

| | | | | | |
|---|---|---|---|---|---|
| Q06323 | PSME1 | Proteasome activator complex subunit 1 | -0.047492345 | 0.19716835 | Immune response |
| Q06546 | GABPA | GA-binding protein alpha chain | -0.097129186 | -0.521393458 | Cellular metabolism |
| Q06587 | RING1 | E3 ubiquitin-protein ligase RING1 | 0.002772013 | 0 | Cellular metabolism |
| Q06609-3 | RAD51 | DNA repair protein RAD51 homolog 1 | -1.20819219 | 0 | Cellular metabolism |
| Q06830 | PRDX1 | Peroxiredoxin-1 | -0.058678945 | 0.193771362 | Immune response |
| Q07021 | C1QBP | Complement component 1 Q subcomponent-binding protein, mitochondrial | -0.286471049 | 0.781965892 | Immune response |
| Q07065 | CKAP4 | Cytoskeleton-associated protein 4 | 1.378355662 | 1.25158755 | Other |
| Q07666 | KHDRBS1 | KH domain-containing, RNA-binding, signal transduction-associated protein 1 | 0.977533976 | 0.500406265 | Cellular metabolism |
| Q07812-5 | BAX | Apoptosis regulator BAX | 0.069409053 | -0.06029892 | Lipid metabolism |
| Q07864 | POLE | DNA polymerase epsilon catalytic subunit A | -0.621325175 | 0 | Cellular metabolism |
| Q07866-6 | KLC1 | Kinesin light chain 1 | 0.159836451 | -0.357318242 | Transport |
| Q07954 | LRP1 | Prolow-density lipoprotein receptor-related protein 1 | 0.465778987 | -2.159653982 | Lipid metabolism |
| Q07960 | ARHGAP1 | Rho GTPase-activating protein 1 | -0.096918106 | -0.011516571 | Other |
| Q08170 | SRSF4 | Serine/arginine-rich splicing factor 4 | -0.00466156 | 1.923623403 | Transport |
| Q08209-2 | PPP3CA | Serine/threonine-protein phosphatase 2B catalytic subunit alpha isoform | -0.126422246 | -0.713585536 | Immune response |
| Q08211 | DHX9 | ATP-dependent RNA helicase A | 0.071545919 | 0.601641973 | Immune response |
| Q08257 | CRYZ | Quinone oxidoreductase | 0.06533432 | -0.085466385 | Cellular metabolism |
| Q08378 | GOLGA3 | Golgin subfamily A member 3 | 0.077269872 | -1.142758052 | Transport |
| Q08380 | LGALS3BP | Galectin-3-binding protein | -0.406539281 | 0.392452876 | Transport |
| Q08554-2 | DSC1 | Desmocollin-1 | 0.584775925 | 1.416156769 | Other |
| Q08623 | HDHD1 | Pseudouridine-5-phosphatase | -0.052347819 | -0.733656565 | Cellular metabolism |
| Q08752 | PPID | Peptidyl-prolyl cis-trans isomerase D | -0.134911855 | -0.382725398 | Transport |
| Q08945 | SSRP1 | FACT complex subunit SSRP1 | -0.038024902 | -0.376657486 | Cellular metabolism |
| Q08J23 | NSUN2 | tRNA (cytosine(34)-C(5))-methyltransferase | -0.229680379 | -0.889668147 | Cellular metabolism |
| Q09028-3 | RBBP4 | Histone-binding protein RBBP4 | -0.102373123 | -0.071152369 | Cellular metabolism |
| Q09161 | NCBP1 | Nuclear cap-binding protein subunit 1 | 0.236000061 | -0.811637878 | Transport |
| Q09666 | AHNAK | Neuroblast differentiation-associated protein AHNAK | -0.392583847 | -0.540823619 | Other |
| Q0JRZ9 | FCHO2 | F-BAR domain only protein 2 | 0.373598735 | 0 | Transport |
| Q0VDF9 | HSPA14 | Heat shock 70 kDa protein 14 | -0.056781133 | -0.7695268 | Cellular metabolism |
| Q0VDG4-2 | SCRN3 | Secernin-3 | -0.336970647 | -1.352563858 | Transport |
| Q0ZGT2-4 | NEXN | Nexilin | 0.359531403 | -0.310829163 | Cellular metabolism |
| Q10471 | GALNT2 | Polypeptide N-acetylgalactosaminyltransferase 2 | 0 | 0.777028402 | Cellular metabolism |
| Q10713 | PMPCA | Mitochondrial-processing peptidase subunit alpha | -0.154591242 | 0.545153936 | Transport |
| Q12765 | SCRN1 | Secernin-1 | -0.045780818 | -0.577390671 | Transport |

| | | | | | |
|---|---|---|---|---|---|
| Q12768 | KIAA0196 | WASH complex subunit strumpellin | -0.133252462 | -1.22739919 | Transport |
| Q12789-3 | GTF3C1 | General transcription factor 3C polypeptide 1 | -0.236918132 | 0 | Cellular metabolism |
| Q12792 | TWF1 | Twinfilin-1 | -0.013924281 | -0.356981277 | Other |
| Q12797 | ASPH | Aspartyl/asparaginyl beta-hydroxylase | 1.002157211 | -0.079557419 | Transport |
| Q12872-2 | SFSWAP | Splicing factor, suppressor of white-apricot homolog | 0.227720896 | -2.294043859 | Cellular metabolism |
| Q12874 | SF3A3 | Splicing factor 3A subunit 3 | 0.014186223 | 0.634087245 | Cellular metabolism |
| Q12882 | DPYD | Dihydropyrimidine dehydrogenase [NADP(+)] | 0.112914403 | 0 | Cellular metabolism |
| Q12888 | TP53BP1 | Tumor suppressor p53-binding protein 1 | 0.027798971 | -0.675283432 | Cellular metabolism |
| Q12899 | TRIM26 | Tripartite motif-containing protein 26 | 1.805610657 | 1.857708613 | Immune response |
| Q12904 | AIMP1 | Aminoacyl tRNA synthase complex-interacting multifunctional protein 1 | 0.265570323 | -1.140035629 | Cellular metabolism |
| Q12905 | ILF2 | Interleukin enhancer-binding factor 2 | 0.153528849 | 0.635660172 | Immune response |
| Q12906-7 | ILF3 | Interleukin enhancer-binding factor 3 | 0.366298676 | 0.332473755 | Cellular metabolism |
| Q12907 | LMAN2 | Vesicular integral-membrane protein VIP36 | 1.430070877 | 1.73976326 | Transport |
| Q12931-2 | TRAP1 | Heat shock protein 75 kDa, mitochondrial | -0.169001897 | 0.258886337 | Cellular metabolism |
| Q12933-3 | TRAF2 | TNF receptor-associated factor 2 | 0.027317683 | -0.280666987 | Immune response |
| Q12965 | MYO1E | Unconventional myosin-Ie | -0.025879542 | -1.891649246 | Transport |
| Q13033-2 | STRN3 | Striatin-3 | 0.152135213 | -1.576934179 | Other |
| Q13042-3 | CDC16 | Cell division cycle protein 16 homolog | 0.062980016 | 0 | Cellular metabolism |
| Q13043 | STK4 | Serine/threonine-protein kinase 4 | -0.453118006 | -2.363189697 | Cellular metabolism |
| Q13045 | FLII | Protein flightless-1 homolog | -0.060008367 | -0.559710821 | Cellular metabolism |
| Q13049 | TRIM32 | E3 ubiquitin-protein ligase TRIM32 | -0.099380493 | -0.511804581 | Immune response |
| Q13057 | COASY | Bifunctional coenzyme A synthase | -0.100738525 | 0.167966207 | Cellular metabolism |
| Q13085 | ACACA | Acetyl-CoA carboxylase 1 | 0.220549901 | -1.173480352 | Lipid metabolism |
| Q13107-2 | USP4 | Ubiquitin carboxyl-terminal hydrolase 4 | 0.217667262 | -1.554803848 | Cellular metabolism |
| Q13112 | CHAF1B | Chromatin assembly factor 1 subunit B | 0.545488993 | -1.016311646 | Cellular metabolism |
| Q13123 | IK | Protein Red | -0.542313258 | 0 | Immune response |
| Q13131 | PRKAA1 | 5-AMP-activated protein kinase catalytic subunit alpha-1 | 0.079579671 | -0.84710439 | Lipid metabolism |
| Q13137 | CALCOCO2 | Calcium-binding and coiled-coil domain-containing protein 2 | -0.966292063 | 0.281667709 | Other |
| Q13148 | TARDBP | TAR DNA-binding protein 43 | 0.399638494 | 1.219584783 | Cellular metabolism |
| Q13151 | HNRNPA0 | Heterogeneous nuclear ribonucleoprotein A0 | 0.099002838 | 0.728845596 | Cellular metabolism |
| Q13153 | PAK1 | Serine/threonine-protein kinase PAK 1 | -0.252449671 | -0.285088857 | Immune response |
| Q13155 | AIMP2 | Aminoacyl tRNA synthase complex-interacting multifunctional protein 2 | 0.244925181 | 1.408567429 | Cellular metabolism |
| Q13158 | FADD | FAS-associated death domain protein | 0.386891683 | -1.341255188 | Immune response |
| Q13162 | PRDX4 | Peroxiredoxin-4 | 0.17915535 | 0.00949351 | Cellular metabolism |
| Q13164 | MAPK7 | Mitogen-activated protein kinase 7 | 0.085866292 | 0.630294164 | Immune response |

| | | | | | |
|---|---|---|---|---|---|
| Q13177 | PAK2 | Serine/threonine-protein kinase PAK 2 | -0.120429993 | -0.606363932 | Immune response |
| Q13185 | CBX3 | Chromobox protein homolog 3 | -0.182091395 | -0.011774063 | Cellular metabolism |
| Q13188-2 | STK3 | Serine/threonine-protein kinase 3 | -0.168626785 | -0.817014694 | Cellular metabolism |
| Q13200 | PSMD2 | 26S proteasome non-ATPase regulatory subunit 2 | -0.246851603 | -0.566766103 | Immune response |
| Q13228 | SELENBP1 | Selenium-binding protein 1 | 0.04276975 | -0.706120173 | Transport |
| Q13232 | NME3 | Nucleoside diphosphate kinase 3 | -0.579426448 | 0.305706024 | Cellular metabolism |
| Q13242 | SRSF9 | Serine/arginine-rich splicing factor 9 | 0.010067622 | -0.175977707 | Transport |
| Q13243 | SRSF5 | Serine/arginine-rich splicing factor 5 | 0.300647736 | -0.837280909 | Transport |
| Q13257 | MAD2L1 | Mitotic spindle assembly checkpoint protein MAD2A | -0.231525421 | -0.439980825 | Cellular metabolism |
| Q13263 | TRIM28 | Transcription intermediary factor 1-beta | 0.073132197 | -0.547300339 | Immune response |
| Q13283 | G3BP1 | Ras GTPase-activating protein-binding protein 1 | -0.113669078 | 0.334290187 | Transport |
| Q13287 | NMI | N-myc-interactor | -0.171678543 | -0.221357346 | Cellular metabolism |
| Q13308-4 | PTK7 | Inactive tyrosine-protein kinase 7 | -0.035547892 | -1.583526611 | Other |
| Q13325 | IFIT5 | Interferon-induced protein with tetratricopeptide repeats 5 | 0.616629918 | 1.637673696 | Immune response |
| Q13363-2 | CTBP1 | C-terminal-binding protein 1 | 0.028645833 | -0.904220581 | Cellular metabolism |
| Q13371-2 | PDCL | Phosducin-like protein | -0.12132899 | -1.676561356 | Cellular metabolism |
| Q13404 | UBE2V1 | Ubiquitin-conjugating enzyme E2 variant 1 | 0.009902318 | -0.185215632 | Immune response |
| Q13405 | MRPL49 | 39S ribosomal protein L49, mitochondrial | 0.154978434 | -0.703802109 | Cellular metabolism |
| Q13409 | DYNC1I2 | Cytoplasmic dynein 1 intermediate chain 2 | 0.1282444 | -0.309604645 | Transport |
| Q13423 | NNT | NAD(P) transhydrogenase, mitochondrial | 0 | 1.115813573 | Transport |
| Q13426 | XRCC4 | DNA repair protein XRCC4 | -0.248307546 | -0.490694046 | Cellular metabolism |
| Q13428-3 | TCOF1 | Treacle protein | 0.275372823 | -0.094786962 | Cellular metabolism |
| Q13432 | UNC119 | Protein unc-119 homolog A | -0.122959773 | 0 | Transport |
| Q13435 | SF3B2 | Splicing factor 3B subunit 2 | 0.029986064 | -0.024481455 | Cellular metabolism |
| Q13439-3 | GOLGA4 | Golgin subfamily A member 4 | 0.520746867 | -0.931149801 | Transport |
| Q13442 | PDAP1 | 28 kDa heat- and acid-stable phosphoprotein | 0.276427587 | -0.813875198 | Other |
| Q13445 | TMED1 | Transmembrane emp24 domain-containing protein 1 | 0 | 0.70364062 | Transport |
| Q13451 | FKBP5 | Peptidyl-prolyl cis-trans isomerase FKBP5 | 0.054115295 | -0.833343506 | Cellular metabolism |
| Q13464 | ROCK1 | Rho-associated protein kinase 1 | 1.013093313 | -1.34851265 | Cellular metabolism |
| Q13492 | PICALM | Phosphatidylinositol-binding clathrin assembly protein | -0.728000641 | 0.22577095 | Transport |
| Q13501 | SQSTM1 | Sequestosome-1 | -0.583028793 | 0.077469508 | Transport |
| Q13505-3 | MTX1 | Metaxin-1 | 0 | 1.714920044 | Transport |
| Q13526 | PIN1 | Peptidyl-prolyl cis-trans isomerase NIMA-interacting 1 | -0.092271169 | -0.388851802 | Immune response |

| | | | | | |
|---|---|---|---|---|---|
| Q13546 | RIPK1 | Receptor-interacting serine/threonine-protein kinase 1 | -0.254744212 | -0.886219025 | Immune response |
| Q13547 | HDAC1 | Histone deacetylase 1 | -0.034456889 | 0.33937645 | Cellular metabolism |
| Q13561 | DCTN2 | Dynactin subunit 2 | 0.207721074 | 0.001998266 | Transport |
| Q13564-2 | NAE1 | NEDD8-activating enzyme E1 regulatory subunit | -0.111583074 | -1.052665075 | Cellular metabolism |
| Q13572 | ITPK1 | Inositol-tetrakisphosphate 1-kinase | -0.214305242 | 0 | Cellular metabolism |
| Q13573 | SNW1 | SNW domain-containing protein 1 | 0.052762349 | -0.681662877 | Cellular metabolism |
| Q13595-4 | TRA2A | Transformer-2 protein homolog alpha | 0.192010244 | -0.327915827 | Cellular metabolism |
| Q13596 | SNX1 | Sorting nexin-1 | 0.069660823 | -0.34536171 | Transport |
| Q13610 | PWP1 | Periodic tryptophan protein 1 homolog | 0.29650561 | -0.66564242 | Cellular metabolism |
| Q13616 | CUL1 | Cullin-1 | 0.004947662 | -0.343095144 | Immune response |
| Q13618 | CUL3 | Cullin-3 | 0.001881917 | -0.455920537 | Transport |
| Q13619 | CUL4A | Cullin-4A | 0.156066259 | -0.543678284 | Cellular metabolism |
| Q13630 | TSTA3 | GDP-L-fucose synthase | 0.055577596 | -0.553705215 | Cellular metabolism |
| Q13636 | RAB31 | Ras-related protein Rab-31 | 0 | 2.898110708 | Transport |
| Q13637 | RAB32 | Ras-related protein Rab-32 | 0.023399989 | 1.029208501 | Transport |
| Q13642-1 | FHL1 | Four and a half LIM domains protein 1 | 0.028659185 | -2.201330185 | Other |
| Q13643 | FHL3 | Four and a half LIM domains protein 3 | -0.102153778 | -0.734524409 | Other |
| Q13671 | RIN1 | Ras and Rab interactor 1 | 0.193519592 | -0.588876088 | Transport |
| Q13740-2 | ALCAM | CD166 antigen | -0.44170634 | -0.1130174 | Other |
| Q13813-3 | SPTAN1 | Spectrin alpha chain, non-erythrocytic 1 | -0.185136795 | -0.67913119 | Other |
| Q13823 | GNL2 | Nucleolar GTP-binding protein 2 | 0.165430069 | 0 | Other |
| Q13835-2 | PKP1 | Plakophilin-1 | 1.097592672 | 1.306034088 | Other |
| Q13838-2 | DDX39B | Spliceosome RNA helicase DDX39B | -0.005584081 | 0.046709061 | Transport |
| Q13867 | BLMH | Bleomycin hydrolase | 0.0401872 | 0.108469009 | Cellular metabolism |
| Q13868 | EXOSC2 | Exosome complex component RRP4 | 0.325925827 | 0.140536626 | Cellular metabolism |
| Q6P1K8 | GTF2H2C | General transcription factor IIH subunit 2-like protein | -0.496782303 | -0.854272207 | Transport |
| Q13895 | BYSL | Bystin | -0.158080419 | 0.519408544 | Cellular metabolism |
| Q13907 | IDI1 | Isopentenyl-diphosphate Delta-isomerase 1 | 0.08685112 | -1.050685883 | Lipid metabolism |
| Q13951 | CBFB | Core-binding factor subunit beta | -0.232365926 | -0.30868276 | Cellular metabolism |
| Q13951-2 | CBFB | Core-binding factor subunit beta | -0.102645238 | -0.896223068 | Cellular metabolism |
| Q14008-2 | CKAP5 | Cytoskeleton-associated protein 5 | 0.252465566 | 0.607683818 | Transport |
| Q14011 | CIRBP | Cold-inducible RNA-binding protein | 0.357848485 | 0.469654719 | Cellular metabolism |
| Q14012 | CAMK1 | Calcium/calmodulin-dependent protein kinase type 1 | -0.679908752 | -0.208485285 | Transport |
| Q14019 | COTL1 | Coactosin-like protein | -0.062900543 | -0.830652873 | Lipid metabolism |
| Q14103-3 | HNRNPD | Heterogeneous nuclear ribonucleoprotein D0 | 0.251349131 | 0.155134201 | Cellular metabolism |

| | | | | | |
|---|---|---|---|---|---|
| Q14116-2 | IL18 | Interleukin-18 | -0.740173976 | -1.017142614 | Immune response |
| Q14134-2 | TRIM29 | Tripartite motif-containing protein 29 | 0 | 1.465651194 | Cellular metabolism |
| Q14137 | BOP1 | Ribosome biogenesis protein BOP1 | 1.336688995 | -0.00719897 | Cellular metabolism |
| Q14139-2 | UBE4A | Ubiquitin conjugation factor E4 A | -0.03924942 | -1.344242096 | Cellular metabolism |
| Q14145 | KEAP1 | Kelch-like ECH-associated protein 1 | 0.309569677 | 0.295220693 | Cellular metabolism |
| Q14152 | EIF3A | Eukaryotic translation initiation factor 3 subunit A | 0.210213343 | -0.331406911 | Cellular metabolism |
| Q14166 | TTLL12 | Tubulin--tyrosine ligase-like protein 12 | -0.009333293 | -0.861063004 | Cellular metabolism |
| Q14181 | POLA2 | DNA polymerase alpha subunit B | 0.146745682 | -0.624628703 | Transport |
| Q14192 | FHL2 | Four and a half LIM domains protein 2 | -0.080404282 | -0.896270116 | Lipid metabolism |
| Q14195-2 | DPYSL3 | Dihydropyrimidinase-related protein 3 | 0.099599202 | -0.562892914 | Other |
| Q14204 | DYNC1H1 | Cytoplasmic dynein 1 heavy chain 1 | -0.091233571 | 0.094057719 | Transport |
| Q14232 | EIF2B1 | Translation initiation factor eIF-2B subunit alpha | -0.246056875 | -0.266862233 | Cellular metabolism |
| Q14240 | EIF4A2 | Eukaryotic initiation factor 4A-II | -0.392604192 | 0.190401713 | Cellular metabolism |
| Q14247 | CTTN | Src substrate cortactin | -0.011978785 | -0.550995509 | Transport |
| Q14257 | RCN2 | Reticulocalbin-2 | 0.448361079 | -0.58063062 | Other |
| Q14258 | TRIM25 | E3 ubiquitin/ISG15 ligase TRIM25 | -0.144650777 | 1.44889005 | Immune response |
| Q14315 | FLNC | Filamin-C | -0.003005981 | -0.506107966 | Other |
| Q14318 | FKBP8 | Peptidyl-prolyl cis-trans isomerase FKBP8 | -0.141166051 | 0.078360875 | Cellular metabolism |
| Q14320 | FAM50A | Protein FAM50A | 0.194510142 | 0.326831818 | Other |
| Q14331 | FRG1 | Protein FRG1 | 0.091890971 | -1.142505646 | Cellular metabolism |
| Q14353 | GAMT | Guanidinoacetate N-methyltransferase | -0.158602397 | -0.111391703 | Cellular metabolism |
| Q14444-2 | CAPRIN1 | Caprin-1 | -0.175912221 | -0.067964554 | Other |
| Q14498 | RBM39 | RNA-binding protein 39 | -0.441302617 | 1.546320597 | Cellular metabolism |
| Q14558-2 | PRPSAP1 | Phosphoribosyl pyrophosphate synthase-associated protein 1 | 0.136988322 | -0.554334641 | Cellular metabolism |
| Q14566 | MCM6 | DNA replication licensing factor MCM6 | 0.015033086 | -0.488310496 | Cellular metabolism |
| Q14573 | ITPR3 | Inositol 1,4,5-trisphosphate receptor type 3 | 0.240182241 | -1.689133962 | Immune response |
| Q14574-2 | DSC3 | Desmocollin-3 | 0.843571345 | 0.942195892 | Other |
| Q14651 | PLS1 | Plastin-1 | -0.049342473 | -2.489079793 | Other |
| Q14653 | IRF3 | Interferon regulatory factor 3 | -0.010183334 | -2.217066447 | Immune response |
| Q14657 | LAGE3 | EKC/KEOPS complex subunit LAGE3 | 0.261920929 | 0 | Cellular metabolism |
| Q5T1Z4 | PUM1 | Pumilio homolog 1 | 0.50679334 | -1.734271367 | Transport |
| Q14677-3 | CLINT1 | Clathrin interactor 1 | 0.099257151 | 0.521250407 | Transport |
| Q14683 | SMC1A | Structural maintenance of chromosomes protein 1A | 0.083366394 | 0.210472743 | Cellular metabolism |
| Q14684-2 | RRP1B | Ribosomal RNA processing protein 1 homolog B | 1.296619415 | 0.266571681 | Cellular metabolism |

| | | | | | |
|---|---|---|---|---|---|
| Q14691 | GINS1 | DNA replication complex GINS protein PSF1 | 0.099251429 | -1.638975143 | Cellular metabolism |
| Q14693 | LPIN1 | Phosphatidate phosphatase LPIN1 | -0.373945236 | 0 | Lipid metabolism |
| Q14694 | USP10 | Ubiquitin carboxyl-terminal hydrolase 10 | 0.070871353 | 0.954018911 | Cellular metabolism |
| Q14696 | MESDC2 | LDLR chaperone MESD | 0.2989947 | -0.112008413 | Cellular metabolism |
| Q14697-2 | GANAB | Neutral alpha-glucosidase AB | 0.289038976 | -0.10994339 | Cellular metabolism |
| Q14738 | PPP2R5D | Serine/threonine-protein phosphatase 2A 56 kDa regulatory subunit delta isoform | -0.031665166 | -2.038988749 | Immune response |
| Q14764 | MVP | Major vault protein | -0.09725825 | -0.333365122 | Transport |
| Q14789-4 | GOLGB1 | Golgin subfamily B member 1 | 0 | -0.006031672 | Other |
| Q14790-2 | CASP8 | Caspase-8 | -0.338434855 | 0 | Immune response |
| Q147X3 | NAA30 | N-alpha-acetyltransferase 30 | 0.039412181 | -0.935525258 | Cellular metabolism |
| Q14847 | LASP1 | LIM and SH3 domain protein 1 | 0.003485998 | -0.309034348 | Transport |
| Q14914 | PTGR1 | Prostaglandin reductase 1 | 0.012395859 | -0.578229268 | Lipid metabolism |
| Q14966-5 | ZNF638 | Zinc finger protein 638 | 0.302638372 | 0.326953888 | Cellular metabolism |
| Q14974 | KPNB1 | Importin subunit beta-1 | -0.046606064 | -0.079069773 | Transport |
| Q14978-3 | NOLC1 | Nucleolar and coiled-body phosphoprotein 1 | -0.080224355 | 0.100858688 | Cellular metabolism |
| Q14980-2 | NUMA1 | Nuclear mitotic apparatus protein 1 | 0.65774409 | 0.223597844 | Other |
| Q14997 | PSME4 | Proteasome activator complex subunit 4 | 0.042163849 | 0.271405538 | Immune response |
| Q14C86-3 | GAPVD1 | GTPase-activating protein and VPS9 domain-containing protein 1 | 0.023003896 | -1.108318965 | Transport |
| Q14CX7 | NAA25 | N-alpha-acetyltransferase 25, NatB auxiliary subunit | -0.490557353 | -0.158714294 | Other |
| Q15006 | EMC2 | ER membrane protein complex subunit 2 | 0 | 1.776264826 | Other |
| Q15007 | WTAP | Pre-mRNA-splicing regulator WTAP | 0.749774933 | -0.147280375 | Cellular metabolism |
| Q15008 | PSMD6 | 26S proteasome non-ATPase regulatory subunit 6 | 0.010238012 | -0.732432048 | Immune response |
| Q15018 | FAM175B | BRISC complex subunit Abro1 | -0.175760905 | -1.291778564 | Other |
| Q15019-2 | 37500 | Septin-2 | -0.418294271 | 0.207182566 | Other |
| Q15021 | NCAPD2 | Condensin complex subunit 1 | -0.28399086 | -0.015935898 | Cellular metabolism |
| Q15024 | EXOSC7 | Exosome complex component RRP42 | -0.075328191 | -0.265429815 | Cellular metabolism |
| Q15029 | EFTUD2 | 116 kDa U5 small nuclear ribonucleoprotein component | 0.465796153 | -0.168478648 | Cellular metabolism |
| Q15036-2 | SNX17 | Sorting nexin-17 | -1.279159546 | -1.19740804 | Lipid metabolism |
| Q15042 | RAB3GAP1 | Rab3 GTPase-activating protein catalytic subunit | -0.219395955 | -1.633879344 | Other |
| Q15046 | KARS | Lysine--tRNA ligase | 0.122830073 | 0.159496307 | Cellular metabolism |
| Q15048 | LRRC14 | Leucine-rich repeat-containing protein 14 | 0.483057022 | -0.654722214 | Other |
| Q15050 | RRS1 | Ribosome biogenesis regulatory protein homolog | 0 | 3.074335098 | Other |
| Q15054 | POLD3 | DNA polymerase delta subunit 3 | -0.139059067 | -1.728001277 | Cellular metabolism |
| Q15056 | EIF4H | Eukaryotic translation initiation factor 4H | 0.09368515 | -0.287776947 | Cellular metabolism |

| | | | | | |
|---|---|---|---|---|---|
| Q15057 | ACAP2 | Arf-GAP with coiled-coil, ANK repeat and PH domain-containing protein 2 | -0.092013677 | -0.046810786 | Other |
| Q15061 | WDR43 | WD repeat-containing protein 43 | -0.95290184 | -3.107026418 | Other |
| Q15067-3 | ACOX1 | Peroxisomal acyl-coenzyme A oxidase 1 | 0 | 1.759806951 | Lipid metabolism |
| Q15075 | EEA1 | Early endosome antigen 1 | -0.329288483 | -0.374251048 | Transport |
| Q15084-3 | PDIA6 | Protein disulfide-isomerase A6 | 0.449895859 | 0.530339559 | Transport |
| Q15102 | PAFAH1B3 | Platelet-activating factor acetylhydrolase IB subunit gamma | -0.076221466 | -1.36250178 | Lipid metabolism |
| Q15121-2 | PEA15 | Astrocytic phosphoprotein PEA-15 | -0.327062607 | -0.616994858 | Transport |
| Q15126 | PMVK | Phosphomevalonate kinase | -0.208854675 | -0.452878316 | Lipid metabolism |
| Q15149 | PLEC | Plectin | 0.345163345 | -0.224696477 | Other |
| Q15181 | PPA1 | Inorganic pyrophosphatase | -0.284805934 | -0.610304515 | Cellular metabolism |
| Q15185-4 | PTGES3 | Prostaglandin E synthase 3 | -0.114679972 | -0.541378021 | Lipid metabolism |
| Q15208 | STK38 | Serine/threonine-protein kinase 38 | -0.192889531 | -0.945760091 | Cellular metabolism |
| Q15257 | PPP2R4 | Serine/threonine-protein phosphatase 2A activator | -0.067564011 | -0.897092819 | Cellular metabolism |
| Q15276 | RABEP1 | Rab GTPase-binding effector protein 1 | -0.001850128 | -0.721217473 | Transport |
| Q15291-2 | RBBP5 | Retinoblastoma-binding protein 5 | -0.04690361 | 0 | Cellular metabolism |
| Q15293 | RCN1 | Reticulocalbin-1 | 0.23050944 | -0.364741643 | Other |
| Q15311 | RALBP1 | RalA-binding protein 1 | -0.354728063 | -1.180988312 | Transport |
| Q15334 | LLGL1 | Lethal(2) giant larvae protein homolog 1 | -0.799212774 | -0.471310933 | Transport |
| Q15363 | TMED2 | Transmembrane emp24 domain-containing protein 2 | 1.92547671 | 2.45897611 | Transport |
| Q15365 | PCBP1 | Poly(rC)-binding protein 1 | 0.018597921 | 0.330968221 | Cellular metabolism |
| Q15366 | PCBP2 | Poly(rC)-binding protein 2 | 0.085123062 | 0.827876409 | Immune response |
| Q15382 | RHEB | GTP-binding protein Rheb | -0.035387039 | -1.160291036 | Cellular metabolism |
| Q15386 | UBE3C | Ubiquitin-protein ligase E3C | -0.116828283 | -1.084700902 | Cellular metabolism |
| Q15388 | TOMM20 | Mitochondrial import receptor subunit TOM20 homolog | -0.658346176 | 1.197495143 | Transport |
| Q15393 | SF3B3 | Splicing factor 3B subunit 3 | 0.167948405 | -0.015263875 | Cellular metabolism |
| Q15398 | DLGAP5 | Disks large-associated protein 5 | -0.678923925 | -2.106434504 | Cellular metabolism |
| Q15404 | RSU1 | Ras suppressor protein 1 | -0.244035085 | -1.387233734 | Other |
| Q15417 | CNN3 | Calponin-3 | 0.06620725 | -0.198848089 | Other |
| Q15424-2 | SAFB | Scaffold attachment factor B1 | 0.912926356 | 0.323466619 | Cellular metabolism |
| Q15427 | SF3B4 | Splicing factor 3B subunit 4 | 0.304465612 | 0.232100169 | Cellular metabolism |
| Q15428 | SF3A2 | Splicing factor 3A subunit 2 | 0.108757019 | -4.375895818 | Cellular metabolism |
| Q15435 | PPP1R7 | Protein phosphatase 1 regulatory subunit 7 | 0.045525869 | -0.587376912 | Other |
| Q15438-2 | CYTH1 | Cytohesin-1 | -0.285467148 | 0 | Transport |
| Q15459 | SF3A1 | Splicing factor 3A subunit 1 | 0.3401076 | -0.015367508 | Cellular metabolism |

| | | | | | |
|---|---|---|---|---|---|
| Q15477 | SKIV2L | Helicase SKI2W | 0.08986028 | 0 | Other |
| Q15527 | SURF2 | Surfeit locus protein 2 | 0.283032099 | -1.160673777 | Other |
| Q15555-5 | MAPRE2 | Microtubule-associated protein RP/EB family member 2 | -0.879654566 | -0.47183609 | Other |
| Q15628 | TRADD | Tumor necrosis factor receptor type 1-associated DEATH domain protein | 0.265138626 | 0 | Other |
| Q15631 | TSN | Translin | 0.212399801 | 0.631427129 | Cellular metabolism |
| Q15637-4 | SF1 | Splicing factor 1 | 0.573959351 | 0.746011098 | Cellular metabolism |
| Q15642-2 | TRIP10 | Cdc42-interacting protein 4 | 0.431310018 | -0.757726669 | Transport |
| Q15645 | TRIP13 | Pachytene checkpoint protein 2 homolog | -0.093917847 | -0.277137121 | Cellular metabolism |
| Q15653-2 | NFKBIB | NF-kappa-B inhibitor beta | -1.297478994 | -0.900307337 | Immune response |
| Q15654 | TRIP6 | Thyroid receptor-interacting protein 6 | -0.225860596 | -1.283311844 | Cellular metabolism |
| Q15691 | MAPRE1 | Microtubule-associated protein RP/EB family member 1 | 0.066876729 | -0.561103185 | Other |
| Q15717 | ELAVL1 | ELAV-like protein 1 | 0.107081095 | -0.034671148 | Cellular metabolism |
| Q15746-9 | MYLK | Myosin light chain kinase, smooth muscle | -0.400168101 | -0.544769287 | Cellular metabolism |
| Q15750 | TAB1 | TGF-beta-activated kinase 1 and MAP3K7-binding protein 1 | 0.253095627 | -1.358590444 | Immune response |
| Q15758 | SLC1A5 | Neutral amino acid transporter B(0) | -0.931573868 | -0.363740285 | Transport |
| Q15773 | MLF2 | Myeloid leukemia factor 2 | -0.621180216 | 0 | Other |
| Q15785 | TOMM34 | Mitochondrial import receptor subunit TOM34 | 0.04082044 | -1.364910762 | Transport |
| Q15796-2 | SMAD2 | Mothers against decapentaplegic homolog 2 | 0.696879705 | 0 | Transport |
| Q15813 | TBCE | Tubulin-specific chaperone E | -0.076675415 | -1.304217656 | Cellular metabolism |
| Q15814 | TBCC | Tubulin-specific chaperone C | 0.253147125 | 0.195554733 | Cellular metabolism |
| Q15819 | UBE2V2 | Ubiquitin-conjugating enzyme E2 variant 2 | 0.018209457 | -0.388638814 | Cellular metabolism |
| Q15828 | CST6 | Cystatin-M | 0 | 1.400784175 | Other |
| Q15836 | VAMP3 | Vesicle-associated membrane protein 3 | 0 | 2.774852117 | Transport |
| Q15843 | NEDD8 | NEDD8 | 0.047643661 | -0.30208079 | Cellular metabolism |
| Q15907 | RAB11B | Ras-related protein Rab-11B | -0.163349152 | 1.82980601 | Transport |
| Q15942 | ZYX | Zyxin | -0.250065486 | -0.399132411 | Other |
| Q16186 | ADRM1 | Proteasomal ubiquitin receptor ADRM1 | 0.246192296 | -0.102458318 | Cellular metabolism |
| Q16204 | CCDC6 | Coiled-coil domain-containing protein 6 | 0.039773941 | -1.147204717 | Other |
| Q16222 | UAP1 | UDP-N-acetylhexosamine pyrophosphorylase | -0.106789907 | -0.286511739 | Cellular metabolism |
| Q16342-3 | PDCD2 | Programmed cell death protein 2 | -0.535984675 | 0 | Other |
| Q16352 | INA | Alpha-internexin | 0.265045802 | -0.559642792 | Other |
| Q16401-2 | PSMD5 | 26S proteasome non-ATPase regulatory subunit 5 | 0.212100983 | -0.157498678 | Immune response |
| Q16512 | PKN1 | Serine/threonine-protein kinase N1 | 0.035254796 | -0.730056763 | Cellular metabolism |
| Q16513-3 | PKN2 | Serine/threonine-protein kinase N2 | -0.470201492 | -0.605405807 | Cellular metabolism |

| | | | | | |
|---|---|---|---|---|---|
| Q16531 | DDB1 | DNA damage-binding protein 1 | -0.101015091 | -0.553104401 | Cellular metabolism |
| Q16537 | PPP2R5E | Serine/threonine-protein phosphatase 2A 56 kDa regulatory subunit epsilon isoform | -0.231440862 | -1.021087646 | Other |
| Q16539 | MAPK14 | Mitogen-activated protein kinase 14 | -0.036996206 | -1.365999222 | Immune response |
| Q16543 | CDC37 | Hsp90 co-chaperone Cdc37 | 0.014884949 | -0.631468455 | Transport |
| Q16566 | CAMK4 | Calcium/calmodulin-dependent protein kinase type IV | -0.167393366 | -1.239050547 | Immune response |
| Q16576 | RBBP7 | Histone-binding protein RBBP7 | -0.167277018 | -0.754640579 | Cellular metabolism |
| Q16610-2 | ECM1 | Extracellular matrix protein 1 | -1.850863775 | 0.754487991 | Other |
| Q16623 | STX1A | Syntaxin-1A | 0.418969472 | 0 | Transport |
| Q16629-4 | SRSF7 | Serine/arginine-rich splicing factor 7 | 0.348279317 | 0.025901794 | Transport |
| Q16643 | DBN1 | Drebrin | 0.048037847 | -0.437376658 | Other |
| Q16644 | MAPKAPK3 | MAP kinase-activated protein kinase 3 | -0.46156311 | -0.320627213 | Immune response |
| Q16658 | FSCN1 | Fascin | 0.053070068 | -0.450129827 | Other |
| Q16666-3 | IFI16 | Gamma-interferon-inducible protein 16 | -1.101717631 | 2.12625885 | Immune response |
| Q16698-2 | DECR1 | 2,4-dienoyl-CoA reductase, mitochondrial | 0 | 0.622454961 | Lipid metabolism |
| Q16762 | TST | Thiosulfate sulfurtransferase | -0.095314026 | 0 | Transport |
| Q16763 | UBE2S | Ubiquitin-conjugating enzyme E2 S | -0.553084691 | -2.123224258 | Cellular metabolism |
| Q16774 | GUK1 | Guanylate kinase | -0.414276759 | -2.297887166 | Transport |
| Q16777 | HIST2H2AC | Histone H2A type 2-C | 0.570433935 | 0.803986231 | Other |
| Q16795 | NDUFA9 | NADH dehydrogenase [ubiquinone] 1 alpha subcomplex subunit 9, mitochondrial | 0 | 1.626614253 | Lipid metabolism |
| Q16822 | PCK2 | Phosphoenolpyruvate carboxykinase [GTP], mitochondrial | 0.134981155 | -0.253953298 | Cellular metabolism |
| Q16831 | UPP1 | Uridine phosphorylase 1 | 0.230816523 | -0.190619787 | Cellular metabolism |
| Q16851 | UGP2 | UTP--glucose-1-phosphate uridylyltransferase | 0.053953171 | -0.312032064 | Cellular metabolism |
| Q16864 | ATP6V1F | V-type proton ATPase subunit F | -0.630125046 | 0 | Transport |
| Q16891-2 | IMMT | MICOS complex subunit MIC60 | 0 | 0.757996241 | Other |
| Q1KMD3 | HNRNPUL2 | Heterogeneous nuclear ribonucleoprotein U-like protein 2 | 0.074653625 | -2.531866709 | Other |
| Q29RF7 | PDS5A | Sister chromatid cohesion protein PDS5 homolog A | 0.380398432 | -2.181646983 | Other |
| Q2NL82 | TSR1 | Pre-rRNA-processing protein TSR1 homolog | -0.327865601 | -0.523430506 | Other |
| Q2TAA2 | IAH1 | Isoamyl acetate-hydrolyzing esterase 1 homolog | 0.439612707 | -0.158372879 | Lipid metabolism |
| Q2TAL8 | QRICH1 | Glutamine-rich protein 1 | -0.051320394 | -0.544150035 | Other |
| Q2TAM5 | RELA | Transcription factor p65 | -0.352496465 | -0.325869878 | Immune response |
| Q2TAY7 | SMU1 | WD40 repeat-containing protein SMU1 | -0.19849968 | 0.398192724 | Other |
| Q2TAZ0 | ATG2A | Autophagy-related protein 2 homolog A | 0.789826075 | 2.082326889 | Cellular metabolism |
| Q32MZ4-3 | LRRFIP1 | Leucine-rich repeat flightless-interacting protein 1 | 0.105148315 | -0.178537369 | Immune response |

| | | | | | |
|---|---|---|---|---|---|
| Q32P28-4 | LEPRE1 | Prolyl 3-hydroxylase 1 | 0.365278244 | -0.945628484 | Cellular metabolism |
| Q32P41 | TRMT5 | tRNA (guanine(37)-N1)-methyltransferase | -0.154243469 | -0.501473109 | Other |
| Q3B726 | TWISTNB | DNA-directed RNA polymerase I subunit RPA43 | 0.068766276 | 0 | Cellular metabolism |
| Q3KQU3-2 | MAP7D1 | MAP7 domain-containing protein 1 | 0.209970474 | 1.574918747 | Other |
| Q3KQV9 | UAP1L1 | UDP-N-acetylhexosamine pyrophosphorylase-like protein 1 | -0.25514857 | -1.174152374 | Other |
| Q3YEC7 | RABL6 | Rab-like protein 6 | 0.181571325 | 0.491956711 | Other |
| Q3ZCM7 | TUBB8 | Tubulin beta-8 chain | 0.759577433 | 0 | Other |
| Q3ZCQ8 | TIMM50 | Mitochondrial import inner membrane translocase subunit TIM50 | 0.870394389 | 0.97718811 | Transport |
| Q3ZCW2 | LGALSL | Galectin-related protein | -0.153176626 | 0.277241389 | Other |
| Q49AR2 | C5orf22 | UPF0489 protein C5orf22 | -0.352886836 | 0.021225611 | Other |
| Q4G0J3 | LARP7 | La-related protein 7 | 0.032876968 | -0.811319351 | Cellular metabolism |
| Q4J6C6-4 | PREPL | Prolyl endopeptidase-like | -0.736007055 | -2.285486221 | Other |
| Q4V328 | GRIPAP1 | GRIP1-associated protein 1 | -0.241380056 | -0.327466965 | Other |
| Q4VC31 | CCDC58 | Coiled-coil domain-containing protein 58 | -0.555139542 | -0.351649602 | Other |
| Q52LG2 | KRTAP13-2 | Keratin-associated protein 13-2 | 0 | -0.063522975 | Other |
| Q52LJ0 | FAM98B | Protein FAM98B | 0.137961706 | 0.318612417 | Other |
| Q52LW3 | ARHGAP29 | Rho GTPase-activating protein 29 | -0.073340734 | -1.153673808 | Other |
| Q53EL6-2 | PDCD4 | Programmed cell death protein 4 | -1.206315994 | -0.964687347 | Other |
| Q53ET0 | CRTC2 | CREB-regulated transcription coactivator 2 | 0.30467097 | 0 | Cellular metabolism |
| Q53F19 | C17orf85 | Uncharacterized protein C17orf85 | -0.184429169 | 0.159400304 | Other |
| Q53FA7 | TP53I3 | Quinone oxidoreductase PIG3 | -0.024230321 | -0.289820989 | Cellular metabolism |
| Q53FT3 | C11orf73 | Protein Hikeshi | -0.440889994 | -0.224914551 | Transport |
| Q53GG5-2 | PDLIM3 | PDZ and LIM domain protein 3 | -0.450621923 | 0 | Other |
| Q53GQ0 | HSD17B12 | Very-long-chain 3-oxoacyl-CoA reductase | 0.543097178 | 1.85783577 | Lipid metabolism |
| Q53GT1 | KLHL22 | Kelch-like protein 22 | 0.005942027 | -0.56538264 | Cellular metabolism |
| Q53H82 | LACTB2 | Beta-lactamase-like protein 2 | -0.142894745 | -1.760923386 | Other |
| Q53LP3 | SOWAHC | Ankyrin repeat domain-containing protein SOWAHC | -0.055679321 | 0 | Other |
| Q53RT3 | ASPRV1 | Retroviral-like aspartic protease 1 | -1.103521983 | 0 | Other |
| Q53T59 | HS1BP3 | HCLS1-binding protein 3 | -0.143324534 | -0.899023692 | Other |
| Q562R1 | ACTBL2 | Beta-actin-like protein 2 | -1.155510585 | 0 | Other |
| Q56VL3 | OCIAD2 | OCIA domain-containing protein 2 | 0 | 1.398738225 | Other |
| Q58FF6 | HSP90AB4P | Putative heat shock protein HSP 90-beta 4 | 0.153496424 | 0.524595896 | Cellular metabolism |
| Q58FF8 | HSP90AB2P | Putative heat shock protein HSP 90-beta 2 | -0.045969009 | 1.176554998 | Cellular metabolism |
| Q58WW2 | DCAF6 | DDB1- and CUL4-associated factor 6 | -0.486170451 | 0 | Cellular metabolism |
| Q5BKZ1 | ZNF326 | DBIRD complex subunit ZNF326 | 0.869204839 | 0.020303726 | Cellular metabolism |

| | | | | | |
|---|---|---|---|---|---|
| Q5EBL4 | RILPL1 | RILP-like protein 1 | 0.065797806 | -1.203006109 | Transport |
| Q5F1R6 | DNAJC21 | DnaJ homolog subfamily C member 21 | -0.184919357 | 0 | Cellular metabolism |
| Q5GLZ8-3 | HERC4 | Probable E3 ubiquitin-protein ligase HERC4 | 0.032379786 | -1.898892085 | Cellular metabolism |
| Q5H909 | MAGED2 | Melanoma-associated antigen D2 | -0.318284353 | -0.63226064 | Other |
| Q5JSH3 | WDR44 | WD repeat-containing protein 44 | -0.167980194 | -0.702953339 | Other |
| Q5JTD0-2 | TJAP1 | Tight junction-associated protein 1 | -0.086008708 | -2.066036224 | Other |
| Q5JTH9-2 | RRP12 | RRP12-like protein | 0.002524058 | 0.078900655 | Other |
| Q5JTJ3-3 | COA6 | Cytochrome c oxidase assembly factor 6 homolog | -0.045182546 | -0.385735194 | Other |
| Q5JTV1 | GMEB2 | Glucocorticoid modulatory element-binding protein 2 | 0.795107524 | 0 | Cellular metabolism |
| Q5JVF3-3 | PCID2 | PCI domain-containing protein 2 | -0.006263097 | -0.738911947 | Cellular metabolism |
| Q5JY65 | CRNKL1 | Crooked neck-like protein 1 | 0.060513814 | -1.236864726 | Cellular metabolism |
| Q5MIZ7-2 | SMEK2 | Serine/threonine-protein phosphatase 4 regulatory subunit 3B | -0.204181035 | -2.381533305 | Cellular metabolism |
| Q5QJ74 | TBCEL | Tubulin-specific chaperone cofactor E-like protein | -0.696989059 | 0.230920792 | Other |
| Q5QPL9 | RALY | RNA-binding protein Raly | 0.444278717 | 0.339798609 | Cellular metabolism |
| Q5QPM7 | PSMF1 | Proteasome inhibitor PI31 subunit | -0.271051407 | -0.383406957 | Immune response |
| Q5R372-4 | RABGAP1L | Rab GTPase-activating protein 1-like | -0.932060877 | 0 | Other |
| Q5R3I4 | TTC38 | Tetratricopeptide repeat protein 38 | -0.41574796 | 0 | Other |
| Q5RKV6 | EXOSC6 | Exosome complex component MTR3 | 0.724096298 | 1.239173253 | Cellular metabolism |
| Q5SRE5 | NUP188 | Nucleoporin NUP188 homolog | -0.586761475 | -1.409988403 | Transport |
| Q5SSJ5 | HP1BP3 | Heterochromatin protein 1-binding protein 3 | -1.312842687 | -0.492448807 | Other |
| Q5STZ8 | ABCF1 | ATP-binding cassette sub-family F member 1 | -0.039028168 | -0.261170705 | Transport |
| Q5SW79 | CEP170 | Centrosomal protein of 170 kDa | 0.514657974 | -0.161284765 | Other |
| Q5SY16 | NOL9 | Polynucleotide 5-hydroxyl-kinase NOL9 | 0.104756037 | 0 | Cellular metabolism |
| Q5T0D9 | TPRG1L | Tumor protein p63-regulated gene 1-like protein | 0.402673086 | 1.164717992 | Other |
| Q5T123 | SH3BGRL3 | SH3 domain-binding glutamic acid-rich-like protein 3 | -0.037848155 | -0.175048192 | Other |
| Q5T1M5 | FKBP15 | FK506-binding protein 15 | -0.375271479 | 0.303077698 | Transport |
| Q5T4S7-3 | UBR4 | E3 ubiquitin-protein ligase UBR4 | -0.192467372 | -0.867907206 | Cellular metabolism |
| Q5T6F2 | UBAP2 | Ubiquitin-associated protein 2 | -0.082420349 | -1.100969315 | Other |
| Q5T6V5 | C9orf64 | UPF0553 protein C9orf64 | -0.131345749 | -1.274024328 | Other |
| Q5T749 | KPRP | Keratinocyte proline-rich protein | -0.641412099 | -0.259637833 | Other |
| Q5T750 | XP32 | Skin-specific protein 32 | 0 | 0.34139506 | Other |
| Q8N4J0 | C9orf41 | UPF0586 protein C9orf41 | -0.340796153 | 0 | Other |
| Q5TC12-3 | ATPAF1 | ATP synthase mitochondrial F1 complex assembly factor 1 | -0.050322215 | 0.334756215 | Other |
| Q5TDH0 | DDI2 | Protein DDI1 homolog 2 | -0.468130747 | -0.589758555 | Other |

| | | | | | |
|---|---|---|---|---|---|
| Q5TEC6 | HIST2H3PS2 | Histone H3 | 2.264222463 | -0.540826162 | Other |
| Q5TFE4 | NT5DC1 | 5-nucleotidase domain-containing protein 1 | -0.201066971 | -1.314288457 | Cellular metabolism |
| Q5VSL9 | STRIP1 | Striatin-interacting protein 1 | -0.461257299 | -0.579561869 | Other |
| Q5VTR2 | RNF20 | E3 ubiquitin-protein ligase BRE1A | -0.171939214 | -0.272187551 | Cellular metabolism |
| Q5VVQ6-2 | YOD1 | Ubiquitin thioesterase OTU1 | 0.207997004 | -0.545186996 | Cellular metabolism |
| Q5VW32 | BROX | BRO1 domain-containing protein BROX | 0.050429662 | -0.135587056 | Other |
| Q5VW36 | FOCAD | Focadhesin | 0.181978226 | 0 | Other |
| Q5VWJ9 | SNX30 | Sorting nexin-30 | -0.187928518 | 0 | Transport |
| Q5VWZ2 | LYPLAL1 | Lysophospholipase-like protein 1 | -0.193323771 | -0.496403376 | Cellular metabolism |
| Q5VZ89-5 | DENND4C | DENN domain-containing protein 4C | -0.754837672 | 0.096812566 | Transport |
| Q5VZE5 | NAA35 | N-alpha-acetyltransferase 35, NatC auxiliary subunit | 0.370300929 | 0 | Other |
| Q5W0V3 | FAM160B1 | Protein FAM160B1 | -0.38421758 | -1.272176107 | Other |
| Q60FE5 | FLNA | | 0.457466761 | 0 | Transport |
| Q63ZY3-3 | KANK2 | KN motif and ankyrin repeat domain-containing protein 2 | 0 | -1.381711324 | Cellular metabolism |
| Q641Q2-2 | FAM21A | WASH complex subunit FAM21A | -0.579840978 | -1.390565236 | Transport |
| Q643R3 | LPCAT4 | Lysophospholipid acyltransferase LPCAT4 | 0.737259547 | 4.331912994 | Lipid metabolism |
| Q658Y4 | FAM91A1 | Protein FAM91A1 | 0.236033758 | 0 | Other |
| Q66K14-2 | TBC1D9B | TBC1 domain family member 9B | 0.022152583 | -1.965356827 | Other |
| Q66K74-2 | MAP1S | Microtubule-associated protein 1S | -0.213742574 | 0 | Transport |
| Q66LE6 | PPP2R2D | Serine/threonine-protein phosphatase 2A 55 kDa regulatory subunit B delta isoform | 0.468797684 | 0.591071447 | Other |
| Q684P5-2 | RAP1GAP2 | Rap1 GTPase-activating protein 2 | 0.289016088 | -1.539299011 | Other |
| Q68CQ4 | DIEXF | Digestive organ expansion factor homolog | 0.058847427 | -1.139573415 | Cellular metabolism |
| Q68CZ2 | TNS3 | Tensin-3 | -0.062412262 | 0.212623596 | Other |
| Q68DQ2 | CRYBG3 | Very large A-kinase anchor protein | -0.6452624 | 0 | Other |
| Q68E01-3 | INTS3 | Integrator complex subunit 3 | 0.515464147 | 0 | Cellular metabolism |
| Q69YN2 | CWF19L1 | CWF19-like protein 1 | -0.1437397 | 0.043858846 | Other |
| Q6AI08 | HEATR6 | HEAT repeat-containing protein 6 | -0.512307485 | -0.478035609 | Other |
| Q6AI12 | ANKRD40 | Ankyrin repeat domain-containing protein 40 | 0.143934886 | -0.962741216 | Other |
| Q6DD87 | ZNF787 | Zinc finger protein 787 | 0.152442932 | 0.042903264 | Cellular metabolism |
| Q6DD88 | ATL3 | Atlastin-3 | 0.563508352 | 0.181361516 | Other |
| Q6DHV7-2 | ADAL | Adenosine deaminase-like protein | -0.231662114 | 0 | Cellular metabolism |
| Q6DKJ4 | NXN | Nucleoredoxin | -0.091319402 | -0.46291097 | Other |
| Q6FI81 | CIAPIN1 | Anamorsin | -0.23411115 | -0.30112648 | Cellular metabolism |
| Q6FIF0-2 | ZFAND6 | AN1-type zinc finger protein 6 | -0.585556666 | -1.862622579 | Transport |
| Q6GMV2 | SMYD5 | SET and MYND domain-containing protein 5 | -0.296929042 | -0.835924784 | Other |

| | | | | | |
|---|---|---|---|---|---|
| Q6GMV3 | PTRHD1 | Putative peptidyl-tRNA hydrolase PTRHD1 | -0.330067952 | 0 | Other |
| Q6IA69 | NADSYN1 | Glutamine-dependent NAD(+) synthetase | -0.131336848 | 0.24536705 | Cellular metabolism |
| Q6IA86-2 | ELP2 | Elongator complex protein 2 | -0.231423696 | -1.217880249 | Cellular metabolism |
| Q6IBS0 | TWF2 | Twinfilin-2 | -0.153999964 | -0.434980392 | Immune response |
| Q6IN85 | SMEK1 | Serine/threonine-protein phosphatase 4 regulatory subunit 3A | -0.015189489 | -0.365891774 | Cellular metabolism |
| Q6IQ49 | SDE2 | Protein SDE2 homolog | 0.356571833 | 2.849335353 | Other |
| Q6L8Q7 | PDE12 | 2,5-phosphodiesterase 12 | -0.179672877 | -0.142154058 | Cellular metabolism |
| Q6NUK1 | SLC25A24 | Calcium-binding mitochondrial carrier protein SCaMC-1 | 0 | 1.173400879 | Transport |
| Q6NUQ1 | RINT1 | RAD50-interacting protein 1 | 0.025058746 | 0.149219513 | Transport |
| Q6NXE6-2 | ARMC6 | Armadillo repeat-containing protein 6 | -0.120819092 | -1.094636917 | Other |
| Q6NXT1 | ANKRD54 | Ankyrin repeat domain-containing protein 54 | -0.08793513 | 0 | Transport |
| Q6NZI2 | PTRF | Polymerase I and transcript release factor | -0.535791397 | -0.268548965 | Cellular metabolism |
| Q6NZY4 | ZCCHC8 | Zinc finger CCHC domain-containing protein 8 | 0.173070908 | -0.262786229 | Cellular metabolism |
| Q6P179-3 | ERAP2 | Endoplasmic reticulum aminopeptidase 2 | 0.426118215 | -0.353049596 | Other |
| Q6P1J9 | CDC73 | Parafibromin | 0.123189926 | -0.05382665 | Cellular metabolism |
| Q6P1K2-4 | PMF1 | Polyamine-modulated factor 1 | -0.638243357 | 0.382516225 | Cellular metabolism |
| Q6P1Q9 | METTL2B | Methyltransferase-like protein 2B | -0.794627508 | -1.262086233 | Cellular metabolism |
| Q6P1R4 | DUS1L | tRNA-dihydrouridine(16/17) synthase [NAD(P)(+)]-like | -0.612193425 | -2.011107127 | Other |
| Q6P2E9 | EDC4 | Enhancer of mRNA-decapping protein 4 | 0.421972911 | -0.950423559 | Cellular metabolism |
| Q6P2P2 | PRMT9 | Putative protein arginine N-methyltransferase 9 | -0.835717519 | -1.976151148 | Cellular metabolism |
| Q6P2Q9 | PRPF8 | Pre-mRNA-processing-splicing factor 8 | -0.072940826 | -0.351715724 | Cellular metabolism |
| Q6P3X3 | TTC27 | Tetratricopeptide repeat protein 27 | -0.072076162 | -1.257404327 | Other |
| Q6P4I2 | WDR73 | WD repeat-containing protein 73 | -0.161259333 | -0.371301651 | Other |
| Q6P587 | FAHD1 | Acylpyruvase FAHD1, mitochondrial | 0.171504339 | 0.014392217 | Other |
| Q6P996-4 | PDXDC1 | Pyridoxal-dependent decarboxylase domain-containing protein 1 | -0.017659505 | -1.324012756 | Cellular metabolism |
| Q6P9B6 | TLDC1 | TLD domain-containing protein 1 | -0.257101695 | -0.396940867 | Other |
| Q6PCE3 | PGM2L1 | Glucose 1,6-bisphosphate synthase | -0.17407163 | 0.579951604 | Cellular metabolism |
| Q6PD62 | CTR9 | RNA polymerase-associated protein CTR9 homolog | 0.166957219 | 0.363765717 | Cellular metabolism |
| Q6PD74 | AAGAB | Alpha- and gamma-adaptin-binding protein p34 | -0.2978007 | -1.060085932 | Transport |
| Q6PJG6 | BRAT1 | BRCA1-associated ATM activator 1 | -0.386049271 | -0.820163091 | Other |
| Q6PKG0 | LARP1 | La-related protein 1 | -0.105798086 | -0.09086291 | Other |
| Q6PL24 | TMED8 | Protein TMED8 | -0.173668543 | -1.979929606 | Transport |
| Q6QNY0 | BLOC1S3 | Biogenesis of lysosome-related organelles complex 1 subunit 3 | 0.640441895 | 0 | Transport |
| Q6SZW1 | SARM1 | Sterile alpha and TIR motif-containing protein 1 | 0.482629776 | 0 | Immune response |

| | | | | | |
|---|---|---|---|---|---|
| Q6UN15-4 | FIP1L1 | Pre-mRNA 3-end-processing factor FIP1 | -0.703868866 | 0.036652883 | Cellular metabolism |
| Q6UUV9-3 | CRTC1 | CREB-regulated transcription coactivator 1 | -0.001768748 | 0 | Cellular metabolism |
| Q6UVK1 | CSPG4 | Chondroitin sulfate proteoglycan 4 | -0.1939888 | -0.944069544 | Cellular metabolism |
| Q6UWP2 | DHRS11 | Dehydrogenase/reductase SDR family member 11 | -0.607229233 | -0.361769994 | Other |
| Q6UWP8 | SBSN | Suprabasin | 0.468786875 | 0.976900736 | Other |
| Q6UX04 | CWC27 | Peptidyl-prolyl cis-trans isomerase CWC27 homolog | 0.090606054 | -1.549646378 | Cellular metabolism |
| Q6UXH1-4 | CRELD2 | Cysteine-rich with EGF-like domain protein 2 | 0.197102865 | -0.623202006 | Other |
| Q6UXN9 | WDR82 | WD repeat-containing protein 82 | -0.080709457 | 0.043777466 | Cellular metabolism |
| Q6V1X1-3 | DPP8 | Dipeptidyl peptidase 8 | -0.199530919 | 0.530628204 | Immune response |
| Q6VY07 | PACS1 | Phosphofurin acidic cluster sorting protein 1 | -0.206361135 | -0.080956777 | Transport |
| Q6WKZ4-3 | RAB11FIP1 | Rab11 family-interacting protein 1 | -0.243287404 | 2.467510223 | Transport |
| Q6XQN6 | NAPRT | Nicotinate phosphoribosyltransferase | 0.008660634 | -1.123778025 | Cellular metabolism |
| Q6XZF7 | DNMBP | Dynamin-binding protein | 0.07483991 | 0.402874629 | Other |
| Q6Y7W6 | GIGYF2 | PERQ amino acid-rich with GYF domain-containing protein 2 | 0.253814697 | -1.361063004 | Cellular metabolism |
| Q6YHU6-3 | THADA | Thyroid adenoma-associated protein | -0.401374181 | -1.425489426 | Other |
| Q6YP21-3 | CCBL2 | Kynurenine--oxoglutarate transaminase 3 | -0.079151789 | -0.355541865 | Cellular metabolism |
| Q6ZS17-2 | FAM65A | Protein FAM65A | -0.20675532 | 0.069671631 | Other |
| Q6ZUT6-2 | C15orf52 | Uncharacterized protein C15orf52 | 0.15715917 | 1.573168437 | Other |
| Q6ZVM7 | TOM1L2 | TOM1-like protein 2 | -0.124621073 | -1.525714874 | Transport |
| Q6ZVX7 | NCCRP1 | F-box only protein 50 | -0.241783778 | 3.0242939 | Other |
| Q70CQ2-3 | USP34 | Ubiquitin carboxyl-terminal hydrolase 34 | -1.247950236 | 0 | Cellular metabolism |
| Q70UQ0-4 | IKBIP | Inhibitor of nuclear factor kappa-B kinase-interacting protein | -0.246518453 | 1.254441579 | Other |
| Q712K3 | UBE2R2 | Ubiquitin-conjugating enzyme E2 R2 | -0.322125753 | -0.85294342 | Cellular metabolism |
| Q71RC2-3 | LARP4 | La-related protein 4 | 0.117314021 | -0.448689779 | Other |
| Q7KZ85 | SUPT6H | Transcription elongation factor SPT6 | 0.028299967 | -0.585136414 | Transport |
| Q7KZF4 | SND1 | Staphylococcal nuclease domain-containing protein 1 | 0.11240832 | -0.266557693 | Cellular metabolism |
| Q7L014 | DDX46 | Probable ATP-dependent RNA helicase DDX46 | 0.525400798 | -0.934298833 | Cellular metabolism |
| Q7L0Y3 | TRMT10C | Mitochondrial ribonuclease P protein 1 | -0.187648773 | 0.128454844 | Cellular metabolism |
| Q7L1Q6 | BZW1 | Basic leucine zipper and W2 domain-containing protein 1 | -0.045763016 | -0.259823481 | Cellular metabolism |
| Q7L1T6 | CYB5R4 | Cytochrome b5 reductase 4 | 0.145068487 | 0 | Transport |
| Q7L266 | ASRGL1 | Isoaspartyl peptidase/L-asparaginase | -0.306352615 | -0.77391688 | Cellular metabolism |
| Q7L273 | KCTD9 | BTB/POZ domain-containing protein KCTD9 | -1.031234741 | 0 | Other |
| Q7L2H7 | EIF3M | Eukaryotic translation initiation factor 3 subunit M | 0.046183904 | -1.602782567 | Other |

| | | | | | |
|---|---|---|---|---|---|
| Q7L4I2-2 | RSRC2 | Arginine/serine-rich coiled-coil protein 2 | -0.066526413 | 0 | Other |
| Q7L523 | RRAGA | Ras-related GTP-binding protein A | -0.232302984 | 0.098864237 | Other |
| Q7L576 | CYFIP1 | Cytoplasmic FMR1-interacting protein 1 | -0.135394414 | -0.507441203 | Immune response |
| Q7L5D6 | GET4 | Golgi to ER traffic protein 4 homolog | -0.220932643 | -0.405793508 | Transport |
| Q7L775 | EPM2AIP1 | EPM2A-interacting protein 1 | 0.009565989 | -0.564182281 | Other |
| Q7L8J4 | SH3BP5L | SH3 domain-binding protein 5-like | -0.046260834 | 0 | Other |
| Q7L8W6 | DPH6 | Diphthine--ammonia ligase | -0.561506271 | 0.716025035 | Cellular metabolism |
| Q7L9L4 | MOB1B | MOB kinase activator 1B | -0.256235123 | -1.333617528 | Cellular metabolism |
| Q7LBC6 | KDM3B | Lysine-specific demethylase 3B | -0.291214625 | 0.179559708 | Cellular metabolism |
| Q7LBR1 | CHMP1B | Charged multivesicular body protein 1b | 0.248243332 | 0.316115061 | Transport |
| Q7LG56 | RRM2B | Ribonucleoside-diphosphate reductase subunit M2 B | -0.240029653 | -2.554835002 | Cellular metabolism |
| Q7RTP6 | MICAL3 | Protein-methionine sulfoxide oxidase MICAL3 | 0.500607808 | 0.26960818 | Transport |
| Q7RTV0 | PHF5A | PHD finger-like domain-containing protein 5A | 0.544101079 | -0.292311351 | Cellular metabolism |
| Q7Z2T5 | TRMT1L | TRMT1-like protein | 0.93326505 | -0.919257482 | Cellular metabolism |
| Q7Z2W4 | ZC3HAV1 | Zinc finger CCCH-type antiviral protein 1 | 0.523173014 | 1.833167394 | Immune response |
| Q7Z2Z2 | EFTUD1 | Elongation factor Tu GTP-binding domain-containing protein 1 | 0.125418345 | -0.828343074 | Other |
| Q7Z392-4 | TRAPPC11 | Trafficking protein particle complex subunit 11 | 0.039986928 | -1.424374262 | Transport |
| Q7Z3B4-2 | NUP54 | Nucleoporin p54 | -0.52202479 | 1.351148605 | Transport |
| Q7Z3T8 | ZFYVE16 | Zinc finger FYVE domain-containing protein 16 | -0.023180644 | -0.796400706 | Transport |
| Q7Z406 | MYH14 | Myosin-14 | 0.077465693 | -2.687325795 | Other |
| Q7Z417 | NUFIP2 | Nuclear fragile X mental retardation-interacting protein 2 | -0.079910278 | -1.963677088 | Other |
| Q7Z422-5 | SZRD1 | SUZ domain-containing protein 1 | -0.480660756 | 0 | Other |
| Q7Z434 | MAVS | Mitochondrial antiviral-signaling protein | 1.056013743 | 0 | Immune response |
| Q7Z4G4-2 | TRMT11 | tRNA (guanine(10)-N2)-methyltransferase homolog | -0.213654836 | 0 | Cellular metabolism |
| Q7Z4H3 | HDDC2 | HD domain-containing protein 2 | -0.11924235 | 0.207979202 | Other |
| Q7Z4H7-3 | HAUS6 | HAUS augmin-like complex subunit 6 | 0.161251704 | -0.604207993 | Other |
| Q7Z4H8 | KDELC2 | KDEL motif-containing protein 2 | 0.296773275 | -1.056541443 | Other |
| Q7Z4Q2 | HEATR3 | HEAT repeat-containing protein 3 | 0.075965246 | -1.391446431 | Other |
| Q7Z4V5 | HDGFRP2 | Hepatoma-derived growth factor-related protein 2 | 0.036233902 | -0.38405927 | Cellular metabolism |
| Q7Z5L9 | IRF2BP2 | Interferon regulatory factor 2-binding protein 2 | 0.366794586 | -1.829414368 | Cellular metabolism |
| Q7Z5R6 | APBB1IP | Amyloid beta A4 precursor protein-binding family B member 1-interacting protein | -0.043641408 | -1.09170723 | Other |
| Q7Z6K3 | PTAR1 | Protein prenyltransferase alpha subunit repeat-containing protein 1 | -0.950918198 | 1.77009964 | Cellular metabolism |
| Q7Z6M1 | RABEPK | Rab9 effector protein with kelch motifs | -0.340174357 | -0.867137273 | Transport |

| | | | | | |
|---|---|---|---|---|---|
| Q7Z6Z7-2 | HUWE1 | E3 ubiquitin-protein ligase HUWE1 | -0.14206632 | -1.490479787 | Cellular metabolism |
| Q7Z7E8 | UBE2Q1 | Ubiquitin-conjugating enzyme E2 Q1 | 0.083958308 | -0.535868327 | Transport |
| Q86SQ9-3 | DHDDS | Dehydrodolichyl diphosphate syntase complex subunit DHDDS | -0.305229187 | 4.027943293 | Lipid metabolism |
| Q86SZ2 | TRAPPC6B | Trafficking protein particle complex subunit 6B | 0.364611944 | 0.462538401 | Transport |
| Q86TB9-4 | PATL1 | Protein PAT1 homolog 1 | 0.632898331 | 0.537272135 | Cellular metabolism |
| Q86TI2 | DPP9 | Dipeptidyl peptidase 9 | 0.002648671 | -0.675725301 | Other |
| Q86TP1 | PRUNE | Protein prune homolog | -0.246925354 | 0 | Cellular metabolism |
| Q86TU7 | SETD3 | Histone-lysine N-methyltransferase setd3 | 0.083680471 | -0.552078883 | Cellular metabolism |
| Q86U28 | ISCA2 | Iron-sulfur cluster assembly 2 homolog, mitochondrial | -1.15700531 | 0 | Cellular metabolism |
| Q86U38 | NOP9 | Nucleolar protein 9 | 0.288756053 | 0 | Other |
| Q86U42-2 | PABPN1 | Polyadenylate-binding protein 2 | 0.190828323 | 1.870049159 | Transport |
| Q86U44 | METTL3 | N6-adenosine-methyltransferase 70 kDa subunit | 0.342369715 | -0.273190816 | Cellular metabolism |
| Q86U90 | YRDC | YrdC domain-containing protein, mitochondrial | -1.065489451 | -0.526724497 | Other |
| Q86UA1 | PRPF39 | Pre-mRNA-processing factor 39 | -0.07773908 | -0.960287094 | Cellular metabolism |
| Q86UA6 | RPAIN | RPA-interacting protein | 0.106536865 | -1.053846359 | Transport |
| Q86UE4 | MTDH | Protein LYRIC | 1.602318446 | -0.383840561 | Other |
| Q86UL3 | AGPAT6 | Glycerol-3-phosphate acyltransferase 4 | 0.445399602 | 1.730294545 | Lipid metabolism |
| Q86UP2-2 | KTN1 | Kinectin | -0.365346909 | 0.813510895 | Other |
| Q86UV5 | USP48 | Ubiquitin carboxyl-terminal hydrolase 48 | 0 | -1.824608485 | Cellular metabolism |
| Q86V21 | AACS | Acetoacetyl-CoA synthetase | 0.033753713 | -0.273187002 | Lipid metabolism |
| Q86V48-2 | LUZP1 | Leucine zipper protein 1 | 1.131251017 | -1.006340663 | Other |
| Q86V81 | ALYREF | THO complex subunit 4 | 0.263701757 | 0.440308889 | Transport |
| Q86VP6 | CAND1 | Cullin-associated NEDD8-dissociated protein 1 | -0.028608322 | -0.257970174 | Cellular metabolism |
| Q86VS8 | HOOK3 | Protein Hook homolog 3 | 0.200651169 | 0.534522374 | Transport |
| Q86W50 | METTL16 | Methyltransferase-like protein 16 | 0.087904612 | -0.908610662 | Cellular metabolism |
| Q86W92-3 | PPFIBP1 | Liprin-beta-1 | 0.015349706 | 0 | Other |
| Q86WN1 | FCHSD1 | F-BAR and double SH3 domains protein 1 | -0.058217367 | 0 | Other |
| Q86WQ0 | NR2C2AP | Nuclear receptor 2C2-associated protein | 0.467650731 | -0.091766357 | Cellular metabolism |
| Q86WR0 | CCDC25 | Coiled-coil domain-containing protein 25 | -0.263608932 | 0.236407598 | Other |
| Q86X27-3 | RALGPS2 | Ras-specific guanine nucleotide-releasing factor RalGPS2 | 0.01472791 | 0 | Other |
| Q86X55-1 | CARM1 | Histone-arginine methyltransferase CARM1 | -0.049430847 | 0.132921219 | Lipid metabolism |
| Q86X76-2 | NIT1 | Nitrilase homolog 1 | -0.095781326 | -0.621199926 | Other |
| Q86X83-2 | COMMD2 | COMM domain-containing protein 2 | -0.096445719 | 0 | Cellular metabolism |
| S4R446 | MEX3D | RNA-binding protein MEX3D | 0.241849264 | -0.419787725 | Cellular metabolism |

| | | | | | |
|---|---|---|---|---|---|
| Q86Y37 | CACUL1 | CDK2-associated and cullin domain-containing protein 1 | -0.251242956 | 0 | Cellular metabolism |
| Q86Y56 | DNAAF5 | Dynein assembly factor 5, axonemal | -0.12664477 | -1.062936147 | Other |
| Q8IU85 | CAMK1D | Calcium/calmodulin-dependent protein kinase type 1D | -1.663286209 | 0 | Other |
| Q8IUF8-4 | MINA | Bifunctional lysine-specific demethylase and histidyl-hydroxylase MINA | 0.841151555 | 0 | Other |
| Q8IUI8 | CRLF3 | Cytokine receptor-like factor 3 | 0.103391647 | -1.085335414 | Other |
| Q8IUR0 | TRAPPC5 | Trafficking protein particle complex subunit 5 | -0.127739588 | 0.060658137 | Transport |
| Q8IUR7-3 | ARMC8 | Armadillo repeat-containing protein 8 | 0.26698939 | -1.081611633 | Other |
| Q8IV08 | PLD3 | Phospholipase D3 | -0.021966298 | 0.010297775 | Lipid metabolism |
| Q8IV48 | ERI1 | 3-5 exoribonuclease 1 | 0.430042267 | 0.467720668 | Cellular metabolism |
| Q8IV50 | LYSMD2 | LysM and putative peptidoglycan-binding domain-containing protein 2 | -0.056362152 | -1.064513524 | Other |
| Q8IVD9 | NUDCD3 | NudC domain-containing protein 3 | -0.197818756 | -1.678781509 | Other |
| Q8IVF2-3 | AHNAK2 | Protein AHNAK2 | 0.639214198 | -2.337247849 | Other |
| Q8IVL6 | LEPREL2 | Prolyl 3-hydroxylase 3 | -0.286876678 | -0.773596446 | Other |
| Q8IVM0 | CCDC50 | Coiled-coil domain-containing protein 50 | -0.048938115 | -1.504053116 | Other |
| Q8IW35 | CEP97 | Centrosomal protein of 97 kDa | -0.482108434 | 0 | Other |
| Q8IW75 | SERPINA12 | Serpin A12 | 0 | 0.089183807 | Other |
| Q8IWB7 | WDFY1 | WD repeat and FYVE domain-containing protein 1 | -0.193309148 | -0.977568944 | Other |
| Q8IWE2 | FAM114A1 | Protein NOXP20 | 0.041789373 | -1.516820908 | Other |
| Q8IWJ2 | GCC2 | GRIP and coiled-coil domain-containing protein 2 | -0.525849024 | -1.779399872 | Transport |
| Q8IWT0-2 | ZBTB8OS | Protein archease | 0.024445216 | -0.352245331 | Cellular metabolism |
| Q8IWV7 | UBR1 | E3 ubiquitin-protein ligase UBR1 | -0.217556636 | 0.354769389 | Cellular metabolism |
| Q8IWV8-4 | UBR2 | E3 ubiquitin-protein ligase UBR2 | -0.808741252 | -1.24321874 | Cellular metabolism |
| Q8IWY9-1 | CDAN1 | Codanin-1 | 0.469228109 | 0 | Other |
| Q8IWZ8 | SUGP1 | SURP and G-patch domain-containing protein 1 | 0 | -0.559824626 | Cellular metabolism |
| Q8IX04-6 | UEVLD | Ubiquitin-conjugating enzyme E2 variant 3 | -0.49346288 | 0 | Transport |
| Q8IX12-2 | CCAR1 | Cell division cycle and apoptosis regulator protein 1 | 0.501931508 | -0.09301122 | Cellular metabolism |
| Q8IX90-3 | SKA3 | Spindle and kinetochore-associated protein 3 | 0.12444814 | 0 | Other |
| Q8IXJ6-4 | SIRT2 | NAD-dependent protein deacetylase sirtuin-2 | -0.461326599 | -1.200785955 | Immune response |
| Q8IXM3 | MRPL41 | 39S ribosomal protein L41, mitochondrial | 0 | -0.814441681 | Cellular metabolism |
| Q8IXT5 | RBM12B | RNA-binding protein 12B | 0.281772614 | -0.116059621 | Other |
| Q8IY47 | KBTBD2 | Kelch repeat and BTB domain-containing protein 2 | -0.540061315 | 0 | Other |
| Q8IY81 | FTSJ3 | pre-rRNA processing protein FTSJ3 | 0.801955541 | -0.956379573 | Cellular metabolism |
| Q8IYB1 | MB21D2 | Protein MB21D2 | 0.132011414 | -2.210236231 | Other |

| | | | | | |
|---|---|---|---|---|---|
| Q8IYB5-3 | SMAP1 | Stromal membrane-associated protein 1 | -0.786094666 | 0 | Other |
| Q8IYB7 | DIS3L2 | DIS3-like exonuclease 2 | 0.235300064 | -2.070709864 | Cellular metabolism |
| Q8IYD1 | GSPT2 | Eukaryotic peptide chain release factor GTP-binding subunit ERF3B | -0.334916433 | -0.672316869 | Cellular metabolism |
| Q8IYR2 | SMYD4 | SET and MYND domain-containing protein 4 | 0.378456116 | -0.419232051 | Other |
| Q8IYS1 | PM20D2 | Peptidase M20 domain-containing protein 2 | -0.246904373 | -1.252538681 | Other |
| Q8IZ07 | ANKRD13A | Ankyrin repeat domain-containing protein 13A | -0.615813573 | -0.718903859 | Other |
| Q8IZ73 | RPUSD2 | RNA pseudouridylate synthase domain-containing protein 2 | 0.221081416 | -0.914933523 | Cellular metabolism |
| Q8IZ81 | ELMOD2 | ELMO domain-containing protein 2 | 1.04108429 | 2.639913559 | Transport |
| Q8IZ83-3 | ALDH16A1 | Aldehyde dehydrogenase family 16 member A1 | -0.059768677 | 0.21539561 | Other |
| Q8IZV5 | RDH10 | Retinol dehydrogenase 10 | 0.020922979 | 2.789551417 | Lipid metabolism |
| Q8N0Z6 | TTC5 | Tetratricopeptide repeat protein 5 | -0.92420133 | -0.712053935 | Cellular metabolism |
| Q8N108-17 | MIER1 | Mesoderm induction early response protein 1 | 0.061844508 | -0.66272227 | Cellular metabolism |
| Q8N122 | RPTOR | Regulatory-associated protein of mTOR | 0.294073105 | 0 | Cellular metabolism |
| Q8N128 | FAM177A1 | Protein FAM177A1 | -0.39358139 | 3.581485748 | Other |
| Q8N129 | CNPY4 | Protein canopy homolog 4 | -0.265638351 | 0 | Other |
| Q8N163 | CCAR2 | Cell cycle and apoptosis regulator protein 2 | 0.581674576 | 0.371767044 | Cellular metabolism |
| Q8N1F7 | NUP93 | Nuclear pore complex protein Nup93 | -0.088087718 | -0.068319321 | Transport |
| Q8N1G2 | CMTR1 | Cap-specific mRNA (nucleoside-2-O-)-methyltransferase 1 | 0.027588526 | 0 | Cellular metabolism |
| Q8N1G4 | LRRC47 | Leucine-rich repeat-containing protein 47 | 0.112653732 | -0.629744212 | Other |
| Q8N1Q1 | CA13 | Carbonic anhydrase 13 | -0.49967893 | 0 | Transport |
| Q8N335 | GPD1L | Glycerol-3-phosphate dehydrogenase 1-like protein | -0.119803747 | -1.743289948 | Lipid metabolism |
| Q8N392 | ARHGAP18 | Rho GTPase-activating protein 18 | -0.301317215 | -0.379831314 | Other |
| Q8N3X1 | FNBP4 | Formin-binding protein 4 | -0.226806641 | 0 | Other |
| Q8N543 | OGFOD1 | Prolyl 3-hydroxylase OGFOD1 | -0.024941127 | -1.094755173 | Cellular metabolism |
| Q8N556 | AFAP1 | Actin filament-associated protein 1 | -0.050237656 | 0.325459798 | Other |
| Q8N573-2 | OXR1 | Oxidation resistance protein 1 | 0.084782283 | -0.594790141 | Other |
| Q8N584 | TTC39C | Tetratricopeptide repeat protein 39C | -0.647578557 | -0.049631755 | Other |
| Q8N5A5-3 | ZGPAT | Zinc finger CCCH-type with G patch domain-containing protein | 0.546718597 | -0.267336528 | Other |
| Q8N5K1 | CISD2 | CDGSH iron-sulfur domain-containing protein 2 | 0 | 2.002351125 | Cellular metabolism |
| Q8N5M4 | TTC9C | Tetratricopeptide repeat protein 9C | -0.149483363 | 0 | Other |
| Q8N668 | COMMD1 | COMM domain-containing protein 1 | -0.262924194 | 0 | Transport |
| Q8N6M0 | OTUD6B | OTU domain-containing protein 6B | -0.345606486 | -0.991303762 | Other |
| Q8N6N3 | C1orf52 | UPF0690 protein C1orf52 | -0.189779917 | 0.045361201 | Other |
| Q8N6R0 | METTL13 | Methyltransferase-like protein 13 | -0.287383397 | 0 | Other |

| | | | | | |
|---|---|---|---|---|---|
| Q8N6T3 | ARFGAP1 | ADP-ribosylation factor GTPase-activating protein 1 | 0.492495219 | 1.882769267 | Transport |
| Q8N766-4 | EMC1 | ER membrane protein complex subunit 1 | 0 | 0.862290064 | Other |
| Q8N7H5 | PAF1 | RNA polymerase II-associated factor 1 homolog | 0.060337702 | -0.509305318 | Cellular metabolism |
| Q8N806 | UBR7 | Putative E3 ubiquitin-protein ligase UBR7 | -0.361915588 | -0.509433111 | Cellular metabolism |
| Q8N8N7 | PTGR2 | Prostaglandin reductase 2 | -0.006584803 | -1.541368484 | Lipid metabolism |
| Q8N8R5 | C2orf69 | UPF0565 protein C2orf69 | -0.088319143 | 0 | Other |
| Q8N8S7-3 | ENAH | Protein enabled homolog | 0.406641006 | -0.696743647 | Transport |
| Q8N999 | C12orf29 | Uncharacterized protein C12orf29 | -0.086395899 | 0.278763453 | Other |
| Q8N9V3 | WDSUB1 | WD repeat, SAM and U-box domain-containing protein 1 | -0.258593241 | 0 | Other |
| Q8NAV1 | PRPF38A | Pre-mRNA-splicing factor 38A | 0.237927755 | 0 | Cellular metabolism |
| Q8NB37 | PDDC1 | Parkinson disease 7 domain-containing protein 1 | -0.38898023 | -0.904998779 | Other |
| Q8NB46 | ANKRD52 | Serine/threonine-protein phosphatase 6 regulatory ankyrin repeat subunit C | -1.269824346 | -3.211023966 | Other |
| Q8NBF2 | NHLRC2 | NHL repeat-containing protein 2 | -0.159631729 | -1.941761653 | Other |
| Q8NBJ5 | COLGALT1 | Procollagen galactosyltransferase 1 | 0.814016978 | 0.637826284 | Other |
| Q8NBQ5 | HSD17B11 | Estradiol 17-beta-dehydrogenase 11 | 0.524435679 | 2.048348109 | Lipid metabolism |
| Q8NBS9 | TXNDC5 | Thioredoxin domain-containing protein 5 | 0.211720785 | -0.160032272 | Transport |
| Q8NBX0 | SCCPDH | Saccharopine dehydrogenase-like oxidoreductase | 0.793271383 | 2.253985087 | Other |
| Q8NC51-3 | SERBP1 | Plasminogen activator inhibitor 1 RNA-binding protein | 0.164562225 | -0.261707306 | Cellular metabolism |
| Q8NC56-2 | LEMD2 | LEM domain-containing protein 2 | 0 | 1.440635045 | Other |
| Q8NCA5 | FAM98A | Protein FAM98A | -0.299968084 | 0 | Other |
| Q8NCC3 | PLA2G15 | Group XV phospholipase A2 | 0.038941701 | -0.636360168 | Lipid metabolism |
| Q8NCE2-3 | MTMR14 | Myotubularin-related protein 14 | -0.743807475 | 0 | Lipid metabolism |
| Q8NCF5 | NFATC2IP | NFATC2-interacting protein | 0.029534022 | -0.819800695 | Other |
| Q8NCW5 | APOA1BP | NAD(P)H-hydrate epimerase | -0.254301071 | -0.677691142 | Cellular metabolism |
| Q8ND24 | RNF214 | RING finger protein 214 | 0.444262822 | 0 | Other |
| Q8ND56-3 | LSM14A | Protein LSM14 homolog A | -0.188836416 | -0.239255905 | Other |
| Q8NDH3-3 | NPEPL1 | Probable aminopeptidase NPEPL1 | 0.061029434 | 0 | Other |
| Q8NDI1-3 | EHBP1 | EH domain-binding protein 1 | -0.422803243 | 0 | Other |
| Q8NDW4 | ZNF248 | Zinc finger protein 248 | 0.618034999 | 0 | Cellular metabolism |
| Q8NEC7 | GSTCD | Glutathione S-transferase C-terminal domain-containing protein | -0.072994232 | 0 | Other |
| Q8NEM2 | SHCBP1 | SHC SH2 domain-binding protein 1 | -0.701402028 | -1.757331212 | Other |
| Q8NEU8 | APPL2 | DCC-interacting protein 13-beta | -0.366195043 | -1.99779892 | Other |
| Q8NEZ2 | VPS37A | Vacuolar protein sorting-associated protein 37A | -0.487048467 | -0.763724009 | Transport |
| Q8NEZ5 | FBXO22 | F-box only protein 22 | -0.431170781 | -0.724145889 | Transport |

| | | | | | |
|---|---|---|---|---|---|
| Q8NFC6 | BOD1L1 | Biorientation of chromosomes in cell division protein 1-like 1 | 0.238745372 | 0 | Other |
| Q8NFF5-3 | FLAD1 | FAD synthase | -0.113345464 | -2.088289897 | Cellular metabolism |
| Q8NFG4 | FLCN | Folliculin | -0.779161453 | 0 | Other |
| Q8NFH3 | NUP43 | Nucleoporin Nup43 | -0.212855657 | 0.197846731 | Transport |
| Q8NFH4 | NUP37 | Nucleoporin Nup37 | -0.216864268 | 0.142501831 | Transport |
| Q8NFT8 | DNER | Delta and Notch-like epidermal growth factor-related receptor | -0.602617264 | -1.514677684 | Transport |
| Q8NG68 | TTL | Tubulin--tyrosine ligase | -0.204182943 | -0.784036001 | Cellular metabolism |
| Q8NHP8 | PLBD2 | Putative phospholipase B-like 2 | -0.031884511 | -0.296386083 | Lipid metabolism |
| Q8TAF3-4 | WDR48 | WD repeat-containing protein 48 | 0.383399963 | 0 | Cellular metabolism |
| Q8TAT6 | NPLOC4 | Nuclear protein localization protein 4 homolog | 0.079374313 | -0.124241511 | Cellular metabolism |
| Q8TB03 | CXorf38 | Uncharacterized protein CXorf38 | -0.215865453 | -0.747386297 | Other |
| Q8TB52 | FBXO30 | F-box only protein 30 | -0.229548136 | 0 | Other |
| Q8TBC4 | UBA3 | NEDD8-activating enzyme E1 catalytic subunit | 0.104484558 | -1.573599497 | Immune response |
| Q8TBE9 | NANP | N-acylneuraminate-9-phosphatase | -0.051916122 | 0 | Cellular metabolism |
| Q8TBX8-3 | PIP4K2C | Phosphatidylinositol 5-phosphate 4-kinase type-2 gamma | 0.189996084 | -0.007048925 | Other |
| Q8TBZ6 | TRMT10A | tRNA methyltransferase 10 homolog A | -0.252307892 | 0 | Cellular metabolism |
| Q8TC07-2 | TBC1D15 | TBC1 domain family member 15 | 0.02306366 | -0.824630102 | Other |
| Q8TC12 | RDH11 | Retinol dehydrogenase 11 | 0.844935099 | 1.751280467 | Lipid metabolism |
| Q8TCA0 | LRRC20 | Leucine-rich repeat-containing protein 20 | 0.1066583 | -0.235256195 | Other |
| Q8TCD1 | C18orf32 | UPF0729 protein C18orf32 | 0 | 4.967929204 | Other |
| Q8TCD5 | NT5C | 5(3)-deoxyribonucleotidase, cytosolic type | -0.05920283 | -0.818375905 | Cellular metabolism |
| Q8TCE6-2 | FAM45A | Protein FAM45A | -0.02644221 | -2.547993342 | Other |
| Q8TCF1 | ZFAND1 | AN1-type zinc finger protein 1 | -0.085768382 | 0.917761485 | Other |
| Q8TCJ2 | STT3B | Dolichyl-diphosphooligosaccharide--protein glycosyltransferase subunit STT3B | 0 | 0.610673269 | Cellular metabolism |
| Q8TCS8 | PNPT1 | Polyribonucleotide nucleotidyltransferase 1, mitochondrial | 0 | 1.934031169 | Transport |
| Q8TCY9 | URGCP | Up-regulator of cell proliferation | -0.474210103 | 0 | Other |
| Q8TD16 | BICD2 | Protein bicaudal D homolog 2 | -0.176551819 | -0.666945775 | Transport |
| Q8TD55 | PLEKHO2 | Pleckstrin homology domain-containing family O member 2 | 0.065612793 | -1.599333445 | Other |
| Q8TDN6 | BRIX1 | Ribosome biogenesis protein BRX1 homolog | -0.37122345 | 0.507036845 | Other |
| Q8TDQ7 | GNPDA2 | Glucosamine-6-phosphate isomerase 2 | -0.291805267 | -0.85153389 | Cellular metabolism |
| Q8TDX7 | NEK7 | Serine/threonine-protein kinase Nek7 | 0.119174957 | -0.725603739 | Cellular metabolism |
| Q8TDY2-2 | RB1CC1 | RB1-inducible coiled-coil protein 1 | -0.132677078 | -6.088322322 | Cellular metabolism |
| Q8TDZ2 | MICAL1 | Protein-methionine sulfoxide oxidase MICAL1 | -0.159410477 | -0.734532038 | Cellular metabolism |

| | | | | | |
|---|---|---|---|---|---|
| Q8TEA8 | DTD1 | D-tyrosyl-tRNA(Tyr) deacylase 1 | -0.025569916 | -0.658575694 | Cellular metabolism |
| Q8TEQ6 | GEMIN5 | Gem-associated protein 5 | 0.061197917 | -0.80396843 | Cellular metabolism |
| Q8TEX9 | IPO4 | Importin-4 | -0.041618983 | -0.821215312 | Transport |
| Q8TF05-2 | PPP4R1 | Serine/threonine-protein phosphatase 4 regulatory subunit 1 | -0.213722865 | -0.051965714 | Cellular metabolism |
| Q8TF42 | UBASH3B | Ubiquitin-associated and SH3 domain-containing protein B | 0.410788218 | -0.587154388 | Other |
| Q8TF72 | SHROOM3 | Protein Shroom3 | 1.539539973 | -0.729464849 | Other |
| Q8WTS1 | ABHD5 | 1-acylglycerol-3-phosphate O-acyltransferase ABHD5 | 0.259691238 | 1.985891342 | Lipid metabolism |
| Q8WTS6 | SETD7 | Histone-lysine N-methyltransferase SETD7 | -0.354388555 | -2.672465007 | Cellular metabolism |
| Q8WU10 | PYROXD1 | Pyridine nucleotide-disulfide oxidoreductase domain-containing protein 1 | -1.289749146 | 0 | Other |
| Q8WU90 | ZC3H15 | Zinc finger CCCH domain-containing protein 15 | -0.055756251 | -2.374465307 | Other |
| Q8WUA2 | PPIL4 | Peptidyl-prolyl cis-trans isomerase-like 4 | 0.030131022 | -0.606209437 | Cellular metabolism |
| Q8WUD1 | RAB2B | Ras-related protein Rab-2B | 0 | 3.750553767 | Transport |
| Q8WUF5 | PPP1R13L | RelA-associated inhibitor | -0.460732778 | 0.971825282 | Cellular metabolism |
| Q8WUI4-5 | HDAC7 | Histone deacetylase 7 | -0.634196599 | -0.109809875 | Cellular metabolism |
| Q8WUM0 | NUP133 | Nuclear pore complex protein Nup133 | -0.189584096 | -0.310898463 | Transport |
| Q8WUM4 | PDCD6IP | Programmed cell death 6-interacting protein | -0.059609095 | -0.327058156 | Transport |
| Q8WUP2 | FBLIM1 | Filamin-binding LIM protein 1 | 0.18651708 | -2.182114919 | Other |
| Q8WUX2 | CHAC2 | Putative glutathione-specific gamma-glutamylcyclotransferase 2 | -0.257614772 | -1.889738719 | Other |
| Q8WUX9 | CHMP7 | Charged multivesicular body protein 7 | -0.458109538 | -0.263618469 | Transport |
| Q8WV07 | ORAOV1 | Oral cancer-overexpressed protein 1 | -0.503510157 | 0 | Other |
| Q8WV41 | SNX33 | Sorting nexin-33 | 0.073539734 | 0 | Transport |
| Q8WV99 | ZFAND2B | AN1-type zinc finger protein 2B | -0.180845896 | -0.980642319 | Transport |
| Q8WVC0 | LEO1 | RNA polymerase-associated protein LEO1 | -0.054621379 | -0.541823069 | Cellular metabolism |
| Q8WVJ2 | NUDCD2 | NudC domain-containing protein 2 | 0.121631622 | -1.210942586 | Other |
| Q8WVT3 | TRAPPC12 | Trafficking protein particle complex subunit 12 | 0.882764816 | 0.697858175 | Transport |
| Q8WVV4-1 | POF1B | Protein POF1B | -0.940034866 | 1.26556015 | Other |
| Q8WVY7 | UBLCP1 | Ubiquitin-like domain-containing CTD phosphatase 1 | -0.094093323 | -0.397332509 | Other |
| Q8WW01 | TSEN15 | tRNA-splicing endonuclease subunit Sen15 | 0.123904546 | 0 | Cellular metabolism |
| Q8WW12 | PCNP | PEST proteolytic signal-containing nuclear protein | -0.213209152 | -0.423282623 | Cellular metabolism |
| Q8WW59 | SPRYD4 | SPRY domain-containing protein 4 | 0.025457382 | 0.330612183 | Other |
| Q8WWH5 | TRUB1 | Probable tRNA pseudouridine synthase 1 | -0.12214152 | -3.785128276 | Cellular metabolism |
| Q8WWM7-6 | ATXN2L | Ataxin-2-like protein | 0.026402156 | -0.823467255 | Other |
| Q8WWY3 | PRPF31 | U4/U6 small nuclear ribonucleoprotein Prp31 | -0.16873614 | 0.807985942 | Cellular metabolism |
| Q8WX93-3 | PALLD | Palladin | 0.052656809 | -1.215031306 | Other |

| | | | | | |
|---|---|---|---|---|---|
| Q8WXA9-2 | SREK1 | Splicing regulatory glutamine/lysine-rich protein 1 | 0.52595075 | -0.998654683 | Cellular metabolism |
| Q8WXD5 | GEMIN6 | Gem-associated protein 6 | -0.225388209 | 0 | Cellular metabolism |
| Q8WXF1 | PSPC1 | Paraspeckle component 1 | 1.012792587 | 0.309191386 | Cellular metabolism |
| Q8WXX5 | DNAJC9 | DnaJ homolog subfamily C member 9 | -0.581033071 | 0.253587087 | Other |
| Q8WZ82 | OVCA2 | Ovarian cancer-associated gene 2 protein | -0.68164444 | -0.112284978 | Other |
| Q8WZA0-2 | LZIC | Protein LZIC | -0.538223902 | -1.078289032 | Other |
| Q8WZA9 | IRGQ | Immunity-related GTPase family Q protein | 0.16284434 | -0.748820623 | Other |
| Q92484-2 | SMPDL3A | Acid sphingomyelinase-like phosphodiesterase 3a | 0.318390528 | 0 | Lipid metabolism |
| Q92522 | H1FX | Histone H1x | 0.320858002 | 1.811506271 | Other |
| Q92541 | RTF1 | RNA polymerase-associated protein RTF1 homolog | -0.192431768 | -0.596595764 | Cellular metabolism |
| Q92542-2 | NCSTN | Nicastrin | -0.696231842 | 0.749863307 | Cellular metabolism |
| Q92551 | IP6K1 | Inositol hexakisphosphate kinase 1 | -0.193721771 | -0.287668228 | Lipid metabolism |
| Q92572 | AP3S1 | AP-3 complex subunit sigma-1 | -0.048521042 | -0.004770279 | Transport |
| Q92597 | NDRG1 | Protein NDRG1 | -0.261517207 | -0.517830531 | Other |
| Q92598 | HSPH1 | Heat shock protein 105 kDa | -0.213486989 | -0.90961647 | Transport |
| Q92600 | RQCD1 | Cell differentiation protein RCD1 homolog | 0.189008713 | 0 | Cellular metabolism |
| Q92609 | TBC1D5 | TBC1 domain family member 5 | 0 | 1.150787354 | Transport |
| Q92615 | LARP4B | La-related protein 4B | -0.173039754 | -0.14401118 | Other |
| Q92616 | GCN1L1 | Translational activator GCN1 | 0.107304891 | 0.230766932 | Cellular metabolism |
| Q92620 | DHX38 | Pre-mRNA-splicing factor ATP-dependent RNA helicase PRP16 | -0.61060969 | -1.004034042 | Transport |
| Q92621 | NUP205 | Nuclear pore complex protein Nup205 | -0.84698232 | 0.118634542 | Transport |
| Q92636 | NSMAF | Protein FAN | -0.23170344 | 0 | Lipid metabolism |
| Q92688-2 | ANP32B | Acidic leucine-rich nuclear phosphoprotein 32 family member B | 0.293064117 | -0.482508977 | Other |
| Q92696 | RABGGTA | Geranylgeranyl transferase type-2 subunit alpha | -0.066179911 | -0.309511185 | Cellular metabolism |
| Q92733 | PRCC | Proline-rich protein PRCC | 0.171620687 | -0.153987249 | Other |
| Q92747 | ARPC1A | Actin-related protein 2/3 complex subunit 1A | 0.124247869 | 0 | Immune response |
| Q92769-3 | HDAC2 | Histone deacetylase 2 | -0.100770315 | 0.102551142 | Cellular metabolism |
| Q92783-2 | STAM | Signal transducing adapter molecule 1 | 0.010113398 | -0.932028453 | Transport |
| Q92804-2 | TAF15 | TATA-binding protein-associated factor 2N | 0.188774109 | -0.442087173 | Other |
| Q92817 | EVPL | Envoplakin | -0.593829473 | -0.200353622 | Cellular metabolism |
| Q92820 | GGH | Gamma-glutamyl hydrolase | -0.040995916 | -0.425127029 | Cellular metabolism |
| Q92845 | KIFAP3 | Kinesin-associated protein 3 | 0.228512446 | -0.735874812 | Transport |
| Q92871 | PMM1 | Phosphomannomutase 1 | -0.314939499 | -1.502955755 | Cellular metabolism |
| Q92876 | KLK6 | Kallikrein-6 | 0 | -0.044438044 | Cellular metabolism |

| | | | | | |
|---|---|---|---|---|---|
| Q92878 | RAD50 | DNA repair protein RAD50 | 0.244858424 | 0.642639796 | Cellular metabolism |
| Q92882 | OSTF1 | Osteoclast-stimulating factor 1 | -0.220633189 | -0.321974436 | Other |
| Q92889 | ERCC4 | DNA repair endonuclease XPF | -1.251731873 | 0 | Cellular metabolism |
| Q92890 | UFD1L | Ubiquitin fusion degradation protein 1 homolog | -0.03285408 | -0.183970769 | Cellular metabolism |
| Q92896 | GLG1 | Golgi apparatus protein 1 | -1.364760081 | 0.714330037 | Other |
| Q92900-2 | UPF1 | Regulator of nonsense transcripts 1 | 0.916573842 | 0.874943415 | Transport |
| Q92905 | COPS5 | COP9 signalosome complex subunit 5 | 0.001332601 | 0.057336171 | Cellular metabolism |
| Q92917 | GPKOW | G patch domain and KOW motifs-containing protein | 0.021684647 | -1.113020579 | Other |
| Q92925-3 | SMARCD2 | SWI/SNF-related matrix-associated actin-dependent regulator of chromatin subfamily D member 2 | 0 | -1.601390203 | Cellular metabolism |
| Q92934 | BAD | Bcl2-associated agonist of cell death | 0.373411814 | 1.143625895 | Immune response |
| Q92968 | PEX13 | Peroxisomal membrane protein PEX13 | 0 | 2.459808985 | Lipid metabolism |
| Q92973-2 | TNPO1 | Transportin-1 | -0.20516332 | -0.246765773 | Transport |
| Q92990 | GLMN | Glomulin | 0.050671895 | -0.440227509 | Other |
| Q93008 | USP9X | Probable ubiquitin carboxyl-terminal hydrolase FAF-X | -0.121195475 | -1.073818843 | Cellular metabolism |
| Q93034 | CUL5 | Cullin-5 | -0.105353038 | -2.224002202 | Transport |
| Q93050-1 | ATP6V0A1 | V-type proton ATPase 116 kDa subunit a isoform 1 | 0 | -2.275386175 | Transport |
| Q93052 | LPP | Lipoma-preferred partner | -0.211760839 | -1.926165899 | Other |
| Q93100-4 | PHKB | Phosphorylase b kinase regulatory subunit beta | 0.147095362 | 0 | Cellular metabolism |
| Q969E4 | TCEAL3 | Transcription elongation factor A protein-like 3 | -0.232346217 | 0.382945379 | Cellular metabolism |
| Q969E8 | TSR2 | Pre-rRNA-processing protein TSR2 homolog | -1.213881175 | -2.679185867 | Cellular metabolism |
| Q969F1 | GTF3C6 | General transcription factor 3C polypeptide 6 | 0.370461146 | 0 | Cellular metabolism |
| Q969G6 | RFK | Riboflavin kinase | -0.580756505 | 0 | Cellular metabolism |
| Q969H8 | MYDGF | Myeloid-derived growth factor | 0.160569509 | -0.525231043 | Cellular metabolism |
| Q969K3 | RNF34 | E3 ubiquitin-protein ligase RNF34 | -0.508652369 | -2.395719528 | Transport |
| Q969S3 | ZNF622 | Zinc finger protein 622 | 0.839811961 | 0.814078649 | Other |
| Q969Z0-2 | TBRG4 | Protein TBRG4 | -0.264713287 | -0.23840332 | Cellular metabolism |
| Q96A33-2 | CCDC47 | Coiled-coil domain-containing protein 47 | 1.407904943 | 1.231653849 | Other |
| Q96A49 | SYAP1 | Synapse-associated protein 1 | -0.113702138 | -0.592178981 | Other |
| Q96A65 | EXOC4 | Exocyst complex component 4 | 0 | -1.651637395 | Transport |
| Q96AB6 | NTAN1 | Protein N-terminal asparagine amidohydrolase | -0.269199371 | -0.73938942 | Other |
| Q96AC1 | FERMT2 | Fermitin family homolog 2 | -0.294873556 | -0.87505722 | Other |
| Q96AD5 | PNPLA2 | Patatin-like phospholipase domain-containing protein 2 | -0.499617259 | 1.487378438 | Lipid metabolism |
| Q96AG4 | LRRC59 | Leucine-rich repeat-containing protein 59 | 0.667291641 | 0.874684652 | Other |
| Q96AT1 | KIAA1143 | Uncharacterized protein KIAA1143 | 0.079057693 | 0.407859166 | Other |
| Q96AT9 | RPE | Ribulose-phosphate 3-epimerase | 0.017737707 | -1.391899745 | Cellular metabolism |

| | | | | | |
|---|---|---|---|---|---|
| Q96AY3 | FKBP10 | Peptidyl-prolyl cis-trans isomerase FKBP10 | -0.062657038 | -1.168251673 | Cellular metabolism |
| Q96B23-2 | C18orf25 | Uncharacterized protein C18orf25 | -0.031613668 | -1.193969727 | Other |
| Q96B26 | EXOSC8 | Exosome complex component RRP43 | 0.066114426 | 0 | Cellular metabolism |
| Q96B36 | AKT1S1 | Proline-rich AKT1 substrate 1 | -0.063254674 | -0.584533691 | Immune response |
| Q96B96-2 | TMEM159 | Promethin | 0.350661596 | 1.833368937 | Other |
| Q96BJ3 | AIDA | Axin interactor, dorsalization-associated protein | -0.415887833 | -2.403112411 | Other |
| Q96BM9 | ARL8A | ADP-ribosylation factor-like protein 8A | 0 | 1.933207194 | Other |
| Q96BN8 | OTULIN | Ubiquitin thioesterase otulin | 0.35080719 | 0.345564524 | Immune response |
| Q96BT7 | ALKBH8 | Alkylated DNA repair protein alkB homolog 8 | 0.212696075 | 0 | Cellular metabolism |
| Q96BX8 | MOB3A | MOB kinase activator 3A | 0.031604131 | 0 | Other |
| Q96BY7 | ATG2B | Autophagy-related protein 2 homolog B | 0.017588298 | 0.266742706 | Cellular metabolism |
| Q96C19 | EFHD2 | EF-hand domain-containing protein D2 | -0.0090669 | -0.400121689 | Other |
| Q96C23 | GALM | Aldose 1-epimerase | -0.158416748 | -1.534041087 | Cellular metabolism |
| Q96C86 | DCPS | m7GpppX diphosphatase | 0.020217896 | -0.593211492 | Cellular metabolism |
| Q96C90 | PPP1R14B | Protein phosphatase 1 regulatory subunit 14B | -0.714632034 | -0.117403666 | Immune response |
| Q96CN7 | ISOC1 | Isochorismatase domain-containing protein 1 | -0.118832906 | -1.0385348 | Other |
| Q96CN9 | GCC1 | GRIP and coiled-coil domain-containing protein 1 | -1.080363592 | 0 | Transport |
| Q96CT7 | CCDC124 | Coiled-coil domain-containing protein 124 | -0.133464177 | 0.027653376 | Other |
| Q96CV9 | OPTN | Optineurin | -0.166223526 | -0.312992732 | Transport |
| Q96CW6 | SLC7A6OS | Probable RNA polymerase II nuclear localization protein SLC7A6OS | 0.390289307 | 0.084451675 | Transport |
| Q96CX2 | KCTD12 | BTB/POZ domain-containing protein KCTD12 | -0.006587346 | 0 | Other |
| Q96D09 | GPRASP2 | G-protein coupled receptor-associated sorting protein 2 | -0.133727392 | 0 | Other |
| Q96D15 | RCN3 | Reticulocalbin-3 | 0.190960566 | 0 | Other |
| Q96D71-3 | REPS1 | RalBP1-associated Eps domain-containing protein 1 | -0.26526769 | -1.531572978 | Transport |
| Q96DF8 | DGCR14 | Protein DGCR14 | 0.152482986 | 0 | Cellular metabolism |
| Q96DG6 | CMBL | Carboxymethylenebutenolidase homolog | -0.035611471 | -0.392596563 | Other |
| Q96DI7 | SNRNP40 | U5 small nuclear ribonucleoprotein 40 kDa protein | 0.339531581 | 0.187764486 | Cellular metabolism |
| Q96E22 | NUS1 | Dehydrodolichyl diphosphate syntase complex subunit NUS1 | 2.221925735 | 5.47211647 | Transport |
| Q96EA4 | SPDL1 | Protein Spindly | -0.384395599 | 1.332979202 | Other |
| Q96EB6 | SIRT1 | NAD-dependent protein deacetylase sirtuin-1 | -0.252871831 | -1.478459676 | Lipid metabolism |
| Q96EE3 | SEH1L | Nucleoporin SEH1 | -0.47824351 | 0.290091832 | Transport |
| Q96EI5 | TCEAL4 | Transcription elongation factor A protein-like 4 | -0.615339279 | -2.215240479 | Cellular metabolism |
| Q96EK6 | GNPNAT1 | Glucosamine 6-phosphate N-acetyltransferase | 0.165814718 | 0 | Cellular metabolism |
| Q96EK9 | KTI12 | Protein KTI12 homolog | 0.011084874 | 0.007718404 | Other |
| Q96EM0 | L3HYPDH | Trans-3-hydroxy-L-proline dehydratase | -0.445412954 | -0.922880809 | Other |

| | | | | | |
|---|---|---|---|---|---|
| Q96EQ0 | SGTB | Small glutamine-rich tetratricopeptide repeat-containing protein beta | 0.078780492 | -1.09597079 | Other |
| Q96ER3 | SAAL1 | Protein SAAL1 | -0.350889842 | -1.03250885 | Other |
| Q96EY1-3 | DNAJA3 | DnaJ homolog subfamily A member 3, mitochondrial | 0.225337982 | 0.930058161 | Cellular metabolism |
| Q96EY8 | MMAB | Cob(I)yrinic acid a,c-diamide adenosyltransferase, mitochondrial | 0 | 0.55871582 | Cellular metabolism |
| Q96F24 | NRBF2 | Nuclear receptor-binding factor 2 | -0.064641317 | 0 | Cellular metabolism |
| Q96F63 | CCDC97 | Coiled-coil domain-containing protein 97 | 0.269765854 | 0 | Other |
| Q96F85 | CNRIP1 | CB1 cannabinoid receptor-interacting protein 1 | -0.001440048 | -0.884109497 | Other |
| Q96F86 | EDC3 | Enhancer of mRNA-decapping protein 3 | -0.804342906 | 0 | Cellular metabolism |
| Q96FN4 | CPNE2 | Copine-2 | 0.02100118 | -0.784538269 | Other |
| Q96FQ6 | S100A16 | Protein S100-A16 | -1.020524979 | -0.218528112 | Other |
| Q96FX7 | TRMT61A | tRNA (adenine(58)-N(1))-methyltransferase catalytic subunit TRMT61A | 0.504521052 | -0.182312012 | Cellular metabolism |
| Q96G03 | PGM2 | Phosphoglucomutase-2 | -0.093883514 | -0.380129496 | Cellular metabolism |
| Q96G28 | CFAP36 | Cilia- and flagella-associated protein 36 | -0.529681524 | 0.270186742 | Other |
| Q96G46 | DUS3L | tRNA-dihydrouridine(47) synthase [NAD(P)(+)]-like | -0.203509649 | -1.199573517 | Other |
| Q96G61 | NUDT11 | Diphosphoinositol polyphosphate phosphohydrolase 3-beta | -0.395258586 | 0 | Other |
| Q96GA3 | LTV1 | Protein LTV1 homolog | 0.025754293 | 0 | Other |
| Q96GA7 | SDSL | Serine dehydratase-like | -0.122991562 | 0 | Cellular metabolism |
| Q96GD0 | PDXP | Pyridoxal phosphate phosphatase | 0.006692886 | -1.049411138 | Cellular metabolism |
| Q96GG9 | DCUN1D1 | DCN1-like protein 1 | 0.245681763 | -0.547425588 | Other |
| Q96GK7 | FAHD2A | Fumarylacetoacetate hydrolase domain-containing protein 2A | 0.037613551 | -0.652793884 | Other |
| Q96GM8 | TOE1 | Target of EGR1 protein 1 | 0.424373627 | 2.161059697 | Other |
| Q96GS4 | C17orf59 | Uncharacterized protein C17orf59 | -0.107477188 | 1.205452601 | Other |
| Q96GX5-2 | MASTL | Serine/threonine-protein kinase greatwall | -0.857777913 | 0.415118535 | Other |
| Q96GX9 | APIP | Methylthioribulose-1-phosphate dehydratase | 0.182699203 | -0.012952805 | Cellular metabolism |
| Q96HA8 | WDYHV1 | Protein N-terminal glutamine amidohydrolase | -0.569648743 | -1.306355794 | Cellular metabolism |
| Q96HC4 | PDLIM5 | PDZ and LIM domain protein 5 | -0.047971725 | -0.349781672 | Other |
| Q96HE7 | ERO1L | ERO1-like protein alpha | 0.130671819 | -0.124755224 | Transport |
| Q96HE9 | PRR11 | Proline-rich protein 11 | -0.060407639 | 2.399184545 | Cellular metabolism |
| Q96HR8 | NAF1 | H/ACA ribonucleoprotein complex non-core subunit NAF1 | -0.075910568 | -1.293063482 | Cellular metabolism |
| Q96HS1 | PGAM5 | Serine/threonine-protein phosphatase PGAM5, mitochondrial | 0 | 0.225767771 | Cellular metabolism |
| Q96I24 | FUBP3 | Far upstream element-binding protein 3 | -0.861796697 | -1.39504687 | Cellular metabolism |
| Q96I25 | RBM17 | Splicing factor 45 | 0.949820836 | -0.459185918 | Cellular metabolism |
| Q96I99 | SUCLG2 | Succinyl-CoA ligase [GDP-forming] subunit beta, mitochondrial | -0.368018468 | -0.432738622 | Cellular metabolism |

| | | | | | |
|---|---|---|---|---|---|
| Q96IF1 | AJUBA | LIM domain-containing protein ajuba | -0.529404958 | 0 | Lipid metabolism |
| Q96IJ6 | GMPPA | Mannose-1-phosphate guanyltransferase alpha | -0.12535731 | -0.639799754 | Cellular metabolism |
| Q96IU4 | ABHD14B | Alpha/beta hydrolase domain-containing protein 14B | -0.200797399 | -0.512556712 | Other |
| Q96IV0 | NGLY1 | Peptide-N(4)-(N-acetyl-beta-glucosaminyl)asparagine amidase | -0.558354696 | 0.357577642 | Cellular metabolism |
| Q96IZ0 | PAWR | PRKC apoptosis WT1 regulator protein | -0.926894506 | -0.683322906 | Cellular metabolism |
| Q96J01 | THOC3 | THO complex subunit 3 | -0.378755569 | -0.832919439 | Transport |
| Q96J92-3 | WNK4 | Serine/threonine-protein kinase WNK4 | -0.081616084 | 0 | Transport |
| Q96JB2 | COG3 | Conserved oligomeric Golgi complex subunit 3 | -0.283508937 | 0 | Transport |
| Q96JB5 | CDK5RAP3 | CDK5 regulatory subunit-associated protein 3 | 0 | 0.609956741 | Cellular metabolism |
| Q96JH7 | VCPIP1 | Deubiquitinating protein VCIP135 | -0.049044291 | -0.206615448 | Cellular metabolism |
| Q96JJ7 | TMX3 | Protein disulfide-isomerase TMX3 | 0 | 1.771596909 | Cellular metabolism |
| Q96JY6 | PDLIM2 | PDZ and LIM domain protein 2 | -0.666377385 | 0 | Other |
| Q96K17 | BTF3L4 | Transcription factor BTF3 homolog 4 | -0.169496536 | -0.593571981 | Other |
| Q96K76-4 | USP47 | Ubiquitin carboxyl-terminal hydrolase 47 | -0.191340129 | -0.728280385 | Cellular metabolism |
| Q96KB5 | PBK | Lymphokine-activated killer T-cell-originated protein kinase | -0.143176397 | -0.938041687 | Other |
| Q96KP4 | CNDP2 | Cytosolic non-specific dipeptidase | -0.264030457 | -0.71455574 | Cellular metabolism |
| Q96L92-3 | SNX27 | Sorting nexin-27 | -0.656270345 | -0.016639074 | Transport |
| Q96L93-2 | KIF16B | Kinesin-like protein KIF16B | 0.034330368 | 2.560387929 | Transport |
| Q96LD4 | TRIM47 | Tripartite motif-containing protein 47 | -0.077634811 | 0 | Other |
| Q96M27 | PRRC1 | Protein PRRC1 | -0.346794128 | 0.187643051 | Other |
| Q96MW1 | CCDC43 | Coiled-coil domain-containing protein 43 | -0.31046168 | -0.497227351 | Other |
| Q96MX6-2 | WDR92 | WD repeat-containing protein 92 | 0.751808802 | -1.822812398 | Cellular metabolism |
| Q96N66 | MBOAT7 | Lysophospholipid acyltransferase 7 | 0.890110652 | 3.347091039 | Lipid metabolism |
| Q96N67 | DOCK7 | Dedicator of cytokinesis protein 7 | 0.431365331 | -0.008476257 | Other |
| Q96NB3 | ZNF830 | Zinc finger protein 830 | 1.316035589 | 0 | Cellular metabolism |
| Q96NY7-2 | CLIC6 | Chloride intracellular channel protein 6 | -0.089829763 | 0 | Transport |
| Q96P16 | RPRD1A | Regulation of nuclear pre-mRNA domain-containing protein 1A | 0.007194519 | -2.570228577 | Cellular metabolism |
| Q96P47 | AGAP3 | Arf-GAP with GTPase, ANK repeat and PH domain-containing protein 3 | -0.009284973 | 0 | Transport |
| Q96P48-3 | ARAP1 | Arf-GAP with Rho-GAP domain, ANK repeat and PH domain-containing protein 1 | -0.356602987 | -0.044917425 | Other |
| Q96P63-2 | SERPINB12 | Serpin B12 | 0.425605774 | 1.248601913 | Other |
| Q96P70 | IPO9 | Importin-9 | -0.044374466 | -0.116845449 | Transport |
| Q96PK6 | RBM14 | RNA-binding protein 14 | 0.459086736 | 0.390708288 | Cellular metabolism |
| Q96PY5-3 | FMNL2 | Formin-like protein 2 | 1.162825267 | 0 | Other |
| Q96PZ0 | PUS7 | Pseudouridylate synthase 7 homolog | -0.075843175 | -0.49607722 | Cellular metabolism |

| | | | | | |
|---|---|---|---|---|---|
| Q96Q11-2 | TRNT1 | CCA tRNA nucleotidyltransferase 1, mitochondrial | -0.041224162 | -0.130965551 | Transport |
| Q96QA5 | GSDMA | Gasdermin-A | 0.394312541 | 0.729115804 | Other |
| Q96QB1-4 | DLC1 | Rho GTPase-activating protein 7 | -0.355380376 | 0 | Other |
| Q96QC0 | PPP1R10 | Serine/threonine-protein phosphatase 1 regulatory subunit 10 | 0.369937261 | 0 | Transport |
| Q96QK1 | VPS35 | Vacuolar protein sorting-associated protein 35 | 0.035415649 | -0.724515915 | Transport |
| Q96QR8 | PURB | Transcriptional activator protein Pur-beta | 0.826913198 | 0.373528163 | Cellular metabolism |
| Q96QU8 | XPO6 | Exportin-6 | -0.426317215 | -0.441469193 | Transport |
| Q96R05 | RBP7 | Retinoid-binding protein 7 | -0.332247416 | -1.718304316 | Other |
| Q96R06 | SPAG5 | Sperm-associated antigen 5 | -1.030115128 | -0.738419851 | Other |
| Q96RG2 | PASK | PAS domain-containing serine/threonine-protein kinase | -0.175434113 | 0 | Cellular metabolism |
| Q96RP9 | GFM1 | Elongation factor G, mitochondrial | -0.194239934 | 0.656806946 | Cellular metabolism |
| Q96RR4-2 | CAMKK2 | Calcium/calmodulin-dependent protein kinase kinase 2 | -0.099273046 | 0 | Cellular metabolism |
| Q96RS0 | TGS1 | Trimethylguanosine synthase | 0.274700801 | -0.591635386 | Lipid metabolism |
| Q96RS6 | NUDCD1 | NudC domain-containing protein 1 | -0.065446854 | 0.000710169 | Other |
| Q96RU2-2 | USP28 | Ubiquitin carboxyl-terminal hydrolase 28 | -0.49225235 | 0.738470078 | Cellular metabolism |
| Q96S44 | TP53RK | TP53-regulating kinase | 0.008180618 | -0.411089579 | Cellular metabolism |
| Q96S55 | WRNIP1 | ATPase WRNIP1 | 0.016851425 | -0.512748082 | Cellular metabolism |
| Q96S59 | RANBP9 | Ran-binding protein 9 | 0.058677038 | -1.721682866 | Other |
| Q96S82 | UBL7 | Ubiquitin-like protein 7 | 0.218681971 | 0 | Other |
| Q96SB3 | PPP1R9B | Neurabin-2 | 0.37834994 | 1.477470398 | Cellular metabolism |
| Q96ST2-2 | IWS1 | Protein IWS1 homolog | 0.045259476 | -1.005500158 | Transport |
| Q96SU4-7 | OSBPL9 | Oxysterol-binding protein-related protein 9 | -0.110914866 | -1.908543905 | Transport |
| Q96SZ5 | ADO | 2-aminoethanethiol dioxygenase | -0.085618973 | 0.202679952 | Other |
| Q96T23 | RSF1 | Remodeling and spacing factor 1 | -0.180928548 | -0.64040184 | Cellular metabolism |
| Q96T51 | RUFY1 | RUN and FYVE domain-containing protein 1 | -0.474875768 | -1.050958633 | Transport |
| Q96T76 | MMS19 | MMS19 nucleotide excision repair protein homolog | -0.326424917 | -0.180870056 | Cellular metabolism |
| Q96TA1-2 | FAM129B | Niban-like protein 1 | 0.122962952 | -0.203097026 | Other |
| Q96TC7 | RMDN3 | Regulator of microtubule dynamics protein 3 | 0 | 1.28320694 | Other |
| Q99426 | TBCB | Tubulin-folding cofactor B | -0.136526108 | -0.444965998 | Cellular metabolism |
| Q99436 | PSMB7 | Proteasome subunit beta type-7 | 0.028160095 | 0.12316068 | Immune response |
| Q99447 | PCYT2 | Ethanolamine-phosphate cytidylyltransferase | -0.07396253 | -1.645298004 | Lipid metabolism |
| Q99459 | CDC5L | Cell division cycle 5-like protein | 0.98903211 | -2.271975199 | Cellular metabolism |
| Q99460 | PSMD1 | 26S proteasome non-ATPase regulatory subunit 1 | -0.118083318 | -0.791567485 | Immune response |
| Q99470 | SDF2 | Stromal cell-derived factor 2 | 1.014888763 | 0 | Cellular metabolism |

| | | | | | |
|---|---|---|---|---|---|
| Q99471 | PFDN5 | Prefoldin subunit 5 | -0.870011012 | -1.490058899 | Cellular metabolism |
| Q99497 | PARK7 | Protein deglycase DJ-1 | 0.036959966 | -0.231964747 | Transport |
| Q99519 | NEU1 | Sialidase-1 | -2.214363098 | -1.69394811 | Lipid metabolism |
| Q99536 | VAT1 | Synaptic vesicle membrane protein VAT-1 homolog | 0.05352656 | 1.749153137 | Other |
| Q99541 | PLIN2 | Perilipin-2 | -0.178200404 | 2.094742457 | Lipid metabolism |
| Q99543 | DNAJC2 | DnaJ homolog subfamily C member 2 | 0.218887965 | -0.708827972 | Cellular metabolism |
| Q99567 | NUP88 | Nuclear pore complex protein Nup88 | -0.358309428 | 0.30961736 | Transport |
| Q99570 | PIK3R4 | Phosphoinositide 3-kinase regulatory subunit 4 | -0.065973282 | -0.713820775 | Immune response |
| Q99575 | POP1 | Ribonucleases P/MRP protein subunit POP1 | 0.601601283 | -1.018597285 | Cellular metabolism |
| Q99584 | S100A13 | Protein S100-A13 | -0.212250392 | 0.128852844 | Transport |
| Q99598 | TSNAX | Translin-associated protein X | 0.261393865 | -0.501533508 | Transport |
| Q99611 | SEPHS2 | Selenide, water dikinase 2 | -0.701493581 | 0 | Cellular metabolism |
| Q99613 | EIF3C | Eukaryotic translation initiation factor 3 subunit C | 0.116156896 | -0.643508911 | Cellular metabolism |
| Q99614 | TTC1 | Tetratricopeptide repeat protein 1 | -0.16135025 | -1.505585353 | Cellular metabolism |
| Q99623 | PHB2 | Prohibitin-2 | 0.054496765 | 1.199457169 | Cellular metabolism |
| Q99638 | RAD9A | Cell cycle checkpoint control protein RAD9A | 0.063675563 | 0 | Cellular metabolism |
| Q99640-3 | PKMYT1 | Membrane-associated tyrosine- and threonine-specific cdc2-inhibitory kinase | 1.43244044 | 4.041166306 | Other |
| Q99653 | CHP1 | Calcineurin B homologous protein 1 | 0.151138941 | 1.829808553 | Transport |
| Q99661 | KIF2C | Kinesin-like protein KIF2C | -0.917292277 | -0.012839635 | Other |
| Q99704 | DOK1 | Docking protein 1 | -0.437060674 | 0 | Other |
| Q99714 | HSD17B10 | 3-hydroxyacyl-CoA dehydrogenase type-2 | -0.042001088 | 0.682643255 | Lipid metabolism |
| Q99733 | NAP1L4 | Nucleosome assembly protein 1-like 4 | 0.028711955 | -0.166040421 | Other |
| Q99747 | NAPG | Gamma-soluble NSF attachment protein | -0.457568487 | 0.968835195 | Transport |
| Q99829 | CPNE1 | Copine-1 | -0.18608284 | -0.739764531 | Lipid metabolism |
| Q99832 | CCT7 | T-complex protein 1 subunit eta | 0.0849425 | 0.300231934 | Cellular metabolism |
| Q99836-5 | MYD88 | Myeloid differentiation primary response protein MyD88 | 0.598958333 | -0.525411606 | Immune response |
| Q99873-3 | PRMT1 | Protein arginine N-methyltransferase 1 | 0.099930445 | -0.091385523 | Cellular metabolism |
| Q99961 | SH3GL1 | Endophilin-A2 | 0.167854945 | -1.199625651 | Transport |
| Q9BPW8 | NIPSNAP1 | Protein NipSnap homolog 1 | 0.096396764 | -0.873682658 | Other |
| Q9BPX3 | NCAPG | Condensin complex subunit 3 | -0.029454549 | 0.478039424 | Cellular metabolism |
| Q9BPX5 | ARPC5L | Actin-related protein 2/3 complex subunit 5-like protein | 0.086118062 | 0.916178385 | Other |
| Q9BPX7 | C7orf25 | UPF0415 protein C7orf25 | 0.00200208 | -0.077266693 | Other |
| Q9BQ52 | ELAC2 | Zinc phosphodiesterase ELAC protein 2 | -0.014190038 | -1.563057582 | Cellular metabolism |

| | | | | | |
|---|---|---|---|---|---|
| Q9BQ61 | C19orf43 | Uncharacterized protein C19orf43 | 0.100599925 | 0.114730199 | Other |
| Q9BQ67 | GRWD1 | Glutamate-rich WD repeat-containing protein 1 | -0.149024963 | -0.142233531 | Other |
| Q9BQ70 | TCF25 | Transcription factor 25 | -1.050350825 | -1.046593984 | Cellular metabolism |
| Q9BQA1 | WDR77 | Methylosome protein 50 | 0.096364339 | -0.361239115 | Cellular metabolism |
| Q9BQC3 | DPH2 | Diphthamide biosynthesis protein 2 | 0.024819692 | -0.547119141 | Cellular metabolism |
| Q9BQE3 | TUBA1C | Tubulin alpha-1C chain | 0.12785848 | 1.969779968 | Transport |
| Q9BQG0 | MYBBP1A | Myb-binding protein 1A | 0.128880819 | 0.905675888 | Transport |
| Q9BQQ3 | GORASP1 | Golgi reassembly-stacking protein 1 | -0.851348877 | -0.832578023 | Transport |
| Q9BQS8 | FYCO1 | FYVE and coiled-coil domain-containing protein 1 | -0.177471161 | -1.07253774 | Transport |
| Q9BR61 | ACBD6 | Acyl-CoA-binding domain-containing protein 6 | -0.289536158 | -1.771263758 | Other |
| Q9BR76 | CORO1B | Coronin-1B | -0.030459086 | -0.546478271 | Other |
| Q9BRA2 | TXNDC17 | Thioredoxin domain-containing protein 17 | -0.159203211 | -0.076100032 | Other |
| Q9BRF8 | CPPED1 | Serine/threonine-protein phosphatase CPPED1 | -0.218507131 | -1.186418533 | Other |
| Q9BRG1 | VPS25 | Vacuolar protein-sorting-associated protein 25 | -0.250150681 | -1.500248591 | Transport |
| Q9BRP1 | PDCD2L | Programmed cell death protein 2-like | 0.116406759 | -0.38839976 | Other |
| Q9BRP4 | PAAF1 | Proteasomal ATPase-associated factor 1 | -0.121408463 | -0.223616282 | Other |
| Q9BRP8-2 | WIBG | Partner of Y14 and mago | 0.238256454 | 1.190931956 | Cellular metabolism |
| Q9BRS2 | RIOK1 | Serine/threonine-protein kinase RIO1 | 0.249741872 | -0.889470418 | Other |
| Q9BRT3 | MIEN1 | Migration and invasion enhancer 1 | -0.116728465 | -0.490997314 | Other |
| Q9BRT6 | LLPH | Protein LLP homolog | 0.948188782 | -0.885152181 | Other |
| Q9BRX2 | PELO | Protein pelota homolog | 0.166041056 | 0.232315699 | Cellular metabolism |
| Q9BRX5 | GINS3 | DNA replication complex GINS protein PSF3 | 0.019619624 | -0.920921326 | Cellular metabolism |
| Q9BRX8-2 | FAM213A | Redox-regulatory protein FAM213A | 0 | 1.695607503 | Other |
| Q9BS26 | ERP44 | Endoplasmic reticulum resident protein 44 | -0.022035599 | -0.107845942 | Cellular metabolism |
| Q9BS40 | LXN | Latexin | -0.056879679 | 0.162960052 | Other |
| Q9BSB4 | ATG101 | Autophagy-related protein 101 | -0.64940834 | 0 | Other |
| Q9BSD7 | NTPCR | Cancer-related nucleoside-triphosphatase | 0.078627904 | -0.960000992 | Other |
| Q9BSJ8-2 | ESYT1 | Extended synaptotagmin-1 | 0.710982005 | 0.798569361 | Transport |
| Q9BSL1 | UBAC1 | Ubiquitin-associated domain-containing protein 1 | -0.031714122 | -1.990659714 | Cellular metabolism |
| Q9BSY9 | DESI2 | Desumoylating isopeptidase 2 | 0.272239049 | 0 | Other |
| Q9BT09 | CNPY3 | Protein canopy homolog 3 | -0.179553986 | -2.597456614 | Immune response |
| Q9BT25-2 | HAUS8 | HAUS augmin-like complex subunit 8 | -0.487758636 | 0 | Other |
| Q9BT73 | PSMG3 | Proteasome assembly chaperone 3 | 0.164784114 | 0 | Other |
| Q9BT78 | COPS4 | COP9 signalosome complex subunit 4 | 0.336708069 | -0.051383336 | Cellular metabolism |
| Q9BTE3-2 | MCMBP | Mini-chromosome maintenance complex-binding protein | -0.266352971 | -0.706384659 | Cellular metabolism |

| | | | | | |
|---|---|---|---|---|---|
| Q9BTE7 | DCUN1D5 | DCN1-like protein 5 | 0.043878555 | -0.114760081 | Other |
| Q9BTL3 | FAM103A1 | RNMT-activating mini protein | -0.014625549 | 0 | Cellular metabolism |
| Q9BTT0 | ANP32E | Acidic leucine-rich nuclear phosphoprotein 32 family member E | 0.115022024 | -1.223526001 | Other |
| Q9BTV4 | TMEM43 | Transmembrane protein 43 | 0.632817586 | 1.509281794 | Other |
| Q9BTX7 | TTPAL | Alpha-tocopherol transfer protein-like | -0.973615646 | 0 | Other |
| Q9BTY2 | FUCA2 | Plasma alpha-L-fucosidase | 0.021796544 | -0.127545039 | Cellular metabolism |
| Q9BTY7 | HGH1 | Protein HGH1 homolog | 0.327505747 | -1.067618688 | Other |
| Q9BU02 | THTPA | Thiamine-triphosphatase | 0.195381165 | 0 | Cellular metabolism |
| Q9BU89 | DOHH | Deoxyhypusine hydroxylase | -0.086675008 | 0 | Cellular metabolism |
| Q9BUA3 | C11orf84 | Uncharacterized protein C11orf84 | 0.123998642 | 0 | Other |
| Q9BUF5 | TUBB6 | Tubulin beta-6 chain | -0.138048172 | -0.513692856 | Cellular metabolism |
| Q9BUH6 | C9orf142 | Protein PAXX | -0.741114934 | -1.416686376 | Cellular metabolism |
| Q9BUI4 | POLR3C | DNA-directed RNA polymerase III subunit RPC3 | -0.167933146 | -0.69328626 | Immune response |
| Q9BUK6-2 | MSTO1 | Protein misato homolog 1 | -0.238129298 | 0 | Other |
| Q9BUL5-4 | PHF23 | PHD finger protein 23 | 0.042682012 | -0.708960215 | Other |
| Q9BUL8 | PDCD10 | Programmed cell death protein 10 | 0.086081823 | 0.003317515 | Other |
| Q9BUQ8 | DDX23 | Probable ATP-dependent RNA helicase DDX23 | 0.805542628 | -0.583287557 | Cellular metabolism |
| Q9BUR4 | WRAP53 | Telomerase Cajal body protein 1 | -0.045377096 | 0.132935842 | Cellular metabolism |
| Q9BUT1 | BDH2 | 3-hydroxybutyrate dehydrogenase type 2 | 0.099794388 | 0 | Lipid metabolism |
| Q9BV20 | MRI1 | Methylthioribose-1-phosphate isomerase | -0.132732391 | -2.157768885 | Cellular metabolism |
| Q9BV44 | THUMPD3 | THUMP domain-containing protein 3 | -0.539090474 | -0.153430939 | Other |
| Q9BV57 | ADI1 | 1,2-dihydroxy-3-keto-5-methylthiopentene dioxygenase | -1.09616216 | -1.383988063 | Cellular metabolism |
| Q9BV73 | CEP250 | Centrosome-associated protein CEP250 | 0.004273097 | -0.11815389 | Other |
| Q9BV86 | NTMT1 | N-terminal Xaa-Pro-Lys N-methyltransferase 1 | 0.043397903 | -0.714197795 | Cellular metabolism |
| Q9BVA0 | KATNB1 | Katanin p80 WD40 repeat-containing subunit B1 | 0.195034027 | 0 | Transport |
| Q9BVA1 | TUBB2B | Tubulin beta-2B chain | 0.04511706 | 0.60189565 | Cellular metabolism |
| Q9BVG4 | PBDC1 | Protein PBDC1 | 0.032792409 | -0.91305987 | Other |
| Q9BVI4 | NOC4L | Nucleolar complex protein 4 homolog | 0.646888733 | 0 | Cellular metabolism |
| Q9BVJ7 | DUSP23 | Dual specificity protein phosphatase 23 | 0.002599716 | 0 | Other |
| Q9BVK6 | TMED9 | Transmembrane emp24 domain-containing protein 9 | 0.372895559 | 1.352249781 | Transport |
| Q9BVM2 | DPCD | Protein DPCD | -0.182841619 | 0 | Other |
| Q9BVP2-2 | GNL3 | Guanine nucleotide-binding protein-like 3 | 0.321174622 | 0.913337708 | Other |
| Q9BVS4-2 | RIOK2 | Serine/threonine-protein kinase RIO2 | 0.55779012 | 0 | Other |
| Q9BVW5 | TIPIN | TIMELESS-interacting protein | 0.404834112 | -0.152414958 | Cellular metabolism |
| Q9BW19 | KIFC1 | Kinesin-like protein KIFC1 | 0.096803665 | 0 | Other |
| Q9BW27 | NUP85 | Nuclear pore complex protein Nup85 | -0.433980942 | 0.548720042 | Transport |

| | | | | | |
|---|---|---|---|---|---|
| Q9BW61 | DDA1 | DET1- and DDB1-associated protein 1 | -0.678763072 | -0.217350006 | Other |
| Q9BW62 | KATNAL1 | Katanin p60 ATPase-containing subunit A-like 1 | -0.019131343 | 0 | Other |
| Q9BW71 | HIRIP3 | HIRA-interacting protein 3 | 0.048433304 | -1.403071086 | Other |
| Q9BW83 | IFT27 | Intraflagellar transport protein 27 homolog | -0.180988312 | -0.568508148 | Transport |
| Q9BW85 | CCDC94 | Coiled-coil domain-containing protein 94 | -0.200338364 | -0.932935715 | Other |
| Q9BW91-2 | NUDT9 | ADP-ribose pyrophosphatase, mitochondrial | -0.374176025 | -1.667863846 | Cellular metabolism |
| Q9BWD1 | ACAT2 | Acetyl-CoA acetyltransferase, cytosolic | -0.381823858 | -1.04979833 | Lipid metabolism |
| Q9BWF3 | RBM4 | RNA-binding protein 4 | 0.366716385 | -1.317302068 | Cellular metabolism |
| Q9BWH6 | RPAP1 | RNA polymerase II-associated protein 1 | -0.204924266 | 0 | Cellular metabolism |
| Q9BX40 | LSM14B | Protein LSM14 homolog B | -0.245368958 | 0 | Other |
| Q9BX84-7 | TRPM6 | Transient receptor potential cation channel subfamily M member 6 | 1.821994146 | -0.432167689 | Transport |
| Q9BXB4 | OSBPL11 | Oxysterol-binding protein-related protein 11 | -0.054005941 | -0.156476974 | Transport |
| Q9BXJ9 | NAA15 | N-alpha-acetyltransferase 15, NatA auxiliary subunit | 0.021527608 | -0.322576523 | Cellular metabolism |
| Q9BXL6-3 | CARD14 | Caspase recruitment domain-containing protein 14 | -0.237123489 | 0 | Other |
| Q9BXP5-4 | SRRT | Serrate RNA effector molecule homolog | 0.1509552 | -0.174596151 | Cellular metabolism |
| Q9BXR0 | QTRT1 | Queuine tRNA-ribosyltransferase | -0.034006755 | -0.139710108 | Cellular metabolism |
| Q9BXS5-2 | AP1M1 | AP-1 complex subunit mu-1 | -0.081219991 | 0.104788462 | Transport |
| Q9BXS6-7 | NUSAP1 | Nucleolar and spindle-associated protein 1 | -4.58515358 | 0 | Cellular metabolism |
| Q9BXW6 | OSBPL1A | Oxysterol-binding protein-related protein 1 | 0.117357254 | -1.958617528 | Lipid metabolism |
| Q9BXW9-3 | FANCD2 | Fanconi anemia group D2 protein | -0.049785614 | -0.855602264 | Cellular metabolism |
| Q9BY32 | ITPA | Inosine triphosphate pyrophosphatase | 0.244087855 | -0.980438232 | Cellular metabolism |
| Q9BY43-2 | CHMP4A | Charged multivesicular body protein 4a | -0.312629064 | -0.962673187 | Transport |
| Q9BY44-3 | EIF2A | Eukaryotic translation initiation factor 2A | 0.097517649 | -0.481527964 | Cellular metabolism |
| Q9BYB4 | GNB1L | Guanine nucleotide-binding protein subunit beta-like protein 1 | -0.438755671 | -0.8069369 | Other |
| Q9BYD2 | MRPL9 | 39S ribosomal protein L9, mitochondrial | 0 | 1.414995193 | Cellular metabolism |
| Q9BYG3 | NIFK | MKI67 FHA domain-interacting nucleolar phosphoprotein | 1.228639603 | 1.264533361 | Cellular metabolism |
| Q9BYN0 | SRXN1 | Sulfiredoxin-1 | -0.601590474 | 0 | Other |
| Q9BYT8 | NLN | Neurolysin, mitochondrial | -0.307729085 | -0.935147603 | Other |
| Q9BYX2 | TBC1D2 | TBC1 domain family member 2A | 0.374012629 | 0.157402039 | Other |
| Q9BZ23-3 | PANK2 | Pantothenate kinase 2, mitochondrial | -0.018571854 | -0.669285456 | Cellular metabolism |
| Q9BZD4 | NUF2 | Kinetochore protein Nuf2 | -0.814303716 | 0 | Other |
| Q9BZE1 | MRPL37 | 39S ribosomal protein L37, mitochondrial | 0 | 0.372119904 | Cellular metabolism |
| Q9BZE2 | PUS3 | tRNA pseudouridine(38/39) synthase | 0.049163818 | 0 | Cellular metabolism |

| | | | | | |
|---|---|---|---|---|---|
| Q9BZE4 | GTPBP4 | Nucleolar GTP-binding protein 1 | 1.032423019 | 0.539721807 | Other |
| Q9BZE9 | ASPSCR1 | Tether containing UBX domain for GLUT4 | -0.112142563 | -1.47659874 | Transport |
| Q9BZF3-4 | OSBPL6 | Oxysterol-binding protein-related protein 6 | 0.242879232 | 0 | Transport |
| Q9BZG1 | RAB34 | Ras-related protein Rab-34 | 0.497343699 | 1.632897695 | Transport |
| Q9BZH6 | WDR11 | WD repeat-containing protein 11 | 0.322779338 | 0 | Other |
| Q9BZK7 | TBL1XR1 | F-box-like/WD repeat-containing protein TBL1XR1 | 0.198936462 | -1.949572881 | Lipid metabolism |
| Q9BZL6-2 | PRKD2 | Serine/threonine-protein kinase D2 | 0.403184891 | 0 | Cellular metabolism |
| Q9BZQ8 | FAM129A | Protein Niban | -0.148742676 | -0.29315122 | Other |
| Q9BZV1 | UBXN6 | UBX domain-containing protein 6 | -0.832135518 | 0 | Other |
| Q9BZX2 | UCK2 | Uridine-cytidine kinase 2 | -0.717459997 | 0.397481283 | Cellular metabolism |
| Q9BZZ5-2 | API5 | Apoptosis inhibitor 5 | 0.137429555 | -0.2575442 | Other |
| Q9C035-2 | TRIM5 | Tripartite motif-containing protein 5 | 0.318811417 | 0 | Immune response |
| Q9C037-2 | TRIM4 | E3 ubiquitin-protein ligase TRIM4 | -0.113637924 | 0 | Immune response |
| Q9C0C2 | TNKS1BP1 | 182 kDa tankyrase-1-binding protein | -0.032639821 | -1.104617437 | Cellular metabolism |
| Q9C0C9 | UBE2O | E2/E3 hybrid ubiquitin-protein ligase UBE2O | -0.191152573 | -1.030672073 | Transport |
| Q9C0D3 | ZYG11B | Protein zyg-11 homolog B | 0.234401703 | 0 | Other |
| Q9C0E2 | XPO4 | Exportin-4 | -0.065166473 | -0.063100815 | Transport |
| Q9C0J8 | WDR33 | pre-mRNA 3 end processing protein WDR33 | 0.933509191 | -0.568865458 | Cellular metabolism |
| Q9GZL7 | WDR12 | Ribosome biogenesis protein WDR12 | 0.066753387 | -1.240673065 | Cellular metabolism |
| Q9GZN8-2 | C20orf27 | UPF0687 protein C20orf27 | -0.006467819 | -1.303514481 | Other |
| Q9GZP4 | PITHD1 | PITH domain-containing protein 1 | -0.11042277 | -2.5233078 | Other |
| Q9GZS1-1 | POLR1E | DNA-directed RNA polymerase I subunit RPA49 | -0.776647568 | 0 | Cellular metabolism |
| Q9GZT3-2 | SLIRP | SRA stem-loop-interacting RNA-binding protein, mitochondrial | 1.099318822 | -0.478396734 | Cellular metabolism |
| Q9GZT4 | SRR | Serine racemase | -0.126321157 | 0.35111618 | Cellular metabolism |
| Q9GZT8-2 | NIF3L1 | NIF3-like protein 1 | -0.362435659 | -0.023412704 | Cellular metabolism |
| Q9GZT9-2 | EGLN1 | Egl nine homolog 1 | 0.011231105 | 0 | Cellular metabolism |
| Q9GZU8 | FAM192A | Protein FAM192A | 0.0148983 | -0.433858236 | Other |
| Q9GZX9 | TWSG1 | Twisted gastrulation protein homolog 1 | 0.033102036 | 0 | Other |
| Q9GZZ8 | LACRT | Extracellular glycoprotein lacritin | 0 | 1.569657008 | Transport |
| Q9GZZ9 | UBA5 | Ubiquitin-like modifier-activating enzyme 5 | 0.011086782 | -1.78200531 | Cellular metabolism |
| Q9H008 | LHPP | Phospholysine phosphohistidine inorganic pyrophosphate phosphatase | -0.208430608 | 1.121056875 | Cellular metabolism |
| Q9H074 | PAIP1 | Polyadenylate-binding protein-interacting protein 1 | 0.016351064 | -0.240020116 | Cellular metabolism |
| Q9H078-2 | CLPB | Caseinolytic peptidase B protein homolog | 0.099669139 | 0.280754089 | Other |
| Q9H098-2 | FAM107B | Protein FAM107B | -0.656756719 | -2.849524816 | Other |
| Q9H0A0-2 | NAT10 | N-acetyltransferase 10 | 0.166169484 | 0.460400263 | Other |
| Q9H0A8 | COMMD4 | COMM domain-containing protein 4 | -0.262124379 | 0 | Cellular metabolism |

| | | | | | |
|---|---|---|---|---|---|
| Q9H0B6 | KLC2 | Kinesin light chain 2 | 0.153576533 | -0.628309886 | Transport |
| Q9H0C8 | ILKAP | Integrin-linked kinase-associated serine/threonine phosphatase 2C | 0.033409754 | 0.884636561 | Cellular metabolism |
| Q9H0D6 | XRN2 | 5-3 exoribonuclease 2 | -0.021534602 | -1.237291972 | Cellular metabolism |
| Q9H0E2 | TOLLIP | Toll-interacting protein | -0.137192408 | 0.099601746 | Immune response |
| Q9H0P0-1 | NT5C3A | Cytosolic 5-nucleotidase 3A | 0.813891729 | 1.509741465 | Cellular metabolism |
| Q9H0Q0 | FAM49A | Protein FAM49A | -1.035032908 | -0.6347332 | Other |
| Q9H0S4 | DDX47 | Probable ATP-dependent RNA helicase DDX47 | 0.1355896 | 0.18245697 | Cellular metabolism |
| Q9H0U4 | RAB1B | Ras-related protein Rab-1B | 0.081494649 | 2.008636475 | Transport |
| Q9H0V9-3 | LMAN2L | VIP36-like protein | 0.990219116 | 0 | Transport |
| Q9H0W8-2 | SMG9 | Protein SMG9 | -0.467498779 | -2.29847463 | Cellular metabolism |
| Q9H0W9-2 | C11orf54 | Ester hydrolase C11orf54 | -0.191338221 | -0.879865011 | Other |
| Q9H1A4 | ANAPC1 | Anaphase-promoting complex subunit 1 | -0.026330948 | 0 | Cellular metabolism |
| Q9H1D9 | POLR3F | DNA-directed RNA polymerase III subunit RPC6 | -0.128756205 | 1.480237961 | Immune response |
| Q9H1E3 | NUCKS1 | Nuclear ubiquitous casein and cyclin-dependent kinase substrate 1 | 0.102240245 | -0.259606043 | Other |
| Q9H1Y0 | ATG5 | Autophagy protein 5 | 0.205067317 | 0.486602147 | Immune response |
| Q9H1Z4 | WDR13 | WD repeat-containing protein 13 | 0.046889623 | 0 | Other |
| Q9H223 | EHD4 | EH domain-containing protein 4 | -0.119171143 | 0.746346156 | Transport |
| Q9H269 | VPS16 | Vacuolar protein sorting-associated protein 16 homolog | 0.00201416 | 0.26597023 | Transport |
| Q9H2C0 | GAN | Gigaxonin | -1.019429525 | -2.031813939 | Cellular metabolism |
| Q9H2G2 | SLK | STE20-like serine/threonine-protein kinase | -0.074658076 | -0.334984461 | Cellular metabolism |
| Q9H2H8 | PPIL3 | Peptidyl-prolyl cis-trans isomerase-like 3 | 0.04156812 | -1.374166489 | Cellular metabolism |
| Q9H2J4 | PDCL3 | Phosducin-like protein 3 | -0.381231944 | -0.70058314 | Cellular metabolism |
| Q9H2L5-3 | RASSF4 | Ras association domain-containing protein 4 | 0.236366272 | 0 | Other |
| Q9H2M9 | RAB3GAP2 | Rab3 GTPase-activating protein non-catalytic subunit | 0.003540039 | -0.745600382 | Transport |
| Q9H2P9 | DPH5 | Diphthine synthase | 0.089629491 | -0.093568166 | Cellular metabolism |
| Q9H2U1-3 | DHX36 | ATP-dependent RNA helicase DHX36 | 0.017742157 | -0.140078863 | Immune response |
| Q9H2U2 | PPA2 | Inorganic pyrophosphatase 2, mitochondrial | -0.190183004 | 0.052930832 | Cellular metabolism |
| Q9H307 | PNN | Pinin | 0.183539708 | -0.586941401 | Cellular metabolism |
| Q9H3G5 | CPVL | Probable serine carboxypeptidase CPVL | 0.150467555 | -0.243917465 | Other |
| Q9H3H3-1 | C11orf68 | UPF0696 protein C11orf68 | 0.081486384 | -0.993436813 | Other |
| Q9H3K6 | BOLA2 | BolA-like protein 2 | -0.032477697 | -1.517325719 | Other |
| Q9H3N1 | TMX1 | Thioredoxin-related transmembrane protein 1 | 0.932092031 | 1.633225123 | Cellular metabolism |
| Q9H3P7 | ACBD3 | Golgi resident protein GCP60 | -0.103254954 | -0.640207926 | Lipid metabolism |
| Q9H3Q1 | CDC42EP4 | Cdc42 effector protein 4 | -0.234642665 | 0.544237773 | Other |

| | | | | | |
|---|---|---|---|---|---|
| Q9H3S7 | PTPN23 | Tyrosine-protein phosphatase non-receptor type 23 | 0.031754812 | -0.498001099 | Transport |
| Q9H3U1 | UNC45A | Protein unc-45 homolog A | -0.158016841 | -0.758527756 | Cellular metabolism |
| Q9H410-4 | DSN1 | Kinetochore-associated protein DSN1 homolog | -0.843649546 | -1.194836934 | Other |
| Q9H444 | CHMP4B | Charged multivesicular body protein 4b | -0.05327034 | -0.336909612 | Transport |
| Q9H467 | CUEDC2 | CUE domain-containing protein 2 | -0.638331095 | -0.724872589 | Other |
| Q9H477 | RBKS | Ribokinase | -0.172993342 | -0.172579447 | Cellular metabolism |
| Q9H488 | POFUT1 | GDP-fucose protein O-fucosyltransferase 1 | 0.112179438 | 0.617718379 | Cellular metabolism |
| Q9H490-2 | PIGU | Phosphatidylinositol glycan anchor biosynthesis class U protein | 0 | 1.20063146 | Lipid metabolism |
| Q9H4A4 | RNPEP | Aminopeptidase B | -0.046739578 | -0.631340663 | Lipid metabolism |
| Q9H4A5 | GOLPH3L | Golgi phosphoprotein 3-like | -0.675950368 | -0.227802912 | Other |
| Q9H4A6 | GOLPH3 | Golgi phosphoprotein 3 | -0.334269206 | -1.645130793 | Transport |
| Q9H4L4 | SENP3 | Sentrin-specific protease 3 | 0.547449112 | -0.549057007 | Other |
| Q9H4L5-4 | OSBPL3 | Oxysterol-binding protein-related protein 3 | -0.210532506 | -1.272054672 | Transport |
| Q9H4L7 | SMARCAD1 | SWI/SNF-related matrix-associated actin-dependent regulator of chromatin subfamily A containing DEAD/H box 1 | -0.200838089 | -1.005370458 | Cellular metabolism |
| Q9H4M9 | EHD1 | EH domain-containing protein 1 | 0.176115036 | 0.539112091 | Transport |
| Q9H501 | ESF1 | ESF1 homolog | 0.793777466 | 1.025166194 | Cellular metabolism |
| Q9H583 | HEATR1 | HEAT repeat-containing protein 1 | 0 | 1.007767359 | Cellular metabolism |
| Q9H5V8 | CDCP1 | CUB domain-containing protein 1 | 0 | 0.497946421 | Other |
| Q9H5X1 | FAM96A | MIP18 family protein FAM96A | -1.241709391 | 0 | Cellular metabolism |
| Q9H6K1 | C6orf106 | Uncharacterized protein C6orf106 | -0.982556025 | 0 | Cellular metabolism |
| Q9H6N6 | MYH16 | Putative uncharacterized protein MYH16 | 2.685211182 | 5.532729467 | Other |
| Q9H6Q4 | NARFL | Cytosolic Fe-S cluster assembly factor NARFL | 0.06804657 | 0 | Cellular metabolism |
| Q9H6S3 | EPS8L2 | Epidermal growth factor receptor kinase substrate 8-like protein 2 | 0.284139633 | -0.411646525 | Other |
| Q9H6T3 | RPAP3 | RNA polymerase II-associated protein 3 | 0.126703262 | -0.610253652 | Other |
| Q9H6Y2 | WDR55 | WD repeat-containing protein 55 | -0.113842646 | -1.186190287 | Cellular metabolism |
| Q9H6Z4-3 | RANBP3 | Ran-binding protein 3 | -0.002490997 | -0.397561391 | Transport |
| Q9H788-2 | SH2D4A | SH2 domain-containing protein 4A | 0.179557165 | 0 | Other |
| Q9H7B4 | SMYD3 | Histone-lysine N-methyltransferase SMYD3 | -0.098081589 | 0 | Other |
| Q9H7C4-2 | SYNC | Syncoilin | -0.357088725 | 1.142148336 | Other |
| Q9H7C9 | AAMDC | Mth938 domain-containing protein | -0.225404739 | -1.567296346 | Other |
| Q9H7X7 | IFT22 | Intraflagellar transport protein 22 homolog | -0.366522471 | 0 | Transport |
| Q9H814 | PHAX | Phosphorylated adapter RNA export protein | -0.130882899 | -1.930082321 | Transport |
| Q9H832 | UBE2Z | Ubiquitin-conjugating enzyme E2 Z | -0.117176056 | -0.353244146 | Cellular metabolism |

| | | | | | |
|---|---|---|---|---|---|
| Q9H871 | RMND5A | Protein RMD5 homolog A | -0.819253922 | 0 | Other |
| Q9H875 | PRKRIP1 | PRKR-interacting protein 1 | 0.652330399 | -0.678417206 | Other |
| Q9H8H0 | NOL11 | Nucleolar protein 11 | -0.065716426 | -0.157665253 | Cellular metabolism |
| Q9H8M7 | FAM188A | Protein FAM188A | -0.429719289 | -0.93367068 | Other |
| Q9H8W4 | PLEKHF2 | Pleckstrin homology domain-containing family F member 2 | 0.338627497 | 0.343135834 | Transport |
| Q9H8Y8-3 | GORASP2 | Golgi reassembly-stacking protein 2 | -0.064378103 | -0.050468445 | Other |
| Q9H900 | ZWILCH | Protein zwilch homolog | -0.081457138 | 0 | Other |
| Q9H910 | HN1L | Hematological and neurological expressed 1-like protein | -0.443548838 | -0.229417801 | Other |
| Q9H974 | QTRTD1 | Queuine tRNA-ribosyltransferase subunit QTRTD1 | 0.003448486 | -2.00575765 | Cellular metabolism |
| Q9H977-4 | WDR54 | WD repeat-containing protein 54 | -0.618904114 | -1.00642395 | Other |
| Q9H993 | ARMT1 | Protein-glutamate O-methyltransferase | -0.004025777 | -0.488309224 | Cellular metabolism |
| Q9H999 | PANK3 | Pantothenate kinase 3 | -0.631053289 | 0 | Cellular metabolism |
| Q9H9A6 | LRRC40 | Leucine-rich repeat-containing protein 40 | -0.028373082 | -0.890548706 | Other |
| Q9H9B4 | SFXN1 | Sideroflexin-1 | 0 | 2.444971085 | Transport |
| Q9H9Q2 | COPS7B | COP9 signalosome complex subunit 7b | 0.182509104 | -0.119273504 | Cellular metabolism |
| Q9H9T3 | ELP3 | Elongator complex protein 3 | -0.041662216 | -0.595089595 | Cellular metabolism |
| Q9HA47-3 | UCK1 | Uridine-cytidine kinase 1 | -0.034677505 | -1.691994349 | Cellular metabolism |
| Q9HA64 | FN3KRP | Ketosamine-3-kinase | -0.380572001 | -0.168838501 | Cellular metabolism |
| Q9HA65-3 | TBC1D17 | TBC1 domain family member 17 | -0.608694712 | 0 | Transport |
| Q9HAB8 | PPCS | Phosphopantothenate--cysteine ligase | -0.046700795 | -0.839722315 | Cellular metabolism |
| Q9HAV0 | GNB4 | Guanine nucleotide-binding protein subunit beta-4 | 0 | 2.559118907 | Cellular metabolism |
| Q9HAV4 | XPO5 | Exportin-5 | -0.096511205 | -0.516195297 | Transport |
| Q9HAV7 | GRPEL1 | GrpE protein homolog 1, mitochondrial | -0.311308543 | 0.332492193 | Transport |
| Q9HAW4-3 | CLSPN | Claspin | -0.664809545 | -1.899787903 | Cellular metabolism |
| Q9HB07 | C12orf10 | UPF0160 protein MYG1, mitochondrial | 0.079648336 | -0.4449145 | Other |
| Q9HB40 | SCPEP1 | Retinoid-inducible serine carboxypeptidase | -0.182027817 | -0.106635412 | Lipid metabolism |
| Q9HB71 | CACYBP | Calcyclin-binding protein | 0.107352575 | -0.006284714 | Other |
| Q9HB90 | RRAGC | Ras-related GTP-binding protein C | -0.150725683 | -0.47065417 | Cellular metabolism |
| Q9HBH5 | RDH14 | Retinol dehydrogenase 14 | 1.027914047 | 3.146547318 | Cellular metabolism |
| Q9HBK9 | AS3MT | Arsenite methyltransferase | -0.068968455 | -0.655181249 | Cellular metabolism |
| Q9HBL8 | NMRAL1 | NmrA-like family domain-containing protein 1 | -0.236054103 | -0.340606689 | Other |
| Q9HC38-2 | GLOD4 | Glyoxalase domain-containing protein 4 | -0.297628403 | -0.553777059 | Other |
| Q9HCE0 | EPG5 | Ectopic P granules protein 5 homolog | -0.332759221 | -0.382329941 | Transport |

| | | | | | |
|---|---|---|---|---|---|
| Q9HCE1 | MOV10 | Putative helicase MOV-10 | 1.028495153 | 0 | Immune response |
| Q9HCJ6 | VAT1L | Synaptic vesicle membrane protein VAT-1 homolog-like | -0.131878535 | 0 | Other |
| Q9HCN4 | GPN1 | GPN-loop GTPase 1 | -0.421161652 | 0 | Other |
| Q9HCN8 | SDF2L1 | Stromal cell-derived factor 2-like protein 1 | 0.283183416 | -1.374371847 | Cellular metabolism |
| Q9HCS7 | XAB2 | Pre-mRNA-splicing factor SYF1 | 1.042749405 | -0.830092112 | Cellular metabolism |
| Q9HCU5 | PREB | Prolactin regulatory element-binding protein | 0 | 2.07762591 | Transport |
| Q9HCY8 | S100A14 | Protein S100-A14 | 0 | 0.714611689 | Other |
| Q9HD15 | SRA1 | Steroid receptor RNA activator 1 | 0.269699097 | -1.253619512 | Cellular metabolism |
| Q9HD26-2 | GOPC | Golgi-associated PDZ and coiled-coil motif-containing protein | -0.473908106 | -0.940465927 | Transport |
| Q9HD33 | MRPL47 | 39S ribosomal protein L47, mitochondrial | 0 | 1.160638173 | Cellular metabolism |
| Q9HD45 | TM9SF3 | Transmembrane 9 superfamily member 3 | 0 | 1.595573425 | Other |
| Q9NNW5 | WDR6 | WD repeat-containing protein 6 | -0.301295598 | 0.314154943 | Other |
| Q9NP61 | ARFGAP3 | ADP-ribosylation factor GTPase-activating protein 3 | -0.28883934 | -0.953366597 | Transport |
| Q9NP77 | SSU72 | RNA polymerase II subunit A C-terminal domain phosphatase SSU72 | -0.114011129 | 0.25630633 | Cellular metabolism |
| Q9NP97 | DYNLRB1 | Dynein light chain roadblock-type 1 | -0.069114685 | -1.082605998 | Transport |
| Q9NPA8-2 | ENY2 | Transcription and mRNA export factor ENY2 | -0.60411644 | 0 | Transport |
| Q9NPD3 | EXOSC4 | Exosome complex component RRP41 | 0.226741155 | -0.16952769 | Cellular metabolism |
| Q9NPD8 | UBE2T | Ubiquitin-conjugating enzyme E2 T | -0.568637212 | -1.534393946 | Cellular metabolism |
| Q9NPE3 | NOP10 | H/ACA ribonucleoprotein complex subunit 3 | 0.480155945 | 0 | Cellular metabolism |
| Q9NPF4 | OSGEP | Probable tRNA N6-adenosine threonylcarbamoyltransferase | 0.032037099 | -0.664393743 | Cellular metabolism |
| Q9NPH2-2 | ISYNA1 | Inositol-3-phosphate synthase 1 | -0.089307149 | -0.605257034 | Lipid metabolism |
| Q9NPI6 | DCP1A | mRNA-decapping enzyme 1A | 0.447205226 | 0 | Cellular metabolism |
| Q9NPQ8-4 | RIC8A | Synembryn-A | -0.226170222 | -0.58354187 | Other |
| Q9NQ29-2 | LUC7L | Putative RNA-binding protein Luc7-like 1 | 0.128131231 | -0.014389674 | Other |
| Q9NQ48 | LZTFL1 | Leucine zipper transcription factor-like protein 1 | 0.07975769 | -1.325702031 | Other |
| Q9NQ88 | TIGAR | Fructose-2,6-bisphosphatase TIGAR | 0.008412679 | -0.415754318 | Cellular metabolism |
| Q9NQC3-2 | RTN4 | Reticulon-4 | 0.561284383 | 1.023888906 | Other |
| Q9NQC7-2 | CYLD | Ubiquitin carboxyl-terminal hydrolase CYLD | 0.017237345 | -2.246594111 | Immune response |
| Q9NQE9 | HINT3 | Histidine triad nucleotide-binding protein 3 | -0.214185715 | 1.70553716 | Other |
| Q9NQG5 | RPRD1B | Regulation of nuclear pre-mRNA domain-containing protein 1B | 0.113838196 | -0.279467901 | Cellular metabolism |
| Q9NQR4 | NIT2 | Omega-amidase NIT2 | -0.124212265 | -1.152974447 | Cellular metabolism |
| Q9NQT4 | EXOSC5 | Exosome complex component RRP46 | 0.711332957 | 0 | Cellular metabolism |
| Q9NQT8 | KIF13B | Kinesin-like protein KIF13B | -0.598143895 | -0.770657857 | Transport |
| Q9NQW6 | ANLN | Actin-binding protein anillin | -0.718672434 | -1.801367442 | Other |
| Q9NQW7-3 | XPNPEP1 | Xaa-Pro aminopeptidase 1 | -0.298111598 | -0.5672671 | Cellular metabolism |

| | | | | | |
|---|---|---|---|---|---|
| Q9NQX4 | MYO5C | Unconventional myosin-Vc | 0.366027832 | 0 | Other |
| Q9NQZ2 | UTP3 | Something about silencing protein 10 | 2.830470403 | -0.976865133 | Cellular metabolism |
| Q9NR09 | BIRC6 | Baculoviral IAP repeat-containing protein 6 | -0.186026255 | 0 | Cellular metabolism |
| Q9NR12 | PDLIM7 | PDZ and LIM domain protein 7 | 0.748657862 | 0.249490102 | Transport |
| Q9NR19 | ACSS2 | Acetyl-coenzyme A synthetase, cytoplasmic | -0.088331858 | -0.844760259 | Lipid metabolism |
| Q9NR30 | DDX21 | Nucleolar RNA helicase 2 | 0.418634415 | 0.332471212 | Cellular metabolism |
| Q9NR31 | SAR1A | GTP-binding protein SAR1a | -0.127438227 | -0.00743103 | Transport |
| Q9NR45 | NANS | Sialic acid synthase | -0.081242879 | -0.336951574 | Lipid metabolism |
| Q9NR50 | EIF2B3 | Translation initiation factor eIF-2B subunit gamma | -0.134256999 | -0.784475962 | Cellular metabolism |
| Q9NRD1 | FBXO6 | F-box only protein 6 | -0.179796855 | 0 | Cellular metabolism |
| Q9NRF9 | POLE3 | DNA polymerase epsilon subunit 3 | -0.10580953 | -1.460414886 | Cellular metabolism |
| Q9NRG0 | CHRAC1 | Chromatin accessibility complex protein 1 | -0.041531245 | -0.083307902 | Cellular metabolism |
| Q9NRG1 | PRTFDC1 | Phosphoribosyltransferase domain-containing protein 1 | -0.130686442 | 0 | Cellular metabolism |
| Q9NRG4 | SMYD2 | N-lysine methyltransferase SMYD2 | -1.084621429 | 0 | Cellular metabolism |
| Q9NRL3 | STRN4 | Striatin-4 | -0.130282084 | -1.437421799 | Other |
| Q9NRN7 | AASDHPPT | L-aminoadipate-semialdehyde dehydrogenase-phosphopantetheinyl transferase | 0.388899485 | -0.59333547 | Lipid metabolism |
| Q9NRR5 | UBQLN4 | Ubiquilin-4 | -0.123001734 | 0 | Other |
| Q9NRS6 | SNX15 | Sorting nexin-15 | 0.125066121 | -0.006867091 | Transport |
| Q9NRV9 | HEBP1 | Heme-binding protein 1 | -0.277011236 | -0.329163233 | Other |
| Q9NRW3 | APOBEC3C | DNA dC->dU-editing enzyme APOBEC-3C | 0.038547516 | 0 | Immune response |
| Q9NRX1 | PNO1 | RNA-binding protein PNO1 | -0.255198797 | -1.183570862 | Other |
| Q9NRX4 | PHPT1 | 14 kDa phosphohistidine phosphatase | 0.151773453 | -0.779556274 | Cellular metabolism |
| Q9NRY4 | ARHGAP35 | Rho GTPase-activating protein 35 | -0.661536535 | 0 | Cellular metabolism |
| Q9NRY5 | FAM114A2 | Protein FAM114A2 | -0.091379801 | -1.258700689 | Other |
| Q9NS86 | LANCL2 | LanC-like protein 2 | -0.126955032 | -0.038535436 | Other |
| Q9NS87-2 | KIF15 | Kinesin-like protein KIF15 | 0.167093913 | 0 | Other |
| Q9NS91 | RAD18 | E3 ubiquitin-protein ligase RAD18 | -0.196637472 | -1.026004155 | Cellular metabolism |
| Q9NSC5-2 | HOMER3 | Homer protein homolog 3 | -0.137034734 | 0.491266886 | Transport |
| Q9NSD9 | FARSB | Phenylalanine--tRNA ligase beta subunit | 0.463711421 | 1.295850118 | Cellular metabolism |
| Q9NSE4 | IARS2 | Isoleucine--tRNA ligase, mitochondrial | -0.32183075 | -0.67913119 | Cellular metabolism |
| Q9NSI2 | FAM207A | Protein FAM207A | -0.729520162 | -1.376366297 | Other |
| Q9NSK0 | KLC4 | Kinesin light chain 4 | 0.474484126 | -0.829924266 | Other |
| Q9NSV4-6 | DIAPH3 | Protein diaphanous homolog 3 | 0.06094869 | 0.239400228 | Other |
| Q9NT62 | ATG3 | Ubiquitin-like-conjugating enzyme ATG3 | -0.161186854 | -0.292265574 | Transport |

| | | | | | |
|---|---|---|---|---|---|
| Q9NTJ3 | SMC4 | Structural maintenance of chromosomes protein 4 | 0.110255559 | 0.201583862 | Cellular metabolism |
| Q9NTX5-6 | ECHDC1 | Ethylmalonyl-CoA decarboxylase | -0.086544673 | 0.048619588 | Other |
| Q9NTZ6 | RBM12 | RNA-binding protein 12 | 0.169790904 | -0.206530253 | Other |
| Q9NUJ1 | ABHD10 | Mycophenolic acid acyl-glucuronide esterase, mitochondrial | -1.116000493 | 0.005414963 | Cellular metabolism |
| Q9NUJ3 | TCP11L1 | T-complex protein 11-like protein 1 | -0.046027501 | 0 | Other |
| Q9NUM4 | TMEM106B | Transmembrane protein 106B | 0 | -1.880763372 | Transport |
| Q9NUP7 | TRMT13 | tRNA:m(4)X modification enzyme TRM13 homolog | -0.334632874 | -1.226584117 | Cellular metabolism |
| Q9NUQ3 | TXLNG | Gamma-taxilin | 0.419087728 | 0 | Cellular metabolism |
| Q9NUQ6-2 | SPATS2L | SPATS2-like protein | -0.033100128 | -1.79433314 | Other |
| Q9NUQ8 | ABCF3 | ATP-binding cassette sub-family F member 3 | -0.060825348 | -1.996148427 | Other |
| Q9NUQ9 | FAM49B | Protein FAM49B | 0.061384837 | -0.874914169 | Other |
| Q9NUY8 | TBC1D23 | TBC1 domain family member 23 | -0.423149109 | -1.003977458 | Other |
| Q9NV35 | NUDT15 | Probable 8-oxo-dGTP diphosphatase NUDT15 | -0.19822375 | -1.176876068 | Cellular metabolism |
| Q9NVD7 | PARVA | Alpha-parvin | -0.275162379 | -0.462332408 | Other |
| Q9NVE7 | PANK4 | Pantothenate kinase 4 | -0.157721837 | -0.489183426 | Cellular metabolism |
| Q9NVG8 | TBC1D13 | TBC1 domain family member 13 | -0.083824158 | -1.108425776 | Other |
| Q9NVI7-2 | ATAD3A | ATPase family AAA domain-containing protein 3A | 0.483644485 | 0.68316714 | Other |
| Q9NVJ2 | ARL8B | ADP-ribosylation factor-like protein 8B | -0.065068563 | 1.508574168 | Other |
| Q9NVM4-3 | PRMT7 | Protein arginine N-methyltransferase 7 | -0.2960186 | 0 | Cellular metabolism |
| Q9NVM6 | DNAJC17 | DnaJ homolog subfamily C member 17 | 0.696467717 | -0.307905197 | Other |
| Q9NVP1 | DDX18 | ATP-dependent RNA helicase DDX18 | 0.678655624 | 1.27269872 | Cellular metabolism |
| Q9NVP2 | ASF1B | Histone chaperone ASF1B | -0.42960612 | -0.064336777 | Cellular metabolism |
| Q9NVQ4 | FAIM | Fas apoptotic inhibitory molecule 1 | -0.341108958 | -0.190718969 | Other |
| Q9NVR0 | KLHL11 | Kelch-like protein 11 | -0.77982076 | 0 | Other |
| Q9NVR5 | DNAAF2 | Protein kintoun | -0.412719727 | 0 | Other |
| Q9NVS9 | PNPO | Pyridoxine-5-phosphate oxidase | -0.465180715 | -0.772113164 | Cellular metabolism |
| Q9NVT9 | ARMC1 | Armadillo repeat-containing protein 1 | 0.489282608 | -0.022070567 | Transport |
| Q9NVZ3 | NECAP2 | Adaptin ear-binding coat-associated protein 2 | 0.124050776 | -0.528242747 | Transport |
| Q9NW64 | RBM22 | Pre-mRNA-splicing factor RBM22 | 0.189546585 | 0.286099116 | Transport |
| Q9NW68-2 | BSDC1 | BSD domain-containing protein 1 | -0.54028511 | -0.552930832 | Other |
| Q9NW82 | WDR70 | WD repeat-containing protein 70 | -0.399742762 | 0.568166097 | Other |
| Q9NWM8 | FKBP14 | Peptidyl-prolyl cis-trans isomerase FKBP14 | 0.191748301 | -2.092490514 | Cellular metabolism |
| Q9NWS0 | PIH1D1 | PIH1 domain-containing protein 1 | 0.047952652 | -0.623532613 | Other |
| Q9NWT1 | PAK1IP1 | p21-activated protein kinase-interacting protein 1 | 1.07621638 | 0 | Other |

| | | | | | |
|---|---|---|---|---|---|
| Q9NWT6 | HIF1AN | Hypoxia-inducible factor 1-alpha inhibitor | -0.459896088 | -0.698370616 | Cellular metabolism |
| Q9NWV4 | C1orf123 | UPF0587 protein C1orf123 | -0.226413091 | -1.343235016 | Other |
| Q9NWV8 | BABAM1 | BRISC and BRCA1-A complex member 1 | -0.273979823 | -0.647176107 | Cellular metabolism |
| Q9NWX6 | THG1L | Probable tRNA(His) guanylyltransferase | -0.038387934 | -1.288740794 | Cellular metabolism |
| Q9NWY4 | C4orf27 | UPF0609 protein C4orf27 | -0.352438609 | -2.973033905 | Other |
| Q9NWZ5-4 | UCKL1 | Uridine-cytidine kinase-like 1 | -0.413726807 | 0 | Cellular metabolism |
| Q9NX07 | TRNAU1AP | tRNA selenocysteine 1-associated protein 1 | -0.058284124 | 0 | Cellular metabolism |
| Q9NX24 | NHP2 | H/ACA ribonucleoprotein complex subunit 2 | -0.213950475 | -1.153713862 | Cellular metabolism |
| X6R8U7 | FAM206A | Protein Simiate | 0.523692449 | 0 | Other |
| Q9NX40 | OCIAD1 | OCIA domain-containing protein 1 | 0 | -0.318613052 | Other |
| Q9NX46 | ADPRHL2 | Poly(ADP-ribose) glycohydrolase ARH3 | 0.103321075 | -0.219472249 | Other |
| Q9NX55 | HYPK | Huntingtin-interacting protein K | 0.093011856 | 0 | Other |
| Q9NX58 | LYAR | Cell growth-regulating nucleolar protein | -0.567059199 | -0.457525253 | Other |
| Q9NX62 | IMPAD1 | Inositol monophosphatase 3 | 0 | 1.014334361 | Lipid metabolism |
| Q9NXF1 | TEX10 | Testis-expressed sequence 10 protein | 0.882533391 | 0 | Other |
| Q9NXF7 | DCAF16 | DDB1- and CUL4-associated factor 16 | -0.13262558 | 0 | Cellular metabolism |
| Q9NXG2 | THUMPD1 | THUMP domain-containing protein 1 | -0.050505956 | -0.580658595 | Other |
| Q9NXH9-2 | TRMT1 | tRNA (guanine(26)-N(2))-dimethyltransferase | -0.255968094 | -0.542139053 | Other |
| Q9NXJ5 | PGPEP1 | Pyroglutamyl-peptidase 1 | -0.540348689 | 0 | Other |
| Q9NXK8 | FBXL12 | F-box/LRR-repeat protein 12 | 0.063280106 | 0 | Cellular metabolism |
| Q9NXN4-2 | GDAP2 | Ganglioside-induced differentiation-associated protein 2 | -0.864730199 | -1.299933116 | Other |
| Q9NXR7-4 | BRE | BRCA1-A complex subunit BRE | 0.476714452 | -1.478707631 | Cellular metabolism |
| Q9NXV2 | KCTD5 | BTB/POZ domain-containing protein KCTD5 | -0.215765635 | -0.262257258 | Other |
| Q9NXV6 | CDKN2AIP | CDKN2A-interacting protein | -0.205748876 | 0.325138728 | Other |
| Q9NY12-2 | GAR1 | H/ACA ribonucleoprotein complex subunit 1 | 0.361531576 | 0.072116852 | Transport |
| Q9NY27 | PPP4R2 | Serine/threonine-protein phosphatase 4 regulatory subunit 2 | -0.283319473 | -0.405977249 | Cellular metabolism |
| Q9NY33 | DPP3 | Dipeptidyl peptidase 3 | 0.162323634 | -1.233367284 | Other |
| Q9NYA1 | SPHK1 | Sphingosine kinase 1 | -0.781859716 | -0.605869929 | Lipid metabolism |
| Q9NYB0 | TERF2IP | Telomeric repeat-binding factor 2-interacting protein 1 | -0.411687851 | -0.902364731 | Cellular metabolism |
| Q9NYL9 | TMOD3 | Tropomodulin-3 | 0.092802048 | -0.699342092 | Other |
| Q9NYP9 | MIS18A | Protein Mis18-alpha | -0.248374939 | 0 | Other |
| Q9NYU2-2 | UGGT1 | UDP-glucose:glycoprotein glucosyltransferase 1 | 0.221000036 | 0.01782163 | Cellular metabolism |
| Q9NZ08 | ERAP1 | Endoplasmic reticulum aminopeptidase 1 | 0.027929306 | -0.259309769 | Other |
| Q9NZ09-2 | UBAP1 | Ubiquitin-associated protein 1 | -0.295621236 | 0 | Transport |

| | | | | | |
|---|---|---|---|---|---|
| Q9NZ32 | ACTR10 | Actin-related protein 10 | -0.125579198 | 1.509634654 | Other |
| Q9NZ63 | C9orf78 | Uncharacterized protein C9orf78 | -0.114458084 | 0 | Other |
| Q9NZB2 | FAM120A | Constitutive coactivator of PPAR-gamma-like protein 1 | 1.268604279 | -0.68391482 | Other |
| Q9NZD2 | GLTP | Glycolipid transfer protein | 0.184061686 | -0.730718613 | Lipid metabolism |
| Q9NZD8 | SPG21 | Maspardin | -0.331600189 | -0.612716675 | Other |
| Q9NZH8 | IL36G | Interleukin-36 gamma | 0 | 2.100091298 | Immune response |
| Q9NZI8 | IGF2BP1 | Insulin-like growth factor 2 mRNA-binding protein 1 | 0.672185262 | -1.269437154 | Transport |
| Q9NZJ4 | SACS | Sacsin | 0.265922546 | 0 | Cellular metabolism |
| Q9NZL4 | HSPBP1 | Hsp70-binding protein 1 | -0.009967168 | -0.311274211 | Cellular metabolism |
| Q9NZL9 | MAT2B | Methionine adenosyltransferase 2 subunit beta | 0.000924428 | -0.31921196 | Cellular metabolism |
| Q9NZM1-6 | MYOF | Myoferlin | -0.348576864 | 1.334411621 | Other |
| Q9NZN4 | EHD2 | EH domain-containing protein 2 | -1.230421702 | 0 | Transport |
| Q9NZQ3-3 | NCKIPSD | NCK-interacting protein with SH3 domain | 0.504429499 | 0 | Immune response |
| Q9NZT1 | CALML5 | Calmodulin-like protein 5 | -1.195037842 | 0.846790314 | Other |
| Q9NZT2-2 | OGFR | Opioid growth factor receptor | 0.027812322 | 0.678124746 | Other |
| Q9NZU5-2 | LMCD1 | LIM and cysteine-rich domains protein 1 | 0.191453298 | -0.592975616 | Cellular metabolism |
| Q9NZZ3 | CHMP5 | Charged multivesicular body protein 5 | -0.138700485 | -0.261494954 | Transport |
| Q9P000 | COMMD9 | COMM domain-containing protein 9 | -0.33103625 | -1.204105377 | Transport |
| Q9P013 | CWC15 | Spliceosome-associated protein CWC15 homolog | -0.019823074 | -0.22898229 | Cellular metabolism |
| Q9P016 | THYN1 | Thymocyte nuclear protein 1 | -0.022378922 | 0.077180227 | Other |
| Q9P0J7 | KCMF1 | E3 ubiquitin-protein ligase KCMF1 | 0.170856476 | -2.358222961 | Other |
| Q9P0K7-4 | RAI14 | Ankycorbin | -0.188559214 | 0 | Other |
| Q9P0L0 | VAPA | Vesicle-associated membrane protein-associated protein A | 0.561822891 | 1.583744049 | Lipid metabolism |
| Q9P1F3 | ABRACL | Costars family protein ABRACL | -0.450152715 | 0.311222076 | Other |
| Q9P253 | VPS18 | Vacuolar protein sorting-associated protein 18 homolog | -0.306379318 | -1.223470688 | Transport |
| Q9P258 | RCC2 | Protein RCC2 | -0.177083333 | 0.302141825 | Other |
| Q9P2B4 | CTTNBP2NL | CTTNBP2 N-terminal-like protein | 0.212655385 | -0.839746475 | Cellular metabolism |
| Q9P2D7-6 | DNAH1 | Dynein heavy chain 1, axonemal | -0.925982157 | -0.548553467 | Other |
| Q9P2E3 | ZNFX1 | NFX1-type zinc finger-containing protein 1 | 0.903221766 | 3.336798986 | Other |
| Q9P2E9 | RRBP1 | Ribosome-binding protein 1 | 1.768044154 | 0.725269953 | Transport |
| Q9P2J3 | KLHL9 | Kelch-like protein 9 | -0.439851125 | -0.167961756 | Cellular metabolism |
| Q9P2K6 | KLHL42 | Kelch-like protein 42 | 0.067211787 | 0 | Cellular metabolism |
| Q9P2N5 | RBM27 | RNA-binding protein 27 | 0.265126546 | 0 | Cellular metabolism |
| Q9P2R3 | ANKFY1 | Rabankyrin-5 | 0.079505285 | -1.004898071 | Transport |
| Q9UBB4 | ATXN10 | Ataxin-10 | -0.455869039 | -0.649787903 | Other |

| | | | | | |
|---|---|---|---|---|---|
| Q9UBB6-2 | NCDN | Neurochondrin | -0.058959961 | -0.921243668 | Other |
| Q9UBB9 | TFIP11 | Tuftelin-interacting protein 11 | -0.630233765 | 0.823855718 | Cellular metabolism |
| Q9UBC2-3 | EPS15L1 | Epidermal growth factor receptor substrate 15-like 1 | 0.019964854 | -0.286636353 | Transport |
| Q9UBE0 | SAE1 | SUMO-activating enzyme subunit 1 | -0.167482376 | -0.319073995 | Cellular metabolism |
| Q9UBF2 | COPG2 | Coatomer subunit gamma-2 | 0.568058014 | -1.33041954 | Lipid metabolism |
| Q9UBF6 | RNF7 | RING-box protein 2 | -0.049301783 | -2.428652445 | Cellular metabolism |
| Q9UBG0 | MRC2 | C-type mannose receptor 2 | 0 | -0.17970403 | Transport |
| Q9UBI1 | COMMD3 | COMM domain-containing protein 3 | 0.188505173 | 0 | Transport |
| Q9UBI6 | GNG12 | Guanine nucleotide-binding protein G(I)/G(S)/G(O) subunit gamma-12 | 0 | -0.02360789 | Cellular metabolism |
| Q9UBK8 | MTRR | Methionine synthase reductase | -0.419294993 | -0.441399256 | Cellular metabolism |
| Q9UBM7 | DHCR7 | 7-dehydrocholesterol reductase | 0 | 2.979146322 | Lipid metabolism |
| Q9UBP6 | METTL1 | tRNA (guanine-N(7)-)-methyltransferase | 0.279013316 | -1.528832118 | Cellular metabolism |
| Q9UBQ0 | VPS29 | Vacuolar protein sorting-associated protein 29 | 0.107757568 | -0.279819489 | Transport |
| Q9UBQ5-2 | EIF3K | Eukaryotic translation initiation factor 3 subunit K | -0.012444178 | -0.315233231 | Cellular metabolism |
| Q9UBQ7 | GRHPR | Glyoxylate reductase/hydroxypyruvate reductase | -0.176460902 | -1.067605972 | Transport |
| Q9UBS4 | DNAJB11 | DnaJ homolog subfamily B member 11 | 0.218135834 | 0.624582926 | Cellular metabolism |
| Q9UBS8 | RNF14 | E3 ubiquitin-protein ligase RNF14 | -0.596476237 | -0.266647339 | Cellular metabolism |
| Q9UBT2 | UBA2 | SUMO-activating enzyme subunit 2 | 0.009389242 | -0.427076976 | Cellular metabolism |
| Q9UBT7 | CTNNAL1 | Alpha-catulin | -3.314463298 | 0 | Other |
| Q9UBU9 | NXF1 | Nuclear RNA export factor 1 | -0.003096263 | 0.002246221 | Transport |
| Q9UBV8 | PEF1 | Peflin | -0.026985804 | 1.328813553 | Other |
| Q9UBW8 | COPS7A | COP9 signalosome complex subunit 7a | 0.243489583 | -1.161876043 | Cellular metabolism |
| Q9UDT6-2 | CLIP2 | CAP-Gly domain-containing linker protein 2 | -0.711294174 | 0 | Other |
| Q9UDY2 | TJP2 | Tight junction protein ZO-2 | 0.0598526 | -0.146433512 | Cellular metabolism |
| Q9UDY4 | DNAJB4 | DnaJ homolog subfamily B member 4 | 0.366324107 | -1.795800527 | Cellular metabolism |
| Q9UDY8 | MALT1 | Mucosa-associated lymphoid tissue lymphoma translocation protein 1 | -0.224468231 | -1.259255727 | Immune response |
| Q9UEE5 | STK17A | Serine/threonine-protein kinase 17A | -1.588479996 | 0 | Cellular metabolism |
| Q9UEE9 | CFDP1 | Craniofacial development protein 1 | -0.50426356 | -2.80995369 | Other |
| Q9UER7 | DAXX | Death domain-associated protein 6 | 0.198767344 | -0.424847921 | Cellular metabolism |
| Q9UET6 | FTSJ1 | Putative tRNA (cytidine(32)/guanosine(34)-2-O)-methyltransferase | -0.065642039 | -0.942310333 | Cellular metabolism |
| Q9UEW8-2 | STK39 | STE20/SPS1-related proline-alanine-rich protein kinase | -0.190832774 | -1.105187734 | Cellular metabolism |
| Q9UEY8 | ADD3 | Gamma-adducin | 0.099753062 | -0.977858861 | Transport |
| Q9UG56-2 | PISD | Phosphatidylserine decarboxylase proenzyme | 0.317390442 | 2.495550791 | Lipid metabolism |

| | | | | | |
|---|---|---|---|---|---|
| Q9UG63 | ABCF2 | ATP-binding cassette sub-family F member 2 | -0.0033741 | -0.038942973 | Transport |
| Q9UGI8-2 | TES | Testin | -0.327329 | -0.826601028 | Other |
| Q9UGP4 | LIMD1 | LIM domain-containing protein 1 | -0.288696289 | -0.885435104 | Cellular metabolism |
| Q9UGV2-3 | NDRG3 | Protein NDRG3 | -0.022250493 | -1.632340749 | Other |
| Q9UH62 | ARMCX3 | Armadillo repeat-containing X-linked protein 3 | -0.254359563 | -0.749132792 | Other |
| Q9UH65 | SWAP70 | Switch-associated protein 70 | 0.161060333 | -0.528326035 | Cellular metabolism |
| Q9UHA4 | LAMTOR3 | Ragulator complex protein LAMTOR3 | 0.003477732 | 0.40459315 | Other |
| Q9UHB4-3 | NDOR1 | NADPH-dependent diflavin oxidoreductase 1 | -0.514328003 | 0 | Cellular metabolism |
| Q9UHB9 | SRP68 | Signal recognition particle subunit SRP68 | -0.04518954 | -0.56575648 | Transport |
| Q9UHD1 | CHORDC1 | Cysteine and histidine-rich domain-containing protein 1 | -0.231496811 | -1.528956095 | Cellular metabolism |
| Q9UHD8 | 40057 | Septin-9 | 0.307583491 | -0.656553904 | Other |
| Q9UHD9 | UBQLN2 | Ubiquilin-2 | 0.114526113 | 0.097911835 | Other |
| Q9UHG3-2 | PCYOX1 | Prenylcysteine oxidase 1 | -0.593609492 | 1.576374054 | Cellular metabolism |
| Q9UHI6 | DDX20 | Probable ATP-dependent RNA helicase DDX20 | 0.872143428 | -0.479903539 | Cellular metabolism |
| Q9UHL4 | DPP7 | Dipeptidyl peptidase 2 | 0.230770111 | -0.293198903 | Other |
| Q9UHP3 | USP25 | Ubiquitin carboxyl-terminal hydrolase 25 | -0.333239237 | -0.020151774 | Cellular metabolism |
| Q9UHR6 | ZNHIT2 | Zinc finger HIT domain-containing protein 2 | 0.168854396 | 0 | Other |
| Q9UHV9 | PFDN2 | Prefoldin subunit 2 | -0.254061381 | -0.23161761 | Cellular metabolism |
| Q9UHW5 | GPN3 | GPN-loop GTPase 3 | -0.375301361 | -1.159620285 | Other |
| Q9UHY7 | ENOPH1 | Enolase-phosphatase E1 | -0.260814667 | 2.203223546 | Cellular metabolism |
| Q9UI12-2 | ATP6V1H | V-type proton ATPase subunit H | -0.251113256 | -0.291128794 | Transport |
| Q9UI15 | TAGLN3 | Transgelin-3 | 0.657559077 | -1.010976791 | Other |
| Q9UI26 | IPO11 | Importin-11 | -0.125284195 | -2.073738098 | Transport |
| Q9UI42-2 | CPA4 | Carboxypeptidase A4 | -0.804119746 | 0.59444809 | Cellular metabolism |
| Q9UIA9 | XPO7 | Exportin-7 | 0.188109716 | -0.576043447 | Transport |
| Q9UIC8 | LCMT1 | Leucine carboxyl methyltransferase 1 | -0.20964241 | -1.159420013 | Cellular metabolism |
| Q9UID3 | VPS51 | Vacuolar protein sorting-associated protein 51 homolog | -1.543793996 | 0 | Transport |
| Q9UII2 | ATPIF1 | ATPase inhibitor, mitochondrial | -0.621064504 | -1.864794413 | Cellular metabolism |
| Q9UIJ7 | AK3 | GTP:AMP phosphotransferase AK3, mitochondrial | -0.056856155 | -0.291755676 | Cellular metabolism |
| Q9UIV1-2 | CNOT7 | CCR4-NOT transcription complex subunit 7 | -0.068958918 | 0.153636932 | Cellular metabolism |
| Q9UIV8-2 | SERPINB13 | Serpin B13 | 0 | -0.386253357 | Other |
| Q9UJ68-2 | MSRA | Mitochondrial peptide methionine sulfoxide reductase | -0.096811295 | -0.865835826 | Cellular metabolism |
| Q9UJ70 | NAGK | N-acetyl-D-glucosamine kinase | -0.222675323 | -0.20506223 | Cellular metabolism |

| | | | | | |
|---|---|---|---|---|---|
| Q9UJA5 | TRMT6 | tRNA (adenine(58)-N(1))-methyltransferase non-catalytic subunit TRM6 | 0.063407262 | -0.426530838 | Cellular metabolism |
| Q9UJC5 | SH3BGRL2 | SH3 domain-binding glutamic acid-rich-like protein 2 | -0.122112274 | 0 | Other |
| Q9UJS0-2 | SLC25A13 | Calcium-binding mitochondrial carrier protein Aralar2 | 0 | 1.175113678 | Transport |
| Q9UJU6 | DBNL | Drebrin-like protein | 0.008797963 | -0.474442164 | Transport |
| Q9UJV9 | DDX41 | Probable ATP-dependent RNA helicase DDX41 | 0.625727971 | 0.75557518 | Immune response |
| Q9UJW0 | DCTN4 | Dynactin subunit 4 | 0.059224447 | -1.207607269 | Other |
| Q9UJX2 | CDC23 | Cell division cycle protein 23 homolog | -0.079229355 | 0 | Cellular metabolism |
| Q9UJX3-2 | ANAPC7 | Anaphase-promoting complex subunit 7 | -0.223812103 | -0.181824366 | Cellular metabolism |
| Q9UJY4 | GGA2 | ADP-ribosylation factor-binding protein GGA2 | -0.241346359 | -0.575532913 | Transport |
| Q9UJY5-4 | GGA1 | ADP-ribosylation factor-binding protein GGA1 | -0.363227208 | -1.757550557 | Transport |
| Q9UJZ1-2 | STOML2 | Stomatin-like protein 2, mitochondrial | -0.051825205 | 0.874468486 | Transport |
| Q9UK39 | CCRN4L | Nocturnin | 0.452084859 | 0 | Cellular metabolism |
| Q9UK41 | VPS28 | Vacuolar protein sorting-associated protein 28 homolog | 0.028879801 | -1.516546885 | Transport |
| Q9UK58-4 | CCNL1 | Cyclin-L1 | 0.759543101 | -0.05510203 | Cellular metabolism |
| Q9UK59 | DBR1 | Lariat debranching enzyme | 0.036229451 | 0.672517141 | Cellular metabolism |
| Q9UK76 | HN1 | Hematological and neurological expressed 1 protein | -0.359646479 | -0.035483042 | Other |
| Q9UKD2 | MRTO4 | mRNA turnover protein 4 homolog | 0.366414388 | 0.194746017 | Other |
| Q9UKF6 | CPSF3 | Cleavage and polyadenylation specificity factor subunit 3 | -0.629164378 | -1.367848714 | Transport |
| Q9UKG1 | APPL1 | DCC-interacting protein 13-alpha | 0.118759155 | -0.685757319 | Other |
| Q9UKK3 | PARP4 | Poly [ADP-ribose] polymerase 4 | -0.589331945 | -0.706062317 | Transport |
| Q9UKK6 | NXT1 | NTF2-related export protein 1 | 0.440803528 | 2.697455088 | Transport |
| Q9UKL0 | RCOR1 | REST corepressor 1 | 0.275709152 | -1.730646769 | Cellular metabolism |
| Q9UKN8 | GTF3C4 | General transcription factor 3C polypeptide 4 | -0.075732549 | -0.725200017 | Cellular metabolism |
| Q9UKR5 | C14orf1 | Probable ergosterol biosynthetic protein 28 | 0.559404373 | 1.681943893 | Lipid metabolism |
| Q9UKV8-2 | AGO2 | Protein argonaute-2 | -0.466036479 | 0 | Immune response |
| Q9UKX7 | NUP50 | Nuclear pore complex protein Nup50 | 0.065506617 | -0.506256739 | Transport |
| Q9UKY7 | CDV3 | Protein CDV3 homolog | 0.074042638 | -0.282612483 | Other |
| Q9UL15 | BAG5 | BAG family molecular chaperone regulator 5 | -0.004989624 | 0.805372874 | Cellular metabolism |
| Q9UL63 | MKLN1 | Muskelin | -0.408193588 | 0 | Other |
| Q9ULC4 | MCTS1 | Malignant T-cell-amplified sequence 1 | -0.423542023 | -0.562976837 | Cellular metabolism |
| Q9ULR0 | ISY1 | Pre-mRNA-splicing factor ISY1 homolog | -0.125963847 | -0.646169027 | Cellular metabolism |
| Q9ULT8 | HECTD1 | E3 ubiquitin-protein ligase HECTD1 | -0.416028976 | -1.229204814 | Cellular metabolism |
| Q9ULV4-3 | CORO1C | Coronin-1C | -0.003891627 | -0.671334585 | Transport |
| Q9ULW0 | TPX2 | Targeting protein for Xklp2 | 0.270452499 | 0.132356008 | Other |

| | | | | | |
|---|---|---|---|---|---|
| Q9ULZ3-2 | PYCARD | Apoptosis-associated speck-like protein containing a CARD | -0.735937754 | 0 | Immune response |
| Q9UM22 | EPDR1 | Mammalian ependymin-related protein 1 | 0 | -0.047283173 | Other |
| Q9UMS4 | PRPF19 | Pre-mRNA-processing factor 19 | 0.831000646 | 0.096105576 | Lipid metabolism |
| Q9UMX0 | UBQLN1 | Ubiquilin-1 | -0.152842204 | -0.474990845 | Cellular metabolism |
| Q9UMX5 | NENF | Neudesin | 0.136304855 | -0.14503479 | Other |
| Q9UMZ2-6 | SYNRG | Synergin gamma | -0.994572322 | -0.703125 | Transport |
| Q9UN36 | NDRG2 | Protein NDRG2 | -0.908224106 | 0 | Other |
| Q9UN37 | VPS4A | Vacuolar protein sorting-associated protein 4A | -0.303036372 | 0.29360199 | Transport |
| Q9UN86 | G3BP2 | Ras GTPase-activating protein-binding protein 2 | -0.216580073 | 1.124815623 | Transport |
| Q9UNE7 | STUB1 | E3 ubiquitin-protein ligase CHIP | -0.163763682 | -1.094698588 | Cellular metabolism |
| Q9UNF0-2 | PACSIN2 | Protein kinase C and casein kinase substrate in neurons protein 2 | -0.248573303 | -0.681616465 | Transport |
| Q9UNH6-2 | SNX7 | Sorting nexin-7 | -0.198022207 | 0 | Transport |
| Q9UNH7 | SNX6 | Sorting nexin-6 | 0.039772669 | -0.368484497 | Transport |
| Q9UNI6 | DUSP12 | Dual specificity protein phosphatase 12 | 0.212739309 | -0.112188975 | Cellular metabolism |
| Q9UNM6 | PSMD13 | 26S proteasome non-ATPase regulatory subunit 13 | 0.01385053 | 0.004609426 | Immune response |
| Q9UNN5 | FAF1 | FAS-associated factor 1 | -0.133054097 | -1.454031626 | Cellular metabolism |
| Q9UNP9-2 | PPIE | Peptidyl-prolyl cis-trans isomerase E | -0.102776845 | 0.947509766 | Cellular metabolism |
| Q9UNS1-2 | TIMELESS | Protein timeless homolog | -0.590868632 | -0.764484406 | Cellular metabolism |
| Q9UNS2 | COPS3 | COP9 signalosome complex subunit 3 | -0.011551539 | -0.291968028 | Cellular metabolism |
| Q9UNT1-2 | RABL2B | Rab-like protein 2B | 0.045078913 | 0 | Transport |
| Q9UNW1 | MINPP1 | Multiple inositol polyphosphate phosphatase 1 | -0.14991188 | -0.999148051 | Cellular metabolism |
| Q9UNZ2 | NSFL1C | NSFL1 cofactor p47 | 0.010564168 | -0.218685786 | Cellular metabolism |
| Q9UPN7 | PPP6R1 | Serine/threonine-protein phosphatase 6 regulatory subunit 1 | -0.590274811 | -0.93589592 | Other |
| Q9UPN9-2 | TRIM33 | E3 ubiquitin-protein ligase TRIM33 | 0.222069422 | 0.467669169 | Cellular metabolism |
| Q9UPU5 | USP24 | Ubiquitin carboxyl-terminal hydrolase 24 | 0.175771077 | -0.881605148 | Cellular metabolism |
| Q9UPU7-2 | TBC1D2B | TBC1 domain family member 2B | -0.341584524 | 0 | Other |
| Q9UPV0-2 | CEP164 | Centrosomal protein of 164 kDa | -0.317897797 | -0.244723002 | Cellular metabolism |
| Q9UQ35 | SRRM2 | Serine/arginine repetitive matrix protein 2 | 0.272172928 | -0.872519811 | Cellular metabolism |
| Q9UQ80 | PA2G4 | Proliferation-associated protein 2G4 | 0.223453522 | 0.036827087 | Cellular metabolism |
| Q9UQE7 | SMC3 | Structural maintenance of chromosomes protein 3 | 0.418678919 | -1.201627096 | Cellular metabolism |
| Q9Y223 | GNE | Bifunctional UDP-N-acetylglucosamine 2-epimerase/N-acetylmannosamine kinase | -0.080038706 | -1.457417806 | Cellular metabolism |
| Q9Y224 | C14orf166 | UPF0568 protein C14orf166 | -0.423283895 | 0.481210073 | Transport |
| Q9Y230 | RUVBL2 | RuvB-like 2 | 0.105872472 | -0.175690969 | Cellular metabolism |

| | | | | | |
|---|---|---|---|---|---|
| Q9Y237 | PIN4 | Peptidyl-prolyl cis-trans isomerase NIMA-interacting 4 | -0.231948217 | 0 | Cellular metabolism |
| Q9Y262 | EIF3L | Eukaryotic translation initiation factor 3 subunit L | 0.23428154 | -0.286776861 | Other |
| Q9Y263 | PLAA | Phospholipase A-2-activating protein | 0.052125295 | -0.516054789 | Lipid metabolism |
| Q9Y265 | RUVBL1 | RuvB-like 1 | -0.060365041 | 0.215588888 | Cellular metabolism |
| Q9Y266 | NUDC | Nuclear migration protein nudC | -0.112909953 | -0.35033226 | Other |
| Q9Y277 | VDAC3 | Voltage-dependent anion-selective channel protein 3 | 0 | 0.722075144 | Transport |
| Q9Y281 | CFL2 | Cofilin-2 | -0.200129191 | -0.803393046 | Other |
| Q9Y285 | FARSA | Phenylalanine--tRNA ligase alpha subunit | 0.39556249 | 0.196280162 | Cellular metabolism |
| Q9Y287 | ITM2B | Integral membrane protein 2B | -0.399828593 | 1.199063619 | Other |
| Q9Y294 | ASF1A | Histone chaperone ASF1A | -0.152008692 | -0.362354279 | Cellular metabolism |
| Q9Y295 | DRG1 | Developmentally-regulated GTP-binding protein 1 | -0.170078278 | -0.664709091 | Cellular metabolism |
| Q9Y296 | TRAPPC4 | Trafficking protein particle complex subunit 4 | -0.485242844 | 0 | Transport |
| Q9Y2A7 | NCKAP1 | Nck-associated protein 1 | 0.305793762 | -0.401465098 | Immune response |
| Q9Y2D5 | AKAP2 | A-kinase anchor protein 2 | -0.055258433 | -0.429767609 | Other |
| Q9Y2G5 | POFUT2 | GDP-fucose protein O-fucosyltransferase 2 | -0.517066956 | 0.694352468 | Cellular metabolism |
| Q9Y2H0-1 | DLGAP4 | Disks large-associated protein 4 | -0.36631012 | -0.225415548 | Other |
| Q9Y2J2-2 | EPB41L3 | Band 4.1-like protein 3 | 0.046751022 | -6.38178889 | Other |
| Q9Y2L1 | DIS3 | Exosome complex exonuclease RRP44 | 0.150487264 | 0.269947688 | Cellular metabolism |
| Q9Y2Q5-2 | LAMTOR2 | Ragulator complex protein LAMTOR2 | -0.104094823 | 0 | Cellular metabolism |
| Q9Y2Q9 | MRPS28 | 28S ribosomal protein S28, mitochondrial | 0 | 0.347873052 | Cellular metabolism |
| Q9Y2S6 | TMA7 | Translation machinery-associated protein 7 | 0.116666158 | -0.736005147 | Other |
| Q9Y2S7 | POLDIP2 | Polymerase delta-interacting protein 2 | -0.087544759 | -0.170055389 | Other |
| Q9Y2T2 | AP3M1 | AP-3 complex subunit mu-1 | 0.01810201 | -0.069995244 | Transport |
| Q9Y2T3 | GDA | Guanine deaminase | 0.511203766 | 0 | Cellular metabolism |
| Q9Y2U5 | MAP3K2 | Mitogen-activated protein kinase kinase kinase 2 | -0.684562047 | 0 | Cellular metabolism |
| Q9Y2V2 | CARHSP1 | Calcium-regulated heat stable protein 1 | 0.047134399 | -1.494519552 | Cellular metabolism |
| Q9Y2V7 | COG6 | Conserved oligomeric Golgi complex subunit 6 | 0.667151769 | 1.05380249 | Transport |
| Q9Y2W2 | WBP11 | WW domain-binding protein 11 | 0.155058543 | 0.179997762 | Cellular metabolism |
| Q9Y2X3 | NOP58 | Nucleolar protein 58 | 0.131175995 | -0.262846629 | Transport |
| Q9Y2Y0-2 | ARL2BP | ADP-ribosylation factor-like protein 2-binding protein | -0.661553065 | -0.16591835 | Cellular metabolism |
| Q9Y2Z0-2 | SUGT1 | Suppressor of G2 allele of SKP1 homolog | -0.221160889 | -0.721897761 | Immune response |
| Q9Y303 | AMDHD2 | Putative N-acetylglucosamine-6-phosphate deacetylase | -0.296246847 | 0.115371704 | Cellular metabolism |
| Q9Y305-2 | ACOT9 | Acyl-coenzyme A thioesterase 9, mitochondrial | 0.287371318 | 0.275332769 | Cellular metabolism |
| Q9Y314 | NOSIP | Nitric oxide synthase-interacting protein | -0.279640834 | -0.448961258 | Other |

| | | | | | |
|---|---|---|---|---|---|
| Q9Y315 | DERA | Deoxyribose-phosphate aldolase | 0.111166636 | -0.064736048 | Cellular metabolism |
| Q9Y316 | MEMO1 | Protein MEMO1 | -0.119936625 | 0.000738144 | Other |
| Q9Y333 | LSM2 | U6 snRNA-associated Sm-like protein LSm2 | 0.293886185 | 0.038024902 | Cellular metabolism |
| Q9Y376 | CAB39 | Calcium-binding protein 39 | -0.330201467 | -0.821878433 | Other |
| Q9Y383 | LUC7L2 | Putative RNA-binding protein Luc7-like 2 | 0.293458939 | -0.146398544 | Other |
| Q9Y3A6 | TMED5 | Transmembrane emp24 domain-containing protein 5 | 0 | 0.663219452 | Transport |
| Q9Y3B4 | SF3B6 | Splicing factor 3B subunit 6 | 0.180176417 | 1.664482752 | Cellular metabolism |
| Q9Y3B7 | MRPL11 | 39S ribosomal protein L11, mitochondrial | 0 | -0.510819117 | Cellular metabolism |
| Q9Y3B9 | RRP15 | RRP15-like protein | -0.149887721 | -0.027372996 | Other |
| Q9Y3C4-3 | TPRKB | EKC/KEOPS complex subunit TPRKB | -0.140642166 | 0 | Cellular metabolism |
| Q9Y3C6 | PPIL1 | Peptidyl-prolyl cis-trans isomerase-like 1 | 0.30256017 | -0.830194473 | Cellular metabolism |
| Q9Y3C8 | UFC1 | Ubiquitin-fold modifier-conjugating enzyme 1 | -0.161835988 | -2.06325531 | Cellular metabolism |
| Q9Y3D6 | FIS1 | Mitochondrial fission 1 protein | 0.035830816 | 0.949078242 | Transport |
| Q9Y3D8 | AK6 | Adenylate kinase isoenzyme 6 | -0.020879745 | 0.445788066 | Cellular metabolism |
| Q9Y3E1 | HDGFRP3 | Hepatoma-derived growth factor-related protein 3 | -2.563597361 | 0.944342295 | Other |
| Q9Y3E7 | CHMP3 | Charged multivesicular body protein 3 | -0.22505188 | -0.232211431 | Transport |
| Q9Y3F4-2 | STRAP | Serine-threonine kinase receptor-associated protein | 0.133756638 | -0.448015849 | Other |
| Q9Y3I0 | RTCB | tRNA-splicing ligase RtcB homolog | -0.066740036 | 0.001092911 | Cellular metabolism |
| Q9Y3I1 | FBXO7 | F-box only protein 7 | -0.224419912 | -0.237591426 | Transport |
| Q9Y3L5 | RAP2C | Ras-related protein Rap-2c | -0.313047409 | -0.184342066 | Transport |
| Q9Y3P9 | RABGAP1 | Rab GTPase-activating protein 1 | -0.030483246 | -0.901215871 | Other |
| Q9Y3Q8 | TSC22D4 | TSC22 domain family protein 4 | -0.061084112 | -0.278372447 | Cellular metabolism |
| Q9Y3R4 | NEU2 | Sialidase-2 | 0 | 1.829270045 | Lipid metabolism |
| Q9Y3S2 | ZNF330 | Zinc finger protein 330 | 0.131344477 | -0.220941544 | Other |
| Q9Y3T9 | NOC2L | Nucleolar complex protein 2 homolog | 0.698874156 | 0.177398046 | Transport |
| Q9Y3U8 | RPL36 | 60S ribosomal protein L36 | 0.631744385 | -0.12641271 | Transport |
| Q9Y3Z3 | SAMHD1 | Deoxynucleoside triphosphate triphosphohydrolase SAMHD1 | 0.107469559 | 0.934901555 | Immune response |
| Q9Y450 | HBS1L | HBS1-like protein | 0.041407267 | -1.183314006 | Cellular metabolism |
| Q9Y478 | PRKAB1 | 5-AMP-activated protein kinase subunit beta-1 | 0.103887558 | 0.802597046 | Lipid metabolism |
| Q9Y490 | TLN1 | Talin-1 | -0.072094599 | -0.523617427 | Transport |
| Q9Y4B5-3 | MTCL1 | Microtubule cross-linking factor 1 | 0.253014247 | 0.112577438 | Other |
| Q9Y4B6-3 | VPRBP | Protein VPRBP | -0.478122075 | 0 | Cellular metabolism |
| Q9Y4C2-2 | TCAF1 | TRPM8 channel-associated factor 1 | -1.341856003 | -0.828927994 | Transport |
| Q9Y4E8 | USP15 | Ubiquitin carboxyl-terminal hydrolase 15 | -0.200520833 | -0.351435979 | Cellular metabolism |

| | | | | | |
|---|---|---|---|---|---|
| Q9Y4G6 | TLN2 | Talin-2 | -1.258832296 | 0 | Other |
| Q9Y4K0 | LOXL2 | Lysyl oxidase homolog 2 | -1.758871714 | 0 | Cellular metabolism |
| Q9Y4K4 | MAP4K5 | Mitogen-activated protein kinase kinase kinase kinase 5 | 1.132656097 | -0.524218241 | Cellular metabolism |
| Q9Y4P1-6 | ATG4B | Cysteine protease ATG4B | -0.215516408 | -0.819131215 | Transport |
| Q9Y4P8-4 | WIPI2 | WD repeat domain phosphoinositide-interacting protein 2 | 0.140959422 | 1.085378011 | Cellular metabolism |
| Q9Y4P8-2 | WIPI2 | WD repeat domain phosphoinositide-interacting protein 2 | 0.053643545 | 0 | Cellular metabolism |
| Q9Y4R8 | TELO2 | Telomere length regulation protein TEL2 homolog | -0.497221629 | -0.582946777 | Other |
| Q9Y4W2-2 | LAS1L | Ribosomal biogenesis protein LAS1L | 0.093128204 | -1.018283208 | Cellular metabolism |
| Q9Y4W6 | AFG3L2 | AFG3-like protein 2 | 0 | 1.101818085 | Cellular metabolism |
| Q9Y4X5 | ARIH1 | E3 ubiquitin-protein ligase ARIH1 | -0.235018412 | -0.526164373 | Cellular metabolism |
| V9GZ56 | LSM4 | U6 snRNA-associated Sm-like protein LSm4 | 0.533959707 | -0.618822734 | Cellular metabolism |
| Q9Y508 | RNF114 | E3 ubiquitin-protein ligase RNF114 | -0.341849645 | -0.620111465 | Cellular metabolism |
| Q9Y547 | HSPB11 | Intraflagellar transport protein 25 homolog | -0.380110423 | 0.088000615 | Transport |
| Q9Y570 | PPME1 | Protein phosphatase methylesterase 1 | -0.161160151 | -1.318468094 | Cellular metabolism |
| Q9Y575-2 | ASB3 | Ankyrin repeat and SOCS box protein 3 | -0.473323822 | 0 | Cellular metabolism |
| Q9Y5A9 | YTHDF2 | YTH domain-containing family protein 2 | -0.069609324 | 0.146500905 | Immune response |
| Q9Y5B8-2 | NME7 | Nucleoside diphosphate kinase 7 | -0.328631719 | -1.718779882 | Transport |
| Q9Y5B9 | SUPT16H | FACT complex subunit SPT16 | -0.161249161 | -0.383444468 | Cellular metabolism |
| Q9Y5K5-2 | UCHL5 | Ubiquitin carboxyl-terminal hydrolase isozyme L5 | 0.010666529 | 0.586403529 | Cellular metabolism |
| Q9Y5K6 | CD2AP | CD2-associated protein | -0.076294581 | 0.130371094 | Other |
| Q9Y5K8 | ATP6V1D | V-type proton ATPase subunit D | -0.173278809 | -1.148091634 | Transport |
| Q9Y5L0-3 | TNPO3 | Transportin-3 | -0.072566986 | -0.743996302 | Transport |
| Q9Y5L4 | TIMM13 | Mitochondrial import inner membrane translocase subunit Tim13 | -0.065200806 | 0.426770528 | Transport |
| Q9Y5M8 | SRPRB | Signal recognition particle receptor subunit beta | 0.59292539 | 1.129131317 | Transport |
| Q9Y5N5-2 | N6AMT1 | HemK methyltransferase family member 2 | -0.338847478 | 0 | Cellular metabolism |
| Q9Y5P6 | GMPPB | Mannose-1-phosphate guanyltransferase beta | 0.229895274 | -0.851309458 | Cellular metabolism |
| Q9Y5Q8-2 | GTF3C5 | General transcription factor 3C polypeptide 5 | 0.355946223 | 0.015769958 | Cellular metabolism |
| Q9Y5R8 | TRAPPC1 | Trafficking protein particle complex subunit 1 | -0.319086711 | 0 | Transport |
| Q9Y5S9 | RBM8A | RNA-binding protein 8A | -0.087003708 | -0.176277161 | Transport |
| Q9Y5T5-2 | USP16 | Ubiquitin carboxyl-terminal hydrolase 16 | -0.409399668 | 0 | Cellular metabolism |
| Q9Y5U2-2 | TSSC4 | Protein TSSC4 | 0.24822108 | -1.096985499 | Other |
| Q9Y5X2 | SNX8 | Sorting nexin-8 | 0.200749079 | 0.975033442 | Transport |
| Q9Y5X3 | SNX5 | Sorting nexin-5 | -0.154167175 | -0.689196269 | Transport |

| | | | | | |
|---|---|---|---|---|---|
| Q9Y5Y2 | NUBP2 | Cytosolic Fe-S cluster assembly factor NUBP2 | -0.227031072 | -0.170128504 | Cellular metabolism |
| Q9Y5Z4 | HEBP2 | Heme-binding protein 2 | 0.096960704 | -0.511957804 | Other |
| Q9Y608-4 | LRRFIP2 | Leucine-rich repeat flightless-interacting protein 2 | -0.05625089 | -0.187684377 | Other |
| Q9Y613 | FHOD1 | FH1/FH2 domain-containing protein 1 | 0.188203176 | 0 | Other |
| Q9Y617 | PSAT1 | Phosphoserine aminotransferase | -0.036656698 | -1.096462886 | Cellular metabolism |
| Q9Y657 | SPIN1 | Spindlin-1 | -0.37437884 | 0.114829381 | Cellular metabolism |
| Q9Y678 | COPG1 | Coatomer subunit gamma-1 | -0.100980123 | -0.659122467 | Transport |
| Q9Y680-3 | FKBP7 | Peptidyl-prolyl cis-trans isomerase FKBP7 | -0.140088399 | 0 | Cellular metabolism |
| Q9Y696 | CLIC4 | Chloride intracellular channel protein 4 | -0.45562617 | -0.444358826 | Transport |
| Q9Y6A4 | CFAP20 | Cilia- and flagella-associated protein 20 | -0.018814723 | -0.550186157 | Cellular metabolism |
| Q9Y6A5 | TACC3 | Transforming acidic coiled-coil-containing protein 3 | -0.378862381 | -0.8288517 | Other |
| Q9Y6B6 | SAR1B | GTP-binding protein SAR1b | -0.112058004 | -0.620632807 | Transport |
| Q9Y6D6 | ARFGEF1 | Brefeldin A-inhibited guanine nucleotide-exchange protein 1 | -0.003145854 | -1.627852122 | Transport |
| Q9Y6D9 | MAD1L1 | Mitotic spindle assembly checkpoint protein MAD1 | 0.465703964 | 0.763475418 | Other |
| Q9Y6G9 | DYNC1LI1 | Cytoplasmic dynein 1 light intermediate chain 1 | 0.10351181 | -0.029809952 | Transport |
| Q9Y6H1 | CHCHD2 | Coiled-coil-helix-coiled-coil-helix domain-containing protein 2 | -1.903194427 | -0.501785914 | Cellular metabolism |
| Q9Y6I3-3 | EPN1 | Epsin-1 | -0.006201426 | 0.241162618 | Transport |
| Q9Y6K9-2 | IKBKG | NF-kappa-B essential modulator | -0.085069021 | -2.323764165 | Immune response |
| Q9Y6N5 | SQRDL | Sulfide:quinone oxidoreductase, mitochondrial | 0 | 1.506532033 | Cellular metabolism |
| Q9Y6W5 | WASF2 | Wiskott-Aldrich syndrome protein family member 2 | -0.520668666 | -0.320847193 | Immune response |
| Q9Y6Y0 | IVNS1ABP | Influenza virus NS1A-binding protein | -0.848788579 | -1.357397715 | Cellular metabolism |
| Q9Y6Y8 | SEC23IP | SEC23-interacting protein | -0.052139282 | -0.693941116 | Transport |
| S4R3H4 | ACIN1 | Apoptotic chromatin condensation inducer in the nucleus | -0.082501094 | -0.201570511 | Cellular metabolism |
| S4R3N1 | HSPE1-MOB4 | MOB-like protein phocein | -0.063262939 | 0.159543355 | Transport |
