## Supplementary material for "Virally induced lipid droplets are a platform for innate immune signalling complexes": S Table 3

### Table S3: LD Proteins identified and their regulation in Astrocytes infected with ZIKV

| Protein IDs | Gene Names | Protein Descriptions | Log2FC (ZIKV 24hpi vs Control) | Log2FC (ZIKV 8hpi vs Control) |
|---|---|---|---|---|
| **Proteins upregulated following ZIKV infection (24hrs)** | | | | |
| A0A669KB53 | STAT1 | Signal transducer and activator of transcription | 1.786000888 | 0.221436 |
| A0A7P0T855 | IFIT3 | Interferon-induced protein with tetratricopeptide repeats 3 | 3.1630222 | -0.373457 |
| E9PK67 | PARP10 | Poly [ADP-ribose] polymerase | 3.143637101 | 0.175685 |
| E9PK91 | BCLAF1 | Bcl-2-associated transcription factor 1 | 7.739313165 | 0 |
| G3V150 | B3GAT3 | Galactosylgalactosylxylosylprotein 3-beta-glucuronosyltransferase | 9.314172029 | 0 |
| J3KQL8 | APOL2 | Apolipoprotein L2 | 1.278439204 | 0.0788997 |
| K7EKI0 | EVPL | Envoplakin | 4.426320791 | 0 |
| O14933 | UBE2L6 | Ubiquitin/ISG15-conjugating enzyme E2 L6 | 1.095478058 | 0.484787 |
| O95786 | DDX58 | Antiviral innate immune response receptor RIG-I | 2.69200786 | 0.324314 |
| P05161 | ISG15 | Ubiquitin-like protein ISG15 | 3.238688151 | 0.476218 |
| P09914 | IFIT1 | Interferon-induced protein with tetratricopeptide repeats 1 | 4.702760696 | 1.13822 |
| P19474 | TRIM21 | E3 ubiquitin-protein ligase TRIM21 | 1.220693509 | 0.0991888 |
| P29590 | PML | Protein PML | 1.132013798 | 0.159978 |
| P32455 | GBP1 | Guanylate-binding protein 1 | 1.264704784 | -0.0863787 |
| Q00978 | IRF9 | Interferon regulatory factor 9 | 2.15659229 | 0.294558 |
| Q13325 | IFIT5 | Interferon-induced protein with tetratricopeptide repeats 5 | 1.187128703 | 0.724266 |
| Q5EBM0 | CMPK2 | UMP-CMP kinase 2, mitochondrial | 3.841128031 | 0 |
| Q8TCB0 | IFI44 | Interferon-induced protein 44 | 3.677878698 | 0.158234 |
| Q8WXG1 | RSAD2 | Radical S-adenosyl methionine domain-containing protein 2 | 5.519729932 | 0 |
| Q9BQE5 | APOL2 | Apolipoprotein L2 | 0.831963062 | 0.0902853 |
| Q9BYX4 | IFIH1 | Interferon-induced helicase C domain-containing protein 1 | 2.324022373 | -0.622855 |
| Q9Y3Z3 | SAMHD1 | Deoxynucleoside triphosphate triphosphohydrolase SAMHD1 | 1.139678081 | 0.153357 |
| **Proteins downregulated following ZIKV infection (24hrs)** | | | | |
| A0A0J9YWQ2 | PTDSS2 | L-serine-phosphatidylethanolamine phosphatidyltransferase (Fragment) | -2.093715191 | -0.864916 |
| A0A2R8Y518 | TBC1D24 | TBC1 domain family member 24 (Fragment) | -1.163009961 | -0.586502 |
| A0A804HHX0 | BSCL2 | Seipin | -2.149681886 | -0.816231 |
| A6NHL2 | TUBAL3 | Tubulin alpha chain-like 3 | -2.167627811 | -4.50237 |
| C9JZB0 | INPP5K | Inositol polyphosphate 5-phosphatase K (Fragment) | -2.351265192 | -0.357906 |
| E9PHI4 | SUN1 | SUN domain-containing protein 1 | -2.108109395 | 0.371393 |
| F5H6T0 | ABHD4 | (Lyso)-N-acylphosphatidylethanolamine lipase (Fragment) | -2.292742729 | -0.321578 |
| F8VSA1 | TARBP2 | RISC-loading complex subunit TARBP2 (Fragment) | -1.31394426 | -2.21537 |
| K7EK01 | VMP1 | Vacuole membrane protein 1 (Fragment) | -3.164499362 | 0.365029 |
| O75554 | WBP4 | WW domain-binding protein 4 | -1.112925212 | -2.03895 |
| O94915 | FRYL | Protein furry homolog-like | -0.544808547 | -2.78032 |
| P01833 | PIGR | Polymeric immunoglobulin receptor | -1.761735837 | -4.4263 |
| P17066 | HSPA6 | Heat shock 70 kDa protein 6 | -2.902829568 | 0.260303 |
| P82970 | HMGN5 | High mobility group nucleosome-binding domain-containing protein 5 | -2.988786062 | 0 |
| Q7LGA3 | HS2ST1 | Heparan sulfate 2-O-sulfotransferase 1 | -2.88745443 | 0.73856 |
| Q8N5M9 | JAGN1 | Protein jagunal homolog 1 | -4.139576515 | -0.781759 |
| Q99767 | APBA2 | Amyloid-beta A4 precursor protein-binding family A member 2 | -3.555211782 | -1.74115 |
| Q9BW62 | KATNAL1 | Katanin p60 ATPase-containing subunit A-like 1 | -0.884017547 | -0.426958 |
| Q9H1I8 | ASCC2 | Activating signal cointegrator 1 complex subunit 2 | -2.049494267 | -0.199719 |
| Q9H553 | ALG2 | Alpha-1,3/1,6-mannosyltransferase ALG2 | -1.455616633 | 0.0509965 |
| Q9H875 | PRKRIP1 | PRKR-interacting protein 1 | -2.674875259 | 0 |
| Q9NRG9 | AAAS | Aladin | -1.613782088 | -0.145578 |
| Q9NSV4 | DIAPH3 | Protein diaphanous homolog 3 | -1.310214837 | -0.263368 |
| **Proteins present/unchanged** | | | | |
| A0A024R4E5 | HDLBP | High density lipoprotein binding protein (Vigilin), isoform CRA_a | -0.42253081 | -0.064833 |
| A0A024R5G9 | CTSF | Cathepsin F | -0.486051957 | -0.475324 |
| A0A024R6I7 | SERPINA1 | Alpha-1-antitrypsin | -0.104571422 | -0.422489 |
| A0A024R7E8 | ELOF1 | Transcription elongation factor 1 homolog | -0.739141067 | 0.593805 |
| A0A024RA52 | PSMA2 | Proteasome subunit alpha type | -0.144051472 | 0.11978 |
| A0A024RBG1 | NUDT4B | Diphosphoinositol polyphosphate phosphohydrolase NUDT4B | -0.007428885 | -0.082551 |
| A0A024RCQ8 | PPT2 | HCG1999928, isoform CRA_a | -0.654887358 | 0.333566 |
| A0A024RCR6 | BAG6 | BCL2-associated athanogene 6 | 0.17121315 | -0.117184 |
| A0A075B6Q2 | FNBP1L | Formin-binding protein 1-like (Fragment) | 1.727155447 | 0 |
| A0A075B6T4 | KBTBD4 | Kelch repeat and BTB domain-containing protein 4 | -0.143869321 | -1.28591 |
| A0A075B716 | RPS17 | 40S ribosomal protein S17 | -0.497846047 | -0.0812155 |
| A0A075B730 | EPPK1 | Epiplakin | 0 | -0.578883 |
| A0A075B749 | PDE4DIP | Myomegalin | 0.40034771 | -0.0698484 |
| A0A075B784 | USP36 | Ubiquitin carboxyl-terminal hydrolase | -0.584647099 | 0.21756 |
| A0A075B7B5 | SRGAP2 | SLIT-ROBO Rho GTPase-activating protein 2 | -0.080765009 | -0.322813 |
| A0A087WSV2 | | Biogenesis of lysosome-related organelles complex 1 subunit 1 (Fragment) | 1.688123544 | 0 |

| | | | | |
|---|---|---|---|---|
| A0A087WSW7 | HELLS | Helicase, lymphoid-specific, isoform CRA_d | 0.34842068 | 0 |
| A0A087WSW9 | TXNRD1 | Thioredoxin-disulfide reductase | -0.078398228 | -0.210824 |
| A0A087WT12 | GPX4 | Glutathione peroxidase | -0.251900593 | 0.125774 |
| A0A087WT20 | DCAF13 | DDB1- and CUL4-associated factor 13 | -1.076944192 | -0.318667 |
| A0A087WT99 | C11orf54 | Ester hydrolase C11orf54 | 0.709142764 | -1.2939 |
| A0A087WTA5 | EIF2B4 | Translation initiation factor eIF-2B subunit delta | -0.714248816 | 0.394897 |
| A0A087WTA8 | COL1A2 | Collagen alpha-2(I) chain | -0.711564938 | -0.439888 |
| A0A087WTC6 | MND1 | Meiotic nuclear division protein 1 homolog | -0.196010749 | -0.619385 |
| A0A087WTM5 | RNF34 | E3 ubiquitin-protein ligase RNF34 | -0.884259462 | 2.1746 |
| A0A087WTU3 | TEX264 | Testis-expressed protein 264 | -1.569614331 | -0.0204051 |
| A0A087WTU9 | ALMS1 | Alstrom syndrome protein 1 | -0.561461767 | -1.00413 |
| A0A087WTV6 | PYCR2 | Pyrroline-5-carboxylate reductase | -1.718885899 | -1.17003 |
| A0A087WTW0 | UHRF1 | RING-type E3 ubiquitin transferase | -1.122967164 | 0.378203 |
| A0A087WTY9 | MRFAP1 | MORF4 family-associated protein 1 | -0.464458068 | 0 |
| A0A087WU02 | ERGIC2 | Endoplasmic reticulum-Golgi intermediate compartment protein 2 | -1.721195539 | -1.72829 |
| A0A087WU13 | CD9 | Tetraspanin | -2.322664817 | 1.22872 |
| A0A087WU53 | MAGT1 | Dolichyl-diphosphooligosaccharide--protein glycosyltransferase subunit MAGT1 | -1.271516482 | 0.187705 |
| A0A087WU80 | TNFAIP3 | Ubiquitinyl hydrolase 1 | -0.227277915 | 0.0911903 |
| A0A087WUB9 | CTNNBL1 | Beta-catenin-like protein 1 | 0.027903795 | 0.039058 |
| A0A087WUC2 | PCDHGC4 | Protocadherin gamma-C4 | -1.559509198 | 1.47201 |
| A0A087WUC6 | SPCS2 | Signal peptidase complex subunit 2 | -0.975070318 | 0.219755 |
| A0A087WUD3 | OSTC | Oligosaccharyltransferase complex subunit | -1.644385576 | 0.694421 |
| A0A087WUE4 | LENG8 | Leukocyte receptor cluster member 8 | -0.532923619 | -1.52683 |
| A0A087WUK2 | HNRNPDL | Heterogeneous nuclear ribonucleoprotein D-like | -0.367968798 | 0.40607 |
| A0A087WUL4 | PEG10 | Retrotransposon-derived protein PEG10 | -0.172757943 | -0.2542 |
| A0A087WUT0 | MYEF2 | Myelin expression factor 2 | -0.168137312 | 0 |
| A0A087WUT6 | EIF5B | Eukaryotic translation initiation factor 5B | -0.210644404 | -0.395397 |
| A0A087WUW6 | IKBKG | NF-kappa-B essential modulator | -0.11098218 | -0.00438849 |
| A0A087WUY7 | PIAS2 | E3 SUMO-protein ligase PIAS2 (Fragment) | 0.052172343 | 0.336944 |
| A0A087WV25 | CEP43 | Centrosomal protein 43 | 0.150152365 | 0.110984 |
| A0A087WV34 | GLMP | Glycosylated lysosomal membrane protein | -2.283329407 | 2.85508 |
| A0A087WV57 | ZMYND8 | Protein kinase C-binding protein 1 | 0.660206715 | 0 |
| A0A087WV97 | SMAP2 | Stromal membrane-associated protein 2 | -0.020715872 | 0.00178623 |
| A0A087WVA1 | SELENOT | Selenoprotein T | -1.708867709 | 0.656702 |
| A0A087WVA8 | TEX2 | Testis-expressed protein 2 | -0.469551086 | 0 |
| A0A087WVF7 | ITSN2 | Intersectin-2 | 0.174467643 | -0.166084 |
| A0A087WVM4 | MTHFD1L | Formate--tetrahydrofolate ligase | -0.969640811 | 0.374406 |
| A0A087WVQ6 | CLTC | Clathrin heavy chain | 0.0937651 | -0.0652046 |
| A0A087WVW4 | TGM5 | Protein-glutamine gamma-glutamyltransferase 5 | -0.400981665 | 0.792664 |
| A0A087WW06 | TTC28 | Tetratricopeptide repeat protein 28 | 0.139877796 | -1.03176 |
| A0A087WW18 | ZBTB8OS | Protein archease (Fragment) | 0.419948498 | 0.530019 |
| A0A087WW40 | SH3GLB1 | Endophilin-B1 | -0.136642297 | 0.0294476 |
| A0A087WW43 | ITIH3 | Inter-alpha-trypsin inhibitor heavy chain H3 | -0.000975053 | 0.287693 |
| A0A087WW58 | SELENOI | Ethanolaminephosphotransferase 1 | -1.165000995 | 1.64403 |
| A0A087WW66 | PSMD1 | 26S proteasome non-ATPase regulatory subunit 1 | -0.268590927 | -0.0569757 |
| A0A087WW78 | GPR89B | Golgi pH regulator B (Fragment) | -0.741010984 | 0.239877 |
| A0A087WW88 | CHMP2B | Charged multivesicular body protein 2b | -0.854676882 | 0.0540657 |
| A0A087WWA3 | KIF1B | Kinesin-like protein KIF1B | -0.027329524 | 0.00882467 |
| A0A087WWF6 | POLD2 | DNA polymerase delta subunit 2 | -0.08591636 | 0.167885 |
| A0A087WWF8 | MUCL1 | Mucin-like protein 1 | -0.700689395 | 0.2363 |
| A0A087WWG1 | SSH3 | Protein phosphatase Slingshot homolog 3 (Fragment) | 0.148593108 | -0.0753965 |
| A0A087WWM4 | GMPR2 | GMP reductase | -0.147286654 | 0.134442 |
| A0A087WWP4 | RBM15 | RNA-binding protein 15 | -0.636658112 | 0.0773376 |
| A0A087WWS1 | THOC1 | THO complex subunit 1 | -1.083181381 | 0.219658 |
| A0A087WWZ2 | PCIF1 | mRNA (2'-O-methyladenosine-N(6)-)-methyltransferase (Fragment) | -0.006162087 | 0.565509 |
| A0A087WX29 | TARDBP | TAR DNA-binding protein 43 (Fragment) | -0.376271089 | 0.275042 |
| A0A087WXB0 | SCAMP1 | Secretory carrier-associated membrane protein | 0.954472701 | 1.93425 |
| A0A087WXC5 | NDUFA10 | NADH dehydrogenase [ubiquinone] 1 alpha subcomplex subunit 10, mitochondrial | -1.204637686 | -0.00772699 |
| A0A087WXK8 | FAM83D | Chromosome 20 open reading frame 129 | -0.830632607 | -0.0971769 |
| A0A087WXL6 | VPS11 | Vacuolar protein sorting-associated protein 11 homolog | -0.228493134 | 0.0906795 |
| A0A087WXM6 | RPL17 | 60S ribosomal protein L17 (Fragment) | -1.189886888 | -0.324256 |
| A0A087WXP2 | NFIB | CCAAT-box-binding transcription factor | -0.872614702 | 0.643415 |
| A0A087WXR7 | RNASEH2B | Ribonuclease H2 subunit B | 0.02015241 | -0.672039 |
| A0A087WXU0 | RMND1 | Required for meiotic nuclear division protein 1 homolog | 0 | 2.43897 |
| A0A087WXU3 | ESYT2 | Extended synaptotagmin-2 | -1.264080048 | -0.0614163 |
| A0A087WXV6 | OTUD5 | Deubiquitinating enzyme A | -1.427519242 | -1.17741 |
| A0A087WXX9 | PRKAA2 | 5'-AMP-activated protein kinase catalytic subunit alpha-2 | -0.678049485 | -0.400029 |
| A0A087WY55 | VTA1 | Chromosome 6 open reading frame 55, isoform CRA_b | -0.035910765 | -0.0586383 |

| | | | | |
|---|---|---|---|---|
| A0A087WY58 | NEK3 | Serine/threonine-protein kinase Nek3 | 0.469965537 | -1.84073 |
| A0A087WY67 | GSTT2 | Glutathione transferase | -0.285346508 | 0.0261154 |
| A0A087WY73 | | Uncharacterized protein (Fragment) | -0.016489903 | 2.42264 |
| A0A087WYD6 | TAP2 | Antigen peptide transporter 2 | 0.458831708 | -0.774385 |
| A0A087WYE2 | CERCAM | Inactive glycosyltransferase 25 family member 3 | -1.477282365 | 0 |
| A0A087WYF7 | APOOL | MICOS complex subunit | -1.539931774 | 1.00297 |
| A0A087WYI0 | INTS11 | Integrator complex subunit 11 | -6.4373E-06 | 1.38463 |
| A0A087WYN9 | DHX29 | ATP-dependent RNA helicase DHX29 | -0.796756506 | -0.135195 |
| A0A087WYR0 | SRP19 | Signal recognition particle 19 kDa protein | -0.222820361 | 0.354444 |
| A0A087WYV9 | SYNJ2BP-COX16 | Cytochrome c oxidase assembly protein COX16 homolog, mitochondrial | -1.953989108 | -0.0769297 |
| A0A087WZ13 | RAVER1 | Ribonucleoprotein PTB-binding 1 | -0.833704631 | 0.202619 |
| A0A087WZ32 | PLEKHA2 | Pleckstrin homology domain-containing family A member 2 | -0.377853711 | 0.144705 |
| A0A087WZM2 | RNASET2 | Ribonuclease T2 | 0.755164067 | 0.603254 |
| A0A087WZN1 | IDH3B | Isocitrate dehydrogenase [NAD] subunit, mitochondrial | -0.715347052 | -0.199702 |
| A0A087WZT0 | SAMD4B | Protein Smaug homolog 2 | 0.921076059 | 0 |
| A0A087WZT2 | METTL7B | Thiol S-methyltransferase METTL7B | -1.235914389 | -0.0788479 |
| A0A087WZT3 | BOLA2B | BolA-like protein 2 | 0.001007875 | 0.3769 |
| A0A087WZU2 | CDK16 | Cyclin-dependent kinase 16 | -0.047795296 | -0.214441 |
| A0A087WZV0 | PPIP5K2 | Inositol hexakisphosphate and diphosphoinositol-pentakisphosphate kinase | -0.459925254 | -0.977836 |
| A0A087WZX0 | BCL9L | B-cell CLL/lymphoma 9-like protein | 0.267091831 | -0.0169719 |
| A0A087WZZ4 | RHAG | Ammonium transporter Rh type A | 0.69483916 | 0.090517 |
| A0A087X011 | KIF3A | Kinesin-like protein | -0.072011391 | -0.0139745 |
| A0A087X085 | JADE2 | E3 ubiquitin-protein ligase Jade-2 | 0.427164237 | 0.855087 |
| A0A087X091 | WDR13 | WD repeat-containing protein 13 (Fragment) | 0.725055456 | -1.76573 |
| A0A087X0D5 | CTSH | Pro-cathepsin H | -0.493854602 | 0.2073 |
| A0A087X0H9 | RBM26 | RNA-binding protein 26 | 0.38983047 | 0 |
| A0A087X0L7 | OXA1L | Mitochondrial inner membrane protein OXA1L | 0 | -1.18039 |
| A0A087X0M4 | SLC4A1AP | Kanadaptin | 0.001696587 | -0.229487 |
| A0A087X0P4 | PBX1 | Pre-B-cell leukemia transcription factor 1 | 0.887897253 | 0 |
| A0A087X0Q1 | YTHDF3 | YTH domain-containing family protein 3 | -0.546956301 | 0.0180194 |
| A0A087X0Q9 | SELENOO | Selenoprotein O | -0.084424734 | 2.80179 |
| A0A087X0R6 | SNX12 | Sorting nexin-12 | 0.071377754 | -0.0618788 |
| A0A087X0V5 | OAS2 | 2'-5' oligoadenylate synthase (Fragment) | 0.506870826 | -0.892588 |
| A0A087X0W7 | ACOT2 | Acyl-coenzyme A thioesterase 2, mitochondrial | -0.144543568 | -0.165083 |
| A0A087X0W8 | RELA | Transcription factor p65 | 0.184152444 | 0.0269744 |
| A0A087X0W9 | OTUD6B | Ubiquitinyl hydrolase 1 | -0.111095349 | 0.128178 |
| A0A087X0X3 | HNRNPM | Heterogeneous nuclear ribonucleoprotein M | -0.598930041 | 0.456104 |
| A0A087X117 | NOMO1 | Nodal modulator 1 | -0.850920757 | 0.392573 |
| A0A087X142 | SEPTIN8 | Septin | -0.284705162 | 0.241198 |
| A0A087X1A8 | ATXN3 | Ubiquitinyl hydrolase 1 | 0.054138581 | 0.522262 |
| A0A087X1E4 | ARFIP2 | Arfaptin-2 | -0.381361802 | 0.11559 |
| A0A087X1G1 | NDUFA5 | Complex I subunit B13 | 0 | 2.46404 |
| A0A087X1G7 | SELENOF | Selenoprotein F | -0.801900387 | 1.42084 |
| A0A087X1K9 | LYPLA1 | Acyl-protein thioesterase 1 | 0.014998754 | 0.15941 |
| A0A087X1S3 | HUWE1 | E3 ubiquitin-protein ligase HUWE1 (Fragment) | -0.061305205 | 0.832392 |
| A0A087X1U5 | RPS6KC1 | Ribosomal protein S6 kinase delta-1 | 0.161055803 | -0.370489 |
| A0A087X1Z3 | PSME2 | Proteasome activator complex subunit 2 | 0.07737716 | -0.0315758 |
| A0A087X208 | AGRN | Agrin | -0.243490775 | 1.0068 |
| A0A087X211 | CIP2A | Protein CIP2A | 0.10761404 | -0.0969512 |
| A0A087X256 | WASHC4 | WASH complex subunit 4 | 0.064218601 | -0.125751 |
| A0A087X2B1 | RBFOX1 | RNA binding protein fox-1 homolog 1 | -0.90369757 | 0.0723286 |
| A0A087X2D5 | MRPL45 | 39S ribosomal protein L45, mitochondrial | -3.199347536 | 1.1427 |
| A0A087X2I1 | PSMC6 | 26S proteasome regulatory subunit 10B | -0.199107726 | -0.108284 |
| A0A096LNH6 | DOCK1 | Dedicator of cytokinesis protein 1 | -1.052754243 | 0.850535 |
| A0A096LNU0 | MCC | Colorectal mutant cancer protein | -0.269694249 | -0.38958 |
| A0A096LNV3 | RNF114 | E3 ubiquitin-protein ligase RNF114 (Fragment) | -0.041051229 | 0.156978 |
| A0A096LNY6 | ADSL | Adenylosuccinate lyase | -0.008254766 | 0.116495 |
| A0A096LP10 | ALG13 | N-acetylglucosaminyldiphosphodolichol N-acetylglucosaminyltransferase (Fragment) | -0.074733575 | -0.25813 |
| A0A096LP25 | AAK1 | AP2-associated protein kinase 1 | -0.08144323 | 0.0885849 |
| A0A096LPC5 | WASHC2C | WASH complex subunit 2C | 0.064089378 | -0.0404431 |
| A0A096LPI6 | | Utp12 domain-containing protein (Fragment) | -0.306719224 | 0.0627384 |
| A0A0A0MQR2 | RTF2 | Replication termination factor 2 | -0.32118686 | 0.217893 |
| A0A0A0MQS1 | PYCR3 | Pyrroline-5-carboxylate reductase | -0.546368281 | 0.0662921 |
| A0A0A0MQS2 | CLASRP | CLK4-associating serine/arginine-rich protein | -0.772666613 | -0.0124499 |
| A0A0A0MQU0 | WDR18 | WD repeat-containing protein 18 (Fragment) | -0.508470456 | 0.313597 |
| A0A0A0MQU4 | SCLY | Selenocysteine lyase | -0.133447011 | 0.204574 |
| A0A0A0MQW3 | SERPINB13 | Serpin B13 | -1.414925973 | 1.26543 |
| A0A0A0MQX1 | MYO10 | Unconventional myosin-X | -0.886165857 | -0.00760206 |

| | | | | |
|---|---|---|---|---|
| A0A0A0MQX7 | SPOCK1 | Testican-1 | -1.65791297 | -1.93117 |
| A0A0A0MQX8 | MBNL1 | Muscleblind-like protein 1 | -0.627746662 | -0.0657832 |
| A0A0A0MR09 | PTPN9 | Tyrosine-protein phosphatase non-receptor type 9 | -0.9772319 | -0.10622 |
| A0A0A0MR13 | GNAS | Guanine nucleotide-binding protein G(s) subunit alpha isoforms short | -0.773322344 | -0.0274728 |
| A0A0A0MR33 | GSTZ1 | Maleylacetoacetate isomerase | -0.843022823 | -2.30029 |
| A0A0A0MR87 | STIL | SCL-interrupting locus protein | 0.705617428 | 0 |
| A0A0A0MRA8 | EPB41L3 | Band 4.1-like protein 3 | -0.630853891 | 0.773839 |
| A0A0A0MRB7 | TCF3 | Transcription factor E2-alpha | -0.221108039 | 0.154222 |
| A0A0A0MRE3 | SYNE2 | Nesprin-2 | -0.122486671 | -2.11521 |
| A0A0A0MRE5 | ASAP1 | Arf-GAP with SH3 domain, ANK repeat and PH domain-containing protein 1 | -0.317826986 | 0.000973066 |
| A0A0A0MRF6 | AKAP9 | A-kinase anchor protein 9 | -1.218245745 | 0.183235 |
| A0A0A0MRI1 | GPALPP1 | GPALPP motifs-containing protein 1 | -0.239775896 | 0.1092 |
| A0A0A0MRJ3 | NAV1 | Neuron navigator 1 | -0.279892365 | -1.82351 |
| A0A0A0MRJ4 | TNNT2 | Troponin T, cardiac muscle | -1.924271901 | -1.02371 |
| A0A0A0MRJ7 | F5 | Coagulation factor V | -0.695640008 | 0.796898 |
| A0A0A0MRM8 | MYO6 | Unconventional myosin-6 | -0.040728569 | 0.0100451 |
| A0A0A0MRM9 | NOLC1 | Nucleolar and coiled-body phosphoprotein 1 (Fragment) | -0.092558463 | 0.0235402 |
| A0A0A0MRP6 | SMARCA1 | Probable global transcription activator SNF2L1 | -0.115679661 | 0.160296 |
| A0A0A0MRR3 | TAF8 | TBP-associated factor 8 | -0.321970145 | -0.222378 |
| A0A0A0MRR5 | UPRT | Uracil phosphoribosyltransferase homolog | -0.19849817 | 0.00493781 |
| A0A0A0MRT6 | ABI1 | Abl interactor 1 | -0.202387412 | -0.0078249 |
| A0A0A0MRV4 | PRXL2B | Prostamide/prostaglandin F synthase | 0.13276879 | 0.357323 |
| A0A0A0MRW7 | APTX | Aprataxin | -0.684162458 | -0.394493 |
| A0A0A0MRY0 | TBC1D22A | TBC1 domain family member 22A | -0.522838831 | -0.348006 |
| A0A0A0MRZ4 | TNIP1 | TNFAIP3-interacting protein 1 | -0.234172424 | 0.555402 |
| A0A0A0MS08 | IGHG1 | Immunoglobulin heavy constant gamma 1 (Fragment) | -0.804508924 | 0.703903 |
| A0A0A0MS45 | COG4 | Component of oligomeric Golgi complex 4 | -0.493591309 | 0.150525 |
| A0A0A0MS54 | PRKACB | cAMP-dependent protein kinase | -0.533418179 | -2.87535 |
| A0A0A0MS79 | NCKAP5 | Nck-associated protein 5 | -0.113964955 | 0.183605 |
| A0A0A0MS99 | ABCC1 | ABC-type glutathione-S-conjugate transporter | -0.64086175 | 0.428768 |
| A0A0A0MSA7 | EIF4G3 | Eukaryotic translation initiation factor 4 gamma 3 | -0.499869823 | 0.162625 |
| A0A0A0MSA9 | PVR | Poliovirus receptor | -1.885664622 | 0.401392 |
| A0A0A0MSB8 | EXOC7 | Exocyst complex component 7 | -0.164849361 | -0.107589 |
| A0A0A0MSI2 | PHC2 | Polyhomeotic-like protein 2 | -1.147357702 | 0 |
| A0A0A0MSI8 | EXOC5 | Exocyst complex component 5 | -0.289521297 | 0.225945 |
| A0A0A0MSJ9 | RASA2 | Ras GTPase-activating protein 2 | -0.736459414 | -0.00487137 |
| A0A0A0MSK4 | GPSM1 | G-protein-signaling modulator 1 | 0.016317685 | 0.0151288 |
| A0A0A0MSK6 | ARHGAP5 | Rho GTPase-activating protein 5 | -0.438446204 | 0.540979 |
| A0A0A0MSV9 | TAPBP | Tapasin | -1.211374124 | 0.0312405 |
| A0A0A0MSX9 | IARS1 | Isoleucyl-tRNA synthetase | -0.221354723 | -0.0631345 |
| A0A0A0MT00 | NFYC | Nuclear transcription factor Y subunit gamma (Fragment) | -0.188622157 | 0.394222 |
| A0A0A0MT26 | ATP1A3 | Sodium/potassium-transporting ATPase subunit alpha-3 | 0.406485478 | -0.559793 |
| A0A0A0MT32 | LIPA | Lysosomal acid lipase/cholesteryl ester hydrolase | -0.29576691 | -0.739429 |
| A0A0A0MT33 | SCAF8 | SR-related and CTD-associated factor 8 | -1.025114218 | 0.945335 |
| A0A0A0MT61 | ATG4A | Cysteine protease (Fragment) | -1.289428234 | 0 |
| A0A0A0MT83 | IVD | Butyryl-CoA dehydrogenase | -0.136692206 | 0.144611 |
| A0A0A0MTB8 | WDR36 | WD repeat-containing protein 36 | -0.667251428 | 0.297445 |
| A0A0A0MTB9 | RBP1 | Retinol-binding protein 1 | -0.008983612 | -1.08626 |
| A0A0A0MTH3 | ILK | 59 kDa serine/threonine-protein kinase | -0.125357151 | 0.142486 |
| A0A0A0MTH9 | BTAF1 | TATA-binding protein-associated factor 172 | -0.39234972 | -0.279072 |
| A0A0A0MTI1 | DHCR24 | Delta(24)-sterol reductase | -1.5243934 | 0.0197922 |
| A0A0A0MTI6 | ELOVL5 | Elongation of very long chain fatty acids protein | 0.985423009 | 0.45996 |
| A0A0A0MTJ9 | NCEH1 | Neutral cholesterol ester hydrolase 1 | -1.311672131 | 0.215332 |
| A0A0A0MTL5 | SKP2 | S-phase kinase-associated protein 2 | 0.185501258 | -0.42476 |
| A0A0A0MTN0 | CUL2 | Cullin-2 | -0.009460926 | -0.198027 |
| A0A0A0MTQ5 | ATP5MJ | ATP synthase subunit ATP5MJ, mitochondrial | 1.049842993 | 0 |
| A0A0A0MTS7 | TTN | Titin | -0.813963413 | 0.627768 |
| A0A0A6YYA0 | TMED7-TICAM2 | Protein TMED7-TICAM2 | -1.524989128 | 0.255944 |
| A0A0A6YYC0 | RPS6KA4 | Non-specific serine/threonine protein kinase (Fragment) | -0.663568179 | -1.09689 |
| A0A0A6YYH1 | C15orf38-AP3S2 | Arpin | 0.237775564 | 0.144145 |
| A0A0A6YYI3 | PPAN-P2RY11 | Protein PPAN-P2RY11 | -2.340735316 | -0.0451488 |
| A0A0A6YYL4 | CORO7-PAM16 | Coronin | 0.08419776 | 0.104529 |
| A0A0B4J1S8 | PI4KB | Phosphatidylinositol 4-kinase beta | -0.137079795 | -0.195016 |
| A0A0B4J1W0 | MED24 | Mediator complex subunit 24 | 1.243654092 | 2.70602 |
| A0A0B4J1Z1 | SRSF7 | Serine/arginine-rich-splicing factor 7 | 0.148883502 | -0.261359 |
| A0A0B4J294 | TRAPPC2L | Trafficking protein particle complex subunit 2-like protein | -0.355493148 | -0.216875 |
| A0A0B4J2A0 | | FGGY_N domain-containing protein | 0.159447352 | 0.178683 |

| | | | | |
|---|---|---|---|---|
| A0A0B4J2B4 | RPS15 | 40S ribosomal protein S15 | -0.357082685 | -0.291425 |
| A0A0B4J2B5 | IGHV3OR16-9 | Immunoglobulin heavy variable 3/OR16-9 (non-functional) (Fragment) | -0.437567631 | 0.122316 |
| A0A0B4J2E5 | LOC102724159 | Utp12 domain-containing protein | -0.887461901 | -0.421357 |
| A0A0C4DFM1 | TM9SF4 | Transmembrane 9 superfamily member | -1.098593076 | 0.51897 |
| A0A0C4DFM7 | TUT4 | Terminal uridylyltransferase 4 | -0.055154006 | -0.127174 |
| A0A0C4DFN3 | MGLL | Monoglyceride lipase | -0.482160012 | -0.481264 |
| A0A0C4DFP4 | NGLY1 | N-glycanase 1 (Fragment) | 0.156262636 | 0.00304953 |
| A0A0C4DFS0 | PBLD | MAWD binding protein, isoform CRA_d | -1.940946937 | 0 |
| A0A0C4DFT3 | DLG1 | Disks large homolog 1 | -0.289304256 | -0.00113106 |
| A0A0C4DFW4 | TEX30 | Testis-expressed protein 30 | -1.01219364 | 0 |
| A0A0C4DFX9 | NELFA | Negative elongation factor A | 0.028914849 | 0.180676 |
| A0A0C4DFZ1 | STX1A | Syntaxin-1A | 0.275588512 | -0.47111 |
| A0A0C4DFZ2 | ARSA | Arylsulfatase A | -0.090990543 | -0.387436 |
| A0A0C4DG17 | RPSA | 40S ribosomal protein SA | 0.301935355 | -0.0240949 |
| A0A0C4DG21 | ABI2 | Abl interactor 2 | -0.013321956 | 0.349447 |
| A0A0C4DG52 | WDR64 | WD repeat-containing protein 64 | -0.281274478 | -0.128035 |
| A0A0C4DG62 | ARL6IP4 | ADP-ribosylation factor-like protein 6-interacting protein 4 | 0.077749173 | -0.345328 |
| A0A0C4DG88 | POU2F1 | POU domain protein (Fragment) | -0.907456716 | -2.17762 |
| A0A0C4DG89 | DDX46 | RNA helicase | 0.013833523 | -0.233953 |
| A0A0C4DGA2 | ECI2 | Enoyl-CoA delta isomerase 2 | -2.726670822 | 0.673487 |
| A0A0C4DGB9 | PPA2 | Inorganic diphosphatase | -0.435080926 | -0.365373 |
| A0A0C4DGE4 | STX3 | Syntaxin-3 (Fragment) | -0.279767513 | -1.36334 |
| A0A0C4DGG1 | PACSIN3 | Protein kinase C and casein kinase substrate in neurons protein 3 (Fragment) | -0.455687364 | -0.0562668 |
| A0A0C4DGG9 | CHD4 | DNA helicase | -0.387119373 | 0.0160821 |
| A0A0C4DGH3 | NUMBL | Numb-like protein | -0.809782227 | 0 |
| A0A0C4DGN4 | ZG16B | Zymogen granule protein 16 homolog B | -0.72799778 | 0.940804 |
| A0A0C4DGN6 | GIT1 | ARF GTPase-activating protein GIT1 | -0.746515274 | -0.274253 |
| A0A0C4DGP0 | PITPNC1 | Cytoplasmic phosphatidylinositol transfer protein 1 | -0.423072179 | -0.974344 |
| A0A0C4DGQ5 | CAPNS1 | Calcium-activated neutral proteinase small subunit | 0.045531114 | 0.214169 |
| A0A0C4DGQ8 | DHRS7B | Dehydrogenase/reductase SDR family member 7B | -0.450500886 | -0.175846 |
| A0A0C4DGS1 | DDOST | Dolichyl-diphosphooligosaccharide--protein glycosyltransferase 48 kDa subunit | -1.123343309 | 0.235068 |
| A0A0C4DGU2 | MMACHC | Alkylcobalamin:glutathione S-alkyltransferase | -0.72639362 | -0.92766 |
| A0A0C4DGV4 | LAMTOR5 | Late endosomal/lysosomal adaptor and MAPK and MTOR activator 5 | 0.297338486 | 0.158644 |
| A0A0C4DGV9 | LIG4 | DNA ligase | -0.439773083 | -0.15423 |
| A0A0D9SEI3 | CDK11B | Cyclin-dependent kinase 11B | 0.013374329 | -0.108159 |
| A0A0D9SEI9 | SLC25A22 | Mitochondrial glutamate carrier 1 (Fragment) | -0.507161061 | 0.126847 |
| A0A0D9SEN1 | FAP | Prolyl endopeptidase FAP | -0.506542365 | -0.0442266 |
| A0A0D9SEU5 | RANBP10 | Ran-binding protein 10 | -0.41913112 | 0.114305 |
| A0A0D9SF44 | R3HCC1 | R3H and coiled-coil domain-containing protein 1 | 0.441555818 | -0.242363 |
| A0A0D9SF53 | DDX3X | RNA helicase | -0.54742233 | -0.0702807 |
| A0A0D9SFE4 | DNM1 | Dynamin GTPase | 0.628413121 | 0.850781 |
| A0A0D9SFG6 | CYTH2 | Cytohesin-2 | 0.169290304 | 0.131631 |
| A0A0D9SFL2 | PNKP | Bifunctional polynucleotide phosphatase/kinase | -0.179507573 | -0.0150541 |
| A0A0D9SFS3 | OGDH | Oxoglutarate dehydrogenase (succinyl-transferring) | -0.78195858 | 0.247515 |
| A0A0D9SG71 | UBE2A | Ubiquitin-conjugating enzyme E2 A | -1.080045541 | 0.248503 |
| A0A0D9SG72 | STXBP1 | Syntaxin-binding protein 1 | -0.039522966 | -0.351424 |
| A0A0D9SGE8 | PHF6 | PHD finger protein 6 | -0.98258098 | -0.191132 |
| A0A0D9SGJ6 | SYNJ1 | Phosphoinositide 5-phosphatase | 1.79005448 | -2.78691 |
| A0A0G2JH68 | DIAPH1 | Protein diaphanous homolog 1 | 0.091011286 | -0.0601454 |
| A0A0G2JIC6 | MRPS18B | 28S ribosomal protein S18b, mitochondrial | -2.23894453 | 2.37863 |
| A0A0G2JIW1 | HSPA1B | Heat shock 70 kDa protein 1B | 0.184126059 | -0.126042 |
| A0A0G2JJA7 | PSMB9 | Proteasome subunit beta | 0.932547967 | -0.792077 |
| A0A0G2JJD3 | ABHD16A | Alpha/beta hydrolase domain-containing protein 16A | -1.380602439 | -0.592119 |
| A0A0G2JK44 | BRD2 | Bromodomain-containing protein 2 | 0.120555083 | -0.146921 |
| A0A0G2JLB3 | GBA | Glucosylceramidase | -0.206059933 | -0.0990442 |
| A0A0G2JMB2 | IGHA2 | Immunoglobulin heavy constant alpha 2 (Fragment) | -0.374125004 | 0.274966 |
| A0A0G2JNB4 | PRB3 | Basic salivary proline-rich protein 3 | -0.294577678 | -1.15307 |
| A0A0G2JNH5 | RAD17 | Cell cycle checkpoint protein RAD17 | 0.292937001 | 0 |
| A0A0G2JNZ2 | SCRIB | Protein scribble homolog | -0.189304193 | 0.290589 |
| A0A0G2JPA8 | SERPINF2 | Alpha-2-antiplasmin | -0.773673058 | 0.121918 |
| A0A0G2JPR0 | C4A | Complement C4-A | -0.088579337 | 0.378556 |
| A0A0G2JQ41 | ABR | Active breakpoint cluster region-related protein (Fragment) | 0.000138124 | -0.261422 |
| A0A0G2JQ92 | TAMM41 | CDP-diacylglycerol synthase (Fragment) | 0 | 3.1168 |
| A0A0G2JQT1 | CYFIP1 | Cytoplasmic FMR1-interacting protein 1 (Fragment) | -0.630753756 | 0 |
| A0A0J9YVP6 | PUF60 | 60 kDa poly(U)-binding-splicing factor (Fragment) | 0.093688806 | 0.00494607 |
| A0A0J9YWB6 | CTDP1 | RNA polymerase II subunit A C-terminal domain phosphatase (Fragment) | 0.200038671 | -0.169333 |

| | | | | |
|---|---|---|---|---|
| A0A0J9YXF2 | PON2 | Paraoxonase | -1.747332811 | -0.339899 |
| A0A0J9YXM6 | WDR81 | WD repeat-containing protein 81 | 0.763492107 | 1.65675 |
| A0A0J9YXU7 | PON2 | Paraoxonase (Fragment) | -1.528036515 | -0.407542 |
| A0A0J9YYC8 | PRSS2 | Trypsin-2 | 0.109932502 | -0.706065 |
| A0A0R4J2E8 | MATR3 | Matrin-3 | -0.653321822 | 0.324557 |
| A0A0R4J2F2 | CLDND1 | Claudin domain-containing protein 1 | -0.433541377 | 0 |
| A0A0R4J2F3 | POLR1D | DNA-directed RNA polymerase I subunit D | -0.177695354 | 0.573783 |
| A0A0R4J2F6 | IFRD2 | Interferon-related developmental regulator 2 | -0.089537144 | -1.84697 |
| A0A0S2Z5C6 | FPGT | Fucose-1-phosphate guanylyltransferase (Fragment) | -0.289082766 | -1.31934 |
| A0A0S2Z5D0 | CLN6 | Ceroid-lipofuscinosis neuronal 6 late infantile variant isoform 2 | -0.546697696 | 0 |
| A0A0U1RQC9 | TP53 | Cellular tumor antigen p53 | -0.212285757 | 0.0691353 |
| A0A0U1RQD2 | CCDC149 | Coiled-coil domain-containing protein 149 | -0.139573971 | 0.492698 |
| A0A0U1RQH9 | CSNK1G1 | Casein kinase I isoform gamma-1 | -2.192094723 | 0.440328 |
| A0A0U1RQM0 | | PCNA-associated factor | -0.754867792 | -0.41855 |
| A0A0U1RQP1 | DNM1 | Dynamin GTPase | -0.137796481 | 0.162611 |
| A0A0U1RQQ9 | SCYL2 | SCY1-like protein 2 | -0.651091655 | 0.246158 |
| A0A0U1RR06 | WDR45 | WD repeat domain phosphoinositide-interacting protein 4 | -0.473133485 | -0.00637627 |
| A0A0U1RRB6 | EXOC6B | Exocyst complex component | 0.191304207 | -0.230323 |
| A0A0U1RRM1 | GATAD2B | Transcriptional repressor p66-beta | -0.506682634 | -0.222251 |
| A0A0U1RRM4 | PTBP1 | Polypyrimidine tract-binding protein 1 | -0.508172115 | 0.303026 |
| A0A0U1RRM6 | ENAH | Protein enabled homolog | -0.096299887 | 0.0238123 |
| A0A140T913 | HLA-A | HLA class I histocompatibility antigen, A alpha chain | -1.129534403 | -0.0913329 |
| A0A140T921 | HLA-C | HLA class I histocompatibility antigen, C alpha chain | -0.741278648 | 0.581271 |
| A0A140T997 | HLA-B | HLA class I histocompatibility antigen, B alpha chain (Fragment) | -0.86247619 | 0.0234321 |
| A0A140T9M0 | HLA-C | HLA class I histocompatibility antigen, C alpha chain | -0.892374754 | 0.12969 |
| A0A140T9T7 | TAP1 | Antigen peptide transporter 1 | -1.786468506 | -0.245465 |
| A0A140TA73 | SNTB2 | Beta-2-syntrophin (Fragment) | 0.16106534 | -1.41592 |
| A0A163QEF0 | SLC1A2 | Amino acid transporter | 0.443959475 | -0.562873 |
| A0A182DWF2 | TXNRD2 | Thioredoxin-disulfide reductase | 0.059250673 | 0.679189 |
| A0A1B0GTB0 | ATP6AP2 | ATPase H(+)-transporting lysosomal accessory protein 2 (Fragment) | 0.896668434 | 0 |
| A0A1B0GTB8 | CPT2 | Carnitine O-palmitoyltransferase 2, mitochondrial | -1.933842262 | -1.03649 |
| A0A1B0GTC3 | FTO | Alpha-ketoglutarate-dependent dioxygenase FTO | -0.0962708 | 0.120722 |
| A0A1B0GTF3 | HSPA12A | Heat shock 70 kDa protein 12A | 0.254926999 | -0.416538 |
| A0A1B0GTL5 | RAB11FIP5 | Rab11 family-interacting protein 5 | -0.614716609 | -0.0922502 |
| A0A1B0GTM3 | ASAH1 | Ceramidase | -0.151321173 | 0.0231183 |
| A0A1B0GTU9 | CHD2 | DNA helicase (Fragment) | -0.587362925 | 2.25195 |
| A0A1B0GU86 | ACY1 | N-acyl-L-amino-acid amidohydrolase | -0.136292775 | -0.160763 |
| A0A1B0GUA3 | KIFBP | KIF-binding protein | -0.024014314 | -0.099549 |
| A0A1B0GUS7 | UNC13B | Protein unc-13 homolog B | 0.11802268 | -0.49847 |
| A0A1B0GV05 | RAP1GAP2 | Rap1 GTPase-activating protein 2 | -0.294118325 | 0.107783 |
| A0A1B0GV47 | KIF21A | Kinesin-like protein KIF21A (Fragment) | 0.055009524 | -0.111831 |
| A0A1B0GV70 | TEPSIN | AP-4 complex accessory subunit Tepsin | 0.457469702 | 0 |
| A0A1B0GVD5 | CTSD | Cathepsin D | -0.026318073 | 0.084439 |
| A0A1B0GVE3 | CAPN15 | Calpain-15 (Fragment) | -3.029107014 | 1.15812 |
| A0A1B0GVI3 | KRT10 | Keratin, type I cytoskeletal 10 | -0.230047862 | -0.096529 |
| A0A1B0GVS8 | KIF16B | Kinesin-like protein KIF16B (Fragment) | -0.844676097 | -0.289838 |
| A0A1B0GW05 | DPY19L1 | Probable C-mannosyltransferase DPY19L1 | -0.322944482 | 0 |
| A0A1B0GWG0 | EPB41 | Band 4.1 (Fragment) | -0.565502644 | -0.821943 |
| A0A1B0GWI2 | CLN5 | Ceroid-lipofuscinosis neuronal protein 5 (Fragment) | -0.166645845 | -0.128074 |
| A0A1B0GX65 | UBE2E3 | Ubiquitin-conjugating enzyme E2 E3 (Fragment) | -0.056651433 | 0.377604 |
| A0A1B0GXE7 | ORMDL1 | ORM1-like protein 1 | -3.005705198 | -0.689636 |
| A0A1B0GXL9 | TFEB | Transcription factor EB (Fragment) | 0.416298548 | 0 |
| A0A1C7CYX9 | DPYSL2 | Dihydropyrimidinase-related protein 2 | 0.043274244 | -0.0537491 |
| A0A1P0AYU5 | SFXN3 | Sidoreflexin | -1.447053274 | 0.59556 |
| A0A1W2PNP0 | PIGT | GPI transamidase component PIG-T (Fragment) | 1.237567902 | -0.0694272 |
| A0A1W2PNR9 | IER3IP1 | Immediate early response 3-interacting protein 1 | -0.538687189 | 0.893815 |
| A0A1W2PNV4 | | Uncharacterized protein | -0.437310378 | -0.329106 |
| A0A1W2PNV7 | STRADA | STE20-related kinase adapter protein alpha | 0.210503658 | -0.419786 |
| A0A1W2PNY5 | ELP4 | Elongator complex protein 4 | 0.230054935 | -2.25846 |
| A0A1W2PPG1 | GOSR2 | Golgi SNAP receptor complex member 2 | -1.341985146 | 0.577589 |
| A0A1W2PPH1 | ME2 | Malic enzyme | -0.65754056 | 0.0810328 |
| A0A1W2PPS8 | ERGIC3 | Endoplasmic reticulum-Golgi intermediate compartment protein 3 | -0.798748891 | 0.34849 |
| A0A1W2PPU0 | PRKCH | Protein kinase C eta type | 0.195230484 | -1.1004 |
| A0A1W2PPZ5 | TCEA1 | Transcription elongation factor | -0.042190552 | 0.0856921 |
| A0A1W2PQS6 | RPS10-NUDT3 | RPS10-NUDT3 readthrough | -0.252281507 | -0.237236 |
| A0A1W2PR09 | NECAP1 | Adaptin ear-binding coat-associated protein 1 | -0.444391251 | -0.737844 |
| A0A1W2PR17 | PFKFB3 | 6-phosphofructo-2-kinase/fructose-2,6-bisphosphatase 3 | 0.149152358 | -1.2986 |
| A0A1W2PR85 | SYNJ2 | Phosphoinositide 5-phosphatase | -0.464718262 | -0.597916 |
| A0A1W2PRB8 | | Cofactor required for Sp1 transcriptional activation subunit 6 | -0.539491971 | -1.45433 |
| A0A1W2PRS1 | SCARB2 | Lysosome membrane protein 2 (Fragment) | -1.155037244 | 0.0739269 |

| | | | | |
|---|---|---|---|---|
| A0A1W2PRV5 | SMN2 | Survival motor neuron protein | -1.485134919 | -1.90394 |
| A0A1W2PS05 | PIK3C3 | Phosphatidylinositol 3-kinase catalytic subunit type 3 | -0.035416603 | 0.18234 |
| A0A1W2PS85 | BIVM-ERCC5 | BIVM-ERCC5 readthrough | -0.17678078 | -0.147062 |
| A0A1X7SBZ2 | DDX17 | RNA helicase | -0.482069016 | 0.242263 |
| A0A286YEY1 | IGHA1 | Immunoglobulin heavy constant alpha 1 (Fragment) | -0.446785371 | 0.449174 |
| A0A286YEY4 | IGHG2 | Immunoglobulin heavy constant gamma 2 (Fragment) | -1.366241137 | -0.0323121 |
| A0A286YF17 | MTHFR | Methylenetetrahydrofolate reductase | -0.234710614 | -0.119131 |
| A0A286YF22 | PHGDH | D-3-phosphoglycerate dehydrogenase | -0.08705012 | 0.0673447 |
| A0A286YFF7 | PPT1 | Palmitoyl-protein hydrolase 1 | -0.003801584 | 0.30945 |
| A0A2Q3DP09 | ARMC9 | LisH domain-containing protein ARMC9 | -0.177786032 | -0.214539 |
| A0A2Q3DQE3 | CAMK2G | Calcium/calmodulin-dependent protein kinase | -0.252917131 | -0.654454 |
| A0A2R8Y2Y1 | PNISR | Arginine/serine-rich protein PNISR | -0.126242479 | -0.673424 |
| A0A2R8Y359 | SLC33A1 | Acetyl-coenzyme A transporter 1 | -0.486013571 | 0.450691 |
| A0A2R8Y3N3 | HARS2 | Histidine--tRNA ligase | -0.173091809 | -0.14568 |
| A0A2R8Y432 | CEP78 | Centrosomal protein of 78 kDa (Fragment) | -0.697458426 | -0.0458625 |
| A0A2R8Y455 | | COMM domain-containing protein | 0.317520459 | -0.20339 |
| A0A2R8Y473 | ABCB7 | Iron-sulfur clusters transporter ABCB7, mitochondrial | -0.613199353 | -0.0934879 |
| A0A2R8Y484 | CD47 | Integrin-associated protein (Fragment) | -1.043583155 | 0.280382 |
| A0A2R8Y4P4 | SMARCA4 | Transcription activator BRG1 | -0.713326931 | 0.298772 |
| A0A2R8Y4T1 | TNS1 | Tensin-1 | -0.067682346 | 0.129903 |
| A0A2R8Y4T4 | SMARCE1 | SWI/SNF-related matrix-associated actin-dependent regulator of chromatin subfamily E member 1 | -0.886520068 | -0.121715 |
| A0A2R8Y4W2 | EPS8 | Epidermal growth factor receptor kinase substrate 8 | -0.208698988 | -0.169505 |
| A0A2R8Y4W8 | CCDC88A | Girdin | -0.743789991 | -0.667505 |
| A0A2R8Y543 | CTNNB1 | Catenin beta-1 | -0.701663574 | 0.160857 |
| A0A2R8Y566 | RELCH | RAB11-binding protein RELCH | 0.134781202 | -0.121016 |
| A0A2R8Y575 | PLEKHM2 | Pleckstrin homology domain-containing family M member 2 | 0.633740664 | 0 |
| A0A2R8Y5A6 | ATXN2 | Ataxin-2 | -1.1228532 | 0.151061 |
| A0A2R8Y5C4 | TNKS1BP1 | 182 kDa tankyrase-1-binding protein (Fragment) | -0.247424682 | -0.0970303 |
| A0A2R8Y5F1 | TSC2 | Tuberin | -0.661531528 | -0.413637 |
| A0A2R8Y5K2 | C9orf72 | Guanine nucleotide exchange C9orf72 | 0.295537233 | -0.00179927 |
| A0A2R8Y5L0 | TSEN15 | tRNA-splicing endonuclease subunit Sen15 | -0.779948195 | 1.68184 |
| A0A2R8Y5N2 | TSC1 | Hamartin | -0.012381156 | -0.384916 |
| A0A2R8Y635 | ATP13A3 | Polyamine-transporting ATPase 13A3 | -0.604442437 | 0 |
| A0A2R8Y653 | UCK2 | Uridine/cytidine kinase | 0.180538177 | 0.130789 |
| A0A2R8Y661 | RAPGEF2 | Cyclic nucleotide ras GEF | -0.110433896 | -0.158565 |
| A0A2R8Y6I6 | DYRK1A | Dual-specificity kinase | 0.459038496 | -2.46015 |
| A0A2R8Y6W8 | ERCC3 | DNA excision repair protein ERCC-3 | 4.848046819 | -1.68387 |
| A0A2R8Y6Y7 | SUCLA2 | Succinate--CoA ligase [ADP-forming] subunit beta, mitochondrial | -0.625681241 | -0.0205339 |
| A0A2R8Y706 | DYNC1H1 | Cytoplasmic dynein 1 heavy chain 1 | -0.215428352 | -0.00727145 |
| A0A2R8Y7J6 | CDIN1 | CDAN1-interacting nuclease 1 | 1.200969299 | 0.676229 |
| A0A2R8Y7R9 | PPFIA1 | Liprin-alpha-1 (Fragment) | -0.754841169 | -0.425633 |
| A0A2R8Y7X9 | | GLOBIN domain-containing protein | 1.056633472 | -0.642025 |
| A0A2R8Y811 | RPS14 | 40S ribosomal protein S14 (Fragment) | -0.362508774 | -0.280495 |
| A0A2R8Y840 | CHD8 | DNA helicase (Fragment) | -0.030115366 | 0.00413577 |
| A0A2R8Y849 | RPS24 | 40S ribosomal protein S24 | -1.035981099 | -0.339758 |
| A0A2R8Y880 | RBBP6 | E3 ubiquitin-protein ligase RBBP6 (Fragment) | -0.078704039 | 0.806212 |
| A0A2R8Y891 | PFKM | ATP-dependent 6-phosphofructokinase | -0.345170498 | 0.110824 |
| A0A2R8Y8E0 | BRAF | Non-specific serine/threonine protein kinase | -0.084996541 | -0.19245 |
| A0A2R8YCL1 | GRB10 | Growth factor receptor-bound protein 10 | 0.663676023 | 0 |
| A0A2R8YD45 | TPP1 | Tripeptidyl aminopeptidase | -0.208160718 | -0.0820115 |
| A0A2R8YD49 | OSBPL2 | Oxysterol-binding protein | -0.560479244 | -0.303844 |
| A0A2R8YD50 | HSD17B4 | Peroxisomal multifunctional enzyme type 2 | -0.918260177 | 0.353228 |
| A0A2R8YD58 | CSNK2A1 | Non-specific serine/threonine protein kinase | 0.083752473 | -0.153643 |
| A0A2R8YDE6 | HDAC6 | Histone deacetylase 6 | -1.146143834 | -0.636228 |
| A0A2R8YE10 | VPS45 | Vacuolar protein sorting-associated protein 45 | -0.45977815 | 0.014185 |
| A0A2R8YE77 | CASK | Calcium/calmodulin-dependent serine protein kinase (MAGUK family), isoform CRA_c | -0.093007485 | 0.105361 |
| A0A2R8YEM2 | CCNH | Cyclin-H | -0.232981761 | -0.144959 |
| A0A2R8YF04 | OTOGL | Otogelin-like protein | -1.586518645 | -1.97682 |
| A0A2R8YF87 | VPS33A | Vacuolar protein sorting-associated protein 33A | -0.082375447 | -0.0133406 |
| A0A2R8YFB8 | RAB15 | Ras-related protein Rab-15 | -1.537451903 | 0.428017 |
| A0A2R8YFH5 | SEC23B | Protein transport protein SEC23 | -0.039834261 | -0.0417951 |
| A0A2R8YFX5 | NAV3 | Neuron navigator 3 | 0.088916143 | 0 |
| A0A2R8YG42 | ARHGEF7 | Rho guanine nucleotide exchange factor 7 | -0.57875371 | -0.625239 |
| A0A2R8YGH5 | AP1S1 | AP complex subunit sigma | -0.101788918 | -0.761159 |
| A0A2R8YH90 | TPM4 | Tropomyosin alpha-4 chain (Fragment) | 0.39765954 | 0.825246 |
| A0A2U3TZH3 | EEF1A2 | Elongation factor 1-alpha | -0.816382011 | -2.15066 |
| A0A2U3TZL5 | CD59 | CD59 glycoprotein (Fragment) | -0.93890206 | 1.36668 |
| A0A2U3TZV8 | PLCH1 | Phosphoinositide phospholipase C | -0.064586242 | -0.0070378 |
| A0A2Y9D025 | NDUFA6 | Complex I-B14 | -1.389151812 | 0.381576 |

| | | | | |
|---|---|---|---|---|
| A0A384DVK4 | R3HCC1L | Coiled-coil domain-containing protein R3HCC1L | 0.184639374 | 0 |
| A0A384DVL6 | NCOR2 | Nuclear receptor corepressor 2 | -0.223675489 | -1.37608 |
| A0A384DVU0 | PNPLA6 | Patatin-like phospholipase domain-containing protein 6 | -0.309776306 | 0 |
| A0A3B3IRI9 | HDAC8 | Histone deacetylase | 0.473226468 | -0.399934 |
| A0A3B3IRJ9 | INPP5F | Phosphatidylinositide phosphatase SAC2 | -0.221822103 | -0.0131381 |
| A0A3B3IRL2 | CREG1 | Cellular repressor of E1A-stimulated genes 1, isoform CRA_a | 1.92100215 | 0 |
| A0A3B3IRL5 | IMPDH1 | Inosine-5'-monophosphate dehydrogenase (Fragment) | -0.045498292 | -0.28015 |
| A0A3B3IRM4 | XYLB | Xylulose kinase | -0.201040824 | -0.577525 |
| A0A3B3IRR6 | C8orf33 | UPF0488 protein C8orf33 | -0.368507067 | -0.238251 |
| A0A3B3IRT8 | SSR1 | Translocon-associated protein subunit alpha | -0.984954516 | 0.279088 |
| A0A3B3IS71 | RB1 | Retinoblastoma-associated protein | -0.891359965 | -0.0022405 |
| A0A3B3ISF9 | ECE1 | Endothelin-converting enzyme 1 | -2.391077201 | 0.295144 |
| A0A3B3ISG5 | IDE | Insulin-degrading enzyme | 0.004133224 | 0.0769971 |
| A0A3B3ISP1 | ANKS1A | Ankyrin repeat and SAM domain-containing protein 1A | -0.14716061 | -0.0656052 |
| A0A3B3ISQ4 | EPS8L2 | Epidermal growth factor receptor kinase substrate 8-like protein 2 | 0.209012747 | 0.0145191 |
| A0A3B3ISS6 | GPNMB | Transmembrane glycoprotein NMB | -0.186266661 | 0.500066 |
| A0A3B3ISV4 | VKORC1L1 | Vitamin-K-epoxide reductase (warfarin-sensitive) | -0.411390543 | -1.11125 |
| A0A3B3ISV5 | | Uncharacterized protein | -0.675732533 | -0.0567252 |
| A0A3B3ISW7 | VPS36 | Vacuolar protein-sorting-associated protein 36 | 0.220377048 | -0.803954 |
| A0A3B3ITD5 | PARP14 | Protein mono-ADP-ribosyltransferase PARP14 | -0.422524254 | 0 |
| A0A3B3ITG3 | DAB2IP | Disabled homolog 2-interacting protein | 0.300330559 | -3.46139 |
| A0A3B3ITH3 | IMPACT | Protein IMPACT | -0.360986233 | 0.126533 |
| A0A3B3ITP9 | ITM2B | Integral membrane protein 2 | 0 | -0.541998 |
| A0A3B3ITT1 | RAB12 | Ras-related protein Rab-12 | -0.922577778 | -0.067502 |
| A0A3B3ITT5 | RPL29 | 60S ribosomal protein L29 | -0.947553078 | 0.401097 |
| A0A3B3ITW1 | GSK3B | Glycogen synthase kinase-3 beta | -0.566308101 | -0.260092 |
| A0A3B3ITW3 | ABCD3 | ATP-binding cassette sub-family D member 3 | -1.825798591 | 0.258172 |
| A0A3B3ITX4 | | Uncharacterized protein | -1.254550219 | 0.482698 |
| A0A3B3ITZ9 | THRAP3 | Thyroid hormone receptor-associated protein 3 | -0.119434039 | -0.0501359 |
| A0A3B3IU55 | STON2 | Stonin-2 | -0.658317804 | 0.266546 |
| A0A3B3IU69 | EML1 | Echinoderm microtubule-associated protein-like 1 | -0.19839565 | -0.072854 |
| A0A3B3IUA2 | SNU13 | Ribonucleoprotein | -0.394929727 | -0.358852 |
| A0A3B3IUB1 | EIF2B5 | eIF-2B GDP-GTP exchange factor subunit epsilon | 0.072898229 | 0.149586 |
| A0A3B3IUB5 | HM13 | Minor histocompatibility antigen H13 | -1.879142364 | 0.179029 |
| A0A3B3IUC4 | GLA | Alpha-galactosidase | -0.205609719 | -0.192701 |
| A0A3B3IUD5 | QPCT | Glutaminyl-peptide cyclotransferase (Fragment) | -0.779130061 | 1.03194 |
| A0A3B3IUD7 | PDCD11 | Programmed cell death 11, isoform CRA_a | -1.749694268 | 0.353621 |
| A0A3F2YNY7 | HMBS | Hydroxymethylbilane synthase | 0.182177623 | 0.00776641 |
| A0A494BZT8 | TACC3 | Transforming acidic coiled-coil-containing protein 3 | 0.05270497 | 0.336199 |
| A0A494BZV2 | MPRIP | Myosin phosphatase Rho-interacting protein | -0.276799599 | 0.178967 |
| A0A494BZY7 | EEF1AKMT2 | EEF1A lysine methyltransferase 2 | 0.261501074 | 0.368527 |
| A0A494C039 | HYOU1 | Hypoxia up-regulated protein 1 | 0.034801006 | 0.0259075 |
| A0A494C050 | NKAP | NF-kappa-B-activating protein (Fragment) | -1.321429968 | -0.132108 |
| A0A494C066 | KYAT1 | Kynurenine--oxoglutarate transaminase 1 | 0.027772427 | 0.839658 |
| A0A494C068 | PRKAG2 | 5'-AMP-activated protein kinase subunit gamma-2 | 0.009427547 | -0.277848 |
| A0A494C0B4 | RAD23A | UV excision repair protein RAD23 homolog A | 0.08808279 | 0.409489 |
| A0A494C0C9 | CHN1 | N-chimaerin | -0.351368348 | -0.753355 |
| A0A494C0J7 | | TGc domain-containing protein | -0.044536511 | 0.107985 |
| A0A494C0L6 | TYMP | Thymidine phosphorylase (Fragment) | 1.027788877 | -2.68554 |
| A0A494C0N6 | ACOT8 | Acyl-coenzyme A thioesterase 8 | -2.649463733 | -1.63442 |
| A0A494C0R8 | CLUH | Clustered mitochondria protein homolog | -0.023900191 | -0.0842272 |
| A0A494C0W0 | PARN | Poly(A)-specific ribonuclease PARN | -0.073507071 | -0.0297028 |
| A0A494C0Z5 | RAB33B | Ras-related protein Rab-33B | -1.008479675 | -0.132648 |
| A0A494C105 | MFSD1 | Major facilitator superfamily domain-containing protein 1 | -1.148098469 | 0.1015 |
| A0A494C108 | KRI1 | Protein KRI1 homolog | -0.09756279 | -0.163363 |
| A0A494C164 | STAT2 | Signal transducer and activator of transcription | 0.733680566 | -2.86929 |
| A0A494C1A0 | ACTN2 | Alpha-actinin-2 (Fragment) | -0.144125382 | -0.349878 |
| A0A494C1J1 | SPECC1L | Cytospin-A | -1.393604517 | 0.181557 |
| A0A494C1K3 | GTF2I | General transcription factor II-I | -0.684568246 | 0.242561 |
| A0A494C1L1 | LBR | 3-beta-hydroxysterol Delta (14)-reductase | -2.235568364 | 0.0943448 |
| A0A494C1L5 | LRBA | Lipopolysaccharide-responsive and beige-like anchor protein | 0.030086199 | -0.0269245 |
| A0A494C1N0 | FKBP2 | Peptidylprolyl isomerase | -0.368550777 | -0.041378 |
| A0A494C1S7 | GTF3C3 | General transcription factor 3C polypeptide 3 | -0.420679013 | -0.0211163 |
| A0A499FI31 | SART3 | Squamous cell carcinoma antigen recognized by T-cells 3 | -0.166291952 | -0.01756 |
| A0A499FIY8 | MYADM | Myeloid-associated differentiation marker (Fragment) | -1.292617718 | -0.330797 |
| A0A499FIZ0 | WDR26 | WD repeat-containing protein 26 | 0.032996257 | -0.136838 |
| A0A499FJK2 | TGFB1 | Transforming growth factor beta | -0.722471078 | -0.145452 |
| A0A4W8VX11 | PCM1 | Pericentriolar material 1 protein | -0.132846038 | 0.0135393 |
| A0A590TQL1 | ACD | Adrenocortical dysplasia homolog (Mouse), isoform CRA_a | -0.531683683 | 0.229405 |
| A0A590UJ43 | CSNK1A1 | Casein kinase I isoform alpha | -0.551783244 | 0.116558 |
| A0A590UJ73 | TSEN34 | tRNA-splicing endonuclease subunit Sen34 | -0.042851845 | 0.0328604 |

| | | | | |
|---|---|---|---|---|
| A0A590UJF2 | POT1 | Protection of telomeres protein 1 | -0.872900724 | 0 |
| A0A590UJK4 | SRSF6 | Serine/arginine-rich-splicing factor 6 | 0.08826081 | 0.0426025 |
| A0A590UJL0 | STS | Steryl-sulfatase | 0 | -1.2484 |
| A0A5F9UK94 | AMPD2 | AMP deaminase | 0.245963176 | 0.00146612 |
| A0A5F9UP49 | SDF4 | 45 kDa calcium-binding protein | 0.330621401 | -0.0963008 |
| A0A5F9UY30 | TGOLN2 | Trans-Golgi network integral membrane protein 2 | -1.780380805 | -0.252355 |
| A0A5F9YFS9 | SPCS1 | Microsomal signal peptidase 12 kDa subunit | -1.347202778 | 0.86855 |
| A0A5F9ZGX3 | ANK2 | Ankyrin-2 | -0.122370958 | 0.491159 |
| A0A5F9ZH07 | JAK1 | Tyrosine-protein kinase (Fragment) | -1.291546663 | 2.73512 |
| A0A5F9ZH21 | CAMK2A | Calcium/calmodulin-dependent protein kinase | 0.481283903 | -0.224794 |
| A0A5F9ZHB6 | PRIM1 | DNA primase | -0.316450516 | -0.410014 |
| A0A5F9ZHI5 | ALDH3A2 | Aldehyde dehydrogenase | -1.18126448 | 0.158124 |
| A0A5F9ZHN7 | SACM1L | Phosphatidylinositol-3-phosphatase SAC1 | -1.67641902 | 0.19669 |
| A0A5F9ZHQ7 | HAUS3 | HAUS augmin-like complex subunit 3 | 0.820240895 | 0.511638 |
| A0A5F9ZHR7 | ACP2 | Lysosomal acid phosphatase | -0.999616861 | -0.33643 |
| A0A5F9ZHU7 | PRIM2 | DNA primase large subunit | -0.311613083 | -0.222644 |
| A0A5F9ZHY7 | BRI3BP | BRI3-binding protein | -0.726362387 | -0.727042 |
| A0A5F9ZI21 | ARHGEF2 | Guanine nucleotide exchange factor H1 | -0.788025777 | -0.28948 |
| A0A5F9ZI63 | EED | Polycomb protein EED | -0.424323559 | -0.172429 |
| A0A5H1ZRP4 | NELFB | Negative elongation factor B | -0.269022385 | -0.0263224 |
| A0A5K1VW28 | ETNK1 | Ethanolamine kinase 1 | -0.187146346 | -0.054752 |
| A0A669KAU7 | CUL7 | Cullin-7 (Fragment) | -0.100848754 | -0.622897 |
| A0A669KAW1 | FUT8 | Alpha-(1,6)-fucosyltransferase | -2.502719402 | -0.95765 |
| A0A669KAX3 | WFS1 | Wolframin | -1.947698116 | 0.0179307 |
| A0A669KAX4 | CUL4B | Cullin-4B | 0.052714666 | 0.0586748 |
| A0A669KAX9 | | Ribosomal_L2_C domain-containing protein (Fragment) | -1.170322736 | 0.873222 |
| A0A669KAY5 | DUS4L-BCAP29 | DUS4L-BCAP29 readthrough | -1.525707523 | 1.25311 |
| A0A669KB03 | MLH1 | DNA mismatch repair protein Mlh1 | -1.062190135 | 0.315719 |
| A0A669KB05 | CTNND1 | Catenin delta-1 | -1.040493727 | 0.484167 |
| A0A669KB10 | GBF1 | Golgi-specific brefeldin A-resistance guanine nucleotide exchange factor 1 | -0.105336428 | -0.0732781 |
| A0A669KB12 | EP300 | Histone acetyltransferase | 0 | 0.992648 |
| A0A669KB41 | SNCA | Alpha-synuclein | 0.081709067 | -0.21247 |
| A0A669KBE6 | TMEM30A | Cell cycle control protein | -2.014920076 | -0.032148 |
| A0A669KBL1 | USP7 | Ubiquitin carboxyl-terminal hydrolase 7 | 0.076446772 | 0.00965087 |
| A0A669KBL4 | AEBP2 | Zinc finger protein AEBP2 | 0.270265818 | 0.52958 |
| A0A6I8PIV1 | LARS1 | Leucyl-tRNA synthetase | 0.117226283 | -0.000661214 |
| A0A6I8PIX7 | SVIL | Supervillin | -0.845257998 | -0.143144 |
| A0A6I8PLD9 | KRTCAP2 | Dolichyl-diphosphooligosaccharide--protein glycosyltransferase subunit KCP2 | -1.897442381 | 0.479569 |
| A0A6I8PRS5 | GPATCH11 | Coiled-coil domain-containing protein 75 | -0.179561059 | 0.123495 |
| A0A6I8PS01 | MSANTD2 | Myb/SANT-like DNA-binding domain-containing protein 2 | 1.739459674 | 0 |
| A0A6I8PTZ8 | AGO1 | Protein argonaute-1 | -0.729246696 | -0.501736 |
| A0A6Q8JGF9 | DPH1 | 2-(3-amino-3-carboxypropyl)histidine synthase subunit 1 | -1.893554846 | -2.20076 |
| A0A6Q8KRG2 | GOLGA2 | Golgin subfamily A member 2 | 0.10679237 | -0.0302277 |
| A0A6Q8PEV5 | CEP41 | Centrosomal protein of 41 kDa | -0.832407554 | -0.989632 |
| A0A6Q8PEX8 | GDAP1 | Ganglioside-induced differentiation-associated protein 1 | -1.424363693 | 0 |
| A0A6Q8PEY3 | NDE1 | Nuclear distribution protein nudE homolog 1 | -0.33004721 | -0.54976 |
| A0A6Q8PEZ3 | PTBP2 | Polypyrimidine tract-binding protein 2 | -0.525343736 | 0.169576 |
| A0A6Q8PF05 | MSI2 | RNA-binding protein Musashi homolog 2 (Fragment) | -1.726416469 | 0.151559 |
| A0A6Q8PF35 | CHMP1A | Charged multivesicular body protein 1a | -1.003605525 | -0.428813 |
| A0A6Q8PF94 | MID1 | E3 ubiquitin-protein ligase Midline-1 | -0.017020146 | -0.721694 |
| A0A6Q8PFA6 | KIF2A | Kinesin-like protein | -0.982515653 | -0.363863 |
| A0A6Q8PFE1 | AIFM1 | Apoptosis-inducing factor 1, mitochondrial | -0.531310558 | 0.0359236 |
| A0A6Q8PFI7 | NEDD4L | E3 ubiquitin-protein ligase | 0.004692872 | -0.0754712 |
| A0A6Q8PFJ0 | LMNA | Prelamin-A/C | -0.47781992 | 0.400192 |
| A0A6Q8PFJ4 | MFN2 | Mitofusin-2 | -1.218203863 | 0.698867 |
| A0A6Q8PFL5 | LRSAM1 | E3 ubiquitin-protein ligase LRSAM1 | 0.005223592 | -0.0456429 |
| A0A6Q8PFU3 | PAFAH1B1 | Platelet-activating factor acetylhydrolase IB subunit alpha | -0.049929937 | -0.119559 |
| A0A6Q8PFY3 | FANCD2 | Fanconi anemia group D2 protein (Fragment) | 0.181658347 | -0.575394 |
| A0A6Q8PFY7 | TFG | Protein TFG | -0.071121534 | -0.0550028 |
| A0A6Q8PG35 | SLC52A2 | Riboflavin transporter | -2.180466255 | -0.767589 |
| A0A6Q8PGB0 | POLR2A | DNA-directed RNA polymerase subunit | -0.061620235 | -0.170003 |
| A0A6Q8PGN3 | INF2 | Inverted formin-2 | -0.05980587 | 0.124543 |
| A0A6Q8PGN5 | AARS1 | Alanine--tRNA ligase | -0.022528807 | -0.00283305 |
| A0A6Q8PGU6 | SPATA5 | Vesicle-fusing ATPase | 0.157423019 | -0.0595741 |
| A0A6Q8PH09 | FANCI | Fanconi anemia group I protein | -0.52097551 | 0.479143 |
| A0A6Q8PH21 | SLC12A6 | Solute carrier family 12 member 6 | -2.100879431 | 2.0531 |
| A0A6Q8PHA5 | HARS1 | Histidine--tRNA ligase, cytoplasmic | 0.070384979 | 0.0039959 |
| A0A6Q8PHK0 | TRIP12 | E3 ubiquitin-protein ligase TRIP12 | -0.375487963 | 0.0468146 |
| A0A6Q8PHP0 | NT5C2 | Cytosolic purine 5'-nucleotidase | 0.07844305 | -0.31303 |

| | | | | |
|---|---|---|---|---|
| A0A7I2R591 | POLR3G | DNA-directed RNA polymerase III subunit RPC7 | 0.527471145 | -1.4849 |
| A0A7I2V268 | PLAT | Tissue-type plasminogen activator | -0.649298271 | 1.85752 |
| A0A7I2V2C6 | RSL24D1 | Probable ribosome biogenesis protein RLP24 | 0 | 0.159012 |
| A0A7I2V2D8 | HGS | Hepatocyte growth factor-regulated tyrosine kinase substrate | 0.146366119 | -0.014431 |
| A0A7I2V2E1 | ADAM9 | Disintegrin and metalloproteinase domain-containing protein 9 | -1.386760314 | 1.19838 |
| A0A7I2V2F9 | ITGB1 | Integrin beta | -1.321089347 | 0.560772 |
| A0A7I2V2G1 | STAT3 | Signal transducer and activator of transcription | -0.940079848 | -2.16462 |
| A0A7I2V2G2 | HSPA9 | 75 kDa glucose-regulated protein | -0.233547052 | 0.0372442 |
| A0A7I2V2K5 | NACA | Nascent polypeptide-associated complex subunit alpha | 0.18537728 | -0.00865936 |
| A0A7I2V2M2 | ACBD5 | Acyl-CoA-binding domain-containing protein 5 | -1.712696711 | 0.00456238 |
| A0A7I2V2Q8 | CTSC | Cathepsin C | -0.175826867 | 0.0841767 |
| A0A7I2V2R3 | HNRNPA3 | Heterogeneous nuclear ribonucleoprotein A3 | -0.656835953 | 0.262541 |
| A0A7I2V2R5 | DPP4 | Dipeptidyl peptidase 4 | -0.998159488 | -0.00778135 |
| A0A7I2V2T3 | IMPDH2 | Inosine-5'-monophosphate dehydrogenase | -0.082834562 | -0.00815646 |
| A0A7I2V2U2 | SGTA | Small glutamine-rich tetratricopeptide repeat-containing protein alpha | 0.209299962 | 0.75688 |
| A0A7I2V2U6 | MTHFD2 | Methenyltetrahydrofolate cyclohydrolase | -1.76302743 | -2.95504 |
| A0A7I2V351 | KPNA2 | Importin subunit alpha | 0.09993124 | -0.0132771 |
| A0A7I2V383 | BCL2L2-PABPN1 | BCL2L2-PABPN1 readthrough | -0.324917078 | 0.0492903 |
| A0A7I2V3B4 | SCFD1 | Sec1 family domain-containing protein 1 | 0.104290326 | -0.233035 |
| A0A7I2V3I0 | EIF3E | Eukaryotic translation initiation factor 3 subunit E | 0.107485771 | -0.284738 |
| A0A7I2V3R8 | HNRNPA1 | Helix-destabilizing protein | -0.509450436 | 0.235293 |
| A0A7I2V3V0 | STAT3 | Signal transducer and activator of transcription | 0.149837891 | -0.0931327 |
| A0A7I2V416 | SPECC1 | Cytospin-B | -0.991206249 | 0 |
| A0A7I2V428 | NCL | Nucleolin | -0.005293687 | -0.179634 |
| A0A7I2V430 | DDX1 | ATP-dependent RNA helicase DDX1 | -0.422958215 | 0.031833 |
| A0A7I2V490 | DNMT1 | DNA (cytosine-5)-methyltransferase | -0.635720094 | 0.172233 |
| A0A7I2V4I6 | HNRNPA2B1 | Heterogeneous nuclear ribonucleoproteins A2/B1 | -0.508937518 | 0.315722 |
| A0A7I2V4K5 | CTSA | Carboxypeptidase | -0.128420273 | 0.184891 |
| A0A7I2V4L6 | YKT6 | Synaptobrevin homolog YKT6 | -0.007833004 | -0.216267 |
| A0A7I2V4L7 | PABPC4 | Polyadenylate-binding protein | -0.189390898 | -0.0726039 |
| A0A7I2V4T6 | NOMO3 | Nodal modulator 3 | 4.510168513 | -6.07745 |
| A0A7I2V4W4 | TBK1 | Serine/threonine-protein kinase TBK1 | -0.044509013 | -0.276378 |
| A0A7I2V535 | NONO | Non-POU domain-containing octamer-binding protein | -0.713724057 | 0.443022 |
| A0A7I2V539 | NMT1 | Glycylpeptide N-tetradecanoyltransferase | -0.01077191 | -0.0639381 |
| A0A7I2V597 | BCL2L1 | Apoptosis regulator Bcl-X | -0.558413426 | -0.594372 |
| A0A7I2V5C6 | PITRM1 | Pitrilysin metalloproteinase 1 | -0.497824113 | 0.11727 |
| A0A7I2V5D2 | MKRN2 | Probable E3 ubiquitin-protein ligase makorin-2 | -0.563907623 | -0.247789 |
| A0A7I2V5H3 | PRKRA | Interferon-inducible double-stranded RNA-dependent protein kinase activator A | -2.077498198 | 1.88478 |
| A0A7I2V5J8 | NPM1 | Nucleophosmin | -2.523499966 | -0.896177 |
| A0A7I2V5M3 | CTSL | Procathepsin L | -0.040996075 | 0.26032 |
| A0A7I2V5Q5 | PPIG | Peptidyl-prolyl cis-trans isomerase G | -0.687565724 | 0.2315 |
| A0A7I2V5R1 | NEK9 | Serine/threonine-protein kinase Nek9 | 0.140989939 | 0.172811 |
| A0A7I2V5T4 | ACO2 | Aconitate hydratase, mitochondrial | -0.191526095 | -0.00744152 |
| A0A7I2V5T7 | PTGES2 | Microsomal prostaglandin E synthase 2 | -1.42547369 | 0.096186 |
| A0A7I2V5U5 | TJP1 | Tight junction protein ZO-1 | 0.035749992 | 0.0832783 |
| A0A7I2V5Y5 | KIF23 | Kinesin-like protein | -1.967873931 | 2.9558 |
| A0A7I2Y1C3 | EIF4G1 | Eukaryotic translation initiation factor 4 gamma 1 | -0.195170244 | -0.1992 |
| A0A7I2YBM0 | DICER1 | Endoribonuclease Dicer | -0.614397605 | 3.11261 |
| A0A7I2YQ74 | UGP2 | UTP--glucose-1-phosphate uridylyltransferase | 0.408193588 | 0.273449 |
| A0A7I2YQ75 | USP40 | Ubiquitin carboxyl-terminal hydrolase 40 | -0.119710763 | -0.445501 |
| A0A7I2YQ78 | CSNK2B | Casein kinase II subunit beta | -0.068701744 | 0.344625 |
| A0A7I2YQF8 | EIF3I | Eukaryotic translation initiation factor 3 subunit I | 0.106351535 | -0.162011 |
| A0A7I2YQG2 | RPL19 | Ribosomal protein L19 | -0.777533213 | -0.0263402 |
| A0A7I2YQN2 | SYNCRIP | Heterogeneous nuclear ribonucleoprotein Q | 0.141430298 | 0.439425 |
| A0A7I2YQU6 | WASH6P | WAS protein family homolog 6 | 0.060818513 | -0.0431108 |
| A0A7N4I394 | PRPF40A | Pre-mRNA-processing factor 40 homolog A | 0.013882081 | 0.0338373 |
| A0A7P0MKV3 | MRPS22 | 28S ribosomal protein S22, mitochondrial | -0.915603717 | -0.458588 |
| A0A7P0MUT5 | CHEK2 | Serine/threonine-protein kinase Chk2 | -0.680917581 | 0.18035 |
| A0A7P0T7V2 | OAS3 | 2'-5'-oligoadenylate synthase 3 | -1.634942671 | 0 |
| A0A7P0T7Z2 | ARFGEF2 | Brefeldin A-inhibited guanine nucleotide-exchange protein 2 | -0.209019423 | 0.0945082 |
| A0A7P0T824 | VRK1 | Serine/threonine-protein kinase VRK1 | -1.293100039 | 0.187984 |
| A0A7P0T841 | TSEN2 | tRNA-splicing endonuclease subunit Sen2 | -0.804338694 | 0.847292 |
| A0A7P0T868 | AATF | Protein AATF | -1.772214333 | 1.00595 |
| A0A7P0T890 | DST | Dystonin | -0.229117393 | -0.046868 |
| A0A7P0T893 | AKAP8 | A-kinase anchor protein 8 | -0.12251099 | -0.0688938 |
| A0A7P0T8C7 | SNX9 | Sorting nexin-9 | -0.001085917 | 0.0171013 |
| A0A7P0T8F3 | ALS2 | Alsin | -0.690379937 | -0.871015 |
| A0A7P0T8I6 | CTSZ | Cathepsin X | 0.204595486 | -0.604589 |

| | | | | |
|---|---|---|---|---|
| A0A7P0T8L1 | DENND5A | DENN domain-containing protein 5A | -0.145890792 | 0.535669 |
| A0A7P0T8N8 | TCERG1 | Transcription elongation regulator 1 | -0.866156816 | 0.264341 |
| A0A7P0T901 | ALG5 | Dolichyl-phosphate beta-glucosyltransferase | -0.914175113 | -0.103822 |
| A0A7P0T904 | CTSS | Cathepsin S | -0.239366213 | 0.747771 |
| A0A7P0T913 | SCARB1 | Scavenger receptor class B member 1 | -0.125105619 | 0.561227 |
| A0A7P0T915 | SIRT5 | NAD-dependent protein deacylase sirtuin-5, mitochondrial | 0.075584968 | 0.21132 |
| A0A7P0T937 | CANX | Calnexin | -1.207417488 | 0.0501026 |
| A0A7P0T943 | FAM126A | Hyccin | -0.602684736 | -1.53364 |
| A0A7P0T947 | RBPJ | Recombining-binding protein suppressor of hairless (Fragment) | -0.473708789 | -0.318842 |
| A0A7P0T9B2 | VPS53 | Vacuolar protein sorting-associated protein 53 homolog | -2.416678468 | -2.33623 |
| A0A7P0T9D5 | SIGMAR1 | Sigma non-opioid intracellular receptor 1 | -1.480093241 | -0.183978 |
| A0A7P0T9D8 | NAXD | ATP-dependent (S)-NAD(P)H-hydrate dehydratase | 0.304694255 | -0.174089 |
| A0A7P0T9D9 | WDR62 | WD repeat-containing protein 62 | 0.063250701 | -0.151254 |
| A0A7P0T9E7 | CRAT | Carnitine O-acetyltransferase | -1.418447971 | 1.92729 |
| A0A7P0T9G4 | ERCC6 | Chimeric ERCC6-PGBD3 protein | -0.196534793 | -0.631714 |
| A0A7P0T9G7 | MTR | Methionine synthase | -0.148814758 | 0.190988 |
| A0A7P0T9I9 | NAMPT | Nicotinamide phosphoribosyltransferase | 0.083579063 | -0.13592 |
| A0A7P0T9J1 | DIMT1 | rRNA adenine N(6)-methyltransferase | -1.346688191 | -0.9002 |
| A0A7P0T9J6 | SMPD4 | Sphingomyelin phosphodiesterase 4 | -1.868688901 | 0.517735 |
| A0A7P0T9P2 | MICALL1 | MICAL-like protein 1 | -0.419678291 | -1.24349 |
| A0A7P0T9T8 | DDX20 | RNA helicase | -0.124905904 | -0.345169 |
| A0A7P0T9W1 | DBT | Lipoamide acyltransferase component of branched-chain alpha-keto acid dehydrogenase complex, mitochondrial | 0.063538909 | 0 |
| A0A7P0TA35 | P4HB | Protein disulfide-isomerase | -0.006124655 | 0.0579211 |
| A0A7P0TA70 | AAR2 | AAR2 splicing factor homolog | -0.660149654 | -0.147722 |
| A0A7P0TA76 | ACSL3 | Fatty acid CoA ligase Acsl3 | -1.022706827 | -0.348415 |
| A0A7P0TA78 | HTT | Huntingtin | -0.227612336 | -0.154879 |
| A0A7P0TA85 | BSG | Basigin | -1.16242377 | 0.355062 |
| A0A7P0TAE3 | AKAP8L | A-kinase anchor protein 8-like | -0.073292176 | 0.0352519 |
| A0A7P0TAG1 | DLAT | Dihydrolipoamide acetyltransferase component of pyruvate dehydrogenase complex | -0.702656507 | -0.0576626 |
| A0A7P0TAN7 | SLC3A2 | 4F2 cell-surface antigen heavy chain | -0.812917312 | 0.0787226 |
| A0A7P0TAP9 | USP13 | Ubiquitin carboxyl-terminal hydrolase | 0.115071456 | 0.0944983 |
| A0A7P0TAQ9 | DNAJC10 | DnaJ homolog subfamily C member 10 | -0.883182208 | 0.010006 |
| A0A7P0TAW3 | VCP | 15S Mg(2+)-ATPase p97 subunit | -0.112817764 | -5.72E-06 |
| A0A7P0TBA9 | EIF2AK2 | Interferon-induced, double-stranded RNA-activated protein kinase | 0.696128686 | 0.167875 |
| A0A7P0Z415 | CTSV | Cathepsin L2 | -0.284902334 | 0.516147 |
| A0A7P0Z497 | PPIB | Peptidyl-prolyl cis-trans isomerase | -0.574911276 | 0.0606346 |
| A0A7P0Z4C3 | DCTN1 | Dynactin subunit 1 | 0.068548838 | -0.0771948 |
| A0A7P0Z4F9 | ADAR | Double-stranded RNA-specific adenosine deaminase | -0.634670019 | 0.233922 |
| A0A804C9D6 | KLHL5 | Kelch-like protein 5 | 0.335554997 | -0.0269092 |
| A0A804F6T5 | PKM | Pyruvate kinase | -0.2362655 | 0.195397 |
| A0A804HHU9 | STAMBP | STAM-binding protein | -0.004306237 | 0.148171 |
| A0A804HHW7 | TBCD | Tubulin-specific chaperone D | -0.136193832 | -0.235736 |
| A0A804HI25 | DHCR7 | 7-dehydrocholesterol reductase | -1.42111063 | 0.312485 |
| A0A804HI27 | TUBGCP2 | Gamma-tubulin complex component | -0.223368963 | 0.545081 |
| A0A804HI56 | SLC25A25 | Calcium-binding mitochondrial carrier protein SCaMC-2 | -1.470266898 | -0.29799 |
| A0A804HI79 | RP9 | Retinitis pigmentosa 9 protein | -3.34941411 | 0.778521 |
| A0A804HIC4 | EVI5 | Ecotropic viral integration site 5 protein homolog | -0.085140864 | 0.0600011 |
| A0A804HIG4 | LRPPRC | Leucine-rich PPR motif-containing protein, mitochondrial | -0.770076672 | 0.134135 |
| A0A804HII9 | ACTN1 | Alpha-actinin-1 | 0.115775108 | -0.135798 |
| A0A804HIS2 | MAP4K5 | Mitogen-activated protein kinase kinase kinase kinase 5 | -0.707797686 | -1.03182 |
| A0A804HIU8 | EXT2 | Exostosin-2 | 6.032466213 | 0 |
| A0A804HJ24 | MCCC2 | 3-methylcrotonyl-CoA carboxylase 2 | -0.760610183 | 0.171525 |
| A0A804HJ86 | UBA5 | UFM1-activating enzyme | -0.02578338 | 0.0951118 |
| A0A804HJA4 | ECPAS | Proteasome adapter and scaffold protein ECM29 | -0.592490991 | 0.04365 |
| A0A804HJB7 | TACO1 | Translational activator of cytochrome c oxidase 1 | -1.006941636 | -0.578931 |
| A0A804HJG7 | CLASP2 | CLIP-associating protein 2 | -0.401036024 | -0.0672461 |
| A0A804HJL6 | ALG1 | Chitobiosyldiphosphodolichol beta-mannosyltransferase | -1.942580938 | 0.0631253 |
| A0A804HK53 | ERCC2 | General transcription and DNA repair factor IIH helicase subunit XPD | -0.595801512 | -0.737341 |
| A0A804HKA1 | | Calcipressin-1 (Fragment) | 0.642325242 | -0.0268475 |
| A0A804HKE7 | MAP4 | Microtubule-associated protein | -0.604583104 | 0.303563 |
| A0A804HKH9 | AIP | Peptidylprolyl isomerase (Fragment) | 0.109414895 | 0.0195662 |
| A0A804HKV3 | ACTA1 | Actin, alpha skeletal muscle | -0.162053585 | -0.440198 |
| A0A804HL87 | RAPGEF1 | Rap guanine nucleotide exchange factor 1 | 0.157449087 | 0.301567 |
| A0A8C8KBL6 | ITGA6 | Integrin alpha-6 | -0.887411714 | 0.431025 |
| A0A8C8MQ70 | UBAP2 | Ubiquitin-associated protein 2 | -0.236406008 | 0.147822 |
| A0A8C8MSB2 | PDHX | Pyruvate dehydrogenase protein X component, mitochondrial | -0.522953749 | -0.23932 |
| A0A8C8MUD1 | AIMP1 | Aminoacyl tRNA synthase complex-interacting multifunctional protein 1 | -0.097619613 | 0.139167 |

| | | | | |
|---|---|---|---|---|
| A0A8C8PZE2 | C19orf12 | Protein C19orf12 (Fragment) | 0.856770118 | -0.418625 |
| A0A8C8QSB1 | HLCS | Biotin--protein ligase | -0.057611942 | -0.197238 |
| A0AVT1 | UBA6 | Ubiquitin-like modifier-activating enzyme 6 | 0.057456175 | -0.0767167 |
| A0JNW5 | UHRF1BP1L | UHRF1-binding protein 1-like | 0.057911237 | -0.177118 |
| A0MZ66 | SHTN1 | Shootin-1 | -0.316296816 | -0.380275 |
| A1L0T0 | ILVBL | 2-hydroxyacyl-CoA lyase 2 | -3.014614741 | -3.61155 |
| A1X283 | SH3PXD2B | SH3 and PX domain-containing protein 2B | -0.773887157 | 0.0124108 |
| A2A283 | SYNGR1 | Synaptogyrin-1 | 0.192459106 | 0.179022 |
| A2A2V1 | PRNP | Major prion protein (Fragment) | -3.00138247 | 1.44205 |
| A2IDC6 | MRPL28 | 39S ribosomal protein L28, mitochondrial (Fragment) | -1.298509757 | 0.861408 |
| A2RRP1 | NBAS | NBAS subunit of NRZ tethering complex | -1.309476852 | -0.447546 |
| A2RUC4 | TYW5 | tRNA wybutosine-synthesizing protein 5 | -0.267341693 | -0.298077 |
| A3KN83 | SBNO1 | Protein strawberry notch homolog 1 | -0.104539792 | 0.0252234 |
| A5PLN9 | TRAPPC13 | Trafficking protein particle complex subunit 13 | -0.608518442 | 0.535218 |
| A5YKK6 | CNOT1 | CCR4-NOT transcription complex subunit 1 | -0.387920777 | -0.202912 |
| A6NCC9 | TSSC4 | Protein TSSC4 | -0.170533339 | 0.754331 |
| A6NCE7 | MAP1LC3B2 | Microtubule-associated proteins 1A/1B light chain 3 beta 2 | -1.59794871 | -0.48315 |
| A6ND22 | MRPS16 | 28S ribosomal protein S16, mitochondrial | -2.010292768 | 0.114583 |
| A6ND36 | FAM83G | Protein FAM83G | -0.020932039 | -0.624495 |
| A6NDG6 | PGP | Glycerol-3-phosphate phosphatase | 0.087905566 | -0.108349 |
| A6NDU8 | RIMOC1 | RAB7A-interacting MON1-CCZ1 complex subunit 1 | -0.865942319 | -0.0726461 |
| A6NED2 | RCCD1 | RCC1 domain-containing protein 1 | -0.396807432 | -1.9431 |
| A6NEM2 | HCFC1 | Host cell factor 1 | -0.306057851 | 0.331171 |
| A6NEM5 | PIGK | GPI-anchor transamidase | -1.823446035 | -0.0867964 |
| A6NF51 | BPNT1 | 3'(2'),5'-bisphosphate nucleotidase 1 | 0.004003445 | 0.040062 |
| A6NFX8 | NUDT5 | ADP-sugar pyrophosphatase | 0.118787766 | 0.216368 |
| A6NG10 | WBP2 | WW domain-binding protein 2 | 0.142768542 | -0.0513554 |
| A6NGJ0 | DYNLT3 | Dynein light chain Tctex-type 3 | -0.912793001 | 0.376261 |
| A6NGP5 | JPT2 | Jupiter microtubule-associated homolog 2 | -0.278495391 | 0.420157 |
| A6NGQ4 | ATG4C | Cysteine protease (Fragment) | 1.666629513 | 0.247181 |
| A6NGW1 | YIF1A | Protein YIF1 | -2.694240808 | 1.98624 |
| A6NHR9 | SMCHD1 | Structural maintenance of chromosomes flexible hinge domain-containing protein 1 | -1.960371017 | 0.191668 |
| A6NHX0 | CASTOR2 | Cytosolic arginine sensor for mTORC1 subunit 2 | -0.358346701 | -1.06616 |
| A6NIH7 | UNC119B | Protein unc-119 homolog B | -0.193375985 | 0.467027 |
| A6NIZ0 | NDEL1 | Nuclear distribution protein nudE-like 1 | -0.633355459 | -0.33571 |
| A6NLH6 | CNIH4 | Protein cornichon homolog 4 | -0.497379224 | 0.229482 |
| A6NMH8 | CD81 | CD81 antigen | -0.545019229 | -0.313148 |
| A6NML8 | DIAPH2 | Diaphanous homolog 2 (Drosophila), isoform CRA_c | 0.125614564 | 0 |
| A6NMQ1 | POLA1 | DNA polymerase | -0.343385458 | -2.55001 |
| A6NMQ3 | ENSA | Alpha-endosulfine | 0.036336342 | -0.305039 |
| A6NN40 | SHROOM1 | Protein Shroom1 | -0.354658127 | -0.476715 |
| A6NN50 | OBSL1 | Obscurin-like protein 1 | -0.531366428 | 0.296144 |
| A6NNY8 | USP27X | Ubiquitin carboxyl-terminal hydrolase 27 | -1.142142534 | -1.30528 |
| A6PVK5 | F13A1 | Coagulation factor XIII A chain (Fragment) | -0.723579407 | -2.08581 |
| A6PW57 | PIP5K1A | Phosphatidylinositol 4-phosphate 5-kinase type-1 alpha | 0 | -0.107655 |
| A6ZKI3 | RTL8C | Retrotransposon Gag-like protein 8C | -0.133545558 | -0.125618 |
| A7E2V4 | ZSWIM8 | Zinc finger SWIM domain-containing protein 8 | -0.324694157 | -0.91086 |
| A8DPD7 | ZYG11B | Protein zyg-11 homolog B | -1.223413507 | 2.76758 |
| A8K2U0 | A2ML1 | Alpha-2-macroglobulin-like protein 1 | -0.457193295 | 0.128563 |
| A8K7I4 | CLCA1 | Calcium-activated chloride channel regulator 1 | 0.118768295 | 0 |
| A8MQ02 | AFDN | Afadin | -0.520801306 | -0.450085 |
| A8MQB8 | FMR1 | Synaptic functional regulator FMR1 | -1.434722026 | 2.28259 |
| A8MT37 | GSK3A | [Tau protein] kinase | -0.192505519 | -0.23401 |
| A8MT40 | PDPR | Pyruvate dehydrogenase phosphatase regulatory subunit, mitochondrial | -1.932934682 | 1.39251 |
| A8MTY9 | VPS26C | Vacuolar protein sorting-associated protein 26C | -0.519387801 | 0.725355 |
| A8MU27 | SUMO3 | Small ubiquitin-related modifier 3 | 0.150533994 | 0.999435 |
| A8MUM1 | EIPR1 | EARP and GARP complex-interacting protein 1 | -0.014057716 | -0.17398 |
| A8MV73 | ATF7IP | Activating transcription factor 7-interacting protein 1 (Fragment) | -0.728284995 | 1.3957 |
| A8MWD9 | SNRPGP15 | Putative small nuclear ribonucleoprotein G-like protein 15 | -0.548987468 | 0.182151 |
| A8MX12 | MYOM1 | Myomesin-1 | 0.014930964 | -0.335151 |
| A8MXB7 | SNX24 | Sorting nexin-24 | -0.40009435 | 2.37103 |
| A8MXQ1 | PTTG1IP | Pituitary tumor-transforming gene 1 protein-interacting protein | -0.499848525 | 1.50777 |
| A8MXV4 | NUDT19 | Acyl-coenzyme A diphosphatase NUDT19 | -1.416442633 | 0.350554 |
| A8MYR4 | GTF2A2 | Transcription initiation factor IIA gamma chain | 0.251844724 | -0.967955 |
| A8MZH3 | MBP | Myelin basic protein | 0.435985247 | -0.43564 |
| A9Z1X7 | SRRM1 | Serine/arginine repetitive matrix protein 1 | -0.633362532 | -0.738579 |
| B0QY16 | KIAA0930 | Uncharacterized protein KIAA0930 (Fragment) | -0.22334369 | 0.114881 |
| B0QY89 | EIF3L | Eukaryotic translation initiation factor 3 subunit L | 0.059121768 | -0.276133 |
| B0QYD3 | APOBEC3B | DNA dC->dU-editing enzyme APOBEC-3B | -0.671499729 | 1.26148 |

| | | | | |
|---|---|---|---|---|
| B0QYK0 | EWSR1 | RNA-binding protein EWS | -0.368097544 | -0.163355 |
| B0QYN7 | UBE2I | SUMO-conjugating enzyme UBC9 | -0.208409786 | 0.301398 |
| B0QZ18 | CPNE1 | Copine-1 | 0.098558744 | -0.204567 |
| B0YIW2 | APOC3 | Apolipoprotein C-III | -0.013186614 | 2.66895 |
| B1AH77 | RAC2 | Ras-related C3 botulinum toxin substrate 2 | -0.616526286 | 0.195785 |
| B1AH87 | TSPO | Peripheral-type benzodiazepine receptor (Fragment) | -1.590275923 | -0.0902913 |
| B1AKC9 | EPHB2 | Receptor protein-tyrosine kinase | -0.665011962 | 0.431224 |
| B1AKN3 | RERE | Arginine-glutamic acid dipeptide (RE) repeats, isoform CRA_b | -0.399819613 | 0 |
| B1AKV2 | UQCC1 | Ubiquinol-cytochrome-c reductase complex assembly factor 1 | -2.076441407 | 3.82193 |
| B1AKY9 | ATP1A2 | Sodium/potassium-transporting ATPase subunit alpha | 0.402934869 | -0.312469 |
| B1AL05 | MRPL43 | 39S ribosomal protein L43, mitochondrial | 0 | -1.17682 |
| B1AL69 | CDC37L1 | Hsp90 co-chaperone Cdc37-like 1 | -0.19465065 | 0.361579 |
| B1ALA9 | PRPS1 | Ribose-phosphate pyrophosphokinase 1 | -0.208013137 | -0.119583 |
| B1ALB4 | SMG7 | Protein SMG7 (Fragment) | -0.519162178 | 0.70228 |
| B1ALD0 | AP4B1 | AP complex subunit beta | -0.482892354 | 0.321916 |
| B1ALY0 | PALM2AKAP2 | PALM2 and AKAP2 fusion (Fragment) | -0.002790292 | 0 |
| B1AMU4 | EXOSC1 | Exosome complex component CSL4 | -0.477587938 | 0.350044 |
| B1AN48 | SPRR3 | Small proline-rich protein 3 (Fragment) | 0.303597172 | 0 |
| B1ANG9 | GUK1 | Guanylate kinase | -0.388050954 | -0.448711 |
| B1AP46 | EXOC6 | Exocyst complex component 6 | 0.263305505 | 0.414768 |
| B2R5W2 | HNRNPC | Heterogeneous nuclear ribonucleoproteins C1/C2 | -0.487958113 | 0.245786 |
| B2WTI3 | JMJD6 | Bifunctional arginine demethylase and lysyl-hydroxylase JMJD6 | -0.197425524 | -0.20809 |
| B3KQ30 | ISCU | Iron-sulfur cluster assembly enzyme ISCU, mitochondrial | -0.620958408 | -0.0272163 |
| B3KS98 | EIF3H | Eukaryotic translation initiation factor 3 subunit H | 0.073359013 | -0.335686 |
| B3KSI3 | BCAT2 | Branched-chain-amino-acid aminotransferase | -0.111770709 | -0.251661 |
| B3KSP9 | PTGES3L-AARSD1 | PTGES3L-AARSD1 readthrough | 0.14854455 | 0.239926 |
| B3KTM8 | MORF4L1 | Mortality factor 4-like protein 1 | -0.898683469 | 0.422431 |
| B3KVL5 | ZCCHC10 | Zinc finger CCHC domain-containing protein 10 | 0.697476188 | -1.71093 |
| B3KVZ3 | CENPH | Centromere protein H | -0.532725334 | 0.101298 |
| B3KY94 | CDIPT | CDP-diacylglycerol--inositol 3-phosphatidyltransferase | -1.687042316 | -1.15622 |
| B4DDF4 | CNN2 | Calponin | 0.052555402 | 0.256557 |
| B4DDH2 | STK33 | Serine/threonine-protein kinase 33 | 0.610714912 | 0 |
| B4DE15 | SLC35A2 | UDP-galactose translocator | -1.36943837 | 0 |
| B4DEN4 | ZFAND2B | AN1-type zinc finger protein 2B | -0.820086797 | 0.491645 |
| B4DFA5 | LTO1 | Protein LTO1 homolog | 0.562748591 | 0 |
| B4DGI9 | TCF12 | Transcription factor 12 (Fragment) | -0.594274203 | -0.383886 |
| B4DJL6 | INTS14 | Integrator complex subunit 14 | 0.594125668 | 0 |
| B4DJV2 | CS | Citrate synthase | -0.520009518 | 0.264089 |
| B4DK69 | AKR1C2 | Aldo-keto reductase family 1 member C2 | -0.974583069 | 0.456053 |
| B4DK89 | TRIT1 | cDNA FLJ60623, highly similar to tRNA isopentenyltransferase, mitochondrial | -0.790878455 | -1.40866 |
| B4DLN1 | | cDNA FLJ60124, highly similar to Mitochondrial dicarboxylate carrier | 0.133925517 | 0.58705 |
| B4DQK9 | MTMR14 | Myotubularin-related protein 14 | 0.77691261 | 1.562 |
| B4DQT1 | MAEA | E3 ubiquitin-protein transferase MAEA | -1.447747628 | -0.0971533 |
| B4DR61 | SEC61A1 | Protein transport protein Sec61 subunit alpha isoform 1 | -1.534352303 | 0.146662 |
| B4DR80 | STK24 | Serine/threonine-protein kinase 24 | 0.160730282 | -0.00800196 |
| B4DUD2 | TMBIM1 | Protein lifeguard 3 | 0 | -0.288192 |
| B4DVL2 | NAAA | N-acylethanolamine-hydrolyzing acid amidase | -0.34024636 | -1.11894 |
| B4DXZ6 | FXR1 | Fragile X mental retardation syndrome-related protein 1 | -0.414373954 | -0.092116 |
| B4DY09 | ILF2 | Interleukin enhancer-binding factor 2 | -0.487139861 | -0.00933584 |
| B4DYU2 | COG8 | Conserved oligomeric Golgi complex subunit 8 | 0.642710924 | -3.16648 |
| B4E0T2 | PEX5 | PTS1-BP | 0.201245387 | -0.262333 |
| B4E0Y9 | STK26 | Serine/threonine-protein kinase 26 | 0.135767619 | 0.00696945 |
| B4E128 | SLFN5 | Schlafen family member 5 | -0.405122201 | -0.333498 |
| B4E1G1 | DERL1 | Derlin | -1.492338498 | -0.193067 |
| B4E1Q4 | RIOK3 | Serine/threonine-protein kinase RIO3 | 0.014950832 | -0.34401 |
| B4E2Q0 | ATP2C1 | Calcium-transporting ATPase | -2.157360554 | -0.627703 |
| B4E321 | OS9 | Protein OS-9 | -1.296033303 | 0 |
| B4E3T4 | RBPMS | HCG2043421, isoform CRA_c | -0.727539937 | 0.515916 |
| B5MBZ0 | EML4 | Echinoderm microtubule-associated protein-like 4 | -0.578140895 | 0.0481666 |
| B5MC14 | IL6 | Interleukin-6 | 0.908976158 | 0 |
| B5MCF4 | FGF1 | Fibroblast growth factor 1 | -0.01839757 | 1.96916 |
| B5MCF9 | PES1 | Pescadillo homolog | -0.549030224 | -0.175919 |
| B5MCT9 | STXBP4 | Syntaxin-binding protein 4 | -0.198630651 | 1.82504 |
| B5MCX4 | TRIB2 | Tribbles homolog 2 | -0.122933229 | 0.869991 |
| B5MD46 | TBC1D10A | TBC1 domain family member 10A | -0.472201188 | -0.818927 |
| B5MDE0 | RFT1 | Protein RFT1 homolog | -2.004174471 | -0.154744 |
| B5MDF5 | RAN | GTP-binding nuclear protein Ran | -0.067381859 | -0.0726515 |
| B5MDU6 | LDAH | Lipid droplet-associated hydrolase | -1.108130455 | -0.150014 |

| | | | | |
|---|---|---|---|---|
| B5ME97 | SEPTIN10 | Septin 10, isoform CRA_c | -0.811099529 | 0.41547 |
| B7WPG3 | HNRNPLL | Heterogeneous nuclear ribonucleoprotein L-like | -0.717707554 | -0.821067 |
| B7WPL0 | RIC8B | Synembryn-B | -1.46166269 | 0.887177 |
| B7Z2A9 | ATP6V0A1 | V-type proton ATPase subunit a | -1.084722519 | 0.309063 |
| B7Z2Y2 | COG2 | Component of oligomeric Golgi complex 2 | 0.669729789 | -0.0208864 |
| B7Z3B9 | GAB1 | GRB2-associated-binding protein 1 | -1.800613801 | 0.543907 |
| B7Z4C8 | RPL31 | 60S ribosomal protein L31 | -1.376597484 | -0.477386 |
| B7Z637 | ARMC8 | Armadillo repeat-containing protein 8 | -0.042042494 | -0.678616 |
| B7Z6D5 | DDX27 | RNA helicase | -1.180452665 | 0.340551 |
| B7Z6Q0 | NPRL3 | GATOR complex protein NPRL3 | -0.943181554 | -0.706442 |
| B7Z7F3 | RANBP3 | Ran-binding protein 3 | 0.237385233 | 0 |
| B7Z8L1 | PPHLN1 | Periphilin-1 | -0.955899 | 1.46673 |
| B7ZC38 | SH3GLB2 | Endophilin-B2 | -0.147535324 | 0.150282 |
| B7ZL14 | FNBP1 | FNBP1 protein | 0.233946482 | -0.024449 |
| B8ZZC8 | METTL5 | rRNA N6-adenosine-methyltransferase METTL5 | -0.066315413 | 0.154311 |
| B8ZZD4 | TAX1BP1 | Tax1-binding protein 1 | -0.169246753 | -0.0919015 |
| B8ZZF8 | LSM5 | U6 snRNA-associated Sm-like protein LSm5 | -0.292468389 | 1.04225 |
| B8ZZG1 | PALS2 | Membrane protein, palmitoylated 6 (MAGUK p55 subfamily member 6), isoform CRA_a | -0.442895095 | -0.0339851 |
| B8ZZL5 | MITD1 | MIT domain-containing protein 1 (Fragment) | -2.040759087 | -1.605 |
| B8ZZN6 | SUMO1 | Small ubiquitin-related modifier 1 | 0.075737317 | 0.27114 |
| B8ZZQ6 | PTMA | Prothymosin alpha | 0.380106688 | 0.27657 |
| B8ZZS4 | ANKZF1 | Ankyrin repeat and zinc finger domain-containing protein 1 | -0.850268284 | -0.889128 |
| B8ZZU8 | ELOB | Elongin-B | -0.023346583 | 0.621089 |
| B8ZZW8 | FCHSD2 | F-BAR and double SH3 domains protein 2 | -0.475992441 | 1.94786 |
| B8ZZX6 | STEAP3 | Metalloreductase STEAP3 | -1.246262232 | 0.114785 |
| C4AM86 | KRT35 | Keratin, type I cuticular Ha5 | -0.704089483 | 3.12194 |
| C4AMC7 | WASH3P | Putative WAS protein family homolog 3 | 0.008976539 | -0.0357736 |
| C9IY66 | BTN2A2 | Butyrophilin subfamily 2 member A2 (Fragment) | -1.332387924 | -2.47738 |
| C9IYN7 | ELP6 | Elongator complex protein 6 | -0.789406697 | -2.16784 |
| C9IZG4 | CUTA | Protein CutA | 0.814408859 | 0.864028 |
| C9IZL5 | AP4M1 | AP-4 complex subunit mu-1 | -0.607476473 | 0.467117 |
| C9J0I5 | SLC12A9 | Solute carrier family 12 member 9 (Fragment) | 0 | -0.287389 |
| C9J0I9 | ZC3HC1 | Nuclear-interacting partner of ALK | 0.1307048 | 0.13355 |
| C9J0J7 | PFN2 | Profilin | -0.145092885 | 0.0247717 |
| C9J0K6 | SRI | Sorcin | -0.028831323 | 0.593964 |
| C9J126 | CDH2 | Cadherin-2 | -1.324898958 | 0.505993 |
| C9J1C6 | RNF181 | E3 ubiquitin-protein ligase RNF181 | -0.630656083 | 0 |
| C9J1V9 | EEF1E1-BLOC1S5 | EEF1E1-BLOC1S5 readthrough (NMD candidate) | -0.747696797 | -0.473905 |
| C9J2P0 | UBE2E1 | Ubiquitin-conjugating enzyme E2 E1 (Fragment) | 0.056655327 | -0.219611 |
| C9J2V2 | IKBKG | NF-kappa-B essential modulator (Fragment) | -1.265437921 | 0 |
| C9J2Y9 | POLR2B | DNA-directed RNA polymerase subunit beta | -0.024065336 | 0.0527725 |
| C9J384 | CMSS1 | Protein CMSS1 (Fragment) | -1.464237372 | -0.803549 |
| C9J3G2 | CBWD2 | COBW domain-containing protein 2 | 0.293619712 | 0.171388 |
| C9J4W5 | EIF5A2 | Eukaryotic translation initiation factor 5A (Fragment) | -1.081857284 | -0.207352 |
| C9J4Z3 | RPL37A | 60S ribosomal protein L37a | -1.91989255 | 0.370991 |
| C9J5J4 | LIN9 | Protein lin-9 homolog | 0.671076457 | -0.968953 |
| C9J712 | PFN2 | Profilin | 0.162733714 | 0.204536 |
| C9J880 | GLT8D1 | Glycosyltransferase 8 domain-containing protein 1 (Fragment) | 0.109734376 | -1.08927 |
| C9J8E1 | MAPKAPK3 | MAP kinase-activated protein kinase 3 (Fragment) | -0.019455115 | -0.120925 |
| C9J9P8 | UBE2F | NEDD8-conjugating enzyme UBE2F (Fragment) | -0.288759708 | -0.241398 |
| C9JA08 | NMD3 | 60S ribosomal export protein NMD3 | -0.374166727 | -0.206926 |
| C9JA28 | SSR3 | Signal sequence receptor subunit gamma | -0.970325867 | 0.72203 |
| C9JAW5 | | HIG1 domain-containing protein | -1.459907055 | 0.681391 |
| C9JB30 | MAPRE3 | Microtubule-associated protein RP/EB family member 3 (Fragment) | 0.011758248 | -0.0930354 |
| C9JB84 | IFT57 | Intraflagellar transport protein 57 homolog (Fragment) | -0.483543555 | 2.16375 |
| C9JBG5 | SIDT2 | SID1 transmembrane family member 2 | -0.489697059 | 0 |
| C9JBI3 | PSPH | O-phosphoserine phosphohydrolase (Fragment) | 0.08678627 | -0.108726 |
| C9JBY0 | BRD9 | Bromodomain-containing protein 9 (Fragment) | -1.061884324 | 0.130472 |
| C9JC84 | FGG | Fibrinogen gamma chain | -0.436831077 | -0.165744 |
| C9JCC6 | DRAP1 | Dr1-associated corepressor | -0.131028255 | -0.091437 |
| C9JDE9 | ACAA1 | 3-ketoacyl-CoA thiolase, peroxisomal | -0.463494778 | -1.19422 |
| C9JEH3 | AAMP | Angio-associated migratory cell protein | -0.12301143 | 0.251626 |
| C9JEZ4 | CDC42EP3 | Cdc42 effector protein 3 (Fragment) | -1.232745091 | -2.30805 |
| C9JF17 | APOD | Apolipoprotein D (Fragment) | -0.602255106 | -0.059996 |
| C9JFE4 | GPS1 | COP9 signalosome complex subunit 1 | -0.073808829 | -0.0510782 |
| C9JFR7 | CYCS | Cytochrome c (Fragment) | -0.680658897 | 0.138789 |
| C9JFV4 | PELP1 | Modulator of non-genomic activity of estrogen receptor | -1.497318188 | 0.535206 |
| C9JIF9 | APEH | Acyl-peptide hydrolase | 0.061595281 | 0.0421041 |

| | | | | |
|---|---|---|---|---|
| C9JIZ6 | PSAP | Prosaposin | -0.06605045 | 0.233752 |
| C9JJ19 | MRPS34 | 28S ribosomal protein S34, mitochondrial | -1.228324493 | -0.377173 |
| C9JJN9 | VPS8 | Vacuolar protein sorting-associated protein 8 homolog | -0.163672447 | -1.22365 |
| C9JK30 | SH3BP5 | SH3 domain-binding protein 5 (Fragment) | 0.089310726 | -0.172705 |
| C9JKL2 | STARD3NL | STARD3 N-terminal-like protein | -1.898473342 | -1.20514 |
| C9JLF4 | EXD2 | Exonuclease 3'-5' domain-containing protein 2 (Fragment) | 0.049217463 | -2.14069 |
| C9JLU1 | POLR2H | DNA-directed RNA polymerases I, II, and III subunit RPABC3 (Fragment) | -0.127937873 | 0.0798054 |
| C9JLV4 | APAF1 | Apoptotic protease-activating factor 1 | -0.121348302 | -0.354744 |
| C9JLW8 | MCRIP1 | Mapk-regulated corepressor-interacting protein 1 | -0.458239158 | 0 |
| C9JNW5 | RPL24 | 60S ribosomal protein L24 | -1.323523919 | -0.140716 |
| C9JQ41 | MIX23 | Coiled-coil domain-containing protein 58 | -0.131972392 | 0.983733 |
| C9JQD4 | PPIH | Peptidyl-prolyl cis-trans isomerase | -0.068264325 | 0.178445 |
| C9JQG9 | AGBL5 | Cytosolic carboxypeptidase-like protein 5 (Fragment) | -1.662493308 | 0 |
| C9JRZ6 | CHCHD3 | MICOS complex subunit MIC19 | -0.998957157 | 0.517582 |
| C9JSK5 | PRKAR1B | cAMP-dependent protein kinase type I-beta regulatory subunit (Fragment) | 0.092630863 | 0.553941 |
| C9JW74 | MEST | Mesoderm-specific transcript homolog protein | -1.625255823 | 1.62326 |
| C9JWV4 | NHEJ1 | Non-homologous end-joining factor 1 (Fragment) | -0.443940322 | 0.0425968 |
| C9JX88 | PSMC2 | 26S proteasome AAA-ATPase subunit RPT1 | -0.034197013 | -0.0732956 |
| C9JXM3 | RABL3 | Rab-like protein 3 | -1.293637991 | -0.550055 |
| C9JYM0 | POP7 | Ribonuclease P (Fragment) | -0.296144088 | -0.482384 |
| C9JYQ9 | RPL22L1 | 60S ribosomal protein L22-like 1 | -1.061816374 | 0.607341 |
| C9JZJ1 | STK17B | Serine/threonine-protein kinase 17B (Fragment) | -0.279512008 | -1.30153 |
| C9JZW7 | ADPRH | [Protein ADP-ribosylarginine] hydrolase (Fragment) | -1.075359503 | 0.03989 |
| C9JZY6 | UBE2H | Ubiquitin-conjugating enzyme E2 H (Fragment) | -0.170969725 | -0.302674 |
| C9K0J5 | RAPH1 | Ras association (RalGDS/AF-6) and pleckstrin homology domains 1, isoform CRA_b | -0.446485519 | -0.758123 |
| D3DPS3 | MAP7D1 | Arginine/proline rich coiled-coil 1, isoform CRA_b | -0.643083731 | -0.286513 |
| D3DQV9 | EIF4G2 | Eukaryotic translation initiation factor 4 gamma 2 (Fragment) | -0.462022305 | -0.0990267 |
| D3YTA9 | PPP3R1 | Calcineurin subunit B type 1 | -0.384059191 | -0.393663 |
| D3YTB1 | RPL32 | 60S ribosomal protein L32 (Fragment) | -1.267770529 | 0.0877355 |
| D6R938 | CAMK2D | Calcium/calmodulin-dependent protein kinase | -0.107865493 | 0.214863 |
| D6R945 | TTC29 | Tetratricopeptide repeat protein 29 (Fragment) | 1.783695459 | -0.879575 |
| D6R992 | RELB | Transcription factor RelB | 0.150316477 | -0.107868 |
| D6R995 | SH3BP2 | SH3 domain-binding protein 2 (Fragment) | -0.273170074 | 0 |
| D6R9D2 | GPM6A | Neuronal membrane glycoprotein M6-a (Fragment) | 1.740386963 | 0.0261974 |
| D6R9P3 | HNRNPAB | Heterogeneous nuclear ribonucleoprotein A/B | -0.646103462 | 0.0788269 |
| D6RA47 | MFSD10 | Major facilitator superfamily domain-containing protein 10 | -0.045388699 | -2.76444 |
| D6RAA5 | TACC2 | Transforming acidic coiled-coil-containing protein 2 | -0.703769843 | 0.401913 |
| D6RAR4 | HGFAC | Hepatocyte growth factor activator | -0.676835616 | 0 |
| D6RB01 | FAM169A | Soluble lamin-associated protein of 75 kDa | 0.017178694 | -1.49777 |
| D6RB40 | AGTRAP | Type-1 angiotensin II receptor-associated protein | -2.350191116 | 0.285018 |
| D6RCT7 | KIF15 | Kinesin-like protein KIF15 (Fragment) | -0.19003137 | 0.118723 |
| D6RD17 | JCHAIN | Immunoglobulin J chain (Fragment) | -1.891068061 | 0.764014 |
| D6RD46 | LIMCH1 | LIM and calponin homology domains-containing protein 1 | 0.04029425 | 0.862264 |
| D6REA1 | SIL1 | Nucleotide exchange factor SIL1 | -0.275063435 | -3.20274 |
| D6RER5 | SEPTIN11 | Septin | -0.462696155 | 0.00379276 |
| D6REX3 | SEC31A | Protein transport protein Sec31A | 0.06722784 | -0.0512454 |
| D6RF48 | STX18 | Syntaxin-18 | -1.08356142 | 1.7213 |
| D6RF69 | MGAT1 | Alpha-1,3-mannosyl-glycoprotein 2-beta-N-acetylglucosaminyltransferase (Fragment) | -4.0813423 | -1.60315 |
| D6RFI3 | YIPF3 | Protein YIPF3 (Fragment) | -0.714274963 | -3.05768 |
| D6RGK3 | TAF9 | Transcription initiation factor TFIID subunit 9 (Fragment) | -0.40195926 | -0.830958 |
| D6RGK9 | CNOT6L | CCR4-NOT transcription complex subunit 6-like (Fragment) | 0.598170201 | -0.0107892 |
| D6RGV5 | COX7A2 | Cytochrome c oxidase subunit 7A2, mitochondrial (Fragment) | -3.858730247 | 0.861163 |
| D6RGX2 | UFSP2 | Ufm1-specific protease 2 | -1.437359095 | -2.25358 |
| D6RHD0 | ELOVL7 | Elongation of very long chain fatty acids protein | -0.001801173 | 0 |
| D6RJC3 | INPP4B | Phosphatidylinositol-3,4-bisphosphate 4-phosphatase (Fragment) | 0.216925065 | 0.01052 |
| D6W601 | ADAT3 | HCG22695, isoform CRA_a | -0.6404562 | -0.239543 |
| D9J307 | HLA-B | HLA class I antigen | -0.551315626 | 0.277717 |
| E2QRF9 | GMNN | Geminin (Fragment) | -0.200228532 | -0.44393 |
| E5RFC6 | SNAP91 | Clathrin coat assembly protein AP180 (Fragment) | 1.066206694 | -0.628394 |
| E5RFZ8 | SLC39A14 | Metal cation symporter ZIP14 (Fragment) | -4.686071013 | 0.567252 |
| E5RG17 | TATDN1 | Putative deoxyribonuclease TATDN1 (Fragment) | -0.19263649 | 0.241591 |
| E5RG25 | UBE2W | Ubiquitin-conjugating enzyme E2 W | -0.15001901 | -1.11695 |
| E5RGC1 | FBXW11 | F-box/WD repeat-containing protein 11 | -0.520075083 | -0.0229321 |
| E5RGQ3 | RWDD1 | RWD domain-containing protein 1 (Fragment) | -0.20289286 | -2.31598 |
| E5RGS2 | REEP4 | Receptor expression-enhancing protein | -1.885341724 | -0.564689 |
| E5RGS4 | PFDN1 | Prefoldin subunit 1 | 0.72681133 | 0.382512 |
| E5RGS7 | | LIM zinc-binding domain-containing protein | -0.377794425 | 0 |

| | | | | |
|---|---|---|---|---|
| E5RGX5 | STMN2 | Stathmin | 0.099809488 | -0.0888583 |
| E5RH81 | CA1 | Carbonic anhydrase (Fragment) | 0.4696455 | -0.981772 |
| E5RHG8 | ELOC | Elongin-C (Fragment) | 0.217707316 | 0.659425 |
| E5RHW4 | ERLIN2 | Erlin-2 (Fragment) | -1.036015272 | 0.260472 |
| E5RI99 | RPL30 | 60S ribosomal protein L30 (Fragment) | -1.06531771 | 0.297092 |
| E5RIA4 | DNPEP | Aspartyl aminopeptidase | -0.082704624 | -0.240074 |
| E5RIG2 | CSMD1 | CUB and sushi domain-containing protein 1 | 0.398894787 | -0.792945 |
| E5RIH5 | TTI2 | TELO2-interacting protein 2 | -0.002387842 | -1.05711 |
| E5RJR5 | SKP1 | S-phase kinase-associated protein 1 | 0.026570082 | 0.420705 |
| E5RK69 | ANXA6 | Annexin | -0.026377598 | -0.0489629 |
| E7EM57 | G6PD | Glucose-6-phosphate 1-dehydrogenase (Fragment) | -0.343030214 | 1.08297 |
| E7EM64 | COPS6 | COP9 signalosome complex subunit 6 | 0.01443704 | -0.0253633 |
| E7EMS2 | NPC2 | Epididymal secretory protein E1 | -0.12968715 | 0.239153 |
| E7EN19 | MAP4K4 | Mitogen-activated protein kinase kinase kinase kinase 4 | -0.562882264 | 0.118214 |
| E7ENR5 | IMP4 | U3 small nucleolar ribonucleoprotein protein IMP4 (Fragment) | -2.1613729 | 4.85896 |
| E7ENY0 | ADD1 | Alpha-adducin | 0.003515402 | -0.0140476 |
| E7EPD8 | TDP1 | Tyrosyl-DNA phosphodiesterase 1 | -0.181581895 | -0.259287 |
| E7EPK1 | SEPTIN7 | Septin | -0.509003639 | -0.0832526 |
| E7EPN9 | PRRC2C | Protein PRRC2C | -0.459040801 | -0.232526 |
| E7EPP6 | CHEK1 | Serine/threonine-protein kinase Chk1 | -0.762744268 | -0.199068 |
| E7EPT4 | NDUFV2 | NADH dehydrogenase [ubiquinone] flavoprotein 2, mitochondrial | -0.886257251 | 0.869772 |
| E7EQ47 | RHCE | Blood group Rh(CE) polypeptide | 0.986022234 | -0.750306 |
| E7EQ69 | NAA50 | N-alpha-acetyltransferase 50 | -0.09995993 | 0.248619 |
| E7EQ72 | TMED2 | Transmembrane emp24 domain-containing protein 2 (Fragment) | -1.580279589 | -0.879444 |
| E7EQB2 | LTF | Lactotransferrin (Fragment) | -0.514759461 | 0.543225 |
| E7EQU1 | HMGB3 | High mobility group protein B3 (Fragment) | -0.137475411 | 0.0955877 |
| E7EQY4 | MTA3 | Metastasis-associated protein MTA3 | 0.066562017 | -0.479814 |
| E7ERK9 | EIF2B4 | Translation initiation factor eIF-2B subunit delta | -0.186019977 | -0.272289 |
| E7ERR0 | KLHDC3 | Kelch domain-containing protein 3 | -0.773175796 | 0 |
| E7ERS3 | ZC3H18 | Zinc finger CCCH domain-containing protein 18 | -0.060047309 | -0.0612224 |
| E7ES96 | PSEN1 | Presenilin | -1.426902533 | 0.785029 |
| E7ESL0 | MRPL22 | 39S ribosomal protein L22, mitochondrial | 0 | -0.0133362 |
| E7ESZ3 | PLEKHG2 | Pleckstrin homology domain-containing family G member 2 | -1.218830427 | 0 |
| E7ET40 | PLAU | Urokinase-type plasminogen activator | -1.251370748 | -0.254436 |
| E7ETU5 | RBMS1 | RNA-binding motif, single-stranded-interacting protein 1 | -0.128465255 | 0.127399 |
| E7ETU7 | MRPL3 | 39S ribosomal protein L3, mitochondrial | -1.529486259 | 1.33545 |
| E7ETZ4 | BZW2 | Basic leucine zipper and W2 domain-containing protein 2 (Fragment) | 0.023080667 | -0.137868 |
| E7EU81 | GOLGB1 | Golgin subfamily B member 1 (Fragment) | -1.091724634 | -2.25638 |
| E7EUK8 | CDK14 | Cyclin-dependent kinase 14 | -0.224676291 | 1.00123 |
| E7EUP3 | KAT7 | Histone acetyltransferase | -0.767519156 | -2.08145 |
| E7EVG2 | PBRM1 | Protein polybromo-1 (Fragment) | 0 | -4.5173 |
| E7EVY3 | CAST | Calpain inhibitor | 1.749915918 | -2.16168 |
| E7EW05 | SDAD1 | Protein SDA1 | -1.13688 | -0.738268 |
| E7EWP0 | NDUFB5 | Complex I-SGDH | -1.700579643 | 1.6597 |
| E7EWR4 | CSTF2 | Cleavage stimulation factor subunit 2 | -0.226545731 | 0.608431 |
| E7EX17 | EIF4B | Eukaryotic translation initiation factor 4B | 0.044680754 | 0.0207539 |
| E7EX40 | RAB11FIP1 | Rab11 family-interacting protein 1 | -1.069163322 | 0 |
| E7EX70 | POLR1E | DNA-directed RNA polymerase I subunit RPA49 | -2.877039274 | 1.22539 |
| E9PC47 | LIMK1 | LIM domain kinase 1 | -0.264748732 | 0.106553 |
| E9PC84 | TNC | Tenascin | -1.67086633 | -0.0142174 |
| E9PCD7 | MAN2B2 | Alpha-mannosidase | -0.412733873 | 0.645859 |
| E9PCH4 | | Uncharacterized protein | -0.365837574 | -2.05467 |
| E9PCW1 | GOSR1 | Golgi SNAP receptor complex member 1 | -2.007475456 | -0.00546106 |
| E9PD68 | CRMP1 | Dihydropyrimidinase-related protein 1 | 1.087307215 | -0.18089 |
| E9PDF6 | MYO1B | Unconventional myosin-Ib | -0.488375028 | 0.280257 |
| E9PE17 | MRPS17 | 28S ribosomal protein S17, mitochondrial (Fragment) | -1.3092858 | -0.696916 |
| E9PE96 | PCLO | Protein piccolo | -1.784680764 | -0.418197 |
| E9PEI0 | CDCA2 | Cell division cycle-associated protein 2 | -0.468236128 | 0.438292 |
| E9PEP6 | NRP1 | Neuropilin | -0.272282124 | -0.121142 |
| E9PER6 | PDPK1 | 3-phosphoinositide-dependent protein kinase 1 | 0.106905619 | -0.0690012 |
| E9PF10 | NUP155 | Nuclear pore complex protein Nup155 | -0.006917159 | 0.164876 |
| E9PF16 | ACSF2 | Medium-chain acyl-CoA ligase ACSF2, mitochondrial | -1.029585679 | 0.534207 |
| E9PF18 | HADH | Hydroxyacyl-coenzyme A dehydrogenase, mitochondrial | -1.157556852 | 0.189766 |
| E9PF19 | TBL2 | Transducin beta-like protein 2 | -1.5430065 | 0.273676 |
| E9PF46 | ACYP2 | Acylphosphatase | 0.238066753 | -0.461116 |
| E9PFH1 | RND3 | Rho-related GTP-binding protein RhoE | -0.575573047 | 0.26678 |
| E9PFK9 | RABGEF1 | Rab5 GDP/GTP exchange factor | -0.134281953 | -0.0280094 |
| E9PFR3 | PPP2R5D | Serine/threonine-protein phosphatase 2A 56 kDa regulatory subunit | -0.112149 | -0.250554 |
| E9PGC0 | RASA1 | Ras GTPase-activating protein 1 | 0.139962196 | -0.0021197 |

| | | | | |
|---|---|---|---|---|
| E9PGC5 | PTPRK | Protein-tyrosine-phosphatase | -0.005051851 | 2.04827 |
| E9PGC8 | MAP1A | Microtubule-associated protein 1A | -0.617304564 | -0.120094 |
| E9PGM9 | RBM6 | RNA-binding protein 6 | -0.560974717 | 0 |
| E9PGT1 | TSN | Component 3 of promoter of RISC | 0.059367498 | -0.157152 |
| E9PGT3 | RPS6KA1 | Ribosomal protein S6 kinase | -0.124855995 | 0.0114094 |
| E9PGT6 | COPS8 | COP9 signalosome complex subunit 8 | -0.030779441 | 0.0599842 |
| E9PH64 | NDUFB9 | NADH dehydrogenase [ubiquinone] 1 beta subcomplex subunit 9 | -1.531842788 | 0.443317 |
| E9PHV5 | ITPRID2 | Protein ITPRID2 | -0.324325562 | 0.293643 |
| E9PHZ5 | CTSB | Cathepsin B | -0.225800037 | 0.0420761 |
| E9PI70 | CHID1 | Chitinase domain-containing protein 1 (Fragment) | -0.599526882 | 1.25103 |
| E9PIM6 | THY1 | Thy-1 antigen (Fragment) | -1.65504535 | 0.113311 |
| E9PIT3 | F2 | Coagulation factor II | -1.239849647 | -0.634815 |
| E9PIV9 | CYB5R2 | NADH-cytochrome b5 reductase 2 (Fragment) | -0.6350317 | 0.344041 |
| E9PIW9 | SAMD1 | Sterile alpha motif domain-containing protein 1 | -0.712228457 | -0.0615309 |
| E9PJ40 | EBAG9 | Receptor-binding cancer antigen-expressed on SiSo cells (Fragment) | -0.866906484 | 0.0481908 |
| E9PJ42 | TMEM41B | Transmembrane protein 41B | -1.526345094 | -0.47237 |
| E9PJ55 | TCP11L1 | T-complex protein 11-like protein 1 | 0.173416456 | -0.174068 |
| E9PJD7 | CYHR1 | Cysteine and histidine-rich protein 1 | -0.148730516 | -0.147002 |
| E9PJJ5 | KCTD21 | BTB/POZ domain-containing protein KCTD21 (Fragment) | 0.225420713 | -0.660067 |
| E9PJM3 | FBXO3 | F-box only protein 3 | -0.199489196 | 0.343642 |
| E9PK00 | UVRAG | UV radiation resistance-associated gene protein (Fragment) | 0.119976679 | -0.0477854 |
| E9PK54 | HSPA8 | Heat shock cognate 71 kDa protein (Fragment) | 1.129199584 | 2.42114 |
| E9PKF6 | PPP6R3 | Serine/threonine-protein phosphatase 6 regulatory subunit 3 | -0.040157 | -0.336588 |
| E9PKG1 | PRMT1 | Protein arginine N-methyltransferase 1 | -0.020166079 | -0.230876 |
| E9PKT4 | TMEM123 | Porimin | -1.944765449 | 3.64046 |
| E9PKV2 | MRPL17 | 39S ribosomal protein L17, mitochondrial (Fragment) | -1.375771443 | 0.173096 |
| E9PL57 | NEDD8-MDP1 | NEDD8-MDP1 readthrough (Fragment) | -0.568420013 | 1.39551 |
| E9PLG8 | GPAA1 | Glycosylphosphatidylinositol anchor attachment 1 protein (Fragment) | -0.587279399 | 0.0348066 |
| E9PLI6 | EIF1AD | Eukaryotic translation initiation factor 1A domain-containing protein (Fragment) | 0.056836764 | -0.222272 |
| E9PLK3 | NPEPPS | Aminopeptidase | 0.099522114 | -0.0894674 |
| E9PLN8 | | VKc domain-containing protein (Fragment) | -1.814765294 | 1.22276 |
| E9PLP8 | CSTF3 | Cleavage stimulation factor subunit 3 | -0.518390179 | 0.287155 |
| E9PLX2 | RNF141 | RING finger protein 141 (Fragment) | -1.153944413 | -0.644059 |
| E9PLY5 | MACF1 | Microtubule-actin cross-linking factor 1, isoforms 1/2/3/5 (Fragment) | -0.811014136 | 0 |
| E9PM62 | WLS | Protein wntless homolog (Fragment) | -2.237593015 | 0.27496 |
| E9PM92 | C11orf58 | Small acidic protein | -0.032578707 | -0.0599302 |
| E9PMH5 | BIRC2 | Baculoviral IAP repeat-containing protein 2 | 0.142957131 | 0.441585 |
| E9PMI6 | CLNS1A | Chloride channel, nucleotide sensitive 1A | 0.023304939 | 0.427274 |
| E9PMJ2 | FAM118B | Protein FAM118B (Fragment) | -0.024597724 | 0.128223 |
| E9PMQ6 | HSF1 | Heat shock factor protein 1 | 0.187582731 | -0.169112 |
| E9PMR4 | CD151 | Tetraspanin | -1.660857995 | 1.1314 |
| E9PMR6 | ARHGEF12 | Rho guanine nucleotide exchange factor 12 | -0.03705438 | -0.39672 |
| E9PMS6 | LMO7 | LIM domain only protein 7 | -0.429764589 | 0.507248 |
| E9PN70 | TRAPPC4 | Trafficking protein particle complex subunit | -0.365439812 | -0.111587 |
| E9PN81 | RNASEH2C | Ribonuclease H2 subunit C | -0.440982819 | 0.371495 |
| E9PNF7 | PRCP | Lysosomal Pro-X carboxypeptidase | 0.110328356 | -0.00350889 |
| E9PNN3 | PTS | 6-pyruvoyl tetrahydrobiopterin synthase | 0.116225243 | 0.186534 |
| E9PP40 | LRRC14 | Leucine-rich repeat-containing protein 14 (Fragment) | 0.438056628 | -2.30927 |
| E9PPJ1 | KNL1 | Kinetochore scaffold 1 | -0.70834748 | 0.176172 |
| E9PPW7 | NDUFS8 | Complex I-23kD (Fragment) | -1.245549202 | 3.2432 |
| E9PQA6 | MVB12A | Multivesicular body subunit 12A | -0.419969241 | -0.282463 |
| E9PQG4 | PDE4DIP | Myomegalin | -0.154449304 | -1.1105 |
| E9PQY2 | PFDN4 | Prefoldin subunit 4 | -0.457777739 | 1.17082 |
| E9PR11 | NRBP2 | Nuclear receptor-binding protein 2 (Fragment) | 0.858311176 | -2.30293 |
| E9PR30 | FAU | 40S ribosomal protein S30 | -1.11405158 | -0.763144 |
| E9PR76 | MAF1 | Repressor of RNA polymerase III transcription MAF1 homolog (Fragment) | 0.034961939 | 0.0764573 |
| E9PRF4 | SETDB1 | Histone-lysine N-methyltransferase SETDB1 (Fragment) | -0.416987658 | 0 |
| E9PRG8 | C11orf98 | Uncharacterized protein C11orf98 | -0.195300897 | -0.0653804 |
| E9PS17 | SCYL1 | N-terminal kinase-like protein | -0.085114876 | 0.107375 |
| E9PS62 | FRMD8 | FERM domain-containing protein 8 (Fragment) | 0.785843054 | -1.27257 |
| E9PS74 | SLC43A3 | Solute carrier family 43 member 3 (Fragment) | 0 | 0.663096 |
| E9PSE3 | IRF7 | Interferon regulatory factor 7 | 1.711402973 | 0 |
| E9PSI1 | | Transmembrane 9 superfamily member | -0.120182037 | -0.509317 |
| F1DAL7 | NR2F1 | COUP transcription factor 1 | -1.373878519 | 0 |
| F2Z2T2 | XPA | DNA repair protein-complementing XP-A cells | -0.229520162 | 0 |
| F2Z2W7 | TRMT2A | tRNA (uracil-5-)-methyltransferase homolog A | -0.062718948 | -0.114244 |

| | | | | |
|---|---|---|---|---|
| F2Z2X0 | NME4 | Nucleoside diphosphate kinase | 0 | -0.453154 |
| F2Z3J9 | PTGR1 | 15-oxoprostaglandin 13-reductase | -0.027955532 | -0.118687 |
| F5GWI4 | ADA | Adenosine deaminase | 0.00845178 | 0.628299 |
| F5GWI9 | WASHC3 | Coiled-coil domain-containing protein 53 | -0.607307514 | 0.125223 |
| F5GWT4 | WNK1 | Non-specific serine/threonine protein kinase | -0.063549836 | -0.093744 |
| F5GXE4 | ATE1 | Arginyl-tRNA--protein transferase 1 | -0.132173141 | 0.0283941 |
| F5GXJ1 | NDUFS7 | Complex I-20kD | -1.884426435 | 0.0706418 |
| F5GXU9 | BCKDHA | 2-oxoisovalerate dehydrogenase subunit alpha (Fragment) | -0.273956617 | -2.03952 |
| F5GXX5 | DAD1 | Dolichyl-diphosphooligosaccharide--protein glycosyltransferase subunit DAD1 | -0.563191732 | 0.154849 |
| F5GYG5 | REXO2 | Oligoribonuclease, mitochondrial | -0.335203966 | 0.332882 |
| F5GYQ1 | ATP6V0D1 | V-type proton ATPase subunit | -0.089097023 | 0.528979 |
| F5GZ78 | PXN | Paxillin | 0.08255291 | 0.132329 |
| F5GZ90 | DTL | Denticleless protein homolog | -0.321338495 | -0.245062 |
| F5H0C4 | PAAF1 | Proteasomal ATPase-associated factor 1 | 0.141463041 | -0.117287 |
| F5H0F9 | ANAPC5 | Anaphase-promoting complex subunit 5 | -0.482494275 | 0.364431 |
| F5H0R1 | TUT1 | RNA-binding motif protein 21 | -0.630336126 | 2.45629 |
| F5H148 | RRN3 | RNA polymerase I-specific transcription initiation factor RRN3 | -0.219904343 | -1.11844 |
| F5H170 | DNAJA4 | DnaJ homolog subfamily A member 4 | -0.231021166 | -0.280582 |
| F5H199 | MSRB3 | Peptide-methionine (R)-S-oxide reductase (Fragment) | 0.260205666 | 0.939202 |
| F5H1U9 | MPDZ | Multiple PDZ domain protein | -1.050211271 | 0.569422 |
| F5H1Y0 | RAD51AP1 | RAD51-associated protein 1 | -0.307912747 | -0.133191 |
| F5H1Z6 | STARD10 | START domain-containing protein 10 (Fragment) | -0.198014895 | -0.113075 |
| F5H2A4 | HMGA2 | High mobility group protein HMGI-C | -0.858351946 | 0.12877 |
| F5H2C9 | NAA40 | N-alpha-acetyltransferase 40 | -0.187127908 | 0.516579 |
| F5H2F4 | MTHFD1 | C-1-tetrahydrofolate synthase, cytoplasmic | -0.368092537 | 0.255198 |
| F5H365 | SEC23A | Protein transport protein SEC23 | -0.209106922 | -0.00657272 |
| F5H3P7 | PPP1R37 | Protein phosphatase 1 regulatory subunit 37 (Fragment) | 0.529396454 | -1.0847 |
| F5H3Y4 | DHX37 | Probable ATP-dependent RNA helicase DHX37 (Fragment) | 0.266381105 | -1.58813 |
| F5H4B6 | ALDH16A1 | Aldehyde dehydrogenase family 16 member A1 | -0.05046018 | -0.0819352 |
| F5H4M7 | TMED3 | Transmembrane emp24 domain-containing protein 3 | -1.278665066 | 0.90887 |
| F5H4R1 | MAP3K4 | Mitogen-activated protein kinase kinase kinase 4 | -0.384762128 | -0.175485 |
| F5H4V9 | PDCD2 | Programmed cell death protein 2 | -0.107673963 | -0.38673 |
| F5H523 | KLHL42 | Kelch-like protein 42 | -0.216287692 | 0.186795 |
| F5H5D3 | TUBA1C | Tubulin alpha chain | -0.878736178 | -0.0974649 |
| F5H5I6 | GRSF1 | G-rich sequence factor 1 | -0.015563965 | 0.0324033 |
| F5H619 | HEATR5A | HEAT repeat-containing protein 5A | 0.196504196 | 3.58322 |
| F5H625 | DHX40 | Probable ATP-dependent RNA helicase DHX40 (Fragment) | 1.30552268 | -0.425122 |
| F5H658 | DHX8 | RNA helicase | -0.556811492 | -0.149706 |
| F5H6B6 | TRPT1 | 2'-phosphotransferase | 0.192228556 | 0 |
| F5H6E2 | MYO1C | Unconventional myosin-Ic | -0.835208416 | 0.122384 |
| F5H6I7 | ATL3 | Atlastin-3 | -0.133395831 | 0.733251 |
| F5H6Z0 | CDK17 | Cyclin-dependent kinase 17 | -0.158481121 | 0.156379 |
| F5H702 | MRPL48 | 39S ribosomal protein L48, mitochondrial | -3.688711286 | 2.09936 |
| F5H7J9 | LRP6 | Low-density lipoprotein receptor-related protein | 0.245114406 | 0 |
| F5H7W8 | C12orf43 | Protein CUSTOS | 0.285941442 | 0.181321 |
| F5H8F7 | ASH2L | Set1/Ash2 histone methyltransferase complex subunit ASH2 | -0.200451215 | 0.238256 |
| F5H8H2 | MVK | Mevalonate kinase | -0.26994427 | 0.14586 |
| F6SS63 | TAB3 | TAK1-binding protein 3, isoform CRA_c | -0.697775443 | -0.789829 |
| F6TX30 | ASB6 | Ankyrin repeat and SOCS box protein 6 | -0.372311115 | -0.184988 |
| F6UJY1 | HYI | Putative hydroxypyruvate isomerase | -1.100445429 | 0.323589 |
| F6V107 | PICK1 | PRKCA-binding protein (Fragment) | -0.288707733 | 0 |
| F6VRR5 | POLDIP3 | Polymerase delta-interacting protein 3 | -1.389457305 | 0.462059 |
| F6VZ39 | RBM38 | RNA-binding protein 38 | -0.146448215 | 0.0808144 |
| F8VP47 | RPH3A | Rabphilin-3A | 0.717452129 | 2.61202 |
| F8VPD4 | CAD | CAD protein | -0.150813977 | -0.217175 |
| F8VPI1 | TMBIM6 | Bax inhibitor 1 (Fragment) | -1.302527229 | -0.553894 |
| F8VQD9 | ATG101 | Autophagy-related protein 101 (Fragment) | -0.338520845 | 0.0145543 |
| F8VQR7 | CSRP2 | Cysteine and glycine-rich protein 2 | -0.996964455 | 0.0677996 |
| F8VRE8 | TMEM19 | Transmembrane protein 19 (Fragment) | -1.988448461 | -0.292577 |
| F8VSL3 | NFYB | CAAT box DNA-binding protein subunit B | 1.054405689 | 1.43316 |
| F8VU90 | FKBP11 | Peptidylprolyl isomerase | -1.528799772 | 0.566272 |
| F8VUB4 | CNOT2 | CCR4-NOT transcription complex subunit 2 | -0.888413588 | 0.04999 |
| F8VV56 | CD63 | CD63 antigen | -2.297903538 | 0.04498 |
| F8VV59 | NAP1L1 | Nucleosome assembly protein 1-like 1 | 0.098813375 | -0.230644 |
| F8VVA7 | COPZ1 | Coatomer subunit zeta | 0.443174124 | 0.328384 |
| F8VXC8 | SMARCC2 | SWI/SNF complex subunit SMARCC2 | -0.636682908 | 0.723939 |
| F8VZA2 | LETMD1 | LETM1 domain-containing protein 1 (Fragment) | -1.148248593 | -0.858647 |
| F8W038 | C17orf49 | Chromosome 17 open reading frame 49 | 0.072654724 | 0.572014 |
| F8W111 | CPM | Carboxypeptidase M (Fragment) | 0.074085871 | 0.13397 |
| F8W1A4 | AK2 | Adenylate kinase 2, mitochondrial | 0.046354135 | 0.0891336 |

| | | | | |
|---|---|---|---|---|
| F8W1F5 | FMNL3 | Formin-like protein 3 | -0.898171504 | 0.0468715 |
| F8W1J4 | IFT81 | Intraflagellar transport protein 81 homolog (Fragment) | 1.263519883 | 0 |
| F8W1R7 | MYL6 | Myosin light polypeptide 6 | 0.167795499 | 0.777109 |
| F8W1T6 | RBMS2 | RNA-binding motif, single-stranded-interacting protein 2 | -0.975708326 | 1.1643 |
| F8W543 | NAP1L1 | Nucleosome assembly protein 1-like 1 | 0.563413461 | 0.435435 |
| F8W696 | APOA1 | Apolipoprotein A-I | 0.965151588 | 0 |
| F8W7C6 | RPL10 | 60S ribosomal protein L10 | -1.546040416 | -0.651393 |
| F8W7Q4 | FAM162A | Protein FAM162A | -1.686790466 | -0.552639 |
| F8W7U0 | ITSN1 | Intersectin-1 | 0.127676328 | -0.0370725 |
| F8W7U3 | WASHC2A | WASH complex subunit 2A | -0.662927628 | 3.91475 |
| F8W8D3 | SLBP | Histone RNA hairpin-binding protein | -0.205077807 | -0.384431 |
| F8W8I6 | TIA1 | Nucleolysin TIA-1 isoform p40 | -0.532447338 | -1.39419 |
| F8W914 | RTN4 | Reticulon | -0.979442517 | 0 |
| F8W930 | IGF2BP2 | Insulin-like growth factor 2 mRNA-binding protein 2 | -0.848200878 | 0.133816 |
| F8W9P4 | PIK3CD | Phosphatidylinositol-4,5-bisphosphate 3-kinase | -0.329291582 | -0.661694 |
| F8W9X7 | CCDC93 | Coiled-coil domain-containing protein 93 | -0.055527369 | 0.171307 |
| F8WA11 | CLASP1 | CLIP-associating protein 1 | -0.616991917 | 0.505682 |
| F8WA39 | MTMR1 | Phosphatidylinositol-3,5-bisphosphate 3-phosphatase | 0.166723092 | 0.180915 |
| F8WBA3 | PRKD1 | Serine/threonine-protein kinase | -0.06632177 | -1.05724 |
| F8WCD0 | RNF149 | E3 ubiquitin-protein ligase RNF149 | -0.926028887 | -2.86954 |
| F8WCJ7 | SEC62 | Translocation protein SEC62 | -1.449759324 | 0.0215575 |
| F8WE88 | MYO5A | Unconventional myosin-Va | -0.443629583 | -0.949238 |
| F8WEC0 | MMADHC | Cobalamin-trafficking protein CblD | -0.021226645 | -0.0418606 |
| F8WEF8 | ESS2 | Splicing factor ESS-2 homolog | 0.429380735 | -0.889449 |
| F8WJN3 | CPSF6 | Cleavage and polyadenylation specificity factor subunit 6 | -0.863530159 | 0.201109 |
| G0XQ39 | STIM1 | STIM1L | -1.751387199 | -3.36703 |
| G3V0I5 | NDUFV1 | NADH dehydrogenase [ubiquinone] flavoprotein 1, mitochondrial | -0.967881918 | 0.155484 |
| G3V169 | CASP1 | Caspase 1, apoptosis-related cysteine peptidase (Interleukin 1, beta, convertase), isoform CRA_c | 0.438953956 | 0.297804 |
| G3V180 | DPP3 | Dipeptidyl peptidase 3 | 0.160626094 | 0.0468559 |
| G3V1A6 | GSDMD | Gasdermin domain containing 1, isoform CRA_d | 0.253623009 | 0.57266 |
| G3V1P3 | BORCS5 | BLOC-1-related complex subunit 5 | -0.429136197 | -1.62809 |
| G3V1P5 | MED15 | Mediator of RNA polymerase II transcription subunit 15 | 0.166725636 | -2.17878 |
| G3V1R2 | RTN4IP1 | Reticulon-4-interacting protein 1, mitochondrial | 0.098924915 | -2.11935 |
| G3V1S4 | FCF1 | Chromosome 14 open reading frame 111, isoform CRA_a | -0.694137573 | -0.964259 |
| G3V1S7 | HAS1 | Hyaluronan synthase 1 | 0.363145034 | -0.522425 |
| G3V1U5 | GOLT1B | Golgi transport 1 homolog B (S. cerevisiae), isoform CRA_c | -1.637353261 | -1.89679 |
| G3V235 | CCNK | Cyclin-K (Fragment) | -0.407037735 | -0.203411 |
| G3V256 | ZC3H14 | Zinc finger CCCH domain-containing protein 14 | -0.350393852 | 0.436251 |
| G3V287 | SPATA7 | Spermatogenesis-associated protein 7 | 0.436699231 | -0.240859 |
| G3V2D5 | ZFP36L1 | mRNA decay activator protein ZFP36L1 (Fragment) | -0.236744006 | -0.425931 |
| G3V2M0 | DNAL1 | Dynein axonemal light chain 1 | -0.410574039 | 0 |
| G3V3S1 | DCAF5 | DDB1- and CUL4-associated factor 5 (Fragment) | -0.17650334 | 0 |
| G3V4K3 | VIPAS39 | Spermatogenesis-defective protein 39 homolog | -0.661680778 | -2.2034 |
| G3V4M9 | ITPK1 | Inositol-1,3,4-trisphosphate 5/6-kinase | 0.190358798 | 0.0841996 |
| G3V529 | DDX24 | RNA helicase | -0.520388603 | -0.0449254 |
| G3V570 | MAX | Protein max (Fragment) | -0.256825924 | 0.55251 |
| G3V583 | FAM177A1 | Protein FAM177A1 (Fragment) | -0.220639706 | 0.286811 |
| G3V5N8 | ZFYVE1 | Zinc finger FYVE domain-containing protein 1 | -0.35138615 | -0.136045 |
| G3V5R9 | KLC1 | Kinesin light chain | -0.173505942 | -0.0330181 |
| G3V5T9 | CDK2 | Cyclin-dependent kinase 2 | -0.199435711 | -0.290133 |
| G3V5Z7 | PSMA6 | Proteasome subunit alpha type | -0.356411298 | 0.0692024 |
| G3XAH6 | PAPOLA | Poly(A) polymerase | 0.007653157 | 0.113492 |
| G3XAI2 | LAMB1 | Laminin subunit beta-1 | -0.031508764 | 0.349096 |
| G3XAI9 | FAM133B | Protein FAM133B | -0.020856222 | -0.694906 |
| G3XAN4 | TRAM1 | Translocating chain-associated membrane protein 1 | -1.747449716 | -0.890369 |
| G5E931 | BLOC1S5 | Biogenesis of lysosome-related organelles complex 1 subunit 5 | -1.099741141 | 1.24779 |
| G5E975 | SMARCB1 | SWI/SNF-related matrix-associated actin-dependent regulator of chromatin subfamily B member 1 | -0.974740585 | 0.476783 |
| G5E9A6 | USP11 | Ubiquitin carboxyl-terminal hydrolase | 0.065693617 | -0.429761 |
| G5E9C8 | SOS1 | Son of sevenless homolog 1 | 0.311322133 | -2.77172 |
| G5E9L0 | ARFGAP2 | ADP-ribosylation factor GTPase-activating protein 2 | 0.072811445 | 0.0954126 |
| G5E9W3 | CPSF3 | Cleavage and polyadenylation specific factor 3, 73kDa, isoform CRA_b | -0.448558172 | -0.879441 |
| G5E9Y1 | APBB2 | Amyloid beta (A4) protein-binding, family B, member 2 (Fe65-like), isoform CRA_c | -0.028384368 | -0.203299 |
| G5E9Z2 | CLPTM1L | Cisplatin resistance related protein CRR9p, isoform CRA_c | 0.045363744 | 2.36551 |
| G5EA06 | MRPS27 | 28S ribosomal protein S27, mitochondrial | -1.157429298 | 0.600581 |
| G5EA30 | CELF1 | CUG triplet repeat, RNA binding protein 1, isoform CRA_c | -1.329144637 | -2.59749 |
| G5EA37 | WIPI1 | WD repeat domain phosphoinositide-interacting protein 1 | 0.178691467 | -0.032684 |
| G5EA42 | TMOD2 | Tropomodulin 2 (Neuronal), isoform CRA_a | -1.421792984 | 1.81114 |

| | | | | |
|---|---|---|---|---|
| G8JL96 | PTPRS | Protein-tyrosine-phosphatase | -0.742242972 | -1.43091 |
| G8JLB3 | PUS1 | tRNA pseudouridine synthase A (Fragment) | -0.012717724 | -0.00242043 |
| G8JLB6 | HNRNPH1 | Heterogeneous nuclear ribonucleoprotein H | -0.121375879 | 0.153647 |
| G8JLD3 | ERC1 | ELKS/Rab6-interacting/CAST family member 1 | -0.189198812 | -0.291053 |
| G8JLG1 | SMC1A | Structural maintenance of chromosomes protein | -0.559820493 | 0.391919 |
| G8JLG2 | CDSN | Corneodesmosin | -0.022985379 | 0.192425 |
| H0UI80 | NELFCD | Negative elongation factor C/D | -0.349998236 | 0.0290101 |
| H0UID0 | RAD51D | DNA repair protein RAD51 homolog 4 | -0.219826937 | 0.793981 |
| H0Y368 | DPM1 | Dolichol-phosphate mannosyltransferase subunit 1 | -1.451378028 | 0.348895 |
| H0Y3G1 | PXDC1 | PX domain-containing protein 1 (Fragment) | 0.653256178 | 0 |
| H0Y4E8 | SIK3 | Non-specific serine/threonine protein kinase | -0.588021119 | -1.63391 |
| H0Y552 | TBC1D8B | TBC1 domain family member 8B (Fragment) | 0.906493028 | 4.33014 |
| H0Y5C6 | FLNA | Filamin-A (Fragment) | -0.054252942 | 0 |
| H0Y5D8 | ARHGAP12 | Rho GTPase-activating protein 12 (Fragment) | -0.387735287 | 0 |
| H0Y614 | UFM1 | Ubiquitin-fold modifier 1 (Fragment) | -0.333942254 | -0.188859 |
| H0Y6A0 | ARFGAP3 | ADP-ribosylation factor GTPase-activating protein 3 (Fragment) | -0.176718235 | 0.466602 |
| H0Y6U5 | NCAPG2 | Condensin-2 complex subunit G2 (Fragment) | 1.141340971 | 2.8848 |
| H0Y6V6 | LMBR1 | Limb region 1 protein homolog (Fragment) | 0 | -0.636274 |
| H0Y6Y0 | BMT2 | S-adenosylmethionine sensor upstream of mTORC1 (Fragment) | 0.305279493 | -0.517793 |
| H0Y760 | RAD54L2 | Helicase ARIP4 (Fragment) | -0.371276299 | -0.777273 |
| H0Y7F7 | CEP350 | Centrosome-associated protein 350 (Fragment) | 0 | 0.653308 |
| H0Y831 | MEDAG | Mesenteric estrogen-dependent adipogenesis protein (Fragment) | -0.206225395 | 0.813515 |
| H0Y8B3 | AHCYL2 | Adenosylhomocysteinase 3 (Fragment) | 2.742706779 | 0 |
| H0Y8C3 | MTCH1 | Mitochondrial carrier homolog 1 (Fragment) | -1.385673205 | -1.11924 |
| H0Y8P0 | TBC1D1 | TBC1 domain family member 1 (Fragment) | -0.136059046 | -1.48865 |
| H0Y8X4 | DNPH1 | 2'-deoxynucleoside 5'-phosphate N-hydrolase 1 (Fragment) | -0.07632192 | 0.202472 |
| H0Y928 | TXNDC15 | Thioredoxin domain-containing protein 15 (Fragment) | 0.765211105 | 0 |
| H0Y938 | COPB2 | Coatomer subunit beta' | -0.009698709 | -0.257299 |
| H0Y966 | CEP192 | Centrosomal protein of 192 kDa (Fragment) | -0.283657153 | -0.413485 |
| H0Y987 | PGM3 | Phosphoacetylglucosamine mutase | 0.036836306 | 0.0651464 |
| H0Y993 | DEK | DEK oncogene (DNA binding), isoform CRA_b | 0.142384052 | 0.00503604 |
| H0Y9X1 | TMA16 | Translation machinery-associated protein 16 (Fragment) | -0.088981152 | -0.374951 |
| H0YAN8 | ARHGEF10 | Rho guanine nucleotide exchange factor 10 (Fragment) | -0.118569851 | -0.104772 |
| H0YB91 | IDS | Iduronate 2-sulfatase (Fragment) | -2.197398901 | 0 |
| H0YD03 | RAB3IP | Rab-3A-interacting protein (Fragment) | -0.503730377 | 0.631789 |
| H0YD48 | PICALM | Phosphatidylinositol-binding clathrin assembly protein (Fragment) | -0.58893315 | -1.68547 |
| H0YD93 | RUFY2 | RUN and FYVE domain-containing protein 2 (Fragment) | -0.628127575 | 0.346639 |
| H0YDK7 | RAB30 | Ras-related protein Rab-30 (Fragment) | -0.948269208 | -0.0474011 |
| H0YDT6 | EIF3F | Eukaryotic translation initiation factor 3 subunit F | 0.027417024 | 0.125734 |
| H0YDU8 | PPP5C | Serine/threonine-protein phosphatase (Fragment) | -0.022547245 | -0.00563876 |
| H0YEB6 | ZNRD2 | Protein ZNRD2 (Fragment) | 0.360432784 | -0.400118 |
| H0YF29 | C8orf82 | UPF0598 protein C8orf82 (Fragment) | -2.791054279 | 8.10284 |
| H0YFD6 | HADHA | Enoyl-CoA hydratase | -1.050168912 | 0.327209 |
| H0YG27 | TRIM65 | Tripartite motif-containing protein 65 (Fragment) | -0.564020554 | 0.996392 |
| H0YGN5 | | Cysteine desulfurase, mitochondrial (Fragment) | -2.716954708 | -0.38985 |
| H0YHS6 | YARS2 | Tyrosine--tRNA ligase (Fragment) | -2.328055223 | 0.123197 |
| H0YI09 | METTL7A | Methyltransferase-like protein 7A (Fragment) | -0.99996225 | -0.139435 |
| H0YJ75 | PPP2R5C | Serine/threonine-protein phosphatase 2A 56 kDa regulatory subunit gamma isoform (Fragment) | -0.309808175 | 0.0919701 |
| H0YJ90 | PIP4P1 | Phosphatidylinositol-4,5-bisphosphate 4-phosphatase (Fragment) | -0.603686412 | -0.291432 |
| H0YK61 | EMC4 | ER membrane protein complex subunit 4 | -4.150526365 | -2.09029 |
| H0YKJ4 | TPM1 | Tropomyosin 1 (Alpha), isoform CRA_b | 0.270257235 | -0.0960531 |
| H0YKJ9 | TSPAN3 | Tetraspanin-3 | -1.252798955 | 0.202635 |
| H0YKN8 | TLE3 | Transducin-like enhancer protein 3 | -0.09952275 | 0.0132736 |
| H0YKY0 | CINP | Cyclin-dependent kinase 2-interacting protein | 0.302195628 | 0.93817 |
| H0YL99 | MRPS11 | 28S ribosomal protein S11, mitochondrial | -2.003140728 | 0.867113 |
| H0YLG5 | RMDN3 | Regulator of microtubule dynamics protein 3 (Fragment) | -1.24662598 | 0.974987 |
| H0YLI7 | AP3S2 | AP-3 complex subunit sigma-2 | 0.375455379 | 0.863923 |
| H0YLM8 | DMXL2 | DmX-like protein 2 | -0.277932008 | -0.632411 |
| H0YLY0 | CDC42BPB | Non-specific serine/threonine protein kinase (Fragment) | -0.750294526 | -1.03611 |
| H0YM23 | ANKRD17 | Ankyrin repeat domain-containing protein 17 (Fragment) | -0.141779105 | 0.223911 |
| H0YMD1 | LDLR | Low-density lipoprotein receptor | -1.116830428 | -0.711083 |
| H0YMV8 | RPS27L | 40S ribosomal protein S27 | -0.907710314 | -0.342712 |
| H0YMW2 | AKAP13 | A-kinase anchor protein 13 (Fragment) | -0.482379198 | 0.67535 |
| H0YNA7 | SPPL2A | Signal peptide peptidase-like 2A (Fragment) | -2.077066263 | -1.03965 |
| H0YNG3 | SEC11A | Signal peptidase complex catalytic subunit SEC11 | -1.132181883 | 0.244584 |
| H0YNP9 | HDDC3 | Guanosine-3',5'-bis(diphosphate) 3'-pyrophosphohydrolase MESH1 | -0.201348464 | -1.63489 |
| H0YNU7 | CERS2 | Ceramide synthase 2 | -1.206875801 | 0.354451 |
| H3BLU7 | AKR7A2 | Aflatoxin B1 aldehyde reductase member 2 (Fragment) | 0.085295836 | -0.0658576 |
| H3BLV9 | SRPK1 | SRSF protein kinase 1 (Fragment) | -0.718742847 | -0.352295 |

| | | | | |
|---|---|---|---|---|
| H3BM14 | NUB1 | NEDD8 ultimate buster 1 | 0.29334863 | -0.471162 |
| H3BM21 | | Integrin beta (Fragment) | -0.559290489 | -0.667041 |
| H3BM39 | KDM8 | Bifunctional peptidase and arginyl-hydroxylase JMJD5 (Fragment) | -0.587723573 | -1.15578 |
| H3BMD8 | ARPP19 | cAMP-regulated phosphoprotein 19 | -0.554999193 | -0.559512 |
| H3BMF4 | SPNS1 | Protein spinster homolog 1 | -1.487767577 | 0.361434 |
| H3BML4 | | Ig-like domain-containing protein (Fragment) | -0.419384877 | 1.51238 |
| H3BMM9 | RNPS1 | RNA-binding protein with serine-rich domain 1 (Fragment) | -0.358131806 | -0.295815 |
| H3BMU1 | IST1 | IST1 homolog (Fragment) | -0.884723425 | 0.0635115 |
| H3BN14 | CALB2 | Calretinin (Fragment) | 0.720170418 | -1.54087 |
| H3BN93 | SCAMP2 | Secretory carrier-associated membrane protein | -1.22605598 | -1.69923 |
| H3BNQ5 | CLK3 | Dual-specificity protein kinase CLK3 (Fragment) | -1.590464036 | 0.193086 |
| H3BNT2 | COQ9 | Ubiquinone biosynthesis protein (Fragment) | 0.336356839 | -0.249698 |
| H3BNT4 | MPHOSPH6 | M-phase phosphoprotein 6 | -0.737435818 | 1.5389 |
| H3BNX8 | COX5A | Cytochrome c oxidase polypeptide Va | -0.570613941 | 0.147083 |
| H3BP16 | HAUS2 | HAUS augmin-like complex subunit 2 | 0.042504152 | 0.215989 |
| H3BP20 | HEXA | Beta-hexosaminidase | -0.177702188 | -0.165922 |
| H3BPE1 | MACF1 | Microtubule-actin cross-linking factor 1, isoforms 1/2/3/5 | -0.225625356 | -0.0484924 |
| H3BPE7 | FUS | RNA-binding protein FUS | -0.434250514 | -0.0107333 |
| H3BPG6 | ZNF598 | RING-type E3 ubiquitin transferase | 0.492450158 | 0.706781 |
| H3BPJ2 | ARL6IP1 | ADP-ribosylation factor-like protein 6-interacting protein 1 (Fragment) | -1.189798911 | -0.382874 |
| H3BPK3 | HAGH | Hydroxyacylglutathione hydrolase, mitochondrial (Fragment) | -0.247052908 | -0.184219 |
| H3BPU4 | TUBGCP4 | Gamma-tubulin complex component (Fragment) | -0.248411576 | 0.610506 |
| H3BQ89 | PHKB | Phosphorylase b kinase regulatory subunit (Fragment) | 1.044313033 | 2.72855 |
| H3BQH3 | KLHDC4 | Kelch domain-containing protein 4 (Fragment) | -1.269753059 | 1.1011 |
| H3BRB3 | KIF22 | Kinesin-like protein | -1.826010942 | -0.675645 |
| H3BRF9 | ZFYVE19 | Abscission/NoCut checkpoint regulator | 0.047886054 | -0.00420411 |
| H3BRG3 | RPUSD1 | RNA pseudouridylate synthase domain-containing protein 1 (Fragment) | -0.223343054 | -1.54813 |
| H3BRL3 | UBFD1 | Ubiquitin domain-containing protein UBFD1 | 0.169991732 | -0.101664 |
| H3BRQ0 | PPCDC | Phosphopantothenoylcysteine decarboxylase | -0.900239547 | 0.168986 |
| H3BRU6 | PCBP2 | Poly(rC)-binding protein 2 (Fragment) | -2.411024491 | 2.21077 |
| H3BS02 | BCKDK | Protein-serine/threonine kinase | -1.413212299 | 0.519874 |
| H3BS74 | ARMC5 | Armadillo repeat-containing protein 5 (Fragment) | 0.260814349 | 1.09979 |
| H3BSW6 | CTU2 | Cytoplasmic tRNA 2-thiolation protein 2 | 0.41421477 | -0.321467 |
| H3BU49 | ARL2BP | ADP-ribosylation factor-like protein 2-binding protein | 0.160725832 | 0.0674594 |
| H3BU86 | STX16-NPEPL1 | STX16-NPEPL1 readthrough (NMD candidate) | 0 | 0.988914 |
| H3BUH9 | GCSH | Glycine cleavage system H protein, mitochondrial (Fragment) | 0.208008766 | -0.451122 |
| H3BUR4 | NOB1 | RNA-binding protein NOB1 | -0.385887702 | 0.0100304 |
| H3BUX2 | CYB5B | Cytochrome b5 type B | -0.99712793 | -0.448865 |
| H3BVI6 | CNN2 | Calponin | -0.218492826 | -1.30642 |
| H6UMI1 | GABARAP | GABARAP-a | -0.280634085 | -0.270849 |
| H7BXD5 | GCA | Grancalcin | -0.121325175 | -0.302076 |
| H7BXP1 | NKIRAS2 | NF-kappa-B inhibitor-interacting Ras-like protein 2 | -1.454944253 | 0 |
| H7BXY3 | DHX30 | RNA helicase | -1.045108954 | 0.215868 |
| H7BYF2 | NUP58 | Nucleoporin p58/p45 (Fragment) | -0.093688965 | -0.0550782 |
| H7BYT1 | CSNK1D | Casein kinase I isoform delta | -1.08639582 | -0.308017 |
| H7BYV6 | BIN3 | Bridging integrator 3 | 0.278874 | -0.544122 |
| H7BZF8 | HAUS7 | HAUS augmin-like complex subunit 7 (Fragment) | -0.155303955 | 0 |
| H7BZJ3 | PDIA3 | Protein disulfide-isomerase A3 (Fragment) | 0.287753423 | 0.322546 |
| H7BZK4 | SLC20A1 | Sodium-dependent phosphate transporter 1 (Fragment) | -1.231309255 | -0.0842409 |
| H7BZW6 | SAP18 | Histone deacetylase complex subunit SAP18 (Fragment) | -0.162863811 | -0.200754 |
| H7C096 | BNIP2 | BCL2/adenovirus E1B 19 kDa protein-interacting protein 2 (Fragment) | -1.172911882 | -2.17611 |
| H7C0D5 | YIPF4 | Protein YIPF4 (Fragment) | -0.836254994 | 0.10875 |
| H7C144 | ACTN4 | Alpha-actinin-4 | -0.900795221 | -0.971544 |
| H7C1J4 | UHRF1BP1 | Chromosome 6 open reading frame 107, isoform CRA_b | 0.5117987 | 0.712291 |
| H7C1U3 | CC2D1B | Coiled-coil and C2 domain-containing protein 1B (Fragment) | -0.112129132 | -0.371941 |
| H7C1U8 | APOO | MICOS complex subunit (Fragment) | -1.456947962 | 0.40175 |
| H7C1X9 | CNOT10 | CCR4-NOT transcription complex subunit 10 (Fragment) | 1.308561007 | -0.500098 |
| H7C276 | PLCL2 | Phosphoinositide phospholipase C (Fragment) | 0.05954965 | -1.26049 |
| H7C2Q8 | EBNA1BP2 | EBNA1 binding protein 2, isoform CRA_d | -1.536870162 | -0.265729 |
| H7C3D5 | RETREG2 | Reticulophagy regulator 2 (Fragment) | -1.112264792 | -0.131756 |
| H7C3L3 | ANKRD54 | Ankyrin repeat domain-containing protein 54 (Fragment) | -0.387397607 | 1.18153 |
| H7C3M7 | FARP2 | FERM, ARHGEF and pleckstrin domain-containing protein 2 (Fragment) | -2.340379238 | -1.75981 |
| H7C4F6 | KPNA4 | Importin subunit alpha | 0.04140679 | -0.12182 |
| H7C4W1 | ITGB5 | Integrin beta (Fragment) | -0.272094846 | 0 |
| H7C4W3 | SYP | Synaptophysin (Fragment) | 0.340381384 | -0.940743 |
| H7C5E4 | XRN1 | 5'-3' exoribonuclease 1 (Fragment) | 0.106594404 | -0.559229 |

| | | | | |
|---|---|---|---|---|
| H9ZYI9 | GABPB1 | GA binding protein transcription factor beta subunit 1 transcript variant gamma-2 | -0.186079979 | -0.258 |
| I3L0H8 | DDX19A | ATP-dependent RNA helicase DDX19A | 0.055976868 | 0.0235669 |
| I3L0N3 | NSF | Vesicle-fusing ATPase | -0.206986825 | -0.219335 |
| I3L0R8 | DERL2 | Derlin (Fragment) | -2.487548908 | 0.491969 |
| I3L0X5 | SPAG7 | Sperm-associated antigen 7 | -0.107823372 | -0.199925 |
| I3L1D9 | CTDNEP1 | CTD nuclear envelope phosphatase 1 (Fragment) | -1.284921964 | -0.300808 |
| I3L1P8 | SLC25A11 | Mitochondrial 2-oxoglutarate/malate carrier protein (Fragment) | -1.294185321 | 0.369077 |
| I3L1Q3 | ELP5 | Elongator complex protein 5 (Fragment) | -0.31578207 | 0.589906 |
| I3L1Y9 | FLYWCH2 | FLYWCH family member 2 | 0.003920317 | 0.121559 |
| I3L295 | MPDU1 | Mannose-P-dolichol utilization defect 1 isoform 2 | -1.549813509 | -0.125737 |
| I3L2C7 | GEMIN4 | Gem-associated protein 4 | -1.207582434 | -0.835389 |
| I3L2X1 | MRPL20 | 39S ribosomal protein L20, mitochondrial (Fragment) | 0.349962155 | 0.886099 |
| I3L397 | EIF5A | Eukaryotic translation initiation factor 5A (Fragment) | -0.034080346 | 0.132974 |
| I3L448 | ATPAF1 | ATP synthase mitochondrial F1 complex assembly factor 1 | -0.271271149 | -0.0232627 |
| I3L4C2 | BAIAP2 | Brain-specific angiogenesis inhibitor 1-associated protein 2 | -0.560525338 | -0.452577 |
| I3L4I4 | TOM1L1 | TOM1-like protein 1 | 0.090093215 | 0.114849 |
| I3L4N8 | ACTG1 | Actin, cytoplasmic 2 | 0.203175386 | 0.639414 |
| I3VM54 | KDM2A | Lysine-specific demethylase 2A | -0.933841864 | -0.175124 |
| J3KMZ8 | DPF2 | Zinc finger protein ubi-d4 | -0.073377132 | -0.417988 |
| J3KN01 | AFDN | Afadin | -0.101826509 | 0.00429598 |
| J3KNB4 | CAMP | Cathelicidin antimicrobial peptide | 1.154016972 | -0.967215 |
| J3KND1 | SAAL1 | Protein SAAL1 | 0.317668597 | -2.3741 |
| J3KNF4 | CCS | Superoxide dismutase copper chaperone | -0.03597943 | 0.134302 |
| J3KNN3 | PHKG2 | Phosphorylase kinase | -0.529940049 | 0.533515 |
| J3KNN7 | BRAP | BRCA1-associated protein | 0.949493229 | 0 |
| J3KNZ9 | FBRS | Probable fibrosin-1 | -0.869860888 | -0.758517 |
| J3KP15 | SRSF2 | Serine/arginine-rich splicing factor 2 (Fragment) | 0.088283539 | -0.0722504 |
| J3KPD3 | RBM7 | RNA binding motif protein 7, isoform CRA_c | -0.850121578 | -0.665446 |
| J3KPH2 | ALOXE3 | Arachidonate lipoxygenase 3, isoform CRA_a | -0.001101812 | -2.49018 |
| J3KPJ3 | CAMKK1 | Calcium/calmodulin-dependent protein kinase kinase 1 | -0.146278779 | -0.731982 |
| J3KPL8 | APBB1 | Amyloid beta precursor protein-binding family B member 1 | -0.847626289 | -3.04657 |
| J3KPP4 | LUC7L3 | Cisplatin resistance-associated overexpressed protein, isoform CRA_b | -0.077284813 | -0.384624 |
| J3KPS0 | DNAJB12 | DnaJ (Hsp40) homolog, subfamily B, member 12, isoform CRA_c | -2.66991353 | 0.376697 |
| J3KPY9 | ANTXR2 | Anthrax toxin receptor | -1.519514561 | 0.735752 |
| J3KQ32 | OLA1 | Obg-like ATPase 1 | -0.059119542 | -0.163298 |
| J3KQ34 | COPS7B | COP9 signalosome complex subunit 7b | -0.053613742 | -0.284831 |
| J3KQ48 | PTRH2 | Aminoacyl-tRNA hydrolase | -0.898461739 | -0.0248467 |
| J3KQ72 | FBXO6 | F-box only protein 6 (Fragment) | 0.78591466 | 2.40595 |
| J3KQA0 | SYT1 | Synaptotagmin | 0.685676654 | 0.910188 |
| J3KQJ1 | SUMF2 | Inactive C-alpha-formylglycine-generating enzyme 2 | -0.258626064 | 0.0318095 |
| J3KQN4 | RPL36A | 60S ribosomal protein L36a | -1.424430052 | 0.0469055 |
| J3KR35 | CCDC12 | Coiled-coil domain containing 12, isoform CRA_a | -0.903848966 | -1.39936 |
| J3KR44 | OTUB1 | Ubiquitin thioesterase | 0.271268209 | 0.305088 |
| J3KRD0 | AMZ2 | Archaemetzincin-2 (Fragment) | -0.6083076 | 0.0364358 |
| J3KRN4 | MAP3K3 | Mitogen-activated protein kinase kinase kinase 3 | 0.271815697 | 0.736512 |
| J3KRP9 | TANC2 | Protein TANC2 (Fragment) | -0.686941544 | 0.462836 |
| J3KS15 | MRPL58 | Peptidyl-tRNA hydrolase ICT1, mitochondrial (Fragment) | -1.834198554 | -1.50835 |
| J3KS22 | DCXR | L-xylulose reductase (Fragment) | -0.248403231 | 0.00323741 |
| J3KSL8 | TRAPPC8 | Trafficking protein particle complex subunit 8 | -0.485426585 | -0.816863 |
| J3KT68 | TMEM97 | Sigma intracellular receptor 2 | -1.786972761 | 0.622597 |
| J3KTA1 | FBXL20 | F-box and leucine-rich repeat protein 20, isoform CRA_a | -1.266753515 | 0.0379664 |
| J3KTA4 | DDX5 | DEAD box protein 5 | -0.560156822 | 0.171629 |
| J3KTE2 | DNHD1 | Dynein heavy chain domain-containing protein 1 (Fragment) | -0.508883794 | 1.24656 |
| J3KTF5 | ECSIT | Evolutionarily conserved signaling intermediate in Toll pathway, mitochondrial (Fragment) | -2.789100806 | 1.10752 |
| J3KTI2 | MIEN1 | Migration and invasion enhancer 1 | 0.281247218 | 0.345395 |
| J3KTL2 | SRSF1 | Serine/arginine-rich splicing factor 1 | 0.060675303 | -0.123163 |
| J3KTM6 | SLC16A3 | Monocarboxylate transporter 4 | -2.112706741 | 0.107953 |
| J3KTN0 | EIF4A1 | Eukaryotic initiation factor 4A-I (Fragment) | -2.158136884 | 0.327801 |
| J3QK89 | CHERP | Calcium homeostasis endoplasmic reticulum protein | -0.282337666 | -0.0423714 |
| J3QKK3 | RAD51C | DNA repair protein RAD51 homolog 3 | 0.444039106 | 0.363697 |
| J3QL56 | SCO1 | Protein SCO1 homolog, mitochondrial | -1.579247077 | -0.720578 |
| J3QL71 | SCRN2 | Secernin-2 | -0.188538233 | -0.150522 |
| J3QLE5 | SNRPN | Small nuclear ribonucleoprotein-associated protein N (Fragment) | -0.183237871 | -0.0791114 |
| J3QLN8 | AURKB | Aurora kinase (Fragment) | -1.835488081 | -1.36788 |
| J3QLR8 | MRPS23 | 28S ribosomal protein S23, mitochondrial | -1.409329414 | 0.253272 |
| J3QLS3 | MRPS7 | 28S ribosomal protein S7, mitochondrial | -2.174348752 | 0.493929 |
| J3QQ67 | RPL18 | 60S ribosomal protein L18 (Fragment) | -0.909362952 | 0.314974 |
| J3QQJ0 | SAP30BP | SAP30-binding protein (Fragment) | 0.204390446 | 0.0857194 |

| | | | | |
|---|---|---|---|---|
| J3QQM5 | LIMD2 | LIM domain-containing protein 2 (Fragment) | -1.786797126 | -0.882846 |
| J3QQW9 | SUZ12 | Polycomb protein SUZ12 | -0.137914658 | 0.0601552 |
| J3QQZ9 | PNPO | Pyridoxal 5'-phosphate synthase (Fragment) | 0.094267289 | -0.182819 |
| J3QR07 | YTHDC1 | YTH domain-containing protein 1 | 0.022571484 | -0.306339 |
| J3QRD3 | ADAP2 | Arf-GAP with dual PH domain-containing protein 2 (Fragment) | 0.529672305 | 0 |
| J3QRG6 | CDKN2A | Cyclin-dependent kinase inhibitor 2A | -0.234217882 | -0.157976 |
| J3QRI1 | TIMM21 | Mitochondrial import inner membrane translocase subunit Tim21 (Fragment) | 0 | 1.72277 |
| J3QRS3 | MYL12A | Myosin regulatory light chain 12A | -0.375929832 | -1.08238 |
| J3QRS9 | ZNF207 | BUB3-interacting and GLEBS motif-containing protein ZNF207 | -0.364603519 | 0.15626 |
| J3QSF7 | MPO | Myeloperoxidase (Fragment) | 0.778512637 | 0.194838 |
| J3QT28 | BUB3 | Mitotic checkpoint protein BUB3 (Fragment) | -0.076882998 | 0.105867 |
| J3QTA2 | BAG1 | BAG family molecular chaperone regulator 1 | 0.879985889 | -0.455655 |
| J9JIC5 | C17orf75 | Protein Njmu-R1 | -0.26693209 | -0.226961 |
| K4DI92 | RWDD4 | RWD domain containing 4A | 0.131292979 | -0.112224 |
| K7EIN2 | NUDT16L1 | Tudor-interacting repair regulator protein (Fragment) | 0.607316732 | 0 |
| K7EIU8 | SMAD4 | Mothers against decapentaplegic homolog | -0.59134531 | 0.13704 |
| K7EIX0 | PLEKHF1 | Pleckstrin homology domain-containing family F member 1 (Fragment) | 0.35032018 | 0.0732002 |
| K7EIY6 | RNF126 | RING-type E3 ubiquitin transferase (Fragment) | -0.139674664 | -0.083602 |
| K7EJ34 | RETREG3 | Reticulophagy regulator 3 | -1.73533233 | 0.736039 |
| K7EJA6 | TPGS2 | Tubulin polyglutamylase complex subunit 2 (Fragment) | 1.519405365 | 0.148343 |
| K7EJB9 | CALR | Calreticulin | -0.610328595 | -0.505933 |
| K7EJE8 | LONP1 | Lon protease homolog, mitochondrial | -0.40241154 | 0.192184 |
| K7EK00 | FAM210A | Protein FAM210A (Fragment) | -1.848335663 | 0 |
| K7EK07 | H3-3B | Histone H3 (Fragment) | -0.724906445 | 0.0201187 |
| K7EK33 | DAZAP1 | DAZ-associated protein 1 | -0.59904178 | 0.188785 |
| K7EK78 | | Complex I-B14.7 | -0.358122031 | 2.27959 |
| K7EKB9 | TCF3 | Transcription factor E2-alpha (Fragment) | 0.642289559 | -0.815148 |
| K7EKI4 | MRPL4 | 39S ribosomal protein L4, mitochondrial | -1.456436237 | -1.73897 |
| K7EKI8 | PPL | Periplakin | -2.664459308 | -4.7819 |
| K7EKQ2 | CDC37 | Hsp90 co-chaperone Cdc37 (Fragment) | -1.624709288 | -0.590467 |
| K7EL68 | CDC37 | Hsp90 co-chaperone Cdc37 (Fragment) | -0.109697898 | -0.329825 |
| K7EL81 | PPP1R12C | Protein phosphatase 1 regulatory subunit 12C (Fragment) | -0.665837447 | 0.517695 |
| K7ELE3 | TLE5 | TLE family member 5 (Fragment) | 2.228825569 | 1.14963 |
| K7ELL7 | PRKCSH | Glucosidase 2 subunit beta | -0.086959362 | 0.0842603 |
| K7ELM1 | ANKRD27 | Ankyrin repeat domain-containing protein 27 | -0.09429733 | 0 |
| K7ELQ9 | TMEM205 | Transmembrane protein 205 (Fragment) | -1.945515871 | -0.0744263 |
| K7ELX9 | INO80C | INO80 complex subunit C | 0.170330365 | 0.318016 |
| K7EM18 | EIF1 | Eukaryotic translation initiation factor 1 | -0.26587224 | -0.20424 |
| K7EMC7 | CYGB | Cytoglobin | 0.60223182 | -0.318262 |
| K7EMS3 | KRT19 | Keratin, type I cytoskeletal 19 (Fragment) | -0.444653829 | -1.13458 |
| K7ENF0 | NARS1 | Asparagine--tRNA ligase, cytoplasmic | -0.526918093 | 0.413564 |
| K7ENI6 | TMEM256-PLSCR3 | TMEM256-PLSCR3 readthrough (NMD candidate) | 0.023281256 | 1.70425 |
| K7ENR6 | PSMG2 | Proteasome assembly chaperone 2 | -0.025412083 | -0.0829964 |
| K7EP14 | ERCC1 | DNA excision repair protein ERCC-1 (Fragment) | -0.783189376 | 0.0913181 |
| K7EP59 | STK11 | Non-specific serine/threonine protein kinase | -1.016096751 | -1.08429 |
| K7EP90 | RBM42 | RNA-binding motif protein 42 | -0.562814395 | -0.682795 |
| K7EQB1 | STX8 | Syntaxin-8 | -1.49320821 | 0 |
| K7ER90 | EIF3G | Eukaryotic translation initiation factor 3 subunit G (Fragment) | 0.165065845 | -0.146062 |
| K7ERA3 | P3H4 | Endoplasmic reticulum protein SC65 (Fragment) | -0.59256204 | -0.625374 |
| K7ERC8 | KDSR | 3-ketodihydrosphingosine reductase | -1.655560652 | -0.565291 |
| K7ERD7 | POP4 | Ribonuclease P protein subunit p29 (Fragment) | -0.672116597 | 0.104125 |
| K7ESN3 | GHDC | GH3 domain-containing protein | -0.745128592 | 0 |
| M0QXX4 | KPTN | KICSTOR complex protein kaptin (Fragment) | 0.009937366 | -0.546109 |
| M0QXZ5 | ZNF428 | Zinc finger protein 428 (Fragment) | 0.201164405 | 1.33478 |
| M0QXZ8 | SYDE1 | Rho GTPase-activating protein SYDE1 | -0.087394079 | 0 |
| M0QY97 | ZC3H4 | Zinc finger CCCH domain-containing protein 4 (Fragment) | -0.71610872 | -0.225527 |
| M0QYC5 | SIN3B | Paired amphipathic helix protein Sin3b | -0.623228391 | 0 |
| M0QYM7 | TUBB4A | Tubulin beta-4A chain (Fragment) | -0.436011632 | -1.14221 |
| M0QYP8 | LIPE | Hormone-sensitive lipase (Fragment) | -0.654183785 | 0 |
| M0QYT5 | USE1 | USE1-like protein (Fragment) | -1.132540862 | 0 |
| M0QYZ2 | AP2S1 | AP complex subunit sigma | -0.695673863 | -0.0258176 |
| M0QZ66 | EMP3 | Epithelial membrane protein 3 (Fragment) | -1.027270555 | 0.903266 |
| M0QZM1 | HNRNPM | Heterogeneous nuclear ribonucleoprotein M (Fragment) | -0.649745305 | 1.34207 |
| M0QZQ2 | MED25 | Mediator complex subunit 25 | 0.199398994 | 0.652184 |
| M0QZR4 | ARHGEF1 | Rho guanine nucleotide exchange factor 1 | -0.057335059 | -0.0673164 |
| M0R050 | EXOSC5 | Exosome complex component RRP46 | -2.027278264 | -0.617041 |
| M0R083 | CD37 | Leukocyte antigen CD37 (Fragment) | -2.933188955 | -1.58122 |
| M0R0C6 | KHSRP | Far upstream element-binding protein 2 | -0.426057021 | 0.404415 |

| | | | | |
|---|---|---|---|---|
| M0R0F0 | RPS5 | 40S ribosomal protein S5 (Fragment) | -0.283803463 | -0.143759 |
| M0R0P8 | MYO9B | Unconventional myosin-IXb | -0.353244146 | -0.274278 |
| M0R117 | RPL18A | 60S ribosomal protein L18a | -1.703864415 | -0.0139128 |
| M0R132 | KLK10 | Kallikrein-10 (Fragment) | -0.5359056 | -0.644536 |
| M0R1E3 | NAT14 | Probable N-acetyltransferase 14 (Fragment) | 1.1762E-05 | 0 |
| M0R208 | CLPP | ATP-dependent Clp protease proteolytic subunit | -0.428155661 | 0.615328 |
| M0R2B7 | POLD1 | DNA polymerase | -0.143716415 | 0.1409 |
| M0R2C6 | | Seryl-tRNA synthetase (Fragment) | -0.663614114 | 0.329833 |
| M0R2Z9 | SUGP2 | SURP and G-patch domain-containing protein 2 | -1.047715664 | -0.162331 |
| M0R366 | FSD1 | Fibronectin type III and SPRY domain-containing protein 1 | -1.019907157 | 1.17833 |
| O00115 | DNASE2 | Deoxyribonuclease-2-alpha | -0.188922246 | 0.10517 |
| O00116 | AGPS | Alkyldihydroxyacetonephosphate synthase, peroxisomal | -1.247130871 | 0.125039 |
| O00148 | DDX39A | ATP-dependent RNA helicase DDX39A | 0.176719824 | -0.143681 |
| O00151 | PDLIM1 | PDZ and LIM domain protein 1 | 0.140046438 | 0.176394 |
| O00154-4 | ACOT7 | Isoform 4 of Cytosolic acyl coenzyme A thioester hydrolase | 0.059449673 | -0.0923888 |
| O00161 | SNAP23 | Synaptosomal-associated protein 23 | -1.185709794 | 0.0682036 |
| O00165 | HAX1 | HCLS1-associated protein X-1 | -0.177255551 | -0.0813182 |
| O00178 | GTPBP1 | GTP-binding protein 1 | -0.186112086 | -0.222199 |
| O00186 | STXBP3 | Syntaxin-binding protein 3 | -1.945220391 | 0.23608 |
| O00194 | RAB27B | Ras-related protein Rab-27B | -0.849368731 | -0.013265 |
| O00203 | AP3B1 | AP-3 complex subunit beta-1 | 0.027383804 | -0.224716 |
| O00204 | SULT2B1 | Sulfotransferase 2B1 | 0.363112926 | 0 |
| O00214 | LGALS8 | Galectin-8 | -0.711428245 | 0.215033 |
| O00221 | NFKBIE | NF-kappa-B inhibitor epsilon | 0.002982696 | 0.766391 |
| O00231 | PSMD11 | 26S proteasome non-ATPase regulatory subunit 11 | -0.08424902 | 0.0238501 |
| O00232 | PSMD12 | 26S proteasome non-ATPase regulatory subunit 12 | -0.240597486 | -0.151272 |
| O00233 | PSMD9 | 26S proteasome non-ATPase regulatory subunit 9 | 0.125108878 | 0.178616 |
| O00244 | ATOX1 | Copper transport protein ATOX1 | -0.429664612 | 0.279063 |
| O00264 | PGRMC1 | Membrane-associated progesterone receptor component 1 | -0.780312141 | 0.86356 |
| O00267-2 | SUPT5H | Isoform 2 of Transcription elongation factor SPT5 | 0.016960939 | 0.0350876 |
| O00273 | DFFA | DNA fragmentation factor subunit alpha | 0.03768762 | 0.298324 |
| O00273-2 | DFFA | Isoform DFF35 of DNA fragmentation factor subunit alpha | -0.270859321 | 0.535748 |
| O00291 | HIP1 | Huntingtin-interacting protein 1 | -0.906516314 | -0.24293 |
| O00299 | CLIC1 | Chloride intracellular channel protein 1 | 0.065499624 | 0.301856 |
| O00391 | QSOX1 | Sulfhydryl oxidase 1 | -0.741859833 | 0.0157102 |
| O00399 | DCTN6 | Dynactin subunit 6 | -0.208300511 | -0.0869935 |
| O00410 | IPO5 | Importin-5 | -0.309579213 | -0.179815 |
| O00418 | EEF2K | Eukaryotic elongation factor 2 kinase | -0.265377045 | -0.237335 |
| O00425 | IGF2BP3 | Insulin-like growth factor 2 mRNA-binding protein 3 | -0.61418581 | 0.039676 |
| O00429-3 | DNM1L | Isoform 2 of Dynamin-1-like protein | -0.095114946 | 1.0271 |
| O00429-4 | DNM1L | Isoform 3 of Dynamin-1-like protein | 0.024278164 | -0.0322049 |
| O00442 | RTCA | RNA 3'-terminal phosphate cyclase | -0.437238614 | -0.695016 |
| O00443 | PIK3C2A | Phosphatidylinositol 4-phosphate 3-kinase C2 domain-containing subunit alpha | -0.513910929 | 1.09191 |
| O00459 | PIK3R2 | Phosphatidylinositol 3-kinase regulatory subunit beta | -0.736172517 | 2.67023 |
| O00461 | GOLIM4 | Golgi integral membrane protein 4 | -1.631487926 | 0.741252 |
| O00462 | MANBA | Beta-mannosidase | -0.300670306 | -0.0984608 |
| O00469-2 | PLOD2 | Isoform 2 of Procollagen-lysine,2-oxoglutarate 5-dioxygenase 2 | -0.091102759 | 0.138775 |
| O00483 | NDUFA4 | Cytochrome c oxidase subunit NDUFA4 | -0.419579109 | -0.84134 |
| O00487 | PSMD14 | 26S proteasome non-ATPase regulatory subunit 14 | -0.379206022 | 0.0563717 |
| O00499-11 | BIN1 | Isoform BIN1+12A of Myc box-dependent-interacting protein 1 | -0.002958457 | 0.129635 |
| O00505 | KPNA3 | Importin subunit alpha-4 | 0.096103112 | -0.0239328 |
| O00506 | STK25 | Serine/threonine-protein kinase 25 | -0.38157018 | 0.646049 |
| O00507 | USP9Y | Probable ubiquitin carboxyl-terminal hydrolase FAF-Y | -1.714901447 | -0.556582 |
| O00560 | SDCBP | Syntenin-1 | -0.44339784 | 0.244764 |
| O00560-2 | SDCBP | Isoform 2 of Syntenin-1 | -0.681651274 | 0.396549 |
| O00562 | PITPNM1 | Membrane-associated phosphatidylinositol transfer protein 1 | -0.266864459 | -0.703898 |
| O00566 | MPHOSPH10 | U3 small nucleolar ribonucleoprotein protein MPP10 | -2.320331415 | 0.683766 |
| O00567 | NOP56 | Nucleolar protein 56 | -0.40029494 | 0.0053587 |
| O00592 | PODXL | Podocalyxin | -1.135222673 | 0.516836 |
| O00622 | CCN1 | CCN family member 1 | -0.529788733 | 0.702589 |
| O00625 | PIR | Pirin | -0.544296026 | -0.345748 |
| O00743 | PPP6C | Serine/threonine-protein phosphatase 6 catalytic subunit | -0.128613154 | -0.105721 |
| O00748 | CES2 | Cocaine esterase | 0.144181967 | 0 |
| O00754 | MAN2B1 | Lysosomal alpha-mannosidase | -0.034287771 | 0.287074 |
| O00762 | UBE2C | Ubiquitin-conjugating enzyme E2 C | -0.431639433 | 0.694696 |
| O00764 | PDXK | Pyridoxal kinase | 0.112040361 | -0.0490023 |
| O14530 | TXNDC9 | Thioredoxin domain-containing protein 9 | -0.277406216 | -0.0355781 |
| O14531 | DPYSL4 | Dihydropyrimidinase-related protein 4 | 0.312973261 | -0.65041 |
| O14545 | TRAFD1 | TRAF-type zinc finger domain-containing protein 1 | 0.321839333 | -0.184172 |
| O14548 | COX7A2L | Cytochrome c oxidase subunit 7A-related protein, mitochondrial | -1.27363054 | 0 |

| | | | | |
|---|---|---|---|---|
| O14579 | COPE | Coatomer subunit epsilon | 0.167501132 | 0.339303 |
| O14613 | CDC42EP2 | Cdc42 effector protein 2 | 0.278507471 | 0 |
| O14617 | AP3D1 | AP-3 complex subunit delta-1 | -0.317033132 | -0.67527 |
| O14633 | LCE2B | Late cornified envelope protein 2B | -1.234559576 | 0 |
| O14641 | DVL2 | Segment polarity protein dishevelled homolog DVL-2 | 0.000137488 | 0.0170403 |
| O14646 | CHD1 | Chromodomain-helicase-DNA-binding protein 1 | -0.091251532 | -0.141905 |
| O14656 | TOR1A | Torsin-1A | -1.029215574 | -0.148521 |
| O14657 | TOR1B | Torsin-1B | -0.720442851 | 0.120445 |
| O14672 | ADAM10 | Disintegrin and metalloproteinase domain-containing protein 10 | -1.161134561 | 0.0307744 |
| O14715 | RGPD8 | RANBP2-like and GRIP domain-containing protein 8 | -0.637629747 | -0.528397 |
| O14732 | IMPA2 | Inositol monophosphatase 2 | -0.94311738 | -0.96189 |
| O14737 | PDCD5 | Programmed cell death protein 5 | 0.223827362 | 0.216858 |
| O14744 | PRMT5 | Protein arginine N-methyltransferase 5 | 0.001914501 | -0.165307 |
| O14745 | SLC9A3R1 | Na(+)/H(+) exchange regulatory cofactor NHE-RF1 | 0.092651208 | -0.0649153 |
| O14777 | NDC80 | Kinetochore protein NDC80 homolog | -0.190916618 | 0.040871 |
| O14787 | TNPO2 | Transportin-2 | -0.252167463 | -0.60775 |
| O14802 | POLR3A | DNA-directed RNA polymerase III subunit RPC1 | 0.104058027 | 1.96001 |
| O14818 | PSMA7 | Proteasome subunit alpha type-7 | -0.091671626 | -0.148697 |
| O14828 | SCAMP3 | Secretory carrier-associated membrane protein 3 | -1.716491063 | 0.38622 |
| O14841 | OPLAH | 5-oxoprolinase | -0.873660405 | -0.482967 |
| O14880 | MGST3 | Microsomal glutathione S-transferase 3 | -1.966540416 | -0.052693 |
| O14907 | TAX1BP3 | Tax1-binding protein 3 | -0.038186868 | 0.763462 |
| O14908 | GIPC1 | PDZ domain-containing protein GIPC1 | -0.009971062 | 0.208459 |
| O14920 | IKBKB | Inhibitor of nuclear factor kappa-B kinase subunit beta | -0.316159328 | 0.0587905 |
| O14925 | TIMM23 | Mitochondrial import inner membrane translocase subunit Tim23 | -1.771750291 | 0.00308545 |
| O14929 | HAT1 | Histone acetyltransferase type B catalytic subunit | 0.203310172 | -0.0545031 |
| O14949 | UQCRQ | Cytochrome b-c1 complex subunit 8 | -1.180510044 | 0.284081 |
| O14965 | AURKA | Aurora kinase A | -1.353513877 | 0.37171 |
| O14966 | RAB29 | Ras-related protein Rab-7L1 | -0.89506348 | -0.164024 |
| O14967 | CLGN | Calmegin | -1.800003052 | -0.297671 |
| O14974 | PPP1R12A | Protein phosphatase 1 regulatory subunit 12A | 0.169372797 | -0.00339413 |
| O14976 | GAK | Cyclin-G-associated kinase | 0.191565673 | 0.0869242 |
| O14980 | XPO1 | Exportin-1 | -0.182929834 | -0.31509 |
| O15027 | SEC16A | Protein transport protein Sec16A | 0.139482975 | 0.224662 |
| O15031 | PLXNB2 | Plexin-B2 | -1.606471459 | 0.0716098 |
| O15040 | TECPR2 | Tectonin beta-propeller repeat-containing protein 2 | -2.290268262 | 0.846581 |
| O15042 | U2SURP | U2 snRNP-associated SURP motif-containing protein | -0.182819049 | 0.0342315 |
| O15061 | SYNM | Synemin | -0.302623987 | -0.0786603 |
| O15066 | KIF3B | Kinesin-like protein KIF3B | -0.089863936 | 0.106473 |
| O15067 | PFAS | Phosphoribosylformylglycinamidine synthase | 0.024102052 | 0.0382353 |
| O15069 | NACAD | NAC-alpha domain-containing protein 1 | -0.171027501 | 0.092377 |
| O15084 | ANKRD28 | Serine/threonine-protein phosphatase 6 regulatory ankyrin repeat subunit A | -0.337103685 | 0.00754038 |
| O15085 | ARHGEF11 | Rho guanine nucleotide exchange factor 11 | 0.62436835 | 1.67779 |
| O15111 | CHUK | Inhibitor of nuclear factor kappa-B kinase subunit alpha | -0.147968292 | -1.86714 |
| O15116 | LSM1 | U6 snRNA-associated Sm-like protein LSm1 | 0.644999345 | 0.580016 |
| O15118 | NPC1 | NPC intracellular cholesterol transporter 1 | -1.237497965 | 0.216843 |
| O15121 | DEGS1 | Sphingolipid delta(4)-desaturase DES1 | -0.010395686 | -4.99015 |
| O15143 | ARPC1B | Actin-related protein 2/3 complex subunit 1B | -0.211155176 | -0.123048 |
| O15144 | ARPC2 | Actin-related protein 2/3 complex subunit 2 | -0.276384433 | -0.0853341 |
| O15145 | ARPC3 | Actin-related protein 2/3 complex subunit 3 | -0.356915951 | -0.135434 |
| O15160 | POLR1C | DNA-directed RNA polymerases I and III subunit RPAC1 | -0.00537475 | 0.163579 |
| O15164 | TRIM24 | Transcription intermediary factor 1-alpha | 0.042049964 | 0.122521 |
| O15173 | PGRMC2 | Membrane-associated progesterone receptor component 2 | -1.1006066 | 0.451473 |
| O15211 | RGL2 | Ral guanine nucleotide dissociation stimulator-like 2 | -0.119223515 | -0.237319 |
| O15212 | PFDN6 | Prefoldin subunit 6 | 0.350050131 | 0.134993 |
| O15213 | WDR46 | WD repeat-containing protein 46 | -0.472607931 | -1.27368 |
| O15228 | GNPAT | Dihydroxyacetone phosphate acyltransferase | -0.702508132 | 0 |
| O15230 | LAMA5 | Laminin subunit alpha-5 | -1.206911087 | 1.26223 |
| O15231 | ZNF185 | Zinc finger protein 185 | 0.168994268 | 0.25347 |
| O15231-3 | ZNF185 | Isoform 3 of Zinc finger protein 185 | 0.088818789 | 0.394813 |
| O15231-6 | ZNF185 | Isoform 6 of Zinc finger protein 185 | 0.100441853 | 0.0253197 |
| O15234 | CASC3 | Protein CASC3 | 2.763705174 | 0 |
| O15254 | ACOX3 | Peroxisomal acyl-coenzyme A oxidase 3 | -0.176031828 | 0 |
| O15258 | RER1 | Protein RER1 | -1.229771932 | 0.253215 |
| O15260 | SURF4 | Surfeit locus protein 4 | -1.788097223 | 0.0991586 |
| O15264 | MAPK13 | Mitogen-activated protein kinase 13 | -0.54220589 | 0.714139 |
| O15269 | SPTLC1 | Serine palmitoyltransferase 1 | -1.38229084 | 0.165038 |
| O15270 | SPTLC2 | Serine palmitoyltransferase 2 | -2.013830026 | 0.52721 |
| O15294 | OGT | UDP-N-acetylglucosamine--peptide N-acetylglucosaminyltransferase 110 kDa subunit | 0.165620486 | -0.0161877 |

| | | | | |
|---|---|---|---|---|
| O15294-3 | OGT | Isoform 1 of UDP-N-acetylglucosamine--peptide N-acetylglucosaminyltransferase 110 kDa subunit | -0.45714728 | -0.795712 |
| O15305 | PMM2 | Phosphomannomutase 2 | 0.135084947 | 0.0499312 |
| O15355 | PPM1G | Protein phosphatase 1G | 0.169898192 | 0.0934747 |
| O15357 | INPPL1 | Phosphatidylinositol 3,4,5-trisphosphate 5-phosphatase 2 | 0.079725345 | -0.213744 |
| O15371 | EIF3D | Eukaryotic translation initiation factor 3 subunit D | -0.073581219 | -0.233793 |
| O15397 | IPO8 | Importin-8 | 0.056934357 | -0.284496 |
| O15400 | STX7 | Syntaxin-7 | -0.464346647 | 0.312332 |
| O15446 | POLR1G | DNA-directed RNA polymerase I subunit RPA34 | -2.189594269 | 0.984498 |
| O15460-2 | P4HA2 | Isoform IIa of Prolyl 4-hydroxylase subunit alpha-2 | -0.210509777 | 0.178684 |
| O15511 | ARPC5 | Actin-related protein 2/3 complex subunit 5 | -0.629792849 | -0.0212466 |
| O15541 | RNF113A | E3 ubiquitin-protein ligase RNF113A | -0.43434914 | -0.29325 |
| O43143 | DHX15 | ATP-dependent RNA helicase DHX15 | -0.064758778 | -0.0409355 |
| O43148 | RNMT | mRNA cap guanine-N7 methyltransferase | -0.079788367 | -0.0019118 |
| O43149 | ZZEF1 | Zinc finger ZZ-type and EF-hand domain-containing protein 1 | -1.297786236 | 0 |
| O43150 | ASAP2 | Arf-GAP with SH3 domain, ANK repeat and PH domain-containing protein 2 | -0.139461358 | -0.755717 |
| O43156 | TTI1 | TELO2-interacting protein 1 homolog | -0.205732346 | -0.0948903 |
| O43164 | PJA2 | E3 ubiquitin-protein ligase Praja-2 | -0.733992577 | 0.784125 |
| O43167 | ZBTB24 | Zinc finger and BTB domain-containing protein 24 | -2.637863914 | 2.4855 |
| O43172 | PRPF4 | U4/U6 small nuclear ribonucleoprotein Prp4 | -0.46470348 | 0.130091 |
| O43187 | IRAK2 | Interleukin-1 receptor-associated kinase-like 2 | 0.15583396 | 0.307284 |
| O43237 | DYNC1LI2 | Cytoplasmic dynein 1 light intermediate chain 2 | -0.215296189 | 0.348254 |
| O43242 | PSMD3 | 26S proteasome non-ATPase regulatory subunit 3 | -0.153512001 | -0.161198 |
| O43252 | PAPSS1 | Bifunctional 3'-phosphoadenosine 5'-phosphosulfate synthase 1 | 0.108466705 | -0.0583566 |
| O43264 | ZW10 | Centromere/kinetochore protein zw10 homolog | -0.193682273 | -0.206617 |
| O43281 | EFS | Embryonal Fyn-associated substrate | -0.743592421 | -0.595391 |
| O43290 | SART1 | U4/U6.U5 tri-snRNP-associated protein 1 | -0.64325889 | 0.0830243 |
| O43293 | DAPK3 | Death-associated protein kinase 3 | -2.766783237 | 0.213597 |
| O43294 | TGFB1I1 | Transforming growth factor beta-1-induced transcript 1 protein | -0.033116897 | 0.293878 |
| O43299 | AP5Z1 | AP-5 complex subunit zeta-1 | -0.190864881 | 0.00601355 |
| O43318 | MAP3K7 | Mitogen-activated protein kinase kinase kinase 7 | -0.071236213 | 0.173163 |
| O43353 | RIPK2 | Receptor-interacting serine/threonine-protein kinase 2 | 0.352035205 | 0.0681534 |
| O43390 | HNRNPR | Heterogeneous nuclear ribonucleoprotein R | -0.625068823 | 0.498947 |
| O43395 | PRPF3 | U4/U6 small nuclear ribonucleoprotein Prp3 | -0.324108839 | -0.340699 |
| O43396 | TXNL1 | Thioredoxin-like protein 1 | -0.055414995 | 0.1092 |
| O43399-2 | TPD52L2 | Isoform 2 of Tumor protein D54 | -0.879042069 | -1.15652 |
| O43399 | TPD52L2 | Tumor protein D54 | -0.591855367 | -0.803029 |
| O43427 | FIBP | Acidic fibroblast growth factor intracellular-binding protein | -0.606899341 | 0.713643 |
| O43464 | HTRA2 | Serine protease HTRA2, mitochondrial | -0.061059316 | -0.188843 |
| O43491 | EPB41L2 | Band 4.1-like protein 2 | -0.442706108 | 0.124585 |
| O43524 | FOXO3 | Forkhead box protein O3 | -0.897914807 | 0.292261 |
| O43570 | CA12 | Carbonic anhydrase 12 | -1.353138169 | 0.382273 |
| O43572 | AKAP10 | A-kinase anchor protein 10, mitochondrial | -0.753899813 | -0.602738 |
| O43583 | DENR | Density-regulated protein | 0.197296143 | -0.0348794 |
| O43592 | XPOT | Exportin-T | -0.221659263 | -0.474734 |
| O43615 | TIMM44 | Mitochondrial import inner membrane translocase subunit TIM44 | -0.970297496 | 0.432618 |
| O43617 | TRAPPC3 | Trafficking protein particle complex subunit 3 | 0.041148424 | 0.393646 |
| O43633 | CHMP2A | Charged multivesicular body protein 2a | -0.728641907 | 0.0158777 |
| O43639 | NCK2 | Cytoplasmic protein NCK2 | 0.067615668 | -0.0886691 |
| O43657 | TSPAN6 | Tetraspanin-6 | -1.908113798 | -0.273629 |
| O43660 | PLRG1 | Pleiotropic regulator 1 | -1.007218202 | -0.0466894 |
| O43663 | PRC1 | Protein regulator of cytokinesis 1 | -0.241284768 | 0.501825 |
| O43665 | RGS10 | Regulator of G-protein signaling 10 | -0.127055089 | -0.258663 |
| O43676 | NDUFB3 | NADH dehydrogenase [ubiquinone] 1 beta subcomplex subunit 3 | -2.511215766 | 1.54255 |
| O43678 | NDUFA2 | NADH dehydrogenase [ubiquinone] 1 alpha subcomplex subunit 2 | -1.453037103 | -0.232972 |
| O43681 | GET3 | ATPase GET3 | -0.11253039 | -0.394703 |
| O43683 | BUB1 | Mitotic checkpoint serine/threonine-protein kinase BUB1 | 0.150794188 | -0.208108 |
| O43707 | ACTN4 | Alpha-actinin-4 | 0.175108115 | -0.100475 |
| O43709 | BUD23 | Probable 18S rRNA (guanine-N(7))-methyltransferase | -1.136076132 | -0.356345 |
| O43719 | HTATSF1 | HIV Tat-specific factor 1 | -0.091775974 | -0.10323 |
| O43731 | KDELR3 | ER lumen protein-retaining receptor 3 | -1.396573702 | 0 |
| O43741 | PRKAB2 | 5'-AMP-activated protein kinase subunit beta-2 | 1.400880496 | -3.00084 |
| O43747 | AP1G1 | AP-1 complex subunit gamma-1 | -0.288483858 | -0.466671 |
| O43776 | NARS1 | Asparagine--tRNA ligase, cytoplasmic | -0.363302072 | 0.177958 |
| O43809 | NUDT21 | Cleavage and polyadenylation specificity factor subunit 5 | -0.548070828 | 0.421826 |
| O43813 | LANCL1 | Glutathione S-transferase LANCL1 | -0.362791061 | 0.128318 |
| O43815 | STRN | Striatin | 0.121471246 | -0.0109018 |
| O43818 | RRP9 | U3 small nucleolar RNA-interacting protein 2 | -0.110365868 | 0.0367985 |
| O43847 | NRDC | Nardilysin | 0.009912491 | 0.0650256 |
| O43852-2 | CALU | Isoform 2 of Calumenin | -0.637911638 | -0.0296793 |

| | | | | |
|---|---|---|---|---|
| O43852 | CALU | Calumenin | 0.184068203 | 0.159316 |
| O43854 | EDIL3 | EGF-like repeat and discoidin I-like domain-containing protein 3 | -1.510548115 | 1.20087 |
| O43865 | AHCYL1 | S-adenosylhomocysteine hydrolase-like protein 1 | -0.018551509 | 0.125814 |
| O43896 | KIF1C | Kinesin-like protein KIF1C | -0.204500755 | 1.12785 |
| O43920 | NDUFS5 | NADH dehydrogenase [ubiquinone] iron-sulfur protein 5 | -1.463432471 | 0.350227 |
| O43924 | PDE6D | Retinal rod rhodopsin-sensitive cGMP 3',5'-cyclic phosphodiesterase subunit delta | -0.268455426 | -0.343395 |
| O43929 | ORC4 | Origin recognition complex subunit 4 | -1.078782876 | 0 |
| O60216 | RAD21 | Double-strand-break repair protein rad21 homolog | -1.716290236 | 1.88077 |
| O60231 | DHX16 | Pre-mRNA-splicing factor ATP-dependent RNA helicase DHX16 | -0.059074879 | -0.124166 |
| O60237 | PPP1R12B | Protein phosphatase 1 regulatory subunit 12B | -0.583604852 | -1.51111 |
| O60256 | PRPSAP2 | Phosphoribosyl pyrophosphate synthase-associated protein 2 | -0.193722328 | 0.0478166 |
| O60264 | SMARCA5 | SWI/SNF-related matrix-associated actin-dependent regulator of chromatin subfamily A member 5 | -0.254696131 | 0.0482006 |
| O60271-4 | SPAG9 | Isoform 4 of C-Jun-amino-terminal kinase-interacting protein 4 | 0.15759476 | 0.00512441 |
| O60287 | URB1 | Nucleolar pre-ribosomal-associated protein 1 | 0.3046333 | -0.991689 |
| O60294 | LCMT2 | tRNA wybutosine-synthesizing protein 4 | -0.152500788 | 0.515862 |
| O60306 | AQR | RNA helicase aquarius | -0.421280861 | 0.246532 |
| O60313-2 | OPA1 | Isoform 2 of Dynamin-like 120 kDa protein, mitochondrial | -0.649380048 | 0.0488574 |
| O60318 | MCM3AP | Germinal-center associated nuclear protein | -0.138000488 | 0.902945 |
| O60341 | KDM1A | Lysine-specific histone demethylase 1A | -0.399516503 | -0.191421 |
| O60343 | TBC1D4 | TBC1 domain family member 4 | 0.28925395 | 0.00568453 |
| O60347 | TBC1D12 | TBC1 domain family member 12 | -1.253260056 | -2.0887 |
| O60361 | NME2P1 | Putative nucleoside diphosphate kinase | 0.787675778 | 1.81395 |
| O60443 | GSDME | Gasdermin-E | 0.031258424 | -0.0911013 |
| O60462 | NRP2 | Neuropilin-2 | -0.005069892 | 0.127849 |
| O60488 | ACSL4 | Long-chain-fatty-acid--CoA ligase 4 | -1.043706258 | -0.0440896 |
| O60493 | SNX3 | Sorting nexin-3 | -0.065264702 | 0.101173 |
| O60502 | OGA | Protein O-GlcNAcase | -0.068433444 | 0.23682 |
| O60504 | SORBS3 | Vinexin | -0.170923154 | 0.0475938 |
| O60506 | SYNCRIP | Heterogeneous nuclear ribonucleoprotein Q | -0.53829813 | -0.0553761 |
| O60518 | RANBP6 | Ran-binding protein 6 | -0.19042627 | -0.430918 |
| O60524 | NEMF | Ribosome quality control complex subunit NEMF | -0.179684242 | -0.148504 |
| O60543 | CIDEA | Cell death activator CIDE-A | -1.317048152 | -0.313611 |
| O60547 | GMDS | GDP-mannose 4,6 dehydratase | -0.05416139 | 0.108287 |
| O60551 | NMT2 | Glycylpeptide N-tetradecanoyltransferase 2 | -0.061250846 | -0.310431 |
| O60563 | CCNT1 | Cyclin-T1 | -1.048010826 | -0.31038 |
| O60565 | GREM1 | Gremlin-1 | -2.152873993 | -0.190264 |
| O60566 | BUB1B | Mitotic checkpoint serine/threonine-protein kinase BUB1 beta | -0.013597647 | -0.00947285 |
| O60568 | PLOD3 | Multifunctional procollagen lysine hydroxylase and glycosyltransferase LH3 | -0.53305157 | 0.0950813 |
| O60573 | EIF4E2 | Eukaryotic translation initiation factor 4E type 2 | -1.36504372 | -0.740089 |
| O60645 | EXOC3 | Exocyst complex component 3 | 0.009992917 | -0.828228 |
| O60664 | PLIN3 | Perilipin-3 | -0.41888984 | -0.161599 |
| O60664-3 | PLIN3 | Isoform 3 of Perilipin-3 | -1.315393925 | -0.102223 |
| O60671 | RAD1 | Cell cycle checkpoint protein RAD1 | -0.01128912 | -0.0916541 |
| O60678 | PRMT3 | Protein arginine N-methyltransferase 3 | 0.019324541 | 0.070185 |
| O60684 | KPNA6 | Importin subunit alpha-7 | -0.063182672 | 0.00980568 |
| O60701 | UGDH | UDP-glucose 6-dehydrogenase | 0.102838834 | -0.102406 |
| O60711 | LPXN | Leupaxin | -0.268168688 | -0.0920372 |
| O60725 | ICMT | Protein-S-isoprenylcysteine O-methyltransferase | -1.416954041 | 0.894359 |
| O60749 | SNX2 | Sorting nexin-2 | 0.176086426 | -0.0580037 |
| O60763 | USO1 | General vesicular transport factor p115 | 0.122157415 | 0.0377649 |
| O60783 | MRPS14 | 28S ribosomal protein S14, mitochondrial | 0.637996912 | 1.99698 |
| O60784 | TOM1 | Target of Myb protein 1 | 0.229649464 | 0.370774 |
| O60826 | CCDC22 | Coiled-coil domain-containing protein 22 | 0.054694812 | -0.438252 |
| O60828 | PQBP1 | Polyglutamine-binding protein 1 | -0.283002774 | -0.118153 |
| O60830 | TIMM17B | Mitochondrial import inner membrane translocase subunit Tim17-B | 0 | -7.24276 |
| O60831 | PRAF2 | PRA1 family protein 2 | -0.745761077 | 0.589507 |
| O60832 | DKC1 | H/ACA ribonucleoprotein complex subunit DKC1 | -0.458120108 | -0.131458 |
| O60869 | EDF1 | Endothelial differentiation-related factor 1 | -0.530613581 | -0.0430803 |
| O60870 | KIN | DNA/RNA-binding protein KIN17 | -0.10228006 | 0.988854 |
| O60884 | DNAJA2 | DnaJ homolog subfamily A member 2 | -0.013460954 | 0.122111 |
| O60885 | BRD4 | Bromodomain-containing protein 4 | -0.150883516 | 0.0301695 |
| O60921 | HUS1 | Checkpoint protein HUS1 | -0.199693998 | -0.3787 |
| O60927 | PPP1R11 | E3 ubiquitin-protein ligase PPP1R11 | -0.927672942 | -0.350317 |
| O60934 | NBN | Nibrin | -0.42793417 | 0.24038 |
| O60942 | RNGTT | mRNA-capping enzyme | -0.404201508 | -0.0679741 |
| O75083 | WDR1 | WD repeat-containing protein 1 | 0.044874032 | -0.031496 |
| O75113 | N4BP1 | NEDD4-binding protein 1 | 0.772906701 | 0.0378138 |

| | | | | |
|---|---|---|---|---|
| O75116 | ROCK2 | Rho-associated protein kinase 2 | -0.227086147 | -0.115663 |
| O75131 | CPNE3 | Copine-3 | 0.046194553 | 0.00726 |
| O75143 | ATG13 | Autophagy-related protein 13 | -0.055079699 | -0.281334 |
| O75146 | HIP1R | Huntingtin-interacting protein 1-related protein | -0.420095682 | 0.273179 |
| O75150 | RNF40 | E3 ubiquitin-protein ligase BRE1B | -0.319043239 | 0.594443 |
| O75153 | CLUH | Clustered mitochondria protein homolog | -1.191459102 | -0.793168 |
| O75155 | CAND2 | Cullin-associated NEDD8-dissociated protein 2 | 0.311331828 | 0.00588894 |
| O75157 | TSC22D2 | TSC22 domain family protein 2 | 0.189770937 | 0.0785793 |
| O75164 | KDM4A | Lysine-specific demethylase 4A | -0.385485013 | -0.595091 |
| O75165 | DNAJC13 | DnaJ homolog subfamily C member 13 | -1.094911496 | -0.0964464 |
| O75170 | PPP6R2 | Serine/threonine-protein phosphatase 6 regulatory subunit 2 | -0.003312588 | -0.0702734 |
| O75175 | CNOT3 | CCR4-NOT transcription complex subunit 3 | -0.353380283 | 0.282785 |
| O75190 | DNAJB6 | DnaJ homolog subfamily B member 6 | -0.074388027 | 0.0532904 |
| O75223 | GGCT | Gamma-glutamylcyclotransferase | 0.006874561 | -0.00868734 |
| O75306 | NDUFS2 | NADH dehydrogenase [ubiquinone] iron-sulfur protein 2, mitochondrial | -1.125313361 | -0.719045 |
| O75312 | ZPR1 | Zinc finger protein ZPR1 | 0.056244214 | 0.0828816 |
| O75323 | NIPSNAP2 | Protein NipSnap homolog 2 | -2.291321596 | 0.0200837 |
| O75326 | SEMA7A | Semaphorin-7A | -0.831859986 | 0.0380898 |
| O75330 | HMMR | Hyaluronan mediated motility receptor | -0.062759638 | -2.56711 |
| O75340 | PDCD6 | Programmed cell death protein 6 | -0.052909374 | 0.131005 |
| O75342 | ALOX12B | Arachidonate 12-lipoxygenase, 12R-type | -0.099186341 | -0.202366 |
| O75347 | TBCA | Tubulin-specific chaperone A | -0.020052751 | -0.0728493 |
| O75348 | ATP6V1G1 | V-type proton ATPase subunit G 1 | -0.091255267 | 0.124705 |
| O75351 | VPS4B | Vacuolar protein sorting-associated protein 4B | 0.11550204 | -0.0339667 |
| O75367 | MACROH2A1 | Core histone macro-H2A.1 | -1.012225548 | 0.241528 |
| O75368 | SH3BGRL | SH3 domain-binding glutamic acid-rich-like protein | 0.081187089 | 0.620087 |
| O75369-8 | FLNB | Isoform 8 of Filamin-B | 0.162222068 | 0.0231469 |
| O75380 | NDUFS6 | NADH dehydrogenase [ubiquinone] iron-sulfur protein 6, mitochondrial | -0.475141128 | 0.393373 |
| O75381 | PEX14 | Peroxisomal membrane protein PEX14 | -0.483845075 | 0.327321 |
| O75396 | SEC22B | Vesicle-trafficking protein SEC22b | -0.98248299 | 0.179732 |
| O75410 | TACC1 | Transforming acidic coiled-coil-containing protein 1 | 0.260137955 | 0.111577 |
| O75419 | CDC45 | Cell division control protein 45 homolog | -1.161127647 | 0.049037 |
| O75420 | GIGYF1 | GRB10-interacting GYF protein 1 | 3.186571519 | 1.09528 |
| O75431 | MTX2 | Metaxin-2 | -1.592657129 | 1.9505 |
| O75436 | VPS26A | Vacuolar protein sorting-associated protein 26A | 0.03688701 | -0.0377846 |
| O75439 | PMPCB | Mitochondrial-processing peptidase subunit beta | -0.380082607 | 0.308431 |
| O75449 | KATNA1 | Katanin p60 ATPase-containing subunit A1 | -0.616408348 | -0.00262896 |
| O75475 | PSIP1 | PC4 and SFRS1-interacting protein | -0.753358285 | 0.729472 |
| O75477 | ERLIN1 | Erlin-1 | -1.342078606 | 0.451723 |
| O75489 | NDUFS3 | NADH dehydrogenase [ubiquinone] iron-sulfur protein 3, mitochondrial | -0.854177793 | 0.583953 |
| O75494 | SRSF10 | Serine/arginine-rich splicing factor 10 | -0.20863692 | 0.0130199 |
| O75531 | BANF1 | Barrier-to-autointegration factor | -1.727641463 | 0.137331 |
| O75533 | SF3B1 | Splicing factor 3B subunit 1 | -0.335671663 | 0.0345097 |
| O75534 | CSDE1 | Cold shock domain-containing protein E1 | -0.265880028 | -0.0348914 |
| O75582 | RPS6KA5 | Ribosomal protein S6 kinase alpha-5 | -0.343224923 | 0.13818 |
| O75593 | FOXH1 | Forkhead box protein H1 | 0.498926202 | 0 |
| O75607 | NPM3 | Nucleoplasmin-3 | -0.134546439 | -0.0593847 |
| O75616 | ERAL1 | GTPase Era, mitochondrial | -0.146956841 | 0.413364 |
| O75635 | SERPINB7 | Serpin B7 | -0.136718035 | 1.95321 |
| O75643 | SNRNP200 | U5 small nuclear ribonucleoprotein 200 kDa helicase | -0.168069045 | 0.0544462 |
| O75663 | TIPRL | TIP41-like protein | 0.018399556 | -0.0415996 |
| O75683 | SURF6 | Surfeit locus protein 6 | -2.147705128 | 0.142997 |
| O75688 | PPM1B | Protein phosphatase 1B | 0.145450751 | -0.0231031 |
| O75691 | UTP20 | Small subunit processome component 20 homolog | -0.16326348 | 0.192465 |
| O75695 | RP2 | Protein XRP2 | -0.874970277 | -0.168037 |
| O75717 | WDHD1 | WD repeat and HMG-box DNA-binding protein 1 | -0.025548538 | 0.0148201 |
| O75718 | CRTAP | Cartilage-associated protein | -0.13741374 | 0.360122 |
| O75746 | SLC25A12 | Calcium-binding mitochondrial carrier protein Aralar1 | -1.051602999 | 0.716163 |
| O75792 | RNASEH2A | Ribonuclease H2 subunit A | -0.1612312 | 0.5477 |
| O75794 | CDC123 | Cell division cycle protein 123 homolog | -0.096734285 | 0.0766726 |
| O75815 | BCAR3 | Breast cancer anti-estrogen resistance protein 3 | -0.359060923 | 0.0540698 |
| O75818 | RPP40 | Ribonuclease P protein subunit p40 | -0.788944483 | 0.371169 |
| O75821 | EIF3G | Eukaryotic translation initiation factor 3 subunit G | -0.018229485 | -0.0852044 |
| O75822 | EIF3J | Eukaryotic translation initiation factor 3 subunit J | 0.240558942 | -0.148788 |
| O75828 | CBR3 | Carbonyl reductase [NADPH] 3 | 0.002058983 | 0.0810413 |
| O75832 | PSMD10 | 26S proteasome non-ATPase regulatory subunit 10 | -0.063211838 | 0.292668 |
| O75843 | AP1G2 | AP-1 complex subunit gamma-like 2 | -0.123270591 | -0.250463 |
| O75844 | ZMPSTE24 | CAAX prenyl protease 1 homolog | -1.51016434 | 0.349151 |

| | | | | |
|---|---|---|---|---|
| O75874 | IDH1 | Isocitrate dehydrogenase [NADP] cytoplasmic | -0.026928266 | -0.00546837 |
| O75882 | ATRN | Attractin | -0.268281857 | -0.778384 |
| O75884 | RBBP9 | Serine hydrolase RBBP9 | -0.093268156 | 0.263531 |
| O75886 | STAM2 | Signal transducing adapter molecule 2 | 0.290708303 | 0.0368811 |
| O75896 | TUSC2 | Tumor suppressor candidate 2 | -0.755968491 | 0.201433 |
| O75911 | DHRS3 | Short-chain dehydrogenase/reductase 3 | -0.988011042 | -0.0639954 |
| O75915 | ARL6IP5 | PRA1 family protein 3 | -0.98385795 | 0.0249583 |
| O75923 | DYSF | Dysferlin | -0.82925272 | -0.0543858 |
| O75934 | BCAS2 | Pre-mRNA-splicing factor SPF27 | -0.039809942 | 0.878869 |
| O75935 | DCTN3 | Dynactin subunit 3 | 0.368099928 | 0.265479 |
| O75937 | DNAJC8 | DnaJ homolog subfamily C member 8 | -0.102501392 | 0.029631 |
| O75940 | SMNDC1 | Survival of motor neuron-related-splicing factor 30 | -0.22060434 | 0.0681372 |
| O75947 | ATP5PD | ATP synthase subunit d, mitochondrial | -1.170817852 | 0.0574547 |
| O75955 | FLOT1 | Flotillin-1 | -0.934097449 | 0.0835374 |
| O75962 | TRIO | Triple functional domain protein | -0.32599028 | 0.43597 |
| O75976 | CPD | Carboxypeptidase D | -1.055589596 | 0.766128 |
| O76003 | GLRX3 | Glutaredoxin-3 | 0.118936539 | 0.071249 |
| O76009 | KRT33A | Keratin, type I cuticular Ha3-I | -1.818628629 | 2.83257 |
| O76013 | KRT36 | Keratin, type I cuticular Ha6 | -1.162036101 | 2.19919 |
| O76021 | RSL1D1 | Ribosomal L1 domain-containing protein 1 | -0.72337532 | 0.195222 |
| O76031 | CLPX | ATP-dependent Clp protease ATP-binding subunit clpX-like, mitochondrial | -0.582511187 | 0.310845 |
| O76054 | SEC14L2 | SEC14-like protein 2 | 0.019616048 | 0.0431547 |
| O76061 | STC2 | Stanniocalcin-2 | -1.102256298 | 0.0483147 |
| O76070 | SNCG | Gamma-synuclein | -1.353382428 | 2.84181 |
| O76071 | CIAO1 | Probable cytosolic iron-sulfur protein assembly protein CIAO1 | -0.147549073 | -0.00847149 |
| O76080 | ZFAND5 | AN1-type zinc finger protein 5 | -1.015449762 | -0.259375 |
| O76094 | SRP72 | Signal recognition particle subunit SRP72 | -0.155477842 | -0.364843 |
| O76095 | JTB | Protein JTB | -0.411338886 | 0.392278 |
| O94760 | DDAH1 | N(G),N(G)-dimethylarginine dimethylaminohydrolase 1 | 0.100663026 | -0.141223 |
| O94763 | URI1 | Unconventional prefoldin RPB5 interactor 1 | -0.78948156 | 0.164467 |
| O94776 | MTA2 | Metastasis-associated protein MTA2 | -0.07917285 | 0.166087 |
| O94782 | USP1 | Ubiquitin carboxyl-terminal hydrolase 1 | -0.276805162 | -0.46419 |
| O94804 | STK10 | Serine/threonine-protein kinase 10 | -0.487672726 | -0.247992 |
| O94806 | PRKD3 | Serine/threonine-protein kinase D3 | 0.135318041 | -0.663851 |
| O94808 | GFPT2 | Glutamine--fructose-6-phosphate aminotransferase [isomerizing] 2 | -0.351466656 | 0.0312134 |
| O94817 | ATG12 | Ubiquitin-like protein ATG12 | -0.269096136 | 0.0522302 |
| O94819 | KBTBD11 | Kelch repeat and BTB domain-containing protein 11 | 0.014481147 | -0.913006 |
| O94822 | LTN1 | E3 ubiquitin-protein ligase listerin | -0.527831395 | 0.102455 |
| O94826 | TOMM70 | Mitochondrial import receptor subunit TOM70 | -1.005869706 | 0.375913 |
| O94829 | IPO13 | Importin-13 | 0.119071166 | -0.382472 |
| O94830 | DDHD2 | Phospholipase DDHD2 | -0.035647392 | 0.104166 |
| O94842 | TOX4 | TOX high mobility group box family member 4 | -0.257958412 | 0.174685 |
| O94851 | MICAL2 | [F-actin]-monooxygenase MICAL2 | -1.23464632 | -0.228203 |
| O94855 | SEC24D | Protein transport protein Sec24D | -0.092193762 | -0.103415 |
| O94874 | UFL1 | E3 UFM1-protein ligase 1 | -1.10875535 | 0.467241 |
| O94888 | UBXN7 | UBX domain-containing protein 7 | 0.071399609 | 0.034859 |
| O94889 | KLHL18 | Kelch-like protein 18 | 0.187109311 | -0.164271 |
| O94903 | PLPBP | Pyridoxal phosphate homeostasis protein | 0.148578405 | 0.0354538 |
| O94906 | PRPF6 | Pre-mRNA-processing factor 6 | -0.438636382 | -0.123452 |
| O94915-2 | FRYL | Isoform 2 of Protein furry homolog-like | -3.483776649 | 0 |
| O94919 | ENDOD1 | Endonuclease domain-containing 1 protein | -0.98170956 | 0.257909 |
| O94925-3 | GLS | Isoform 3 of Glutaminase kidney isoform, mitochondrial | -0.576594989 | -0.0983111 |
| O94927 | HAUS5 | HAUS augmin-like complex subunit 5 | -0.110165358 | -0.0599241 |
| O94941 | UBOX5 | RING finger protein 37 | 1.092011849 | 0 |
| O94952 | FBXO21 | F-box only protein 21 | -0.525045554 | -0.777519 |
| O94966-6 | USP19 | Isoform 6 of Ubiquitin carboxyl-terminal hydrolase 19 | 0.030886571 | -0.089215 |
| O94967 | WDR47 | WD repeat-containing protein 47 | 0.164755265 | -1.31732 |
| O94973 | AP2A2 | AP-2 complex subunit alpha-2 | -0.587210019 | -0.177952 |
| O94992 | HEXIM1 | Protein HEXIM1 | -0.197940509 | -0.0362968 |
| O95059 | RPP14 | Ribonuclease P protein subunit p14 | -0.735926469 | -1.25717 |
| O95067 | CCNB2 | G2/mitotic-specific cyclin-B2 | -0.955351035 | -0.134633 |
| O95071 | UBR5 | E3 ubiquitin-protein ligase UBR5 | -0.216207663 | 0.257998 |
| O95104 | SCAF4 | SR-related and CTD-associated factor 4 | -0.468136152 | -0.288101 |
| O95139 | NDUFB6 | NADH dehydrogenase [ubiquinone] 1 beta subcomplex subunit 6 | -0.724247853 | 0.881857 |
| O95149 | SNUPN | Snurportin-1 | 0.008100828 | -0.16091 |
| O95155 | UBE4B | Ubiquitin conjugation factor E4 B | 0.038487434 | -0.11426 |
| O95159 | ZFPL1 | Zinc finger protein-like 1 | -1.100111643 | 0.263426 |
| O95163 | ELP1 | Elongator complex protein 1 | 0.107347647 | -0.353992 |
| O95168 | NDUFB4 | NADH dehydrogenase [ubiquinone] 1 beta subcomplex subunit 4 | -1.376089811 | 0.17461 |

| | | | | |
|---|---|---|---|---|
| O95182 | NDUFA7 | NADH dehydrogenase [ubiquinone] 1 alpha subcomplex subunit 7 | -1.891133149 | -0.492501 |
| O95197-3 | RTN3 | Isoform 3 of Reticulon-3 | -1.225004991 | -0.410423 |
| O95202 | LETM1 | Mitochondrial proton/calcium exchanger protein | -1.309700569 | 0.575006 |
| O95208 | EPN2 | Epsin-2 | -0.237505754 | -0.327045 |
| O95210 | STBD1 | Starch-binding domain-containing protein 1 | -1.700805346 | 0.0527693 |
| O95218 | ZRANB2 | Zinc finger Ran-binding domain-containing protein 2 | -0.093352238 | -0.222186 |
| O95219 | SNX4 | Sorting nexin-4 | 0.286017497 | -0.00974305 |
| O95229 | ZWINT | ZW10 interactor | -0.070732594 | -0.149341 |
| O95235 | KIF20A | Kinesin-like protein KIF20A | -3.505628626 | 0.427535 |
| O95239 | KIF4A | Chromosome-associated kinesin KIF4A | -1.266505559 | -0.989023 |
| O95273 | CCNDBP1 | Cyclin-D1-binding protein 1 | 0.037458579 | -0.0778236 |
| O95274 | LYPD3 | Ly6/PLAUR domain-containing protein 3 | -1.538584312 | 0.983594 |
| O95292 | VAPB | Vesicle-associated membrane protein-associated protein B/C | -1.393781424 | 0.0241896 |
| O95297 | MPZL1 | Myelin protein zero-like protein 1 | -2.113048395 | 2.65666 |
| O95298 | NDUFC2 | NADH dehydrogenase [ubiquinone] 1 subunit C2 | -2.953341921 | 0.631282 |
| O95302 | FKBP9 | Peptidyl-prolyl cis-trans isomerase FKBP9 | 0.118198554 | 0.281534 |
| O95336 | PGLS | 6-phosphogluconolactonase | 0.128173033 | 0.440639 |
| O95340 | PAPSS2 | Bifunctional 3'-phosphoadenosine 5'-phosphosulfate synthase 2 | 0.040200154 | 0.0591189 |
| O95347 | SMC2 | Structural maintenance of chromosomes protein 2 | -0.958669265 | 0.228124 |
| O95352 | ATG7 | Ubiquitin-like modifier-activating enzyme ATG7 | 0.117280245 | -0.154006 |
| O95361 | TRIM16 | Tripartite motif-containing protein 16 | -0.211620569 | -0.0399596 |
| O95372 | LYPLA2 | Acyl-protein thioesterase 2 | 0.059427023 | 0.461107 |
| O95373 | IPO7 | Importin-7 | -0.07874918 | -0.115299 |
| O95376 | ARIH2 | E3 ubiquitin-protein ligase ARIH2 | -0.0276951 | 0.249173 |
| O95379 | TNFAIP8 | Tumor necrosis factor alpha-induced protein 8 | -0.48026228 | -0.0903336 |
| O95391 | SLU7 | Pre-mRNA-splicing factor SLU7 | -0.173358838 | 0.0366729 |
| O95396 | MOCS3 | Adenylyltransferase and sulfurtransferase MOCS3 | -0.352183421 | 0.144791 |
| O95400 | CD2BP2 | CD2 antigen cytoplasmic tail-binding protein 2 | -0.113725265 | 0.340828 |
| O95405 | ZFYVE9 | Zinc finger FYVE domain-containing protein 9 | 0.008730809 | 0.99146 |
| O95415 | BRI3 | Brain protein I3 | -2.395694097 | 0.683519 |
| O95429 | BAG4 | BAG family molecular chaperone regulator 4 | -0.266938289 | -0.600206 |
| O95433 | AHSA1 | Activator of 90 kDa heat shock protein ATPase homolog 1 | 0.139619827 | -0.360919 |
| O95456 | PSMG1 | Proteasome assembly chaperone 1 | 0.034915606 | -0.129033 |
| O95466 | FMNL1 | Formin-like protein 1 | -0.003123681 | -0.88435 |
| O95470 | SGPL1 | Sphingosine-1-phosphate lyase 1 | -1.611065149 | 0.511006 |
| O95478 | NSA2 | Ribosome biogenesis protein NSA2 homolog | 0 | 3.19386 |
| O95479 | H6PD | GDH/6PGL endoplasmic bifunctional protein | -1.060002724 | 1.79808 |
| O95486 | SEC24A | Protein transport protein Sec24A | -0.190440098 | 0.244995 |
| O95487 | SEC24B | Protein transport protein Sec24B | -0.103358984 | -0.141538 |
| O95544 | NADK | NAD kinase | -0.532152573 | 0 |
| O95551 | TDP2 | Tyrosyl-DNA phosphodiesterase 2 | 0.146688938 | 0.23028 |
| O95571 | ETHE1 | Persulfide dioxygenase ETHE1, mitochondrial | -0.373185873 | -0.00247002 |
| O95602 | POLR1A | DNA-directed RNA polymerase I subunit RPA1 | -0.944913228 | -0.123524 |
| O95628 | CNOT4 | CCR4-NOT transcription complex subunit 4 | 0.086488644 | 0.262494 |
| O95671 | ASMTL | Probable bifunctional dTTP/UTP pyrophosphatase/methyltransferase protein | -0.02762723 | 0.117701 |
| O95685 | PPP1R3D | Protein phosphatase 1 regulatory subunit 3D | 0.442727327 | -2.07377 |
| O95714 | HERC2 | E3 ubiquitin-protein ligase HERC2 | -1.346093973 | -1.50247 |
| O95716 | RAB3D | Ras-related protein Rab-3D | -0.421484311 | -0.309258 |
| O95721 | SNAP29 | Synaptosomal-associated protein 29 | -0.317779462 | 0.0985781 |
| O95747 | OXSR1 | Serine/threonine-protein kinase OSR1 | 0.124552886 | -0.186399 |
| O95749 | GGPS1 | Geranylgeranyl pyrophosphate synthase | -0.284215053 | -0.0783717 |
| O95755 | RAB36 | Ras-related protein Rab-36 | -1.739923159 | 0.169879 |
| O95757 | HSPA4L | Heat shock 70 kDa protein 4L | 0.093075832 | -0.072307 |
| O95758 | PTBP3 | Polypyrimidine tract-binding protein 3 | 1.235979398 | -0.147316 |
| O95777 | LSM8 | U6 snRNA-associated Sm-like protein LSm8 | -0.122724533 | -0.327816 |
| O95782 | AP2A1 | AP-2 complex subunit alpha-1 | -0.586154143 | -0.223166 |
| O95793 | STAU1 | Double-stranded RNA-binding protein Staufen homolog 1 | -0.854929129 | 0.1131 |
| O95801 | TTC4 | Tetratricopeptide repeat protein 4 | 0.089469274 | 0.0921815 |
| O95807 | TMEM50A | Transmembrane protein 50A | -1.239345551 | 1.75858 |
| O95816 | BAG2 | BAG family molecular chaperone regulator 2 | -0.337478558 | 0.000625292 |
| O95817 | BAG3 | BAG family molecular chaperone regulator 3 | -0.006945292 | -0.0190325 |
| O95825 | CRYZL1 | Quinone oxidoreductase-like protein 1 | -0.16268452 | 0.162131 |
| O95833 | CLIC3 | Chloride intracellular channel protein 3 | -1.205613295 | 0.973649 |
| O95834 | EML2 | Echinoderm microtubule-associated protein-like 2 | 0.008097013 | -0.246564 |
| O95848 | NUDT14 | Uridine diphosphate glucose pyrophosphatase NUDT14 | -0.085186799 | -0.732726 |
| O95861 | BPNT1 | 3'(2'),5'-bisphosphate nucleotidase 1 | -0.08516407 | -0.228181 |
| O95865 | DDAH2 | N(G),N(G)-dimethylarginine dimethylaminohydrolase 2 | 0.149300734 | 0.254801 |
| O95881 | TXNDC12 | Thioredoxin domain-containing protein 12 | -0.072252512 | 0.288193 |
| O95905 | ECD | Protein ecdysoneless homolog | -0.119204601 | 0.144436 |
| O95983 | MBD3 | Methyl-CpG-binding domain protein 3 | -0.753131469 | 0.0977201 |

| | | | | |
|---|---|---|---|---|
| O95989 | NUDT3 | Diphosphoinositol polyphosphate phosphohydrolase 1 | 0.186199983 | 0.0828597 |
| O95997 | PTTG1 | Securin | -0.642636696 | -0.132085 |
| O95999 | BCL10 | B-cell lymphoma/leukemia 10 | 0.295425812 | 0.379181 |
| O96000 | NDUFB10 | NADH dehydrogenase [ubiquinone] 1 beta subcomplex subunit 10 | -1.925309022 | 0.598718 |
| O96005 | CLPTM1 | Cleft lip and palate transmembrane protein 1 | -1.464104652 | 0.288616 |
| O96007 | MOCS2 | Molybdopterin synthase catalytic subunit | -0.176936229 | -0.825379 |
| O96008 | TOMM40 | Mitochondrial import receptor subunit TOM40 homolog | -2.039093177 | 0.191313 |
| O96013 | PAK4 | Serine/threonine-protein kinase PAK 4 | -1.55181702 | -0.36942 |
| O96019 | ACTL6A | Actin-like protein 6A | -0.016227643 | -0.0870422 |
| P00156 | MT-CYB | Cytochrome b | 0 | -0.594971 |
| P00167 | CYB5A | Cytochrome b5 | -0.54129076 | 1.88244 |
| P00338 | LDHA | L-lactate dehydrogenase A chain | -0.025241693 | -0.261342 |
| P00367 | GLUD1 | Glutamate dehydrogenase 1, mitochondrial | -0.778930346 | 0.262163 |
| P00374 | DHFR | Dihydrofolate reductase | -0.031002045 | 0.107665 |
| P00387 | CYB5R3 | NADH-cytochrome b5 reductase 3 | -0.859211445 | -0.16283 |
| P00390 | GSR | Glutathione reductase, mitochondrial | 0.015987237 | -0.0787659 |
| P00403 | MT-CO2 | Cytochrome c oxidase subunit 2 | -1.17550842 | 0.218182 |
| P00414 | MT-CO3 | Cytochrome c oxidase subunit 3 | -2.756172816 | -0.73939 |
| P00441 | SOD1 | Superoxide dismutase [Cu-Zn] | 0.173183759 | 0.34338 |
| P00491 | PNP | Purine nucleoside phosphorylase | -0.061184088 | -0.229996 |
| P00492 | HPRT1 | Hypoxanthine-guanine phosphoribosyltransferase | 0.006237189 | 0.0828743 |
| P00505 | GOT2 | Aspartate aminotransferase, mitochondrial | -0.798503876 | 0.052297 |
| P00533 | EGFR | Epidermal growth factor receptor | -1.009574095 | 0.490021 |
| P00558 | PGK1 | Phosphoglycerate kinase 1 | 0.057844798 | -0.0624758 |
| P00568 | AK1 | Adenylate kinase isoenzyme 1 | 0.043263753 | 0.0654189 |
| P00846 | MT-ATP6 | ATP synthase subunit a | -2.055364688 | 0.307495 |
| P00918 | CA2 | Carbonic anhydrase 2 | 0.289644957 | -0.282541 |
| P00966 | ASS1 | Argininosuccinate synthase | -0.155009826 | -0.00747172 |
| P01008 | SERPINC1 | Antithrombin-III | 0.029924234 | -0.0677675 |
| P01023 | A2M | Alpha-2-macroglobulin | 0.110988617 | -0.0935446 |
| P01024 | C3 | Complement C3 | -0.567145427 | -0.20533 |
| P01031 | C5 | Complement C5 | -1.905034781 | 0.206005 |
| P01033 | TIMP1 | Metalloproteinase inhibitor 1 | -0.322934389 | -0.387476 |
| P01036 | CST4 | Cystatin-S | -0.95704476 | 1.38372 |
| P01037 | CST1 | Cystatin-SN | -1.968863885 | 1.17459 |
| P01040 | CSTA | Cystatin-A | -0.143111785 | 0.416554 |
| P01111 | NRAS | GTPase NRas | 0.229869525 | 0.1283 |
| P01112 | HRAS | GTPase HRas | -0.287464778 | 0.178988 |
| P01116-2 | KRAS | Isoform 2B of GTPase KRas | -1.135569652 | 0.832907 |
| P01583 | IL1A | Interleukin-1 alpha | -0.978598436 | 2.00476 |
| P01584 | IL1B | Interleukin-1 beta | 0.221994718 | 0.638857 |
| P01834 | IGKC | Immunoglobulin kappa constant | -0.359772682 | -0.0245778 |
| P02042 | HBD | Hemoglobin subunit delta | 0.672357639 | -0.524004 |
| P02452 | COL1A1 | Collagen alpha-1(I) chain | -0.3546772 | 1.11557 |
| P02461 | COL3A1 | Collagen alpha-1(III) chain | -0.970299641 | 0.498889 |
| P02533 | KRT14 | Keratin, type I cytoskeletal 14 | -0.273659865 | -0.213154 |
| P02538 | KRT6A | Keratin, type II cytoskeletal 6A | -1.301658948 | 0.120495 |
| P02545-2 | LMNA | Isoform C of Prelamin-A/C | -1.049654563 | -0.112672 |
| P02549 | SPTA1 | Spectrin alpha chain, erythrocytic 1 | 0.829059442 | -0.82336 |
| P02649 | APOE | Apolipoprotein E | -0.606069803 | 0.122978 |
| P02671 | FGA | Fibrinogen alpha chain | 0.541719755 | -0.865906 |
| P02675 | FGB | Fibrinogen beta chain | 0.576471567 | -0.90635 |
| P02730 | SLC4A1 | Band 3 anion transport protein | 0.593644063 | -0.59009 |
| P02748 | C9 | Complement component C9 | -0.993603627 | 0.465466 |
| P02751-1 | FN1 | Isoform 1 of Fibronectin | -0.250435034 | 0.270248 |
| P02765 | AHSG | Alpha-2-HS-glycoprotein | -0.180286328 | 1.21849 |
| P02768 | ALB | Albumin | -0.072944323 | -0.316764 |
| P02775 | PPBP | Platelet basic protein | 0.989638329 | -0.400375 |
| P02776 | PF4 | Platelet factor 4 | 1.339981318 | -2.81303 |
| P02786 | TFRC | Transferrin receptor protein 1 | -0.9203403 | -0.03829 |
| P02787 | TF | Serotransferrin | 0.310215791 | -0.158017 |
| P02792 | FTL | Ferritin light chain | 0.581964731 | -0.502004 |
| P02794 | FTH1 | Ferritin heavy chain | 0.398255905 | 0.090366 |
| P02795 | MT2A | Metallothionein-2 | -0.553278764 | 0.186526 |
| P03891 | MT-ND2 | NADH-ubiquinone oxidoreductase chain 2 | 0 | 0.0271702 |
| P03915 | MT-ND5 | NADH-ubiquinone oxidoreductase chain 5 | -1.956204335 | -2.38197 |
| P03973 | SLPI | Antileukoproteinase | -0.985488256 | 0.827903 |
| P04004 | VTN | Vitronectin | -0.566381613 | 0.210519 |
| P04040 | CAT | Catalase | -0.157359759 | 0.0253359 |
| P04049 | RAF1 | RAF proto-oncogene serine/threonine-protein kinase | -0.314687808 | -0.782383 |
| P04066 | FUCA1 | Tissue alpha-L-fucosidase | -0.227965117 | 0.381584 |

| | | | | |
|---|---|---|---|---|
| P04075 | ALDOA | Fructose-bisphosphate aldolase A | -0.122582277 | 0.112149 |
| P04080 | CSTB | Cystatin-B | 0.171578884 | 0.144608 |
| P04083 | ANXA1 | Annexin A1 | 0.163863182 | 0.0233409 |
| P04114 | APOB | Apolipoprotein B-100 | -0.059570313 | 0.0691481 |
| P04150 | NR3C1 | Glucocorticoid receptor | -0.044141293 | 0.00267092 |
| P04179 | SOD2 | Superoxide dismutase [Mn], mitochondrial | -0.189974785 | 0.140656 |
| P04181 | OAT | Ornithine aminotransferase, mitochondrial | -0.314006805 | -0.0683775 |
| P04183 | TK1 | Thymidine kinase, cytosolic | -0.784490188 | 0.565579 |
| P04259 | KRT6B | Keratin, type II cytoskeletal 6B | -0.839207172 | 0.137594 |
| P04264 | KRT1 | Keratin, type II cytoskeletal 1 | -0.197428862 | -0.086305 |
| P04279 | SEMG1 | Semenogelin-1 | 1.328878323 | 0.309054 |
| P04350 | TUBB4A | Tubulin beta-4A chain | -0.13379097 | 0.0588077 |
| P04406 | GAPDH | Glyceraldehyde-3-phosphate dehydrogenase | -0.246353626 | -0.295143 |
| P04406-2 | GAPDH | Isoform 2 of Glyceraldehyde-3-phosphate dehydrogenase | -0.334874312 | 0 |
| P04424 | ASL | Argininosuccinate lyase | 0.157119592 | -0.179976 |
| P04439 | HLA-A | HLA class I histocompatibility antigen, A alpha chain | -0.894009272 | 0.172037 |
| P04733 | MT1F | Metallothionein-1F | -0.457010746 | 0.178363 |
| P04792 | HSPB1 | Heat shock protein beta-1 | 0.165992101 | 0.0813859 |
| P04818 | TYMS | Thymidylate synthase | -0.334474802 | -0.364135 |
| P04843 | RPN1 | Dolichyl-diphosphooligosaccharide--protein glycosyltransferase subunit 1 | -1.202992042 | 0.197765 |
| P04844 | RPN2 | Dolichyl-diphosphooligosaccharide--protein glycosyltransferase subunit 2 | -1.124376297 | -0.125498 |
| P04899 | GNAI2 | Guanine nucleotide-binding protein G(i) subunit alpha-2 | -0.988814751 | 0.186687 |
| P04908 | H2AC4 | Histone H2A type 1-B/E | -0.714946429 | 0.224194 |
| P05023 | ATP1A1 | Sodium/potassium-transporting ATPase subunit alpha-1 | -0.873623451 | 0.0361684 |
| P05026 | ATP1B1 | Sodium/potassium-transporting ATPase subunit beta-1 | -0.077927033 | -0.399437 |
| P05067 | APP | Amyloid-beta precursor protein | -0.000431299 | -0.020614 |
| P05089 | ARG1 | Arginase-1 | -0.418100437 | 0.21648 |
| P05091 | ALDH2 | Aldehyde dehydrogenase, mitochondrial | -0.407708168 | 0.364762 |
| P05109 | S100A8 | Protein S100-A8 | -0.265673637 | 0.247578 |
| P05114 | HMGN1 | Non-histone chromosomal protein HMG-14 | -0.634042978 | 0.403512 |
| P05120 | SERPINB2 | Plasminogen activator inhibitor 2 | 0.191716353 | 0.0157948 |
| P05121 | SERPINE1 | Plasminogen activator inhibitor 1 | -0.336657206 | 0.0855961 |
| P05141 | SLC25A5 | ADP/ATP translocase 2 | -1.328926404 | 0.302203 |
| P05198 | EIF2S1 | Eukaryotic translation initiation factor 2 subunit 1 | -0.038208961 | -0.110416 |
| P05204 | HMGN2 | Non-histone chromosomal protein HMG-17 | -0.778270324 | 0.104249 |
| P05362 | ICAM1 | Intercellular adhesion molecule 1 | -0.910423835 | 0.433227 |
| P05386 | RPLP1 | 60S acidic ribosomal protein P1 | -1.788626591 | 0.804501 |
| P05387 | RPLP2 | 60S acidic ribosomal protein P2 | -0.652360598 | 0.253317 |
| P05388 | RPLP0 | 60S acidic ribosomal protein P0 | -0.581734498 | 0.0204322 |
| P05412 | JUN | Transcription factor Jun | -0.457741896 | 0.641772 |
| P05423 | POLR3D | DNA-directed RNA polymerase III subunit RPC4 | -2.103741407 | 2.357 |
| P05455 | SSB | Lupus La protein | -0.04813242 | -0.0718892 |
| P05783 | KRT18 | Keratin, type I cytoskeletal 18 | 0.171168486 | -0.0786146 |
| P05787 | KRT8 | Keratin, type II cytoskeletal 8 | 0.028217872 | -0.422851 |
| P06132 | UROD | Uroporphyrinogen decarboxylase | -0.071902593 | 0.0722154 |
| P06396-2 | GSN | Isoform 2 of Gelsolin | 0.105444908 | -0.0212065 |
| P06493 | CDK1 | Cyclin-dependent kinase 1 | -0.162885666 | -0.118799 |
| P06576 | ATP5F1B | ATP synthase subunit beta, mitochondrial | -0.542185465 | 0.0616163 |
| P06702 | S100A9 | Protein S100-A9 | -0.013922373 | 0.317671 |
| P06703 | S100A6 | Protein S100-A6 | 0.170414448 | 0.0505276 |
| P06727 | APOA4 | Apolipoprotein A-IV | -0.826758146 | -1.73378 |
| P06730 | EIF4E | Eukaryotic translation initiation factor 4E | -0.142450333 | -0.479521 |
| P06732 | CKM | Creatine kinase M-type | 0.024633249 | -1.19082 |
| P06733 | ENO1 | Alpha-enolase | 0.098405043 | -0.0452404 |
| P06737 | PYGL | Glycogen phosphorylase, liver form | 0.072435379 | -0.204609 |
| P06744 | GPI | Glucose-6-phosphate isomerase | -0.001271089 | -0.21888 |
| P06748 | NPM1 | Nucleophosmin | 0.28268528 | -0.103565 |
| P06748-3 | NPM1 | Isoform 3 of Nucleophosmin | 0.149369558 | 0.331205 |
| P06753-2 | TPM3 | Isoform 2 of Tropomyosin alpha-3 chain | 0.192760944 | 0.0771701 |
| P06756 | ITGAV | Integrin alpha-V | -1.486176093 | 0.115951 |
| P06899 | H2BC11 | Histone H2B type 1-J | -0.725540956 | 0.0355701 |
| P07093-2 | SERPINE2 | Isoform 2 of Glia-derived nexin | 0.064695676 | 0.358764 |
| P07099 | EPHX1 | Epoxide hydrolase 1 | -1.676449935 | -0.170923 |
| P07108 | DBI | Acyl-CoA-binding protein | -0.008079767 | 0.744092 |
| P07195 | LDHB | L-lactate dehydrogenase B chain | 0.11665074 | -0.100282 |
| P07203 | GPX1 | Glutathione peroxidase 1 | 0.072860718 | 0.122346 |
| P07205 | PGK2 | Phosphoglycerate kinase 2 | -0.717599312 | -1.59963 |
| P07305 | H1-0 | Histone H1.0 | -1.119257768 | 0.311715 |
| P07311 | ACYP1 | Acylphosphatase-1 | -0.037530661 | 0.0107632 |

| | | | | |
|---|---|---|---|---|
| P07355 | ANXA2 | Annexin A2 | 0.181883971 | -0.0351391 |
| P07384 | CAPN1 | Calpain-1 catalytic subunit | 0.013055484 | -0.0319589 |
| P07437 | TUBB | Tubulin beta chain | -0.303265254 | 0.253384 |
| P07451 | CA3 | Carbonic anhydrase 3 | 0.342160861 | -0.426671 |
| P07476 | IVL | Involucrin | -0.817323844 | 0.923823 |
| P07686 | HEXB | Beta-hexosaminidase subunit beta | -0.051287969 | 0.0521653 |
| P07737 | PFN1 | Profilin-1 | 0.087262948 | -4.32E-05 |
| P07738 | BPGM | Bisphosphoglycerate mutase | 0.020359993 | -0.35374 |
| P07741 | APRT | Adenine phosphoribosyltransferase | 0.121703784 | 0.178075 |
| P07814 | EPRS1 | Bifunctional glutamate/proline--tRNA ligase | -0.351581653 | -0.0584733 |
| P07900 | HSP90AA1 | Heat shock protein HSP 90-alpha | 0.064065933 | -0.0939827 |
| P07902 | GALT | Galactose-1-phosphate uridylyltransferase | -0.017010689 | -0.21525 |
| P07951 | TPM2 | Tropomyosin beta chain | 0.181004524 | 0.410051 |
| P07951-2 | TPM2 | Isoform 2 of Tropomyosin beta chain | 0.119126717 | 0.23199 |
| P07954-2 | FH | Isoform Cytoplasmic of Fumarate hydratase, mitochondrial | -0.072944641 | -0.0527763 |
| P07996 | THBS1 | Thrombospondin-1 | 0.24298962 | -0.0760504 |
| P08133 | ANXA6 | Annexin A6 | 0.142270247 | -0.0282027 |
| P08134 | RHOC | Rho-related GTP-binding protein RhoC | -0.223250707 | 0.853997 |
| P08236 | GUSB | Beta-glucuronidase | -1.079969883 | -0.582763 |
| P08238 | HSP90AB1 | Heat shock protein HSP 90-beta | 0.11005497 | 0.00647799 |
| P08240 | SRPRA | Signal recognition particle receptor subunit alpha | -0.533413728 | 0.298069 |
| P08243 | ASNS | Asparagine synthetase [glutamine-hydrolyzing] | 0.04055198 | -0.0417423 |
| P08253 | MMP2 | 72 kDa type IV collagenase | -1.462589741 | 0.605013 |
| P08473 | MME | Neprilysin | -0.861740112 | 0.328477 |
| P08476 | INHBA | Inhibin beta A chain | 0.106895645 | 3.51892 |
| P08514 | ITGA2B | Integrin alpha-IIb | 0.710502307 | -0.761437 |
| P08559 | PDHA1 | Pyruvate dehydrogenase E1 component subunit alpha, somatic form, mitochondrial | -0.586779515 | 0.265968 |
| P08572 | COL4A2 | Collagen alpha-2(IV) chain | -1.954400976 | 0.707126 |
| P08574 | CYC1 | Cytochrome c1, heme protein, mitochondrial | -1.806793451 | 0.896758 |
| P08579 | SNRPB2 | U2 small nuclear ribonucleoprotein B'' | -0.193597635 | -0.114621 |
| P08582 | MELTF | Melanotransferrin | -1.596811215 | 1.26418 |
| P08621 | SNRNP70 | U1 small nuclear ribonucleoprotein 70 kDa | 0.137818019 | -0.375939 |
| P08648 | ITGA5 | Integrin alpha-5 | -1.791923682 | 0.469795 |
| P08670 | VIM | Vimentin | 0.007979552 | -0.138759 |
| P08727 | KRT19 | Keratin, type I cytoskeletal 19 | -0.547200918 | -0.949969 |
| P08729 | KRT7 | Keratin, type II cytoskeletal 7 | 0.13537852 | -0.0926482 |
| P08754 | GNAI3 | Guanine nucleotide-binding protein G(i) subunit alpha-3 | -0.993037701 | -0.0215673 |
| P08758 | ANXA5 | Annexin A5 | 0.106105328 | -0.0140864 |
| P08779 | KRT16 | Keratin, type I cytoskeletal 16 | -0.772823016 | 0.0955969 |
| P09012 | SNRPA | U1 small nuclear ribonucleoprotein A | 0.094047546 | -0.141663 |
| P09038-2 | FGF2 | Isoform 3 of Fibroblast growth factor 2 | -0.445892175 | 0.0595121 |
| P09104 | ENO2 | Gamma-enolase | -0.021753788 | 0.0233695 |
| P09211 | GSTP1 | Glutathione S-transferase P | 0.009969393 | 0.0183786 |
| P09234 | SNRPC | U1 small nuclear ribonucleoprotein C | -0.470825354 | -0.081598 |
| P09382 | LGALS1 | Galectin-1 | -0.004722595 | 0.268276 |
| P09417 | QDPR | Dihydropteridine reductase | -0.049622854 | 0.0632585 |
| P09429 | HMGB1 | High mobility group protein B1 | 0.082883358 | 0.110133 |
| P09486 | SPARC | SPARC | -0.14125371 | 1.01896 |
| P09488 | GSTM1 | Glutathione S-transferase Mu 1 | -0.203439792 | -0.0921383 |
| P09493-4 | TPM1 | Isoform 4 of Tropomyosin alpha-1 chain | 0.251971722 | 0.0107918 |
| P09493-8 | TPM1 | Isoform 8 of Tropomyosin alpha-1 chain | 0.292443752 | 0.0847178 |
| P09496-2 | CLTA | Isoform Non-brain of Clathrin light chain A | 0.230776946 | 0.610521 |
| P09497-2 | CLTB | Isoform Non-brain of Clathrin light chain B | 0.046832641 | 0.125449 |
| P09525 | ANXA4 | Annexin A4 | 0.117150148 | -0.104143 |
| P09543 | CNP | 2',3'-cyclic-nucleotide 3'-phosphodiesterase | -0.753079176 | 0.249749 |
| P09601 | HMOX1 | Heme oxygenase 1 | -1.101924102 | 0.0652409 |
| P09622 | DLD | Dihydrolipoyl dehydrogenase, mitochondrial | -0.420954148 | 0.139287 |
| P09661 | SNRPA1 | U2 small nuclear ribonucleoprotein A' | -0.026510874 | -0.0452944 |
| P09669 | COX6C | Cytochrome c oxidase subunit 6C | -0.99408563 | 0.171957 |
| P09874 | PARP1 | Poly [ADP-ribose] polymerase 1 | -0.602324804 | 0.203858 |
| P09913 | IFIT2 | Interferon-induced protein with tetratricopeptide repeats 2 | 4.149381955 | 0.291233 |
| P09936 | UCHL1 | Ubiquitin carboxyl-terminal hydrolase isozyme L1 | 0.005551974 | 0.214568 |
| P09960 | LTA4H | Leukotriene A-4 hydrolase | -0.004598776 | -0.0506274 |
| P09972 | ALDOC | Fructose-bisphosphate aldolase C | 0.171402613 | -0.0442327 |
| P0C0S5 | H2AZ1 | Histone H2A.Z | -0.693807443 | 0.0608527 |
| P0C0S8 | H2AC11 | Histone H2A type 1 | -1.418237766 | 0.160784 |
| P0CAP2 | POLR2M | DNA-directed RNA polymerase II subunit GRINL1A | -0.20837005 | 0.147806 |
| P0CG38 | POTEI | POTE ankyrin domain family member I | -0.684724331 | -0.108373 |
| P0DI81 | TRAPPC2 | Trafficking protein particle complex subunit 2 | -1.326754093 | -0.040278 |
| P0DOY2 | IGLC2 | Immunoglobulin lambda constant 2 | -0.995929003 | -0.586225 |

| | | | | |
|---|---|---|---|---|
| P0DP23 | CALM1 | Calmodulin-1 | 0.879505078 | 0.533764 |
| P0DPD7 | EEF1AKMT4 | EEF1A lysine methyltransferase 4 | 0.344802618 | 0.124052 |
| P0DPH7 | TUBA3C | Tubulin alpha-3C chain | 0.68718791 | -1.28655 |
| P0DTE7 | AMY1B | Alpha-amylase 1B | -0.55473129 | 1.3312 |
| P10155 | RO60 | RNA-binding protein RO60 | -0.026786884 | 0.00390816 |
| P10253 | GAA | Lysosomal alpha-glucosidase | 0.104714076 | -0.069555 |
| P10301 | RRAS | Ras-related protein R-Ras | -1.479017178 | -0.0478805 |
| P10398 | ARAF | Serine/threonine-protein kinase A-Raf | -0.5043571 | 0.0937303 |
| P10412 | H1-4 | Histone H1.4 | -0.585629543 | 0.176423 |
| P10586 | PTPRF | Receptor-type tyrosine-protein phosphatase F | -0.797895988 | -1.25821 |
| P10599 | TXN | Thioredoxin | 0.108387311 | -0.00618362 |
| P10606 | COX5B | Cytochrome c oxidase subunit 5B, mitochondrial | -1.82701381 | 0 |
| P10620 | MGST1 | Microsomal glutathione S-transferase 1 | -1.510579745 | 0.384599 |
| P10643 | C7 | Complement component C7 | -1.982391556 | 0.771743 |
| P10644 | PRKAR1A | cAMP-dependent protein kinase type I-alpha regulatory subunit | 0.034471989 | 0.0245444 |
| P10746 | UROS | Uroporphyrinogen-III synthase | -0.382012049 | -0.0152171 |
| P10768 | ESD | S-formylglutathione hydrolase | -0.061664899 | 0.0604172 |
| P10809 | HSPD1 | 60 kDa heat shock protein, mitochondrial | -0.137801329 | 0.171887 |
| P10909 | CLU | Clusterin | -0.496578455 | 0.112462 |
| P11021 | HSPA5 | Endoplasmic reticulum chaperone BiP | -0.068809986 | 0.0846081 |
| P11047 | LAMC1 | Laminin subunit gamma-1 | -0.072669506 | 0.360876 |
| P11142 | HSPA8 | Heat shock cognate 71 kDa protein | 0.189909299 | -0.0316099 |
| P11166 | SLC2A1 | Solute carrier family 2, facilitated glucose transporter member 1 | -0.36057965 | -0.608206 |
| P11169 | SLC2A3 | Solute carrier family 2, facilitated glucose transporter member 3 | 0.032207648 | 0.916937 |
| P11172 | UMPS | Uridine 5'-monophosphate synthase | -0.012991428 | -0.0319271 |
| P11177 | PDHB | Pyruvate dehydrogenase E1 component subunit beta, mitochondrial | -0.563275337 | 0.197848 |
| P11216 | PYGB | Glycogen phosphorylase, brain form | -0.010399342 | -0.228433 |
| P11217 | PYGM | Glycogen phosphorylase, muscle form | 0.094412009 | -0.0292053 |
| P11233 | RALA | Ras-related protein Ral-A | -1.064023097 | 0.213309 |
| P11234 | RALB | Ras-related protein Ral-B | -1.313186963 | 0.179531 |
| P11274 | BCR | Breakpoint cluster region protein | -0.458651702 | -0.0709143 |
| P11277 | SPTB | Spectrin beta chain, erythrocytic | 0.658489943 | -0.599075 |
| P11279 | LAMP1 | Lysosome-associated membrane glycoprotein 1 | -0.501810869 | -0.0208569 |
| P11310 | ACADM | Medium-chain specific acyl-CoA dehydrogenase, mitochondrial | -0.412467082 | 0.234657 |
| P11387 | TOP1 | DNA topoisomerase 1 | -0.538823287 | 0.548321 |
| P11388 | TOP2A | DNA topoisomerase 2-alpha | -1.094813983 | 0.319751 |
| P11413 | G6PD | Glucose-6-phosphate 1-dehydrogenase | -0.025643826 | -0.094389 |
| P11441 | UBL4A | Ubiquitin-like protein 4A | 0.220649799 | 0.816173 |
| P11717 | IGF2R | Cation-independent mannose-6-phosphate receptor | -0.277913411 | 0.357034 |
| P11766 | ADH5 | Alcohol dehydrogenase class-3 | 0.036744595 | -0.214444 |
| P11802 | CDK4 | Cyclin-dependent kinase 4 | -0.048190435 | -0.0654494 |
| P11908 | PRPS2 | Ribose-phosphate pyrophosphokinase 2 | -0.148196379 | -0.142893 |
| P11940 | PABPC1 | Polyadenylate-binding protein 1 | -0.244217555 | -0.0517324 |
| P12004 | PCNA | Proliferating cell nuclear antigen | 0.031411807 | -0.125477 |
| P12109 | COL6A1 | Collagen alpha-1(VI) chain | -0.135646979 | 0.135049 |
| P12110 | COL6A2 | Collagen alpha-2(VI) chain | -0.365103563 | 0.100394 |
| P12235 | SLC25A4 | ADP/ATP translocase 1 | -1.270302931 | 0.246882 |
| P12236 | SLC25A6 | ADP/ATP translocase 3 | -1.11548535 | 0.620474 |
| P12270 | TPR | Nucleoprotein TPR | 0.140186946 | 0.230326 |
| P12273 | PIP | Prolactin-inducible protein | -0.381676992 | 0.379696 |
| P12277 | CKB | Creatine kinase B-type | 0.303967794 | -0.0167306 |
| P12429 | ANXA3 | Annexin A3 | 0.259254138 | 0.136633 |
| P12532 | CKMT1A | Creatine kinase U-type, mitochondrial | 0.435066303 | 0.276238 |
| P12931 | SRC | Proto-oncogene tyrosine-protein kinase Src | -0.305992047 | -0.067325 |
| P12955 | PEPD | Xaa-Pro dipeptidase | 0.103999774 | 0.081267 |
| P12956 | XRCC6 | X-ray repair cross-complementing protein 6 | -0.012197177 | 0.0808347 |
| P13010 | XRCC5 | X-ray repair cross-complementing protein 5 | -0.026649634 | 0.106443 |
| P13073 | COX4I1 | Cytochrome c oxidase subunit 4 isoform 1, mitochondrial | -1.037851334 | 0.706816 |
| P13284 | IFI30 | Gamma-interferon-inducible lysosomal thiol reductase | -0.760762334 | 0 |
| P13473 | LAMP2 | Lysosome-associated membrane glycoprotein 2 | -0.422667821 | -0.017369 |
| P13473-2 | LAMP2 | Isoform LAMP-2B of Lysosome-associated membrane glycoprotein 2 | -0.493190527 | 0.391489 |
| P13489 | RNH1 | Ribonuclease inhibitor | -0.383336862 | 0.158079 |
| P13498 | CYBA | Cytochrome b-245 light chain | -3.47570994 | 0.984074 |
| P13639 | EEF2 | Elongation factor 2 | -0.098004977 | 0.0363833 |
| P13646 | KRT13 | Keratin, type I cytoskeletal 13 | -0.086168766 | 2.34558 |
| P13647 | KRT5 | Keratin, type II cytoskeletal 5 | -0.324690501 | -0.201688 |
| P13667 | PDIA4 | Protein disulfide-isomerase A4 | 0.070566495 | 0.081974 |
| P13674 | P4HA1 | Prolyl 4-hydroxylase subunit alpha-1 | -0.158767859 | 0.130613 |
| P13674-2 | P4HA1 | Isoform 2 of Prolyl 4-hydroxylase subunit alpha-1 | 0.094012181 | -0.331907 |

| | | | | |
|---|---|---|---|---|
| P13693 | TPT1 | Translationally-controlled tumor protein | 0.201432705 | -0.00820605 |
| P13716 | ALAD | Delta-aminolevulinic acid dehydratase | -0.165788174 | -0.226203 |
| P13726 | F3 | Tissue factor | -1.963143547 | 0 |
| P13796 | LCP1 | Plastin-2 | 0.331690868 | 0.117032 |
| P13797 | PLS3 | Plastin-3 | 0.156274954 | -0.0744737 |
| P13804 | ETFA | Electron transfer flavoprotein subunit alpha, mitochondrial | -0.351580302 | -0.0146004 |
| P13807 | GYS1 | Glycogen [starch] synthase, muscle | -1.349607468 | -0.0733678 |
| P13861 | PRKAR2A | cAMP-dependent protein kinase type II-alpha regulatory subunit | 0.139685949 | -0.0461159 |
| P13929 | ENO3 | Beta-enolase | -0.01634113 | 0.616718 |
| P13984 | GTF2F2 | General transcription factor IIF subunit 2 | 0.155510426 | -0.235928 |
| P14151 | SELL | L-selectin | -1.416729371 | 0.105738 |
| P14174 | MIF | Macrophage migration inhibitory factor | -0.605940819 | 0.360512 |
| P14324 | FDPS | Farnesyl pyrophosphate synthase | 0.126557986 | -0.202923 |
| P14373 | TRIM27 | Zinc finger protein RFP | -1.99525555 | -3.82437 |
| P14550 | AKR1A1 | Aldo-keto reductase family 1 member A1 | 0.114284833 | -0.0311775 |
| P14618 | PKM | Pyruvate kinase PKM | 0.011083126 | -0.0645459 |
| P14625 | HSP90B1 | Endoplasmin | 0.030549685 | -0.0455411 |
| P14635 | CCNB1 | G2/mitotic-specific cyclin-B1 | -1.301539342 | -0.284123 |
| P14649 | MYL6B | Myosin light chain 6B | -0.471044381 | -0.492682 |
| P14854 | COX6B1 | Cytochrome c oxidase subunit 6B1 | 0.017631451 | 0.847497 |
| P14866 | HNRNPL | Heterogeneous nuclear ribonucleoprotein L | -1.028321425 | 0.47277 |
| P14868 | DARS1 | Aspartate--tRNA ligase, cytoplasmic | -0.152051608 | -0.0177116 |
| P14921 | ETS1 | Protein C-ets-1 | -0.473356088 | -1.31308 |
| P14923 | JUP | Junction plakoglobin | -0.381002426 | 0.345304 |
| P14927 | UQCRB | Cytochrome b-c1 complex subunit 7 | -1.519189835 | 0.398804 |
| P15104 | GLUL | Glutamine synthetase | -0.14970994 | -0.324737 |
| P15121 | AKR1B1 | Aldo-keto reductase family 1 member B1 | -0.037165324 | 0.0513808 |
| P15144 | ANPEP | Aminopeptidase N | -0.245178699 | -0.0488071 |
| P15170-2 | GSPT1 | Isoform 2 of Eukaryotic peptide chain release factor GTP-binding subunit ERF3A | 0.025517146 | 0.00516701 |
| P15291 | B4GALT1 | Beta-1,4-galactosyltransferase 1 | 0.868626515 | 0 |
| P15311 | EZR | Ezrin | -0.26788648 | -0.0318171 |
| P15374 | UCHL3 | Ubiquitin carboxyl-terminal hydrolase isozyme L3 | 0.155676842 | 0.564839 |
| P15407 | FOSL1 | Fos-related antigen 1 | -0.763624112 | 0.830684 |
| P15531 | NME1 | Nucleoside diphosphate kinase A | 0.263949871 | -0.0245508 |
| P15559 | NQO1 | NAD(P)H dehydrogenase [quinone] 1 | -0.087639173 | -0.207406 |
| P15586 | GNS | N-acetylglucosamine-6-sulfatase | 0.056306521 | 0.186036 |
| P15848 | ARSB | Arylsulfatase B | 0.006470044 | 0.0916459 |
| P15880 | RPS2 | 40S ribosomal protein S2 | -0.434390704 | -0.455952 |
| P15924 | DSP | Desmoplakin | -0.290772835 | 0.376051 |
| P15927 | RPA2 | Replication protein A 32 kDa subunit | -0.392376026 | -0.0371704 |
| P16035 | TIMP2 | Metalloproteinase inhibitor 2 | 0.224470297 | 0.043135 |
| P16070 | CD44 | CD44 antigen | -1.090250651 | 0.72995 |
| P16083 | NQO2 | Ribosyldihydronicotinamide dehydrogenase [quinone] | -0.15702184 | 0.13704 |
| P16152 | CBR1 | Carbonyl reductase [NADPH] 1 | -0.071837425 | -0.0747522 |
| P16157 | ANK1 | Ankyrin-1 | 0.823680957 | -0.467629 |
| P16220 | CREB1 | Cyclic AMP-responsive element-binding protein 1 | -0.425578674 | 0.591236 |
| P16278 | GLB1 | Beta-galactosidase | -0.09885176 | -0.284559 |
| P16298 | PPP3CB | Serine/threonine-protein phosphatase 2B catalytic subunit beta isoform | 0.245037079 | -0.0475801 |
| P16333 | NCK1 | Cytoplasmic protein NCK1 | 0.03042992 | -0.0342166 |
| P16383 | GCFC2 | Intron Large complex component GCFC2 | -0.329314947 | -0.201849 |
| P16401 | H1-5 | Histone H1.5 | -0.659612497 | 0.141251 |
| P16402 | H1-3 | Histone H1.3 | -0.467968464 | 0.0880076 |
| P16403 | H1-2 | Histone H1.2 | -0.455207109 | 0.428685 |
| P16435 | POR | NADPH--cytochrome P450 reductase | -1.360841115 | 0.288287 |
| P16452 | EPB42 | Protein 4.2 | 0.664031188 | -0.595237 |
| P16615 | ATP2A2 | Sarcoplasmic/endoplasmic reticulum calcium ATPase 2 | -1.214647373 | 0.251937 |
| P16930 | FAH | Fumarylacetoacetase | 0.055179278 | 0.105087 |
| P16949 | STMN1 | Stathmin | 0.18711551 | 0.0146834 |
| P16989 | YBX3 | Y-box-binding protein 3 | -0.484791358 | -0.252083 |
| P17050 | NAGA | Alpha-N-acetylgalactosaminidase | -0.063884735 | 0.123727 |
| P17096 | HMGA1 | High mobility group protein HMG-I/HMG-Y | -0.835753918 | 0.0726048 |
| P17096-2 | HMGA1 | Isoform HMG-Y of High mobility group protein HMG-I/HMG-Y | -0.979797602 | 0.133394 |
| P17152 | TMEM11 | Transmembrane protein 11, mitochondrial | -1.957955758 | -1.16684 |
| P17174 | GOT1 | Aspartate aminotransferase, cytoplasmic | 0.147205194 | -0.0230573 |
| P17252 | PRKCA | Protein kinase C alpha type | 0.045204242 | -0.0932614 |
| P17301 | ITGA2 | Integrin alpha-2 | -0.821924369 | 0.248439 |
| P17302 | GJA1 | Gap junction alpha-1 protein | -0.167621215 | 0.508086 |
| P17405 | SMPD1 | Sphingomyelin phosphodiesterase | -0.515763919 | -0.164061 |
| P17480 | UBTF | Nucleolar transcription factor 1 | 0.279338121 | 0.0978661 |

| | | | | |
|---|---|---|---|---|
| P17535 | JUND | Transcription factor JunD | -0.070187251 | 0 |
| P17568 | NDUFB7 | NADH dehydrogenase [ubiquinone] 1 beta subcomplex subunit 7 | -3.130632798 | 1.36522 |
| P17600 | SYN1 | Synapsin-1 | 0.690481742 | -0.909298 |
| P17612 | PRKACA | cAMP-dependent protein kinase catalytic subunit alpha | -0.021163464 | -0.318521 |
| P17655 | CAPN2 | Calpain-2 catalytic subunit | 0.002078374 | -0.0136967 |
| P17676 | CEBPB | CCAAT/enhancer-binding protein beta | -0.915765206 | 0.191143 |
| P17706 | PTPN2 | Tyrosine-protein phosphatase non-receptor type 2 | -0.566768328 | 1.19763 |
| P17812 | CTPS1 | CTP synthase 1 | 0.036970456 | -0.00772476 |
| P17813 | ENG | Endoglin | -1.712211927 | 1.15165 |
| P17858 | PFKL | ATP-dependent 6-phosphofructokinase, liver type | -0.245498419 | -0.201633 |
| P17900 | GM2A | Ganglioside GM2 activator | -0.467229525 | -0.0949532 |
| P17931 | LGALS3 | Galectin-3 | -0.617289066 | 0.487342 |
| P17980 | PSMC3 | 26S proteasome regulatory subunit 6A | -0.024817944 | 0.0156962 |
| P17987 | TCP1 | T-complex protein 1 subunit alpha | -0.049889882 | -0.161066 |
| P18031 | PTPN1 | Tyrosine-protein phosphatase non-receptor type 1 | -0.95100681 | 0.229712 |
| P18077 | RPL35A | 60S ribosomal protein L35a | -1.375471354 | 0.033707 |
| P18085 | ARF4 | ADP-ribosylation factor 4 | 0.056230545 | -0.478583 |
| P18124 | RPL7 | 60S ribosomal protein L7 | -1.147166252 | 0.141826 |
| P18206 | VCL | Vinculin | 0.089044253 | -0.0600923 |
| P18583 | SON | Protein SON | -0.385912021 | -0.343264 |
| P18615 | NELFE | Negative elongation factor E | -0.218082984 | 0.133981 |
| P18669 | PGAM1 | Phosphoglycerate mutase 1 | 0.003062725 | -0.237937 |
| P18754 | RCC1 | Regulator of chromosome condensation | -0.792787155 | 0.333409 |
| P18858 | LIG1 | DNA ligase 1 | 0.00118343 | -0.0714095 |
| P18887 | XRCC1 | DNA repair protein XRCC1 | -0.062438806 | 0.0414632 |
| P19012 | KRT15 | Keratin, type I cytoskeletal 15 | -3.385417302 | -0.856585 |
| P19013 | KRT4 | Keratin, type II cytoskeletal 4 | 0.083157142 | 1.21241 |
| P19174-2 | PLCG1 | Isoform 2 of 1-phosphatidylinositol 4,5-bisphosphate phosphodiesterase gamma-1 | 0.035181522 | 0.0457252 |
| P19367 | HK1 | Hexokinase-1 | -0.237421672 | 0.0987072 |
| P19387 | POLR2C | DNA-directed RNA polymerase II subunit RPB3 | 0.028143883 | 0.175261 |
| P19388 | POLR2E | DNA-directed RNA polymerases I, II, and III subunit RPABC1 | 0.067961295 | 0.0829382 |
| P19623 | SRM | Spermidine synthase | 0.178130945 | 0.0152454 |
| P19784 | CSNK2A2 | Casein kinase II subunit alpha' | -0.120029132 | -0.0973647 |
| P19823 | ITIH2 | Inter-alpha-trypsin inhibitor heavy chain H2 | 0.382832607 | -0.33888 |
| P19838 | NFKB1 | Nuclear factor NF-kappa-B p105 subunit | -0.006932259 | -0.00431538 |
| P20020 | ATP2B1 | Plasma membrane calcium-transporting ATPase 1 | -1.386351585 | -0.0633036 |
| P20042 | EIF2S2 | Eukaryotic translation initiation factor 2 subunit 2 | -0.370616674 | -0.0839017 |
| P20073 | ANXA7 | Annexin A7 | 0.067378998 | 0.0590604 |
| P20248 | CCNA2 | Cyclin-A2 | -1.441062133 | 0.564189 |
| P20290 | BTF3 | Transcription factor BTF3 | -0.10580794 | -0.294431 |
| P20290-2 | BTF3 | Isoform 2 of Transcription factor BTF3 | -0.144491434 | 0.0490233 |
| P20336 | RAB3A | Ras-related protein Rab-3A | -0.359338681 | -0.743316 |
| P20337 | RAB3B | Ras-related protein Rab-3B | -0.655064662 | -0.189148 |
| P20338 | RAB4A | Ras-related protein Rab-4A | -0.445857048 | -0.055974 |
| P20339 | RAB5A | Ras-related protein Rab-5A | -0.515555223 | 0.0695712 |
| P20340 | RAB6A | Ras-related protein Rab-6A | -0.040100416 | 0.0974719 |
| P20591 | MX1 | Interferon-induced GTP-binding protein Mx1 | 2.578252713 | 0.513686 |
| P20618 | PSMB1 | Proteasome subunit beta type-1 | -0.105221907 | -0.170662 |
| P20645 | M6PR | Cation-dependent mannose-6-phosphate receptor | -1.606033643 | 0.333942 |
| P20700 | LMNB1 | Lamin-B1 | -0.241853476 | 0.193872 |
| P20742 | PZP | Pregnancy zone protein | -0.399588823 | 3.13849 |
| P20749 | BCL3 | B-cell lymphoma 3 protein | -0.414654573 | -1.75568 |
| P20809-2 | IL11 | Isoform 2 of Interleukin-11 | -0.507911364 | -0.00795492 |
| P20810-5 | CAST | Isoform 5 of Calpastatin | -1.076089223 | 0.657261 |
| P20810 | CAST | Calpastatin | -0.03127408 | 0.287667 |
| P20930 | FLG | Filaggrin | -0.970850309 | 0.799701 |
| P20933 | AGA | N(4)-(beta-N-acetylglucosaminyl)-L-asparaginase | -0.803067843 | -0.61037 |
| P20962 | PTMS | Parathymosin | 0.018925269 | 0.3118 |
| P21266 | GSTM3 | Glutathione S-transferase Mu 3 | -0.159747283 | 0.0839154 |
| P21281 | ATP6V1B2 | V-type proton ATPase subunit B, brain isoform | 0.123990695 | 0.122617 |
| P21283 | ATP6V1C1 | V-type proton ATPase subunit C 1 | -0.093508879 | -0.115004 |
| P21291 | CSRP1 | Cysteine and glycine-rich protein 1 | -0.610253016 | 0.632101 |
| P21333 | FLNA | Filamin-A | 0.139589469 | -0.00698598 |
| P21399 | ACO1 | Cytoplasmic aconitate hydratase | -0.006510099 | -0.0459105 |
| P21589 | NT5E | 5'-nucleotidase | -1.613930543 | 0.503764 |
| P21796 | VDAC1 | Voltage-dependent anion-selective channel protein 1 | -1.005082448 | 0.26593 |
| P21912 | SDHB | Succinate dehydrogenase [ubiquinone] iron-sulfur subunit, mitochondrial | -0.83783412 | 0.63697 |
| P21964 | COMT | Catechol O-methyltransferase | -0.865571022 | 0.940096 |
| P21980 | TGM2 | Protein-glutamine gamma-glutamyltransferase 2 | 0.127805869 | 0.0667439 |

| | | | | |
|---|---|---|---|---|
| P22033 | MMUT | Methylmalonyl-CoA mutase, mitochondrial | 0.254774253 | -0.382929 |
| P22059 | OSBP | Oxysterol-binding protein 1 | 0.063963652 | -0.245967 |
| P22061-2 | PCMT1 | Isoform 2 of Protein-L-isoaspartate(D-aspartate) O-methyltransferase | -0.113040129 | -0.178995 |
| P22087 | FBL | rRNA 2'-O-methyltransferase fibrillarin | -0.395715396 | 0.0635726 |
| P22102 | GART | Trifunctional purine biosynthetic protein adenosine-3 | 0.035720348 | 0.0155341 |
| P22234 | PAICS | Multifunctional protein ADE2 | -0.044843356 | -0.0249322 |
| P22307 | SCP2 | Sterol carrier protein 2 | -0.326127291 | -0.0168753 |
| P22314 | UBA1 | Ubiquitin-like modifier-activating enzyme 1 | 0.149497509 | -0.0644499 |
| P22392-2 | NME2 | Isoform 3 of Nucleoside diphosphate kinase B | -0.029674053 | -0.121593 |
| P22528 | SPRR1B | Cornifin-B | -1.779274543 | -0.268522 |
| P22532 | SPRR2D | Small proline-rich protein 2D | -1.065392971 | 0.12319 |
| P22670 | RFX1 | MHC class II regulatory factor RFX1 | -0.234548489 | -1.56096 |
| P22681 | CBL | E3 ubiquitin-protein ligase CBL | -0.21423769 | 0.0487773 |
| P22695 | UQCRC2 | Cytochrome b-c1 complex subunit 2, mitochondrial | -1.208112081 | 0.432849 |
| P22735 | TGM1 | Protein-glutamine gamma-glutamyltransferase K | -0.336280982 | 0.296604 |
| P23141 | CES1 | Liver carboxylesterase 1 | -0.934755762 | 0 |
| P23142 | FBLN1 | Fibulin-1 | -0.441377878 | 0.316418 |
| P23246 | SFPQ | Splicing factor, proline- and glutamine-rich | -0.810387373 | 0.298898 |
| P23258 | TUBG1 | Tubulin gamma-1 chain | -0.326631228 | 0.258883 |
| P23381 | WARS1 | Tryptophan--tRNA ligase, cytoplasmic | 0.209601402 | -0.0443071 |
| P23396 | RPS3 | 40S ribosomal protein S3 | -0.301741123 | -0.290346 |
| P23443 | RPS6KB1 | Ribosomal protein S6 kinase beta-1 | -0.357046445 | -0.0733906 |
| P23490 | LORICRIN | Loricrin | -0.869430304 | -1.87345 |
| P23497 | SP100 | Nuclear autoantigen Sp-100 | 0.482328892 | 0.0764923 |
| P23511 | NFYA | Nuclear transcription factor Y subunit alpha | 0.263270855 | -0.185199 |
| P23526 | AHCY | Adenosylhomocysteinase | 0.041664282 | -0.0365804 |
| P23528 | CFL1 | Cofilin-1 | -0.055052121 | 0.225524 |
| P23610 | F8A1 | 40-kDa huntingtin-associated protein | -0.083798091 | 0.587226 |
| P23634 | ATP2B4 | Plasma membrane calcium-transporting ATPase 4 | -2.00592343 | 0.74628 |
| P23743 | DGKA | Diacylglycerol kinase alpha | -0.014160713 | 0.0279894 |
| P23919 | DTYMK | Thymidylate kinase | -0.057403088 | -0.0802962 |
| P23921 | RRM1 | Ribonucleoside-diphosphate reductase large subunit | 0.018354575 | -0.060262 |
| P24158 | PRTN3 | Myeloblastin | 0.540989637 | -0.0661195 |
| P24385 | CCND1 | G1/S-specific cyclin-D1 | -0.021533887 | 0.275279 |
| P24386 | CHM | Rab proteins geranylgeranyltransferase component A 1 | -0.08978645 | -0.00174332 |
| P24390 | KDELR1 | ER lumen protein-retaining receptor 1 | -3.140742779 | -1.09169 |
| P24534 | EEF1B2 | Elongation factor 1-beta | 0.231954575 | 0.411055 |
| P24539 | ATP5PB | ATP synthase F(0) complex subunit B1, mitochondrial | -1.114692529 | 0.2393 |
| P24666 | ACP1 | Low molecular weight phosphotyrosine protein phosphatase | -0.00665315 | 0.186871 |
| P24752 | ACAT1 | Acetyl-CoA acetyltransferase, mitochondrial | -0.300317923 | 0.0888179 |
| P24844 | MYL9 | Myosin regulatory light polypeptide 9 | -0.052302519 | -0.421051 |
| P25098 | GRK2 | Beta-adrenergic receptor kinase 1 | -0.579312007 | -0.439468 |
| P25205 | MCM3 | DNA replication licensing factor MCM3 | -0.125894229 | 0.0303847 |
| P25311 | AZGP1 | Zinc-alpha-2-glycoprotein | -0.620580276 | 0.192582 |
| P25325-2 | MPST | Isoform 2 of 3-mercaptopyruvate sulfurtransferase | -0.125051022 | -0.0052948 |
| P25398 | RPS12 | 40S ribosomal protein S12 | -0.683645089 | 0.436724 |
| P25490 | YY1 | Transcriptional repressor protein YY1 | 0.520946423 | -1.35874 |
| P25685 | DNAJB1 | DnaJ homolog subfamily B member 1 | -0.39275527 | 0.1064 |
| P25686 | DNAJB2 | DnaJ homolog subfamily B member 2 | -0.144222021 | 0.256123 |
| P25705 | ATP5F1A | ATP synthase subunit alpha, mitochondrial | -0.722740014 | 0.222363 |
| P25786 | PSMA1 | Proteasome subunit alpha type-1 | -0.101641814 | -0.0414804 |
| P25788 | PSMA3 | Proteasome subunit alpha type-3 | -0.180676142 | -0.118875 |
| P25789 | PSMA4 | Proteasome subunit alpha type-4 | -0.105123043 | -0.127168 |
| P26006 | ITGA3 | Integrin alpha-3 | -1.355362813 | 0.479447 |
| P26022 | PTX3 | Pentraxin-related protein PTX3 | -0.971991539 | -4.9966 |
| P26038 | MSN | Moesin | -0.104764779 | 0.00309372 |
| P26196 | DDX6 | Probable ATP-dependent RNA helicase DDX6 | -0.773698886 | -0.0419817 |
| P26368 | U2AF2 | Splicing factor U2AF 65 kDa subunit | -0.259084622 | -0.0115058 |
| P26373 | RPL13 | 60S ribosomal protein L13 | -1.091445049 | 0.19183 |
| P26374 | CHML | Rab proteins geranylgeranyltransferase component A 2 | -0.155139605 | -0.088453 |
| P26447 | S100A4 | Protein S100-A4 | -0.208086014 | -0.605392 |
| P26583 | HMGB2 | High mobility group protein B2 | -0.50660793 | 0.203338 |
| P26639 | TARS1 | Threonine--tRNA ligase 1, cytoplasmic | 0.003674666 | -0.14138 |
| P26640 | VARS1 | Valine--tRNA ligase | 0.070302804 | 0.012956 |
| P26641 | EEF1G | Elongation factor 1-gamma | -0.055519581 | -0.00981267 |
| P27105 | STOM | Stomatin | -0.129225016 | -0.236006 |
| P27144 | AK4 | Adenylate kinase 4, mitochondrial | -0.623836358 | -0.0770804 |
| P27348 | YWHAQ | 14-3-3 protein theta | -0.026813189 | 0.459579 |
| P27361 | MAPK3 | Mitogen-activated protein kinase 3 | 0.073043108 | 0.0122706 |
| P27449 | ATP6V0C | V-type proton ATPase 16 kDa proteolipid subunit c | -1.58005619 | 1.79888 |

| | | | | |
|---|---|---|---|---|
| P27469 | G0S2 | G0/G1 switch protein 2 | 0.078487873 | 1.00964 |
| P27482 | CALML3 | Calmodulin-like protein 3 | -0.411276897 | 0.000687917 |
| P27635 | RPL10 | 60S ribosomal protein L10 | -1.056783994 | -0.0885941 |
| P27694 | RPA1 | Replication protein A 70 kDa DNA-binding subunit | -0.132187605 | -0.0835915 |
| P27695 | APEX1 | DNA-(apurinic or apyrimidinic site) endonuclease | -0.173935413 | 0.124974 |
| P27701 | CD82 | CD82 antigen | -2.241890828 | 3.02293 |
| P27707 | DCK | Deoxycytidine kinase | 0.132448832 | -0.0265837 |
| P27797 | CALR | Calreticulin | -0.09437116 | 0.0304616 |
| P27986 | PIK3R1 | Phosphatidylinositol 3-kinase regulatory subunit alpha | -1.130824645 | -0.304382 |
| P28062 | PSMB8 | Proteasome subunit beta type-8 | 0.118955612 | 0.0336631 |
| P28066 | PSMA5 | Proteasome subunit alpha type-5 | -0.069306215 | 0.158423 |
| P28070 | PSMB4 | Proteasome subunit beta type-4 | -0.233583291 | -0.0890242 |
| P28072 | PSMB6 | Proteasome subunit beta type-6 | -0.084856987 | 0.0224183 |
| P28074 | PSMB5 | Proteasome subunit beta type-5 | -0.183947722 | -0.130849 |
| P28161 | GSTM2 | Glutathione S-transferase Mu 2 | 0.02758201 | -0.0340614 |
| P28300 | LOX | Protein-lysine 6-oxidase | -1.536514362 | 0.348014 |
| P28331 | NDUFS1 | NADH-ubiquinone oxidoreductase 75 kDa subunit, mitochondrial | -0.843076388 | -0.0878439 |
| P28482 | MAPK1 | Mitogen-activated protein kinase 1 | 0.079862118 | -0.00989978 |
| P28799 | GRN | Progranulin | -0.42944932 | 0.13647 |
| P28838-2 | LAP3 | Isoform 2 of Cytosol aminopeptidase | 0.695894241 | -0.0795199 |
| P29034 | S100A2 | Protein S100-A2 | 2.256056388 | -0.388961 |
| P29083 | GTF2E1 | General transcription factor IIE subunit 1 | -0.016963164 | 0.236082 |
| P29084 | GTF2E2 | Transcription initiation factor IIE subunit beta | -0.332541943 | -1.06065 |
| P29144 | TPP2 | Tripeptidyl-peptidase 2 | -0.372946501 | 0.0583849 |
| P29218 | IMPA1 | Inositol monophosphatase 1 | 0.17318519 | -0.0451787 |
| P29279 | CCN2 | CCN family member 2 | -0.164513906 | 0.563714 |
| P29317 | EPHA2 | Ephrin type-A receptor 2 | -0.282028755 | -0.0799662 |
| P29353 | SHC1 | SHC-transforming protein 1 | 0.050906976 | -0.174404 |
| P29373 | CRABP2 | Cellular retinoic acid-binding protein 2 | -0.448253791 | 0.212051 |
| P29400 | COL4A5 | Collagen alpha-5(IV) chain | -0.783979098 | -0.672827 |
| P29401 | TKT | Transketolase | 0.080892245 | -0.0635974 |
| P29508 | SERPINB3 | Serpin B3 | -1.317148924 | -0.634799 |
| P29692 | EEF1D | Elongation factor 1-delta | 0.241602103 | 0.327824 |
| P29692-3 | EEF1D | Isoform 3 of Elongation factor 1-delta | -0.236132145 | 0.168865 |
| P29692-4 | EEF1D | Isoform 4 of Elongation factor 1-delta | 0.428389907 | 0 |
| P29966 | MARCKS | Myristoylated alanine-rich C-kinase substrate | 0.33230861 | 0.262454 |
| P29992 | GNA11 | Guanine nucleotide-binding protein subunit alpha-11 | -0.907689253 | -0.135698 |
| P30038 | ALDH4A1 | Delta-1-pyrroline-5-carboxylate dehydrogenase, mitochondrial | -0.003499031 | -0.288161 |
| P30040 | ERP29 | Endoplasmic reticulum resident protein 29 | 0.138304392 | -0.060524 |
| P30041 | PRDX6 | Peroxiredoxin-6 | 0.17774423 | 0.0361061 |
| P30043 | BLVRB | Flavin reductase (NADPH) | -0.094999472 | -0.00987752 |
| P30044-2 | PRDX5 | Isoform Cytoplasmic+peroxisomal of Peroxiredoxin-5, mitochondrial | 0.104044914 | 0.335542 |
| P30046 | DDT | D-dopachrome decarboxylase | -0.252434572 | 0.258093 |
| P30048 | PRDX3 | Thioredoxin-dependent peroxide reductase, mitochondrial | -0.065039953 | 0.0845318 |
| P30049 | ATP5F1D | ATP synthase subunit delta, mitochondrial | -0.843713522 | 0.166776 |
| P30050 | RPL12 | 60S ribosomal protein L12 | -1.174687783 | 0.140481 |
| P30084 | ECHS1 | Enoyl-CoA hydratase, mitochondrial | -0.463094711 | -0.0488478 |
| P30085 | CMPK1 | UMP-CMP kinase | 0.009441058 | -0.15504 |
| P30086 | PEBP1 | Phosphatidylethanolamine-binding protein 1 | 0.122464816 | -0.103293 |
| P30101 | PDIA3 | Protein disulfide-isomerase A3 | 0.005347093 | 0.145861 |
| P30153 | PPP2R1A | Serine/threonine-protein phosphatase 2A 65 kDa regulatory subunit A alpha isoform | -0.054028193 | -0.193747 |
| P30154 | PPP2R1B | Serine/threonine-protein phosphatase 2A 65 kDa regulatory subunit A beta isoform | 0.008315245 | 0.0246096 |
| P30260 | CDC27 | Cell division cycle protein 27 homolog | -0.785988251 | 0.578889 |
| P30307 | CDC25C | M-phase inducer phosphatase 3 | -0.219434023 | 0.532826 |
| P30405 | PPIF | Peptidyl-prolyl cis-trans isomerase F, mitochondrial | -0.961881161 | 0.195199 |
| P30414 | NKTR | NK-tumor recognition protein | -1.192248344 | -3.47268 |
| P30519 | HMOX2 | Heme oxygenase 2 | -1.082699617 | 0.170945 |
| P30520 | ADSS2 | Adenylosuccinate synthetase isozyme 2 | 0.101577123 | -0.036864 |
| P30530 | AXL | Tyrosine-protein kinase receptor UFO | -1.017291625 | 0.939666 |
| P30533 | LRPAP1 | Alpha-2-macroglobulin receptor-associated protein | -0.182838122 | 0.0505727 |
| P30622 | CLIP1 | CAP-Gly domain-containing linker protein 1 | 0.060494105 | 0.184746 |
| P30711 | GSTT1 | Glutathione S-transferase theta-1 | -0.0071582 | 0.548771 |
| P30740 | SERPINB1 | Leukocyte elastase inhibitor | 0.019958019 | 0.0571079 |
| P30837 | ALDH1B1 | Aldehyde dehydrogenase X, mitochondrial | -0.586268584 | -0.0226736 |
| P31025 | LCN1 | Lipocalin-1 | 0.187821865 | 1.38993 |
| P31040 | SDHA | Succinate dehydrogenase [ubiquinone] flavoprotein subunit, mitochondrial | -0.738347133 | 0.212907 |
| P31146 | CORO1A | Coronin-1A | 0.212108374 | -0.0629654 |

| | | | | |
|---|---|---|---|---|
| P31150 | GDI1 | Rab GDP dissociation inhibitor alpha | -0.107838949 | 0.132388 |
| P31151 | S100A7 | Protein S100-A7 | -1.434266726 | 0.358837 |
| P31153 | MAT2A | S-adenosylmethionine synthase isoform type-2 | 0.01695315 | -0.0299454 |
| P31323 | PRKAR2B | cAMP-dependent protein kinase type II-beta regulatory subunit | -0.643408696 | -2.89914 |
| P31327 | CPS1 | Carbamoyl-phosphate synthase [ammonia], mitochondrial | -0.743305922 | 0 |
| P31350 | RRM2 | Ribonucleoside-diphosphate reductase subunit M2 | -0.145366828 | -0.0713056 |
| P31689 | DNAJA1 | DnaJ homolog subfamily A member 1 | -0.011097749 | -0.0593866 |
| P31749 | AKT1 | RAC-alpha serine/threonine-protein kinase | 0.006216764 | 0.0513579 |
| P31751 | AKT2 | RAC-beta serine/threonine-protein kinase | 0.060831785 | 0.050518 |
| P31930 | UQCRC1 | Cytochrome b-c1 complex subunit 1, mitochondrial | -0.914339383 | 0.332738 |
| P31937 | HIBADH | 3-hydroxyisobutyrate dehydrogenase, mitochondrial | 0.140912056 | 0.0749356 |
| P31939 | ATIC | Bifunctional purine biosynthesis protein ATIC | 0.127121925 | -0.00712458 |
| P31942 | HNRNPH3 | Heterogeneous nuclear ribonucleoprotein H3 | -0.354375601 | 0.237237 |
| P31944 | CASP14 | Caspase-14 | -0.268315077 | 1.37932 |
| P31946 | YWHAB | 14-3-3 protein beta/alpha | 0.101731777 | -0.0993849 |
| P31946-2 | YWHAB | Isoform Short of 14-3-3 protein beta/alpha | -0.285573959 | -0.147246 |
| P31947 | SFN | 14-3-3 protein sigma | -0.289943695 | 1.27688 |
| P31948 | STIP1 | Stress-induced-phosphoprotein 1 | 0.10894982 | -0.00811577 |
| P31949 | S100A11 | Protein S100-A11 | -0.100076516 | 0.62378 |
| P32019 | INPP5B | Type II inositol 1,4,5-trisphosphate 5-phosphatase | -0.392560879 | 0.463235 |
| P32119 | PRDX2 | Peroxiredoxin-2 | 0.001687527 | 0.15498 |
| P32121 | ARRB2 | Beta-arrestin-2 | -0.146812677 | 0.55723 |
| P32189 | GK | Glycerol kinase | -0.402335087 | 0.329153 |
| P32321 | DCTD | Deoxycytidylate deaminase | -0.155025641 | 0.274583 |
| P32456 | GBP2 | Guanylate-binding protein 2 | 0.524153074 | -0.308858 |
| P32856 | STX2 | Syntaxin-2 | -0.376471678 | 0.455304 |
| P32926 | DSG3 | Desmoglein-3 | 0.20184048 | 1.33259 |
| P32929 | CTH | Cystathionine gamma-lyase | -1.521733443 | 0.408706 |
| P32969 | RPL9 | 60S ribosomal protein L9 | -0.662830273 | 0.0120751 |
| P33121 | ACSL1 | Long-chain-fatty-acid--CoA ligase 1 | -1.292073727 | -0.133492 |
| P33176 | KIF5B | Kinesin-1 heavy chain | -0.168257872 | 0.00791295 |
| P33316 | DUT | Deoxyuridine 5'-triphosphate nucleotidohydrolase, mitochondrial | 0.116310438 | 0.0669994 |
| P33316-2 | DUT | Isoform 2 of Deoxyuridine 5'-triphosphate nucleotidohydrolase, mitochondrial | 0.308289925 | 0.16142 |
| P33552 | CKS2 | Cyclin-dependent kinases regulatory subunit 2 | -1.372913758 | 0.560023 |
| P33981 | TTK | Dual specificity protein kinase TTK | -0.194069544 | -0.629306 |
| P33991 | MCM4 | DNA replication licensing factor MCM4 | 0.038289388 | 0.00232379 |
| P33992 | MCM5 | DNA replication licensing factor MCM5 | 0.029304345 | -0.270456 |
| P33993 | MCM7 | DNA replication licensing factor MCM7 | 0.057970842 | -0.0054156 |
| P34059 | GALNS | N-acetylgalactosamine-6-sulfatase | -0.845105489 | -0.926019 |
| P34810 | CD68 | Macrosialin | -2.047450225 | 0.00376987 |
| P34896 | SHMT1 | Serine hydroxymethyltransferase, cytosolic | 0.071453174 | 0.172599 |
| P34897 | SHMT2 | Serine hydroxymethyltransferase, mitochondrial | -0.680116494 | 0.261139 |
| P34932 | HSPA4 | Heat shock 70 kDa protein 4 | 0.101081053 | -0.027633 |
| P34949 | MPI | Mannose-6-phosphate isomerase | -0.386080901 | 0.0385507 |
| P35221 | CTNNA1 | Catenin alpha-1 | -0.390440623 | 0.0611684 |
| P35232 | PHB1 | Prohibitin 1 | -1.059310834 | 0.467342 |
| P35237 | SERPINB6 | Serpin B6 | -0.051129659 | -0.154212 |
| P35240 | NF2 | Merlin | -0.25085028 | -0.751216 |
| P35241 | RDX | Radixin | -0.224725723 | 0.110604 |
| P35244 | RPA3 | Replication protein A 14 kDa subunit | -0.01224184 | 0.679057 |
| P35249 | RFC4 | Replication factor C subunit 4 | -0.191793124 | 0.259064 |
| P35250 | RFC2 | Replication factor C subunit 2 | -0.159922997 | 0.02049 |
| P35251 | RFC1 | Replication factor C subunit 1 | -1.305436293 | 1.41548 |
| P35268 | RPL22 | 60S ribosomal protein L22 | -0.296020667 | 0.327749 |
| P35269 | GTF2F1 | General transcription factor IIF subunit 1 | 0.00027744 | -0.189456 |
| P35270 | SPR | Sepiapterin reductase | -0.02983586 | 0.173612 |
| P35520 | CBS | Cystathionine beta-synthase | -0.165925821 | -0.211823 |
| P35527 | KRT9 | Keratin, type I cytoskeletal 9 | -0.351871332 | -0.181087 |
| P35555 | FBN1 | Fibrillin-1 | -1.296051343 | 0.555186 |
| P35568 | IRS1 | Insulin receptor substrate 1 | -0.268207947 | 0.432583 |
| P35573 | AGL | Glycogen debranching enzyme | 0.227688869 | 0.0254227 |
| P35579 | MYH9 | Myosin-9 | -0.222325166 | -0.0374247 |
| P35580 | MYH10 | Myosin-10 | -0.363151073 | 0.0112753 |
| P35610 | SOAT1 | Sterol O-acyltransferase 1 | -2.351177454 | -0.204496 |
| P35612 | ADD2 | Beta-adducin | -0.003044287 | -0.168497 |
| P35658-3 | NUP214 | Isoform 3 of Nuclear pore complex protein Nup214 | 0.15118885 | 0.120278 |
| P35754 | GLRX | Glutaredoxin-1 | -0.457300742 | 0.23745 |
| P35790 | CHKA | Choline kinase alpha | -0.402129253 | 0.292031 |
| P35813 | PPM1A | Protein phosphatase 1A | 0.116729021 | -0.0920674 |
| P35908 | KRT2 | Keratin, type II cytoskeletal 2 epidermal | -0.127347787 | -0.228012 |

| | | | | |
|---|---|---|---|---|
| P35914 | HMGCL | Hydroxymethylglutaryl-CoA lyase, mitochondrial | -1.027471383 | -0.704726 |
| P36404 | ARL2 | ADP-ribosylation factor-like protein 2 | 0.015668154 | 0.16313 |
| P36405 | ARL3 | ADP-ribosylation factor-like protein 3 | 0.135023435 | -0.113906 |
| P36507 | MAP2K2 | Dual specificity mitogen-activated protein kinase kinase 2 | 0.128814538 | -0.0422363 |
| P36542 | ATP5F1C | ATP synthase subunit gamma, mitochondrial | -1.124692599 | 0.400925 |
| P36543 | ATP6V1E1 | V-type proton ATPase subunit E 1 | 0.001309951 | -0.104043 |
| P36551 | CPOX | Oxygen-dependent coproporphyrinogen-III oxidase, mitochondrial | -0.261075974 | 0.0568256 |
| P36578 | RPL4 | 60S ribosomal protein L4 | -1.208209753 | 0.272161 |
| P36639 | NUDT1 | Oxidized purine nucleoside triphosphate hydrolase | 0.143604358 | 0.172537 |
| P36871 | PGM1 | Phosphoglucomutase-1 | 0.153937181 | -0.0760161 |
| P36873 | PPP1CC | Serine/threonine-protein phosphatase PP1-gamma catalytic subunit | -0.289633433 | 0.181824 |
| P36915 | GNL1 | Guanine nucleotide-binding protein-like 1 | -0.053331057 | -0.133892 |
| P36954 | POLR2I | DNA-directed RNA polymerase II subunit RPB9 | -0.404591084 | 1.00005 |
| P36957 | DLST | Dihydrolipoyllysine-residue succinyltransferase component of 2-oxoglutarate dehydrogenase complex, mitochondrial | -0.810188055 | 0.551571 |
| P37108 | SRP14 | Signal recognition particle 14 kDa protein | -0.267443577 | -0.24439 |
| P37198 | NUP62 | Nuclear pore glycoprotein p62 | 0.116587241 | -0.123536 |
| P37235 | HPCAL1 | Hippocalcin-like protein 1 | -0.032071034 | 0.452279 |
| P37268 | FDFT1 | Squalene synthase | -1.122529666 | 0.339851 |
| P37802 | TAGLN2 | Transgelin-2 | 0.106088797 | 0.0771586 |
| P37837 | TALDO1 | Transaldolase | 0.155590057 | -0.0410029 |
| P38117 | ETFB | Electron transfer flavoprotein subunit beta | -0.298569679 | 0.0678959 |
| P38159 | RBMX | RNA-binding motif protein, X chromosome | -0.836897135 | 0.438882 |
| P38432 | COIL | Coilin | -0.194299777 | -0.379007 |
| P38435 | GGCX | Vitamin K-dependent gamma-carboxylase | -1.170742432 | 0.118961 |
| P38606 | ATP6V1A | V-type proton ATPase catalytic subunit A | 0.032022794 | -0.188716 |
| P38919 | EIF4A3 | Eukaryotic initiation factor 4A-III | -0.304114342 | -0.0626717 |
| P39019 | RPS19 | 40S ribosomal protein S19 | -0.198554675 | -0.50986 |
| P39023 | RPL3 | 60S ribosomal protein L3 | -1.281443278 | -0.131887 |
| P39687 | ANP32A | Acidic leucine-rich nuclear phosphoprotein 32 family member A | -0.132633209 | 0.00550143 |
| P39748 | FEN1 | Flap endonuclease 1 | -0.090906302 | -0.0818996 |
| P40121 | CAPG | Macrophage-capping protein | 0.238481204 | 0.147999 |
| P40123 | CAP2 | Adenylyl cyclase-associated protein 2 | 0.131011804 | -0.100116 |
| P40189 | IL6ST | Interleukin-6 receptor subunit beta | -0.017524878 | 0.160424 |
| P40222 | TXLNA | Alpha-taxilin | -0.651973963 | 0.0606505 |
| P40227 | CCT6A | T-complex protein 1 subunit zeta | -0.064933618 | -0.133113 |
| P40261 | NNMT | Nicotinamide N-methyltransferase | 0.08275938 | 0.0814362 |
| P40337 | VHL | von Hippel-Lindau disease tumor suppressor | -0.818939845 | 0.168684 |
| P40429 | RPL13A | 60S ribosomal protein L13a | -1.279100815 | 0.128436 |
| P40616 | ARL1 | ADP-ribosylation factor-like protein 1 | -0.309963703 | -0.190584 |
| P40818 | USP8 | Ubiquitin carboxyl-terminal hydrolase 8 | -1.158059676 | 0.315759 |
| P40855 | PEX19 | Peroxisomal biogenesis factor 19 | 1.308112224 | 0.503359 |
| P40925 | MDH1 | Malate dehydrogenase, cytoplasmic | 0.023899873 | -0.118611 |
| P40926 | MDH2 | Malate dehydrogenase, mitochondrial | -0.25799195 | 0.236714 |
| P40937 | RFC5 | Replication factor C subunit 5 | -0.299526374 | -0.209516 |
| P40938 | RFC3 | Replication factor C subunit 3 | -0.682733138 | 0.0724042 |
| P41091 | EIF2S3 | Eukaryotic translation initiation factor 2 subunit 3 | -0.214502176 | -0.0541331 |
| P41208 | CETN2 | Centrin-2 | -0.412500858 | -0.012523 |
| P41214 | EIF2D | Eukaryotic translation initiation factor 2D | -0.810390234 | -0.546084 |
| P41223 | BUD31 | Protein BUD31 homolog | -1.904951572 | 0.367621 |
| P41226 | UBA7 | Ubiquitin-like modifier-activating enzyme 7 | 1.427345514 | -0.629633 |
| P41227 | NAA10 | N-alpha-acetyltransferase 10 | 0.083508333 | 0.00174014 |
| P41229 | KDM5C | Lysine-specific demethylase 5C | -0.035523891 | -0.0711006 |
| P41236 | PPP1R2 | Protein phosphatase inhibitor 2 | 0.289945682 | 0.731709 |
| P41240 | CSK | Tyrosine-protein kinase CSK | -0.131900311 | 0.0715663 |
| P41250-2 | GARS1 | Isoform 2 of Glycine--tRNA ligase | 0.03407224 | -0.0237579 |
| P41743 | PRKCI | Protein kinase C iota type | -0.19587938 | -0.487662 |
| P42025 | ACTR1B | Beta-centractin | 0.170653105 | -0.768469 |
| P42126 | ECI1 | Enoyl-CoA delta isomerase 1, mitochondrial | -0.446857929 | 0.128598 |
| P42166 | TMPO | Lamina-associated polypeptide 2, isoform alpha | -1.083908478 | 0.537498 |
| P42167 | TMPO | Lamina-associated polypeptide 2, isoforms beta/gamma | -0.637833834 | 0.282425 |
| P42285 | MTREX | Exosome RNA helicase MTR4 | -0.386731704 | -0.00370979 |
| P42338 | PIK3CB | Phosphatidylinositol 4,5-bisphosphate 3-kinase catalytic subunit beta isoform | -1.016629616 | 0.817439 |
| P42345 | MTOR | Serine/threonine-protein kinase mTOR | -0.95440491 | 0.0836706 |
| P42356 | PI4KA | Phosphatidylinositol 4-kinase alpha | -0.659952323 | 0.907716 |
| P42357 | HAL | Histidine ammonia-lyase | -0.563282331 | 1.00406 |
| P42566 | EPS15 | Epidermal growth factor receptor substrate 15 | 0.204676708 | -0.182853 |
| P42574 | CASP3 | Caspase-3 | -0.098969142 | -0.295324 |
| P42575 | CASP2 | Caspase-2 | -0.068101088 | 0.0405916 |

| | | | | |
|---|---|---|---|---|
| P42677 | RPS27 | 40S ribosomal protein S27 | -0.769185384 | -0.385256 |
| P42695 | NCAPD3 | Condensin-2 complex subunit D3 | -0.661769112 | 0.796022 |
| P42696 | RBM34 | RNA-binding protein 34 | -2.436802904 | 0.844122 |
| P42765 | ACAA2 | 3-ketoacyl-CoA thiolase, mitochondrial | -0.382981221 | 0.159985 |
| P42766 | RPL35 | 60S ribosomal protein L35 | -1.119437774 | -0.260613 |
| P43121 | MCAM | Cell surface glycoprotein MUC18 | -1.061173439 | 0.399609 |
| P43246 | MSH2 | DNA mismatch repair protein Msh2 | 0.092782656 | 0.0911582 |
| P43304 | GPD2 | Glycerol-3-phosphate dehydrogenase, mitochondrial | -1.942280769 | -0.596506 |
| P43487 | RANBP1 | Ran-specific GTPase-activating protein | -0.022206624 | -0.0935707 |
| P43686 | PSMC4 | 26S proteasome regulatory subunit 6B | -0.228003899 | 0.00663503 |
| P43897 | TSFM | Elongation factor Ts, mitochondrial | -0.69487373 | 0.342064 |
| P45877 | PPIC | Peptidyl-prolyl cis-trans isomerase C | -0.981630643 | -2.02091 |
| P45880 | VDAC2 | Voltage-dependent anion-selective channel protein 2 | -1.24889334 | 0.377103 |
| P45954 | ACADSB | Short/branched chain specific acyl-CoA dehydrogenase, mitochondrial | -0.860738357 | 0.885396 |
| P45973 | CBX5 | Chromobox protein homolog 5 | -0.024715185 | 0.0652695 |
| P45974 | USP5 | Ubiquitin carboxyl-terminal hydrolase 5 | 0.112023989 | -0.114192 |
| P45983-3 | MAPK8 | Isoform 3 of Mitogen-activated protein kinase 8 | -0.764041026 | 0.638368 |
| P45984 | MAPK9 | Mitogen-activated protein kinase 9 | -0.008989414 | -0.125964 |
| P45985 | MAP2K4 | Dual specificity mitogen-activated protein kinase kinase 4 | -0.108286699 | -0.0654383 |
| P46013 | MKI67 | Proliferation marker protein Ki-67 | -1.398649772 | 0.591997 |
| P46060 | RANGAP1 | Ran GTPase-activating protein 1 | 0.1096193 | 0.142433 |
| P46063 | RECQL | ATP-dependent DNA helicase Q1 | -0.743340572 | 0.114091 |
| P46087 | NOP2 | Probable 28S rRNA (cytosine(4447)-C(5))-methyltransferase | -0.773862521 | 0.489974 |
| P46100 | ATRX | Transcriptional regulator ATRX | 0.784229596 | 0 |
| P46108 | CRK | Adapter molecule crk | 0.139426072 | -0.0570443 |
| P46108-2 | CRK | Isoform Crk-I of Adapter molecule crk | 0.304447174 | -0.220463 |
| P46109 | CRKL | Crk-like protein | 0.18294239 | -0.0575571 |
| P46734 | MAP2K3 | Dual specificity mitogen-activated protein kinase kinase 3 | -0.046703021 | 0.188982 |
| P46734-2 | MAP2K3 | Isoform 1 of Dual specificity mitogen-activated protein kinase kinase 3 | -0.169011037 | 0.266148 |
| P46736 | BRCC3 | Lys-63-specific deubiquitinase BRCC36 | -0.264355342 | -0.159367 |
| P46776 | RPL27A | 60S ribosomal protein L27a | -1.103135427 | 0.104755 |
| P46777 | RPL5 | 60S ribosomal protein L5 | -0.176696301 | -0.53522 |
| P46778 | RPL21 | 60S ribosomal protein L21 | -1.553955317 | -0.347506 |
| P46779 | RPL28 | 60S ribosomal protein L28 | -1.276073933 | -0.20852 |
| P46781 | RPS9 | 40S ribosomal protein S9 | -0.173576037 | -0.376453 |
| P46783 | RPS10 | 40S ribosomal protein S10 | -0.190670331 | -0.202321 |
| P46821 | MAP1B | Microtubule-associated protein 1B | -0.442562262 | 0.158191 |
| P46926 | GNPDA1 | Glucosamine-6-phosphate isomerase 1 | -0.052193801 | -0.241553 |
| P46934-4 | NEDD4 | Isoform 4 of E3 ubiquitin-protein ligase NEDD4 | 0.000883738 | -0.113406 |
| P46937 | YAP1 | Transcriptional coactivator YAP1 | 0.353248517 | 0.252046 |
| P46939 | UTRN | Utrophin | -1.203113159 | -0.542079 |
| P46940 | IQGAP1 | Ras GTPase-activating-like protein IQGAP1 | 0.018937429 | 0.0130501 |
| P46976 | GYG1 | Glycogenin-1 | -0.277474801 | -0.246407 |
| P46977 | STT3A | Dolichyl-diphosphooligosaccharide--protein glycosyltransferase subunit STT3A | -1.390214125 | 0.262691 |
| P47224 | RABIF | Guanine nucleotide exchange factor MSS4 | -0.053087552 | 0.463802 |
| P47712 | PLA2G4A | Cytosolic phospholipase A2 | -0.069703023 | -0.803934 |
| P47755 | CAPZA2 | F-actin-capping protein subunit alpha-2 | 0.010766347 | 0.0351251 |
| P47756-2 | CAPZB | Isoform 2 of F-actin-capping protein subunit beta | 0.080909729 | 0.0193501 |
| P47813 | EIF1AX | Eukaryotic translation initiation factor 1A, X-chromosomal | -0.229757627 | 0.139489 |
| P47895 | ALDH1A3 | Aldehyde dehydrogenase family 1 member A3 | 0.081878026 | -0.0662308 |
| P47897 | QARS1 | Glutamine--tRNA ligase | -0.120240529 | -0.0376765 |
| P47929 | LGALS7 | Galectin-7 | -0.733900388 | 1.03486 |
| P47974 | ZFP36L2 | mRNA decay activator protein ZFP36L2 | -0.585220098 | -1.31039 |
| P47985 | UQCRFS1 | Cytochrome b-c1 complex subunit Rieske, mitochondrial | -0.979847272 | 0.346127 |
| P48047 | ATP5PO | ATP synthase subunit O, mitochondrial | -1.080716451 | 0.527376 |
| P48059-3 | LIMS1 | Isoform 3 of LIM and senescent cell antigen-like-containing domain protein 1 | -0.190803607 | 0.148616 |
| P48066 | SLC6A11 | Sodium- and chloride-dependent GABA transporter 3 | -0.007369995 | -0.944617 |
| P48147 | PREP | Prolyl endopeptidase | -0.193890492 | -0.089894 |
| P48163 | ME1 | NADP-dependent malic enzyme | -0.173488855 | 0.0609849 |
| P48200 | IREB2 | Iron-responsive element-binding protein 2 | 0.12011075 | 0.774362 |
| P48307 | TFPI2 | Tissue factor pathway inhibitor 2 | -0.557031075 | 0.63534 |
| P48426 | PIP4K2A | Phosphatidylinositol 5-phosphate 4-kinase type-2 alpha | 0.078098774 | 0.382392 |
| P48436 | SOX9 | Transcription factor SOX-9 | -0.058368127 | -0.473476 |
| P48444 | ARCN1 | Coatomer subunit delta | 0.132317066 | 0.00224431 |
| P48449 | LSS | Lanosterol synthase | -0.961106936 | -0.149766 |
| P48454 | PPP3CC | Serine/threonine-protein phosphatase 2B catalytic subunit gamma isoform | -0.185272455 | -0.12457 |

| | | | | |
|---|---|---|---|---|
| P48506 | GCLC | Glutamate--cysteine ligase catalytic subunit | 0.016071637 | 0.0823806 |
| P48507 | GCLM | Glutamate--cysteine ligase regulatory subunit | -0.071543058 | -0.409715 |
| P48556 | PSMD8 | 26S proteasome non-ATPase regulatory subunit 8 | -0.155091763 | 0.0637881 |
| P48594 | SERPINB4 | Serpin B4 | -1.102409601 | 0.0968119 |
| P48634 | PRRC2A | Protein PRRC2A | -0.977194468 | -1.27792 |
| P48637 | GSS | Glutathione synthetase | 0.037916025 | -0.117787 |
| P48643 | CCT5 | T-complex protein 1 subunit epsilon | -0.070773602 | 0.00933011 |
| P48651 | PTDSS1 | Phosphatidylserine synthase 1 | -3.007256031 | -0.50844 |
| P48668 | KRT6C | Keratin, type II cytoskeletal 6C | -2.767449617 | -1.93184 |
| P48681 | NES | Nestin | 0.222103596 | 0.117239 |
| P48723 | HSPA13 | Heat shock 70 kDa protein 13 | -0.426142375 | 0.144917 |
| P48735 | IDH2 | Isocitrate dehydrogenase [NADP], mitochondrial | -0.797040065 | 0.369264 |
| P48739 | PITPNB | Phosphatidylinositol transfer protein beta isoform | 0.10118564 | -0.0396061 |
| P49006 | MARCKSL1 | MARCKS-related protein | -0.089033445 | 0.413721 |
| P49137 | MAPKAPK2 | MAP kinase-activated protein kinase 2 | -0.061563094 | 0.0396388 |
| P49189 | ALDH9A1 | 4-trimethylaminobutyraldehyde dehydrogenase | 0.106927872 | 0.138621 |
| P49207 | RPL34 | 60S ribosomal protein L34 | -1.608252923 | -0.205102 |
| P49247 | RPIA | Ribose-5-phosphate isomerase | -0.227760235 | 0.11946 |
| P49257 | LMAN1 | Protein ERGIC-53 | -0.985720158 | 0.331992 |
| P49321 | NASP | Nuclear autoantigenic sperm protein | 0.27178065 | -0.066939 |
| P49327 | FASN | Fatty acid synthase | 0.018112659 | -0.113419 |
| P49354 | FNTA | Protein farnesyltransferase/geranylgeranyltransferase type-1 subunit alpha | 0.02660807 | -0.332719 |
| P49356 | FNTB | Protein farnesyltransferase subunit beta | -0.160529137 | 0.171426 |
| P49366 | DHPS | Deoxyhypusine synthase | 0.101039966 | -0.256533 |
| P49368 | CCT3 | T-complex protein 1 subunit gamma | -0.024385611 | -0.210454 |
| P49406 | MRPL19 | 39S ribosomal protein L19, mitochondrial | -1.247117678 | -0.234057 |
| P49407 | ARRB1 | Beta-arrestin-1 | -0.263866822 | 0.0909392 |
| P49411 | TUFM | Elongation factor Tu, mitochondrial | -0.649628242 | 0.162962 |
| P49418 | AMPH | Amphiphysin | 0.11905098 | -0.421123 |
| P49419-2 | ALDH7A1 | Isoform 2 of Alpha-aminoadipic semialdehyde dehydrogenase | -0.009731929 | 0.076184 |
| P49427 | CDC34 | Ubiquitin-conjugating enzyme E2 R1 | -0.018258095 | 0.279525 |
| P49441 | INPP1 | Inositol polyphosphate 1-phosphatase | -0.454651515 | 1.39248 |
| P49454 | CENPF | Centromere protein F | 5.150284688 | -1.36431 |
| P49458 | SRP9 | Signal recognition particle 9 kDa protein | -0.604950905 | 0.354116 |
| P49585 | PCYT1A | Choline-phosphate cytidylyltransferase A | 0.000293414 | -0.0719433 |
| P49589-3 | CARS1 | Isoform 3 of Cysteine--tRNA ligase, cytoplasmic | 0.05950133 | -0.0778573 |
| P49591 | SARS1 | Serine--tRNA ligase, cytoplasmic | -0.242434184 | 0.0944684 |
| P49593 | PPM1F | Protein phosphatase 1F | 0.158866882 | 0.079423 |
| P49662 | CASP4 | Caspase-4 | 0.11468641 | -0.0226383 |
| P49720 | PSMB3 | Proteasome subunit beta type-3 | -0.477015495 | -0.253591 |
| P49721 | PSMB2 | Proteasome subunit beta type-2 | -0.329717716 | 0.0953875 |
| P49736 | MCM2 | DNA replication licensing factor MCM2 | 0.037897746 | -0.029988 |
| P49748 | ACADVL | Very long-chain specific acyl-CoA dehydrogenase, mitochondrial | -0.957895279 | 0.117802 |
| P49750 | YLPM1 | YLP motif-containing protein 1 | -0.341779947 | 0.131477 |
| P49754 | VPS41 | Vacuolar protein sorting-associated protein 41 homolog | -1.618031224 | -0.00508785 |
| P49755 | TMED10 | Transmembrane emp24 domain-containing protein 10 | -1.166840394 | -0.092082 |
| P49756 | RBM25 | RNA-binding protein 25 | -0.019777139 | 0.0559069 |
| P49757 | NUMB | Protein numb homolog | -1.345074733 | 0.651612 |
| P49770 | EIF2B2 | Translation initiation factor eIF-2B subunit beta | -0.065460364 | -0.128044 |
| P49773 | HINT1 | Adenosine 5'-monophosphoramidase HINT1 | -0.4888738 | -0.0603065 |
| P49790 | NUP153 | Nuclear pore complex protein Nup153 | -0.035448949 | 0.342361 |
| P49792 | RANBP2 | E3 SUMO-protein ligase RanBP2 | -0.083075841 | 0.394453 |
| P49862 | KLK7 | Kallikrein-7 | -1.073448181 | 0.597617 |
| P49903 | SEPHS1 | Selenide, water dikinase 1 | 0.12012887 | -0.401652 |
| P49914 | MTHFS | 5-formyltetrahydrofolate cyclo-ligase | 0.125082652 | 0.106498 |
| P49915 | GMPS | GMP synthase [glutamine-hydrolyzing] | 0.083498478 | 0.00643476 |
| P49916 | LIG3 | DNA ligase 3 | -1.91264677 | 0.456887 |
| P49959 | MRE11 | Double-strand break repair protein MRE11 | -0.033280611 | -0.000728289 |
| P50148 | GNAQ | Guanine nucleotide-binding protein G(q) subunit alpha | -1.369444211 | 0.164299 |
| P50213 | IDH3A | Isocitrate dehydrogenase [NAD] subunit alpha, mitochondrial | -0.270598809 | -0.00988166 |
| P50281 | MMP14 | Matrix metalloproteinase-14 | -0.89648215 | 0.212391 |
| P50395 | GDI2 | Rab GDP dissociation inhibitor beta | 0.015197754 | -0.0719357 |
| P50402 | EMD | Emerin | -0.957080046 | 0.196174 |
| P50416 | CPT1A | Carnitine O-palmitoyltransferase 1, liver isoform | -1.126255751 | 0.140397 |
| P50452 | SERPINB8 | Serpin B8 | 0.078523477 | 0.00658099 |
| P50453 | SERPINB9 | Serpin B9 | 0.148626328 | 0.129749 |
| P50454 | SERPINH1 | Serpin H1 | -0.689685822 | 0.193521 |
| P50479 | PDLIM4 | PDZ and LIM domain protein 4 | -0.026000341 | 0.22831 |
| P50502 | ST13 | Hsc70-interacting protein | 0.099881331 | -0.0118968 |
| P50548 | ERF | ETS domain-containing transcription factor ERF | 0.212113221 | -0.334967 |

| | | | | |
|---|---|---|---|---|
| P50552 | VASP | Vasodilator-stimulated phosphoprotein | -0.215844631 | -0.0792058 |
| P50570-2 | DNM2 | Isoform 2 of Dynamin-2 | -0.299537182 | 0.32129 |
| P50570 | DNM2 | Dynamin-2 | -0.399067879 | 0.252123 |
| P50579 | METAP2 | Methionine aminopeptidase 2 | -0.349100987 | -0.0585136 |
| P50583 | NUDT2 | Bis(5'-nucleosyl)-tetraphosphatase [asymmetrical] | -0.192561467 | -0.0122045 |
| P50613 | CDK7 | Cyclin-dependent kinase 7 | -0.291924795 | 0.0820065 |
| P50748 | KNTC1 | Kinetochore-associated protein 1 | -0.02200071 | -0.31298 |
| P50750 | CDK9 | Cyclin-dependent kinase 9 | -0.222669919 | -0.222489 |
| P50914 | RPL14 | 60S ribosomal protein L14 | -1.369108041 | -0.0841122 |
| P50990 | CCT8 | T-complex protein 1 subunit theta | 0.010778586 | -0.236775 |
| P50991 | CCT4 | T-complex protein 1 subunit delta | -0.190948963 | -0.218088 |
| P50995 | ANXA11 | Annexin A11 | 0.116128445 | -0.0158304 |
| P51116 | FXR2 | Fragile X mental retardation syndrome-related protein 2 | -0.841983716 | -0.00416756 |
| P51148 | RAB5C | Ras-related protein Rab-5C | -0.578989188 | 0.0317059 |
| P51149 | RAB7A | Ras-related protein Rab-7a | -0.710311413 | 0.0612545 |
| P51151 | RAB9A | Ras-related protein Rab-9A | -0.343341827 | 0.0590026 |
| P51153 | RAB13 | Ras-related protein Rab-13 | -1.003855785 | 0.0771796 |
| P51157 | RAB28 | Ras-related protein Rab-28 | -0.247009675 | -0.820775 |
| P51159 | RAB27A | Ras-related protein Rab-27A | -0.817721844 | -0.201183 |
| P51178 | PLCD1 | 1-phosphatidylinositol 4,5-bisphosphate phosphodiesterase delta-1 | -0.43558534 | -1.61319 |
| P51398 | DAP3 | 28S ribosomal protein S29, mitochondrial | -1.339530945 | 0.532591 |
| P51452 | DUSP3 | Dual specificity protein phosphatase 3 | 0.081472079 | 0.0730184 |
| P51553 | IDH3G | Isocitrate dehydrogenase [NAD] subunit gamma, mitochondrial | -0.729613225 | -0.19598 |
| P51570 | GALK1 | Galactokinase | 0.161802133 | -0.00371234 |
| P51571 | SSR4 | Translocon-associated protein subunit delta | -1.401270469 | 0.550159 |
| P51572 | BCAP31 | B-cell receptor-associated protein 31 | -1.050283591 | 0.348774 |
| P51580 | TPMT | Thiopurine S-methyltransferase | -0.08033967 | -0.121697 |
| P51608 | MECP2 | Methyl-CpG-binding protein 2 | -0.719959418 | 0.801263 |
| P51636 | CAV2 | Caveolin-2 | -2.435323715 | -1.62869 |
| P51665 | PSMD7 | 26S proteasome non-ATPase regulatory subunit 7 | -0.364280462 | -0.219945 |
| P51668 | UBE2D1 | Ubiquitin-conjugating enzyme E2 D1 | 0.124210676 | -0.133467 |
| P51687 | SUOX | Sulfite oxidase, mitochondrial | -0.359102885 | -0.58689 |
| P51688 | SGSH | N-sulphoglucosamine sulphohydrolase | -0.404233138 | -1.44142 |
| P51692 | STAT5B | Signal transducer and activator of transcription 5B | -0.012608131 | -0.0366217 |
| P51809 | VAMP7 | Vesicle-associated membrane protein 7 | -1.05135711 | -2.48689 |
| P51812 | RPS6KA3 | Ribosomal protein S6 kinase alpha-3 | -0.009113471 | -0.0719045 |
| P51858 | HDGF | Hepatoma-derived growth factor | 0.289502462 | 0.0803846 |
| P51911 | CNN1 | Calponin-1 | -0.205179453 | -0.0756674 |
| P51948 | MNAT1 | CDK-activating kinase assembly factor MAT1 | -0.156826337 | 0.291571 |
| P51970 | NDUFA8 | NADH dehydrogenase [ubiquinone] 1 alpha subcomplex subunit 8 | -1.066927354 | -0.325753 |
| P51991-2 | HNRNPA3 | Isoform 2 of Heterogeneous nuclear ribonucleoprotein A3 | -1.190704664 | -0.298236 |
| P52209 | PGD | 6-phosphogluconate dehydrogenase, decarboxylating | -0.105850856 | -0.115676 |
| P52294 | KPNA1 | Importin subunit alpha-5 | -0.008025487 | -0.259664 |
| P52298 | NCBP2 | Nuclear cap-binding protein subunit 2 | -0.140190125 | 0.287333 |
| P52306 | RAP1GDS1 | Rap1 GTPase-GDP dissociation stimulator 1 | -1.610867818 | -0.620309 |
| P52306-5 | RAP1GDS1 | Isoform 5 of Rap1 GTPase-GDP dissociation stimulator 1 | 0.048086325 | -0.116295 |
| P52565 | ARHGDIA | Rho GDP-dissociation inhibitor 1 | 0.264625072 | -0.12986 |
| P52566 | ARHGDIB | Rho GDP-dissociation inhibitor 2 | 0.224190871 | -0.261312 |
| P52594 | AGFG1 | Arf-GAP domain and FG repeat-containing protein 1 | -0.261739016 | -0.114431 |
| P52597 | HNRNPF | Heterogeneous nuclear ribonucleoprotein F | 0.126692295 | 0.00168737 |
| P52655 | GTF2A1 | Transcription initiation factor IIA subunit 1 | -1.722308238 | 0.873932 |
| P52701 | MSH6 | DNA mismatch repair protein Msh6 | -0.055522283 | 0.0560913 |
| P52732 | KIF11 | Kinesin-like protein KIF11 | -0.079387665 | -0.102282 |
| P52735 | VAV2 | Guanine nucleotide exchange factor VAV2 | -0.245033662 | -0.333616 |
| P52756 | RBM5 | RNA-binding protein 5 | -1.043406804 | -1.29228 |
| P52758 | RIDA | 2-iminobutanoate/2-iminopropanoate deaminase | -0.273453712 | -0.205919 |
| P52788 | SMS | Spermine synthase | 0.147732258 | -0.0373453 |
| P52789 | HK2 | Hexokinase-2 | 0.138851802 | 0.119798 |
| P52888 | THOP1 | Thimet oligopeptidase | 0.090474765 | 0.0862433 |
| P52907 | CAPZA1 | F-actin-capping protein subunit alpha-1 | 0.006236871 | -0.0927118 |
| P52943 | CRIP2 | Cysteine-rich protein 2 | -0.858825048 | -0.0898285 |
| P52948 | NUP98 | Nuclear pore complex protein Nup98-Nup96 | -0.09978048 | 0.244129 |
| P53004 | BLVRA | Biliverdin reductase A | 0.132091681 | -0.13588 |
| P53007 | SLC25A1 | Tricarboxylate transport protein, mitochondrial | -1.399582624 | 0.0907234 |
| P53350 | PLK1 | Serine/threonine-protein kinase PLK1 | -0.979619185 | 0.275922 |
| P53367 | ARFIP1 | Arfaptin-1 | 0.203717947 | -0.0901759 |
| P53367-2 | ARFIP1 | Isoform A of Arfaptin-1 | -0.253760576 | -0.162572 |
| P53384 | NUBP1 | Cytosolic Fe-S cluster assembly factor NUBP1 | 0.05529658 | -0.108993 |
| P53396 | ACLY | ATP-citrate synthase | -0.111057123 | 0.0138372 |
| P53582 | METAP1 | Methionine aminopeptidase 1 | -0.468651692 | 0.410237 |

| | | | | |
|---|---|---|---|---|
| P53597 | SUCLG1 | Succinate--CoA ligase [ADP/GDP-forming] subunit alpha, mitochondrial | -0.433537404 | 0.405924 |
| P53602 | MVD | Diphosphomevalonate decarboxylase | 0.403975964 | 0.107212 |
| P53609 | PGGT1B | Geranylgeranyl transferase type-1 subunit beta | 0.087447564 | 0.0269527 |
| P53611 | RABGGTB | Geranylgeranyl transferase type-2 subunit beta | 0.034594297 | 0.231452 |
| P53618 | COPB1 | Coatomer subunit beta | -0.041273912 | -0.218877 |
| P53621 | COPA | Coatomer subunit alpha | 0.000447591 | -0.194153 |
| P53677 | AP3M2 | AP-3 complex subunit mu-2 | -0.181877454 | -0.388843 |
| P53701 | HCCS | Holocytochrome c-type synthase | -1.742257754 | 0.248176 |
| P53778 | MAPK12 | Mitogen-activated protein kinase 12 | -1.554139694 | -1.61356 |
| P53814 | SMTN | Smoothelin | -0.209826152 | 0.160178 |
| P53985 | SLC16A1 | Monocarboxylate transporter 1 | -0.838930527 | -0.202617 |
| P53992 | SEC24C | Protein transport protein Sec24C | -0.348031521 | 0.137727 |
| P53999 | SUB1 | Activated RNA polymerase II transcriptional coactivator p15 | 0.041601022 | -0.176009 |
| P54136 | RARS1 | Arginine--tRNA ligase, cytoplasmic | -0.10456419 | 0.018376 |
| P54136-2 | RARS1 | Isoform Monomeric of Arginine--tRNA ligase, cytoplasmic | -1.061790307 | 0.0463171 |
| P54259 | ATN1 | Atrophin-1 | 0.340107361 | 0.829755 |
| P54577 | YARS1 | Tyrosine--tRNA ligase, cytoplasmic | 0.005712668 | -0.00509262 |
| P54578 | USP14 | Ubiquitin carboxyl-terminal hydrolase 14 | 0.070390066 | -0.100381 |
| P54619 | PRKAG1 | 5'-AMP-activated protein kinase subunit gamma-1 | -0.028966188 | -0.0222893 |
| P54652 | HSPA2 | Heat shock-related 70 kDa protein 2 | 0.061382135 | -0.118748 |
| P54687 | BCAT1 | Branched-chain-amino-acid aminotransferase, cytosolic | 0.149624983 | -0.00424957 |
| P54709 | ATP1B3 | Sodium/potassium-transporting ATPase subunit beta-3 | -1.333281438 | 0.341742 |
| P54727 | RAD23B | UV excision repair protein RAD23 homolog B | 0.140219688 | 0.133113 |
| P54802 | NAGLU | Alpha-N-acetylglucosaminidase | -1.027727524 | -0.571706 |
| P54886 | ALDH18A1 | Delta-1-pyrroline-5-carboxylate synthase | -0.424816211 | 0.02239 |
| P54920 | NAPA | Alpha-soluble NSF attachment protein | -0.012943586 | 0.014253 |
| P55000 | SLURP1 | Secreted Ly-6/uPAR-related protein 1 | 0.71864899 | 0 |
| P55010 | EIF5 | Eukaryotic translation initiation factor 5 | 0.011209329 | -0.0676994 |
| P55011 | SLC12A2 | Solute carrier family 12 member 2 | -1.296813011 | -0.605164 |
| P55036 | PSMD4 | 26S proteasome non-ATPase regulatory subunit 4 | -0.315951268 | 0.0817337 |
| P55039 | DRG2 | Developmentally-regulated GTP-binding protein 2 | -0.085310539 | -0.235361 |
| P55040 | GEM | GTP-binding protein GEM | -0.420711358 | -0.0792952 |
| P55042 | RRAD | GTP-binding protein RAD | -0.330527226 | -0.0321296 |
| P55060 | CSE1L | Exportin-2 | -0.101181825 | -0.132102 |
| P55081 | MFAP1 | Microfibrillar-associated protein 1 | -0.185598532 | 0.112535 |
| P55084 | HADHB | Trifunctional enzyme subunit beta, mitochondrial | -0.810635805 | 0.253284 |
| P55145 | MANF | Mesencephalic astrocyte-derived neurotrophic factor | -0.775478522 | 0.0312678 |
| P55210 | CASP7 | Caspase-7 | 0.561682383 | 0.131709 |
| P55212 | CASP6 | Caspase-6 | 0.012156566 | -0.21444 |
| P55263 | ADK | Adenosine kinase | -0.025080522 | 0.0909087 |
| P55268 | LAMB2 | Laminin subunit beta-2 | 0.612895966 | 0.407287 |
| P55290 | CDH13 | Cadherin-13 | 0.143606901 | -0.42685 |
| P55327-2 | TPD52 | Isoform 2 of Tumor protein D52 | -0.053988377 | 0.0183719 |
| P55735 | SEC13 | Protein SEC13 homolog | -0.174945196 | -0.220345 |
| P55769 | SNU13 | NHP2-like protein 1 | -0.49919947 | 0.287811 |
| P55786-2 | NPEPPS | Isoform 2 of Puromycin-sensitive aminopeptidase | 0.312877019 | 0.174717 |
| P55789 | GFER | FAD-linked sulfhydryl oxidase ALR | -0.463342587 | 0.28004 |
| P55795 | HNRNPH2 | Heterogeneous nuclear ribonucleoprotein H2 | -0.467649778 | 0.474054 |
| P55809 | OXCT1 | Succinyl-CoA:3-ketoacid coenzyme A transferase 1, mitochondrial | -0.2814641 | -0.0177021 |
| P55884 | EIF3B | Eukaryotic translation initiation factor 3 subunit B | 0.062051137 | -0.192173 |
| P55957 | BID | BH3-interacting domain death agonist | 0.293975512 | 0.472572 |
| P56134 | ATP5MF | ATP synthase subunit f, mitochondrial | -1.316804488 | -0.521077 |
| P56181-2 | NDUFV3 | Isoform 2 of NADH dehydrogenase [ubiquinone] flavoprotein 3, mitochondrial | -1.483486295 | 0.496335 |
| P56182 | RRP1 | Ribosomal RNA processing protein 1 homolog A | -1.105425994 | -0.0481567 |
| P56192 | MARS1 | Methionine--tRNA ligase, cytoplasmic | -0.378455242 | -0.129675 |
| P56282 | POLE2 | DNA polymerase epsilon subunit 2 | -0.354864438 | 0 |
| P56377 | AP1S2 | AP-1 complex subunit sigma-2 | 0.152074337 | -0.456985 |
| P56381 | ATP5F1E | ATP synthase subunit epsilon, mitochondrial | -1.236243884 | 0.396473 |
| P56385 | ATP5ME | ATP synthase subunit e, mitochondrial | -0.508994818 | 0.784633 |
| P56537 | EIF6 | Eukaryotic translation initiation factor 6 | 0.009119352 | 0.166643 |
| P56545 | CTBP2 | C-terminal-binding protein 2 | 0.088011742 | -0.0594571 |
| P56937 | HSD17B7 | 3-keto-steroid reductase/17-beta-hydroxysteroid dehydrogenase 7 | -0.814782143 | -0.253575 |
| P56945 | BCAR1 | Breast cancer anti-estrogen resistance protein 1 | 0.09416008 | 0.052762 |
| P57076 | CFAP298 | Cilia- and flagella-associated protein 298 | 0.11382095 | -0.959366 |
| P57081 | WDR4 | tRNA (guanine-N(7)-)-methyltransferase non-catalytic subunit WDR4 | -0.048926433 | -0.119723 |
| P57088 | TMEM33 | Transmembrane protein 33 | -1.718729019 | -0.168844 |
| P57729 | RAB38 | Ras-related protein Rab-38 | -1.120942195 | -0.770257 |

| | | | | |
|---|---|---|---|---|
| P57740 | NUP107 | Nuclear pore complex protein Nup107 | 0.008708239 | 0.251086 |
| P57768 | SNX16 | Sorting nexin-16 | -0.307390451 | -0.219732 |
| P57772 | EEFSEC | Selenocysteine-specific elongation factor | -1.333395402 | 2.81466 |
| P58004 | SESN2 | Sestrin-2 | -0.787573814 | -2.86173 |
| P58546 | MTPN | Myotrophin | 0.160252094 | 0.543387 |
| P58876 | H2BC5 | Histone H2B type 1-D | -0.868341446 | 0.128286 |
| P59665 | DEFA1 | Neutrophil defensin 1 | -0.320138534 | -0.367456 |
| P59998 | ARPC4 | Actin-related protein 2/3 complex subunit 4 | -0.086414496 | 0.10608 |
| P60059 | SEC61G | Protein transport protein Sec61 subunit gamma | -0.586274624 | -0.587254 |
| P60174 | TPI1 | Triosephosphate isomerase | 0.073022207 | 0.0709712 |
| P60201 | PLP1 | Myelin proteolipid protein | 0.369155486 | -0.365547 |
| P60468 | SEC61B | Protein transport protein Sec61 subunit beta | -1.387686888 | 0.135989 |
| P60484 | PTEN | Phosphatidylinositol 3,4,5-trisphosphate 3-phosphatase and dual-specificity protein phosphatase PTEN | 0.061134895 | -0.501186 |
| P60510 | PPP4C | Serine/threonine-protein phosphatase 4 catalytic subunit | -0.093853553 | -0.150123 |
| P60520 | GABARAPL2 | Gamma-aminobutyric acid receptor-associated protein-like 2 | -0.337570826 | -0.158311 |
| P60602 | ROMO1 | Reactive oxygen species modulator 1 | -2.09930555 | 0.933346 |
| P60604 | UBE2G2 | Ubiquitin-conjugating enzyme E2 G2 | -0.87126112 | 0.315203 |
| P60608 | ERVFC1-1 | Endogenous retrovirus group FC1 member 1 Env polyprotein | -1.723416646 | 6.47081 |
| P60709 | ACTB | Actin, cytoplasmic 1 | 0.46126461 | 1.44036 |
| P60763 | RAC3 | Ras-related C3 botulinum toxin substrate 3 | -2.370503028 | 0.425743 |
| P60842 | EIF4A1 | Eukaryotic initiation factor 4A-I | -0.050144196 | -0.284387 |
| P60866 | RPS20 | 40S ribosomal protein S20 | -0.405227423 | -0.47182 |
| P60903 | S100A10 | Protein S100-A10 | -0.594467084 | -0.286237 |
| P60953 | CDC42 | Cell division control protein 42 homolog | -0.383455435 | 0.304354 |
| P60981 | DSTN | Destrin | -0.28892231 | 0.512651 |
| P60983 | GMFB | Glia maturation factor beta | 0.077857892 | 0.292278 |
| P61006 | RAB8A | Ras-related protein Rab-8A | -0.976972659 | -0.24809 |
| P61009 | SPCS3 | Signal peptidase complex subunit 3 | -1.353642861 | 0.215343 |
| P61011 | SRP54 | Signal recognition particle 54 kDa protein | -0.135509253 | -0.183531 |
| P61018 | RAB4B | Ras-related protein Rab-4B | -0.779371977 | 0.105845 |
| P61019 | RAB2A | Ras-related protein Rab-2A | -0.779753526 | 0.00297991 |
| P61020 | RAB5B | Ras-related protein Rab-5B | -0.784190496 | -0.0368767 |
| P61024 | CKS1B | Cyclin-dependent kinases regulatory subunit 1 | -0.713922421 | 0.168272 |
| P61026 | RAB10 | Ras-related protein Rab-10 | -0.750211318 | -0.217805 |
| P61077 | UBE2D3 | Ubiquitin-conjugating enzyme E2 D3 | 0.263222059 | 0.201366 |
| P61081 | UBE2M | NEDD8-conjugating enzyme Ubc12 | 0.191327254 | 0.0158583 |
| P61086 | UBE2K | Ubiquitin-conjugating enzyme E2 K | 0.050928593 | -0.0381069 |
| P61088 | UBE2N | Ubiquitin-conjugating enzyme E2 N | 0.036184311 | 0.275302 |
| P61106 | RAB14 | Ras-related protein Rab-14 | -0.57744503 | 0.185804 |
| P61158 | ACTR3 | Actin-related protein 3 | -0.307575703 | 0.0636387 |
| P61160 | ACTR2 | Actin-related protein 2 | -0.285923322 | 0.0135911 |
| P61163 | ACTR1A | Alpha-centractin | -0.076188723 | -0.176577 |
| P61201 | COPS2 | COP9 signalosome complex subunit 2 | 0.04995807 | -0.33631 |
| P61221 | ABCE1 | ATP-binding cassette sub-family E member 1 | -0.465650717 | 0.0477352 |
| P61224 | RAP1B | Ras-related protein Rap-1b | -0.535368919 | 0.127759 |
| P61225 | RAP2B | Ras-related protein Rap-2b | -0.6749355 | 0.0206655 |
| P61247 | RPS3A | 40S ribosomal protein S3a | -0.376473586 | -0.27317 |
| P61254 | RPL26 | 60S ribosomal protein L26 | -1.221061865 | -0.0269883 |
| P61266 | STX1B | Syntaxin-1B | -0.481188138 | -0.218821 |
| P61289 | PSME3 | Proteasome activator complex subunit 3 | 0.257780711 | 0.427883 |
| P61313 | RPL15 | 60S ribosomal protein L15 | -1.01731356 | 0.0700477 |
| P61326 | MAGOH | Protein mago nashi homolog | -0.475634416 | -0.178493 |
| P61353 | RPL27 | 60S ribosomal protein L27 | -1.290590604 | 0.10592 |
| P61457 | PCBD1 | Pterin-4-alpha-carbinolamine dehydratase | 0.078428745 | 0.0270281 |
| P61586 | RHOA | Transforming protein RhoA | -0.146685918 | 0.00355466 |
| P61599 | NAA20 | N-alpha-acetyltransferase 20 | -1.028182983 | 0.426119 |
| P61604 | HSPE1 | 10 kDa heat shock protein, mitochondrial | -0.331538995 | 0.405673 |
| P61626 | LYZ | Lysozyme C | -0.650571664 | 0.559075 |
| P61758 | VBP1 | Prefoldin subunit 3 | 0.1816535 | -0.0886408 |
| P61769 | B2M | Beta-2-microglobulin | -0.269358714 | 0.356302 |
| P61956 | SUMO2 | Small ubiquitin-related modifier 2 | 0.184898535 | 0.293 |
| P61962 | DCAF7 | DDB1- and CUL4-associated factor 7 | -0.269234578 | 0.0665998 |
| P61964 | WDR5 | WD repeat-containing protein 5 | -0.213711182 | 0.149359 |
| P61970 | NUTF2 | Nuclear transport factor 2 | 0.139582713 | 0.813246 |
| P61978-3 | HNRNPK | Isoform 3 of Heterogeneous nuclear ribonucleoprotein K | 0.029884179 | 0.11145 |
| P61981 | YWHAG | 14-3-3 protein gamma | 0.264207522 | 0.180005 |
| P62070 | RRAS2 | Ras-related protein R-Ras2 | -0.168671211 | -0.0996297 |
| P62081 | RPS7 | 40S ribosomal protein S7 | -0.422665914 | -0.451894 |
| P62136 | PPP1CA | Serine/threonine-protein phosphatase PP1-alpha catalytic subunit | -0.005508105 | 0.0517578 |
| P62140 | PPP1CB | Serine/threonine-protein phosphatase PP1-beta catalytic subunit | -0.176083724 | -0.192548 |

| | | | | |
|---|---|---|---|---|
| P62166 | NCS1 | Neuronal calcium sensor 1 | 0.151406844 | 0.373398 |
| P62191 | PSMC1 | 26S proteasome regulatory subunit 4 | -0.153305372 | -0.106259 |
| P62195 | PSMC5 | 26S proteasome regulatory subunit 8 | -0.192508698 | -0.0971247 |
| P62241 | RPS8 | 40S ribosomal protein S8 | -0.529802004 | -0.444002 |
| P62244 | RPS15A | 40S ribosomal protein S15a | -0.473537604 | -0.343443 |
| P62249 | RPS16 | 40S ribosomal protein S16 | -0.285501003 | -0.394756 |
| P62253 | UBE2G1 | Ubiquitin-conjugating enzyme E2 G1 | 0.138050556 | -0.0458943 |
| P62258 | YWHAE | 14-3-3 protein epsilon | 0.395136992 | 0.322606 |
| P62266 | RPS23 | 40S ribosomal protein S23 | -0.483721574 | -0.636368 |
| P62269 | RPS18 | 40S ribosomal protein S18 | -0.111842314 | -0.256758 |
| P62273 | RPS29 | 40S ribosomal protein S29 | -0.447058757 | -0.815535 |
| P62277 | RPS13 | 40S ribosomal protein S13 | -0.328499953 | -0.485244 |
| P62280 | RPS11 | 40S ribosomal protein S11 | -0.444929441 | -0.379622 |
| P62304 | SNRPE | Small nuclear ribonucleoprotein E | 0.089044333 | -0.797354 |
| P62306 | SNRPF | Small nuclear ribonucleoprotein F | -0.453566551 | -0.152951 |
| P62310 | LSM3 | U6 snRNA-associated Sm-like protein LSm3 | -0.312452555 | 0.887947 |
| P62312 | LSM6 | U6 snRNA-associated Sm-like protein LSm6 | -0.150978883 | -0.206479 |
| P62314 | SNRPD1 | Small nuclear ribonucleoprotein Sm D1 | -0.397212823 | -0.990475 |
| P62316 | SNRPD2 | Small nuclear ribonucleoprotein Sm D2 | -0.116353671 | -0.361268 |
| P62318 | SNRPD3 | Small nuclear ribonucleoprotein Sm D3 | 0.188952287 | -0.0290597 |
| P62328 | TMSB4X | Thymosin beta-4 | 0.074374755 | -0.397945 |
| P62330 | ARF6 | ADP-ribosylation factor 6 | -0.598520358 | -0.233162 |
| P62380 | TBPL1 | TATA box-binding protein-like 1 | -0.162377437 | 0.460373 |
| P62424 | RPL7A | 60S ribosomal protein L7a | -1.241320769 | 0.0288022 |
| P62487 | POLR2G | DNA-directed RNA polymerase II subunit RPB7 | -0.201826096 | -0.20941 |
| P62495 | ETF1 | Eukaryotic peptide chain release factor subunit 1 | -0.298828761 | -0.0559966 |
| P62633-4 | CNBP | Isoform 4 of CCHC-type zinc finger nucleic acid binding protein | -0.768738031 | 0.695599 |
| P62633-5 | CNBP | Isoform 5 of CCHC-type zinc finger nucleic acid binding protein | 0.409192721 | 0 |
| P62633 | CNBP | CCHC-type zinc finger nucleic acid binding protein | -0.077177842 | 0.454316 |
| P62699 | YPEL5 | Protein yippee-like 5 | -0.035049677 | -0.0994294 |
| P62701 | RPS4X | 40S ribosomal protein S4, X isoform | -0.279401143 | -0.405628 |
| P62714 | PPP2CB | Serine/threonine-protein phosphatase 2A catalytic subunit beta isoform | -0.541230838 | -0.452131 |
| P62736 | ACTA2 | Actin, aortic smooth muscle | 0.383142312 | 0.466059 |
| P62750 | RPL23A | 60S ribosomal protein L23a | -1.102577607 | 0.0847209 |
| P62753 | RPS6 | 40S ribosomal protein S6 | -0.183506807 | -0.378497 |
| P62760 | VSNL1 | Visinin-like protein 1 | -0.45414408 | 0.00167179 |
| P62805 | H4C1 | Histone H4 | -0.767324607 | -0.23303 |
| P62820 | RAB1A | Ras-related protein Rab-1A | -0.480373065 | -0.00567118 |
| P62829 | RPL23 | 60S ribosomal protein L23 | -0.789770047 | -0.0716441 |
| P62834 | RAP1A | Ras-related protein Rap-1A | -0.939650853 | 0.232043 |
| P62837 | UBE2D2 | Ubiquitin-conjugating enzyme E2 D2 | -0.171422005 | -0.315564 |
| P62851 | RPS25 | 40S ribosomal protein S25 | -0.379655043 | -0.40706 |
| P62854 | RPS26 | 40S ribosomal protein S26 | -0.688245455 | -0.590778 |
| P62857 | RPS28 | 40S ribosomal protein S28 | -0.335720857 | -0.862871 |
| P62873 | GNB1 | Guanine nucleotide-binding protein G(I)/G(S)/G(T) subunit beta-1 | -1.11224548 | 0.627138 |
| P62877 | RBX1 | E3 ubiquitin-protein ligase RBX1 | -0.503650427 | 0.203066 |
| P62879 | GNB2 | Guanine nucleotide-binding protein G(I)/G(S)/G(T) subunit beta-2 | -0.972024202 | 0.170554 |
| P62891 | RPL39 | 60S ribosomal protein L39 | -1.74060758 | -2.67008 |
| P62906 | RPL10A | 60S ribosomal protein L10a | -1.286343336 | 0.189821 |
| P62913 | RPL11 | 60S ribosomal protein L11 | -1.399523099 | -0.178787 |
| P62913-2 | RPL11 | Isoform 2 of 60S ribosomal protein L11 | -0.762255907 | 0.0729027 |
| P62917 | RPL8 | 60S ribosomal protein L8 | -1.271781365 | -0.0568495 |
| P62937 | PPIA | Peptidyl-prolyl cis-trans isomerase A | 0.044239998 | -0.0205523 |
| P62937-2 | PPIA | Isoform 2 of Peptidyl-prolyl cis-trans isomerase A | -0.53926007 | 0.53349 |
| P62942 | FKBP1A | Peptidyl-prolyl cis-trans isomerase FKBP1A | -0.043978691 | 0.312431 |
| P62979 | RPS27A | Ubiquitin-40S ribosomal protein S27a | -0.243236383 | 0.0519346 |
| P62993 | GRB2 | Growth factor receptor-bound protein 2 | 0.069819132 | 0.108034 |
| P62995 | TRA2B | Transformer-2 protein homolog beta | -0.108686765 | 0.0149139 |
| P63000 | RAC1 | Ras-related C3 botulinum toxin substrate 1 | -0.78991286 | 0.124367 |
| P63010 | AP2B1 | AP-2 complex subunit beta | -0.506301721 | -0.043684 |
| P63096 | GNAI1 | Guanine nucleotide-binding protein G(i) subunit alpha-1 | -3.132942597 | 0.320885 |
| P63104 | YWHAZ | 14-3-3 protein zeta/delta | 0.094966412 | 0.114059 |
| P63151 | PPP2R2A | Serine/threonine-protein phosphatase 2A 55 kDa regulatory subunit B alpha isoform | 0.136742274 | 0.00237846 |
| P63167 | DYNLL1 | Dynein light chain 1, cytoplasmic | -0.388570944 | 0.00324726 |
| P63172 | DYNLT1 | Dynein light chain Tctex-type 1 | -0.244054159 | 0.0182708 |
| P63173 | RPL38 | 60S ribosomal protein L38 | -0.48203826 | -0.255241 |
| P63220 | RPS21 | 40S ribosomal protein S21 | -0.11982139 | -0.100325 |
| P63244 | RACK1 | Receptor of activated protein C kinase 1 | -0.113505205 | -0.237068 |
| P63261 | ACTG1 | Actin, cytoplasmic 2 | 0.265034358 | -0.0759621 |

| | | | | |
|---|---|---|---|---|
| P63272 | SUPT4H1 | Transcription elongation factor SPT4 | -0.032068968 | 0.0624669 |
| P63313 | TMSB10 | Thymosin beta-10 | -0.377771934 | 0.0181894 |
| P67775 | PPP2CA | Serine/threonine-protein phosphatase 2A catalytic subunit alpha isoform | -0.286824544 | -0.0853659 |
| P67809 | YBX1 | Y-box-binding protein 1 | -0.431265434 | -0.315908 |
| P67936 | TPM4 | Tropomyosin alpha-4 chain | 0.273580551 | 0.00627391 |
| P68036 | UBE2L3 | Ubiquitin-conjugating enzyme E2 L3 | -0.073037148 | -0.0429236 |
| P68104 | EEF1A1 | Elongation factor 1-alpha 1 | -0.240095615 | 0.0379906 |
| P68363 | TUBA1B | Tubulin alpha-1B chain | -0.010866801 | -0.249697 |
| P68366 | TUBA4A | Tubulin alpha-4A chain | 0.177735329 | 0.103813 |
| P68371 | TUBB4B | Tubulin beta-4B chain | -0.036735853 | -0.0167166 |
| P68402 | PAFAH1B2 | Platelet-activating factor acetylhydrolase IB subunit alpha2 | -0.071748575 | 0.021678 |
| P68543 | UBXN2A | UBX domain-containing protein 2A | 0.907589118 | 0 |
| P68871 | HBB | Hemoglobin subunit beta | 0.418710232 | -0.467215 |
| P69905 | HBA1 | Hemoglobin subunit alpha | 0.46796147 | -0.565036 |
| P78316 | NOP14 | Nucleolar protein 14 | -0.798087915 | -0.442912 |
| P78318 | IGBP1 | Immunoglobulin-binding protein 1 | 0.341606379 | 0.204594 |
| P78345 | RPP38 | Ribonuclease P protein subunit p38 | -0.573141336 | -0.393605 |
| P78346 | RPP30 | Ribonuclease P protein subunit p30 | -0.161837419 | -0.262331 |
| P78356 | PIP4K2B | Phosphatidylinositol 5-phosphate 4-kinase type-2 beta | 0.018207709 | 0.447047 |
| P78357 | CNTNAP1 | Contactin-associated protein 1 | -1.000561396 | -0.354947 |
| P78362 | SRPK2 | SRSF protein kinase 2 | -0.366014163 | -0.273628 |
| P78371 | CCT2 | T-complex protein 1 subunit beta | -0.106190364 | -0.174617 |
| P78385 | KRT83 | Keratin, type II cuticular Hb3 | -0.210922321 | 2.4206 |
| P78386 | KRT85 | Keratin, type II cuticular Hb5 | 0.732705593 | -0.250179 |
| P78406 | RAE1 | mRNA export factor RAE1 | -0.329504808 | 0.300005 |
| P78417 | GSTO1 | Glutathione S-transferase omega-1 | 0.044960181 | -0.244411 |
| P78527 | PRKDC | DNA-dependent protein kinase catalytic subunit | -0.721355438 | 0.117221 |
| P78560 | CRADD | Death domain-containing protein CRADD | 0.540216366 | 0 |
| P80188 | LCN2 | Neutrophil gelatinase-associated lipocalin | -1.199677308 | 0.441778 |
| P80217 | IFI35 | Interferon-induced 35 kDa protein | 0.850145658 | 0.283507 |
| P80303 | NUCB2 | Nucleobindin-2 | -0.177835226 | 0.144952 |
| P80723 | BASP1 | Brain acid soluble protein 1 | -0.370604515 | 0.555123 |
| P81605 | DCD | Dermcidin | -0.128799518 | 0.185734 |
| P82094 | TMF1 | TATA element modulatory factor | 0.405680974 | 0.0224867 |
| P82673 | MRPS35 | 28S ribosomal protein S35, mitochondrial | -0.272934079 | -0.453513 |
| P82675 | MRPS5 | 28S ribosomal protein S5, mitochondrial | 0.63495032 | 0.437953 |
| P82909 | MRPS36 | Alpha-ketoglutarate dehydrogenase component 4 | -1.277192672 | 0.756076 |
| P82921 | MRPS21 | 28S ribosomal protein S21, mitochondrial | 0 | 3.26767 |
| P82932 | MRPS6 | 28S ribosomal protein S6, mitochondrial | -1.380365531 | -1.85125 |
| P82933 | MRPS9 | 28S ribosomal protein S9, mitochondrial | -1.139367421 | -0.0729955 |
| P82979 | SARNP | SAP domain-containing ribonucleoprotein | -0.139630397 | 0.238582 |
| P83111 | LACTB | Serine beta-lactamase-like protein LACTB, mitochondrial | -1.892684539 | 0.148318 |
| P83436 | COG7 | Conserved oligomeric Golgi complex subunit 7 | -0.760742108 | -1.40961 |
| P83876 | TXNL4A | Thioredoxin-like protein 4A | -0.963867029 | -0.396918 |
| P83916 | CBX1 | Chromobox protein homolog 1 | -0.221429904 | -0.0543509 |
| P84022 | SMAD3 | Mothers against decapentaplegic homolog 3 | 0.102838119 | -0.182322 |
| P84077 | ARF1 | ADP-ribosylation factor 1 | -0.1885554 | -0.271966 |
| P84085 | ARF5 | ADP-ribosylation factor 5 | -0.144199451 | -0.171449 |
| P84090 | ERH | Enhancer of rudimentary homolog | -0.211113532 | 0.499217 |
| P84095 | RHOG | Rho-related GTP-binding protein RhoG | -1.078174273 | 0.0407639 |
| P84101 | SERF2 | Small EDRK-rich factor 2 | -0.455302795 | -0.209227 |
| P84103 | SRSF3 | Serine/arginine-rich splicing factor 3 | -0.041986624 | -0.0273457 |
| P85037 | FOXK1 | Forkhead box protein K1 | -0.315363566 | -0.0924037 |
| P86790 | CCZ1B | Vacuolar fusion protein CCZ1 homolog B | -0.367142598 | -0.178041 |
| P98082 | DAB2 | Disabled homolog 2 | 0.371510029 | 0.218539 |
| P98170 | XIAP | E3 ubiquitin-protein ligase XIAP | 0.196034114 | 0.0154746 |
| P98175-5 | RBM10 | Isoform 5 of RNA-binding protein 10 | 0.051343362 | -0.0915257 |
| P98179 | RBM3 | RNA-binding protein 3 | -0.569462299 | 0.302288 |
| Q00013 | MPP1 | 55 kDa erythrocyte membrane protein | 0.058410168 | 0.104501 |
| Q00059 | TFAM | Transcription factor A, mitochondrial | -1.168099324 | 0.330403 |
| Q00169 | PITPNA | Phosphatidylinositol transfer protein alpha isoform | 0.06071504 | -0.212343 |
| Q00325-2 | SLC25A3 | Isoform B of Phosphate carrier protein, mitochondrial | -1.403301954 | 0.213276 |
| Q00403 | GTF2B | Transcription initiation factor IIB | 0.210519075 | -0.282731 |
| Q00534 | CDK6 | Cyclin-dependent kinase 6 | -0.037853241 | 0.0782916 |
| Q00535 | CDK5 | Cyclin-dependent-like kinase 5 | 0.011887709 | -0.316815 |
| Q00577 | PURA | Transcriptional activator protein Pur-alpha | -0.214806636 | -0.0208832 |
| Q00587 | CDC42EP1 | Cdc42 effector protein 1 | -1.160100381 | -3.52725 |
| Q00653 | NFKB2 | Nuclear factor NF-kappa-B p100 subunit | 0.115238587 | 0.0878703 |
| Q00653-4 | NFKB2 | Isoform 4 of Nuclear factor NF-kappa-B p100 subunit | -1.095327457 | -0.674189 |
| Q00688 | FKBP3 | Peptidyl-prolyl cis-trans isomerase FKBP3 | -0.382155259 | -0.332318 |

| | | | | |
|---|---|---|---|---|
| Q00765 | REEP5 | Receptor expression-enhancing protein 5 | -1.186914762 | 0.19049 |
| Q00796 | SORD | Sorbitol dehydrogenase | -0.666585763 | 0.187497 |
| Q00839 | HNRNPU | Heterogeneous nuclear ribonucleoprotein U | -0.274032275 | 0.0909017 |
| Q01081 | U2AF1 | Splicing factor U2AF 35 kDa subunit | -0.443156481 | 0.410026 |
| Q01082 | SPTBN1 | Spectrin beta chain, non-erythrocytic 1 | 0.077765465 | -0.0531801 |
| Q01085 | TIAL1 | Nucleolysin TIAR | -0.604477247 | -0.0630169 |
| Q01105 | SET | Protein SET | 0.1875693 | 0.074777 |
| Q01105-2 | SET | Isoform 2 of Protein SET | 0.585210959 | -0.646349 |
| Q01415 | GALK2 | N-acetylgalactosamine kinase | 0.199688991 | 0.17768 |
| Q01432 | AMPD3 | AMP deaminase 3 | -0.07403334 | 0.076258 |
| Q01459 | CTBS | Di-N-acetylchitobiase | -1.060363849 | -0.685009 |
| Q01469 | FABP5 | Fatty acid-binding protein 5 | -0.328173637 | 0.571375 |
| Q01518 | CAP1 | Adenylyl cyclase-associated protein 1 | 0.143973033 | -0.0170555 |
| Q01546 | KRT76 | Keratin, type II cytoskeletal 2 oral | -2.330908616 | -2.98566 |
| Q01581 | HMGCS1 | Hydroxymethylglutaryl-CoA synthase, cytoplasmic | 0.284385125 | -0.00946045 |
| Q01650 | SLC7A5 | Large neutral amino acids transporter small subunit 1 | -1.747970343 | -0.641378 |
| Q01658 | DR1 | Protein Dr1 | 0.152611017 | 0.572109 |
| Q01780 | EXOSC10 | Exosome component 10 | -0.621400356 | -0.0816911 |
| Q01804 | OTUD4 | OTU domain-containing protein 4 | -1.082251549 | -0.781338 |
| Q01813 | PFKP | ATP-dependent 6-phosphofructokinase, platelet type | -0.214705785 | -0.0771224 |
| Q01968 | OCRL | Inositol polyphosphate 5-phosphatase OCRL | -0.062871774 | -0.0586818 |
| Q01970 | PLCB3 | 1-phosphatidylinositol 4,5-bisphosphate phosphodiesterase beta-3 | -0.109122912 | -0.300755 |
| Q01995 | TAGLN | Transgelin | 0.218631268 | -0.0583324 |
| Q02388 | COL7A1 | Collagen alpha-1(VII) chain | -1.630763769 | 0.436662 |
| Q02413 | DSG1 | Desmoglein-1 | -0.219444513 | -0.113766 |
| Q02539 | H1-1 | Histone H1.1 | -2.24817427 | 1.21677 |
| Q02750 | MAP2K1 | Dual specificity mitogen-activated protein kinase kinase 1 | 0.133250872 | -0.0929063 |
| Q02790 | FKBP4 | Peptidyl-prolyl cis-trans isomerase FKBP4 | 0.027891795 | -0.0260773 |
| Q02809 | PLOD1 | Procollagen-lysine,2-oxoglutarate 5-dioxygenase 1 | -0.691627343 | 0.240843 |
| Q02818 | NUCB1 | Nucleobindin-1 | -0.1743776 | 0.21314 |
| Q02878 | RPL6 | 60S ribosomal protein L6 | -1.009261211 | 0.217564 |
| Q02880 | TOP2B | DNA topoisomerase 2-beta | -1.018747886 | 0.0836417 |
| Q02952 | AKAP12 | A-kinase anchor protein 12 | 0.070842107 | 0.0535202 |
| Q03013 | GSTM4 | Glutathione S-transferase Mu 4 | -0.131073554 | -0.713422 |
| Q03111 | MLLT1 | Protein ENL | -0.105891228 | 0.000543276 |
| Q03135 | CAV1 | Caveolin-1 | -1.60818553 | -0.165968 |
| Q03252 | LMNB2 | Lamin-B2 | -0.206455549 | 0.208238 |
| Q03405 | PLAUR | Urokinase plasminogen activator surface receptor | -1.231528441 | 0.828649 |
| Q03405-2 | PLAUR | Isoform 2 of Urokinase plasminogen activator surface receptor | -1.66676712 | 1.05324 |
| Q03701 | CEBPZ | CCAAT/enhancer-binding protein zeta | -1.653007428 | -0.590872 |
| Q04323 | UBXN1 | UBX domain-containing protein 1 | 0.159580072 | -0.0116933 |
| Q04446 | GBE1 | 1,4-alpha-glucan-branching enzyme | 0.021011194 | -0.0803064 |
| Q04637-7 | EIF4G1 | Isoform 7 of Eukaryotic translation initiation factor 4 gamma 1 | -0.074556112 | -0.00585524 |
| Q04695 | KRT17 | Keratin, type I cytoskeletal 17 | -0.449652195 | -0.197329 |
| Q04721 | NOTCH2 | Neurogenic locus notch homolog protein 2 | -0.152914604 | 0.845034 |
| Q04724 | TLE1 | Transducin-like enhancer protein 1 | 0.240804036 | 0.0411539 |
| Q04760 | GLO1 | Lactoylglutathione lyase | 0.140924295 | 0.288781 |
| Q04828 | AKR1C1 | Aldo-keto reductase family 1 member C1 | -0.104349613 | 0.10174 |
| Q04837 | SSBP1 | Single-stranded DNA-binding protein, mitochondrial | -0.886718909 | 0.17088 |
| Q04864 | REL | Proto-oncogene c-Rel | -0.457698584 | 0.135426 |
| Q04917 | YWHAH | 14-3-3 protein eta | 0.026486715 | -0.000781377 |
| Q04941 | PLP2 | Proteolipid protein 2 | -0.610096296 | 0.363779 |
| Q05048 | CSTF1 | Cleavage stimulation factor subunit 1 | -0.484527985 | 0.0486546 |
| Q05086 | UBE3A | Ubiquitin-protein ligase E3A | 0.083628496 | -0.0277306 |
| Q05209 | PTPN12 | Tyrosine-protein phosphatase non-receptor type 12 | 0.129122496 | 0.198142 |
| Q05397 | PTK2 | Focal adhesion kinase 1 | -0.02412955 | -0.0301698 |
| Q05469 | LIPE | Hormone-sensitive lipase | -0.804095904 | -0.201566 |
| Q05519 | SRSF11 | Serine/arginine-rich splicing factor 11 | 0.161942403 | -0.87052 |
| Q05655 | PRKCD | Protein kinase C delta type | -0.761554718 | 0.584647 |
| Q05682-4 | CALD1 | Isoform 4 of Caldesmon | 0.233282963 | 0.261283 |
| Q05682 | CALD1 | Caldesmon | -0.041142623 | 0.0963249 |
| Q05932 | FPGS | Folylpolyglutamate synthase, mitochondrial | -1.351098061 | -1.06319 |
| Q05D32 | CTDSPL2 | CTD small phosphatase-like protein 2 | 0 | -2.13749 |
| Q06124-1 | PTPN11 | Isoform 2 of Tyrosine-protein phosphatase non-receptor type 11 | 0.110412598 | -0.0378653 |
| Q06190 | PPP2R3A | Serine/threonine-protein phosphatase 2A regulatory subunit B'' subunit alpha | 0.106219451 | -0.0836579 |
| Q06203 | PPAT | Amidophosphoribosyltransferase | 0.149456024 | -0.0550162 |
| Q06210 | GFPT1 | Glutamine--fructose-6-phosphate aminotransferase [isomerizing] 1 | -0.218873819 | -0.0295563 |
| Q06265 | EXOSC9 | Exosome complex component RRP45 | -0.196302891 | 0.579613 |

| | | | | |
|---|---|---|---|---|
| Q06323 | PSME1 | Proteasome activator complex subunit 1 | 0.029624621 | -0.0127017 |
| Q06481 | APLP2 | Amyloid beta precursor like protein 2 | 0.031767607 | -0.071565 |
| Q06546 | GABPA | GA-binding protein alpha chain | -0.002664248 | 0.178229 |
| Q06587 | RING1 | E3 ubiquitin-protein ligase RING1 | -0.069945971 | 0.160573 |
| Q06609 | RAD51 | DNA repair protein RAD51 homolog 1 | -0.859829823 | -1.09309 |
| Q06830 | PRDX1 | Peroxiredoxin-1 | -0.1063447 | 0.279248 |
| Q07021 | C1QBP | Complement component 1 Q subcomponent-binding protein, mitochondrial | -0.654168447 | -0.0781571 |
| Q07065 | CKAP4 | Cytoskeleton-associated protein 4 | -1.159414291 | 0.33666 |
| Q07617 | SPAG1 | Sperm-associated antigen 1 | -0.017021894 | -1.49405 |
| Q07666 | KHDRBS1 | KH domain-containing, RNA-binding, signal transduction-associated protein 1 | -0.485737403 | 0.251352 |
| Q07812 | BAX | Apoptosis regulator BAX | -0.173128208 | 0.237523 |
| Q07820 | MCL1 | Induced myeloid leukemia cell differentiation protein Mcl-1 | -0.289397001 | -0.618979 |
| Q07864 | POLE | DNA polymerase epsilon catalytic subunit A | -0.615327756 | -0.391449 |
| Q07954 | LRP1 | Prolow-density lipoprotein receptor-related protein 1 | -0.346966346 | 0.149779 |
| Q07960 | ARHGAP1 | Rho GTPase-activating protein 1 | -0.284245809 | 0.197641 |
| Q08170 | SRSF4 | Serine/arginine-rich splicing factor 4 | 0.12821881 | -0.0418847 |
| Q08209-2 | PPP3CA | Isoform 2 of Serine/threonine-protein phosphatase 2B catalytic subunit alpha isoform | 0.106802146 | -0.0304782 |
| Q08211 | DHX9 | ATP-dependent RNA helicase A | -0.382824421 | 0.134229 |
| Q08257 | CRYZ | Quinone oxidoreductase | -0.662492275 | 0.323675 |
| Q08378 | GOLGA3 | Golgin subfamily A member 3 | -0.024621248 | 0.0158539 |
| Q08380 | LGALS3BP | Galectin-3-binding protein | 0.292760928 | 0.27494 |
| Q08431 | MFGE8 | Lactadherin | -1.208216667 | 0.0783647 |
| Q08554 | DSC1 | Desmocollin-1 | -0.369870663 | -0.0406205 |
| Q08623 | PUDP | Pseudouridine-5'-phosphatase | 0.011565129 | 0.0906671 |
| Q08752 | PPID | Peptidyl-prolyl cis-trans isomerase D | 0.06166331 | -0.159709 |
| Q08945 | SSRP1 | FACT complex subunit SSRP1 | -0.15679582 | -0.0693652 |
| Q08AD1 | CAMSAP2 | Calmodulin-regulated spectrin-associated protein 2 | -0.354068995 | -0.722906 |
| Q08AE8 | SPIRE1 | Protein spire homolog 1 | -0.612195571 | -3.47425 |
| Q08AM6 | VAC14 | Protein VAC14 homolog | -1.375369549 | 0.256227 |
| Q08J23 | NSUN2 | RNA cytosine C(5)-methyltransferase NSUN2 | -0.03734684 | 0.0468473 |
| Q09028-2 | RBBP4 | Isoform 2 of Histone-binding protein RBBP4 | 0.601332823 | 0.209768 |
| Q09028 | RBBP4 | Histone-binding protein RBBP4 | -0.014326572 | -0.117232 |
| Q09161 | NCBP1 | Nuclear cap-binding protein subunit 1 | -0.307867765 | -0.283912 |
| Q09666 | AHNAK | Neuroblast differentiation-associated protein AHNAK | -0.456775824 | 0.15507 |
| Q0JRZ9 | FCHO2 | F-BAR domain only protein 2 | -0.174475511 | -0.489872 |
| Q0VDF9 | HSPA14 | Heat shock 70 kDa protein 14 | 0.006818136 | 0.0555356 |
| Q0VDG4 | SCRN3 | Secernin-3 | -0.1552821 | -0.535485 |
| Q0VGL1 | LAMTOR4 | Ragulator complex protein LAMTOR4 | -0.1526951 | 0.105498 |
| Q0ZGT2 | NEXN | Nexilin | 0.269923687 | 0.0929982 |
| Q10469 | MGAT2 | Alpha-1,6-mannosyl-glycoprotein 2-beta-N-acetylglucosaminyltransferase | -2.122987111 | -2.39186 |
| Q10471 | GALNT2 | Polypeptide N-acetylgalactosaminyltransferase 2 | -0.922105074 | 0.290273 |
| Q10472 | GALNT1 | Polypeptide N-acetylgalactosaminyltransferase 1 | 0 | -2.45229 |
| Q10567-3 | AP1B1 | Isoform C of AP-1 complex subunit beta-1 | -0.278478305 | 0.611227 |
| Q10567 | AP1B1 | AP-1 complex subunit beta-1 | -0.132483403 | 0.145295 |
| Q10570 | CPSF1 | Cleavage and polyadenylation specificity factor subunit 1 | -0.643301964 | 0.460598 |
| Q10713 | PMPCA | Mitochondrial-processing peptidase subunit alpha | -0.19742918 | 0.00693099 |
| Q12765 | SCRN1 | Secernin-1 | 0.064424197 | 0.137119 |
| Q12768 | WASHC5 | WASH complex subunit 5 | 0.093120575 | -0.603384 |
| Q12769 | NUP160 | Nuclear pore complex protein Nup160 | -0.08211263 | 0.302523 |
| Q12788 | TBL3 | Transducin beta-like protein 3 | -0.917809486 | 0.548951 |
| Q12789 | GTF3C1 | General transcription factor 3C polypeptide 1 | 0.626940211 | 2.17956 |
| Q12791-6 | KCNMA1 | Isoform 6 of Calcium-activated potassium channel subunit alpha-1 | -1.45936052 | -0.0469983 |
| Q12792 | TWF1 | Twinfilin-1 | 0.178950151 | -0.238241 |
| Q12797 | ASPH | Aspartyl/asparaginyl beta-hydroxylase | -1.301478148 | 0.657803 |
| Q12800 | TFCP2 | Alpha-globin transcription factor CP2 | -0.405517101 | -0.887797 |
| Q12834 | CDC20 | Cell division cycle protein 20 homolog | -1.698883216 | -1.04313 |
| Q12841 | FSTL1 | Follistatin-related protein 1 | 0.066381852 | 0.403616 |
| Q12846 | STX4 | Syntaxin-4 | -1.327435811 | 0.0045414 |
| Q12851 | MAP4K2 | Mitogen-activated protein kinase kinase kinase kinase 2 | -0.827806075 | -1.12396 |
| Q12860 | CNTN1 | Contactin-1 | 0.652146737 | -1.10812 |
| Q12872 | SFSWAP | Splicing factor, suppressor of white-apricot homolog | 0.080616395 | 0.164003 |
| Q12874 | SF3A3 | Splicing factor 3A subunit 3 | -0.092936277 | -0.0826162 |
| Q12882 | DPYD | Dihydropyrimidine dehydrogenase [NADP(+)] | 0.864923636 | 0 |
| Q12888 | TP53BP1 | TP53-binding protein 1 | 0.010633866 | 0.189095 |
| Q12893 | TMEM115 | Transmembrane protein 115 | -0.265948852 | -2.03498 |
| Q12899 | TRIM26 | Tripartite motif-containing protein 26 | 0.279542446 | -0.721848 |
| Q12906-7 | ILF3 | Isoform 7 of Interleukin enhancer-binding factor 3 | -0.319350322 | 0.167875 |

| | | | | |
|---|---|---|---|---|
| Q12907 | LMAN2 | Vesicular integral-membrane protein VIP36 | -1.111162663 | 0.503476 |
| Q12931 | TRAP1 | Heat shock protein 75 kDa, mitochondrial | -0.089100917 | 0.0585601 |
| Q12933 | TRAF2 | TNF receptor-associated factor 2 | -0.10197711 | 0.0571518 |
| Q12965 | MYO1E | Unconventional myosin-Ie | -0.285876989 | -0.0262893 |
| Q12972 | PPP1R8 | Nuclear inhibitor of protein phosphatase 1 | -0.109075546 | -0.00396633 |
| Q12974 | PTP4A2 | Protein tyrosine phosphatase type IVA 2 | -0.753991683 | 0.581752 |
| Q12981 | BNIP1 | Vesicle transport protein SEC20 | -0.688424587 | 0.575994 |
| Q13011 | ECH1 | Delta(3,5)-Delta(2,4)-dienoyl-CoA isomerase, mitochondrial | -1.07965517 | 0.0512772 |
| Q13015 | MLLT11 | Protein AF1q | -0.911777258 | -0.0251028 |
| Q13033 | STRN3 | Striatin-3 | 0.035008272 | -0.0157429 |
| Q13033-2 | STRN3 | Isoform Alpha of Striatin-3 | 0.321304878 | -0.343767 |
| Q13042 | CDC16 | Cell division cycle protein 16 homolog | -0.782142162 | -1.8901 |
| Q13043 | STK4 | Serine/threonine-protein kinase 4 | -0.018586397 | 0.00110976 |
| Q13045 | FLII | Protein flightless-1 homolog | 0.093286991 | 0.0342223 |
| Q13045-2 | FLII | Isoform 2 of Protein flightless-1 homolog | -0.791675409 | -0.600121 |
| Q13049 | TRIM32 | E3 ubiquitin-protein ligase TRIM32 | 0.08468016 | 0.0207411 |
| Q13057 | COASY | Bifunctional coenzyme A synthase | 0.080337604 | 0.01708 |
| Q13085 | ACACA | Acetyl-CoA carboxylase 1 | -0.236647288 | -0.00266202 |
| Q13107 | USP4 | Ubiquitin carboxyl-terminal hydrolase 4 | -0.334094604 | 0.125618 |
| Q13111 | CHAF1A | Chromatin assembly factor 1 subunit A | 0.02074035 | 0.292663 |
| Q13112 | CHAF1B | Chromatin assembly factor 1 subunit B | -0.090241432 | 0.0177081 |
| Q13115 | DUSP4 | Dual specificity protein phosphatase 4 | 1.599694729 | 3.43837 |
| Q13123 | IK | Protein Red | -0.513174693 | 0.189741 |
| Q13126 | MTAP | S-methyl-5'-thioadenosine phosphorylase | 0.026115576 | -0.104139 |
| Q13131 | PRKAA1 | 5'-AMP-activated protein kinase catalytic subunit alpha-1 | -0.057158311 | -6.80E-05 |
| Q13137 | CALCOCO2 | Calcium-binding and coiled-coil domain-containing protein 2 | 0.16630276 | -0.00319831 |
| Q13151 | HNRNPA0 | Heterogeneous nuclear ribonucleoprotein A0 | -0.580292702 | 0.185079 |
| Q13153 | PAK1 | Serine/threonine-protein kinase PAK 1 | 0.077946345 | 0.0754569 |
| Q13155 | AIMP2 | Aminoacyl tRNA synthase complex-interacting multifunctional protein 2 | -0.472322464 | -0.144139 |
| Q13158 | FADD | FAS-associated death domain protein | 0.115377267 | 0.440617 |
| Q13162 | PRDX4 | Peroxiredoxin-4 | -0.107271989 | 0.101339 |
| Q13164 | MAPK7 | Mitogen-activated protein kinase 7 | -0.067575216 | -0.0204684 |
| Q13177 | PAK2 | Serine/threonine-protein kinase PAK 2 | 0.079231739 | -0.147233 |
| Q13185 | CBX3 | Chromobox protein homolog 3 | 0.034197966 | 0.0658487 |
| Q13188 | STK3 | Serine/threonine-protein kinase 3 | 0.415107489 | 0.151395 |
| Q13190 | STX5 | Syntaxin-5 | -0.557299296 | -0.328569 |
| Q13200 | PSMD2 | 26S proteasome non-ATPase regulatory subunit 2 | -0.047144254 | -0.0613098 |
| Q13206 | DDX10 | Probable ATP-dependent RNA helicase DDX10 | -1.383686701 | 0.769473 |
| Q13217 | DNAJC3 | DnaJ homolog subfamily C member 3 | -0.810321013 | 0.111679 |
| Q13228 | SELENBP1 | Methanethiol oxidase | 0.052805901 | -0.221023 |
| Q13232 | NME3 | Nucleoside diphosphate kinase 3 | -0.104869127 | -0.21894 |
| Q13242 | SRSF9 | Serine/arginine-rich splicing factor 9 | -0.009720008 | -0.0149822 |
| Q13243 | SRSF5 | Serine/arginine-rich splicing factor 5 | 0.037532171 | 0.127487 |
| Q13257 | MAD2L1 | Mitotic spindle assembly checkpoint protein MAD2A | 0.051678658 | -0.0657883 |
| Q13263 | TRIM28 | Transcription intermediary factor 1-beta | -0.015303135 | 0.0787551 |
| Q13283 | G3BP1 | Ras GTPase-activating protein-binding protein 1 | -0.301787059 | -0.247002 |
| Q13287 | NMI | N-myc-interactor | 0.598744869 | -0.53031 |
| Q13308 | PTK7 | Inactive tyrosine-protein kinase 7 | -0.534141541 | 0.304204 |
| Q13330-3 | MTA1 | Isoform 3 of Metastasis-associated protein MTA1 | -0.157426357 | 0.171456 |
| Q13363 | CTBP1 | C-terminal-binding protein 1 | 0.138526281 | 0.0364234 |
| Q13371 | PDCL | Phosducin-like protein | -0.553845882 | -0.409359 |
| Q13395 | TARBP1 | Probable methyltransferase TARBP1 | 0.164366086 | -0.055226 |
| Q13404 | UBE2V1 | Ubiquitin-conjugating enzyme E2 variant 1 | 0.125872612 | -0.0208289 |
| Q13405 | MRPL49 | 39S ribosomal protein L49, mitochondrial | -0.411658565 | 1.28266 |
| Q13409 | DYNC1I2 | Cytoplasmic dynein 1 intermediate chain 2 | -0.048181057 | 0.162758 |
| Q13416 | ORC2 | Origin recognition complex subunit 2 | -0.380383253 | 0.320168 |
| Q13423 | NNT | NAD(P) transhydrogenase, mitochondrial | -1.002729813 | -0.012675 |
| Q13426 | XRCC4 | DNA repair protein XRCC4 | -0.176239014 | -0.10395 |
| Q13428-3 | TCOF1 | Isoform 3 of Treacle protein | -0.672470093 | -1.07249 |
| Q13428 | TCOF1 | Treacle protein | -0.10487024 | 0.151534 |
| Q13432 | UNC119 | Protein unc-119 homolog A | 0.083925724 | -0.0771033 |
| Q13435 | SF3B2 | Splicing factor 3B subunit 2 | -0.275515874 | -0.141543 |
| Q13439 | GOLGA4 | Golgin subfamily A member 4 | 0.149507046 | 0.0388568 |
| Q13442 | PDAP1 | 28 kDa heat- and acid-stable phosphoprotein | -0.217753728 | 0.122742 |
| Q13445 | TMED1 | Transmembrane emp24 domain-containing protein 1 | -1.922643344 | -1.05651 |
| Q13451 | FKBP5 | Peptidyl-prolyl cis-trans isomerase FKBP5 | -0.001442909 | 0.0423708 |
| Q13454-2 | TUSC3 | Isoform 2 of Tumor suppressor candidate 3 | -1.581420422 | 0.351735 |
| Q13464 | ROCK1 | Rho-associated protein kinase 1 | 0.106749852 | -0.459574 |
| Q13488 | TCIRG1 | V-type proton ATPase 116 kDa subunit a 3 | 0 | 2.28704 |
| Q13492 | PICALM | Phosphatidylinositol-binding clathrin assembly protein | -0.533040444 | 0.126857 |

| | | | | |
|---|---|---|---|---|
| Q13501 | SQSTM1 | Sequestosome-1 | -0.097962856 | 0.24593 |
| Q13505 | MTX1 | Metaxin-1 | -0.985939821 | 0.0341444 |
| Q13509 | TUBB3 | Tubulin beta-3 chain | 0.043363889 | 0.117039 |
| Q13523 | PRPF4B | Serine/threonine-protein kinase PRP4 homolog | -0.304622491 | -1.71723 |
| Q13526 | PIN1 | Peptidyl-prolyl cis-trans isomerase NIMA-interacting 1 | -0.102395535 | -0.000874519 |
| Q13546 | RIPK1 | Receptor-interacting serine/threonine-protein kinase 1 | 0.118504604 | 0.191698 |
| Q13547 | HDAC1 | Histone deacetylase 1 | -0.341840823 | -0.137308 |
| Q13554 | CAMK2B | Calcium/calmodulin-dependent protein kinase type II subunit beta | 1.654160023 | -1.56434 |
| Q13561 | DCTN2 | Dynactin subunit 2 | 0.103738785 | -0.153267 |
| Q13564 | NAE1 | NEDD8-activating enzyme E1 regulatory subunit | 0.050810655 | -0.0938479 |
| Q13573 | SNW1 | SNW domain-containing protein 1 | -0.297786395 | 0.204553 |
| Q13595 | TRA2A | Transformer-2 protein homolog alpha | 0.014052709 | -0.206134 |
| Q13596 | SNX1 | Sorting nexin-1 | 0.219739278 | -0.0613238 |
| Q13601 | KRR1 | KRR1 small subunit processome component homolog | -2.58775115 | -0.502876 |
| Q13610 | PWP1 | Periodic tryptophan protein 1 homolog | -0.791063309 | 0.11254 |
| Q13614 | MTMR2 | Myotubularin-related protein 2 | -0.616501729 | -1.28548 |
| Q13615 | MTMR3 | Myotubularin-related protein 3 | -0.224593163 | -0.184606 |
| Q13616 | CUL1 | Cullin-1 | 0.085976442 | 0.0747353 |
| Q13618 | CUL3 | Cullin-3 | -0.056242387 | -0.0676966 |
| Q13619 | CUL4A | Cullin-4A | 0.029326042 | -0.136701 |
| Q13620-3 | CUL4B | Isoform 3 of Cullin-4B | 1.95373257 | 0.0493972 |
| Q13625 | TP53BP2 | Apoptosis-stimulating of p53 protein 2 | -0.247986476 | -0.100536 |
| Q13630 | GFUS | GDP-L-fucose synthase | 0.120961189 | -0.21358 |
| Q13636 | RAB31 | Ras-related protein Rab-31 | -0.896498998 | 0.00314649 |
| Q13637 | RAB32 | Ras-related protein Rab-32 | -0.322742939 | -0.0644957 |
| Q13641 | TPBG | Trophoblast glycoprotein | -1.045569261 | 0.377643 |
| Q13642-1 | FHL1 | Isoform 1 of Four and a half LIM domains protein 1 | -0.375159184 | 0.359755 |
| Q13643 | FHL3 | Four and a half LIM domains protein 3 | 0.185898463 | 0.0093476 |
| Q13671 | RIN1 | Ras and Rab interactor 1 | -0.133061091 | 0.0936902 |
| Q13724 | MOGS | Mannosyl-oligosaccharide glucosidase | -1.131122986 | -0.285049 |
| Q13740 | ALCAM | CD166 antigen | -0.40658776 | 0.423966 |
| Q13753 | LAMC2 | Laminin subunit gamma-2 | -1.147654136 | 0.415301 |
| Q13813 | SPTAN1 | Spectrin alpha chain, non-erythrocytic 1 | 0.168074767 | 0.0197938 |
| Q13813-3 | SPTAN1 | Isoform 3 of Spectrin alpha chain, non-erythrocytic 1 | 0.203789155 | -1.29641 |
| Q13823 | GNL2 | Nucleolar GTP-binding protein 2 | -0.170047283 | 0.152752 |
| Q13835 | PKP1 | Plakophilin-1 | -0.356739442 | 0.0914847 |
| Q13838 | DDX39B | Spliceosome RNA helicase DDX39B | -0.19351538 | 0.0189362 |
| Q13867 | BLMH | Bleomycin hydrolase | -0.213848432 | 0.462289 |
| Q13868 | EXOSC2 | Exosome complex component RRP4 | -0.167454004 | -0.54136 |
| Q13885 | TUBB2A | Tubulin beta-2A chain | 0.032699267 | -0.527017 |
| Q13888 | GTF2H2 | General transcription factor IIH subunit 2 | -0.929213206 | -0.114535 |
| Q13895 | BYSL | Bystin | -0.49146533 | -0.324244 |
| Q13907 | IDI1 | Isopentenyl-diphosphate Delta-isomerase 1 | 0.072434982 | 0.0166324 |
| Q13951 | CBFB | Core-binding factor subunit beta | 0.118691762 | 0.132041 |
| Q13951-2 | CBFB | Isoform 2 of Core-binding factor subunit beta | -0.017955065 | 0.180741 |
| Q14008 | CKAP5 | Cytoskeleton-associated protein 5 | -0.694305658 | -0.0687256 |
| Q14011 | CIRBP | Cold-inducible RNA-binding protein | -0.495449622 | 0.102876 |
| Q14012 | CAMK1 | Calcium/calmodulin-dependent protein kinase type 1 | 0.040494283 | 0.0663176 |
| Q14019 | COTL1 | Coactosin-like protein | 0.144352595 | -0.0594406 |
| Q14103 | HNRNPD | Heterogeneous nuclear ribonucleoprotein D0 | -0.46419239 | 0.146977 |
| Q14116 | IL18 | Interleukin-18 | 0.21741724 | 1.05361 |
| Q14126 | DSG2 | Desmoglein-2 | -1.173768997 | -0.135085 |
| Q14137 | BOP1 | Ribosome biogenesis protein BOP1 | -0.049758673 | 0.23697 |
| Q14139 | UBE4A | Ubiquitin conjugation factor E4 A | -0.26194946 | -0.223766 |
| Q14145 | KEAP1 | Kelch-like ECH-associated protein 1 | -0.065461 | 0.00707245 |
| Q14149 | MORC3 | MORC family CW-type zinc finger protein 3 | -1.146657864 | 0 |
| Q14151 | SAFB2 | Scaffold attachment factor B2 | -0.024858554 | 0.213016 |
| Q14152 | EIF3A | Eukaryotic translation initiation factor 3 subunit A | -0.093567848 | -0.353922 |
| Q14157-1 | UBAP2L | Isoform 2 of Ubiquitin-associated protein 2-like | -1.649636348 | -1.24944 |
| Q14157 | UBAP2L | Ubiquitin-associated protein 2-like | -0.11835138 | -0.00967598 |
| Q14161 | GIT2 | ARF GTPase-activating protein GIT2 | -0.87750721 | 0.244877 |
| Q14165 | MLEC | Malectin | -1.21780324 | 0.254691 |
| Q14166 | TTLL12 | Tubulin--tyrosine ligase-like protein 12 | 0.102326552 | -0.0646248 |
| Q14181 | POLA2 | DNA polymerase alpha subunit B | 0.029198726 | 0.206581 |
| Q14192 | FHL2 | Four and a half LIM domains protein 2 | -0.121592045 | 0.276534 |
| Q14195-2 | DPYSL3 | Isoform LCRMP-4 of Dihydropyrimidinase-related protein 3 | 0.038079262 | -0.0204201 |
| Q14210 | LY6D | Lymphocyte antigen 6D | -2.06486702 | -0.237652 |
| Q14232 | EIF2B1 | Translation initiation factor eIF-2B subunit alpha | -0.085825602 | -0.353772 |
| Q14240 | EIF4A2 | Eukaryotic initiation factor 4A-II | -0.056821982 | -0.536291 |
| Q14247 | CTTN | Src substrate cortactin | 0.182351748 | 0.0944424 |
| Q14247-2 | CTTN | Isoform 2 of Src substrate cortactin | -0.637399832 | 0.213848 |

| | | | | |
|---|---|---|---|---|
| Q14254 | FLOT2 | Flotillin-2 | -0.776530981 | 0.134144 |
| Q14257 | RCN2 | Reticulocalbin-2 | -0.192897002 | 1.15315 |
| Q14258 | TRIM25 | E3 ubiquitin/ISG15 ligase TRIM25 | 0.255069097 | -0.0514921 |
| Q14315 | FLNC | Filamin-C | 0.158062935 | -0.00389417 |
| Q14318-2 | FKBP8 | Isoform 2 of Peptidyl-prolyl cis-trans isomerase FKBP8 | -0.58185037 | 0.660966 |
| Q14320 | FAM50A | Protein FAM50A | -0.331834714 | -0.112536 |
| Q14331 | FRG1 | Protein FRG1 | -0.357745012 | 0.117439 |
| Q14332 | FZD2 | Frizzled-2 | -1.68363142 | 1.92348 |
| Q14344 | GNA13 | Guanine nucleotide-binding protein subunit alpha-13 | -0.361061414 | -1.42794 |
| Q14353 | GAMT | Guanidinoacetate N-methyltransferase | -0.372096697 | 0.692717 |
| Q14376 | GALE | UDP-glucose 4-epimerase | 0.021633705 | 0.0304708 |
| Q14444 | CAPRIN1 | Caprin-1 | -0.176416556 | -0.0690943 |
| Q14498 | RBM39 | RNA-binding protein 39 | -0.29376475 | -0.184027 |
| Q14525 | KRT33B | Keratin, type I cuticular Ha3-II | 0.322146058 | 1.90706 |
| Q14534 | SQLE | Squalene monooxygenase | -0.944648186 | -0.0260207 |
| Q14554 | PDIA5 | Protein disulfide-isomerase A5 | -0.919404348 | 0.0214507 |
| Q14558 | PRPSAP1 | Phosphoribosyl pyrophosphate synthase-associated protein 1 | -0.286101103 | -0.677009 |
| Q14566 | MCM6 | DNA replication licensing factor MCM6 | 0.12789313 | -0.117561 |
| Q14573 | ITPR3 | Inositol 1,4,5-trisphosphate receptor type 3 | -1.226124287 | -0.481399 |
| Q14574 | DSC3 | Desmocollin-3 | -0.321844101 | -0.0519212 |
| Q14596 | NBR1 | Next to BRCA1 gene 1 protein | -1.397680124 | 0.220373 |
| Q14644 | RASA3 | Ras GTPase-activating protein 3 | -1.379495144 | -2.51313 |
| Q14651 | PLS1 | Plastin-1 | 0.045309146 | -0.101696 |
| Q14653 | IRF3 | Interferon regulatory factor 3 | -0.154201428 | 0.0472504 |
| Q14671-2 | PUM1 | Isoform 2 of Pumilio homolog 1 | -0.208364805 | 0.0144278 |
| Q14676 | MDC1 | Mediator of DNA damage checkpoint protein 1 | 0.039737225 | 0.411771 |
| Q14677-3 | CLINT1 | Isoform 3 of Clathrin interactor 1 | 0.058039824 | -0.109969 |
| Q14684 | RRP1B | Ribosomal RNA processing protein 1 homolog B | -0.976841927 | 0.167574 |
| Q14691 | GINS1 | DNA replication complex GINS protein PSF1 | -0.115788539 | -0.0555614 |
| Q14692 | BMS1 | Ribosome biogenesis protein BMS1 homolog | -0.767361879 | -0.102687 |
| Q14693 | LPIN1 | Phosphatidate phosphatase LPIN1 | 0.030870756 | 0.502902 |
| Q14694 | USP10 | Ubiquitin carboxyl-terminal hydrolase 10 | -0.249666532 | -0.0423514 |
| Q14696 | MESD | LRP chaperone MESD | -0.15978686 | 0.0706555 |
| Q14697 | GANAB | Neutral alpha-glucosidase AB | -0.11570549 | 0.0869395 |
| Q14697-2 | GANAB | Isoform 2 of Neutral alpha-glucosidase AB | -0.33259503 | -0.427691 |
| Q14699 | RFTN1 | Raftlin | -1.128813187 | 1.11768 |
| Q14764 | MVP | Major vault protein | 0.224358241 | 0.157204 |
| Q14790-9 | CASP8 | Isoform 9 of Caspase-8 | -0.239227136 | 0.390122 |
| Q147X3 | NAA30 | N-alpha-acetyltransferase 30 | 0.105898539 | 0.00778357 |
| Q14847 | LASP1 | LIM and SH3 domain protein 1 | 0.127026081 | 0.178729 |
| Q14934 | NFATC4 | Nuclear factor of activated T-cells, cytoplasmic 4 | 0.174612164 | 3.41749 |
| Q14966 | ZNF638 | Zinc finger protein 638 | -0.454043547 | 0.217841 |
| Q14974 | KPNB1 | Importin subunit beta-1 | -0.048947652 | -0.216813 |
| Q14980 | NUMA1 | Nuclear mitotic apparatus protein 1 | -0.283874512 | 0.109829 |
| Q14997 | PSME4 | Proteasome activator complex subunit 4 | -0.51551652 | -0.428756 |
| Q14C86-2 | GAPVD1 | Isoform 2 of GTPase-activating protein and VPS9 domain-containing protein 1 | -0.05856212 | -0.0517658 |
| Q14CN4-2 | KRT72 | Isoform 2 of Keratin, type II cytoskeletal 72 | -0.97576499 | -1.18646 |
| Q14CX7 | NAA25 | N-alpha-acetyltransferase 25, NatB auxiliary subunit | -0.384859562 | 0.126095 |
| Q15003 | NCAPH | Condensin complex subunit 2 | -0.726912816 | 0.23967 |
| Q15006 | EMC2 | ER membrane protein complex subunit 2 | -1.118104537 | 0.510396 |
| Q15007 | WTAP | Pre-mRNA-splicing regulator WTAP | 0.289755265 | 0.237171 |
| Q15008 | PSMD6 | 26S proteasome non-ATPase regulatory subunit 6 | -0.170037111 | -0.172053 |
| Q15013 | MAD2L1BP | MAD2L1-binding protein | -0.490816434 | -0.104246 |
| Q15014 | MORF4L2 | Mortality factor 4-like protein 2 | -0.550609191 | -0.713635 |
| Q15018 | ABRAXAS2 | BRISC complex subunit Abraxas 2 | 0.114517689 | 0.208456 |
| Q15019 | SEPTIN2 | Septin-2 | -0.370381276 | 0.0343339 |
| Q15021 | NCAPD2 | Condensin complex subunit 1 | -0.676788886 | -0.33147 |
| Q15024 | EXOSC7 | Exosome complex component RRP42 | -0.218416214 | -0.095396 |
| Q15029 | EFTUD2 | 116 kDa U5 small nuclear ribonucleoprotein component | -0.120242278 | 0.0849247 |
| Q15031 | LARS2 | Leucine--tRNA ligase, mitochondrial | -2.248195569 | -0.83219 |
| Q15036 | SNX17 | Sorting nexin-17 | -0.048924367 | -0.0327772 |
| Q15042 | RAB3GAP1 | Rab3 GTPase-activating protein catalytic subunit | 0.141067505 | -0.0862077 |
| Q15046 | KARS1 | Lysine--tRNA ligase | -0.156922102 | -0.0710039 |
| Q15050 | RRS1 | Ribosome biogenesis regulatory protein homolog | -1.063538154 | 0.760033 |
| Q15054 | POLD3 | DNA polymerase delta subunit 3 | -0.489107291 | 0.0222677 |
| Q15056 | EIF4H | Eukaryotic translation initiation factor 4H | -0.062832197 | -0.147648 |
| Q15056-2 | EIF4H | Isoform Short of Eukaryotic translation initiation factor 4H | 0.02120169 | 0.480251 |
| Q15057 | ACAP2 | Arf-GAP with coiled-coil, ANK repeat and PH domain-containing protein 2 | -0.052247047 | -0.284157 |
| Q15059 | BRD3 | Bromodomain-containing protein 3 | 0.088238001 | 0.133289 |

| | | | | |
|---|---|---|---|---|
| Q15061 | WDR43 | WD repeat-containing protein 43 | -1.79642566 | -0.0473261 |
| Q15067 | ACOX1 | Peroxisomal acyl-coenzyme A oxidase 1 | -1.598845402 | 0.906018 |
| Q15075 | EEA1 | Early endosome antigen 1 | 0.491265297 | -0.0487219 |
| Q15084 | PDIA6 | Protein disulfide-isomerase A6 | 0.075774034 | -0.280139 |
| Q15102 | PAFAH1B3 | Platelet-activating factor acetylhydrolase IB subunit alpha1 | 0.119801203 | 0.189249 |
| Q15118 | PDK1 | [Pyruvate dehydrogenase (acetyl-transferring)] kinase isozyme 1, mitochondrial | -0.490221739 | -4.58676 |
| Q15121 | PEA15 | Astrocytic phosphoprotein PEA-15 | -0.075497627 | 0.695556 |
| Q15125 | EBP | 3-beta-hydroxysteroid-Delta(8),Delta(7)-isomerase | -2.348662853 | -0.438213 |
| Q15126 | PMVK | Phosphomevalonate kinase | 0.145177126 | -0.00311438 |
| Q15149 | PLEC | Plectin | 0.092847665 | -0.0344289 |
| Q15149-3 | PLEC | Isoform 3 of Plectin | 0.127014081 | -0.225452 |
| Q15149-4 | PLEC | Isoform 4 of Plectin | -0.0204041 | -0.744743 |
| Q15172 | PPP2R5A | Serine/threonine-protein phosphatase 2A 56 kDa regulatory subunit alpha isoform | 0.305368662 | -1.84437 |
| Q15173 | PPP2R5B | Serine/threonine-protein phosphatase 2A 56 kDa regulatory subunit beta isoform | -0.022897641 | -0.109314 |
| Q15181 | PPA1 | Inorganic pyrophosphatase | 0.062546571 | -0.265098 |
| Q15185 | PTGES3 | Prostaglandin E synthase 3 | -0.049723466 | -0.0636635 |
| Q15208 | STK38 | Serine/threonine-protein kinase 38 | -0.057824453 | 0.242125 |
| Q15257 | PTPA | Serine/threonine-protein phosphatase 2A activator | 0.112002055 | -0.0972137 |
| Q15276 | RABEP1 | Rab GTPase-binding effector protein 1 | 0.081413746 | -0.101342 |
| Q15286 | RAB35 | Ras-related protein Rab-35 | -1.360610723 | 0.235871 |
| Q15291 | RBBP5 | Retinoblastoma-binding protein 5 | -0.370343288 | 0.121066 |
| Q15293 | RCN1 | Reticulocalbin-1 | -0.048367977 | 0.333652 |
| Q15311 | RALBP1 | RalA-binding protein 1 | -0.792299112 | 0.111954 |
| Q15323 | KRT31 | Keratin, type I cuticular Ha1 | -1.883154988 | 1.00215 |
| Q15334 | LLGL1 | Lethal(2) giant larvae protein homolog 1 | -0.283981244 | 0.153596 |
| Q15365 | PCBP1 | Poly(rC)-binding protein 1 | -0.313594977 | -0.0976512 |
| Q15366 | PCBP2 | Poly(rC)-binding protein 2 | -0.172971567 | -0.219786 |
| Q15382 | RHEB | GTP-binding protein Rheb | 0.142413616 | -0.0226237 |
| Q15386 | UBE3C | Ubiquitin-protein ligase E3C | -0.060698112 | -0.140915 |
| Q15388 | TOMM20 | Mitochondrial import receptor subunit TOM20 homolog | -1.192817847 | 0.23445 |
| Q15393 | SF3B3 | Splicing factor 3B subunit 3 | -0.174083233 | -0.126325 |
| Q15397 | PUM3 | Pumilio homolog 3 | -1.03438131 | 0.499361 |
| Q15398-3 | DLGAP5 | Isoform 3 of Disks large-associated protein 5 | -0.182534059 | 0.149927 |
| Q15404 | RSU1 | Ras suppressor protein 1 | 0.046861092 | 0.0304327 |
| Q15417 | CNN3 | Calponin-3 | 0.091045856 | 0.0781345 |
| Q15417-2 | CNN3 | Isoform 2 of Calponin-3 | -0.698786259 | 0.992802 |
| Q15424 | SAFB | Scaffold attachment factor B1 | -0.574128628 | 0.433085 |
| Q15427 | SF3B4 | Splicing factor 3B subunit 4 | -0.229654074 | -0.12936 |
| Q15428 | SF3A2 | Splicing factor 3A subunit 2 | -0.249832471 | 0.28826 |
| Q15435 | PPP1R7 | Protein phosphatase 1 regulatory subunit 7 | 0.315911134 | -0.162891 |
| Q15438 | CYTH1 | Cytohesin-1 | -0.269813458 | -0.428678 |
| Q15459 | SF3A1 | Splicing factor 3A subunit 1 | -0.087246021 | 0.192087 |
| Q15477 | SKIV2L | Helicase SKI2W | -0.721100012 | -1.93958 |
| Q15527 | SURF2 | Surfeit locus protein 2 | -0.558465004 | 0.419054 |
| Q15545 | TAF7 | Transcription initiation factor TFIID subunit 7 | 0.466206868 | -0.194203 |
| Q15555 | MAPRE2 | Microtubule-associated protein RP/EB family member 2 | -0.01657335 | 0.148067 |
| Q15582 | TGFBI | Transforming growth factor-beta-induced protein ig-h3 | -0.884000937 | 0.255578 |
| Q15599 | SLC9A3R2 | Na(+)/H(+) exchange regulatory cofactor NHE-RF2 | -0.52250576 | -0.0583169 |
| Q15628 | TRADD | Tumor necrosis factor receptor type 1-associated DEATH domain protein | -0.27596426 | -0.0309324 |
| Q15637 | SF1 | Splicing factor 1 | -0.425871134 | 0.00799751 |
| Q15642-2 | TRIP10 | Isoform 2 of Cdc42-interacting protein 4 | 0.173913558 | -0.163736 |
| Q15643 | TRIP11 | Thyroid receptor-interacting protein 11 | -0.228799979 | 0.040377 |
| Q15645 | TRIP13 | Pachytene checkpoint protein 2 homolog | 0.111656984 | -0.188392 |
| Q15648 | MED1 | Mediator of RNA polymerase II transcription subunit 1 | -1.063414256 | 1.61812 |
| Q15653 | NFKBIB | NF-kappa-B inhibitor beta | -0.491357644 | 0.101511 |
| Q15654 | TRIP6 | Thyroid receptor-interacting protein 6 | -0.206779083 | 0.0451902 |
| Q15678 | PTPN14 | Tyrosine-protein phosphatase non-receptor type 14 | -1.76744318 | -1.62885 |
| Q15691 | MAPRE1 | Microtubule-associated protein RP/EB family member 1 | 0.266880989 | -0.0903123 |
| Q15717 | ELAVL1 | ELAV-like protein 1 | -0.473524253 | 0.123991 |
| Q15738 | NSDHL | Sterol-4-alpha-carboxylate 3-dehydrogenase, decarboxylating | -0.973084132 | -0.125773 |
| Q15742 | NAB2 | NGFI-A-binding protein 2 | 0.064700445 | -0.0355943 |
| Q15746 | MYLK | Myosin light chain kinase, smooth muscle | -0.095056534 | -0.0171639 |
| Q15750 | TAB1 | TGF-beta-activated kinase 1 and MAP3K7-binding protein 1 | 0.01293095 | -0.104877 |
| Q15758 | SLC1A5 | Neutral amino acid transporter B(0) | -1.185212851 | -0.647422 |
| Q15773 | MLF2 | Myeloid leukemia factor 2 | -0.474245707 | -0.514424 |
| Q15785 | TOMM34 | Mitochondrial import receptor subunit TOM34 | -0.691341559 | 0.11771 |
| Q15796 | SMAD2 | Mothers against decapentaplegic homolog 2 | -0.12337176 | 0.164466 |

| | | | | |
|---|---|---|---|---|
| Q15797 | SMAD1 | Mothers against decapentaplegic homolog 1 | 0.056530952 | 0 |
| Q15800 | MSMO1 | Methylsterol monooxygenase 1 | -1.743164539 | -0.0651007 |
| Q15813 | TBCE | Tubulin-specific chaperone E | 0.105201403 | -0.0663112 |
| Q15814 | TBCC | Tubulin-specific chaperone C | 0.037634452 | 0.00472991 |
| Q15819 | UBE2V2 | Ubiquitin-conjugating enzyme E2 variant 2 | 0.16275231 | 0.128327 |
| Q15828 | CST6 | Cystatin-M | -0.604564428 | -0.322873 |
| Q15833 | STXBP2 | Syntaxin-binding protein 2 | 0.283277114 | -0.165765 |
| Q15836 | VAMP3 | Vesicle-associated membrane protein 3 | -1.592588743 | -0.00314681 |
| Q15843 | NEDD8 | NEDD8 | -0.036262353 | 0.00186666 |
| Q15904 | ATP6AP1 | V-type proton ATPase subunit S1 | -1.134318113 | -2.46779 |
| Q15907 | RAB11B | Ras-related protein Rab-11B | -0.252396266 | -0.027455 |
| Q15910 | EZH2 | Histone-lysine N-methyltransferase EZH2 | 0.433662415 | 0 |
| Q15942 | ZYX | Zyxin | 0.239146233 | 0.0189807 |
| Q16143 | SNCB | Beta-synuclein | -0.014039755 | 0.129364 |
| Q16186 | ADRM1 | Proteasomal ubiquitin receptor ADRM1 | -0.027814229 | -0.0364958 |
| Q16204 | CCDC6 | Coiled-coil domain-containing protein 6 | 0.158558528 | 0.0845979 |
| Q16222 | UAP1 | UDP-N-acetylhexosamine pyrophosphorylase | 0.150416056 | 0.0883993 |
| Q16254 | E2F4 | Transcription factor E2F4 | 0.742607196 | 0 |
| Q16352 | INA | Alpha-internexin | 0.197900772 | 0.0229966 |
| Q16401 | PSMD5 | 26S proteasome non-ATPase regulatory subunit 5 | 0.07096386 | -0.139969 |
| Q16512 | PKN1 | Serine/threonine-protein kinase N1 | 0.132114887 | -0.462018 |
| Q16513 | PKN2 | Serine/threonine-protein kinase N2 | -0.268704971 | -0.095575 |
| Q16531 | DDB1 | DNA damage-binding protein 1 | -0.065353711 | 0.0791006 |
| Q16537 | PPP2R5E | Serine/threonine-protein phosphatase 2A 56 kDa regulatory subunit epsilon isoform | 0.028676271 | -0.0684903 |
| Q16539 | MAPK14 | Mitogen-activated protein kinase 14 | 0.011055787 | -0.121876 |
| Q16539-2 | MAPK14 | Isoform CSBP1 of Mitogen-activated protein kinase 14 | -0.835153818 | 0.153534 |
| Q16540 | MRPL23 | 39S ribosomal protein L23, mitochondrial | -1.159203132 | -0.0602156 |
| Q16543 | CDC37 | Hsp90 co-chaperone Cdc37 | 0.086407344 | 0.0709559 |
| Q16563 | SYPL1 | Synaptophysin-like protein 1 | -2.011172533 | -0.14414 |
| Q16566 | CAMK4 | Calcium/calmodulin-dependent protein kinase type IV | -0.025025686 | 0.0455869 |
| Q16576 | RBBP7 | Histone-binding protein RBBP7 | 0.046832403 | -0.00451024 |
| Q16610 | ECM1 | Extracellular matrix protein 1 | -0.379046599 | 0.450167 |
| Q16643 | DBN1 | Drebrin | 0.056791464 | 0.0894915 |
| Q16656 | NRF1 | Nuclear respiratory factor 1 | -0.181175788 | 0.0737454 |
| Q16658 | FSCN1 | Fascin | 0.034700235 | -0.0210908 |
| Q16666 | IFI16 | Gamma-interferon-inducible protein 16 | -0.17991209 | 0.787192 |
| Q16698 | DECR1 | 2,4-dienoyl-CoA reductase [(3E)-enoyl-CoA-producing], mitochondrial | -1.443146865 | 0.650371 |
| Q16706 | MAN2A1 | Alpha-mannosidase 2 | -1.648346345 | 0.0389929 |
| Q16719 | KYNU | Kynureninase | -0.234465599 | 0.762066 |
| Q16739 | UGCG | Ceramide glucosyltransferase | -0.515360991 | 0.802548 |
| Q16762 | TST | Thiosulfate sulfurtransferase | -0.06090196 | 0.129744 |
| Q16763 | UBE2S | Ubiquitin-conjugating enzyme E2 S | -0.288616975 | 0.0882222 |
| Q16774 | GUK1 | Guanylate kinase | 0.113722324 | -0.134225 |
| Q16795 | NDUFA9 | NADH dehydrogenase [ubiquinone] 1 alpha subcomplex subunit 9, mitochondrial | -1.581447919 | -0.0142476 |
| Q16822 | PCK2 | Phosphoenolpyruvate carboxykinase [GTP], mitochondrial | -1.622740547 | -0.114064 |
| Q16831 | UPP1 | Uridine phosphorylase 1 | -0.407292604 | 0.167255 |
| Q16850 | CYP51A1 | Lanosterol 14-alpha demethylase | -1.478303432 | 0.330324 |
| Q16851 | UGP2 | UTP--glucose-1-phosphate uridylyltransferase | 0.005251249 | -0.125437 |
| Q16854 | DGUOK | Deoxyguanosine kinase, mitochondrial | 0.132484436 | 2.88434 |
| Q16864 | ATP6V1F | V-type proton ATPase subunit F | 0.404149691 | 0.874258 |
| Q16878 | CDO1 | Cysteine dioxygenase type 1 | -0.675146023 | -0.273817 |
| Q16891 | IMMT | MICOS complex subunit MIC60 | -1.242570798 | 0.500403 |
| Q1KMD3 | HNRNPUL2 | Heterogeneous nuclear ribonucleoprotein U-like protein 2 | -0.366461754 | -0.0364606 |
| Q24JP5 | TMEM132A | Transmembrane protein 132A | -0.080445369 | -1.26918 |
| Q29RF7 | PDS5A | Sister chromatid cohesion protein PDS5 homolog A | -0.952936967 | 0.204668 |
| Q2KHT3 | CLEC16A | Protein CLEC16A | -0.423806349 | -0.694271 |
| Q2NKX8 | ERCC6L | DNA excision repair protein ERCC-6-like | 0.009145657 | 0.00482051 |
| Q2NL82 | TSR1 | Pre-rRNA-processing protein TSR1 homolog | -0.423609893 | -0.542439 |
| Q2TAA2 | IAH1 | Isoamyl acetate-hydrolyzing esterase 1 homolog | -0.064745585 | 0.0506334 |
| Q2TAL8 | QRICH1 | Transcriptional regulator QRICH1 | 0.160520554 | 0.0555639 |
| Q2TAM9 | TUSC1 | Tumor suppressor candidate gene 1 protein | -0.354184786 | -1.49898 |
| Q2TAY7 | SMU1 | WD40 repeat-containing protein SMU1 | -0.565229416 | 0.585867 |
| Q2TAZ0 | ATG2A | Autophagy-related protein 2 homolog A | -0.853491704 | -0.142797 |
| Q32MZ4-3 | LRRFIP1 | Isoform 3 of Leucine-rich repeat flightless-interacting protein 1 | -0.040799856 | 0.11104 |
| Q32MZ4-4 | LRRFIP1 | Isoform 4 of Leucine-rich repeat flightless-interacting protein 1 | -0.092533271 | -0.200575 |
| Q32NB8 | PGS1 | CDP-diacylglycerol--glycerol-3-phosphate 3-phosphatidyltransferase, mitochondrial | -1.096627871 | -0.326099 |
| Q32P28 | P3H1 | Prolyl 3-hydroxylase 1 | 0.05345583 | -0.0438353 |

| | | | | |
|---|---|---|---|---|
| Q32P41 | TRMT5 | tRNA (guanine(37)-N1)-methyltransferase | -0.124853134 | 0.121148 |
| Q3B726 | POLR1F | DNA-directed RNA polymerase I subunit RPA43 | -0.873853683 | -0.563842 |
| Q3KQV9 | UAP1L1 | UDP-N-acetylhexosamine pyrophosphorylase-like protein 1 | 0.069824537 | -0.0358734 |
| Q3LIE5 | ADPRM | Manganese-dependent ADP-ribose/CDP-alcohol diphosphatase | -0.368887981 | 0.633143 |
| Q3LXA3 | TKFC | Triokinase/FMN cyclase | 0.197248459 | 0.118806 |
| Q3MHD2 | LSM12 | Protein LSM12 homolog | -0.46808521 | 0.173609 |
| Q3MII6 | TBC1D25 | TBC1 domain family member 25 | -0.136521975 | 0.160425 |
| Q3MIT2 | PUS10 | tRNA pseudouridine synthase Pus10 | 0.313481569 | -0.422855 |
| Q3SXM5 | HSDL1 | Inactive hydroxysteroid dehydrogenase-like protein 1 | -1.045158704 | -0.135844 |
| Q3SY84 | KRT71 | Keratin, type II cytoskeletal 71 | 0.250858148 | 0.131557 |
| Q3YEC7 | RABL6 | Rab-like protein 6 | -0.06102705 | -0.531462 |
| Q3ZAQ7 | VMA21 | Vacuolar ATPase assembly integral membrane protein VMA21 | -1.484798749 | 3.54702 |
| Q3ZCQ8 | TIMM50 | Mitochondrial import inner membrane translocase subunit TIM50 | -0.900505304 | 0.442418 |
| Q3ZCW2 | LGALSL | Galectin-related protein | -0.953843117 | 0.741276 |
| Q495W5 | FUT11 | Alpha-(1,3)-fucosyltransferase 11 | 0.233876864 | -0.614974 |
| Q49AR2 | C5orf22 | UPF0489 protein C5orf22 | -0.055976947 | -2.90163 |
| Q4G0F5 | VPS26B | Vacuolar protein sorting-associated protein 26B | 0.168367863 | -0.150468 |
| Q4G0J3 | LARP7 | La-related protein 7 | -0.068551302 | 0.0504649 |
| Q4G0N4 | NADK2 | NAD kinase 2, mitochondrial | -0.259429375 | -0.636488 |
| Q4G148 | GXYLT1 | Glucoside xylosyltransferase 1 | -0.003395637 | -0.542085 |
| Q4J6C6 | PREPL | Prolyl endopeptidase-like | -0.233215491 | 0.127577 |
| Q4KMP7 | TBC1D10B | TBC1 domain family member 10B | 0 | 3.52875 |
| Q4LE39 | ARID4B | AT-rich interactive domain-containing protein 4B | 0.030896664 | -0.990175 |
| Q4V328 | GRIPAP1 | GRIP1-associated protein 1 | 0.231850068 | 0.132993 |
| Q52LJ0 | FAM98B | Protein FAM98B | -0.503440857 | 0.0159044 |
| Q52LW3 | ARHGAP29 | Rho GTPase-activating protein 29 | -0.104062955 | 0.269722 |
| Q53EL6 | PDCD4 | Programmed cell death protein 4 | 0.077727715 | -0.000104904 |
| Q53EP0 | FNDC3B | Fibronectin type III domain-containing protein 3B | -1.846393694 | 0.580849 |
| Q53ET0 | CRTC2 | CREB-regulated transcription coactivator 2 | -0.151573181 | -1.45437 |
| Q53EU6 | GPAT3 | Glycerol-3-phosphate acyltransferase 3 | -0.918017228 | -0.411992 |
| Q53EZ4 | CEP55 | Centrosomal protein of 55 kDa | -1.005607684 | 0.414611 |
| Q53F19 | NCBP3 | Nuclear cap-binding protein subunit 3 | -0.038281361 | 0.10003 |
| Q53FA7 | TP53I3 | Quinone oxidoreductase PIG3 | -0.20714736 | 0.394918 |
| Q53FT3 | HIKESHI | Protein Hikeshi | -0.189139684 | -0.485379 |
| Q53G59 | KLHL12 | Kelch-like protein 12 | 0.544309696 | -1.38569 |
| Q53GA4 | PHLDA2 | Pleckstrin homology-like domain family A member 2 | -0.440346479 | 0.105179 |
| Q53GG5-2 | PDLIM3 | Isoform 2 of PDZ and LIM domain protein 3 | 0.213997523 | -0.377873 |
| Q53GQ0 | HSD17B12 | Very-long-chain 3-oxoacyl-CoA reductase | -1.708569288 | 0.225362 |
| Q53GS9 | USP39 | U4/U6.U5 tri-snRNP-associated protein 2 | -0.684596221 | -0.374309 |
| Q53GT1 | KLHL22 | Kelch-like protein 22 | -0.193178177 | -0.259743 |
| Q53H12 | AGK | Acylglycerol kinase, mitochondrial | -0.755691528 | 0.327723 |
| Q53H82 | LACTB2 | Endoribonuclease LACTB2 | 0.051107645 | 0.198222 |
| Q53HC5 | KLHL26 | Kelch-like protein 26 | 0.472827514 | -0.698609 |
| Q53HL2 | CDCA8 | Borealin | -0.786060572 | -0.617855 |
| Q53LP3 | SOWAHC | Ankyrin repeat domain-containing protein SOWAHC | -0.876635313 | -0.516294 |
| Q53RE8 | ANKRD39 | Ankyrin repeat domain-containing protein 39 | -0.030972878 | 1.97146 |
| Q53RT3 | ASPRV1 | Retroviral-like aspartic protease 1 | -0.624655763 | 2.34787 |
| Q53S33 | BOLA3 | BolA-like protein 3 | -2.340160728 | 0 |
| Q53T59 | HS1BP3 | HCLS1-binding protein 3 | 0.074830214 | 0.0695257 |
| Q562R1 | ACTBL2 | Beta-actin-like protein 2 | 0.01331981 | 0.762956 |
| Q56VL3 | OCIAD2 | OCIA domain-containing protein 2 | -1.752250751 | 0.709942 |
| Q58FF6 | HSP90AB4P | Putative heat shock protein HSP 90-beta 4 | -0.575772603 | -0.348989 |
| Q58WW2 | DCAF6 | DDB1- and CUL4-associated factor 6 | -0.205545743 | 0.677015 |
| Q5BJH7 | YIF1B | Protein YIF1B | -3.358186285 | -0.0398102 |
| Q5BKZ1 | ZNF326 | DBIRD complex subunit ZNF326 | 0.103501002 | 0.336869 |
| Q5D862 | FLG2 | Filaggrin-2 | -0.533198118 | -0.0845677 |
| Q5EBL4 | RILPL1 | RILP-like protein 1 | -0.042596102 | 0.251006 |
| Q5EBL8 | PDZD11 | PDZ domain-containing protein 11 | -0.57616361 | 0.698571 |
| Q5F1R6 | DNAJC21 | DnaJ homolog subfamily C member 21 | -0.24854819 | -0.422725 |
| Q5GLZ8 | HERC4 | Probable E3 ubiquitin-protein ligase HERC4 | -0.020828724 | -0.0984586 |
| Q5JS37 | NHLRC3 | NHL repeat-containing protein 3 | 0.171723445 | -0.279459 |
| Q5JS54 | PSMG4 | Proteasome assembly chaperone 4 | 0.362122814 | 0.731712 |
| Q5JSH3 | WDR44 | WD repeat-containing protein 44 | 0.190202316 | 0.0499983 |
| Q5JST6 | EFHC2 | EF-hand domain-containing family member C2 | -1.756855567 | -0.817046 |
| Q5JTD0 | TJAP1 | Tight junction-associated protein 1 | -0.203757524 | 0.0446879 |
| Q5JTH9 | RRP12 | RRP12-like protein | -0.867835522 | 0.0147308 |
| Q5JTJ3 | COA6 | Cytochrome c oxidase assembly factor 6 homolog | -1.569402774 | 1.04349 |
| Q5JTV1 | GMEB2 | Glucocorticoid modulatory element binding protein 2, isoform CRA_a | -0.361228466 | 0.228341 |
| Q5JTV8-3 | TOR1AIP1 | Isoform 3 of Torsin-1A-interacting protein 1 | -0.962018013 | 0.143109 |
| Q5JTZ9 | AARS2 | Alanine--tRNA ligase, mitochondrial | -0.453136921 | 0.377912 |

| | | | | |
|---|---|---|---|---|
| Q5JVF3 | PCID2 | PCI domain-containing protein 2 | -0.48159639 | 0.297634 |
| Q5JVZ5 | ELMO2 | Engulfment and cell motility protein 2 | -0.014048894 | 0.126462 |
| Q5JW54 | DSN1 | Kinetochore-associated protein DSN1 homolog | -0.278949976 | -0.0283216 |
| Q5JWT2 | MED23 | Mediator complex subunit 23 | -0.174196959 | -1.4602 |
| Q5JY65 | CRNKL1 | Crooked neck-like protein 1 | -0.146838109 | 0.0459855 |
| Q5JZH3 | SNX21 | Sorting nexin-21 | -1.233206828 | 3.37684 |
| Q5MIZ7 | PPP4R3B | Serine/threonine-protein phosphatase 4 regulatory subunit 3B | 0.029727697 | -0.222841 |
| Q5NDL2 | EOGT | EGF domain-specific O-linked N-acetylglucosamine transferase | -0.132428885 | -0.915053 |
| Q5QJ74 | TBCEL | Tubulin-specific chaperone cofactor E-like protein | 0.12904652 | -0.118675 |
| Q5QJE6 | DNTTIP2 | Deoxynucleotidyltransferase terminal-interacting protein 2 | -0.86232837 | 0.707301 |
| Q5QPA5 | MRPS18A | 39S ribosomal protein S18a, mitochondrial (Fragment) | -1.594758193 | 1.9572 |
| Q5QPM7 | PSMF1 | Proteasome inhibitor PI31 subunit | 0.164335092 | 0.308133 |
| Q5R372 | RABGAP1L | Rab GTPase-activating protein 1-like | -0.088809093 | -0.812783 |
| Q5R3I4 | TTC38 | Tetratricopeptide repeat protein 38 | 0.160889149 | -1.59051 |
| Q5RI15 | COX20 | Cytochrome c oxidase assembly protein COX20, mitochondrial | -1.826173306 | 0.287279 |
| Q5RKV6 | EXOSC6 | Exosome complex component MTR3 | -0.462983767 | 0.0776065 |
| Q5SRE5 | NUP188 | Nucleoporin NUP188 | -0.227936268 | 0.101804 |
| Q5SSJ5 | HP1BP3 | Heterochromatin protein 1-binding protein 3 | -0.389168262 | 0.0823669 |
| Q5STZ8 | ABCF1 | ATP-binding cassette sub-family F member 1 | -0.204280853 | -0.0891504 |
| Q5SW79 | CEP170 | Centrosomal protein of 170 kDa | -0.511569659 | 0.148764 |
| Q5SW79-2 | CEP170 | Isoform 2 of Centrosomal protein of 170 kDa | -0.945838451 | -1.68424 |
| Q5SW96 | LDLRAP1 | Low density lipoprotein receptor adapter protein 1 | 0.343464295 | 3.62153 |
| Q5SWA1 | PPP1R15B | Protein phosphatase 1 regulatory subunit 15B | 0.118544102 | -0.138095 |
| Q5SWX8 | ODR4 | Protein odr-4 homolog | -0.963528951 | 1.14161 |
| Q5SY16 | NOL9 | Polynucleotide 5'-hydroxyl-kinase NOL9 | -0.93287317 | 0.0418622 |
| Q5SZE1 | CERS2 | Ceramide synthase 2 (Fragment) | -1.531592131 | -0.162927 |
| Q5T0D9 | TPRG1L | Tumor protein p63-regulated gene 1-like protein | -0.969651699 | 0.0711231 |
| Q5T123 | SH3BGRL3 | SH3 domain-binding glutamic acid-rich-like protein 3 | -0.268629392 | -0.246112 |
| Q5T160 | RARS2 | Probable arginine--tRNA ligase, mitochondrial | -0.632027785 | 0.515744 |
| Q5T1M5 | FKBP15 | FK506-binding protein 15 | -0.09703207 | -0.161843 |
| Q5T280 | SPOUT1 | Putative methyltransferase C9orf114 | -0.954120239 | -1.34485 |
| Q5T2E6 | ARMH3 | Armadillo-like helical domain-containing protein 3 | -0.133764664 | 0.213647 |
| Q5T447 | HECTD3 | E3 ubiquitin-protein ligase HECTD3 | -0.124635061 | -0.306716 |
| Q5T4K2 | SLC39A1 | Zinc transporter ZIP1 (Fragment) | -1.146067381 | 0 |
| Q5T4S7 | UBR4 | E3 ubiquitin-protein ligase UBR4 | -0.132259369 | -0.272714 |
| Q5T5C0 | STXBP5 | Syntaxin-binding protein 5 | -0.086753527 | -0.191513 |
| Q5T5P2 | KIAA1217 | Sickle tail protein homolog | -0.822624922 | -0.608696 |
| Q5T6V5 | C9orf64 | Queuosine salvage protein | -0.031383355 | 0.029576 |
| Q5T749 | KPRP | Keratinocyte proline-rich protein | -0.299872875 | 0.208924 |
| Q5T750 | XP32 | Skin-specific protein 32 | -0.348531167 | -0.576871 |
| Q5T753 | LCE1E | Late cornified envelope protein 1E | -1.157926083 | 0 |
| Q5T9A4 | ATAD3B | ATPase family AAA domain-containing protein 3B | 0 | 0.712464 |
| Q5TA50 | CPTP | Ceramide-1-phosphate transfer protein | 0.042135 | -0.181791 |
| Q5TAQ0 | UQCC2 | Mitochondrial nucleoid factor 1 | -2.100770791 | 1.81892 |
| Q5TAQ9 | DCAF8 | DDB1- and CUL4-associated factor 8 | 0.265345971 | -0.612949 |
| Q5TBP9 | LSM14B | Protein LSM14 homolog B (Fragment) | -0.794638157 | -0.946884 |
| Q5TDE2 | TMEM53 | Transmembrane protein 53 | -1.39784495 | -0.763837 |
| Q5TDE7 | RNF220 | E3 ubiquitin-protein ligase RNF220 (Fragment) | -0.035533587 | -0.191659 |
| Q5TDF0 | NTPCR | Cancer-related nucleoside-triphosphatase | -0.231755416 | 0.443568 |
| Q5TDH0 | DDI2 | Protein DDI1 homolog 2 | 0.11928757 | 0.0279795 |
| Q5TEJ8 | THEMIS2 | Protein THEMIS2 | 0.610738118 | -1.70006 |
| Q5TFE4 | NT5DC1 | 5'-nucleotidase domain-containing protein 1 | 0.021488031 | 0.0303253 |
| Q5THJ4 | VPS13D | Vacuolar protein sorting-associated protein 13D | 3.394834042 | 0 |
| Q5U5Q3 | MEX3C | RNA-binding E3 ubiquitin-protein ligase MEX3C | -0.10872817 | 0.24627 |
| Q5U5X0 | LYRM7 | Complex III assembly factor LYRM7 | -1.174429973 | 0.636988 |
| Q5UIP0 | RIF1 | Telomere-associated protein RIF1 | -1.343814055 | -0.396334 |
| Q5VSL9 | STRIP1 | Striatin-interacting protein 1 | -0.034015258 | -0.0056893 |
| Q5VST6 | ABHD17B | Alpha/beta hydrolase domain-containing protein 17B | 0 | -6.54153 |
| Q5VT52 | RPRD2 | Regulation of nuclear pre-mRNA domain-containing protein 2 | -0.230570078 | 0.0225795 |
| Q5VTR2 | RNF20 | E3 ubiquitin-protein ligase BRE1A | -0.676210244 | 0.0752834 |
| Q5VVQ6 | YOD1 | Ubiquitin thioesterase OTU1 | 0.222744226 | 0.23909 |
| Q5VW32 | BROX | BRO1 domain-containing protein BROX | -0.045448542 | 0.00514444 |
| Q5VW36 | FOCAD | Focadhesin | -0.338025252 | -0.448333 |
| Q5VW52 | GPAM | Glycerol-3-phosphate acyltransferase 1, mitochondrial | -0.689132929 | 0.186526 |
| Q5VWJ9 | SNX30 | Sorting nexin-30 | 0.489203612 | -0.16131 |
| Q5VWU8 | PARD3 | Partitioning defective 3 homolog | -0.632985989 | 0.0680281 |
| Q5VWZ2 | LYPLAL1 | Lysophospholipase-like protein 1 | 0.089589516 | -0.125086 |
| Q5VY60 | HAUS6 | HAUS augmin-like complex subunit 6 | -0.123342673 | -0.19694 |
| Q5VZ72 | IZUMO3 | Izumo sperm-egg fusion protein 3 | -0.994104465 | -0.170615 |
| Q5VZ89 | DENND4C | DENN domain-containing protein 4C | 0.086916049 | -0.597216 |
| Q5VZE5 | NAA35 | N-alpha-acetyltransferase 35, NatC auxiliary subunit | -0.252810081 | 0.0326773 |

| | | | | |
|---|---|---|---|---|
| Q5W0C6 | TOR3A | Torsin-3A (Fragment) | 0.349455674 | -1.04159 |
| Q5W0V3 | FHIP2A | FHF complex subunit HOOK interacting protein 2A | 0.128359 | -0.123038 |
| Q5W111 | SPRYD7 | SPRY domain-containing protein 7 | -0.851215839 | -0.0460215 |
| Q5XKE5 | KRT79 | Keratin, type II cytoskeletal 79 | -0.874821027 | -0.277031 |
| Q5XPI4 | RNF123 | E3 ubiquitin-protein ligase RNF123 | -0.212255796 | -0.455514 |
| Q5XUX1 | FBXW9 | F-box/WD repeat-containing protein 9 | -0.166629712 | -0.839833 |
| Q60FE5 | FLNA | Filamin-A | 0.03935051 | -0.0166976 |
| Q63ZY3 | KANK2 | KN motif and ankyrin repeat domain-containing protein 2 | 0.018269857 | -0.126327 |
| Q643R3 | LPCAT4 | Lysophospholipid acyltransferase LPCAT4 | -0.980366389 | -0.0460739 |
| Q658K8 | EEF1DP3 | Putative elongation factor 1-delta-like protein | 0.078456879 | 0.227861 |
| Q658Y4 | FAM91A1 | Protein FAM91A1 | -0.199610472 | -0.396765 |
| Q66K14 | TBC1D9B | TBC1 domain family member 9B | 0.24777627 | 0.0383485 |
| Q66K74 | MAP1S | Microtubule-associated protein 1S | -0.799730778 | -0.742811 |
| Q66LE6 | PPP2R2D | Serine/threonine-protein phosphatase 2A 55 kDa regulatory subunit B delta isoform | -0.038818041 | -1.20907 |
| Q676U5 | ATG16L1 | Autophagy-related protein 16-1 | 0.113779942 | -0.155291 |
| Q68CQ4 | UTP25 | U3 small nucleolar RNA-associated protein 25 homolog | -0.184756676 | -0.167006 |
| Q68CZ2 | TNS3 | Tensin-3 | 0.134698788 | 0.0686687 |
| Q68D91 | MBLAC2 | Acyl-coenzyme A thioesterase MBLAC2 | -2.254438639 | 0.0623865 |
| Q68DQ2 | CRYBG3 | Very large A-kinase anchor protein | 0.074998299 | -0.000967979 |
| Q68E01 | INTS3 | Integrator complex subunit 3 | -1.321873188 | -0.211832 |
| Q68EM7 | ARHGAP17 | Rho GTPase-activating protein 17 | -1.427875916 | 0.522581 |
| Q69YN2 | CWF19L1 | CWF19-like protein 1 | -0.047862291 | -0.163506 |
| Q6AI08 | HEATR6 | HEAT repeat-containing protein 6 | -0.150352637 | -0.368175 |
| Q6AWC2 | WWC2 | Protein WWC2 | -0.96058766 | 0 |
| Q6DD87 | ZNF787 | Zinc finger protein 787 | -0.971754471 | 1.42851 |
| Q6DD88 | ATL3 | Atlastin-3 | -1.038962762 | 0.166333 |
| Q6DHV7 | ADAL | Adenosine deaminase-like protein | 0.253238281 | -0.60715 |
| Q6DKI1 | RPL7L1 | 60S ribosomal protein L7-like 1 | -3.136398077 | -0.721599 |
| Q6DKJ4 | NXN | Nucleoredoxin | 0.01174434 | 0.20956 |
| Q6DKK2 | TTC19 | Tetratricopeptide repeat protein 19, mitochondrial | 0.170483033 | -0.415048 |
| Q6FI81 | CIAPIN1 | Anamorsin | 0.073221525 | 0.117362 |
| Q6FIF0 | ZFAND6 | AN1-type zinc finger protein 6 | -0.319641272 | 0.498409 |
| Q6GMV2 | SMYD5 | Histone-lysine N-trimethyltransferase SMYD5 | -0.060690165 | -0.0315091 |
| Q6GQQ9 | OTUD7B | OTU domain-containing protein 7B | 0.044279099 | -0.268133 |
| Q6IA69 | NADSYN1 | Glutamine-dependent NAD(+) synthetase | -0.016568184 | -0.109385 |
| Q6IA86 | ELP2 | Elongator complex protein 2 | -0.213596026 | -0.336214 |
| Q6IAA8 | LAMTOR1 | Ragulator complex protein LAMTOR1 | -1.10277462 | 0.188331 |
| Q6IBS0 | TWF2 | Twinfilin-2 | -0.095934709 | -0.102619 |
| Q6ICB0 | DESI1 | Desumoylating isopeptidase 1 | 0.054741383 | 0.690822 |
| Q6IN85 | PPP4R3A | Serine/threonine-protein phosphatase 4 regulatory subunit 3A | 0.152024905 | -0.0576935 |
| Q6IPR3 | TYW3 | tRNA wybutosine-synthesizing protein 3 homolog | 0.006709814 | -1.49998 |
| Q6IQ49 | SDE2 | Replication stress response regulator SDE2 | -0.09484005 | -0.224889 |
| Q6KB66 | KRT80 | Keratin, type II cytoskeletal 80 | -0.184645017 | 0.0231578 |
| Q6L8Q7 | PDE12 | 2',5'-phosphodiesterase 12 | -0.019923608 | -0.0968049 |
| Q6NTF9 | RHBDD2 | Rhomboid domain-containing protein 2 | -0.820985635 | 0 |
| Q6NUK1 | SLC25A24 | Calcium-binding mitochondrial carrier protein SCaMC-1 | -1.232468287 | 0.524547 |
| Q6NUM9 | RETSAT | All-trans-retinol 13,14-reductase | -2.305059234 | 2.00569 |
| Q6NUQ1 | RINT1 | RAD50-interacting protein 1 | -0.210774183 | -0.384404 |
| Q6NUQ4 | TMEM214 | Transmembrane protein 214 | -2.051135381 | -0.259758 |
| Q6NVY1 | HIBCH | 3-hydroxyisobutyryl-CoA hydrolase, mitochondrial | 0.174054543 | -0.45342 |
| Q6NW34 | NEPRO | Nucleolus and neural progenitor protein | -0.477381468 | -0.0721356 |
| Q6NXE6 | ARMC6 | Armadillo repeat-containing protein 6 | 0.015877883 | 0.135528 |
| Q6NYC8 | PPP1R18 | Phostensin | 0.151197751 | 0.0519505 |
| Q6NZI2 | CAVIN1 | Caveolae-associated protein 1 | -0.658584277 | -0.09373 |
| Q6NZY4 | ZCCHC8 | Zinc finger CCHC domain-containing protein 8 | 0.254322767 | 0.127632 |
| Q6P179 | ERAP2 | Endoplasmic reticulum aminopeptidase 2 | -0.034294446 | 0.0916977 |
| Q6P1J9 | CDC73 | Parafibromin | 0.000975291 | -0.0533034 |
| Q6P1K2 | PMF1 | Polyamine-modulated factor 1 | -0.645481904 | 1.52356 |
| Q6P1L8 | MRPL14 | 39S ribosomal protein L14, mitochondrial | -1.365134478 | 0.605291 |
| Q6P1M0 | SLC27A4 | Long-chain fatty acid transport protein 4 | -1.892537912 | 0.28259 |
| Q6P1N0 | CC2D1A | Coiled-coil and C2 domain-containing protein 1A | -0.140352329 | -0.0653513 |
| Q6P1Q9 | METTL2B | tRNA N(3)-methylcytidine methyltransferase METTL2B | -0.003728867 | 0.0277681 |
| Q6P1R4 | DUS1L | tRNA-dihydrouridine(16/17) synthase [NAD(P)(+)]-like | 0.291172822 | -0.645456 |
| Q6P1S2 | C3orf33 | Protein C3orf33 | -1.725817283 | 0.625144 |
| Q6P2E9 | EDC4 | Enhancer of mRNA-decapping protein 4 | -0.429012775 | -0.164193 |
| Q6P2P2 | PRMT9 | Protein arginine N-methyltransferase 9 | -0.166429361 | -0.504099 |
| Q6P2Q9 | PRPF8 | Pre-mRNA-processing-splicing factor 8 | -0.498984178 | -0.0808163 |
| Q6P3S6 | FBXO42 | F-box only protein 42 | 0.215006669 | 0.481028 |
| Q6P3X3 | TTC27 | Tetratricopeptide repeat protein 27 | -0.06822443 | 0.00808938 |
| Q6P4A8 | PLBD1 | Phospholipase B-like 1 | -0.633697271 | 0.255129 |

| | | | | |
|---|---|---|---|---|
| Q6P4I2 | WDR73 | WD repeat-containing protein 73 | -0.575132449 | -0.750592 |
| Q6P587 | FAHD1 | Acylpyruvase FAHD1, mitochondrial | 0.094991128 | 0.288829 |
| Q6P6C2 | ALKBH5 | RNA demethylase ALKBH5 | -0.130863349 | 0.428781 |
| Q6P996 | PDXDC1 | Pyridoxal-dependent decarboxylase domain-containing protein 1 | 0.027384281 | -0.102864 |
| Q6P9B6 | MEAK7 | MTOR-associated protein MEAK7 | -0.365824461 | 0.11599 |
| Q6PCE3 | PGM2L1 | Glucose 1,6-bisphosphate synthase | 0.084876299 | -0.00392246 |
| Q6PD62 | CTR9 | RNA polymerase-associated protein CTR9 homolog | -0.418916941 | -0.0212221 |
| Q6PD74 | AAGAB | Alpha- and gamma-adaptin-binding protein p34 | -0.335408211 | 0.0856886 |
| Q6PGP7 | TTC37 | Tetratricopeptide repeat protein 37 | -0.836714745 | -0.380267 |
| Q6PHR2 | ULK3 | Serine/threonine-protein kinase ULK3 | -0.321364244 | -0.652161 |
| Q6PI48 | DARS2 | Aspartate--tRNA ligase, mitochondrial | -1.216390292 | -0.0943604 |
| Q6PI78 | TMEM65 | Transmembrane protein 65 | -2.498262882 | -0.863975 |
| Q6PID6 | TTC33 | Tetratricopeptide repeat protein 33 | -0.999654849 | 0 |
| Q6PIJ6 | FBXO38 | F-box only protein 38 | -2.127400716 | -0.014486 |
| Q6PIW4 | FIGNL1 | Fidgetin-like protein 1 | 0.004454613 | -0.147682 |
| Q6PJG6 | BRAT1 | BRCA1-associated ATM activator 1 | -0.015721083 | -0.225115 |
| Q6PKG0 | LARP1 | La-related protein 1 | -0.378919125 | -0.487775 |
| Q6PL18 | ATAD2 | ATPase family AAA domain-containing protein 2 | -0.535815001 | -0.507526 |
| Q6PL24 | TMED8 | Protein TMED8 | -0.105371634 | 0.113381 |
| Q6QNY0 | BLOC1S3 | Biogenesis of lysosome-related organelles complex 1 subunit 3 | 0.260540167 | -0.0686928 |
| Q6RFH5 | WDR74 | WD repeat-containing protein 74 | -1.638052622 | -0.000221252 |
| Q6SZW1 | SARM1 | NAD(+) hydrolase SARM1 | 0.031200091 | -0.142641 |
| Q6ULP2 | AFTPH | Aftiphilin | 0.539093812 | 3.8408 |
| Q6UN15 | FIP1L1 | Pre-mRNA 3'-end-processing factor FIP1 | -0.547982613 | 0.466016 |
| Q6UUV9 | CRTC1 | CREB-regulated transcription coactivator 1 | -0.044853528 | 0.505551 |
| Q6UVK1 | CSPG4 | Chondroitin sulfate proteoglycan 4 | -0.401607593 | 0.332424 |
| Q6UVY6 | MOXD1 | DBH-like monooxygenase protein 1 | -0.869218667 | -2.57237 |
| Q6UW63 | POGLUT2 | Protein O-glucosyltransferase 2 | 0.317957719 | 1.17515 |
| Q6UWP2 | DHRS11 | Dehydrogenase/reductase SDR family member 11 | -0.154846986 | -0.0988299 |
| Q6UWP8 | SBSN | Suprabasin | -0.607435067 | 0.311087 |
| Q6UWV6 | ENPP7 | Ectonucleotide pyrophosphatase/phosphodiesterase family member 7 | -1.555570682 | 0 |
| Q6UX04 | CWC27 | Spliceosome-associated protein CWC27 homolog | -0.328640064 | -0.0412699 |
| Q6UXH1 | CRELD2 | Protein disulfide isomerase CRELD2 | -0.131875436 | -0.329051 |
| Q6UXN9 | WDR82 | WD repeat-containing protein 82 | -0.214292049 | -0.130836 |
| Q6V1X1 | DPP8 | Dipeptidyl peptidase 8 | -0.001902501 | -0.271186 |
| Q6VY07 | PACS1 | Phosphofurin acidic cluster sorting protein 1 | -0.097058853 | -0.0245399 |
| Q6XQN6 | NAPRT | Nicotinate phosphoribosyltransferase | 0.169305086 | -0.00655206 |
| Q6XZF7 | DNMBP | Dynamin-binding protein | -0.199438016 | 0.0888974 |
| Q6Y2X3 | DNAJC14 | DnaJ homolog subfamily C member 14 | -1.151075363 | -0.180626 |
| Q6Y7W6 | GIGYF2 | GRB10-interacting GYF protein 2 | -0.305581967 | -0.171045 |
| Q6YHK3 | CD109 | CD109 antigen | -0.937960863 | 3.18813 |
| Q6YHU6 | THADA | Thyroid adenoma-associated protein | -0.269291242 | -0.320957 |
| Q6YN16 | HSDL2 | Hydroxysteroid dehydrogenase-like protein 2 | -0.744817813 | 0.341212 |
| Q6YP21 | KYAT3 | Kynurenine--oxoglutarate transaminase 3 | 0.137675444 | 0.0871356 |
| Q6ZNF0 | ACP7 | Acid phosphatase type 7 | -1.715738932 | 1.88601 |
| Q6ZRP7 | QSOX2 | Sulfhydryl oxidase 2 | -0.421106021 | -0.321508 |
| Q6ZS17 | RIPOR1 | Rho family-interacting cell polarization regulator 1 | -0.234778404 | 0.282392 |
| Q6ZS86 | GK5 | Putative glycerol kinase 5 | -1.272693237 | -0.879298 |
| Q6ZT12 | UBR3 | E3 ubiquitin-protein ligase UBR3 | 0.589740912 | -2.89134 |
| Q6ZTN6 | ANKRD13D | Ankyrin repeat domain-containing protein 13D | 0.520446142 | -1.83774 |
| Q6ZUT6 | CCDC9B | Coiled-coil domain-containing protein 9B | -0.716216087 | 0.00981299 |
| Q6ZVM7 | TOM1L2 | TOM1-like protein 2 | 0.127520482 | 0.201127 |
| Q6ZVX7 | NCCRP1 | F-box only protein 50 | -0.519263585 | 0.0859133 |
| Q6ZW49 | PAXIP1 | PAX-interacting protein 1 | -0.819717487 | -0.51401 |
| Q6ZXV5 | TMTC3 | Protein O-mannosyl-transferase TMTC3 | -0.940491199 | -0.294757 |
| Q709C8 | VPS13C | Vacuolar protein sorting-associated protein 13C | -1.104105473 | 0.0567048 |
| Q70CQ2 | USP34 | Ubiquitin carboxyl-terminal hydrolase 34 | 0.009866556 | 0.414829 |
| Q70IA6 | MOB2 | MOB kinase activator 2 | 0.111953656 | 0.0892328 |
| Q70UQ0 | IKBIP | Inhibitor of nuclear factor kappa-B kinase-interacting protein | -1.475203435 | 0.189924 |
| Q70UQ0-4 | IKBIP | Isoform 4 of Inhibitor of nuclear factor kappa-B kinase-interacting protein | -1.424902201 | 0.418431 |
| Q70Z53 | FRA10AC1 | Protein FRA10AC1 | -0.653446992 | 0 |
| Q712K3 | UBE2R2 | Ubiquitin-conjugating enzyme E2 R2 | 0.09184583 | 0.0423985 |
| Q71RC2 | LARP4 | La-related protein 4 | -0.330943425 | -0.216153 |
| Q7KZ85 | SUPT6H | Transcription elongation factor SPT6 | -0.039098978 | -0.118492 |
| Q7KZF4 | SND1 | Staphylococcal nuclease domain-containing protein 1 | -0.47546188 | -0.143566 |
| Q7L099 | RUFY3 | Protein RUFY3 | -0.042484125 | -0.667121 |
| Q7L0Y3 | TRMT10C | tRNA methyltransferase 10 homolog C | -1.053825537 | -0.170999 |
| Q7L1Q6 | BZW1 | eIF5-mimic protein 2 | -0.108698845 | -0.0453599 |
| Q7L1T6 | CYB5R4 | Cytochrome b5 reductase 4 | -0.052214225 | -0.421243 |

| | | | | |
|---|---|---|---|---|
| Q7L266 | ASRGL1 | Isoaspartyl peptidase/L-asparaginase | 0.505987167 | 0 |
| Q7L273 | KCTD9 | BTB/POZ domain-containing protein KCTD9 | -1.769105514 | 0.0432673 |
| Q7L2H7 | EIF3M | Eukaryotic translation initiation factor 3 subunit M | -0.142149687 | -0.229836 |
| Q7L2J0 | MEPCE | 7SK snRNA methylphosphate capping enzyme | -0.332773606 | -0.643088 |
| Q7L4I2 | RSRC2 | Arginine/serine-rich coiled-coil protein 2 | -0.236838261 | -0.227336 |
| Q7L523 | RRAGA | Ras-related GTP-binding protein A | -0.152824879 | -0.148964 |
| Q7L576 | CYFIP1 | Cytoplasmic FMR1-interacting protein 1 | -0.476111809 | 0.179195 |
| Q7L5D6 | GET4 | Golgi to ER traffic protein 4 homolog | 0.035038869 | -0.0913506 |
| Q7L5L3 | GDPD3 | Lysophospholipase D GDPD3 | -0.864628394 | 0.0114183 |
| Q7L5N7 | LPCAT2 | Lysophosphatidylcholine acyltransferase 2 | -1.092504422 | -0.232738 |
| Q7L775 | EPM2AIP1 | EPM2A-interacting protein 1 | -0.130387227 | -0.101974 |
| Q7L7V1 | DHX32 | Putative pre-mRNA-splicing factor ATP-dependent RNA helicase DHX32 | -0.546919664 | 0.980378 |
| Q7L7X3 | TAOK1 | Serine/threonine-protein kinase TAO1 | -0.620573521 | 0.602615 |
| Q7L8J4 | SH3BP5L | SH3 domain-binding protein 5-like | 0.056673447 | -0.0702496 |
| Q7L8W6 | DPH6 | Diphthine--ammonia ligase | -0.368124326 | 1.07314 |
| Q7L9L4 | MOB1B | MOB kinase activator 1B | 0.119265874 | -0.027715 |
| Q7LBC6 | KDM3B | Lysine-specific demethylase 3B | 0.032916228 | -0.0191603 |
| Q7LBR1 | CHMP1B | Charged multivesicular body protein 1b | -0.669531027 | -0.120002 |
| Q7LG56 | RRM2B | Ribonucleoside-diphosphate reductase subunit M2 B | -0.073956887 | 0.0356684 |
| Q7RTP6 | MICAL3 | [F-actin]-monooxygenase MICAL3 | -0.009299914 | 0.206099 |
| Q7RTS9 | DYM | Dymeclin | -0.030542612 | -0.273575 |
| Q7RTV0 | PHF5A | PHD finger-like domain-containing protein 5A | -0.158043941 | 0.152675 |
| Q7RTV5 | PRXL2C | Peroxiredoxin-like 2C | -0.957611322 | -0.155854 |
| Q7Z2T5 | TRMT1L | TRMT1-like protein | -0.093521595 | -0.0509701 |
| Q7Z2W4 | ZC3HAV1 | Zinc finger CCCH-type antiviral protein 1 | -0.025095145 | 0.0358648 |
| Q7Z2W9 | MRPL21 | 39S ribosomal protein L21, mitochondrial | -1.813144684 | -0.16302 |
| Q7Z2X4 | PID1 | PTB-containing, cubilin and LRP1-interacting protein | -0.649711172 | 0.0610712 |
| Q7Z2Z2 | EFL1 | Elongation factor-like GTPase 1 | 0.06311059 | -0.0550871 |
| Q7Z392 | TRAPPC11 | Trafficking protein particle complex subunit 11 | -0.497429212 | -0.378699 |
| Q7Z3B4 | NUP54 | Nucleoporin p54 | 0.093496561 | -0.0164849 |
| Q7Z3C6 | ATG9A | Autophagy-related protein 9A | 0.193729719 | -2.12729 |
| Q7Z3J2 | VPS35L | VPS35 endosomal protein-sorting factor-like | -0.373503049 | -0.529486 |
| Q7Z3K3 | POGZ | Pogo transposable element with ZNF domain | -0.725609938 | 0.135524 |
| Q7Z3K6 | MIER3 | Mesoderm induction early response protein 3 | -0.500960191 | 0.784248 |
| Q7Z3T8 | ZFYVE16 | Zinc finger FYVE domain-containing protein 16 | 0.082780759 | 0.136876 |
| Q7Z3U7-5 | MON2 | Isoform 5 of Protein MON2 homolog | 0.086075147 | -0.208944 |
| Q7Z3V4 | UBE3B | Ubiquitin-protein ligase E3B | -0.713285605 | -0.346178 |
| Q7Z3Y7 | KRT28 | Keratin, type I cytoskeletal 28 | 0.29540503 | -1.91106 |
| Q7Z3Y8 | KRT27 | Keratin, type I cytoskeletal 27 | -0.855119785 | -0.828968 |
| Q7Z3Z0 | KRT25 | Keratin, type I cytoskeletal 25 | -0.08937184 | -0.451866 |
| Q7Z417 | NUFIP2 | Nuclear fragile X mental retardation-interacting protein 2 | -0.52285401 | 0.337851 |
| Q7Z422 | SZRD1 | SUZ domain-containing protein 1 | 0.297333082 | -0.089351 |
| Q7Z434 | MAVS | Mitochondrial antiviral-signaling protein | -0.676711559 | 0.131831 |
| Q7Z4F1 | LRP10 | Low-density lipoprotein receptor-related protein 10 | 0 | -0.192357 |
| Q7Z4G4 | TRMT11 | tRNA (guanine(10)-N2)-methyltransferase homolog | -0.375300646 | -0.573256 |
| Q7Z4H3 | HDDC2 | 5'-deoxynucleotidase HDDC2 | 0.089867036 | 0.368496 |
| Q7Z4H8 | POGLUT3 | Protein O-glucosyltransferase 3 | -0.297889868 | 0.0845896 |
| Q7Z4Q2 | HEATR3 | HEAT repeat-containing protein 3 | -0.179692825 | -0.0280689 |
| Q7Z4V5 | HDGFL2 | Hepatoma-derived growth factor-related protein 2 | 0.006117821 | -0.10353 |
| Q7Z5G4 | GOLGA7 | Golgin subfamily A member 7 | -1.468294144 | -0.138951 |
| Q7Z5H4 | VN1R5 | Vomeronasal type-1 receptor 5 | -0.13287727 | 0.278593 |
| Q7Z5K2 | WAPL | Wings apart-like protein homolog | -0.599997123 | 1.96689 |
| Q7Z5L9 | IRF2BP2 | Interferon regulatory factor 2-binding protein 2 | -0.167247613 | 0.302269 |
| Q7Z5R6 | APBB1IP | Amyloid beta A4 precursor protein-binding family B member 1-interacting protein | -0.433477879 | -0.291946 |
| Q7Z6B0 | CCDC91 | Coiled-coil domain-containing protein 91 | -0.08966732 | -0.139341 |
| Q7Z6J0 | SH3RF1 | E3 ubiquitin-protein ligase SH3RF1 | 0.198165099 | 0.120761 |
| Q7Z6J8 | UBE3D | E3 ubiquitin-protein ligase E3D | -0.447311163 | 0 |
| Q7Z6K3 | PTAR1 | Protein prenyltransferase alpha subunit repeat-containing protein 1 | 0.157820384 | -0.327936 |
| Q7Z6L1 | TECPR1 | Tectonin beta-propeller repeat-containing protein 1 | -1.639935096 | 1.00933 |
| Q7Z6M1 | RABEPK | Rab9 effector protein with kelch motifs | -0.570593834 | 0.0248111 |
| Q7Z6V5 | ADAT2 | tRNA-specific adenosine deaminase 2 | -0.92256856 | 1.2856 |
| Q7Z6Z7 | HUWE1 | E3 ubiquitin-protein ligase HUWE1 | -0.046100299 | -0.112877 |
| Q7Z794 | KRT77 | Keratin, type II cytoskeletal 1b | -0.036890825 | 0.0586405 |
| Q7Z7E8 | UBE2Q1 | Ubiquitin-conjugating enzyme E2 Q1 | 0.0694019 | -0.0887731 |
| Q7Z7F7 | MRPL55 | 39S ribosomal protein L55, mitochondrial | 0 | 0.644418 |
| Q7Z7H5 | TMED4 | Transmembrane emp24 domain-containing protein 4 | -1.582309882 | -0.0680027 |
| Q7Z7K6 | CENPV | Centromere protein V | -0.994294167 | 0.422431 |
| Q7Z7L7 | ZER1 | Protein zer-1 homolog | -0.333046595 | 0.30939 |

| | | | | |
|---|---|---|---|---|
| Q7Z7L8 | C11orf96 | Uncharacterized protein C11orf96 | 0.160784721 | -0.118551 |
| Q86SE9 | PCGF5 | Polycomb group RING finger protein 5 | 1.198274533 | 0 |
| Q86SF2 | GALNT7 | N-acetylgalactosaminyltransferase 7 | -1.043206453 | 0.0999037 |
| Q86SK9 | SCD5 | Stearoyl-CoA desaturase 5 | 1.800089558 | -0.96379 |
| Q86SQ9 | DHDDS | Dehydrodolichyl diphosphate synthase complex subunit DHDDS | -0.976339658 | -0.0518392 |
| Q86SX6 | GLRX5 | Glutaredoxin-related protein 5, mitochondrial | -0.17321078 | 0.577321 |
| Q86SZ2 | TRAPPC6B | Trafficking protein particle complex subunit 6B | -0.661556244 | -0.00842253 |
| Q86TB9 | PATL1 | Protein PAT1 homolog 1 | -0.407317638 | -0.362257 |
| Q86TI2 | DPP9 | Dipeptidyl peptidase 9 | 0.053686539 | 0.00669511 |
| Q86TP1 | PRUNE1 | Exopolyphosphatase PRUNE1 | -0.111145973 | -0.130388 |
| Q86TU7 | SETD3 | Actin-histidine N-methyltransferase | -0.050319115 | -0.126555 |
| Q86U28 | ISCA2 | Iron-sulfur cluster assembly 2 homolog, mitochondrial | 0.332942963 | -0.620471 |
| Q86U38 | NOP9 | Nucleolar protein 9 | -0.833857298 | -1.4115 |
| Q86U44 | METTL3 | N6-adenosine-methyltransferase catalytic subunit | 0.044397672 | 0.105311 |
| Q86U90 | YRDC | YrdC domain-containing protein, mitochondrial | -0.038412094 | 0.413941 |
| Q86UA1 | PRPF39 | Pre-mRNA-processing factor 39 | 0.046438138 | -0.075422 |
| Q86UA6 | RPAIN | RPA-interacting protein | -1.117676973 | -0.0508814 |
| Q86UE4 | MTDH | Protein LYRIC | -0.909723997 | 0.293586 |
| Q86UL3 | GPAT4 | Glycerol-3-phosphate acyltransferase 4 | -1.141431332 | -0.134359 |
| Q86UP2 | KTN1 | Kinectin | -0.916052898 | 0.23256 |
| Q86US8 | SMG6 | Telomerase-binding protein EST1A | -0.795410633 | -0.177884 |
| Q86UU1 | PHLDB1 | Pleckstrin homology-like domain family B member 1 | -0.109730164 | -0.237132 |
| Q86UV5 | USP48 | Ubiquitin carboxyl-terminal hydrolase 48 | 0.168026129 | -0.0738382 |
| Q86UX7 | FERMT3 | Fermitin family homolog 3 | 0.897535801 | -0.125603 |
| Q86V21 | AACS | Acetoacetyl-CoA synthetase | 0.235451937 | 0.0269785 |
| Q86V48 | LUZP1 | Leucine zipper protein 1 | -1.013214986 | 0.147968 |
| Q86V81 | ALYREF | THO complex subunit 4 | -0.598510265 | 0.0207319 |
| Q86V85 | GPR180 | Integral membrane protein GPR180 | -0.657436053 | 0.868395 |
| Q86V87 | FHIP2B | FHF complex subunit HOOK interacting protein 2B | 1.119710048 | -2.73615 |
| Q86VI3 | IQGAP3 | Ras GTPase-activating-like protein IQGAP3 | 0.429043214 | -0.271478 |
| Q86VP6 | CAND1 | Cullin-associated NEDD8-dissociated protein 1 | -0.074779669 | -0.0948537 |
| Q86VS8 | HOOK3 | Protein Hook homolog 3 | -0.01492842 | 0.0796057 |
| Q86VW0 | SESTD1 | SEC14 domain and spectrin repeat-containing protein 1 | -0.341771205 | -0.565595 |
| Q86VY9 | TMEM200A | Transmembrane protein 200A | 0.636958361 | 0 |
| Q86W42 | THOC6 | THO complex subunit 6 homolog | -0.793529908 | -0.0155296 |
| Q86W50 | METTL16 | RNA N6-adenosine-methyltransferase METTL16 | -0.632531484 | -0.19381 |
| Q86W56 | PARG | Poly(ADP-ribose) glycohydrolase | -0.167585929 | 0.33019 |
| Q86W92 | PPFIBP1 | Liprin-beta-1 | -0.85658137 | -0.268673 |
| Q86WJ1 | CHD1L | Chromodomain-helicase-DNA-binding protein 1-like | -0.460635662 | 0.176457 |
| Q86WN1 | FCHSD1 | F-BAR and double SH3 domains protein 1 | 0.184263786 | -0.0309976 |
| Q86WQ0 | NR2C2AP | Nuclear receptor 2C2-associated protein | -0.024485509 | -0.105991 |
| Q86WR0 | CCDC25 | Coiled-coil domain-containing protein 25 | 0.036191146 | -0.334727 |
| Q86WX3 | RPS19BP1 | Active regulator of SIRT1 | -2.622386018 | 0.325033 |
| Q86X02 | CDR2L | Cerebellar degeneration-related protein 2-like | 0.057499886 | 3.42026 |
| Q86X27 | RALGPS2 | Ras-specific guanine nucleotide-releasing factor RalGPS2 | 1.023189068 | 0 |
| Q86X55 | CARM1 | Histone-arginine methyltransferase CARM1 | 0.002328396 | -0.264417 |
| Q86X76 | NIT1 | Deaminated glutathione amidase | -0.199459314 | -0.0307741 |
| Q86X83 | COMMD2 | COMM domain-containing protein 2 | -0.18494153 | 0.377294 |
| Q86XL3 | ANKLE2 | Ankyrin repeat and LEM domain-containing protein 2 | -0.960878213 | 0.347382 |
| Q86XN8 | MEX3D | RNA-binding protein MEX3D | -0.307669004 | -0.0409063 |
| Q86XP3 | DDX42 | ATP-dependent RNA helicase DDX42 | -0.109867414 | 0.121365 |
| Q86XZ4 | SPATS2 | Spermatogenesis-associated serine-rich protein 2 | -0.893560092 | -0.000188192 |
| Q86Y37 | CACUL1 | CDK2-associated and cullin domain-containing protein 1 | 0.106572469 | 0.453712 |
| Q86Y38 | XYLT1 | Xylosyltransferase 1 | -0.019388199 | -0.36187 |
| Q86Y56 | DNAAF5 | Dynein axonemal assembly factor 5 | 0.031109571 | -0.00557613 |
| Q86Y82 | STX12 | Syntaxin-12 | -0.617451429 | 0.926148 |
| Q86YJ7 | ANKRD13B | Ankyrin repeat domain-containing protein 13B | -0.509437799 | -1.85367 |
| Q86YL5 | TDRP | Testis development-related protein | -0.590757052 | 0.321866 |
| Q86YN1 | DOLPP1 | Dolichyldiphosphatase 1 | -1.639978806 | 0 |
| Q86YP4 | GATAD2A | Transcriptional repressor p66-alpha | -0.51183327 | 0.432124 |
| Q86YQ8 | CPNE8 | Copine-8 | -0.183816036 | 0.0344133 |
| Q86YS6 | RAB43 | Ras-related protein Rab-43 | -0.370623032 | -0.071126 |
| Q86YS7 | C2CD5 | C2 domain-containing protein 5 | -0.296445688 | -0.834133 |
| Q86YT6 | MIB1 | E3 ubiquitin-protein ligase MIB1 | -0.290807724 | -0.788424 |
| Q86YV9 | HPS6 | Hermansky-Pudlak syndrome 6 protein | 0.253153483 | -1.98017 |
| Q86YZ3 | HRNR | Hornerin | -0.052948078 | -0.106923 |
| Q8IU81 | IRF2BP1 | Interferon regulatory factor 2-binding protein 1 | -0.596316099 | 0.627124 |
| Q8IU85 | CAMK1D | Calcium/calmodulin-dependent protein kinase type 1D | 0.140249968 | -0.400091 |
| Q8IUC4 | RHPN2 | Rhophilin-2 | -0.456501245 | 0.761494 |
| Q8IUE6 | H2AC21 | Histone H2A type 2-B | -0.837316513 | -0.401121 |
| Q8IUF8 | RIOX2 | Ribosomal oxygenase 2 | -0.720490376 | -0.572376 |

| | | | | |
|---|---|---|---|---|
| Q8IUI8 | CRLF3 | Cytokine receptor-like factor 3 | 0.031802098 | 0.0723534 |
| Q8IUI8-2 | CRLF3 | Isoform 2 of Cytokine receptor-like factor 3 | -0.471007268 | 1.00478 |
| Q8IUQ4 | SIAH1 | E3 ubiquitin-protein ligase SIAH1 | -1.095033884 | -0.860496 |
| Q8IUR0 | TRAPPC5 | Trafficking protein particle complex subunit 5 | -0.267018 | -0.253474 |
| Q8IUS5 | EPHX4 | Epoxide hydrolase 4 | -0.982922792 | -0.07938 |
| Q8IV08 | PLD3 | 5'-3' exonuclease PLD3 | 0.005885124 | 0.0324631 |
| Q8IV38 | ANKMY2 | Ankyrin repeat and MYND domain-containing protein 2 | -0.024987221 | -0.0360184 |
| Q8IV48 | ERI1 | 3'-5' exoribonuclease 1 | -0.219782909 | 0.287136 |
| Q8IV50 | LYSMD2 | LysM and putative peptidoglycan-binding domain-containing protein 2 | -1.614780029 | -0.115206 |
| Q8IVD9 | NUDCD3 | NudC domain-containing protein 3 | 0.059574048 | -0.261231 |
| Q8IVF2 | AHNAK2 | Protein AHNAK2 | -0.479534388 | 0.18288 |
| Q8IVL6 | P3H3 | Prolyl 3-hydroxylase 3 | -1.101810694 | -2.33578 |
| Q8IVM0 | CCDC50 | Coiled-coil domain-containing protein 50 | -0.065597455 | -0.230587 |
| Q8IW35 | CEP97 | Centrosomal protein of 97 kDa | -0.145866871 | -0.0284379 |
| Q8IW75 | SERPINA12 | Serpin A12 | -0.10740757 | -0.162724 |
| Q8IWA0 | WDR75 | WD repeat-containing protein 75 | -1.479264577 | -2.26174 |
| Q8IWB1 | ITPRIP | Inositol 1,4,5-trisphosphate receptor-interacting protein | -1.582647244 | -2.94612 |
| Q8IWB7 | WDFY1 | WD repeat and FYVE domain-containing protein 1 | -0.126863877 | -0.0672197 |
| Q8IWE2 | FAM114A1 | Protein NOXP20 | -0.030239503 | -0.175054 |
| Q8IWJ2 | GCC2 | GRIP and coiled-coil domain-containing protein 2 | 0.151844581 | 0.0512946 |
| Q8IWT6 | LRRC8A | Volume-regulated anion channel subunit LRRC8A | -1.562648535 | 0.00158723 |
| Q8IWV7 | UBR1 | E3 ubiquitin-protein ligase UBR1 | -0.067836364 | -0.199121 |
| Q8IWV8 | UBR2 | E3 ubiquitin-protein ligase UBR2 | -0.269848665 | 0.512608 |
| Q8IWY9 | CDAN1 | Codanin-1 | -0.208834092 | 0.516257 |
| Q8IWZ3 | ANKHD1 | Ankyrin repeat and KH domain-containing protein 1 | 0.155370871 | -0.362385 |
| Q8IWZ6 | BBS7 | Bardet-Biedl syndrome 7 protein | -0.080272834 | -0.987123 |
| Q8IWZ8 | SUGP1 | SURP and G-patch domain-containing protein 1 | -0.038283189 | 0.149277 |
| Q8IX04 | UEVLD | Ubiquitin-conjugating enzyme E2 variant 3 | -0.041969697 | -0.240073 |
| Q8IX12 | CCAR1 | Cell division cycle and apoptosis regulator protein 1 | 0.09691755 | -0.0208925 |
| Q8IX90 | SKA3 | Spindle and kinetochore-associated protein 3 | -0.216403087 | -0.309667 |
| Q8IXI1 | RHOT2 | Mitochondrial Rho GTPase 2 | -0.036909382 | 0.496632 |
| Q8IXJ6 | SIRT2 | NAD-dependent protein deacetylase sirtuin-2 | -0.041789611 | 0.132567 |
| Q8IXM3 | MRPL41 | 39S ribosomal protein L41, mitochondrial | -1.120589256 | -0.0722284 |
| Q8IXT5 | RBM12B | RNA-binding protein 12B | 0.072534402 | -0.0313835 |
| Q8IXW5 | RPAP2 | Putative RNA polymerase II subunit B1 CTD phosphatase RPAP2 | -0.782146295 | 0.283409 |
| Q8IY47 | KBTBD2 | Kelch repeat and BTB domain-containing protein 2 | -0.241846323 | -0.481372 |
| Q8IY81 | FTSJ3 | pre-rRNA 2'-O-ribose RNA methyltransferase FTSJ3 | -0.626652161 | 0.564711 |
| Q8IY95 | TMEM192 | Transmembrane protein 192 | -0.149316947 | -0.900706 |
| Q8IYB1 | MB21D2 | Nucleotidyltransferase MB21D2 | -0.037405729 | 0.0363814 |
| Q8IYB5 | SMAP1 | Stromal membrane-associated protein 1 | -0.148552895 | -0.231902 |
| Q8IYB7 | DIS3L2 | DIS3-like exonuclease 2 | -0.444360654 | -0.222705 |
| Q8IYB8 | SUPV3L1 | ATP-dependent RNA helicase SUPV3L1, mitochondrial | -1.225419362 | -0.709284 |
| Q8IYD1 | GSPT2 | Eukaryotic peptide chain release factor GTP-binding subunit ERF3B | 0.476067861 | -0.211219 |
| Q8IYI6 | EXOC8 | Exocyst complex component 8 | 0.208890835 | -0.306892 |
| Q8IYM9 | TRIM22 | E3 ubiquitin-protein ligase TRIM22 | 1.009602865 | 3.97818 |
| Q8IYQ7 | THNSL1 | Threonine synthase-like 1 | 0.510204713 | -0.862751 |
| Q8IYR2 | SMYD4 | SET and MYND domain-containing protein 4 | 0.051430464 | 2.25608 |
| Q8IYS1 | PM20D2 | Xaa-Arg dipeptidase | -0.39283673 | 0.635373 |
| Q8IYS2 | KIAA2013 | Uncharacterized protein KIAA2013 | -2.229324182 | 0.00755183 |
| Q8IYU2 | HACE1 | E3 ubiquitin-protein ligase HACE1 | -0.22324117 | 0.174815 |
| Q8IZ07 | ANKRD13A | Ankyrin repeat domain-containing protein 13A | 0.001114289 | 0.0931114 |
| Q8IZ13 | ZBED8 | Protein ZBED8 | 0.286951105 | 0 |
| Q8IZ21 | PHACTR4 | Phosphatase and actin regulator 4 | -1.261549791 | 0.728907 |
| Q8IZ52 | CHPF | Chondroitin sulfate synthase 2 | -0.88493077 | -1.01214 |
| Q8IZ73 | RPUSD2 | RNA pseudouridylate synthase domain-containing protein 2 | -0.164355199 | -0.152753 |
| Q8IZ81 | ELMOD2 | ELMO domain-containing protein 2 | -1.053439299 | -0.164107 |
| Q8IZV5 | RDH10 | Retinol dehydrogenase 10 | -1.3686951 | 0.104618 |
| Q8N0T1 | RBIS | Ribosomal biogenesis factor | -1.24564751 | 0.181636 |
| Q8N0X7 | SPART | Spartin | -0.441688855 | -0.00233396 |
| Q8N0Z6 | TTC5 | Tetratricopeptide repeat protein 5 | 0.061443329 | -0.188061 |
| Q8N108 | MIER1 | Mesoderm induction early response protein 1 | 0.054812193 | -0.470637 |
| Q8N122 | RPTOR | Regulatory-associated protein of mTOR | 0.081133366 | -0.527391 |
| Q8N138 | ORMDL3 | ORM1-like protein 3 | -1.675644 | -0.311583 |
| Q8N163 | CCAR2 | Cell cycle and apoptosis regulator protein 2 | 0.175751925 | -0.144091 |
| Q8N183 | NDUFAF2 | NADH dehydrogenase [ubiquinone] 1 alpha subcomplex assembly factor 2 | -1.466444969 | 0.415026 |
| Q8N1F7 | NUP93 | Nuclear pore complex protein Nup93 | 0.16804266 | 0.0971724 |
| Q8N1G2 | CMTR1 | Cap-specific mRNA (nucleoside-2'-O-)-methyltransferase 1 | 0.13698562 | 0.062185 |
| Q8N1G4 | LRRC47 | Leucine-rich repeat-containing protein 47 | 0.098940452 | 0.0265233 |
| Q8N1N4 | KRT78 | Keratin, type II cytoskeletal 78 | -0.234513521 | 0.153515 |

| | | | | |
|---|---|---|---|---|
| Q8N1Q1 | CA13 | Carbonic anhydrase 13 | -0.550813039 | -0.615589 |
| Q8N201 | INTS1 | Integrator complex subunit 1 | 0.072125038 | 0.814785 |
| Q8N2K0 | ABHD12 | Lysophosphatidylserine lipase ABHD12 | -2.005544901 | 0.340422 |
| Q8N2W9 | PIAS4 | E3 SUMO-protein ligase PIAS4 | 1.119148811 | 0 |
| Q8N2Z9 | CENPS | Centromere protein S | -0.363128821 | 2.29905 |
| Q8N302 | AGGF1 | Angiogenic factor with G patch and FHA domains 1 | 0.100457748 | -0.40158 |
| Q8N335 | GPD1L | Glycerol-3-phosphate dehydrogenase 1-like protein | -0.121827761 | 0.314553 |
| Q8N357 | SLC35F6 | Solute carrier family 35 member F6 | -1.749760588 | 0.278603 |
| Q8N392 | ARHGAP18 | Rho GTPase-activating protein 18 | 0.138339202 | 0.036898 |
| Q8N3C0 | ASCC3 | Activating signal cointegrator 1 complex subunit 3 | -0.323179881 | -0.659511 |
| Q8N3D4 | EHBP1L1 | EH domain-binding protein 1-like protein 1 | -0.571323951 | 0.406908 |
| Q8N3R9 | PALS1 | Protein PALS1 | -0.388048649 | 0.119857 |
| Q8N3U4 | STAG2 | Cohesin subunit SA-2 | -2.082921664 | -0.635117 |
| Q8N3V7 | SYNPO | Synaptopodin | -0.995856206 | 0.791426 |
| Q8N3X1 | FNBP4 | Formin-binding protein 4 | 0.065953732 | -0.0182896 |
| Q8N488 | RYBP | RING1 and YY1-binding protein | -0.426098982 | -1.09555 |
| Q8N490-2 | PNKD | Isoform 2 of Probable hydrolase PNKD | 0 | 0.589644 |
| Q8N4J0 | CARNMT1 | Carnosine N-methyltransferase | 0.137630304 | -0.48781 |
| Q8N4Q0 | ZADH2 | Prostaglandin reductase 3 | 0.159831524 | -0.631322 |
| Q8N4V1 | MMGT1 | ER membrane protein complex subunit 5 | -1.395197233 | 0.73659 |
| Q8N543 | OGFOD1 | Prolyl 3-hydroxylase OGFOD1 | -0.206824621 | -0.293466 |
| Q8N556 | AFAP1 | Actin filament-associated protein 1 | -0.355348825 | 0.128594 |
| Q8N573 | OXR1 | Oxidation resistance protein 1 | 0.160282056 | 0.043561 |
| Q8N584 | TTC39C | Tetratricopeptide repeat protein 39C | -0.136359851 | 0.0152505 |
| Q8N5A5 | ZGPAT | Zinc finger CCCH-type with G patch domain-containing protein | -0.086215576 | -0.57018 |
| Q8N5I4 | DHRSX | Dehydrogenase/reductase SDR family member on chromosome X | -0.924166838 | -0.184508 |
| Q8N5I9 | NOPCHAP1 | NOP protein chaperone 1 | -0.332330783 | -2.01503 |
| Q8N5K1 | CISD2 | CDGSH iron-sulfur domain-containing protein 2 | -1.226884524 | 0.111135 |
| Q8N5L8 | RPP25L | Ribonuclease P protein subunit p25-like protein | 0.012348175 | -1.49585 |
| Q8N5M1 | ATPAF2 | ATP synthase mitochondrial F1 complex assembly factor 2 | 0.050296466 | 0.536268 |
| Q8N5M4 | TTC9C | Tetratricopeptide repeat protein 9C | -0.156343778 | 0.198666 |
| Q8N612 | FHIP1B | FHF complex subunit HOOK interacting protein 1B | -0.180328846 | -2.01295 |
| Q8N653 | LZTR1 | Leucine-zipper-like transcriptional regulator 1 | -0.415768623 | 0.136371 |
| Q8N668 | COMMD1 | COMM domain-containing protein 1 | -0.037527402 | -0.0956227 |
| Q8N684 | CPSF7 | Cleavage and polyadenylation specificity factor subunit 7 | -0.728048245 | 0.309284 |
| Q8N6I1 | EID2 | EP300-interacting inhibitor of differentiation 2 | 0.108780861 | 0.282662 |
| Q8N6N3 | C1orf52 | UPF0690 protein C1orf52 | -0.231049856 | -0.429957 |
| Q8N6R0 | METTL13 | eEF1A lysine and N-terminal methyltransferase | -0.321638107 | -0.384519 |
| Q8N6T3 | ARFGAP1 | ADP-ribosylation factor GTPase-activating protein 1 | -0.323711713 | -0.336596 |
| Q8N766 | EMC1 | ER membrane protein complex subunit 1 | -1.314563115 | 0.139875 |
| Q8N7H5 | PAF1 | RNA polymerase II-associated factor 1 homolog | 0.01466012 | -0.0642271 |
| Q8N806 | UBR7 | Putative E3 ubiquitin-protein ligase UBR7 | -0.119487206 | 0.546386 |
| Q8N8A6 | DDX51 | ATP-dependent RNA helicase DDX51 | -1.922018528 | 0.751674 |
| Q8N8L6 | ARL10 | ADP-ribosylation factor-like protein 10 | -0.861926715 | -0.0580924 |
| Q8N8N7 | PTGR2 | Prostaglandin reductase 2 | 0.260471423 | 0.0777569 |
| Q8N8R5 | C2orf69 | Mitochondrial protein C2orf69 | -0.211277803 | -0.310169 |
| Q8N961 | ABTB2 | Ankyrin repeat and BTB/POZ domain-containing protein 2 | 0.03831172 | -2.20851 |
| Q8N999 | C12orf29 | Uncharacterized protein C12orf29 | -0.105380535 | -0.00508404 |
| Q8N9B5 | JMY | Junction-mediating and -regulatory protein | 0.067081849 | -0.191035 |
| Q8N9V3 | WDSUB1 | WD repeat, SAM and U-box domain-containing protein 1 | -1.005379279 | -3.13409 |
| Q8NAV1 | PRPF38A | Pre-mRNA-splicing factor 38A | -0.050024668 | -0.258787 |
| Q8NB37 | GATD1 | Glutamine amidotransferase-like class 1 domain-containing protein 1 | -0.099179188 | 0.12353 |
| Q8NB46 | ANKRD52 | Serine/threonine-protein phosphatase 6 regulatory ankyrin repeat subunit C | -0.279031197 | -0.486261 |
| Q8NBF2 | NHLRC2 | NHL repeat-containing protein 2 | 0.176248709 | -0.0192887 |
| Q8NBF6 | AVL9 | Late secretory pathway protein AVL9 homolog | 0.125311454 | -0.225823 |
| Q8NBJ4 | GOLM1 | Golgi membrane protein 1 | -0.434472084 | 1.89561 |
| Q8NBJ5 | COLGALT1 | Procollagen galactosyltransferase 1 | -0.793301741 | 0.0310815 |
| Q8NBK3 | SUMF1 | Formylglycine-generating enzyme | -0.110470851 | 0.0212943 |
| Q8NBL1 | POGLUT1 | Protein O-glucosyltransferase 1 | 0.854947646 | -0.0854203 |
| Q8NBQ5 | HSD17B11 | Estradiol 17-beta-dehydrogenase 11 | -0.869359175 | -0.261738 |
| Q8NBS9 | TXNDC5 | Thioredoxin domain-containing protein 5 | -0.093376478 | 0.171792 |
| Q8NBT2 | SPC24 | Kinetochore protein Spc24 | 0.549829324 | -0.451276 |
| Q8NBU5 | ATAD1 | Outer mitochondrial transmembrane helix translocase | -1.915217956 | 0.515338 |
| Q8NBX0 | SCCPDH | Saccharopine dehydrogenase-like oxidoreductase | -0.778433164 | -0.106449 |
| Q8NC51 | SERBP1 | Plasminogen activator inhibitor 1 RNA-binding protein | -0.510001103 | -0.17782 |
| Q8NC51-3 | SERBP1 | Isoform 3 of Plasminogen activator inhibitor 1 RNA-binding protein | -0.027320306 | 0.204286 |
| Q8NC56 | LEMD2 | LEM domain-containing protein 2 | -1.704618295 | 0.14611 |
| Q8NCA5 | FAM98A | Protein FAM98A | -1.006037792 | -0.587846 |

| | | | | |
|---|---|---|---|---|
| Q8NCC3 | PLA2G15 | Phospholipase A2 group XV | -0.115765413 | -0.256202 |
| Q8NCE2 | MTMR14 | Myotubularin-related protein 14 | -0.277874947 | -0.0374845 |
| Q8NCF5 | NFATC2IP | NFATC2-interacting protein | 0.087967952 | -0.091609 |
| Q8NCW5 | NAXE | NAD(P)H-hydrate epimerase | 0.019247929 | 0.220459 |
| Q8ND04 | SMG8 | Nonsense-mediated mRNA decay factor SMG8 | -0.502524932 | -0.301258 |
| Q8ND24 | RNF214 | RING finger protein 214 | 0.045078278 | -0.0930109 |
| Q8ND56 | LSM14A | Protein LSM14 homolog A | -0.530000051 | -0.0892528 |
| Q8ND76 | CCNY | Cyclin-Y | -0.132660309 | 0.945628 |
| Q8ND90 | PNMA1 | Paraneoplastic antigen Ma1 | 0.10690848 | 0.861348 |
| Q8NDC0 | MAPK1IP1L | MAPK-interacting and spindle-stabilizing protein-like | 0.823705236 | 0 |
| Q8NDH3 | NPEPL1 | Probable aminopeptidase NPEPL1 | 0.094720125 | 0.404154 |
| Q8NDI1 | EHBP1 | EH domain-binding protein 1 | -0.285315275 | -0.23669 |
| Q8NE86 | MCU | Calcium uniporter protein, mitochondrial | 0.340681156 | -1.85112 |
| Q8NEC7 | GSTCD | Glutathione S-transferase C-terminal domain-containing protein | -0.443642775 | -0.391898 |
| Q8NEM2 | SHCBP1 | SHC SH2 domain-binding protein 1 | -0.08585159 | 0.0126276 |
| Q8NEN9 | PDZD8 | PDZ domain-containing protein 8 | 0.037704945 | 0 |
| Q8NEU8 | APPL2 | DCC-interacting protein 13-beta | 0.092590412 | 0.159115 |
| Q8NEW0 | SLC30A7 | Zinc transporter 7 | -1.678252935 | -0.357231 |
| Q8NEX9 | SDR9C7 | Short-chain dehydrogenase/reductase family 9C member 7 | -0.691347917 | -0.575681 |
| Q8NEZ2 | VPS37A | Vacuolar protein sorting-associated protein 37A | 0.162985404 | 0.062575 |
| Q8NEZ5 | FBXO22 | F-box only protein 22 | -0.099143982 | -0.158494 |
| Q8NF37 | LPCAT1 | Lysophosphatidylcholine acyltransferase 1 | -0.972411791 | -0.0848026 |
| Q8NFA0 | USP32 | Ubiquitin carboxyl-terminal hydrolase 32 | -0.456522703 | -0.988189 |
| Q8NFF5 | FLAD1 | FAD synthase | -0.203162034 | 0.302821 |
| Q8NFG4 | FLCN | Folliculin | -0.229970932 | 0.00590038 |
| Q8NFH3 | NUP43 | Nucleoporin Nup43 | -0.003652334 | -0.0600748 |
| Q8NFH4 | NUP37 | Nucleoporin Nup37 | -0.28177913 | 0.205831 |
| Q8NFH5 | NUP35 | Nucleoporin NUP35 | -0.0925831 | 0.143701 |
| Q8NFQ8 | TOR1AIP2 | Torsin-1A-interacting protein 2 | -0.656287273 | 0.212849 |
| Q8NFT8 | DNER | Delta and Notch-like epidermal growth factor-related receptor | 0.136895736 | 1.23592 |
| Q8NFV4 | ABHD11 | Protein ABHD11 | -1.002852837 | -0.481032 |
| Q8NFW8 | CMAS | N-acylneuraminate cytidylyltransferase | -1.546260516 | -1.55633 |
| Q8NG11 | TSPAN14 | Tetraspanin-14 | 0.235130628 | -0.350621 |
| Q8NG68 | TTL | Tubulin--tyrosine ligase | -0.424206972 | -0.12672 |
| Q8NHH9 | ATL2 | Atlastin-2 | -0.837557952 | -0.177192 |
| Q8NHP8 | PLBD2 | Putative phospholipase B-like 2 | -0.076628526 | 0.090325 |
| Q8NI08 | NCOA7 | Nuclear receptor coactivator 7 | -0.327437162 | 0 |
| Q8NI27 | THOC2 | THO complex subunit 2 | -1.53402102 | 1.68847 |
| Q8TAD8 | SNIP1 | Smad nuclear-interacting protein 1 | -0.031390985 | 1.30336 |
| Q8TAE8 | GADD45GIP1 | Growth arrest and DNA damage-inducible proteins-interacting protein 1 | -1.220703443 | -1.66954 |
| Q8TAF3 | WDR48 | WD repeat-containing protein 48 | -0.223127683 | -0.157892 |
| Q8TAP6 | CEP76 | Centrosomal protein of 76 kDa | -0.934509436 | -3.17612 |
| Q8TAT6 | NPLOC4 | Nuclear protein localization protein 4 homolog | -0.315594991 | 0.0738608 |
| Q8TB03 | CXorf38 | Uncharacterized protein CXorf38 | 0.188545465 | -0.5367 |
| Q8TB52 | FBXO30 | F-box only protein 30 | -0.138090372 | 0.228356 |
| Q8TB61 | SLC35B2 | Adenosine 3'-phospho 5'-phosphosulfate transporter 1 | -2.053215504 | 0.286991 |
| Q8TBC4 | UBA3 | NEDD8-activating enzyme E1 catalytic subunit | -0.038516363 | 0.0494715 |
| Q8TBE9 | NANP | N-acylneuraminate-9-phosphatase | -0.786373456 | -0.56798 |
| Q8TBQ9 | TMEM167A | Protein kish-A | -1.121756792 | 0.28861 |
| Q8TBX8 | PIP4K2C | Phosphatidylinositol 5-phosphate 4-kinase type-2 gamma | -0.11644419 | -0.0474405 |
| Q8TBZ6 | TRMT10A | tRNA methyltransferase 10 homolog A | -0.509304126 | -0.939804 |
| Q8TC07 | TBC1D15 | TBC1 domain family member 15 | 0.109271526 | 0.169331 |
| Q8TC12 | RDH11 | Retinol dehydrogenase 11 | -1.380903641 | 0.0792742 |
| Q8TCA0 | LRRC20 | Leucine-rich repeat-containing protein 20 | 0.581928333 | -0.018141 |
| Q8TCD1 | C18orf32 | UPF0729 protein C18orf32 | -0.899939617 | 0.201537 |
| Q8TCD5 | NT5C | 5'(3')-deoxyribonucleotidase, cytosolic type | -0.07708931 | 0.110206 |
| Q8TCE6 | DENND10 | DENN domain-containing protein 10 | -0.106638749 | 0.191924 |
| Q8TCF1 | ZFAND1 | AN1-type zinc finger protein 1 | -0.085384051 | 0.269948 |
| Q8TCJ2 | STT3B | Dolichyl-diphosphooligosaccharide--protein glycosyltransferase subunit STT3B | -1.688680013 | 0.219925 |
| Q8TCS8 | PNPT1 | Polyribonucleotide nucleotidyltransferase 1, mitochondrial | 0.110304912 | 0.361614 |
| Q8TCU6 | PREX1 | Phosphatidylinositol 3,4,5-trisphosphate-dependent Rac exchanger 1 protein | -0.117240349 | -0.149226 |
| Q8TCY9 | URGCP | Up-regulator of cell proliferation | -0.560959578 | -0.509798 |
| Q8TD16 | BICD2 | Protein bicaudal D homolog 2 | 0.105800867 | -0.094245 |
| Q8TD30 | GPT2 | Alanine aminotransferase 2 | -1.113720417 | -0.194049 |
| Q8TD55 | PLEKHO2 | Pleckstrin homology domain-containing family O member 2 | 0.02414163 | 0.0405108 |
| Q8TDD1 | DDX54 | ATP-dependent RNA helicase DDX54 | 0 | 1.69626 |
| Q8TDN2 | KCNV2 | Potassium voltage-gated channel subfamily V member 2 | -2.093781948 | 3.05421 |
| Q8TDN6 | BRIX1 | Ribosome biogenesis protein BRX1 homolog | -0.699990829 | -0.0358556 |

| | | | | |
|---|---|---|---|---|
| Q8TDQ7 | GNPDA2 | Glucosamine-6-phosphate isomerase 2 | -0.213357528 | 0.0421441 |
| Q8TDX7 | NEK7 | Serine/threonine-protein kinase Nek7 | -0.074999889 | -0.112851 |
| Q8TDY2 | RB1CC1 | RB1-inducible coiled-coil protein 1 | -0.822029352 | -0.562812 |
| Q8TDZ2 | MICAL1 | [F-actin]-monooxygenase MICAL1 | 0.043123563 | -0.125445 |
| Q8TEA1 | NSUN6 | tRNA (cytosine(72)-C(5))-methyltransferase NSUN6 | -2.387299776 | 1.01397 |
| Q8TEA8 | DTD1 | D-aminoacyl-tRNA deacylase 1 | -0.142061234 | -0.282141 |
| Q8TEB1 | DCAF11 | DDB1- and CUL4-associated factor 11 | -0.227305253 | -2.17306 |
| Q8TED0 | UTP15 | U3 small nucleolar RNA-associated protein 15 homolog | -0.953123887 | -0.210048 |
| Q8TED1 | GPX8 | Probable glutathione peroxidase 8 | -1.368782202 | 0.44733 |
| Q8TEQ0 | SNX29 | Sorting nexin-29 | -0.155563037 | -0.95454 |
| Q8TEQ6 | GEMIN5 | Gem-associated protein 5 | -0.070200284 | -0.152975 |
| Q8TER5 | ARHGEF40 | Rho guanine nucleotide exchange factor 40 | -1.481655121 | -2.5723 |
| Q8TEV9 | SMCR8 | Guanine nucleotide exchange protein SMCR8 | -0.313073397 | -0.288081 |
| Q8TEX9 | IPO4 | Importin-4 | -0.130771955 | -0.0102297 |
| Q8TEY7 | USP33 | Ubiquitin carboxyl-terminal hydrolase 33 | -0.647259235 | -0.324926 |
| Q8TF05 | PPP4R1 | Serine/threonine-protein phosphatase 4 regulatory subunit 1 | 0.083908876 | 0.0181885 |
| Q8TF42 | UBASH3B | Ubiquitin-associated and SH3 domain-containing protein B | -0.121539672 | 0.137412 |
| Q8TF66 | LRRC15 | Leucine-rich repeat-containing protein 15 | -0.441826026 | 0 |
| Q8WTS1 | ABHD5 | 1-acylglycerol-3-phosphate O-acyltransferase ABHD5 | -1.149223646 | 0.174217 |
| Q8WTS6 | SETD7 | Histone-lysine N-methyltransferase SETD7 | -0.049563805 | -0.0616112 |
| Q8WTT2 | NOC3L | Nucleolar complex protein 3 homolog | -0.549879869 | -0.227691 |
| Q8WTW3 | COG1 | Conserved oligomeric Golgi complex subunit 1 | 0.050060431 | -0.292076 |
| Q8WU10 | PYROXD1 | Pyridine nucleotide-disulfide oxidoreductase domain-containing protein 1 | 0.036798636 | 0.361001 |
| Q8WU76 | SCFD2 | Sec1 family domain-containing protein 2 | 0.789624373 | 0.271258 |
| Q8WU90 | ZC3H15 | Zinc finger CCCH domain-containing protein 15 | -1.041298628 | -1.45268 |
| Q8WUA2 | PPIL4 | Peptidyl-prolyl cis-trans isomerase-like 4 | -0.210804303 | -0.0693998 |
| Q8WUD1 | RAB2B | Ras-related protein Rab-2B | -1.124041716 | -0.00320371 |
| Q8WUF5 | PPP1R13L | RelA-associated inhibitor | 0.065544923 | -0.00806109 |
| Q8WUF8 | FAM172A | Cotranscriptional regulator FAM172A | -0.498032649 | -0.401617 |
| Q8WUH1 | CHURC1 | Protein Churchill | -0.029056787 | 0.566129 |
| Q8WUH2 | TGFBRAP1 | Transforming growth factor-beta receptor-associated protein 1 | -0.747979959 | -0.595976 |
| Q8WUH6 | TMEM263 | Transmembrane protein 263 | -1.136258682 | 0.0546239 |
| Q8WUI4 | HDAC7 | Histone deacetylase 7 | -0.046899557 | 0.0870616 |
| Q8WUJ0 | STYX | Serine/threonine/tyrosine-interacting protein | 0.835309029 | 0.208268 |
| Q8WUM0 | NUP133 | Nuclear pore complex protein Nup133 | -0.072190921 | 0.138424 |
| Q8WUM4 | PDCD6IP | Programmed cell death 6-interacting protein | 0.130388419 | 0.0389703 |
| Q8WUP2 | FBLIM1 | Filamin-binding LIM protein 1 | -0.011758566 | 0.193168 |
| Q8WUQ7 | CACTIN | Cactin | -0.090356429 | -0.45244 |
| Q8WUX2 | CHAC2 | Glutathione-specific gamma-glutamylcyclotransferase 2 | -0.160560608 | -0.26708 |
| Q8WUX9 | CHMP7 | Charged multivesicular body protein 7 | 0.169830958 | -0.301592 |
| Q8WV24 | PHLDA1 | Pleckstrin homology-like domain family A member 1 | -1.147377491 | -0.684943 |
| Q8WV41 | SNX33 | Sorting nexin-33 | -0.003613234 | 0.259388 |
| Q8WVC0 | LEO1 | RNA polymerase-associated protein LEO1 | -0.056810459 | -0.196454 |
| Q8WVD3 | RNF138 | E3 ubiquitin-protein ligase RNF138 | 0.096244017 | 0.110575 |
| Q8WVF1-3 | OSCP1 | Isoform 3 of Protein OSCP1 | -0.388451576 | -0.111132 |
| Q8WVJ2 | NUDCD2 | NudC domain-containing protein 2 | 0.103148301 | 0.293344 |
| Q8WVQ1 | CANT1 | Soluble calcium-activated nucleotidase 1 | -0.99463733 | -1.20131 |
| Q8WVT3 | TRAPPC12 | Trafficking protein particle complex subunit 12 | -0.616470337 | -0.749768 |
| Q8WVV4 | POF1B | Protein POF1B | -0.959672133 | -0.302187 |
| Q8WVX9 | FAR1 | Fatty acyl-CoA reductase 1 | 0.711204211 | -0.0864449 |
| Q8WVY7 | UBLCP1 | Ubiquitin-like domain-containing CTD phosphatase 1 | 0.08390824 | 0.0976855 |
| Q8WW12 | PCNP | PEST proteolytic signal-containing nuclear protein | 0.180485249 | -0.0533381 |
| Q8WW59 | SPRYD4 | SPRY domain-containing protein 4 | -0.677228769 | -0.5299 |
| Q8WWH5 | TRUB1 | Probable tRNA pseudouridine synthase 1 | -0.173252424 | -0.0445213 |
| Q8WWK9 | CKAP2 | Cytoskeleton-associated protein 2 | 0.402745167 | -0.600365 |
| Q8WWM7 | ATXN2L | Ataxin-2-like protein | -0.218595982 | -0.0592445 |
| Q8WWN8 | ARAP3 | Arf-GAP with Rho-GAP domain, ANK repeat and PH domain-containing protein 3 | -0.139939149 | -0.714068 |
| Q8WWY3 | PRPF31 | U4/U6 small nuclear ribonucleoprotein Prp31 | -0.093157291 | -0.0163333 |
| Q8WX93 | PALLD | Palladin | 0.103240728 | -0.0500094 |
| Q8WXA9-2 | SREK1 | Isoform 2 of Splicing regulatory glutamine/lysine-rich protein 1 | -0.234384537 | -1.31656 |
| Q8WXD5 | GEMIN6 | Gem-associated protein 6 | -0.378006538 | 0.18329 |
| Q8WXF1 | PSPC1 | Paraspeckle component 1 | -1.564844767 | 0.12692 |
| Q8WXX5 | DNAJC9 | DnaJ homolog subfamily C member 9 | 0.681837638 | 0.156057 |
| Q8WYL5 | SSH1 | Protein phosphatase Slingshot homolog 1 | 0.122142156 | -0.423715 |
| Q8WYP5 | AHCTF1 | Protein ELYS | 0.062211434 | -0.190405 |
| Q8WZ82 | OVCA2 | Esterase OVCA2 | -0.210163434 | 0.518499 |
| Q8WZA0 | LZIC | Protein LZIC | 0.143263261 | 0.282781 |
| Q8WZA9 | IRGQ | Immunity-related GTPase family Q protein | 0.174556096 | 0.126731 |
| Q92484 | SMPDL3A | Acid sphingomyelinase-like phosphodiesterase 3a | -0.076056639 | 3.04767 |

| | | | | |
|---|---|---|---|---|
| Q92503 | SEC14L1 | SEC14-like protein 1 | 0.119785627 | -4.98498 |
| Q92520 | FAM3C | Protein FAM3C | -2.213010073 | 0.738441 |
| Q92522 | H1-10 | Histone H1.10 | -1.216283798 | 0.00421079 |
| Q92526 | CCT6B | T-complex protein 1 subunit zeta-2 | -0.714054187 | -0.529607 |
| Q92536 | SLC7A6 | Y+L amino acid transporter 2 | 1.099077304 | 3.54129 |
| Q92541 | RTF1 | RNA polymerase-associated protein RTF1 homolog | -0.00611194 | -0.0457738 |
| Q92542 | NCSTN | Nicastrin | -0.602225939 | 0.275411 |
| Q92551 | IP6K1 | Inositol hexakisphosphate kinase 1 | -0.069681565 | -0.354168 |
| Q92572 | AP3S1 | AP-3 complex subunit sigma-1 | 0.24276797 | 0.117261 |
| Q92575 | UBXN4 | UBX domain-containing protein 4 | -1.109907548 | -0.127942 |
| Q92597 | NDRG1 | Protein NDRG1 | -0.079199314 | -0.0264746 |
| Q92598 | HSPH1 | Heat shock protein 105 kDa | 0.042265256 | 0.0157458 |
| Q92600 | CNOT9 | CCR4-NOT transcription complex subunit 9 | -2.045907736 | -1.9963 |
| Q92609 | TBC1D5 | TBC1 domain family member 5 | 0.273030281 | -0.00598621 |
| Q92615 | LARP4B | La-related protein 4B | -0.487446706 | -0.417955 |
| Q92616 | GCN1 | eIF-2-alpha kinase activator GCN1 | -0.155939261 | -0.355537 |
| Q92620 | DHX38 | Pre-mRNA-splicing factor ATP-dependent RNA helicase PRP16 | -0.188584169 | 0.020956 |
| Q92621 | NUP205 | Nuclear pore complex protein Nup205 | -0.133236488 | 0.00806268 |
| Q92626 | PXDN | Peroxidasin homolog | -0.823264837 | -0.0124779 |
| Q92636 | NSMAF | Protein FAN | 0.394756953 | 0.108714 |
| Q92665 | MRPS31 | 28S ribosomal protein S31, mitochondrial | -1.861821651 | -0.924258 |
| Q92688 | ANP32B | Acidic leucine-rich nuclear phosphoprotein 32 family member B | -0.157132467 | -0.155212 |
| Q92692 | NECTIN2 | Nectin-2 | -0.885267417 | 0.264081 |
| Q92696 | RABGGTA | Geranylgeranyl transferase type-2 subunit alpha | 0.094364484 | 0.0469672 |
| Q92733 | PRCC | Proline-rich protein PRCC | -0.002225161 | -0.213214 |
| Q92747 | ARPC1A | Actin-related protein 2/3 complex subunit 1A | -0.297470649 | -0.685572 |
| Q92759 | GTF2H4 | General transcription factor IIH subunit 4 | -0.483997186 | 0.041976 |
| Q92769 | HDAC2 | Histone deacetylase 2 | -0.314136346 | 0.0149657 |
| Q92777 | SYN2 | Synapsin-2 | 0.641118447 | -1.53718 |
| Q92783 | STAM | Signal transducing adapter molecule 1 | 0.167165915 | 0.0330741 |
| Q92797 | SYMPK | Symplekin | -0.754155477 | -0.8373 |
| Q92804 | TAF15 | TATA-binding protein-associated factor 2N | -0.492220163 | -0.0744559 |
| Q92805 | GOLGA1 | Golgin subfamily A member 1 | -0.225269397 | -0.0737178 |
| Q92820 | GGH | Gamma-glutamyl hydrolase | -0.20577081 | -0.117023 |
| Q92845 | KIFAP3 | Kinesin-associated protein 3 | -0.277157863 | -0.639755 |
| Q92871 | PMM1 | Phosphomannomutase 1 | 0.065091292 | 0.191497 |
| Q92878 | RAD50 | DNA repair protein RAD50 | 0.055395047 | 0.162587 |
| Q92882 | OSTF1 | Osteoclast-stimulating factor 1 | -0.097111702 | -0.186237 |
| Q92889 | ERCC4 | DNA repair endonuclease XPF | -1.703423897 | 0.19471 |
| Q92890 | UFD1 | Ubiquitin recognition factor in ER-associated degradation protein 1 | -0.291941961 | 0.125065 |
| Q92896 | GLG1 | Golgi apparatus protein 1 | -0.974354506 | 0.11143 |
| Q92900 | UPF1 | Regulator of nonsense transcripts 1 | -0.415698131 | 0.118947 |
| Q92905 | COPS5 | COP9 signalosome complex subunit 5 | 0.097917318 | -0.260874 |
| Q92917 | GPKOW | G-patch domain and KOW motifs-containing protein | -0.078668674 | -0.0297546 |
| Q92922 | SMARCC1 | SWI/SNF complex subunit SMARCC1 | -1.45051686 | -0.442068 |
| Q92930 | RAB8B | Ras-related protein Rab-8B | -0.813080231 | 0.0254987 |
| Q92934 | BAD | Bcl2-associated agonist of cell death | -1.202490966 | -0.336194 |
| Q92947 | GCDH | Glutaryl-CoA dehydrogenase, mitochondrial | -1.639229695 | -0.985007 |
| Q92968 | PEX13 | Peroxisomal membrane protein PEX13 | -1.060324669 | -0.0618331 |
| Q92973 | TNPO1 | Transportin-1 | -0.13601017 | -0.284774 |
| Q92979 | EMG1 | Ribosomal RNA small subunit methyltransferase NEP1 | -0.51144592 | 0.302433 |
| Q92989 | CLP1 | Polyribonucleotide 5'-hydroxyl-kinase Clp1 | -0.003201882 | -0.127704 |
| Q92990 | GLMN | Glomulin | -0.045734326 | -0.562065 |
| Q92997 | DVL3 | Segment polarity protein dishevelled homolog DVL-3 | 0.330227375 | 0 |
| Q93008 | USP9X | Probable ubiquitin carboxyl-terminal hydrolase FAF-X | -0.014998277 | -0.134569 |
| Q93034 | CUL5 | Cullin-5 | 0.170129538 | -0.0389338 |
| Q93052 | LPP | Lipoma-preferred partner | 0.015361389 | 0.0583216 |
| Q93083 | MT1L | Metallothionein-1L | -0.498512824 | 0 |
| Q969E2 | SCAMP4 | Secretory carrier-associated membrane protein 4 | -3.347274502 | -1.63509 |
| Q969E4 | TCEAL3 | Transcription elongation factor A protein-like 3 | 0.1106565 | 0.24677 |
| Q969E8 | TSR2 | Pre-rRNA-processing protein TSR2 homolog | -0.860836744 | -0.0527604 |
| Q969F9 | HPS3 | Hermansky-Pudlak syndrome 3 protein | -0.210001071 | 0.289567 |
| Q969G6 | RFK | Riboflavin kinase | -0.488937855 | -0.139843 |
| Q969H6 | POP5 | Ribonuclease P/MRP protein subunit POP5 | -0.721010764 | -0.244843 |
| Q969H8 | MYDGF | Myeloid-derived growth factor | 0.170330842 | 0.117278 |
| Q969L2 | MAL2 | Protein MAL2 | -0.115311861 | -1.61927 |
| Q969M3 | YIPF5 | Protein YIPF5 | -1.968525728 | 0.449034 |
| Q969Q0 | RPL36AL | 60S ribosomal protein L36a-like | -1.432491461 | -0.157675 |
| Q969Q5 | RAB24 | Ras-related protein Rab-24 | -0.874366442 | 0.130901 |
| Q969R8 | ITFG2 | KICSTOR complex protein ITFG2 | 0.442431609 | -0.190939 |

| | | | | |
|---|---|---|---|---|
| Q969S2 | NEIL2 | Endonuclease 8-like 2 | 0.093884865 | -0.0387837 |
| Q969S3 | ZNF622 | Zinc finger protein 622 | 0.036566655 | 0.118593 |
| Q969S9 | GFM2 | Ribosome-releasing factor 2, mitochondrial | -0.679956436 | 0.116071 |
| Q969T7 | NT5C3B | 7-methylguanosine phosphate-specific 5'-nucleotidase | 0.133633137 | -0.0544246 |
| Q969V3 | NCLN | Nicalin | -1.547086 | 0.612548 |
| Q969X5 | ERGIC1 | Endoplasmic reticulum-Golgi intermediate compartment protein 1 | -1.35934782 | 0.176871 |
| Q969X6 | UTP4 | U3 small nucleolar RNA-associated protein 4 homolog | -0.914940039 | 0.299511 |
| Q969Z0 | TBRG4 | FAST kinase domain-containing protein 4 | -0.831407309 | -0.368449 |
| Q96A19 | CCDC102A | Coiled-coil domain-containing protein 102A | -0.824945211 | -0.796334 |
| Q96A33 | CCDC47 | PAT complex subunit CCDC47 | -1.093597889 | 0.574865 |
| Q96A35 | MRPL24 | 39S ribosomal protein L24, mitochondrial | -0.862170776 | -0.57688 |
| Q96A49 | SYAP1 | Synapse-associated protein 1 | 0.181236267 | -0.0199089 |
| Q96A65 | EXOC4 | Exocyst complex component 4 | -0.461616278 | -0.0452271 |
| Q96AB3 | ISOC2 | Isochorismatase domain-containing protein 2 | -0.172642787 | 0.814888 |
| Q96AB6 | NTAN1 | Protein N-terminal asparagine amidohydrolase | 0.078204075 | 0.147875 |
| Q96AC1 | FERMT2 | Fermitin family homolog 2 | 0.016961177 | -0.0822608 |
| Q96AD5 | PNPLA2 | Patatin-like phospholipase domain-containing protein 2 | -0.945903142 | 0.0343984 |
| Q96AE4-2 | FUBP1 | Isoform 2 of Far upstream element-binding protein 1 | -0.309916019 | 0.368569 |
| Q96AG4 | LRRC59 | Leucine-rich repeat-containing protein 59 | -1.069399198 | 0.486785 |
| Q96AH0 | NABP1 | SOSS complex subunit B2 | -0.556007067 | 0.0480935 |
| Q96AJ9 | VTI1A | Vesicle transport through interaction with t-SNAREs homolog 1A | -0.440556924 | -0.76651 |
| Q96AQ6 | PBXIP1 | Pre-B-cell leukemia transcription factor-interacting protein 1 | -0.763530334 | -0.80826 |
| Q96AT1 | KIAA1143 | Uncharacterized protein KIAA1143 | -0.100228707 | -0.235243 |
| Q96AT9 | RPE | Ribulose-phosphate 3-epimerase | 0.305715481 | -0.760064 |
| Q96AY3 | FKBP10 | Peptidyl-prolyl cis-trans isomerase FKBP10 | -0.01399676 | 0.204166 |
| Q96B23-2 | C18orf25 | Isoform 2 of Uncharacterized protein C18orf25 | -0.536386569 | 0.957551 |
| Q96B26 | EXOSC8 | Exosome complex component RRP43 | -0.348293781 | -0.110884 |
| Q96B36 | AKT1S1 | Proline-rich AKT1 substrate 1 | -0.02096041 | 0.54971 |
| Q96B96-2 | LDAF1 | Isoform 2 of Lipid droplet assembly factor 1 | -0.702586174 | 0.0150019 |
| Q96B97 | SH3KBP1 | SH3 domain-containing kinase-binding protein 1 | -0.859920422 | 0.736014 |
| Q96BH1 | RNF25 | E3 ubiquitin-protein ligase RNF25 | -0.335109552 | 0.0483815 |
| Q96BJ3 | AIDA | Axin interactor, dorsalization-associated protein | 0.026254336 | -0.0864741 |
| Q96BM9 | ARL8A | ADP-ribosylation factor-like protein 8A | -0.761353254 | -0.0288397 |
| Q96BN8 | OTULIN | Ubiquitin thioesterase otulin | -0.035388549 | -0.156364 |
| Q96BP3 | PPWD1 | Peptidylprolyl isomerase domain and WD repeat-containing protein 1 | -0.415662527 | -0.258887 |
| Q96BR5 | COA7 | Cytochrome c oxidase assembly factor 7 | -0.341388464 | 0.146428 |
| Q96BT7 | ALKBH8 | Alkylated DNA repair protein alkB homolog 8 | -0.832399607 | -1.19998 |
| Q96BX8 | MOB3A | MOB kinase activator 3A | 0.074370543 | -0.198738 |
| Q96BY7 | ATG2B | Autophagy-related protein 2 homolog B | -0.068806092 | -0.493299 |
| Q96C01 | FAM136A | Protein FAM136A | -0.281827609 | 0.440116 |
| Q96C19 | EFHD2 | EF-hand domain-containing protein D2 | 0.128219922 | -0.0632362 |
| Q96C23 | GALM | Galactose mutarotase | 0.059280475 | -0.636475 |
| Q96C86 | DCPS | m7GpppX diphosphatase | 0.243201097 | -0.119621 |
| Q96C90 | PPP1R14B | Protein phosphatase 1 regulatory subunit 14B | 0.618846258 | 0.0814966 |
| Q96C92 | ENTR1 | Endosome-associated-trafficking regulator 1 | -1.323943535 | 0 |
| Q96CN4 | EVI5L | EVI5-like protein | 5.515758758 | 0 |
| Q96CN7 | ISOC1 | Isochorismatase domain-containing protein 1 | 0.087821802 | 0.0378246 |
| Q96CN9 | GCC1 | GRIP and coiled-coil domain-containing protein 1 | -0.113895893 | -0.282777 |
| Q96CS2 | HAUS1 | HAUS augmin-like complex subunit 1 | -0.161672115 | -0.331754 |
| Q96CS3 | FAF2 | FAS-associated factor 2 | -0.943090916 | -0.0398127 |
| Q96CT7 | CCDC124 | Coiled-coil domain-containing protein 124 | -0.093349775 | 0.0704352 |
| Q96CV9 | OPTN | Optineurin | 0.318546613 | 0.0965109 |
| Q96CW1 | AP2M1 | AP-2 complex subunit mu | -0.417080005 | -0.145551 |
| Q96CW5 | TUBGCP3 | Gamma-tubulin complex component 3 | -0.069343805 | 2.5039 |
| Q96CW6 | SLC7A6OS | Probable RNA polymerase II nuclear localization protein SLC7A6OS | -0.705774387 | 0.401519 |
| Q96CX2 | KCTD12 | BTB/POZ domain-containing protein KCTD12 | -0.467094421 | -2.66966 |
| Q96D09 | GPRASP2 | G-protein coupled receptor-associated sorting protein 2 | -0.206004302 | -0.627673 |
| Q96D15 | RCN3 | Reticulocalbin-3 | -0.043510993 | 0.693902 |
| Q96D71 | REPS1 | RalBP1-associated Eps domain-containing protein 1 | 0.030451218 | -0.192654 |
| Q96DA6 | DNAJC19 | Mitochondrial import inner membrane translocase subunit TIM14 | -2.030388991 | 4.41351 |
| Q96DB5 | RMDN1 | Regulator of microtubule dynamics protein 1 | -0.360103051 | 0.0603952 |
| Q96DC9 | OTUB2 | Ubiquitin thioesterase OTUB2 | -0.203395685 | -0.0906353 |
| Q96DE0 | NUDT16 | U8 snoRNA-decapping enzyme | -0.170315107 | 0.063736 |
| Q96DG6 | CMBL | Carboxymethylenebutenolidase homolog | 0.137766679 | 0.104179 |
| Q96DI7 | SNRNP40 | U5 small nuclear ribonucleoprotein 40 kDa protein | -0.293347677 | -0.00545216 |
| Q96DV4 | MRPL38 | 39S ribosomal protein L38, mitochondrial | -2.719620069 | 1.09237 |
| Q96DZ1 | ERLEC1 | Endoplasmic reticulum lectin 1 | -1.132149696 | 0.174343 |
| Q96E09 | PABIR1 | PPP2R1A-PPP2R2A-interacting phosphatase regulator 1 | -0.171396017 | 0.0194699 |
| Q96E11 | MRRF | Ribosome-recycling factor, mitochondrial | -0.975357771 | -1.2096 |
| Q96E22 | NUS1 | Dehydrodolichyl diphosphate synthase complex subunit NUS1 | -1.222371976 | -0.128744 |

| | | | | |
|---|---|---|---|---|
| Q96EA4 | SPDL1 | Protein Spindly | -0.649742126 | -0.122272 |
| Q96EB6 | SIRT1 | NAD-dependent protein deacetylase sirtuin-1 | -0.194611549 | 0.0552794 |
| Q96EE3 | SEH1L | Nucleoporin SEH1 | -0.560352802 | -0.0813046 |
| Q96EI5 | TCEAL4 | Transcription elongation factor A protein-like 4 | 0.222829421 | 0.287462 |
| Q96EK6 | GNPNAT1 | Glucosamine 6-phosphate N-acetyltransferase | -0.380205154 | -0.0840877 |
| Q96EK9 | KTI12 | Protein KTI12 homolog | -0.305504084 | -0.384571 |
| Q96EL3 | MRPL53 | 39S ribosomal protein L53, mitochondrial | 0.4888436 | 2.96854 |
| Q96EM0 | L3HYPDH | Trans-3-hydroxy-L-proline dehydratase | 0.23193121 | -0.504802 |
| Q96EN8 | MOCOS | Molybdenum cofactor sulfurase | 0.400015116 | -3.7202 |
| Q96EP0 | RNF31 | E3 ubiquitin-protein ligase RNF31 | 0.290743351 | 0.195948 |
| Q96EQ0 | SGTB | Small glutamine-rich tetratricopeptide repeat-containing protein beta | 0.116197189 | 0.51366 |
| Q96ER3 | SAAL1 | Protein SAAL1 | -0.707237959 | 0.256877 |
| Q96EU7 | C1GALT1C1 | C1GALT1-specific chaperone 1 | 0.222908417 | 0 |
| Q96EV8 | DTNBP1 | Dysbindin | -0.498200893 | -0.947183 |
| Q96EW2 | HSPBAP1 | HSPB1-associated protein 1 | -0.110775312 | 0.95601 |
| Q96EX3 | DYNC2I2 | Cytoplasmic dynein 2 intermediate chain 2 | -0.44270134 | -0.539116 |
| Q96EY1 | DNAJA3 | DnaJ homolog subfamily A member 3, mitochondrial | -0.988022089 | 0.062569 |
| Q96EY7 | PTCD3 | Pentatricopeptide repeat domain-containing protein 3, mitochondrial | -1.235944192 | 0.0598127 |
| Q96EY8 | MMAB | Corrinoid adenosyltransferase | -1.521674633 | -0.414632 |
| Q96F24 | NRBF2 | Nuclear receptor-binding factor 2 | -0.069576899 | 0.426833 |
| Q96F44 | TRIM11 | E3 ubiquitin-protein ligase TRIM11 | -0.820401271 | 0.0301215 |
| Q96F63 | CCDC97 | Coiled-coil domain-containing protein 97 | -0.796225627 | 1.26895 |
| Q96F85 | CNRIP1 | CB1 cannabinoid receptor-interacting protein 1 | 0.116979917 | -0.117056 |
| Q96F86 | EDC3 | Enhancer of mRNA-decapping protein 3 | -1.077263395 | 0.0650228 |
| Q96FN4 | CPNE2 | Copine-2 | -0.036159356 | 0.0114257 |
| Q96FQ6 | S100A16 | Protein S100-A16 | -0.182727893 | 0.115724 |
| Q96FS4 | SIPA1 | Signal-induced proliferation-associated protein 1 | -0.068840981 | -0.18268 |
| Q96FX7 | TRMT61A | tRNA (adenine(58)-N(1))-methyltransferase catalytic subunit TRMT61A | -0.155399164 | 0.0323029 |
| Q96FZ2 | HMCES | Abasic site processing protein HMCES | -0.818127076 | -0.0137933 |
| Q96FZ7 | CHMP6 | Charged multivesicular body protein 6 | -0.996881882 | 0.166001 |
| Q96G03 | PGM2 | Phosphoglucomutase-2 | 0.058291435 | -0.0702432 |
| Q96G28 | CFAP36 | Cilia- and flagella-associated protein 36 | -0.055783908 | -0.539661 |
| Q96G46 | DUS3L | tRNA-dihydrouridine(47) synthase [NAD(P)(+)]-like | -0.161849658 | -0.593911 |
| Q96G61 | NUDT11 | Diphosphoinositol polyphosphate phosphohydrolase 3-beta | -0.004918496 | 0.00767072 |
| Q96GA3 | LTV1 | Protein LTV1 homolog | -0.251089493 | -0.286256 |
| Q96GA7 | SDSL | Serine dehydratase-like | -0.032068094 | -0.161193 |
| Q96GD0 | PDXP | Chronophin | 0.118473291 | -0.196682 |
| Q96GG9 | DCUN1D1 | DCN1-like protein 1 | 0.107208252 | 0.0286849 |
| Q96GK7 | FAHD2A | Fumarylacetoacetate hydrolase domain-containing protein 2A | -0.071804285 | 0.127683 |
| Q96GM8 | TOE1 | Target of EGR1 protein 1 | -0.740000963 | 0.101936 |
| Q96GS4 | BORCS6 | BLOC-1-related complex subunit 6 | 0.61017712 | -0.274329 |
| Q96GX5 | MASTL | Serine/threonine-protein kinase greatwall | 0.855216901 | 0.153065 |
| Q96GX9 | APIP | Methylthioribulose-1-phosphate dehydratase | 0.182436546 | 0.0606724 |
| Q96H20 | SNF8 | Vacuolar-sorting protein SNF8 | -0.276998917 | -0.365079 |
| Q96H79 | ZC3HAV1L | Zinc finger CCCH-type antiviral protein 1-like | -0.105983257 | 1.09631 |
| Q96HA7 | TONSL | Tonsoku-like protein | -0.428586801 | -0.444027 |
| Q96HA8 | NTAQ1 | Protein N-terminal glutamine amidohydrolase | 0.564982891 | -0.254681 |
| Q96HC4 | PDLIM5 | PDZ and LIM domain protein 5 | -0.136631489 | 0.0697864 |
| Q96HE7 | ERO1A | ERO1-like protein alpha | -0.166448911 | 0.0610326 |
| Q96HE9 | PRR11 | Proline-rich protein 11 | -1.141007662 | -0.0810239 |
| Q96HJ9 | FMC1 | Protein FMC1 homolog | -1.589346965 | 0.342517 |
| Q96HR8 | NAF1 | H/ACA ribonucleoprotein complex non-core subunit NAF1 | -0.141296387 | 0.224843 |
| Q96HS1 | PGAM5 | Serine/threonine-protein phosphatase PGAM5, mitochondrial | -1.297465404 | 0.470976 |
| Q96HY6 | DDRGK1 | DDRGK domain-containing protein 1 | -1.373369376 | 0.499331 |
| Q96I24 | FUBP3 | Far upstream element-binding protein 3 | -0.66030256 | 0.271049 |
| Q96I25 | RBM17 | Splicing factor 45 | -0.172004143 | 0.198371 |
| Q96I51 | RCC1L | RCC1-like G exchanging factor-like protein | -0.879998883 | 1.07705 |
| Q96I99 | SUCLG2 | Succinate--CoA ligase [GDP-forming] subunit beta, mitochondrial | -0.245109717 | 0.237646 |
| Q96IF1 | AJUBA | LIM domain-containing protein ajuba | -0.776275237 | 0.464537 |
| Q96II8 | LRCH3 | DISP complex protein LRCH3 | -0.695621332 | 0.138202 |
| Q96IJ6 | GMPPA | Mannose-1-phosphate guanyltransferase alpha | 0.161136389 | -0.0321608 |
| Q96IU4 | ABHD14B | Protein ABHD14B | 0.209492922 | -0.0869484 |
| Q96IX5 | ATP5MK | ATP synthase membrane subunit K, mitochondrial | -2.184073766 | 0.354064 |
| Q96IY1 | NSL1 | Kinetochore-associated protein NSL1 homolog | -0.371816953 | -0.646458 |
| Q96IZ0 | PAWR | PRKC apoptosis WT1 regulator protein | -0.566612959 | 0.084967 |
| Q96IZ6 | METTL2A | tRNA N(3)-methylcytidine methyltransferase METTL2A | 0.063955625 | 0.273662 |
| Q96J01 | THOC3 | THO complex subunit 3 | -0.100126584 | 0.0270233 |
| Q96J02 | ITCH | E3 ubiquitin-protein ligase Itchy homolog | -0.329911073 | 0.0757929 |

| | | | | |
|---|---|---|---|---|
| Q96J92 | WNK4 | Serine/threonine-protein kinase WNK4 | 0.201802651 | 0.308414 |
| Q96JA1 | LRIG1 | Leucine-rich repeats and immunoglobulin-like domains protein 1 | -0.125144482 | -1.06798 |
| Q96JA3 | PLEKHA8 | Pleckstrin homology domain-containing family A member 8 | 1.198927244 | 0 |
| Q96JB2 | COG3 | Conserved oligomeric Golgi complex subunit 3 | -0.164406061 | 0.398585 |
| Q96JB5 | CDK5RAP3 | CDK5 regulatory subunit-associated protein 3 | -1.494071802 | -0.115163 |
| Q96JC9 | EAF1 | ELL-associated factor 1 | -1.668144902 | 0 |
| Q96JG6 | VPS50 | Syndetin | 0.185121218 | -0.114398 |
| Q96JH7 | VCPIP1 | Deubiquitinating protein VCPIP1 | 0.052451054 | -0.0451059 |
| Q96JJ7 | TMX3 | Protein disulfide-isomerase TMX3 | -1.244501909 | 0.372233 |
| Q96JM3 | CHAMP1 | Chromosome alignment-maintaining phosphoprotein 1 | -2.994889657 | -2.45862 |
| Q96JY6 | PDLIM2 | PDZ and LIM domain protein 2 | -0.081584533 | 0.407614 |
| Q96K17 | BTF3L4 | Transcription factor BTF3 homolog 4 | 0.085891088 | -0.446674 |
| Q96K37 | SLC35E1 | Solute carrier family 35 member E1 | -1.884236733 | 0.961026 |
| Q96K76-4 | USP47 | Isoform 4 of Ubiquitin carboxyl-terminal hydrolase 47 | -0.027362108 | -0.136105 |
| Q96KB5 | PBK | Lymphokine-activated killer T-cell-originated protein kinase | 0.146929423 | -0.0326646 |
| Q96KC2 | ARL5B | ADP-ribosylation factor-like protein 5B | 0.356037855 | 0.428886 |
| Q96KC8 | DNAJC1 | DnaJ homolog subfamily C member 1 | -0.875586669 | 1.89933 |
| Q96KP4 | CNDP2 | Cytosolic non-specific dipeptidase | 0.074776808 | 0.0847365 |
| Q96KR1 | ZFR | Zinc finger RNA-binding protein | -1.191695134 | 0.854587 |
| Q96L46 | CAPNS2 | Calpain small subunit 2 | -0.362900893 | 0 |
| Q96L92 | SNX27 | Sorting nexin-27 | 0.262363911 | -0.275288 |
| Q96L93-2 | KIF16B | Isoform 2 of Kinesin-like protein KIF16B | -0.697657665 | -0.177035 |
| Q96LB3 | IFT74 | Intraflagellar transport protein 74 homolog | -0.011553605 | 0.390216 |
| Q96LD4 | TRIM47 | E3 ubiquitin-protein ligase TRIM47 | -0.07411019 | -0.357935 |
| Q96LD8 | SENP8 | Sentrin-specific protease 8 | 0.469609698 | 3.88698 |
| Q96LJ7 | DHRS1 | Dehydrogenase/reductase SDR family member 1 | -1.033869425 | 0.263269 |
| Q96M27 | PRRC1 | Protein PRRC1 | -0.072346926 | -0.211031 |
| Q96ME1-4 | FBXL18 | Isoform 4 of F-box/LRR-repeat protein 18 | -0.259823879 | -0.62929 |
| Q96MG7 | NSMCE3 | Non-structural maintenance of chromosomes element 3 homolog | 1.2496291 | 0.480316 |
| Q96MW1 | CCDC43 | Coiled-coil domain-containing protein 43 | 0.258663177 | -0.315559 |
| Q96MX6 | DNAAF10 | Dynein axonemal assembly factor 10 | -0.38705039 | -1.10514 |
| Q96N66 | MBOAT7 | Lysophospholipid acyltransferase 7 | -1.534020503 | 0.365586 |
| Q96N67 | DOCK7 | Dedicator of cytokinesis protein 7 | -0.429154714 | 0.0769978 |
| Q96NC0 | ZMAT2 | Zinc finger matrin-type protein 2 | 0.021143516 | -0.0331955 |
| Q96NE9 | FRMD6 | FERM domain-containing protein 6 | -0.828862429 | -0.0868168 |
| Q96NY7 | CLIC6 | Chloride intracellular channel protein 6 | -0.276147525 | -0.248322 |
| Q96P16 | RPRD1A | Regulation of nuclear pre-mRNA domain-containing protein 1A | 0.059704542 | 0.0129242 |
| Q96P47 | AGAP3 | Arf-GAP with GTPase, ANK repeat and PH domain-containing protein 3 | -0.562613885 | -0.3889 |
| Q96P48 | ARAP1 | Arf-GAP with Rho-GAP domain, ANK repeat and PH domain-containing protein 1 | 0.203690449 | 0.0547326 |
| Q96P53 | WDFY2 | WD repeat and FYVE domain-containing protein 2 | -0.219993512 | 0.114712 |
| Q96P63 | SERPINB12 | Serpin B12 | -0.180172761 | -0.0912905 |
| Q96P70 | IPO9 | Importin-9 | -0.029187997 | -0.202375 |
| Q96PD2 | DCBLD2 | Discoidin, CUB and LCCL domain-containing protein 2 | -2.253601948 | 0.705009 |
| Q96PE3 | INPP4A | Inositol polyphosphate-4-phosphatase type I A | 0.106824954 | -0.344177 |
| Q96PK6 | RBM14 | RNA-binding protein 14 | -0.640344938 | 0.44925 |
| Q96PM5 | RCHY1 | RING finger and CHY zinc finger domain-containing protein 1 | -1.139765541 | 0 |
| Q96PP9 | GBP4 | Guanylate-binding protein 4 | 0.661436717 | 0 |
| Q96PU8 | QKI | Protein quaking | -0.614966631 | 0.139778 |
| Q96PY5 | FMNL2 | Formin-like protein 2 | -0.866312544 | 0 |
| Q96PZ0 | PUS7 | Pseudouridylate synthase 7 homolog | 0.263683319 | 0.254245 |
| Q96Q11 | TRNT1 | CCA tRNA nucleotidyltransferase 1, mitochondrial | -0.005713065 | -0.302434 |
| Q96QA5 | GSDMA | Gasdermin-A | -0.278796434 | -0.324778 |
| Q96QB1-6 | DLC1 | Isoform 6 of Rho GTPase-activating protein 7 | -0.190475543 | 0.0900065 |
| Q96QC0 | PPP1R10 | Serine/threonine-protein phosphatase 1 regulatory subunit 10 | 0.016753912 | -0.123044 |
| Q96QD8 | SLC38A2 | Sodium-coupled neutral amino acid transporter 2 | -1.69972229 | 0.158066 |
| Q96QG7 | MTMR9 | Myotubularin-related protein 9 | -0.263823827 | -0.10419 |
| Q96QK1 | VPS35 | Vacuolar protein sorting-associated protein 35 | -0.044477145 | -0.291955 |
| Q96QR8 | PURB | Transcriptional activator protein Pur-beta | -0.310753743 | -0.0798241 |
| Q96QU8 | XPO6 | Exportin-6 | 0.033869823 | -0.0874914 |
| Q96R05 | RBP7 | Retinoid-binding protein 7 | 0.085106929 | 0.863419 |
| Q96R06 | SPAG5 | Sperm-associated antigen 5 | -0.584840377 | -0.0517985 |
| Q96RE7 | NACC1 | Nucleus accumbens-associated protein 1 | 0.205539862 | 0.337281 |
| Q96RF0 | SNX18 | Sorting nexin-18 | 0.156836033 | -0.173673 |
| Q96RG2 | PASK | PAS domain-containing serine/threonine-protein kinase | 0.697945197 | -1.32852 |
| Q96RL7 | VPS13A | Vacuolar protein sorting-associated protein 13A | -0.703153849 | 0.085043 |
| Q96RM1 | SPRR2F | Small proline-rich protein 2F | -2.881013493 | 0 |
| Q96RP9 | GFM1 | Elongation factor G, mitochondrial | -0.214228153 | -0.00587908 |
| Q96RR4 | CAMKK2 | Calcium/calmodulin-dependent protein kinase kinase 2 | -0.664812406 | -0.150496 |
| Q96RS0 | TGS1 | Trimethylguanosine synthase | -0.241949876 | 0.262825 |

| | | | | |
|---|---|---|---|---|
| Q96RS6 | NUDCD1 | NudC domain-containing protein 1 | 0.106147448 | -0.0329024 |
| Q96RT1 | ERBIN | Erbin | -0.528154294 | 0.414743 |
| Q96RU2-2 | USP28 | Isoform 2 of Ubiquitin carboxyl-terminal hydrolase 28 | 0.218778451 | -0.0413974 |
| Q96S19 | METTL26 | Methyltransferase-like 26 | -0.027277788 | -0.0530399 |
| Q96S44 | TP53RK | EKC/KEOPS complex subunit TP53RK | -0.034655412 | -0.360144 |
| Q96S52 | PIGS | GPI transamidase component PIG-S | -1.42019852 | -1.16284 |
| Q96S55 | WRNIP1 | ATPase WRNIP1 | -0.188485622 | -0.165286 |
| Q96S59 | RANBP9 | Ran-binding protein 9 | 0.172313293 | -0.596546 |
| Q96S82 | UBL7 | Ubiquitin-like protein 7 | -0.524644613 | 0.931525 |
| Q96S94 | CCNL2 | Cyclin-L2 | 0.857985338 | 1.40203 |
| Q96SB3 | PPP1R9B | Neurabin-2 | -0.138665199 | 0.0242586 |
| Q96ST2 | IWS1 | Protein IWS1 homolog | -0.225785494 | -0.000143369 |
| Q96ST3 | SIN3A | Paired amphipathic helix protein Sin3a | -0.384169579 | 0.0623678 |
| Q96SU4 | OSBPL9 | Oxysterol-binding protein-related protein 9 | -0.096338431 | -0.0145569 |
| Q96SZ5 | ADO | 2-aminoethanethiol dioxygenase | -0.018888791 | 0.176303 |
| Q96T23 | RSF1 | Remodeling and spacing factor 1 | 0.303470373 | -0.240677 |
| Q96T51 | RUFY1 | RUN and FYVE domain-containing protein 1 | -0.038182497 | 0.0657495 |
| Q96T76 | MMS19 | MMS19 nucleotide excision repair protein homolog | -0.121550083 | -0.291523 |
| Q96TA1 | NIBAN2 | Protein Niban 2 | -0.012199402 | -0.0581163 |
| Q96TA2 | YME1L1 | ATP-dependent zinc metalloprotease YME1L1 | -0.995572249 | 0.498113 |
| Q99426 | TBCB | Tubulin-folding cofactor B | 0.096845945 | 0.135317 |
| Q99436 | PSMB7 | Proteasome subunit beta type-7 | -0.259154638 | -0.041261 |
| Q99447 | PCYT2 | Ethanolamine-phosphate cytidylyltransferase | -0.052029451 | 0.0801455 |
| Q99459 | CDC5L | Cell division cycle 5-like protein | -0.574788014 | 0.243566 |
| Q99470 | SDF2 | Stromal cell-derived factor 2 | -1.270763318 | -0.144027 |
| Q99471 | PFDN5 | Prefoldin subunit 5 | 0.08607014 | 0.253457 |
| Q99487 | PAFAH2 | Platelet-activating factor acetylhydrolase 2, cytoplasmic | -1.453940233 | -0.801188 |
| Q99496 | RNF2 | E3 ubiquitin-protein ligase RING2 | -0.414566517 | -0.0146221 |
| Q99497 | PARK7 | Parkinson disease protein 7 | 0.153063456 | 0.230461 |
| Q99504 | EYA3 | Eyes absent homolog 3 | 0.025172393 | -0.168667 |
| Q99519 | NEU1 | Sialidase-1 | -0.24167641 | 0.14849 |
| Q99536 | VAT1 | Synaptic vesicle membrane protein VAT-1 homolog | -0.295293967 | -0.0677681 |
| Q99538 | LGMN | Legumain | -0.270294587 | 0.891897 |
| Q99541 | PLIN2 | Perilipin-2 | -0.808222294 | -0.175681 |
| Q99543 | DNAJC2 | DnaJ homolog subfamily C member 2 | -0.457236528 | -0.404034 |
| Q99567 | NUP88 | Nuclear pore complex protein Nup88 | -0.088114977 | 0.212752 |
| Q99570 | PIK3R4 | Phosphoinositide 3-kinase regulatory subunit 4 | -0.050263325 | -0.0938635 |
| Q99575 | POP1 | Ribonucleases P/MRP protein subunit POP1 | -0.3665452 | -0.336467 |
| Q99584 | S100A13 | Protein S100-A13 | -0.055644115 | 0.480375 |
| Q99598 | TSNAX | Translin-associated protein X | 0.037988106 | -0.200284 |
| Q99608 | NDN | Necdin | -0.797783613 | -0.610946 |
| Q99611 | SEPHS2 | Selenide, water dikinase 2 | -0.313940684 | 0.553523 |
| Q99613 | EIF3C | Eukaryotic translation initiation factor 3 subunit C | 0.085702419 | -0.185635 |
| Q99614 | TTC1 | Tetratricopeptide repeat protein 1 | 0.124014854 | 0.0356159 |
| Q99615 | DNAJC7 | DnaJ homolog subfamily C member 7 | 0.072163423 | 0.13453 |
| Q99623 | PHB2 | Prohibitin-2 | -1.017037789 | 0.362143 |
| Q99633 | PRPF18 | Pre-mRNA-splicing factor 18 | -0.937728167 | -2.34925 |
| Q99638 | RAD9A | Cell cycle checkpoint control protein RAD9A | -1.113962491 | -3.64385 |
| Q99640 | PKMYT1 | Membrane-associated tyrosine- and threonine-specific cdc2-inhibitory kinase | -1.297861814 | -0.270027 |
| Q99653 | CHP1 | Calcineurin B homologous protein 1 | -0.627239545 | 0.256837 |
| Q99661 | KIF2C | Kinesin-like protein KIF2C | -0.564817667 | 0.16987 |
| Q99704 | DOK1 | Docking protein 1 | -0.109323422 | -0.33574 |
| Q99714 | HSD17B10 | 3-hydroxyacyl-CoA dehydrogenase type-2 | -0.485801299 | 0.0215667 |
| Q99717 | SMAD5 | Mothers against decapentaplegic homolog 5 | -0.494513909 | -0.55004 |
| Q99733 | NAP1L4 | Nucleosome assembly protein 1-like 4 | 0.091558615 | 0.0981223 |
| Q99735 | MGST2 | Microsomal glutathione S-transferase 2 | -1.047865788 | 0.0767275 |
| Q99747 | NAPG | Gamma-soluble NSF attachment protein | 0.021297296 | -0.161819 |
| Q99797 | MIPEP | Mitochondrial intermediate peptidase | -1.715223948 | 0 |
| Q99805 | TM9SF2 | Transmembrane 9 superfamily member 2 | -1.606757561 | 0.0898259 |
| Q99808 | SLC29A1 | Equilibrative nucleoside transporter 1 | -2.140467167 | 2.01184 |
| Q99816 | TSG101 | Tumor susceptibility gene 101 protein | 0.10130628 | -0.0427608 |
| Q99832 | CCT7 | T-complex protein 1 subunit eta | -0.089715799 | -0.0657622 |
| Q99836 | MYD88 | Myeloid differentiation primary response protein MyD88 | 0.670337995 | -0.0277418 |
| Q99961 | SH3GL1 | Endophilin-A2 | 0.097415765 | -0.0378971 |
| Q9BPW8 | NIPSNAP1 | Protein NipSnap homolog 1 | -1.049902836 | 0.0753406 |
| Q9BPX3 | NCAPG | Condensin complex subunit 3 | -0.545045614 | -0.132198 |
| Q9BPX5 | ARPC5L | Actin-related protein 2/3 complex subunit 5-like protein | -0.365859906 | -0.321493 |
| Q9BPX7 | C7orf25 | UPF0415 protein C7orf25 | 0.200532913 | -0.258796 |
| Q9BQ04 | RBM4B | RNA-binding protein 4B | -1.002202431 | -3.07814 |
| Q9BQ39 | DDX50 | ATP-dependent RNA helicase DDX50 | -1.656242669 | 0.303795 |

| | | | | |
|---|---|---|---|---|
| Q9BQ50 | TREX2 | Three prime repair exonuclease 2 | -0.298818827 | 0 |
| Q9BQ51 | PDCD1LG2 | Programmed cell death 1 ligand 2 | -1.242776712 | 0 |
| Q9BQ52 | ELAC2 | Zinc phosphodiesterase ELAC protein 2 | -0.213188807 | 0.0152283 |
| Q9BQ61 | TRIR | Telomerase RNA component interacting RNase | -0.20432361 | 0.19798 |
| Q9BQ67 | GRWD1 | Glutamate-rich WD repeat-containing protein 1 | 0.061702728 | 0.0799497 |
| Q9BQ70 | TCF25 | Transcription factor 25 | -0.135955095 | -0.324055 |
| Q9BQA1 | WDR77 | Methylosome protein 50 | -0.132116636 | 0.0354163 |
| Q9BQC3 | DPH2 | 2-(3-amino-3-carboxypropyl)histidine synthase subunit 2 | 0.195110957 | 0.45944 |
| Q9BQC6 | MRPL57 | Ribosomal protein 63, mitochondrial | -0.790390968 | 0 |
| Q9BQG0 | MYBBP1A | Myb-binding protein 1A | -1.020742416 | -0.0182187 |
| Q9BQP7 | MGME1 | Mitochondrial genome maintenance exonuclease 1 | 0.331018209 | -0.65976 |
| Q9BQQ3 | GORASP1 | Golgi reassembly-stacking protein 1 | 0.072149118 | 1.75439 |
| Q9BQS8 | FYCO1 | FYVE and coiled-coil domain-containing protein 1 | -0.087478956 | 0.0517991 |
| Q9BR61 | ACBD6 | Acyl-CoA-binding domain-containing protein 6 | -0.047516187 | 0.298342 |
| Q9BR76 | CORO1B | Coronin-1B | 0.142903646 | 0.0492166 |
| Q9BRA2 | TXNDC17 | Thioredoxin domain-containing protein 17 | -0.112033367 | 0.524721 |
| Q9BRF8 | CPPED1 | Serine/threonine-protein phosphatase CPPED1 | 0.232942104 | -0.041268 |
| Q9BRG1 | VPS25 | Vacuolar protein-sorting-associated protein 25 | -0.183159033 | -0.406609 |
| Q9BRJ6 | C7orf50 | Uncharacterized protein C7orf50 | -0.801803192 | -0.0438499 |
| Q9BRP1 | PDCD2L | Programmed cell death protein 2-like | -0.050042709 | 0.185889 |
| Q9BRP8 | PYM1 | Partner of Y14 and mago | -0.645619154 | -0.185555 |
| Q9BRQ8 | AIFM2 | Ferroptosis suppressor protein 1 | -0.944096247 | -0.570688 |
| Q9BRR6 | ADPGK | ADP-dependent glucokinase | -1.252719084 | 0.168118 |
| Q9BRS2 | RIOK1 | Serine/threonine-protein kinase RIO1 | -0.244861364 | 0.0654348 |
| Q9BRT6 | LLPH | Protein LLP homolog | -1.638343811 | 1.30371 |
| Q9BRT9 | GINS4 | DNA replication complex GINS protein SLD5 | 0.378533999 | 0.619138 |
| Q9BRX2 | PELO | Protein pelota homolog | -0.040785074 | -0.0190802 |
| Q9BRX5 | GINS3 | DNA replication complex GINS protein PSF3 | 0.119478226 | 0.157868 |
| Q9BRX8 | PRXL2A | Peroxiredoxin-like 2A | -1.766064485 | 0.236984 |
| Q9BRZ2 | TRIM56 | E3 ubiquitin-protein ligase TRIM56 | -0.519051393 | -1.98817 |
| Q9BS26 | ERP44 | Endoplasmic reticulum resident protein 44 | -0.115942637 | 0.326845 |
| Q9BS40 | LXN | Latexin | 0.07420675 | 0.585917 |
| Q9BSH5 | HDHD3 | Haloacid dehalogenase-like hydrolase domain-containing protein 3 | -0.823050499 | 0 |
| Q9BSJ8-2 | ESYT1 | Isoform 2 of Extended synaptotagmin-1 | -1.041417758 | 0.254875 |
| Q9BSL1 | UBAC1 | Ubiquitin-associated domain-containing protein 1 | -0.486564159 | -1.14379 |
| Q9BSU1 | PHAF1 | Phagosome assembly factor 1 | -0.231696685 | 0.253224 |
| Q9BSY9 | DESI2 | Deubiquitinase DESI2 | -1.323664824 | 0.230631 |
| Q9BT09 | CNPY3 | Protein canopy homolog 3 | -0.174229463 | 0.468605 |
| Q9BT25 | HAUS8 | HAUS augmin-like complex subunit 8 | -0.739945809 | 0.277224 |
| Q9BT67 | NDFIP1 | NEDD4 family-interacting protein 1 | -1.367379308 | 0 |
| Q9BT73 | PSMG3 | Proteasome assembly chaperone 3 | -0.616114457 | -0.48261 |
| Q9BT78 | COPS4 | COP9 signalosome complex subunit 4 | -0.002154827 | -0.00397746 |
| Q9BTC0 | DIDO1 | Death-inducer obliterator 1 | -1.359878222 | -0.0712061 |
| Q9BTE1 | DCTN5 | Dynactin subunit 5 | -0.133354743 | 0.228971 |
| Q9BTE3 | MCMBP | Mini-chromosome maintenance complex-binding protein | -0.058460951 | -0.00433954 |
| Q9BTE3-2 | MCMBP | Isoform 2 of Mini-chromosome maintenance complex-binding protein | -0.519784768 | 0 |
| Q9BTE7 | DCUN1D5 | DCN1-like protein 5 | -0.244818926 | 0.103669 |
| Q9BTM9 | URM1 | Ubiquitin-related modifier 1 | -0.384877761 | -0.586723 |
| Q9BTT0 | ANP32E | Acidic leucine-rich nuclear phosphoprotein 32 family member E | -0.084902287 | -0.0448119 |
| Q9BTU6 | PI4K2A | Phosphatidylinositol 4-kinase type 2-alpha | -1.571290175 | -0.0326951 |
| Q9BTV4 | TMEM43 | Transmembrane protein 43 | -1.560442448 | 0.0497675 |
| Q9BTV6 | DPH7 | Diphthine methyltransferase | -0.526223898 | 2.2185 |
| Q9BTX1 | NDC1 | Nucleoporin NDC1 | 0.242309093 | 0.0443821 |
| Q9BTX7 | TTPAL | Alpha-tocopherol transfer protein-like | -0.500305971 | -0.375559 |
| Q9BTY2 | FUCA2 | Plasma alpha-L-fucosidase | -0.633207957 | -0.164084 |
| Q9BTY7 | HGH1 | Protein HGH1 homolog | 0.25949796 | 0.821615 |
| Q9BTZ2 | DHRS4 | Dehydrogenase/reductase SDR family member 4 | -0.725958506 | 0.123162 |
| Q9BU02 | THTPA | Thiamine-triphosphatase | 0.375860453 | 1.69187 |
| Q9BU23 | LMF2 | Lipase maturation factor 2 | -1.810503761 | 0.368176 |
| Q9BU61 | NDUFAF3 | NADH dehydrogenase [ubiquinone] 1 alpha subcomplex assembly factor 3 | -0.927657684 | 0.414133 |
| Q9BU89 | DOHH | Deoxyhypusine hydroxylase | 0.176831007 | 0.186665 |
| Q9BUA3 | SPINDOC | Spindlin interactor and repressor of chromatin-binding protein | 0.089137475 | -0.341322 |
| Q9BUB7 | TMEM70 | Transmembrane protein 70, mitochondrial | -3.574865103 | 3.0607 |
| Q9BUF5 | TUBB6 | Tubulin beta-6 chain | -0.0393068 | 0.110792 |
| Q9BUH6 | PAXX | Protein PAXX | -0.033227523 | 0.137304 |
| Q9BUI4 | POLR3C | DNA-directed RNA polymerase III subunit RPC3 | 0.057778041 | 0.0438913 |
| Q9BUJ2 | HNRNPUL1 | Heterogeneous nuclear ribonucleoprotein U-like protein 1 | -0.257391373 | 0.14713 |
| Q9BUK6 | MSTO1 | Protein misato homolog 1 | -0.073521217 | 0.0729262 |
| Q9BUL5 | PHF23 | PHD finger protein 23 | -0.1781768 | -0.0389175 |

| | | | | |
|---|---|---|---|---|
| Q9BUL8 | PDCD10 | Programmed cell death protein 10 | 0.060474237 | 0.00556215 |
| Q9BUL9 | RPP25 | Ribonuclease P protein subunit p25 | 0.889582396 | 1.59724 |
| Q9BUP3 | HTATIP2 | Oxidoreductase HTATIP2 | -1.02297775 | 0.017031 |
| Q9BUQ8 | DDX23 | Probable ATP-dependent RNA helicase DDX23 | -0.611183087 | 0.0695305 |
| Q9BUR4 | WRAP53 | Telomerase Cajal body protein 1 | 0.140175819 | 0.182645 |
| Q9BUT1 | BDH2 | 3-hydroxybutyrate dehydrogenase type 2 | 0.263785203 | 0.897545 |
| Q9BV20 | MRI1 | Methylthioribose-1-phosphate isomerase | 0.140654643 | -0.0165094 |
| Q9BV36 | MLPH | Melanophilin | -0.324185054 | 0.100181 |
| Q9BV36-3 | MLPH | Isoform 3 of Melanophilin | -0.920434952 | 0.0870361 |
| Q9BV44 | THUMPD3 | THUMP domain-containing protein 3 | 0.176216841 | -0.0210648 |
| Q9BV57 | ADI1 | 1,2-dihydroxy-3-keto-5-methylthiopentene dioxygenase | -0.011128823 | 0.438527 |
| Q9BV73 | CEP250 | Centrosome-associated protein CEP250 | 0.5456779 | 2.10885 |
| Q9BV81 | EMC6 | ER membrane protein complex subunit 6 | -2.008030574 | 0.621535 |
| Q9BV86 | NTMT1 | N-terminal Xaa-Pro-Lys N-methyltransferase 1 | -0.065504233 | 0.418288 |
| Q9BVA0 | KATNB1 | Katanin p80 WD40 repeat-containing subunit B1 | -0.069448709 | -0.0227207 |
| Q9BVA1 | TUBB2B | Tubulin beta-2B chain | 0.077144782 | -0.246321 |
| Q9BVC4 | MLST8 | Target of rapamycin complex subunit LST8 | -1.433476845 | -0.210075 |
| Q9BVC6 | TMEM109 | Transmembrane protein 109 | -1.270473639 | 0.222174 |
| Q9BVG4 | PBDC1 | Protein PBDC1 | 0.029530366 | 0.240211 |
| Q9BVI4 | NOC4L | Nucleolar complex protein 4 homolog | -1.309242566 | -0.591658 |
| Q9BVJ6 | UTP14A | U3 small nucleolar RNA-associated protein 14 homolog A | -1.094551206 | 0.229128 |
| Q9BVJ7 | DUSP23 | Dual specificity protein phosphatase 23 | -0.030703386 | -0.100919 |
| Q9BVK6 | TMED9 | Transmembrane emp24 domain-containing protein 9 | -1.291920106 | 0.253589 |
| Q9BVM2 | DPCD | Protein DPCD | -0.038334846 | -0.548327 |
| Q9BVM4 | GGACT | Gamma-glutamylaminecyclotransferase | 0.525714199 | 0 |
| Q9BVP2 | GNL3 | Guanine nucleotide-binding protein-like 3 | -0.783902804 | 0.259963 |
| Q9BVQ7 | SPATA5L1 | Spermatogenesis-associated protein 5-like protein 1 | -0.044248422 | -0.365155 |
| Q9BVS4 | RIOK2 | Serine/threonine-protein kinase RIO2 | -1.199604193 | -0.376048 |
| Q9BVW5 | TIPIN | TIMELESS-interacting protein | 0.236455202 | 0.683945 |
| Q9BW19 | KIFC1 | Kinesin-like protein KIFC1 | -2.635716558 | -1.04788 |
| Q9BW27 | NUP85 | Nuclear pore complex protein Nup85 | -0.367331266 | 0.337275 |
| Q9BW60 | ELOVL1 | Elongation of very long chain fatty acids protein 1 | -3.302666465 | 0.309763 |
| Q9BW61 | DDA1 | DET1- and DDB1-associated protein 1 | -1.188428322 | 1.06884 |
| Q9BW71 | HIRIP3 | HIRA-interacting protein 3 | -0.550827265 | 0.251634 |
| Q9BW83 | IFT27 | Intraflagellar transport protein 27 homolog | -0.044495185 | 0.0853888 |
| Q9BW85 | YJU2 | Splicing factor YJU2 | -0.645045757 | 0.251823 |
| Q9BW91 | NUDT9 | ADP-ribose pyrophosphatase, mitochondrial | -0.161296368 | -0.0285568 |
| Q9BWD1 | ACAT2 | Acetyl-CoA acetyltransferase, cytosolic | 0.173140208 | -0.143389 |
| Q9BWF3 | RBM4 | RNA-binding protein 4 | -0.076715708 | 0.166685 |
| Q9BWH6 | RPAP1 | RNA polymerase II-associated protein 1 | -0.369102796 | -0.611418 |
| Q9BX68 | HINT2 | Adenosine 5'-monophosphoramidase HINT2 | -0.808659633 | 2.51964 |
| Q9BX70 | BTBD2 | BTB/POZ domain-containing protein 2 | -1.013315161 | 0 |
| Q9BXB4 | OSBPL11 | Oxysterol-binding protein-related protein 11 | -0.09734424 | -0.00465488 |
| Q9BXJ9 | NAA15 | N-alpha-acetyltransferase 15, NatA auxiliary subunit | 0.03784736 | -0.0740719 |
| Q9BXK5 | BCL2L13 | Bcl-2-like protein 13 | -0.917764425 | 0.452252 |
| Q9BXP5 | SRRT | Serrate RNA effector molecule homolog | 0.02519687 | 0.0949198 |
| Q9BXR0 | QTRT1 | Queuine tRNA-ribosyltransferase catalytic subunit 1 | 0.121881326 | 0.213613 |
| Q9BXS5 | AP1M1 | AP-1 complex subunit mu-1 | 0.032509009 | -0.217663 |
| Q9BXS6 | NUSAP1 | Nucleolar and spindle-associated protein 1 | -0.603828192 | -0.387216 |
| Q9BXW6 | OSBPL1A | Oxysterol-binding protein-related protein 1 | -0.76078399 | -2.35684 |
| Q9BXW7 | HDHD5 | Haloacid dehalogenase-like hydrolase domain-containing 5 | -0.577888648 | -0.0792128 |
| Q9BXY0 | MAK16 | Protein MAK16 homolog | -0.846509457 | 0.00633589 |
| Q9BY32 | ITPA | Inosine triphosphate pyrophosphatase | 0.103461583 | 0.0487556 |
| Q9BY43 | CHMP4A | Charged multivesicular body protein 4a | 0.297005653 | -0.224843 |
| Q9BY44 | EIF2A | Eukaryotic translation initiation factor 2A | -0.190196117 | 0.00759602 |
| Q9BY78 | RNF26 | E3 ubiquitin-protein ligase RNF26 | -0.962334792 | -0.419167 |
| Q9BYB4 | GNB1L | Guanine nucleotide-binding protein subunit beta-like protein 1 | -0.465435982 | 0.567845 |
| Q9BYD1 | MRPL13 | 39S ribosomal protein L13, mitochondrial | -2.778954665 | -1.48329 |
| Q9BYD2 | MRPL9 | 39S ribosomal protein L9, mitochondrial | -2.067102631 | -1.00995 |
| Q9BYD6 | MRPL1 | 39S ribosomal protein L1, mitochondrial | -0.096488476 | -0.257476 |
| Q9BYG3 | NIFK | MKI67 FHA domain-interacting nucleolar phosphoprotein | -1.346419573 | 0.478849 |
| Q9BYJ9 | YTHDF1 | YTH domain-containing family protein 1 | -0.510556777 | -0.237677 |
| Q9BYM8 | RBCK1 | RanBP-type and C3HC4-type zinc finger-containing protein 1 | 0.458987236 | 0.100517 |
| Q9BYN0 | SRXN1 | Sulfiredoxin-1 | -0.790962299 | -0.821997 |
| Q9BYN8 | MRPS26 | 28S ribosomal protein S26, mitochondrial | 0.17533648 | -3.05852 |
| Q9BYT8 | NLN | Neurolysin, mitochondrial | 0.057710091 | -0.227667 |
| Q9BYX2 | TBC1D2 | TBC1 domain family member 2A | -0.195987463 | -0.183108 |
| Q9BZ23 | PANK2 | Pantothenate kinase 2, mitochondrial | -0.0253733 | -0.321535 |
| Q9BZD4 | NUF2 | Kinetochore protein Nuf2 | -0.226693471 | -0.354298 |
| Q9BZE1 | MRPL37 | 39S ribosomal protein L37, mitochondrial | -0.890868823 | -2.75048 |
| Q9BZE2 | PUS3 | tRNA pseudouridine(38/39) synthase | -0.148861249 | -4.48013 |

| | | | | |
|---|---|---|---|---|
| Q9BZE4 | GTPBP4 | GTP-binding protein 4 | -1.042907635 | 0.127509 |
| Q9BZE9 | ASPSCR1 | Tether containing UBX domain for GLUT4 | -0.069011688 | 0.164152 |
| Q9BZF1 | OSBPL8 | Oxysterol-binding protein-related protein 8 | -1.124847571 | -0.291792 |
| Q9BZF3 | OSBPL6 | Oxysterol-binding protein-related protein 6 | -0.068938971 | -0.787426 |
| Q9BZG1 | RAB34 | Ras-related protein Rab-34 | -0.703638713 | 0.353971 |
| Q9BZH6 | WDR11 | WD repeat-containing protein 11 | -1.004799128 | -1.10231 |
| Q9BZK7 | TBL1XR1 | F-box-like/WD repeat-containing protein TBL1XR1 | -0.195165396 | -0.139468 |
| Q9BZL1 | UBL5 | Ubiquitin-like protein 5 | -0.305323919 | 0.782058 |
| Q9BZL6 | PRKD2 | Serine/threonine-protein kinase D2 | -0.286801418 | 0.00247224 |
| Q9BZQ8 | NIBAN1 | Protein Niban 1 | -0.095452785 | 0.193722 |
| Q9BZV1 | UBXN6 | UBX domain-containing protein 6 | -0.128621578 | -0.056016 |
| Q9BZZ5 | API5 | Apoptosis inhibitor 5 | -0.005912463 | -0.0507018 |
| Q9BZZ5-2 | API5 | Isoform 2 of Apoptosis inhibitor 5 | 0.443458239 | -0.0214326 |
| Q9C000 | NLRP1 | NACHT, LRR and PYD domains-containing protein 1 | -0.43107303 | 3.52078 |
| Q9C002 | NMES1 | Normal mucosa of esophagus-specific gene 1 protein | -4.412487912 | 0 |
| Q9C035 | TRIM5 | Tripartite motif-containing protein 5 | 0.967847586 | 0.257744 |
| Q9C037 | TRIM4 | E3 ubiquitin-protein ligase TRIM4 | -2.543888489 | 0.526331 |
| Q9C075 | KRT23 | Keratin, type I cytoskeletal 23 | -0.590857506 | -0.770452 |
| Q9C0B7 | TANGO6 | Transport and Golgi organization protein 6 homolog | 0.230537812 | -0.896118 |
| Q9C0C2 | TNKS1BP1 | 182 kDa tankyrase-1-binding protein | -0.033178568 | 0.121756 |
| Q9C0C9 | UBE2O | (E3-independent) E2 ubiquitin-conjugating enzyme | 0.001390457 | 0.0345341 |
| Q9C0E2 | XPO4 | Exportin-4 | -0.122706175 | -0.133855 |
| Q9C0E8 | LNPK | Endoplasmic reticulum junction formation protein lunapark | -0.878683567 | 0.430637 |
| Q9C0I1 | MTMR12 | Myotubularin-related protein 12 | -0.255242427 | -1.03823 |
| Q9C0J8 | WDR33 | pre-mRNA 3' end processing protein WDR33 | 0.470053752 | 0 |
| Q9GZL7 | WDR12 | Ribosome biogenesis protein WDR12 | -0.185650746 | -0.153128 |
| Q9GZN1 | ACTR6 | Actin-related protein 6 | -0.620420615 | 0 |
| Q9GZN8 | C20orf27 | UPF0687 protein C20orf27 | -0.081760248 | -0.278936 |
| Q9GZP4 | PITHD1 | PITH domain-containing protein 1 | -0.189968109 | -0.0466366 |
| Q9GZP8 | IMUP | Immortalization up-regulated protein | -0.86209019 | 0 |
| Q9GZS3 | WDR61 | WD repeat-containing protein 61 | -0.374599695 | -0.174509 |
| Q9GZT3 | SLIRP | SRA stem-loop-interacting RNA-binding protein, mitochondrial | -1.841139634 | 0.10533 |
| Q9GZT4 | SRR | Serine racemase | 0.095370372 | 0.0536432 |
| Q9GZT8 | NIF3L1 | NIF3-like protein 1 | -0.288982948 | -0.351934 |
| Q9GZT9 | EGLN1 | Egl nine homolog 1 | -0.68526268 | 0.608511 |
| Q9GZU8 | PSME3IP1 | PSME3-interacting protein | 0.122945627 | -0.230731 |
| Q9GZX9 | TWSG1 | Twisted gastrulation protein homolog 1 | -0.099662701 | -0.0553818 |
| Q9GZY8 | MFF | Mitochondrial fission factor | -2.689484119 | 4.98972 |
| Q9H061 | TMEM126A | Transmembrane protein 126A | -1.554547946 | 0.169257 |
| Q9H074 | PAIP1 | Polyadenylate-binding protein-interacting protein 1 | -0.114769538 | -0.100606 |
| Q9H074-3 | PAIP1 | Isoform 3 of Polyadenylate-binding protein-interacting protein 1 | -0.658006032 | -1.47944 |
| Q9H078 | CLPB | Caseinolytic peptidase B protein homolog | -0.764153639 | -0.231105 |
| Q9H089 | LSG1 | Large subunit GTPase 1 homolog | -0.495682081 | -0.0200469 |
| Q9H0A0 | NAT10 | RNA cytidine acetyltransferase | -0.620627403 | 0.115629 |
| Q9H0A8 | COMMD4 | COMM domain-containing protein 4 | -0.137281338 | -0.176208 |
| Q9H0B6 | KLC2 | Kinesin light chain 2 | -0.168483178 | -0.0419143 |
| Q9H0C8 | ILKAP | Integrin-linked kinase-associated serine/threonine phosphatase 2C | -0.10104386 | -0.127483 |
| Q9H0D6 | XRN2 | 5'-3' exoribonuclease 2 | -0.237744172 | 0.0305471 |
| Q9H0E2 | TOLLIP | Toll-interacting protein | -0.038994869 | 0.321396 |
| Q9H0F6 | SHARPIN | Sharpin | 0.745866537 | 0.773051 |
| Q9H0F7 | ARL6 | ADP-ribosylation factor-like protein 6 | -0.39314429 | 0.386896 |
| Q9H0H5 | RACGAP1 | Rac GTPase-activating protein 1 | -1.391344786 | -0.136237 |
| Q9H0K1 | SIK2 | Serine/threonine-protein kinase SIK2 | -0.58561341 | 1.36602 |
| Q9H0K6 | PUS7L | Pseudouridylate synthase 7 homolog-like protein | -0.335447868 | 0.0236588 |
| Q9H0P0 | NT5C3A | Cytosolic 5'-nucleotidase 3A | 0.598711888 | -0.0319529 |
| Q9H0Q0 | CYRIA | CYFIP-related Rac1 interactor A | 0.097992659 | -0.858023 |
| Q9H0R4 | HDHD2 | Haloacid dehalogenase-like hydrolase domain-containing protein 2 | -0.277142763 | -0.167783 |
| Q9H0S4 | DDX47 | Probable ATP-dependent RNA helicase DDX47 | -0.541802406 | 0.0832138 |
| Q9H0U4 | RAB1B | Ras-related protein Rab-1B | -0.676709652 | 0.232145 |
| Q9H0V9 | LMAN2L | VIP36-like protein | 0.174131234 | 0.192383 |
| Q9H0W8 | SMG9 | Nonsense-mediated mRNA decay factor SMG9 | -0.530734857 | 0.709969 |
| Q9H1A3 | METTL9 | Protein-L-histidine N-pros-methyltransferase | -0.516432047 | -0.0315723 |
| Q9H1A4 | ANAPC1 | Anaphase-promoting complex subunit 1 | -0.213368018 | 0.560888 |
| Q9H1B7 | IRF2BPL | Probable E3 ubiquitin-protein ligase IRF2BPL | -0.769607147 | -0.817328 |
| Q9H1D9 | POLR3F | DNA-directed RNA polymerase III subunit RPC6 | -0.160081228 | -0.069891 |
| Q9H1E1 | RNASE7 | Ribonuclease 7 | -1.54143699 | 0.867919 |
| Q9H1E3 | NUCKS1 | Nuclear ubiquitous casein and cyclin-dependent kinase substrate 1 | -0.098206043 | -0.154678 |
| Q9H1E5 | TMX4 | Thioredoxin-related transmembrane protein 4 | -0.421563864 | 0.491346 |
| Q9H1H9 | KIF13A | Kinesin-like protein KIF13A | -0.006803195 | -0.0227944 |
| Q9H1Y0 | ATG5 | Autophagy protein 5 | 0.123370409 | -0.088528 |

| | | | | |
|---|---|---|---|---|
| Q9H1Z4 | WDR13 | WD repeat-containing protein 13 | 0.015666326 | -0.264358 |
| Q9H223 | EHD4 | EH domain-containing protein 4 | -0.02484266 | 0.0389798 |
| Q9H267 | VPS33B | Vacuolar protein sorting-associated protein 33B | -0.147878965 | -0.442911 |
| Q9H269 | VPS16 | Vacuolar protein sorting-associated protein 16 homolog | -0.071040154 | 0.0191402 |
| Q9H2C0 | GAN | Gigaxonin | 0.249339978 | -0.200676 |
| Q9H2D6 | TRIOBP | TRIO and F-actin-binding protein | -0.677323977 | 0.00400543 |
| Q9H2G2-2 | SLK | Isoform 2 of STE20-like serine/threonine-protein kinase | -0.104454438 | 0.175334 |
| Q9H2H8 | PPIL3 | Peptidyl-prolyl cis-trans isomerase-like 3 | -0.182976484 | 0.205643 |
| Q9H2J4 | PDCL3 | Phosducin-like protein 3 | -0.09969155 | -0.114898 |
| Q9H2K8 | TAOK3 | Serine/threonine-protein kinase TAO3 | 0 | 0.0952104 |
| Q9H2M9 | RAB3GAP2 | Rab3 GTPase-activating protein non-catalytic subunit | 0.095140775 | -0.111368 |
| Q9H2P0 | ADNP | Activity-dependent neuroprotector homeobox protein | -0.698681434 | -0.0702597 |
| Q9H2P9 | DPH5 | Diphthine methyl ester synthase | 0.027301311 | -0.198815 |
| Q9H2U1 | DHX36 | ATP-dependent DNA/RNA helicase DHX36 | -0.918475707 | -0.395817 |
| Q9H2U2 | PPA2 | Inorganic pyrophosphatase 2, mitochondrial | -0.283858776 | -0.0479441 |
| Q9H2W6 | MRPL46 | 39S ribosomal protein L46, mitochondrial | -1.144099553 | -0.497124 |
| Q9H307 | PNN | Pinin | -0.597683827 | 0.0123294 |
| Q9H330 | TMEM245 | Transmembrane protein 245 | -2.279513359 | 0.277786 |
| Q9H3G5 | CPVL | Probable serine carboxypeptidase CPVL | 0.040489515 | -0.00227197 |
| Q9H3H3 | C11orf68 | UPF0696 protein C11orf68 | 0.011781693 | -0.067097 |
| Q9H3K2 | GHITM | Growth hormone-inducible transmembrane protein | -1.38176767 | 1.48288 |
| Q9H3M7 | TXNIP | Thioredoxin-interacting protein | -0.138843973 | 0 |
| Q9H3N1 | TMX1 | Thioredoxin-related transmembrane protein 1 | -1.159859498 | 0.143678 |
| Q9H3P7 | ACBD3 | Golgi resident protein GCP60 | 0.132939974 | -0.15377 |
| Q9H3Q1 | CDC42EP4 | Cdc42 effector protein 4 | 0.04541254 | -0.249314 |
| Q9H3S7 | PTPN23 | Tyrosine-protein phosphatase non-receptor type 23 | 0.096385002 | 0.0592686 |
| Q9H3U1 | UNC45A | Protein unc-45 homolog A | 0.016236464 | -0.0283076 |
| Q9H3U1-2 | UNC45A | Isoform 2 of Protein unc-45 homolog A | -0.099076112 | -0.50327 |
| Q9H3Z4 | DNAJC5 | DnaJ homolog subfamily C member 5 | -1.51923728 | -0.182243 |
| Q9H425 | C1orf198 | Uncharacterized protein C1orf198 | -1.36740303 | -0.885918 |
| Q9H444 | CHMP4B | Charged multivesicular body protein 4b | -0.049394925 | -0.250934 |
| Q9H467 | CUEDC2 | CUE domain-containing protein 2 | 0.140672843 | 1.07316 |
| Q9H477 | RBKS | Ribokinase | 0.366381884 | 0 |
| Q9H479 | FN3K | Fructosamine-3-kinase | -0.159295241 | 0.5282 |
| Q9H488 | POFUT1 | GDP-fucose protein O-fucosyltransferase 1 | -0.130176385 | 0.335824 |
| Q9H490 | PIGU | Phosphatidylinositol glycan anchor biosynthesis class U protein | 4.40890038 | 0 |
| Q9H492 | MAP1LC3A | Microtubule-associated proteins 1A/1B light chain 3A | -0.693573236 | -0.150984 |
| Q9H4A4 | RNPEP | Aminopeptidase B | 0.127096335 | -0.149156 |
| Q9H4A5 | GOLPH3L | Golgi phosphoprotein 3-like | 0.186713616 | -0.0447842 |
| Q9H4A6 | GOLPH3 | Golgi phosphoprotein 3 | -0.225284656 | -0.0139173 |
| Q9H4B7 | TUBB1 | Tubulin beta-1 chain | -0.144788663 | -0.707929 |
| Q9H4L4 | SENP3 | Sentrin-specific protease 3 | -1.11437122 | 0.00383059 |
| Q9H4L5 | OSBPL3 | Oxysterol-binding protein-related protein 3 | -0.244657278 | -0.0481224 |
| Q9H4L7 | SMARCAD1 | SWI/SNF-related matrix-associated actin-dependent regulator of chromatin subfamily A containing DEAD/H box 1 | -0.306337516 | -0.0165373 |
| Q9H4M9 | EHD1 | EH domain-containing protein 1 | -0.060695648 | -0.15118 |
| Q9H501 | ESF1 | ESF1 homolog | -0.456202586 | -0.0990861 |
| Q9H583 | HEATR1 | HEAT repeat-containing protein 1 | -1.087215662 | 0.251076 |
| Q9H5N1 | RABEP2 | Rab GTPase-binding effector protein 2 | 0.424587091 | 0.111903 |
| Q9H5Q4 | TFB2M | Dimethyladenosine transferase 2, mitochondrial | -0.066636086 | 0.839851 |
| Q9H5V8 | CDCP1 | CUB domain-containing protein 1 | -1.258949121 | 0.507418 |
| Q9H5V9 | STEEP1 | STING ER exit protein | -0.52052021 | 1.08324 |
| Q9H5X1 | CIAO2A | Cytosolic iron-sulfur assembly component 2A | -0.814833721 | 0.45764 |
| Q9H6D7 | HAUS4 | HAUS augmin-like complex subunit 4 | 0.100987514 | -0.160563 |
| Q9H6E4 | CCDC134 | Coiled-coil domain-containing protein 134 | -2.081059933 | -2.14607 |
| Q9H6F5 | CCDC86 | Coiled-coil domain-containing protein 86 | 0.127872388 | 0.0555112 |
| Q9H6K1 | ILRUN | Protein ILRUN | -1.167474747 | 1.16246 |
| Q9H6Q4 | CIAO3 | Cytosolic iron-sulfur assembly component 3 | -0.103974422 | 0.488379 |
| Q9H6R4 | NOL6 | Nucleolar protein 6 | -1.594021956 | 0.728285 |
| Q9H6T3 | RPAP3 | RNA polymerase II-associated protein 3 | 0.161196073 | 0.0365861 |
| Q9H6U6 | BCAS3 | BCAS3 microtubule associated cell migration factor | -0.937987487 | -1.2174 |
| Q9H6W3 | RIOX1 | Ribosomal oxygenase 1 | -1.49395593 | 0.119763 |
| Q9H6Y2 | WDR55 | WD repeat-containing protein 55 | 0.125201305 | -0.237419 |
| Q9H6Z4 | RANBP3 | Ran-binding protein 3 | 0.136579196 | 0.0903533 |
| Q9H773 | DCTPP1 | dCTP pyrophosphatase 1 | 0.94907395 | -0.147468 |
| Q9H788 | SH2D4A | SH2 domain-containing protein 4A | -0.194212039 | -0.0375102 |
| Q9H7B2 | RPF2 | Ribosome production factor 2 homolog | -0.311878045 | -0.258577 |
| Q9H7B4 | SMYD3 | Histone-lysine N-methyltransferase SMYD3 | 0.114873568 | -0.482235 |
| Q9H7C4 | SYNC | Syncoilin | 0.296193918 | 0.142081 |
| Q9H7C9 | AAMDC | Mth938 domain-containing protein | 0.056089958 | 0.411304 |
| Q9H7S9 | ZNF703 | Zinc finger protein 703 | -1.442529202 | -0.435417 |

| | | | | |
|---|---|---|---|---|
| Q9H7X7 | IFT22 | Intraflagellar transport protein 22 homolog | -0.794034799 | 1.0654 |
| Q9H813 | PACC1 | Proton-activated chloride channel | -3.26772968 | 1.55015 |
| Q9H814 | PHAX | Phosphorylated adapter RNA export protein | -0.056574504 | 0.182094 |
| Q9H832 | UBE2Z | Ubiquitin-conjugating enzyme E2 Z | -0.12223204 | 0.182852 |
| Q9H840 | GEMIN7 | Gem-associated protein 7 | 1.075055599 | 0 |
| Q9H845 | ACAD9 | Complex I assembly factor ACAD9, mitochondrial | -1.229148865 | 0.217991 |
| Q9H853 | TUBA4B | Putative tubulin-like protein alpha-4B | 0.706923485 | -5.06067 |
| Q9H857 | NT5DC2 | 5'-nucleotidase domain-containing protein 2 | -1.238369147 | -0.205243 |
| Q9H871 | RMND5A | E3 ubiquitin-protein transferase RMND5A | 0.399820646 | 0 |
| Q9H8H0 | NOL11 | Nucleolar protein 11 | -0.461025476 | 0.286908 |
| Q9H8K7 | PAAT | ATPase PAAT | -0.23012042 | 0.206559 |
| Q9H8M7 | MINDY3 | Ubiquitin carboxyl-terminal hydrolase MINDY-3 | 0.029930353 | 0.198099 |
| Q9H8S9 | MOB1A | MOB kinase activator 1A | 0.929118395 | 0.960175 |
| Q9H8U3 | ZFAND3 | AN1-type zinc finger protein 3 | -0.577326536 | -1.08179 |
| Q9H8V3 | ECT2 | Protein ECT2 | -0.043570518 | 0.348922 |
| Q9H8W4 | PLEKHF2 | Pleckstrin homology domain-containing family F member 2 | -0.381760438 | -0.499458 |
| Q9H8Y8 | GORASP2 | Golgi reassembly-stacking protein 2 | 0.017875036 | 0.17822 |
| Q9H900 | ZWILCH | Protein zwilch homolog | -0.052973668 | 0.0503893 |
| Q9H944 | MED20 | Mediator of RNA polymerase II transcription subunit 20 | -0.321916739 | -2.93219 |
| Q9H974 | QTRT2 | Queuine tRNA-ribosyltransferase accessory subunit 2 | -0.362540563 | -0.207136 |
| Q9H977 | WDR54 | WD repeat-containing protein 54 | -0.133137782 | -0.164044 |
| Q9H993 | ARMT1 | Damage-control phosphatase ARMT1 | -0.022207022 | -0.246388 |
| Q9H999 | PANK3 | Pantothenate kinase 3 | -0.142519236 | -0.294181 |
| Q9H9A6 | LRRC40 | Leucine-rich repeat-containing protein 40 | 0.097250144 | -0.0948356 |
| Q9H9B4 | SFXN1 | Sideroflexin-1 | -1.316347758 | 0.298043 |
| Q9H9F9 | ACTR5 | Actin-related protein 5 | -0.443887949 | -1.10017 |
| Q9H9J2 | MRPL44 | 39S ribosomal protein L44, mitochondrial | -2.4999547 | 3.86476 |
| Q9H9T3 | ELP3 | Elongator complex protein 3 | -0.008700768 | -0.197232 |
| Q9H9Y2 | RPF1 | Ribosome production factor 1 | -0.523853739 | -1.96379 |
| Q9H9Y6 | POLR1B | DNA-directed RNA polymerase I subunit RPA2 | 0.187862714 | -0.319349 |
| Q9HA47 | UCK1 | Uridine-cytidine kinase 1 | 0.1335543 | -0.227464 |
| Q9HA64 | FN3KRP | Ketosamine-3-kinase | 0.126686017 | 0.16057 |
| Q9HA65 | TBC1D17 | TBC1 domain family member 17 | 0.274404685 | -0.0309842 |
| Q9HA77 | CARS2 | Probable cysteine--tRNA ligase, mitochondrial | -2.141577562 | -0.156674 |
| Q9HAB8 | PPCS | Phosphopantothenate--cysteine ligase | 0.220750411 | -0.151203 |
| Q9HAC8 | UBTD1 | Ubiquitin domain-containing protein 1 | -0.214980364 | 0 |
| Q9HAD4 | WDR41 | WD repeat-containing protein 41 | 0.062329531 | -0.182014 |
| Q9HAN9 | NMNAT1 | Nicotinamide/nicotinic acid mononucleotide adenylyltransferase 1 | -0.725221952 | 1.20511 |
| Q9HAT2 | SIAE | Sialate O-acetylesterase | -0.354442676 | 0.936537 |
| Q9HAU5 | UPF2 | Regulator of nonsense transcripts 2 | -0.003976345 | 1.46969 |
| Q9HAV0 | GNB4 | Guanine nucleotide-binding protein subunit beta-4 | -1.805505435 | 0.288309 |
| Q9HAV4 | XPO5 | Exportin-5 | -0.307966312 | -0.0674057 |
| Q9HAV7 | GRPEL1 | GrpE protein homolog 1, mitochondrial | 0.061376731 | -0.00726446 |
| Q9HAW4 | CLSPN | Claspin | 0.004077752 | -0.0454979 |
| Q9HB07 | MYG1 | MYG1 exonuclease | 0.074753602 | -0.154922 |
| Q9HB20 | PLEKHA3 | Pleckstrin homology domain-containing family A member 3 | -0.37790966 | -0.987769 |
| Q9HB40 | SCPEP1 | Retinoid-inducible serine carboxypeptidase | -0.262325923 | 0.0103674 |
| Q9HB71 | CACYBP | Calcyclin-binding protein | 0.001734416 | -0.0637716 |
| Q9HB90 | RRAGC | Ras-related GTP-binding protein C | 0.132525126 | -0.143806 |
| Q9HBH5 | RDH14 | Retinol dehydrogenase 14 | -1.130713781 | 0.0790555 |
| Q9HBK9 | AS3MT | Arsenite methyltransferase | 0.012004217 | -0.171853 |
| Q9HBL7 | PLGRKT | Plasminogen receptor (KT) | -1.387304703 | -1.66009 |
| Q9HBL8 | NMRAL1 | NmrA-like family domain-containing protein 1 | -0.136048237 | 0.115669 |
| Q9HBM1 | SPC25 | Kinetochore protein Spc25 | -0.033275286 | -0.129114 |
| Q9HC07 | TMEM165 | Transmembrane protein 165 | -2.067167679 | -0.456567 |
| Q9HC38-2 | GLOD4 | Isoform 2 of Glyoxalase domain-containing protein 4 | -0.063585599 | 0.112348 |
| Q9HCD5 | NCOA5 | Nuclear receptor coactivator 5 | 1.106630961 | 1.73518 |
| Q9HCE0 | EPG5 | Ectopic P granules protein 5 homolog | -0.384884357 | -0.174164 |
| Q9HCE1 | MOV10 | Helicase MOV-10 | -0.299404224 | 0.265786 |
| Q9HCE5 | METTL14 | N6-adenosine-methyltransferase non-catalytic subunit | -0.193263292 | -0.0435127 |
| Q9HCG8 | CWC22 | Pre-mRNA-splicing factor CWC22 homolog | -0.285564979 | 0.0935109 |
| Q9HCJ6 | VAT1L | Synaptic vesicle membrane protein VAT-1 homolog-like | 0.278100729 | -0.325945 |
| Q9HCN4 | GPN1 | GPN-loop GTPase 1 | -0.336790323 | -0.0675815 |
| Q9HCN8 | SDF2L1 | Stromal cell-derived factor 2-like protein 1 | -0.290409247 | -0.350303 |
| Q9HCS7 | XAB2 | Pre-mRNA-splicing factor SYF1 | -0.378686746 | 0.0962213 |
| Q9HCU5 | PREB | Prolactin regulatory element-binding protein | -1.420583328 | 0.0830679 |
| Q9HCY8 | S100A14 | Protein S100-A14 | -0.841829459 | 0.214467 |
| Q9HD15 | SRA1 | Steroid receptor RNA activator 1 | 0.059783459 | -0.0100943 |
| Q9HD20 | ATP13A1 | Endoplasmic reticulum transmembrane helix translocase | -1.539484501 | 0.284307 |
| Q9HD26 | GOPC | Golgi-associated PDZ and coiled-coil motif-containing protein | 0.108298461 | -0.0219371 |
| Q9HD33 | MRPL47 | 39S ribosomal protein L47, mitochondrial | -0.34422938 | -0.316915 |

| | | | | |
|---|---|---|---|---|
| Q9HD45 | TM9SF3 | Transmembrane 9 superfamily member 3 | -1.117580175 | 0.287775 |
| Q9HDC9 | APMAP | Adipocyte plasma membrane-associated protein | -1.355473836 | 0.295916 |
| Q9NNW5 | WDR6 | WD repeat-containing protein 6 | -0.001530011 | -0.162975 |
| Q9NP55 | BPIFA1 | BPI fold-containing family A member 1 | 1.666903098 | 4.29759 |
| Q9NP61 | ARFGAP3 | ADP-ribosylation factor GTPase-activating protein 3 | -0.18792359 | 0.132518 |
| Q9NP72 | RAB18 | Ras-related protein Rab-18 | -0.604833444 | -0.0673771 |
| Q9NP77 | SSU72 | RNA polymerase II subunit A C-terminal domain phosphatase SSU72 | 0.040672938 | -0.112705 |
| Q9NP92 | MRPS30 | 39S ribosomal protein S30, mitochondrial | -1.333754738 | 0 |
| Q9NP97 | DYNLRB1 | Dynein light chain roadblock-type 1 | 0.387732665 | 0.663091 |
| Q9NPA0 | EMC7 | ER membrane protein complex subunit 7 | -2.406157335 | -1.77814 |
| Q9NPA8 | ENY2 | Transcription and mRNA export factor ENY2 | 0.202117364 | 0.0684586 |
| Q9NPB8 | GPCPD1 | Glycerophosphocholine phosphodiesterase GPCPD1 | -0.110307852 | 0.499221 |
| Q9NPD3 | EXOSC4 | Exosome complex component RRP41 | -0.142196178 | -0.11132 |
| Q9NPD8 | UBE2T | Ubiquitin-conjugating enzyme E2 T | 0.026800712 | -0.0679859 |
| Q9NPE3 | NOP10 | H/ACA ribonucleoprotein complex subunit 3 | -1.115278641 | -0.479625 |
| Q9NPF4 | OSGEP | Probable tRNA N6-adenosine threonylcarbamoyltransferase | 0.111168305 | -0.0321662 |
| Q9NPH0 | ACP6 | Lysophosphatidic acid phosphatase type 6 | 0.441675822 | -0.178885 |
| Q9NPH2 | ISYNA1 | Inositol-3-phosphate synthase 1 | -0.217781862 | -0.127171 |
| Q9NPI6 | DCP1A | mRNA-decapping enzyme 1A | -0.312491576 | -0.349713 |
| Q9NPJ3 | ACOT13 | Acyl-coenzyme A thioesterase 13 | 0 | -0.190708 |
| Q9NPP4 | NLRC4 | NLR family CARD domain-containing protein 4 | -1.708287001 | -0.603272 |
| Q9NPQ8 | RIC8A | Synembryn-A | 0.080585003 | -0.0713603 |
| Q9NQ29 | LUC7L | Putative RNA-binding protein Luc7-like 1 | -0.005786657 | -0.222328 |
| Q9NQ48 | LZTFL1 | Leucine zipper transcription factor-like protein 1 | 0.166698138 | -0.159212 |
| Q9NQ50 | MRPL40 | 39S ribosomal protein L40, mitochondrial | -1.113395294 | 0.596617 |
| Q9NQ88 | TIGAR | Fructose-2,6-bisphosphatase TIGAR | 0.146677017 | 0.167017 |
| Q9NQC3 | RTN4 | Reticulon-4 | -0.993935744 | 0.0617682 |
| Q9NQC3-5 | RTN4 | Isoform B2 of Reticulon-4 | -1.079754988 | 0 |
| Q9NQC7 | CYLD | Ubiquitin carboxyl-terminal hydrolase CYLD | 0.119110584 | -0.383873 |
| Q9NQE9 | HINT3 | Adenosine 5'-monophosphoramidase HINT3 | -0.890291373 | 0.624171 |
| Q9NQG5 | RPRD1B | Regulation of nuclear pre-mRNA domain-containing protein 1B | 0.164550463 | -0.293985 |
| Q9NQR4 | NIT2 | Omega-amidase NIT2 | 0.083061695 | -0.00431569 |
| Q9NQS1 | AVEN | Cell death regulator Aven | 0 | -0.777383 |
| Q9NQS7 | INCENP | Inner centromere protein | 0.089264711 | 0.142349 |
| Q9NQT5 | EXOSC3 | Exosome complex component RRP40 | 1.393754403 | 0.019112 |
| Q9NQT8 | KIF13B | Kinesin-like protein KIF13B | -0.228723367 | -0.0674264 |
| Q9NQW6 | ANLN | Anillin | -0.905864159 | 0.561151 |
| Q9NQW7-3 | XPNPEP1 | Isoform 3 of Xaa-Pro aminopeptidase 1 | -0.017502785 | -0.0387723 |
| Q9NQX3 | GPHN | Gephyrin | 0.268724759 | 0.10857 |
| Q9NQX7-2 | ITM2C | Isoform 2 of Integral membrane protein 2C | -0.592611869 | -1.52274 |
| Q9NQZ2 | UTP3 | Something about silencing protein 10 | -1.705187639 | 1.60003 |
| Q9NQZ5 | STARD7 | StAR-related lipid transfer protein 7, mitochondrial | 0.114063501 | 0.146225 |
| Q9NR12 | PDLIM7 | PDZ and LIM domain protein 7 | -0.360740105 | 0.149843 |
| Q9NR19 | ACSS2 | Acetyl-coenzyme A synthetase, cytoplasmic | 0.023358186 | 0.191932 |
| Q9NR28 | DIABLO | Diablo IAP-binding mitochondrial protein | -0.19326369 | 0.852439 |
| Q9NR30 | DDX21 | Nucleolar RNA helicase 2 | -0.7810359 | 0.350886 |
| Q9NR31 | SAR1A | GTP-binding protein SAR1a | -0.338064353 | 0.12852 |
| Q9NR45 | NANS | Sialic acid synthase | 0.13676405 | -0.0742842 |
| Q9NR50 | EIF2B3 | Translation initiation factor eIF-2B subunit gamma | -0.287062407 | -0.0258338 |
| Q9NRF2 | SH2B1 | SH2B adapter protein 1 | -0.407837868 | -0.587886 |
| Q9NRF8 | CTPS2 | CTP synthase 2 | -0.012998501 | -0.00336393 |
| Q9NRF9 | POLE3 | DNA polymerase epsilon subunit 3 | 0.161084493 | 0.115495 |
| Q9NRG0 | CHRAC1 | Chromatin accessibility complex protein 1 | 0.305456082 | 0.162861 |
| Q9NRG1 | PRTFDC1 | Phosphoribosyltransferase domain-containing protein 1 | 0.108134667 | -0.0774921 |
| Q9NRG4 | SMYD2 | N-lysine methyltransferase SMYD2 | -0.423517704 | 0.0770248 |
| Q9NRH1 | YAE1 | Protein YAE1 homolog | -0.168998321 | 0.0372969 |
| Q9NRL3 | STRN4 | Striatin-4 | 0.003359238 | -0.040657 |
| Q9NRN7 | AASDHPPT | L-aminoadipate-semialdehyde dehydrogenase-phosphopantetheinyl transferase | 0.022696972 | 0.0049607 |
| Q9NRR5 | UBQLN4 | Ubiquilin-4 | -0.402912935 | -0.0669429 |
| Q9NRS6 | SNX15 | Sorting nexin-15 | 0.089492957 | 0.242166 |
| Q9NRV9 | HEBP1 | Heme-binding protein 1 | 0.037797769 | -0.0621351 |
| Q9NRW3 | APOBEC3C | DNA dC->dU-editing enzyme APOBEC-3C | -1.10589838 | 0.603784 |
| Q9NRX1 | PNO1 | RNA-binding protein PNO1 | -2.516307751 | -0.66641 |
| Q9NRX4 | PHPT1 | 14 kDa phosphohistidine phosphatase | -0.461237431 | 0.0579109 |
| Q9NRX5 | SERINC1 | Serine incorporator 1 | -2.544492126 | 0.472772 |
| Q9NRY4 | ARHGAP35 | Rho GTPase-activating protein 35 | -0.3192722 | 0.417399 |
| Q9NRY5 | FAM114A2 | Protein FAM114A2 | -0.067282677 | -0.227801 |
| Q9NS00 | C1GALT1 | Glycoprotein-N-acetylgalactosamine 3-beta-galactosyltransferase 1 | -0.657953699 | 0.928987 |

| | | | | |
|---|---|---|---|---|
| Q9NS69 | TOMM22 | Mitochondrial import receptor subunit TOM22 homolog | -0.908462842 | 0 |
| Q9NS86 | LANCL2 | LanC-like protein 2 | -0.117711862 | -0.0715666 |
| Q9NS91 | RAD18 | E3 ubiquitin-protein ligase RAD18 | -0.436952035 | -0.0271215 |
| Q9NSB2 | KRT84 | Keratin, type II cuticular Hb4 | -1.093211889 | 0.115296 |
| Q9NSB4 | KRT82 | Keratin, type II cuticular Hb2 | -1.51532348 | 3.67692 |
| Q9NSC5 | HOMER3 | Homer protein homolog 3 | -0.068846623 | -0.362676 |
| Q9NSD9 | FARSB | Phenylalanine--tRNA ligase beta subunit | -0.334781249 | 0.0987161 |
| Q9NSE4 | IARS2 | Isoleucine--tRNA ligase, mitochondrial | -0.429798603 | 0.313037 |
| Q9NSI2 | SLX9 | Ribosome biogenesis protein SLX9 homolog | 0.100955963 | 0.631916 |
| Q9NSK0 | KLC4 | Kinesin light chain 4 | 0.139961322 | 0.936062 |
| Q9NSP4 | CENPM | Centromere protein M | -0.640185118 | 0.143837 |
| Q9NST1 | PNPLA3 | 1-acylglycerol-3-phosphate O-acyltransferase PNPLA3 | -0.962319136 | 0.0415198 |
| Q9NSY1 | BMP2K | BMP-2-inducible protein kinase | -0.450150331 | -0.376751 |
| Q9NT62 | ATG3 | Ubiquitin-like-conjugating enzyme ATG3 | 0.247512499 | 0.00762749 |
| Q9NTI5 | PDS5B | Sister chromatid cohesion protein PDS5 homolog B | -1.244416157 | -0.990192 |
| Q9NTJ3 | SMC4 | Structural maintenance of chromosomes protein 4 | -0.895807028 | 0.091589 |
| Q9NTJ4 | MAN2C1 | Alpha-mannosidase 2C1 | -0.315348466 | 0.23039 |
| Q9NTX5 | ECHDC1 | Ethylmalonyl-CoA decarboxylase | -0.265761455 | 0.0471465 |
| Q9NTZ6 | RBM12 | RNA-binding protein 12 | -0.172372421 | 0.0030276 |
| Q9NU22 | MDN1 | Midasin | -0.609152079 | 3.645 |
| Q9NUG6 | PDRG1 | p53 and DNA damage-regulated protein 1 | 0.412195365 | -0.195144 |
| Q9NUJ1 | ABHD10 | Palmitoyl-protein thioesterase ABHD10, mitochondrial | -1.081348419 | -0.132331 |
| Q9NUL5 | SHFL | Shiftless antiviral inhibitor of ribosomal frameshifting protein | 2.571864764 | -2.52928 |
| Q9NUL7 | DDX28 | Probable ATP-dependent RNA helicase DDX28 | 0 | 0.802566 |
| Q9NUM4 | TMEM106B | Transmembrane protein 106B | -0.808758179 | -0.165596 |
| Q9NUP7 | TRMT13 | tRNA:m(4)X modification enzyme TRM13 homolog | -1.217772086 | 1.0265 |
| Q9NUP9 | LIN7C | Protein lin-7 homolog C | -0.21459492 | -0.695048 |
| Q9NUQ2 | AGPAT5 | 1-acyl-sn-glycerol-3-phosphate acyltransferase epsilon | -1.998685519 | 1.16334 |
| Q9NUQ3 | TXLNG | Gamma-taxilin | -0.802363714 | -0.183956 |
| Q9NUQ6 | SPATS2L | SPATS2-like protein | -1.105133931 | -0.121031 |
| Q9NUQ8 | ABCF3 | ATP-binding cassette sub-family F member 3 | 0.240412553 | 0.080081 |
| Q9NUQ9 | CYRIB | CYFIP-related Rac1 interactor B | -0.021863461 | -0.181548 |
| Q9NUS5 | AP5S1 | AP-5 complex subunit sigma-1 | -0.707606951 | 0.0338291 |
| Q9NUU6 | OTULINL | Inactive ubiquitin thioesterase OTULINL | -1.54192249 | -0.467441 |
| Q9NUY8 | TBC1D23 | TBC1 domain family member 23 | 0.022524357 | -0.00630824 |
| Q9NV31 | IMP3 | U3 small nucleolar ribonucleoprotein protein IMP3 | 0.502332528 | -0.104546 |
| Q9NV35 | NUDT15 | Nucleotide triphosphate diphosphatase NUDT15 | -0.730924209 | 0.0293366 |
| Q9NV56 | MRGBP | MRG/MORF4L-binding protein | -0.66741991 | 1.02151 |
| Q9NV70 | EXOC1 | Exocyst complex component 1 | -0.682041248 | 0.0153662 |
| Q9NVA4 | TMEM184C | Transmembrane protein 184C | -1.874553521 | 0 |
| Q9NVD7 | PARVA | Alpha-parvin | -0.052830458 | -0.104009 |
| Q9NVE7 | PANK4 | 4'-phosphopantetheine phosphatase | -0.016889731 | -0.106982 |
| Q9NVG8 | TBC1D13 | TBC1 domain family member 13 | 0.12246863 | 0.0688591 |
| Q9NVH1 | DNAJC11 | DnaJ homolog subfamily C member 11 | -0.822046041 | 0.29837 |
| Q9NVH2 | INTS7 | Integrator complex subunit 7 | -0.942576806 | -0.465838 |
| Q9NVI7-2 | ATAD3A | Isoform 2 of ATPase family AAA domain-containing protein 3A | -1.198569457 | 0.426765 |
| Q9NVJ2 | ARL8B | ADP-ribosylation factor-like protein 8B | -0.643205325 | 0.379151 |
| Q9NVM4 | PRMT7 | Protein arginine N-methyltransferase 7 | -0.342876355 | -0.134271 |
| Q9NVM6 | DNAJC17 | DnaJ homolog subfamily C member 17 | -0.260195255 | 0.432287 |
| Q9NVM9 | INTS13 | Integrator complex subunit 13 | -0.301142057 | 0.0236432 |
| Q9NVP1 | DDX18 | ATP-dependent RNA helicase DDX18 | -0.486950318 | 0.0708253 |
| Q9NVP2 | ASF1B | Histone chaperone ASF1B | -0.249812126 | 0.0564133 |
| Q9NVQ4 | FAIM | Fas apoptotic inhibitory molecule 1 | 0.168919166 | -0.493616 |
| Q9NVR0 | KLHL11 | Kelch-like protein 11 | -0.453008652 | 0.350622 |
| Q9NVR5 | DNAAF2 | Protein kintoun | -0.518320004 | -0.607625 |
| Q9NVT9 | ARMC1 | Armadillo repeat-containing protein 1 | 0.06999445 | -0.535773 |
| Q9NVV4 | MTPAP | Poly(A) RNA polymerase, mitochondrial | 2.140549262 | 4.7229 |
| Q9NVX2 | NLE1 | Notchless protein homolog 1 | -0.578854005 | -0.43047 |
| Q9NVZ3 | NECAP2 | Adaptin ear-binding coat-associated protein 2 | -0.399763187 | -0.30968 |
| Q9NW08 | POLR3B | DNA-directed RNA polymerase III subunit RPC2 | 0.105009238 | -0.230485 |
| Q9NW13 | RBM28 | RNA-binding protein 28 | -0.616981586 | -1.5663 |
| Q9NW15 | ANO10 | Anoctamin-10 | -1.693126361 | -1.47715 |
| Q9NW64 | RBM22 | Pre-mRNA-splicing factor RBM22 | -0.068198522 | -0.603994 |
| Q9NW68 | BSDC1 | BSD domain-containing protein 1 | 0.329662402 | 0.406173 |
| Q9NW75 | GPATCH2 | G patch domain-containing protein 2 | -0.458167315 | -0.654053 |
| Q9NW82 | WDR70 | WD repeat-containing protein 70 | 0.10450983 | 0.039911 |
| Q9NWB6 | ARGLU1 | Arginine and glutamate-rich protein 1 | -0.009666761 | -0.389796 |
| Q9NWH9 | SLTM | SAFB-like transcription modulator | -0.888691505 | -0.0663268 |
| Q9NWK9 | ZNHIT6 | Box C/D snoRNA protein 1 | 0.366120497 | -1.46575 |
| Q9NWM8 | FKBP14 | Peptidyl-prolyl cis-trans isomerase FKBP14 | -0.015941143 | -0.0501067 |
| Q9NWS0 | PIH1D1 | PIH1 domain-containing protein 1 | -0.120427211 | 0.554515 |

| | | | | |
|---|---|---|---|---|
| Q9NWT1 | PAK1IP1 | p21-activated protein kinase-interacting protein 1 | -1.888035377 | -1.95115 |
| Q9NWT6 | HIF1AN | Hypoxia-inducible factor 1-alpha inhibitor | 0.114813805 | -0.103808 |
| Q9NWV4 | CZIB | CXXC motif containing zinc binding protein | -0.105912606 | -0.410169 |
| Q9NWV8 | BABAM1 | BRISC and BRCA1-A complex member 1 | 0.302670081 | 0.396986 |
| Q9NWX6 | THG1L | Probable tRNA(His) guanylyltransferase | 0.054196755 | -0.198197 |
| Q9NWY4 | HPF1 | Histone PARylation factor 1 | -0.116700093 | 0.220842 |
| Q9NWZ3 | IRAK4 | Interleukin-1 receptor-associated kinase 4 | -0.32531627 | 0.070831 |
| Q9NWZ5 | UCKL1 | Uridine-cytidine kinase-like 1 | -0.112398068 | 0.126804 |
| Q9NX01 | TXNL4B | Thioredoxin-like protein 4B | 0.369741758 | 0.971665 |
| Q9NX08 | COMMD8 | COMM domain-containing protein 8 | -0.332839251 | 1.01706 |
| Q9NX20 | MRPL16 | 39S ribosomal protein L16, mitochondrial | -0.323196093 | 0.742382 |
| Q9NX24 | NHP2 | H/ACA ribonucleoprotein complex subunit 2 | -0.209386746 | 0.207031 |
| Q9NX38 | ABITRAM | Protein Abitram | 0.15717713 | 0.452079 |
| Q9NX40 | OCIAD1 | OCIA domain-containing protein 1 | -0.731352091 | 0.881429 |
| Q9NX46 | ADPRS | ADP-ribosylhydrolase ARH3 | 0.229640643 | -0.00395616 |
| Q9NX47 | MARCHF5 | E3 ubiquitin-protein ligase MARCHF5 | -0.687719027 | -0.181835 |
| Q9NX55 | HYPK | Huntingtin-interacting protein K | -0.087967157 | -0.495341 |
| Q9NX58 | LYAR | Cell growth-regulating nucleolar protein | -0.506256978 | 0.00408936 |
| Q9NX62 | BPNT2 | Golgi-resident adenosine 3',5'-bisphosphate 3'-phosphatase | -2.035669764 | 0.545775 |
| Q9NX74 | DUS2 | tRNA-dihydrouridine(20) synthase [NAD(P)+]-like | 0.002226353 | 0.0109336 |
| Q9NX76 | CMTM6 | CKLF-like MARVEL transmembrane domain-containing protein 6 | -1.263628483 | 1.16712 |
| Q9NXF1 | TEX10 | Testis-expressed protein 10 | -2.627818624 | -0.0695429 |
| Q9NXF7 | DCAF16 | DDB1- and CUL4-associated factor 16 | -0.001474142 | -0.395664 |
| Q9NXG2 | THUMPD1 | THUMP domain-containing protein 1 | 0.108245691 | 0.0270182 |
| Q9NXG6 | P4HTM | Transmembrane prolyl 4-hydroxylase | -1.929836392 | 0.349955 |
| Q9NXH8 | TOR4A | Torsin-4A | -1.20562609 | -0.58602 |
| Q9NXH9 | TRMT1 | tRNA (guanine(26)-N(2))-dimethyltransferase | -0.072972218 | -0.0239347 |
| Q9NXJ5 | PGPEP1 | Pyroglutamyl-peptidase 1 | -0.984135469 | -1.04086 |
| Q9NXK8 | FBXL12 | F-box/LRR-repeat protein 12 | -0.465545495 | -1.2971 |
| Q9NXN4 | GDAP2 | Ganglioside-induced differentiation-associated protein 2 | -0.163050731 | 0.368657 |
| Q9NXR7 | BABAM2 | BRISC and BRCA1-A complex member 2 | -0.392700354 | 0.577881 |
| Q9NXV2 | KCTD5 | BTB/POZ domain-containing protein KCTD5 | -0.041126331 | 0.175378 |
| Q9NXV6 | CDKN2AIP | CDKN2A-interacting protein | -0.583004157 | 0.275 |
| Q9NXW9 | ALKBH4 | Alpha-ketoglutarate-dependent dioxygenase alkB homolog 4 | 0.171973626 | -0.218153 |
| Q9NY12 | GAR1 | H/ACA ribonucleoprotein complex subunit 1 | -0.366408666 | 0.020607 |
| Q9NY27 | PPP4R2 | Serine/threonine-protein phosphatase 4 regulatory subunit 2 | -0.028827429 | 0.1923 |
| Q9NY93 | DDX56 | Probable ATP-dependent RNA helicase DDX56 | -1.351083199 | 0.251403 |
| Q9NYA1-2 | SPHK1 | Isoform 2 of Sphingosine kinase 1 | -0.173769236 | -0.621915 |
| Q9NYB0 | TERF2IP | Telomeric repeat-binding factor 2-interacting protein 1 | -1.129725218 | -2.17871 |
| Q9NYF8 | BCLAF1 | Bcl-2-associated transcription factor 1 | -0.259431839 | -0.10967 |
| Q9NYH9 | UTP6 | U3 small nucleolar RNA-associated protein 6 homolog | -1.434607267 | 0.367649 |
| Q9NYJ8 | TAB2 | TGF-beta-activated kinase 1 and MAP3K7-binding protein 2 | -0.303335905 | -0.674473 |
| Q9NYK5 | MRPL39 | 39S ribosomal protein L39, mitochondrial | -1.721479416 | -0.338084 |
| Q9NYL2-2 | MAP3K20 | Isoform 2 of Mitogen-activated protein kinase kinase kinase 20 | -0.023114125 | 0.039086 |
| Q9NYL9 | TMOD3 | Tropomodulin-3 | 0.261553923 | -0.0664768 |
| Q9NYP9 | MIS18A | Protein Mis18-alpha | -2.500175476 | 0.425498 |
| Q9NYQ8 | FAT2 | Protocadherin Fat 2 | -0.346392234 | -1.6112 |
| Q9NYS0 | NKIRAS1 | NF-kappa-B inhibitor-interacting Ras-like protein 1 | -0.388329347 | -0.126811 |
| Q9NYU1 | UGGT2 | UDP-glucose:glycoprotein glucosyltransferase 2 | -1.199949463 | 0 |
| Q9NYU2 | UGGT1 | UDP-glucose:glycoprotein glucosyltransferase 1 | -0.186077436 | 0.0374212 |
| Q9NYY8 | FASTKD2 | FAST kinase domain-containing protein 2, mitochondrial | -1.353731871 | -1.23371 |
| Q9NZ01 | TECR | Very-long-chain enoyl-CoA reductase | -1.259899775 | -0.0763763 |
| Q9NZ08 | ERAP1 | Endoplasmic reticulum aminopeptidase 1 | 0.038127184 | 0.209622 |
| Q9NZ09 | UBAP1 | Ubiquitin-associated protein 1 | 0.268361092 | 0.144383 |
| Q9NZ32 | ACTR10 | Actin-related protein 10 | -0.167077223 | -0.0950998 |
| Q9NZ45 | CISD1 | CDGSH iron-sulfur domain-containing protein 1 | -0.795970758 | 0.188717 |
| Q9NZ52 | GGA3 | ADP-ribosylation factor-binding protein GGA3 | -0.540857077 | -1.56961 |
| Q9NZ63 | C9orf78 | Telomere length and silencing protein 1 homolog | -0.240676721 | 0.0218105 |
| Q9NZB2 | FAM120A | Constitutive coactivator of PPAR-gamma-like protein 1 | -0.462710063 | 0.320492 |
| Q9NZD2 | GLTP | Glycolipid transfer protein | -0.01588273 | -0.0480124 |
| Q9NZD8 | SPG21 | Maspardin | -0.084587336 | -0.102174 |
| Q9NZH8 | IL36G | Interleukin-36 gamma | 0.474379619 | 0.270557 |
| Q9NZI7 | UBP1 | Upstream-binding protein 1 | 0.32867988 | -1.52785 |
| Q9NZI8 | IGF2BP1 | Insulin-like growth factor 2 mRNA-binding protein 1 | -0.751895428 | -0.0573982 |
| Q9NZJ9 | NUDT4 | Diphosphoinositol polyphosphate phosphohydrolase 2 | -0.375989755 | -0.862484 |
| Q9NZL4 | HSPBP1 | Hsp70-binding protein 1 | -0.243225098 | -0.00402514 |
| Q9NZL9 | MAT2B | Methionine adenosyltransferase 2 subunit beta | 0.071375052 | -0.0644417 |
| Q9NZL9-2 | MAT2B | Isoform 2 of Methionine adenosyltransferase 2 subunit beta | -0.457484643 | -0.297655 |
| Q9NZM1 | MYOF | Myoferlin | -1.486946821 | 0.227507 |
| Q9NZM5 | NOP53 | Ribosome biogenesis protein NOP53 | 0 | -2.74371 |
| Q9NZN3 | EHD3 | EH domain-containing protein 3 | 0.307136536 | -1.04563 |

| | | | | |
|---|---|---|---|---|
| Q9NZN4 | EHD2 | EH domain-containing protein 2 | -0.67103672 | 0.12045 |
| Q9NZP5 | OR5AC2 | Olfactory receptor 5AC2 | -1.795022647 | 1.49923 |
| Q9NZQ3 | NCKIPSD | NCK-interacting protein with SH3 domain | -0.468628089 | -0.565434 |
| Q9NZT1 | CALML5 | Calmodulin-like protein 5 | -0.04808569 | 0.280952 |
| Q9NZT2 | OGFR | Opioid growth factor receptor | 0.577368418 | 0.116266 |
| Q9NZU5 | LMCD1 | LIM and cysteine-rich domains protein 1 | 0.179075638 | -0.0123323 |
| Q9NZV1 | CRIM1 | Cysteine-rich motor neuron 1 protein | -0.328718821 | 0.927634 |
| Q9NZZ3 | CHMP5 | Charged multivesicular body protein 5 | 0.078827222 | -0.203602 |
| Q9P000 | COMMD9 | COMM domain-containing protein 9 | -0.061765194 | -0.207461 |
| Q9P013 | CWC15 | Spliceosome-associated protein CWC15 homolog | 0.212763151 | 0.177778 |
| Q9P015 | MRPL15 | 39S ribosomal protein L15, mitochondrial | -1.145491044 | 0.036972 |
| Q9P016 | THYN1 | Thymocyte nuclear protein 1 | -0.450385094 | 0.481538 |
| Q9P032 | NDUFAF4 | NADH dehydrogenase [ubiquinone] 1 alpha subcomplex assembly factor 4 | -1.984217564 | 1.49963 |
| Q9P035 | HACD3 | Very-long-chain (3R)-3-hydroxyacyl-CoA dehydratase 3 | -1.535236279 | 0.10226 |
| Q9P0I2 | EMC3 | ER membrane protein complex subunit 3 | -1.732316891 | -0.116358 |
| Q9P0J0 | NDUFA13 | NADH dehydrogenase [ubiquinone] 1 alpha subcomplex subunit 13 | -0.978360494 | 0.665795 |
| Q9P0J1 | PDP1 | [Pyruvate dehydrogenase [acetyl-transferring]]-phosphatase 1, mitochondrial | 0.085643371 | -2.58976 |
| Q9P0J7 | KCMF1 | E3 ubiquitin-protein ligase KCMF1 | -0.557352622 | -0.821174 |
| Q9P0K7 | RAI14 | Ankycorbin | -1.106382529 | 0.305117 |
| Q9P0L0 | VAPA | Vesicle-associated membrane protein-associated protein A | -1.124674241 | 0.10452 |
| Q9P0M6 | MACROH2A2 | Core histone macro-H2A.2 | -0.942075173 | -0.217769 |
| Q9P0S9 | TMEM14C | Transmembrane protein 14C | -2.264150143 | -0.277329 |
| Q9P0V3 | SH3BP4 | SH3 domain-binding protein 4 | -1.510357618 | 0.58076 |
| Q9P107 | GMIP | GEM-interacting protein | -1.768449982 | 0.809874 |
| Q9P1F3 | ABRACL | Costars family protein ABRACL | -1.150217851 | 3.61706 |
| Q9P1Z2 | CALCOCO1 | Calcium-binding and coiled-coil domain-containing protein 1 | 0.107666254 | -0.341402 |
| Q9P253 | VPS18 | Vacuolar protein sorting-associated protein 18 homolog | -0.0562102 | 0.28393 |
| Q9P258 | RCC2 | Protein RCC2 | -0.758579413 | 0.185521 |
| Q9P265 | DIP2B | Disco-interacting protein 2 homolog B | -1.008903583 | -0.508916 |
| Q9P270 | SLAIN2 | SLAIN motif-containing protein 2 | -0.055313349 | 0.143157 |
| Q9P287 | BCCIP | BRCA2 and CDKN1A-interacting protein | -0.096057653 | 0.0585108 |
| Q9P2B4 | CTTNBP2NL | CTTNBP2 N-terminal-like protein | 0.113556385 | -0.125764 |
| Q9P2D0-3 | IBTK | Isoform 3 of Inhibitor of Bruton tyrosine kinase | -1.522703727 | 0.139423 |
| Q9P2D3 | HEATR5B | HEAT repeat-containing protein 5B | -0.09001565 | -0.721665 |
| Q9P2E3 | ZNFX1 | NFX1-type zinc finger-containing protein 1 | -0.01047039 | -2.80612 |
| Q9P2E9 | RRBP1 | Ribosome-binding protein 1 | -1.413800001 | 0.138614 |
| Q9P2G1 | ANKIB1 | Ankyrin repeat and IBR domain-containing protein 1 | 0.508303404 | -0.476423 |
| Q9P2I0 | CPSF2 | Cleavage and polyadenylation specificity factor subunit 2 | -1.377050718 | 0.306156 |
| Q9P2J3 | KLHL9 | Kelch-like protein 9 | -0.85668389 | 0.214105 |
| Q9P2K8 | EIF2AK4 | eIF-2-alpha kinase GCN2 | 0.092197498 | -0.212403 |
| Q9P2N5 | RBM27 | RNA-binding protein 27 | -0.249941985 | 0.80665 |
| Q9P2Q2 | FRMD4A | FERM domain-containing protein 4A | -1.230206331 | -2.37642 |
| Q9P2R3 | ANKFY1 | Rabankyrin-5 | 0.022724152 | -0.0816005 |
| Q9UBB4 | ATXN10 | Ataxin-10 | -0.091291746 | 0.113575 |
| Q9UBB5 | MBD2 | Methyl-CpG-binding domain protein 2 | -2.213617166 | -2.22384 |
| Q9UBB6-2 | NCDN | Isoform 2 of Neurochondrin | -0.108734926 | -0.0841233 |
| Q9UBB6 | NCDN | Neurochondrin | -0.678509235 | 0.219991 |
| Q9UBB9 | TFIP11 | Tuftelin-interacting protein 11 | -0.132194757 | 0.216129 |
| Q9UBC2 | EPS15L1 | Epidermal growth factor receptor substrate 15-like 1 | 0.20702521 | 0.0188297 |
| Q9UBD5 | ORC3 | Origin recognition complex subunit 3 | -1.885411183 | -1.79862 |
| Q9UBE0 | SAE1 | SUMO-activating enzyme subunit 1 | 0.011459986 | -0.0893612 |
| Q9UBF2 | COPG2 | Coatomer subunit gamma-2 | 0.07851243 | -0.00822481 |
| Q9UBF6 | RNF7 | RING-box protein 2 | -0.461216609 | 0.793354 |
| Q9UBG0 | MRC2 | C-type mannose receptor 2 | -1.016986211 | -1.16646 |
| Q9UBG3 | CRNN | Cornulin | -1.41941158 | 2.42666 |
| Q9UBI1 | COMMD3 | COMM domain-containing protein 3 | -0.297180971 | -0.149326 |
| Q9UBI6 | GNG12 | Guanine nucleotide-binding protein G(I)/G(S)/G(O) subunit gamma-12 | 0 | -0.170997 |
| Q9UBK7 | RABL2A | Rab-like protein 2A | -1.244258483 | 0.430216 |
| Q9UBK8 | MTRR | Methionine synthase reductase | -0.164014339 | 0.140304 |
| Q9UBK9 | UXT | Protein UXT | -0.226371845 | 0.0212129 |
| Q9UBP6 | METTL1 | tRNA (guanine-N(7)-)-methyltransferase | -0.004746119 | 0.112165 |
| Q9UBQ0 | VPS29 | Vacuolar protein sorting-associated protein 29 | -0.136882464 | -0.0701021 |
| Q9UBQ5 | EIF3K | Eukaryotic translation initiation factor 3 subunit K | -0.239738941 | 0.105639 |
| Q9UBQ6 | EXTL2 | Exostosin-like 2 | -0.781914234 | 0.100715 |
| Q9UBQ7 | GRHPR | Glyoxylate reductase/hydroxypyruvate reductase | -0.048598448 | -0.0298322 |
| Q9UBS4 | DNAJB11 | DnaJ homolog subfamily B member 11 | -0.299084981 | 0.112158 |
| Q9UBS8 | RNF14 | E3 ubiquitin-protein ligase RNF14 | 0.035010974 | 0.0636549 |

| | | | | |
|---|---|---|---|---|
| Q9UBT2 | UBA2 | SUMO-activating enzyme subunit 2 | 0.080535412 | 0.077865 |
| Q9UBU9 | NXF1 | Nuclear RNA export factor 1 | 0.02746733 | -0.0293032 |
| Q9UBV8 | PEF1 | Peflin | -0.265526136 | 0.5688 |
| Q9UBW8 | COPS7A | COP9 signalosome complex subunit 7a | 0.226619562 | -0.0317307 |
| Q9UBX3 | SLC25A10 | Mitochondrial dicarboxylate carrier | 0.672817866 | 0.378537 |
| Q9UDT6 | CLIP2 | CAP-Gly domain-containing linker protein 2 | -0.067229112 | 0.334833 |
| Q9UDW1 | UQCR10 | Cytochrome b-c1 complex subunit 9 | -1.022934834 | -0.258747 |
| Q9UDY2 | TJP2 | Tight junction protein ZO-2 | 0.017939885 | 0.0611935 |
| Q9UDY4 | DNAJB4 | DnaJ homolog subfamily B member 4 | -0.319825172 | 0.121117 |
| Q9UDY8 | MALT1 | Mucosa-associated lymphoid tissue lymphoma translocation protein 1 | 0.095770597 | 0.118973 |
| Q9UEE5 | STK17A | Serine/threonine-protein kinase 17A | -0.365171671 | -0.217024 |
| Q9UEE9 | CFDP1 | Craniofacial development protein 1 | -0.05401667 | -0.0176786 |
| Q9UER7 | DAXX | Death domain-associated protein 6 | -0.566366673 | 0.336748 |
| Q9UET6 | FTSJ1 | Putative tRNA (cytidine(32)/guanosine(34)-2'-O)-methyltransferase | -0.074026982 | 0.0715685 |
| Q9UEW8 | STK39 | STE20/SPS1-related proline-alanine-rich protein kinase | 0.034181595 | 0.0605793 |
| Q9UEY8 | ADD3 | Gamma-adducin | -0.238965829 | -0.238496 |
| Q9UFC0 | LRWD1 | Leucine-rich repeat and WD repeat-containing protein 1 | -2.930097183 | 1.0125 |
| Q9UFN0 | NIPSNAP3A | Protein NipSnap homolog 3A | -0.805871169 | 0.357043 |
| Q9UFW8 | CGGBP1 | CGG triplet repeat-binding protein 1 | -0.841823022 | -0.177028 |
| Q9UG56-2 | PISD | Isoform 2 of Phosphatidylserine decarboxylase proenzyme, mitochondrial | -1.006583214 | -0.051829 |
| Q9UG63 | ABCF2 | ATP-binding cassette sub-family F member 2 | -0.327002128 | -0.821427 |
| Q9UGI8 | TES | Testin | 0.15770491 | 0.104873 |
| Q9UGP4 | LIMD1 | LIM domain-containing protein 1 | 0.033865531 | 0.156799 |
| Q9UGP8 | SEC63 | Translocation protein SEC63 homolog | -1.671814839 | 0.060407 |
| Q9UGR2 | ZC3H7B | Zinc finger CCCH domain-containing protein 7B | -0.789833466 | -2.22159 |
| Q9UGV2 | NDRG3 | Protein NDRG3 | 0.037873745 | -0.0698325 |
| Q9UH62 | ARMCX3 | Armadillo repeat-containing X-linked protein 3 | -0.272698959 | 0.0532379 |
| Q9UH65 | SWAP70 | Switch-associated protein 70 | 0.010725419 | -0.00351556 |
| Q9UH99 | SUN2 | SUN domain-containing protein 2 | -0.304699659 | 0.798814 |
| Q9UHA4 | LAMTOR3 | Ragulator complex protein LAMTOR3 | -1.362286806 | -2.31137 |
| Q9UHB4 | NDOR1 | NADPH-dependent diflavin oxidoreductase 1 | -0.261956533 | 0.60097 |
| Q9UHB6 | LIMA1 | LIM domain and actin-binding protein 1 | 0.199852864 | 0.281754 |
| Q9UHB6-4 | LIMA1 | Isoform 4 of LIM domain and actin-binding protein 1 | 0.101138035 | -0.275914 |
| Q9UHB7 | AFF4 | AF4/FMR2 family member 4 | -1.039753596 | 0.219101 |
| Q9UHB9 | SRP68 | Signal recognition particle subunit SRP68 | 0.087339719 | -0.198961 |
| Q9UHD1 | CHORDC1 | Cysteine and histidine-rich domain-containing protein 1 | -0.116601944 | -0.12741 |
| Q9UHD8 | SEPTIN9 | Septin-9 | -0.621340434 | -0.0250152 |
| Q9UHD9 | UBQLN2 | Ubiquilin-2 | 0.142619212 | -0.434635 |
| Q9UHG3 | PCYOX1 | Prenylcysteine oxidase 1 | -1.350919247 | 0.314252 |
| Q9UHL4 | DPP7 | Dipeptidyl peptidase 2 | 0.157123804 | -0.0520716 |
| Q9UHP3 | USP25 | Ubiquitin carboxyl-terminal hydrolase 25 | 0.43266654 | -0.642863 |
| Q9UHQ9 | CYB5R1 | NADH-cytochrome b5 reductase 1 | -2.145056804 | 0.672622 |
| Q9UHR6 | ZNHIT2 | Zinc finger HIT domain-containing protein 2 | -0.285449107 | 0.285657 |
| Q9UHV9 | PFDN2 | Prefoldin subunit 2 | 0.448103269 | 0.252177 |
| Q9UHW5 | GPN3 | GPN-loop GTPase 3 | -0.216614326 | 2.92885 |
| Q9UHX1-5 | PUF60 | Isoform 5 of Poly(U)-binding-splicing factor PUF60 | 0.240165313 | 0.911666 |
| Q9UHY1 | NRBP1 | Nuclear receptor-binding protein | 0.202125311 | -0.0546983 |
| Q9UHY7 | ENOPH1 | Enolase-phosphatase E1 | 0.004800479 | 0.00252183 |
| Q9UI09 | NDUFA12 | NADH dehydrogenase [ubiquinone] 1 alpha subcomplex subunit 12 | -1.53636233 | 0.171051 |
| Q9UI12 | ATP6V1H | V-type proton ATPase subunit H | -0.023689906 | -0.144859 |
| Q9UI15 | TAGLN3 | Transgelin-3 | 0.197274049 | 0.0396932 |
| Q9UI26 | IPO11 | Importin-11 | -0.078998645 | -0.132149 |
| Q9UI30 | TRMT112 | Multifunctional methyltransferase subunit TRM112-like protein | -0.32149148 | 0.819458 |
| Q9UI42 | CPA4 | Carboxypeptidase A4 | -0.120758772 | 0.429468 |
| Q9UIA9 | XPO7 | Exportin-7 | -0.041114171 | -0.203931 |
| Q9UIC8 | LCMT1 | Leucine carboxyl methyltransferase 1 | 0.084941626 | -0.016065 |
| Q9UID3 | VPS51 | Vacuolar protein sorting-associated protein 51 homolog | -0.555221001 | 0.0130682 |
| Q9UIG0 | BAZ1B | Tyrosine-protein kinase BAZ1B | -1.070836306 | 0.499798 |
| Q9UII2 | ATP5IF1 | ATPase inhibitor, mitochondrial | -1.103760481 | 0.280485 |
| Q9UIJ7 | AK3 | GTP:AMP phosphotransferase AK3, mitochondrial | -0.338353395 | -0.157271 |
| Q9UIQ6 | LNPEP | Leucyl-cystinyl aminopeptidase | -0.644364993 | 0.421517 |
| Q9UIV1 | CNOT7 | CCR4-NOT transcription complex subunit 7 | -0.474535942 | 0.109815 |
| Q9UJ68 | MSRA | Mitochondrial peptide methionine sulfoxide reductase | -0.000280698 | -0.381569 |
| Q9UJ70 | NAGK | N-acetyl-D-glucosamine kinase | 0.083595594 | -0.122216 |
| Q9UJ83 | HACL1 | 2-hydroxyacyl-CoA lyase 1 | -0.187059323 | 0.599507 |
| Q9UJA5 | TRMT6 | tRNA (adenine(58)-N(1))-methyltransferase non-catalytic subunit TRM6 | -0.086040258 | -0.247638 |

| | | | | |
|---|---|---|---|---|
| Q9UJC5 | SH3BGRL2 | SH3 domain-binding glutamic acid-rich-like protein 2 | -1.582395236 | 0.584897 |
| Q9UJK0 | TSR3 | 18S rRNA aminocarboxypropyltransferase | -0.228981733 | 0.450135 |
| Q9UJP4 | KLHL21 | Kelch-like protein 21 | -0.434736649 | 0.0539595 |
| Q9UJS0 | SLC25A13 | Calcium-binding mitochondrial carrier protein Aralar2 | -0.696810643 | 0.101859 |
| Q9UJU6 | DBNL | Drebrin-like protein | 0.243214766 | 0.0492102 |
| Q9UJU6-2 | DBNL | Isoform 2 of Drebrin-like protein | -2.29297026 | 0.391631 |
| Q9UJV9 | DDX41 | Probable ATP-dependent RNA helicase DDX41 | -0.074369589 | 0.136185 |
| Q9UJW0 | DCTN4 | Dynactin subunit 4 | -0.010467688 | -0.152972 |
| Q9UJX2 | CDC23 | Cell division cycle protein 23 homolog | -0.387779951 | 0.247888 |
| Q9UJX3 | ANAPC7 | Anaphase-promoting complex subunit 7 | -0.719131549 | 0.530919 |
| Q9UJX5 | ANAPC4 | Anaphase-promoting complex subunit 4 | -0.942550262 | 0.320302 |
| Q9UJX6 | ANAPC2 | Anaphase-promoting complex subunit 2 | 0.37572066 | -2.08899 |
| Q9UJY4 | GGA2 | ADP-ribosylation factor-binding protein GGA2 | -0.004940748 | -0.0339788 |
| Q9UJY5 | GGA1 | ADP-ribosylation factor-binding protein GGA1 | 0.306564331 | 0.0320292 |
| Q9UJZ1 | STOML2 | Stomatin-like protein 2, mitochondrial | -0.931699753 | 0.282405 |
| Q9UK23 | NAGPA | N-acetylglucosamine-1-phosphodiester alpha-N-acetylglucosaminidase | 0.404753288 | 0.450853 |
| Q9UK39 | NOCT | Nocturnin | -0.268371105 | 0.225394 |
| Q9UK41 | VPS28 | Vacuolar protein sorting-associated protein 28 homolog | -0.253824234 | 0.390538 |
| Q9UK45 | LSM7 | U6 snRNA-associated Sm-like protein LSm7 | -0.829085906 | 0.476615 |
| Q9UK58 | CCNL1 | Cyclin-L1 | -0.524288336 | -0.528885 |
| Q9UK59 | DBR1 | Lariat debranching enzyme | -0.194021066 | -0.0657352 |
| Q9UK73 | FEM1B | Protein fem-1 homolog B | -1.019506296 | 0 |
| Q9UK76 | JPT1 | Jupiter microtubule associated homolog 1 | -0.400913715 | 1.09661 |
| Q9UK97 | FBXO9 | F-box only protein 9 | 0.159347057 | 2.31994 |
| Q9UKC9 | FBXL2 | F-box/LRR-repeat protein 2 | 6.756603003 | 0 |
| Q9UKD2 | MRTO4 | mRNA turnover protein 4 homolog | -0.20340244 | -0.233003 |
| Q9UKG1 | APPL1 | DCC-interacting protein 13-alpha | 0.010183891 | -0.102482 |
| Q9UKJ3 | GPATCH8 | G patch domain-containing protein 8 | 0.008770784 | 0.156274 |
| Q9UKK3 | PARP4 | Protein mono-ADP-ribosyltransferase PARP4 | 0.208902518 | -0.00714493 |
| Q9UKK6 | NXT1 | NTF2-related export protein 1 | -0.438601812 | -0.971592 |
| Q9UKL0 | RCOR1 | REST corepressor 1 | -0.194587708 | 0.120456 |
| Q9UKL6 | PCTP | Phosphatidylcholine transfer protein | 0.526362578 | -4.05734 |
| Q9UKM9 | RALY | RNA-binding protein Raly | -0.699044625 | 0.525084 |
| Q9UKN8 | GTF3C4 | General transcription factor 3C polypeptide 4 | -0.18142271 | -0.0733836 |
| Q9UKR5 | ERG28 | Ergosterol biosynthetic protein 28 homolog | -1.722812096 | -0.0904668 |
| Q9UKT7 | FBXL3 | F-box/LRR-repeat protein 3 | -1.125294328 | -0.3943 |
| Q9UKU7 | ACAD8 | Isobutyryl-CoA dehydrogenase, mitochondrial | -1.691551288 | 1.36904 |
| Q9UKV5 | AMFR | E3 ubiquitin-protein ligase AMFR | -1.939371347 | 0.0291411 |
| Q9UKV8 | AGO2 | Protein argonaute-2 | 0 | 2.16546 |
| Q9UKX7 | NUP50 | Nuclear pore complex protein Nup50 | 0.067154884 | 0.15073 |
| Q9UKY7 | CDV3 | Protein CDV3 homolog | 0.144569874 | -0.0984071 |
| Q9UKZ1 | CNOT11 | CCR4-NOT transcription complex subunit 11 | -0.665675243 | 3.24604 |
| Q9UL15 | BAG5 | BAG family molecular chaperone regulator 5 | 0.142234166 | -0.144479 |
| Q9UL25 | RAB21 | Ras-related protein Rab-21 | -0.684843302 | -0.0131741 |
| Q9UL26 | RAB22A | Ras-related protein Rab-22A | -1.010071198 | 0.0962884 |
| Q9UL63 | MKLN1 | Muskelin | 0.069003979 | -2.60861 |
| Q9ULC3 | RAB23 | Ras-related protein Rab-23 | -1.866002361 | 0.165319 |
| Q9ULC4 | MCTS1 | Malignant T-cell-amplified sequence 1 | -0.124546051 | -0.0163746 |
| Q9ULJ3 | ZBTB21 | Zinc finger and BTB domain-containing protein 21 | -0.487354 | 0 |
| Q9ULJ6 | ZMIZ1 | Zinc finger MIZ domain-containing protein 1 | -0.943993886 | -0.66312 |
| Q9ULJ7 | ANKRD50 | Ankyrin repeat domain-containing protein 50 | 0.520345291 | 0.12809 |
| Q9ULQ0 | STRIP2 | Striatin-interacting protein 2 | -0.052990278 | -0.896173 |
| Q9ULR0 | ISY1 | Pre-mRNA-splicing factor ISY1 homolog | -0.439695915 | -0.17543 |
| Q9ULT8 | HECTD1 | E3 ubiquitin-protein ligase HECTD1 | -0.000389973 | 0.0255658 |
| Q9ULV4 | CORO1C | Coronin-1C | 0.022682985 | 0.0913556 |
| Q9ULW0 | TPX2 | Targeting protein for Xklp2 | -1.952437401 | -0.0439835 |
| Q9ULW3 | ABT1 | Activator of basal transcription 1 | -1.041433652 | 0 |
| Q9ULX9 | MAFF | Transcription factor MafF | -1.67992576 | -1.53673 |
| Q9ULZ3 | PYCARD | Apoptosis-associated speck-like protein containing a CARD | 0.810074488 | -0.0292428 |
| Q9UM00 | TMCO1 | Calcium load-activated calcium channel | -1.529171944 | 0.37088 |
| Q9UM22 | EPDR1 | Mammalian ependymin-related protein 1 | 0.15356644 | -0.0581719 |
| Q9UMR2 | DDX19B | ATP-dependent RNA helicase DDX19B | -0.668063879 | 2.04484 |
| Q9UMS0 | NFU1 | NFU1 iron-sulfur cluster scaffold homolog, mitochondrial | 0.284067233 | 0.0629161 |
| Q9UMS4 | PRPF19 | Pre-mRNA-processing factor 19 | -0.422384501 | 0.146324 |
| Q9UMX0 | UBQLN1 | Ubiquilin-1 | 0.043447495 | -0.00045077 |
| Q9UMX1 | SUFU | Suppressor of fused homolog | 0.561816613 | -1.14051 |
| Q9UMX5 | NENF | Neudesin | -0.436753194 | -0.285262 |
| Q9UMY1 | NOL7 | Nucleolar protein 7 | -0.198938688 | 0.451881 |
| Q9UMZ2 | SYNRG | Synergin gamma | 0.192408641 | 0.220409 |
| Q9UN37 | VPS4A | Vacuolar protein sorting-associated protein 4A | 0.144519965 | -0.197345 |

| | | | | |
|---|---|---|---|---|
| Q9UN86 | G3BP2 | Ras GTPase-activating protein-binding protein 2 | -0.472768545 | -0.082833 |
| Q9UNE7 | STUB1 | E3 ubiquitin-protein ligase CHIP | 0.01830252 | 0.0383155 |
| Q9UNF0 | PACSIN2 | Protein kinase C and casein kinase substrate in neurons protein 2 | -0.117886225 | -0.191369 |
| Q9UNF1 | MAGED2 | Melanoma-associated antigen D2 | 0.087537209 | -0.372336 |
| Q9UNH6 | SNX7 | Sorting nexin-7 | 0.159239531 | -0.82045 |
| Q9UNH7 | SNX6 | Sorting nexin-6 | 0.166574558 | 0.00837866 |
| Q9UNI6 | DUSP12 | Dual specificity protein phosphatase 12 | 0.015243133 | -0.0864798 |
| Q9UNM6 | PSMD13 | 26S proteasome non-ATPase regulatory subunit 13 | -0.242408117 | -0.115407 |
| Q9UNN5 | FAF1 | FAS-associated factor 1 | 0.072689692 | 0.0260595 |
| Q9UNP9 | PPIE | Peptidyl-prolyl cis-trans isomerase E | -0.467839162 | 0.0595932 |
| Q9UNS1 | TIMELESS | Protein timeless homolog | 0.113540252 | 0.0429831 |
| Q9UNS2 | COPS3 | COP9 signalosome complex subunit 3 | -0.008550008 | 0.216925 |
| Q9UNW1 | MINPP1 | Multiple inositol polyphosphate phosphatase 1 | -0.147181193 | 0.139458 |
| Q9UNX3 | RPL26L1 | 60S ribosomal protein L26-like 1 | -1.472874959 | -0.104386 |
| Q9UNX4 | WDR3 | WD repeat-containing protein 3 | -0.988124609 | 0.298433 |
| Q9UNZ2 | NSFL1C | NSFL1 cofactor p47 | 0.277368069 | 0.0492045 |
| Q9UNZ2-5 | NSFL1C | Isoform 3 of NSFL1 cofactor p47 | 0.186143955 | 0.601246 |
| Q9UP65 | PLA2G4C | Cytosolic phospholipase A2 gamma | -1.612623374 | -0.818094 |
| Q9UP83 | COG5 | Conserved oligomeric Golgi complex subunit 5 | -0.079412858 | 0.193271 |
| Q9UPM8 | AP4E1 | AP-4 complex subunit epsilon-1 | -0.03193442 | -0.0556428 |
| Q9UPN7 | PPP6R1 | Serine/threonine-protein phosphatase 6 regulatory subunit 1 | -0.092861493 | -0.187486 |
| Q9UPN9 | TRIM33 | E3 ubiquitin-protein ligase TRIM33 | 0.030004899 | -0.210605 |
| Q9UPR3 | SMG5 | Nonsense-mediated mRNA decay factor SMG5 | -0.131438732 | -1.20414 |
| Q9UPU5 | USP24 | Ubiquitin carboxyl-terminal hydrolase 24 | -0.042109569 | -0.0760746 |
| Q9UPU7 | TBC1D2B | TBC1 domain family member 2B | 0.138819536 | -0.194866 |
| Q9UPZ3 | HPS5 | Hermansky-Pudlak syndrome 5 protein | 0.343533913 | -1.71525 |
| Q9UQ03 | CORO2B | Coronin-2B | -1.879238526 | 0 |
| Q9UQ13 | SHOC2 | Leucine-rich repeat protein SHOC-2 | -0.320054173 | 1.2018 |
| Q9UQ35 | SRRM2 | Serine/arginine repetitive matrix protein 2 | 0.142148495 | -0.161399 |
| Q9UQ80 | PA2G4 | Proliferation-associated protein 2G4 | -0.418331623 | -0.0839895 |
| Q9UQE7 | SMC3 | Structural maintenance of chromosomes protein 3 | -0.719964504 | 0.309384 |
| Q9UQQ2 | SH2B3 | SH2B adapter protein 3 | 0.496341944 | 0 |
| Q9Y217 | MTMR6 | Myotubularin-related protein 6 | -0.756632725 | -0.217237 |
| Q9Y221 | NIP7 | 60S ribosome subunit biogenesis protein NIP7 homolog | -0.760515531 | 0.300169 |
| Q9Y223 | GNE | Bifunctional UDP-N-acetylglucosamine 2-epimerase/N-acetylmannosamine kinase | 0.023477952 | -0.0396922 |
| Q9Y224 | RTRAF | RNA transcription, translation and transport factor protein | -0.315798362 | 0.581964 |
| Q9Y230 | RUVBL2 | RuvB-like 2 | 0.081302484 | 0.0930379 |
| Q9Y237 | PIN4 | Peptidyl-prolyl cis-trans isomerase NIMA-interacting 4 | -1.191687425 | 0.571602 |
| Q9Y243 | AKT3 | RAC-gamma serine/threonine-protein kinase | 0.129750172 | -0.192275 |
| Q9Y248 | GINS2 | DNA replication complex GINS protein PSF2 | -0.014451186 | 0 |
| Q9Y259 | CHKB | Choline/ethanolamine kinase | 0.193386714 | 0.217786 |
| Q9Y263 | PLAA | Phospholipase A-2-activating protein | 0.053569953 | -0.0343736 |
| Q9Y265 | RUVBL1 | RuvB-like 1 | -0.051598867 | 0.116451 |
| Q9Y266 | NUDC | Nuclear migration protein nudC | 0.180461566 | 0.0128498 |
| Q9Y276 | BCS1L | Mitochondrial chaperone BCS1 | 0.003814697 | 0.00351238 |
| Q9Y277 | VDAC3 | Voltage-dependent anion-selective channel protein 3 | -0.901885748 | 0.382324 |
| Q9Y281 | CFL2 | Cofilin-2 | 0.062023958 | 0.138916 |
| Q9Y285 | FARSA | Phenylalanine--tRNA ligase alpha subunit | -0.64858381 | -0.0180175 |
| Q9Y294 | ASF1A | Histone chaperone ASF1A | -0.031848113 | -0.198721 |
| Q9Y295 | DRG1 | Developmentally-regulated GTP-binding protein 1 | -0.538322608 | 0.0879869 |
| Q9Y2A7 | NCKAP1 | Nck-associated protein 1 | -0.405424992 | -0.227514 |
| Q9Y2B0 | CNPY2 | Protein canopy homolog 2 | 0.067888101 | 0.770426 |
| Q9Y2C4 | EXOG | Nuclease EXOG, mitochondrial | -0.068828265 | 0.129102 |
| Q9Y2D5-6 | AKAP2 | Isoform 4 of A-kinase anchor protein 2 | 0.263304154 | 0.170403 |
| Q9Y2G5 | POFUT2 | GDP-fucose protein O-fucosyltransferase 2 | -0.521354198 | 0.414378 |
| Q9Y2H0 | DLGAP4 | Disks large-associated protein 4 | -0.051949581 | 0.031456 |
| Q9Y2H1 | STK38L | Serine/threonine-protein kinase 38-like | 0.203548431 | 0.216075 |
| Q9Y2H6 | FNDC3A | Fibronectin type-III domain-containing protein 3A | -1.172600627 | -3.69503 |
| Q9Y2I1 | NISCH | Nischarin | 0.069894393 | -0.00995541 |
| Q9Y2J4 | AMOTL2 | Angiomotin-like protein 2 | -0.024172068 | 0.22597 |
| Q9Y2K6 | USP20 | Ubiquitin carboxyl-terminal hydrolase 20 | -1.630501548 | 2.52763 |
| Q9Y2L1 | DIS3 | Exosome complex exonuclease RRP44 | -0.225121498 | 0.00879447 |
| Q9Y2L9 | LRCH1 | Leucine-rich repeat and calponin homology domain-containing protein 1 | 1.516496619 | 0 |
| Q9Y2P8 | RCL1 | RNA 3'-terminal phosphate cyclase-like protein | -1.645132701 | -0.41655 |
| Q9Y2Q3 | GSTK1 | Glutathione S-transferase kappa 1 | -0.934821844 | 0.685785 |
| Q9Y2Q5 | LAMTOR2 | Ragulator complex protein LAMTOR2 | -1.990266482 | -2.28199 |
| Q9Y2Q9 | MRPS28 | 28S ribosomal protein S28, mitochondrial | -3.357818127 | 1.23786 |
| Q9Y2R0 | COA3 | Cytochrome c oxidase assembly factor 3 homolog, mitochondrial | 0 | -1.36646 |
| Q9Y2R4 | DDX52 | Probable ATP-dependent RNA helicase DDX52 | -1.315301498 | 1.22072 |

| | | | | |
|---|---|---|---|---|
| Q9Y2S6 | TMA7 | Translation machinery-associated protein 7 | -0.970335007 | -0.353163 |
| Q9Y2S7 | POLDIP2 | Polymerase delta-interacting protein 2 | -0.502125978 | -0.217985 |
| Q9Y2T2 | AP3M1 | AP-3 complex subunit mu-1 | 0.00530084 | -0.00938352 |
| Q9Y2U5 | MAP3K2 | Mitogen-activated protein kinase kinase kinase 2 | -0.394265572 | -0.508085 |
| Q9Y2U8 | LEMD3 | Inner nuclear membrane protein Man1 | -0.010173639 | 0.268475 |
| Q9Y2U9 | KLHDC2 | Kelch domain-containing protein 2 | -0.018332561 | -0.223438 |
| Q9Y2V2 | CARHSP1 | Calcium-regulated heat-stable protein 1 | -0.007895947 | -0.306512 |
| Q9Y2V7 | COG6 | Conserved oligomeric Golgi complex subunit 6 | 0.258124431 | 0.216064 |
| Q9Y2W2 | WBP11 | WW domain-binding protein 11 | -0.561786254 | 0.12182 |
| Q9Y2X3 | NOP58 | Nucleolar protein 58 | -0.38621823 | -0.0216767 |
| Q9Y2Z0 | SUGT1 | Protein SGT1 homolog | 0.051385244 | -0.0268784 |
| Q9Y303 | AMDHD2 | N-acetylglucosamine-6-phosphate deacetylase | 0.08961908 | -0.281537 |
| Q9Y305 | ACOT9 | Acyl-coenzyme A thioesterase 9, mitochondrial | -0.786264022 | 0.243665 |
| Q9Y314 | NOSIP | Nitric oxide synthase-interacting protein | -0.512313922 | 0.207527 |
| Q9Y315 | DERA | Deoxyribose-phosphate aldolase | -0.339848042 | -0.858532 |
| Q9Y316 | MEMO1 | Protein MEMO1 | 0.000564655 | -0.0477638 |
| Q9Y320 | TMX2 | Thioredoxin-related transmembrane protein 2 | -2.590431889 | 0.0212873 |
| Q9Y333 | LSM2 | U6 snRNA-associated Sm-like protein LSm2 | -0.579868952 | 0.016531 |
| Q9Y376 | CAB39 | Calcium-binding protein 39 | -0.022598743 | -0.21083 |
| Q9Y383 | LUC7L2 | Putative RNA-binding protein Luc7-like 2 | -0.237288555 | 0.0445652 |
| Q9Y394 | DHRS7 | Dehydrogenase/reductase SDR family member 7 | -1.286679586 | 0.00207074 |
| Q9Y399 | MRPS2 | 28S ribosomal protein S2, mitochondrial | -1.372179031 | 0.893998 |
| Q9Y3A3 | MOB4 | MOB-like protein phocein | -0.221024195 | -0.0734901 |
| Q9Y3A4 | RRP7A | Ribosomal RNA-processing protein 7 homolog A | 0.073160569 | 0.34386 |
| Q9Y3A5 | SBDS | Ribosome maturation protein SBDS | -0.422997872 | 0.0301339 |
| Q9Y3A6 | TMED5 | Transmembrane emp24 domain-containing protein 5 | 0.077062766 | 0.154335 |
| Q9Y3B4 | SF3B6 | Splicing factor 3B subunit 6 | -0.406685829 | -0.211237 |
| Q9Y3B7 | MRPL11 | 39S ribosomal protein L11, mitochondrial | -0.753121932 | 0.664708 |
| Q9Y3B8-3 | REXO2 | Isoform 3 of Oligoribonuclease, mitochondrial | 0.023798307 | 0.229229 |
| Q9Y3B9 | RRP15 | RRP15-like protein | -0.08470273 | -0.0661195 |
| Q9Y3C1 | NOP16 | Nucleolar protein 16 | -0.785528342 | 0.06427 |
| Q9Y3C4 | TPRKB | EKC/KEOPS complex subunit TPRKB | -0.827316443 | -0.0852032 |
| Q9Y3C6 | PPIL1 | Peptidyl-prolyl cis-trans isomerase-like 1 | -0.06662027 | 0.0734072 |
| Q9Y3C8 | UFC1 | Ubiquitin-fold modifier-conjugating enzyme 1 | -0.106444518 | 0.0821501 |
| Q9Y3D6 | FIS1 | Mitochondrial fission 1 protein | -0.536636114 | -0.347725 |
| Q9Y3D8 | AK6 | Adenylate kinase isoenzyme 6 | 0.375456095 | 0.799051 |
| Q9Y3E1 | HDGFL3 | Hepatoma-derived growth factor-related protein 3 | -0.054145257 | 0.118411 |
| Q9Y3E7 | CHMP3 | Charged multivesicular body protein 3 | -0.225365559 | -0.00103092 |
| Q9Y3F4 | STRAP | Serine-threonine kinase receptor-associated protein | -0.075579166 | -0.126441 |
| Q9Y3I0 | RTCB | RNA-splicing ligase RtcB homolog | -0.382764022 | -0.0627308 |
| Q9Y3I1 | FBXO7 | F-box only protein 7 | -0.279785872 | 0.00483259 |
| Q9Y3M8 | STARD13 | StAR-related lipid transfer protein 13 | -0.225346009 | 0 |
| Q9Y3P9 | RABGAP1 | Rab GTPase-activating protein 1 | -0.000940959 | 0.0566527 |
| Q9Y3Q8 | TSC22D4 | TSC22 domain family protein 4 | 0.305280368 | -0.0151974 |
| Q9Y3R4 | NEU2 | Sialidase-2 | -1.771921158 | 0 |
| Q9Y3R5 | DOP1B | Protein dopey-2 | -0.5684038 | -3.59742 |
| Q9Y3S2 | ZNF330 | Zinc finger protein 330 | -0.656848351 | -0.182553 |
| Q9Y3T9 | NOC2L | Nucleolar complex protein 2 homolog | -0.897499164 | 0.299791 |
| Q9Y3U8 | RPL36 | 60S ribosomal protein L36 | -1.431477467 | -0.407572 |
| Q9Y3Y2 | CHTOP | Chromatin target of PRMT1 protein | -0.982459625 | 1.02528 |
| Q9Y446 | PKP3 | Plakophilin-3 | 0.11941727 | -0.359527 |
| Q9Y450 | HBS1L | HBS1-like protein | -0.025789181 | -0.0709 |
| Q9Y478 | PRKAB1 | 5'-AMP-activated protein kinase subunit beta-1 | -0.848132213 | -0.730976 |
| Q9Y487 | ATP6V0A2 | V-type proton ATPase 116 kDa subunit a 2 | 0.451124191 | 0.134667 |
| Q9Y490 | TLN1 | Talin-1 | 0.098679543 | -0.071043 |
| Q9Y4B6 | DCAF1 | DDB1- and CUL4-associated factor 1 | -0.937416951 | -3.29545 |
| Q9Y4C2 | TCAF1 | TRPM8 channel-associated factor 1 | -0.076040188 | -0.0853586 |
| Q9Y4C8 | RBM19 | Probable RNA-binding protein 19 | -1.166599989 | 0.385189 |
| Q9Y4E8 | USP15 | Ubiquitin carboxyl-terminal hydrolase 15 | 0.28320384 | 0.0714353 |
| Q9Y4G6 | TLN2 | Talin-2 | 0.184703509 | -0.0301278 |
| Q9Y4H2 | IRS2 | Insulin receptor substrate 2 | -0.168032726 | -0.0891962 |
| Q9Y4K0 | LOXL2 | Lysyl oxidase homolog 2 | -0.284844081 | 0.228628 |
| Q9Y4K3 | TRAF6 | TNF receptor-associated factor 6 | -0.063803991 | -0.00974401 |
| Q9Y4P1 | ATG4B | Cysteine protease ATG4B | -0.110196431 | -0.0348597 |
| Q9Y4P8 | WIPI2 | WD repeat domain phosphoinositide-interacting protein 2 | -0.107269446 | -0.369095 |
| Q9Y4R8 | TELO2 | Telomere length regulation protein TEL2 homolog | -0.120934327 | -0.100776 |
| Q9Y4W2 | LAS1L | Ribosomal biogenesis protein LAS1L | -0.443877856 | 0.0984688 |
| Q9Y4W6 | AFG3L2 | AFG3-like protein 2 | -1.433205843 | 0.788449 |
| Q9Y4X5 | ARIH1 | E3 ubiquitin-protein ligase ARIH1 | 0.021999041 | 0.178391 |
| Q9Y4Z0 | LSM4 | U6 snRNA-associated Sm-like protein LSm4 | -0.454108238 | 0.0259899 |
| Q9Y512 | SAMM50 | Sorting and assembly machinery component 50 homolog | -1.355228265 | -0.0115795 |

| | | | | |
|---|---|---|---|---|
| Q9Y535 | POLR3H | DNA-directed RNA polymerase III subunit RPC8 | -0.385660609 | 0 |
| Q9Y546 | LRRC42 | Leucine-rich repeat-containing protein 42 | 0.868527412 | 1.3177 |
| Q9Y547 | HSPB11 | Intraflagellar transport protein 25 homolog | -0.049394449 | -2.93407 |
| Q9Y570 | PPME1 | Protein phosphatase methylesterase 1 | -0.010210037 | -0.078688 |
| Q9Y575 | ASB3 | Ankyrin repeat and SOCS box protein 3 | -0.187237819 | -0.627588 |
| Q9Y576 | ASB1 | Ankyrin repeat and SOCS box protein 1 | -0.784254551 | -2.70339 |
| Q9Y587 | AP4S1 | AP-4 complex subunit sigma-1 | 0.275985638 | -1.05648 |
| Q9Y5A9 | YTHDF2 | YTH domain-containing family protein 2 | -0.266182025 | 0.00965405 |
| Q9Y5B6 | PAXBP1 | PAX3- and PAX7-binding protein 1 | -1.643667698 | -2.07881 |
| Q9Y5B8 | NME7 | Nucleoside diphosphate kinase 7 | 0.053072214 | -0.00496006 |
| Q9Y5B9 | SUPT16H | FACT complex subunit SPT16 | 0.060209433 | -0.0577876 |
| Q9Y5J1 | UTP18 | U3 small nucleolar RNA-associated protein 18 homolog | -0.460664988 | 0.22598 |
| Q9Y5K5 | UCHL5 | Ubiquitin carboxyl-terminal hydrolase isozyme L5 | -0.026262363 | -0.100292 |
| Q9Y5K6 | CD2AP | CD2-associated protein | -0.053078016 | 0.260096 |
| Q9Y5K8 | ATP6V1D | V-type proton ATPase subunit D | -0.079245647 | -0.353582 |
| Q9Y5L0 | TNPO3 | Transportin-3 | -0.283908526 | -0.264167 |
| Q9Y5L2 | HILPDA | Hypoxia-inducible lipid droplet-associated protein | -0.885793845 | -0.88285 |
| Q9Y5L4 | TIMM13 | Mitochondrial import inner membrane translocase subunit Tim13 | -0.913350423 | 0.772264 |
| Q9Y5M8 | SRPRB | Signal recognition particle receptor subunit beta | -1.545567036 | 0.23997 |
| Q9Y5N5 | N6AMT1 | Methyltransferase N6AMT1 | 0.286658843 | 0.214391 |
| Q9Y5P4 | CERT1 | Ceramide transfer protein | -0.078657707 | -0.136583 |
| Q9Y5P6 | GMPPB | Mannose-1-phosphate guanyltransferase beta | 0.078039487 | -0.0999559 |
| Q9Y5Q8 | GTF3C5 | General transcription factor 3C polypeptide 5 | -0.025471369 | 0.287043 |
| Q9Y5R8 | TRAPPC1 | Trafficking protein particle complex subunit 1 | -0.17773064 | -0.179708 |
| Q9Y5S9 | RBM8A | RNA-binding protein 8A | -0.088626067 | -0.162465 |
| Q9Y5V3 | MAGED1 | Melanoma-associated antigen D1 | -0.479803244 | -0.767568 |
| Q9Y5W9 | SNX11 | Sorting nexin-11 | -0.222472668 | 0.52077 |
| Q9Y5X2 | SNX8 | Sorting nexin-8 | -0.645429373 | 0.229278 |
| Q9Y5X3 | SNX5 | Sorting nexin-5 | 0.142741283 | 0.137042 |
| Q9Y5Y2 | NUBP2 | Cytosolic Fe-S cluster assembly factor NUBP2 | -0.097498655 | -0.524758 |
| Q9Y5Z4 | HEBP2 | Heme-binding protein 2 | 0.197494507 | 0.264443 |
| Q9Y608-4 | LRRFIP2 | Isoform 4 of Leucine-rich repeat flightless-interacting protein 2 | 0.281919003 | -0.210281 |
| Q9Y613 | FHOD1 | FH1/FH2 domain-containing protein 1 | -0.586684465 | -0.202948 |
| Q9Y617 | PSAT1 | Phosphoserine aminotransferase | 0.024339835 | -0.134448 |
| Q9Y639 | NPTN | Neuroplastin | -0.84625109 | 0.276237 |
| Q9Y646 | CPQ | Carboxypeptidase Q | -0.0339516 | 0.244694 |
| Q9Y657 | SPIN1 | Spindlin-1 | -0.651618878 | -0.343716 |
| Q9Y678 | COPG1 | Coatomer subunit gamma-1 | -0.02028656 | -0.0306587 |
| Q9Y679 | AUP1 | Lipid droplet-regulating VLDL assembly factor AUP1 | -0.783657869 | 0.00249736 |
| Q9Y680 | FKBP7 | Peptidyl-prolyl cis-trans isomerase FKBP7 | -0.413943847 | 0.0473601 |
| Q9Y689 | ARL5A | ADP-ribosylation factor-like protein 5A | 0.012016058 | -1.1601 |
| Q9Y696 | CLIC4 | Chloride intracellular channel protein 4 | 0.005027294 | 0.0139942 |
| Q9Y6A4 | CFAP20 | Cilia- and flagella-associated protein 20 | 0.0504632 | -0.202061 |
| Q9Y6B6 | SAR1B | GTP-binding protein SAR1b | -0.020032883 | -0.0368029 |
| Q9Y6C9 | MTCH2 | Mitochondrial carrier homolog 2 | -1.074519396 | 0.403259 |
| Q9Y6D6 | ARFGEF1 | Brefeldin A-inhibited guanine nucleotide-exchange protein 1 | -0.043392022 | -0.0541226 |
| Q9Y6D9 | MAD1L1 | Mitotic spindle assembly checkpoint protein MAD1 | 0.175396522 | 0.157024 |
| Q9Y6G5 | COMMD10 | COMM domain-containing protein 10 | 0.088400602 | 0.446803 |
| Q9Y6G9 | DYNC1LI1 | Cytoplasmic dynein 1 light intermediate chain 1 | -0.165347656 | 0.180336 |
| Q9Y6H1 | CHCHD2 | Coiled-coil-helix-coiled-coil-helix domain-containing protein 2 | -1.866196632 | 0.274173 |
| Q9Y6I3 | EPN1 | Epsin-1 | 0.151305119 | 0.114281 |
| Q9Y6J9 | TAF6L | TAF6-like RNA polymerase II p300/CBP-associated factor-associated factor 65 kDa subunit 6L | -0.136017561 | 0.48851 |
| Q9Y6M5 | SLC30A1 | Zinc transporter 1 | -1.02902468 | -0.583794 |
| Q9Y6N5 | SQOR | Sulfide:quinone oxidoreductase, mitochondrial | -1.230683883 | 0.411612 |
| Q9Y6W3 | CAPN7 | Calpain-7 | -0.471737544 | -0.875626 |
| Q9Y6W5 | WASF2 | Wiskott-Aldrich syndrome protein family member 2 | -0.342676163 | 0.314821 |
| Q9Y6Y0 | IVNS1ABP | Influenza virus NS1A-binding protein | -0.209566673 | 0.305836 |
| Q9Y6Y8 | SEC23IP | SEC23-interacting protein | 0.027348121 | -0.225889 |
| R4GNH3 | PSMC3 | 26S proteasome regulatory subunit 6A | -0.281907638 | 0.484744 |
| S4R3H4 | ACIN1 | Apoptotic chromatin condensation inducer in the nucleus | 0.010220766 | -0.126923 |
| U3KQU7 | COG5 | Conserved oligomeric Golgi complex subunit 5 (Fragment) | 0.08840092 | -3.13422 |
