## Supplementary material for "Virally induced lipid droplets are a platform for innate immune signalling complexes": S Table 4

### LD fraction lipidomic data set

| Class | Metabolite name | Adduct type | Average Rt(min) | Average Mz | 24 hrs data set: lipid intensities normalized to LDs protein amount µg | | | | | | 24 hrs data set: Relative abundance of lipid intensities | | | | | | | | | | | | 8 hrs data set: lipid intensities normalized to LDs protein amount µg | | | | | | 8 hrs data set: Relative abundance of lipid intensities | | | | | | | | | | | |
|---|---|---|---|---|---|---|---|---|---|---|---|---|---|---|---|---|---|---|---|---|---|---|---|---|---|---|---|---|---|---|---|---|---|---|---|---|---|---|---|---|
| | | | | | Control at pmol/ug | Control at pmol/ug | Control at pmol/ug | dsRNA at pmol/ug | dsRNA at pmol/ug | dsRNA at pmol/ug | control % | control % | control % | dsRNA % | dsRNA % | dsRNA % | avg control % | RSD control % | avg dsRNA % | RSD dsRNA % | P value | LOG2FC | Control at pmol/ug | Control at pmol/ug | Control at pmol/ug | dsRNA at pmol/ug | dsRNA at pmol/ug | dsRNA at pmol/ug | control % | control % | control % | dsRNA % | dsRNA % | dsRNA % | avg control % | RSD control % | avg dsRNA % | RSD dsRNA % | P value | LOG2FC |
| PC | PC 27:1 | [M+H]+ | 12.166 | 662.47565 | 0.01957929 | 0.019780738 | 0.017832912 | 0.01134819 | 0.012222485 | 0.010937448 | 5.28129E-05 | 5.05563E-05 | 3.43937E-05 | 1.26512E-05 | 1.29686E-05 | 1.24035E-05 | 4.5921E-05 | 8.20292E-06 | 1.26744E-05 | 2.313E-07 | 0.029031881 | -1.857230433 | 0.001272423 | 0.001500927 | 0.001316453 | 0.001363622 | 0.001624104 | 0.002099481 | 5.95072E-06 | 5.43504E-06 | 1.05346E-05 | 3.80625E-06 | 4.50136E-06 | 6.52669E-06 | 7.3068E-06 | 2.29211E-06 | 4.94477E-06 | 1.15402E-06 | 0.285270957 | -0.563336896 |
| PC | PC 28:0 | [M+H]+ | 13.896 | 678.50732 | 0.004541418 | 0.005288154 | 0.005889699 | 0.007115454 | 0.006542921 | 0.005771966 | 1.225E-05 | 1.35156E-05 | 1.13592E-05 | 7.93245E-06 | 6.94235E-06 | 6.54564E-06 | 1.2375E-05 | 8.84771E-07 | 7.14015E-06 | 5.83179E-07 | 0.003715538 | -0.793397042 | 0.002122898 | 0.002515981 | 0.00129337 | 0.004165283 | 0.003686102 | 0.003226894 | 9.92812E-06 | 9.11067E-06 | 1.03499E-05 | 1.16265E-05 | 1.02164E-05 | 1.00315E-05 | 9.79623E-06 | 5.14439E-07 | 1.06248E-05 | 7.12309E-07 | 0.259665415 | 0.117133832 |
| PC | PC 30:0 | [M+H]+ | 14.837 | 706.53772 | 0.077374673 | 0.083745076 | 0.092184662 | 0.124123503 | 0.105722833 | 0.098081109 | 0.000208709 | 0.000214039 | 0.000177793 | 0.000138375 | 0.000112177 | 0.000111228 | 0.00020018 | 1.59789E-05 | 0.000120593 | 1.25797E-05 | 0.006088796 | -0.731149554 | 0.030663735 | 0.039290994 | 0.019652081 | 0.064202266 | 0.055772832 | 0.051049492 | 0.000143405 | 0.000142277 | 0.000157261 | 0.000179206 | 0.00015458 | 0.000158698 | 0.000147648 | 6.81342E-06 | 0.000164161 | 1.07704E-05 | 0.153839166 | 0.152955672 |
| PC | PC 30:1 | [M+H]+ | 14.06 | 704.52246 | 0.017081764 | 0.015769746 | 0.01986216 | 0.032818879 | 0.036385912 | 0.026587745 | 4.60761E-05 | 4.03049E-05 | 3.83074E-05 | 3.65871E-05 | 3.86072E-05 | 3.01516E-05 | 4.15628E-05 | 3.29394E-06 | 3.51153E-05 | 3.60547E-06 | 0.135864395 | -0.2431932 | 0.007773437 | 0.009664183 | 0.004787872 | 0.013510539 | 0.01486932 | 0.012837561 | 3.63539E-05 | 3.49952E-05 | 3.83139E-05 | 3.77117E-05 | 4.12118E-05 | 3.99083E-05 | 3.65543E-05 | 1.36225E-06 | 3.96106E-05 | 1.44433E-06 | 0.095296198 | 0.11584383 |
| PC | PC 31:0 | [M+H]+ | 15.295 | 720.55292 | 0.013809228 | 0.018237885 | 0.01748388 | 0.042085576 | 0.038891772 | 0.029955628 | 3.72488E-05 | 4.6613E-05 | 3.37205E-05 | 4.69178E-05 | 4.1266E-05 | 3.39709E-05 | 3.91941E-05 | 5.44011E-06 | 4.07183E-05 | 5.29973E-06 | 0.790654883 | 0.05503825 | 0.007970017 | 0.01097811 | 0.004879231 | 0.01786403 | 0.015677425 | 0.01405601 | 3.72732E-05 | 3.97531E-05 | 3.9045E-05 | 4.98635E-05 | 4.34515E-05 | 4.36961E-05 | 3.86904E-05 | 1.04297E-06 | 4.56704E-05 | 2.96666E-06 | 0.066493778 | 0.239281691 |
| PC | PC 31:1 | [M+H]+ | 15.764 | 718.53711 | 0.038252817 | 0.042615336 | 0.056329506 | 0.088565222 | 0.081193373 | 0.07354945 | 0.000103183 | 0.000108918 | 0.000108641 | 9.87342E-05 | 8.61501E-05 | 8.34081E-05 | 0.000106914 | 2.64065E-06 | 8.94308E-05 | 6.67309E-06 | 0.050741763 | -0.257603118 | 0.018076819 | 0.024565036 | 0.011870364 | 0.033930637 | 0.035150534 | 0.030329588 | 8.45395E-05 | 8.8953E-05 | 9.499E-05 | 9.47099E-05 | 9.74231E-05 | 9.4286E-05 | 8.94942E-05 | 4.28351E-06 | 9.5473E-05 | 1.38975E-06 | 0.178868179 | 0.09329906 |
| PC | PC 32:0\|PC 16:0_16:0 | [M+H]+ | 15.953 | 734.56848 | 0.162744904 | 0.199729082 | 0.232523617 | 0.257459825 | 0.199939555 | 0.188886127 | 0.000438986 | [illegible] | [illegible] | [illegible] | [illegible] | [illegible] | [illegible] | [illegible] | [illegible] | [illegible] | [illegible] | [illegible] | [illegible] | [illegible] | [illegible] | [illegible] | [illegible] | [illegible] | [illegible] | [illegible] | [illegible] | [illegible] | [illegible] | [illegible] | [illegible] | [illegible] | [illegible] | [illegible] | [illegible] | [illegible] | [illegible] |
| PC | PC 32:1\|PC 16:0_16:1 | [M+H]+ | 15.037 | 732.55237 | 0.225870972 | 0.256686916 | 0.312486838 | 0.608507848 | 0.621279002 | 0.521398757 | 0.000609262 | [illegible] | [illegible] | [illegible] | [illegible] | [illegible] | [illegible] | [illegible] | [illegible] | [illegible] | [illegible] | [illegible] | [illegible] | [illegible] | [illegible] | [illegible] | [illegible] | [illegible] | [illegible] | [illegible] | [illegible] | [illegible] | [illegible] | [illegible] | [illegible] | [illegible] | [illegible] | [illegible] | [illegible] | [illegible] | [illegible] |
| PC | PC 32:2 | [M+H]+ | 14.247 | 730.53687 | 0.022467244 | 0.023587314 | 0.030526574 | 0.062865584 | 0.0588808 | 0.051126221 | 6.06029E-05 | [illegible] | [illegible] | [illegible] | [illegible] | [illegible] | [illegible] | [illegible] | [illegible] | [illegible] | [illegible] | [illegible] | [illegible] | [illegible] | [illegible] | [illegible] | [illegible] | [illegible] | [illegible] | [illegible] | [illegible] | [illegible] | [illegible] | [illegible] | [illegible] | [illegible] | [illegible] | [illegible] | [illegible] | [illegible] | [illegible] |
| PC | PC 33:2 | [M+H]+ | 14.699 | 744.55328 | 0.014606994 | 0.014498606 | 0.021284031 | 0.03911337 | 0.040593308 | 0.032871963 | 3.94007E-05 | [illegible] | [illegible] | [illegible] | [illegible] | [illegible] | [illegible] | [illegible] | [illegible] | [illegible] | [illegible] | [illegible] | [illegible] | [illegible] | [illegible] | [illegible] | [illegible] | [illegible] | [illegible] | [illegible] | [illegible] | [illegible] | [illegible] | [illegible] | [illegible] | [illegible] | [illegible] | [illegible] | [illegible] | [illegible] | [illegible] |
| PC | PC 34:1\|PC 16:0_18:1 | [M+H]+ | 16.072 | 760.58337 | 1.032062897 | 1.135699011 | 1.321702471 | 2.662035471 | 2.505406676 | 2.210709037 | 0.002783873 | [illegible] | [illegible] | [illegible] | [illegible] | [illegible] | [illegible] | [illegible] | [illegible] | [illegible] | [illegible] | [illegible] | [illegible] | [illegible] | [illegible] | [illegible] | [illegible] | [illegible] | [illegible] | [illegible] | [illegible] | [illegible] | [illegible] | [illegible] | [illegible] | [illegible] | [illegible] | [illegible] | [illegible] | [illegible] | [illegible] |
| PC | PC 34:2\|PC 16:1_18:1 | [M+H]+ | 15.191 | 758.56903 | 0.235281479 | 0.254503392 | 0.336353677 | 0.673397536 | 0.665536647 | 0.577319551 | 0.000634645 | [illegible] | [illegible] | [illegible] | [illegible] | [illegible] | [illegible] | [illegible] | [illegible] | [illegible] | [illegible] | [illegible] | [illegible] | [illegible] | [illegible] | [illegible] | [illegible] | [illegible] | [illegible] | [illegible] | [illegible] | [illegible] | [illegible] | [illegible] | [illegible] | [illegible] | [illegible] | [illegible] | [illegible] | [illegible] | [illegible] |
| PC | PC 35:1 | [M+H]+ | 16.679 | 774.59955 | 0.084716358 | [illegible] | [illegible] | [illegible] | [illegible] | [illegible] | [illegible] | [illegible] | [illegible] | [illegible] | [illegible] | [illegible] | [illegible] | [illegible] | [illegible] | [illegible] | [illegible] | [illegible] | [illegible] | [illegible] | [illegible] | [illegible] | [illegible] | [illegible] | [illegible] | [illegible] | [illegible] | [illegible] | [illegible] | [illegible] | [illegible] | [illegible] | [illegible] | [illegible] | [illegible] | [illegible] | [illegible] |
| PC | PC 35:2 | [M+H]+ | 15.572 | 772.58685 | 0.035321383 | [illegible] | [illegible] | [illegible] | [illegible] | [illegible] | [illegible] | [illegible] | [illegible] | [illegible] | [illegible] | [illegible] | [illegible] | [illegible] | [illegible] | [illegible] | [illegible] | [illegible] | [illegible] | [illegible] | [illegible] | [illegible] | [illegible] | [illegible] | [illegible] | [illegible] | [illegible] | [illegible] | [illegible] | [illegible] | [illegible] | [illegible] | [illegible] | [illegible] | [illegible] | [illegible] | [illegible] |
| PC | PC 35:3 | [M+H]+ | 14.856 | 770.57025 | 0.005038923 | [illegible] | [illegible] | [illegible] | [illegible] | [illegible] | [illegible] | [illegible] | [illegible] | [illegible] | [illegible] | [illegible] | [illegible] | [illegible] | [illegible] | [illegible] | [illegible] | [illegible] | [illegible] | [illegible] | [illegible] | [illegible] | [illegible] | [illegible] | [illegible] | [illegible] | [illegible] | [illegible] | [illegible] | [illegible] | [illegible] | [illegible] | [illegible] | [illegible] | [illegible] | [illegible] | [illegible] |
| PC | PC 36:1\|PC 18:0_18:1 | [M+H]+ | 17.317 | 788.61621 | 0.28835142 | [illegible] | [illegible] | [illegible] | [illegible] | [illegible] | [illegible] | [illegible] | [illegible] | [illegible] | [illegible] | [illegible] | [illegible] | [illegible] | [illegible] | [illegible] | [illegible] | [illegible] | [illegible] | [illegible] | [illegible] | [illegible] | [illegible] | [illegible] | [illegible] | [illegible] | [illegible] | [illegible] | [illegible] | [illegible] | [illegible] | [illegible] | [illegible] | [illegible] | [illegible] | [illegible] | [illegible] |
| PC | PC 36:2\|PC 18:1_18:1 | [M+H]+ | 16.182 | 786.59943 | 0.433039042 | [illegible] | [illegible] | [illegible] | [illegible] | [illegible] | [illegible] | [illegible] | [illegible] | [illegible] | [illegible] | [illegible] | [illegible] | [illegible] | [illegible] | [illegible] | [illegible] | [illegible] | [illegible] | [illegible] | [illegible] | [illegible] | [illegible] | [illegible] | [illegible] | [illegible] | [illegible] | [illegible] | [illegible] | [illegible] | [illegible] | [illegible] | [illegible] | [illegible] | [illegible] | [illegible] | [illegible] |
| PC | PC 36:3 | [M+H]+ | 15.286 | 784.58453 | 0.11684463 | [illegible] | [illegible] | [illegible] | [illegible] | [illegible] | [illegible] | [illegible] | [illegible] | [illegible] | [illegible] | [illegible] | [illegible] | [illegible] | [illegible] | [illegible] | [illegible] | [illegible] | [illegible] | [illegible] | [illegible] | [illegible] | [illegible] | [illegible] | [illegible] | [illegible] | [illegible] | [illegible] | [illegible] | [illegible] | [illegible] | [illegible] | [illegible] | [illegible] | [illegible] | [illegible] | [illegible] |
| PC | PC 36:4\|PC 16:0_20:4 | [M+H]+ | 15.052 | 782.5672 | 0.12148424 | [illegible] | [illegible] | [illegible] | [illegible] | [illegible] | [illegible] | [illegible] | [illegible] | [illegible] | [illegible] | [illegible] | [illegible] | [illegible] | [illegible] | [illegible] | [illegible] | [illegible] | [illegible] | [illegible] | [illegible] | [illegible] | [illegible] | [illegible] | [illegible] | [illegible] | [illegible] | [illegible] | [illegible] | [illegible] | [illegible] | [illegible] | [illegible] | [illegible] | [illegible] | [illegible] | [illegible] |
| PC | PC 36:5 | [M+H]+ | 14.361 | 780.55255 | 0.027121923 | [illegible] | [illegible] | [illegible] | [illegible] | [illegible] | [illegible] | [illegible] | [illegible] | [illegible] | [illegible] | [illegible] | [illegible] | [illegible] | [illegible] | [illegible] | [illegible] | [illegible] | [illegible] | [illegible] | [illegible] | [illegible] | [illegible] | [illegible] | [illegible] | [illegible] | [illegible] | [illegible] | [illegible] | [illegible] | [illegible] | [illegible] | [illegible] | [illegible] | [illegible] | [illegible] | [illegible] |
| PC | PC 38:2 | [M+H]+ | 17.387 | 814.63129 | 0.019805471 | [illegible] | [illegible] | [illegible] | [illegible] | [illegible] | [illegible] | [illegible] | [illegible] | [illegible] | [illegible] | [illegible] | [illegible] | [illegible] | [illegible] | [illegible] | [illegible] | [illegible] | [illegible] | [illegible] | [illegible] | [illegible] | [illegible] | [illegible] | [illegible] | [illegible] | [illegible] | [illegible] | [illegible] | [illegible] | [illegible] | [illegible] | [illegible] | [illegible] | [illegible] | [illegible] | [illegible] |
| PC | PC 38:3 | [M+H]+ | 16.702 | 812.61389 | 0.062276287 | [illegible] | [illegible] | [illegible] | [illegible] | [illegible] | [illegible] | [illegible] | [illegible] | [illegible] | [illegible] | [illegible] | [illegible] | [illegible] | [illegible] | [illegible] | [illegible] | [illegible] | [illegible] | [illegible] | [illegible] | [illegible] | [illegible] | [illegible] | [illegible] | [illegible] | [illegible] | [illegible] | [illegible] | [illegible] | [illegible] | [illegible] | [illegible] | [illegible] | [illegible] | [illegible] | [illegible] |
| PC | PC 38:4\|PC 18:0_20:4 | [M+H]+ | 16.18 | 810.59949 | 0.106547839 | [illegible] | [illegible] | [illegible] | [illegible] | [illegible] | [illegible] | [illegible] | [illegible] | [illegible] | [illegible] | [illegible] | [illegible] | [illegible] | [illegible] | [illegible] | [illegible] | [illegible] | [illegible] | [illegible] | [illegible] | [illegible] | [illegible] | [illegible] | [illegible] | [illegible] | [illegible] | [illegible] | [illegible] | [illegible] | [illegible] | [illegible] | [illegible] | [illegible] | [illegible] | [illegible] | [illegible] |
| PC | PC 38:5 | [M+H]+ | 15.139 | 808.58331 | 0.105024203 | [illegible] | [illegible] | [illegible] | [illegible] | [illegible] | [illegible] | [illegible] | [illegible] | [illegible] | [illegible] | [illegible] | [illegible] | [illegible] | [illegible] | [illegible] | [illegible] | [illegible] | [illegible] | [illegible] | [illegible] | [illegible] | [illegible] | [illegible] | [illegible] | [illegible] | [illegible] | [illegible] | [illegible] | [illegible] | [illegible] | [illegible] | [illegible] | [illegible] | [illegible] | [illegible] | [illegible] |
| PC | PC 38:6 | [M+H]+ | 14.78 | 806.57001 | 0.053549121 | [illegible] | [illegible] | [illegible] | [illegible] | [illegible] | [illegible] | [illegible] | [illegible] | [illegible] | [illegible] | [illegible] | [illegible] | [illegible] | [illegible] | [illegible] | [illegible] | [illegible] | [illegible] | [illegible] | [illegible] | [illegible] | [illegible] | [illegible] | [illegible] | [illegible] | [illegible] | [illegible] | [illegible] | [illegible] | [illegible] | [illegible] | [illegible] | [illegible] | [illegible] | [illegible] | [illegible] |
| PC | PC 38:7 | [M+H]+ | 14.014 | 804.55298 | 0.007279602 | [illegible] | [illegible] | [illegible] | [illegible] | [illegible] | [illegible] | [illegible] | [illegible] | [illegible] | [illegible] | [illegible] | [illegible] | [illegible] | [illegible] | [illegible] | [illegible] | [illegible] | [illegible] | [illegible] | [illegible] | [illegible] | [illegible] | [illegible] | [illegible] | [illegible] | [illegible] | [illegible] | [illegible] | [illegible] | [illegible] | [illegible] | [illegible] | [illegible] | [illegible] | [illegible] | [illegible] |
| PC | PC 40:4 | [M+H]+ | 16.934 | 838.63086 | 0.018832366 | [illegible] | [illegible] | [illegible] | [illegible] | [illegible] | [illegible] | [illegible] | [illegible] | [illegible] | [illegible] | [illegible] | [illegible] | [illegible] | [illegible] | [illegible] | [illegible] | [illegible] | [illegible] | [illegible] | [illegible] | [illegible] | [illegible] | [illegible] | [illegible] | [illegible] | [illegible] | [illegible] | [illegible] | [illegible] | [illegible] | [illegible] | [illegible] | [illegible] | [illegible] | [illegible] | [illegible] |
| PC | PC 40:5 | [M+H]+ | 16.213 | 836.61639 | 0.038368492 | [illegible] | [illegible] | [illegible] | [illegible] | [illegible] | [illegible] | [illegible] | [illegible] | [illegible] | [illegible] | [illegible] | [illegible] | [illegible] | [illegible] | [illegible] | [illegible] | [illegible] | [illegible] | [illegible] | [illegible] | [illegible] | [illegible] | [illegible] | [illegible] | [illegible] | [illegible] | [illegible] | [illegible] | [illegible] | [illegible] | [illegible] | [illegible] | [illegible] | [illegible] | [illegible] | [illegible] |
| PC | PC 40:6 | [M+H]+ | 15.9 | 834.6004 | 0.023614032 | [illegible] | [illegible] | [illegible] | [illegible] | [illegible] | [illegible] | [illegible] | [illegible] | [illegible] | [illegible] | [illegible] | [illegible] | [illegible] | [illegible] | [illegible] | [illegible] | [illegible] | [illegible] | [illegible] | [illegible] | [illegible] | [illegible] | [illegible] | [illegible] | [illegible] | [illegible] | [illegible] | [illegible] | [illegible] | [illegible] | [illegible] | [illegible] | [illegible] | [illegible] | [illegible] | [illegible] |
| PC | PC 40:7 | [M+H]+ | 14.873 | 832.58533 | 0.02170464 | [illegible] | [illegible] | [illegible] | [illegible] | [illegible] | [illegible] | [illegible] | [illegible] | [illegible] | [illegible] | [illegible] | [illegible] | [illegible] | [illegible] | [illegible] | [illegible] | [illegible] | [illegible] | [illegible] | [illegible] | [illegible] | [illegible] | [illegible] | [illegible] | [illegible] | [illegible] | [illegible] | [illegible] | [illegible] | [illegible] | [illegible] | [illegible] | [illegible] | [illegible] | [illegible] | [illegible] |
| PC | PC 42:6 | [M+H]+ | 16.242 | 862.63068 | 0.005975723 | [illegible] | [illegible] | [illegible] | [illegible] | [illegible] | [illegible] | [illegible] | [illegible] | [illegible] | [illegible] | [illegible] | [illegible] | [illegible] | [illegible] | [illegible] | [illegible] | [illegible] | [illegible] | [illegible] | [illegible] | [illegible] | [illegible] | [illegible] | [illegible] | [illegible] | [illegible] | [illegible] | [illegible] | [illegible] | [illegible] | [illegible] | [illegible] | [illegible] | [illegible] | [illegible] | [illegible] |
| PC | PC O-32:1 | [M+H]+ | 17.419 | 718.57037 | 0.023591635 | [illegible] | [illegible] | [illegible] | [illegible] | [illegible] | [illegible] | [illegible] | [illegible] | [illegible] | [illegible] | [illegible] | [illegible] | [illegible] | [illegible] | [illegible] | [illegible] | [illegible] | [illegible] | [illegible] | [illegible] | [illegible] | [illegible] | [illegible] | [illegible] | [illegible] | [illegible] | [illegible] | [illegible] | [illegible] | [illegible] | [illegible] | [illegible] | [illegible] | [illegible] | [illegible] | [illegible] |
| PC | PC O-34:0 | [M+H]+ | 18.281 | 748.62115 | 0.00341121 | [illegible] | [illegible] | [illegible] | [illegible] | [illegible] | [illegible] | [illegible] | [illegible] | [illegible] | [illegible] | [illegible] | [illegible] | [illegible] | [illegible] | [illegible] | [illegible] | [illegible] | [illegible] | [illegible] | [illegible] | [illegible] | [illegible] | [illegible] | [illegible] | [illegible] | [illegible] | [illegible] | [illegible] | [illegible] | [illegible] | [illegible] | [illegible] | [illegible] | [illegible] | [illegible] | [illegible] |
| PC | PC O-37:8 | [M+H]+ | 16.5 | 774.54083 | 0.006145545 | [illegible] | [illegible] | [illegible] | [illegible] | [illegible] | [illegible] | [illegible] | [illegible] | [illegible] | [illegible] | [illegible] | [illegible] | [illegible] | [illegible] | [illegible] | [illegible] | [illegible] | [illegible] | [illegible] | [illegible] | [illegible] | [illegible] | [illegible] | [illegible] | [illegible] | [illegible] | [illegible] | [illegible] | [illegible] | [illegible] | [illegible] | [illegible] | [illegible] | [illegible] | [illegible] | [illegible] |
| PC | PC O-38:4 | [M+H]+ | 17.114 | 796.62134 | 0.007561967 | [illegible] | [illegible] | [illegible] | [illegible] | [illegible] | [illegible] | [illegible] | [illegible] | [illegible] | [illegible] | [illegible] | [illegible] | [illegible] | [illegible] | [illegible] | [illegible] | [illegible] | [illegible] | [illegible] | [illegible] | [illegible] | [illegible] | [illegible] | [illegible] | [illegible] | [illegible] | [illegible] | [illegible] | [illegible] | [illegible] | [illegible] | [illegible] | [illegible] | [illegible] | [illegible] | [illegible] |
| PC | PC O-54:2 | [M+H]+ | 21.27 | 1024.89954 | 0.013845235 | [illegible] | [illegible] | [illegible] | [illegible] | [illegible] | [illegible] | [illegible] | [illegible] | [illegible] | [illegible] | [illegible] | [illegible] | [illegible] | [illegible] | [illegible] | [illegible] | [illegible] | [illegible] | [illegible] | [illegible] | [illegible] | [illegible] | [illegible] | [illegible] | [illegible] | [illegible] | [illegible] | [illegible] | [illegible] | [illegible] | [illegible] | [illegible] | [illegible] | [illegible] | [illegible] | [illegible] |
| PE | PE 32:0\|PE 16:0_16:0 | [M-H]- | 14.835 | 690.50787 | 0.0127483 | [illegible] | [illegible] | [illegible] | [illegible] | [illegible] | [illegible] | [illegible] | [illegible] | [illegible] | [illegible] | [illegible] | [illegible] | [illegible] | [illegible] | [illegible] | [illegible] | [illegible] | [illegible] | [illegible] | [illegible] | [illegible] | [illegible] | [illegible] | [illegible] | [illegible] | [illegible] | [illegible] | [illegible] | [illegible] | [illegible] | [illegible] | [illegible] | [illegible] | [illegible] | [illegible] |
| PE | PE 32:1\|PE 14:0_18:1 | [M-H]- | 14.761 | 688.49091 | 0.009935186 | [illegible] | [illegible] | [illegible] | [illegible] | [illegible] | [illegible] | [illegible] | [illegible] | [illegible] | [illegible] | [illegible] | [illegible] | [illegible] | [illegible] | [illegible] | [illegible] | [illegible] | [illegible] | [illegible] | [illegible] | [illegible] | [illegible] | [illegible] | [illegible] | [illegible] | [illegible] | [illegible] | [illegible] | [illegible] | [illegible] | [illegible] | [illegible] | [illegible] | [illegible] | [illegible] |
| PE | PE 33:1\|PE 16:0_17:1 | [M-H]- | 15.186 | 702.50726 | 0.002094292 | [illegible] | [illegible] | [illegible] | [illegible] | [illegible] | [illegible] | [illegible] | [illegible] | [illegible] | [illegible] | [illegible] | [illegible] | [illegible] | [illegible] | [illegible] | [illegible] | [illegible] | [illegible] | [illegible] | [illegible] | [illegible] | [illegible] | [illegible] | [illegible] | [illegible] | [illegible] | [illegible] | [illegible] | [illegible] | [illegible] | [illegible] | [illegible] | [illegible] | [illegible] | [illegible] |
| PE | PE 34:0\|PE 16:0_18:0 | [M-H]- | 15.941 | 718.53711 | 0.035245889 | [illegible] | [illegible] | [illegible] | [illegible] | [illegible] | [illegible] | [illegible] | [illegible] | [illegible] | [illegible] | [illegible] | [illegible] | [illegible] | [illegible] | [illegible] | [illegible] | [illegible] | [illegible] | [illegible] | [illegible] | [illegible] | [illegible] | [illegible] | [illegible] | [illegible] | [illegible] | [illegible] | [illegible] | [illegible] | [illegible] | [illegible] | [illegible] | [illegible] | [illegible] | [illegible] |
| PE | PE 34:1\|PE 16:0_18:1 | [M-H]- | 15.004 | 716.52246 | 0.039671873 | [illegible] | [illegible] | [illegible] | [illegible] | [illegible] | [illegible] | [illegible] | [illegible] | [illegible] | [illegible] | [illegible] | [illegible] | [illegible] | [illegible] | [illegible] | [illegible] | [illegible] | [illegible] | [illegible] | [illegible] | [illegible] | [illegible] | [illegible] | [illegible] | [illegible] | [illegible] | [illegible] | [illegible] | [illegible] | [illegible] | [illegible] | [illegible] | [illegible] | [illegible] | [illegible] |
| PE | PE 34:1\|PE 18:0_16:1 | [M-H]- | 15.735 | 716.52246 | 0.080782809 | [illegible] | [illegible] | [illegible] | [illegible] | [illegible] | [illegible] | [illegible] | [illegible] | [illegible] | [illegible] | [illegible] | [illegible] | [illegible] | [illegible] | [illegible] | [illegible] | [illegible] | [illegible] | [illegible] | [illegible] | [illegible] | [illegible] | [illegible] | [illegible] | [illegible] | [illegible] | [illegible] | [illegible] | [illegible] | [illegible] | [illegible] | [illegible] | [illegible] | [illegible] | [illegible] |
| PE | PE 34:2\|PE 16:1_18:1 | [M-H]- | 14.859 | 714.50854 | 0.023311286 | [illegible] | [illegible] | [illegible] | [illegible] | [illegible] | [illegible] | [illegible] | [illegible] | [illegible] | [illegible] | [illegible] | [illegible] | [illegible] | [illegible] | [illegible] | [illegible] | [illegible] | [illegible] | [illegible] | [illegible] | [illegible] | [illegible] | [illegible] | [illegible] | [illegible] | [illegible] | [illegible] | [illegible] | [illegible] | [illegible] | [illegible] | [illegible] | [illegible] | [illegible] | [illegible] |
| PE | PE 35:1\|PE 17:0_18:1 | [M-H]- | 16.264 | 730.5376 | 0.021277837 | [illegible] | [illegible] | [illegible] | [illegible] | [illegible] | [illegible] | [illegible] | [illegible] | [illegible] | [illegible] | [illegible] | [illegible] | [illegible] | [illegible] | [illegible] | [illegible] | [illegible] | [illegible] | [illegible] | [illegible] | [illegible] | [illegible] | [illegible] | [illegible] | [illegible] | [illegible] | [illegible] | [illegible] | [illegible] | [illegible] | [illegible] | [illegible] | [illegible] | [illegible] | [illegible] |
| PE | PE 35:2\|PE 17:1_18:1 | [M-H]- | 15.338 | 728.52332 | 0.003662257 | [illegible] | [illegible] | [illegible] | [illegible] | [illegible] | [illegible] | [illegible] | [illegible] | [illegible] | [illegible] | [illegible] | [illegible] | [illegible] | [illegible] | [illegible] | [illegible] | [illegible] | [illegible] | [illegible] | [illegible] | [illegible] | [illegible] | [illegible] | [illegible] | [illegible] | [illegible] | [illegible] | [illegible] | [illegible] | [illegible] | [illegible] | [illegible] | [illegible] | [illegible] | [illegible] |
| PE | PE 36:1\|PE 18:0_18:1 | [M-H]- | 16.79 | 744.55359 | 0.253177964 | [illegible] | [illegible] | [illegible] | [illegible] | [illegible] | [illegible] | [illegible] | [illegible] | [illegible] | [illegible] | [illegible] | [illegible] | [illegible] | [illegible] | [illegible] | [illegible] | [illegible] | [illegible] | [illegible] | [illegible] | [illegible] | [illegible] | [illegible] | [illegible] | [illegible] | [illegible] | [illegible] | [illegible] | [illegible] | [illegible] | [illegible] | [illegible] | [illegible] | [illegible] | [illegible] |
| PE | PE 36:2\|PE 18:0_18:2 | [M-H]- | 15.851 | 742.53711 | 0.126563682 | [illegible] | [illegible] | [illegible] | [illegible] | [illegible] | [illegible] | [illegible] | [illegible] | [illegible] | [illegible] | [illegible] | [illegible] | [illegible] | [illegible] | [illegible] | [illegible] | [illegible] | [illegible] | [illegible] | [illegible] | [illegible] | [illegible] | [illegible] | [illegible] | [illegible] | [illegible] | [illegible] | [illegible] | [illegible] | [illegible] | [illegible] | [illegible] | [illegible] | [illegible] | [illegible] |
| PE | PE 36:3\|PE 18:1_18:2 | [M-H]- | 15.102 | 740.5213 | 0.016941144 | [illegible] | [illegible] | [illegible] | [illegible] | [illegible] | [illegible] | [illegible] | [illegible] | [illegible] | [illegible] | [illegible] | [illegible] | [illegible] | [illegible] | [illegible] | [illegible] | [illegible] | [illegible] | [illegible] | [illegible] | [illegible] | [illegible] | [illegible] | [illegible] | [illegible] | [illegible] | [illegible] | [illegible] | [illegible] | [illegible] | [illegible] | [illegible] | [illegible] | [illegible] | [illegible] |
| PE | PE 36:4\|PE 16:0_20:4 | [M-H]- | 14.864 | 738.50732 | 0.021507098 | [illegible] | [illegible] | [illegible] | [illegible] | [illegible] | [illegible] | [illegible] | [illegible] | [illegible] | [illegible] | [illegible] | [illegible] | [illegible] | [illegible] | [illegible] | [illegible] | [illegible] | [illegible] | [illegible] | [illegible] | [illegible] | [illegible] | [illegible] | [illegible] | [illegible] | [illegible] | [illegible] | [illegible] | [illegible] | [illegible] | [illegible] | [illegible] | [illegible] | [illegible] | [illegible] |
| PE | PE 37:1\|PE 18:0_19:1 | [M-H]- | 16.676 | 758.57013 | 0.012563207 | [illegible] | [illegible] | [illegible] | [illegible] | [illegible] | [illegible] | [illegible] | [illegible] | [illegible] | [illegible] | [illegible] | [illegible] | [illegible] | [illegible] | [illegible] | [illegible] | [illegible] | [illegible] | [illegible] | [illegible] | [illegible] | [illegible] | [illegible] | [illegible] | [illegible] | [illegible] | [illegible] | [illegible] | [illegible] | [illegible] | [illegible] | [illegible] | [illegible] | [illegible] | [illegible] |
| PE | PE 37:2\|PE 18:1_19:1 | [M-H]- | 16.316 | 756.55469 | 0.005391546 | [illegible] | [illegible] | [illegible] | [illegible] | [illegible] | [illegible] | [illegible] | [illegible] | [illegible] | [illegible] | [illegible] | [illegible] | [illegible] | [illegible] | [illegible] | [illegible] | [illegible] | [illegible] | [illegible] | [illegible] | [illegible] | [illegible] | [illegible] | [illegible] | [illegible] | [illegible] | [illegible] | [illegible] | [illegible] | [illegible] | [illegible] | [illegible] | [illegible] | [illegible] | [illegible] |
| PE | PE 38:1\|PE 18:0_20:1 | [M-H]- | 17.31 | 772.58466 | 0.048643531 | [illegible] | [illegible] | [illegible] | [illegible] | [illegible] | [illegible] | [illegible] | [illegible] | [illegible] | [illegible] | [illegible] | [illegible] | [illegible] | [illegible] | [illegible] | [illegible] | [illegible] | [illegible] | [illegible] | [illegible] | [illegible] | [illegible] | [illegible] | [illegible] | [illegible] | [illegible] | [illegible] | [illegible] | [illegible] | [illegible] | [illegible] | [illegible] | [illegible] | [illegible] | [illegible] |
| PE | PE 38:2\|PE 18:0_20:2 | [M-H]- | 16.176 | 770.56824 | 0.038649142 | [illegible] | [illegible] | [illegible] | [illegible] | [illegible] | [illegible] | [illegible] | [illegible] | [illegible] | [illegible] | [illegible] | [illegible] | [illegible] | [illegible] | [illegible] | [illegible] | [illegible] | [illegible] | [illegible] | [illegible] | [illegible] | [illegible] | [illegible] | [illegible] | [illegible] | [illegible] | [illegible] | [illegible] | [illegible] | [illegible] | [illegible] | [illegible] | [illegible] | [illegible] | [illegible] |
| PE | PE 38:3\|PE 18:0_20:3 | [M-H]- | 16.302 | 768.55402 | 0.04732219 | [illegible] | [illegible] | [illegible] | [illegible] | [illegible] | [illegible] | [illegible] | [illegible] | [illegible] | [illegible] | [illegible] | [illegible] | [illegible] | [illegible] | [illegible] | [illegible] | [illegible] | [illegible] | [illegible] | [illegible] | [illegible] | [illegible] | [illegible] | [illegible] | [illegible] | [illegible] | [illegible] | [illegible] | [illegible] | [illegible] | [illegible] | [illegible] | [illegible] | [illegible] | [illegible] |
| PE | PE 38:4\|PE 18:0_20:4 | [M-H]- | 15.888 | 766.53632 | 0.201079925 | [illegible] | [illegible] | [illegible] | [illegible] | [illegible] | [illegible] | [illegible] | [illegible] | [illegible] | [illegible] | [illegible] | [illegible] | [illegible] | [illegible] | [illegible] | [illegible] | [illegible] | [illegible] | [illegible] | [illegible] | [illegible] | [illegible] | [illegible] | [illegible] | [illegible] | [illegible] | [illegible] | [illegible] | [illegible] | [illegible] | [illegible] | [illegible] | [illegible] | [illegible] | [illegible] |
| PE | PE 38:5\|PE 18:1_20:4 | [M-H]- | 14.972 | 764.52081 | 0.056779434 | [illegible] | [illegible] | [illegible] | [illegible] | [illegible] | [illegible] | [illegible] | [illegible] | [illegible] | [illegible] | [illegible] | [illegible] | [illegible] | [illegible] | [illegible] | [illegible] | [illegible] | [illegible] | [illegible] | [illegible] | [illegible] | [illegible] | [illegible] | [illegible] | [illegible] | [illegible] | [illegible] | [illegible] | [illegible] | [illegible] | [illegible] | [illegible] | [illegible] | [illegible] | [illegible] |
| PE | PE 38:6\|PE 16:0_22:6 | [M-H]- | 14.653 | 762.5083 | 0.02356449 | [illegible] | [illegible] | [illegible] | [illegible] | [illegible] | [illegible] | [illegible] | [illegible] | [illegible] | [illegible] | [illegible] | [illegible] | [illegible] | [illegible] | [illegible] | [illegible] | [illegible] | [illegible] | [illegible] | [illegible] | [illegible] | [illegible] | [illegible] | [illegible] | [illegible] | [illegible] | [illegible] | [illegible] | [illegible] | [illegible] | [illegible] | [illegible] | [illegible] | [illegible] | [illegible] |
| PE | PE 39:3\|PE 18:0_21:3 | [M-H]- | 16.896 | 782.56885 | 0.001671891 | [illegible] | [illegible] | [illegible] | [illegible] | [illegible] | [illegible] | [illegible] | [illegible] | [illegible] | [illegible] | [illegible] | [illegible] | [illegible] | [illegible] | [illegible] | [illegible] | [illegible] | [illegible] | [illegible] | [illegible] | [illegible] | [illegible] | [illegible] | [illegible] | [illegible] | [illegible] | [illegible] | [illegible] | [illegible] | [illegible] | [illegible] | [illegible] | [illegible] | [illegible] | [illegible] |
| PE | PE 40:4\|PE 18:0_22:4 | [M-H]- | 16.622 | 794.56976 | 0.057189728 | [illegible] | [illegible] | [illegible] | [illegible] | [illegible] | [illegible] | [illegible] | [illegible] | [illegible] | [illegible] | [illegible] | [illegible] | [illegible] | [illegible] | [illegible] | [illegible] | [illegible] | [illegible] | [illegible] | [illegible] | [illegible] | [illegible] | [illegible] | [illegible] | [illegible] | [illegible] | [illegible] | [illegible] | [illegible] | [illegible] | [illegible] | [illegible] | [illegible] | [illegible] | [illegible] |
| PE | PE 40:5\|PE 18:0_22:5 | [M-H]- | 15.953 | 792.55316 | 0.079480732 | [illegible] | [illegible] | [illegible] | [illegible] | [illegible] | [illegible] | [illegible] | [illegible] | [illegible] | [illegible] | [illegible] | [illegible] | [illegible] | [illegible] | [illegible] | [illegible] | [illegible] | [illegible] | [illegible] | [illegible] | [illegible] | [illegible] | [illegible] | [illegible] | [illegible] | [illegible] | [illegible] | [illegible] | [illegible] | [illegible] | [illegible] | [illegible] | [illegible] | [illegible] | [illegible] |
| PE | PE 40:6\|PE 18:0_22:6 | [M-H]- | 15.657 | 790.53894 | 0.072554962 | [illegible] | [illegible] | [illegible] | [illegible] | [illegible] | [illegible] | [illegible] | [illegible] | [illegible] | [illegible] | [illegible] | [illegible] | [illegible] | [illegible] | [illegible] | [illegible] | [illegible] | [illegible] | [illegible] | [illegible] | [illegible] | [illegible] | [illegible] | [illegible] | [illegible] | [illegible] | [illegible] | [illegible] | [illegible] | [illegible] | [illegible] | [illegible] | [illegible] | [illegible] | [illegible] |
| PE | PE 42:1\|PE 24:0_18:1 | [M-H]- | 20.094 | 828.64874 | 0.005281352 | [illegible] | [illegible] | [illegible] | [illegible] | [illegible] | [illegible] | [illegible] | [illegible] | [illegible] | [illegible] | [illegible] | [illegible] | [illegible] | [illegible] | [illegible] | [illegible] | [illegible] | [illegible] | [illegible] | [illegible] | [illegible] | [illegible] | [illegible] | [illegible] | [illegible] | [illegible] | [illegible] | [illegible] | [illegible] | [illegible] | [illegible] | [illegible] | [illegible] | [illegible] | [illegible] |
| PE | PE 42:4\|PE 18:0_24:4 | [M-H]- | 17.423 | 822.59357 | 0.075675667 | [illegible] | [illegible] | [illegible] | [illegible] | [illegible] | [illegible] | [illegible] | [illegible] | [illegible] | [illegible] | [illegible] | [illegible] | [illegible] | [illegible] | [illegible] | [illegible] | [illegible] | [illegible] | [illegible] | [illegible] | [illegible] | [illegible] | [illegible] | [illegible] | [illegible] | [illegible] | [illegible] | [illegible] | [illegible] | [illegible] | [illegible] | [illegible] | [illegible] | [illegible] | [illegible] |
| PE | PE 44:1\|PE 26:0_18:1 | [M-H]- | 21.13 | 856.68018 | 0.001484415 | [illegible] | [illegible] | [illegible] | [illegible] | [illegible] | [illegible] | [illegible] | [illegible] | [illegible] | [illegible] | [illegible] | [illegible] | [illegible] | [illegible] | [illegible] | [illegible] | [illegible] | [illegible] | [illegible] | [illegible] | [illegible] | [illegible] | [illegible] | [illegible] | [illegible] | [illegible] | [illegible] | [illegible] | [illegible] | [illegible] | [illegible] | [illegible] | [illegible] | [illegible] | [illegible] |
| PE | PE 44:4\|PE 24:0_20:4 | [M-H]- | 19.17 | 850.63141 | 0.000686988 | [illegible] | [illegible] | [illegible] | [illegible] | [illegible] | [illegible] | [illegible] | [illegible] | [illegible] | [illegible] | [illegible] | [illegible] | [illegible] | [illegible] | [illegible] | [illegible] | [illegible] | [illegible] | [illegible] | [illegible] | [illegible] | [illegible] | [illegible] | [illegible] | [illegible] | [illegible] | [illegible] | [illegible] | [illegible] | [illegible] | [illegible] | [illegible] | [illegible] | [illegible] | [illegible] |
| PE | PE O-34:1\|PE O-18:1_16:0 | [M-H]- | 17.354 | 702.54224 | 0.006425397 | [illegible] | [illegible] | [illegible] | [illegible] | [illegible] | [illegible] | [illegible] | [illegible] | [illegible] | [illegible] | [illegible] | [illegible] | [illegible] | [illegible] | [illegible] | [illegible] | [illegible] | [illegible] | [illegible] | [illegible] | [illegible] | [illegible] | [illegible] | [illegible] | [illegible] | [illegible] | [illegible] | [illegible] | [illegible] | [illegible] | [illegible] | [illegible] | [illegible] | [illegible] | [illegible] |
| PE | PE O-34:2\|PE O-16:1_18:1 | [M-H]- | 16.37 | 700.52673 | 0.02148077 | [illegible] | [illegible] | [illegible] | [illegible] | [illegible] | [illegible] | [illegible] | [illegible] | [illegible] | [illegible] | [illegible] | [illegible] | [illegible] | [illegible] | [illegible] | [illegible] | [illegible] | [illegible] | [illegible] | [illegible] | [illegible] | [illegible] | [illegible] | [illegible] | [illegible] | [illegible] | [illegible] | [illegible] | [illegible] | [illegible] | [illegible] | [illegible] | [illegible] | [illegible] | [illegible] |
| PE | PE O-36:2\|PE O-18:1_18:1 | [M-H]- | 17.468 | 728.55994 | 0.010831664 | [illegible] | [illegible] | [illegible] | [illegible] | [illegible] | [illegible] | [illegible] | [illegible] | [illegible] | [illegible] | [illegible] | [illegible] | [illegible] | [illegible] | [illegible] | [illegible] | [illegible] | [illegible] | [illegible] | [illegible] | [illegible] | [illegible] | [illegible] | [illegible] | [illegible] | [illegible] | [illegible] | [illegible] | [illegible] | [illegible] | [illegible] | [illegible] | [illegible] | [illegible] | [illegible] |
| PE | PE O-36:3\|PE O-18:2_18:1 | [M-H]- | 16.503 | 726.5423 | 0.007953172 | [illegible] | [illegible] | [illegible] | [illegible] | [illegible] | [illegible] | [illegible] | [illegible] | [illegible] | [illegible] | [illegible] | [illegible] | [illegible] | [illegible] | [illegible] | [illegible] | [illegible] | [illegible] | [illegible] | [illegible] | [illegible] | [illegible] | [illegible] | [illegible] | [illegible] | [illegible] | [illegible] | [illegible] | [illegible] | [illegible] | [illegible] | [illegible] | [illegible] | [illegible] | [illegible] |
| PE | PE O-36:5\|PE O-16:1_20:4 | [M-H]- | 15.413 | 722.51166 | 0.031923984 | [illegible] | [illegible] | [illegible] | [illegible] | [illegible] | [illegible] | [illegible] | [illegible] | [illegible] | [illegible] | [illegible] | [illegible] | [illegible] | [illegible] | [illegible] | [illegible] | [illegible] | [illegible] | [illegible] | [illegible] | [illegible] | [illegible] | [illegible] | [illegible] | [illegible] | [illegible] | [illegible] | [illegible] | [illegible] | [illegible] | [illegible] | [illegible] | [illegible] | [illegible] | [illegible] |
| PE | PE O-36:6\|PE O-16:1_20:5 | [M-H]- | 14.778 | 720.49579 | 0.003798565 | [illegible] | [illegible] | [illegible] | [illegible] | [illegible] | [illegible] | [illegible] | [illegible] | [illegible] | [illegible] | [illegible] | [illegible] | [illegible] | [illegible] | [illegible] | [illegible] | [illegible] | [illegible] | [illegible] | [illegible] | [illegible] | [illegible] | [illegible] | [illegible] | [illegible] | [illegible] | [illegible] | [illegible] | [illegible] | [illegible] | [illegible] | [illegible] | [illegible] | [illegible] | [illegible] |
| PE | PE O-37:5\|PE O-17:1_20:4 | [M-H]- | 15.935 | 736.52887 | 0.004820734 | [illegible] | [illegible] | [illegible] | [illegible] | [illegible] | [illegible] | [illegible] | [illegible] | [illegible] | [illegible] | [illegible] | [illegible] | [illegible] | [illegible] | [illegible] | [illegible] | [illegible] | [illegible] | [illegible] | [illegible] | [illegible] | [illegible] | [illegible] | [illegible] | [illegible] | [illegible] | [illegible] | [illegible] | [illegible] | [illegible] | [illegible] | [illegible] | [illegible] | [illegible] | [illegible] |
| PE | PE O-38:5\|PE O-18:1_20:4 | [M-H]- | 16.494 | 750.54419 | 0.082770651 | [illegible] | [illegible] | [illegible] | [illegible] | [illegible] | [illegible] | [illegible] | [illegible] | [illegible] | [illegible] | [illegible] | [illegible] | [illegible] | [illegible] | [illegible] | [illegible] | [illegible] | [illegible] | [illegible] | [illegible] | [illegible] | [illegible] | [illegible] | [illegible] | [illegible] | [illegible] | [illegible] | [illegible] | [illegible] | [illegible] | [illegible] | [illegible] | [illegible] | [illegible] | [illegible] |
| PE | PE O-38:6\|PE O-16:1_22:5 | [M-H]- | 15.523 | 748.52576 | 0.042180263 | [illegible] | [illegible] | [illegible] | [illegible] | [illegible] | [illegible] | [illegible] | [illegible] | [illegible] | [illegible] | [illegible] | [illegible] | [illegible] | [illegible] | [illegible] | [illegible] | [illegible] | [illegible] | [illegible] | [illegible] | [illegible] | [illegible] | [illegible] | [illegible] | [illegible] | [illegible] | [illegible] | [illegible] | [illegible] | [illegible] | [illegible] | [illegible] | [illegible] | [illegible] | [illegible] |
| PE | PE O-38:7\|PE O-16:1_22:6 | [M-H]- | 15.169 | 746.51263 | 0.027218981 | [illegible] | [illegible] | [illegible] | [illegible] | [illegible] | [illegible] | [illegible] | [illegible] | [illegible] | [illegible] | [illegible] | [illegible] | [illegible] | [illegible] | [illegible] | [illegible] | [illegible] | [illegible] | [illegible] | [illegible] | [illegible] | [illegible] | [illegible] | [illegible] | [illegible] | [illegible] | [illegible] | [illegible] | [illegible] | [illegible] | [illegible] | [illegible] | [illegible] | [illegible] | [illegible] |
| PE | PE O-39:6\|PE O-17:1_22:5 | [M-H]- | 16.015 | 762.54437 | 0.004187687 | [illegible] | [illegible] | [illegible] | [illegible] | [illegible] | [illegible] | [illegible] | [illegible] | [illegible] | [illegible] | [illegible] | [illegible] | [illegible] | [illegible] | [illegible] | [illegible] | [illegible] | [illegible] | [illegible] | [illegible] | [illegible] | [illegible] | [illegible] | [illegible] | [illegible] | [illegible] | [illegible] | [illegible] | [illegible] | [illegible] | [illegible] | [illegible] | [illegible] | [illegible] | [illegible] |
| PE | PE O-40:4\|PE O-18:0_22:4 | [M-H]- | 17.43 | 780.59009 | 0.004363473 | [illegible] | [illegible] | [illegible] | [illegible] | [illegible] | [illegible] | [illegible] | [illegible] | [illegible] | [illegible] | [illegible] | [illegible] | [illegible] | [illegible] | [illegible] | [illegible] | [illegible] | [illegible] | [illegible] | [illegible] | [illegible] | [illegible] | [illegible] | [illegible] | [illegible] | [illegible] | [illegible] | [illegible] | [illegible] | [illegible] | [illegible] | [illegible] | [illegible] | [illegible] | [illegible] |
| PE | PE O-40:5\|PE O-18:1_22:4 | [M-H]- | 17.258 | 778.57367 | 0.027849158 | [illegible] | [illegible] | [illegible] | [illegible] | [illegible] | [illegible] | [illegible] | [illegible] | [illegible] | [illegible] | [illegible] | [illegible] | [illegible] | [illegible] | [illegible] | [illegible] | [illegible] | [illegible] | [illegible] | [illegible] | [illegible] | [illegible] | [illegible] | [illegible] | [illegible] | [illegible] | [illegible] | [illegible] | [illegible] | [illegible] | [illegible] | [illegible] | [illegible] | [illegible] | [illegible] |
| PE | PE O-40:6\|PE O-18:1_22:5 | [M-H]- | 16.557 | 776.55737 | 0.050375304 | [illegible] | [illegible] | [illegible] | [illegible] | [illegible] | [illegible] | [illegible] | [illegible] | [illegible] | [illegible] | [illegible] | [illegible] | [illegible] | [illegible] | [illegible] | [illegible] | [illegible] | [illegible] | [illegible] | [illegible] | [illegible] | [illegible] | [illegible] | [illegible] | [illegible] | [illegible] | [illegible] | [illegible] | [illegible] | [illegible] | [illegible] | [illegible] | [illegible] | [illegible] | [illegible] |
| PE | PE O-40:7\|PE O-18:1_22:6 | [M-H]- | 16.233 | 774.54242 | 0.038976961 | [illegible] | [illegible] | [illegible] | [illegible] | [illegible] | [illegible] | [illegible] | [illegible] | [illegible] | [illegible] | [illegible] | [illegible] | [illegible] | [illegible] | [illegible] | [illegible] | [illegible] | [illegible] | [illegible] | [illegible] | [illegible] | [illegible] | [illegible] | [illegible] | [illegible] | [illegible] | [illegible] | [illegible] | [illegible] | [illegible] | [illegible] | [illegible] | [illegible] | [illegible] | [illegible] |
| PE | PE O-40:8\|PE O-18:2_22:6 | [M-H]- | 15.266 | 772.52777 | 0.008738189 | [illegible] | [illegible] | [illegible] | [illegible] | [illegible] | [illegible] | [illegible] | [illegible] | [illegible] | [illegible] | [illegible] | [illegible] | [illegible] | [illegible] | [illegible] | [illegible] | [illegible] | [illegible] | [illegible] | [illegible] | [illegible] | [illegible] | [illegible] | [illegible] | [illegible] | [illegible] | [illegible] | [illegible] | [illegible] | [illegible] | [illegible] | [illegible] | [illegible] | [illegible] | [illegible] |
| PE | PE O-42:7\|PE O-20:1_22:6 | [M-H]- | 17.309 | 802.57458 | 0.000903016 | [illegible] | [illegible] | [illegible] | [illegible] | [illegible] | [illegible] | [illegible] | [illegible] | [illegible] | [illegible] | [illegible] | [illegible] | [illegible] | [illegible] | [illegible] | [illegible] | [illegible] | [illegible] | [illegible] | [illegible] | [illegible] | [illegible] | [illegible] | [illegible] | [illegible] | [illegible] | [illegible] | [illegible] | [illegible] | [illegible] | [illegible] | [illegible] | [illegible] | [illegible] | [illegible] |
| PS | PS 34:1\|PS 16:0_18:1 | [M-H]- | 13.595 | 760.5116 | 0.016578828 | [illegible] | [illegible] | [illegible] | [illegible] | [illegible] | [illegible] | [illegible] | [illegible] | [illegible] | [illegible] | [illegible] | [illegible] | [illegible] | [illegible] | [illegible] | [illegible] | [illegible] | [illegible] | [illegible] | [illegible] | [illegible] | [illegible] | [illegible] | [illegible] | [illegible] | [illegible] | [illegible] | [illegible] | [illegible] | [illegible] | [illegible] | [illegible] | [illegible] | [illegible] | [illegible] |
| PS | PS 36:1\|PS 18:0_18:1 | [M-H]- | 14.429 | 788.54376 | 0.109601858 | [illegible] | [illegible] | [illegible] | [illegible] | [illegible] | [illegible] | [illegible] | [illegible] | [illegible] | [illegible] | [illegible] | [illegible] | [illegible] | [illegible] | [illegible] | [illegible] | [illegible] | [illegible] | [illegible] | [illegible] | [illegible] | [illegible] | [illegible] | [illegible] | [illegible] | [illegible] | [illegible] | [illegible] | [illegible] | [illegible] | [illegible] | [illegible] | [illegible] | [illegible] | [illegible] |
| PS | PS 36:2\|PS 18:0_18:2 | [M-H]- | 13.788 | 786.52679 | 0.017967504 | [illegible] | [illegible] | [illegible] | [illegible] | [illegible] | [illegible] | [illegible] | [illegible] | [illegible] | [illegible] | [illegible] | [illegible] | [illegible] | [illegible] | [illegible] | [illegible] | [illegible] | [illegible] | [illegible] | [illegible] | [illegible] | [illegible] | [illegible] | [illegible] | [illegible] | [illegible] | [illegible] | [illegible] | [illegible] | [illegible] | [illegible] | [illegible] | [illegible] | [illegible] | [illegible] |
| PS | PS 38:3\|PS 18:0_20:3 | [M-H]- | 14.069 | 812.54382 | 0.016154772 | [illegible] | [illegible] | [illegible] | [illegible] | [illegible] | [illegible] | [illegible] | [illegible] | [illegible] | [illegible] | [illegible] | [illegible] | [illegible] | [illegible] | [illegible] | [illegible] | [illegible] | [illegible] | [illegible] | [illegible] | [illegible] | [illegible] | [illegible] | [illegible] | [illegible] | [illegible] | [illegible] | [illegible] | [illegible] | [illegible] | [illegible] | [illegible] | [illegible] | [illegible] | [illegible] |
| PI | PI 32:1\|PI 16:0_16:1 | [M-H]- | 12.749 | 807.50055 | 0.00268775 | [illegible] | [illegible] | [illegible] | [illegible] | [illegible] | [illegible] | [illegible] | [illegible] | [illegible] | [illegible] | [illegible] | [illegible] | [illegible] | [illegible] | [illegible] | [illegible] | [illegible] | [illegible] | [illegible] | [illegible] | [illegible] | [illegible] | [illegible] | [illegible] | [illegible] | [illegible] | [illegible] | [illegible] | [illegible] | [illegible] | [illegible] | [illegible] | [illegible] | [illegible] | [illegible] |
| PI | PI 34:0\|PI 16:0_18:0 | [M-H]- | 13.996 | 837.54889 | 0.006660561 | [illegible] | [illegible] | [illegible] | [illegible] | [illegible] | [illegible] | [illegible] | [illegible] | [illegible] | [illegible] | [illegible] | [illegible] | [illegible] | [illegible] | [illegible] | [illegible] | [illegible] | [illegible] | [illegible] | [illegible] | [illegible] | [illegible] | [illegible] | [illegible] | [illegible] | [illegible] | [illegible] | [illegible] | [illegible] | [illegible] | [illegible] | [illegible] | [illegible] | [illegible] | [illegible] |
| PI | PI 34:1\|PI 16:0_18:1 | [M-H]- | 13.38 | 835.53247 | 0.027389183 | [illegible] | [illegible] | [illegible] | [illegible] | [illegible] | [illegible] | [illegible] | [illegible] | [illegible] | [illegible] | [illegible] | [illegible] | [illegible] | [illegible] | [illegible] | [illegible] | [illegible] | [illegible] | [illegible] | [illegible] | [illegible] | [illegible] | [illegible] | [illegible] | [illegible] | [illegible] | [illegible] | [illegible] | [illegible] | [illegible] | [illegible] | [illegible] | [illegible] | [illegible] | [illegible] |
| PI | PI 34:2\|PI 16:1_18:1 | [M-H]- | 12.861 | 833.51685 | 0.011656274 | [illegible] | [illegible] | [illegible] | [illegible] | [illegible] | [illegible] | [illegible] | [illegible] | [illegible] | [illegible] | [illegible] | [illegible] | [illegible] | [illegible] | [illegible] | [illegible] | [illegible] | [illegible] | [illegible] | [illegible] | [illegible] | [illegible] | [illegible] | [illegible] | [illegible] | [illegible] | [illegible] | [illegible] | [illegible] | [illegible] | [illegible] | [illegible] | [illegible] | [illegible] | [illegible] |
| PI | PI 34:3\|PI 16:0_18:3 | [M-H]- | 12.663 | 831.50183 | 0.001850122 | [illegible] | [illegible] | [illegible] | [illegible] | [illegible] | [illegible] | [illegible] | [illegible] | [illegible] | [illegible] | [illegible] | [illegible] | [illegible] | [illegible] | [illegible] | [illegible] | [illegible] | [illegible] | [illegible] | [illegible] | [illegible] | [illegible] | [illegible] | [illegible] | [illegible] | [illegible] | [illegible] | [illegible] | [illegible] | [illegible] | [illegible] | [illegible] | [illegible] | [illegible] | [illegible] |
| PI | PI 35:1 | [M-H]- | 13.754 | 849.55029 | 0.004287615 | [illegible] | [illegible] | [illegible] | [illegible] | [illegible] | [illegible] | [illegible] | [illegible] | [illegible] | [illegible] | [illegible] | [illegible] | [illegible] | [illegible] | [illegible] | [illegible] | [illegible] | [illegible] | [illegible] | [illegible] | [illegible] | [illegible] | [illegible] | [illegible] | [illegible] | [illegible] | [illegible] | [illegible] | [illegible] | [illegible] | [illegible] | [illegible] | [illegible] | [illegible] | [illegible] |
| PI | PI 35:2\|PI 17:1_18:1 | [M-H]- | 13.169 | 847.5307 | 0.00111822 | [illegible] | [illegible] | [illegible] | [illegible] | [illegible] | [illegible] | [illegible] | [illegible] | [illegible] | [illegible] | [illegible] | [illegible] | [illegible] | [illegible] | [illegible] | [illegible] | [illegible] | [illegible] | [illegible] | [illegible] | [illegible] | [illegible] | [illegible] | [illegible] | [illegible] | [illegible] | [illegible] | [illegible] | [illegible] | [illegible] | [illegible] | [illegible] | [illegible] | [illegible] | [illegible] |
| PI | PI 36:1\|PI 18:0_18:1 | [M-H]- | 14.128 | 863.56299 | 0.035241275 | [illegible] | [illegible] | [illegible] | [illegible] | [illegible] | [illegible] | [illegible] | [illegible] | [illegible] | [illegible] | [illegible] | [illegible] | [illegible] | [illegible] | [illegible] | [illegible] | [illegible] | [illegible] | [illegible] | [illegible] | [illegible] | [illegible] | [illegible] | [illegible] | [illegible] | [illegible] | [illegible] | [illegible] | [illegible] | [illegible] | [illegible] | [illegible] | [illegible] | [illegible] | [illegible] |
| PI | PI 36:2\|PI 18:1_18:1 | [M-H]- | 13.568 | 861.54761 | 0.044004422 | [illegible] | [illegible] | [illegible] | [illegible] | [illegible] | [illegible] | [illegible] | [illegible] | [illegible] | [illegible] | [illegible] | [illegible] | [illegible] | [illegible] | [illegible] | [illegible] | [illegible] | [illegible] | [illegible] | [illegible] | [illegible] | [illegible] | [illegible] | [illegible] | [illegible] | [illegible] | [illegible] | [illegible] | [illegible] | [illegible] | [illegible] | [illegible] | [illegible] | [illegible] | [illegible] |
| PI | PI 36:3\|PI 18:1_18:2 | [M-H]- | 12.993 | 859.53192 | 0.021911295 | [illegible] | [illegible] | [illegible] | [illegible] | [illegible] | [illegible] | [illegible] | [illegible] | [illegible] | [illegible] | [illegible] | [illegible] | [illegible] | [illegible] | [illegible] | [illegible] | [illegible] | [illegible] | [illegible] | [illegible] | [illegible] | [illegible] | [illegible] | [illegible] | [illegible] | [illegible] | [illegible] | [illegible] | [illegible] | [illegible] | [illegible] | [illegible] | [illegible] | [illegible] | [illegible] |
| PI | PI 36:4\|PI 16:0_20:4 | [M-H]- | 12.849 | 857.51538 | 0.015100314 | [illegible] | [illegible] | [illegible] | [illegible] | [illegible] | [illegible] | [illegible] | [illegible] | [illegible] | [illegible] | [illegible] | [illegible] | [illegible] | [illegible] | [illegible] | [illegible] | [illegible] | [illegible] | [illegible] | [illegible] | [illegible] | [illegible] | [illegible] | [illegible] | [illegible] | [illegible] | [illegible] | [illegible] | [illegible] | [illegible] | [illegible] | [illegible] | [illegible] | [illegible] | [illegible] |
| PI | PI 36:5\|PI 16:1_20:4 | [M-H]- | 12.42 | 855.50024 | 0.000850319 | [illegible] | [illegible] | [illegible] | [illegible] | [illegible] | [illegible] | [illegible] | [illegible] | [illegible] | [illegible] | [illegible] | [illegible] | [illegible] | [illegible] | [illegible] | [illegible] | [illegible] | [illegible] | [illegible] | [illegible] | [illegible] | [illegible] | [illegible] | [illegible] | [illegible] | [illegible] | [illegible] | [illegible] | [illegible] | [illegible] | [illegible] | [illegible] | [illegible] | [illegible] | [illegible] |
| PI | PI 37:2\|PI 18:1_19:1 | [M-H]- | 13.862 | 875.56287 | 0.001545165 | [illegible] | [illegible] | [illegible] | [illegible] | [illegible] | [illegible] | [illegible] | [illegible] | [illegible] | [illegible] | [illegible] | [illegible] | [illegible] | [illegible] | [illegible] | [illegible] | [illegible] | [illegible] | [illegible] | [illegible] | [illegible] | [illegible] | [illegible] | [illegible] | [illegible] | [illegible] | [illegible] | [illegible] | [illegible] | [illegible] | [illegible] | [illegible] | [illegible] | [illegible] | [illegible] |
| PI | PI 37:4\|PI 17:0_20:4 | [M-H]- | 13.191 | 871.53351 | 0.003903502 | [illegible] | [illegible] | [illegible] | [illegible] | [illegible] | [illegible] | [illegible] | [illegible] | [illegible] | [illegible] | [illegible] | [illegible] | [illegible] | [illegible] | [illegible] | [illegible] | [illegible] | [illegible] | [illegible] | [illegible] | [illegible] | [illegible] | [illegible] | [illegible] | [illegible] | [illegible] | [illegible] | [illegible] | [illegible] | [illegible] | [illegible] | [illegible] | [illegible] | [illegible] | [illegible] |
| PI | PI 38:3\|PI 18:0_20:3 | [M-H]- | 13.801 | 887.55914 | 0.036267332 | [illegible] | [illegible] | [illegible] | [illegible] | [illegible] | [illegible] | [illegible] | [illegible] | [illegible] | [illegible] | [illegible] | [illegible] | [illegible] | [illegible] | [illegible] | [illegible] | [illegible] | [illegible] | [illegible] | [illegible] | [illegible] | [illegible] | [illegible] | [illegible] | [illegible] | [illegible] | [illegible] | [illegible] | [illegible] | [illegible] | [illegible] | [illegible] | [illegible] | [illegible] | [illegible] |
| PI | PI 38:4\|PI 18:0_20:4 | [M-H]- | 13.535 | 885.54761 | 0.448251769 | [illegible] | [illegible] | [illegible] | [illegible] | [illegible] | [illegible] | [illegible] | [illegible] | [illegible] | [illegible] | [illegible] | [illegible] | [illegible] | [illegible] | [illegible] | [illegible] | [illegible] | [illegible] | [illegible] | [illegible] | [illegible] | [illegible] | [illegible] | [illegible] | [illegible] | [illegible] | [illegible] | [illegible] | [illegible] | [illegible] | [illegible] | [illegible] | [illegible] | [illegible] | [illegible] |
| PI | PI 38:5\|PI 18:1_20:4 | [M-H]- | 12.92 | 883.53107 | 0.059418877 | [illegible] | [illegible] | [illegible] | [illegible] | [illegible] | [illegible] | [illegible] | [illegible] | [illegible] | [illegible] | [illegible] | [illegible] | [illegible] | [illegible] | [illegible] | [illegible] | [illegible] | [illegible] | [illegible] | [illegible] | [illegible] | [illegible] | [illegible] | [illegible] | [illegible] | [illegible] | [illegible] | [illegible] | [illegible] | [illegible] | [illegible] | [illegible] | [illegible] | [illegible] | [illegible] |
| PI | PI 38:6\|PI 16:0_22:6 | [M-H]- | 12.675 | 881.51813 | 0.004456498 | [illegible] | [illegible] | [illegible] | [illegible] | [illegible] | [illegible] | [illegible] | [illegible] | [illegible] | [illegible] | [illegible] | [illegible] | [illegible] | [illegible] | [illegible] | [illegible] | [illegible] | [illegible] | [illegible] | [illegible] | [illegible] | [illegible] | [illegible] | [illegible] | [illegible] | [illegible] | [illegible] | [illegible] | [illegible] | [illegible] | [illegible] | [illegible] | [illegible] | [illegible] | [illegible] |
| PI | PI 39:3\|PI 18:0_21:3 | [M-H]- | 14.271 | 901.5813 | 0.001682951 | [illegible] | [illegible] | [illegible] | [illegible] | [illegible] | [illegible] | [illegible] | [illegible] | [illegible] | [illegible] | [illegible] | [illegible] | [illegible] | [illegible] | [illegible] | [illegible] | [illegible] | [illegible] | [illegible] | [illegible] | [illegible] | [illegible] | [illegible] | [illegible] | [illegible] | [illegible] | [illegible] | [illegible] | [illegible] | [illegible] | [illegible] | [illegible] | [illegible] | [illegible] | [illegible] |
| PI | PI 39:4\|PI 19:0_20:4 | [M-H]- | 13.919 | 899.56458 | 0.00200527 | [illegible] | [illegible] | [illegible] | [illegible] | [illegible] | [illegible] | [illegible] | [illegible] | [illegible] | [illegible] | [illegible] | [illegible] | [illegible] | [illegible] | [illegible] | [illegible] | [illegible] | [illegible] | [illegible] | [illegible] | [illegible] | [illegible] | [illegible] | [illegible] | [illegible] | [illegible] | [illegible] | [illegible] | [illegible] | [illegible] | [illegible] | [illegible] | [illegible] | [illegible] | [illegible] |
| PI | PI 40:4\|PI 18:0_22:4 | [M-H]- | 14.077 | 913.58081 | 0.013988697 | [illegible] | [illegible] | [illegible] | [illegible] | [illegible] | [illegible] | [illegible] | [illegible] | [illegible] | [illegible] | [illegible] | [illegible] | [illegible] | [illegible] | [illegible] | [illegible] | [illegible] | [illegible] | [illegible] | [illegible] | [illegible] | [illegible] | [illegible] | [illegible] | [illegible] | [illegible] | [illegible] | [illegible] | [illegible] | [illegible] | [illegible] | [illegible] | [illegible] | [illegible] | [illegible] |
| PI | PI 40:5\|PI 18:0_22:5 | [M-H]- | 13.578 | 911.56494 | 0.023512188 | [illegible] | [illegible] | [illegible] | [illegible] | [illegible] | [illegible] | [illegible] | [illegible] | [illegible] | [illegible] | [illegible] | [illegible] | [illegible] | [illegible] | [illegible] | [illegible] | [illegible] | [illegible] | [illegible] | [illegible] | [illegible] | [illegible] | [illegible] | [illegible] | [illegible] | [illegible] | [illegible] | [illegible] | [illegible] | [illegible] | [illegible] | [illegible] | [illegible] | [illegible] | [illegible] |
| PI | PI 40:6\|PI 18:0_22:6 | [M-H]- | 13.388 | 909.54712 | 0.009665958 | [illegible] | [illegible] | [illegible] | [illegible] | [illegible] | [illegible] | [illegible] | [illegible] | [illegible] | [illegible] | [illegible] | [illegible] | [illegible] | [illegible] | [illegible] | [illegible] | [illegible] | [illegible] | [illegible] | [illegible] | [illegible] | [illegible] | [illegible] | [illegible] | [illegible] | [illegible] | [illegible] | [illegible] | [illegible] | [illegible] | [illegible] | [illegible] | [illegible] | [illegible] | [illegible] |
| PI | PI 40:6\|PI 18:1_22:5 | [M-H]- | 13.017 | 909.55518 | 0.00999777 | [illegible] | [illegible] | [illegible] | [illegible] | [illegible] | [illegible] | [illegible] | [illegible] | [illegible] | [illegible] | [illegible] | [illegible] | [illegible] | [illegible] | [illegible] | [illegible] | [illegible] | [illegible] | [illegible] | [illegible] | [illegible] | [illegible] | [illegible] | [illegible] | [illegible] | [illegible] | [illegible] | [illegible] | [illegible] | [illegible] | [illegible] | [illegible] | [illegible] | [illegible] | [illegible] |
| PI | PI 40:8\|PI 20:4_20:4 | [M-H]- | 12.446 | 905.51807 | 0.000518941 | [illegible] | [illegible] | [illegible] | [illegible] | [illegible] | [illegible] | [illegible] | [illegible] | [illegible] | [illegible] | [illegible] | [illegible] | [illegible] | [illegible] | [illegible] | [illegible] | [illegible] | [illegible] | [illegible] | [illegible] | [illegible] | [illegible] | [illegible] | [illegible] | [illegible] | [illegible] | [illegible] | [illegible] | [illegible] | [illegible] | [illegible] | [illegible] | [illegible] | [illegible] | [illegible] |
| LPC | LPC 15:0/0:0 | [M+H]+ | 10.121 | 482.32382 | 0.021534707 | [illegible] | [illegible] | [illegible] | [illegible] | [illegible] | [illegible] | [illegible] | [illegible] | [illegible] | [illegible] | [illegible] | [illegible] | [illegible] | [illegible] | [illegible] | [illegible] | [illegible] | [illegible] | [illegible] | [illegible] | [illegible] | [illegible] | [illegible] | [illegible] | [illegible] | [illegible] | [illegible] | [illegible] | [illegible] | [illegible] | [illegible] | [illegible] | [illegible] | [illegible] | [illegible] |
| LPC | LPC 16:0/0:0 | [M+H]+ | 10.699 | 496.3396 | 0.029590434 | [illegible] | [illegible] | [illegible] | [illegible] | [illegible] | [illegible] | [illegible] | [illegible] | [illegible] | [illegible] | [illegible] | [illegible] | [illegible] | [illegible] | [illegible] | [illegible] | [illegible] | [illegible] | [illegible] | [illegible] | [illegible] | [illegible] | [illegible] | [illegible] | [illegible] | [illegible] | [illegible] | [illegible] | [illegible] | [illegible] | [illegible] | [illegible] | [illegible] | [illegible] | [illegible] |
| LPC | LPC 18:0/0:0 | [M+H]+ | 11.477 | 524.37036 | 0.021016745 | [illegible] | [illegible] | [illegible] | [illegible] | [illegible] | [illegible] | [illegible] | [illegible] | [illegible] | [illegible] | [illegible] | [illegible] | [illegible] | [illegible] | [illegible] | [illegible] | [illegible] | [illegible] | [illegible] | [illegible] | [illegible] | [illegible] | [illegible] | [illegible] | [illegible] | [illegible] | [illegible] | [illegible] | [illegible] | [illegible] | [illegible] | [illegible] | [illegible] | [illegible] | [illegible] |
| LPC | LPC 18:1/0:0 | [M+H]+ | 10.872 | 522.3559 | 0.019567779 | [illegible] | [illegible] | [illegible] | [illegible] | [illegible] | [illegible] | [illegible] | [illegible] | [illegible] | [illegible] | [illegible] | [illegible] | [illegible] | [illegible] | [illegible] | [illegible] | [illegible] | [illegible] | [illegible] | [illegible] | [illegible] | [illegible] | [illegible] | [illegible] | [illegible] | [illegible] | [illegible] | [illegible] | [illegible] | [illegible] | [illegible] | [illegible] | [illegible] | [illegible] | [illegible] |
| LPC | LPC 20:1 | [M+H]+ | 12.039 | 550.38678 | 0.009873565 | [illegible] | [illegible] | [illegible] | [illegible] | [illegible] | [illegible] | [illegible] | [illegible] | [illegible] | [illegible] | [illegible] | [illegible] | [illegible] | [illegible] | [illegible] | [illegible] | [illegible] | [illegible] | [illegible] | [illegible] | [illegible] | [illegible] | [illegible] | [illegible] | [illegible] | [illegible] | [illegible] | [illegible] | [illegible] | [illegible] | [illegible] | [illegible] | [illegible] | [illegible] | [illegible] |
| LPC | LPC 26:0/0:0 | [M+H]+ | 15.216 | 636.49634 | 0.324103898 | [illegible] | [illegible] | [illegible] | [illegible] | [illegible] | [illegible] | [illegible] | [illegible] | [illegible] | [illegible] | [illegible] | [illegible] | [illegible] | [illegible] | [illegible] | [illegible] | [illegible] | [illegible] | [illegible] | [illegible] | [illegible] | [illegible] | [illegible] | [illegible] | [illegible] | [illegible] | [illegible] | [illegible] | [illegible] | [illegible] | [illegible] | [illegible] | [illegible] | [illegible] | [illegible] |
| LPE | LPE 18:0 | [M-H]- | 11.294 | 480.30878 | 0.006284881 | [illegible] | [illegible] | [illegible] | [illegible] | [illegible] | [illegible] | [illegible] | [illegible] | [illegible] | [illegible] | [illegible] | [illegible] | [illegible] | [illegible] | [illegible] | [illegible] | [illegible] | [illegible] | [illegible] | [illegible] | [illegible] | [illegible] | [illegible] | [illegible] | [illegible] | [illegible] | [illegible] | [illegible] | [illegible] | [illegible] | [illegible] | [illegible] | [illegible] | [illegible] | [illegible] |
| LPE | LPE 20:4 | [M-H]- | 9.659 | 500.27823 | 0.001781656 | [illegible] | [illegible] | [illegible] | [illegible] | [illegible] | [illegible] | [illegible] | [illegible] | [illegible] | [illegible] | [illegible] | [illegible] | [illegible] | [illegible] | [illegible] | [illegible] | [illegible] | [illegible] | [illegible] | [illegible] | [illegible] | [illegible] | [illegible] | [illegible] | [illegible] | [illegible] | [illegible] | [illegible] | [illegible] | [illegible] | [illegible] | [illegible] | [illegible] | [illegible] | [illegible] |
| LPE | LPE 22:6 | [M-H]- | 9.643 | 524.27814 | 0.001354187 | [illegible] | [illegible] | [illegible] | [illegible] | [illegible] | [illegible] | [illegible] | [illegible] | [illegible] | [illegible] | [illegible] | [illegible] | [illegible] | [illegible] | [illegible] | [illegible] | [illegible] | [illegible] | [illegible] | [illegible] | [illegible] | [illegible] | [illegible] | [illegible] | [illegible] | [illegible] | [illegible] | [illegible] | [illegible] | [illegible] | [illegible] | [illegible] | [illegible] | [illegible] | [illegible] |
| LPE | LPE O-18:1 | [M-H]- | 17.413 | 464.31372 | 0.028141887 | [illegible] | [illegible] | [illegible] | [illegible] | [illegible] | [illegible] | [illegible] | [illegible] | [illegible] | [illegible] | [illegible] | [illegible] | [illegible] | [illegible] | [illegible] | [illegible] | [illegible] | [illegible] | [illegible] | [illegible] | [illegible] | [illegible] | [illegible] | [illegible] | [illegible] | [illegible] | [illegible] | [illegible] | [illegible] | [illegible] | [illegible] | [illegible] | [illegible] | [illegible] | [illegible] |
| LPE | LPE-N (FA)40:7\|LPE-N (FA 18:1)22:6 | [M-H]- | 14.76 | 788.52356 | 0.024803043 | [illegible] | [illegible] | [illegible] | [illegible] | [illegible] | [illegible] | [illegible] | [illegible] | [illegible] | [illegible] | [illegible] | [illegible] | [illegible] | [illegible] | [illegible] | [illegible] | [illegible] | [illegible] | [illegible] | [illegible] | [illegible] | [illegible] | [illegible] | [illegible] | [illegible] | [illegible] | [illegible] | [illegible] | [illegible] | [illegible] | [illegible] | [illegible] | [illegible] | [illegible] | [illegible] |
| TG | TG 26:0\|TG 8:0_8:0_10:0 | [M+NH4]+ | 14.094 | 516.42523 | 0.019027205 | [illegible] | [illegible] | [illegible] | [illegible] | [illegible] | [illegible] | [illegible] | [illegible] | [illegible] | [illegible] | [illegible] | [illegible] | [illegible] | [illegible] | [illegible] | [illegible] | [illegible] | [illegible] | [illegible] | [illegible] | [illegible] | [illegible] | [illegible] | [illegible] | [illegible] | [illegible] | [illegible] | [illegible] | [illegible] | [illegible] | [illegible] | [illegible] | [illegible] | [illegible] | [illegible] |
| TG | TG 27:0\|TG 9:0_9:0_9:0 | [M+NH4]+ | 13.739 | 530.44177 | 0.00350489 | [illegible] | [illegible] | [illegible] | [illegible] | [illegible] | [illegible] | [illegible] | [illegible] | [illegible] | [illegible] | [illegible] | [illegible] | [illegible] | [illegible] | [illegible] | [illegible] | [illegible] | [illegible] | [illegible] | [illegible] | [illegible] | [illegible] | [illegible] | [illegible] | [illegible] | [illegible] | [illegible] | [illegible] | [illegible] | [illegible] | [illegible] | [illegible] | [illegible] | [illegible] | [illegible] |
| TG | TG 28:0\|TG 8:0_10:0_10:0 | [M+NH4]+ | 15.013 | 544.4566 | 0.010874646 | [illegible] | [illegible] | [illegible] | [illegible] | [illegible] | [illegible] | [illegible] | [illegible] | [illegible] | [illegible] | [illegible] | [illegible] | [illegible] | [illegible] | [illegible] | [illegible] | [illegible] | [illegible] | [illegible] | [illegible] | [illegible] | [illegible] | [illegible] | [illegible] | [illegible] | [illegible] | [illegible] | [illegible] | [illegible] | [illegible] | [illegible] | [illegible] | [illegible] | [illegible] | [illegible] |
| TG | TG 30:0\|TG 8:0_10:0_12:0 | [M+NH4]+ | 16.015 | 572.48828 | 0.005178532 | [illegible] | [illegible] | [illegible] | [illegible] | [illegible] | [illegible] | [illegible] | [illegible] | [illegible] | [illegible] | [illegible] | [illegible] | [illegible] | [illegible] | [illegible] | [illegible] | [illegible] | [illegible] | [illegible] | [illegible] | [illegible] | [illegible] | [illegible] | [illegible] | [illegible] | [illegible] | [illegible] | [illegible] | [illegible] | [illegible] | [illegible] | [illegible] | [illegible] | [illegible] | [illegible] |
| TG | TG 34:0\|TG 8:0_12:0_14:0 | [M+NH4]+ | 18.223 | 628.55017 | 0.01665893 | [illegible] | [illegible] | [illegible] | [illegible] | [illegible] | [illegible] | [illegible] | [illegible] | [illegible] | [illegible] | [illegible] | [illegible] | [illegible] | [illegible] | [illegible] | [illegible] | [illegible] | [illegible] | [illegible] | [illegible] | [illegible] | [illegible] | [illegible] | [illegible] | [illegible] | [illegible] | [illegible] | [illegible] | [illegible] | [illegible] | [illegible] | [illegible] | [illegible] | [illegible] | [illegible] |
| TG | TG 36:0\|TG 8:0_10:0_18:0 | [M+NH4]+ | 19.254 | 656.58368 | 0.022293285 | [illegible] | [illegible] | [illegible] | [illegible] | [illegible] | [illegible] | [illegible] | [illegible] | [illegible] | [illegible] | [illegible] | [illegible] | [illegible] | [illegible] | [illegible] | [illegible] | [illegible] | [illegible] | [illegible] | [illegible] | [illegible] | [illegible] | [illegible] | [illegible] | [illegible] | [illegible] | [illegible] | [illegible] | [illegible] | [illegible] | [illegible] | [illegible] | [illegible] | [illegible] | [illegible] |
| TG | TG 38:0\|TG 12:0_12:0_14:0 | [M+NH4]+ | 20.162 | 684.6134 | 0.020053386 | [illegible] | [illegible] | [illegible] | [illegible] | [illegible] | [illegible] | [illegible] | [illegible] | [illegible] | [illegible] | [illegible] | [illegible] | [illegible] | [illegible] | [illegible] | [illegible] | [illegible] | [illegible] | [illegible] | [illegible] | [illegible] | [illegible] | [illegible] | [illegible] | [illegible] | [illegible] | [illegible] | [illegible] | [illegible] | [illegible] | [illegible] | [illegible] | [illegible] | [illegible] | [illegible] |
| TG | TG 40:0\|TG 10:0_14:0_16:0 | [M+NH4]+ | 21.078 | 712.64563 | 0.013255627 | [illegible] | [illegible] | [illegible] | [illegible] | [illegible] | [illegible] | [illegible] | [illegible] | [illegible] | [illegible] | [illegible] | [illegible] | [illegible] | [illegible] | [illegible] | [illegible] | [illegible] | [illegible] | [illegible] | [illegible] | [illegible] | [illegible] | [illegible] | [illegible] | [illegible] | [illegible] | [illegible] | [illegible] | [illegible] | [illegible] | [illegible] | [illegible] | [illegible] | [illegible] | [illegible] |
| TG | TG 42:0\|TG 12:0_14:0_16:0 | [M+NH4]+ | 21.916 | 740.67682 | 0.016315979 | [illegible] | [illegible] | [illegible] | [illegible] | [illegible] | [illegible] | [illegible] | [illegible] | [illegible] | [illegible] | [illegible] | [illegible] | [illegible] | [illegible] | [illegible] | [illegible] | [illegible] | [illegible] | [illegible] | [illegible] | [illegible] | [illegible] | [illegible] | [illegible] | [illegible] | [illegible] | [illegible] | [illegible] | [illegible] | [illegible] | [illegible] | [illegible] | [illegible] | [illegible] | [illegible] |
| TG | TG 42:1\|TG 12:0_14:0_16:1 | [M+NH4]+ | 21.205 | 738.65997 | 0.003585129 | [illegible] | [illegible] | [illegible] | [illegible] | [illegible] | [illegible] | [illegible] | [illegible] | [illegible] | [illegible] | [illegible] | [illegible] | [illegible] | [illegible] | [illegible] | [illegible] | [illegible] | [illegible] | [illegible] | [illegible] | [illegible] | [illegible] | [illegible] | [illegible] | [illegible] | [illegible] | [illegible] | [illegible] | [illegible] | [illegible] | [illegible] | [illegible] | [illegible] | [illegible] | [illegible] |
| TG | TG 44:0\|TG 14:0_14:0_16:0 | [M+NH4]+ | 22.681 | 768.70734 | 0.03517044 | [illegible] | [illegible] | [illegible] | [illegible] | [illegible] | [illegible] | [illegible] | [illegible] | [illegible] | [illegible] | [illegible] | [illegible] | [illegible] | [illegible] | [illegible] | [illegible] | [illegible] | [illegible] | [illegible] | [illegible] | [illegible] | [illegible] | [illegible] | [illegible] | [illegible] | [illegible] | [illegible] | [illegible] | [illegible] | [illegible] | [illegible] | [illegible] | [illegible] | [illegible] | [illegible] |
| TG | TG 44:1\|TG 12:0_16:0_16:1 | [M+NH4]+ | 22.006 | 766.69159 | 0.023881341 | [illegible] | [illegible] | [illegible] | [illegible] | [illegible] | [illegible] | [illegible] | [illegible] | [illegible] | [illegible] | [illegible] | [illegible] | [illegible] | [illegible] | [illegible] | [illegible] | [illegible] | [illegible] | [illegible] | [illegible] | [illegible] | [illegible] | [illegible] | [illegible] | [illegible] | [illegible] | [illegible] | [illegible] | [illegible] | [illegible] | [illegible] | [illegible] | [illegible] | [illegible] | [illegible] |
| TG | TG 44:2\|TG 12:0_14:0_18:2 | [M+NH4]+ | 21.344 | 764.67554 | 0.008583697 | [illegible] | [illegible] | [illegible] | [illegible] | [illegible] | [illegible] | [illegible] | [illegible] | [illegible] | [illegible] | [illegible] | [illegible] | [illegible] | [illegible] | [illegible] | [illegible] | [illegible] | [illegible] | [illegible] | [illegible] | [illegible] | [illegible] | [illegible] | [illegible] | [illegible] | [illegible] | [illegible] | [illegible] | [illegible] | [illegible] | [illegible] | [illegible] | [illegible] | [illegible] | [illegible] |
| TG | TG 44:3\|TG 14:0_14:0_16:3 | [M+NH4]+ | 20.922 | 762.65894 | 0.001789444 | [illegible] | [illegible] | [illegible] | [illegible] | [illegible] | [illegible] | [illegible] | [illegible] | [illegible] | [illegible] | [illegible] | [illegible] | [illegible] | [illegible] | [illegible] | [illegible] | [illegible] | [illegible] | [illegible] | [illegible] | [illegible] | [illegible] | [illegible] | [illegible] | [illegible] | [illegible] | [illegible] | [illegible] | [illegible] | [illegible] | [illegible] | [illegible] | [illegible] | [illegible] | [illegible] |
| TG | TG 45:0\|TG 14:0_15:0_16:0 | [M+NH4]+ | 23.031 | 782.72284 | 0.060937916 | [illegible] | [illegible] | [illegible] | [illegible] | [illegible] | [illegible] | [illegible] | [illegible] | [illegible] | [illegible] | [illegible] | [illegible] | [illegible] | [illegible] | [illegible] | [illegible] | [illegible] | [illegible] | [illegible] | [illegible] | [illegible] | [illegible] | [illegible] | [illegible] | [illegible] | [illegible] | [illegible] | [illegible] | [illegible] | [illegible] | [illegible] | [illegible] | [illegible] | [illegible] | [illegible] |

| | | | | | |
|---|---|---|---|---|---|
| TG | TG 45:1\|TG 14:0_15:0_16:1 | [M+NH4]+ | 22.37 | 780.7077 | 0.02094053 |
| TG | TG 46:0\|TG 14:0_16:0_16:0 | [M+NH4]+ | 23.382 | 796.73932 | 0.106788635 |
| TG | TG 46:1\|TG 14:0_16:0_16:1 | [M+NH4]+ | 22.753 | 794.72369 | 0.148046022 |
| TG | TG 46:2\|TG 12:0_16:1_18:1 | [M+NH4]+ | 22.116 | 792.70697 | 0.066245696 |
| TG | TG 46:3\|TG 14:0_16:1_16:2 | [M+NH4]+ | 21.829 | 790.69189 | 0.020151172 |
| TG | TG 46:4\|TG 14:0_16:1_16:3 | [M+NH4]+ | 21.145 | 788.67566 | 0.005872335 |
| TG | TG 47:0\|TG 14:0_16:0_17:0 | [M+NH4]+ | 23.726 | 810.75391 | 0.052775692 |
| TG | TG 47:1\|TG 15:0_16:0_16:1 | [M+NH4]+ | 23.103 | 808.73761 | 0.084680575 |
| TG | TG 47:2\|TG 14:0_16:1_17:1 | [M+NH4]+ | 22.49 | 806.72272 | 0.048796777 |
| TG | TG 47:4\|TG 14:0_17:1_16:3 | [M+NH4]+ | 21.567 | 802.69098 | 0.003878666 |
| TG | TG 48:0\|TG 14:0_16:0_18:0 | [M+NH4]+ | 24.061 | 824.77008 | 0.267247162 |
| TG | TG 48:1\|TG 14:0_16:0_18:1 | [M+NH4]+ | 23.441 | 822.75488 | 0.662174988 |
| TG | TG 48:2\|TG 16:0_16:1_16:1 | [M+NH4]+ | 22.849 | 820.73792 | 0.449948339 |
| TG | TG 48:3\|TG 16:0_16:1_16:2 | [M+NH4]+ | 22.266 | 818.72198 | 0.167699231 |
| TG | TG 48:4\|TG 16:0_16:1_16:3 | [M+NH4]+ | 21.958 | 816.70721 | 0.068724813 |
| TG | TG 48:5\|TG 16:1_16:1_16:3 | [M+NH4]+ | 21.374 | 814.69055 | 0.017262427 |
| TG | TG 48:6\|TG 16:1_16:2_16:3 | [M+NH4]+ | 20.83 | 812.67511 | 0.005390388 |
| TG | TG 49:0\|TG 15:0_16:0_18:0 | [M+NH4]+ | 24.364 | 838.78357 | 0.077442142 |
| TG | TG 49:1\|TG 15:0_16:0_18:1 | [M+NH4]+ | 23.771 | 836.76904 | 0.295141971 |
| TG | TG 49:2\|TG 16:0_16:1_17:1 | [M+NH4]+ | 23.289 | 834.75342 | 1.098077532 |
| TG | TG 49:3\|TG 15:1_16:1_18:1 | [M+NH4]+ | 22.615 | 832.73975 | 0.085625603 |
| TG | TG 49:4\|TG 16:0_17:1_16:3 | [M+NH4]+ | 22.331 | 830.72095 | 0.032972353 |
| TG | TG 49:5\|TG 16:1_17:1_16:3 | [M+NH4]+ | 21.724 | 828.7066 | 0.009015033 |
| TG | TG 50:0\|TG 16:0_16:0_18:0 | [M+NH4]+ | 24.678 | 852.80072 | 0.385982783 |
| TG | TG 50:1\|TG 16:0_18:0_16:1 | [M+NH4]+ | 24.089 | 850.78528 | 1.911389292 |
| TG | TG 50:2\|TG 16:0_16:1_18:1 | [M+NH4]+ | 23.523 | 848.77008 | 1.865479302 |
| TG | TG 50:3\|TG 16:1_16:1_18:1 | [M+NH4]+ | 22.951 | 846.75269 | 0.899007281 |
| TG | TG 50:4\|TG 16:1_18:1_16:2 | [M+NH4]+ | 22.699 | 844.73901 | 0.412576051 |
| TG | TG 50:5\|TG 16:1_16:2_18:2 | [M+NH4]+ | 22.107 | 842.72278 | 0.167429433 |
| TG | TG 50:6\|TG 12:0_16:1_22:5 | [M+NH4]+ | 21.581 | 840.70715 | 0.0476992 |
| TG | TG 50:6\|TG 14:0_16:2_20:4 | [M+NH4]+ | 21.581 | 840.70715 | 0.048069756 |
| TG | TG 50:7\|TG 16:1_16:3_18:3 | [M+NH4]+ | 21.078 | 838.69226 | 0.010143099 |
| TG | TG 50:8\|TG 14:0_16:3_20:5 | [M+NH4]+ | 20.525 | 836.67487 | 0.001316263 |
| TG | TG 51:0\|TG 16:0_17:0_18:0 | [M+NH4]+ | 24.961 | 866.81439 | 0.067703648 |
| TG | TG 51:1\|TG 16:0_17:0_18:1 | [M+NH4]+ | 24.398 | 864.80054 | 0.460505809 |
| TG | TG 51:2\|TG 16:0_17:1_18:1 | [M+NH4]+ | 23.818 | 862.78351 | 0.691695505 |
| TG | TG 51:3\|TG 16:1_17:1_18:1 | [M+NH4]+ | 23.274 | 860.77026 | 0.347814048 |
| TG | TG 51:4\|TG 16:1_17:1_18:2 | [M+NH4]+ | 22.905 | 858.75372 | 0.140514408 |
| TG | TG 51:5\|TG 17:1_18:1_16:3 | [M+NH4]+ | 22.408 | 856.73749 | 0.06218209 |
| TG | TG 51:6\|TG 17:1_18:2_16:3 | [M+NH4]+ | 21.878 | 854.72174 | 0.021143374 |
| TG | TG 52:0\|TG 16:0_18:0_18:0 | [M+NH4]+ | 25.277 | 880.8299 | 0.251613124 |
| TG | TG 52:1\|TG 16:0_18:0_18:1 | [M+NH4]+ | 24.703 | 878.81628 | 2.077846295 |
| TG | TG 52:2\|TG 16:0_18:0_18:2 | [M+NH4]+ | 24.131 | 876.79889 | 4.156550925 |
| TG | TG 52:3\|TG 16:1_18:1_18:1 | [M+NH4]+ | 23.605 | 874.78442 | 2.542402327 |
| TG | TG 52:4\|TG 18:0_18:1_16:3 | [M+NH4]+ | 23.142 | 872.76727 | 1.497234104 |
| TG | TG 52:5\|TG 16:1_18:1_18:3 | [M+NH4]+ | 22.766 | 870.75287 | 0.854607077 |
| TG | TG 52:6\|TG 16:0_16:1_20:5 | [M+NH4]+ | 22.24 | 868.73907 | 0.385587923 |
| TG | TG 52:7\|TG 14:0_16:1_22:6 | [M+NH4]+ | 21.768 | 866.72192 | 0.117283396 |
| TG | TG 52:8\|TG 16:1_16:3_20:4 | [M+NH4]+ | 21.371 | 864.70837 | 0.026159237 |
| TG | TG 52:9\|TG 14:0_16:3_22:6 | [M+NH4]+ | 20.835 | 862.69165 | 0.005809885 |
| TG | TG 53:0\|TG 16:0_18:0_19:0 | [M+NH4]+ | 25.518 | 894.84985 | 0.021516003 |
| TG | TG 53:1\|TG 17:0_18:0_18:1 | [M+NH4]+ | 24.987 | 892.83063 | 0.22642713 |
| TG | TG 53:2\|TG 18:0_17:1_18:1 | [M+NH4]+ | 24.43 | 890.8147 | 0.636765693 |
| TG | TG 53:3\|TG 17:1_18:1_18:1 | [M+NH4]+ | 23.895 | 888.79846 | 0.579662399 |
| TG | TG 53:4\|TG 17:1_18:1_18:2 | [M+NH4]+ | 23.37 | 886.78601 | 0.364604242 |
| TG | TG 53:5\|TG 15:0_16:0_22:5 | [M+NH4]+ | 23.107 | 884.7688 | 0.248875712 |
| TG | TG 53:6\|TG 15:0_16:0_22:6 | [M+NH4]+ | 22.591 | 882.75232 | 0.127190237 |
| TG | TG 53:7\|TG 16:1_17:1_20:5 | [M+NH4]+ | 22.156 | 880.73846 | 0.044162496 |
| TG | TG 54:0\|TG 16:0_18:0_20:0 | [M+NH4]+ | 25.805 | 908.86163 | 0.047395668 |
| TG | TG 54:1\|TG 16:0_18:0_20:1 | [M+NH4]+ | 25.288 | 906.84619 | 0.567835771 |
| TG | TG 54:10\|TG 16:1_16:3_22:6 | [M+NH4]+ | 20.93 | 888.70599 | 0.011827944 |
| TG | TG 54:2\|TG 16:0_18:1_20:1 | [M+NH4]+ | 24.73 | 904.83075 | 2.079261163 |
| TG | TG 54:3\|TG 18:1_18:1_18:1 | [M+NH4]+ | 24.2 | 902.81372 | 2.883211972 |
| TG | TG 54:4\|TG 18:1_18:1_18:2 | [M+NH4]+ | 23.735 | 900.79993 | 2.712829473 |
| TG | TG 54:5\|TG 16:0_16:1_22:4 | [M+NH4]+ | 23.434 | 898.78577 | 2.636575977 |
| TG | TG 54:6\|TG 16:1_18:2_20:3 | [M+NH4]+ | 22.905 | 896.76794 | 1.701272658 |
| TG | TG 54:7\|TG 16:1_18:2_20:4 | [M+NH4]+ | 22.52 | 894.75293 | 0.79663472 |
| TG | TG 54:8\|TG 16:0_16:2_22:6 | [M+NH4]+ | 21.955 | 892.73755 | 0.221420366 |
| TG | TG 54:9\|TG 16:1_18:4_20:4 | [M+NH4]+ | 21.432 | 890.72192 | 0.039747346 |
| TG | TG 55:0\|TG 15:0_16:0_24:0 | [M+NH4]+ | 26.044 | 922.87775 | 0.008878992 |
| TG | TG 55:1\|TG 18:0_19:0_18:1 | [M+NH4]+ | 25.538 | 920.86426 | 0.042285022 |
| TG | TG 55:10\|TG 17:1_16:3_22:6 | [M+NH4]+ | 21.352 | 902.72241 | 0.002702817 |
| TG | TG 55:2\|TG 18:0_18:1_19:1 | [M+NH4]+ | 25 | 918.84613 | 0.155350928 |
| TG | TG 55:3\|TG 18:1_18:1_19:1 | [M+NH4]+ | 24.484 | 916.83087 | 0.269678381 |
| TG | TG 55:4\|TG 18:1_19:1_18:2 | [M+NH4]+ | 24.067 | 914.81519 | 0.35550713 |
| TG | TG 55:5\|TG 16:0_17:0_22:5 | [M+NH4]+ | 23.741 | 912.79803 | 0.410152641 |
| TG | TG 55:6\|TG 17:1_18:2_20:3 | [M+NH4]+ | 23.154 | 910.78436 | 0.38758182 |
| TG | TG 55:7\|TG 16:0_17:1_22:6 | [M+NH4]+ | 22.872 | 908.76978 | 0.208810227 |
| TG | TG 55:8\|TG 17:1_18:2_20:5 | [M+NH4]+ | 22.268 | 906.75458 | 0.063937116 |
| TG | TG 55:9\|TG 15:0_20:4_20:5 | [M+NH4]+ | 21.813 | 904.73877 | 0.013823375 |
| TG | TG 56:0\|TG 16:0_18:0_22:0 | [M+NH4]+ | 26.319 | 936.89331 | 0.023521787 |
| TG | TG 56:1\|TG 16:0_18:0_22:1 | [M+NH4]+ | 25.812 | 934.87982 | 0.125921219 |
| TG | TG 56:10\|TG 16:1_20:4_20:5 | [M+NH4]+ | 21.72 | 916.73773 | 0.103690639 |
| TG | TG 56:11\|TG 18:2_16:3_22:6 | [M+NH4]+ | 21.173 | 914.72205 | 0.018985347 |
| TG | TG 56:2\|TG 18:0_18:1_20:1 | [M+NH4]+ | 25.3 | 932.86163 | 0.322261661 |
| TG | TG 56:3\|TG 18:1_18:1_20:1 | [M+NH4]+ | 24.792 | 930.84607 | 0.632270166 |
| TG | TG 56:4\|TG 18:1_20:1_18:2 | [M+NH4]+ | 24.4 | 928.83258 | 1.390657694 |
| TG | TG 56:5\|TG 16:0_18:1_22:4 | [M+NH4]+ | 23.92 | 926.815 | 1.512691392 |
| TG | TG 56:6\|TG 16:0_18:0_22:6 | [M+NH4]+ | 23.47 | 924.79968 | 2.53098994 |
| TG | TG 56:6\|TG 18:1_18:2_20:3 | [M+NH4]+ | 23.469 | 924.79993 | 2.957645661 |
| TG | TG 56:7\|TG 16:0_18:1_22:6 | [M+NH4]+ | 23.209 | 922.78418 | 1.37567474 |
| TG | TG 56:7\|TG 16:1_18:1_22:5 | [M+NH4]+ | 22.96 | 922.78369 | 0.975487734 |
| TG | TG 56:8\|TG 16:1_18:2_22:5 | [M+NH4]+ | 22.646 | 920.77063 | 1.078847206 |
| TG | TG 56:9\|TG 16:0_20:4_20:5 | [M+NH4]+ | 22.204 | 918.75269 | 0.366005828 |
| TG | TG 57:0\|TG 16:0_17:0_24:0 | [M+NH4]+ | 26.522 | 950.91174 | 0.00855274 |
| TG | TG 57:1\|TG 16:0_17:0_24:1 | [M+NH4]+ | 26.06 | 948.89215 | 0.023806967 |
| TG | TG 57:10\|TG 15:0_20:4_22:6 | [M+NH4]+ | 22.093 | 930.75415 | 0.008606499 |
| TG | TG 57:10\|TG 17:1_20:4_20:5 | [M+NH4]+ | 21.911 | 930.75415 | 0.014017678 |
| TG | TG 57:2\|TG 16:0_18:1_23:1 | [M+NH4]+ | 25.543 | 946.87897 | 0.047832741 |
| TG | TG 57:3\|TG 18:1_19:1_20:1 | [M+NH4]+ | 25.038 | 944.86444 | 0.058309932 |
| TG | TG 57:4\|TG 18:1_19:1_20:2 | [M+NH4]+ | 24.658 | 942.84546 | 0.118089458 |
| TG | TG 57:5\|TG 17:0_18:0_22:5 | [M+NH4]+ | 24.22 | 940.83142 | 0.237412621 |
| TG | TG 57:6\|TG 17:1_18:1_22:4 | [M+NH4]+ | 23.765 | 938.81677 | 0.318160064 |
| TG | TG 57:7\|TG 17:0_18:1_22:6 | [M+NH4]+ | 23.539 | 936.80096 | 0.113733681 |
| TG | TG 57:7\|TG 17:1_18:1_22:5 | [M+NH4]+ | 23.226 | 936.80145 | 0.228510473 |
| TG | TG 57:8\|TG 17:1_18:1_22:6 | [M+NH4]+ | 22.934 | 934.784 | 0.181957771 |
| TG | TG 57:9\|TG 17:0_20:4_20:5 | [M+NH4]+ | 22.4 | 932.76971 | 0.066449632 |
| TG | TG 58:0\|TG 16:0_18:0_24:0 | [M+NH4]+ | 26.777 | 964.92529 | 0.023773181 |
| TG | TG 58:1\|TG 16:0_18:0_24:1 | [M+NH4]+ | 26.315 | 962.90906 | 0.11614169 |
| TG | TG 58:10\|TG 16:0_20:4_22:6 | [M+NH4]+ | 22.489 | 944.76703 | 0.226603556 |
| TG | TG 58:10\|TG 16:1_20:4_22:5 | [M+NH4]+ | 22.251 | 944.76703 | 0.344038545 |
| TG | TG 58:11\|TG 16:1_20:4_22:6 | [M+NH4]+ | 21.953 | 942.7514 | 0.183490551 |
| TG | TG 58:12\|TG 16:1_20:5_22:6 | [M+NH4]+ | 21.355 | 940.7384 | 0.036600506 |
| TG | TG 58:13\|TG 16:3_20:4_22:6 | [M+NH4]+ | 20.953 | 938.72412 | 0.002590606 |
| TG | TG 58:2\|TG 16:0_18:1_24:1 | [M+NH4]+ | 25.804 | 960.89514 | 0.216298459 |
| TG | TG 58:3\|TG 16:1_20:1_22:1 | [M+NH4]+ | 25.345 | 958.87952 | 0.208814185 |
| TG | TG 58:4\|TG 16:0_20:2_22:2 | [M+NH4]+ | 24.945 | 956.8623 | 0.349561563 |
| TG | TG 58:5\|TG 16:0_18:1_24:4 | [M+NH4]+ | 24.532 | 954.84753 | 0.926515515 |
| TG | TG 58:6\|TG 18:1_18:1_22:4 | [M+NH4]+ | 24.049 | 952.83124 | 1.712783164 |
| TG | TG 58:7\|TG 18:0_18:1_22:6 | [M+NH4]+ | 23.87 | 950.81335 | 0.563695475 |
| TG | TG 58:7\|TG 18:1_18:2_22:4 | [M+NH4]+ | 23.551 | 950.81335 | 1.241396279 |
| TG | TG 58:8\|TG 18:1_18:2_22:5 | [M+NH4]+ | 23.254 | 948.79822 | 1.75186057 |
| TG | TG 58:9\|TG 18:2_20:3_20:4 | [M+NH4]+ | 22.759 | 946.7851 | 1.118931684 |
| TG | TG 59:0\|TG 16:0_17:0_26:0 | [M+NH4]+ | 26.968 | 978.94257 | 0.004531521 |
| TG | TG 59:1\|TG 16:0_18:0_25:1 | [M+NH4]+ | 26.539 | 976.92731 | 0.023443316 |
| TG | TG 59:10\|TG 17:1_20:4_22:5 | [M+NH4]+ | 22.508 | 958.78387 | 0.068860215 |
| TG | TG 59:11\|TG 17:1_20:4_22:6 | [M+NH4]+ | 22.182 | 956.76703 | 0.03208643 |
| TG | TG 59:12\|TG 17:1_20:5_22:6 | [M+NH4]+ | 21.676 | 954.7525 | 0.005306605 |
| TG | TG 59:2\|TG 16:0_18:1_25:1 | [M+NH4]+ | 26.043 | 974.9101 | 0.04778579 |
| TG | TG 59:3\|TG 17:1_18:1_24:1 | [M+NH4]+ | 25.579 | 972.89508 | 0.043190643 |
| TG | TG 59:5\|TG 17:0_18:1_24:4 | [M+NH4]+ | 24.732 | 968.86426 | 0.076034294 |
| TG | TG 59:6\|TG 18:1_19:1_22:4 | [M+NH4]+ | 24.301 | 966.84698 | 0.131103597 |
| TG | TG 59:7\|TG 18:1_19:1_22:5 | [M+NH4]+ | 23.821 | 964.83276 | 0.168238102 |
| TG | TG 59:8\|TG 18:1_19:1_22:6 | [M+NH4]+ | 23.358 | 962.81708 | 0.074564824 |
| TG | TG 59:9\|TG 17:0_20:4_22:5 | [M+NH4]+ | 23.031 | 960.79919 | 0.096527072 |
| TG | TG 60:0\|TG 16:0_18:0_26:0 | [M+NH4]+ | 27.201 | 992.95667 | 0.013860727 |
| TG | TG 60:1\|TG 16:0_18:0_26:1 | [M+NH4]+ | 26.772 | 990.93988 | 0.106483881 |
| TG | TG 60:10\|TG 16:0_22:4_22:6 | [M+NH4]+ | 22.825 | 972.80145 | 0.888187941 |
| TG | TG 60:11\|TG 16:0_22:5_22:6 | [M+NH4]+ | 22.53 | 970.78479 | 0.602382563 |
| TG | TG 60:11\|TG 16:1_22:5_22:5 | [M+NH4]+ | 22.53 | 970.78479 | 0.604945237 |
| TG | TG 60:12\|TG 16:0_22:6_22:6 | [M+NH4]+ | 22.196 | 968.77045 | 0.265693875 |
| TG | TG 60:13\|TG 18:2_20:5_22:6 | [M+NH4]+ | 21.574 | 966.75421 | 0.053416539 |

| | | | | | | | | | | | | | | | | | | | | | | | | | | | | | | | | | | | |
|---|---|---|---|---|---|---|---|---|---|---|---|---|---|---|---|---|---|---|---|---|---|---|---|---|---|---|---|---|---|---|---|---|---|---|---|
| TG 45:1\|TG 14:0_15:0_16:1 | 0.027323017 | 0.029028307 | 0.041858071 | 0.05768093 | 0.04385105 | 5.64847E-05 | 6.98331E-05 | 5.59858E-05 | 4.66642E-05 | 6.12022E-05 | 4.97289E-05 | 6.07679E-05 | 6.41331E-06 | 5.25318E-05 | 6.25731E-06 | 0.263510669 | -0.210119064 | 0.003293712 | 0.004868977 | 0.002482376 | 0.012615576 | 0.012311593 | 0.010401252 | 1.54036E-05 | 1.76312E-05 | 1.98647E-05 | 3.52136E-05 | 3.41228E-05 | 3.23345E-05 | 1.76332E-05 | 1.82121E-06 | 3.38903E-05 | 1.18681E-06 | 0.000963148 | 0.942581414 |
| TG 46:0\|TG 14:0_16:0_16:0 | 0.12894319 | 0.135786027 | 0.346489584 | 0.351456825 | 0.315704852 | 0.00028805 | 0.000329557 | 0.000261886 | 0.000386273 | 0.000372913 | 0.000358022 | 0.000293164 | 2.78626E-05 | 0.000372403 | 1.15391E-05 | 0.041318781 | 0.345153323 | 0.000890901 | 0.001356642 | 0.000649683 | 0.004848434 | 0.004881996 | 0.004258074 | 4.16646E-06 | 4.91257E-06 | 5.19895E-06 | 1.35333E-05 | 1.35309E-05 | 1.32371E-05 | 4.75932E-06 | 4.35216E-07 | 1.34338E-05 | 1.39063E-07 | 0.000488187 | 1.497038321 |
| TG 46:1\|TG 14:0_16:0_16:1 | 0.162193346 | 0.212059098 | 0.567262748 | 0.661672786 | 0.553238333 | 0.000399337 | 0.000414539 | 0.000408991 | 0.000632396 | 0.000702067 | 0.000627395 | 0.000407622 | 6.28107E-06 | 0.000653952 | 3.40832E-05 | 0.007797033 | 0.68195177 | 0.012611704 | 0.015629607 | 0.009768652 | 0.025223418 | 0.028741384 | 0.025067321 | 5.89808E-05 | 5.65967E-05 | 7.81715E-05 | 7.04056E-05 | 7.96595E-05 | 7.79271E-05 | 6.4583E-05 | 9.65766E-06 | 7.59974E-05 | 4.01678E-06 | 0.231241186 | 0.234794961 |
| TG 46:2\|TG 12:0_16:1_18:1 | 0.071111249 | 0.099459992 | 0.18843423 | 0.217713085 | 0.18717508 | 0.00017869 | 0.000181749 | 0.000191825 | 0.00021007 | 0.000231004 | 0.000212264 | 0.000184088 | 5.6115E-06 | 0.000217779 | 9.394E-06 | 0.018754678 | 0.242472867 | 0.007438907 | 0.010715147 | 0.005650636 | 0.027449183 | 0.026280577 | 0.02248875 | 3.47894E-05 | 3.88009E-05 | 4.5218E-05 | 7.66183E-05 | 7.28392E-05 | 6.99111E-05 | 3.96027E-05 | 4.29504E-06 | 7.31229E-05 | 2.74555E-06 | 0.001555553 | 0.88472206 |
| TG 46:3\|TG 14:0_16:1_16:2 | 0.021063381 | 0.031473583 | 0.05934845 | 0.065849371 | 0.056361124 | 5.43555E-05 | 5.38345E-05 | 6.07019E-05 | 6.61628E-05 | 6.98693E-05 | 6.39158E-05 | 5.62973E-05 | 3.12179E-06 | 6.66493E-05 | 2.45475E-06 | 0.023180746 | 0.243523681 | 0.038639654 | 0.051358278 | 0.026149753 | 0.105724239 | 0.099314037 | 0.082953077 | 0.000180705 | 0.000185975 | 0.000209258 | 0.000295106 | 0.000275258 | 0.000257877 | 0.000191979 | 1.24056E-05 | 0.000276081 | 1.52095E-05 | 0.004251193 | 0.52413924 |
| TG 46:4\|TG 14:0_16:1_16:3 | 0.005743954 | 0.010190629 | 0.018879162 | 0.022087695 | 0.01785453 | 1.584E-05 | 1.46806E-05 | 1.96543E-05 | 2.10469E-05 | 2.34361E-05 | 2.02478E-05 | 1.67249E-05 | 2.12475E-06 | 2.15769E-05 | 1.35452E-06 | 0.063158865 | 0.367486216 | 0.074062431 | 0.113002006 | 0.048489467 | 0.292200605 | 0.307830712 | 0.254606499 | 0.000346366 | 0.000409194 | 0.000388026 | 0.000815613 | 0.000853183 | 0.000791499 | 0.000381196 | 2.61001E-05 | 0.000820098 | 2.53812E-05 | 6.97941E-05 | 1.105265276 |
| TG 47:0\|TG 14:0_16:0_17:0 | 0.066896885 | 0.063694394 | 0.194745508 | 0.212096363 | 0.178520067 | 0.000142356 | 0.000170977 | 0.000122845 | 0.000217106 | 0.000225044 | 0.000202449 | 0.000145393 | 1.97669E-05 | 0.000214866 | 9.35949E-06 | 0.02287387 | 0.563483121 | 0.029880288 | 0.04553899 | 0.020885512 | 0.125233193 | 0.125068712 | 0.108187383 | 0.000139741 | 0.000164902 | 0.000167132 | 0.000349561 | 0.00034664 | 0.000336324 | 0.000157258 | 1.24202E-05 | 0.000344175 | 5.6782E-06 | 0.000455727 | 1.13000636 |
| TG 47:1\|TG 15:0_16:0_16:1 | 0.099285273 | 0.128477316 | 0.342361995 | 0.416476446 | 0.342890528 | 0.000228416 | 0.000253757 | 0.000247789 | 0.000381672 | 0.000441902 | 0.000388852 | 0.000243321 | 1.0817E-05 | 0.000404142 | 2.68606E-05 | 0.006757757 | 0.732001293 | 0.00988982 | 0.015271004 | 0.006833842 | 0.050177813 | 0.051851731 | 0.045109602 | 4.62515E-05 | 5.52982E-05 | 5.46863E-05 | 0.00014006 | 0.000143712 | 0.000140233 | 5.20787E-05 | 4.12797E-06 | 0.000141335 | 1.68228E-06 | 0.000231948 | 1.440354755 |
| TG 47:2\|TG 14:0_16:1_17:1 | 0.055331239 | 0.070162917 | 0.135564806 | 0.172413516 | 0.142117867 | 0.000131624 | 0.000141417 | 0.000135321 | 0.00015113 | 0.000182939 | 0.000161167 | 0.000136121 | 4.03806E-06 | 0.000165079 | 1.32771E-05 | 0.080259323 | 0.278269881 | 0.003445326 | 0.005524349 | 0.002412861 | 0.020964694 | 0.021689422 | 0.018560634 | 1.61127E-05 | 2.00043E-05 | 1.93084E-05 | 5.85183E-05 | 6.01143E-05 | 5.76997E-05 | 1.84751E-05 | 1.69449E-06 | 5.87774E-05 | 1.00265E-06 | 4.95408E-05 | 1.66967724 |
| TG 47:4\|TG 14:0_17:1_16:3 | 0.004078516 | 0.006627708 | 0.013125265 | 0.01456404 | 0.01246112 | 1.04623E-05 | 1.0424E-05 | 1.27826E-05 | 1.46323E-05 | 1.54531E-05 | 1.41314E-05 | 1.1223E-05 | 1.10295E-06 | 1.4739E-05 | 5.44838E-07 | 0.028648926 | 0.393180326 | 0.017604699 | 0.024556594 | 0.012903471 | 0.049623739 | 0.049067265 | 0.042829427 | 8.23316E-05 | 8.89224E-05 | 0.000103257 | 0.000138514 | 0.000135995 | 0.000133144 | 9.15037E-05 | 8.73567E-06 | 0.000135884 | 2.19337E-06 | 0.014470677 | 0.570475942 |
| TG 48:0\|TG 14:0_16:0_18:0 | 0.317059109 | 0.332523444 | 1.205322174 | 1.253587885 | 1.071719728 | 0.000720869 | [illegible] | [illegible] | [illegible] | [illegible] | [illegible] | [illegible] | [illegible] | [illegible] | [illegible] | [illegible] | [illegible] | [illegible] | [illegible] | [illegible] | [illegible] | [illegible] | [illegible] | [illegible] | [illegible] | [illegible] | [illegible] | [illegible] | [illegible] | [illegible] | [illegible] | [illegible] | [illegible] | [illegible] | [illegible] |
| TG 48:1\|TG 14:0_16:0_18:1 | 0.738089526 | 0.959369123 | 2.882194379 | 3.013013956 | 2.622383853 | 0.001786142 | [illegible] | [illegible] | [illegible] | [illegible] | [illegible] | [illegible] | [illegible] | [illegible] | [illegible] | [illegible] | [illegible] | [illegible] | [illegible] | [illegible] | [illegible] | [illegible] | [illegible] | [illegible] | [illegible] | [illegible] | [illegible] | [illegible] | [illegible] | [illegible] | [illegible] | [illegible] | [illegible] | [illegible] | [illegible] |
| TG 48:2\|TG 16:0_16:1_16:1 | 0.488346383 | 0.687167612 | 1.890743519 | 2.232768138 | 1.884116572 | 0.001213685 | [illegible] | [illegible] | [illegible] | [illegible] | [illegible] | [illegible] | [illegible] | [illegible] | [illegible] | [illegible] | [illegible] | [illegible] | [illegible] | [illegible] | [illegible] | [illegible] | [illegible] | [illegible] | [illegible] | [illegible] | [illegible] | [illegible] | [illegible] | [illegible] | [illegible] | [illegible] | [illegible] | [illegible] | [illegible] |
| TG 48:3\|TG 16:0_16:1_16:2 | 0.176740302 | 0.263011331 | 0.595458458 | 0.689610666 | 0.586321595 | 0.00045235 | [illegible] | [illegible] | [illegible] | [illegible] | [illegible] | [illegible] | [illegible] | [illegible] | [illegible] | [illegible] | [illegible] | [illegible] | [illegible] | [illegible] | [illegible] | [illegible] | [illegible] | [illegible] | [illegible] | [illegible] | [illegible] | [illegible] | [illegible] | [illegible] | [illegible] | [illegible] | [illegible] | [illegible] | [illegible] |
| TG 48:4\|TG 16:0_16:1_16:3 | 0.068905506 | 0.110303697 | 0.251017033 | 0.289665782 | 0.238348709 | 0.000185377 | [illegible] | [illegible] | [illegible] | [illegible] | [illegible] | [illegible] | [illegible] | [illegible] | [illegible] | [illegible] | [illegible] | [illegible] | [illegible] | [illegible] | [illegible] | [illegible] | [illegible] | [illegible] | [illegible] | [illegible] | [illegible] | [illegible] | [illegible] | [illegible] | [illegible] | [illegible] | [illegible] | [illegible] | [illegible] |
| TG 48:5\|TG 16:1_16:1_16:3 | 0.017031716 | 0.030674362 | 0.06896607 | 0.076838869 | 0.062396972 | 4.65635E-05 | [illegible] | [illegible] | [illegible] | [illegible] | [illegible] | [illegible] | [illegible] | [illegible] | [illegible] | [illegible] | [illegible] | [illegible] | [illegible] | [illegible] | [illegible] | [illegible] | [illegible] | [illegible] | [illegible] | [illegible] | [illegible] | [illegible] | [illegible] | [illegible] | [illegible] | [illegible] | [illegible] | [illegible] | [illegible] |

[illegible]

| | | | | | | | | |
|---|---|---|---|---|---|---|---|---|
| TG | TG 60:15\|TG 16:3_22:6_22:6 | [M+NH4]+ | 20.686 | 962.72461 | 0.000748865 | 0.000474008 | 0.00137703 | [illegible] |
| TG | TG 60:2\|TG 16:0_18:1_26:1 | [M+NH4]+ | 26.286 | 988.92505 | 0.212175274 | 0.245922943 | 0.299895323 | [illegible] |
| TG | TG 60:3\|TG 24:0_18:1_18:2 | [M+NH4]+ | 25.833 | 986.90808 | 0.206749532 | 0.230669329 | 0.299320636 | [illegible] |
| TG | TG 60:4\|TG 18:1_24:1_18:2 | [M+NH4]+ | 25.419 | 984.89307 | 0.193717277 | 0.212689313 | 0.274745072 | [illegible] |
| TG | TG 60:5\|TG 16:0_18:1_26:4 | [M+NH4]+ | 24.99 | 982.87598 | 0.272901635 | 0.30040646 | 0.394101109 | [illegible] |
| TG | TG 60:6\|TG 18:1_18:1_24:4 | [M+NH4]+ | 24.571 | 980.86096 | 0.564412124 | 0.592069728 | 0.823279893 | [illegible] |
| TG | TG 60:7\|TG 18:1_18:2_24:4 | [M+NH4]+ | 24.136 | 978.84851 | 0.791797331 | 0.798653109 | 1.136335213 | [illegible] |
| TG | TG 60:8\|TG 18:1_20:3_22:4 | [M+NH4]+ | 23.699 | 976.83307 | 0.864521013 | 0.863692753 | 1.295304184 | [illegible] |
| TG | TG 60:9\|TG 18:1_20:3_22:5 | [M+NH4]+ | 23.283 | 974.81628 | 0.710257117 | 0.792417423 | 1.311625122 | [illegible] |
| TG | TG 61:1\|TG 17:0_26:0_18:1 | [M+NH4]+ | 26.971 | 1004.95673 | 0.011295708 | 0.014177196 | 0.015633721 | [illegible] |
| TG | TG 61:10\|TG 17:1_22:4_22:5 | [M+NH4]+ | 23.163 | 986.81647 | 0.052189215 | 0.048954244 | 0.082035493 | [illegible] |
| TG | TG 61:12\|TG 17:1_22:5_22:6 | [M+NH4]+ | 22.226 | 982.78503 | 0.020617732 | 0.017815066 | 0.034018484 | [illegible] |
| TG | TG 61:13\|TG 17:1_22:6_22:6 | [M+NH4]+ | 21.95 | 980.77026 | 0.008017147 | 0.006905689 | 0.014594588 | [illegible] |
| TG | TG 61:2\|TG 16:0_18:1_27:1 | [M+NH4]+ | 26.513 | 1002.94171 | 0.032798093 | 0.038542193 | 0.047201954 | [illegible] |
| TG | TG 61:3\|TG 17:1_18:1_26:1 | [M+NH4]+ | 26.057 | 1000.92419 | 0.036487898 | 0.041506377 | 0.054738071 | [illegible] |
| TG | TG 61:4\|TG 18:1_25:1_18:2 | [M+NH4]+ | 25.646 | 998.90851 | 0.030661659 | 0.03116635 | 0.044388749 | [illegible] |
| TG | TG 61:5\|TG 18:0_17:1_26:4 | [M+NH4]+ | 25.268 | 996.89496 | 0.02304072 | 0.030901314 | 0.041745664 | [illegible] |
| TG | TG 61:6\|TG 17:0_18:1_26:5 | [M+NH4]+ | 24.831 | 994.87866 | 0.043547126 | 0.044572219 | 0.063962852 | [illegible] |
| TG | TG 61:7\|TG 17:0_18:1_26:6 | [M+NH4]+ | 24.393 | 992.86432 | 0.061315719 | 0.062325948 | 0.091820373 | [illegible] |
| TG | TG 62:0\|TG 16:0_18:0_28:0 | [M+NH4]+ | 27.593 | 1020.9873 | 0.002915939 | 0.003523859 | 0.003256553 | [illegible] |
| TG | TG 62:1\|TG 18:0_26:0_18:1 | [M+NH4]+ | 27.196 | 1018.97186 | 0.030482075 | 0.036578765 | 0.038710684 | [illegible] |
| TG | TG 62:10\|TG 18:1_22:4_22:5 | [M+NH4]+ | 23.341 | 1000.83124 | 0.446008913 | 0.425270197 | 0.650061482 | [illegible] |
| TG | TG 62:11\|TG 18:1_22:4_22:6 | [M+NH4]+ | 23.053 | 998.81531 | 0.502241456 | 0.457287693 | 0.766977913 | [illegible] |
| TG | TG 62:12\|TG 18:1_22:5_22:6 | [M+NH4]+ | 22.567 | 996.79755 | 0.335436597 | 0.300082099 | 0.547585311 | [illegible] |
| TG | TG 62:13\|TG 20:3_20:4_22:6 | [M+NH4]+ | 22.309 | 994.78589 | 0.164289233 | 0.140422624 | 0.281419277 | [illegible] |
| TG | TG 62:14\|TG 20:4_20:4_22:6 | [M+NH4]+ | 21.747 | 992.7677 | 0.047656315 | 0.038571334 | 0.086324655 | [illegible] |
| TG | TG 62:15\|TG 20:4_20:5_22:6 | [M+NH4]+ | 21.243 | 990.75574 | 0.008599811 | 0.006710895 | 0.015976104 | [illegible] |
| TG | TG 62:16\|TG 20:5_20:5_22:6 | [M+NH4]+ | 20.682 | 988.73718 | 0.000623171 | 0.000481635 | 0.001352034 | [illegible] |
| TG | TG 62:2\|TG 18:0_18:1_26:1 | [M+NH4]+ | 26.758 | 1016.95807 | 0.106458886 | 0.123714555 | 0.143547595 | [illegible] |
| TG | TG 62:3\|TG 26:0_18:1_18:2 | [M+NH4]+ | 26.288 | 1014.94153 | 0.141695494 | 0.160242996 | 0.203199745 | [illegible] |
| TG | TG 62:4\|TG 18:1_18:1_26:2 | [M+NH4]+ | 25.894 | 1012.92664 | 0.142851617 | 0.153396253 | 0.204516686 | [illegible] |
| TG | TG 62:5\|TG 18:1_18:2_26:2 | [M+NH4]+ | 25.492 | 1010.90912 | 0.170368579 | 0.179153945 | 0.240861469 | [illegible] |
| TG | TG 62:6\|TG 18:1_18:2_26:3 | [M+NH4]+ | 25.044 | 1008.89331 | 0.10313933 | 0.115781768 | 0.162294623 | [illegible] |
| TG | TG 62:7\|TG 18:1_18:2_26:4 | [M+NH4]+ | 24.644 | 1006.87958 | 0.285122755 | 0.289157154 | 0.416694896 | [illegible] |
| TG | TG 62:8\|TG 18:1_18:2_26:5 | [M+NH4]+ | 24.24 | 1004.86316 | 0.328968611 | 0.339666032 | 0.495127422 | [illegible] |
| TG | TG 62:9\|TG 18:1_20:3_24:5 | [M+NH4]+ | 23.822 | 1002.84802 | 0.422902325 | 0.42992923 | 0.612108794 | [illegible] |
| TG | TG 63:1\|TG 18:0_27:0_18:1 | [M+NH4]+ | 27.373 | 1032.98804 | 0.002020132 | 0.002768543 | 0.00262282 | [illegible] |
| TG | TG 63:2\|TG 26:0_18:1_19:1 | [M+NH4]+ | 26.947 | 1030.97253 | 0.008875804 | 0.01048699 | 0.012378644 | [illegible] |
| TG | TG 63:3\|TG 18:1_19:1_26:1 | [M+NH4]+ | 26.51 | 1028.9585 | 0.014941479 | 0.01673156 | 0.021545298 | [illegible] |
| TG | TG 63:4\|TG 18:1_27:1_18:2 | [M+NH4]+ | 26.132 | 1026.94153 | 0.015854195 | 0.017655897 | 0.02304278 | [illegible] |
| TG | TG 63:5\|TG 18:1_25:1_20:3 | [M+NH4]+ | 25.751 | 1024.92603 | 0.020377874 | 0.021301793 | 0.030201477 | [illegible] |
| TG | TG 63:6\|TG 16:0_25:1_22:5 | [M+NH4]+ | 25.446 | 1022.91211 | 0.020254447 | 0.02157688 | 0.030771535 | [illegible] |
| TG | TG 64:1\|TG 16:0_24:0_24:1 | [M+NH4]+ | 27.588 | 1047.00439 | 0.004603132 | 0.006011557 | 0.005992574 | [illegible] |
| TG | TG 64:10\|TG 18:1_22:4_24:5 | [M+NH4]+ | 23.93 | 1028.86096 | 0.20557354 | 0.205731429 | 0.285698048 | [illegible] |
| TG | TG 64:11\|TG 18:1_20:4_26:6 | [M+NH4]+ | 23.542 | 1026.84875 | 0.16874803 | 0.192500135 | 0.258837617 | [illegible] |
| TG | TG 64:12\|TG 18:1_22:5_24:6 | [M+NH4]+ | 23.14 | 1024.83374 | 0.157233077 | 0.14541267 | 0.233867586 | [illegible] |
| TG | TG 64:13\|TG 20:4_22:4_22:5 | [M+NH4]+ | 22.704 | 1022.81549 | 0.10579199 | 0.095228535 | 0.165464897 | [illegible] |
| TG | TG 64:14\|TG 20:4_22:4_22:6 | [M+NH4]+ | 22.184 | 1020.79919 | 0.055428563 | 0.047702277 | 0.091245147 | [illegible] |
| TG | TG 64:15\|TG 20:4_22:5_22:6 | [M+NH4]+ | 21.833 | 1018.78412 | 0.026022286 | 0.020947254 | 0.042311503 | [illegible] |
| TG | TG 64:16\|TG 20:4_22:6_22:6 | [M+NH4]+ | 21.542 | 1016.77094 | 0.010783359 | 0.008243314 | 0.019067881 | [illegible] |
| TG | TG 64:17\|TG 20:5_22:6_22:6 | [M+NH4]+ | 20.998 | 1014.75494 | 0.001344421 | 0.001087849 | 0.002674608 | [illegible] |
| TG | TG 64:2\|TG 18:0_18:1_28:1 | [M+NH4]+ | 27.163 | 1044.98853 | 0.01738075 | 0.020426165 | 0.022969133 | [illegible] |
| TG | TG 64:3\|TG 18:1_20:1_26:1 | [M+NH4]+ | 26.734 | 1042.96948 | 0.035207227 | 0.039243023 | 0.049138998 | [illegible] |
| TG | TG 64:4\|TG 18:1_26:1_20:2 | [M+NH4]+ | 26.374 | 1040.9563 | 0.063641534 | 0.069456107 | 0.09022046 | [illegible] |
| TG | TG 64:5\|TG 18:1_26:1_20:3 | [M+NH4]+ | 26.035 | 1038.93982 | 0.096352539 | 0.104514801 | 0.145189463 | [illegible] |
| TG | TG 64:6\|TG 18:1_26:1_20:4 | [M+NH4]+ | 25.708 | 1036.9248 | 0.12773736 | 0.133552482 | 0.18534024 | [illegible] |
| TG | TG 64:7\|TG 16:0_26:1_22:6 | [M+NH4]+ | 25.281 | 1034.91113 | 0.104955005 | 0.108535612 | 0.156790387 | [illegible] |
| TG | TG 64:8\|TG 18:1_20:3_26:4 | [M+NH4]+ | 24.776 | 1032.8949 | 0.109800463 | 0.135792958 | 0.189641824 | [illegible] |
| TG | TG 64:9\|TG 18:1_22:4_24:4 | [M+NH4]+ | 24.364 | 1030.87927 | 0.164071394 | 0.1660407 | 0.235290085 | [illegible] |
| TG | TG 65:3\|TG 18:1_20:1_27:1 | [M+NH4]+ | 26.934 | 1056.98718 | 0.003485723 | 0.00391412 | 0.004817206 | [illegible] |
| TG | TG 66:1\|TG 16:0_24:0_26:1 | [M+NH4]+ | 27.967 | 1075.03564 | 0.001615074 | 0.001982094 | 0.002073884 | [illegible] |
| TG | TG 66:10\|TG 18:1_22:4_26:5 | [M+NH4]+ | 24.43 | 1056.8949 | 0.081996447 | 0.082755737 | 0.116438455 | [illegible] |
| TG | TG 66:11\|TG 18:1_22:5_26:5 | [M+NH4]+ | 23.987 | 1054.87805 | 0.091851594 | 0.093553188 | 0.128994965 | [illegible] |
| TG | TG 66:15\|TG 22:4_22:5_22:6 | [M+NH4]+ | 22.376 | 1046.8136 | 0.021492253 | 0.019115097 | 0.032530269 | [illegible] |
| TG | TG 66:16\|TG 22:5_22:5_22:6 | [M+NH4]+ | 21.861 | 1044.79822 | 0.006047212 | 0.004567138 | 0.009177541 | [illegible] |
| TG | TG 66:17\|TG 22:5_22:6_22:6 | [M+NH4]+ | 21.577 | 1042.78418 | 0.004306181 | 0.003371031 | 0.007230305 | [illegible] |
| TG | TG 66:2\|TG 24:0_18:1_24:1 | [M+NH4]+ | 27.541 | 1073.01855 | 0.004714559 | 0.00535817 | 0.006327115 | [illegible] |
| TG | TG 66:3\|TG 18:1_22:1_26:1 | [M+NH4]+ | 27.14 | 1071.00244 | 0.010003577 | 0.011567134 | 0.013960211 | [illegible] |
| TG | TG 66:4\|TG 18:1_26:1_22:2 | [M+NH4]+ | 26.796 | 1068.98718 | 0.018250858 | 0.021045021 | 0.026383324 | [illegible] |
| TG | TG 66:5\|TG 26:0_18:1_22:4 | [M+NH4]+ | 26.593 | 1066.97302 | 0.039776995 | 0.0430549 | 0.056623787 | [illegible] |
| TG | TG 66:7\|TG 18:1_26:1_22:5 | [M+NH4]+ | 25.722 | 1062.9386 | 0.081523427 | 0.083758049 | 0.117780465 | [illegible] |
| TG | TG 66:8\|TG 18:1_26:1_22:6 | [M+NH4]+ | 25.328 | 1060.92761 | 0.063918573 | 0.06929486 | 0.093055429 | [illegible] |
| TG | TG 66:9\|TG 18:1_22:4_26:4 | [M+NH4]+ | 24.863 | 1058.90845 | 0.058417651 | 0.061023301 | 0.086453808 | [illegible] |
| TG | TG 68:2\|TG 24:0_18:1_26:1 | [M+NH4]+ | 27.92 | 1101.05322 | 0.002280015 | 0.002998499 | 0.003084003 | [illegible] |
| TG | TG 68:3\|TG 18:1_24:1_26:1 | [M+NH4]+ | 27.512 | 1099.03577 | 0.003589833 | 0.004038023 | 0.004808888 | [illegible] |
| TG | TG 68:4\|TG 24:1_26:1_18:2 | [M+NH4]+ | 27.159 | 1097.01904 | 0.006032522 | 0.006897623 | 0.008417306 | [illegible] |
| TG | TG 68:5\|TG 18:1_26:1_24:3 | [M+NH4]+ | 26.881 | 1095.00305 | 0.012523138 | 0.014082617 | 0.017561835 | [illegible] |
| TG | TG 68:6\|TG 18:1_26:1_24:4 | [M+NH4]+ | 26.533 | 1092.98767 | 0.022354873 | 0.024903863 | 0.032462658 | [illegible] |
| TG | TG 68:7\|TG 18:1_26:1_24:5 | [M+NH4]+ | 26.145 | 1090.97302 | 0.027846833 | 0.03052536 | 0.040850047 | [illegible] |
| TG | TG 68:8\|TG 18:1_26:2_24:5 | [M+NH4]+ | 25.762 | 1088.95435 | 0.03123047 | 0.032456797 | 0.042253228 | [illegible] |
| TG | TG 68:9\|TG 18:1_26:2_24:6 | [M+NH4]+ | 25.386 | 1086.94116 | 0.021810112 | 0.02354529 | 0.031949262 | [illegible] |
| TG | TG 70:3\|TG 18:1_26:1_26:1 | [M+NH4]+ | 27.887 | 1127.06641 | 0.000590465 | 0.000734376 | 0.000795949 | [illegible] |
| TG | TG 70:6\|TG 18:1_26:1_26:4 | [M+NH4]+ | 26.967 | 1121.01746 | 0.002435677 | 0.00490599 | 0.00623028 | [illegible] |
| TG | TG 70:7\|TG 18:1_26:1_26:5 | [M+NH4]+ | 26.54 | 1119.00525 | 0.003322673 | 0.006140847 | 0.007767797 | [illegible] |
| TG | TG O-46:1\|TG O-12:0_16:0_18:1 | [M+NH4]+ | 23.8 | 780.7428 | 0.011326385 | 0.011767165 | 0.016399987 | [illegible] |
| TG | TG O-50:0\|TG O-16:0_16:0_18:0 | [M+NH4]+ | 25.638 | 838.82269 | 0.015533414 | 0.018500358 | 0.021966839 | [illegible] |
| TG | TG O-50:1\|TG O-16:0_16:0_18:1 | [M+NH4]+ | 25.06 | 836.80481 | 0.047228581 | 0.050509005 | 0.062064747 | [illegible] |
| TG | TG O-50:2\|TG O-18:1_16:0_16:1 | [M+NH4]+ | 24.543 | 834.79114 | 0.025538113 | 0.026977627 | 0.036324657 | [illegible] |
| TG | TG O-51:1\|TG O-19:1_16:0_16:0 | [M+NH4]+ | 25.355 | 850.82141 | 0.009691779 | 0.011785762 | 0.013680662 | [illegible] |
| TG | TG O-52:0\|TG O-18:0_16:0_18:0 | [M+NH4]+ | 26.159 | 866.85205 | 0.010512329 | 0.012610839 | 0.013954503 | [illegible] |
| TG | TG O-52:1\|TG O-18:0_16:0_18:1 | [M+NH4]+ | 25.636 | 864.83807 | 0.054270055 | 0.062875912 | 0.073101607 | [illegible] |
| TG | TG O-52:2\|TG O-18:1_16:0_18:1 | [M+NH4]+ | 25.061 | 862.82092 | 0.061394463 | 0.067285857 | 0.087858141 | [illegible] |
| TG | TG O-52:3\|TG O-18:1_16:0_18:2 | [M+NH4]+ | 24.58 | 860.80737 | 0.030839634 | 0.032835193 | 0.044783616 | [illegible] |
| TG | TG O-54:1\|TG O-20:0_16:0_18:1 | [M+NH4]+ | 26.155 | 892.87 | 0.016634576 | 0.018659751 | 0.022909344 | [illegible] |
| TG | TG O-54:2\|TG O-20:1_16:0_18:1 | [M+NH4]+ | 25.667 | 890.85394 | 0.021105093 | 0.022611234 | 0.029750111 | [illegible] |
| TG | TG O-54:3\|TG O-18:0_16:0_20:3 | [M+NH4]+ | 25.302 | 888.83783 | 0.024529891 | 0.027101334 | 0.035825906 | [illegible] |
| TG | TG O-54:4\|TG O-16:0_16:0_22:4 | [M+NH4]+ | 24.917 | 886.82098 | 0.035658353 | 0.037153712 | 0.052437939 | [illegible] |
| TG | TG O-54:5\|TG O-16:0_16:0_22:5 | [M+NH4]+ | 24.424 | 884.80414 | 0.03891743 | 0.037511187 | 0.058461096 | [illegible] |
| TG | TG O-56:1\|TG O-22:0_16:0_18:1 | [M+NH4]+ | 26.636 | 920.90173 | 0.014114012 | 0.015813612 | 0.019583357 | [illegible] |
| TG | TG O-56:2\|TG O-22:1_16:0_18:1 | [M+NH4]+ | 26.159 | 918.88348 | 0.015467229 | 0.01705143 | 0.023180379 | [illegible] |
| TG | TG O-56:3\|TG O-18:0_16:0_22:3 | [M+NH4]+ | 25.758 | 916.8681 | 0.014979323 | 0.015736563 | 0.021684892 | [illegible] |
| TG | TG O-56:4\|TG O-18:0_16:0_22:4 | [M+NH4]+ | 25.421 | 914.85199 | 0.026318087 | 0.028419074 | 0.038192402 | [illegible] |
| TG | TG O-56:5\|TG O-18:0_16:0_22:5 | [M+NH4]+ | 25.016 | 912.83875 | 0.055276683 | 0.055344173 | 0.077645084 | [illegible] |
| TG | TG O-56:6\|TG O-18:1_16:0_22:5 | [M+NH4]+ | 24.435 | 910.82111 | 0.047946176 | 0.047989743 | 0.069559943 | [illegible] |
| TG | TG O-58:4\|TG O-18:0_18:1_22:3 | [M+NH4]+ | 25.887 | 942.88342 | 0.022719853 | 0.023740287 | 0.031621063 | [illegible] |
| TG | TG O-58:5\|TG O-18:0_18:1_22:4 | [M+NH4]+ | 25.523 | 940.87006 | 0.037052546 | 0.040700075 | 0.053124271 | [illegible] |
| TG | TG O-58:6\|TG O-18:0_18:1_22:5 | [M+NH4]+ | 25.019 | 938.8515 | 0.041427301 | 0.043279245 | 0.058229431 | [illegible] |
| TG | TG O-60:6\|TG O-22:1_16:0_22:5 | [M+NH4]+ | 25.572 | 966.8833 | 0.020192121 | 0.021433437 | 0.030679862 | [illegible] |
| DG | DG 32:0\|DG 16:0_16:0 | [M+NH4]+ | 17.59 | 586.54028 | 0.079949179 | 0.174921827 | 0.122688204 | [illegible] |
| DG | DG 32:1\|DG 16:0_16:1 | [M+NH4]+ | 16.651 | 584.52344 | 0.058586659 | 0.072777783 | 0.096768493 | [illegible] |
| DG | DG 33:0\|DG 16:0_17:0 | [M+NH4]+ | 18.089 | 600.55573 | 0.016796556 | 0.042366295 | 0.021876126 | [illegible] |
| DG | DG 33:1\|DG 16:0_17:1 | [M+NH4]+ | 17.188 | 598.53949 | 0.018411096 | 0.027494953 | 0.031266849 | [illegible] |
| DG | DG 34:0\|DG 16:0_18:0 | [M+NH4]+ | 18.656 | 614.57074 | 0.098699796 | 0.426788019 | 0.193354093 | [illegible] |
| DG | DG 34:1\|DG 16:0_18:1 | [M+NH4]+ | 17.697 | 612.55542 | 0.275268298 | 0.344143157 | 0.408839298 | [illegible] |
| DG | DG 34:2\|DG 16:0_18:2 | [M+NH4]+ | 17.14 | 610.53998 | 5.00386267 | 5.587351635 | 5.803498465 | [illegible] |
| DG | DG 35:0\|DG 17:0_18:0 | [M+NH4]+ | 19.182 | 628.58759 | 0.004251147 | 0.035351959 | 0.013828754 | [illegible] |
| DG | DG 35:1\|DG 17:0_18:1 | [M+NH4]+ | 18.228 | 626.57092 | 0.025153721 | 0.032459353 | 0.041422415 | [illegible] |
| DG | DG 35:2\|DG 17:1_18:1 | [M+NH4]+ | 17.324 | 624.5545 | 0.029477119 | 0.037537791 | 0.040976629 | [illegible] |
| DG | DG 36:0\|DG 18:0_18:0 | [M+NH4]+ | 19.688 | 642.6026 | 0.061840342 | 0.447404688 | 0.165544409 | [illegible] |
| DG | DG 36:1\|DG 16:0_20:1 | [M+NH4]+ | 18.754 | 640.58685 | 0.095337426 | 0.129851685 | 0.137879928 | [illegible] |
| DG | DG 36:2\|DG 18:1_18:1 | [M+NH4]+ | 17.827 | 638.57031 | 0.217936372 | 0.300499185 | 0.324265671 | [illegible] |
| DG | DG 36:3\|DG 18:1_18:2 | [M+NH4]+ | 17.029 | 636.55615 | 0.096668653 | 0.126008505 | 0.148727416 | [illegible] |
| DG | DG 36:4\|DG 16:0_20:4 | [M+NH4]+ | 16.66 | 634.53961 | 0.033876943 | 0.035237116 | 0.048773986 | [illegible] |
| DG | DG 36:5\|DG 16:1_20:4 | [M+NH4]+ | 15.857 | 632.52338 | 0.014951395 | 0.017711889 | 0.029277306 | [illegible] |
| DG | DG 37:2\|DG 18:1_19:1 | [M+NH4]+ | 18.323 | 652.58777 | 0.007541786 | 0.009833694 | 0.014554049 | [illegible] |
| DG | DG 38:1\|DG 16:0_22:1 | [M+NH4]+ | 19.711 | 668.61761 | 0.004725457 | 0.00313793 | 0.002162739 | [illegible] |
| DG | DG 38:2\|DG 18:1_20:1 | [M+NH4]+ | 18.811 | 666.60309 | 0.016223959 | 0.021831276 | 0.028392201 | [illegible] |
| DG | DG 38:3\|DG 18:0_20:3 | [M+NH4]+ | 18.259 | 664.58655 | 0.03090236 | 0.038141092 | 0.050801968 | [illegible] |
| DG | DG 38:4\|DG 18:0_20:4 | [M+NH4]+ | 17.842 | 662.57025 | 0.055200949 | 0.058112314 | 0.090665236 | [illegible] |
| DG | DG 38:5\|DG 16:0_22:5 | [M+NH4]+ | 16.836 | 660.55579 | 0.095859802 | 0.086711721 | 0.173266506 | [illegible] |
| DG | DG 38:6\|DG 16:0_22:6 | [M+NH4]+ | 16.45 | 658.54114 | 0.037940868 | 0.043357983 | 0.066589909 | [illegible] |
| DG | DG 38:6\|DG 16:1_22:5 | [M+NH4]+ | 16.451 | 658.53931 | 0.03611279 | 0.041929618 | 0.064159962 | [illegible] |
| DG | DG 40:4\|DG 18:0_22:4 | [M+NH4]+ | 18.597 | 690.60254 | 0.012443929 | 0.013280013 | 0.018064154 | [illegible] |
| DG | DG 40:5\|DG 18:0_22:5 | [M+NH4]+ | 17.875 | 688.58612 | 0.027765767 | 0.028062027 | 0.038786143 | [illegible] |
| DG | DG 40:5\|DG 18:1_22:4 | [M+NH4]+ | 17.637 | 688.58771 | 0.027947227 | 0.034739179 | 0.04818852 | [illegible] |
| DG | DG 40:6\|DG 18:0_22:6 | [M+NH4]+ | 17.562 | 686.57214 | 0.017830439 | 0.019486165 | 0.030484629 | [illegible] |
| DG | DG 40:7\|DG 18:1_22:6 | [M+NH4]+ | 16.584 | 684.55402 | 0.027531081 | 0.024566323 | 0.050442436 | [illegible] |
| DG | DG 42:2\|DG 18:1_24:1 | [M+NH4]+ | 20.743 | 722.66431 | 0.005621809 | 0.006914949 | 0.008004686 | [illegible] |
| SM | SM 24:3;2O | [M+H]+ | 10.154 | 559.38373 | 0.937625742 | 0.797020684 | 0.592738739 | [illegible] |
| SM | SM 32:1;2O | [M+H]+ | 13.739 | 675.5426 | 0.003464037 | 0.004016979 | 0.003974362 | [illegible] |
| SM | SM 32:5;2O | [M+H]+ | 11.335 | 667.479 | 0.036447382 | 0.039632154 | 0.027956825 | [illegible] |
| SM | SM 34:0;2O | [M+H]+ | 14.988 | 705.59039 | 0.008979099 | 0.010919348 | 0.009553552 | [illegible] |

| | | | | | | | | | | | | |
|---|---|---|---|---|---|---|---|---|---|---|---|---|
| SM | SM 34:1;2O | [M+H]+ | 14.632 | 703.5755 | 0.068959666 | 0.086423316 | 0.078326168 | 0.095805572 | 0.070346858 | 0.064300789 | 0.000186011 | [illegible] |
| SM | SM 34:5;3O | [M+H]+ | 12.752 | 711.51099 | 0.076471763 | 0.088964859 | 0.087673713 | 0.073552346 | 0.076808773 | 0.058339737 | 0.000206274 | [illegible] |
| SM | SM 36:1;2O | [M+H]+ | 15.689 | 731.60632 | 0.018472575 | 0.017227609 | 0.021216862 | 0.01542098 | 0.015781094 | 0.01519408 | 4.98277E-05 | [illegible] |
| SM | SM 39:4;3O | [M+H]+ | 17.46 | 783.59637 | 0.138699938 | 0.150577491 | 0.154851356 | 0.131775094 | 0.097034691 | 0.1008009 | 0.000374127 | [illegible] |
| SM | SM 40:1;2O | [M+H]+ | 18.156 | 787.66687 | 0.011917428 | 0.017287252 | 0.014018637 | 0.019541289 | 0.013900159 | 0.012649345 | 3.21459E-05 | [illegible] |
| SM | SM 42:1;2O | [M+H]+ | 19.521 | 815.69952 | 0.021359589 | 0.024341588 | 0.017336983 | 0.035238488 | 0.022021194 | 0.019312249 | 5.76151E-05 | [illegible] |
| SM | SM 42:3;2O | [M+H]+ | 15.37 | 811.66663 | 3.131226143 | 3.297357496 | 3.399932457 | 2.927945293 | 2.550584453 | 2.562439317 | 0.00844613 | [illegible] |
| Cer | Cer 31:1;2O\|Cer 15:1;2O/16:0 | [M+H-H2O]+ | 14.251 | 478.46182 | 0.243631381 | 0.265751149 | 0.249370901 | 0.232607664 | 0.186192147 | 0.216843176 | 0.000657168 | [illegible] |
| Cer | Cer 33:1;2O\|Cer 15:1;2O/18:0 | [M+H-H2O]+ | 15.205 | 506.49213 | 0.190765992 | 0.201977218 | 0.192116635 | 0.173112609 | 0.154472116 | 0.152344549 | 0.00051457 | [illegible] |
| Cer | Cer 34:1;2O\|Cer 18:1;2O/16:0 | [M+H-H2O]+ | 15.688 | 520.5083 | 0.031336986 | 0.036263815 | 0.048805689 | 0.095938579 | 0.074198389 | 0.056815178 | 8.4528E-05 | [illegible] |
| Cer | Cer 35:1;2O\|Cer 17:1;2O/18:0 | [M+H-H2O]+ | 16.219 | 534.52319 | 0.069281814 | 0.073860274 | 0.07372001 | 0.066701128 | 0.04800844 | 0.055931512 | 0.00018688 | [illegible] |
| Cer | Cer 39:1;2O\|Cer 15:1;2O/24:0 | [M+H-H2O]+ | 18.532 | 590.58569 | 1.404790699 | 1.413332458 | 1.398883372 | 1.235941777 | 1.077658651 | 1.115470354 | 0.003789265 | [illegible] |
| Cer | Cer 40:1;2O\|Cer 18:1;2O/22:0 | [M+H-H2O]+ | 18.913 | 604.60303 | 0.039530189 | 0.035632685 | 0.039289907 | 0.070733722 | 0.063983086 | 0.056192901 | 0.000106628 | [illegible] |
| Cer | Cer 41:1;2O\|Cer 17:1;2O/24:0 | [M+H-H2O]+ | 19.46 | 618.61865 | 0.074474928 | 0.080168347 | 0.079561721 | 0.08525507 | 0.075240312 | 0.071270248 | 0.000200888 | [illegible] |
| Cer | Cer 41:2;2O\|Cer 17:1;2O/24:1 | [M+H-H2O]+ | 18.379 | 616.60229 | 0.104887533 | 0.110599125 | 0.112350831 | 0.09722486 | 0.092155035 | 0.091216704 | 0.000282922 | [illegible] |
| Cer | Cer 42:1;2O\|Cer 18:1;2O/24:0 | [M+H-H2O]+ | 19.935 | 632.63458 | 0.069262862 | 0.067904543 | 0.106619815 | 0.168637005 | 0.142138816 | 0.120908937 | 0.000186829 | [illegible] |
| Cer | Cer 44:1;2O\|Cer 18:1;2O/26:0 | [M+H-H2O]+ | 20.874 | 660.66461 | 0.026281079 | 0.044669066 | 0.031411585 | 0.046623029 | 0.040270129 | 0.032823377 | 7.08903E-05 | [illegible] |
| Cer | Cer 44:1;3O\|Cer 20:1;2O/24:0;O | [M+H-H2O]+ | 19.321 | 676.66113 | 0.114310087 | 0.107017044 | 0.094999503 | 0.099105198 | 0.105343438 | 0.073973465 | 0.000308339 | [illegible] |
| Cer | Cer 46:1;3O\|Cer 20:1;2O/26:0;O | [M+H-H2O]+ | 21.153 | 704.69336 | 0.018870403 | 0.025955774 | 0.01739193 | 0.022478434 | 0.018430408 | 0.014753768 | 5.09008E-05 | [illegible] |
| CE | CE 18:1 | [M+NH4]+ | 25.234 | 668.63287 | 5.001605635 | 5.703124857 | 7.431715901 | 14.82695886 | 14.55004472 | 13.91569548 | 0.013491268 | [illegible] |
| CE | CE 18:2 | [M+NH4]+ | 24.67 | 666.6189 | 7.917021864 | 8.51511304 | 10.83421202 | 15.58713913 | 15.4138004 | 16.41052331 | 0.021355275 | [illegible] |
| CE | CE 19:1 | [M+NH4]+ | 25.64 | 682.64868 | 0.485334063 | 0.584229054 | 0.710151097 | 1.243543048 | 1.42517078 | 1.383442374 | 0.001309134 | [illegible] |
| CE | CE 20:1 | [M+NH4]+ | 25.991 | 696.66565 | 2.477039972 | 2.705377005 | 3.393753561 | 5.984989498 | 6.239196024 | 6.487219853 | 0.006681536 | [illegible] |
| CE | CE 20:2 | [M+NH4]+ | 25.378 | 694.64886 | 3.25927607 | 3.729935626 | 4.446644054 | 8.690236828 | 8.827566285 | 8.932410089 | 0.00879153 | [illegible] |
| CE | CE 20:3 | [M+NH4]+ | 24.784 | 692.63367 | 22.05204948 | 24.07892379 | 28.30904879 | 42.99712546 | 43.55488998 | 44.85432154 | 0.059482921 | [illegible] |
| CE | CE 20:4 | [M+NH4]+ | 24.265 | 690.61737 | 28.31611348 | 29.0617256 | 39.23231219 | 72.19499331 | 69.8378254 | 69.85388813 | 0.076379528 | [illegible] |
| CE | CE 20:5 | [M+NH4]+ | 23.711 | 688.60242 | 15.37918404 | 16.15881019 | 21.59299256 | 26.5158572 | 26.85785595 | 26.51599007 | 0.041483617 | [illegible] |
| CE | CE 22:0 | [M+NH4]+ | 26.627 | 726.7121 | 0.022071986 | 0.024520425 | 0.032138699 | 0.078919928 | 0.077407702 | 0.077483679 | 5.95367E-05 | [illegible] |
| CE | CE 22:1 | [M+NH4]+ | 26.626 | 724.69708 | 1.104218785 | 1.20346497 | 1.559755502 | 3.041693666 | 3.273684252 | 3.25739042 | 0.002978506 | [illegible] |
| CE | CE 22:2 | [M+NH4]+ | 26.059 | 722.67999 | 1.378443282 | 1.540978693 | 2.071636237 | 3.308922957 | 3.506515123 | 3.655314795 | 0.003718196 | [illegible] |
| CE | CE 22:3 | [M+NH4]+ | 25.479 | 720.66614 | 4.027933899 | 4.229508536 | 5.931717343 | 10.1013606 | 10.11808144 | 10.44511922 | 0.010864898 | [illegible] |
| CE | CE 22:4 | [M+NH4]+ | 24.9 | 718.65033 | 16.91752464 | 17.23408728 | 23.59389482 | 52.55536142 | 53.44431637 | 54.40620619 | 0.045633118 | [illegible] |
| CE | CE 22:5 | [M+NH4]+ | 24.355 | 716.63263 | 28.19247032 | 28.12671494 | 39.67629231 | 57.01832706 | 56.53007033 | 57.9256964 | 0.076046015 | [illegible] |
| CE | CE 22:6 | [M+NH4]+ | 23.944 | 714.61835 | 47.45847698 | 48.4713008 | 64.81100272 | 60.35446597 | 62.74251307 | 62.55362814 | 0.128013898 | [illegible] |
| CE | CE 23:1 | [M+NH4]+ | 26.926 | 738.71112 | 0.154362112 | 0.180770215 | 0.240304547 | 0.37097349 | 0.413793232 | 0.42482326 | 0.000416374 | [illegible] |
| CE | CE 24:0 | [M+NH4]+ | 27.845 | 754.7442 | 0.292523291 | 0.350400979 | 0.391201562 | 0.618885633 | 0.664973141 | 0.741022925 | 0.000789049 | [illegible] |
| CE | CE 24:1 | [M+NH4]+ | 27.183 | 752.72791 | 2.759890587 | 2.94097877 | 3.920099923 | 5.854609346 | 6.234833448 | 6.650530318 | 0.007444494 | [illegible] |
| CE | CE 24:2 | [M+NH4]+ | 26.663 | 750.71143 | 1.595125512 | 1.738683944 | 2.28887523 | 3.228168579 | 3.374617755 | 3.482028312 | 0.004302671 | [illegible] |
| CE | CE 24:3 | [M+NH4]+ | 26.146 | 748.69482 | 3.301567887 | 3.545620782 | 4.923340421 | 5.884041552 | 6.171389913 | 6.443818614 | 0.008905608 | [illegible] |
| CE | CE 24:4 | [M+NH4]+ | 25.591 | 746.68103 | 7.718954052 | 8.210664325 | 11.16312927 | 23.38290457 | 24.65862114 | 25.41166719 | 0.02082101 | [illegible] |
| CE | CE 24:5 | [M+NH4]+ | 25.015 | 744.66418 | 12.42803325 | 13.12422771 | 17.13299764 | 26.07469192 | 24.5908853 | 26.01070214 | 0.03352322 | [illegible] |
| CE | CE 24:6 | [M+NH4]+ | 24.514 | 742.6474 | 11.5709205 | 12.1122103 | 15.78196385 | 7.568442742 | 6.79968374 | 7.57040609 | 0.031211255 | [illegible] |
| CE | CE 25:0 | [M+NH4]+ | 28.125 | 768.75848 | 0.036019229 | 0.04233627 | 0.047301849 | 0.040109835 | 0.056865776 | 0.058109352 | 9.71578E-05 | [illegible] |
| CE | CE 25:1 | [M+NH4]+ | 27.466 | 766.74286 | 0.442827908 | 0.48348359 | 0.663450311 | 0.774567902 | 0.83787879 | 0.885749338 | 0.001194478 | [illegible] |
| CE | CE 26:0 | [M+NH4]+ | 28.399 | 782.77576 | 0.550890653 | 0.698690925 | 0.755349226 | 0.662002693 | 0.722129265 | 0.815619018 | 0.001485966 | [illegible] |
| CE | CE 26:1 | [M+NH4]+ | 27.707 | 780.75745 | 3.58988436 | 3.841194816 | 5.305343658 | 5.035303423 | 5.276912113 | 5.748926244 | 0.009683309 | [illegible] |
| CE | CE 26:2 | [M+NH4]+ | 27.196 | 778.7439 | 2.824718255 | 3.064111222 | 4.096181119 | 4.040119447 | 4.280086208 | 4.522817222 | 0.007619359 | [illegible] |
| CE | CE 26:3 | [M+NH4]+ | 26.721 | 776.72675 | 4.360042859 | 4.669641469 | 6.302018753 | 5.408180867 | 5.61076089 | 5.676066501 | 0.011760725 | [illegible] |
| CE | CE 26:4 | [M+NH4]+ | 26.225 | 774.71155 | 6.180310472 | 6.871785325 | 8.514956688 | 9.638143577 | 9.526164306 | 10.28675793 | 0.016670692 | [illegible] |
| CE | CE 26:5 | [M+NH4]+ | 25.711 | 772.6955 | 10.78922804 | 11.78276227 | 14.89656201 | 9.835385037 | 9.788322932 | 10.4666679 | 0.029102728 | [illegible] |
| CE | CE 26:6 | [M+NH4]+ | 25.061 | 770.68158 | 6.025904709 | 6.755151331 | 7.645991992 | 3.046859797 | 2.797945772 | 3.066614465 | 0.0162542 | [illegible] |
| CE | CE 28:1 | [M+NH4]+ | 28.259 | 808.7901 | 0.537155476 | 0.607607801 | 0.770407553 | 0.600312608 | 0.654595027 | 0.697521038 | 0.001448916 | [illegible] |
| CE | CE 28:2 | [M+NH4]+ | 27.738 | 806.77496 | 0.527375528 | 0.585784043 | 0.766405879 | 0.614361943 | 0.642378965 | 0.658427358 | 0.001422536 | [illegible] |
| CE | CE 28:3 | [M+NH4]+ | 27.265 | 804.75818 | 0.610244982 | 0.664081259 | 0.873725273 | 0.647256092 | 0.671133709 | 0.70395445 | 0.001646067 | [illegible] |
| CE | CE 28:4 | [M+NH4]+ | 26.797 | 802.74286 | 1.034381694 | 1.109882376 | 1.468040663 | 1.287808077 | 1.323211652 | 1.341796585 | 0.002790128 | [illegible] |
| CE | CE 28:5 | [M+NH4]+ | 26.302 | 800.72595 | 2.5701197 | 2.812473759 | 3.537401586 | 2.168921225 | 2.199605271 | 2.246433639 | 0.006932609 | [illegible] |
| CE | CE 28:6 | [M+NH4]+ | 25.822 | 798.71057 | 2.385550282 | 2.633485208 | 3.181315076 | 1.270590877 | 1.259978646 | 1.388345937 | 0.006434753 | [illegible] |
| CE | CE 30:1 | [M+NH4]+ | 28.853 | 836.82104 | 0.058078264 | 0.084609877 | 0.083930526 | 0.073121602 | 0.096603629 | 0.099912606 | 0.00015666 | [illegible] |
| CE | CE 30:2 | [M+NH4]+ | 28.283 | 834.80731 | 0.082400946 | 0.092017274 | 0.103522699 | 0.124769309 | 0.138202537 | 0.133025418 | 0.000222267 | [illegible] |
| CE | CE 30:3 | [M+NH4]+ | 27.778 | 832.79102 | 0.125781684 | 0.126643054 | 0.178479147 | 0.195829911 | 0.229219897 | 0.221861362 | 0.000339282 | [illegible] |
| CE | CE 30:4 | [M+NH4]+ | 27.317 | 830.77557 | 0.263899263 | 0.278521927 | 0.34872991 | 0.451204903 | 0.506527858 | 0.495383858 | 0.000711839 | [illegible] |
| CE | CE 30:5 | [M+NH4]+ | 26.852 | 828.75781 | 1.303834737 | 1.498125909 | 1.766235586 | 1.770957309 | 1.813567893 | 1.838980692 | 0.003516947 | [illegible] |
| CE | CE 30:6 | [M+NH4]+ | 26.427 | 826.74426 | 1.600875054 | 1.74501665 | 2.084324728 | 1.650496344 | 1.700038564 | 1.79159396 | 0.00431818 | [illegible] |
| CE | CE 32:4 | [M+NH4]+ | 27.816 | 858.80426 | 0.103492471 | 0.119401353 | 0.144848212 | 0.201563226 | 0.229198701 | 0.235526937 | 0.000279159 | [illegible] |
| CE | CE 32:5 | [M+NH4]+ | 27.366 | 856.79132 | 0.61851198 | 0.667678724 | 0.824439067 | 1.144985352 | 1.242411668 | 1.231787193 | 0.001668366 | [illegible] |
| CE | CE 32:6 | [M+NH4]+ | 26.966 | 854.77203 | 1.1594924 | 1.279223635 | 1.564132477 | 1.965013527 | 2.006178319 | 1.989563503 | 0.0031276 | [illegible] |
| CE | CE 34:5 | [M+NH4]+ | 27.865 | 884.82178 | 0.065602822 | 0.071746147 | 0.08742772 | 0.123184118 | 0.149700272 | 0.128142564 | 0.000176956 | [illegible] |
| CE | CE 34:6 | [M+NH4]+ | 27.468 | 882.80475 | 0.235768304 | 0.270820253 | 0.312329661 | 0.415496286 | 0.460980604 | 0.442577766 | 0.000635958 | [illegible] |

Whole Cell Lysate lipidomic data set

| | | | | | 24 hrs data set | | | | | | | |
|---|---|---|---|---|---|---|---|---|---|---|---|---|
| | | | | | lipid intensities normalized to LDs protein amount µg | | | | | | Relative abundance of lipid intensities | |
| Class | Metabolite name | Adduct type | Average Rt(min) | Average Mz | Control at pmol/ug | Control at pmol/ug | Control at pmol/ug | dsRNA at pmol/ug | dsRNA at pmol/ug | dsRNA at pmol/ug | control % | [illegible] |
| PC | PC 28:0\|PC 14:0_14:0 | [M+H]+ | 13.847 | 678.50513 | 0.078945111 | 0.082133089 | 0.068642729 | 0.028855144 | 0.022047438 | 0.023005223 | 0.00075496 | [illegible] |
| PC | PC 29:0 | [M+H]+ | 14.304 | 692.52362 | 0.049579041 | 0.051260182 | 0.047224301 | 0.030969744 | 0.023873394 | 0.023576879 | 0.00047413 | [illegible] |
| PC | PC 30:0\|PC 14:0_16:0 | [M+H]+ | 14.776 | 706.53735 | 0.69030725 | 0.749650752 | 0.674591676 | 0.417149274 | 0.302934965 | 0.299688555 | 0.00660143 | [illegible] |
| PC | PC 30:1\|PC 14:0_16:1 | [M+H]+ | 14.015 | 704.52264 | 0.228002344 | 0.194017901 | 0.183547079 | 0.105475277 | 0.072428015 | 0.074299185 | 0.0021804 | [illegible] |
| PC | PC 31:0\|PC 15:0_16:0 | [M+H]+ | 15.27 | 720.55255 | 0.212221504 | 0.22100763 | 0.19901223 | 0.182119117 | 0.129024654 | 0.12033465 | 0.00202948 | [illegible] |
| PC | PC 31:1 | [M+H]+ | 14.465 | 718.53687 | 0.125045737 | 0.125817512 | 0.125649827 | 0.092488845 | 0.06779868 | 0.066882761 | 0.00119582 | [illegible] |
| PC | PC 32:0\|PC 16:0_16:0 | [M+H]+ | 15.817 | 734.56897 | 1.978030051 | 2.20681238 | 1.713035321 | 1.28529368 | 0.905847331 | 0.949309421 | 0.01891598 | [illegible] |
| PC | PC 32:1\|PC 16:0_16:1 | [M+H]+ | 14.945 | 732.55267 | 2.40190413 | 2.495513836 | 2.464815404 | 1.776090416 | 1.319876263 | 1.29491595 | 0.0229695 | [illegible] |
| PC | PC 32:2\|PC 16:0_16:2 | [M+H]+ | 14.195 | 730.53717 | 0.317848991 | 0.295684961 | 0.312296955 | 0.239213354 | 0.177884238 | 0.183526754 | 0.0030396 | [illegible] |
| PC | PC 33:1\|PC 15:0_18:1 | [M+H]+ | 15.357 | 746.56934 | 0.724471307 | 0.771827284 | 0.808920332 | 0.635668967 | 0.444757912 | 0.424241386 | 0.00692815 | [illegible] |
| PC | PC 33:2\|PC 16:1_17:1 | [M+H]+ | 14.609 | 744.55273 | 0.146337778 | 0.147496355 | 0.154120653 | 0.101363276 | 0.074957461 | 0.074242359 | 0.00139943 | [illegible] |
| PC | PC 34:0\|PC 16:0_18:0 | [M+H]+ | 17.057 | 762.59979 | 1.196179616 | 1.290952799 | 1.082174842 | 0.899308512 | 0.656573993 | 0.684775354 | 0.01143911 | [illegible] |
| PC | PC 34:1\|PC 16:0_18:1 | [M+H]+ | 15.911 | 760.58374 | 9.713088442 | 9.271573818 | 10.37609216 | 7.385232579 | 5.366709668 | 5.50977161 | 0.09288663 | [illegible] |
| PC | PC 34:2\|PC 16:0_18:2 | [M+H]+ | 15.076 | 758.5686 | 2.305686248 | 2.335344902 | 2.391428049 | 1.666874416 | 1.260585463 | 1.267415147 | 0.02204936 | [illegible] |
| PC | PC 35:1\|PC 17:0_18:1 | [M+H]+ | 16.63 | 774.59949 | 0.752083908 | 0.747504726 | 0.796809555 | 0.687617518 | 0.564092426 | 0.573353968 | 0.00719221 | [illegible] |
| PC | PC 35:2\|PC 17:1_18:1 | [M+H]+ | 15.553 | 772.58344 | 0.447988218 | 0.428886209 | 0.457802021 | 0.328013257 | 0.232640651 | 0.24318341 | 0.00428413 | [illegible] |
| PC | PC 35:3\|PC 17:1_18:2 | [M+H]+ | 14.797 | 770.56995 | 0.077899291 | 0.073867229 | 0.081464934 | 0.051326178 | 0.03811484 | 0.037589122 | 0.00074495 | [illegible] |
| PC | PC 35:5\|PC 15:1_20:4 | [M+H]+ | 14.951 | 766.53802 | 0.267555588 | 0.273653106 | 0.28721305 | 0.209339115 | 0.152623342 | 0.147850917 | 0.00255864 | [illegible] |
| PC | PC 36:0\|PC 18:0_18:0 | [M+H]+ | 18.455 | 790.62946 | 0.071692891 | 0.07753375 | 0.063692057 | 0.054071816 | 0.040838516 | 0.040314633 | 0.0006856 | [illegible] |
| PC | PC 36:1\|PC 18:0_18:1 | [M+H]+ | 17.177 | 788.61646 | 2.803291739 | 2.951883902 | 2.922388522 | 2.12170594 | 1.633436126 | 1.706307578 | 0.02680798 | [illegible] |
| PC | PC 36:2\|PC 18:1_18:1 | [M+H]+ | 16.041 | 786.59967 | 3.694147893 | 3.737011843 | 4.005969605 | 3.051388249 | 2.008452757 | 2.113472852 | 0.03532728 | [illegible] |
| PC | PC 36:3\|PC 18:1_18:2 | [M+H]+ | 15.252 | 784.58429 | 1.109438656 | 1.103017009 | 1.161269058 | 0.842493574 | 0.61854051 | 0.626358213 | 0.0106096 | [illegible] |
| PC | PC 36:4\|PC 16:0_20:4 | [M+H]+ | 14.974 | 782.57025 | 1.167907031 | 1.260163731 | 1.295042757 | 0.872874263 | 0.687776895 | 0.690072513 | 0.01116874 | [illegible] |
| PC | PC 36:5\|PC 16:1_20:4 | [M+H]+ | 14.339 | 780.55273 | 0.324972038 | 0.314184876 | 0.315046421 | 0.179337939 | 0.021570336 | 0.025344088 | 0.00310772 | [illegible] |
| PC | PC 37:2\|PC 18:1_19:1 | [M+H]+ | 16.683 | 800.61438 | 0.157763163 | 0.142310973 | 0.161921965 | 0.125036668 | 0.089365354 | 0.094696902 | 0.0015087 | [illegible] |
| PC | PC 37:5 | [M+H]+ | 14.606 | 794.56757 | 0.060656801 | 0.05921893 | 0.063076475 | 0.048390156 | 0.034074727 | 0.032792493 | 0.00058006 | [illegible] |
| PC | PC 38:1 | [M+H]+ | 18.476 | 816.64795 | 0.078211236 | 0.080918532 | 0.075845651 | 0.043366955 | 0.038067545 | 0.039299169 | 0.00074794 | [illegible] |
| PC | PC 38:2\|PC 18:1_20:1 | [M+H]+ | 17.229 | 814.63171 | 0.240350738 | 0.24360857 | 0.246095174 | 0.158718566 | 0.120076817 | 0.124525096 | 0.00229848 | [illegible] |
| PC | PC 38:3\|PC 18:0_20:3 | [M+H]+ | 16.654 | 812.61694 | 0.511862409 | 0.513150013 | 0.52406896 | 0.419110095 | 0.292636618 | 0.301528086 | 0.00489496 | [illegible] |
| PC | PC 38:4\|PC 16:0_22:4 | [M+H]+ | 16.066 | 810.59955 | 0.99834921 | 1.076106799 | 1.151099285 | 0.937886452 | 0.687714202 | 0.707226864 | 0.00954725 | [illegible] |
| PC | PC 38:4\|PC 18:1_20:3 | [M+H]+ | 16.066 | 810.59924 | 0.871082917 | 0.88991008 | 0.937627963 | 0.830372077 | 0.604983625 | 0.610312622 | 0.0083302 | [illegible] |
| PC | PC 38:5\|PC 18:1_20:4 | [M+H]+ | 15.051 | 808.58356 | 1.148235408 | 1.15552003 | 1.20854624 | 0.851828046 | 0.663610971 | 0.624745376 | 0.01098062 | [illegible] |
| PC | PC 38:6\|PC 16:0_22:6 | [M+H]+ | 14.728 | 806.56915 | 0.468093727 | 0.584965156 | 0.458627924 | 0.271165218 | 0.219099867 | 0.210480017 | 0.0044764 | [illegible] |
| PC | PC 38:7\|PC 16:1_22:6 | [M+H]+ | 13.963 | 804.55316 | 0.077835601 | 0.077801224 | 0.074278266 | 0.039232404 | 0.03219855 | 0.03163732 | 0.00074434 | [illegible] |
| PC | PC 40:2 | [M+H]+ | 18.553 | 842.66388 | 0.032854064 | 0.021201882 | 0.024545425 | 0.014868895 | 0.013936195 | 0.01516717 | 0.00031418 | [illegible] |
| PC | PC 40:4 | [M+H]+ | 16.827 | 838.63147 | 0.178667239 | 0.174165196 | 0.177181719 | 0.157086602 | 0.117821844 | 0.117381328 | 0.0017086 | [illegible] |
| PC | PC 40:5\|PC 18:0_22:5 | [M+H]+ | 16.178 | 836.61664 | 0.307586255 | 0.307652032 | 0.366707682 | 0.304581246 | 0.163484252 | 0.17209151 | 0.00294146 | [illegible] |
| PC | PC 40:6\|PC 18:0_22:6 | [M+H]+ | 15.787 | 834.60077 | 0.240563703 | 0.242549933 | 0.237455613 | 0.168711877 | 0.12731457 | 0.132090967 | 0.00230052 | [illegible] |
| PC | PC 40:6\|PC 18:1_22:5 | [M+H]+ | 15.155 | 834.6004 | 0.235008247 | 0.234804519 | 0.233242797 | 0.151258349 | 0.1139704 | 0.109911922 | 0.00224739 | [illegible] |
| PC | PC 40:7\|PC 18:1_22:6 | [M+H]+ | 14.845 | 832.58484 | 0.26293974 | 0.257290241 | 0.267094344 | 0.163864727 | 0.129869616 | 0.122513233 | 0.0025145 | [illegible] |
| PC | PC 42:10 | [M+H]+ | 13.989 | 854.56732 | 0.01871847 | 0.014640861 | 0.014324104 | 0.010224803 | 0.008991824 | 0.005041762 | 0.00017901 | [illegible] |
| PC | PC 42:4 | [M+H]+ | 17.856 | 866.66406 | 0.041094277 | 0.041909964 | 0.039718199 | 0.015694458 | 0.016581633 | 0.01054272 | 0.00039299 | [illegible] |
| PC | PC 42:5 | [M+H]+ | 16.971 | 864.64722 | 0.090312282 | 0.067303248 | 0.053281703 | 0.005886388 | 0.006099953 | 0.003563859 | 0.00086366 | [illegible] |
| PC | PC 42:9\|PC 20:4_22:5 | [M+H]+ | 14.319 | 856.5863 | 0.02976415 | 0.027376657 | 0.030191274 | 0.019455246 | 0.014418946 | 0.01510594 | 0.00028464 | [illegible] |
| PC | PC O-32:0 | [M+H]+ | 16.794 | 720.59113 | 0.208102165 | 0.214662269 | 0.185143588 | 0.1623915 | 0.114542457 | 0.12552997 | 0.00199009 | [illegible] |
| PC | PC O-32:1 | [M+H]+ | 16.586 | 718.57361 | 0.107069467 | 0.105318373 | 0.100614685 | 0.099581541 | 0.076716613 | 0.076848241 | 0.00102391 | [illegible] |
| PC | PC O-33:6 | [M+H]+ | 14.787 | 722.5108 | 0.027881968 | 0.025757699 | 0.030722457 | 0.022545495 | 0.016186152 | 0.022464815 | 0.00026664 | [illegible] |
| PC | PC O-34:0 | [M+H]+ | 18.199 | 748.62091 | 0.024861911 | 0.171917681 | 0.145760555 | 0.015221014 | 0.086984752 | 0.093013057 | 0.00023776 | [illegible] |
| PC | PC O-34:2 | [M+H]+ | 16.679 | 744.58893 | 0.062553471 | 0.111952199 | 0.125333628 | 0.122093104 | 0.091814689 | 0.088984949 | 0.0005982 | [illegible] |
| PC | PC O-35:1 | [M+H]+ | 19.535 | 760.62244 | 0.00370401 | 0.003771349 | 0.003016935 | 0.002571257 | 0.001913236 | 0.00198562 | 3.5422E-05 | [illegible] |
| PC | PC O-36:1 | [M+H]+ | 18.223 | 774.63654 | 0.047229459 | 0.048352941 | 0.047123642 | 0.037895431 | 0.02925011 | 0.031334127 | 0.00045166 | [illegible] |
| PC | PC O-36:2 | [M+H]+ | 16.966 | 772.62079 | 0.055620757 | 0.052234381 | 0.020321734 | 0.05018795 | 0.036769314 | 0.038465462 | 0.0005319 | [illegible] |
| PC | PC O-36:5 | [M+H]+ | 15.546 | 766.57495 | 0.081064435 | 0.076336771 | 0.083472957 | 0.093033219 | 0.071817566 | 0.065720961 | 0.00077522 | [illegible] |
| PC | PC O-37:8 | [M+H]+ | 16.464 | 774.5412 | 0.024862012 | 0.026208258 | 0.030775746 | 0.045967568 | 0.031031619 | 0.032391975 | 0.00023776 | [illegible] |
| PC | PC O-38:4 | [M+H]+ | 16.97 | 796.62158 | 0.131067998 | 0.112348789 | 0.136630037 | 0.109042178 | 0.086443037 | 0.090751329 | 0.00125341 | [illegible] |
| PC | PC O-38:5 | [M+H]+ | 15.802 | 794.60425 | 0.144936205 | 0.143917962 | 0.161119787 | 0.163203649 | 0.125964302 | 0.127333888 | 0.00138603 | [illegible] |
| PC | PC O-40:4 | [M+H]+ | 18.303 | 824.65253 | 0.024317591 | 0.022397381 | 0.023914821 | 0.018583838 | 0.016173016 | 0.014867013 | 0.00023255 | [illegible] |
| PC | PC O-40:5 | [M+H]+ | 17.07 | 822.63788 | 0.033070751 | 0.039985224 | 0.034727275 | 0.026360249 | 0.021542914 | 0.025478265 | 0.00031626 | [illegible] |
| PC | PC O-40:6 | [M+H]+ | 16.722 | 820.62207 | 0.062570172 | 0.059024075 | 0.064842997 | 0.065278485 | 0.052430952 | 0.049272667 | 0.00059836 | [illegible] |
| PE | PE 32:0\|PE 16:0_16:0 | [M-H]- | 14.756 | 690.50549 | 0.138973547 | 0.136949787 | 0.128779508 | 0.103450865 | 0.066064466 | 0.071152087 | 0.00132901 | [illegible] |
| PE | PE 32:0\|PE 16:0_16:0 | [M-H]- | 15.568 | 690.50787 | 0.030836343 | 0.032946995 | 0.023405541 | 0.023130602 | 0.017344669 | 0.016141669 | 0.00029489 | [illegible] |
| PE | PE 33:0;2O\|PE 16:0_17:0;2O | [M-H]- | 13.825 | 736.51312 | 0.020394033 | 0.01835469 | 0.016714751 | 0.008352742 | 0.005023447 | 0.005597956 | 0.00019503 | [illegible] |
| PE | PE 33:0\|PE 16:0_17:0 | [M-H]- | 15.23 | 704.52325 | 0.030809588 | 0.031525011 | 0.030723655 | 0.030869531 | 0.020119533 | 0.019211244 | 0.00029463 | [illegible] |
| PE | PE 34:0;O\|PE 18:0_16:0;O | [M-H]- | 14.577 | 734.53473 | 0.021689798 | 0.021481046 | 0.020908887 | 0.03236779 | 0.015465578 | 0.012225611 | 0.00020742 | [illegible] |
| PE | PE 34:0\|PE 16:0_18:0 | [M-H]- | 16.618 | 718.53741 | 0.106955739 | 0.10828492 | 0.08614011 | 0.086146447 | 0.079399496 | 0.073113704 | 0.00102282 | [illegible] |
| PE | PE 34:0\|PE 16:0_18:0 | [M-H]- | 15.792 | 718.53741 | 0.235377376 | 0.255132972 | 0.239265266 | 0.224637187 | 0.17430055 | 0.154615627 | 0.00225092 | [illegible] |
| PE | PE 34:1\|PE 16:0_18:1 | [M-H]- | 15.688 | 716.52277 | 0.98547762 | 0.986596111 | 0.906072297 | 0.813380796 | 0.586125127 | 0.617206169 | 0.00942416 | [illegible] |
| PE | PE 34:2\|PE 16:1_18:1 | [M-H]- | 14.875 | 714.50635 | 0.242779554 | 0.233888077 | 0.217640139 | 0.168093422 | 0.120485885 | 0.12980648 | 0.00232171 | [illegible] |
| PE | PE 34:4\|PE 18:1_16:3 | [M-H]- | 13.958 | 710.47601 | 0.006622372 | 0.006267199 | 0.00591621 | 0.002659107 | 0.001699034 | 0.001890128 | 6.333E-05 | [illegible] |
| PE | PE 35:1\|PE 18:0_17:1 | [M-H]- | 16.227 | 730.53796 | 0.257649593 | 0.257236996 | 0.229634072 | 0.200627978 | 0.16073118 | 0.157659592 | 0.00246391 | [illegible] |
| PE | PE 35:2\|PE 17:1_18:1 | [M-H]- | 15.324 | 728.52325 | 0.095873243 | 0.087442019 | 0.090847169 | 0.071097129 | 0.048746727 | 0.052575096 | 0.00091684 | [illegible] |

| | | | | | | | | |
|---|---|---|---|---|---|---|---|---|
| PE | PE 36:0\|PE 18:0_18:0 | [M-H]- | 17.06 | 746.57013 | 0.209009945 | 0.230172185 | 0.194284241 | [illegible] |
| PE | PE 36:1;4O\|PE 18:1_18:0;4O | [M-H]- | 13.383 | 808.5332 | 0.001472603 | 0.001408409 | 0.001204846 | [illegible] |
| PE | PE 36:1\|PE 18:0_18:1 | [M-H]- | 16.756 | 744.55133 | 3.019204164 | 2.970351688 | 2.702391622 | [illegible] |
| PE | PE 36:1\|PE 18:0_18:1 | [M-H]- | 14.584 | 744.56207 | 0.006897513 | 0.006790214 | 0.005945181 | [illegible] |
| PE | PE 36:2\|PE 18:0_18:2 | [M-H]- | 15.81 | 742.53748 | 1.479094077 | 1.331779521 | 1.388021268 | [illegible] |
| PE | PE 36:2\|PE 18:1_18:1 | [M-H]- | 15.076 | 742.53741 | 0.278346408 | 0.255420865 | 0.273041578 | [illegible] |
| PE | PE 36:3;O\|PE 18:1_18:2;O | [M-H]- | 16.23 | 756.51587 | 0.002239004 | 0.002065523 | 0.001685927 | [illegible] |
| PE | PE 36:3\|PE 18:1_18:2 | [M-H]- | 15.064 | 740.5238 | 0.278477625 | 0.261029442 | 0.262425248 | [illegible] |
| PE | PE 36:4\|PE 16:0_20:4 | [M-H]- | 14.846 | 738.50763 | 0.245085068 | 0.236022591 | 0.225978685 | [illegible] |
| PE | PE 36:5\|PE 16:1_20:4 | [M-H]- | 14.11 | 736.49164 | 0.038987154 | 0.036613196 | 0.036720643 | [illegible] |
| PE | PE 37:1\|PE 18:0_19:1 | [M-H]- | 16.549 | 758.56989 | 0.17842223 | 0.166859137 | 0.165087304 | [illegible] |
| PE | PE 37:2\|PE 18:1_19:1 | [M-H]- | 16.284 | 756.55225 | 0.071030084 | 0.065704918 | 0.062890238 | [illegible] |
| PE | PE 37:2\|PE 18:1_19:1 | [M-H]- | 15.554 | 756.55225 | 0.024089005 | 0.046186374 | 0.056923733 | [illegible] |
| PE | PE 37:4\|PE 17:0_20:4 | [M-H]- | 15.334 | 752.52197 | 0.068134807 | 0.063920893 | 0.070932551 | [illegible] |
| PE | PE 38:1\|PE 18:0_20:1 | [M-H]- | 17.163 | 772.58527 | 0.492880384 | 0.464180568 | 0.464401114 | [illegible] |
| PE | PE 38:2\|PE 18:0_20:2 | [M-H]- | 16.052 | 770.5686 | 0.467519376 | 0.419534241 | 0.454141698 | [illegible] |
| PE | PE 38:2\|PE 18:1_20:1 | [M-H]- | 16.819 | 770.56885 | 0.167642208 | 0.16206143 | 0.158563476 | [illegible] |
| PE | PE 38:3\|PE 18:0_20:3 | [M-H]- | 16.273 | 768.55151 | 0.52377578 | 0.48867501 | 0.464552419 | [illegible] |
| PE | PE 38:4;O\|PE 18:1_20:3;O | [M-H]- | 16.338 | 782.53094 | 0.01344532 | 0.012524827 | 0.012735586 | [illegible] |
| PE | PE 38:4\|PE 18:0_20:4 | [M-H]- | 15.858 | 766.53699 | 2.526634237 | 2.507789906 | 2.439615984 | [illegible] |
| PE | PE 38:5\|PE 18:1_20:4 | [M-H]- | 14.954 | 764.52118 | 0.776524826 | 0.719705834 | 0.738132547 | [illegible] |
| PE | PE 38:6\|PE 16:0_22:6 | [M-H]- | 14.634 | 762.50519 | 0.27566278 | 0.272449175 | 0.259379664 | [illegible] |
| PE | PE 39:3\|PE 18:0_21:3 | [M-H]- | 16.839 | 782.56934 | 0.02612353 | 0.024535881 | 0.02352415 | [illegible] |
| PE | PE 39:4\|PE 17:0_22:4 | [M-H]- | 16.029 | 780.55499 | 0.04932176 | 0.044901545 | 0.044746402 | [illegible] |
| PE | PE 40:1\|PE 22:0_18:1 | [M-H]- | 18.958 | 800.61578 | 0.073196422 | 0.070370677 | 0.069270633 | [illegible] |
| PE | PE 40:4\|PE 18:0_22:4 | [M-H]- | 16.586 | 794.56958 | 0.525478021 | 0.485599342 | 0.482561607 | [illegible] |
| PE | PE 40:4\|PE 18:0_22:4 | [M-H]- | 16.586 | 794.5697 | 0.564535606 | 0.523147843 | 0.52185465 | [illegible] |
| PE | PE 40:6\|PE 18:0_22:6 | [M-H]- | 15.615 | 790.53607 | 0.863569962 | 0.846664671 | 0.826558849 | [illegible] |
| PE | PE 42:1\|PE 24:0_18:1 | [M-H]- | 20.031 | 828.6488 | 0.091190292 | 0.086572038 | 0.077175084 | [illegible] |
| PE | PE 42:2\|PE 18:1_24:1 | [M-H]- | 18.918 | 826.63232 | 0.049270588 | 0.034966094 | 0.03632697 | [illegible] |
| PE | PE 42:4\|PE 18:0_24:4 | [M-H]- | 17.439 | 822.59937 | 0.033046152 | 0.029229146 | 0.031499229 | [illegible] |
| PE | PE 42:6;2O\|PE 20:3_22:3;2O | [M-H]- | 16.284 | 850.55524 | 0.020741562 | 0.019531977 | 0.021056448 | [illegible] |
| PE | PE 42:7;O\|PE 20:4_22:3;O | [M-H]- | 16.455 | 832.5459 | 0.055364315 | 0.050886111 | 0.055499202 | [illegible] |
| PE | PE 42:8;2O\|PE 20:4_22:4;2O | [M-H]- | 14.937 | 846.5246 | 0.016864288 | 0.016595897 | 0.018619883 | [illegible] |
| PE | PE 42:9;O\|PE 22:6_20:3;O | [M-H]- | 15.137 | 828.51447 | 0.011609839 | 0.012262961 | 0.012835174 | [illegible] |
| PE | PE 43:1\|PE 25:0_18:1 | [M-H]- | 20.565 | 842.66357 | 0.006172546 | 0.005840074 | 0.005312596 | [illegible] |
| PE | PE 43:2\|PE 19:1_24:1 | [M-H]- | 19.439 | 840.64838 | 0.005452217 | 0.004822539 | 0.004726523 | [illegible] |
| PE | PE 44:10;2O\|PE 22:6_22:4;2O | [M-H]- | 14.74 | 870.52594 | 0.007573582 | 0.007341021 | 0.009112685 | [illegible] |
| PE | PE 44:2\|PE 18:1_26:1 | [M-H]- | 19.965 | 854.66187 | 0.021713766 | 0.0200251 | 0.01902818 | [illegible] |
| PE | PE 44:4\|PE 24:0_20:4 | [M-H]- | 19.118 | 850.63184 | 0.013066988 | 0.011981073 | 0.012078593 | [illegible] |
| PE | PE 44:7;O\|PE 22:4_22:3;O | [M-H]- | 17.218 | 860.57733 | 0.027687839 | 0.026590741 | 0.028973425 | [illegible] |
| PE | PE 44:8;2O\|PE 22:5_22:3;2O | [M-H]- | 15.938 | 874.55664 | 0.008227791 | 0.009015653 | 0.011082786 | [illegible] |
| PE | PE 44:8;O\|PE 22:5_22:3;O | [M-H]- | 16.525 | 858.56152 | 0.040289851 | 0.037125575 | 0.043113436 | [illegible] |
| PE | PE 46:5\|PE 24:0_22:5 | [M-H]- | 19.145 | 876.64899 | 0.00810725 | 0.007124167 | 0.007752885 | [illegible] |
| PE | PE O-30:0\|PE O-14:0_16:0 | [M-H]- | 14.49 | 648.49634 | 0.004546257 | 0.004353912 | 0.004855356 | [illegible] |
| PE | PE O-32:0\|PE O-16:0_16:0 | [M-H]- | 16.36 | 676.52643 | 0.008210713 | 0.007600163 | 0.006508483 | [illegible] |
| PE | PE O-32:1\|PE O-16:1_16:0 | [M-H]- | 16.22 | 674.51172 | 0.092741374 | 0.091726874 | 0.074929701 | [illegible] |
| PE | PE O-32:2\|PE O-16:1_16:1 | [M-H]- | 15.317 | 672.49725 | 0.021235907 | 0.019404319 | 0.020461389 | [illegible] |
| PE | PE O-33:1\|PE O-17:1_16:0 | [M-H]- | 16.755 | 688.52771 | 0.02083807 | 0.020569147 | 0.019404264 | [illegible] |
| PE | PE O-33:2\|PE O-15:1_18:1 | [M-H]- | 15.8 | 686.51208 | 0.014475186 | 0.013879453 | 0.013362491 | [illegible] |
| PE | PE O-34:1\|PE O-18:1_16:0 | [M-H]- | 17.311 | 702.54205 | 0.177886335 | 0.203341745 | 0.148258106 | [illegible] |
| PE | PE O-34:2\|PE O-18:2_16:0 | [M-H]- | 16.329 | 700.52704 | 0.465201996 | 0.436263915 | 0.406439789 | [illegible] |
| PE | PE O-34:3\|PE O-16:1_18:2 | [M-H]- | 15.535 | 698.51331 | 0.042951461 | 0.040862234 | 0.043045796 | [illegible] |
| PE | PE O-35:2\|PE O-17:1_18:1 | [M-H]- | 16.856 | 714.54224 | 0.042693666 | 0.037419769 | 0.034841971 | [illegible] |
| PE | PE O-35:5\|PE O-15:1_20:4 | [M-H]- | 14.905 | 708.49518 | 0.011099785 | 0.00899967 | 0.010458943 | [illegible] |
| PE | PE O-36:1\|PE O-20:1_16:0 | [M-H]- | 18.43 | 730.57507 | 0.034989656 | 0.034857005 | 0.030101646 | [illegible] |
| PE | PE O-36:1\|PE O-20:1_16:0 | [M-H]- | 16.869 | 730.57507 | 0.074930695 | 0.061242412 | 0.062728368 | [illegible] |
| PE | PE O-36:2\|PE O-18:1_18:1 | [M-H]- | 17.416 | 728.55975 | 0.19032037 | 0.181294133 | 0.177051036 | [illegible] |
| PE | PE O-36:2\|PE O-18:1_18:1 | [M-H]- | 17.416 | 728.55975 | 0.183891867 | 0.1750041 | 0.173109167 | [illegible] |
| PE | PE O-36:3\|PE O-18:2_18:1 | [M-H]- | 16.431 | 726.54224 | 0.128973468 | 0.115646573 | 0.120690457 | [illegible] |
| PE | PE O-36:5\|PE O-16:1_20:4 | [M-H]- | 15.402 | 722.51154 | 0.822876378 | 0.744077484 | 0.793933966 | [illegible] |
| PE | PE O-36:6\|PE O-16:1_20:5 | [M-H]- | 14.776 | 720.49567 | 0.073858802 | 0.069736033 | 0.069832594 | [illegible] |
| PE | PE O-36:7\|PE O-14:1_22:6 | [M-H]- | 14.223 | 718.48053 | 0.005400534 | 0.005999976 | 0.005075792 | [illegible] |
| PE | PE O-37:2\|PE O-19:1_18:1 | [M-H]- | 17.893 | 742.57562 | 0.008563349 | 0.008257426 | 0.008008015 | [illegible] |
| PE | PE O-37:4\|PE O-17:1_20:3 | [M-H]- | 16.389 | 738.54266 | 0.013851678 | 0.011825224 | 0.012067026 | [illegible] |
| PE | PE O-38:1\|PE O-22:1_16:0 | [M-H]- | 19.53 | 758.60632 | 0.011584736 | 0.01138657 | 0.008681702 | [illegible] |
| PE | PE O-38:2\|PE O-18:1_20:1 | [M-H]- | 18.484 | 756.59131 | 0.046130319 | 0.043469824 | 0.041418833 | [illegible] |
| PE | PE O-38:3\|PE O-18:1_20:2 | [M-H]- | 17.548 | 754.57599 | 0.026048238 | 0.024781051 | 0.02556239 | [illegible] |
| PE | PE O-38:5\|PE O-18:1_20:4 | [M-H]- | 16.453 | 750.54175 | 1.312810942 | 1.186144524 | 1.208902159 | [illegible] |
| PE | PE O-38:6\|PE O-18:2_20:4 | [M-H]- | 15.489 | 748.52576 | 1.024692426 | 0.941990737 | 1.032245602 | [illegible] |
| PE | PE O-38:7\|PE O-16:1_22:6 | [M-H]- | 15.162 | 746.50995 | 0.629121628 | 0.597254342 | 0.617374053 | [illegible] |
| PE | PE O-39:4\|PE O-17:1_22:3 | [M-H]- | 17.468 | 766.57367 | 0.018017331 | 0.017074775 | 0.016552538 | [illegible] |
| PE | PE O-39:5\|PE O-17:1_22:4 | [M-H]- | 16.661 | 764.55804 | 0.083365646 | 0.073349071 | 0.078692535 | [illegible] |
| PE | PE O-39:6\|PE O-17:1_22:5 | [M-H]- | 15.984 | 762.54498 | 0.094567018 | 0.084703637 | 0.090250845 | [illegible] |
| PE | PE O-39:7\|PE O-17:1_22:6 | [M-H]- | 15.653 | 760.52808 | 0.078240788 | 0.072577508 | 0.075170197 | [illegible] |
| PE | PE O-39:8\|PE O-17:2_22:6 | [M-H]- | 14.803 | 758.51099 | 0.004724419 | 0.004704301 | 0.005183799 | [illegible] |
| PE | PE O-40:2\|PE O-18:1_22:1 | [M-H]- | 19.542 | 784.62109 | 0.017022754 | 0.016338547 | 0.014759447 | [illegible] |
| PE | PE O-40:3\|PE O-18:1_22:2 | [M-H]- | 18.587 | 782.60443 | 0.017088711 | 0.015640589 | 0.016195422 | [illegible] |
| PE | PE O-40:4\|PE O-18:0_22:4 | [M-H]- | 17.395 | 780.58972 | 0.068199312 | 0.058272639 | 0.069640906 | [illegible] |
| PE | PE O-40:4\|PE O-18:1_22:3 | [M-H]- | 17.974 | 780.58972 | 0.061701871 | 0.057837949 | 0.059120594 | [illegible] |
| PE | PE O-40:5\|PE O-18:1_22:4 | [M-H]- | 17.216 | 778.57397 | 0.542175035 | 0.587234858 | 0.590046607 | [illegible] |
| PE | PE O-40:6\|PE O-18:1_22:5 | [M-H]- | 16.522 | 776.55737 | 1.047623465 | 0.943661567 | 0.975032781 | [illegible] |
| PE | PE O-40:7\|PE O-18:1_22:6 | [M-H]- | 16.201 | 774.54279 | 0.777650712 | 0.73217976 | 0.72490607 | [illegible] |
| PE | PE O-40:7\|PE O-18:2_22:5 | [M-H]- | 16.201 | 774.54297 | 0.717355835 | 0.703751047 | 0.698562884 | [illegible] |
| PE | PE O-40:8\|PE O-18:2_22:6 | [M-H]- | 15.255 | 772.52777 | 0.244245199 | 0.219656511 | 0.248627167 | [illegible] |
| PE | PE O-40:9\|PE O-18:3_22:6 | [M-H]- | 14.572 | 770.51135 | 0.00443969 | 0.004370425 | 0.00519015 | [illegible] |
| PE | PE O-41:6\|PE O-19:1_22:5 | [M-H]- | 16.824 | 790.57507 | 0.019316466 | 0.01671127 | 0.017604247 | [illegible] |
| PE | PE O-42:3\|PE O-24:2_18:1 | [M-H]- | 19.567 | 810.6358 | 0.00401118 | 0.003577948 | 0.003658355 | [illegible] |
| PE | PE O-42:4\|PE O-18:1_24:3 | [M-H]- | 18.838 | 808.61993 | 0.010537394 | 0.009773537 | 0.009894042 | [illegible] |
| PE | PE O-42:6\|PE O-20:1_22:5 | [M-H]- | 17.535 | 804.58923 | 0.109154048 | 0.100763189 | 0.105513879 | [illegible] |
| PE | PE O-42:7\|PE O-18:1_24:6 | [M-H]- | 17.282 | 802.57446 | 0.033411136 | 0.031003765 | 0.030651138 | [illegible] |
| PE | PE O-42:7\|PE O-18:1_24:6 | [M-H]- | 16.861 | 802.57465 | 0.048155218 | 0.047649566 | 0.044365168 | [illegible] |
| PE | PE O-44:6\|PE O-24:2_20:4 | [M-H]- | 18.615 | 832.62006 | 0.009709615 | 0.008587904 | 0.008735253 | [illegible] |
| PE | PE O-44:7\|PE O-22:1_22:6 | [M-H]- | 18.39 | 830.60522 | 0.005782375 | 0.005492287 | 0.00508316 | [illegible] |
| PE | PE O-46:8\|PE O-24:2_22:6 | [M-H]- | 18.339 | 856.62006 | 0.003229199 | 0.002995762 | 0.002812948 | [illegible] |
| PE | PEOH 42:1\|PEtOH 16:0_26:1 | [M-H]- | 18.8 | 813.64001 | 0.185099251 | 0.208141243 | 0.220592528 | [illegible] |
| PS | PS 32:0\|PS 16:0_16:0 | [M-H]- | 13.45 | 734.4978 | 0.028640391 | 0.030032423 | 0.02280642 | [illegible] |
| PS | PS 32:1\|PS 16:0_16:1 | [M-H]- | 12.911 | 732.48175 | 0.037662681 | 0.036124037 | 0.034599978 | [illegible] |
| PS | PS 33:1\|PS 16:0_17:1 | [M-H]- | 13.219 | 746.49908 | 0.023408301 | 0.022264562 | 0.02066991 | [illegible] |
| PS | PS 34:1\|PS 18:0_16:1 | [M-H]- | 13.555 | 760.51202 | 0.630003865 | 0.637164068 | 0.610414231 | [illegible] |
| PS | PS 34:2\|PS 16:1_18:1 | [M-H]- | 13.045 | 758.4975 | 0.048476317 | 0.047020797 | 0.046826144 | [illegible] |
| PS | PS 34:4 | [M-H]- | 12.897 | 754.46472 | 0.002653205 | 0.002213158 | 0.002403762 | [illegible] |
| PS | PS 35:1\|PS 17:0_18:1 | [M-H]- | 13.944 | 774.52832 | 0.188840929 | 0.188124214 | 0.184700832 | [illegible] |
| PS | PS 35:2\|PS 17:1_18:1 | [M-H]- | 13.379 | 772.51294 | 0.019353634 | 0.018814589 | 0.019340095 | [illegible] |
| PS | PS 36:1\|PS 18:0_18:1 | [M-H]- | 14.385 | 788.54138 | 2.228908065 | 2.487337193 | 2.481569766 | [illegible] |
| PS | PS 36:2\|PS 18:0_18:2 | [M-H]- | 13.755 | 786.52728 | 0.530288932 | 0.519812064 | 0.528179176 | [illegible] |
| PS | PS 36:4\|PS 16:0_20:4 | [M-H]- | 12.983 | 782.4964 | 0.016675646 | 0.016900081 | 0.014939539 | [illegible] |
| PS | PS 37:4\|PS 17:0_20:4 | [M-H]- | 13.94 | 796.50909 | 0.014748023 | 0.015614232 | 0.01660787 | [illegible] |
| PS | PS 37:4\|PS 17:0_20:4 | [M-H]- | 13.323 | 796.51208 | 0.006849045 | 0.005771547 | 0.006510753 | [illegible] |
| PS | PS 38:3\|PS 18:0_20:3 | [M-H]- | 14.035 | 812.54108 | 0.431251667 | 0.431876784 | 0.440827576 | [illegible] |
| PS | PS 38:4\|PS 18:0_20:4 | [M-H]- | 14.35 | 810.5246 | 0.133501562 | 0.154151285 | 0.177906186 | [illegible] |
| PS | PS 38:4\|PS 18:0_20:4 | [M-H]- | 13.701 | 810.5274 | 0.347509734 | 0.331702886 | 0.343295067 | [illegible] |
| PS | PS 39:6\|PS 17:0_22:6 | [M-H]- | 13.168 | 820.51416 | 0.021688163 | 0.020302206 | 0.022595067 | [illegible] |
| PS | PS 40:3\|PS 18:0_22:3 | [M-H]- | 14.957 | 840.57294 | 0.827770004 | 0.745810015 | 0.843013799 | [illegible] |
| PS | PS 40:5\|PS 18:0_22:5 | [M-H]- | 13.779 | 836.54181 | 0.657169087 | 0.633872818 | 0.659163977 | [illegible] |
| PS | PS 40:7 | [M-H]- | 13.701 | 832.51044 | 0.027161947 | 0.027927788 | 0.030797114 | [illegible] |
| PS | PS 40:7\|PS 18:1_22:6 | [M-H]- | 12.926 | 832.51044 | 0.037986458 | 0.034259603 | 0.035288439 | [illegible] |
| PS | PS 42:8 | [M-H]- | 13.791 | 858.52448 | 0.045811869 | 0.04860662 | 0.052203478 | [illegible] |
| PS | PS 42:9 | [M-H]- | 13.537 | 856.51001 | 0.065097504 | 0.062444311 | 0.068355793 | [illegible] |
| PI | PI 34:0\|PI 16:0_18:0 | [M-H]- | 293916224 | 227917424 | 104.5692822 | 101.7927681 | 102.1974132 | [illegible] |
| PI | PI 34:2\|PI 16:1_18:1 | [M-H]- | 165963024 | 165311040 | 0.154835043 | 0.122160121 | 0.128304962 | [illegible] |
| PI | PI 35:2\|PI 17:1_18:1 | [M-H]- | 34683464 | 33646544 | 0.087429307 | 0.088604093 | 0.089545298 | [illegible] |
| PI | PI 36:1\|PI 18:0_18:1 | [M-H]- | 384424832 | 479974560 | 0.018271246 | 0.018034013 | 0.018759589 | [illegible] |
| PI | PI 36:2\|PI 18:1_18:1 | [M-H]- | 577641472 | 530187104 | 0.20251497 | 0.257258745 | 0.238299262 | [illegible] |
| PI | PI 36:3\|PI 18:1_18:2 | [M-H]- | 152182464 | 143389392 | 0.30430148 | 0.284171871 | 0.306798846 | [illegible] |
| PI | PI 36:4\|PI 16:0_20:4 | [M-H]- | 217278400 | 211009760 | 0.08016971 | 0.076854438 | 0.081003718 | [illegible] |
| PI | PI 36:5\|PI 16:1_20:4 | [M-H]- | 19196466 | 18844914 | 0.114462243 | 0.113097882 | 0.113181213 | [illegible] |
| PI | PI 37:3\|PI 18:0_19:3 | [M-H]- | 14350329 | 16104369 | 0.010112697 | 0.010100575 | 0.009966326 | [illegible] |
| PI | PI 37:4\|PI 17:0_20:4 | [M-H]- | 75698440 | 71395248 | 0.007559752 | 0.008631686 | 0.006764486 | [illegible] |
| PI | PI 38:3\|PI 18:0_20:3 | [M-H]- | 443932384 | 453770304 | 0.039877932 | 0.03826672 | 0.04199675 | [illegible] |
| PI | PI 38:4\|PI 18:0_20:4 | [M-H]- | 5508634624 | 4141385216 | 0.233863543 | 0.24321368 | 0.229816891 | [illegible] |
| PI | PI 38:4\|PI 18:0_20:4 | [M-H]- | 1896793 | 2013832 | 2.901948267 | 2.219716734 | 2.358686178 | [illegible] |
| PI | PI 38:5\|PI 18:1_20:4 | [M-H]- | 827247744 | 760406848 | 0.00099923 | 0.001079382 | 0.001144179 | [illegible] |
| PI | PI 38:6\|PI 16:0_22:6 | [M-H]- | 64191196 | 63929812 | 0.435794044 | 0.407566 | 0.426838671 | [illegible] |
| PI | PI 38:6\|PI 18:2_20:4 | [M-H]- | 63513772 | 63381404 | 0.033815917 | 0.034265364 | 0.033576764 | [illegible] |
| PI | PI 38:7 | [M-H]- | 2727060 | 2501468 | 0.03345905 | 0.033971426 | 0.033205427 | [illegible] |
| PI | PI 39:3\|PI 18:0_21:3 | [M-H]- | 28052842 | 28702546 | 0.001436615 | 0.001340747 | 0.001500106 | [illegible] |
| PI | PI 39:4\|PI 19:0_20:4 | [M-H]- | 39958960 | 25764044 | 0.014778235 | 0.015384109 | 0.014941924 | [illegible] |
| PI | PI 39:5\|PI 19:1_20:4 | [M-H]- | 37166268 | 35318840 | 0.021050377 | 0.013809119 | 0.016441361 | [illegible] |
| PI | PI 39:6 | [M-H]- | 5948214 | 5378836 | 0.019579187 | 0.018930338 | 0.019690227 | [illegible] |
| PI | PI 40:3\|PI 18:0_22:3 | [M-H]- | 104148416 | 71174368 | 0.003133519 | 0.002882971 | 0.002947522 | [illegible] |
| PI | PI 40:4\|PI 18:0_22:4 | [M-H]- | 203170832 | 199275008 | 0.054865377 | 0.038148331 | 0.039317443 | [illegible] |

| | | | | | | | | |
|---|---|---|---|---|---|---|---|---|
| PI | PI 40:5\|PI 18:0_22:5 | [M-H]- | 316873504 | 213764160 | 0.107030378 | 0.106808241 | 0.114148512 | [illegible] |
| PI | PI 40:6\|PI 18:0_22:6 | [M-H]- | 163303792 | 153511344 | 0.166928936 | 0.114574196 | 0.11417516 | [illegible] |
| PI | PI 40:6\|PI 18:1_22:5 | [M-H]- | 57680864 | 55403248 | 0.086028424 | 0.082279643 | 0.065945709 | [illegible] |
| PI | PI 40:7\|PI 18:2_22:5 | [M-H]- | 55235360 | 53161152 | 0.030386274 | 0.029695261 | 0.028566835 | [illegible] |
| PI | PI 40:8\|PI 20:4_20:4 | [M-H]- | 10125671 | 9450302 | 0.029097983 | 0.028493534 | 0.028984997 | [illegible] |
| PI | PI 42:9\|PI 20:4_22:5 | [M-H]- | 4396988 | 3901534 | 0.005334203 | 0.005065212 | 0.005894589 | [illegible] |
| PI | PI O-38:4\|PI O-18:0_20:4 | [M-H]- | 1483880 | 1769772 | 0.002316333 | 0.00209116 | 0.00251262 | [illegible] |
| PI | PI O-38:5\|PI O-18:1_20:4 | [M-H]- | 3381915 | 2851579 | 0.000781708 | 0.00094857 | 0.001004875 | [illegible] |
| LPC | LPC 18:0/0:0 | [M+H]+ | 11.433 | 524.37054 | 0.001781593 | 0.001528401 | 0.00186528 | [illegible] |
| LPC | LPC 26:0/0:0 | [M+H]+ | 15.099 | 636.49585 | 0.105750469 | 0.108126024 | 0.100107252 | [illegible] |
| LPE | LPE 16:0 | [M-H]- | 10.352 | 452.27823 | 0.050020189 | 0.045478812 | 0.052564751 | [illegible] |
| LPE | LPE 17:0 | [M-H]- | 10.874 | 466.29373 | 0.023225043 | 0.023617196 | 0.022263545 | [illegible] |
| LPE | LPE 18:0 | [M-H]- | 11.302 | 480.30902 | 0.003518646 | 0.003132442 | 0.004155221 | [illegible] |
| LPE | LPE 18:1 | [M-H]- | 10.621 | 478.2937 | 0.247259921 | 0.257276171 | 0.269321316 | [illegible] |
| LPE | LPE 20:1 | [M-H]- | 11.362 | 506.32489 | 0.066505018 | 0.076015053 | 0.069843751 | [illegible] |
| LPE | LPE 22:0 | [M-H]- | 12.738 | 536.37219 | 0.005229037 | 0.005086201 | 0.005653733 | [illegible] |
| LPE | LPE 22:5 | [M-H]- | 10.177 | 526.29321 | 0.000977215 | 0.001067354 | 0.000836542 | [illegible] |
| LPE | LPE 24:4 | [M-H]- | 11.329 | 556.34064 | 0.036523937 | 0.033004157 | 0.028264949 | [illegible] |
| LPE | LPE 24:5 | [M-H]- | 11.013 | 554.32483 | 0.003122317 | 0.003251619 | 0.002905721 | [illegible] |
| LPE | LPE O-16:1 | [M-H]- | 10.771 | 436.2821 | 0.004418597 | 0.003781159 | 0.003119428 | [illegible] |
| LPE | LPE O-17:1 | [M-H]- | 11.211 | 450.29803 | 0.021831297 | 0.022276592 | 0.022572074 | [illegible] |
| LPE | LPE O-18:1 | [M-H]- | 17.374 | 464.31351 | 0.008737736 | 0.008455706 | 0.007927357 | [illegible] |
| LPE | LPE O-18:1 | [M-H]- | 11.615 | 464.31354 | 0.005440925 | 0.005577224 | 0.005640255 | [illegible] |
| LPE | LPE O-18:2 | [M-H]- | 11 | 462.29825 | 0.041103898 | 0.047531112 | 0.040858889 | [illegible] |
| LPE | LPE-N (FA)30:0\|LPE-N (FA 16:0)14:0 | [M-H]- | 13.821 | 662.47552 | 0.016314315 | 0.018940718 | 0.019013157 | [illegible] |
| LPE | LPE-N (FA)35:3\|LPE-N (FA 17:1)18:2 | [M-H]- | 14.624 | 726.50562 | 0.005651851 | 0.005794452 | 0.005149895 | [illegible] |
| TG | TG 40:0\|TG 10:0_14:0_16:0 | [M+NH4]+ | 21.069 | 712.64557 | 0.005952594 | 0.005841002 | 0.005797913 | [illegible] |
| TG | TG 42:0\|TG 12:0_14:0_16:0 | [M+NH4]+ | 21.907 | 740.67706 | 0.002227974 | 0.003570118 | 0.002244417 | [illegible] |
| TG | TG 44:0\|TG 12:0_14:0_18:0 | [M+NH4]+ | 22.667 | 768.70734 | 0.00245777 | 0.003877217 | 0.003207256 | [illegible] |
| TG | TG 44:1\|TG 12:0_16:0_16:1 | [M+NH4]+ | 22.008 | 766.69177 | 0.005078466 | 0.006065813 | 0.0068584 | [illegible] |
| TG | TG 45:0\|TG 14:0_15:0_16:0 | [M+NH4]+ | 23.019 | 782.72321 | 0.002683496 | 0.002419088 | 0.004909419 | [illegible] |
| TG | TG 45:1\|TG 14:0_15:0_16:1 | [M+NH4]+ | 22.341 | 780.70776 | 0.010326714 | 0.010187805 | 0.012486329 | [illegible] |
| TG | TG 46:1\|TG 14:0_16:0_16:1 | [M+NH4]+ | 22.752 | 794.724 | 0.002998073 | 0.002476722 | 0.006612364 | [illegible] |
| TG | TG 46:2\|TG 14:0_14:1_18:1 | [M+NH4]+ | 22.149 | 792.70703 | 0.010549945 | 0.008876673 | 0.015807623 | [illegible] |
| TG | TG 47:0\|TG 14:0_16:0_17:0 | [M+NH4]+ | 23.707 | 810.75452 | 0.004361187 | 0.003829588 | 0.007404441 | [illegible] |
| TG | TG 47:1\|TG 15:0_16:0_16:1 | [M+NH4]+ | 23.088 | 808.73773 | 0.007437217 | 0.007991419 | 0.01044089 | [illegible] |
| TG | TG 47:2\|TG 15:0_15:1_17:1 | [M+NH4]+ | 22.513 | 806.72278 | 0.007455696 | 0.006670987 | 0.013889227 | [illegible] |
| TG | TG 48:0\|TG 14:0_16:0_18:0 | [M+NH4]+ | 24.052 | 824.76971 | 0.004577096 | 0.003747803 | 0.008516151 | [illegible] |
| TG | TG 48:1\|TG 14:0_16:0_18:1 | [M+NH4]+ | 23.437 | 822.75592 | 0.023549952 | 0.026600953 | 0.025179447 | [illegible] |
| TG | TG 48:2\|TG 14:0_16:1_18:1 | [M+NH4]+ | 22.849 | 820.73822 | 0.031091463 | 0.029708791 | 0.037240732 | [illegible] |
| TG | TG 48:3\|TG 14:0_16:1_18:2 | [M+NH4]+ | 22.302 | 818.72229 | 0.021694249 | 0.019643891 | 0.02831169 | [illegible] |
| TG | TG 49:0\|TG 15:0_16:0_18:0 | [M+NH4]+ | 24.344 | 838.78705 | 0.006501597 | 0.005876536 | 0.008630948 | [illegible] |
| TG | TG 49:1\|TG 15:0_16:0_18:1 | [M+NH4]+ | 23.766 | 836.76929 | 0.007590823 | 0.008122808 | 0.009181895 | [illegible] |
| TG | TG 49:2\|TG 16:0_16:1_17:1 | [M+NH4]+ | 23.371 | 834.75378 | 0.015210449 | 0.014298839 | 0.02212947 | [illegible] |
| TG | TG 50:0\|TG 16:0_16:0_18:0 | [M+NH4]+ | 24.67 | 852.80017 | 0.23454263 | 0.228553673 | 0.259976765 | [illegible] |
| TG | TG 50:1\|TG 16:0_16:0_18:1 | [M+NH4]+ | 24.083 | 850.78516 | 0.026672305 | 0.033269306 | 0.02636058 | [illegible] |
| TG | TG 50:2\|TG 16:0_16:1_18:1 | [M+NH4]+ | 23.517 | 848.77094 | 0.092047359 | 0.093374713 | 0.091616704 | [illegible] |
| TG | TG 50:3\|TG 16:1_16:1_18:1 | [M+NH4]+ | 22.949 | 846.75293 | 0.07561022 | 0.071557359 | 0.078681109 | [illegible] |
| TG | TG 50:4\|TG 16:0_18:1_16:3 | [M+NH4]+ | 22.692 | 844.73895 | 0.031539969 | 0.030119549 | 0.035331993 | [illegible] |
| TG | TG 50:5\|TG 14:0_16:1_20:4 | [M+NH4]+ | 22.103 | 842.72272 | 0.008328722 | 0.006996409 | 0.008915111 | [illegible] |
| TG | TG 51:0\|TG 16:0_17:0_18:0 | [M+NH4]+ | 24.949 | 866.81805 | 0.004960496 | 0.004319372 | 0.005386702 | [illegible] |
| TG | TG 51:1\|TG 16:0_17:0_18:1 | [M+NH4]+ | 24.387 | 864.8009 | 0.004858771 | 0.00559365 | 0.005712828 | [illegible] |
| TG | TG 51:2\|TG 16:0_17:1_18:1 | [M+NH4]+ | 23.817 | 862.78381 | 0.021102291 | 0.021526196 | 0.023299949 | [illegible] |
| TG | TG 51:3\|TG 16:1_17:1_18:1 | [M+NH4]+ | 23.261 | 860.77045 | 0.02785364 | 0.025923473 | 0.032997663 | [illegible] |
| TG | TG 52:0\|TG 16:0_18:0_18:0 | [M+NH4]+ | 25.262 | 880.83002 | 0.012709436 | 0.01133494 | 0.014229858 | [illegible] |
| TG | TG 52:1\|TG 16:0_18:0_18:1 | [M+NH4]+ | 24.69 | 878.8161 | 0.01682512 | 0.02094581 | 0.015846921 | [illegible] |
| TG | TG 52:2\|TG 16:0_18:1_18:1 | [M+NH4]+ | 24.12 | 876.80212 | 0.09276791 | 0.09909403 | 0.088243722 | [illegible] |
| TG | TG 52:3\|TG 16:0_18:1_18:2 | [M+NH4]+ | 23.597 | 874.78467 | 0.166751463 | 0.1611734 | 0.166602327 | [illegible] |
| TG | TG 52:4\|TG 16:0_16:0_20:4 | [M+NH4]+ | 23.378 | 872.76807 | 0.094869782 | 0.08815915 | 0.101657021 | [illegible] |
| TG | TG 52:4\|TG 16:1_18:1_18:2 | [M+NH4]+ | 23.082 | 872.77063 | 0.016378921 | 0.01551881 | 0.017714009 | [illegible] |
| TG | TG 52:5\|TG 16:0_16:1_20:4 | [M+NH4]+ | 22.753 | 870.75311 | 0.049983747 | 0.045574128 | 0.054245794 | [illegible] |
| TG | TG 52:6\|TG 14:0_16:1_22:5 | [M+NH4]+ | 22.228 | 868.73901 | 0.02916708 | 0.024954717 | 0.031994775 | [illegible] |
| TG | TG 52:7\|TG 14:0_16:1_22:6 | [M+NH4]+ | 21.773 | 866.72229 | 0.01139361 | 0.010081677 | 0.012927129 | [illegible] |
| TG | TG 53:0\|TG 16:0_18:0_19:0 | [M+NH4]+ | 25.502 | 894.84631 | 0.003185774 | 0.002609685 | 0.003640567 | [illegible] |
| TG | TG 53:1\|TG 17:0_18:0_18:1 | [M+NH4]+ | 24.975 | 892.83099 | 0.002116496 | 0.00221153 | 0.003312713 | [illegible] |
| TG | TG 53:2\|TG 16:0_18:1_19:1 | [M+NH4]+ | 24.42 | 890.81506 | 0.009306406 | 0.009629946 | 0.011086069 | [illegible] |
| TG | TG 53:3\|TG 17:1_18:1_18:1 | [M+NH4]+ | 23.887 | 888.80237 | 0.02393567 | 0.023026564 | 0.024712856 | [illegible] |
| TG | TG 53:5\|TG 16:0_17:1_20:4 | [M+NH4]+ | 23.098 | 884.76904 | 0.020772701 | 0.019250046 | 0.022804274 | [illegible] |
| TG | TG 54:0\|TG 16:0_18:0_20:0 | [M+NH4]+ | 25.792 | 908.86151 | 0.007960805 | 0.006723393 | 0.008759441 | [illegible] |
| TG | TG 54:1\|TG 16:0_18:0_20:1 | [M+NH4]+ | 25.276 | 906.84644 | 0.004577566 | 0.00809949 | 0.005917349 | [illegible] |
| TG | TG 54:2\|TG 16:0_18:1_20:1 | [M+NH4]+ | 24.729 | 904.83044 | 0.029407317 | 0.036906223 | 0.027691378 | [illegible] |
| TG | TG 54:3\|TG 18:1_18:1_18:1 | [M+NH4]+ | 24.184 | 902.81744 | 0.079737104 | 0.081116857 | 0.076830299 | [illegible] |
| TG | TG 54:4\|TG 18:1_18:1_18:2 | [M+NH4]+ | 23.722 | 900.80029 | 0.118391615 | 0.111319134 | 0.120332969 | [illegible] |
| TG | TG 54:5\|TG 16:0_18:1_20:4 | [M+NH4]+ | 23.431 | 898.78339 | 0.079634414 | 0.071419664 | 0.085746614 | [illegible] |
| TG | TG 54:6\|TG 16:1_18:1_20:4 | [M+NH4]+ | 22.895 | 896.76801 | 0.085568633 | 0.075965966 | 0.092977165 | [illegible] |
| TG | TG 54:7\|TG 16:0_16:1_22:6 | [M+NH4]+ | 22.51 | 894.75281 | 0.049034024 | 0.039195867 | 0.054591544 | [illegible] |
| TG | TG 54:8\|TG 16:1_16:1_22:6 | [M+NH4]+ | 21.942 | 892.73749 | 0.025808574 | 0.020841907 | 0.028960528 | [illegible] |
| TG | TG 55:0\|TG 15:0_16:0_24:0 | [M+NH4]+ | 26.01 | 922.87787 | 0.006481598 | 0.004582841 | 0.006675467 | [illegible] |
| TG | TG 55:1\|TG 18:0_19:0_18:1 | [M+NH4]+ | 25.542 | 920.86432 | 0.001663313 | 0.001867091 | 0.003333168 | [illegible] |
| TG | TG 55:2\|TG 18:0_18:1_19:1 | [M+NH4]+ | 24.992 | 918.8465 | 0.002784897 | 0.002520006 | 0.005582979 | [illegible] |
| TG | TG 55:3\|TG 18:1_18:1_19:1 | [M+NH4]+ | 24.475 | 916.83124 | 0.005504537 | 0.005251345 | 0.006680883 | [illegible] |
| TG | TG 55:4\|TG 16:0_18:1_21:3 | [M+NH4]+ | 24.062 | 914.81525 | 0.008954396 | 0.008537041 | 0.009798197 | [illegible] |
| TG | TG 55:5\|TG 17:0_18:1_20:4 | [M+NH4]+ | 23.735 | 912.802 | 0.010965997 | 0.01074454 | 0.011335392 | [illegible] |
| TG | TG 55:6\|TG 17:1_18:1_20:4 | [M+NH4]+ | 23.145 | 910.78448 | 0.013236772 | 0.00530847 | 0.006517608 | [illegible] |
| TG | TG 55:7\|TG 16:0_17:1_22:6 | [M+NH4]+ | 22.863 | 908.76996 | 0.009974724 | 0.00941475 | 0.013377487 | [illegible] |
| TG | TG 56:0\|TG 15:0_16:0_25:0 | [M+NH4]+ | 26.294 | 936.89343 | 0.004446991 | 0.00368527 | 0.005535636 | [illegible] |
| TG | TG 56:1\|TG 18:0_20:0_18:1 | [M+NH4]+ | 25.803 | 934.88025 | 0.002390311 | 0.002924171 | 0.004000813 | [illegible] |
| TG | TG 56:2\|TG 18:0_18:1_20:1 | [M+NH4]+ | 25.287 | 932.86188 | 0.007156811 | 0.007767672 | 0.00950724 | [illegible] |
| TG | TG 56:3\|TG 18:1_18:1_20:1 | [M+NH4]+ | 24.778 | 930.84601 | 0.011521403 | 0.011454663 | 0.012683936 | [illegible] |
| TG | TG 56:4\|TG 18:1_18:1_20:2 | [M+NH4]+ | 24.381 | 928.83264 | 0.021175572 | 0.020049394 | 0.021188089 | [illegible] |
| TG | TG 56:5\|TG 18:0_18:1_20:4 | [M+NH4]+ | 23.954 | 926.81482 | 0.045462621 | 0.041520872 | 0.044533277 | [illegible] |
| TG | TG 56:5\|TG 18:1_18:1_20:3 | [M+NH4]+ | 23.91 | 926.815 | 0.082512736 | 0.073982365 | 0.088210324 | [illegible] |
| TG | TG 56:6\|TG 16:0_18:0_22:6 | [M+NH4]+ | 23.831 | 924.79993 | 0.043276676 | 0.040622341 | 0.051649193 | [illegible] |
| TG | TG 56:6\|TG 16:0_18:1_22:5 | [M+NH4]+ | 23.465 | 924.80054 | 0.014880111 | 0.012409648 | 0.014684818 | [illegible] |
| TG | TG 56:7\|TG 16:0_18:1_22:6 | [M+NH4]+ | 23.201 | 922.78394 | 0.088036604 | 0.073228098 | 0.092916409 | [illegible] |
| TG | TG 56:7\|TG 16:1_18:1_22:5 | [M+NH4]+ | 22.953 | 922.78394 | 0.04782274 | 0.041224431 | 0.052996438 | [illegible] |
| TG | TG 56:8\|TG 16:1_18:1_22:6 | [M+NH4]+ | 22.641 | 920.77075 | 0.030788483 | 0.027769922 | 0.036979849 | [illegible] |
| TG | TG 56:9\|TG 16:0_20:4_20:5 | [M+NH4]+ | 22.199 | 918.75293 | 0.035502375 | 0.027671676 | 0.04052043 | [illegible] |
| TG | TG 56:9\|TG 16:1_18:2_22:6 | [M+NH4]+ | 22.046 | 918.75293 | 0.005870687 | 0.004630945 | 0.007078512 | [illegible] |
| TG | TG 57:0\|TG 16:0_17:0_24:0 | [M+NH4]+ | 26.482 | 950.91187 | 0.004533418 | 0.00379074 | 0.005517624 | [illegible] |
| TG | TG 57:1\|TG 16:0_25:0_16:1 | [M+NH4]+ | 26.062 | 948.89618 | 0.001305394 | 0.001522392 | 0.002529786 | [illegible] |
| TG | TG 57:2\|TG 23:0_16:1_18:1 | [M+NH4]+ | 25.555 | 946.87909 | 0.002064074 | 0.00178316 | 0.004794066 | [illegible] |
| TG | TG 57:5\|TG 17:0_18:1_22:4 | [M+NH4]+ | 24.206 | 940.8316 | 0.002234146 | 0.002115425 | 0.003942719 | [illegible] |
| TG | TG 58:0\|TG 16:0_18:0_24:0 | [M+NH4]+ | 26.75 | 964.92542 | 0.007662156 | 0.006836366 | 0.007952584 | [illegible] |
| TG | TG 58:1\|TG 16:0_18:0_24:1 | [M+NH4]+ | 26.298 | 962.90942 | 0.00170998 | 0.002126696 | 0.002202652 | [illegible] |
| TG | TG 58:10\|TG 16:0_20:4_22:6 | [M+NH4]+ | 22.486 | 944.77118 | 0.00546697 | 0.005657743 | 0.006914595 | [illegible] |
| TG | TG 58:10\|TG 16:1_20:4_22:5 | [M+NH4]+ | 22.24 | 944.77118 | 0.008226425 | 0.006327939 | 0.009575907 | [illegible] |
| TG | TG 58:2\|TG 16:0_18:1_24:1 | [M+NH4]+ | 25.801 | 960.89551 | 0.008849833 | 0.007246165 | 0.010682918 | [illegible] |
| TG | TG 58:3\|TG 16:0_18:1_24:2 | [M+NH4]+ | 25.334 | 958.87994 | 0.008315438 | 0.007981278 | 0.009677682 | [illegible] |
| TG | TG 58:4\|TG 16:0_18:1_24:3 | [M+NH4]+ | 24.936 | 956.86273 | 0.006279651 | 0.006394402 | 0.006706005 | [illegible] |
| TG | TG 58:5\|TG 18:0_18:1_22:4 | [M+NH4]+ | 24.518 | 954.84778 | 0.010573055 | 0.009737373 | 0.010580384 | [illegible] |
| TG | TG 58:6\|TG 18:0_18:1_22:5 | [M+NH4]+ | 24.052 | 952.83124 | 0.029580476 | 0.027566004 | 0.029477229 | [illegible] |
| TG | TG 58:7\|TG 18:0_18:1_22:6 | [M+NH4]+ | 23.865 | 950.81757 | 0.055944942 | 0.049425823 | 0.0580289 | [illegible] |
| TG | TG 58:7\|TG 18:1_18:1_22:5 | [M+NH4]+ | 23.549 | 950.81757 | 0.020293958 | 0.017520906 | 0.020918243 | [illegible] |
| TG | TG 58:8\|TG 18:1_18:1_22:6 | [M+NH4]+ | 23.243 | 948.80219 | 0.044517598 | 0.037650116 | 0.047395704 | [illegible] |
| TG | TG 58:9\|TG 18:1_18:2_22:6 | [M+NH4]+ | 22.752 | 946.7854 | 0.055798158 | 0.044816305 | 0.06108988 | [illegible] |
| TG | TG 59:1\|TG 16:0_25:0_18:1 | [M+NH4]+ | 26.511 | 976.92743 | 0.035162526 | 0.027403464 | 0.03992566 | [illegible] |
| TG | TG 59:2\|TG 16:0_18:1_25:1 | [M+NH4]+ | 26.055 | 974.91022 | 0.001432585 | 0.00143702 | 0.002830201 | [illegible] |
| TG | TG 60:0\|TG 16:0_18:0_26:0 | [M+NH4]+ | 27.176 | 992.95709 | 0.00206429 | 0.001993453 | 0.003432559 | [illegible] |
| TG | TG 60:1\|TG 16:0_18:0_26:1 | [M+NH4]+ | 26.756 | 990.94031 | 0.000782308 | 0.001136232 | 0.001078122 | [illegible] |
| TG | TG 60:10\|TG 16:0_22:4_22:6 | [M+NH4]+ | 22.817 | 972.8017 | 0.00401166 | 0.0045043 | 0.004533689 | [illegible] |
| TG | TG 60:11\|TG 18:1_20:4_22:6 | [M+NH4]+ | 22.523 | 970.7851 | 0.024216425 | 0.012676803 | 0.025954423 | [illegible] |
| TG | TG 60:11\|TG 18:2_20:4_22:5 | [M+NH4]+ | 22.523 | 970.78485 | 0.014368819 | 0.014435809 | 0.015377444 | [illegible] |
| TG | TG 60:12\|TG 16:0_22:6_22:6 | [M+NH4]+ | 22.241 | 968.77069 | 0.016509247 | 0.013375128 | 0.018957446 | [illegible] |
| TG | TG 60:12\|TG 16:1_22:5_22:6 | [M+NH4]+ | 21.969 | 968.77069 | 0.003442794 | 0.002702086 | 0.003789142 | [illegible] |
| TG | TG 60:2\|TG 16:0_18:1_26:1 | [M+NH4]+ | 26.287 | 988.92548 | 0.00409839 | 0.003204284 | 0.00237562 | [illegible] |
| TG | TG 60:3\|TG 16:0_18:1_26:2 | [M+NH4]+ | 25.823 | 986.90851 | 0.007335093 | 0.007816186 | 0.008164282 | [illegible] |
| TG | TG 60:4\|TG 18:1_24:1_18:2 | [M+NH4]+ | 25.422 | 984.89349 | 0.006268937 | 0.006293739 | 0.006615683 | [illegible] |
| TG | TG 60:5\|TG 16:0_18:1_26:4 | [M+NH4]+ | 24.984 | 982.88055 | 0.004798561 | 0.005385191 | 0.005462759 | [illegible] |
| TG | TG 60:6\|TG 16:0_18:1_26:5 | [M+NH4]+ | 24.566 | 980.86139 | 0.00842672 | 0.007709924 | 0.006581389 | [illegible] |
| TG | TG 60:7\|TG 20:1_18:2_22:4 | [M+NH4]+ | 24.11 | 978.84845 | 0.016793225 | 0.015734208 | 0.016766805 | [illegible] |
| TG | TG 60:8\|TG 18:1_18:1_24:6 | [M+NH4]+ | 23.689 | 976.8335 | 0.023939088 | 0.022091867 | 0.024859375 | [illegible] |
| TG | TG 61:2\|TG 16:0_18:1_27:1 | [M+NH4]+ | 26.51 | 1002.94183 | 0.026337391 | 0.022277861 | 0.028119912 | [illegible] |
| TG | TG 61:3\|TG 17:1_18:1_26:1 | [M+NH4]+ | 26.055 | 1000.92456 | 0.001300188 | 0.001273519 | 0.001958859 | [illegible] |
| TG | TG 62:1\|TG 18:0_26:0_18:1 | [M+NH4]+ | 27.175 | 1018.97229 | 0.001143957 | 0.001139208 | 0.001520521 | [illegible] |
| TG | TG 62:11\|TG 18:1_22:4_22:6 | [M+NH4]+ | 23.043 | 998.81555 | 0.001248002 | 0.001477007 | 0.001586306 | [illegible] |
| TG | TG 62:12\|TG 18:1_22:5_22:6 | [M+NH4]+ | 22.562 | 996.802 | 0.013665359 | 0.011059958 | 0.011180235 | [illegible] |
| TG | TG 62:13\|TG 18:1_22:6_22:6 | [M+NH4]+ | 22.299 | 994.78613 | 0.008786998 | 0.006858998 | 0.010021646 | [illegible] |
| TG | TG 62:2\|TG 18:0_18:1_26:1 | [M+NH4]+ | 26.75 | 1016.95844 | 0.002509374 | 0.002013162 | 0.002981132 | [illegible] |
| TG | TG 62:3\|TG 18:1_18:1_26:1 | [M+NH4]+ | 26.284 | 1014.94189 | 0.003348936 | 0.003605957 | 0.003569957 | [illegible] |
| TG | TG 62:4\|TG 18:1_26:1_18:2 | [M+NH4]+ | 25.883 | 1012.92706 | 0.00416553 | 0.00434281 | 0.004350189 | [illegible] |
| TG | TG 62:5\|TG 18:1_24:1_20:3 | [M+NH4]+ | 25.487 | 1010.90955 | 0.003916608 | 0.003943992 | 0.004068524 | [illegible] |

| | | | | | | | | | | | | | | | | | | | | | |
|---|---|---|---|---|---|---|---|---|---|---|---|---|---|---|---|---|---|---|---|---|---|
| TG | TG 62:6\|TG 16:0_18:1_28:5 | [M+NH4]+ | 25.04 | 1008.89374 | 0.003362194 | 0.003253289 | 0.00331857 | 0.004444657 | 0.004081587 | 0.004718971 | 3.2153E-05 | 3.196E-05 | 3.2472E-05 | 4.7074E-05 | 5.3188E-05 | 6.2632E-05 | 3.2195E-05 | 2.1123E-07 | 5.4298E-05 | 6.39981E-06 | 0.03932327 | 0.75406427 |
| TG | TG 62:7\|TG 18:0_18:1_26:6 | [M+NH4]+ | 24.647 | 1006.8797 | 0.003152134 | 0.002877622 | 0.003118761 | 0.005798728 | 0.00388488 | 0.004561994 | 3.0144E-05 | 2.8269E-05 | 3.0517E-05 | 6.1415E-05 | 5.0625E-05 | 6.0548E-05 | 2.96435E-05 | 9.83469E-07 | 5.75294E-05 | 4.89495E-06 | 0.01246086 | 0.956585666 |
| TG | TG 62:8\|TG 18:1_18:1_26:6 | [M+NH4]+ | 24.224 | 1004.8631 | 0.008449511 | 0.007389455 | 0.008662737 | 0.013786475 | 0.010208689 | 0.011535036 | 8.0803E-05 | 7.2593E-05 | 8.4765E-05 | 0.00014601 | 0.00013303 | 0.0001531 | 7.9387E-05 | 5.06892E-06 | 0.000144048 | 8.30844E-06 | 0.001696 | 0.859573616 |
| TG | TG 62:9\|TG 18:1_20:4_24:4 | [M+NH4]+ | 23.835 | 1002.84821 | 0.010384 | 0.009416377 | 0.010605979 | 0.021654096 | 0.017479186 | 0.020065462 | 9.9303E-05 | 9.2505E-05 | 0.00010378 | 0.00022934 | 0.00022778 | 0.00026632 | 9.85291E-05 | 4.63496E-06 | 0.000241144 | 1.78103E-05 | 0.00518039 | 1.291274906 |
| TG | TG O-52:1\|TG O-18:0_16:0_18:1 | [M+NH4]+ | 25.625 | 864.83844 | 0.012851617 | 0.011625387 | 0.007091406 | 0.036263739 | 0.029888951 | 0.032448161 | 0.0001229 | 0.00011421 | 6.9389E-05 | 0.00038407 | 0.00038949 | 0.00043066 | 0.000102165 | 2.34464E-05 | 0.000401409 | 2.08036E-05 | 0.00018984 | 1.974167198 |
| DG | DG 34:0\|DG 16:0_18:0 | [M+NH4]+ | 18.631 | 614.57062 | 0.00184472 | 0.002090295 | 0.00185867 | 0.003243716 | 0.002722216 | 0.00275094 | 1.7641E-05 | 2.0535E-05 | 1.8187E-05 | 3.4355E-05 | 3.5474E-05 | 3.6511E-05 | 1.87877E-05 | 1.25536E-06 | 3.54466E-05 | 8.8077E-07 | 0.00021163 | 0.915863389 |
| DG | DG 34:1\|DG 16:0_18:1 | [M+NH4]+ | 17.687 | 612.55554 | 0.043681306 | 0.054266026 | 0.037824395 | 0.07105302 | 0.040943991 | 0.035614055 | 0.00041773 | 0.0005331 | 0.00037011 | 0.00075253 | 0.00053355 | 0.00047268 | 0.000440313 | 6.84311E-05 | 0.000586255 | 0.000120171 | 0.22756351 | 0.412997649 |
| DG | DG 36:0\|DG 18:0_18:0 | [M+NH4]+ | 19.658 | 642.60388 | 0.174515126 | 0.190076653 | 0.160951443 | 0.129074605 | 0.09448081 | 0.095944905 | 0.00166889 | 0.00186729 | 0.00157491 | 0.00136704 | 0.00123121 | 0.00127341 | 0.001703697 | 0.000121875 | 0.001290554 | 5.67632E-05 | 0.0253518 | -0.40067819 |
| DG | DG 36:2\|DG 18:1_18:1 | [M+NH4]+ | 17.819 | 638.57227 | 0.01551587 | 0.021079437 | 0.018566571 | 0.048146178 | 0.030140721 | 0.020355876 | 0.00014838 | 0.00020708 | 0.00018167 | 0.00050992 | 0.00039277 | 0.00027017 | 0.000179045 | 2.40374E-05 | 0.000390955 | 9.78863E-05 | 0.08456597 | 1.126680708 |
| DG | DG 38:4\|DG 18:0_20:4 | [M+NH4]+ | 17.816 | 662.57208 | 0.08822694 | 0.092982123 | 0.087072392 | 0.073922881 | 0.049502988 | 0.05714184 | 0.00084372 | 0.00091345 | 0.000852 | 0.00078292 | 0.00064509 | 0.00075841 | 0.000869722 | 3.11018E-05 | 0.000728806 | 6.00379E-05 | 0.06012171 | -0.25501824 |
| SM | SM 32:1;2O\|SM 18:1;2O/14:0 | [M+H]+ | 797582016 | 801507392 | 0.084048734 | 0.096962203 | 0.081443166 | 0.056794434 | 0.04054042 | 0.040964866 | 0.00080376 | 0.00095255 | 0.00079692 | 0.00060152 | 0.00052829 | 0.0005437 | 0.000851075 | 7.18042E-05 | 0.000557837 | 3.15195E-05 | 0.01647389 | -0.60944468 |
| SM | SM 33:1;2O | [M+H]+ | 396792416 | 421416224 | 0.27468402 | 0.268827368 | 0.219732597 | 0.160299328 | 0.119104629 | 0.124342387 | 0.00262681 | 0.00264093 | 0.00215008 | 0.00169775 | 0.00155209 | 0.00165031 | 0.002472607 | 0.000228134 | 0.001633383 | 6.0658E-05 | 0.02832185 | -0.59816999 |
| SM | SM 34:0;2O | [M+H]+ | 940455872 | 1043134336 | 0.136653703 | 0.141343942 | 0.11970221 | 0.09790311 | 0.075796981 | 0.078525946 | 0.00130682 | 0.00138855 | 0.00117128 | 0.0010369 | 0.00098773 | 0.00104222 | 0.001288885 | 8.95993E-05 | 0.001022286 | 2.4529E-05 | 0.04386686 | -0.33432462 |
| SM | SM 34:1;2O\|SM 18:1;2O/16:0 | [M+H]+ | 7833466880 | 8406764032 | 0.323889198 | 0.349869584 | 0.334301876 | 0.287367415 | 0.20106031 | 0.20349297 | 0.00309736 | 0.00343708 | 0.00327114 | 0.00304354 | 0.00262008 | 0.00270083 | 0.003268527 | 0.000138699 | 0.002788147 | 0.000183572 | 0.04577989 | -0.22933366 |
| SM | SM 34:2;2O\|SM 18:2;2O/16:0 | [M+H]+ | 869343744 | 859728256 | 2.697814306 | 2.819647413 | 2.435575496 | 2.350541904 | 1.581236667 | 1.647732198 | 0.0257993 | 0.02769988 | 0.02383207 | 0.02489482 | 0.02060557 | 0.02186927 | 0.025777082 | 0.001579106 | 0.022456552 | 0.001799649 | 0.12256174 | -0.19895254 |
| SM | SM 35:1;2O\|SM 18:1;2O/17:0 | [M+H]+ | 285905440 | 293396640 | 0.299398469 | 0.288354775 | 0.195277124 | 0.157617815 | 0.122181955 | 0.136715765 | 0.00286316 | 0.00283276 | 0.00191078 | 0.00166935 | 0.00159219 | 0.00181454 | 0.002535568 | 0.000441964 | 0.001692025 | 9.21793E-05 | 0.10858293 | -0.58355862 |
| SM | SM 36:0;2O | [M+H]+ | 134023328 | 153060032 | 0.098464677 | 0.098405888 | 0.085358621 | 0.078967182 | 0.056332173 | 0.055066707 | 0.00094162 | 0.00096673 | 0.00083523 | 0.00083635 | 0.00073408 | 0.00073086 | 0.000914527 | 5.69989E-05 | 0.000767098 | 4.89854E-05 | 0.05138701 | -0.25361478 |
| SM | SM 36:1;2O\|SM 18:1;2O/18:0 | [M+H]+ | 1030230144 | 1100627456 | 0.046157092 | 0.051336676 | 0.039671918 | 0.039592296 | 0.032426897 | 0.033575237 | 0.0004414 | 0.00050433 | 0.00038819 | 0.00041933 | 0.00042256 | 0.00044562 | 0.000444639 | 4.74677E-05 | 0.000429171 | 1.17077E-05 | 0.69411925 | -0.05108181 |
| SM | SM 37:1;3O | [M+H]+ | 151935104 | 140951184 | 0.354807094 | 0.369152904 | 0.305049037 | 0.291019402 | 0.208592661 | 0.20261081 | 0.00339303 | 0.00362651 | 0.0029849 | 0.00308222 | 0.00271823 | 0.00268912 | 0.003334816 | 0.000265153 | 0.002829856 | 0.00017884 | 0.09877183 | -0.23687835 |
| SM | SM 38:1;2O\|SM 18:1;2O/20:0 | [M+H]+ | 374023232 | 402618112 | 0.052325835 | 0.047275342 | 0.058781064 | 0.091310644 | 0.092781215 | 0.100175722 | 0.00050039 | 0.00046443 | 0.00057517 | 0.00096708 | 0.00120906 | 0.00132957 | 0.000513331 | 4.61274E-05 | 0.001168569 | 0.000150729 | 0.01835627 | 1.186781354 |
| SM | SM 40:0;2O | [M+H]+ | 152031424 | 167212992 | 0.128812088 | 0.135039013 | 0.117640377 | 0.134986241 | 0.087852896 | 0.091331316 | 0.00123183 | 0.00132661 | 0.00115111 | 0.00142965 | 0.00114484 | 0.00121218 | 0.001236517 | 7.17232E-05 | 0.001262223 | 0.000121541 | 0.81220793 | 0.029685245 |
| SM | SM 40:1;2O\|SM 18:1;2O/22:0 | [M+H]+ | 2008743424 | 2210295552 | 0.052359007 | 0.056083611 | 0.042886412 | 0.040811473 | 0.030744583 | 0.031393229 | 0.00050071 | 0.00055096 | 0.00041964 | 0.00043224 | 0.00040064 | 0.00041666 | 0.000490438 | 5.40994E-05 | 0.000416514 | 1.28996E-05 | 0.18781082 | -0.23570464 |
| SM | SM 40:2;2O\|SM 18:1;2O/22:1 | [M+H]+ | 231422400 | 254092880 | 0.691803109 | 0.74133806 | 0.598262409 | 0.529619521 | 0.399025246 | 0.405959965 | 0.00661574 | 0.00728282 | 0.00585399 | 0.00560925 | 0.00519982 | 0.00538804 | 0.006584181 | 0.000583744 | 0.005399037 | 0.000167332 | 0.09319409 | -0.28630206 |
| SM | SM 40:2;2O\|SM 18:1;2O/22:1 | [M+H]+ | 485980096 | 474499712 | 0.079700938 | 0.085223319 | 0.084875186 | 0.091152259 | 0.062423919 | 0.066877715 | 0.00076218 | 0.00083722 | 0.0008305 | 0.0009654 | 0.00081346 | 0.00088762 | 0.00080997 | 3.39015E-05 | 0.00088883 | 6.20343E-05 | 0.20995007 | 0.134040271 |
| SM | SM 41:2;2O | [M+H]+ | 416138816 | 418265152 | 0.167369579 | 0.159148262 | 0.146682419 | 0.141990738 | 0.106532386 | 0.105240182 | 0.00160056 | 0.00156345 | 0.00143529 | 0.00150384 | 0.00138826 | 0.00139678 | 0.0015331 | 7.08053E-05 | 0.001429626 | 5.25914E-05 | 0.17834832 | -0.10081441 |
| SM | SM 42:1;2O\|SM 18:1;2O/24:0 | [M+H]+ | 4044731136 | 4457455616 | 0.143316525 | 0.140287065 | 0.1282695 | 0.107832871 | 0.079137132 | 0.078453097 | 0.00137054 | 0.00137816 | 0.00125511 | 0.00114207 | 0.00103126 | 0.00104126 | 0.001334607 | 5.62951E-05 | 0.001071528 | 5.00463E-05 | 0.00809985 | -0.31674468 |
| SM | SM 42:2;2O\|SM 18:1;2O/24:1 | [M+H]+ | 4619487744 | 4880965120 | 1.392989042 | 1.495040559 | 1.14670482 | 0.904175517 | 0.700276567 | 0.738784218 | 0.01332121 | 0.0146871 | 0.01122049 | 0.00957621 | 0.00912551 | 0.0098054 | 0.013076265 | 0.001425797 | 0.009502374 | 0.00028243 | 0.06608387 | -0.46059057 |
| SM | SM 42:3;2O\|SM 18:1;2O/24:2 | [M+H]+ | 988393216 | 1018307136 | 1.590932892 | 1.637086591 | 1.482838541 | 1.183278849 | 0.854368176 | 0.8985605 | 0.01521415 | 0.01608254 | 0.01450955 | 0.01253222 | 0.01113353 | 0.011926 | 0.015268748 | 0.000643331 | 0.011863918 | 0.000572701 | 0.00522502 | -0.36400132 |
| SM | SM 43:1;2O | [M+H]+ | 109206744 | 128483032 | 0.340398625 | 0.341542485 | 0.315702859 | 0.258614867 | 0.192679742 | 0.193802897 | 0.00325524 | 0.00335527 | 0.00308915 | 0.00273902 | 0.00251087 | 0.00257222 | 0.003233222 | 0.000109756 | 0.002607367 | 9.64002E-05 | 0.00395042 | -0.31037866 |
| SM | SM 43:2;2O | [M+H]+ | 266686048 | 281607456 | 0.037610361 | 0.043093496 | 0.028520483 | 0.018996955 | 0.01723658 | 0.019061442 | 0.00035967 | 0.00042335 | 0.00027907 | 0.0002012 | 0.00022462 | 0.00025299 | 0.000354029 | 5.9034E-05 | 0.000226268 | 2.1176E-05 | 0.07864555 | -0.64583591 |
| SM | SM 44:1;2O | [M+H]+ | 124130904 | 139888592 | 0.091845596 | 0.094451769 | 0.079053773 | 0.0595104 | 0.045484451 | 0.046035798 | 0.00087832 | 0.00092788 | 0.00077354 | 0.00063028 | 0.00059272 | 0.000611 | 0.000859915 | 6.43406E-05 | 0.000611335 | 1.53352E-05 | 0.02656577 | -0.49223136 |
| SM | SM 44:2;2O | [M+H]+ | 281292896 | 313776480 | 0.042750181 | 0.046918946 | 0.03632783 | 0.025259745 | 0.02037738 | 0.021484241 | 0.00040882 | 0.00046093 | 0.00035547 | 0.00026753 | 0.00026554 | 0.00028515 | 0.000408405 | 4.30544E-05 | 0.000272739 | 8.81025E-06 | 0.04202724 | -0.58247699 |
| SM | SM 44:3;2O | [M+H]+ | 133046720 | 134001736 | 0.096876135 | 0.105241331 | 0.0786516 | 0.060263386 | 0.047749247 | 0.046520998 | 0.00092643 | 0.00103388 | 0.0007696 | 0.00063826 | 0.00062223 | 0.00061744 | 0.000909971 | 0.000108515 | 0.000625978 | 8.89945E-06 | 0.0649348 | -0.53970955 |
| Cer | Cer 28:5;4O | [M+H-H2O]+ | 4.081 | 460.34167 | 0.045820752 | 0.044944481 | 0.038051503 | 0.028350857 | 0.020956304 | 0.021330619 | 0.00043819 | 0.00044153 | 0.00037233 | 0.00030027 | 0.00027309 | 0.00028311 | 0.000417349 | 3.18604E-05 | 0.000285487 | 1.12226E-05 | 0.01881025 | -0.54782911 |
| Cer | Cer 34:1;2O\|Cer 18:1;2O/16:0 | [M+H-H2O]+ | 15.658 | 520.50824 | 0.162111572 | 0.129306932 | 0.149561399 | 0.326162972 | 0.260844099 | 0.192983187 | 0.00155028 | 0.0012703 | 0.00146346 | 0.00345442 | 0.00339914 | 0.00256134 | 0.00142801 | 0.000117018 | 0.0031383 | 0.000408597 | 0.02070424 | 1.135977011 |
| Cer | Cer 39:1;2O\|Cer 15:1;2O/24:0 | [M+H-H2O]+ | 18.505 | 590.58783 | 1.107434436 | 0.965832154 | 0.876349616 | 0.965926819 | 0.850099314 | 0.729644964 | 0.01059044 | 0.00948822 | 0.00857507 | 0.01023023 | 0.0110779 | 0.0096841 | 0.009551241 | 0.000823978 | 0.010330741 | 0.000573438 | 0.34064989 | 0.113183575 |
| Cer | Cer 40:1;2O\|Cer 18:1;2O/22:0 | [M+H-H2O]+ | 17.634 | 604.60303 | 0.280056888 | 0.246800118 | 0.287237749 | 0.306447204 | 0.301836176 | 0.332718637 | 0.00267819 | 0.00242453 | 0.00281062 | 0.00324561 | 0.00393332 | 0.00441596 | 0.002637782 | 0.000160187 | 0.003864962 | 0.000480229 | 0.05675693 | 0.551128842 |
| Cer | Cer 40:1;2O\|Cer 18:1;2O/22:0 | [M+H-H2O]+ | 18.886 | 604.60303 | 0.274928269 | 0.228279516 | 0.205928218 | 0.188493092 | 0.170184392 | 0.157223252 | 0.00262915 | 0.00224259 | 0.002015 | 0.00199635 | 0.00221772 | 0.00208672 | 0.002295581 | 0.000253508 | 0.002100264 | 9.08818E-05 | 0.39370374 | -0.1282886 |
| Cer | Cer 41:1;2O\|Cer 18:1;2O/23:0 | [M+H-H2O]+ | 19.412 | 618.6189 | 0.581015344 | 0.482305109 | 0.463545032 | 0.485932096 | 0.424380842 | 0.396230008 | 0.00555627 | 0.00473811 | 0.00453578 | 0.00514655 | 0.00553023 | 0.0052589 | 0.004943387 | 0.000441177 | 0.005311896 | 0.000161056 | 0.36159706 | 0.103727216 |
| Cer | Cer 41:2;2O\|Cer 18:1;2O/23:1 | [M+H-H2O]+ | 18.343 | 616.60223 | 0.175150356 | 0.145346012 | 0.150208697 | 0.158504986 | 0.138584853 | 0.125840666 | 0.00167497 | 0.00142786 | 0.00146979 | 0.00167874 | 0.00180594 | 0.0016702 | 0.001524207 | 0.000107971 | 0.001718294 | 6.20732E-05 | 0.1093649 | 0.17291835 |
| Cer | Cer 42:0;3O\|Cer 18:0;2O/24:0;O | [M+H-H2O]+ | 19.203 | 650.64539 | 0.124937576 | 0.098524147 | 0.111827659 | 0.106916389 | 0.106552671 | 0.093313752 | 0.00119478 | 0.00096789 | 0.00109423 | 0.00113236 | 0.00138852 | 0.00123849 | 0.001085635 | 9.28281E-05 | 0.001253125 | 0.000105086 | 0.16751474 | 0.20699117 |
| Cer | Cer 42:1;2O\|Cer 18:1;2O/24:0 | [M+H-H2O]+ | 18.667 | 632.63434 | 0.079230345 | 0.073728949 | 0.060828192 | 0.084895811 | 0.074384811 | 0.069136518 | 0.00075768 | 0.0007243 | 0.0005952 | 0.00089914 | 0.00096933 | 0.0009176 | 0.000692397 | 7.00643E-05 | 0.000928691 | 2.97084E-05 | 0.0272019 | 0.423600457 |
| Cer | Cer 42:1;2O\|Cer 18:1;2O/24:0 | [M+H-H2O]+ | 17.735 | 632.63434 | 0.829320676 | 0.700113256 | 0.611773214 | 0.417962935 | 0.382141919 | 0.363342041 | 0.00793082 | 0.00687783 | 0.00598619 | 0.00442669 | 0.00497981 | 0.0048224 | 0.006931615 | 0.000794804 | 0.004742964 | 0.000232691 | 0.05049855 | -0.54740256 |
| Cer | Cer 42:1;2O\|Cer 18:1;2O/24:0 | [M+H-H2O]+ | 18.111 | 632.63434 | 0.290705411 | 0.245019637 | 0.211020156 | 0.19554512 | 0.155786282 | 0.15657244 | 0.00278003 | 0.00240704 | 0.00206483 | 0.00207104 | 0.0020301 | 0.00207808 | 0.0024173 | 0.000292068 | 0.002059739 | 2.11565E-05 | 0.22504579 | -0.23093453 |
| Cer | Cer 42:1;2O\|Cer 18:1;2O/24:0 | [M+H-H2O]+ | 19.908 | 632.63434 | 0.467753311 | 0.412399927 | 0.375220062 | 0.323227577 | 0.280878069 | 0.278508113 | 0.00447314 | 0.00405137 | 0.00367152 | 0.00342334 | 0.00366021 | 0.00369646 | 0.004065344 | 0.000327409 | 0.003593332 | 0.000121114 | 0.16833942 | -0.17805519 |
| Cer | Cer 42:1;3O\|Cer 18:1;2O/24:0;O | [M+H-H2O]+ | 18.855 | 648.62762 | 0.999751662 | 0.858263415 | 0.795870442 | 0.847517837 | 0.695979345 | 0.650069854 | 0.00956066 | 0.00843148 | 0.00778758 | 0.00897615 | 0.00906951 | 0.00862795 | 0.00859324 | 0.00073284 | 0.008891203 | 0.00019001 | 0.627876 | 0.049176517 |
| Cer | Cer 42:2;2O\|Cer 18:1;2O/24:1 | [M+H-H2O]+ | 17.04 | 630.61682 | 0.345872096 | 0.267925278 | 0.267238589 | 0.322273153 | 0.273432304 | 0.318144331 | 0.00330759 | 0.00263207 | 0.00261493 | 0.00341323 | 0.00356318 | 0.00422252 | 0.002851526 | 0.00032256 | 0.003732975 | 0.000351532 | 0.05968484 | 0.388591543 |
| Cer | Cer 42:2;2O\|Cer 18:1;2O/24:1 | [M+H-H2O]+ | 17.606 | 630.61896 | 0.224590135 | 0.187443457 | 0.188132892 | 0.183239604 | 0.151022968 | 0.147792952 | 0.00214776 | 0.00184142 | 0.00184088 | 0.00194071 | 0.00196803 | 0.00196156 | 0.001943354 | 0.00014454 | 0.001956764 | 1.16559E-05 | 0.90778046 | 0.009920926 |
| Cer | Cer 42:2;2O\|Cer 18:1;2O/24:1 | [M+H-H2O]+ | 18.855 | 630.61896 | 0.317911537 | 0.264520314 | 0.256562105 | 0.204374431 | 0.177001715 | 0.15038456 | 0.0030402 | 0.00259862 | 0.00251046 | 0.00216455 | 0.00230656 | 0.00199596 | 0.002716424 | 0.000231756 | 0.002155689 | 0.000126959 | 0.05524386 | -0.33355954 |
| Cer | Cer 42:3;2O\|Cer 18:2;2O/24:1 | [M+H-H2O]+ | 18.017 | 628.6026 | 0.816795714 | 0.652263983 | 0.70469388 | 0.713292956 | 0.638598509 | 0.742693784 | 0.00781105 | 0.00640776 | 0.00689542 | 0.00755456 | 0.00832177 | 0.00985729 | 0.007038077 | 0.000581702 | 0.00857787 | 0.000957369 | 0.13876385 | 0.285438274 |
| Cer | Cer 44:1;2O\|Cer 18:1;2O/26:0 | [M+H-H2O]+ | 20.866 | 660.66492 | 0.196786053 | 0.154569259 | 0.179177295 | 0.175467504 | 0.166708845 | 0.143485961 | 0.00188187 | 0.00151847 | 0.00175325 | 0.00185839 | 0.00217243 | 0.0019044 | 0.001717863 | 0.000150453 | 0.001978407 | 0.000138476 | 0.14639523 | 0.203724244 |
| Cer | Cer 44:1;2O\|Cer 18:1;2O/26:0 | [M+H-H2O]+ | 19.673 | 660.66528 | 0.095814906 | 0.095933543 | 0.073672257 | 0.096868865 | 0.056511517 | 0.057136704 | 0.00091628 | 0.00094244 | 0.00072088 | 0.00102595 | 0.00073642 | 0.00075834 | 0.000859868 | 9.88563E-05 | 0.000840235 | 0.000131623 | 0.87484371 | -0.03332231 |
| CE | CE 46:1;4O\|Cer 20:1;3O/26:0;(2OH) | [M+H-H2O]+ | 19.626 | 720.68744 | 0.063824001 | 0.049287203 | 0.044927078 | 0.021957428 | 0.018064564 | 0.019110955 | 0.00061035 | 0.00048419 | 0.00043961 | 0.00023255 | 0.0002354 | 0.00025365 | 0.000511385 | 7.23081E-05 | 0.000240535 | 9.34445E-06 | 0.03204167 | -1.08816141 |
| CE | CE 18:1 | [M+NH4]+ | 25.232 | 668.63348 | 0.303020451 | 0.289552629 | 0.319727955 | 0.359263007 | 0.292409071 | 0.307937847 | 0.0028978 | 0.00284453 | 0.00312853 | 0.00380499 | 0.00381047 | 0.00408706 | 0.002956953 | 0.000123259 | 0.003900839 | 0.000131695 | 0.0018091 | 0.399673053 |
| CE | CE 18:2 | [M+NH4]+ | 24.71 | 666.61896 | 0.640159619 | 0.568074843 | 0.644726352 | 0.617394378 | 0.626214969 | 0.55493251 | 0.00612187 | 0.0055807 | 0.00630864 | 0.00653888 | 0.00816039 | 0.00736526 | 0.006003736 | 0.000308696 | 0.007354845 | 0.000662019 | 0.0841987 | 0.292834401 |
| CE | CE 20:1 | [M+NH4]+ | 26.022 | 696.66595 | 0.084885187 | 0.085407028 | 0.082083281 | 0.108386587 | 0.133387073 | 0.123654108 | 0.00081176 | 0.00083903 | 0.00080318 | 0.00114793 | 0.00173821 | 0.00164118 | 0.000817991 | 1.52824E-05 | 0.001509107 | 0.000258442 | 0.06286111 | 0.883538203 |
| CE | CE 20:3 | [M+NH4]+ | 24.819 | 692.63379 | 0.786278061 | 0.764529387 | 0.781779207 | 1.154022525 | 1.406008567 | 1.215922655 | 0.00751921 | 0.00751065 | 0.0076497 | 0.01222237 | 0.01832212 | 0.01613814 | 0.00755985 | 6.36275E-05 | 0.015560875 | 0.002523445 | 0.04622695 | 1.041493782 |
| CE | CE 20:4 | [M+NH4]+ | 24.298 | 690.61743 | 1.42189911 | 1.263685534 | 1.506780237 | 2.230024118 | 2.659258166 | 2.207102408 | 0.01359767 | 0.0124143 | 0.01474382 | 0.02361841 | 0.03465358 | 0.02929342 | 0.013585264 | 0.000951065 | 0.029188471 | 0.004505704 | 0.03450408 | 1.103356081 |
| CE | CE 20:5 | [M+NH4]+ | 23.731 | 688.6026 | 0.615786961 | 0.543596865 | 0.635085788 | 0.695664384 | 0.853249342 | 0.677482288 | 0.00588879 | 0.00534023 | 0.0062143 | 0.00736785 | 0.01111895 | 0.00899178 | 0.005814443 | 0.000360691 | 0.009159525 | 0.001535965 | 0.08417228 | 0.655631779 |
| CE | CE 22:1 | [M+NH4]+ | 26.653 | 724.69739 | 0.027539342 | 0.023201584 | 0.026261744 | 0.042452948 | 0.050448408 | 0.046372912 | 0.00026336 | 0.00022793 | 0.00025697 | 0.00044962 | 0.00065741 | 0.00061548 | 0.00024942 | 1.54183E-05 | 0.00057417 | 8.97158E-05 | 0.03294692 | 1.202899627 |
| CE | CE 22:2 | [M+NH4]+ | 26.077 | 722.6803 | 0.035946557 | 0.036994001 | 0.032902587 | 0.047301578 | 0.058605607 | 0.048244976 | 0.00034376 | 0.00036342 | 0.00032195 | 0.00050098 | 0.00076371 | 0.00064032 | 0.000343045 | 1.6939E-05 | 0.000635002 | 0.000107326 | 0.05815423 | 0.888364956 |
| CE | CE 22:3 | [M+NH4]+ | 25.494 | 720.66644 | 0.10170903 | 0.094071417 | 0.096487258 | 0.160193199 | 0.178731777 | 0.154812737 | 0.00097265 | 0.00092415 | 0.00094413 | 0.00169662 | 0.00232911 | 0.00205473 | 0.000946973 | 1.99025E-05 | 0.002026819 | 0.000258964 | 0.02697872 | 1.097821573 |
| CE | CE 22:4 | [M+NH4]+ | 24.92 | 718.65057 | 0.49691088 | 0.422064683 | 0.501774445 | 1.097558972 | 1.320338745 | 1.123904961 | 0.00475198 | 0.00414631 | 0.00490985 | 0.01162436 | 0.01720573 | 0.01491685 | 0.004602715 | 0.000329098 | 0.014582312 | 0.002290832 | 0.02344504 | 1.663662422 |
| CE | CE 22:5 | [M+NH4]+ | 24.396 | 716.63293 | 0.89127477 | 0.805839419 | 0.947341929 | 1.387506685 | 1.642020992 | 1.413089098 | 0.00852329 | 0.00791647 | 0.00926973 | 0.01469522 | 0.02139766 | 0.018755 | 0.00856983 | 0.000553443 | 0.01828263 | 0.002756573 | 0.03370766 | 1.093135121 |
| CE | CE 22:6 | [M+NH4]+ | 23.977 | 714.61865 | 1.782615836 | 1.585229408 | 1.82215765 | 1.621756874 | 1.872156075 | 1.564634043 | 0.01704722 | 0.0155731 | 0.01782978 | 0.01717619 | 0.02439662 | 0.02076636 | 0.016816703 | 0.000935594 | 0.020779724 | 0.002947746 | 0.19017416 | 0.305281573 |
| CE | CE 24:3 | [M+NH4]+ | 26.154 | 748.69788 | 0.08217401 | 0.077897836 | 0.080935019 | 0.08109985 | 0.092927207 | 0.089188929 | 0.00078583 | 0.00076526 | 0.00079195 | 0.00085894 | 0.00121096 | 0.00118375 | 0.000781013 | 1.14162E-05 | 0.001084548 | 0.000159918 | 0.11454484 | 0.473675132 |
| CE | CE 24:4 | [M+NH4]+ | 25.603 | 746.68134 | 0.188070465 | 0.170197199 | 0.19418442 | 0.358861886 | 0.414220727 | 0.373826065 | 0.00179852 | 0.001672 | 0.00190009 | 0.00380074 | 0.00539783 | 0.00496155 | 0.001790204 | 9.33048E-05 | 0.004720041 | 0.000674003 | 0.02367535 | 1.398675087 |
| CE | CE 24:5 | [M+NH4]+ | 25.014 | 744.66449 | 0.293279699 | 0.282725646 | 0.300391295 | 0.430488026 | 0.450387941 | 0.417798635 | 0.00280464 | 0.00277746 | 0.00293932 | 0.00455934 | 0.00586914 | 0.00554517 | 0.002840477 | 7.07705E-05 | 0.005324549 | 0.000557014 | 0.02275728 | 0.906526051 |
| CE | CE 24:6 | [M+NH4]+ | 24.544 | 742.65045 | 0.375148045 | 0.362972584 | 0.357825847 | 0.162797757 | 0.173886744 | 0.164597598 | 0.00358755 | 0.0035658 | 0.00350132 | 0.00172421 | 0.00226597 | 0.0021846 | 0.003551558 | 3.66172E-05 | 0.002058258 | 0.000238534 | 0.011097 | -0.78702849 |
| CE | CE 26:1 | [M+NH4]+ | 27.729 | 780.75775 | 0.091530291 | 0.091599797 | 0.086277733 | 0.05967865 | 0.07403153 | 0.066541447 | 0.00087531 | 0.00089987 | 0.00084423 | 0.00063206 | 0.00096473 | 0.00088316 | 0.000873133 | 2.27666E-05 | 0.00082665 | 0.000141566 | 0.68965508 | -0.07892476 |
| CE | CE 26:2 | [M+NH4]+ | 27.224 | 778.74438 | 0.064941482 | 0.064920096 | 0.065436107 | 0.045408882 | 0.05764648 | 0.052960838 | 0.00062104 | 0.00063777 | 0.00064029 | 0.00048093 | 0.00075121 | 0.00070291 | 0.000633032 | 8.54359E-06 | 0.000645018 | 0.000117691 | 0.89882644 | 0.027059868 |
| CE | CE 26:3 | [M+NH4]+ | 26.731 | 776.72711 | 0.106056262 | 0.100305728 | 0.100412994 | 0.062888318 | 0.081425145 | 0.064963697 | 0.00101422 | 0.00098539 | 0.00098254 | 0.00066606 | 0.00106108 | 0.00086222 | 0.00099405 | 1.43096E-05 | 0.000863117 | 0.000161267 | 0.36961297 | -0.2037621 |
| CE | CE 26:4 | [M+NH4]+ | 26.24 | 774.71185 | 0.158380114 | 0.155023353 | 0.148440295 | 0.128743373 | 0.146381875 | 0.134736018 | 0.0015146 | 0.00152293 | 0.00145249 | 0.00136353 | 0.00190755 | 0.00178826 | 0.001496671 | 3.14281E-05 | 0.001686447 | 0.00023347 | 0.36912538 | 0.172230634 |
| CE | CE 26:5 | [M+NH4]+ | 25.719 | 772.69586 | 0.27388476 | 0.280006618 | 0.252745913 | 0.146528204 | 0.155443061 | 0.145468755 | 0.00261917 | 0.00275075 | 0.00247311 | 0.00155189 | 0.00202562 | 0.00193071 | 0.002614345 | 0.000113396 | 0.001836077 | 0.000204649 | 0.0166116 | -0.5098233 |
| CE | CE 28:5 | [M+NH4]+ | 26.307 | 800.72632 | 0.060323553 | 0.07076951 | 0.061434257 | 0.02259021 | 0.026816154 | 0.025373221 | 0.00057688 | 0.00069523 | 0.00060113 | 0.00023926 | 0.00034945 | 0.00033676 | 0.000624414 | 5.10454E-05 | 0.000308489 | 4.92289E-05 | 0.00325849 | -1.01728364 |
| CE | CE 28:6 | [M+NH4]+ | 25.837 | 798.71094 | 0.062062145 | 0.059444727 | 0.049661171 | 0.014894188 | 0.019824789 | 0.016252908 | 0.0005935 | 0.00058398 | 0.00048593 | 0.00015775 | 0.00025834 | 0.00021571 | 0.000554471 | 4.86192E-05 | 0.000210601 | 4.12274E-05 | 0.00176182 | -1.39660144 |
| CE | CE 30:5 | [M+NH4]+ | 26.85 | 828.75818 | 0.028768114 | 0.02941069 | 0.023751824 | 0.018802005 | 0.023102663 | 0.017189826 | 0.00027511 | 0.00028893 | 0.00023241 | 0.00019913 | 0.00030106 | 0.00022815 | 0.000265483 | 2.40559E-05 | 0.00024278 | 4.28771E-05 | 0.55832068 | -0.12896798 |
| CE | CE 30:6 | [M+NH4]+ | 26.433 | 826.74463 | 0.034951034 | 0.038685666 | 0.028099632 | 0.018867548 | 0.023446212 | 0.019740677 | 0.00033424 | 0.00038004 | 0.00027495 | 0.00019983 | 0.00030553 | 0.00026201 | 0.000329745 | 4.30198E-05 | 0.000255789 | 4.33777E-05 | 0.16206962 | -0.36639655 |
| CE | CE 32:6 | [M+NH4]+ | 26.976 | 854.77576 | 0.025615538 | 0.024434645 | 0.020969744 | 0.018306376 | 0.020658389 | 0.019381894 | 0.00024496 | 0.00024004 | 0.00020519 | 0.00019388 | 0.00026921 | 0.00025724 | 0.000230065 | 1.77043E-05 | 0.000240111 | 3.30499E-05 | 0.72949481 | 0.061662906 |

| | | | | | | | | | | | | | | | | | | |
|---|---|---|---|---|---|---|---|---|---|---|---|---|---|---|---|---|---|---|
| TG 62:6\|TG 16:0_18:1_28:5 | 0.001747045 | 0.001853312 | 0.001962528 | 0.001807921 | 0.001884514 | 0.003234528 | 2.7151E-05 | 2.7124E-05 | 2.5464E-05 | 2.4342E-05 | 0.00013638 | 3.6158E-05 | 2.65797E-05 | 7.88934E-07 | 3.00724E-05 | 4.83054E-06 | 0.4144989 | 0.178118113 |
| TG 62:7\|TG 18:0_18:1_26:6 | 0.00215622 | 0.00212558 | 0.002272471 | 0.002081314 | 0.001972102 | 0.003465465 | 3.3509E-05 | 3.1109E-05 | 2.9486E-05 | 2.8023E-05 | 0.00013638 | 3.874E-05 | 3.13681E-05 | 1.65289E-06 | 3.26204E-05 | 4.50563E-06 | 0.74080096 | 0.056474065 |
| TG 62:8\|TG 18:1_18:1_26:6 | 0.002461843 | 0.002495565 | 0.002466369 | 0.00257347 | 0.002305322 | 0.003938565 | 3.8259E-05 | 3.6524E-05 | 3.2001E-05 | 3.4649E-05 | 0.00013638 | 4.4029E-05 | 3.55949E-05 | 2.63783E-06 | 3.83436E-05 | 4.07963E-06 | 0.47544231 | 0.107313936 |
| TG 62:9\|TG 18:1_20:4_24:4 | 0.002117268 | 0.002017133 | 0.002131138 | 0.002245367 | 0.001950285 | 0.00317819 | 3.2904E-05 | 2.9522E-05 | 2.7652E-05 | 3.0232E-05 | 0.00013638 | 3.5529E-05 | 3.0026E-05 | 2.17365E-06 | 3.21715E-05 | 2.38333E-06 | 0.40056793 | 0.099570675 |
| TG O-52:1\|TG O-18:0_16:0_18:1 | 0.001028734 | 0.000849529 | 0.000862061 | 0.000938799 | 0.000737075 | 0.001224193 | 1.5987E-05 | 1.2433E-05 | 1.1185E-05 | 1.264E-05 | 0.00013638 | 1.3685E-05 | 1.32021E-05 | 2.03437E-06 | 1.26494E-05 | 8.41864E-07 | 0.74878552 | -0.06169638 |
| DG 34:0\|DG 16:0_18:0 | 0.220889352 | 0.208230715 | 0.226418913 | 0.119700894 | 0.102361519 | 0.150872451 | 0.0034328 | 0.00304759 | 0.00293782 | 0.00161166 | 0.00013638 | 0.00168658 | 0.003139405 | 0.000212251 | 0.001637463 | 3.47461E-05 | 0.00847403 | -0.93902904 |
| DG 34:1\|DG 16:0_18:1 | 0.050028407 | 0.058128323 | 0.057831763 | 0.03782967 | 0.030850573 | 0.077328314 | 0.00077748 | 0.00085075 | 0.00075037 | 0.00050934 | 0.00013638 | 0.00086444 | 0.000792868 | 4.23961E-05 | 0.000620089 | 0.000173035 | 0.29144251 | -0.35460504 |
| DG 36:0\|DG 18:0_18:0 | 0.113892395 | 0.106881522 | 0.113329147 | 0.061604029 | 0.054227724 | 0.075654027 | 0.00176998 | 0.00156428 | 0.00147046 | 0.00082944 | 0.00013638 | 0.00084572 | 0.001601575 | 0.00012509 | 0.000843429 | 1.06093E-05 | 0.01286328 | -0.92515292 |
| DG 36:2\|DG 18:1_18:1 | 0.022600417 | 0.039484645 | 0.031530715 | 0.016057245 | 0.014988828 | 0.069990192 | 0.00035123 | 0.00057788 | 0.00040912 | 0.0002162 | 0.00013638 | 0.00078241 | 0.000446076 | 9.61515E-05 | 0.000411655 | 0.000262292 | 0.87466134 | -0.11585361 |
| DG 38:4\|DG 18:0_20:4 | 0.083090386 | 0.079678768 | 0.080987099 | 0.047044569 | 0.040477517 | 0.055229311 | 0.00129129 | 0.00116615 | 0.00105082 | 0.00063341 | 0.00013638 | 0.0006174 | 0.001169421 | 9.82009E-05 | 0.000629702 | 8.9239E-06 | 0.01553946 | -0.89305469 |
| SM 32:1;2O\|SM 18:1;2O/14:0 | 0.162894098 | 0.160244168 | 0.206921651 | 0.168164355 | 0.154365288 | 0.191891604 | 0.00253151 | 0.00234528 | 0.00268484 | 0.00226418 | 0.00013638 | 0.00214513 | 0.002520542 | 0.00013884 | 0.002281167 | 0.000118623 | 0.13912741 | -0.14396162 |
| SM 33:1;2O | 0.078519208 | 0.07501972 | 0.099865803 | 0.079822958 | 0.073454353 | 0.088632064 | 0.00122025 | 0.00109796 | 0.00129577 | 0.00107474 | 0.00013638 | 0.0009908 | 0.001204663 | 8.15041E-05 | 0.001074618 | 6.83832E-05 | 0.16109512 | -0.16480596 |
| SM 34:0;2O | 0.167616657 | 0.167424457 | 0.203227156 | 0.149514161 | 0.125579735 | 0.184347503 | 0.0026049 | 0.00245037 | 0.0026369 | 0.00201307 | 0.00013638 | 0.00206079 | 0.002564057 | 8.14443E-05 | 0.002018046 | 3.3058E-05 | 0.00500985 | -0.34546895 |
| SM 34:1;2O\|SM 18:1;2O/16:0 | 1.321193823 | 1.543637146 | 1.878035094 | 1.746918747 | 1.458091526 | 1.843551882 | 0.02053245 | 0.02259216 | 0.02436775 | 0.02352062 | 0.00013638 | 0.02060879 | 0.022497455 | 0.001567186 | 0.022374061 | 0.001266702 | 0.93532597 | -0.00793463 |
| SM 34:2;2O\|SM 18:2;2O/16:0 | 0.205338949 | 0.260377314 | 0.301033346 | 0.243425984 | 0.235189859 | 0.260155009 | 0.00319114 | 0.00381079 | 0.00390595 | 0.0032775 | 0.00013638 | 0.00290823 | 0.00363596 | 0.000316926 | 0.003298155 | 0.000327127 | 0.35348598 | -0.14067718 |
| SM 35:1;2O\|SM 18:1;2O/17:0 | 0.058948216 | 0.058139993 | 0.072689926 | 0.064162324 | 0.053403593 | 0.070385668 | 0.0009161 | 0.00085092 | 0.00094316 | 0.00086389 | 0.00013638 | 0.00078683 | 0.000903394 | 3.8716E-05 | 0.000830947 | 3.24354E-05 | 0.11455284 | -0.12059886 |
| SM 36:0;2O | 0.205174981 | 0.207689535 | 0.259199025 | 0.232115931 | 0.185867881 | 0.247994255 | 0.00318859 | 0.00303967 | 0.00336314 | 0.00312522 | 0.00013638 | 0.00277229 | 0.003197135 | 0.000132193 | 0.002942827 | 0.000144328 | 0.14054118 | -0.11957722 |
| SM 36:1;2O\|SM 18:1;2O/18:0 | 0.028732239 | 0.030009411 | 0.03901029 | 0.03522092 | 0.028705786 | 0.039573109 | 0.00044652 | 0.00043921 | 0.00050616 | 0.00047422 | 0.00013638 | 0.00044238 | 0.000463965 | 2.99882E-05 | 0.000456421 | 1.32652E-05 | 0.76804491 | -0.02365061 |
| SM 37:1;3O | 0.071210998 | 0.073427063 | 0.095142262 | 0.085073068 | 0.071610698 | 0.087499506 | 0.00110668 | 0.00107465 | 0.00123448 | 0.00114543 | 0.00013638 | 0.00097814 | 0.001138605 | 6.90453E-05 | 0.001084269 | 7.53328E-05 | 0.49419091 | -0.07054435 |
| SM 38:1;2O\|SM 18:1;2O/20:0 | 0.028697417 | 0.025275642 | 0.022025588 | 0.053783579 | 0.040748185 | 0.057615642 | 0.00044598 | 0.00036993 | 0.00028578 | 0.00072415 | 0.00013638 | 0.00064408 | 0.000367231 | 6.54279E-05 | 0.000670261 | 3.81069E-05 | 0.00902145 | 0.868036262 |
| SM 40:0;2O | 0.076433394 | 0.098925139 | 0.05515112 | 0.0559255 | 0.044281624 | 0.066728938 | 0.00118784 | 0.00144784 | 0.00071559 | 0.00075298 | 0.00013638 | 0.00074595 | 0.001117089 | 0.000303094 | 0.000732406 | 2.43003E-05 | 0.21388362 | -0.60902863 |
| SM 40:1;2O\|SM 18:1;2O/22:0 | 0.092171292 | 0.093879226 | 0.115339004 | 0.100929259 | 0.082865973 | 0.107005344 | 0.00143242 | 0.00137398 | 0.00149654 | 0.00135892 | 0.00013638 | 0.0011962 | 0.001434314 | 5.00504E-05 | 0.001287278 | 6.78384E-05 | 0.07468969 | -0.15603696 |
| SM 40:2;2O\|SM 18:1;2O/22:1 | 0.41015266 | 0.417326084 | 0.541949254 | 0.465750832 | 0.382296098 | 0.483199838 | 0.00637411 | 0.00610784 | 0.00703186 | 0.0062709 | 0.00013638 | 0.00540162 | 0.006504608 | 0.00038835 | 0.005900326 | 0.000366266 | 0.1849009 | -0.14066742 |
| SM 40:2;2O\|SM 18:1;2O/22:1 | 0.03218624 | 0.034659494 | 0.042645876 | 0.039367078 | 0.033188909 | 0.042300634 | 0.0005002 | 0.00050726 | 0.00055334 | 0.00053004 | 0.00013638 | 0.00047287 | 0.000520267 | 2.35602E-05 | 0.000508757 | 2.55207E-05 | 0.66387165 | -0.03227512 |
| SM 41:2;2O | 0.084560825 | 0.083350387 | 0.102653961 | 0.091286835 | 0.074105938 | 0.099086388 | 0.00131415 | 0.00121989 | 0.00133195 | 0.00122909 | 0.00013638 | 0.00110767 | 0.001288661 | 4.91697E-05 | 0.001168449 | 4.95696E-05 | 0.07160391 | -0.14127819 |
| SM 42:1;2O\|SM 18:1;2O/24:0 | 0.195427428 | 0.202957094 | 0.247409774 | 0.221960042 | 0.180990492 | 0.228257478 | 0.00303711 | 0.00297041 | 0.00321017 | 0.00298848 | 0.00013638 | 0.00255166 | 0.003072564 | 0.000101043 | 0.002798065 | 0.000182676 | 0.15637722 | -0.13501361 |
| SM 42:2;2O\|SM 18:1;2O/24:1 | 0.992188629 | 1.006881751 | 1.219240061 | 1.08064394 | 0.894502994 | 1.17479522 | 0.01541944 | 0.01473638 | 0.0158198 | 0.01454986 | 0.00013638 | 0.01313286 | 0.015325208 | 0.000447294 | 0.013929404 | 0.000591737 | 0.06070813 | -0.13777313 |
| SM 42:3;2O\|SM 18:1;2O/24:2 | 0.807460428 | 0.851798561 | 1.053842206 | 0.904716093 | 0.785353209 | 1.00428814 | 0.01254861 | 0.01246664 | 0.01367374 | 0.01218115 | 0.00013638 | 0.01122678 | 0.01289633 | 0.000550732 | 0.011930748 | 0.000504639 | 0.1420831 | -0.11227608 |
| SM 43:1;2O | 0.057583197 | 0.05838507 | 0.067170962 | 0.061021565 | 0.052487364 | 0.068210789 | 0.00089489 | 0.0008545 | 0.00087155 | 0.0008216 | 0.00013638 | 0.00076252 | 0.000873649 | 1.65543E-05 | 0.000803931 | 2.93886E-05 | 0.05771628 | -0.11998163 |
| SM 43:2;2O | 0.020566418 | 0.02239525 | 0.032809174 | 0.022897821 | 0.020788072 | 0.024617728 | 0.00031962 | 0.00032777 | 0.0004257 | 0.0003083 | 0.00013638 | 0.0002752 | 0.000357697 | 4.82025E-05 | 0.000303768 | 2.17159E-05 | 0.25163054 | -0.23576875 |
| SM 44:1;2O | 0.029791807 | 0.029409062 | 0.031725487 | 0.032959862 | 0.028064957 | 0.035032562 | 0.00046299 | 0.00043042 | 0.00041164 | 0.00044377 | 0.00013638 | 0.00039162 | 0.000435018 | 2.12129E-05 | 0.000425985 | 2.43024E-05 | 0.71268254 | -0.03027023 |
| SM 44:2;2O | 0.058418122 | 0.070508358 | 0.070746088 | 0.062523557 | 0.065786156 | 0.071369681 | 0.00090787 | 0.00103194 | 0.00091794 | 0.00084182 | 0.00013638 | 0.00079783 | 0.000952581 | 5.62635E-05 | 0.000892347 | 0.00010412 | 0.52248944 | -0.09423763 |
| SM 44:3;2O | 0.02668331 | 0.027709733 | 0.035285959 | 0.028393614 | 0.025619329 | 0.031661249 | 0.00041468 | 0.00040555 | 0.00045784 | 0.00038229 | 0.00013638 | 0.00035394 | 0.000426024 | 2.28042E-05 | 0.000380074 | 2.04959E-05 | 0.10219573 | -0.16465206 |
| Cer 28:5;4O | 0.101644801 | 0.094977699 | 0.097654071 | 0.165059846 | 0.124109819 | 0.175874523 | 0.00157964 | 0.00139006 | 0.00126707 | 0.00222238 | 0.00013638 | 0.00196607 | 0.00141226 | 0.000128568 | 0.002048516 | 0.000122992 | 0.00723195 | 0.536573347 |
| Cer 34:1;2O\|Cer 18:1;2O/16:0 | 0.577875542 | 0.665442918 | 0.722423622 | 0.700386208 | 0.630885059 | 0.895129538 | 0.00898067 | 0.0097392 | 0.00937354 | 0.00943004 | 0.00013638 | 0.01000652 | 0.00936447 | 0.000309736 | 0.009795014 | 0.000259159 | 0.20828372 | 0.064850291 |
| Cer 39:1;2O\|Cer 15:1;2O/24:0 | 0.166548734 | 0.167323875 | 0.170728469 | 0.254969492 | 0.210338958 | 0.300215427 | 0.00258831 | 0.0024489 | 0.00221522 | 0.00343293 | 0.00013638 | 0.00335606 | 0.002417476 | 0.000153922 | 0.003368614 | 4.821E-05 | 0.00806038 | 0.478653708 |
| Cer 40:1;2O\|Cer 18:1;2O/22:0 | 0.170037519 | 0.177013862 | 0.209836553 | 0.195522605 | 0.177855259 | 0.242541917 | 0.00264252 | 0.00259072 | 0.00272266 | 0.00263253 | 0.00013638 | 0.00271134 | 0.002651966 | 5.42771E-05 | 0.002716161 | 7.03368E-05 | 0.36805408 | 0.034506841 |
| Cer 40:1;2O\|Cer 18:1;2O/22:0 | 0.323645375 | 0.353511535 | 0.399716522 | 0.427257805 | 0.399247583 | 0.550039773 | 0.00502972 | 0.00517388 | 0.00518638 | 0.00575263 | 0.00013638 | 0.00614881 | 0.00512999 | 7.10857E-05 | 0.006065736 | 0.000229387 | 0.02102891 | 0.241726625 |
| Cer 41:1;2O\|Cer 18:1;2O/23:0 | 0.076490944 | 0.081606685 | 0.08718862 | 0.098713559 | 0.082737947 | 0.117929638 | 0.00118873 | 0.00119437 | 0.00113128 | 0.00132909 | 0.00013638 | 0.00131832 | 0.001171462 | 2.85029E-05 | 0.001317368 | 9.97715E-06 | 0.01143182 | 0.169349029 |
| Cer 41:2;2O\|Cer 18:1;2O/23:1 | 0.104073071 | 0.108951347 | 0.122801377 | 0.135888711 | 0.122203838 | 0.165296226 | 0.00161738 | 0.00159458 | 0.00159336 | 0.00182961 | 0.00013638 | 0.00184782 | 0.001601774 | 1.10479E-05 | 0.00186816 | 4.22952E-05 | 0.00882491 | 0.221947111 |
| Cer 42:0;3O\|Cer 18:0;2O/24:0;O | 0.129289095 | 0.139330129 | 0.151834545 | 0.151288152 | 0.131258277 | 0.178082748 | 0.00200926 | 0.00203919 | 0.00197007 | 0.00203695 | 0.00013638 | 0.00199076 | 0.002006174 | 2.83006E-05 | 0.002032512 | 3.24298E-05 | 0.43648388 | 0.018817073 |
| Cer 42:1;2O\|Cer 18:1;2O/24:0 | 0.125055397 | 0.144183131 | 0.157550799 | 0.166380337 | 0.145605884 | 0.197600956 | 0.00194347 | 0.00211022 | 0.00204424 | 0.00224016 | 0.00013638 | 0.00220895 | 0.002032641 | 6.85682E-05 | 0.002248393 | 3.60408E-05 | 0.02868881 | 0.145538519 |
| Cer 42:1;2O\|Cer 18:1;2O/24:0 | 0.211234778 | 0.202423214 | 0.233830756 | 0.216587342 | 0.193701662 | 0.267377748 | 0.00328276 | 0.0029626 | 0.00303399 | 0.00291615 | 0.00013638 | 0.00298898 | 0.003093116 | 0.000137232 | 0.00298654 | 5.65086E-05 | 0.3931627 | -0.05058599 |
| Cer 42:1;2O\|Cer 18:1;2O/24:0 | 0.504943217 | 0.555526466 | 0.638161318 | 0.609839664 | 0.744307061 | 0.763818827 | 0.00784724 | 0.0081305 | 0.00828023 | 0.00821092 | 0.00013638 | 0.00853861 | 0.008085989 | 0.000179547 | 0.009495526 | 0.001590628 | 0.33646435 | 0.231823761 |
| Cer 42:1;2O\|Cer 18:1;2O/24:0 | 0.449260439 | 0.464958825 | 0.546871573 | 0.534122198 | 0.46450041 | 0.623630169 | 0.00698188 | 0.00680498 | 0.00709573 | 0.00719145 | 0.00013638 | 0.00697147 | 0.006960865 | 0.000119625 | 0.007162557 | 0.000145667 | 0.20739536 | 0.041208064 |
| Cer 42:1;3O\|Cer 18:1;2O/24:0;O | 0.152182114 | 0.164002022 | 0.188600435 | 0.191731283 | 0.158864955 | 0.217215693 | 0.00236504 | 0.00240028 | 0.00244712 | 0.00258148 | 0.00013638 | 0.00242822 | 0.002404144 | 3.36198E-05 | 0.002504953 | 6.25686E-05 | 0.13639226 | 0.059260334 |
| Cer 42:2;2O\|Cer 18:1;2O/24:1 | 0.261769466 | 0.278049614 | 0.317233333 | 0.332517157 | 0.297344446 | 0.412808041 | 0.00406812 | 0.00406944 | 0.00411614 | 0.00447703 | 0.00013638 | 0.00461472 | 0.004084567 | 2.23353E-05 | 0.004593534 | 8.77626E-05 | 0.01075666 | 0.169421229 |
| Cer 42:2;2O\|Cer 18:1;2O/24:1 | 0.599776318 | 0.620041161 | 0.76390772 | 0.776836273 | 0.714206029 | 0.991633769 | 0.00932102 | 0.00907471 | 0.0099118 | 0.01045937 | 0.00013638 | 0.01108532 | 0.009435848 | 0.000351253 | 0.010935691 | 0.000344477 | 0.01254291 | 0.212820391 |
| Cer 42:2;2O\|Cer 18:1;2O/24:1 | 0.199989698 | 0.212603466 | 0.254692513 | 0.237346733 | 0.299362911 | 0.303822671 | 0.00310801 | 0.00311159 | 0.00330467 | 0.00319565 | 0.00013638 | 0.00339639 | 0.003174756 | 9.1874E-05 | 0.003770906 | 0.000676573 | 0.33849114 | 0.248265462 |
| Cer 42:3;2O\|Cer 18:2;2O/24:1 | 0.038094127 | 0.043359936 | 0.043810703 | 0.074022066 | 0.067992796 | 0.083894292 | 0.00059201 | 0.0006346 | 0.00056845 | 0.00099664 | 0.00013638 | 0.00093784 | 0.000598355 | 2.7376E-05 | 0.001002221 | 5.49872E-05 | 0.0028793 | 0.744126884 |
| Cer 44:1;2O\|Cer 18:1;2O/26:0 | 0.051969245 | 0.052588975 | 0.064622434 | 0.068021997 | 0.063626021 | 0.089666305 | 0.00080765 | 0.00076967 | 0.00083848 | 0.00091585 | 0.00013638 | 0.00100237 | 0.000805268 | 2.81418E-05 | 0.000973848 | 4.10106E-05 | 0.01170455 | 0.27422661 |
| Cer 44:1;2O\|Cer 18:1;2O/26:0 | 0.031762249 | 0.034632951 | 0.042047639 | 0.035982228 | 0.031564466 | 0.045883111 | 0.00049361 | 0.00050688 | 0.00054557 | 0.00048447 | 0.00013638 | 0.00051292 | 0.000515354 | 2.20441E-05 | 0.000498377 | 1.16247E-05 | 0.40572123 | -0.04832692 |
| CE 46:1;4O\|Cer 20:1;3O/26:0;(2OH) | 0.062316453 | 0.030710559 | 0.038065778 | 0.059098776 | 0.032815692 | 0.041075899 | 0.00096845 | 0.00044947 | 0.00049391 | 0.00079571 | 0.00013638 | 0.00045918 | 0.000637276 | 0.000234877 | 0.000590788 | 0.000146843 | 0.82615656 | -0.10927718 |
| CE 18:1 | 0.518063868 | 0.576991726 | 0.665740748 | 0.465431398 | 0.387786674 | 0.547339501 | 0.00805114 | 0.00844466 | 0.00863807 | 0.0062666 | 0.00013638 | 0.00611863 | 0.008377959 | 0.000244211 | 0.006166755 | 7.06144E-05 | 0.00357849 | -0.44208726 |
| CE 18:2 | 0.285672417 | 0.296906803 | 0.357955563 | 0.260995666 | 0.200693117 | 0.30150613 | 0.00443959 | 0.00434543 | 0.00464452 | 0.00351406 | 0.00013638 | 0.00337049 | 0.004476512 | 0.000124864 | 0.003349767 | 0.000143358 | 0.00120202 | -0.41831444 |
| CE 20:1 | 0.435974852 | 0.462955996 | 0.534982205 | 0.585443854 | 0.498758596 | 0.833444388 | 0.00677541 | 0.00677567 | 0.00694147 | 0.00788245 | 0.00013638 | 0.00931695 | 0.006830849 | 7.82176E-05 | 0.00835479 | 0.000680386 | 0.08499266 | 0.290538661 |
| CE 20:3 | 1.074959217 | 1.22071177 | 1.390456064 | 1.367896873 | 1.207357897 | 1.945415376 | 0.01670576 | 0.01786593 | 0.01804135 | 0.01841745 | 0.00013638 | 0.0217475 | 0.017537681 | 0.000592598 | 0.019734629 | 0.001445755 | 0.15274993 | 0.17027138 |
| CE 20:4 | 0.595551285 | 0.61648881 | 0.686863273 | 0.745263062 | 0.649945742 | 1.06058082 | 0.00925536 | 0.00902272 | 0.00891214 | 0.01003427 | 0.00013638 | 0.01185607 | 0.00906341 | 0.000143043 | 0.010713131 | 0.000812924 | 0.09878823 | 0.241254341 |
| CE 20:5 | 0.062618522 | 0.059347332 | 0.076308873 | 0.083648878 | 0.065528479 | 0.110305731 | 0.00097314 | 0.00086859 | 0.00099012 | 0.00112625 | 0.00013638 | 0.00123309 | 0.00094395 | 5.37378E-05 | 0.00113089 | 8.16203E-05 | 0.06290826 | 0.260676167 |
| CE 22:1 | 1.272620275 | 1.311682922 | 1.484596029 | 1.194701727 | 1.047230639 | 1.615478243 | 0.01977758 | 0.01919735 | 0.01926283 | 0.01608554 | 0.00013638 | 0.01805919 | 0.019412587 | 0.00025947 | 0.016886199 | 0.00084766 | 0.0421223 | -0.20114779 |
| CE 22:2 | 0.748852851 | 0.778840449 | 0.827895093 | 0.882854893 | 0.741565704 | 1.247572569 | 0.0116378 | 0.01139885 | 0.01074205 | 0.01188681 | 0.00013638 | 0.01394642 | 0.011259565 | 0.000378718 | 0.012509018 | 0.00101945 | 0.21869113 | 0.151817448 |
| CE 22:3 | 0.333954027 | 0.334534415 | 0.377221142 | 0.425815957 | 0.370561753 | 0.611899621 | 0.00518992 | 0.00489613 | 0.00489449 | 0.00573321 | 0.00013638 | 0.00684033 | 0.004993517 | 0.000138882 | 0.006138989 | 0.000497962 | 0.07356236 | 0.297944688 |
| CE 22:4 | 0.066474807 | 0.073477519 | 0.076941657 | 0.068471345 | 0.065821224 | 0.106495342 | 0.00103307 | 0.00107539 | 0.00099833 | 0.0009219 | 0.00013638 | 0.0011905 | 0.001035598 | 3.15119E-05 | 0.001050113 | 0.00010999 | 0.87205804 | 0.020080014 |
| CE 22:5 | 0.024300733 | 0.026769894 | 0.029392686 | 0.030481914 | 0.026653932 | 0.040173269 | 0.00037765 | 0.0003918 | 0.00038137 | 0.00041041 | 0.00013638 | 0.00044909 | 0.000383608 | 5.9854E-06 | 0.000426603 | 1.64066E-05 | 0.05221329 | 0.153263278 |
| CE 22:6 | 0.021496954 | 0.02065168 | 0.026176051 | 0.026381118 | 0.02354619 | 0.036663344 | 0.00033408 | 0.00030225 | 0.00033964 | 0.0003552 | 0.00013638 | 0.00040985 | 0.000325323 | 1.64714E-05 | 0.000378784 | 2.29323E-05 | 0.0613444 | 0.219503164 |
| CE 24:3 | 0.190513831 | 0.201028704 | 0.213259823 | 0.212431548 | 0.180095817 | 0.293354738 | 0.00296074 | 0.00294219 | 0.00276707 | 0.00286019 | 0.00013638 | 0.00327937 | 0.002890002 | 8.72525E-05 | 0.002993169 | 0.000202542 | 0.55999679 | 0.050603015 |
| CE 24:4 | 0.176414735 | 0.178134475 | 0.192268019 | 0.213266104 | 0.178504964 | 0.310367427 | 0.00274163 | 0.00260712 | 0.0024947 | 0.00287143 | 0.00013638 | 0.00346955 | 0.002614484 | 0.000100943 | 0.003051946 | 0.000296192 | 0.16235437 | 0.223203441 |
| CE 24:5 | 0.108992237 | 0.108257452 | 0.119766396 | 0.127812855 | 0.115032325 | 0.205071271 | 0.00169383 | 0.00158442 | 0.00155398 | 0.00172088 | 0.00013638 | 0.00229246 | 0.001610745 | 6.00497E-05 | 0.001942432 | 0.000250407 | 0.19693158 | 0.270135961 |
| CE 24:6 | 0.053921498 | 0.050251243 | 0.057424409 | 0.056971115 | 0.053832829 | 0.084086394 | 0.00083799 | 0.00073546 | 0.00074509 | 0.00076706 | 0.00013638 | 0.00093999 | 0.000772845 | 4.62285E-05 | 0.000851982 | 7.06307E-05 | 0.26590741 | 0.140644214 |
| CE 26:1 | 0.148747382 | 0.148585969 | 0.161415326 | 0.155066941 | 0.137321016 | 0.230371777 | 0.00231166 | 0.00217465 | 0.00209439 | 0.00208783 | 0.00013638 | 0.00257529 | 0.002193566 | 8.97037E-05 | 0.002276183 | 0.000213861 | 0.65282279 | 0.053338502 |
| CE 26:2 | 0.081812834 | 0.095881352 | 0.101680179 | 0.096266807 | 0.083085627 | 0.152542523 | 0.00127144 | 0.00140329 | 0.00131931 | 0.00129614 | 0.00013638 | 0.00170525 | 0.001331347 | 5.44949E-05 | 0.001437192 | 0.000189632 | 0.51738853 | 0.110366452 |
| CE 26:3 | 0.064594211 | 0.066554985 | 0.068491122 | 0.064732601 | 0.058398553 | 0.096668473 | 0.00100385 | 0.00097408 | 0.00088868 | 0.00087156 | 0.00013638 | 0.00108064 | 0.000955535 | 4.88103E-05 | 0.000957699 | 8.92358E-05 | 0.97782987 | 0.003263657 |
| CE 26:4 | 0.040556204 | 0.040405266 | 0.046855504 | 0.044627767 | 0.038441922 | 0.065671282 | 0.00063028 | 0.00059136 | 0.00060796 | 0.00060087 | 0.00013638 | 0.00073413 | 0.000609864 | 1.59459E-05 | 0.000647065 | 6.16022E-05 | 0.48630502 | 0.085423009 |
| CE 26:5 | 0.050600599 | 0.050660676 | 0.053431781 | 0.056704415 | 0.044869611 | 0.076291168 | 0.00078638 | 0.00074145 | 0.00069328 | 0.00076347 | 0.00013638 | 0.00085285 | 0.000740371 | 3.8012E-05 | 0.000774624 | 5.98381E-05 | 0.53823425 | 0.0652481 |
| CE 28:5 | 0.027749919 | 0.031158425 | 0.032335206 | 0.034729369 | 0.030203523 | 0.055131916 | 0.00043126 | 0.00045602 | 0.00041955 | 0.0004676 | 0.00013638 | 0.00061631 | 0.000435612 | 1.52042E-05 | 0.000520064 | 6.81495E-05 | 0.21790383 | 0.255646643 |
| CE 28:6 | 0.034323323 | 0.034606218 | 0.040312445 | 0.033240187 | 0.03159784 | 0.055840855 | 0.00053341 | 0.00050648 | 0.00052306 | 0.00044755 | 0.00013638 | 0.00062424 | 0.000520986 | 1.10907E-05 | 0.000523351 | 7.42812E-05 | 0.96836733 | 0.006535298 |
| CE 30:5 | 0.024258033 | 0.025346818 | 0.028944489 | 0.02326215 | 0.022437872 | 0.036062571 | 0.00037699 | 0.00037097 | 0.00037556 | 0.0003132 | 0.00013638 | 0.00040314 | 0.000374505 | 2.56901E-06 | 0.000356722 | 3.67729E-05 | 0.56489982 | -0.07018651 |
| CE 30:6 | 0.019911068 | 0.020474687 | 0.024220376 | 0.018478396 | 0.016956303 | 0.028636933 | 0.00030943 | 0.00029966 | 0.00031426 | 0.00024879 | 0.00013638 | 0.00032013 | 0.000307786 | 6.07415E-06 | 0.000278769 | 3.02139E-05 | 0.30604754 | -0.14285612 |
| CE 32:6 | 0.017518142 | 0.017360042 | 0.021444955 | 0.017921216 | 0.014549115 | 0.027314345 | 0.00027225 | 0.00025408 | 0.00027825 | 0.00024129 | 0.00013638 | 0.00030534 | 0.000268191 | 1.02777E-05 | 0.000258687 | 3.33444E-05 | 0.73189002 | -0.05205298 |
