## Supplementary figures and images for "Virally induced lipid droplets are a platform for innate immune signalling complexes"

### S1

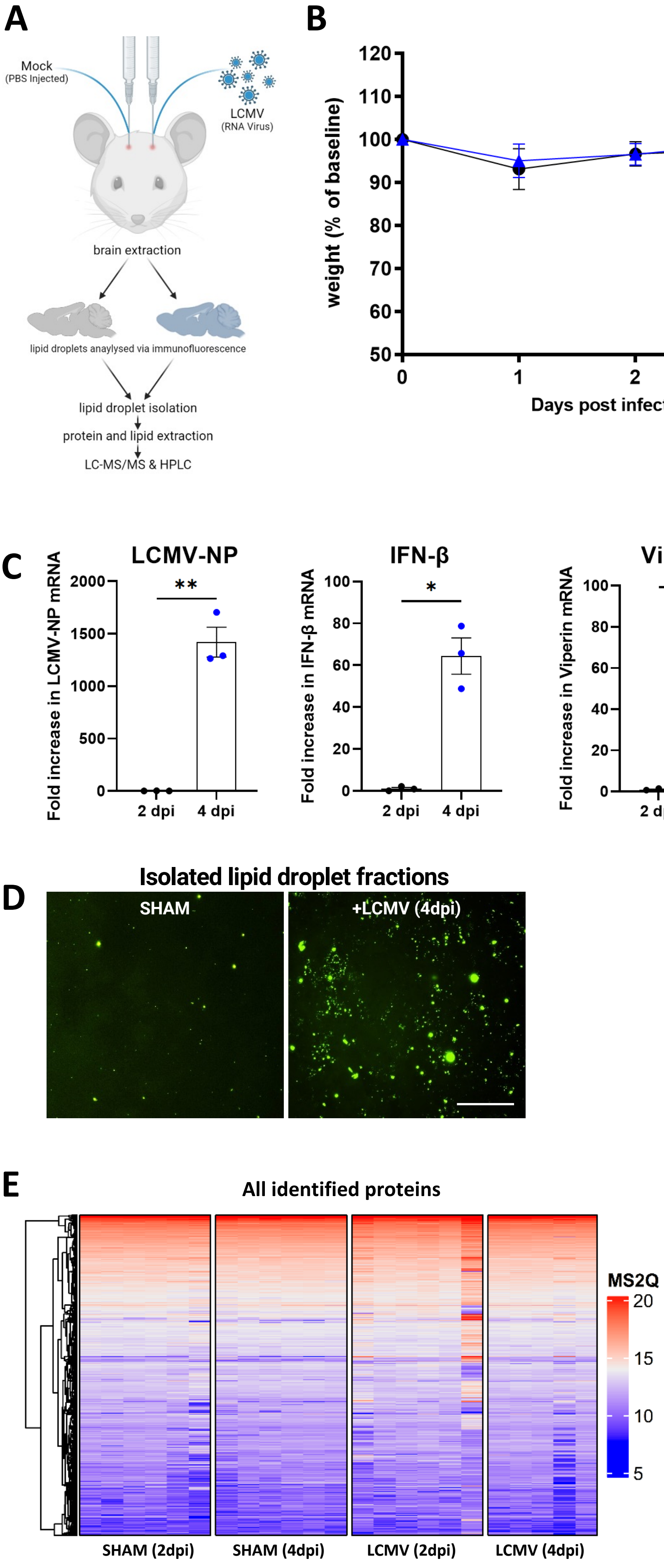

### S2

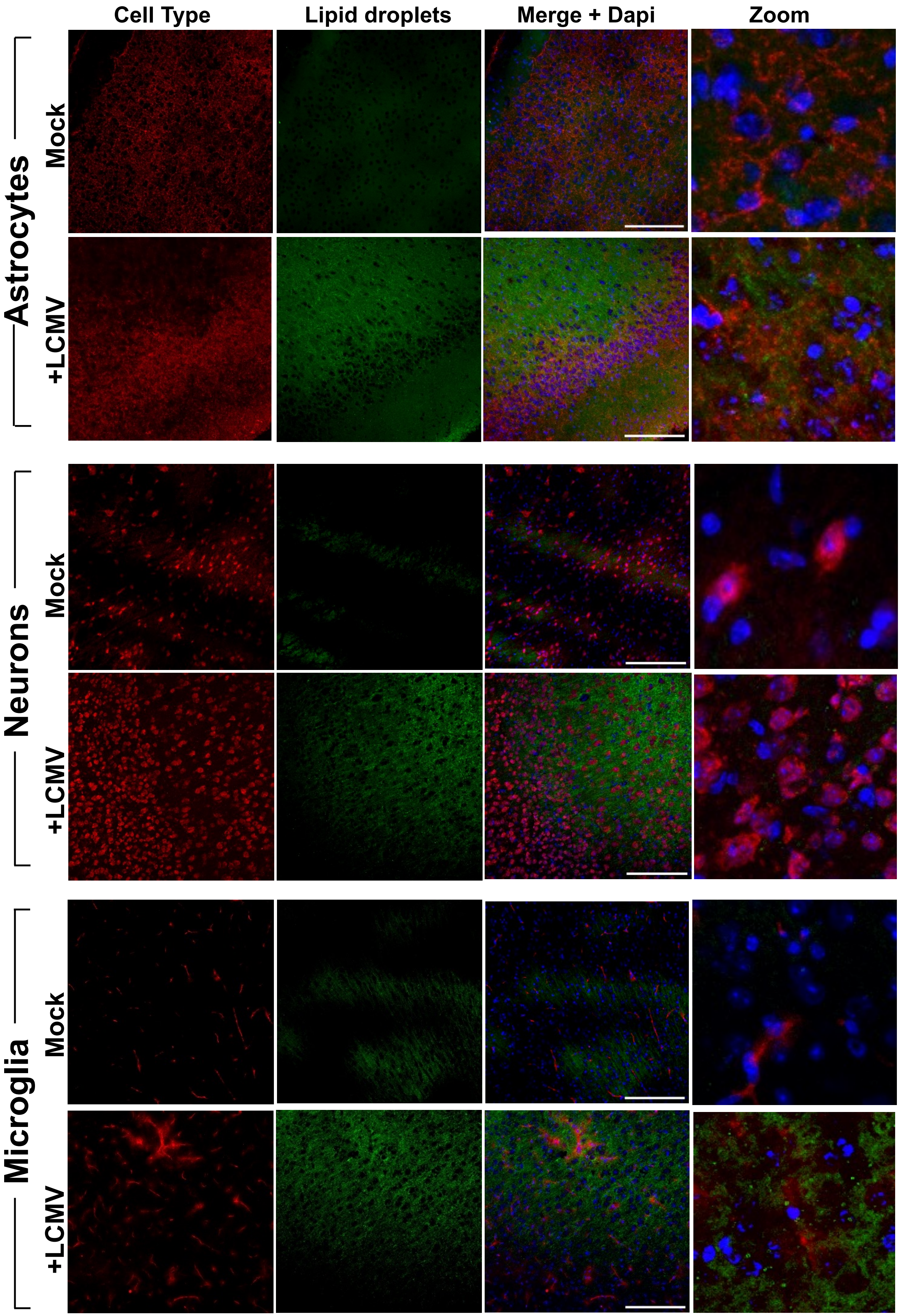

### S3

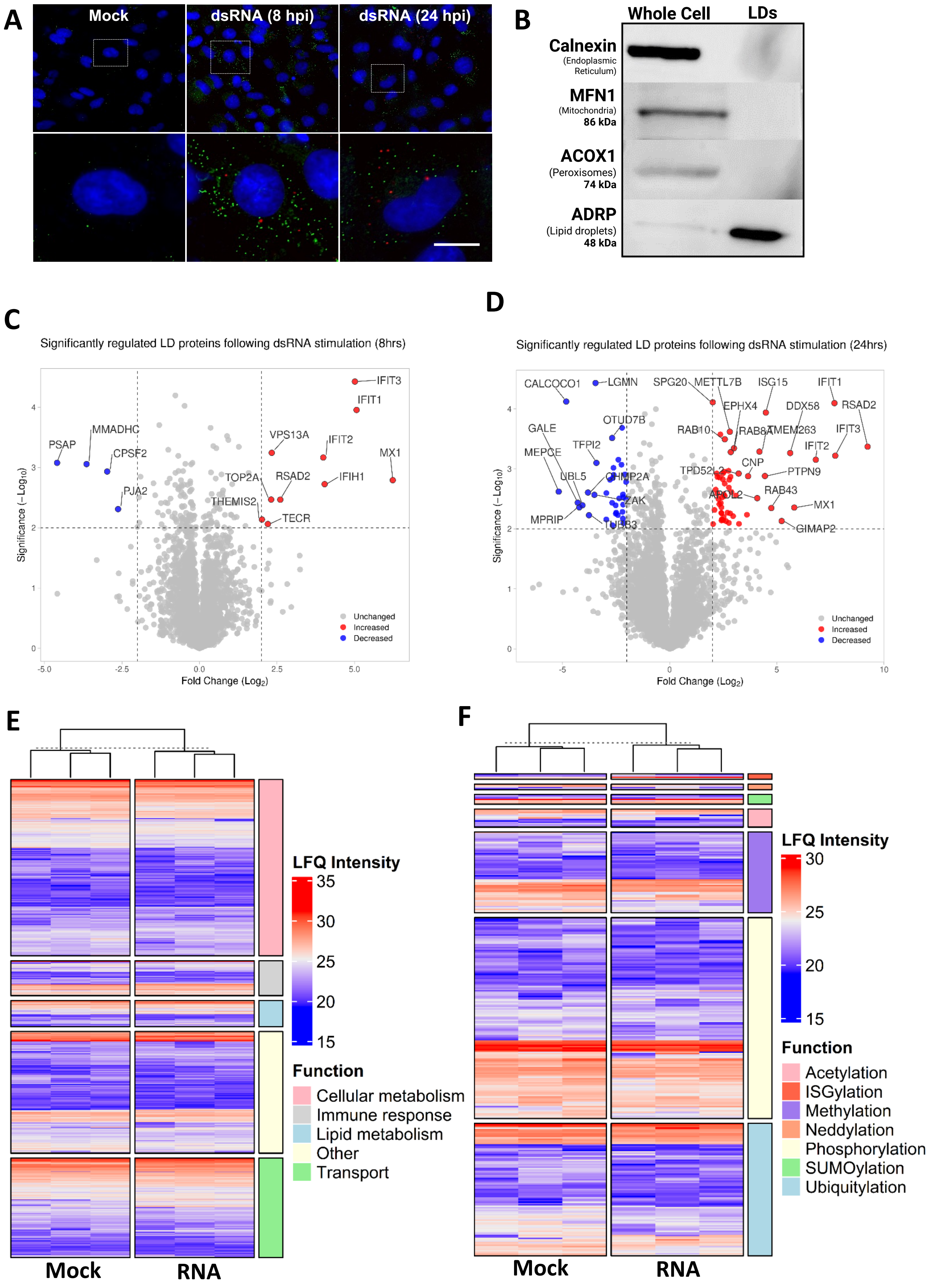

### S4

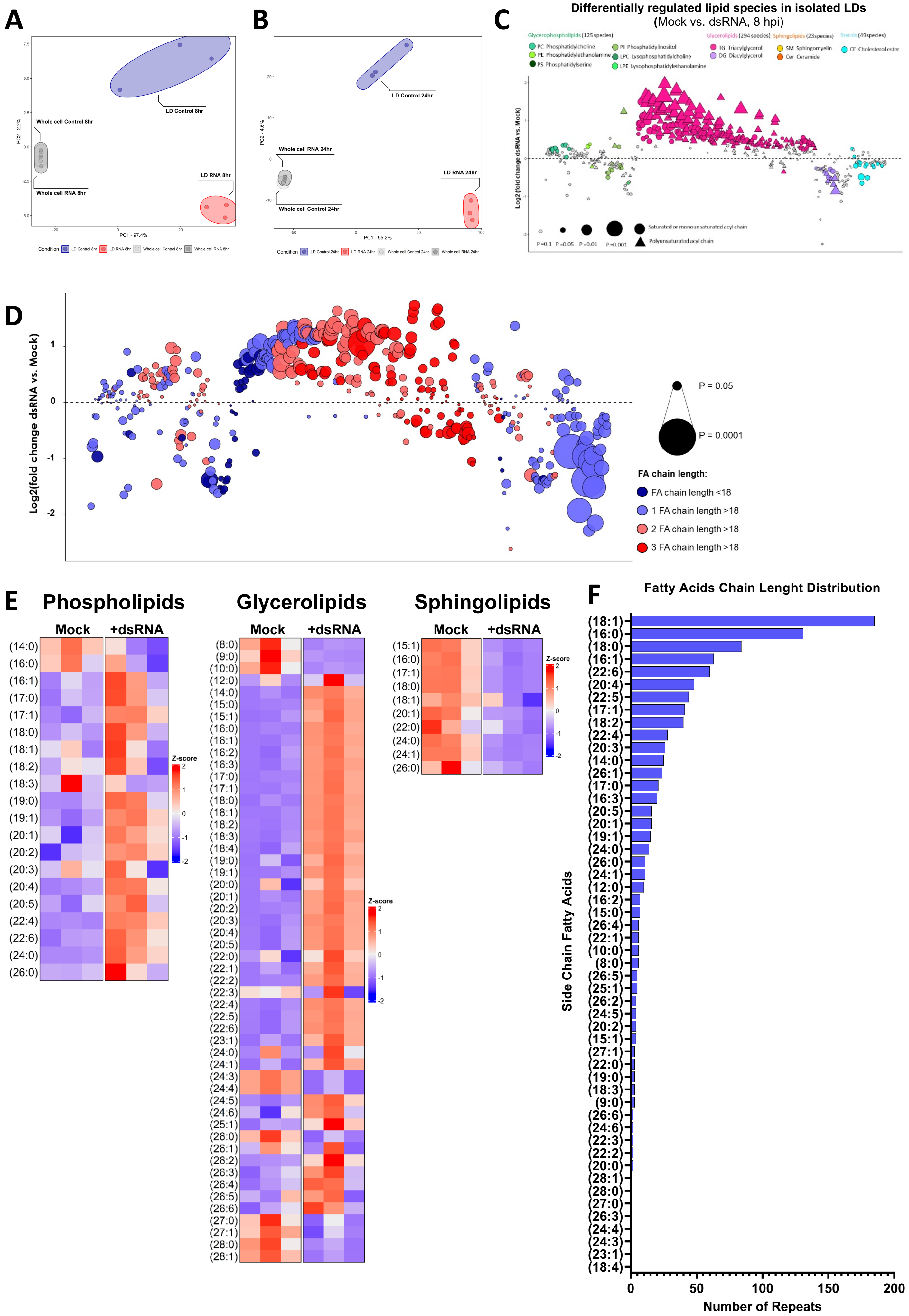

### S5

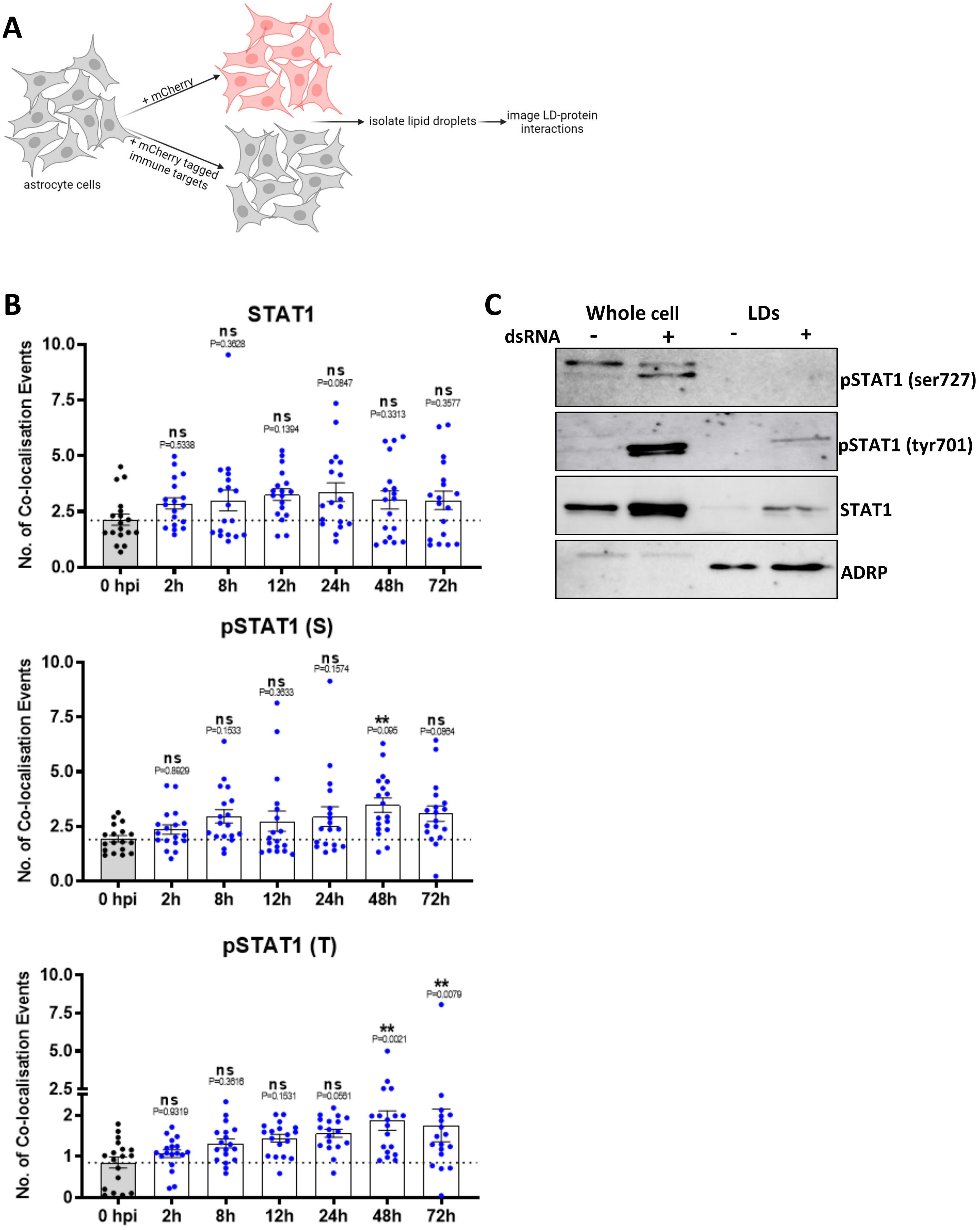

### S6

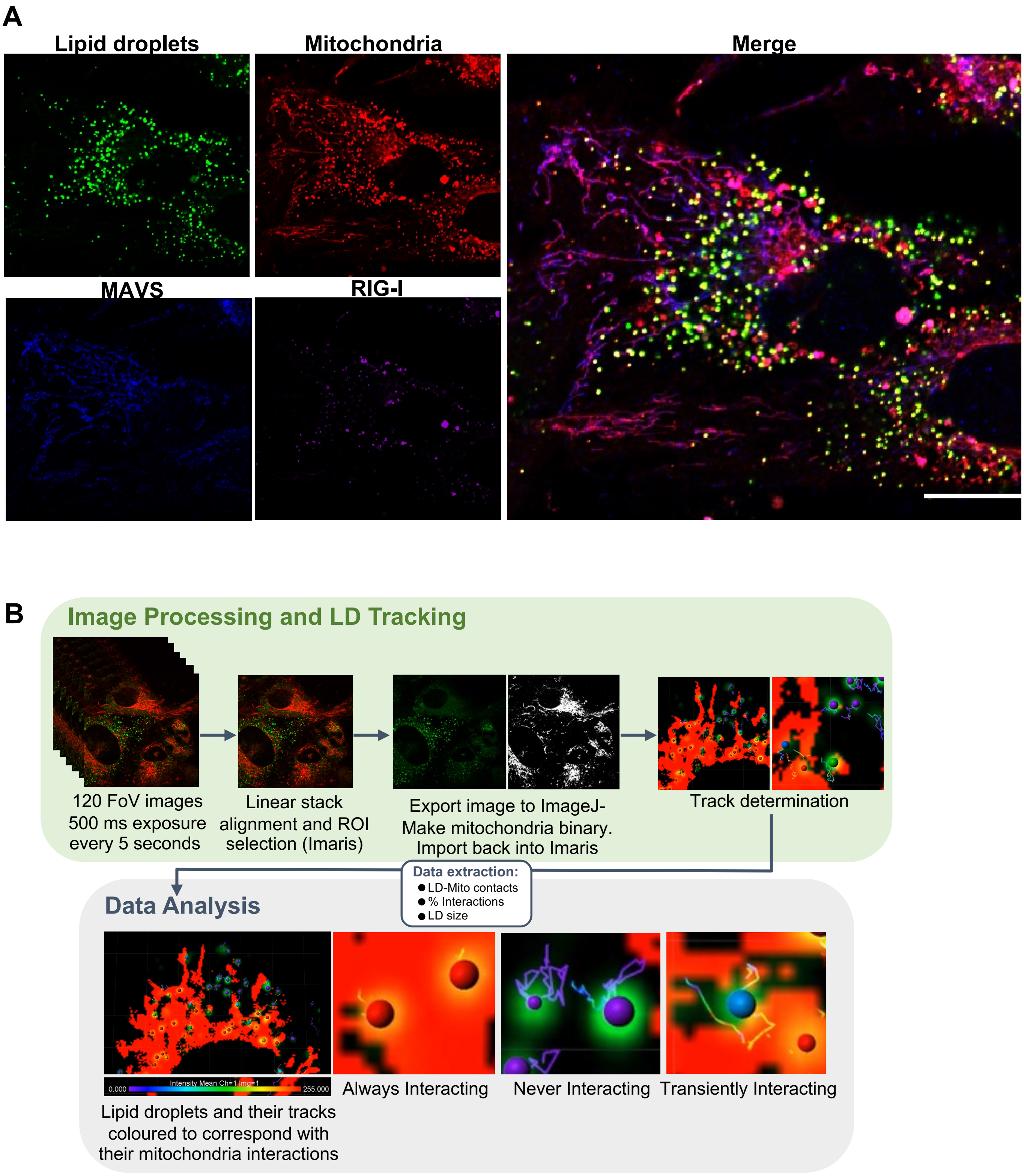

### S7

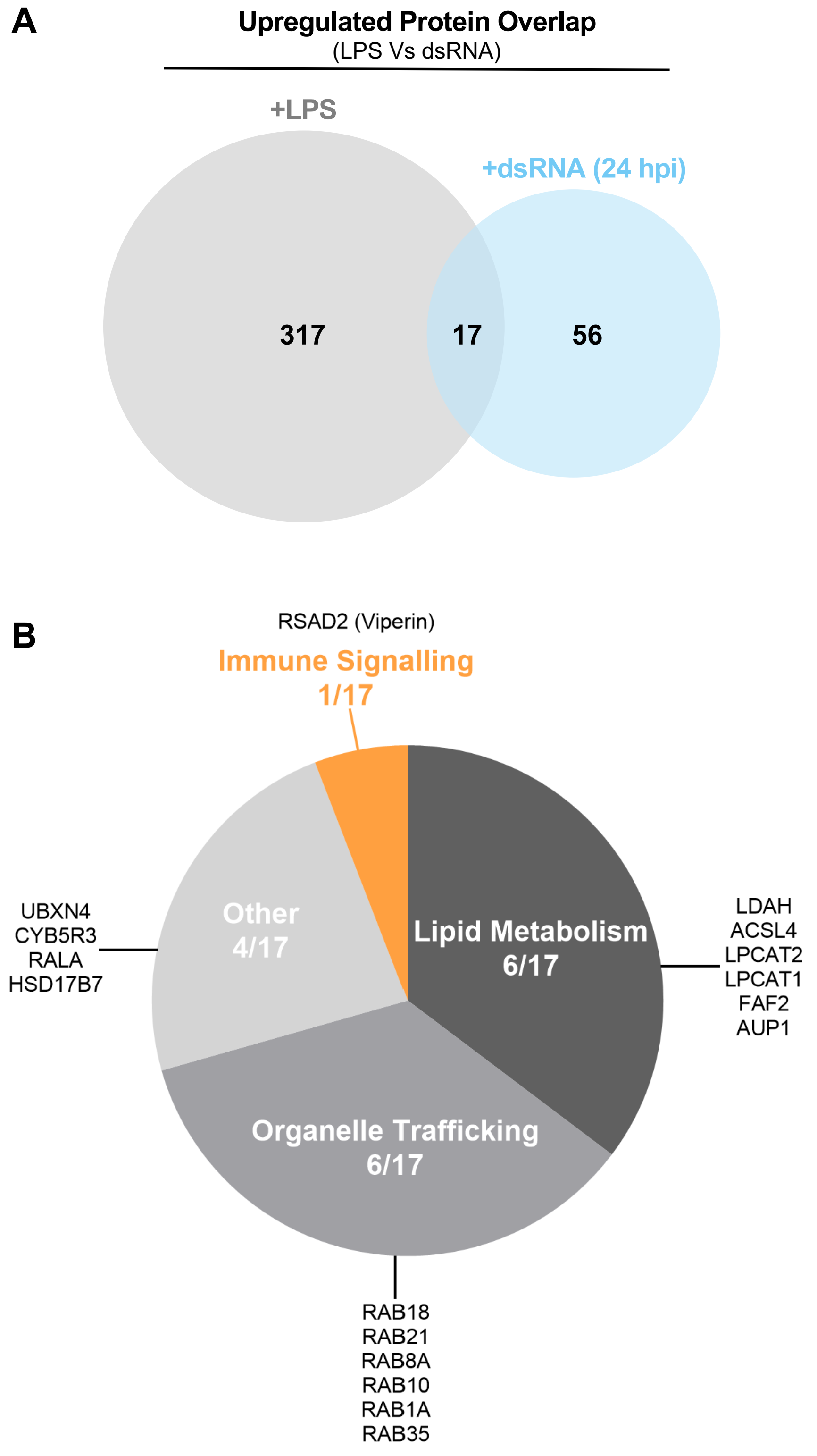
